# Identification of multi-omics molecular signature of acute radiation injury using nonhuman primate model

**DOI:** 10.1101/2021.10.16.464661

**Authors:** Amrita K Cheema, Yaoxiang Li, Joanna Moulton, Michael Girgis, Stephen Y. Wise, Alana Carpenter, Oluseyi O. Fatanmi, Vijay K. Singh

**Author notes:** **Corresponding author:** Vijay K. Singh, Ph.D., Department of Pharmacology and Molecular Therapeutics, F. Edward Hébert School of Medicine, Uniformed Services University of the Health Sciences, Bethesda, MD, USA.

## Abstract

The availability of validated biomarkers to assess radiation exposure and to assist in developing medical countermeasures remains an unmet need. We used a cobalt-60 gamma-irradiated nonhuman primate (NHP) model to delineate a multi-omics-based serum probability index of radiation exposure. Both male and female NHPs were irradiated with different doses ranging from 6.0 to 8.5 Gy, with 0.5 Gy increments between doses. We leveraged high resolution mass spectrometry for analysis of metabolites, lipids, and proteins at 1,2, and 6 days post-irradiation in NHP serum. A logistic regression model was implemented to develop a 4-analyte panel to stratify irradiated NHPs from unirradiated with high accuracy that was agnostic for all doses of γ-rays tested in the study, up to six days after exposure. This panel was comprised of Serpin Family A9, acetylcarnitine, PC (16:0/22:6), and suberylglycine, which showed 2 – 4-fold elevation in serum abundance upon irradiation in NHPs, and can potentially be translated for human use following larger validation studies. Taken together, this study, for the first time, demonstrates the utility of a combinatorial molecular characterization approach using an NHP model for developing minimally invasive assays from small volumes of blood that can be effectively used for radiation exposure assessments.

## Introduction

Nuclear disasters caused by the inadvertent or deliberate release of radioactive materials have far-reaching consequences and pose a significant threat to public health. Due to the catastrophic potential to both property and life, radiological/nuclear threats and accidents remain a priority for major federal entities and public health agencies around the globe (1, 2). Retrospective physical and biological dosimetry techniques in use include dicentrics, translocations, chromosome condensation, electron paramagnetic resonance, thermoluminiscence, hematology, and protein biomarkers among others (3-5). However, these techniques do not provide the required throughput required to screen large populations expeditiously during a mass casualty scenario. Molecular phenotyping technologies, on the other hand, can be used effectively for the development of minimally invasive biomarker assays for rapid radiation exposure assessment in afflicted sub-populations. Biomarkers also help to evaluate the impact of individual doses of radiation on biological systems, thus allowing for the administration of more precise and effective treatment regimens for the exposed population at risk, and therefore underscoring the urgency for biomarker discovery for more comprehensive assessments of ionizing radiation and the associated health risks (6). Further, biomarkers of radiation injury are also informative for assessing countermeasure efficacy that can then be used to convert drug doses deduced from animal studies to those that may be efficacious in humans (6). During the last few decades, several metabolites, lipids, proteins, and miRNA molecules have been studied as dose-dependent putative biomarkers for radiation exposure. However, these biomarker studies have been largely based on murine models and have employed a limited range of radiation exposure conditions, radiation doses, and exposure intensities; generally, these studies and observations have not been validated using higher animal models (7, 8).

In this study, therefore, we leveraged NHPs as a highly translational model system to delineate biomarkers that are indicative of radiation exposure, in compliance with the FDA-MDDT (Food and Drug Administration-Medical Device Development Tools) program. Serum samples from NHPs exposed to six different doses of γ-rays were collected at different time points post-irradiation and analyzed using a combinatorial multi-omics approach that included high resolution mass spectrometry-based proteomics metabolomics, and lipidomics. Our results demonstrate a robust, changes in circulating levels of lipids, metabolites and proteins in a dose and time dependent manner. Moreover, for the first time, we demonstrate the extrapolation of radiation-induced metabolic and proteomic disruptions for the development of a broad range, radiation dose agnostic classification algorithm for the assessment of radiation exposure in the NHP model. Importantly, the multi-analyte panel-based assay described herein can be used up to six days (SD 6) after radiation exposure, underscoring its utility for the screening, triaging, and medical management of individuals exposed to radiation.

## Results

### Exposure to Gamma-Radiation Triggers a Robust Peripheral Metabolomics and Proteomic Response

We have previously reported radiation-induced metabolic changes in mouse (9-14) and NHP models (15, 16). In this study, six NHPs per dose were exposed to gamma-radiation in the range of 6.0 – 8.5 Gy with increments of 0.5 Gy. Serum samples were collected prior to irradiation (n=36), as well as on day 1 (SD 1), SD 2, and SD 6 post-irradiation for metabolomic and lipidomic analyses (Figure 1, panel A). In order to delineate radiation-induced alterations in serum abundance of metabolites and lipids, we compared metabolic profiles of pre-irradiation samples (n=36) with post-irradiation samples for every dose and time point separately (n=108 in total) (Supplementary Table 1, Supplementary Figure 1). Not surprisingly, we observed significant dysregulation of several different classes of metabolites and lipids in a time and dose-dependent manner; a subset (n=129) of which were validated using tandem mass spectrometry (Supplementary Table 2). Overall, we observed dose agnostic dyslipidemia in NHPs upon irradiation; this included an upregulation of glycerophospholipids (PC and PE) and L-carnitine and a concomitant downregulation of Lyso PC and PE classes (Supplementary Table 3). Functional pathway analysis showed dysregulation of fatty acid metabolism, activation, and biosynthesis, specifically in linoleate metabolism across all radiation doses tested (Figure 1, panel B; Supplementary Table 4). Next, we examined alterations in metabolite abundance visualized as a heat map that showed downregulation of amino acid glutamate and derivatized amino acids like phenylacetyl glutamine in a time and dose-dependent manner (Figure 1, panel C).

**Figure 1:**
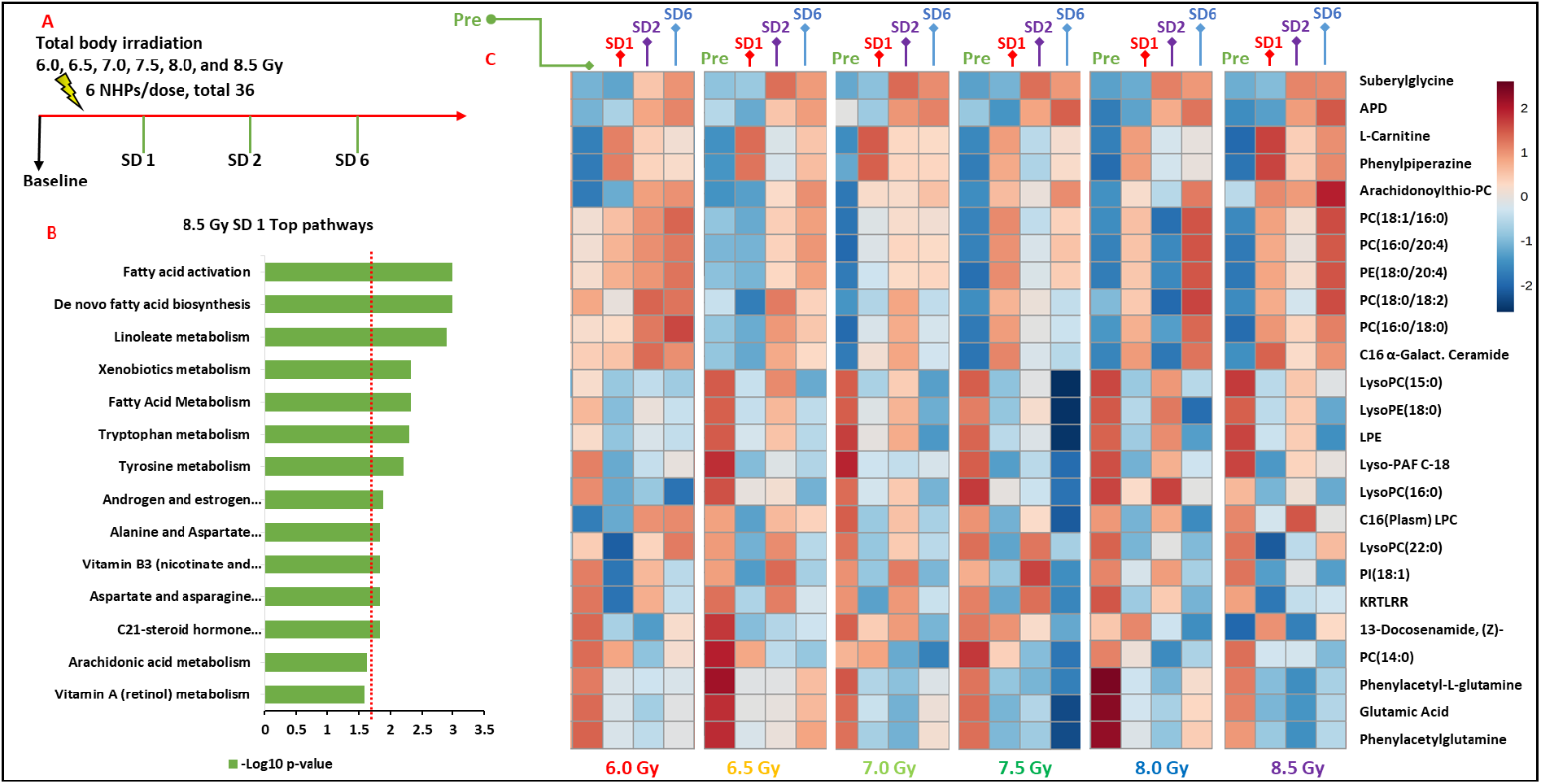
Radiation induced metabolic response in NHPs. Study schema for NHP irradiation and sample collection **(Panel A)**. Pathway analysis showing metabolic perturbations after irradiation in NHPs receiving 8.5 Gy **(Panel B)**. A heat map of most significantly dysregulated metabolites at different doses of radiation, in a time-dependent manner **(Panel C)**.

Next, we examined changes in protein abundance in the same cohort of NHPs by comparing pre and post-irradiation log transformed and TIC normalized liquid chromatography mass spectrometry (LC-MS) data. A total of 752 proteins were identified with more than 97% confidence and a false discovery rate (FDR) value of less than 1% of which 173 proteins met standard filtering and baseline correction criteria. Overall changes in protein abundance were visualized using principal component analysis that showed clear separation between pre-irradiation proteomic profiles as compared to SD 1, SD 2, and SD 6 post-irradiation (Figure 2, Panel A; Supplementary Figures 2 through 6). Proteins contributing to this group separation were examined using volcano plots for each dose and time point (Figure 2 Panels B-D); and an exhaustive list of dysregulated proteins in this survey is presented (Supplementary Table 5). We found 9 dysregulated proteins belonging to immune response and the inflammatory response pathways. Several proteins were upregulated with radiation exposure including lipopolysaccharide (LPS)-binding protein, which showed an increased serum abundance at SD 1 for all doses tested in the study, while the levels of relative abundance remained much higher at SD 6 than pre-irradiation serum abundance (Figure 3, Panel A). LPS binding protein has been previously described as a potential early biomarker for cardiac toxicity after radiation therapy (17) and as a biodosimetric marker of radiotherapy in cancer treatments (18-20). On the other hand, inter-alpha globulin showed a significant decrease after irradiation for all doses with serum abundance that remained significantly dysregulated even after SD 6 from radiation exposure (Figure 3, Panel B,). We also found proteins like C4a anaphylatoxin and Serpin Family A member 9 protein that displayed an oscillatory pattern with respect to serum abundance at SD 6 as compared to SD 1 and 2; interestingly, however, the trend across the three time points was remarkably similar for all doses tested suggesting a commonality of the biologic basis to radiation response (Figure 3, Panels C and D). The relative abundance of Serpin A9, C4 alpha anaphylataxin, C-reactive protein and Haptoglobin were validated using Western blot analysis across different doses and time points (Figure 4, Panels A and B).

**Figure 2:**
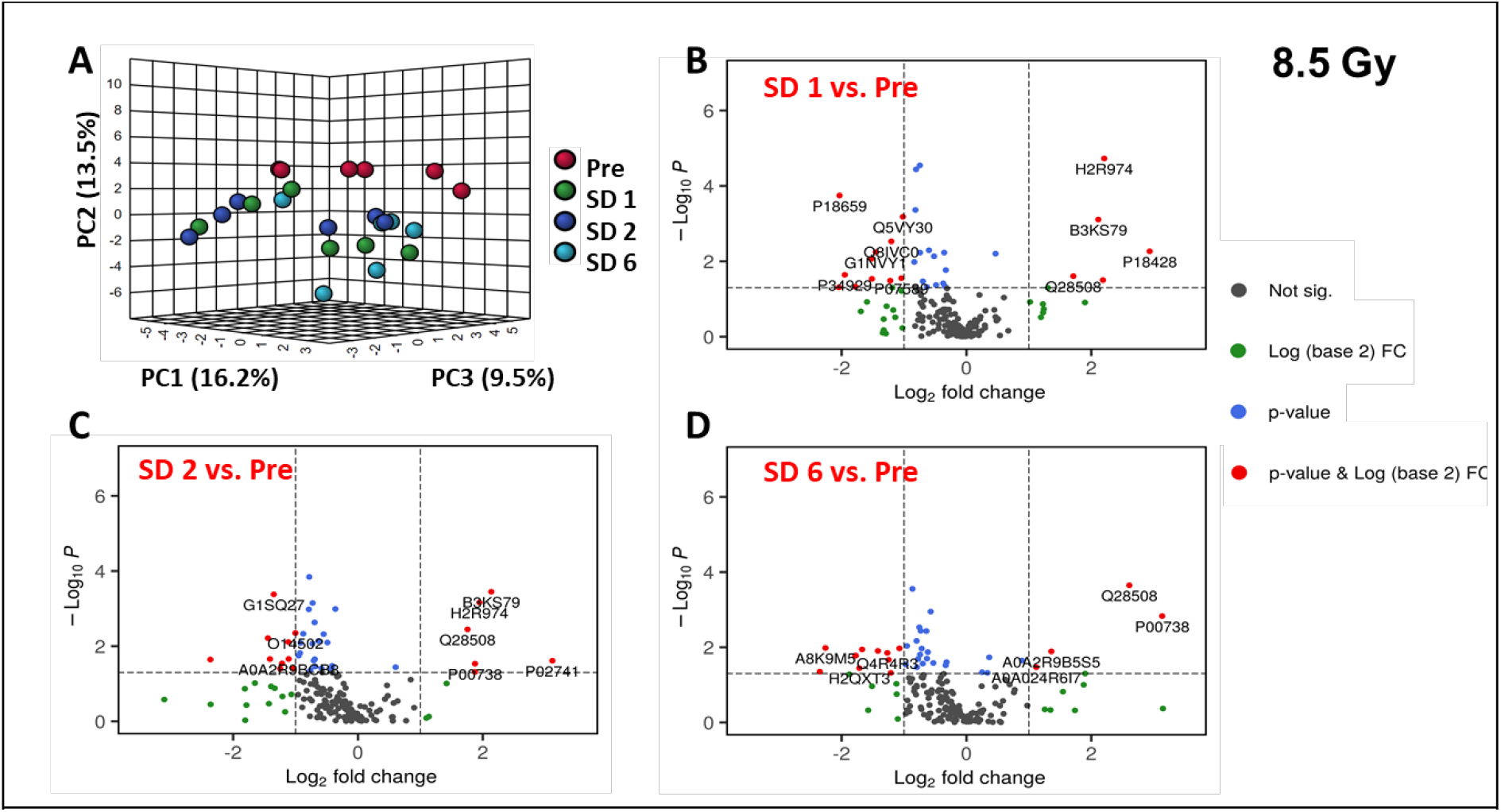
Multivariate analysis show robust alterations in proteomic profiles upon irradiation in NHPs. Principal component analysis comparing overall changes of the proteomic profiles of NHPs exposed to 8.5 Gy as compared to baseline at days 1 (SD 1), 2 (SD 2) and 6 (SD 6) post-irradiation. **(Panel A)**. Significantly dysregulated proteins upon irradiation at 8.5 Gy were visualized as volcano plots wherein dysregulated proteins that met either p-value or fold change cut off criteria were marked in green and blue, respectively, while those that met both were marked in red, and non-significant ones were denoted in black **(Panels B, C, and D)**.

**Figure 3:**
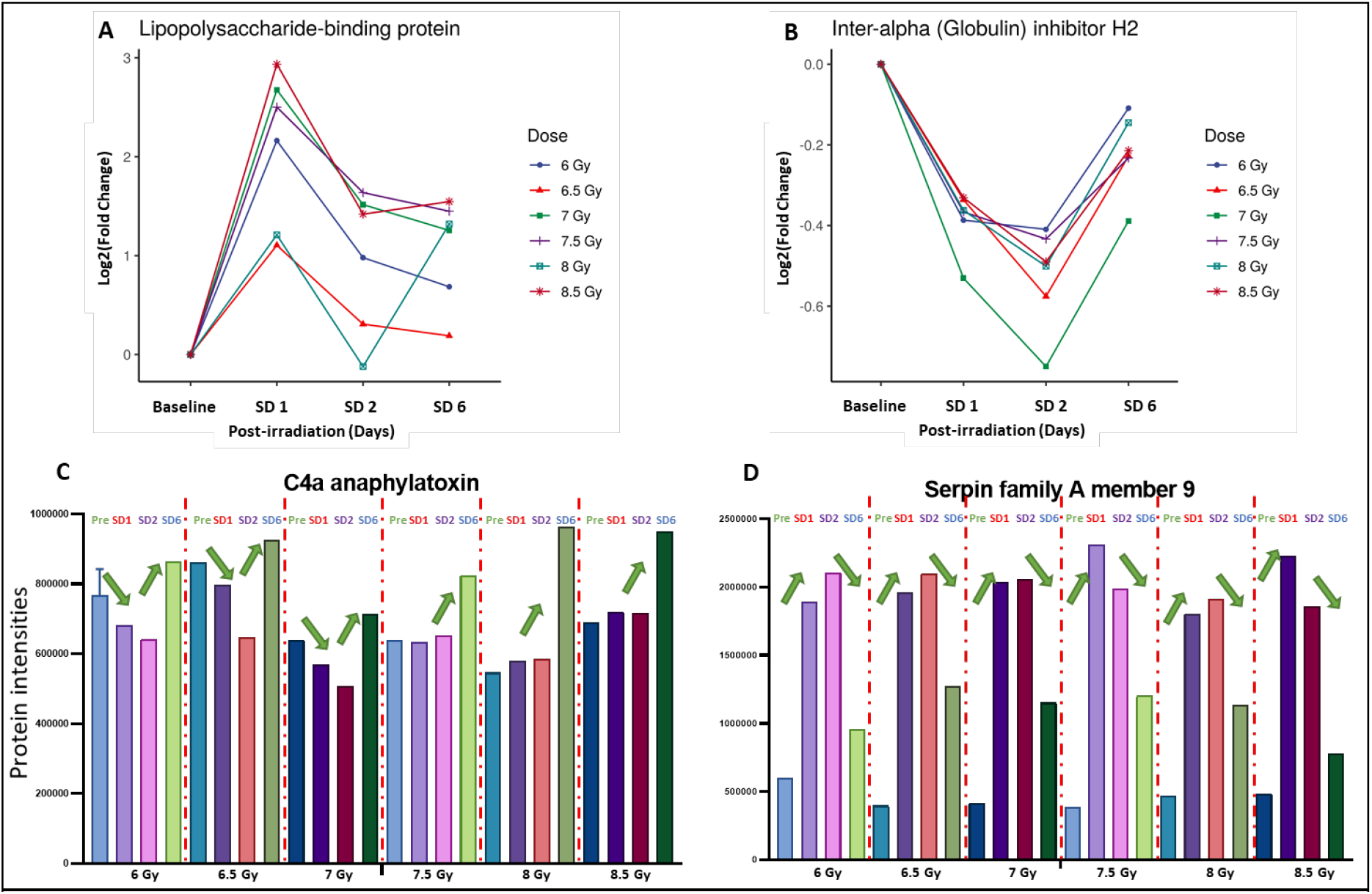
Oscillatory patterns of serum protein abundance in irradiated NHPs. Temporal patterns of protein abundance for all radiation doses as a function of time visualized as trend lines **(Panels A and B)** or as bar graphs **(Panels C and D)** for a subset of proteins.

**Figure 4:**
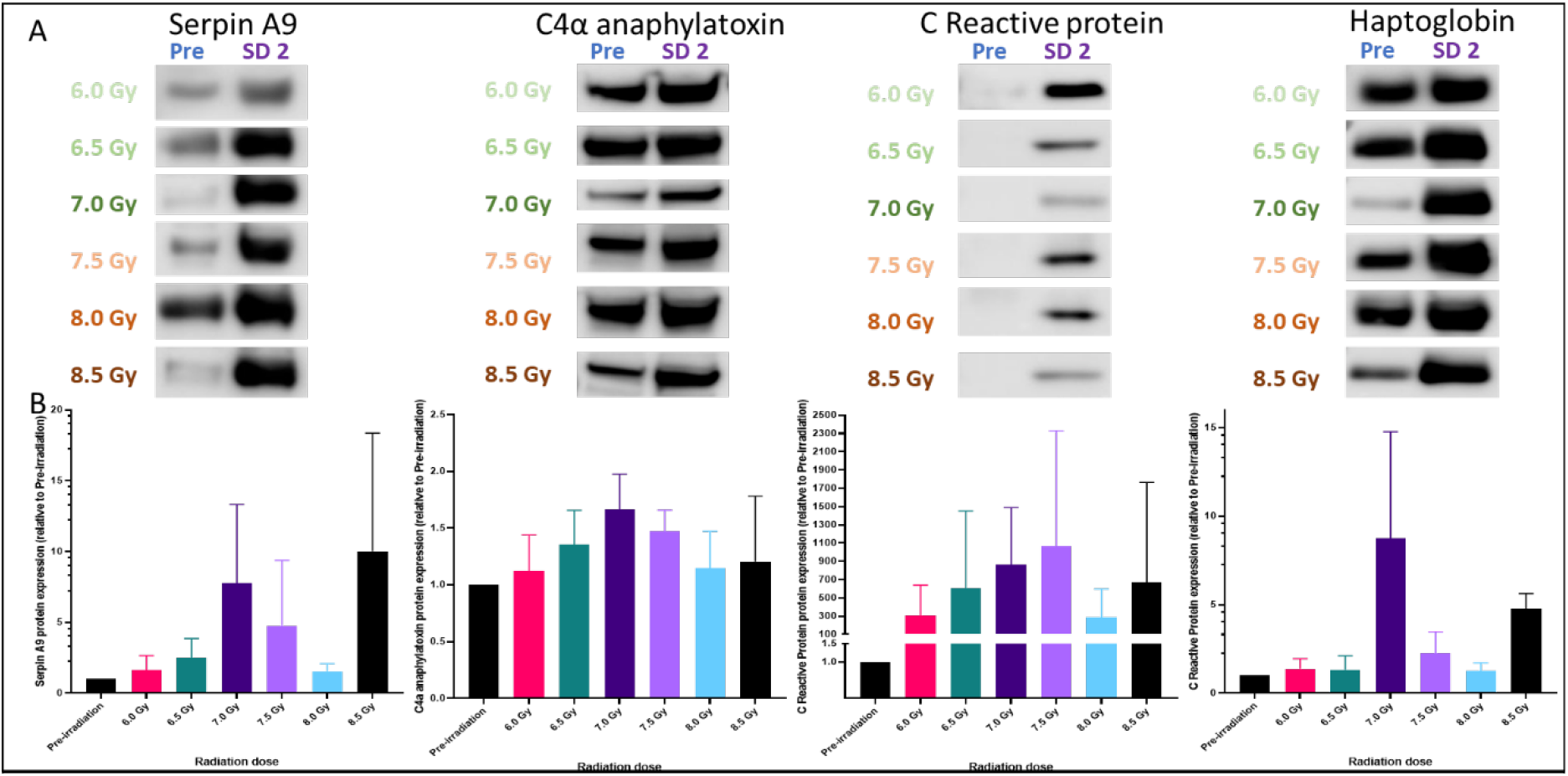
Validation of discovery proteomics data for a select group of proteins using Western blot analysis. Relative serum abundance comparing SD 2 and pre-irradiation samples for different doses (6.0 Gy, 6.5 Gy, 7.0 Gy, 7.5 Gy, 8.0 Gy, and 8.5 Gy) was visualized for Serpin A9, C4α anaphylatoxin, C Reactive protein, and Haptoglobin **(Panel A)**. Protein expression level relative to pre-irradiation was performed with four replicates across all doses for each protein while the quantification and standard deviation was determined using ImageJ software. **(Panel B)**.

### Multi-omics Biomarker Panel for Prediction of Radiation Exposure

The availability of minimally invasive, high throughput, sensitive, and specific biomarker assays for patient triage remains a critical area of need in radiation research. Our molecular characterization using multi-omics approach offers the development of FDA compliant biomarker panels from very small volumes of blood that can be used for rapid testing, triaging, and medical management in the event of a large-scale radiological event. In this study, 10 μl of serum was used for proteomic profiling while 25 μl was used for serum lipidomics and metabolomics yielding high quality discovery data that was used for the development of biomarker panels.

To facilitate prediction feature selection compliant with the FDA-MDDT program, all features (metabolites, lipids, and proteins) were first checked for missing values (less than limit of detection), and features with >20% missing values were excluded from the selection process. Each study includes a set of pooled quality control (QC) duplicates to enable QC-relative standard deviation (RSD) < 15% filter for quality control. Paired *t*-test and Wilcoxon test were applied to further eliminate non-significantly changed features. Elastic net regression were applied for feature pre-selection and biological relevance were also taken into consideration for final selection (21). We used a logistic regression model to fit analytes that could be used for predicting radiation exposure. This resulted in the selection of a 4-analyte biomarker panel that shows a near 100% efficacy of prediction across all doses used. Interestingly, this panel works for all doses and until day 6 after exposure, which suggests a wide utility for assessing exposure 24 h after radiation exposure and up to SD 6 after a radiological event. The panel included PC (16:0/22:6), Acetyl-carnitine, Serpin Family A9, and Suberylglycine. To build the dose-dependent radiation effect prediction model, time points including SD 1, SD 2, and SD 6 of sample collection are combined as an “acute after radiation group (n=18)” for each dose. The performance of this panel was tested for all six doses that yielded a near perfect classification accuracy (Figure 5 panels A-F, Supplementary Table 6). Next, we developed dose specific multi-analyte panels to predict radiation exposure (Figure 6, panels A-F). For example, a four-analyte panel including **PC 18:0/18:2**, albumin, haptoglobin, and LPS-binding protein resulted in high prediction accuracy of exposure to 6.0 Gy AUC (area under the receiver operating characteristic (ROC) curve) ≥ 90%, while C4a anaphylatoxin, angiotensin 1-10, ITIH4 protein, and complement C4-B comprised a high accuracy prediction panel for exposure to 6.5 Gy. Similarly, panels that included haptoglobin, N-acetylmuramoyl-L-alanine amidase, beta actin variant, leucine-rich alpha-2-glycoprotein for 7 Gy; protein S100-A7, protein S100, leucine-rich alpha-2-glycoprotein for 7.5 Gy; epididymis secretory protein Li 37, protein S100, C4a anaphylatoxin for 8.0 Gy; protein S100-A7, and complement C4-B for 8.5 Gy, were developed to predict exposure at specific radiation doses. The index panel for each dose was calculated based on its logistic regression model and probability prediction index (PPI) that were plotted for each individual sample to distinguish between irradiated and control NHPs (pre-irradiation). A threshold calculated index values are associated with classification of irradiated NHPs with confidence transitioning from 90% to 100%. For example, based on calculated PPI in our dataset, a value ≥ 2 represents a true irradiated NHP (at 6.0 Gy) with 90% confidence interval (Figure 6, Panel A). Following validation studies, this approach may be extended as a diagnostic tool for assessment of radiation exposure in the event of a radiological emergency.

**Figure 5:**
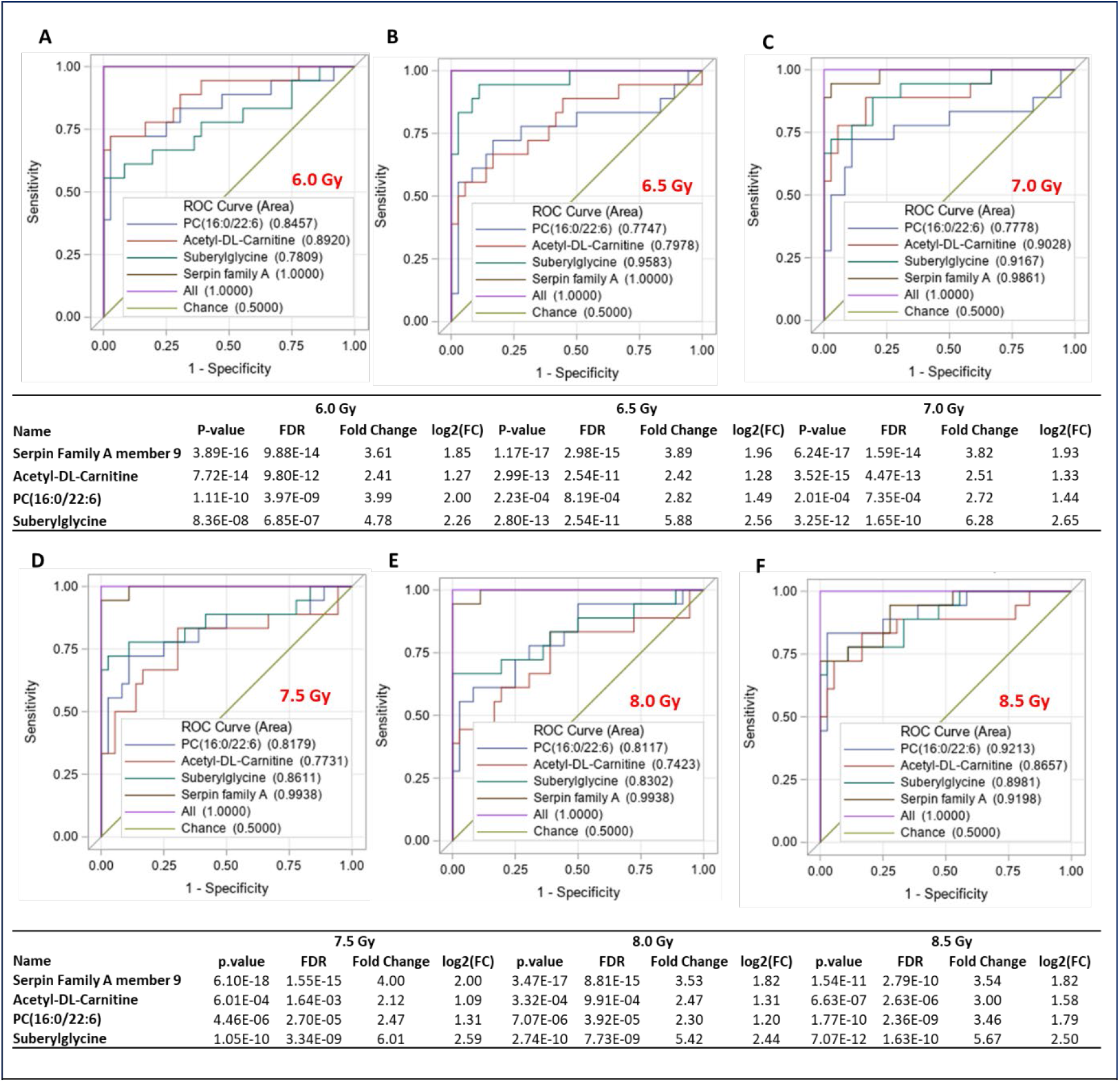
Development of a 4-analyte multi-omics panel for assessment of radiation exposure in NHPs. A logistic regression model was used for the selection of a biomarker panel that showed a very high prediction efficacy across all tested doses **(Panels A through F)**.

**Figure 6:**
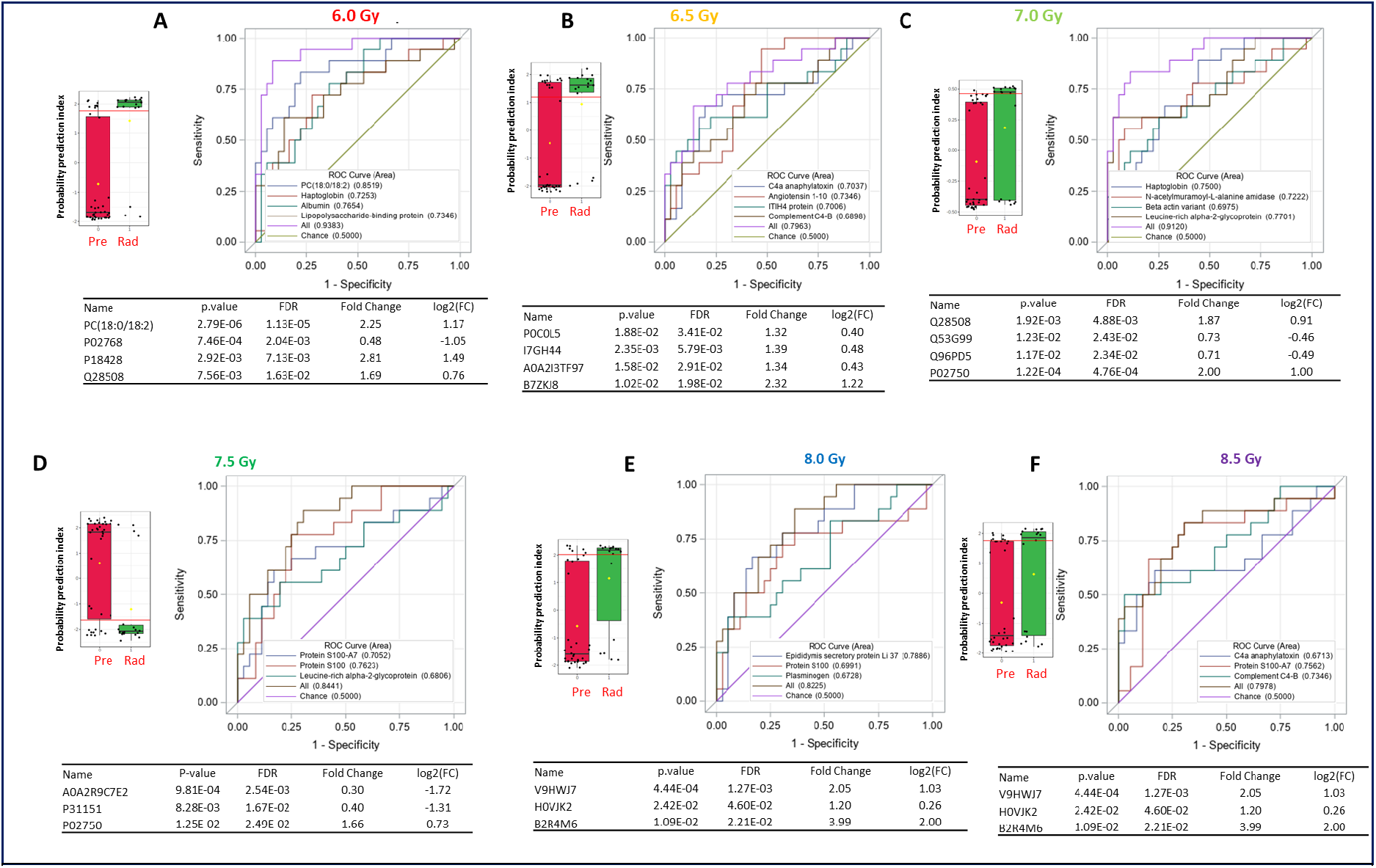
A dose-specific multi-omics biomarker panel to predict the extent of exposure to gamma radiation. A logistic regression model was used for the selection of a biomarker panel that showed a very high prediction efficacy for each dose tested in this study **(Panels A through F)**. Each panel shows a ROC curve showing specificity and sensitivity for the multi-analyte panel described in the table as well as a probability index plot showing threshold for separating irradiated and unirradiated NHPs with >95% confidence interval using the multi-analyte panel described for each dose. Solid red horizontal line within the boxplots represents mean value. Error bars represent standard deviation from mean.

## Discussion

Blood based multi-omics biomarker assays represent a new frontier for the stratification of individuals after exposure to ionizing radiation as compared to conventional techniques (22-24). While several studies have reported metabolite, protein, or miRNA-based biomarkers for predicting radiation exposure, few have been validated in higher animal models for reproducibility and possible clinical translation (6). To develop a potentially translatable prediction model of radiation exposure, we examined serum samples collected from NHPs exposed to a range of gamma-radiation between 6 to 8.5 Gy with increments of 0.5 Gy. We used a high throughput metabolomic, lipidomic, and proteomic profiling approach to create a reliable and reproducible classification model comprised of four analytes that can augment stratification of selectively exposed individuals with high accuracy (AUC > 99%). While there are several reports on the use of omics-based approaches for biomarker development to predict radiation exposure using murine models (25), to the best of our knowledge, this is the first comprehensive multi-omics serum analysis conducted using an NHP model. Our proposed panel encompasses PC (16:0/22:6), Acetyl Carnitine, Suberyl glycine, and Serpin family A protein. PC (16:0/22.6) is a glycerophospholipid in which a phosphorylcholine moiety occupies a glycerol substitution site. It is a key component of the lipid bilayer and a great contributor in metabolism and cellular signaling (26-28). Acetyl carnitine is a source of levocarnitine, an amino acid derivative that facilitates long-chain fatty acid translocation through the mitochondrial membrane to deliver fatty acid substrate for oxidation and subsequent energy production (29). Acetyl carnitine increases in response to exposure as a counter mechanism to ameliorate the damaging effects of ionizing radiation (29-31). Suberyl glycine is an N-acylglycine, which may indicate medium chain acyl-CoA dehydrogenase (MCAD) deficiency leading to a perturbation in fatty acid metabolism (32). Serpins (33), an acronym originated from “Ser-ine P-rotease In-hibitor” (33), is a family of 45-50 kDa proteins that are capable of inhibiting serine and cysteine protease activities irreversibly. They are notable for their unusual mechanism of action in that they irreversibly inhibit their target protease enzyme by undergoing numerous conformational modifications to disrupt its active site. Serpins are key modulators of the inflammatory process; additionally, Serpins9 is known to inhibit Caspase-1 (interleukin-1β-converting enzyme), which is the building block for the caspase mediated inflammatory response (34). Interestingly, we found that the serum levels of Serpin increased significantly 1 – 2 days after exposure, making it a strong indicator of acute and intense radiation exposure. This increase in serpin level can be interpreted as a natural response of the body to the acute inflammation occurring after radiation exposure.

The methodology used for the development of this multi-omics assay is following the FDA’s MDDT program. Moreover, for the first time, we report a 4-analyte Rad-Ex (Radiation Exposure) assay that has been developed in an NHP model with a high translational value. Feature selection methodology used in this study is following FDA biomarker qualification guidelines for test statistics and control of false discovery rate (<5%). The Rad-Ex assay works for males and females with similar efficacy over a wide range of radiation doses (6.0 – 8.5 Gy) that extend from low lethal to supralethal dose levels and over a broad time window after exposure (day 1 to day 6), which underscores the wide applicability of this biomarker assay following analytical and clinical validation studies.

While the study has provided an essential foundation for future developmental and application-type work of a blood based, multi-omics assay that would serve to identify, via a ‘molecular signature’, individuals exposed to hazardous levels of ionizing radiation, there are, nevertheless, some shortcomings that need to be addressed in future studies. For example, despite the robust performance of our classification algorithms, further studies need to be conducted to evaluate the performance of the model in humans either irradiated unwantedly or intentionally as a consequence of radiation therapy. The assay’s specificity relative to the type of radiation exposure (e.g., gamma ray, X-rays, or mixed field gamma-neutron exposures) needs to be developed as well. The impact of basic radiation exposure conditions, e.g., total dose, dose-rate, and radiation quality, all need to be evaluated, along with the biological aspects of partial-body exposures on assay performance. Although the radiation dose range tested here provides a fairly broad range survey of proteomic and metabolomic changes, additional high accuracy predictive panels need to be developed in NHPs, not only for the extremely high, supra-lethal exposures, but more importantly at sublethal/non-lethal levels of radiation exposures.

In conclusion, the results of this study validate the use of metabolomics/lipidomics and proteomic signatures as biomarkers to evaluate the extent of radiation injury which may be used to assess the efficacy of potential radiation countermeasures.

## Material and Methods

### Animals and Animal Care

A total of 36 naïve rhesus macaques (*Macaca mulatta*, Chinese sub-strain, 19 males and 17 females) 2.9 – 7.3 years of age, weighing 3.6 – 7.4 kg, were procured from the National Institutes of Health Animal Center (NIHAC, Poolesville, MD, USA) and maintained in a facility accredited by the Association for Assessment and Accreditation of Laboratory Animal Care (AAALAC)-International. Animals were quarantined for six weeks prior to initiation of the experiment. Animal housing, health monitoring, care, and enrichment during the experimental period have been described earlier (9). Animals were fed a primate diet (Teklad T.2050 diet; Harlan Laboratories Inc., Madison, WI, USA) twice daily with at least six hours between feedings (animals were fed four biscuits each at 7:00 AM and 2:00 PM) and received drinking water *ad libitum*. All procedures involving animals were approved by the Institutional Animal Care and Use Committee (Armed Forces Radiobiology Research Institute Protocol #2015-12-011 and University of Maryland Baltimore Protocol #0581005) and the Department of Defense Animal Care and Use Review Office (ACURO). This study was carried out in strict accordance with the recommendations made in the *Guide for the Care and Use of Laboratory Animals* (35). This study was performed with full supportive care including whole blood transfusion (36).

### Radiation Exposure

For irradiation, two NHPs were placed on the irradiation platform facing away from each other and were exposed to a midline dose (6.0 to 8.5 Gy) ^60^Co γ-radiation at a dose rate of 0.6 Gy/min from both sides (bilateral, simultaneous exposure) as described earlier (9). To minimize radiation-induced vomiting, food was withheld approximately 12 – 18 h prior to irradiation. Approximately 30 – 45 min prior to irradiation, NHPs were administered 10 – 15 mg/kg of ketamine hydrochloride intramuscularly for sedation, then placed in custom-made Plexiglas irradiation boxes and secured in a seated position. To deliver the precise radiation dose, NHP abdominal widths were measured with digital calipers. Animals were observed throughout the irradiation procedure via in-room cameras. After irradiation, the animals were returned to their cages in the housing area and were monitored for recovery from the procedure. The radiation field in the area of the NHP location was uniform within ±1.5%. The dosimetry for photons was based on the alanine/EPR (electron paramagnetic resonance) dosimetry system (37). This is one of the most precise dosimetry techniques at present which is used by national standards laboratories for the most critical measurements and calibrations. Thus, it is one of the very few methods that are used in regular worldwide inter-comparisons of the national standards of Gray.

### Serum Sample Collection

NHP blood samples were collected by venipuncture from the saphenous vein of the lower leg after the site was cleaned using a 70% isopropyl alcohol wipe and dried with sterile gauze as described earlier (38). All animals were restrained using the pole-and-collar method and placed in a chair for blood collection. The blood draw was conducted between 08:00 AM and 10:00 AM, 1 – 3 hours after animals were fed. A 3 mL disposable luer-lock syringe with a 25-gauge needle was used to collect the desired volume of blood. For serum collection, blood samples were transferred to Capiject serum separator tubes (3T-MG; Terumo Medical Corp, Elkton, MD, USA), allowed to clot for 30 min, and then centrifuged for 10 minutes at 400 x g. Serum samples were stored at -70 °C until shipped on dry ice to the Georgetown University Medical Center (Washington, DC, USA) for global metabolomic, lipidomic, and proteomic analysis.

### Western Blot Analysis

Serum samples were depleted of the top 14 interfering high-abundant proteins using the Multiple Affinity Removal Column Human 14 (Agilent, USA). Then, the samples were rebuffered in MPER lysis buffer. Bradford assays were performed on all samples to determine protein concentrations. LDS sample buffer, DTT, and DNase/RNase-free distilled water were added to the lysates (20µg of protein in 30µL total volume), and the samples were boiled for 10 minutes before being separated by electrophoresis using 4 – 12% NuPAGE Bis-Tris gel (Invitrogen). The proteins were then transferred into a PVDF membrane (iBlot 2 Transfer Stacks, Invitrogen) and washed with Ponceau S red stain to confirm the successful transfer of proteins to the membranes. The membranes were washed with TBST and blocked for 1 hour with 1% BSA in TBST. Next, the membranes were incubated for 1.5 hours at room temperature (Serpin A9 dilution 1:1000; cat. no. LC-C406181; LifeSpanBio, C -reactive protein dilution 1:1000; cat. no. 66250-1-Ig; Proteintech, C4 anaphylatoxin dilution 1:1000; cat. no. sc-271181; Santa Cruz Biotech, and haptoglobin dilution 1:1000; cat. no. 66229-1-Ig; Proteintech) with the indicated antibodies. Immune complexes were detected by using horseradish peroxidase-conjugated antibodies to rabbit IgG or mouse (1:5000). Membrane images were captured using Amersham Imager 600 and analyzed using the ImageJ program.

### Proteomics Analysis

A serum volume of 10 µL was diluted ten times with load/wash buffer solution (Agilent Cat.#: 5185-5987). Each sample was filtered through a 0.22-μm spin filter (Agilent Cat.#: 5185-5990) for 2 min at 10,000 × g. Multiple Affinity Removal System (MARS14, Agilent Cat.#: 5188-6557) columns (4.6 × 50 mm) was used to deplete the most abundant 14 proteins in serum according to the manufacturer protocol on a Waters Acquity HPLC system. Protein peaks were selectively collected and transferred to Spin Concentrators, 5 K MWCO, 4 mL capacity (Agilent Cat. #: 5185-5991) and centrifuged at 17,172 × g for 45 min at 4 °C. A volume of 1 mL of LCMS grade water was added to the concentrated sample and centrifuged at 17,172 × g for 45 min at 4 °C to wash out salts and this process was repeated one more time. A volume of 1 mL of 50 mM ammonium bicarbonate solution (pH of approximately 8) was used to rebuffer each sample of concentrated proteins. The final protein concentration was determined using a BCA protein assay kit (Thermo Fisher Scientific Cat. #: 23227). Protein concentration was adjusted to 10 μg/100 μL in 50 mM ammonium bicarbonate buffer, reduced with Dithiothreitol (DTT) to a final concentration of 5 mM for 60 min at 56 °C, and alkylated with iodoacetamide to a final concentration of 15 mM for 30 min at 37 °C in a dark environment with Lys-C (Promega, Cat # V1671) at a 1:100 (wt/wt) enzyme-to-protein ratio for 3 h at 37 °C. Trypsin Gold (Promega, Cat # V5280) was added to a final 1:50 (wt/wt) enzyme-to-protein ratio for overnight digestion at 37 °C for 17 hours to complete the reaction. Trypsin and Lys-C enzymes were deactivated by heating at 90 °C for 10 min, and samples were allowed to cool down to room temperature before being acidified to a pH = 3 using 0.1% trifluoroacetic acid in water. The digested peptides were purified on microspin C18 columns (The Nest Group Inc., HEM S18V) and eluted using 2 × 400 µL of 80% H_2_O acidified by 0.1% formic acid followed by 2 × 400 µL of 50% acetonitrile (ACN)/H_2_O acidified by 0.1% formic acid. The collected fractions were evaporated under vacuum and reconstituted for LC-MS for SWATH analysis.

### Serum Proteomics Using NanoUPLC-MS/MS

A sample pool from all the subjects (with equal amount of proteins combined) was prepared to establish a protein library of all serum proteins detectable with tandem mass-spectrometry (MS/MS). The pooled peptide sample was fractionated with a 2.1 mm × 150 mm column XTerra MS C18 column (Waters) on an Agilent Infinity II HPLC instrument with a Diode Array Detector HS detector in high pH reversed-phase chromatography mode. Solvent A (2% ACN, 4.5 mM ammonium formate, pH =10) and a nonlinear increasing concentration of solvent B (90% ACN, 4.5 mM ammonium formate, pH =10) were used to separate peptides. The 95-min separation LC gradient followed this profile: (min: %B) 0:2; 5:5; 70:50; 75:75; 80:100; 90:2; and 95: 2. The flow rate was 0.2 mL/min. A total of twenty fractions were obtained in a stepwise concatenation strategy, followed by acidification to a final concentration of 0.1% formic acid and dried down with a SpeedVac. Peptides in each fraction were then analyzed with a nanoAcquity UPLC system (Waters) coupled with a TripleTOF 6600 mass spectrometer (Sciex), with similar settings as shown previously (39). Specifically, dried peptides were dissolved into 2% ACN in water with 0.1% formic acid and loaded onto a C18 Trap column (Waters Acquity UPLC Symmetry C18 NanoAcquity 10 K 2G V/M, 100 A, 5 μm, 180 μm × 20 mm) at 15 µL/min for 4 min. Peptides were then separated with an analytical column (Waters Acquity UPLC M-Class, peptide BEH C18 column, 300 A, 1.7 μm, 75 μm × 150 mm) which was temperature controlled at 40 °C. The flow rate was set as 400 nL/min. A 60-min gradient of buffer A (2% ACN, 0.1% formic acid) and buffer B (0.1% formic acid in ACN) was used for separation: 1% buffer B at 0 min, 5% buffer B at 1 min, 35% buffer B at 35 min, 99% buffer B at 37 min, 99% buffer B at 40 min, and 1% buffer B at 40.1 min, with the column equilibrated with 1% buffer B for 20 min. Data were acquired with the TripleTOF 6600 mass spectrometer using an ion spray voltage of 2.3 kV, GS15 psi, GS2 0, CUR 30 psi and an interface heater temperature of 150 °C.

For spectra library generation, the pooled sample (equally combined from 196 samples) was analyzed, and mass spectra were recorded with Analyst TF 1.7 software in DDA mode. Each cycle consisted of a full scan (m/z 400–1800) and fifty data independent acquisitions (DIAs) (m/z 100–2000) in the high sensitivity mode for precursors with a 2 + to 5 + charge state. Rolling collision energy was used. Each sample from the cohort was acquired individually via label-free SWATH DIA to quantify proteins in the samples. For SWATH acquisition, each of the samples were injected into the same NanoUPLC-MS/MS system but acquired by repeatedly cycling through 32 consecutive 25-Da precursor isolation windows, generating time-resolved fragment ion spectra for all the analytes detectable within the 400–1250 m/z precursor ion scope.

### Data Analysis for Proteomics

To generate a spectral library, the data files for DDA of the pooled sample fractions were submitted for simultaneous searches using ProteinPilot version 5.0 software (Sciex) utilizing the Paragon, ProGroup algorithms (40) and the integrated FDR analysis function (41, 42). MS/MS data was searched against the NCBI Uniprot database of Homo sapiens downloaded on May 12th, 2020, and NCBI Uniprot database of rhesus macaque downloaded on May 14th, 2020. Trypsin was selected as the digesting enzyme. Carbamidomethylation was set as a fixed modification on cysteine. Variable peptide modifications included methionine (M) oxidation. Other search parameters including instrument (TripleTOF 6600), ID Focus (Biological modifications), search effort (Thorough), FDR analysis (Yes), and user modified parameter files (No) were optimized accordingly. Proteins were inferred based on the ProGroupTM algorithm associated with the ProteinPilot software. The detected protein threshold in the software was set to the value which corresponded to 1% FDR. Peptides were defined as redundant if they had identical cleavage site(s), amino acid sequence, and modification.

For the label-free SWATH quantification, data from each sample was analyzed by PeakView 2.2 (Sciex), with the following settings: (1) Peptide filter: # of peptides per protein: 6; # of transitions per peptide: 6; peptide confidence threshold: 99%; FDR threshold: 1%; (2) XIC Options: XIC extraction window (min): 5; XIC width (ppm): 75. The peak area of each protein was used for protein level quantification. The retention time was calibrated before processing based on a selected set of reference peptides. Quantifiable peptides were carefully chosen, and the peak intensities were normalized based on total ion current (TIC) for further statistical analysis.

### Sample Preparation for Metabolomics

A volume of 25 μL of each serum sample was mixed with 75 μL of an extraction solvent mixture consisting of 35% water, 25% methanol, and 40% isopropyl alcohol containing internal standards (debrisoquine and 4-nitrobenzoic acid). Samples were vortexed and incubated on ice for 20 minutes. Next, a volume of 100μL of ACN was added to the vials. Samples were vortexed and incubated at -20 °C for 30 minutes. Finally, samples were centrifuged at 15,493 x g for 20 minutes at 4 °C and the supernatants were transferred to mass spectrometry vials for analysis using UHPLC-QTOF-MS. Pooled QC samples were created by combining 5 μL of each sample and were injected every 10 samples during batch acquisition on the LC-MS acquisition.

### Serum Metabolomics using UPLC-QTOF

For metabolomic analysis, a volume of 2 μL of each sample was injected onto a reverse-phase 50 × 2.1 mm Acquity 1.7-μm BEH C18 column at 40 °C column temperature (Waters Corp, Milford, MA) using an Acquity UPLC system (Waters) with a gradient mobile phase consisting of 100% water containing 0.1% formic acid (Solvent A), 100% ACN containing 0.1% formic acid (Solvent B), and 90/10 isopropanol/ACN containing 0.1% formic acid (Solvent D) and resolved for 13 min at a flow rate of 0.4 mL/min. The gradient started with 95% A and 5% B for 0.5 min with a ramp of curve 6. At 8 min, the gradient reached 2% A and 98% B. At 9 min, the gradient shifted to 2% B and 98% D for one minute before starting its return to the initial gradient. The metabolomic analysis of serum samples were also injected onto a BEH C18 column, but at a column temperature of 60 °C. The solvent system and run time were the same, however, the flow rate was set to 0.5 mL/min. The initial conditions for the gradient were 98% A and 2% B and were held for 0.5 min. The gradient shifted to 40% A and 60% at 4 min before reaching 2% A and 98% B at 8 min. At 9.5 min, the gradient was 2% B and 98% D and was held for 1.5 min. After 0.5 min, the composition was 50% A and 50% B. Finally, at 12 min, the gradient returned to initial conditions at 98% A and 2% B and was held for 1 min to reequilibrate the column.

The LC method for the lipidomic analysis for both the heart and the serum samples were the same. The samples were injected onto a 100 × 2.1 mm Acquity 1.7 μm CSH C18 column. The solvents consisted of 50/50 ACN/water (Solvent C) and 90/10 isopropanol/ACN (Solvent D). Both solvents contained 0.1% formic acid and 10 mM ammonium formate. The gradient flowed at 0.45 mL/min and began at 60 °C. After being held for 0.5 min, the gradient shifted to 100% D at 8 min for 0.5 min before returning to the starting gradient.

The column eluent was introduced directly into the mass spectrometer by electrospray. Mass spectrometry was performed on a Waters Xevo G2 QToF MS, operating in either negative (ESI−) or positive (ESI+) electrospray ionization modes with a capillary voltage of 3 kV for positive mode and 2.75 kV for negative mode, and a sampling cone voltage of 30 V in the positive mode and 20 V in the negative mode. The source offset for negative mode was at 80. The desolvation gas flow was 600 L/h and the temperature was set to 500 °C. Cone gas flow was 25 L/h, and the source temperature was 100 °C. Accurate mass was maintained by the introduction of LockSpray interface of Leucine Enkephalin (556.2771 [M+H]+ or 554.2615 [M^−^H]^−^) at a concentration of 2 ng/μL in 50% aqueous ACN and a rate of 20 μL/min. Data were acquired in centroid mode from 50 to 1200 m/z. Pooled QC samples were run throughout the batch to monitor data reproducibility (Supplementary Figures 7-9).

### Statistical Analysis

For metabolomics data, the Isotopologue Parameter Optimization (IPO) (43) R package was first applied to initialize and optimize peak picking parameters. Then, standard XCMS(44) peak pre-processing procedures were applied. Mass spectrometry data, received as mass over charge ratio with retention time, were normalized to internal standards. All metabolomics data were checked for missing values <20% and filtered by QC-RSD <15%. In addition, data was normalized based on the QC-RLSC (QC robust LOESS signal correction (45)). After QC-RLSC, normalized LC-MS data was log transformed and Pareto scaled. The tandem MS spectra of marker candidates were acquired by UPLC-QToF-MS/MS and validated with the NIST 2017 MS/MS standard spectra database. Metabolomic pathway analysis was completed via Mummichog (46) v2.0.6 (URL: http://mummichog.org/) to identify metabolic pathways. All statistical analyses were conducted with the use of SAS software (version 9.4; SAS Institute Inc., Cary; North Carolina, USA) and R (version 4.0.2). Potential group differences were tested using the non-parametric Kruskal-Wallis test for continuous variables and Chi-square test for categorical variables. FDR correcting for multiple testing with Benjamini-Hochberg procedure two-sided for p-values < 0.05 were regarded as statistically significant. Biological relevance and elastic net regression (21) were applied for feature selection and logistic regression model to fit the prediction model as well as calculating the probability prediction index for each sample. The performance of the biomarker prediction panel is evaluated using the AUC (Area under the ROC Curve).

## Author Contributions

Study design: V.K.S., A.K.C. Performance of the study: A.K.C., V.K.S., S.Y.W., O.O.F., A.C. Proteomic data acquisition and analysis: M.G., Y.L. Drafting of the manuscript: A.K.C., V.K.S, A.C., M.G., Y.L., J.M. Revision of manuscript content: A.K.C., V.K.S., Y.L. All authors have read and approved the final submitted manuscript.

## Acknowledgments

The opinions or assertions contained herein are the private views of the authors and are not necessarily those of the Uniformed Services University of the Health Sciences, or the Department of Defense, USA.

## Data Availability

All data generated or analyzed during this study are included in this published article (and its Supplementary Information files).

