## Supplementary file for "Identification of multi-omics molecular signature of acute radiation injury using nonhuman primate model"

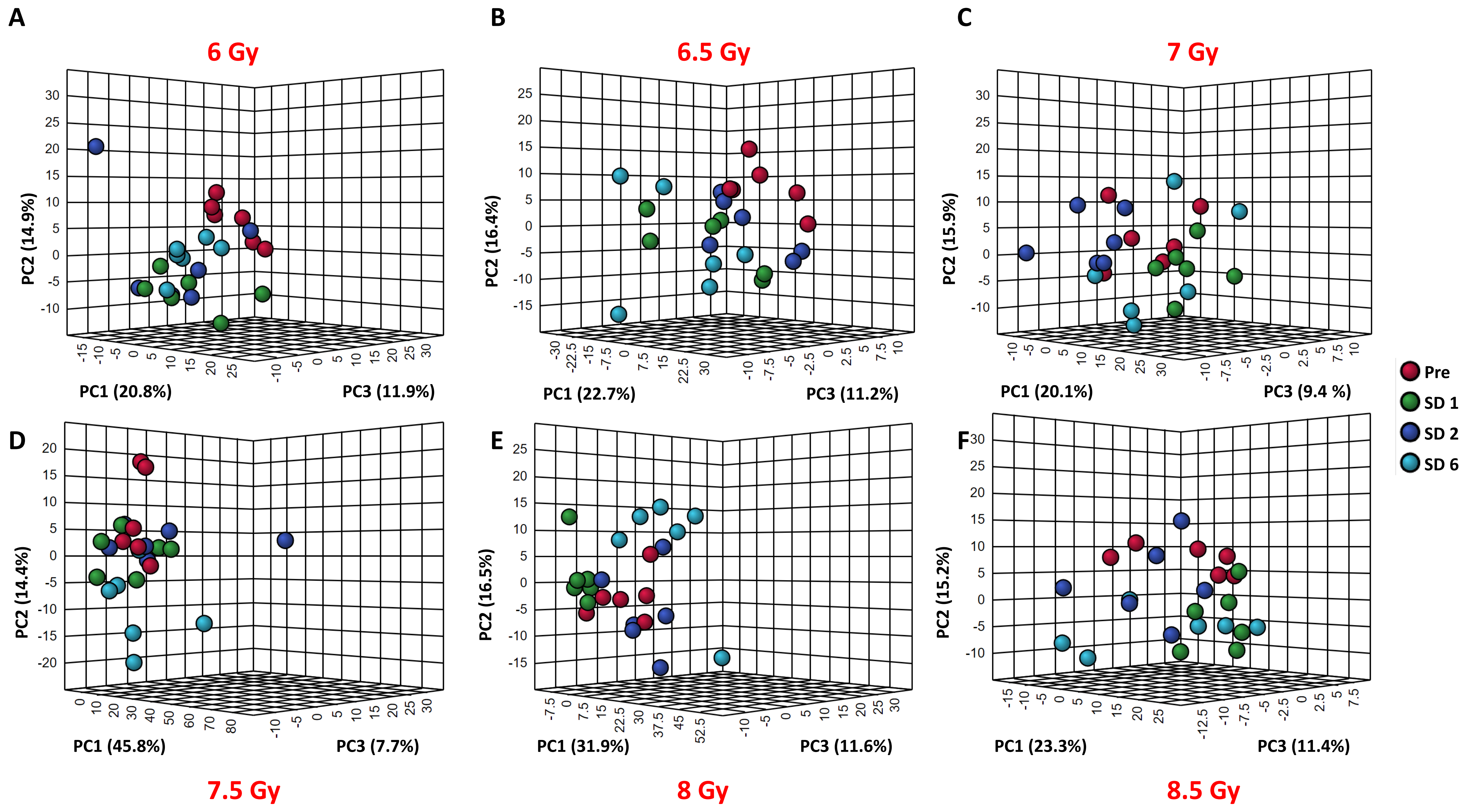
A
6 Gy
PC2 (14.9%)
PC1 (20.8%)
PC3 (11.9%)
B
6.5 Gy
PC2 (16.4%)
PC1 (22.7%)
PC3 (11.2%)
C
7 Gy
PC2 (15.9%)
PC1 (20.1%)
PC3 (9.4 %)
D
PC2 (14.4%)
PC1 (45.8%)
PC3 (7.7%)
7.5 Gy
E
PC2 (16.5%)
PC1 (31.9%)
PC3 (11.6%)
8 Gy
F
PC2 (15.2%)
PC1 (23.3%)
PC3 (11.4%)
8.5 Gy
Pre
SD 1
SD 2
SD 6

**Supp. Figure 1.** Exposure to radiation induces strong dysregulation in metabolomics and lipidomic profiles in NHPs. The first three-principal components three-dimensional score plot for NHPs at SD 1 SD 2, and SD 6 at doses 6.0 Gy, 6.5 Gy, 7.0 Gy, 7.5 Gy, 8.0 Gy, and 8.5 Gy (**Panel A-F**).

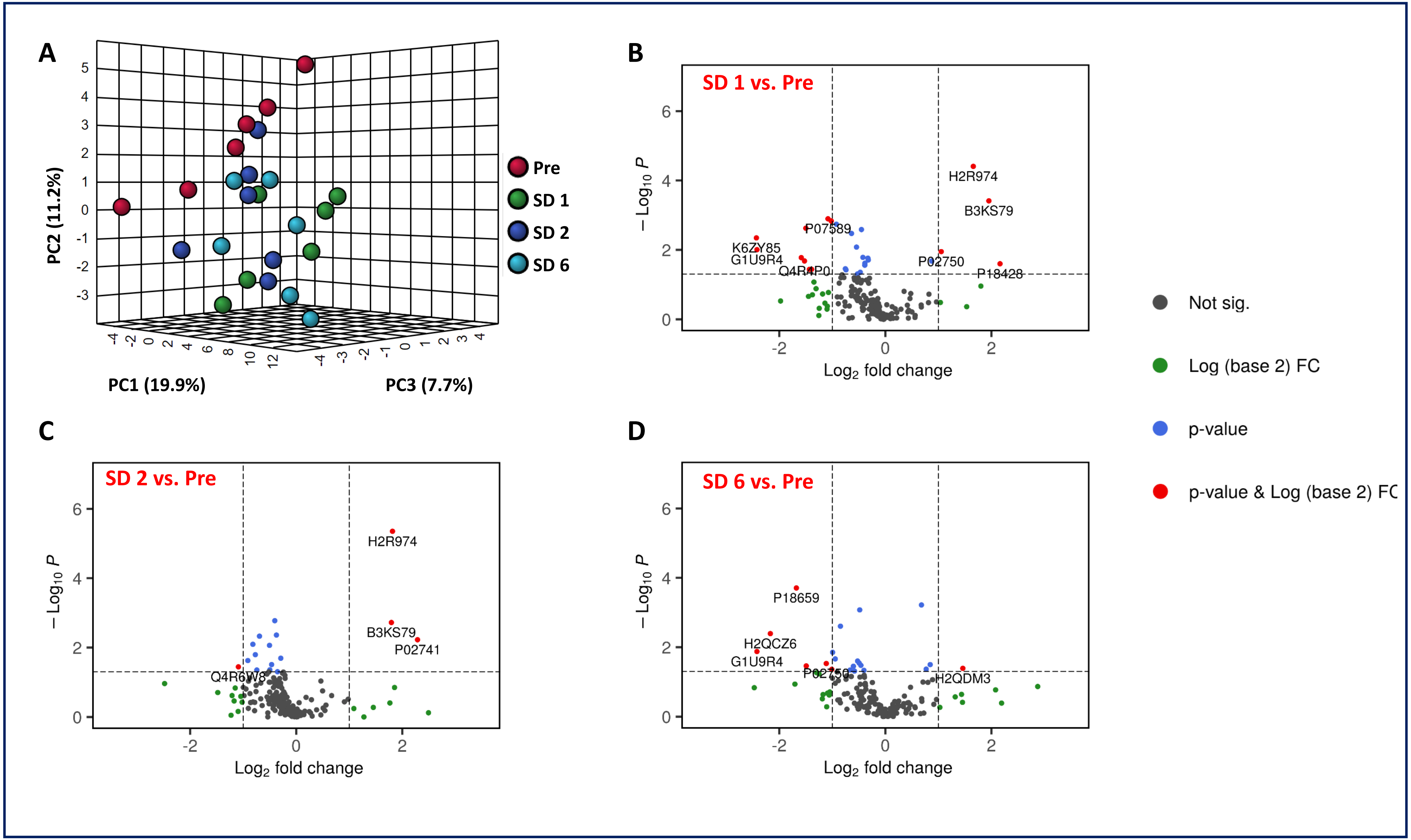
A
B
C
D
PC1 (19.9%)
PC2 (11.2%)
PC3 (7.7%)
Pre
SD 1
SD 2
SD 6
SD 1 vs. Pre
SD 2 vs. Pre
SD 6 vs. Pre
H2R974
B3KS79
P07589
K6ZY85
G1U9R4
Q4R4P0
P02750
P18428
P02741
Q4R6W8
P18659
H2QCZ6
H2QDM3
Log2 fold change
− Log10 P
Not sig.
Log (base 2) FC
p-value
p-value & Log (base 2) FC

**Supp. Figure 2**: Principal component analysis comparing overall changes of proteomic profiles of NHPs exposed **6.0** Gy, as compared to baseline at days 1 (SD 1), 2 (SD 2), and 6 (SD 6) post-irradiation (**Panel A**). Significantly dysregulated proteins upon irradiation at 8.5 Gy were visualized as volcano plots wherein dysregulated proteins that met either p-value or fold change cut off criteria were marked in green and blue, respectively, while those that met both were marked in red, and non-significant ones were denoted in gray (**Panels B, C, and D**).

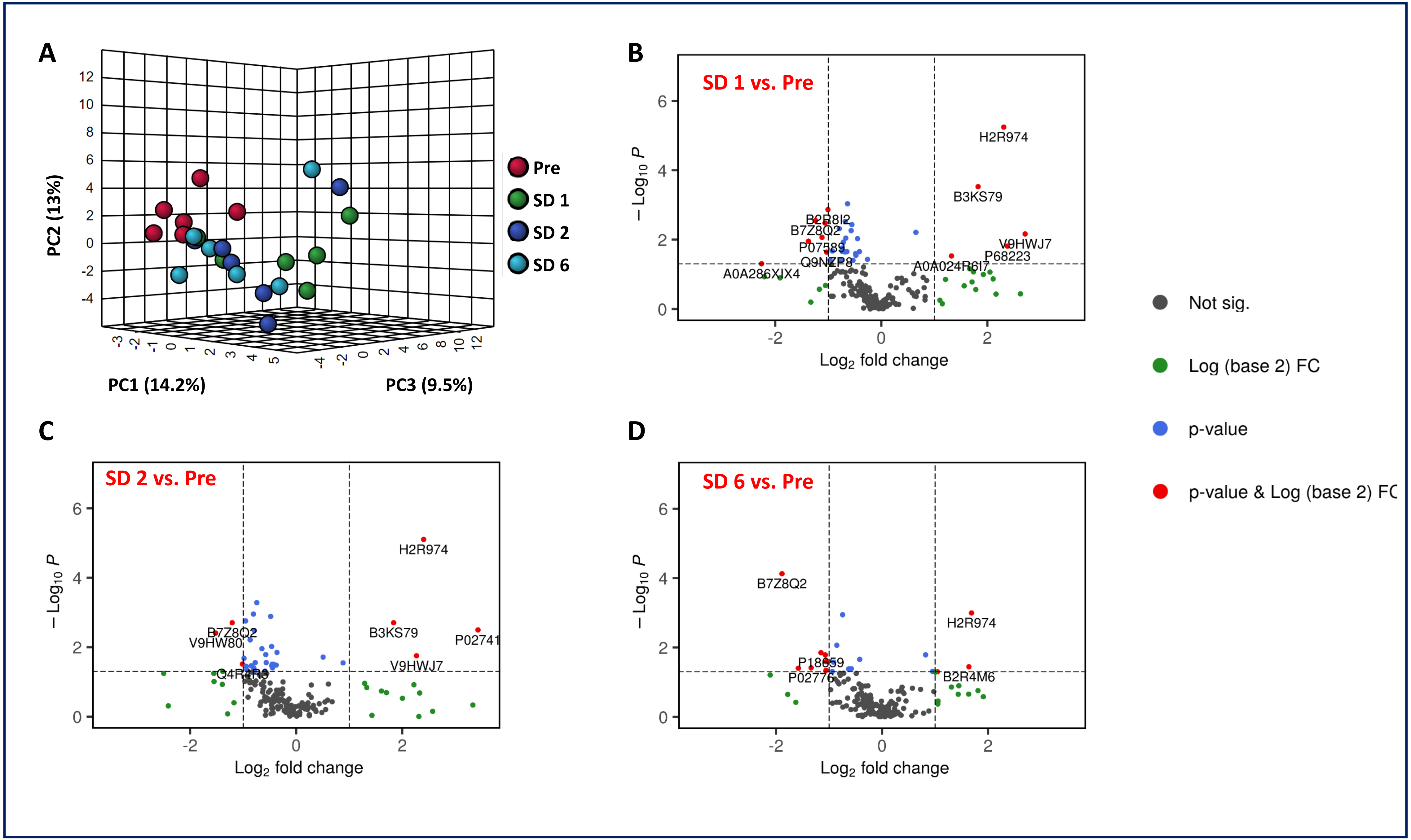

A
PC2 (13%)
PC1 (14.2%)
PC3 (9.5%)
Pre
SD 1
SD 2
SD 6
B
SD 1 vs. Pre
H2R974
B3KS79
B2R8I2
B7Z8Q2
P07589
Q9NZP8
A0A286XJX4
V9HWJ7
P68223
A0A024R6I7
− Log10 P
Log2 fold change
C
SD 2 vs. Pre
H2R974
B3KS79
P02741
V9HWJ7
B7Z8Q2
V9HW80
Q4R4R3
− Log10 P
Log2 fold change
D
SD 6 vs. Pre
B7Z8Q2
H2R974
P18659
P02776
B2R4M6
− Log10 P
Log2 fold change
Not sig.
Log (base 2) FC
p-value
p-value & Log (base 2) FC

**Supp. Figure 3**: Principal component analysis comparing overall changes proteomic profiles of NHPs exposed **6.5** Gy as compared to baseline at days 1 (SD 1), 2 (SD 2), and 6 (SD 6) post-irradiation. (**Panel A**). Significantly dysregulated proteins upon irradiation at 8.5 Gy were visualized as volcano plots wherein dysregulated proteins that met either p-value or fold change cut off criteria were marked in green and blue, respectively, while those that met both were marked in red, and non-significant ones were denoted in gray (**Panels B, C, and D**).

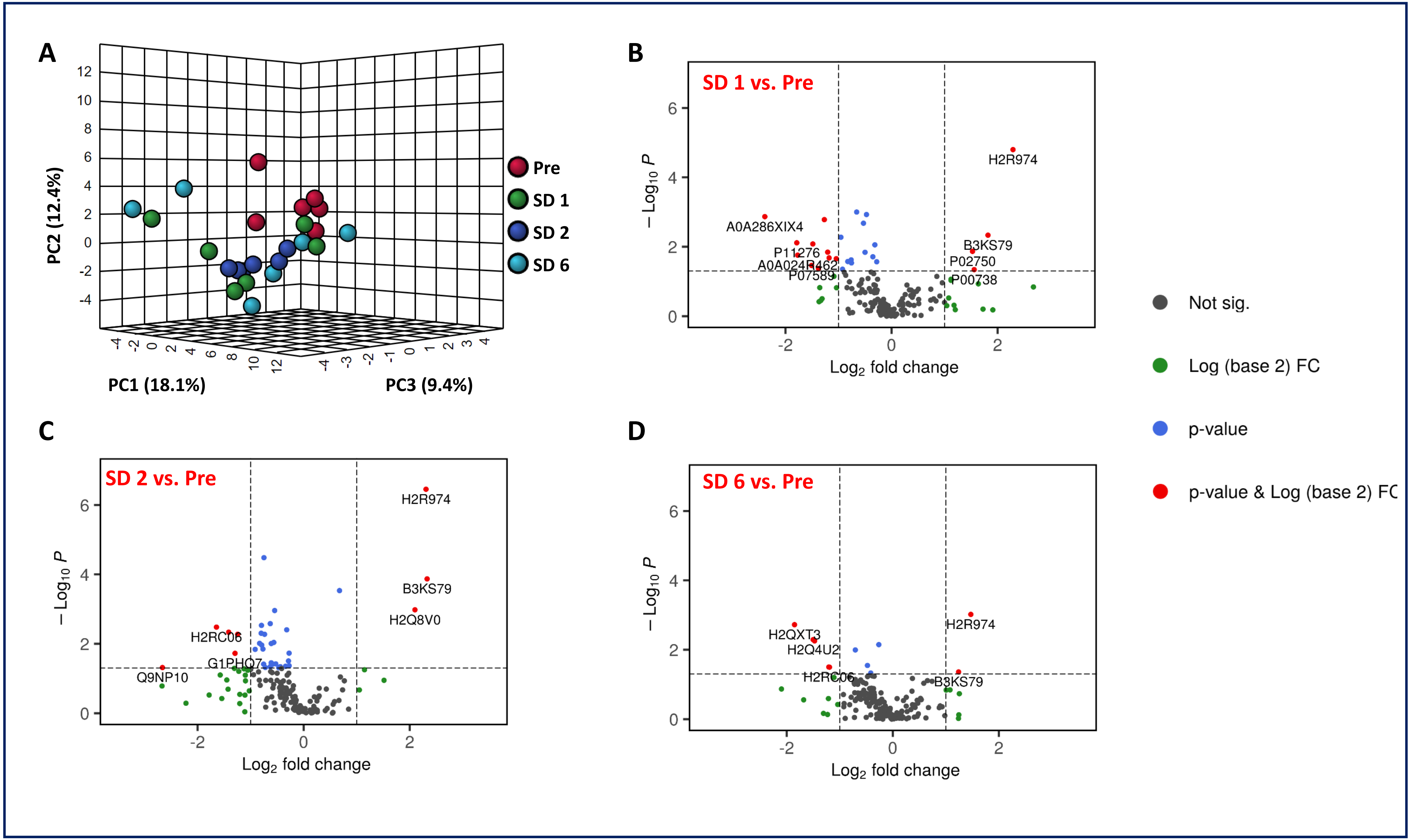

A
PC2 (12.4%)
PC1 (18.1%)
PC3 (9.4%)
Pre
SD 1
SD 2
SD 6
B
SD 1 vs. Pre
H2R974
A0A286XIX4
P11276
A0A024R462
P07589
B3KS79
P02750
P00738
− Log10 P
Log2 fold change
C
SD 2 vs. Pre
H2R974
B3KS79
H2Q8V0
H2RC06
G1PHQ7
Q9NP10
− Log10 P
Log2 fold change
D
SD 6 vs. Pre
H2QXT3
H2Q4U2
H2RC06
H2R974
B3KS79
− Log10 P
Log2 fold change
Not sig.
Log (base 2) FC
p-value
p-value & Log (base 2) FC

**Supp. Figure 4**: Principal component analysis comparing overall changes of proteomic profiles of NHPs exposed **7.0** Gy as compared to baseline at days 1 (SD 1), 2 (SD 2), and 6 (SD 6) post-irradiation. (**Panel A**). Significantly dysregulated proteins upon irradiation at 8.5 Gy were visualized as volcano plots wherein dysregulated proteins that met either p-value or fold change cut off criteria were marked in green and blue, respectively, while those that met both were marked in red, and non-significant ones were denoted in gray (**Panels B, C, and D**).

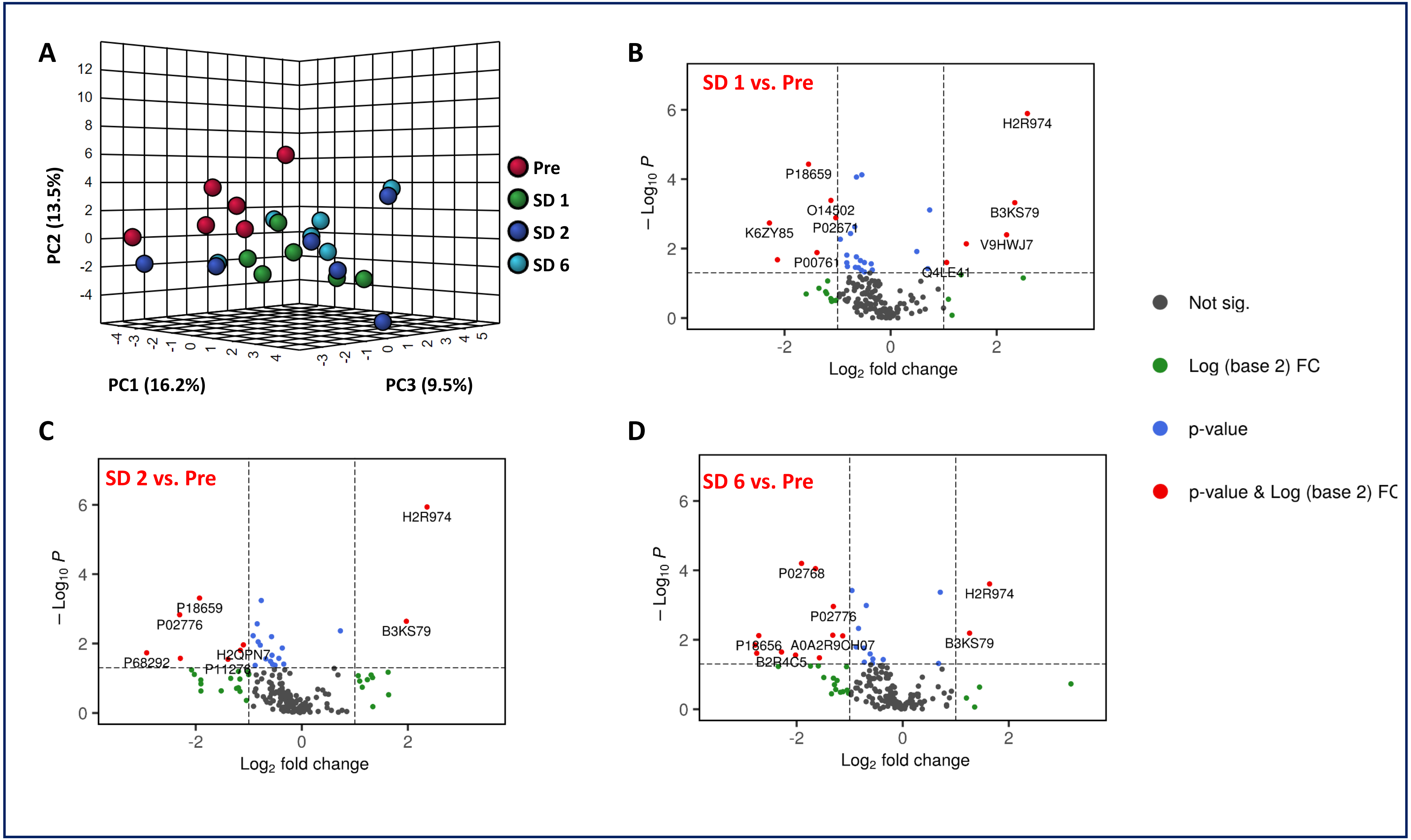
A
PC2 (13.5%)
PC1 (16.2%)
PC3 (9.5%)
Pre
SD 1
SD 2
SD 6
B
SD 1 vs. Pre
H2R974
P18659
O14502
P02671
K6ZY85
B3KS79
V9HWJ7
P00761
Q4LE41
Log2 fold change
−Log10 P
C
SD 2 vs. Pre
H2R974
P18659
P02776
B3KS79
H2QPN7
P68292
P11276
D
SD 6 vs. Pre
P02768
H2R974
P02776
P18656
A0A2R9CH07
B3KS79
B2R4C5
Not sig.
Log (base 2) FC
p-value
p-value & Log (base 2) FC

**Supp. Figure 5**: Principal component analysis comparing overall changes proteomic profiles of NHPs exposed **7.5** Gy as compared to baseline at days 1 (SD 1), 2 (SD 2) and 6 (SD 6) post-irradiation. (**Panel A**). Significantly dysregulated proteins upon irradiation at 8.5 Gy were visualized as volcano plots wherein dysregulated proteins that met either p-value or fold change cut off criteria were marked in green and blue, respectively, while those that met both were marked in red, and non-significant ones were denoted in gray (**Panels B, C, and D**).

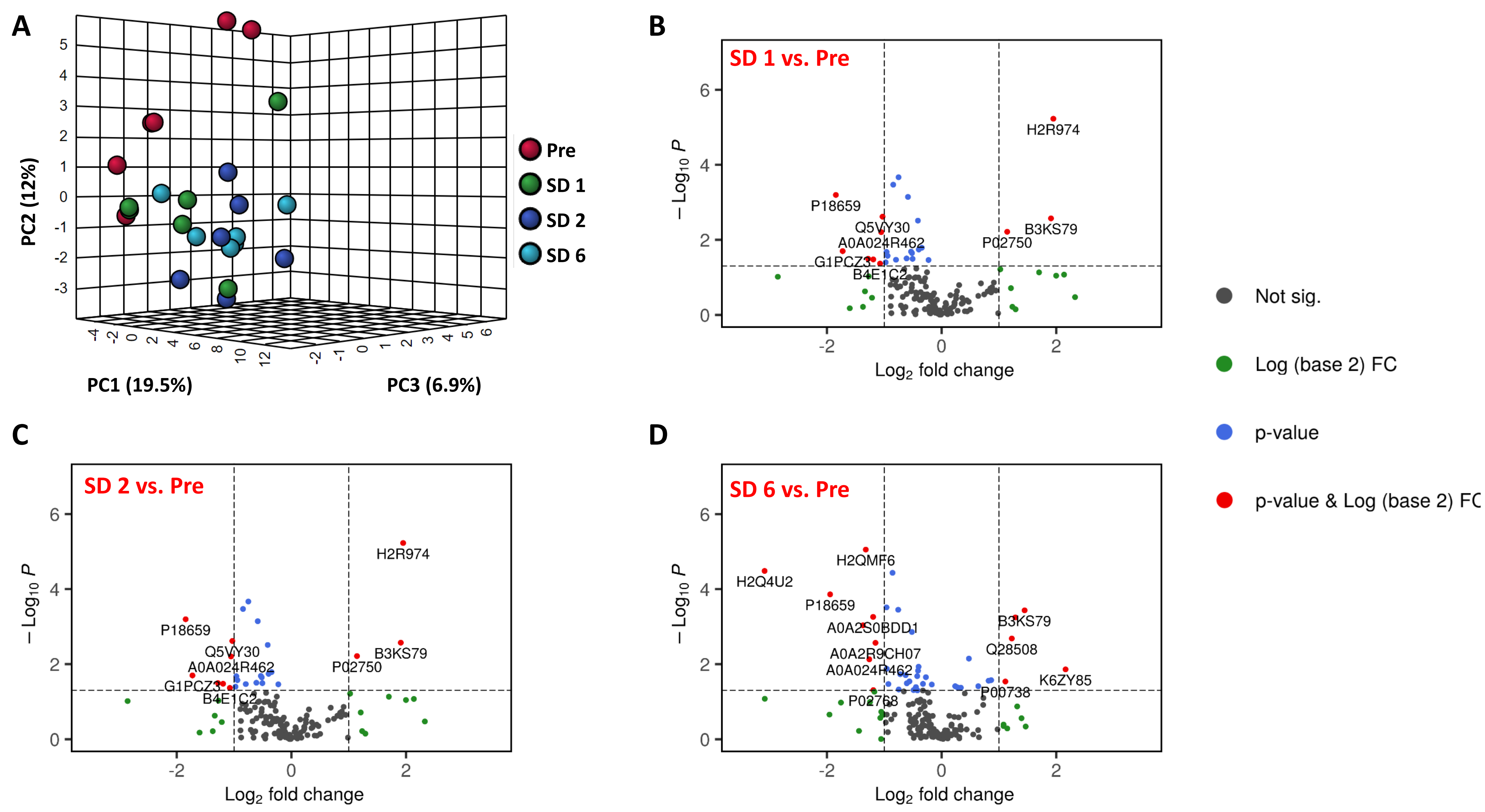

A
PC2 (12%)
PC1 (19.5%)
PC3 (6.9%)
Pre
SD 1
SD 2
SD 6
B
SD 1 vs. Pre
H2R974
P18659
Q5VY30
A0A024R462
G1PCZ3
B4E1C2
B3KS79
P02750
C
SD 2 vs. Pre
H2R974
P18659
Q5VY30
A0A024R462
G1PCZ3
B4E1C2
B3KS79
P02750
D
SD 6 vs. Pre
H2QMF6
H2Q4U2
P18659
A0A2S0BDD1
A0A2R9CH07
A0A024R462
P02768
B3KS79
Q28508
K6ZY85
P00738
− Log10 P
Log2 fold change
Not sig.
Log (base 2) FC
p-value
p-value & Log (base 2) FC

**Supp. Figure 6**: Principal component analysis comparing overall changes proteomic profiles of NHPs exposed **8.0** Gy as compared to baseline at days 1 (SD 1), 2 (SD 2), and 6 (SD 6) post-irradiation. (**Panel A**). Significantly dysregulated proteins upon irradiation at 8.5 Gy were visualized as volcano plots wherein dysregulated proteins that met either p-value or fold change cut off criteria were marked in green and blue, respectively, while those that met both were marked in red, and non-significant ones were denoted in gray (**Panels B, C, and D**).

A

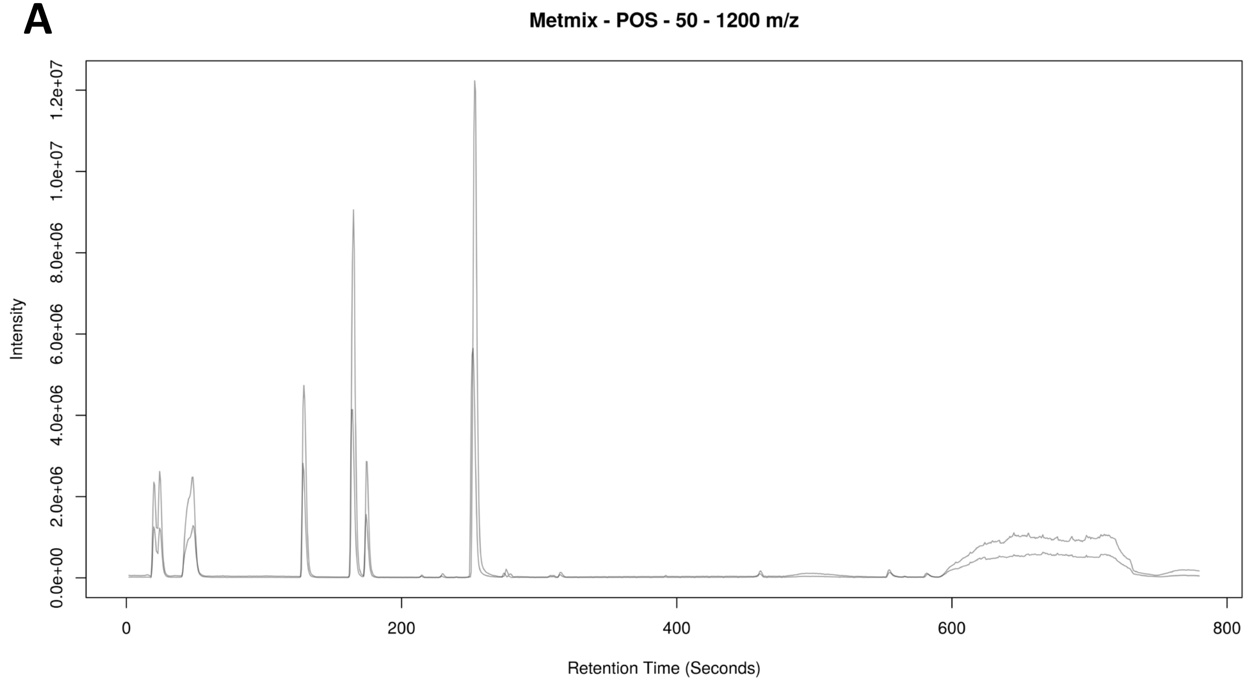
Metmix - POS - 50 - 1200 m/z
Intensity
0.0e+00
2.0e+06
4.0e+06
6.0e+06
8.0e+06
1.0e+07
1.2e+07
0
200
400
600
800
Retention Time (Seconds)

B

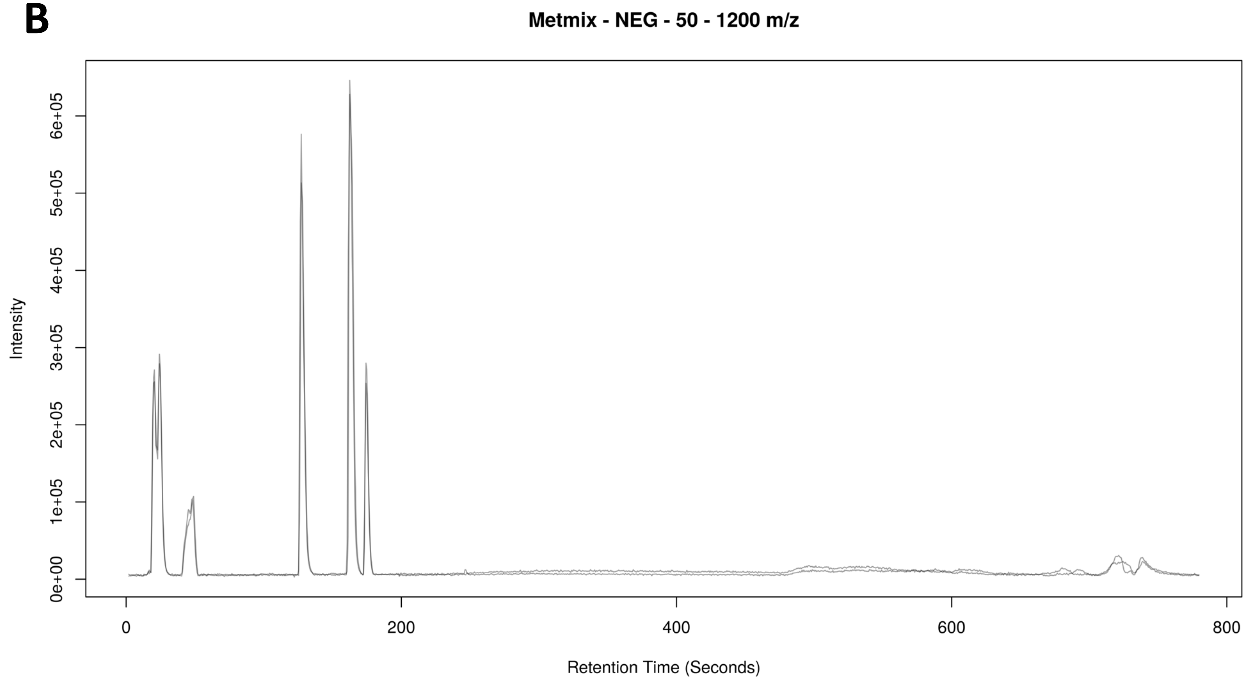
Metmix - NEG - 50 - 1200 m/z
Intensity
0e+00
1e+05
2e+05
3e+05
4e+05
5e+05
6e+05
0
200
400
600
800
Retention Time (Seconds)

**Supp. Figure 7.** Untargeted metabolomics quality control including mass calibrations checks were performed in electrospray (ESI) positive mode (Panel A) and ESI negative mode (Panel B). Metmix shows a mass error window within 5 ppm.

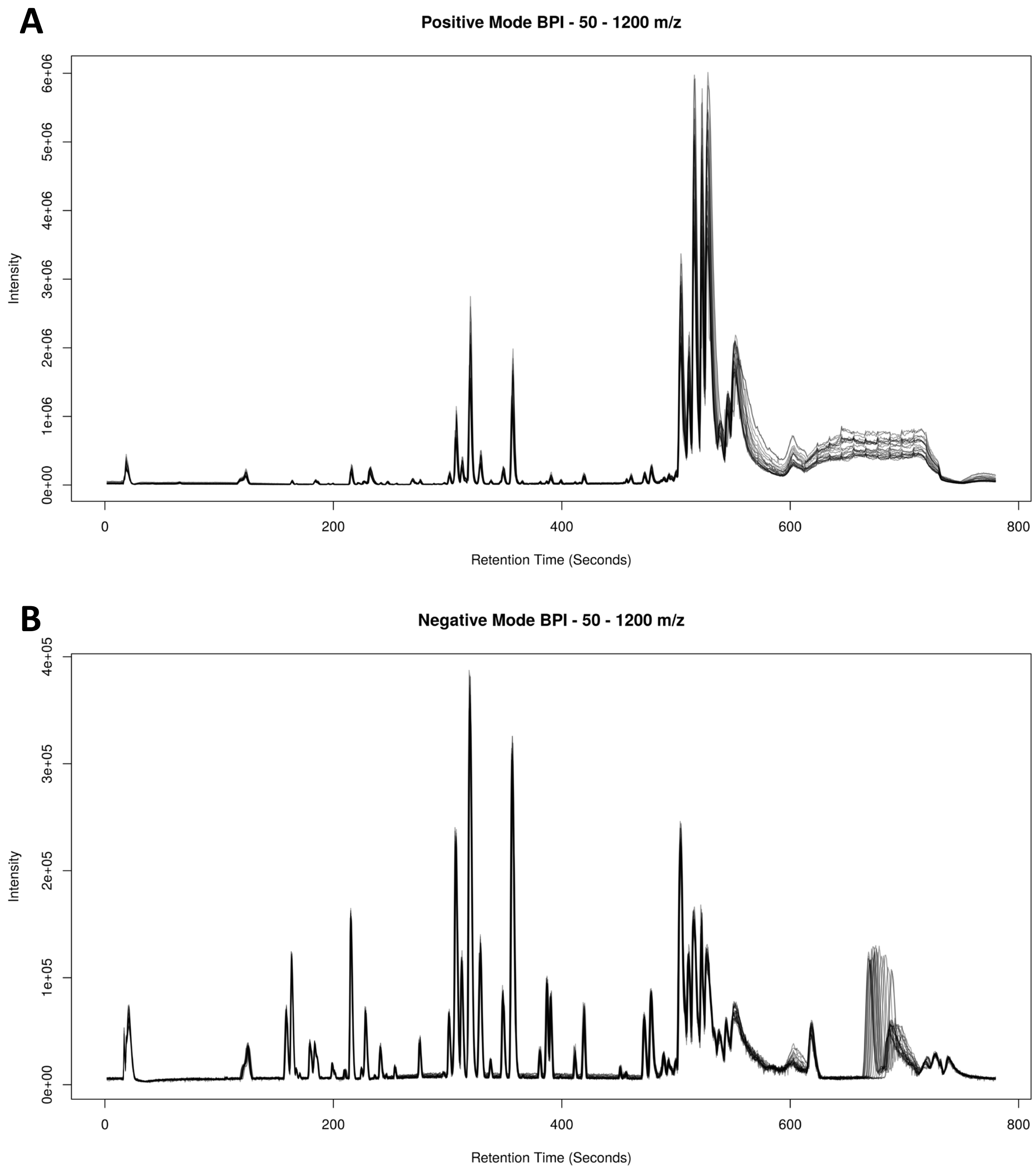
A
Positive Mode BPI - 50 - 1200 m/z
Intensity
0e+00
1e+06
2e+06
3e+06
4e+06
5e+06
6e+06
0
200
400
600
800
Retention Time (Seconds)
B
Negative Mode BPI - 50 - 1200 m/z
Intensity
0e+00
1e+05
2e+05
3e+05
4e+05
0
200
400
600
800
Retention Time (Seconds)

**Supp. Figure 8.** Positive (**Panel A**) and negative (**Panel B**) mode Metabolomics Profiling Quality Control (QC) sample overlays showing minimal shifts in retention time and intensities.

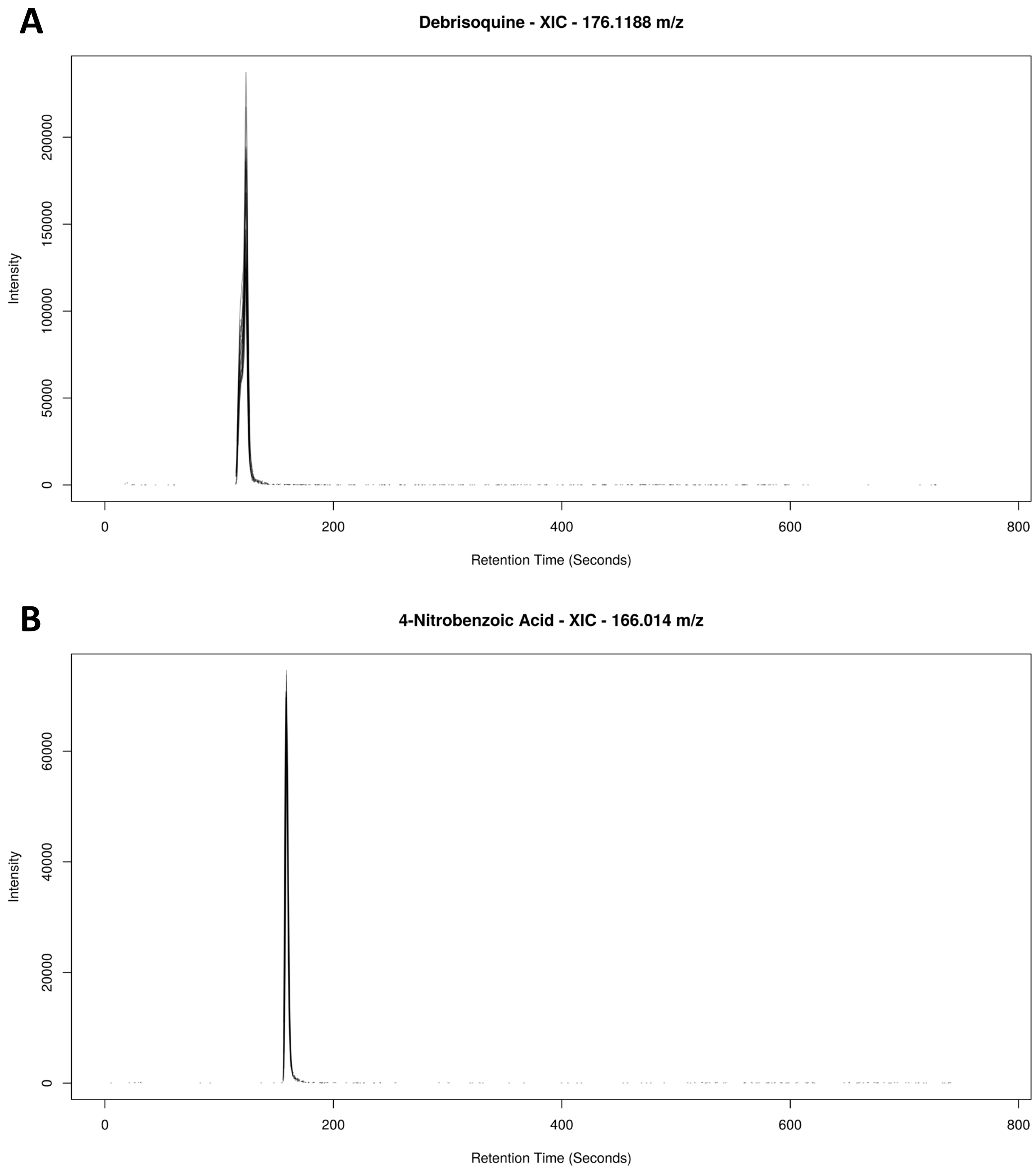
A
Debrisoquine - XIC - 176.1188 m/z
Intensity
0
50000
100000
150000
200000
0
200
400
600
800
Retention Time (Seconds)
B
4-Nitrobenzoic Acid - XIC - 166.014 m/z
Intensity
0
20000
40000
60000
0
200
400
600
800
Retention Time (Seconds)

**Supp. Figure 9.** Internal standard Debrisoquine (**Panel A**) and 4-Nitrobenzoic acid (**Panel B**) overlay showing minimal shifts in retention time and intensities.

### Supplementary Table 1: Metabolite annotations with associated test statistics across different comparative groups using global profiling

| | | | 6 Gy | | | | | | | | | | | |
|---|---|---|---|---|---|---|---|---|---|---|---|---|---|---|
| | | | SD 1 vs. Pre | | | | SD 2 vs. Pre | | | | SD 6 vs. Pre | | | |
| Mode | m/z_RT | Name | p-value | FDR | FC | LOG2FC | p-value | FDR | FC | LOG2FC | p-value | FDR | FC | LOG2FC |
| POS | 162.1127_0.57 | L-Carnitine | 6.6E-06 | 7.4E-03 | 3.11 | ↑ 1.64 | 3.7E-05 | 1.8E-03 | 2.30 | ↑ 1.20 | 1.7E-03 | 1.4E-02 | 2.13 | ↑ 1.09 |
| POS | 163.1155_0.56 | Phenylpiperazine | 1.0E-05 | 7.4E-03 | 3.17 | ↑ 1.67 | 8.6E-05 | 2.2E-03 | 2.29 | ↑ 1.20 | 3.1E-03 | 2.1E-02 | 2.28 | ↑ 1.19 |
| NEG | 861.5497_12.17 | PI(18:1) | 7.0E-05 | 4.6E-03 | 0.41 | ↓ -1.30 | 4.2E-01 | 6.1E-01 | 0.91 | ↓ -0.14 | 1.5E-02 | 1.2E-01 | 0.63 | ↓ -0.67 |
| POS | 580.432_7.28 | LysoPC(22:0) | 8.8E-05 | 1.1E-02 | 0.43 | ↓ -1.22 | 5.9E-01 | 7.3E-01 | 0.92 | ↓ -0.12 | 2.1E-01 | 4.5E-01 | 1.34 | ↑ 0.42 |
| POS | 548.3722_1.53 | Lyso-PAF C-18 | 1.4E-04 | 1.9E-02 | 0.52 | ↓ -0.94 | 2.8E-04 | 4.1E-03 | 0.64 | ↓ -0.65 | 1.7E-02 | 7.0E-02 | 0.73 | ↓ -0.46 |
| POS | 552.4031_6.62 | LysoPC(20:0) | 1.4E-04 | 1.2E-02 | 0.41 | ↓ -1.27 | 9.4E-01 | 9.6E-01 | 1.02 | ↑ 0.03 | 4.8E-01 | 7.0E-01 | 1.22 | ↑ 0.29 |
| POS | 782.5682_8.51 | PC(16:0/18:1) | 2.6E-04 | 1.3E-02 | 0.26 | ↓ -1.97 | 2.2E-01 | 4.2E-01 | 0.74 | ↓ -0.43 | 1.8E-02 | 1.4E-01 | 0.46 | ↓ -1.12 |
| NEG | 790.5343_8.7 | PI(18:0/22:6) | 3.1E-04 | 1.1E-02 | 0.57 | ↓ -0.80 | 7.2E-02 | 2.4E-01 | 0.74 | ↓ -0.44 | 2.3E-02 | 1.5E-01 | 0.64 | ↓ -0.65 |
| POS | 510.3561_5.63 | LysoPC(17:0) | 3.3E-04 | 1.4E-02 | 0.59 | ↓ -0.75 | 7.4E-02 | 2.8E-01 | 0.80 | ↓ -0.33 | 6.8E-03 | 9.2E-02 | 0.67 | ↓ -0.59 |
| POS | 339.3453_7.67 | 9-Oxo-11α,15R-dihydroxyprostan-1-oic acid | 3.4E-04 | 1.4E-02 | 0.26 | ↓ -1.94 | 1.1E-05 | 2.3E-03 | 0.16 | ↓ -2.68 | 1.5E-03 | 4.6E-02 | 0.44 | ↓ -1.20 |
| POS | 338.3423_7.68 | 13-Docosenamide, (Z)- | 3.6E-04 | 1.5E-02 | 0.28 | ↓ -1.82 | 6.3E-06 | 2.3E-03 | 0.17 | ↓ -2.56 | 1.9E-03 | 5.1E-02 | 0.46 | ↓ -1.11 |
| NEG | 554.3457_5.63 | LysoPC(17:0) | 4.2E-04 | 1.2E-02 | 0.58 | ↓ -0.79 | 2.1E-01 | 4.1E-01 | 0.87 | ↓ -0.20 | 7.0E-03 | 8.2E-02 | 0.70 | ↓ -0.52 |
| POS | 608.4631_7.89 | LysoPC(24:0) | 5.8E-04 | 2.0E-02 | 0.45 | ↓ -1.14 | 3.6E-02 | 2.2E-01 | 0.66 | ↓ -0.61 | 2.0E-01 | 4.4E-01 | 1.32 | ↑ 0.40 |
| NEG | 826.5599_8.52 | PC(16:0/20:4) | 6.0E-04 | 1.3E-02 | 0.51 | ↓ -0.97 | 5.7E-01 | 7.2E-01 | 0.94 | ↓ -0.09 | 1.8E-02 | 1.3E-01 | 0.69 | ↓ -0.54 |
| NEG | 596.3925_6.63 | LysoPC(20:0) | 6.1E-04 | 1.3E-02 | 0.44 | ↓ -1.19 | 7.1E-01 | 8.2E-01 | 0.96 | ↓ -0.06 | 5.5E-01 | 7.6E-01 | 1.20 | ↑ 0.26 |
| POS | 770.5677_8.69 | Dihomo-γ-linolenoyl PAF C-16 | 6.5E-04 | 2.1E-02 | 0.41 | ↓ -1.28 | 4.5E-02 | 2.4E-01 | 0.58 | ↓ -0.77 | 7.2E-03 | 9.3E-02 | 0.49 | ↓ -1.04 |
| POS | 532.3383_5.63 | LysoPC(17:0) | 9.0E-04 | 2.4E-02 | 0.59 | ↓ -0.77 | 7.1E-02 | 2.8E-01 | 0.78 | ↓ -0.37 | 3.7E-02 | 2.0E-01 | 0.70 | ↓ -0.51 |
| NEG | 861.5499_11.73 | PI(18:1) | 9.2E-04 | 1.7E-02 | 0.40 | ↓ -1.32 | 3.7E-01 | 5.7E-01 | 0.88 | ↓ -0.19 | 8.9E-03 | 9.3E-02 | 0.53 | ↓ -0.92 |
| POS | 184.0945_0.58 | PC | 9.6E-04 | 3.6E-02 | 3.43 | ↑ 1.78 | 3.7E-03 | 1.8E-02 | 2.86 | ↑ 1.52 | 2.3E-03 | 1.8E-02 | 2.86 | ↑ 1.51 |
| NEG | 788.5249_8.71 | PS(18:0/18:1) | 1.2E-03 | 2.0E-02 | 0.46 | ↓ -1.11 | 5.3E-02 | 2.0E-01 | 0.64 | ↓ -0.66 | 7.6E-02 | 2.8E-01 | 0.63 | ↓ -0.66 |
| POS | 518.3256_4.86 | LysoPC(16:0) | 1.6E-03 | 2.9E-02 | 0.44 | ↓ -1.19 | 1.3E-02 | 1.7E-01 | 0.53 | ↓ -0.93 | 4.4E-04 | 2.0E-02 | 0.35 | ↓ -1.52 |
| NEG | 828.5667_8.52 | C18(Plasm)-20:4 PC | 1.6E-03 | 2.4E-02 | 0.45 | ↓ -1.15 | 5.0E-01 | 6.7E-01 | 0.92 | ↓ -0.12 | 1.1E-02 | 1.0E-01 | 0.62 | ↓ -0.69 |
| NEG | 827.5636_8.52 | KRTLRR | 1.7E-03 | 2.6E-02 | 0.48 | ↓ -1.05 | 5.1E-01 | 6.8E-01 | 0.92 | ↓ -0.11 | 2.2E-02 | 1.5E-01 | 0.67 | ↓ -0.57 |
| NEG | 526.3113_5.06 | LysoPC(15:0) | 2.1E-03 | 2.9E-02 | 0.57 | ↓ -0.81 | 1.9E-01 | 3.9E-01 | 0.84 | ↓ -0.25 | 8.8E-02 | 3.0E-01 | 0.77 | ↓ -0.37 |
| NEG | 594.3771_6.08 | Pyrrolidino PAF C-16 | 2.1E-03 | 2.9E-02 | 0.55 | ↓ -0.86 | 8.7E-01 | 9.2E-01 | 1.03 | ↑ 0.05 | 8.9E-01 | 9.6E-01 | 1.03 | ↑ 0.04 |
| POS | 510.3564_1.56 | LysoPC(17:0) | 2.6E-03 | 5.8E-02 | 0.65 | ↓ -0.63 | 5.4E-01 | 6.0E-01 | 1.07 | ↑ 0.10 | 2.2E-01 | 4.0E-01 | 0.83 | ↓ -0.27 |
| POS | 482.3248_5.06 | LysoPC(15:0) | 2.6E-03 | 3.7E-02 | 0.56 | ↓ -0.83 | 1.1E-01 | 3.2E-01 | 0.76 | ↓ -0.40 | 6.2E-02 | 2.5E-01 | 0.70 | ↓ -0.51 |
| NEG | 554.3463_1.56 | LysoPC(17:0) | 3.0E-03 | 9.4E-02 | 0.64 | ↓ -0.65 | 9.0E-01 | 9.5E-01 | 1.00 | ↑ 0.00 | 1.4E-01 | 4.7E-01 | 0.79 | ↓ -0.34 |
| POS | 552.4037_2.62 | LysoPC(20:0) | 3.0E-03 | 6.1E-02 | 0.52 | ↓ -0.95 | 6.9E-03 | 2.9E-02 | 1.86 | ↑ 0.89 | 4.9E-03 | 2.9E-02 | 2.61 | ↑ 1.38 |
| NEG | 624.4238_7.29 | LysoPC(22:0) | 3.1E-03 | 3.3E-02 | 0.43 | ↓ -1.21 | 8.9E-01 | 9.3E-01 | 0.97 | ↓ -0.04 | 2.8E-01 | 5.6E-01 | 1.36 | ↑ 0.45 |
| POS | 524.3722_5.96 | LysoPC(18:0) | 3.6E-03 | 4.7E-02 | 0.75 | ↓ -0.42 | 7.0E-01 | 8.1E-01 | 1.05 | ↑ 0.07 | 5.0E-01 | 7.1E-01 | 0.95 | ↓ -0.07 |
| NEG | 512.2985_4.81 | LysoPC(14:0) | 4.1E-03 | 3.8E-02 | 0.54 | ↓ -0.89 | 1.2E-01 | 3.2E-01 | 0.74 | ↓ -0.43 | 8.0E-02 | 2.9E-01 | 2.72 | ↑ 1.44 |
| NEG | 652.4552_7.9 | LysoPC(24:0) | 4.2E-03 | 3.8E-02 | 0.54 | ↓ -0.88 | 3.9E-01 | 5.9E-01 | 0.86 | ↓ -0.21 | 3.0E-01 | 5.7E-01 | 1.26 | ↑ 0.34 |
| NEG | 226.0175_2.4 | 4-Quinolinol, 8-methyl-2-(trifluoromethyl)- | 4.2E-03 | 3.8E-02 | 0.34 | ↓ -1.58 | 2.8E-05 | 9.7E-03 | 0.18 | ↓ -2.46 | 2.0E-02 | 1.4E-01 | 0.29 | ↓ -1.76 |
| POS | 516.3104_4.97 | Oleyloxyethylphosphorylcholine | 4.8E-03 | 5.6E-02 | 0.48 | ↓ -1.06 | 1.8E-01 | 3.8E-01 | 0.71 | ↓ -0.49 | 2.7E-01 | 5.1E-01 | 0.76 | ↓ -0.40 |
| POS | 162.1129_0.33 | L-Carnitine | 4.8E-03 | 5.6E-02 | 2.53 | ↑ 1.34 | 5.6E-03 | 1.2E-01 | 2.24 | ↑ 1.16 | 1.2E-02 | 1.2E-01 | 1.78 | ↑ 0.84 |
| POS | 508.3397_5.21 | C18(Plasm) LPC | 4.9E-03 | 5.7E-02 | 0.53 | ↓ -0.90 | 2.3E-01 | 4.2E-01 | 0.78 | ↓ -0.36 | 2.4E-02 | 1.6E-01 | 0.62 | ↓ -0.70 |
| NEG | 512.299_0.99 | LysoPC(14:0) | 5.9E-03 | 1.3E-01 | 0.53 | ↓ -0.91 | 1.7E-01 | 3.6E-01 | 0.74 | ↓ -0.44 | 6.7E-02 | 3.8E-01 | 2.81 | ↑ 1.49 |
| NEG | 816.5759_9.01 | PC(16:0/20:4) | 6.5E-03 | 4.8E-02 | 0.56 | ↓ -0.83 | 1.7E-01 | 3.7E-01 | 0.76 | ↓ -0.40 | 1.1E-02 | 1.0E-01 | 0.57 | ↓ -0.82 |

| | | | | | | | | | | | | | | | | | |
|---|---|---|---|---|---|---|---|---|---|---|---|---|---|---|---|---|---|
| POS | 204.1231_0.58 | Acetyl-DL-carnitine | 8.4E-03 | 1.1E-01 | 4.49 | ↑ | 2.17 | 5.7E-02 | 1.3E-01 | 2.68 | ↑ | 1.42 | 5.0E-02 | 1.5E-01 | 3.19 | ↑ | 1.67 |
| POS | 482.3253_1.13 | LysoPE(18:0) | 9.2E-03 | 1.1E-01 | 0.54 | ↓ | -0.89 | 1.9E-01 | 2.7E-01 | 0.77 | ↓ | -0.38 | 8.2E-02 | 2.2E-01 | 0.68 | ↓ | -0.56 |
| POS | 468.3092_4.81 | LysoPC(14:0) | 1.1E-02 | 8.7E-02 | 0.56 | ↓ | -0.83 | 1.4E-01 | 3.4E-01 | 0.72 | ↓ | -0.48 | 7.1E-02 | 2.7E-01 | 2.69 | ↑ | 1.43 |
| POS | 782.5682_4.96 | PC(16:0/18:1) | 1.3E-02 | 1.3E-01 | 1.34 | ↑ | 0.43 | 6.9E-05 | 2.1E-03 | 3.57 | ↑ | 1.84 | 4.5E-04 | 6.9E-03 | 4.86 | ↑ | 2.28 |
| POS | 530.368_6.36 | C18(Plasm) LPC | 1.4E-02 | 9.5E-02 | 2.88 | ↑ | 1.53 | 6.3E-02 | 2.7E-01 | 2.78 | ↑ | 1.47 | 1.2E-01 | 3.4E-01 | 1.73 | ↑ | 0.79 |
| POS | 530.3694_1.4 | C18(Plasm) LPC | 1.5E-02 | 1.4E-01 | 2.42 | ↑ | 1.28 | 6.1E-02 | 1.3E-01 | 2.36 | ↑ | 1.24 | 2.9E-01 | 4.7E-01 | 1.33 | ↑ | 0.41 |
| NEG | 526.31_1.14 | LysoPC(15:0) | 1.6E-02 | 2.1E-01 | 0.79 | ↓ | -0.35 | 2.0E-01 | 3.9E-01 | 0.87 | ↓ | -0.21 | 1.5E-01 | 4.8E-01 | 0.82 | ↓ | -0.28 |
| NEG | 229.1437_3.65 | Dodecanedioic acid | 1.6E-02 | 8.5E-02 | 0.65 | ↓ | -0.63 | 4.0E-01 | 6.0E-01 | 0.86 | ↓ | -0.21 | 3.8E-01 | 6.4E-01 | 1.33 | ↑ | 0.41 |
| POS | 790.6208_10.03 | PS(18:0/18:1) | 1.7E-02 | 1.0E-01 | 0.79 | ↓ | -0.35 | 5.8E-01 | 7.3E-01 | 0.95 | ↓ | -0.07 | 5.9E-01 | 7.9E-01 | 0.95 | ↓ | -0.07 |
| POS | 204.1141_0.33 | Acetyl-DL-carnitine | 1.8E-02 | 1.1E-01 | 2.71 | ↑ | 1.44 | 7.2E-02 | 2.8E-01 | 2.14 | ↑ | 1.10 | 5.3E-02 | 2.4E-01 | 2.20 | ↑ | 1.14 |
| POS | 316.2485_3.86 | Decanoyl-L-carnitine | 2.2E-02 | 1.2E-01 | 6.36 | ↑ | 2.67 | 1.6E-02 | 1.8E-01 | 3.60 | ↑ | 1.85 | 4.2E-03 | 7.6E-02 | 2.57 | ↑ | 1.36 |
| POS | 468.3099_0.99 | LysoPC(14:0) | 2.2E-02 | 1.7E-01 | 0.55 | ↓ | -0.86 | 1.5E-01 | 2.4E-01 | 0.70 | ↓ | -0.52 | 1.0E-01 | 2.6E-01 | 2.68 | ↑ | 1.42 |
| NEG | 584.3554_4.45 | 2-Thio PAF | 4.0E-02 | 1.4E-01 | 0.59 | ↓ | -0.77 | 5.8E-01 | 7.2E-01 | 1.10 | ↑ | 0.14 | 3.4E-01 | 6.0E-01 | 0.77 | ↓ | -0.38 |
| NEG | 178.0488_0.62 | Hippuric acid | 4.5E-02 | 3.1E-01 | 0.41 | ↓ | -1.28 | 7.5E-03 | 1.9E-01 | 0.28 | ↓ | -1.84 | 3.8E-02 | 3.8E-01 | 0.38 | ↓ | -1.40 |
| POS | 728.5584_10.1 | PE(18:0/18:1) | 4.9E-02 | 1.8E-01 | 0.60 | ↓ | -0.73 | 5.0E-01 | 6.6E-01 | 1.18 | ↑ | 0.24 | 6.9E-02 | 2.6E-01 | 0.62 | ↓ | -0.70 |
| NEG | 452.2775_0.99 | LysoPE(16:0) | 5.1E-02 | 3.3E-01 | 0.67 | ↓ | -0.57 | 1.2E-01 | 3.1E-01 | 0.72 | ↓ | -0.47 | 9.0E-02 | 4.2E-01 | 3.03 | ↑ | 1.60 |
| POS | 257.2267_3.76 | Androst-16-en-3-ol, (3β,5α)- | 5.1E-02 | 1.9E-01 | 1.96 | ↑ | 0.97 | 3.8E-02 | 2.2E-01 | 2.10 | ↑ | 1.07 | 5.1E-02 | 2.3E-01 | 2.35 | ↑ | 1.23 |
| NEG | 862.5542_4.81 | C18(Plasm)-22:6 PC | 5.7E-02 | 3.4E-01 | 0.45 | ↓ | -1.16 | 1.6E-01 | 3.5E-01 | 1.72 | ↑ | 0.78 | 1.7E-01 | 4.8E-01 | 2.54 | ↑ | 1.35 |
| POS | 806.5689_4.84 | Arachidonoylthio-PC | 5.9E-02 | 2.7E-01 | 1.36 | ↑ | 0.44 | 9.3E-06 | 1.1E-03 | 3.49 | ↑ | 1.81 | 3.7E-04 | 6.0E-03 | 4.02 | ↑ | 2.01 |
| NEG | 263.1033_1.9 | L-Glutamine, N2-(2-phenylacetyl)- | 6.1E-02 | 1.8E-01 | 0.42 | ↓ | -1.26 | 2.7E-02 | 1.6E-01 | 0.37 | ↓ | -1.45 | 8.6E-02 | 3.0E-01 | 0.45 | ↓ | -1.15 |
| POS | 504.3088_1.14 | Lysophosphatidylethanolamine | 6.2E-02 | 2.8E-01 | 0.69 | ↓ | -0.53 | 4.0E-01 | 4.7E-01 | 0.84 | ↓ | -0.25 | 2.9E-01 | 4.6E-01 | 0.81 | ↓ | -0.31 |
| POS | 524.3717_1.89 | LysoPC(18:0) | 6.3E-02 | 2.8E-01 | 0.84 | ↓ | -0.26 | 1.7E-02 | 5.4E-02 | 1.56 | ↑ | 0.64 | 9.4E-02 | 2.4E-01 | 1.42 | ↑ | 0.50 |
| NEG | 830.5807_5.1 | PC(18:0/18:2) | 6.5E-02 | 3.5E-01 | 0.59 | ↓ | -0.77 | 6.1E-01 | 7.5E-01 | 1.08 | ↑ | 0.11 | 5.8E-01 | 7.7E-01 | 1.40 | ↑ | 0.48 |
| POS | 544.338_1.41 | LysoPC(18:1) | 7.3E-02 | 3.0E-01 | 0.78 | ↓ | -0.36 | 3.3E-02 | 8.6E-02 | 1.32 | ↑ | 0.40 | 9.5E-02 | 2.4E-01 | 1.41 | ↑ | 0.50 |
| POS | 214.1134_3.03 | Suberylglycine | 7.6E-02 | 2.4E-01 | 0.87 | ↓ | -0.20 | 4.5E-03 | 1.2E-01 | 5.90 | ↑ | 2.56 | 6.8E-10 | 9.2E-07 | 7.01 | ↑ | 2.81 |
| NEG | 768.5551_5.11 | PC(18:0/18:2) | 7.6E-02 | 3.7E-01 | 0.69 | ↓ | -0.54 | 1.2E-01 | 3.1E-01 | 1.29 | ↑ | 0.37 | 5.3E-01 | 7.5E-01 | 1.41 | ↑ | 0.50 |
| POS | 130.0499_1.9 | Glutamic Acid | 8.5E-02 | 2.5E-01 | 0.43 | ↓ | -1.22 | 2.0E-02 | 1.9E-01 | 0.33 | ↓ | -1.60 | 1.2E-01 | 3.3E-01 | 0.47 | ↓ | -1.10 |
| POS | 490.2926_0.99 | LysoPC(14:0) | 1.3E-01 | 3.7E-01 | 0.61 | ↓ | -0.72 | 9.8E-01 | 9.8E-01 | 1.05 | ↑ | 0.06 | 9.0E-02 | 2.3E-01 | 4.07 | ↑ | 2.02 |
| POS | 788.6068_5.47 | PC(18:0/18:1) | 1.6E-01 | 4.0E-01 | 0.75 | ↓ | -0.42 | 7.9E-04 | 6.8E-03 | 2.43 | ↑ | 1.28 | 1.2E-03 | 1.2E-02 | 3.05 | ↑ | 1.61 |
| POS | 287.1015_0.61 | Phenylacetyl-L-glutamine | 1.8E-01 | 4.3E-01 | 0.46 | ↓ | -1.12 | 1.5E-01 | 2.3E-01 | 0.44 | ↓ | -1.19 | 2.0E-01 | 3.7E-01 | 0.48 | ↓ | -1.07 |
| POS | 786.6003_5.47 | 1,2-Dioleoyl-sn-glycero-3-PC | 1.9E-01 | 4.5E-01 | 0.77 | ↓ | -0.37 | 7.6E-04 | 6.7E-03 | 2.52 | ↑ | 1.33 | 1.1E-03 | 1.2E-02 | 3.30 | ↑ | 1.72 |
| POS | 532.3389_1.56 | LysoPC(17:0) | 1.9E-01 | 4.5E-01 | 0.73 | ↓ | -0.45 | 2.2E-02 | 6.4E-02 | 1.58 | ↑ | 0.66 | 4.0E-01 | 5.7E-01 | 1.21 | ↑ | 0.27 |
| POS | 794.5989_5.26 | C18(Plasm)-20:4 PC | 1.9E-01 | 4.5E-01 | 0.76 | ↓ | -0.41 | 3.7E-04 | 4.6E-03 | 2.69 | ↑ | 1.43 | 8.1E-03 | 4.1E-02 | 3.06 | ↑ | 1.62 |
| POS | 421.2528_5.75 | Tris(2-butoxyethyl) phosphate | 2.0E-01 | 4.2E-01 | 1.54 | ↑ | 0.62 | 7.4E-02 | 2.8E-01 | 1.96 | ↑ | 0.97 | 6.8E-01 | 8.4E-01 | 1.05 | ↑ | 0.07 |
| POS | 180.0972_0.33 | N-Benzylglycine methyl ester | 2.2E-01 | 4.5E-01 | 0.77 | ↓ | -0.38 | 2.5E-01 | 4.5E-01 | 0.77 | ↓ | -0.38 | 2.0E-01 | 4.3E-01 | 0.90 | ↓ | -0.16 |
| NEG | 826.5604_4.96 | PC(16:0/20:4) | 2.3E-01 | 6.3E-01 | 1.48 | ↑ | 0.56 | 2.3E-02 | 2.3E-01 | 1.68 | ↑ | 0.75 | 9.8E-02 | 4.2E-01 | 2.07 | ↑ | 1.05 |
| POS | 637.3064_1.8 | Acetylpodocarpic acid anhydride | 2.4E-01 | 5.0E-01 | 1.19 | ↑ | 0.25 | 5.4E-05 | 2.1E-03 | 1.97 | ↑ | 0.98 | 5.6E-06 | 1.6E-03 | 2.16 | ↑ | 1.11 |
| POS | 808.5841_5 | 1,2-Dioleoyl-sn-glycero-3-PC | 2.8E-01 | 5.4E-01 | 0.86 | ↓ | -0.23 | 3.5E-04 | 4.4E-03 | 2.23 | ↑ | 1.16 | 7.6E-03 | 4.0E-02 | 2.59 | ↑ | 1.37 |
| NEG | 766.5393_4.96 | PE(18:0/20:4) | 2.9E-01 | 6.7E-01 | 1.43 | ↑ | 0.52 | 6.6E-02 | 2.7E-01 | 1.60 | ↑ | 0.67 | 1.4E-01 | 4.7E-01 | 2.06 | ↑ | 1.05 |
| NEG | 835.5345_5.54 | PI(16:0/18:1) | 3.1E-01 | 6.8E-01 | 1.47 | ↑ | 0.56 | 1.5E-01 | 3.3E-01 | 1.55 | ↑ | 0.63 | 2.6E-01 | 5.5E-01 | 1.95 | ↑ | 0.96 |
| NEG | 850.5605_4.85 | PC(16;0/22:6) | 3.2E-01 | 6.9E-01 | 1.64 | ↑ | 0.72 | 2.8E-02 | 2.3E-01 | 2.05 | ↑ | 1.03 | 1.6E-01 | 4.8E-01 | 2.50 | ↑ | 1.32 |
| POS | 808.5769_4.83 | 1,2-Dioleoyl-sn-glycero-3-PC | 3.2E-01 | 5.7E-01 | 0.86 | ↓ | -0.22 | 2.3E-04 | 3.7E-03 | 2.22 | ↑ | 1.15 | 7.1E-03 | 3.8E-02 | 2.37 | ↑ | 1.24 |
| POS | 502.3262_1.49 | C16(Plasm) LPC | 3.3E-01 | 5.7E-01 | 1.21 | ↑ | 0.27 | 3.5E-04 | 4.4E-03 | 2.51 | ↑ | 1.33 | 1.5E-04 | 3.6E-03 | 2.55 | ↑ | 1.35 |

| | | | | | | | | | | | | | | | | | |
|---|---|---|---|---|---|---|---|---|---|---|---|---|---|---|---|---|---|
| POS | 678.4682_6.87 | PC(14:0) | 3.5E-01 | 5.7E-01 | 0.74 | ↓ | -0.43 | 4.7E-03 | 1.2E-01 | 0.34 | ↓ | -1.57 | 9.6E-02 | 3.1E-01 | 0.58 | ↓ | -0.79 |
| POS | 546.3536_1.89 | LysoPC(18:0) | 3.6E-01 | 6.0E-01 | 0.93 | ↓ | -0.10 | 1.4E-02 | 4.8E-02 | 1.27 | ↑ | 0.34 | 8.3E-02 | 2.2E-01 | 1.17 | ↑ | 0.22 |
| NEG | 770.571_5.48 | DMPE(18:1/18:1) | 3.8E-01 | 7.3E-01 | 0.79 | ↓ | -0.34 | 3.0E-01 | 4.7E-01 | 1.19 | ↑ | 0.25 | 2.5E-01 | 5.4E-01 | 1.82 | ↑ | 0.86 |
| NEG | 832.5981_5.48 | PC(18:1/18:0) | 4.1E-01 | 7.5E-01 | 0.80 | ↓ | -0.32 | 3.7E-01 | 5.4E-01 | 1.18 | ↑ | 0.24 | 3.0E-01 | 5.8E-01 | 1.68 | ↑ | 0.75 |
| POS | 792.5895_5.18 | Dihomo-γ-linolenoyl PAF C-16 | 4.3E-01 | 6.5E-01 | 0.85 | ↓ | -0.23 | 2.3E-04 | 3.7E-03 | 3.49 | ↑ | 1.80 | 2.4E-03 | 1.8E-02 | 4.90 | ↑ | 2.29 |
| NEG | 794.5685_5.41 | PC(18:1/16:0) | 4.3E-01 | 7.7E-01 | 1.49 | ↑ | 0.58 | 2.9E-02 | 2.3E-01 | 1.73 | ↑ | 0.79 | 1.1E-01 | 4.4E-01 | 2.60 | ↑ | 1.38 |
| POS | 750.5453_5.35 | Palmitoyl Thio-PC | 4.5E-01 | 6.7E-01 | 0.90 | ↓ | -0.14 | 7.8E-04 | 6.8E-03 | 2.57 | ↑ | 1.36 | 5.1E-03 | 3.0E-02 | 3.69 | ↑ | 1.88 |
| POS | 810.5994_5.42 | PC(18:0/18:1) | 4.5E-01 | 6.7E-01 | 1.14 | ↑ | 0.18 | 7.4E-05 | 2.1E-03 | 3.13 | ↑ | 1.64 | 1.1E-03 | 1.2E-02 | 4.45 | ↑ | 2.15 |
| NEG | 832.6078_5.77 | PC(18:1/18:0) | 4.6E-01 | 7.9E-01 | 0.82 | ↓ | -0.28 | 8.8E-02 | 2.9E-01 | 1.55 | ↑ | 0.63 | 3.2E-02 | 3.5E-01 | 2.70 | ↑ | 1.43 |
| NEG | 830.5917_5.48 | PC(18:1/18:1) | 4.6E-01 | 8.0E-01 | 0.84 | ↓ | -0.26 | 2.4E-01 | 4.3E-01 | 1.23 | ↑ | 0.30 | 2.9E-01 | 5.8E-01 | 1.58 | ↑ | 0.66 |
| NEG | 747.5162_5.21 | SM(d18:1/16:0) | 5.1E-01 | 8.2E-01 | 1.71 | ↑ | 0.78 | 6.5E-02 | 2.7E-01 | 2.14 | ↑ | 1.10 | 4.1E-02 | 3.8E-01 | 3.98 | ↑ | 1.99 |
| NEG | 762.5082_4.98 | PE(16:0/22:6) | 5.2E-01 | 8.3E-01 | 1.64 | ↑ | 0.71 | 2.8E-01 | 4.6E-01 | 1.70 | ↑ | 0.76 | 3.2E-01 | 6.0E-01 | 2.73 | ↑ | 1.45 |
| POS | 790.5725_5.24 | Arachidonoyl PAF C-16 | 5.4E-01 | 7.4E-01 | 0.96 | ↓ | -0.05 | 1.9E-04 | 3.1E-03 | 4.52 | ↑ | 2.18 | 4.5E-04 | 7.0E-03 | 4.78 | ↑ | 2.26 |
| NEG | 765.5276_5.16 | SM(d18:1/18:0) | 5.5E-01 | 8.5E-01 | 0.80 | ↓ | -0.33 | 9.5E-01 | 9.7E-01 | 1.00 | ↑ | 0.00 | 6.0E-01 | 7.9E-01 | 1.71 | ↑ | 0.78 |
| NEG | 820.577_5.49 | PC(18:0/18:2) | 5.6E-01 | 8.5E-01 | 0.87 | ↓ | -0.20 | 4.2E-01 | 5.8E-01 | 1.18 | ↑ | 0.24 | 2.2E-01 | 5.3E-01 | 1.65 | ↑ | 0.72 |
| NEG | 790.5397_5.44 | PE(18:0/22:6) | 5.8E-01 | 8.5E-01 | 1.67 | ↑ | 0.74 | 2.8E-01 | 4.6E-01 | 1.83 | ↑ | 0.87 | 2.4E-01 | 5.4E-01 | 2.98 | ↑ | 1.58 |
| NEG | 806.5944_5.72 | PC(16:0/18:0) | 5.9E-01 | 8.5E-01 | 1.30 | ↑ | 0.38 | 1.5E-02 | 2.1E-01 | 2.00 | ↑ | 1.00 | 1.8E-01 | 4.9E-01 | 2.36 | ↑ | 1.24 |
| NEG | 790.541_4.84 | PE(18:0/22:6) | 6.0E-01 | 8.6E-01 | 1.40 | ↑ | 0.48 | 5.6E-02 | 2.6E-01 | 1.86 | ↑ | 0.90 | 1.5E-01 | 4.8E-01 | 2.39 | ↑ | 1.26 |
| NEG | 878.5916_5.31 | PC(18:0/22:6) | 6.5E-01 | 8.8E-01 | 1.39 | ↑ | 0.47 | 1.0E-01 | 3.1E-01 | 1.83 | ↑ | 0.87 | 1.3E-01 | 4.6E-01 | 2.71 | ↑ | 1.44 |
| NEG | 766.5395_5.55 | PE(18:0/24:0) | 6.7E-01 | 8.9E-01 | 1.25 | ↑ | 0.32 | 2.5E-01 | 4.3E-01 | 1.45 | ↑ | 0.53 | 4.3E-01 | 6.7E-01 | 2.11 | ↑ | 1.08 |
| POS | 766.5724_5.16 | Eicosapentaenoyl PAF C-16 | 6.7E-01 | 8.3E-01 | 0.98 | ↓ | -0.03 | 1.3E-04 | 2.5E-03 | 3.61 | ↑ | 1.85 | 1.5E-04 | 3.6E-03 | 5.60 | ↑ | 2.49 |
| POS | 760.5848_5.35 | PC(16:0/18:1) | 6.7E-01 | 8.3E-01 | 0.93 | ↓ | -0.10 | 4.8E-04 | 5.2E-03 | 3.24 | ↑ | 1.69 | 1.1E-04 | 3.3E-03 | 4.89 | ↑ | 2.29 |
| NEG | 776.5592_5.71 | PC(18:1/14:0) | 7.1E-01 | 9.1E-01 | 1.55 | ↑ | 0.63 | 7.9E-02 | 2.8E-01 | 2.17 | ↑ | 1.12 | 9.5E-02 | 4.2E-01 | 3.13 | ↑ | 1.65 |
| NEG | 768.5463_5.54 | PC(18:0/14:0) | 7.5E-01 | 9.3E-01 | 0.86 | ↓ | -0.21 | 8.0E-01 | 8.9E-01 | 0.96 | ↓ | -0.06 | 6.3E-01 | 8.0E-01 | 1.64 | ↑ | 0.71 |
| POS | 782.567_5.35 | PC(16:0/18:1) | 7.7E-01 | 8.9E-01 | 0.96 | ↓ | -0.06 | 4.1E-04 | 4.8E-03 | 2.41 | ↑ | 1.27 | 1.6E-04 | 3.8E-03 | 2.57 | ↑ | 1.36 |
| NEG | 838.5972_5.6 | C18(Plasm)-20:4 PC | 7.7E-01 | 9.4E-01 | 1.32 | ↑ | 0.40 | 9.6E-02 | 3.0E-01 | 1.85 | ↑ | 0.88 | 6.8E-02 | 3.8E-01 | 3.29 | ↑ | 1.72 |
| NEG | 804.5669_5.01 | PC(18:1/16:0) | 7.9E-01 | 9.5E-01 | 1.05 | ↑ | 0.07 | 5.8E-02 | 2.6E-01 | 1.42 | ↑ | 0.50 | 2.0E-01 | 5.2E-01 | 1.60 | ↑ | 0.68 |
| NEG | 802.5603_5.01 | PC(16:0/18:2) | 8.0E-01 | 9.5E-01 | 1.05 | ↑ | 0.07 | 4.6E-02 | 2.5E-01 | 1.38 | ↑ | 0.46 | 1.7E-01 | 4.8E-01 | 1.53 | ↑ | 0.61 |
| NEG | 744.549_5.01 | α-Galactosyl-N-palmitoylsphingosine | 8.4E-01 | 9.6E-01 | 1.11 | ↑ | 0.15 | 4.7E-02 | 2.5E-01 | 1.48 | ↑ | 0.56 | 3.5E-01 | 6.2E-01 | 1.41 | ↑ | 0.49 |
| POS | 768.5893_5.23 | C16-18:1 PC | 8.5E-01 | 9.2E-01 | 1.05 | ↑ | 0.08 | 1.0E-04 | 2.3E-03 | 4.20 | ↑ | 2.07 | 1.1E-03 | 1.2E-02 | 5.41 | ↑ | 2.44 |
| NEG | 774.545_5.65 | C18(Plasm)-22:6 PE | 8.5E-01 | 9.6E-01 | 1.40 | ↑ | 0.49 | 5.5E-02 | 2.5E-01 | 2.38 | ↑ | 1.25 | 1.0E-01 | 4.3E-01 | 3.53 | ↑ | 1.82 |
| NEG | 714.5081_5.16 | PE(16:0/18:2) | 8.6E-01 | 9.7E-01 | 0.86 | ↓ | -0.22 | 5.8E-01 | 7.2E-01 | 1.11 | ↑ | 0.14 | 4.9E-01 | 7.2E-01 | 1.66 | ↑ | 0.73 |
| NEG | 792.5473_5.01 | PC(16:0/18:2) | 8.7E-01 | 9.7E-01 | 1.02 | ↑ | 0.02 | 7.1E-02 | 2.7E-01 | 1.41 | ↑ | 0.50 | 2.3E-01 | 5.4E-01 | 1.60 | ↑ | 0.68 |
| POS | 735.5737_5.29 | Thioetheramidephosphatidylcholine | 8.7E-01 | 9.3E-01 | 1.05 | ↑ | 0.07 | 1.7E-04 | 2.9E-03 | 3.77 | ↑ | 1.91 | 9.1E-04 | 1.1E-02 | 5.52 | ↑ | 2.46 |
| POS | 756.5531_5.29 | PC(16:0/16:0) | 9.0E-01 | 9.5E-01 | 1.18 | ↑ | 0.24 | 1.3E-04 | 2.5E-03 | 3.40 | ↑ | 1.77 | 3.0E-04 | 5.6E-03 | 3.36 | ↑ | 1.75 |
| POS | 184.0734_5 | PC | 9.1E-01 | 9.5E-01 | 1.02 | ↑ | 0.03 | 8.4E-05 | 2.2E-03 | 3.21 | ↑ | 1.68 | 6.3E-04 | 8.6E-03 | 3.67 | ↑ | 1.88 |
| NEG | 806.5831_5.35 | PC(16:0/18:0) | 9.1E-01 | 9.9E-01 | 1.07 | ↑ | 0.09 | 1.9E-02 | 2.2E-01 | 1.80 | ↑ | 0.85 | 2.4E-02 | 3.2E-01 | 2.53 | ↑ | 1.34 |
| POS | 834.5998_5.31 | 1-Docosahexaenoyl-2-stearoyl-sn-glycero-3-p | 9.2E-01 | 9.6E-01 | 1.06 | ↑ | 0.08 | 2.7E-05 | 1.7E-03 | 3.78 | ↑ | 1.92 | 1.7E-03 | 1.4E-02 | 6.44 | ↑ | 2.69 |
| NEG | 834.5429_5.53 | PS(18:0/22:6) | 9.3E-01 | 9.9E-01 | 1.03 | ↑ | 0.05 | 5.6E-01 | 7.1E-01 | 1.10 | ↑ | 0.14 | 6.0E-01 | 7.9E-01 | 1.45 | ↑ | 0.53 |
| NEG | 742.5398_5.01 | PE(18:0/18:2) | 9.4E-01 | 9.9E-01 | 0.98 | ↓ | -0.04 | 3.4E-02 | 2.3E-01 | 1.52 | ↑ | 0.60 | 2.3E-01 | 5.3E-01 | 1.58 | ↑ | 0.66 |
| NEG | 862.5958_5.51 | C18(Plasm)-22:6 PC | 9.5E-01 | 9.9E-01 | 1.22 | ↑ | 0.28 | 2.6E-01 | 4.4E-01 | 1.53 | ↑ | 0.62 | 1.7E-01 | 4.8E-01 | 2.27 | ↑ | 1.18 |
| POS | 758.5692_5.01 | PC(16:0/18:2) | 9.6E-01 | 9.8E-01 | 1.03 | ↑ | 0.04 | 1.7E-04 | 2.9E-03 | 2.89 | ↑ | 1.53 | 3.1E-04 | 5.6E-03 | 3.32 | ↑ | 1.73 |
| POS | 760.5746_5.01 | PC(16:0/18:1) | 9.6E-01 | 9.8E-01 | 1.04 | ↑ | 0.06 | 1.3E-04 | 2.5E-03 | 2.74 | ↑ | 1.45 | 3.1E-04 | 5.6E-03 | 3.03 | ↑ | 1.60 |

| | | | | | | | | | | | | | | | | | |
|---|---|---|---|---|---|---|---|---|---|---|---|---|---|---|---|---|---|
| NEG | 804.576_5.36 | PC(18:1/16:0) | 9.6E-01 | 9.9E-01 | 1.06 | ↑ | 0.08 | 2.4E-02 | 2.3E-01 | 1.66 | ↑ | 0.73 | 3.1E-02 | 3.4E-01 | 2.19 | ↑ | 1.13 |
| POS | 768.5582_5.55 | PE(18:0/18:1) | 9.8E-01 | 9.9E-01 | 1.16 | ↑ | 0.21 | 3.5E-04 | 4.4E-03 | 3.35 | ↑ | 1.74 | 1.2E-02 | 5.3E-02 | 5.53 | ↑ | 2.47 |
| NEG | 738.5085_5.1 | PE(16:0/20:4) | 9.9E-01 | 9.9E-01 | 1.03 | ↑ | 0.04 | 5.0E-01 | 6.5E-01 | 1.20 | ↑ | 0.26 | 3.6E-01 | 6.2E-01 | 2.01 | ↑ | 1.01 |
| NEG | 802.5603_8.81 | PC(16:0/18:2) | 1.0E+00 | 1.0E+00 | 1.00 | ↑ | 0.00 | 1.7E-02 | 1.3E-01 | 1.20 | ↑ | 0.26 | 3.0E-02 | 1.7E-01 | 1.09 | ↑ | 0.13 |

| | | | 6.5 Gy | | | | | | | | | | | | | | |
|---|---|---|---|---|---|---|---|---|---|---|---|---|---|---|---|---|---|
| | | | SD 1 vs. Pre | | | | | SD 2 vs. Pre | | | | | SD 6 vs. Pre | | | | |
| Mode | m/z_RT | Name | p-value | FDR | FC | | LOG2FC | p-value | FDR | FC | | LOG2FC | p-value | FDR | FC | | LOG2FC |
| POS | 162.1127_0.57 | L-Carnitine | 4.3E-04 | 3.3E-02 | 2.84 | ↑ | 1.51 | 4.5E-02 | 1.0E-01 | 1.59 | ↑ | 0.67 | 2.2E-03 | 2.8E-02 | 2.02 | ↑ | 1.01 |
| POS | 163.1155_0.56 | Phenylpiperazine | 6.5E-04 | 3.5E-02 | 2.72 | ↑ | 1.45 | 5.5E-02 | 1.2E-01 | 1.55 | ↑ | 0.63 | 2.1E-03 | 2.8E-02 | 2.06 | ↑ | 1.05 |
| NEG | 861.5497_12.17 | PI(18:1) | 3.8E-03 | 4.0E-02 | 0.50 | ↓ | -0.99 | 3.9E-01 | 6.8E-01 | 1.12 | ↑ | 0.16 | 4.1E-03 | 6.4E-02 | 0.59 | ↓ | -0.76 |
| POS | 580.432_7.28 | LysoPC(22:0) | 2.2E-04 | 1.2E-02 | 0.53 | ↓ | -0.90 | 8.7E-01 | 9.4E-01 | 1.12 | ↑ | 0.16 | 1.9E-03 | 3.1E-02 | 0.56 | ↓ | -0.85 |
| POS | 548.3722_1.53 | Lyso-PAF C-18 | 4.5E-03 | 8.6E-02 | 0.45 | ↓ | -1.14 | 8.2E-03 | 2.9E-02 | 0.55 | ↓ | -0.87 | 4.9E-03 | 3.9E-02 | 0.49 | ↓ | -1.02 |
| POS | 552.4031_6.62 | LysoPC(20:0) | 5.3E-05 | 7.7E-03 | 0.49 | ↓ | -1.03 | 9.5E-01 | 9.7E-01 | 1.07 | ↑ | 0.09 | 8.7E-04 | 2.1E-02 | 0.53 | ↓ | -0.93 |
| POS | 782.5682_8.51 | PC(16:0/18:1) | 4.0E-03 | 5.0E-02 | 0.41 | ↓ | -1.29 | 5.1E-01 | 7.1E-01 | 0.86 | ↓ | -0.21 | 2.4E-02 | 1.2E-01 | 0.50 | ↓ | -1.00 |
| NEG | 790.5343_8.7 | PI(18:0/22:6) | 7.0E-03 | 5.3E-02 | 0.59 | ↓ | -0.77 | 2.6E-01 | 5.7E-01 | 0.81 | ↓ | -0.31 | 1.5E-02 | 1.1E-01 | 0.60 | ↓ | -0.75 |
| POS | 510.3561_5.63 | LysoPC(17:0) | 3.3E-03 | 4.6E-02 | 0.57 | ↓ | -0.81 | 6.8E-02 | 2.8E-01 | 0.75 | ↓ | -0.42 | 4.8E-03 | 4.7E-02 | 0.60 | ↓ | -0.74 |
| POS | 339.3453_7.67 | 9-Oxo-11α,15R-dihydroxyprostan-1-oic acid | 3.1E-05 | 7.7E-03 | 0.17 | ↓ | -2.55 | 3.5E-04 | 1.7E-02 | 0.22 | ↓ | -2.15 | 4.1E-04 | 1.5E-02 | 0.26 | ↓ | -1.95 |
| POS | 338.3423_7.68 | 13-Docosenamide, (Z)- | 2.8E-05 | 7.7E-03 | 0.19 | ↓ | -2.42 | 2.9E-04 | 1.7E-02 | 0.25 | ↓ | -2.00 | 3.2E-04 | 1.3E-02 | 0.28 | ↓ | -1.83 |
| NEG | 554.3457_5.63 | LysoPC(17:0) | 1.1E-02 | 6.8E-02 | 0.61 | ↓ | -0.72 | 3.3E-01 | 6.4E-01 | 0.85 | ↓ | -0.24 | 2.9E-02 | 1.6E-01 | 0.67 | ↓ | -0.57 |
| POS | 608.4631_7.89 | LysoPC(24:0) | 3.3E-02 | 1.7E-01 | 0.70 | ↓ | -0.52 | 6.9E-01 | 8.4E-01 | 0.97 | ↓ | -0.04 | 4.6E-02 | 1.7E-01 | 0.67 | ↓ | -0.57 |
| NEG | 826.5599_8.52 | PC(16:0/20:4) | 3.3E-03 | 3.6E-02 | 0.67 | ↓ | -0.59 | 8.4E-01 | 9.4E-01 | 0.98 | ↓ | -0.03 | 9.1E-02 | 3.0E-01 | 0.78 | ↓ | -0.37 |
| NEG | 596.3925_6.63 | LysoPC(20:0) | 7.1E-06 | 5.5E-03 | 0.46 | ↓ | -1.12 | 6.6E-01 | 8.6E-01 | 0.97 | ↓ | -0.04 | 4.2E-04 | 2.6E-02 | 0.49 | ↓ | -1.03 |
| POS | 770.5677_8.69 | Dihomo-γ-linolenoyl PAF C-16 | 3.7E-03 | 4.7E-02 | 0.57 | ↓ | -0.81 | 1.1E-01 | 3.4E-01 | 0.80 | ↓ | -0.32 | 6.2E-03 | 5.4E-02 | 0.50 | ↓ | -0.99 |
| POS | 532.3383_5.63 | LysoPC(17:0) | 1.7E-02 | 1.2E-01 | 0.59 | ↓ | -0.76 | 1.8E-01 | 4.2E-01 | 0.76 | ↓ | -0.40 | 2.6E-02 | 1.2E-01 | 0.61 | ↓ | -0.71 |
| NEG | 861.5499_11.73 | PI(18:1) | 2.0E-03 | 3.2E-02 | 0.43 | ↓ | -1.22 | 4.2E-01 | 7.1E-01 | 1.11 | ↑ | 0.16 | 1.3E-03 | 4.4E-02 | 0.49 | ↓ | -1.02 |
| POS | 184.0945_0.58 | PC | 1.7E-04 | 2.4E-02 | 3.35 | ↑ | 1.75 | 1.2E-03 | 9.0E-03 | 2.36 | ↑ | 1.24 | 1.9E-04 | 1.3E-02 | 2.83 | ↑ | 1.50 |
| NEG | 788.5249_8.71 | PS(18:0/18:1) | 1.9E-03 | 3.0E-02 | 0.48 | ↓ | -1.07 | 5.8E-01 | 8.1E-01 | 0.87 | ↓ | -0.19 | 6.3E-02 | 2.4E-01 | 0.64 | ↓ | -0.65 |
| POS | 518.3256_4.86 | LysoPC(16:0) | 2.7E-02 | 1.6E-01 | 0.59 | ↓ | -0.76 | 2.0E-02 | 1.6E-01 | 0.60 | ↓ | -0.73 | 8.9E-05 | 7.5E-03 | 0.37 | ↓ | -1.42 |
| NEG | 828.5667_8.52 | C18(Plasm)-20:4 PC | 6.5E-03 | 5.1E-02 | 0.63 | ↓ | -0.66 | 8.2E-01 | 9.3E-01 | 0.97 | ↓ | -0.05 | 8.1E-02 | 2.8E-01 | 0.71 | ↓ | -0.49 |
| NEG | 827.5636_8.52 | KRTLRR | 2.8E-03 | 3.4E-02 | 0.63 | ↓ | -0.67 | 7.4E-01 | 9.0E-01 | 0.97 | ↓ | -0.05 | 5.4E-02 | 2.3E-01 | 0.72 | ↓ | -0.48 |
| NEG | 526.3113_5.06 | LysoPC(15:0) | 6.6E-02 | 2.0E-01 | 0.70 | ↓ | -0.51 | 9.8E-01 | 9.9E-01 | 0.97 | ↓ | -0.04 | 2.4E-02 | 1.4E-01 | 0.65 | ↓ | -0.63 |
| NEG | 594.3771_6.08 | Pyrrolidino PAF C-16 | 9.6E-04 | 2.3E-02 | 0.64 | ↓ | -0.65 | 7.4E-01 | 9.0E-01 | 1.07 | ↑ | 0.09 | 1.7E-03 | 4.9E-02 | 0.64 | ↓ | -0.65 |
| POS | 510.3564_1.56 | LysoPC(17:0) | 4.9E-04 | 3.3E-02 | 0.44 | ↓ | -1.18 | 5.1E-02 | 1.1E-01 | 0.68 | ↓ | -0.56 | 1.0E-02 | 5.7E-02 | 0.57 | ↓ | -0.80 |
| POS | 482.3248_5.06 | LysoPC(15:0) | 1.3E-02 | 1.0E-01 | 0.61 | ↓ | -0.72 | 6.5E-02 | 2.8E-01 | 0.76 | ↓ | -0.39 | 1.6E-03 | 3.0E-02 | 0.53 | ↓ | -0.91 |
| NEG | 554.3463_1.56 | LysoPC(17:0) | 4.2E-04 | 1.3E-02 | 0.48 | ↓ | -1.06 | 5.7E-01 | 6.6E-01 | 0.90 | ↓ | -0.15 | 1.7E-02 | 1.2E-01 | 0.62 | ↓ | -0.68 |
| POS | 552.4037_2.62 | LysoPC(20:0) | 5.4E-03 | 9.6E-02 | 0.46 | ↓ | -1.13 | 3.1E-01 | 4.3E-01 | 1.24 | ↑ | 0.31 | 2.4E-01 | 4.6E-01 | 0.83 | ↓ | -0.27 |
| NEG | 624.4238_7.29 | LysoPC(22:0) | 2.0E-04 | 1.5E-02 | 0.51 | ↓ | -0.98 | 8.6E-01 | 9.5E-01 | 1.17 | ↑ | 0.23 | 5.7E-03 | 7.5E-02 | 0.62 | ↓ | -0.68 |
| POS | 524.3722_5.96 | LysoPC(18:0) | 9.5E-05 | 8.3E-03 | 0.64 | ↓ | -0.64 | 1.3E-01 | 3.6E-01 | 0.90 | ↓ | -0.16 | 6.2E-03 | 5.4E-02 | 0.78 | ↓ | -0.36 |
| NEG | 512.2985_4.81 | LysoPC(14:0) | 2.1E-03 | 3.2E-02 | 0.56 | ↓ | -0.83 | 1.5E-01 | 4.3E-01 | 0.78 | ↓ | -0.36 | 7.0E-03 | 8.1E-02 | 0.54 | ↓ | -0.90 |
| NEG | 652.4552_7.9 | LysoPC(24:0) | 4.1E-02 | 1.5E-01 | 0.73 | ↓ | -0.46 | 9.2E-01 | 9.7E-01 | 1.04 | ↑ | 0.06 | 6.6E-03 | 7.9E-02 | 0.66 | ↓ | -0.60 |
| NEG | 226.0175_2.4 | 4-Quinolinol, 8-methyl-2-(trifluoromethyl)- | 1.1E-01 | 2.6E-01 | 0.59 | ↓ | -0.75 | 5.1E-04 | 1.6E-02 | 0.31 | ↓ | -1.68 | 7.4E-02 | 2.6E-01 | 0.43 | ↓ | -1.22 |
| POS | 516.3104_4.97 | Oleyloxyethylphosphorylcholine | 3.8E-04 | 1.6E-02 | 0.53 | ↓ | -0.92 | 5.5E-03 | 8.3E-02 | 0.67 | ↓ | -0.57 | 8.5E-04 | 2.1E-02 | 0.49 | ↓ | -1.02 |
| POS | 162.1129_0.33 | L-Carnitine | 4.6E-04 | 1.6E-02 | 2.41 | ↑ | 1.27 | 7.6E-03 | 9.7E-02 | 1.92 | ↑ | 0.94 | 8.6E-03 | 6.2E-02 | 1.83 | ↑ | 0.87 |

| | | | | | | | | | | | | | | | | | |
|---|---|---|---|---|---|---|---|---|---|---|---|---|---|---|---|---|---|
| POS | 508.3397_5.21 | C18(Plasm) LPC | 1.2E-02 | 9.6E-02 | 0.64 | ↓ | -0.64 | 5.3E-03 | 8.2E-02 | 0.78 | ↓ | -0.35 | 8.2E-05 | 7.5E-03 | 0.66 | ↓ | -0.60 |
| NEG | 512.299_0.99 | LysoPC(14:0) | 3.2E-03 | 3.9E-02 | 0.53 | ↓ | -0.90 | 2.1E-01 | 3.1E-01 | 0.79 | ↓ | -0.34 | 3.0E-03 | 9.5E-02 | 0.47 | ↓ | -1.09 |
| NEG | 816.5759_9.01 | PC(16:0/20:4) | 1.1E-01 | 2.6E-01 | 0.74 | ↓ | -0.42 | 4.9E-01 | 7.6E-01 | 0.87 | ↓ | -0.19 | 3.6E-02 | 1.8E-01 | 0.65 | ↓ | -0.61 |
| POS | 204.1231_0.58 | Acetyl-DL-carnitine | 5.1E-02 | 3.3E-01 | 2.07 | ↑ | 1.05 | 5.6E-01 | 6.7E-01 | 1.24 | ↑ | 0.31 | 8.2E-02 | 2.2E-01 | 2.40 | ↑ | 1.27 |
| POS | 482.3253_1.13 | LysoPE(18:0) | 4.6E-03 | 8.7E-02 | 0.50 | ↓ | -1.01 | 8.8E-02 | 1.7E-01 | 0.72 | ↓ | -0.47 | 1.6E-03 | 2.4E-02 | 0.47 | ↓ | -1.08 |
| POS | 468.3092_4.81 | LysoPC(14:0) | 7.8E-04 | 2.1E-02 | 0.56 | ↓ | -0.82 | 1.2E-01 | 3.5E-01 | 0.78 | ↓ | -0.37 | 2.7E-03 | 3.6E-02 | 0.47 | ↓ | -1.09 |
| POS | 782.5682_4.96 | PC(16:0/18:1) | 3.4E-01 | 7.5E-01 | 1.21 | ↑ | 0.28 | 7.1E-05 | 3.3E-03 | 2.09 | ↑ | 1.06 | 9.5E-04 | 1.9E-02 | 4.09 | ↑ | 2.03 |
| POS | 530.368_6.36 | C18(Plasm) LPC | 5.1E-02 | 2.2E-01 | 2.90 | ↑ | 1.54 | 1.9E-02 | 1.6E-01 | 3.43 | ↑ | 1.78 | 6.8E-02 | 2.3E-01 | 2.57 | ↑ | 1.36 |
| POS | 530.3694_1.4 | C18(Plasm) LPC | 5.0E-02 | 3.3E-01 | 4.84 | ↑ | 2.27 | 7.1E-03 | 2.6E-02 | 4.05 | ↑ | 2.02 | 1.4E-01 | 3.2E-01 | 3.16 | ↑ | 1.66 |
| NEG | 526.31_1.14 | LysoPC(15:0) | 2.5E-02 | 1.5E-01 | 0.64 | ↓ | -0.65 | 5.5E-01 | 6.4E-01 | 0.88 | ↓ | -0.19 | 4.4E-03 | 9.5E-02 | 0.53 | ↓ | -0.92 |
| NEG | 229.1437_3.65 | Dodecanedioic acid | 5.6E-03 | 4.8E-02 | 0.46 | ↓ | -1.11 | 1.9E-02 | 1.4E-01 | 0.50 | ↓ | -0.99 | 2.7E-02 | 1.5E-01 | 0.54 | ↓ | -0.90 |
| POS | 790.6208_10.03 | PS(18:0/18:1) | 1.1E-01 | 3.4E-01 | 0.85 | ↓ | -0.23 | 4.3E-02 | 2.3E-01 | 1.21 | ↑ | 0.28 | 3.4E-03 | 4.0E-02 | 0.69 | ↓ | -0.54 |
| POS | 204.1141_0.33 | Acetyl-DL-carnitine | 1.4E-01 | 3.8E-01 | 1.70 | ↑ | 0.76 | 1.4E-01 | 3.8E-01 | 1.53 | ↑ | 0.61 | 3.7E-02 | 1.5E-01 | 2.00 | ↑ | 1.00 |
| POS | 316.2485_3.86 | Decanoyl-L-carnitine | 6.9E-02 | 2.6E-01 | 3.67 | ↑ | 1.88 | 4.1E-01 | 6.3E-01 | 1.40 | ↑ | 0.48 | 6.8E-02 | 2.3E-01 | 4.09 | ↑ | 2.03 |
| POS | 468.3099_0.99 | LysoPC(14:0) | 6.5E-04 | 3.5E-02 | 0.46 | ↓ | -1.11 | 3.6E-02 | 8.7E-02 | 0.66 | ↓ | -0.61 | 9.6E-04 | 1.9E-02 | 0.41 | ↓ | -1.29 |
| NEG | 584.3554_4.45 | 2-Thio PAF | 1.8E-03 | 3.0E-02 | 0.48 | ↓ | -1.06 | 9.5E-01 | 9.8E-01 | 0.97 | ↓ | -0.04 | 7.7E-02 | 2.7E-01 | 0.68 | ↓ | -0.56 |
| NEG | 178.0488_0.62 | Hippuric acid | 1.2E-01 | 3.6E-01 | 0.58 | ↓ | -0.78 | 5.0E-03 | 8.0E-02 | 0.38 | ↓ | -1.40 | 1.6E-01 | 3.2E-01 | 0.63 | ↓ | -0.66 |
| POS | 728.5584_10.1 | PE(18:0/18:1) | 6.5E-02 | 2.6E-01 | 0.74 | ↓ | -0.43 | 3.0E-01 | 5.4E-01 | 1.24 | ↑ | 0.31 | 6.7E-04 | 2.0E-02 | 0.38 | ↓ | -1.38 |
| NEG | 452.2775_0.99 | LysoPE(16:0) | 1.2E-02 | 9.3E-02 | 0.65 | ↓ | -0.62 | 7.0E-01 | 7.7E-01 | 0.94 | ↓ | -0.08 | 3.1E-03 | 9.5E-02 | 0.51 | ↓ | -0.96 |
| POS | 257.2267_3.76 | Androst-16-en-3-ol, (3β,5α)- | 1.9E-04 | 1.1E-02 | 2.82 | ↑ | 1.50 | 9.4E-05 | 9.3E-03 | 2.84 | ↑ | 1.51 | 1.2E-03 | 2.6E-02 | 3.20 | ↑ | 1.68 |
| NEG | 862.5542_4.81 | C18(Plasm)-22:6 PC | 3.9E-02 | 1.9E-01 | 0.55 | ↓ | -0.87 | 1.2E-02 | 9.4E-02 | 6.39 | ↑ | 2.68 | 4.9E-02 | 1.6E-01 | 4.17 | ↑ | 2.06 |
| POS | 806.5689_4.84 | Arachidonoylthio-PC | 6.7E-01 | 9.0E-01 | 1.05 | ↑ | 0.07 | 4.3E-04 | 4.9E-03 | 2.13 | ↑ | 1.09 | 1.4E-03 | 2.3E-02 | 3.53 | ↑ | 1.82 |
| NEG | 263.1033_1.9 | L-Glutamine, N2-(2-phenylacetyl)- | 2.7E-03 | 3.4E-02 | 0.42 | ↓ | -1.26 | 2.7E-02 | 1.7E-01 | 0.50 | ↓ | -1.01 | 7.0E-02 | 2.6E-01 | 0.66 | ↓ | -0.60 |
| POS | 504.3088_1.14 | Lysophosphatidylethanolamine | 2.7E-02 | 2.7E-01 | 0.58 | ↓ | -0.79 | 9.2E-02 | 1.7E-01 | 0.71 | ↓ | -0.50 | 3.3E-03 | 3.3E-02 | 0.51 | ↓ | -0.97 |
| POS | 524.3717_1.89 | LysoPC(18:0) | 2.7E-04 | 2.8E-02 | 0.50 | ↓ | -0.99 | 7.2E-02 | 1.4E-01 | 0.84 | ↓ | -0.24 | 8.3E-02 | 2.2E-01 | 0.82 | ↓ | -0.29 |
| NEG | 830.5807_5.1 | PC(18:0/18:2) | 4.0E-02 | 1.9E-01 | 0.55 | ↓ | -0.87 | 5.8E-02 | 1.4E-01 | 2.18 | ↑ | 1.12 | 3.8E-01 | 5.6E-01 | 1.77 | ↑ | 0.83 |
| POS | 544.338_1.41 | LysoPC(18:1) | 6.1E-03 | 1.0E-01 | 0.63 | ↓ | -0.67 | 9.2E-01 | 9.4E-01 | 0.99 | ↓ | -0.01 | 4.7E-03 | 3.9E-02 | 0.68 | ↓ | -0.55 |
| POS | 214.1134_3.03 | Suberylglycine | 5.1E-01 | 7.5E-01 | 1.08 | ↑ | 0.11 | 2.5E-09 | 6.4E-06 | 7.48 | ↑ | 2.90 | 2.5E-08 | 3.2E-05 | 5.37 | ↑ | 2.43 |
| NEG | 768.5551_5.11 | PC(18:0/18:2) | 5.6E-02 | 2.3E-01 | 0.57 | ↓ | -0.81 | 2.5E-02 | 1.1E-01 | 2.31 | ↑ | 1.21 | 2.7E-01 | 4.6E-01 | 1.61 | ↑ | 0.69 |
| POS | 130.0499_1.9 | Glutamic Acid | 4.3E-03 | 5.1E-02 | 0.45 | ↓ | -1.17 | 2.6E-02 | 1.9E-01 | 0.50 | ↓ | -1.01 | 2.4E-02 | 1.2E-01 | 0.59 | ↓ | -0.75 |
| POS | 490.2926_0.99 | LysoPC(14:0) | 6.3E-03 | 1.0E-01 | 0.42 | ↓ | -1.27 | 1.2E-01 | 2.1E-01 | 0.65 | ↓ | -0.61 | 4.6E-03 | 3.8E-02 | 0.41 | ↓ | -1.30 |
| POS | 788.6068_5.47 | PC(18:0/18:1) | 8.1E-01 | 9.4E-01 | 1.06 | ↑ | 0.09 | 1.6E-04 | 3.3E-03 | 2.53 | ↑ | 1.34 | 1.9E-02 | 8.5E-02 | 2.70 | ↑ | 1.43 |
| POS | 287.1015_0.61 | Phenylacetyl-L-glutamine | 2.2E-03 | 6.0E-02 | 0.43 | ↓ | -1.23 | 6.1E-03 | 2.4E-02 | 0.44 | ↓ | -1.17 | 1.6E-02 | 7.6E-02 | 0.55 | ↓ | -0.85 |
| POS | 786.6003_5.47 | 1,2-Dioleoyl-sn-glycero-3-PC | 8.4E-01 | 9.5E-01 | 1.07 | ↑ | 0.10 | 1.6E-04 | 3.3E-03 | 2.49 | ↑ | 1.32 | 1.6E-02 | 7.6E-02 | 2.76 | ↑ | 1.46 |
| POS | 532.3389_1.56 | LysoPC(17:0) | 2.4E-04 | 2.8E-02 | 0.31 | ↓ | -1.68 | 2.2E-02 | 5.9E-02 | 0.55 | ↓ | -0.87 | 4.1E-04 | 1.6E-02 | 0.38 | ↓ | -1.40 |
| POS | 794.5989_5.26 | C18(Plasm)-20:4 PC | 9.7E-01 | 9.9E-01 | 1.22 | ↑ | 0.28 | 3.5E-05 | 3.2E-03 | 3.08 | ↑ | 1.62 | 3.1E-03 | 3.2E-02 | 4.29 | ↑ | 2.10 |
| POS | 421.2528_5.75 | Tris(2-butoxyethyl) phosphate | 4.2E-02 | 2.0E-01 | 2.03 | ↑ | 1.02 | 1.9E-03 | 4.2E-02 | 2.85 | ↑ | 1.51 | 1.3E-02 | 8.1E-02 | 2.15 | ↑ | 1.11 |
| POS | 180.0972_0.33 | N-Benzylglycine methyl ester | 2.8E-01 | 5.6E-01 | 0.71 | ↓ | -0.49 | 6.9E-01 | 8.4E-01 | 0.99 | ↓ | -0.02 | 4.0E-02 | 1.6E-01 | 0.48 | ↓ | -1.05 |
| NEG | 826.5604_4.96 | PC(16:0/20:4) | 9.3E-01 | 9.8E-01 | 1.09 | ↑ | 0.13 | 1.7E-02 | 1.0E-01 | 2.14 | ↑ | 1.10 | 4.0E-03 | 9.5E-02 | 3.00 | ↑ | 1.59 |
| POS | 637.3064_1.8 | Acetylpodocarpic acid anhydride | 8.2E-02 | 3.9E-01 | 0.82 | ↓ | -0.29 | 5.2E-03 | 2.1E-02 | 1.45 | ↑ | 0.54 | 5.4E-05 | 9.9E-03 | 1.65 | ↑ | 0.73 |
| POS | 808.5841_5 | 1,2-Dioleoyl-sn-glycero-3-PC | 8.1E-01 | 9.4E-01 | 0.98 | ↓ | -0.03 | 2.4E-03 | 1.3E-02 | 1.86 | ↑ | 0.89 | 4.1E-03 | 3.7E-02 | 2.89 | ↑ | 1.53 |
| NEG | 766.5393_4.96 | PE(18:0/20:4) | 8.4E-01 | 9.4E-01 | 1.06 | ↑ | 0.08 | 4.5E-02 | 1.3E-01 | 2.05 | ↑ | 1.03 | 1.0E-02 | 1.0E-01 | 2.79 | ↑ | 1.48 |
| NEG | 835.5345_5.54 | PI(16:0/18:1) | 4.9E-01 | 7.5E-01 | 1.48 | ↑ | 0.56 | 1.5E-02 | 9.6E-02 | 3.12 | ↑ | 1.64 | 2.7E-02 | 1.3E-01 | 3.64 | ↑ | 1.86 |

| | | | | | | | | | | | | | | | | | |
|---|---|---|---|---|---|---|---|---|---|---|---|---|---|---|---|---|---|
| NEG | 850.5605_4.85 | PC(16;0/22:6) | 7.7E-01 | 9.2E-01 | 1.03 | ↑ | 0.05 | 4.1E-02 | 1.3E-01 | 2.39 | ↑ | 1.26 | 5.0E-03 | 9.5E-02 | 3.11 | ↑ | 1.64 |
| POS | 808.5769_4.83 | 1,2-Dioleoyl-sn-glycero-3-PC | 3.4E-01 | 7.5E-01 | 0.86 | ↓ | -0.22 | 9.5E-04 | 7.8E-03 | 1.86 | ↑ | 0.90 | 1.6E-03 | 2.5E-02 | 2.73 | ↑ | 1.45 |
| POS | 502.3262_1.49 | C16(Plasm) LPC | 9.5E-04 | 4.0E-02 | 0.49 | ↓ | -1.03 | 6.6E-01 | 7.6E-01 | 0.95 | ↓ | -0.08 | 2.9E-01 | 5.0E-01 | 0.84 | ↓ | -0.25 |
| POS | 678.4682_6.87 | PC(14:0) | 3.4E-02 | 1.8E-01 | 0.59 | ↓ | -0.77 | 4.2E-05 | 6.5E-03 | 0.29 | ↓ | -1.80 | 5.4E-06 | 2.0E-03 | 0.24 | ↓ | -2.06 |
| POS | 546.3536_1.89 | LysoPC(18:0) | 6.5E-04 | 3.5E-02 | 0.63 | ↓ | -0.68 | 1.1E-01 | 1.9E-01 | 0.90 | ↓ | -0.15 | 5.9E-02 | 1.8E-01 | 0.87 | ↓ | -0.21 |
| NEG | 770.571_5.48 | DMPE(18:1/18:1) | 2.4E-01 | 5.2E-01 | 0.77 | ↓ | -0.38 | 8.3E-03 | 8.9E-02 | 2.78 | ↑ | 1.48 | 4.2E-02 | 1.5E-01 | 2.51 | ↑ | 1.33 |
| NEG | 832.5981_5.48 | PC(18:1/18:0) | 3.6E-01 | 6.3E-01 | 0.80 | ↓ | -0.31 | 1.6E-02 | 9.8E-02 | 2.75 | ↑ | 1.46 | 4.3E-02 | 1.5E-01 | 2.62 | ↑ | 1.39 |
| POS | 792.5895_5.18 | Dihomo-γ-linolenoyl PAF C-16 | 7.8E-01 | 9.3E-01 | 1.20 | ↑ | 0.26 | 1.0E-03 | 8.0E-03 | 2.80 | ↑ | 1.49 | 3.9E-03 | 3.6E-02 | 5.59 | ↑ | 2.48 |
| NEG | 794.5685_5.41 | PC(18:1/16:0) | 6.2E-01 | 8.4E-01 | 0.99 | ↓ | -0.01 | 4.0E-02 | 1.3E-01 | 2.39 | ↑ | 1.26 | 1.0E-02 | 1.0E-01 | 3.25 | ↑ | 1.70 |
| POS | 750.5453_5.35 | Palmitoyl Thio-PC | 4.6E-01 | 8.1E-01 | 1.44 | ↑ | 0.52 | 2.0E-04 | 3.4E-03 | 2.79 | ↑ | 1.48 | 2.3E-03 | 2.9E-02 | 6.12 | ↑ | 2.61 |
| POS | 810.5994_5.42 | PC(18:0/18:1) | 4.1E-01 | 8.0E-01 | 1.26 | ↑ | 0.33 | 6.8E-04 | 6.6E-03 | 2.26 | ↑ | 1.18 | 7.7E-04 | 1.9E-02 | 4.08 | ↑ | 2.03 |
| NEG | 832.6078_5.77 | PC(18:1/18:0) | 3.8E-01 | 6.4E-01 | 0.85 | ↓ | -0.24 | 1.9E-02 | 1.0E-01 | 4.49 | ↑ | 2.17 | 6.1E-02 | 1.8E-01 | 3.16 | ↑ | 1.66 |
| NEG | 830.5917_5.48 | PC(18:1/18:1) | 3.0E-01 | 5.7E-01 | 0.80 | ↓ | -0.33 | 2.2E-02 | 1.0E-01 | 2.39 | ↑ | 1.26 | 5.1E-02 | 1.6E-01 | 2.19 | ↑ | 1.13 |
| NEG | 747.5162_5.21 | SM(d18:1/16:0) | 7.5E-01 | 9.0E-01 | 1.55 | ↑ | 0.64 | 2.1E-02 | 1.0E-01 | 5.51 | ↑ | 2.46 | 2.0E-03 | 9.5E-02 | 8.01 | ↑ | 3.00 |
| NEG | 762.5082_4.98 | PE(16:0/22:6) | 4.2E-01 | 6.7E-01 | 1.95 | ↑ | 0.97 | 1.0E-02 | 9.3E-02 | 5.84 | ↑ | 2.55 | 9.7E-03 | 1.0E-01 | 7.20 | ↑ | 2.85 |
| POS | 790.5725_5.24 | Arachidonoyl PAF C-16 | 5.9E-01 | 8.8E-01 | 1.44 | ↑ | 0.52 | 5.8E-05 | 3.2E-03 | 4.43 | ↑ | 2.15 | 1.6E-03 | 2.4E-02 | 5.65 | ↑ | 2.50 |
| NEG | 765.5276_5.16 | SM(d18:1/18:0) | 1.8E-01 | 4.5E-01 | 0.76 | ↓ | -0.40 | 1.9E-02 | 1.0E-01 | 2.46 | ↑ | 1.30 | 7.3E-02 | 2.0E-01 | 2.77 | ↑ | 1.47 |
| NEG | 820.577_5.49 | PC(18:0/18:2) | 2.8E-01 | 5.5E-01 | 0.77 | ↓ | -0.37 | 5.3E-02 | 1.4E-01 | 2.16 | ↑ | 1.11 | 2.5E-02 | 1.2E-01 | 1.97 | ↑ | 0.97 |
| NEG | 790.5397_5.44 | PE(18:0/22:6) | 4.1E-01 | 6.6E-01 | 1.99 | ↑ | 1.00 | 5.9E-03 | 8.0E-02 | 6.50 | ↑ | 2.70 | 4.1E-03 | 9.5E-02 | 6.59 | ↑ | 2.72 |
| NEG | 806.5944_5.72 | PC(16:0/18:0) | 7.3E-01 | 9.0E-01 | 1.21 | ↑ | 0.27 | 1.2E-02 | 9.4E-02 | 3.67 | ↑ | 1.88 | 7.0E-03 | 9.8E-02 | 3.17 | ↑ | 1.66 |
| NEG | 790.541_4.84 | PE(18:0/22:6) | 5.8E-01 | 8.2E-01 | 0.95 | ↓ | -0.07 | 1.7E-02 | 1.0E-01 | 2.43 | ↑ | 1.28 | 5.1E-03 | 9.5E-02 | 3.05 | ↑ | 1.61 |
| NEG | 878.5916_5.31 | PC(18:0/22:6) | 5.6E-01 | 8.0E-01 | 0.94 | ↓ | -0.08 | 4.1E-02 | 1.3E-01 | 2.73 | ↑ | 1.45 | 5.7E-03 | 9.7E-02 | 3.48 | ↑ | 1.80 |
| NEG | 766.5395_5.55 | PE(18:0/24:0) | 6.2E-01 | 8.4E-01 | 1.46 | ↑ | 0.54 | 1.1E-02 | 9.4E-02 | 4.62 | ↑ | 2.21 | 1.4E-02 | 1.1E-01 | 5.30 | ↑ | 2.41 |
| POS | 766.5724_5.16 | Eicosapentaenoyl PAF C-16 | 8.0E-01 | 9.4E-01 | 1.22 | ↑ | 0.29 | 1.0E-03 | 8.1E-03 | 2.86 | ↑ | 1.52 | 2.7E-03 | 3.0E-02 | 5.60 | ↑ | 2.49 |
| POS | 760.5848_5.35 | PC(16:0/18:1) | 6.4E-01 | 8.9E-01 | 1.23 | ↑ | 0.30 | 5.7E-05 | 3.2E-03 | 3.10 | ↑ | 1.63 | 6.4E-03 | 4.3E-02 | 3.23 | ↑ | 1.69 |
| NEG | 776.5592_5.71 | PC(18:1/14:0) | 7.7E-01 | 9.1E-01 | 1.41 | ↑ | 0.50 | 2.8E-02 | 1.1E-01 | 5.26 | ↑ | 2.40 | 5.4E-03 | 9.5E-02 | 6.85 | ↑ | 2.78 |
| NEG | 768.5463_5.54 | PC(18:0/14:0) | 8.4E-01 | 9.4E-01 | 1.27 | ↑ | 0.34 | 2.2E-02 | 1.0E-01 | 3.66 | ↑ | 1.87 | 3.1E-02 | 1.4E-01 | 3.71 | ↑ | 1.89 |
| POS | 782.567_5.35 | PC(16:0/18:1) | 5.0E-01 | 8.3E-01 | 1.29 | ↑ | 0.37 | 5.8E-05 | 3.2E-03 | 2.89 | ↑ | 1.53 | 2.3E-03 | 2.9E-02 | 2.41 | ↑ | 1.27 |
| NEG | 838.5972_5.6 | C18(Plasm)-20:4 PC | 8.9E-01 | 9.6E-01 | 1.18 | ↑ | 0.24 | 6.2E-02 | 1.4E-01 | 3.79 | ↑ | 1.92 | 2.0E-02 | 1.2E-01 | 5.53 | ↑ | 2.47 |
| NEG | 804.5669_5.01 | PC(18:1/16:0) | 6.3E-01 | 8.5E-01 | 0.96 | ↓ | -0.06 | 6.5E-03 | 8.2E-02 | 2.39 | ↑ | 1.25 | 1.7E-02 | 1.2E-01 | 2.23 | ↑ | 1.16 |
| NEG | 802.5603_5.01 | PC(16:0/18:2) | 6.0E-01 | 8.3E-01 | 0.95 | ↓ | -0.08 | 7.4E-03 | 8.6E-02 | 2.09 | ↑ | 1.06 | 1.9E-02 | 1.2E-01 | 1.98 | ↑ | 0.98 |
| NEG | 744.549_5.01 | α-Galactosyl-N-palmitoylsphingosine | 5.6E-01 | 8.0E-01 | 0.91 | ↓ | -0.14 | 1.8E-02 | 1.0E-01 | 2.36 | ↑ | 1.24 | 6.2E-02 | 1.8E-01 | 1.79 | ↑ | 0.84 |
| POS | 768.5893_5.23 | C16-18:1 PC | 7.1E-01 | 9.2E-01 | 1.25 | ↑ | 0.32 | 3.4E-04 | 4.2E-03 | 3.28 | ↑ | 1.72 | 5.0E-03 | 3.9E-02 | 5.05 | ↑ | 2.34 |
| NEG | 774.545_5.65 | C18(Plasm)-22:6 PE | 9.7E-01 | 9.9E-01 | 1.27 | ↑ | 0.34 | 3.0E-02 | 1.2E-01 | 5.90 | ↑ | 2.56 | 6.4E-03 | 9.8E-02 | 6.40 | ↑ | 2.68 |
| NEG | 714.5081_5.16 | PE(16:0/18:2) | 6.9E-01 | 8.8E-01 | 1.23 | ↑ | 0.30 | 6.1E-03 | 8.0E-02 | 3.87 | ↑ | 1.95 | 9.1E-02 | 2.2E-01 | 3.55 | ↑ | 1.83 |
| NEG | 792.5473_5.01 | PC(16:0/18:2) | 2.7E-01 | 5.5E-01 | 0.79 | ↓ | -0.35 | 1.1E-01 | 2.0E-01 | 1.72 | ↑ | 0.78 | 5.4E-02 | 1.7E-01 | 1.98 | ↑ | 0.99 |
| POS | 735.5737_5.29 | Thioetheramidephosphatidylcholine | 1.3E-01 | 4.8E-01 | 1.78 | ↑ | 0.83 | 7.0E-06 | 2.6E-03 | 3.65 | ↑ | 1.87 | 5.3E-03 | 4.0E-02 | 5.02 | ↑ | 2.33 |
| POS | 756.5531_5.29 | PC(16:0/16:0) | 5.7E-02 | 3.4E-01 | 2.04 | ↑ | 1.03 | 2.5E-05 | 3.1E-03 | 3.12 | ↑ | 1.64 | 8.3E-04 | 1.9E-02 | 3.30 | ↑ | 1.72 |
| POS | 184.0734_5 | PC | 9.1E-01 | 9.7E-01 | 1.04 | ↑ | 0.06 | 2.0E-04 | 3.4E-03 | 2.47 | ↑ | 1.30 | 9.5E-03 | 5.5E-02 | 3.02 | ↑ | 1.59 |
| NEG | 806.5831_5.35 | PC(16:0/18:0) | 6.3E-01 | 8.5E-01 | 1.02 | ↑ | 0.03 | 1.2E-02 | 9.4E-02 | 3.71 | ↑ | 1.89 | 4.0E-02 | 1.5E-01 | 2.66 | ↑ | 1.41 |
| POS | 834.5998_5.31 | 1-Docosahexaenoyl-2-stearoyl-sn-glycero-3-p | 8.4E-01 | 9.5E-01 | 1.01 | ↑ | 0.01 | 1.8E-03 | 1.1E-02 | 2.49 | ↑ | 1.32 | 3.0E-03 | 3.2E-02 | 4.82 | ↑ | 2.27 |
| NEG | 834.5429_5.53 | PS(18:0/22:6) | 9.1E-01 | 9.7E-01 | 1.09 | ↑ | 0.12 | 1.0E-02 | 9.3E-02 | 3.10 | ↑ | 1.63 | 3.8E-02 | 1.4E-01 | 3.08 | ↑ | 1.62 |
| NEG | 742.5398_5.01 | PE(18:0/18:2) | 6.5E-01 | 8.6E-01 | 1.00 | ↓ | -0.01 | 5.5E-03 | 8.0E-02 | 2.25 | ↑ | 1.17 | 5.9E-02 | 1.7E-01 | 1.91 | ↑ | 0.93 |

| | | | | | | | | | | | | | | | | | |
|---|---|---|---|---|---|---|---|---|---|---|---|---|---|---|---|---|---|
| NEG | 862.5958_5.51 | C18(Plasm)-22:6 PC | 7.1E-01 | 9.0E-01 | 1.02 | ↑ | 0.03 | 1.2E-01 | 2.1E-01 | 2.73 | ↑ | 1.45 | 2.2E-02 | 1.2E-01 | 3.54 | ↑ | 1.82 |
| POS | 758.5692_5.01 | PC(16:0/18:2) | 7.8E-01 | 9.3E-01 | 1.15 | ↑ | 0.20 | 8.0E-05 | 3.3E-03 | 2.37 | ↑ | 1.25 | 7.2E-03 | 4.5E-02 | 2.78 | ↑ | 1.47 |
| POS | 760.5746_5.01 | PC(16:0/18:1) | 8.1E-01 | 9.4E-01 | 1.13 | ↑ | 0.18 | 1.0E-04 | 3.3E-03 | 2.24 | ↑ | 1.17 | 6.8E-03 | 4.4E-02 | 2.57 | ↑ | 1.36 |
| NEG | 804.576_5.36 | PC(18:1/16:0) | 7.6E-01 | 9.1E-01 | 1.04 | ↑ | 0.06 | 6.1E-03 | 8.0E-02 | 3.40 | ↑ | 1.76 | 2.2E-02 | 1.2E-01 | 2.65 | ↑ | 1.41 |
| POS | 768.5582_5.55 | PE(18:0/18:1) | 2.7E-01 | 6.7E-01 | 1.80 | ↑ | 0.85 | 1.9E-03 | 1.1E-02 | 2.89 | ↑ | 1.53 | 1.1E-02 | 5.9E-02 | 4.13 | ↑ | 2.05 |
| NEG | 738.5085_5.1 | PE(16:0/20:4) | 9.5E-01 | 9.8E-01 | 1.26 | ↑ | 0.33 | 1.3E-02 | 9.4E-02 | 3.66 | ↑ | 1.87 | 2.8E-02 | 1.3E-01 | 4.65 | ↑ | 2.22 |
| NEG | 802.5603_8.81 | PC(16:0/18:2) | 3.6E-01 | 5.4E-01 | 1.03 | ↑ | 0.04 | 2.0E-02 | 1.4E-01 | 1.12 | ↑ | 0.16 | 5.4E-03 | 7.4E-02 | 1.14 | ↑ | 0.18 |

**7 Gy**

| | | | SD 1 vs. Pre | | | | | SD 2 vs. Pre | | | | | SD 6 vs. Pre | | | | |
|---|---|---|---|---|---|---|---|---|---|---|---|---|---|---|---|---|---|
| **Mode** | **m/z_RT** | **Name** | **p-value** | **FDR** | **FC** | | **LOG2FC** | **p-value** | **FDR** | **FC** | | **LOG2FC** | **p-value** | **FDR** | **FC** | | **LOG2FC** |
| POS | 162.1127_0.57 | L-Carnitine | 2.4E-06 | 3.5E-03 | 3.22 | ↑ | 1.69 | 7.2E-05 | 8.5E-03 | 2.03 | ↑ | 1.02 | 1.0E-03 | 5.9E-02 | 2.05 | ↑ | 1.04 |
| POS | 163.1155_0.56 | Phenylpiperazine | 8.5E-05 | 2.0E-02 | 2.64 | ↑ | 1.40 | 4.4E-03 | 2.9E-02 | 1.73 | ↑ | 0.79 | 1.2E-02 | 1.5E-01 | 1.77 | ↑ | 0.83 |
| NEG | 861.5497_12.17 | PI(18:1) | 1.2E-03 | 4.1E-02 | 0.58 | ↓ | -0.78 | 5.7E-01 | 8.1E-01 | 1.07 | ↑ | 0.10 | 6.5E-04 | 2.8E-02 | 0.55 | ↓ | -0.87 |
| POS | 580.432_7.28 | LysoPC(22:0) | 1.7E-02 | 1.4E-01 | 0.52 | ↓ | -0.95 | 4.5E-01 | 6.9E-01 | 0.87 | ↓ | -0.19 | 3.0E-03 | 5.0E-02 | 0.51 | ↓ | -0.98 |
| POS | 548.3722_1.53 | Lyso-PAF C-18 | 4.4E-03 | 1.3E-01 | 0.56 | ↓ | -0.83 | 1.2E-04 | 8.5E-03 | 0.52 | ↓ | -0.94 | 1.8E-02 | 1.7E-01 | 0.60 | ↓ | -0.74 |
| POS | 552.4031_6.62 | LysoPC(20:0) | 2.0E-02 | 1.6E-01 | 0.48 | ↓ | -1.05 | 6.5E-01 | 8.3E-01 | 0.91 | ↓ | -0.14 | 1.6E-02 | 1.1E-01 | 0.51 | ↓ | -0.96 |
| POS | 782.5682_8.51 | PC(16:0/18:1) | 2.8E-04 | 2.4E-02 | 0.35 | ↓ | -1.50 | 9.9E-01 | 1.0E+00 | 0.99 | ↓ | -0.02 | 6.7E-03 | 7.4E-02 | 0.49 | ↓ | -1.02 |
| NEG | 790.5343_8.7 | PI(18:0/22:6) | 8.0E-03 | 8.7E-02 | 0.66 | ↓ | -0.59 | 1.6E-02 | 2.1E-01 | 0.75 | ↓ | -0.41 | 2.0E-02 | 1.3E-01 | 0.75 | ↓ | -0.42 |
| POS | 510.3561_5.63 | LysoPC(17:0) | 2.3E-03 | 5.6E-02 | 0.61 | ↓ | -0.72 | 4.7E-02 | 2.8E-01 | 0.77 | ↓ | -0.37 | 2.9E-03 | 5.0E-02 | 0.63 | ↓ | -0.67 |
| POS | 339.3453_7.67 | 9-Oxo-11α,15R-dihydroxyprostan-1-oic acid | 1.5E-01 | 4.6E-01 | 0.37 | ↓ | -1.42 | 4.3E-01 | 6.8E-01 | 0.49 | ↓ | -1.02 | 5.4E-03 | 6.6E-02 | 0.15 | ↓ | -2.78 |
| POS | 338.3423_7.68 | 13-Docosenamide, (Z)- | 1.4E-01 | 4.5E-01 | 0.38 | ↓ | -1.39 | 4.3E-01 | 6.8E-01 | 0.51 | ↓ | -0.98 | 4.8E-03 | 6.3E-02 | 0.15 | ↓ | -2.71 |
| NEG | 554.3457_5.63 | LysoPC(17:0) | 1.5E-03 | 4.5E-02 | 0.64 | ↓ | -0.64 | 1.2E-01 | 4.5E-01 | 0.84 | ↓ | -0.25 | 2.1E-02 | 1.3E-01 | 0.72 | ↓ | -0.47 |
| POS | 608.4631_7.89 | LysoPC(24:0) | 1.2E-01 | 4.2E-01 | 0.74 | ↓ | -0.44 | 4.4E-01 | 6.8E-01 | 0.90 | ↓ | -0.15 | 6.7E-03 | 7.4E-02 | 0.61 | ↓ | -0.70 |
| NEG | 826.5599_8.52 | PC(16:0/20:4) | 3.1E-04 | 2.7E-02 | 0.60 | ↓ | -0.74 | 9.9E-01 | 1.0E+00 | #N/A | | #N/A | 3.1E-02 | 1.6E-01 | 0.78 | ↓ | -0.36 |
| NEG | 596.3925_6.63 | LysoPC(20:0) | 2.2E-02 | 1.5E-01 | 0.50 | ↓ | -0.99 | 6.0E-01 | 8.2E-01 | 0.90 | ↓ | -0.15 | 5.2E-03 | 6.2E-02 | 0.47 | ↓ | -1.08 |
| POS | 770.5677_8.69 | Dihomo-γ-linolenoyl PAF C-16 | 4.3E-03 | 7.1E-02 | 0.56 | ↓ | -0.85 | 1.1E-01 | 3.9E-01 | 0.79 | ↓ | -0.34 | 7.0E-03 | 7.4E-02 | 0.56 | ↓ | -0.84 |
| POS | 532.3383_5.63 | LysoPC(17:0) | 4.1E-03 | 7.0E-02 | 0.63 | ↓ | -0.66 | 9.7E-02 | 3.6E-01 | 0.82 | ↓ | -0.29 | 8.9E-03 | 8.5E-02 | 0.67 | ↓ | -0.58 |
| NEG | 861.5499_11.73 | PI(18:1) | 2.0E-03 | 4.5E-02 | 0.54 | ↓ | -0.90 | 4.7E-01 | 7.5E-01 | 0.94 | ↓ | -0.08 | 7.3E-05 | 1.6E-02 | 0.45 | ↓ | -1.17 |
| POS | 184.0945_0.58 | PC | 1.5E-05 | 7.2E-03 | 3.86 | ↑ | 1.95 | 8.0E-04 | 1.2E-02 | 2.61 | ↑ | 1.38 | 3.5E-03 | 9.4E-02 | 2.46 | ↑ | 1.30 |
| NEG | 788.5249_8.71 | PS(18:0/18:1) | 1.2E-03 | 4.3E-02 | 0.53 | ↓ | -0.91 | 1.6E-02 | 2.1E-01 | 0.68 | ↓ | -0.56 | 1.6E-02 | 1.1E-01 | 0.65 | ↓ | -0.63 |
| POS | 518.3256_4.86 | LysoPC(16:0) | 4.3E-04 | 2.8E-02 | 0.56 | ↓ | -0.85 | 5.5E-02 | 3.0E-01 | 0.74 | ↓ | -0.43 | 1.5E-03 | 3.6E-02 | 0.45 | ↓ | -1.16 |
| NEG | 828.5667_8.52 | C18(Plasm)-20:4 PC | 1.3E-03 | 4.5E-02 | 0.53 | ↓ | -0.91 | 7.9E-01 | 9.2E-01 | 0.97 | ↓ | -0.04 | 2.1E-02 | 1.3E-01 | 0.68 | ↓ | -0.55 |
| NEG | 827.5636_8.52 | KRTLRR | 9.5E-04 | 3.7E-02 | 0.57 | ↓ | -0.80 | 9.1E-01 | 9.7E-01 | 0.99 | ↓ | -0.02 | 3.4E-02 | 1.7E-01 | 0.73 | ↓ | -0.46 |
| NEG | 526.3113_5.06 | LysoPC(15:0) | 1.0E-02 | 9.3E-02 | 0.67 | ↓ | -0.57 | 2.9E-01 | 6.3E-01 | 0.88 | ↓ | -0.18 | 3.2E-03 | 5.0E-02 | 0.68 | ↓ | -0.56 |
| NEG | 594.3771_6.08 | Pyrrolidino PAF C-16 | 4.9E-02 | 2.2E-01 | 0.60 | ↓ | -0.73 | 5.1E-01 | 7.7E-01 | 0.86 | ↓ | -0.21 | 5.2E-02 | 2.1E-01 | 0.61 | ↓ | -0.72 |
| POS | 510.3564_1.56 | LysoPC(17:0) | 2.0E-03 | 8.2E-02 | 0.57 | ↓ | -0.82 | 1.6E-02 | 6.5E-02 | 0.64 | ↓ | -0.65 | 5.1E-04 | 4.0E-02 | 0.47 | ↓ | -1.08 |
| POS | 482.3248_5.06 | LysoPC(15:0) | 3.9E-03 | 6.9E-02 | 0.63 | ↓ | -0.68 | 1.2E-01 | 4.0E-01 | 0.82 | ↓ | -0.29 | 4.7E-04 | 2.0E-02 | 0.53 | ↓ | -0.92 |
| NEG | 554.3463_1.56 | LysoPC(17:0) | 9.9E-04 | 4.7E-02 | 0.47 | ↓ | -1.08 | 3.8E-02 | 8.6E-02 | 0.72 | ↓ | -0.47 | 2.8E-02 | 1.3E-01 | 0.63 | ↓ | -0.67 |
| POS | 552.4037_2.62 | LysoPC(20:0) | 1.7E-01 | 4.8E-01 | 0.77 | ↓ | -0.37 | 3.7E-01 | 5.9E-01 | 0.88 | ↓ | -0.19 | 1.9E-02 | 1.7E-01 | 0.59 | ↓ | -0.76 |
| NEG | 624.4238_7.29 | LysoPC(22:0) | 2.9E-02 | 1.7E-01 | 0.55 | ↓ | -0.87 | 2.6E-01 | 6.1E-01 | 0.79 | ↓ | -0.34 | 3.1E-03 | 4.9E-02 | 0.52 | ↓ | -0.95 |
| POS | 524.3722_5.96 | LysoPC(18:0) | 7.7E-03 | 9.7E-02 | 0.71 | ↓ | -0.49 | 3.5E-01 | 6.2E-01 | 0.91 | ↓ | -0.13 | 1.3E-01 | 3.1E-01 | 0.85 | ↓ | -0.23 |
| NEG | 512.2985_4.81 | LysoPC(14:0) | 3.7E-03 | 5.7E-02 | 0.57 | ↓ | -0.81 | 2.0E-01 | 5.4E-01 | 0.78 | ↓ | -0.35 | 1.2E-02 | 9.4E-02 | 0.50 | ↓ | -1.00 |
| NEG | 652.4552_7.9 | LysoPC(24:0) | 5.5E-02 | 2.3E-01 | 0.71 | ↓ | -0.50 | 1.9E-01 | 5.4E-01 | 0.83 | ↓ | -0.26 | 9.3E-03 | 8.4E-02 | 0.66 | ↓ | -0.59 |

| | | | | | | | | | | | | | | | | | |
|---|---|---|---|---|---|---|---|---|---|---|---|---|---|---|---|---|---|
| NEG | 226.0175_2.4 | 4-Quinolinol, 8-methyl-2-(trifluoromethyl)- | 1.9E-03 | 4.5E-02 | 0.26 | ↓ | -1.94 | 8.6E-04 | 7.4E-02 | 0.20 | ↓ | -2.31 | 7.1E-03 | 7.1E-02 | 0.35 | ↓ | -1.51 |
| POS | 516.3104_4.97 | Oleyloxyethylphosphorylcholine | 4.0E-03 | 7.0E-02 | 0.63 | ↓ | -0.68 | 1.9E-01 | 4.8E-01 | 0.80 | ↓ | -0.32 | 1.7E-02 | 1.2E-01 | 0.60 | ↓ | -0.73 |
| POS | 162.1129_0.33 | L-Carnitine | 4.0E-05 | 1.1E-02 | 2.85 | ↑ | 1.51 | 9.4E-04 | 4.3E-02 | 2.11 | ↑ | 1.08 | 4.1E-02 | 1.9E-01 | 1.51 | ↑ | 0.60 |
| POS | 508.3397_5.21 | C18(Plasm) LPC | 8.4E-03 | 1.0E-01 | 0.60 | ↓ | -0.75 | 5.4E-01 | 7.5E-01 | 0.90 | ↓ | -0.15 | 4.5E-02 | 2.0E-01 | 0.68 | ↓ | -0.57 |
| NEG | 512.299_0.99 | LysoPC(14:0) | 9.2E-03 | 1.2E-01 | 0.56 | ↓ | -0.83 | 2.5E-01 | 3.8E-01 | 0.81 | ↓ | -0.30 | 8.4E-03 | 9.6E-02 | 0.52 | ↓ | -0.94 |
| NEG | 816.5759_9.01 | PC(16:0/20:4) | 3.6E-02 | 1.9E-01 | 0.68 | ↓ | -0.55 | 2.0E-01 | 5.5E-01 | 0.82 | ↓ | -0.28 | 6.0E-03 | 6.7E-02 | 0.62 | ↓ | -0.70 |
| POS | 204.1231_0.58 | Acetyl-DL-carnitine | 2.4E-04 | 3.6E-02 | 2.62 | ↑ | 1.39 | 3.5E-02 | 1.1E-01 | 1.57 | ↑ | 0.65 | 5.3E-03 | 1.1E-01 | 3.17 | ↑ | 1.66 |
| POS | 482.3253_1.13 | LysoPE(18:0) | 2.7E-04 | 3.6E-02 | 0.55 | ↓ | -0.87 | 6.8E-02 | 1.8E-01 | 0.77 | ↓ | -0.38 | 7.5E-05 | 1.9E-02 | 0.38 | ↓ | -1.40 |
| POS | 468.3092_4.81 | LysoPC(14:0) | 4.1E-03 | 7.0E-02 | 0.59 | ↓ | -0.75 | 6.6E-01 | 8.3E-01 | 0.93 | ↓ | -0.11 | 9.4E-03 | 8.5E-02 | 0.53 | ↓ | -0.91 |
| POS | 782.5682_4.96 | PC(16:0/18:1) | 8.8E-02 | 4.2E-01 | 5.61 | ↑ | 2.49 | 1.7E-03 | 1.7E-02 | 2.62 | ↑ | 1.39 | 2.5E-02 | 1.9E-01 | 4.87 | ↑ | 2.28 |
| POS | 530.368_6.36 | C18(Plasm) LPC | 8.0E-03 | 9.9E-02 | 4.31 | ↑ | 2.11 | 1.3E-03 | 4.9E-02 | 4.89 | ↑ | 2.29 | 1.2E-02 | 9.6E-02 | 4.11 | ↑ | 2.04 |
| POS | 530.3694_1.4 | C18(Plasm) LPC | 6.3E-02 | 3.9E-01 | 3.02 | ↑ | 1.59 | 1.3E-02 | 5.7E-02 | 3.35 | ↑ | 1.75 | 4.6E-02 | 2.4E-01 | 3.10 | ↑ | 1.63 |
| NEG | 526.31_1.14 | LysoPC(15:0) | 2.2E-02 | 1.4E-01 | 0.63 | ↓ | -0.67 | 1.6E-01 | 2.7E-01 | 0.79 | ↓ | -0.34 | 1.5E-03 | 6.3E-02 | 0.53 | ↓ | -0.93 |
| NEG | 229.1437_3.65 | Dodecanedioic acid | 8.4E-03 | 8.9E-02 | 0.43 | ↓ | -1.21 | 2.0E-02 | 2.5E-01 | 0.48 | ↓ | -1.05 | 1.6E-01 | 3.4E-01 | 0.63 | ↓ | -0.66 |
| POS | 790.6208_10.03 | PS(18:0/18:1) | 2.0E-01 | 5.3E-01 | 0.89 | ↓ | -0.16 | 2.8E-01 | 5.7E-01 | 1.10 | ↑ | 0.13 | 1.0E-02 | 8.6E-02 | 0.78 | ↓ | -0.36 |
| POS | 204.1141_0.33 | Acetyl-DL-carnitine | 2.3E-05 | 7.5E-03 | 2.07 | ↑ | 1.05 | 8.5E-03 | 1.3E-01 | 1.45 | ↑ | 0.54 | 1.5E-02 | 1.1E-01 | 1.98 | ↑ | 0.99 |
| POS | 316.2485_3.86 | Decanoyl-L-carnitine | 5.8E-04 | 3.0E-02 | 6.37 | ↑ | 2.67 | 9.6E-02 | 3.6E-01 | 2.08 | ↑ | 1.05 | 4.6E-02 | 2.0E-01 | 7.41 | ↑ | 2.89 |
| POS | 468.3099_0.99 | LysoPC(14:0) | 2.7E-03 | 9.4E-02 | 0.53 | ↓ | -0.91 | 5.4E-02 | 1.5E-01 | 0.67 | ↓ | -0.57 | 5.8E-04 | 4.1E-02 | 0.36 | ↓ | -1.48 |
| NEG | 584.3554_4.45 | 2-Thio PAF | 3.6E-03 | 5.7E-02 | 0.43 | ↓ | -1.21 | 4.2E-01 | 7.1E-01 | 0.82 | ↓ | -0.29 | 1.4E-02 | 1.1E-01 | 0.53 | ↓ | -0.90 |
| NEG | 178.0488_0.62 | Hippuric acid | 4.7E-03 | 9.3E-02 | 0.38 | ↓ | -1.38 | 6.9E-03 | 2.0E-02 | 0.42 | ↓ | -1.26 | 5.0E-01 | 6.0E-01 | 1.05 | ↑ | 0.06 |
| POS | 728.5584_10.1 | PE(18:0/18:1) | 1.2E-02 | 1.2E-01 | 0.54 | ↓ | -0.89 | 8.2E-01 | 9.2E-01 | 1.00 | ↓ | 0.00 | 2.3E-04 | 1.3E-02 | 0.34 | ↓ | -1.54 |
| NEG | 452.2775_0.99 | LysoPE(16:0) | 3.6E-02 | 1.6E-01 | 0.70 | ↓ | -0.52 | 9.1E-01 | 9.4E-01 | 0.98 | ↓ | -0.02 | 2.1E-02 | 1.2E-01 | 0.60 | ↓ | -0.74 |
| POS | 257.2267_3.76 | Androst-16-en-3-ol, (3β,5α)- | 2.7E-02 | 1.9E-01 | 2.43 | ↑ | 1.28 | 3.1E-02 | 2.4E-01 | 2.56 | ↑ | 1.36 | 2.0E-01 | 3.9E-01 | 1.63 | ↑ | 0.71 |
| NEG | 862.5542_4.81 | C18(Plasm)-22:6 PC | 2.4E-01 | 4.1E-01 | 2.96 | ↑ | 1.56 | 2.2E-04 | 1.6E-03 | 7.95 | ↑ | 2.99 | 6.5E-02 | 1.6E-01 | 3.63 | ↑ | 1.86 |
| POS | 806.5689_4.84 | Arachidonoylthio-PC | 3.2E-02 | 2.8E-01 | 3.43 | ↑ | 1.78 | 6.1E-04 | 1.2E-02 | 2.65 | ↑ | 1.41 | 7.0E-03 | 1.3E-01 | 3.53 | ↑ | 1.82 |
| NEG | 263.1033_1.9 | L-Glutamine, N2-(2-phenylacetyl)- | 1.8E-03 | 4.5E-02 | 0.43 | ↓ | -1.23 | 5.0E-04 | 5.9E-02 | 0.36 | ↓ | -1.48 | 7.8E-03 | 7.5E-02 | 0.59 | ↓ | -0.76 |
| POS | 504.3088_1.14 | Lysophosphatidylethanolamine | 2.5E-03 | 8.9E-02 | 0.59 | ↓ | -0.77 | 4.6E-02 | 1.4E-01 | 0.75 | ↓ | -0.41 | 1.0E-05 | 4.9E-03 | 0.38 | ↓ | -1.39 |
| POS | 524.3717_1.89 | LysoPC(18:0) | 3.0E-02 | 2.8E-01 | 0.72 | ↓ | -0.48 | 6.3E-02 | 1.7E-01 | 0.79 | ↓ | -0.34 | 1.2E-01 | 3.3E-01 | 0.78 | ↓ | -0.36 |
| NEG | 830.5807_5.1 | PC(18:0/18:2) | 3.8E-01 | 5.4E-01 | 1.96 | ↑ | 0.97 | 2.2E-04 | 1.6E-03 | 3.05 | ↑ | 1.61 | 2.7E-01 | 3.8E-01 | 2.63 | ↑ | 1.39 |
| POS | 544.338_1.41 | LysoPC(18:1) | 1.8E-02 | 2.2E-01 | 0.66 | ↓ | -0.60 | 6.2E-01 | 8.0E-01 | 0.96 | ↓ | -0.06 | 2.9E-02 | 2.1E-01 | 0.63 | ↓ | -0.66 |
| POS | 214.1134_3.03 | Suberylglycine | 3.8E-02 | 2.3E-01 | 1.29 | ↑ | 0.36 | 1.1E-10 | 2.9E-07 | 10.57 | ↑ | 3.40 | 2.3E-10 | 6.1E-07 | 8.05 | ↑ | 3.01 |
| NEG | 768.5551_5.11 | PC(18:0/18:2) | 2.5E-01 | 4.2E-01 | 1.80 | ↑ | 0.85 | 9.0E-05 | 1.2E-03 | 3.02 | ↑ | 1.59 | 2.9E-01 | 4.0E-01 | 2.19 | ↑ | 1.13 |
| POS | 130.0499_1.9 | Glutamic Acid | 1.5E-03 | 4.9E-02 | 0.47 | ↓ | -1.08 | 4.7E-04 | 3.0E-02 | 0.35 | ↓ | -1.51 | 9.6E-03 | 8.5E-02 | 0.57 | ↓ | -0.80 |
| POS | 490.2926_0.99 | LysoPC(14:0) | 2.1E-03 | 8.2E-02 | 0.46 | ↓ | -1.12 | 1.4E-01 | 3.1E-01 | 0.72 | ↓ | -0.47 | 5.2E-05 | 1.9E-02 | 0.23 | ↓ | -2.11 |
| POS | 788.6068_5.47 | PC(18:0/18:1) | 1.7E-01 | 4.8E-01 | 3.55 | ↑ | 1.83 | 1.4E-03 | 1.5E-02 | 2.98 | ↑ | 1.57 | 1.5E-01 | 3.6E-01 | 3.10 | ↑ | 1.63 |
| POS | 287.1015_0.61 | Phenylacetyl-L-glutamine | 3.7E-03 | 1.2E-01 | 0.44 | ↓ | -1.20 | 1.6E-03 | 1.6E-02 | 0.39 | ↓ | -1.36 | 1.2E-02 | 1.5E-01 | 0.51 | ↓ | -0.97 |
| POS | 786.6003_5.47 | 1,2-Dioleoyl-sn-glycero-3-PC | 1.6E-01 | 4.7E-01 | 3.72 | ↑ | 1.89 | 1.4E-03 | 1.6E-02 | 2.93 | ↑ | 1.55 | 1.3E-01 | 3.5E-01 | 3.23 | ↑ | 1.69 |
| POS | 532.3389_1.56 | LysoPC(17:0) | 1.1E-05 | 7.2E-03 | 0.36 | ↓ | -1.48 | 9.9E-03 | 4.9E-02 | 0.58 | ↓ | -0.78 | 5.5E-06 | 4.1E-03 | 0.26 | ↓ | -1.94 |
| POS | 794.5989_5.26 | C18(Plasm)-20:4 PC | 1.6E-01 | 4.8E-01 | 5.90 | ↑ | 2.56 | 1.8E-03 | 1.7E-02 | 3.49 | ↑ | 1.80 | 1.2E-01 | 3.3E-01 | 4.37 | ↑ | 2.13 |
| POS | 421.2528_5.75 | Tris(2-butoxyethyl) phosphate | 2.8E-02 | 2.0E-01 | 2.13 | ↑ | 1.09 | 1.9E-03 | 5.8E-02 | 2.79 | ↑ | 1.48 | 8.1E-03 | 8.1E-02 | 2.23 | ↑ | 1.16 |
| POS | 180.0972_0.33 | N-Benzylglycine methyl ester | 1.8E-02 | 1.5E-01 | 0.48 | ↓ | -1.05 | 3.6E-01 | 6.3E-01 | 0.76 | ↓ | -0.39 | 1.1E-03 | 3.1E-02 | 0.37 | ↓ | -1.44 |
| NEG | 826.5604_4.96 | PC(16:0/20:4) | 2.6E-02 | 1.5E-01 | 3.26 | ↑ | 1.70 | 1.1E-04 | 1.2E-03 | 3.53 | ↑ | 1.82 | 1.3E-02 | 1.0E-01 | 4.21 | ↑ | 2.07 |
| POS | 637.3064_1.8 | Acetylpodocarpic acid anhydride | 8.5E-02 | 4.2E-01 | 0.77 | ↓ | -0.38 | 1.7E-02 | 6.8E-02 | 1.41 | ↑ | 0.50 | 1.4E-02 | 1.5E-01 | 1.52 | ↑ | 0.60 |

| | | | | | | | | | | | | | | | | | |
|---|---|---|---|---|---|---|---|---|---|---|---|---|---|---|---|---|---|
| POS | 808.5841_5 | 1,2-Dioleoyl-sn-glycero-3-PC | 8.7E-02 | 4.2E-01 | 3.32 | ↑ | 1.73 | 1.9E-04 | 9.4E-03 | 2.50 | ↑ | 1.32 | 5.7E-02 | 2.6E-01 | 3.44 | ↑ | 1.78 |
| NEG | 766.5393_4.96 | PE(18:0/20:4) | 8.7E-02 | 2.2E-01 | 2.72 | ↑ | 1.45 | 6.6E-04 | 2.9E-03 | 3.11 | ↑ | 1.64 | 1.7E-02 | 1.1E-01 | 3.91 | ↑ | 1.97 |
| NEG | 835.5345_5.54 | PI(16:0/18:1) | 1.3E-02 | 1.3E-01 | 5.10 | ↑ | 2.35 | 2.6E-03 | 8.7E-03 | 4.62 | ↑ | 2.21 | 5.8E-02 | 1.6E-01 | 5.01 | ↑ | 2.33 |
| NEG | 850.5605_4.85 | PC(16;0/22:6) | 6.0E-03 | 1.0E-01 | 3.58 | ↑ | 1.84 | 5.6E-05 | 1.2E-03 | 3.79 | ↑ | 1.92 | 6.7E-03 | 9.6E-02 | 4.67 | ↑ | 2.22 |
| POS | 808.5769_4.83 | 1,2-Dioleoyl-sn-glycero-3-PC | 9.2E-02 | 4.2E-01 | 2.51 | ↑ | 1.33 | 3.0E-03 | 2.3E-02 | 2.25 | ↑ | 1.17 | 1.4E-02 | 1.5E-01 | 2.72 | ↑ | 1.45 |
| POS | 502.3262_1.49 | C16(Plasm) LPC | 1.8E-03 | 7.9E-02 | 0.48 | ↓ | -1.07 | 1.4E-01 | 3.1E-01 | 0.75 | ↓ | -0.41 | 4.4E-03 | 1.0E-01 | 0.50 | ↓ | -1.00 |
| POS | 678.4682_6.87 | PC(14:0) | 7.7E-01 | 9.0E-01 | 0.97 | ↓ | -0.05 | 5.0E-03 | 9.7E-02 | 0.34 | ↓ | -1.54 | 1.7E-03 | 3.9E-02 | 0.27 | ↓ | -1.87 |
| POS | 546.3536_1.89 | LysoPC(18:0) | 1.8E-02 | 2.2E-01 | 0.76 | ↓ | -0.40 | 1.4E-01 | 3.0E-01 | 0.86 | ↓ | -0.21 | 1.8E-01 | 3.9E-01 | 0.85 | ↓ | -0.24 |
| NEG | 770.571_5.48 | DMPE(18:1/18:1) | 8.8E-02 | 2.2E-01 | 2.63 | ↑ | 1.40 | 1.1E-04 | 1.2E-03 | 4.35 | ↑ | 2.12 | 6.3E-02 | 1.6E-01 | 3.60 | ↑ | 1.85 |
| NEG | 832.5981_5.48 | PC(18:1/18:0) | 4.8E-02 | 1.8E-01 | 3.06 | ↑ | 1.61 | 5.1E-05 | 1.2E-03 | 4.72 | ↑ | 2.24 | 6.3E-02 | 1.6E-01 | 3.97 | ↑ | 1.99 |
| POS | 792.5895_5.18 | Dihomo-γ-linolenoyl PAF C-16 | 1.5E-01 | 4.7E-01 | 7.35 | ↑ | 2.88 | 9.8E-04 | 1.3E-02 | 3.02 | ↑ | 1.60 | 6.0E-02 | 2.7E-01 | 5.30 | ↑ | 2.40 |
| NEG | 794.5685_5.41 | PC(18:1/16:0) | 4.2E-02 | 1.6E-01 | 3.89 | ↑ | 1.96 | 1.3E-04 | 1.3E-03 | 3.71 | ↑ | 1.89 | 2.0E-02 | 1.2E-01 | 4.74 | ↑ | 2.24 |
| POS | 750.5453_5.35 | Palmitoyl Thio-PC | 9.0E-02 | 4.2E-01 | 8.57 | ↑ | 3.10 | 2.7E-03 | 2.2E-02 | 3.08 | ↑ | 1.62 | 8.1E-02 | 2.9E-01 | 6.96 | ↑ | 2.80 |
| POS | 810.5994_5.42 | PC(18:0/18:1) | 8.3E-02 | 4.2E-01 | 5.75 | ↑ | 2.52 | 4.9E-03 | 3.1E-02 | 2.89 | ↑ | 1.53 | 4.4E-02 | 2.4E-01 | 5.31 | ↑ | 2.41 |
| NEG | 832.6078_5.77 | PC(18:1/18:0) | 3.6E-02 | 1.6E-01 | 4.73 | ↑ | 2.24 | 8.8E-05 | 1.2E-03 | 7.67 | ↑ | 2.94 | 5.0E-02 | 1.5E-01 | 5.33 | ↑ | 2.41 |
| NEG | 830.5917_5.48 | PC(18:1/18:1) | 6.0E-02 | 1.9E-01 | 2.54 | ↑ | 1.34 | 4.2E-05 | 1.2E-03 | 3.88 | ↑ | 1.96 | 5.9E-02 | 1.6E-01 | 3.17 | ↑ | 1.67 |
| NEG | 747.5162_5.21 | SM(d18:1/16:0) | 1.7E-02 | 1.3E-01 | 6.14 | ↑ | 2.62 | 8.9E-05 | 1.2E-03 | 6.90 | ↑ | 2.79 | 7.4E-03 | 9.6E-02 | 9.70 | ↑ | 3.28 |
| NEG | 762.5082_4.98 | PE(16:0/22:6) | 1.1E-02 | 1.3E-01 | 7.78 | ↑ | 2.96 | 1.7E-02 | 4.3E-02 | 5.99 | ↑ | 2.58 | 4.0E-02 | 1.4E-01 | 7.49 | ↑ | 2.90 |
| POS | 790.5725_5.24 | Arachidonoyl PAF C-16 | 1.6E-01 | 4.7E-01 | 6.44 | ↑ | 2.69 | 7.9E-04 | 1.2E-02 | 4.53 | ↑ | 2.18 | 1.0E-01 | 3.1E-01 | 5.21 | ↑ | 2.38 |
| NEG | 765.5276_5.16 | SM(d18:1/18:0) | 3.3E-02 | 1.6E-01 | 3.29 | ↑ | 1.72 | 1.5E-02 | 3.9E-02 | 3.01 | ↑ | 1.59 | 1.1E-01 | 2.1E-01 | 3.73 | ↑ | 1.90 |
| NEG | 820.577_5.49 | PC(18:0/18:2) | 4.0E-02 | 1.6E-01 | 2.50 | ↑ | 1.32 | 4.0E-04 | 2.2E-03 | 3.16 | ↑ | 1.66 | 2.1E-02 | 1.2E-01 | 3.21 | ↑ | 1.68 |
| NEG | 790.5397_5.44 | PE(18:0/22:6) | 8.8E-03 | 1.2E-01 | 8.42 | ↑ | 3.07 | 6.4E-03 | 1.8E-02 | 7.15 | ↑ | 2.84 | 3.1E-02 | 1.3E-01 | 7.87 | ↑ | 2.98 |
| NEG | 806.5944_5.72 | PC(16:0/18:0) | 2.1E-02 | 1.4E-01 | 4.95 | ↑ | 2.31 | 5.0E-05 | 1.2E-03 | 6.43 | ↑ | 2.68 | 1.2E-02 | 1.0E-01 | 4.26 | ↑ | 2.09 |
| NEG | 790.541_4.84 | PE(18:0/22:6) | 6.2E-03 | 1.0E-01 | 3.21 | ↑ | 1.68 | 2.8E-05 | 1.2E-03 | 3.59 | ↑ | 1.84 | 4.4E-03 | 7.8E-02 | 4.63 | ↑ | 2.21 |
| NEG | 878.5916_5.31 | PC(18:0/22:6) | 3.9E-02 | 1.6E-01 | 3.64 | ↑ | 1.86 | 9.4E-05 | 1.2E-03 | 4.58 | ↑ | 2.20 | 1.7E-02 | 1.1E-01 | 5.22 | ↑ | 2.38 |
| NEG | 766.5395_5.55 | PE(18:0/24:0) | 1.3E-02 | 1.3E-01 | 7.87 | ↑ | 2.98 | 3.5E-03 | 1.1E-02 | 7.10 | ↑ | 2.83 | 3.5E-02 | 1.3E-01 | 7.91 | ↑ | 2.98 |
| POS | 766.5724_5.16 | Eicosapentaenoyl PAF C-16 | 2.2E-01 | 5.1E-01 | 8.35 | ↑ | 3.06 | 1.2E-02 | 5.5E-02 | 3.12 | ↑ | 1.64 | 9.8E-02 | 3.1E-01 | 6.17 | ↑ | 2.63 |
| POS | 760.5848_5.35 | PC(16:0/18:1) | 9.9E-02 | 4.3E-01 | 6.11 | ↑ | 2.61 | 4.0E-04 | 1.0E-02 | 3.74 | ↑ | 1.90 | 1.0E-01 | 3.1E-01 | 4.24 | ↑ | 2.08 |
| NEG | 776.5592_5.71 | PC(18:1/14:0) | 6.6E-03 | 1.1E-01 | 6.11 | ↑ | 2.61 | 1.8E-05 | 1.2E-03 | 6.91 | ↑ | 2.79 | 1.6E-02 | 1.0E-01 | 8.19 | ↑ | 3.03 |
| NEG | 768.5463_5.54 | PC(18:0/14:0) | 9.3E-03 | 1.2E-01 | 5.61 | ↑ | 2.49 | 2.4E-03 | 8.0E-03 | 5.48 | ↑ | 2.46 | 3.3E-02 | 1.3E-01 | 5.81 | ↑ | 2.54 |
| POS | 782.567_5.35 | PC(16:0/18:1) | 8.3E-02 | 4.2E-01 | 2.78 | ↑ | 1.47 | 1.6E-04 | 8.5E-03 | 3.35 | ↑ | 1.75 | 1.5E-01 | 3.6E-01 | 2.36 | ↑ | 1.24 |
| NEG | 838.5972_5.6 | C18(Plasm)-20:4 PC | 5.8E-02 | 1.9E-01 | 4.99 | ↑ | 2.32 | 4.8E-04 | 2.4E-03 | 5.18 | ↑ | 2.37 | 3.7E-02 | 1.3E-01 | 6.50 | ↑ | 2.70 |
| NEG | 804.5669_5.01 | PC(18:1/16:0) | 2.9E-02 | 1.6E-01 | 2.47 | ↑ | 1.30 | 4.5E-05 | 1.2E-03 | 3.64 | ↑ | 1.86 | 4.6E-02 | 1.4E-01 | 2.83 | ↑ | 1.50 |
| NEG | 802.5603_5.01 | PC(16:0/18:2) | 3.2E-02 | 1.6E-01 | 2.09 | ↑ | 1.06 | 8.4E-05 | 1.2E-03 | 2.93 | ↑ | 1.55 | 5.3E-02 | 1.5E-01 | 2.37 | ↑ | 1.24 |
| NEG | 744.549_5.01 | α-Galactosyl-N-palmitoylsphingosine | 2.1E-02 | 1.4E-01 | 2.94 | ↑ | 1.55 | 3.8E-04 | 2.1E-03 | 3.74 | ↑ | 1.90 | 6.1E-02 | 1.6E-01 | 2.45 | ↑ | 1.29 |
| POS | 768.5893_5.23 | C16-18:1 PC | 1.6E-01 | 4.8E-01 | 8.68 | ↑ | 3.12 | 1.9E-03 | 1.7E-02 | 3.57 | ↑ | 1.83 | 8.7E-02 | 3.0E-01 | 5.66 | ↑ | 2.50 |
| NEG | 774.545_5.65 | C18(Plasm)-22:6 PE | 1.4E-02 | 1.3E-01 | 5.96 | ↑ | 2.57 | 1.7E-05 | 1.2E-03 | 7.82 | ↑ | 2.97 | 8.6E-03 | 9.6E-02 | 8.55 | ↑ | 3.10 |
| NEG | 714.5081_5.16 | PE(16:0/18:2) | 2.8E-02 | 1.5E-01 | 4.47 | ↑ | 2.16 | 1.1E-02 | 2.9E-02 | 5.41 | ↑ | 2.44 | 1.1E-01 | 2.1E-01 | 3.30 | ↑ | 1.72 |
| NEG | 792.5473_5.01 | PC(16:0/18:2) | 7.2E-02 | 2.1E-01 | 2.03 | ↑ | 1.02 | 3.6E-04 | 2.1E-03 | 2.66 | ↑ | 1.41 | 3.6E-02 | 1.3E-01 | 2.53 | ↑ | 1.34 |
| POS | 735.5737_5.29 | Thioetheramidephosphatidylcholine | 3.6E-02 | 3.0E-01 | 12.79 | ↑ | 3.68 | 1.2E-04 | 8.5E-03 | 4.28 | ↑ | 2.10 | 1.0E-01 | 3.2E-01 | 6.08 | ↑ | 2.61 |
| POS | 756.5531_5.29 | PC(16:0/16:0) | 3.2E-02 | 2.9E-01 | 6.38 | ↑ | 2.67 | 1.9E-05 | 8.5E-03 | 4.26 | ↑ | 2.09 | 8.5E-02 | 2.9E-01 | 3.66 | ↑ | 1.87 |
| POS | 184.0734_5 | PC | 1.1E-01 | 4.3E-01 | 4.02 | ↑ | 2.01 | 3.5E-04 | 1.0E-02 | 3.09 | ↑ | 1.63 | 8.2E-02 | 2.9E-01 | 3.36 | ↑ | 1.75 |
| NEG | 806.5831_5.35 | PC(16:0/18:0) | 3.4E-02 | 1.6E-01 | 4.17 | ↑ | 2.06 | 1.2E-04 | 1.3E-03 | 6.22 | ↑ | 2.64 | 5.3E-02 | 1.5E-01 | 4.43 | ↑ | 2.15 |

| | | | | | | | | | | | | | | | | | |
|---|---|---|---|---|---|---|---|---|---|---|---|---|---|---|---|---|---|
| POS | 834.5998_5.31 | 1-Docosahexaenoyl-2-stearoyl-sn-glycero-3-p | 4.8E-02 | 3.4E-01 | 5.44 | ↑ | 2.44 | 8.5E-04 | 1.2E-02 | 3.52 | ↑ | 1.82 | 1.7E-02 | 1.6E-01 | 5.61 | ↑ | 2.49 |
| NEG | 834.5429_5.53 | PS(18:0/22:6) | 1.8E-02 | 1.3E-01 | 5.28 | ↑ | 2.40 | 3.6E-03 | 1.1E-02 | 4.41 | ↑ | 2.14 | 9.9E-02 | 2.0E-01 | 4.72 | ↑ | 2.24 |
| NEG | 742.5398_5.01 | PE(18:0/18:2) | 3.7E-02 | 1.6E-01 | 2.22 | ↑ | 1.15 | 9.0E-05 | 1.2E-03 | 3.39 | ↑ | 1.76 | 1.2E-01 | 2.2E-01 | 2.34 | ↑ | 1.22 |
| NEG | 862.5958_5.51 | C18(Plasm)-22:6 PC | 3.1E-02 | 1.6E-01 | 3.65 | ↑ | 1.87 | 2.0E-04 | 1.5E-03 | 3.66 | ↑ | 1.87 | 5.1E-02 | 1.5E-01 | 4.41 | ↑ | 2.14 |
| POS | 758.5692_5.01 | PC(16:0/18:2) | 1.2E-01 | 4.4E-01 | 3.34 | ↑ | 1.74 | 2.4E-04 | 9.7E-03 | 2.86 | ↑ | 1.52 | 6.6E-02 | 2.8E-01 | 2.89 | ↑ | 1.53 |
| POS | 760.5746_5.01 | PC(16:0/18:1) | 1.1E-01 | 4.3E-01 | 3.07 | ↑ | 1.62 | 2.2E-04 | 9.6E-03 | 2.76 | ↑ | 1.46 | 6.6E-02 | 2.8E-01 | 2.69 | ↑ | 1.43 |
| NEG | 804.576_5.36 | PC(18:1/16:0) | 3.5E-02 | 1.6E-01 | 3.64 | ↑ | 1.86 | 5.5E-05 | 1.2E-03 | 5.38 | ↑ | 2.43 | 5.3E-02 | 1.5E-01 | 3.91 | ↑ | 1.97 |
| POS | 768.5582_5.55 | PE(18:0/18:1) | 6.1E-02 | 3.8E-01 | 13.03 | ↑ | 3.70 | 2.0E-02 | 7.9E-02 | 3.14 | ↑ | 1.65 | 1.5E-01 | 3.6E-01 | 6.80 | ↑ | 2.76 |
| NEG | 738.5085_5.1 | PE(16:0/20:4) | 4.8E-02 | 1.8E-01 | 4.50 | ↑ | 2.17 | 3.4E-02 | 7.8E-02 | 4.55 | ↑ | 2.18 | 1.0E-01 | 2.0E-01 | 4.22 | ↑ | 2.08 |
| NEG | 802.5603_8.81 | PC(16:0/18:2) | 5.6E-01 | 7.8E-01 | 1.02 | ↑ | 0.03 | 8.1E-03 | 1.7E-01 | 1.16 | ↑ | 0.21 | 3.1E-02 | 1.6E-01 | 1.12 | ↑ | 0.16 |

| | | | | | | | | | | | | | | | | | |
|---|---|---|---|---|---|---|---|---|---|---|---|---|---|---|---|---|---|
| | | | | | | | | | | 7.5 Gy | | | | | | | |
| | | | SD 1 vs. Pre | | | | | SD 2 vs. Pre | | | | | SD 6 vs. Pre | | | | |
| **Mode** | **m/z_RT** | **Name** | **p-value** | **FDR** | **FC** | | **LOG2FC** | **p-value** | **FDR** | **FC** | | **LOG2FC** | **p-value** | **FDR** | **FC** | | **LOG2FC** |
| POS | 162.1127_0.57 | L-Carnitine | 4.8E-04 | 3.6E-02 | 2.56 | ↑ | 1.36 | 1.0E-01 | 3.0E-01 | 1.52 | ↑ | 0.61 | 5.9E-03 | 4.6E-02 | 1.91 | ↑ | 0.93 |
| POS | 163.1155_0.56 | Phenylpiperazine | 8.0E-04 | 3.7E-02 | 2.53 | ↑ | 1.34 | 1.2E-01 | 3.3E-01 | 1.55 | ↑ | 0.63 | 3.5E-03 | 3.3E-02 | 2.02 | ↑ | 1.01 |
| NEG | 861.5497_12.17 | PI(18:1) | 7.2E-03 | 8.9E-02 | 0.62 | ↓ | -0.70 | 2.1E-01 | 7.0E-01 | 1.42 | ↑ | 0.50 | 1.1E-03 | 2.6E-02 | 0.50 | ↓ | -0.99 |
| POS | 580.432_7.28 | LysoPC(22:0) | 3.8E-05 | 1.7E-02 | 0.44 | ↓ | -1.19 | 7.2E-01 | 8.8E-01 | 1.00 | ↑ | 0.00 | 5.3E-03 | 4.2E-02 | 0.48 | ↓ | -1.04 |
| POS | 548.3722_1.53 | Lyso-PAF C-18 | 1.0E-04 | 2.3E-02 | 0.46 | ↓ | -1.13 | 2.6E-03 | 7.9E-02 | 0.58 | ↓ | -0.79 | 1.3E-05 | 2.6E-03 | 0.38 | ↓ | -1.39 |
| POS | 552.4031_6.62 | LysoPC(20:0) | 7.9E-05 | 1.7E-02 | 0.43 | ↓ | -1.21 | 2.4E-01 | 7.8E-01 | 0.82 | ↓ | -0.28 | 1.1E-02 | 6.5E-02 | 0.49 | ↓ | -1.02 |
| POS | 782.5682_8.51 | PC(16:0/18:1) | 3.1E-05 | 1.7E-02 | 0.29 | ↓ | -1.79 | 3.1E-02 | 5.8E-01 | 0.71 | ↓ | -0.48 | 8.0E-05 | 4.2E-03 | 0.30 | ↓ | -1.76 |
| NEG | 790.5343_8.7 | PI(18:0/22:6) | 4.7E-02 | 2.6E-01 | 0.73 | ↓ | -0.46 | 3.8E-01 | 7.0E-01 | 0.91 | ↓ | -0.13 | 1.4E-03 | 3.0E-02 | 0.62 | ↓ | -0.68 |
| POS | 510.3561_5.63 | LysoPC(17:0) | 1.6E-03 | 3.5E-02 | 0.57 | ↓ | -0.80 | 1.5E-02 | 4.4E-01 | 0.56 | ↓ | -0.84 | 3.3E-03 | 3.2E-02 | 0.49 | ↓ | -1.03 |
| POS | 339.3453_7.67 | 9-Oxo-11α,15R-dihydroxyprostan-1-oic acid | 5.5E-01 | 7.6E-01 | 0.82 | ↓ | -0.28 | 1.6E-01 | 7.8E-01 | 0.67 | ↓ | -0.57 | 4.6E-05 | 3.4E-03 | 0.17 | ↓ | -2.58 |
| POS | 338.3423_7.68 | 13-Docosenamide, (Z)- | 5.5E-01 | 7.6E-01 | 0.83 | ↓ | -0.27 | 1.8E-01 | 7.8E-01 | 0.70 | ↓ | -0.51 | 2.7E-05 | 2.7E-03 | 0.19 | ↓ | -2.42 |
| NEG | 554.3457_5.63 | LysoPC(17:0) | 3.9E-03 | 6.5E-02 | 0.61 | ↓ | -0.72 | 2.3E-02 | 6.9E-01 | 0.63 | ↓ | -0.66 | 7.9E-03 | 6.6E-02 | 0.54 | ↓ | -0.88 |
| POS | 608.4631_7.89 | LysoPC(24:0) | 5.5E-05 | 1.7E-02 | 0.50 | ↓ | -1.00 | 7.3E-02 | 7.8E-01 | 0.76 | ↓ | -0.39 | 6.5E-04 | 1.2E-02 | 0.51 | ↓ | -0.98 |
| NEG | 826.5599_8.52 | PC(16:0/20:4) | 2.9E-05 | 1.1E-02 | 0.58 | ↓ | -0.78 | 7.9E-01 | 8.9E-01 | 1.12 | ↑ | 0.16 | 1.3E-03 | 2.9E-02 | 0.60 | ↓ | -0.73 |
| NEG | 596.3925_6.63 | LysoPC(20:0) | 1.0E-03 | 3.6E-02 | 0.44 | ↓ | -1.19 | 3.3E-01 | 7.0E-01 | 0.83 | ↓ | -0.28 | 1.7E-02 | 1.0E-01 | 0.47 | ↓ | -1.09 |
| POS | 770.5677_8.69 | Dihomo-γ-linolenoyl PAF C-16 | 1.4E-03 | 3.4E-02 | 0.43 | ↓ | -1.20 | 7.4E-03 | 3.3E-01 | 0.61 | ↓ | -0.72 | 3.2E-05 | 2.9E-03 | 0.30 | ↓ | -1.75 |
| POS | 532.3383_5.63 | LysoPC(17:0) | 8.2E-04 | 2.9E-02 | 0.59 | ↓ | -0.75 | 1.4E-02 | 4.4E-01 | 0.68 | ↓ | -0.56 | 5.3E-03 | 4.2E-02 | 0.54 | ↓ | -0.89 |
| NEG | 861.5499_11.73 | PI(18:1) | 2.7E-02 | 2.0E-01 | 0.61 | ↓ | -0.71 | 6.2E-01 | 8.0E-01 | 0.90 | ↓ | -0.16 | 2.9E-03 | 4.0E-02 | 0.44 | ↓ | -1.18 |
| POS | 184.0945_0.58 | PC | 1.8E-04 | 2.6E-02 | 2.94 | ↑ | 1.55 | 2.6E-02 | 1.5E-01 | 2.02 | ↑ | 1.01 | 9.4E-04 | 1.8E-02 | 2.73 | ↑ | 1.45 |
| NEG | 788.5249_8.71 | PS(18:0/18:1) | 8.3E-03 | 9.5E-02 | 0.56 | ↓ | -0.83 | 3.4E-02 | 7.0E-01 | 0.69 | ↓ | -0.54 | 7.3E-03 | 6.3E-02 | 0.57 | ↓ | -0.81 |
| POS | 518.3256_4.86 | LysoPC(16:0) | 2.1E-02 | 1.2E-01 | 0.52 | ↓ | -0.94 | 1.1E-02 | 4.2E-01 | 0.47 | ↓ | -1.09 | 4.1E-04 | 9.8E-03 | 0.27 | ↓ | -1.87 |
| NEG | 828.5667_8.52 | C18(Plasm)-20:4 PC | 1.1E-04 | 1.4E-02 | 0.52 | ↓ | -0.95 | 7.9E-01 | 8.9E-01 | 1.16 | ↑ | 0.21 | 1.2E-02 | 8.6E-02 | 0.48 | ↓ | -1.07 |
| NEG | 827.5636_8.52 | KRTLRR | 1.1E-04 | 1.4E-02 | 0.56 | ↓ | -0.85 | 8.5E-01 | 9.2E-01 | 1.09 | ↑ | 0.13 | 4.4E-03 | 4.7E-02 | 0.53 | ↓ | -0.91 |
| NEG | 526.3113_5.06 | LysoPC(15:0) | 1.2E-02 | 1.2E-01 | 0.65 | ↓ | -0.62 | 6.2E-02 | 7.0E-01 | 0.73 | ↓ | -0.46 | 4.2E-03 | 4.7E-02 | 0.56 | ↓ | -0.83 |
| NEG | 594.3771_6.08 | Pyrrolidino PAF C-16 | 2.4E-02 | 1.9E-01 | 0.68 | ↓ | -0.56 | 5.9E-01 | 7.8E-01 | 0.94 | ↓ | -0.09 | 2.1E-02 | 1.2E-01 | 0.58 | ↓ | -0.78 |
| POS | 510.3564_1.56 | LysoPC(17:0) | 1.2E-04 | 2.3E-02 | 0.44 | ↓ | -1.20 | 9.5E-03 | 1.1E-01 | 0.60 | ↓ | -0.74 | 1.5E-05 | 2.6E-03 | 0.27 | ↓ | -1.90 |
| POS | 482.3248_5.06 | LysoPC(15:0) | 2.8E-03 | 4.5E-02 | 0.52 | ↓ | -0.94 | 2.2E-02 | 5.0E-01 | 0.57 | ↓ | -0.82 | 2.5E-04 | 7.4E-03 | 0.33 | ↓ | -1.59 |
| NEG | 554.3463_1.56 | LysoPC(17:0) | 4.2E-03 | 1.3E-02 | 0.55 | ↓ | -0.85 | 4.4E-03 | 1.5E-01 | 0.67 | ↓ | -0.57 | 3.2E-03 | 1.3E-01 | 0.51 | ↓ | -0.97 |
| POS | 552.4037_2.62 | LysoPC(20:0) | 2.7E-03 | 5.7E-02 | 0.49 | ↓ | -1.03 | 7.5E-01 | 8.7E-01 | 1.13 | ↑ | 0.18 | 5.7E-04 | 1.6E-02 | 0.40 | ↓ | -1.32 |
| NEG | 624.4238_7.29 | LysoPC(22:0) | 4.8E-06 | 9.6E-03 | 0.41 | ↓ | -1.27 | 9.3E-01 | 9.7E-01 | 1.09 | ↑ | 0.12 | 3.2E-02 | 1.5E-01 | 0.59 | ↓ | -0.76 |

| | | | | | | | | | | | | | | | | | |
|---|---|---|---|---|---|---|---|---|---|---|---|---|---|---|---|---|---|
| POS | 524.3722_5.96 | LysoPC(18:0) | 2.2E-02 | 1.3E-01 | 0.78 | ↓ | -0.36 | 2.1E-02 | 4.9E-01 | 0.76 | ↓ | -0.40 | 5.3E-02 | 1.7E-01 | 0.78 | ↓ | -0.36 |
| NEG | 512.2985_4.81 | LysoPC(14:0) | 4.5E-03 | 6.9E-02 | 0.64 | ↓ | -0.64 | 5.5E-01 | 7.6E-01 | 0.93 | ↓ | -0.10 | 6.4E-03 | 6.0E-02 | 0.49 | ↓ | -1.02 |
| NEG | 652.4552_7.9 | LysoPC(24:0) | 3.2E-04 | 2.4E-02 | 0.55 | ↓ | -0.86 | 2.5E-01 | 7.0E-01 | 0.86 | ↓ | -0.23 | 3.4E-03 | 4.2E-02 | 0.52 | ↓ | -0.95 |
| NEG | 226.0175_2.4 | 4-Quinolinol, 8-methyl-2-(trifluoromethyl)- | 5.5E-03 | 7.8E-02 | 0.35 | ↓ | -1.52 | 5.4E-04 | 9.8E-02 | 0.18 | ↓ | -2.51 | 7.0E-04 | 2.1E-02 | 0.11 | ↓ | -3.19 |
| POS | 516.3104_4.97 | Oleyloxyethylphosphorylcholine | 2.0E-03 | 3.9E-02 | 0.55 | ↓ | -0.86 | 5.9E-02 | 7.8E-01 | 0.67 | ↓ | -0.57 | 2.4E-03 | 2.8E-02 | 0.49 | ↓ | -1.04 |
| POS | 162.1129_0.33 | L-Carnitine | 1.0E-04 | 1.7E-02 | 2.53 | ↑ | 1.34 | 4.8E-02 | 7.3E-01 | 2.26 | ↑ | 1.18 | 7.7E-02 | 2.1E-01 | 1.73 | ↑ | 0.79 |
| POS | 508.3397_5.21 | C18(Plasm) LPC | 2.7E-03 | 4.5E-02 | 0.53 | ↓ | -0.93 | 1.7E-02 | 4.5E-01 | 0.59 | ↓ | -0.77 | 8.5E-04 | 1.4E-02 | 0.41 | ↓ | -1.28 |
| NEG | 512.299_0.99 | LysoPC(14:0) | 3.8E-03 | 1.2E-02 | 0.57 | ↓ | -0.81 | 9.0E-01 | 9.4E-01 | 0.99 | ↓ | -0.02 | 1.1E-02 | 1.6E-01 | 0.48 | ↓ | -1.05 |
| NEG | 816.5759_9.01 | PC(16:0/20:4) | 1.6E-02 | 1.5E-01 | 0.61 | ↓ | -0.70 | 3.6E-01 | 7.0E-01 | 0.85 | ↓ | -0.23 | 1.1E-03 | 2.6E-02 | 0.44 | ↓ | -1.20 |
| POS | 204.1231_0.58 | Acetyl-DL-carnitine | 3.5E-02 | 1.2E-01 | 1.65 | ↑ | 0.72 | 9.5E-01 | 9.8E-01 | 0.99 | ↓ | -0.02 | 3.2E-03 | 3.1E-02 | 4.12 | ↑ | 2.04 |
| POS | 482.3253_1.13 | LysoPE(18:0) | 6.6E-04 | 3.6E-02 | 0.44 | ↓ | -1.18 | 4.8E-02 | 2.0E-01 | 0.66 | ↓ | -0.59 | 1.6E-05 | 2.6E-03 | 0.23 | ↓ | -2.09 |
| POS | 468.3092_4.81 | LysoPC(14:0) | 1.1E-03 | 3.3E-02 | 0.58 | ↓ | -0.79 | 2.7E-01 | 7.8E-01 | 0.84 | ↓ | -0.24 | 6.6E-03 | 4.9E-02 | 0.47 | ↓ | -1.10 |
| POS | 782.5682_4.96 | PC(16:0/18:1) | 1.9E-02 | 1.0E-01 | 3.24 | ↑ | 1.70 | 3.9E-03 | 8.6E-02 | 1.84 | ↑ | 0.88 | 6.1E-03 | 4.6E-02 | 4.79 | ↑ | 2.26 |
| POS | 530.368_6.36 | C18(Plasm) LPC | 2.4E-02 | 1.3E-01 | 3.66 | ↑ | 1.87 | 4.9E-03 | 2.7E-01 | 4.76 | ↑ | 2.25 | 1.2E-02 | 7.0E-02 | 4.83 | ↑ | 2.27 |
| POS | 530.3694_1.4 | C18(Plasm) LPC | 6.9E-02 | 1.7E-01 | 3.83 | ↑ | 1.94 | 8.5E-03 | 1.1E-01 | 5.11 | ↑ | 2.35 | 4.8E-02 | 1.5E-01 | 3.34 | ↑ | 1.74 |
| NEG | 526.31_1.14 | LysoPC(15:0) | 3.3E-03 | 1.1E-02 | 0.57 | ↓ | -0.82 | 2.3E-02 | 2.0E-01 | 0.68 | ↓ | -0.56 | 8.3E-04 | 5.2E-02 | 0.40 | ↓ | -1.33 |
| NEG | 229.1437_3.65 | Dodecanedioic acid | 3.8E-01 | 6.8E-01 | 0.78 | ↓ | -0.37 | 8.3E-01 | 9.1E-01 | 1.61 | ↑ | 0.68 | 1.2E-01 | 3.1E-01 | 0.63 | ↓ | -0.67 |
| POS | 790.6208_10.03 | PS(18:0/18:1) | 2.6E-02 | 1.4E-01 | 0.78 | ↓ | -0.36 | 1.6E-01 | 7.8E-01 | 0.75 | ↓ | -0.41 | 8.4E-04 | 1.4E-02 | 0.57 | ↓ | -0.80 |
| POS | 204.1141_0.33 | Acetyl-DL-carnitine | 2.9E-03 | 4.5E-02 | 1.56 | ↑ | 0.64 | 3.6E-01 | 7.8E-01 | 1.24 | ↑ | 0.31 | 2.0E-03 | 2.5E-02 | 2.80 | ↑ | 1.49 |
| POS | 316.2485_3.86 | Decanoyl-L-carnitine | 3.3E-02 | 1.6E-01 | 3.57 | ↑ | 1.83 | 1.7E-01 | 7.8E-01 | 1.38 | ↑ | 0.46 | 1.3E-02 | 7.4E-02 | 6.24 | ↑ | 2.64 |
| POS | 468.3099_0.99 | LysoPC(14:0) | 1.9E-03 | 5.0E-02 | 0.53 | ↓ | -0.91 | 6.0E-01 | 7.8E-01 | 0.93 | ↓ | -0.11 | 8.4E-04 | 1.8E-02 | 0.36 | ↓ | -1.46 |
| NEG | 584.3554_4.45 | 2-Thio PAF | 2.2E-03 | 5.1E-02 | 0.58 | ↓ | -0.77 | 3.0E-01 | 7.0E-01 | 0.77 | ↓ | -0.38 | 3.0E-02 | 1.5E-01 | 0.54 | ↓ | -0.88 |
| NEG | 178.0488_0.62 | Hippuric acid | 2.0E-01 | 3.4E-01 | 0.66 | ↓ | -0.59 | 3.7E-03 | 1.5E-01 | 0.32 | ↓ | -1.63 | 2.3E-03 | 1.1E-01 | 0.22 | ↓ | -2.17 |
| POS | 728.5584_10.1 | PE(18:0/18:1) | 1.2E-03 | 3.3E-02 | 0.54 | ↓ | -0.89 | 2.4E-01 | 7.8E-01 | 0.80 | ↓ | -0.32 | 1.6E-05 | 2.1E-03 | 0.26 | ↓ | -1.96 |
| NEG | 452.2775_0.99 | LysoPE(16:0) | 3.4E-03 | 1.1E-02 | 0.62 | ↓ | -0.70 | 5.2E-01 | 7.1E-01 | 1.11 | ↑ | 0.15 | 8.1E-03 | 1.6E-01 | 0.47 | ↓ | -1.08 |
| POS | 257.2267_3.76 | Androst-16-en-3-ol, (3β,5α)- | 7.4E-03 | 7.3E-02 | 2.14 | ↑ | 1.10 | 4.2E-02 | 6.8E-01 | 1.69 | ↑ | 0.76 | 1.2E-01 | 2.6E-01 | 1.42 | ↑ | 0.50 |
| NEG | 862.5542_4.81 | C18(Plasm)-22:6 PC | 8.8E-04 | 4.2E-03 | 4.33 | ↑ | 2.12 | 3.5E-02 | 2.2E-01 | 6.16 | ↑ | 2.62 | 1.8E-01 | 3.3E-01 | 2.44 | ↑ | 1.29 |
| POS | 806.5689_4.84 | Arachidonoylthio-PC | 8.6E-03 | 8.4E-02 | 3.14 | ↑ | 1.65 | 3.7E-04 | 3.6E-02 | 2.08 | ↑ | 1.06 | 1.4E-02 | 7.6E-02 | 4.61 | ↑ | 2.20 |
| NEG | 263.1033_1.9 | L-Glutamine, N2-(2-phenylacetyl)- | 2.9E-03 | 5.9E-02 | 0.34 | ↓ | -1.58 | 4.5E-03 | 3.0E-01 | 0.37 | ↓ | -1.42 | 3.3E-03 | 4.1E-02 | 0.24 | ↓ | -2.04 |
| POS | 504.3088_1.14 | Lysophosphatidylethanolamine | 2.2E-03 | 5.4E-02 | 0.54 | ↓ | -0.90 | 1.3E-02 | 1.1E-01 | 0.61 | ↓ | -0.72 | 8.4E-06 | 2.6E-03 | 0.28 | ↓ | -1.81 |
| POS | 524.3717_1.89 | LysoPC(18:0) | 6.1E-04 | 3.6E-02 | 0.59 | ↓ | -0.76 | 1.4E-01 | 3.5E-01 | 0.83 | ↓ | -0.27 | 7.9E-04 | 1.8E-02 | 0.54 | ↓ | -0.90 |
| NEG | 830.5807_5.1 | PC(18:0/18:2) | 2.9E-03 | 9.8E-03 | 3.35 | ↑ | 1.74 | 1.6E-02 | 1.8E-01 | 2.57 | ↑ | 1.36 | 1.0E-01 | 2.8E-01 | 2.05 | ↑ | 1.03 |
| POS | 544.338_1.41 | LysoPC(18:1) | 5.3E-03 | 7.5E-02 | 0.63 | ↓ | -0.67 | 8.1E-01 | 9.1E-01 | 0.99 | ↓ | -0.01 | 5.8E-03 | 4.6E-02 | 0.54 | ↓ | -0.88 |
| POS | 214.1134_3.03 | Suberylglycine | 1.5E-01 | 3.9E-01 | 1.24 | ↑ | 0.31 | 3.4E-08 | 4.5E-05 | 11.59 | ↑ | 3.54 | 2.8E-10 | 3.7E-07 | 7.98 | ↑ | 3.00 |
| NEG | 768.5551_5.11 | PC(18:0/18:2) | 2.0E-03 | 7.3E-03 | 3.00 | ↑ | 1.59 | 1.6E-02 | 1.8E-01 | 2.22 | ↑ | 1.15 | 1.0E-01 | 2.8E-01 | 1.86 | ↑ | 0.89 |
| POS | 130.0499_1.9 | Glutamic Acid | 4.9E-03 | 5.7E-02 | 0.33 | ↓ | -1.61 | 1.4E-02 | 4.4E-01 | 0.40 | ↓ | -1.31 | 1.8E-03 | 2.4E-02 | 0.21 | ↓ | -2.23 |
| POS | 490.2926_0.99 | LysoPC(14:0) | 2.6E-02 | 1.1E-01 | 0.53 | ↓ | -0.93 | 2.2E-01 | 4.5E-01 | 0.72 | ↓ | -0.48 | 1.5E-03 | 2.0E-02 | 0.27 | ↓ | -1.87 |
| POS | 788.6068_5.47 | PC(18:0/18:1) | 9.4E-02 | 2.0E-01 | 2.80 | ↑ | 1.49 | 1.4E-02 | 1.2E-01 | 1.88 | ↑ | 0.91 | 1.3E-01 | 2.5E-01 | 2.31 | ↑ | 1.21 |
| POS | 287.1015_0.61 | Phenylacetyl-L-glutamine | 9.7E-03 | 8.6E-02 | 0.42 | ↓ | -1.25 | 4.3E-03 | 8.8E-02 | 0.38 | ↓ | -1.38 | 3.2E-03 | 3.1E-02 | 0.32 | ↓ | -1.65 |
| POS | 786.6003_5.47 | 1,2-Dioleoyl-sn-glycero-3-PC | 8.7E-02 | 1.9E-01 | 2.88 | ↑ | 1.53 | 1.3E-02 | 1.1E-01 | 1.85 | ↑ | 0.89 | 1.1E-01 | 2.3E-01 | 2.38 | ↑ | 1.25 |
| POS | 532.3389_1.56 | LysoPC(17:0) | 6.7E-05 | 2.3E-02 | 0.28 | ↓ | -1.85 | 4.0E-03 | 8.6E-02 | 0.45 | ↓ | -1.17 | 3.8E-06 | 2.6E-03 | 0.17 | ↓ | -2.57 |
| POS | 794.5989_5.26 | C18(Plasm)-20:4 PC | 9.1E-02 | 1.9E-01 | 3.10 | ↑ | 1.63 | 1.1E-02 | 1.1E-01 | 2.38 | ↑ | 1.25 | 2.2E-03 | 2.5E-02 | 4.46 | ↑ | 2.16 |
| POS | 421.2528_5.75 | Tris(2-butoxyethyl) phosphate | 1.6E-02 | 1.1E-01 | 2.21 | ↑ | 1.14 | 1.6E-02 | 4.4E-01 | 5.88 | ↑ | 2.56 | 1.5E-03 | 2.0E-02 | 2.52 | ↑ | 1.34 |

| | | | | | | | | | | | | | | | | | |
|---|---|---|---|---|---|---|---|---|---|---|---|---|---|---|---|---|---|
| POS | 180.0972_0.33 | N-Benzylglycine methyl ester | 7.9E-02 | 2.6E-01 | 0.63 | ↓ | -0.67 | 6.2E-01 | 8.4E-01 | 0.88 | ↓ | -0.19 | 6.6E-03 | 4.9E-02 | 0.40 | ↓ | -1.33 |
| NEG | 826.5604_4.96 | PC(16:0/20:4) | 1.1E-04 | 2.3E-03 | 4.06 | ↑ | 2.02 | 1.6E-02 | 1.8E-01 | 2.25 | ↑ | 1.17 | 8.2E-03 | 1.6E-01 | 3.74 | ↑ | 1.90 |
| POS | 637.3064_1.8 | Acetylpodocarpic acid anhydride | 9.4E-02 | 2.0E-01 | 0.78 | ↓ | -0.36 | 3.2E-03 | 8.6E-02 | 1.77 | ↑ | 0.82 | 4.0E-04 | 1.5E-02 | 2.15 | ↑ | 1.10 |
| POS | 808.5841_5 | 1,2-Dioleoyl-sn-glycero-3-PC | 6.7E-02 | 1.7E-01 | 2.43 | ↑ | 1.28 | 6.9E-03 | 1.0E-01 | 2.06 | ↑ | 1.04 | 1.3E-03 | 2.0E-02 | 4.07 | ↑ | 2.02 |
| NEG | 766.5393_4.96 | PE(18:0/20:4) | 8.7E-05 | 2.3E-03 | 4.69 | ↑ | 2.23 | 1.4E-02 | 1.8E-01 | 2.64 | ↑ | 1.40 | 1.4E-02 | 1.7E-01 | 4.34 | ↑ | 2.12 |
| NEG | 835.5345_5.54 | PI(16:0/18:1) | 2.9E-04 | 2.5E-03 | 6.64 | ↑ | 2.73 | 1.5E-01 | 3.3E-01 | 2.76 | ↑ | 1.46 | 1.0E-01 | 2.8E-01 | 2.87 | ↑ | 1.52 |
| NEG | 850.5605_4.85 | PC(16;0/22:6) | 9.0E-04 | 4.3E-03 | 4.42 | ↑ | 2.14 | 9.9E-02 | 2.8E-01 | 2.48 | ↑ | 1.31 | 3.8E-02 | 2.2E-01 | 4.69 | ↑ | 2.23 |
| POS | 808.5769_4.83 | 1,2-Dioleoyl-sn-glycero-3-PC | 4.3E-02 | 1.4E-01 | 2.28 | ↑ | 1.19 | 1.8E-03 | 7.8E-02 | 1.77 | ↑ | 0.82 | 2.7E-02 | 1.1E-01 | 3.14 | ↑ | 1.65 |
| POS | 502.3262_1.49 | C16(Plasm) LPC | 3.2E-03 | 6.1E-02 | 0.49 | ↓ | -1.02 | 2.4E-01 | 4.6E-01 | 0.79 | ↓ | -0.34 | 1.2E-04 | 7.8E-03 | 0.35 | ↓ | -1.53 |
| POS | 678.4682_6.87 | PC(14:0) | 2.2E-02 | 1.3E-01 | 0.47 | ↓ | -1.08 | 2.3E-03 | 1.8E-01 | 0.27 | ↓ | -1.87 | 6.6E-05 | 3.9E-03 | 0.16 | ↓ | -2.63 |
| POS | 546.3536_1.89 | LysoPC(18:0) | 1.2E-03 | 4.2E-02 | 0.72 | ↓ | -0.47 | 3.6E-02 | 1.7E-01 | 0.86 | ↓ | -0.23 | 1.7E-02 | 8.6E-02 | 0.74 | ↓ | -0.43 |
| NEG | 770.571_5.48 | DMPE(18:1/18:1) | 1.7E-03 | 6.5E-03 | 3.83 | ↑ | 1.94 | 7.7E-02 | 2.7E-01 | 2.59 | ↑ | 1.37 | 1.1E-01 | 2.8E-01 | 2.75 | ↑ | 1.46 |
| NEG | 832.5981_5.48 | PC(18:1/18:0) | 2.1E-04 | 2.3E-03 | 4.18 | ↑ | 2.06 | 5.6E-02 | 2.5E-01 | 2.69 | ↑ | 1.43 | 1.1E-01 | 2.8E-01 | 2.73 | ↑ | 1.45 |
| POS | 792.5895_5.18 | Dihomo-γ-linolenoyl PAF C-16 | 3.7E-02 | 1.3E-01 | 4.55 | ↑ | 2.19 | 7.7E-03 | 1.0E-01 | 2.50 | ↑ | 1.32 | 2.7E-03 | 2.8E-02 | 7.14 | ↑ | 2.84 |
| NEG | 794.5685_5.41 | PC(18:1/16:0) | 9.7E-05 | 2.3E-03 | 5.09 | ↑ | 2.35 | 8.6E-02 | 2.7E-01 | 2.45 | ↑ | 1.30 | 1.6E-02 | 1.7E-01 | 4.23 | ↑ | 2.08 |
| POS | 750.5453_5.35 | Palmitoyl Thio-PC | 2.3E-02 | 1.1E-01 | 4.65 | ↑ | 2.22 | 1.5E-02 | 1.2E-01 | 2.51 | ↑ | 1.33 | 5.8E-03 | 4.6E-02 | 8.24 | ↑ | 3.04 |
| POS | 810.5994_5.42 | PC(18:0/18:1) | 1.0E-02 | 8.6E-02 | 3.78 | ↑ | 1.92 | 5.9E-03 | 9.8E-02 | 1.91 | ↑ | 0.93 | 9.7E-03 | 6.0E-02 | 4.89 | ↑ | 2.29 |
| NEG | 832.6078_5.77 | PC(18:1/18:0) | 3.9E-04 | 2.8E-03 | 5.80 | ↑ | 2.54 | 4.2E-02 | 2.3E-01 | 3.63 | ↑ | 1.86 | 1.1E-01 | 2.8E-01 | 2.90 | ↑ | 1.54 |
| NEG | 830.5917_5.48 | PC(18:1/18:1) | 3.0E-04 | 2.6E-03 | 3.45 | ↑ | 1.79 | 4.5E-02 | 2.3E-01 | 2.39 | ↑ | 1.26 | 8.9E-02 | 2.7E-01 | 2.32 | ↑ | 1.21 |
| NEG | 747.5162_5.21 | SM(d18:1/16:0) | 5.3E-05 | 2.3E-03 | 8.62 | ↑ | 3.11 | 8.0E-02 | 2.7E-01 | 6.59 | ↑ | 2.72 | 2.7E-02 | 2.1E-01 | 11.12 | ↑ | 3.47 |
| NEG | 762.5082_4.98 | PE(16:0/22:6) | 4.1E-04 | 2.8E-03 | 13.06 | ↑ | 3.71 | 4.0E-01 | 6.0E-01 | 4.29 | ↑ | 2.10 | 1.2E-01 | 2.8E-01 | 6.47 | ↑ | 2.69 |
| POS | 790.5725_5.24 | Arachidonoyl PAF C-16 | 3.3E-02 | 1.2E-01 | 4.62 | ↑ | 2.21 | 3.7E-03 | 8.6E-02 | 3.22 | ↑ | 1.69 | 4.6E-03 | 3.9E-02 | 4.65 | ↑ | 2.22 |
| NEG | 765.5276_5.16 | SM(d18:1/18:0) | 1.3E-03 | 5.3E-03 | 6.26 | ↑ | 2.65 | 1.8E-01 | 3.6E-01 | 3.09 | ↑ | 1.63 | 1.2E-01 | 2.8E-01 | 3.13 | ↑ | 1.65 |
| NEG | 820.577_5.49 | PC(18:0/18:2) | 1.9E-03 | 7.2E-03 | 3.06 | ↑ | 1.61 | 1.1E-01 | 2.9E-01 | 2.18 | ↑ | 1.13 | 9.6E-02 | 2.7E-01 | 2.75 | ↑ | 1.46 |
| NEG | 790.5397_5.44 | PE(18:0/22:6) | 3.9E-04 | 2.8E-03 | 15.11 | ↑ | 3.92 | 3.0E-01 | 4.9E-01 | 5.39 | ↑ | 2.43 | 8.1E-02 | 2.7E-01 | 8.02 | ↑ | 3.00 |
| NEG | 806.5944_5.72 | PC(16:0/18:0) | 2.5E-04 | 2.4E-03 | 6.07 | ↑ | 2.60 | 5.9E-02 | 2.6E-01 | 3.89 | ↑ | 1.96 | 6.9E-02 | 2.5E-01 | 3.45 | ↑ | 1.79 |
| NEG | 790.541_4.84 | PE(18:0/22:6) | 1.1E-03 | 4.7E-03 | 4.12 | ↑ | 2.04 | 1.0E-01 | 2.8E-01 | 2.58 | ↑ | 1.37 | 5.5E-02 | 2.5E-01 | 4.50 | ↑ | 2.17 |
| NEG | 878.5916_5.31 | PC(18:0/22:6) | 9.3E-04 | 4.3E-03 | 4.90 | ↑ | 2.29 | 1.7E-01 | 3.5E-01 | 3.06 | ↑ | 1.62 | 5.9E-02 | 2.5E-01 | 5.14 | ↑ | 2.36 |
| NEG | 766.5395_5.55 | PE(18:0/24:0) | 1.7E-04 | 2.3E-03 | 10.43 | ↑ | 3.38 | 2.2E-01 | 4.1E-01 | 3.55 | ↑ | 1.83 | 6.6E-02 | 2.5E-01 | 4.09 | ↑ | 2.03 |
| POS | 766.5724_5.16 | Eicosapentaenoyl PAF C-16 | 4.7E-02 | 1.4E-01 | 3.97 | ↑ | 1.99 | 9.1E-03 | 1.1E-01 | 2.23 | ↑ | 1.16 | 3.6E-03 | 3.3E-02 | 4.97 | ↑ | 2.31 |
| POS | 760.5848_5.35 | PC(16:0/18:1) | 5.6E-02 | 1.5E-01 | 3.58 | ↑ | 1.84 | 3.9E-03 | 8.6E-02 | 2.13 | ↑ | 1.09 | 1.1E-01 | 2.3E-01 | 2.75 | ↑ | 1.46 |
| NEG | 776.5592_5.71 | PC(18:1/14:0) | 1.2E-04 | 2.3E-03 | 9.12 | ↑ | 3.19 | 6.4E-02 | 2.7E-01 | 6.49 | ↑ | 2.70 | 2.7E-02 | 2.1E-01 | 9.43 | ↑ | 3.24 |
| NEG | 768.5463_5.54 | PC(18:0/14:0) | 3.2E-04 | 2.6E-03 | 8.05 | ↑ | 3.01 | 2.2E-01 | 4.1E-01 | 3.00 | ↑ | 1.59 | 6.2E-02 | 2.5E-01 | 3.23 | ↑ | 1.69 |
| POS | 782.567_5.35 | PC(16:0/18:1) | 7.5E-02 | 1.8E-01 | 2.33 | ↑ | 1.22 | 7.9E-03 | 1.0E-01 | 2.01 | ↑ | 1.00 | 1.6E-01 | 2.8E-01 | 1.70 | ↑ | 0.76 |
| NEG | 838.5972_5.6 | C18(Plasm)-20:4 PC | 6.3E-05 | 2.3E-03 | 6.19 | ↑ | 2.63 | 7.5E-02 | 2.7E-01 | 3.58 | ↑ | 1.84 | 7.4E-03 | 1.6E-01 | 4.21 | ↑ | 2.07 |
| NEG | 804.5669_5.01 | PC(18:1/16:0) | 8.5E-05 | 2.3E-03 | 3.39 | ↑ | 1.76 | 6.5E-03 | 1.5E-01 | 2.34 | ↑ | 1.23 | 5.6E-02 | 2.5E-01 | 2.43 | ↑ | 1.28 |
| NEG | 802.5603_5.01 | PC(16:0/18:2) | 8.5E-05 | 2.3E-03 | 2.86 | ↑ | 1.52 | 5.9E-03 | 1.5E-01 | 2.05 | ↑ | 1.04 | 5.5E-02 | 2.5E-01 | 2.09 | ↑ | 1.06 |
| NEG | 744.549_5.01 | α-Galactosyl-N-palmitoylsphingosine | 2.9E-05 | 2.3E-03 | 4.28 | ↑ | 2.10 | 9.5E-03 | 1.7E-01 | 2.48 | ↑ | 1.31 | 6.1E-02 | 2.5E-01 | 2.24 | ↑ | 1.16 |
| POS | 768.5893_5.23 | C16-18:1 PC | 3.1E-02 | 1.2E-01 | 4.25 | ↑ | 2.09 | 3.2E-03 | 8.6E-02 | 2.66 | ↑ | 1.41 | 5.9E-03 | 4.6E-02 | 4.76 | ↑ | 2.25 |
| NEG | 774.545_5.65 | C18(Plasm)-22:6 PE | 3.9E-04 | 2.8E-03 | 8.48 | ↑ | 3.08 | 1.2E-01 | 3.0E-01 | 6.54 | ↑ | 2.71 | 4.5E-02 | 2.4E-01 | 8.24 | ↑ | 3.04 |
| NEG | 714.5081_5.16 | PE(16:0/18:2) | 1.1E-04 | 2.3E-03 | 8.04 | ↑ | 3.01 | 9.1E-02 | 2.7E-01 | 3.69 | ↑ | 1.89 | 3.2E-01 | 4.8E-01 | 2.12 | ↑ | 1.08 |
| NEG | 792.5473_5.01 | PC(16:0/18:2) | 6.1E-04 | 3.4E-03 | 3.23 | ↑ | 1.69 | 7.9E-02 | 2.7E-01 | 2.16 | ↑ | 1.11 | 4.7E-02 | 2.4E-01 | 3.21 | ↑ | 1.68 |
| POS | 735.5737_5.29 | Thioetheramidephosphatidylcholine | 3.8E-02 | 1.3E-01 | 5.19 | ↑ | 2.38 | 1.1E-02 | 1.1E-01 | 2.13 | ↑ | 1.09 | 1.3E-01 | 2.6E-01 | 4.42 | ↑ | 2.14 |

| | | | | | | | | | | | | | | | | | |
|---|---|---|---|---|---|---|---|---|---|---|---|---|---|---|---|---|---|
| POS | 756.5531_5.29 | PC(16:0/16:0) | 4.1E-02 | 1.3E-01 | 3.68 | ↑ | 1.88 | 1.1E-02 | 1.1E-01 | 2.21 | ↑ | 1.14 | 1.7E-01 | 3.0E-01 | 2.55 | ↑ | 1.35 |
| POS | 184.0734_5 | PC | 6.7E-02 | 1.7E-01 | 3.03 | ↑ | 1.60 | 3.3E-03 | 8.6E-02 | 2.22 | ↑ | 1.15 | 2.9E-02 | 1.1E-01 | 3.30 | ↑ | 1.72 |
| NEG | 806.5831_5.35 | PC(16:0/18:0) | 2.3E-04 | 2.3E-03 | 5.87 | ↑ | 2.55 | 8.3E-03 | 1.7E-01 | 3.41 | ↑ | 1.77 | 5.4E-02 | 2.5E-01 | 3.29 | ↑ | 1.72 |
| POS | 834.5998_5.31 | 1-Docosahexaenoyl-2-stearoyl-sn-glycero-3-p | 1.5E-02 | 9.6E-02 | 4.13 | ↑ | 2.05 | 1.2E-02 | 1.1E-01 | 2.23 | ↑ | 1.16 | 4.0E-02 | 1.3E-01 | 6.86 | ↑ | 2.78 |
| NEG | 834.5429_5.53 | PS(18:0/22:6) | 1.4E-04 | 2.3E-03 | 7.10 | ↑ | 2.83 | 1.3E-01 | 3.1E-01 | 2.97 | ↑ | 1.57 | 1.1E-01 | 2.8E-01 | 3.01 | ↑ | 1.59 |
| NEG | 742.5398_5.01 | PE(18:0/18:2) | 9.8E-05 | 2.3E-03 | 3.34 | ↑ | 1.74 | 1.8E-02 | 1.9E-01 | 2.37 | ↑ | 1.25 | 1.6E-01 | 3.1E-01 | 2.03 | ↑ | 1.02 |
| NEG | 862.5958_5.51 | C18(Plasm)-22:6 PC | 5.7E-04 | 3.4E-03 | 4.88 | ↑ | 2.29 | 1.7E-01 | 3.5E-01 | 2.77 | ↑ | 1.47 | 6.4E-02 | 2.5E-01 | 3.84 | ↑ | 1.94 |
| POS | 758.5692_5.01 | PC(16:0/18:2) | 6.8E-02 | 1.7E-01 | 2.75 | ↑ | 1.46 | 1.5E-03 | 7.8E-02 | 2.05 | ↑ | 1.03 | 5.7E-02 | 1.6E-01 | 2.61 | ↑ | 1.39 |
| POS | 760.5746_5.01 | PC(16:0/18:1) | 7.9E-02 | 1.8E-01 | 2.52 | ↑ | 1.34 | 2.1E-03 | 7.8E-02 | 1.98 | ↑ | 0.99 | 5.4E-02 | 1.5E-01 | 2.45 | ↑ | 1.29 |
| NEG | 804.576_5.36 | PC(18:1/16:0) | 2.1E-04 | 2.3E-03 | 4.55 | ↑ | 2.19 | 1.4E-02 | 1.8E-01 | 2.67 | ↑ | 1.42 | 7.1E-02 | 2.5E-01 | 2.71 | ↑ | 1.44 |
| POS | 768.5582_5.55 | PE(18:0/18:1) | 5.0E-03 | 7.3E-02 | 7.81 | ↑ | 2.96 | 6.8E-02 | 2.4E-01 | 2.00 | ↑ | 1.00 | 1.0E-01 | 2.2E-01 | 3.96 | ↑ | 1.99 |
| NEG | 738.5085_5.1 | PE(16:0/20:4) | 8.8E-05 | 2.3E-03 | 8.20 | ↑ | 3.04 | 2.0E-01 | 3.8E-01 | 2.89 | ↑ | 1.53 | 1.2E-01 | 2.8E-01 | 3.00 | ↑ | 1.58 |
| NEG | 802.5603_8.81 | PC(16:0/18:2) | 2.8E-01 | 5.9E-01 | 1.05 | ↑ | 0.06 | 2.5E-01 | 7.0E-01 | 5.84 | ↑ | 2.55 | 4.9E-02 | 2.0E-01 | 1.11 | ↑ | 0.16 |

| | | | 8 Gy | | | | | | | | | | | | | | |
|---|---|---|---|---|---|---|---|---|---|---|---|---|---|---|---|---|---|
| | | | SD 1 vs. Pre | | | | | SD 2 vs. Pre | | | | | SD 6 vs. Pre | | | | |
| Mode | m/z_RT | Name | p-value | FDR | FC | | LOG2FC | p-value | FDR | FC | | LOG2FC | p-value | FDR | FC | | LOG2FC |
| POS | 162.1127_0.57 | L-Carnitine | 3.6E-05 | 1.0E-02 | 2.78 | ↑ | 1.47 | 7.4E-03 | 1.4E-01 | 1.80 | ↑ | 0.84 | 3.9E-03 | 2.5E-02 | 1.97 | ↑ | 0.98 |
| POS | 163.1155_0.56 | Phenylpiperazine | 1.4E-05 | 8.5E-03 | 2.97 | ↑ | 1.57 | 2.2E-03 | 8.5E-02 | 1.92 | ↑ | 0.94 | 1.8E-03 | 1.7E-02 | 2.17 | ↑ | 1.12 |
| NEG | 861.5497_12.17 | PI(18:1) | 6.3E-03 | 7.2E-02 | 0.66 | ↓ | -0.60 | 6.1E-01 | 8.6E-01 | 0.94 | ↓ | -0.10 | 9.7E-04 | 2.2E-02 | 0.57 | ↓ | -0.81 |
| POS | 580.432_7.28 | LysoPC(22:0) | 9.0E-05 | 6.5E-03 | 0.37 | ↓ | -1.42 | 7.0E-03 | 1.9E-01 | 0.58 | ↓ | -0.79 | 3.6E-02 | 1.6E-01 | 0.61 | ↓ | -0.72 |
| POS | 548.3722_1.53 | Lyso-PAF C-18 | 1.7E-03 | 3.9E-02 | 0.49 | ↓ | -1.02 | 1.2E-01 | 4.5E-01 | 0.80 | ↓ | -0.31 | 1.8E-02 | 6.6E-02 | 0.62 | ↓ | -0.68 |
| POS | 552.4031_6.62 | LysoPC(20:0) | 9.8E-04 | 2.1E-02 | 0.43 | ↓ | -1.23 | 3.6E-03 | 1.2E-01 | 0.59 | ↓ | -0.75 | 4.6E-04 | 2.0E-02 | 0.46 | ↓ | -1.13 |
| POS | 782.5682_8.51 | PC(16:0/18:1) | 3.3E-04 | 1.1E-02 | 0.36 | ↓ | -1.47 | 1.3E-02 | 2.4E-01 | 0.65 | ↓ | -0.62 | 5.7E-04 | 2.1E-02 | 0.31 | ↓ | -1.68 |
| NEG | 790.5343_8.7 | PI(18:0/22:6) | 3.6E-03 | 5.3E-02 | 0.63 | ↓ | -0.68 | 2.3E-03 | 8.5E-02 | 0.60 | ↓ | -0.73 | 4.8E-02 | 1.7E-01 | 0.67 | ↓ | -0.57 |
| POS | 510.3561_5.63 | LysoPC(17:0) | 1.8E-03 | 2.7E-02 | 0.59 | ↓ | -0.77 | 5.1E-02 | 3.6E-01 | 0.78 | ↓ | -0.36 | 4.8E-04 | 2.0E-02 | 0.56 | ↓ | -0.82 |
| POS | 339.3453_7.67 | 9-Oxo-11α,15R-dihydroxyprostan-1-oic acid | 5.6E-01 | 7.0E-01 | 0.55 | ↓ | -0.87 | 5.5E-01 | 7.9E-01 | 0.25 | ↓ | -2.03 | 1.3E-01 | 3.5E-01 | 0.08 | ↓ | -3.63 |
| POS | 338.3423_7.68 | 13-Docosenamide, (Z)- | 6.2E-01 | 7.4E-01 | 0.54 | ↓ | -0.89 | 5.2E-01 | 7.9E-01 | 0.25 | ↓ | -1.99 | 1.1E-01 | 3.2E-01 | 0.09 | ↓ | -3.48 |
| NEG | 554.3457_5.63 | LysoPC(17:0) | 3.2E-03 | 5.0E-02 | 0.63 | ↓ | -0.66 | 8.4E-02 | 3.9E-01 | 0.80 | ↓ | -0.32 | 1.6E-03 | 3.0E-02 | 0.59 | ↓ | -0.75 |
| POS | 608.4631_7.89 | LysoPC(24:0) | 1.0E-03 | 2.1E-02 | 0.42 | ↓ | -1.25 | 3.6E-02 | 3.1E-01 | 0.64 | ↓ | -0.64 | 6.4E-02 | 2.2E-01 | 0.65 | ↓ | -0.62 |
| NEG | 826.5599_8.52 | PC(16:0/20:4) | 6.0E-05 | 1.1E-02 | 0.62 | ↓ | -0.69 | 1.2E-02 | 1.6E-01 | 0.81 | ↓ | -0.31 | 2.1E-03 | 3.4E-02 | 0.63 | ↓ | -0.66 |
| NEG | 596.3925_6.63 | LysoPC(20:0) | 8.4E-04 | 2.5E-02 | 0.39 | ↓ | -1.37 | 1.6E-03 | 7.9E-02 | 0.57 | ↓ | -0.82 | 2.0E-04 | 1.1E-02 | 0.39 | ↓ | -1.37 |
| POS | 770.5677_8.69 | Dihomo-γ-linolenoyl PAF C-16 | 9.6E-04 | 2.1E-02 | 0.47 | ↓ | -1.08 | 1.2E-01 | 4.8E-01 | 0.73 | ↓ | -0.46 | 7.1E-03 | 5.6E-02 | 0.43 | ↓ | -1.20 |
| POS | 532.3383_5.63 | LysoPC(17:0) | 6.7E-03 | 5.7E-02 | 0.62 | ↓ | -0.69 | 3.1E-01 | 6.9E-01 | 0.89 | ↓ | -0.17 | 1.3E-01 | 3.5E-01 | 0.79 | ↓ | -0.34 |
| NEG | 861.5499_11.73 | PI(18:1) | 2.0E-02 | 1.4E-01 | 0.68 | ↓ | -0.55 | 7.3E-01 | 9.2E-01 | 0.95 | ↓ | -0.07 | 5.8E-04 | 1.7E-02 | 0.51 | ↓ | -0.98 |
| POS | 184.0945_0.58 | PC | 2.9E-06 | 4.2E-03 | 3.73 | ↑ | 1.90 | 1.7E-03 | 6.9E-02 | 2.27 | ↑ | 1.18 | 7.0E-05 | 4.8E-03 | 2.84 | ↑ | 1.51 |
| NEG | 788.5249_8.71 | PS(18:0/18:1) | 3.7E-03 | 5.4E-02 | 0.56 | ↓ | -0.84 | 6.0E-03 | 1.2E-01 | 0.59 | ↓ | -0.77 | 4.0E-02 | 1.5E-01 | 0.61 | ↓ | -0.71 |
| POS | 518.3256_4.86 | LysoPC(16:0) | 2.5E-02 | 1.2E-01 | 0.63 | ↓ | -0.67 | 9.7E-01 | 9.8E-01 | 1.07 | ↑ | 0.09 | 9.1E-02 | 2.8E-01 | 0.73 | ↓ | -0.45 |
| NEG | 828.5667_8.52 | C18(Plasm)-20:4 PC | 2.8E-04 | 1.6E-02 | 0.53 | ↓ | -0.91 | 3.7E-03 | 1.0E-01 | 0.74 | ↓ | -0.43 | 7.2E-04 | 1.9E-02 | 0.51 | ↓ | -0.97 |
| NEG | 827.5636_8.52 | KRTLRR | 1.6E-04 | 1.4E-02 | 0.57 | ↓ | -0.80 | 5.4E-03 | 1.2E-01 | 0.77 | ↓ | -0.37 | 4.5E-04 | 1.6E-02 | 0.54 | ↓ | -0.89 |
| NEG | 526.3113_5.06 | LysoPC(15:0) | 8.4E-03 | 8.2E-02 | 0.67 | ↓ | -0.57 | 7.4E-01 | 9.3E-01 | 0.97 | ↓ | -0.04 | 4.9E-03 | 5.1E-02 | 0.60 | ↓ | -0.75 |
| NEG | 594.3771_6.08 | Pyrrolidino PAF C-16 | 1.0E-03 | 2.7E-02 | 0.58 | ↓ | -0.78 | 1.3E-02 | 1.7E-01 | 0.73 | ↓ | -0.46 | 1.2E-03 | 2.5E-02 | 0.59 | ↓ | -0.77 |
| POS | 510.3564_1.56 | LysoPC(17:0) | 4.1E-04 | 2.0E-02 | 0.44 | ↓ | -1.18 | 2.0E-01 | 5.7E-01 | 0.84 | ↓ | -0.25 | 1.1E-05 | 1.9E-03 | 0.38 | ↓ | -1.41 |
| POS | 482.3248_5.06 | LysoPC(15:0) | 1.5E-03 | 2.5E-02 | 0.56 | ↓ | -0.83 | 2.6E-01 | 6.4E-01 | 0.87 | ↓ | -0.20 | 1.5E-03 | 3.1E-02 | 0.50 | ↓ | -1.00 |

| | | | | | | | | | | | | | | | | | |
|---|---|---|---|---|---|---|---|---|---|---|---|---|---|---|---|---|---|
| NEG | 554.3463_1.56 | LysoPC(17:0) | 1.5E-03 | 5.8E-03 | 0.50 | ↓ | -1.01 | 1.0E-01 | 4.1E-01 | 0.77 | ↓ | -0.38 | 1.1E-01 | 1.7E-01 | 0.77 | ↓ | -0.38 |
| POS | 552.4037_2.62 | LysoPC(20:0) | 1.3E-03 | 3.6E-02 | 0.39 | ↓ | -1.37 | 2.2E-01 | 6.2E-01 | 0.78 | ↓ | -0.35 | 8.4E-02 | 2.1E-01 | 0.65 | ↓ | -0.63 |
| NEG | 624.4238_7.29 | LysoPC(22:0) | 2.4E-04 | 1.6E-02 | 0.44 | ↓ | -1.20 | 9.8E-04 | 7.0E-02 | 0.59 | ↓ | -0.75 | 7.6E-04 | 1.9E-02 | 0.45 | ↓ | -1.14 |
| POS | 524.3722_5.96 | LysoPC(18:0) | 1.1E-02 | 7.6E-02 | 0.75 | ↓ | -0.42 | 4.4E-01 | 7.5E-01 | 0.94 | ↓ | -0.10 | 2.8E-01 | 5.2E-01 | 0.89 | ↓ | -0.17 |
| NEG | 512.2985_4.81 | LysoPC(14:0) | 4.2E-02 | 1.9E-01 | 0.73 | ↓ | -0.45 | 2.3E-01 | 5.9E-01 | 1.31 | ↑ | 0.39 | 7.2E-02 | 2.2E-01 | 0.71 | ↓ | -0.50 |
| NEG | 652.4552_7.9 | LysoPC(24:0) | 5.7E-04 | 2.3E-02 | 0.51 | ↓ | -0.97 | 2.1E-02 | 2.2E-01 | 0.70 | ↓ | -0.52 | 4.8E-03 | 5.0E-02 | 0.51 | ↓ | -0.97 |
| NEG | 226.0175_2.4 | 4-Quinolinol, 8-methyl-2-(trifluoromethyl)- | 1.1E-02 | 9.5E-02 | 0.45 | ↓ | -1.15 | 8.7E-03 | 1.4E-01 | 0.24 | ↓ | -2.07 | 1.5E-03 | 3.0E-02 | 0.17 | ↓ | -2.53 |
| POS | 516.3104_4.97 | Oleyloxyethylphosphorylcholine | 3.9E-04 | 1.2E-02 | 0.46 | ↓ | -1.11 | 3.0E-01 | 6.8E-01 | 0.88 | ↓ | -0.19 | 6.6E-01 | 7.9E-01 | 1.73 | ↑ | 0.79 |
| POS | 162.1129_0.33 | L-Carnitine | 7.2E-06 | 1.4E-03 | 2.89 | ↑ | 1.53 | 8.9E-04 | 4.8E-02 | 1.95 | ↑ | 0.96 | 3.4E-02 | 1.5E-01 | 1.52 | ↑ | 0.60 |
| POS | 508.3397_5.21 | C18(Plasm) LPC | 3.8E-03 | 4.3E-02 | 0.53 | ↓ | -0.92 | 6.9E-01 | 8.6E-01 | 0.94 | ↓ | -0.09 | 4.7E-02 | 1.8E-01 | 0.64 | ↓ | -0.65 |
| NEG | 512.299_0.99 | LysoPC(14:0) | 2.8E-02 | 6.0E-02 | 0.67 | ↓ | -0.59 | 4.3E-01 | 7.7E-01 | 1.37 | ↑ | 0.45 | 1.7E-01 | 2.4E-01 | 0.81 | ↓ | -0.30 |
| NEG | 816.5759_9.01 | PC(16:0/20:4) | 6.1E-03 | 7.1E-02 | 0.68 | ↓ | -0.56 | 2.5E-02 | 2.3E-01 | 0.72 | ↓ | -0.47 | 5.2E-03 | 5.3E-02 | 0.58 | ↓ | -0.80 |
| POS | 204.1231_0.58 | Acetyl-DL-carnitine | 4.9E-03 | 7.0E-02 | 1.42 | ↑ | 0.51 | 7.4E-01 | 9.9E-01 | 1.29 | ↑ | 0.36 | 9.2E-05 | 4.8E-03 | 5.60 | ↑ | 2.48 |
| POS | 482.3253_1.13 | LysoPE(18:0) | 1.6E-03 | 3.9E-02 | 0.48 | ↓ | -1.07 | 5.7E-01 | 9.3E-01 | 0.93 | ↓ | -0.10 | 1.8E-04 | 6.6E-03 | 0.29 | ↓ | -1.77 |
| POS | 468.3092_4.81 | LysoPC(14:0) | 2.6E-02 | 1.2E-01 | 0.71 | ↓ | -0.50 | 2.1E-01 | 5.9E-01 | 1.40 | ↑ | 0.48 | 7.2E-02 | 2.4E-01 | 0.71 | ↓ | -0.49 |
| POS | 782.5682_4.96 | PC(16:0/18:1) | 6.4E-02 | 1.7E-01 | 2.94 | ↑ | 1.56 | 4.1E-02 | 3.1E-01 | 1.57 | ↑ | 0.65 | 1.5E-03 | 1.5E-02 | 4.33 | ↑ | 2.11 |
| POS | 530.368_6.36 | C18(Plasm) LPC | 3.6E-02 | 1.5E-01 | 1.91 | ↑ | 0.94 | 1.5E-03 | 6.7E-02 | 2.98 | ↑ | 1.57 | 2.4E-03 | 3.7E-02 | 5.20 | ↑ | 2.38 |
| POS | 530.3694_1.4 | C18(Plasm) LPC | 4.1E-02 | 1.5E-01 | 2.22 | ↑ | 1.15 | 6.3E-03 | 1.3E-01 | 3.46 | ↑ | 1.79 | 3.6E-02 | 1.1E-01 | 3.06 | ↑ | 1.61 |
| NEG | 526.31_1.14 | LysoPC(15:0) | 8.1E-04 | 3.8E-03 | 0.57 | ↓ | -0.81 | 3.4E-01 | 7.1E-01 | 0.88 | ↓ | -0.18 | 2.1E-02 | 4.2E-02 | 0.63 | ↓ | -0.66 |
| NEG | 229.1437_3.65 | Dodecanedioic acid | 1.5E-02 | 1.1E-01 | 0.58 | ↓ | -0.80 | 7.4E-03 | 1.4E-01 | 0.62 | ↓ | -0.69 | 1.2E-01 | 2.9E-01 | 0.73 | ↓ | -0.45 |
| POS | 790.6208_10.03 | PS(18:0/18:1) | 1.2E-02 | 7.6E-02 | 0.71 | ↓ | -0.49 | 1.0E+00 | 1.0E+00 | 1.02 | ↑ | 0.03 | 6.1E-03 | 5.2E-02 | 0.68 | ↓ | -0.55 |
| POS | 204.1141_0.33 | Acetyl-DL-carnitine | 8.7E-03 | 6.5E-02 | 1.50 | ↑ | 0.59 | 3.3E-01 | 7.0E-01 | 1.26 | ↑ | 0.33 | 5.3E-05 | 7.0E-03 | 3.50 | ↑ | 1.81 |
| POS | 316.2485_3.86 | Decanoyl-L-carnitine | 2.1E-01 | 3.8E-01 | 1.81 | ↑ | 0.85 | 6.6E-01 | 8.5E-01 | 1.20 | ↑ | 0.27 | 8.4E-04 | 2.5E-02 | 6.91 | ↑ | 2.79 |
| POS | 468.3099_0.99 | LysoPC(14:0) | 9.5E-03 | 9.5E-02 | 0.59 | ↓ | -0.77 | 9.1E-01 | 9.9E-01 | 1.09 | ↑ | 0.12 | 1.4E-02 | 5.6E-02 | 0.51 | ↓ | -0.96 |
| NEG | 584.3554_4.45 | 2-Thio PAF | 1.1E-02 | 9.3E-02 | 0.65 | ↓ | -0.61 | 6.4E-01 | 8.8E-01 | 1.09 | ↑ | 0.12 | 1.6E-05 | 3.5E-03 | 0.54 | ↓ | -0.88 |
| NEG | 178.0488_0.62 | Hippuric acid | 3.0E-02 | 6.2E-02 | 0.48 | ↓ | -1.07 | 5.0E-03 | 1.7E-01 | 0.38 | ↓ | -1.39 | 5.0E-02 | 8.4E-02 | 0.49 | ↓ | -1.02 |
| POS | 728.5584_10.1 | PE(18:0/18:1) | 9.8E-02 | 2.6E-01 | 0.61 | ↓ | -0.72 | 4.6E-01 | 7.6E-01 | 0.81 | ↓ | -0.31 | 9.7E-04 | 2.6E-02 | 0.26 | ↓ | -1.96 |
| NEG | 452.2775_0.99 | LysoPE(16:0) | 2.7E-03 | 8.9E-03 | 0.56 | ↓ | -0.84 | 3.4E-01 | 7.1E-01 | 1.36 | ↑ | 0.44 | 1.5E-01 | 2.1E-01 | 0.77 | ↓ | -0.37 |
| POS | 257.2267_3.76 | Androst-16-en-3-ol, (3β,5α)- | 5.6E-04 | 1.6E-02 | 2.77 | ↑ | 1.47 | 1.2E-03 | 5.9E-02 | 2.63 | ↑ | 1.40 | 1.1E-02 | 7.0E-02 | 2.01 | ↑ | 1.01 |
| NEG | 862.5542_4.81 | C18(Plasm)-22:6 PC | 1.1E-03 | 4.6E-03 | 3.57 | ↑ | 1.84 | 8.5E-01 | 9.7E-01 | 1.04 | ↑ | 0.06 | 8.4E-03 | 1.9E-02 | 5.74 | ↑ | 2.52 |
| POS | 806.5689_4.84 | Arachidonoylthio-PC | 4.0E-02 | 1.5E-01 | 2.61 | ↑ | 1.38 | 4.6E-02 | 3.3E-01 | 1.59 | ↑ | 0.67 | 1.3E-03 | 1.4E-02 | 3.80 | ↑ | 1.92 |
| NEG | 263.1033_1.9 | L-Glutamine, N2-(2-phenylacetyl)- | 9.9E-04 | 2.7E-02 | 0.33 | ↓ | -1.60 | 5.4E-04 | 7.0E-02 | 0.25 | ↓ | -1.98 | 2.7E-02 | 1.2E-01 | 0.47 | ↓ | -1.09 |
| POS | 504.3088_1.14 | Lysophosphatidylethanolamine | 2.1E-03 | 4.5E-02 | 0.52 | ↓ | -0.94 | 5.1E-01 | 9.1E-01 | 0.91 | ↓ | -0.13 | 3.2E-04 | 8.1E-03 | 0.43 | ↓ | -1.20 |
| POS | 524.3717_1.89 | LysoPC(18:0) | 3.5E-03 | 5.9E-02 | 0.56 | ↓ | -0.85 | 6.1E-01 | 9.5E-01 | 0.95 | ↓ | -0.07 | 4.8E-03 | 2.8E-02 | 0.63 | ↓ | -0.67 |
| NEG | 830.5807_5.1 | PC(18:0/18:2) | 2.4E-03 | 8.3E-03 | 2.09 | ↑ | 1.06 | 1.1E-01 | 4.2E-01 | 0.65 | ↓ | -0.62 | 6.8E-04 | 3.2E-03 | 3.94 | ↑ | 1.98 |
| POS | 544.338_1.41 | LysoPC(18:1) | 5.5E-05 | 1.1E-02 | 0.60 | ↓ | -0.75 | 8.7E-02 | 4.1E-01 | 0.88 | ↓ | -0.18 | 3.3E-02 | 1.0E-01 | 0.61 | ↓ | -0.72 |
| POS | 214.1134_3.03 | Suberylglycine | 3.7E-01 | 5.5E-01 | 1.11 | ↑ | 0.15 | 1.5E-09 | 3.9E-06 | 9.41 | ↑ | 3.23 | 3.3E-09 | 8.5E-06 | 7.91 | ↑ | 2.98 |
| NEG | 768.5551_5.11 | PC(18:0/18:2) | 1.9E-03 | 6.9E-03 | 2.02 | ↑ | 1.01 | 1.0E-01 | 4.1E-01 | 0.69 | ↓ | -0.54 | 3.8E-04 | 2.6E-03 | 3.83 | ↑ | 1.94 |
| POS | 130.0499_1.9 | Glutamic Acid | 4.6E-04 | 1.4E-02 | 0.30 | ↓ | -1.74 | 4.1E-04 | 2.9E-02 | 0.26 | ↓ | -1.93 | 1.6E-02 | 9.6E-02 | 0.43 | ↓ | -1.22 |
| POS | 490.2926_0.99 | LysoPC(14:0) | 1.5E-01 | 2.5E-01 | 0.72 | ↓ | -0.47 | 5.9E-01 | 9.4E-01 | 0.89 | ↓ | -0.18 | 2.0E-02 | 7.1E-02 | 0.46 | ↓ | -1.11 |
| POS | 788.6068_5.47 | PC(18:0/18:1) | 6.2E-02 | 1.6E-01 | 2.72 | ↑ | 1.44 | 7.2E-02 | 3.9E-01 | 1.48 | ↑ | 0.56 | 1.2E-03 | 1.4E-02 | 3.01 | ↑ | 1.59 |
| POS | 287.1015_0.61 | Phenylacetyl-L-glutamine | 3.7E-03 | 6.0E-02 | 0.33 | ↓ | -1.58 | 5.9E-04 | 3.8E-02 | 0.25 | ↓ | -1.99 | 1.4E-02 | 5.5E-02 | 0.41 | ↓ | -1.29 |
| POS | 786.6003_5.47 | 1,2-Dioleoyl-sn-glycero-3-PC | 6.4E-02 | 1.7E-01 | 2.81 | ↑ | 1.49 | 6.7E-02 | 3.9E-01 | 1.46 | ↑ | 0.54 | 1.6E-03 | 1.6E-02 | 2.99 | ↑ | 1.58 |

| | | | | | | | | | | | | | | | | | |
|---|---|---|---|---|---|---|---|---|---|---|---|---|---|---|---|---|---|
| POS | 532.3389_1.56 | LysoPC(17:0) | 5.8E-04 | 2.1E-02 | 0.33 | ↓ | -1.58 | 5.3E-02 | 3.5E-01 | 0.65 | ↓ | -0.61 | 5.9E-07 | 2.2E-04 | 0.24 | ↓ | -2.05 |
| POS | 794.5989_5.26 | C18(Plasm)-20:4 PC | 6.8E-02 | 1.7E-01 | 3.76 | ↑ | 1.91 | 3.7E-02 | 3.0E-01 | 1.76 | ↑ | 0.82 | 4.0E-04 | 9.1E-03 | 4.47 | ↑ | 2.16 |
| POS | 421.2528_5.75 | Tris(2-butoxyethyl) phosphate | 2.2E-02 | 1.1E-01 | 1.55 | ↑ | 0.63 | 1.7E-06 | 5.6E-04 | 2.98 | ↑ | 1.58 | 1.2E-03 | 2.8E-02 | 2.57 | ↑ | 1.36 |
| POS | 180.0972_0.33 | N-Benzylglycine methyl ester | 1.7E-03 | 2.7E-02 | 0.46 | ↓ | -1.13 | 7.1E-02 | 4.0E-01 | 0.63 | ↓ | -0.67 | 1.0E-03 | 2.6E-02 | 0.34 | ↓ | -1.55 |
| NEG | 826.5604_4.96 | PC(16:0/20:4) | 4.9E-04 | 2.8E-03 | 3.00 | ↑ | 1.59 | 8.6E-01 | 9.7E-01 | 1.05 | ↑ | 0.07 | 2.1E-04 | 2.5E-03 | 5.42 | ↑ | 2.44 |
| POS | 637.3064_1.8 | Acetylpodocarpic acid anhydride | 1.3E-01 | 2.3E-01 | 1.18 | ↑ | 0.24 | 1.4E-06 | 8.2E-04 | 2.39 | ↑ | 1.26 | 5.8E-08 | 8.6E-05 | 2.81 | ↑ | 1.49 |
| POS | 808.5841_5 | 1,2-Dioleoyl-sn-glycero-3-PC | 6.1E-02 | 1.6E-01 | 2.38 | ↑ | 1.25 | 3.6E-02 | 3.0E-01 | 1.55 | ↑ | 0.63 | 5.6E-05 | 4.8E-03 | 4.21 | ↑ | 2.07 |
| NEG | 766.5393_4.96 | PE(18:0/20:4) | 1.6E-04 | 1.9E-03 | 3.23 | ↑ | 1.69 | 7.4E-01 | 9.2E-01 | 1.14 | ↑ | 0.19 | 2.6E-04 | 2.5E-03 | 6.12 | ↑ | 2.61 |
| NEG | 835.5345_5.54 | PI(16:0/18:1) | 1.3E-05 | 1.4E-03 | 7.22 | ↑ | 2.85 | 9.4E-01 | 9.9E-01 | 1.10 | ↑ | 0.14 | 2.5E-04 | 2.5E-03 | 7.82 | ↑ | 2.97 |
| NEG | 850.5605_4.85 | PC(16;0/22:6) | 2.9E-04 | 2.2E-03 | 3.30 | ↑ | 1.72 | 7.5E-01 | 9.3E-01 | 0.98 | ↓ | -0.04 | 2.2E-04 | 2.5E-03 | 6.14 | ↑ | 2.62 |
| POS | 808.5769_4.83 | 1,2-Dioleoyl-sn-glycero-3-PC | 1.4E-01 | 2.4E-01 | 1.96 | ↑ | 0.97 | 2.2E-01 | 6.1E-01 | 1.33 | ↑ | 0.42 | 2.0E-03 | 1.8E-02 | 2.92 | ↑ | 1.54 |
| POS | 502.3262_1.49 | C16(Plasm) LPC | 1.9E-03 | 4.2E-02 | 0.50 | ↓ | -1.00 | 7.6E-01 | 9.9E-01 | 0.95 | ↓ | -0.07 | 2.8E-04 | 7.8E-03 | 0.42 | ↓ | -1.24 |
| POS | 678.4682_6.87 | PC(14:0) | 2.2E-02 | 1.1E-01 | 0.58 | ↓ | -0.78 | 1.4E-04 | 1.3E-02 | 0.27 | ↓ | -1.89 | 6.6E-02 | 2.2E-01 | 0.64 | ↓ | -0.65 |
| POS | 546.3536_1.89 | LysoPC(18:0) | 2.8E-03 | 5.3E-02 | 0.63 | ↓ | -0.66 | 4.3E-01 | 8.5E-01 | 0.93 | ↓ | -0.10 | 9.9E-02 | 2.4E-01 | 0.79 | ↓ | -0.34 |
| NEG | 770.571_5.48 | DMPE(18:1/18:1) | 2.3E-03 | 8.0E-03 | 2.73 | ↑ | 1.45 | 4.8E-01 | 8.0E-01 | 0.75 | ↓ | -0.41 | 6.5E-04 | 3.2E-03 | 4.80 | ↑ | 2.26 |
| NEG | 832.5981_5.48 | PC(18:1/18:0) | 1.3E-03 | 5.3E-03 | 2.80 | ↑ | 1.49 | 4.1E-01 | 7.5E-01 | 0.77 | ↓ | -0.39 | 3.4E-04 | 2.6E-03 | 5.12 | ↑ | 2.36 |
| POS | 792.5895_5.18 | Dihomo-γ-linolenoyl PAF C-16 | 4.4E-02 | 1.5E-01 | 5.28 | ↑ | 2.40 | 3.7E-02 | 3.0E-01 | 1.73 | ↑ | 0.79 | 6.8E-04 | 1.2E-02 | 4.88 | ↑ | 2.29 |
| NEG | 794.5685_5.41 | PC(18:1/16:0) | 4.0E-04 | 2.6E-03 | 3.51 | ↑ | 1.81 | 4.8E-01 | 8.0E-01 | 0.87 | ↓ | -0.20 | 3.3E-04 | 2.6E-03 | 7.07 | ↑ | 2.82 |
| POS | 750.5453_5.35 | Palmitoyl Thio-PC | 1.6E-02 | 1.1E-01 | 6.06 | ↑ | 2.60 | 1.4E-02 | 1.8E-01 | 1.91 | ↑ | 0.93 | 3.8E-04 | 8.9E-03 | 6.09 | ↑ | 2.61 |
| POS | 810.5994_5.42 | PC(18:0/18:1) | 1.4E-02 | 1.1E-01 | 3.82 | ↑ | 1.93 | 4.0E-02 | 3.1E-01 | 1.70 | ↑ | 0.76 | 8.9E-04 | 1.3E-02 | 5.28 | ↑ | 2.40 |
| NEG | 832.6078_5.77 | PC(18:1/18:0) | 6.6E-04 | 3.3E-03 | 3.76 | ↑ | 1.91 | 8.2E-01 | 9.6E-01 | 0.88 | ↓ | -0.18 | 7.8E-04 | 3.4E-03 | 7.02 | ↑ | 2.81 |
| NEG | 830.5917_5.48 | PC(18:1/18:1) | 1.4E-03 | 5.5E-03 | 2.51 | ↑ | 1.33 | 4.7E-01 | 8.0E-01 | 0.80 | ↓ | -0.32 | 4.3E-04 | 2.8E-03 | 4.18 | ↑ | 2.07 |
| NEG | 747.5162_5.21 | SM(d18:1/16:0) | 1.5E-04 | 1.9E-03 | 5.78 | ↑ | 2.53 | 8.3E-01 | 9.6E-01 | 0.94 | ↓ | -0.08 | 4.2E-04 | 2.8E-03 | 11.99 | ↑ | 3.58 |
| NEG | 762.5082_4.98 | PE(16:0/22:6) | 4.5E-05 | 1.6E-03 | 10.69 | ↑ | 3.42 | 5.4E-01 | 8.3E-01 | 0.80 | ↓ | -0.32 | 9.8E-04 | 3.8E-03 | 15.34 | ↑ | 3.94 |
| POS | 790.5725_5.24 | Arachidonoyl PAF C-16 | 3.7E-02 | 1.4E-01 | 4.32 | ↑ | 2.11 | 3.1E-02 | 2.8E-01 | 2.05 | ↑ | 1.03 | 1.2E-03 | 1.4E-02 | 4.99 | ↑ | 2.32 |
| NEG | 765.5276_5.16 | SM(d18:1/18:0) | 6.2E-05 | 1.7E-03 | 4.52 | ↑ | 2.18 | 1.1E-01 | 4.2E-01 | 0.64 | ↓ | -0.63 | 2.4E-03 | 6.9E-03 | 5.72 | ↑ | 2.52 |
| NEG | 820.577_5.49 | PC(18:0/18:2) | 9.5E-05 | 1.7E-03 | 2.84 | ↑ | 1.51 | 9.5E-01 | 9.9E-01 | 1.03 | ↑ | 0.05 | 1.8E-03 | 5.5E-03 | 10.75 | ↑ | 3.43 |
| NEG | 790.5397_5.44 | PE(18:0/22:6) | 4.1E-05 | 1.6E-03 | 11.63 | ↑ | 3.54 | 5.9E-01 | 8.5E-01 | 0.79 | ↓ | -0.35 | 9.2E-04 | 3.7E-03 | 14.69 | ↑ | 3.88 |
| NEG | 806.5944_5.72 | PC(16:0/18:0) | 2.9E-04 | 2.2E-03 | 3.93 | ↑ | 1.97 | 9.0E-01 | 9.8E-01 | 0.95 | ↓ | -0.07 | 1.5E-03 | 4.9E-03 | 6.40 | ↑ | 2.68 |
| NEG | 790.541_4.84 | PE(18:0/22:6) | 1.3E-03 | 5.1E-03 | 2.60 | ↑ | 1.38 | 6.8E-01 | 8.9E-01 | 0.99 | ↓ | -0.02 | 5.4E-04 | 3.0E-03 | 5.52 | ↑ | 2.46 |
| NEG | 878.5916_5.31 | PC(18:0/22:6) | 4.2E-04 | 2.6E-03 | 3.64 | ↑ | 1.86 | 6.8E-01 | 8.9E-01 | 0.92 | ↓ | -0.12 | 3.4E-04 | 2.6E-03 | 7.12 | ↑ | 2.83 |
| NEG | 766.5395_5.55 | PE(18:0/24:0) | 1.4E-05 | 1.4E-03 | 10.32 | ↑ | 3.37 | 6.2E-01 | 8.7E-01 | 0.93 | ↓ | -0.10 | 6.4E-04 | 3.2E-03 | 13.74 | ↑ | 3.78 |
| POS | 766.5724_5.16 | Eicosapentaenoyl PAF C-16 | 3.7E-02 | 1.4E-01 | 4.49 | ↑ | 2.17 | 6.5E-02 | 3.8E-01 | 1.70 | ↑ | 0.76 | 2.1E-03 | 1.8E-02 | 4.63 | ↑ | 2.21 |
| POS | 760.5848_5.35 | PC(16:0/18:1) | 4.3E-02 | 1.5E-01 | 3.10 | ↑ | 1.63 | 5.2E-03 | 1.3E-01 | 1.95 | ↑ | 0.96 | 3.1E-03 | 2.1E-02 | 3.25 | ↑ | 1.70 |
| NEG | 776.5592_5.71 | PC(18:1/14:0) | 5.7E-04 | 3.0E-03 | 6.28 | ↑ | 2.65 | 8.3E-01 | 9.6E-01 | 0.82 | ↓ | -0.29 | 1.4E-03 | 4.7E-03 | 11.54 | ↑ | 3.53 |
| NEG | 768.5463_5.54 | PC(18:0/14:0) | 2.0E-04 | 2.0E-03 | 5.66 | ↑ | 2.50 | 4.1E-01 | 7.5E-01 | 0.76 | ↓ | -0.40 | 4.7E-03 | 1.2E-02 | 7.40 | ↑ | 2.89 |
| POS | 782.567_5.35 | PC(16:0/18:1) | 4.5E-02 | 1.5E-01 | 1.84 | ↑ | 0.88 | 1.2E-02 | 1.7E-01 | 1.77 | ↑ | 0.82 | 1.9E-03 | 1.7E-02 | 2.21 | ↑ | 1.14 |
| NEG | 838.5972_5.6 | C18(Plasm)-20:4 PC | 4.8E-04 | 2.7E-03 | 4.81 | ↑ | 2.27 | 6.8E-01 | 8.9E-01 | 0.86 | ↓ | -0.22 | 2.9E-03 | 7.9E-03 | 8.47 | ↑ | 3.08 |
| NEG | 804.5669_5.01 | PC(18:1/16:0) | 3.0E-04 | 2.3E-03 | 2.41 | ↑ | 1.27 | 8.2E-01 | 9.6E-01 | 0.96 | ↓ | -0.06 | 1.1E-04 | 2.5E-03 | 3.74 | ↑ | 1.90 |
| NEG | 802.5603_5.01 | PC(16:0/18:2) | 3.5E-04 | 2.4E-03 | 2.13 | ↑ | 1.09 | 9.8E-01 | 9.9E-01 | 1.00 | ↓ | 0.00 | 9.2E-05 | 2.5E-03 | 3.30 | ↑ | 1.72 |
| NEG | 744.549_5.01 | α-Galactosyl-N-palmitoylsphingosine | 9.8E-05 | 1.7E-03 | 3.23 | ↑ | 1.69 | 7.8E-01 | 9.5E-01 | 0.95 | ↓ | -0.07 | 1.2E-04 | 2.5E-03 | 3.87 | ↑ | 1.95 |
| POS | 768.5893_5.23 | C16-18:1 PC | 3.3E-02 | 1.4E-01 | 4.72 | ↑ | 2.24 | 1.3E-02 | 1.7E-01 | 1.99 | ↑ | 0.99 | 3.1E-03 | 2.2E-02 | 4.36 | ↑ | 2.13 |
| NEG | 774.545_5.65 | C18(Plasm)-22:6 PE | 2.2E-04 | 2.1E-03 | 5.86 | ↑ | 2.55 | 9.4E-01 | 9.9E-01 | 0.84 | ↓ | -0.25 | 1.6E-03 | 5.3E-03 | 10.66 | ↑ | 3.41 |

| | | | | | | | | | | | | | | | | | |
|---|---|---|---|---|---|---|---|---|---|---|---|---|---|---|---|---|---|
| NEG | 714.5081_5.16 | PE(16:0/18:2) | 1.4E-04 | 1.8E-03 | 5.48 | ↑ | 2.45 | 9.1E-01 | 9.8E-01 | 0.99 | ↓ | -0.02 | 4.2E-03 | 1.1E-02 | 5.10 | ↑ | 2.35 |
| NEG | 792.5473_5.01 | PC(16:0/18:2) | 8.0E-04 | 3.7E-03 | 2.07 | ↑ | 1.05 | 3.5E-01 | 7.1E-01 | 0.86 | ↓ | -0.22 | 4.2E-05 | 2.5E-03 | 4.27 | ↑ | 2.09 |
| POS | 735.5737_5.29 | Thioetheramidephosphatidylcholine | 1.9E-02 | 1.2E-01 | 5.24 | ↑ | 2.39 | 6.7E-03 | 1.4E-01 | 2.07 | ↑ | 1.05 | 6.3E-03 | 3.3E-02 | 4.34 | ↑ | 2.12 |
| POS | 756.5531_5.29 | PC(16:0/16:0) | 1.3E-02 | 1.0E-01 | 3.71 | ↑ | 1.89 | 5.1E-03 | 1.3E-01 | 2.14 | ↑ | 1.09 | 4.6E-03 | 2.7E-02 | 2.94 | ↑ | 1.56 |
| POS | 184.0734_5 | PC | 1.3E-01 | 2.3E-01 | 2.53 | ↑ | 1.34 | 2.2E-02 | 2.3E-01 | 1.68 | ↑ | 0.75 | 7.6E-04 | 1.2E-02 | 3.28 | ↑ | 1.71 |
| NEG | 806.5831_5.35 | PC(16:0/18:0) | 7.5E-06 | 1.4E-03 | 3.76 | ↑ | 1.91 | 8.1E-01 | 9.6E-01 | 1.09 | ↑ | 0.13 | 2.5E-04 | 2.5E-03 | 6.45 | ↑ | 2.69 |
| POS | 834.5998_5.31 | 1-Docosahexaenoyl-2-stearoyl-sn-glycero-3-p | 3.8E-02 | 1.4E-01 | 4.18 | ↑ | 2.06 | 3.2E-02 | 2.9E-01 | 1.78 | ↑ | 0.83 | 1.3E-03 | 1.5E-02 | 5.16 | ↑ | 2.37 |
| NEG | 834.5429_5.53 | PS(18:0/22:6) | 2.7E-05 | 1.5E-03 | 6.01 | ↑ | 2.59 | 4.1E-01 | 7.5E-01 | 0.81 | ↓ | -0.31 | 9.0E-04 | 3.6E-03 | 7.68 | ↑ | 2.94 |
| NEG | 742.5398_5.01 | PE(18:0/18:2) | 2.8E-04 | 2.2E-03 | 2.66 | ↑ | 1.41 | 8.6E-01 | 9.7E-01 | 1.04 | ↑ | 0.06 | 8.6E-05 | 2.5E-03 | 3.88 | ↑ | 1.96 |
| NEG | 862.5958_5.51 | C18(Plasm)-22:6 PC | 8.5E-04 | 3.8E-03 | 3.50 | ↑ | 1.81 | 2.4E-01 | 6.1E-01 | 0.66 | ↓ | -0.59 | 2.9E-03 | 8.0E-03 | 4.87 | ↑ | 2.28 |
| POS | 758.5692_5.01 | PC(16:0/18:2) | 1.0E-01 | 2.1E-01 | 2.42 | ↑ | 1.27 | 7.8E-03 | 1.4E-01 | 1.61 | ↑ | 0.69 | 8.9E-04 | 1.3E-02 | 2.86 | ↑ | 1.51 |
| POS | 760.5746_5.01 | PC(16:0/18:1) | 1.2E-01 | 2.2E-01 | 2.19 | ↑ | 1.13 | 8.6E-03 | 1.5E-01 | 1.57 | ↑ | 0.65 | 8.8E-04 | 1.3E-02 | 2.69 | ↑ | 1.43 |
| NEG | 804.576_5.36 | PC(18:1/16:0) | 7.4E-05 | 1.7E-03 | 2.99 | ↑ | 1.58 | 9.0E-01 | 9.8E-01 | 1.01 | ↑ | 0.02 | 2.9E-04 | 2.5E-03 | 5.12 | ↑ | 2.36 |
| POS | 768.5582_5.55 | PE(18:0/18:1) | 4.7E-03 | 6.8E-02 | 9.91 | ↑ | 3.31 | 1.7E-01 | 5.2E-01 | 1.63 | ↑ | 0.71 | 1.9E-03 | 1.7E-02 | 4.83 | ↑ | 2.27 |
| NEG | 738.5085_5.1 | PE(16:0/20:4) | 4.4E-05 | 1.6E-03 | 7.24 | ↑ | 2.86 | 6.1E-01 | 8.6E-01 | 0.87 | ↓ | -0.20 | 1.3E-03 | 4.5E-03 | 9.86 | ↑ | 3.30 |
| NEG | 802.5603_8.81 | PC(16:0/18:2) | 1.3E-02 | 1.0E-01 | 1.06 | ↑ | 0.08 | 1.5E-03 | 7.9E-02 | 1.13 | ↑ | 0.18 | 2.4E-02 | 1.1E-01 | 1.15 | ↑ | 0.21 |

| | | | | | | | | | | 8.5 Gy | | | | | | | |
|---|---|---|---|---|---|---|---|---|---|---|---|---|---|---|---|---|---|
| | | | SD 1 vs. Pre | | | | | SD 2 vs. Pre | | | | | SD 6 vs. Pre | | | | |
| **Mode** | **m/z_RT** | **Name** | **p-value** | **FDR** | **FC** | | **LOG2FC** | **p-value** | **FDR** | **FC** | | **LOG2FC** | **p-value** | **FDR** | **FC** | | **LOG2FC** |
| POS | 162.1127_0.57 | L-Carnitine | 8.9E-07 | 6.4E-04 | 4.24 | ↑ | 2.08 | 4.7E-06 | 2.5E-03 | 2.55 | ↑ | 1.35 | 2.7E-04 | 1.5E-02 | 3.62 | ↑ | 1.85 |
| POS | 163.1155_0.56 | Phenylpiperazine | 1.3E-06 | 6.4E-04 | 3.89 | ↑ | 1.96 | 5.2E-06 | 2.5E-03 | 2.36 | ↑ | 1.24 | 3.9E-04 | 1.9E-02 | 3.42 | ↑ | 1.77 |
| NEG | 861.5497_12.17 | PI(18:1) | 1.6E-03 | 2.2E-02 | 0.48 | ↓ | -1.07 | 7.0E-02 | 3.4E-01 | 0.70 | ↓ | -0.51 | 2.8E-04 | 1.4E-02 | 0.46 | ↓ | -1.12 |
| POS | 580.432_7.28 | LysoPC(22:0) | 4.2E-04 | 7.8E-03 | 0.35 | ↓ | -1.51 | 3.3E-02 | 2.1E-01 | 0.60 | ↓ | -0.74 | 1.6E-01 | 4.2E-01 | 0.74 | ↓ | -0.44 |
| POS | 548.3722_1.53 | Lyso-PAF C-18 | 1.0E-03 | 3.0E-02 | 0.44 | ↓ | -1.18 | 1.2E-01 | 4.0E-01 | 0.75 | ↓ | -0.41 | 2.6E-03 | 4.1E-02 | 0.62 | ↓ | -0.69 |
| POS | 552.4031_6.62 | LysoPC(20:0) | 1.5E-04 | 7.4E-03 | 0.29 | ↓ | -1.76 | 2.8E-02 | 2.0E-01 | 0.54 | ↓ | -0.89 | 2.1E-01 | 4.8E-01 | 0.74 | ↓ | -0.44 |
| POS | 782.5682_8.51 | PC(16:0/18:1) | 1.9E-03 | 1.4E-02 | 0.35 | ↓ | -1.50 | 1.2E-02 | 1.4E-01 | 0.58 | ↓ | -0.80 | 1.6E-02 | 9.4E-02 | 0.55 | ↓ | -0.87 |
| NEG | 790.5343_8.7 | PI(18:0/22:6) | 3.5E-04 | 1.5E-02 | 0.50 | ↓ | -1.00 | 3.2E-04 | 4.0E-02 | 0.55 | ↓ | -0.87 | 1.1E-03 | 2.4E-02 | 0.60 | ↓ | -0.73 |
| POS | 510.3561_5.63 | LysoPC(17:0) | 6.4E-03 | 2.9E-02 | 0.62 | ↓ | -0.68 | 3.1E-01 | 5.8E-01 | 0.85 | ↓ | -0.24 | 2.1E-02 | 1.1E-01 | 0.70 | ↓ | -0.52 |
| POS | 339.3453_7.67 | 9-Oxo-11α,15R-dihydroxyprostan-1-oic acid | 8.6E-08 | 7.5E-05 | 9.31 | ↑ | 3.22 | 4.5E-01 | 7.0E-01 | 2.53 | ↑ | 1.34 | 4.4E-02 | 1.8E-01 | 14.42 | ↑ | 3.85 |
| POS | 338.3423_7.68 | 13-Docosenamide, (Z)- | 7.7E-08 | 7.5E-05 | 7.59 | ↑ | 2.92 | 6.2E-01 | 8.2E-01 | 2.09 | ↑ | 1.07 | 5.3E-02 | 2.0E-01 | 11.15 | ↑ | 3.48 |
| NEG | 554.3457_5.63 | LysoPC(17:0) | 2.1E-03 | 2.3E-02 | 0.60 | ↓ | -0.74 | 2.9E-01 | 5.9E-01 | 0.85 | ↓ | -0.23 | 3.0E-02 | 1.5E-01 | 0.75 | ↓ | -0.42 |
| POS | 608.4631_7.89 | LysoPC(24:0) | 1.2E-03 | 1.1E-02 | 0.42 | ↓ | -1.26 | 3.9E-02 | 2.3E-01 | 0.61 | ↓ | -0.72 | 1.8E-01 | 4.3E-01 | 0.72 | ↓ | -0.47 |
| NEG | 826.5599_8.52 | PC(16:0/20:4) | 2.8E-03 | 2.6E-02 | 0.59 | ↓ | -0.75 | 4.6E-02 | 2.9E-01 | 0.78 | ↓ | -0.37 | 9.8E-02 | 3.0E-01 | 0.84 | ↓ | -0.25 |
| NEG | 596.3925_6.63 | LysoPC(20:0) | 3.8E-04 | 1.5E-02 | 0.32 | ↓ | -1.65 | 2.8E-02 | 2.5E-01 | 0.53 | ↓ | -0.91 | 1.1E-01 | 3.1E-01 | 0.69 | ↓ | -0.54 |
| POS | 770.5677_8.69 | Dihomo-γ-linolenoyl PAF C-16 | 6.9E-05 | 5.7E-03 | 0.44 | ↓ | -1.18 | 6.3E-04 | 4.0E-02 | 0.57 | ↓ | -0.81 | 5.5E-04 | 1.1E-02 | 0.49 | ↓ | -1.03 |
| POS | 532.3383_5.63 | LysoPC(17:0) | 1.1E-02 | 4.1E-02 | 0.63 | ↓ | -0.66 | 4.6E-01 | 7.1E-01 | 0.89 | ↓ | -0.17 | 2.8E-02 | 1.3E-01 | 0.69 | ↓ | -0.54 |
| NEG | 861.5499_11.73 | PI(18:1) | 2.0E-03 | 2.3E-02 | 0.46 | ↓ | -1.11 | 2.1E-02 | 2.2E-01 | 0.60 | ↓ | -0.74 | 1.3E-04 | 1.0E-02 | 0.41 | ↓ | -1.29 |
| POS | 184.0945_0.58 | PC | 7.0E-07 | 6.4E-04 | 4.73 | ↑ | 2.24 | 3.8E-04 | 4.7E-02 | 2.92 | ↑ | 1.54 | 6.9E-05 | 5.7E-03 | 4.36 | ↑ | 2.13 |
| NEG | 788.5249_8.71 | PS(18:0/18:1) | 4.1E-04 | 1.5E-02 | 0.39 | ↓ | -1.35 | 1.8E-03 | 9.1E-02 | 0.47 | ↓ | -1.09 | 1.5E-02 | 1.0E-01 | 0.59 | ↓ | -0.77 |
| POS | 518.3256_4.86 | LysoPC(16:0) | 3.5E-03 | 2.0E-02 | 0.52 | ↓ | -0.95 | 3.9E-01 | 6.6E-01 | 0.83 | ↓ | -0.27 | 9.7E-03 | 6.9E-02 | 0.52 | ↓ | -0.95 |
| NEG | 828.5667_8.52 | C18(Plasm)-20:4 PC | 1.9E-03 | 2.3E-02 | 0.52 | ↓ | -0.95 | 4.5E-02 | 2.9E-01 | 0.72 | ↓ | -0.47 | 5.6E-02 | 2.2E-01 | 0.75 | ↓ | -0.41 |
| NEG | 827.5636_8.52 | KRTLRR | 2.0E-03 | 2.3E-02 | 0.54 | ↓ | -0.89 | 3.5E-02 | 2.7E-01 | 0.74 | ↓ | -0.43 | 4.8E-02 | 2.0E-01 | 0.77 | ↓ | -0.39 |
| NEG | 526.3113_5.06 | LysoPC(15:0) | 4.7E-03 | 3.3E-02 | 0.64 | ↓ | -0.65 | 1.6E-01 | 4.6E-01 | 0.83 | ↓ | -0.27 | 3.1E-04 | 1.4E-02 | 0.60 | ↓ | -0.74 |

| | | | | | | | | | | | | | | | | | |
|---|---|---|---|---|---|---|---|---|---|---|---|---|---|---|---|---|---|
| NEG | 594.3771_6.08 | Pyrrolidino PAF C-16 | 4.7E-04 | 1.5E-02 | 0.52 | ↓ | -0.93 | 8.0E-02 | 3.5E-01 | 0.75 | ↓ | -0.42 | 2.8E-01 | 5.3E-01 | 0.86 | ↓ | -0.22 |
| POS | 510.3564_1.56 | LysoPC(17:0) | 1.2E-02 | 1.0E-01 | 0.53 | ↓ | -0.92 | 1.7E-01 | 4.2E-01 | 0.75 | ↓ | -0.41 | 2.6E-03 | 4.1E-02 | 0.45 | ↓ | -1.14 |
| POS | 482.3248_5.06 | LysoPC(15:0) | 2.4E-04 | 7.4E-03 | 0.47 | ↓ | -1.10 | 6.6E-02 | 2.8E-01 | 0.73 | ↓ | -0.45 | 9.5E-05 | 4.3E-03 | 0.51 | ↓ | -0.97 |
| NEG | 554.3463_1.56 | LysoPC(17:0) | 3.3E-02 | 6.7E-02 | 0.66 | ↓ | -0.60 | 2.2E-01 | 3.9E-01 | 0.81 | ↓ | -0.30 | 3.3E-01 | 4.4E-01 | 0.86 | ↓ | -0.22 |
| POS | 552.4037_2.62 | LysoPC(20:0) | 3.1E-03 | 5.0E-02 | 0.41 | ↓ | -1.29 | 1.6E-01 | 4.2E-01 | 0.73 | ↓ | -0.46 | 6.5E-01 | 8.5E-01 | 0.99 | ↓ | -0.01 |
| NEG | 624.4238_7.29 | LysoPC(22:0) | 5.3E-05 | 8.6E-03 | 0.31 | ↓ | -1.69 | 1.6E-02 | 2.0E-01 | 0.56 | ↓ | -0.85 | 3.7E-01 | 6.1E-01 | 0.84 | ↓ | -0.25 |
| POS | 524.3722_5.96 | LysoPC(18:0) | 2.4E-02 | 7.1E-02 | 0.76 | ↓ | -0.40 | 9.4E-01 | 9.8E-01 | 1.03 | ↑ | 0.04 | 4.3E-01 | 6.9E-01 | 1.08 | ↑ | 0.11 |
| NEG | 512.2985_4.81 | LysoPC(14:0) | 1.1E-02 | 5.4E-02 | 0.64 | ↓ | -0.65 | 8.0E-01 | 9.2E-01 | 0.99 | ↓ | -0.01 | 9.6E-04 | 2.3E-02 | 0.65 | ↓ | -0.62 |
| NEG | 652.4552_7.9 | LysoPC(24:0) | 9.7E-04 | 1.9E-02 | 0.48 | ↓ | -1.06 | 8.4E-03 | 1.6E-01 | 0.55 | ↓ | -0.87 | 1.1E-01 | 3.2E-01 | 0.72 | ↓ | -0.48 |
| NEG | 226.0175_2.4 | 4-Quinolinol, 8-methyl-2-(trifluoromethyl)- | 2.9E-03 | 2.6E-02 | 0.41 | ↓ | -1.28 | 1.6E-04 | 2.8E-02 | 0.20 | ↓ | -2.31 | 4.2E-03 | 5.3E-02 | 0.30 | ↓ | -1.74 |
| POS | 516.3104_4.97 | Oleyloxyethylphosphorylcholine | 2.7E-03 | 1.7E-02 | 0.50 | ↓ | -0.99 | 4.2E-01 | 6.8E-01 | 0.87 | ↓ | -0.21 | 9.4E-03 | 6.8E-02 | 0.62 | ↓ | -0.70 |
| POS | 162.1129_0.33 | L-Carnitine | 2.4E-07 | 1.6E-04 | 3.61 | ↑ | 1.85 | 2.2E-05 | 5.0E-03 | 2.33 | ↑ | 1.22 | 1.0E-04 | 4.3E-03 | 2.52 | ↑ | 1.33 |
| POS | 508.3397_5.21 | C18(Plasm) LPC | 3.7E-04 | 7.8E-03 | 0.53 | ↓ | -0.91 | 2.4E-01 | 5.1E-01 | 0.85 | ↓ | -0.24 | 1.2E-02 | 7.7E-02 | 0.69 | ↓ | -0.54 |
| NEG | 512.299_0.99 | LysoPC(14:0) | 2.8E-03 | 8.8E-03 | 0.62 | ↓ | -0.70 | 4.7E-01 | 6.1E-01 | 0.92 | ↓ | -0.12 | 1.8E-02 | 4.1E-02 | 0.72 | ↓ | -0.47 |
| NEG | 816.5759_9.01 | PC(16:0/20:4) | 7.7E-03 | 4.3E-02 | 0.67 | ↓ | -0.58 | 6.3E-03 | 1.4E-01 | 0.65 | ↓ | -0.63 | 7.6E-03 | 6.9E-02 | 0.62 | ↓ | -0.70 |
| POS | 204.1231_0.58 | Acetyl-DL-carnitine | 1.2E-02 | 1.0E-01 | 2.33 | ↑ | 1.22 | 1.7E-01 | 4.2E-01 | 1.56 | ↑ | 0.64 | 2.1E-03 | 3.8E-02 | 5.35 | ↑ | 2.42 |
| POS | 482.3253_1.13 | LysoPE(18:0) | 6.4E-03 | 7.2E-02 | 0.51 | ↓ | -0.98 | 5.9E-02 | 3.9E-01 | 0.70 | ↓ | -0.51 | 1.2E-04 | 7.4E-03 | 0.37 | ↓ | -1.44 |
| POS | 468.3092_4.81 | LysoPC(14:0) | 2.8E-03 | 1.7E-02 | 0.59 | ↓ | -0.77 | 9.9E-01 | 1.0E+00 | 1.02 | ↑ | 0.03 | 2.0E-03 | 2.4E-02 | 0.63 | ↓ | -0.66 |
| POS | 782.5682_4.96 | PC(16:0/18:1) | 7.5E-03 | 8.1E-02 | 2.25 | ↑ | 1.17 | 6.5E-02 | 3.9E-01 | 2.48 | ↑ | 1.31 | 7.9E-04 | 2.5E-02 | 3.72 | ↑ | 1.89 |
| POS | 530.368_6.36 | C18(Plasm) LPC | 6.7E-03 | 3.0E-02 | 3.12 | ↑ | 1.64 | 1.8E-03 | 6.2E-02 | 5.80 | ↑ | 2.54 | 5.0E-03 | 4.3E-02 | 4.50 | ↑ | 2.17 |
| POS | 530.3694_1.4 | C18(Plasm) LPC | 1.8E-03 | 3.6E-02 | 4.36 | ↑ | 2.12 | 8.7E-04 | 7.6E-02 | 7.14 | ↑ | 2.84 | 1.6E-02 | 9.8E-02 | 5.51 | ↑ | 2.46 |
| NEG | 526.31_1.14 | LysoPC(15:0) | 1.3E-04 | 3.1E-03 | 0.59 | ↓ | -0.76 | 1.3E-02 | 2.8E-01 | 0.75 | ↓ | -0.42 | 1.0E-02 | 2.6E-02 | 0.67 | ↓ | -0.59 |
| NEG | 229.1437_3.65 | Dodecanedioic acid | 1.4E-02 | 6.6E-02 | 0.53 | ↓ | -0.91 | 3.4E-04 | 4.0E-02 | 0.49 | ↓ | -1.04 | 2.4E-03 | 3.5E-02 | 0.54 | ↓ | -0.89 |
| POS | 790.6208_10.03 | PS(18:0/18:1) | 2.0E-02 | 6.3E-02 | 0.80 | ↓ | -0.32 | 1.6E-01 | 4.2E-01 | 0.87 | ↓ | -0.20 | 5.1E-02 | 1.9E-01 | 0.84 | ↓ | -0.26 |
| POS | 204.1141_0.33 | Acetyl-DL-carnitine | 2.1E-02 | 6.3E-02 | 1.78 | ↑ | 0.83 | 2.2E-01 | 4.9E-01 | 1.43 | ↑ | 0.52 | 1.0E-04 | 4.3E-03 | 2.98 | ↑ | 1.57 |
| POS | 316.2485_3.86 | Decanoyl-L-carnitine | 9.5E-02 | 1.9E-01 | 2.39 | ↑ | 1.26 | 1.0E+00 | 1.0E+00 | 1.58 | ↑ | 0.66 | 1.1E-02 | 7.3E-02 | 3.50 | ↑ | 1.81 |
| POS | 468.3099_0.99 | LysoPC(14:0) | 1.6E-02 | 1.1E-01 | 0.57 | ↓ | -0.81 | 3.8E-01 | 5.7E-01 | 0.88 | ↓ | -0.18 | 3.1E-05 | 2.7E-03 | 0.47 | ↓ | -1.08 |
| NEG | 584.3554_4.45 | 2-Thio PAF | 1.5E-02 | 6.8E-02 | 0.54 | ↓ | -0.88 | 7.9E-01 | 9.2E-01 | 1.00 | ↓ | -0.01 | 6.4E-02 | 2.4E-01 | 0.72 | ↓ | -0.47 |
| NEG | 178.0488_0.62 | Hippuric acid | 1.6E-01 | 2.5E-01 | 0.72 | ↓ | -0.47 | 1.2E-01 | 3.0E-01 | 0.61 | ↓ | -0.70 | 3.8E-01 | 4.8E-01 | 1.17 | ↑ | 0.23 |
| POS | 728.5584_10.1 | PE(18:0/18:1) | 1.6E-02 | 5.5E-02 | 0.53 | ↓ | -0.91 | 4.0E-02 | 2.3E-01 | 0.60 | ↓ | -0.73 | 2.6E-03 | 2.8E-02 | 0.40 | ↓ | -1.31 |
| NEG | 452.2775_0.99 | LysoPE(16:0) | 2.2E-04 | 3.1E-03 | 0.57 | ↓ | -0.81 | 6.4E-01 | 7.4E-01 | 0.96 | ↓ | -0.05 | 7.0E-04 | 3.1E-03 | 0.65 | ↓ | -0.63 |
| POS | 257.2267_3.76 | Androst-16-en-3-ol, (3β,5α)- | 7.1E-02 | 1.5E-01 | 1.96 | ↑ | 0.97 | 8.1E-02 | 3.1E-01 | 1.88 | ↑ | 0.91 | 1.2E-01 | 3.4E-01 | 1.59 | ↑ | 0.67 |
| NEG | 862.5542_4.81 | C18(Plasm)-22:6 PC | 6.9E-03 | 1.7E-02 | 3.24 | ↑ | 1.70 | 5.2E-02 | 2.8E-01 | 5.75 | ↑ | 2.52 | 1.8E-02 | 4.2E-02 | 5.08 | ↑ | 2.34 |
| POS | 806.5689_4.84 | Arachidonoylthio-PC | 2.5E-02 | 1.4E-01 | 1.89 | ↑ | 0.92 | 1.1E-01 | 4.0E-01 | 2.13 | ↑ | 1.09 | 1.4E-03 | 3.2E-02 | 2.76 | ↑ | 1.47 |
| NEG | 263.1033_1.9 | L-Glutamine, N2-(2-phenylacetyl)- | 4.7E-03 | 3.3E-02 | 0.43 | ↓ | -1.22 | 1.9E-03 | 9.2E-02 | 0.33 | ↓ | -1.59 | 3.9E-02 | 1.7E-01 | 0.52 | ↓ | -0.95 |
| POS | 504.3088_1.14 | Lysophosphatidylethanolamine | 1.0E-02 | 9.3E-02 | 0.56 | ↓ | -0.84 | 2.4E-02 | 3.9E-01 | 0.68 | ↓ | -0.56 | 1.7E-05 | 2.0E-03 | 0.37 | ↓ | -1.43 |
| POS | 524.3717_1.89 | LysoPC(18:0) | 4.6E-02 | 1.6E-01 | 0.69 | ↓ | -0.53 | 8.4E-01 | 8.9E-01 | 1.03 | ↑ | 0.05 | 1.2E-01 | 3.3E-01 | 0.79 | ↓ | -0.34 |
| NEG | 830.5807_5.1 | PC(18:0/18:2) | 9.4E-04 | 4.8E-03 | 2.86 | ↑ | 1.52 | 2.2E-01 | 3.9E-01 | 2.76 | ↑ | 1.46 | 2.9E-04 | 1.9E-03 | 5.10 | ↑ | 2.35 |
| POS | 544.338_1.41 | LysoPC(18:1) | 6.9E-04 | 2.8E-02 | 0.57 | ↓ | -0.81 | 1.8E-01 | 4.2E-01 | 0.84 | ↓ | -0.26 | 8.3E-02 | 2.7E-01 | 0.83 | ↓ | -0.27 |
| POS | 214.1134_3.03 | Suberylglycine | 1.8E-01 | 3.0E-01 | 1.17 | ↑ | 0.22 | 2.6E-10 | 7.0E-07 | 8.08 | ↑ | 3.01 | 2.7E-10 | 7.1E-07 | 7.78 | ↑ | 2.96 |
| NEG | 768.5551_5.11 | PC(18:0/18:2) | 5.0E-04 | 4.1E-03 | 2.73 | ↑ | 1.45 | 2.0E-01 | 3.8E-01 | 2.30 | ↑ | 1.20 | 3.5E-04 | 2.0E-03 | 4.34 | ↑ | 2.12 |
| POS | 130.0499_1.9 | Glutamic Acid | 3.1E-04 | 7.8E-03 | 0.36 | ↓ | -1.49 | 3.3E-04 | 2.8E-02 | 0.31 | ↓ | -1.69 | 1.5E-02 | 9.1E-02 | 0.48 | ↓ | -1.05 |
| POS | 490.2926_0.99 | LysoPC(14:0) | 2.1E-01 | 3.4E-01 | 0.75 | ↓ | -0.42 | 5.6E-01 | 7.0E-01 | 0.89 | ↓ | -0.17 | 3.9E-03 | 5.0E-02 | 0.47 | ↓ | -1.08 |

| | | | | | | | | | | | | | | | | | |
|---|---|---|---|---|---|---|---|---|---|---|---|---|---|---|---|---|---|
| POS | 788.6068_5.47 | PC(18:0/18:1) | 9.5E-02 | 2.1E-01 | 1.69 | ↑ | 0.75 | 2.9E-01 | 5.1E-01 | 1.97 | ↑ | 0.98 | 2.3E-02 | 1.2E-01 | 2.43 | ↑ | 1.28 |
| POS | 287.1015_0.61 | Phenylacetyl-L-glutamine | 1.1E-03 | 3.0E-02 | 0.42 | ↓ | -1.25 | 3.7E-04 | 4.7E-02 | 0.34 | ↓ | -1.54 | 1.5E-02 | 9.4E-02 | 0.50 | ↓ | -1.00 |
| POS | 786.6003_5.47 | 1,2-Dioleoyl-sn-glycero-3-PC | 9.7E-02 | 2.1E-01 | 1.72 | ↑ | 0.78 | 2.5E-01 | 4.8E-01 | 2.10 | ↑ | 1.07 | 2.1E-02 | 1.2E-01 | 2.55 | ↑ | 1.35 |
| POS | 532.3389_1.56 | LysoPC(17:0) | 4.1E-03 | 5.6E-02 | 0.43 | ↓ | -1.23 | 5.5E-02 | 3.9E-01 | 0.61 | ↓ | -0.71 | 1.1E-04 | 7.4E-03 | 0.35 | ↓ | -1.51 |
| POS | 794.5989_5.26 | C18(Plasm)-20:4 PC | 3.9E-02 | 1.6E-01 | 2.10 | ↑ | 1.07 | 9.6E-02 | 4.0E-01 | 2.71 | ↑ | 1.44 | 2.8E-03 | 4.2E-02 | 3.54 | ↑ | 1.82 |
| POS | 421.2528_5.75 | Tris(2-butoxyethyl) phosphate | 5.5E-04 | 8.3E-03 | 2.11 | ↑ | 1.08 | 2.9E-03 | 7.8E-02 | 3.26 | ↑ | 1.71 | 4.5E-03 | 4.0E-02 | 2.58 | ↑ | 1.37 |
| POS | 180.0972_0.33 | N-Benzylglycine methyl ester | 1.3E-02 | 4.8E-02 | 0.48 | ↓ | -1.07 | 7.6E-02 | 3.0E-01 | 0.60 | ↓ | -0.73 | 1.9E-02 | 1.0E-01 | 0.43 | ↓ | -1.23 |
| NEG | 826.5604_4.96 | PC(16:0/20:4) | 1.4E-04 | 3.1E-03 | 3.61 | ↑ | 1.85 | 4.7E-02 | 2.8E-01 | 3.12 | ↑ | 1.64 | 2.6E-05 | 6.9E-04 | 5.19 | ↑ | 2.38 |
| POS | 637.3064_1.8 | Acetylpodocarpic acid anhydride | 6.3E-01 | 7.5E-01 | 1.13 | ↑ | 0.17 | 3.9E-05 | 9.6E-03 | 2.39 | ↑ | 1.26 | 4.5E-08 | 3.3E-05 | 2.83 | ↑ | 1.50 |
| POS | 808.5841_5 | 1,2-Dioleoyl-sn-glycero-3-PC | 8.7E-02 | 2.0E-01 | 1.54 | ↑ | 0.62 | 2.1E-01 | 4.5E-01 | 1.88 | ↑ | 0.91 | 4.1E-03 | 5.1E-02 | 3.07 | ↑ | 1.62 |
| NEG | 766.5393_4.96 | PE(18:0/20:4) | 8.0E-05 | 3.1E-03 | 3.86 | ↑ | 1.95 | 3.1E-02 | 2.8E-01 | 3.58 | ↑ | 1.84 | 1.0E-05 | 5.2E-04 | 5.55 | ↑ | 2.47 |
| NEG | 835.5345_5.54 | PI(16:0/18:1) | 7.3E-04 | 4.7E-03 | 3.77 | ↑ | 1.91 | 1.4E-01 | 3.1E-01 | 3.18 | ↑ | 1.67 | 3.0E-03 | 9.6E-03 | 4.22 | ↑ | 2.08 |
| NEG | 850.5605_4.85 | PC(16;0/22:6) | 2.8E-04 | 3.1E-03 | 4.04 | ↑ | 2.01 | 7.6E-02 | 2.8E-01 | 3.91 | ↑ | 1.97 | 9.1E-06 | 5.2E-04 | 5.96 | ↑ | 2.57 |
| POS | 808.5769_4.83 | 1,2-Dioleoyl-sn-glycero-3-PC | 6.0E-02 | 1.8E-01 | 1.57 | ↑ | 0.65 | 1.5E-01 | 4.1E-01 | 1.73 | ↑ | 0.79 | 3.0E-03 | 4.3E-02 | 2.28 | ↑ | 1.19 |
| POS | 502.3262_1.49 | C16(Plasm) LPC | 6.4E-02 | 1.8E-01 | 0.63 | ↓ | -0.66 | 6.0E-01 | 7.3E-01 | 1.18 | ↑ | 0.24 | 5.0E-02 | 1.9E-01 | 0.64 | ↓ | -0.63 |
| POS | 678.4682_6.87 | PC(14:0) | 3.2E-03 | 1.9E-02 | 0.46 | ↓ | -1.12 | 2.2E-02 | 1.8E-01 | 0.52 | ↓ | -0.94 | 4.9E-05 | 3.2E-03 | 0.31 | ↓ | -1.67 |
| POS | 546.3536_1.89 | LysoPC(18:0) | 1.2E-02 | 1.0E-01 | 0.74 | ↓ | -0.43 | 1.3E-01 | 4.0E-01 | 0.87 | ↓ | -0.20 | 1.5E-01 | 3.7E-01 | 0.88 | ↓ | -0.19 |
| NEG | 770.571_5.48 | DMPE(18:1/18:1) | 1.9E-03 | 7.1E-03 | 3.19 | ↑ | 1.67 | 9.4E-02 | 2.8E-01 | 3.80 | ↑ | 1.93 | 6.0E-04 | 2.8E-03 | 5.21 | ↑ | 2.38 |
| NEG | 832.5981_5.48 | PC(18:1/18:0) | 9.1E-04 | 4.7E-03 | 3.53 | ↑ | 1.82 | 9.4E-02 | 2.8E-01 | 3.71 | ↑ | 1.89 | 6.2E-04 | 2.9E-03 | 5.04 | ↑ | 2.33 |
| POS | 792.5895_5.18 | Dihomo-γ-linolenoyl PAF C-16 | 6.1E-02 | 1.8E-01 | 2.23 | ↑ | 1.15 | 1.5E-01 | 4.1E-01 | 2.99 | ↑ | 1.58 | 4.9E-03 | 5.6E-02 | 3.52 | ↑ | 1.81 |
| NEG | 794.5685_5.41 | PC(18:1/16:0) | 6.4E-04 | 4.6E-03 | 4.01 | ↑ | 2.00 | 7.8E-02 | 2.8E-01 | 3.78 | ↑ | 1.92 | 2.3E-04 | 1.7E-03 | 6.48 | ↑ | 2.70 |
| POS | 750.5453_5.35 | Palmitoyl Thio-PC | 5.9E-02 | 1.7E-01 | 2.48 | ↑ | 1.31 | 1.4E-01 | 4.0E-01 | 2.72 | ↑ | 1.44 | 4.2E-03 | 5.2E-02 | 3.43 | ↑ | 1.78 |
| POS | 810.5994_5.42 | PC(18:0/18:1) | 2.0E-02 | 1.3E-01 | 2.21 | ↑ | 1.15 | 1.1E-01 | 4.0E-01 | 2.38 | ↑ | 1.25 | 2.6E-03 | 4.1E-02 | 3.79 | ↑ | 1.92 |
| NEG | 832.6078_5.77 | PC(18:1/18:0) | 1.3E-03 | 5.6E-03 | 4.96 | ↑ | 2.31 | 6.2E-02 | 2.8E-01 | 5.91 | ↑ | 2.56 | 1.0E-03 | 4.1E-03 | 7.14 | ↑ | 2.84 |
| NEG | 830.5917_5.48 | PC(18:1/18:1) | 8.1E-04 | 4.7E-03 | 3.18 | ↑ | 1.67 | 9.3E-02 | 2.8E-01 | 3.22 | ↑ | 1.69 | 2.9E-04 | 1.9E-03 | 4.42 | ↑ | 2.14 |
| NEG | 747.5162_5.21 | SM(d18:1/16:0) | 1.8E-03 | 6.7E-03 | 9.60 | ↑ | 3.26 | 6.0E-02 | 2.8E-01 | 12.17 | ↑ | 3.61 | 8.6E-06 | 5.2E-04 | 11.38 | ↑ | 3.51 |
| NEG | 762.5082_4.98 | PE(16:0/22:6) | 1.6E-03 | 6.1E-03 | 5.31 | ↑ | 2.41 | 1.7E-01 | 3.4E-01 | 4.46 | ↑ | 2.16 | 2.8E-03 | 9.0E-03 | 5.86 | ↑ | 2.55 |
| POS | 790.5725_5.24 | Arachidonoyl PAF C-16 | 2.0E-02 | 1.3E-01 | 2.43 | ↑ | 1.28 | 5.2E-02 | 3.9E-01 | 2.78 | ↑ | 1.47 | 5.6E-03 | 6.0E-02 | 3.57 | ↑ | 1.84 |
| NEG | 765.5276_5.16 | SM(d18:1/18:0) | 2.7E-03 | 8.8E-03 | 3.41 | ↑ | 1.77 | 2.0E-01 | 3.8E-01 | 2.87 | ↑ | 1.52 | 4.2E-03 | 1.2E-02 | 3.90 | ↑ | 1.96 |
| NEG | 820.577_5.49 | PC(18:0/18:2) | 2.6E-03 | 8.6E-03 | 2.74 | ↑ | 1.45 | 8.6E-02 | 2.8E-01 | 3.20 | ↑ | 1.68 | 4.8E-04 | 2.4E-03 | 4.03 | ↑ | 2.01 |
| NEG | 790.5397_5.44 | PE(18:0/22:6) | 8.6E-04 | 4.7E-03 | 7.45 | ↑ | 2.90 | 7.8E-02 | 2.8E-01 | 6.34 | ↑ | 2.66 | 1.3E-03 | 5.1E-03 | 6.73 | ↑ | 2.75 |
| NEG | 806.5944_5.72 | PC(16:0/18:0) | 3.3E-04 | 3.4E-03 | 5.61 | ↑ | 2.49 | 7.8E-02 | 2.8E-01 | 5.20 | ↑ | 2.38 | 5.8E-04 | 2.7E-03 | 5.51 | ↑ | 2.46 |
| NEG | 790.541_4.84 | PE(18:0/22:6) | 1.1E-03 | 5.2E-03 | 3.21 | ↑ | 1.68 | 1.2E-01 | 3.0E-01 | 3.43 | ↑ | 1.78 | 7.5E-05 | 8.6E-04 | 4.80 | ↑ | 2.26 |
| NEG | 878.5916_5.31 | PC(18:0/22:6) | 1.1E-03 | 5.0E-03 | 4.18 | ↑ | 2.06 | 8.0E-02 | 2.8E-01 | 4.78 | ↑ | 2.26 | 8.5E-05 | 9.1E-04 | 6.54 | ↑ | 2.71 |
| NEG | 766.5395_5.55 | PE(18:0/24:0) | 5.1E-04 | 4.1E-03 | 5.99 | ↑ | 2.58 | 7.9E-02 | 2.8E-01 | 4.97 | ↑ | 2.31 | 2.8E-03 | 9.0E-03 | 6.31 | ↑ | 2.66 |
| POS | 766.5724_5.16 | Eicosapentaenoyl PAF C-16 | 7.9E-02 | 1.9E-01 | 2.11 | ↑ | 1.08 | 1.2E-01 | 4.0E-01 | 2.64 | ↑ | 1.40 | 6.8E-03 | 6.6E-02 | 3.45 | ↑ | 1.79 |
| POS | 760.5848_5.35 | PC(16:0/18:1) | 4.3E-02 | 1.6E-01 | 2.25 | ↑ | 1.17 | 8.1E-02 | 3.9E-01 | 3.12 | ↑ | 1.64 | 9.3E-03 | 7.6E-02 | 3.02 | ↑ | 1.59 |
| NEG | 776.5592_5.71 | PC(18:1/14:0) | 6.8E-04 | 4.6E-03 | 8.17 | ↑ | 3.03 | 4.8E-02 | 2.8E-01 | 8.09 | ↑ | 3.02 | 7.8E-05 | 8.7E-04 | 9.97 | ↑ | 3.32 |
| NEG | 768.5463_5.54 | PC(18:0/14:0) | 6.0E-04 | 4.5E-03 | 4.99 | ↑ | 2.32 | 5.0E-02 | 2.8E-01 | 4.40 | ↑ | 2.14 | 2.8E-03 | 9.0E-03 | 5.56 | ↑ | 2.47 |
| POS | 782.567_5.35 | PC(16:0/18:1) | 3.8E-02 | 1.6E-01 | 1.71 | ↑ | 0.78 | 6.8E-02 | 3.9E-01 | 1.85 | ↑ | 0.88 | 1.2E-02 | 8.6E-02 | 1.93 | ↑ | 0.94 |
| NEG | 838.5972_5.6 | C18(Plasm)-20:4 PC | 5.4E-04 | 4.3E-03 | 6.87 | ↑ | 2.78 | 3.4E-02 | 2.8E-01 | 6.82 | ↑ | 2.77 | 7.5E-05 | 8.6E-04 | 8.61 | ↑ | 3.11 |
| NEG | 804.5669_5.01 | PC(18:1/16:0) | 4.0E-04 | 3.8E-03 | 3.22 | ↑ | 1.69 | 8.4E-02 | 2.8E-01 | 2.97 | ↑ | 1.57 | 9.6E-05 | 9.7E-04 | 3.99 | ↑ | 2.00 |
| NEG | 802.5603_5.01 | PC(16:0/18:2) | 2.5E-04 | 3.1E-03 | 2.73 | ↑ | 1.45 | 8.0E-02 | 2.8E-01 | 2.52 | ↑ | 1.33 | 4.7E-05 | 7.9E-04 | 3.40 | ↑ | 1.77 |

| | | | | | | | | | | | | | | | | | |
|---|---|---|---|---|---|---|---|---|---|---|---|---|---|---|---|---|---|
| NEG | 744.549_5.01 | α-Galactosyl-N-palmitoylsphingosine | 1.4E-04 | 3.1E-03 | 3.81 | ↑ | 1.93 | 1.3E-01 | 3.1E-01 | 3.21 | ↑ | 1.68 | 2.3E-04 | 1.7E-03 | 3.05 | ↑ | 1.61 |
| POS | 768.5893_5.23 | C16-18:1 PC | 2.9E-02 | 1.5E-01 | 2.42 | ↑ | 1.28 | 5.6E-02 | 3.9E-01 | 3.28 | ↑ | 1.71 | 4.9E-03 | 5.6E-02 | 3.81 | ↑ | 1.93 |
| NEG | 774.545_5.65 | C18(Plasm)-22:6 PE | 2.8E-03 | 8.9E-03 | 10.79 | ↑ | 3.43 | 7.5E-02 | 2.8E-01 | 12.12 | ↑ | 3.60 | 2.8E-05 | 6.9E-04 | 9.96 | ↑ | 3.32 |
| NEG | 714.5081_5.16 | PE(16:0/18:2) | 6.2E-04 | 4.6E-03 | 4.03 | ↑ | 2.01 | 7.9E-02 | 2.8E-01 | 3.42 | ↑ | 1.77 | 2.2E-02 | 5.0E-02 | 2.93 | ↑ | 1.55 |
| NEG | 792.5473_5.01 | PC(16:0/18:2) | 4.4E-04 | 3.9E-03 | 3.06 | ↑ | 1.61 | 1.3E-01 | 3.1E-01 | 2.87 | ↑ | 1.52 | 1.7E-05 | 6.3E-04 | 4.31 | ↑ | 2.11 |
| POS | 735.5737_5.29 | Thioetheramidephosphatidylcholine | 2.1E-02 | 1.3E-01 | 3.30 | ↑ | 1.72 | 7.9E-02 | 3.9E-01 | 3.56 | ↑ | 1.83 | 1.5E-02 | 9.4E-02 | 3.23 | ↑ | 1.69 |
| POS | 756.5531_5.29 | PC(16:0/16:0) | 1.1E-02 | 9.4E-02 | 2.75 | ↑ | 1.46 | 8.4E-02 | 4.0E-01 | 2.05 | ↑ | 1.03 | 2.3E-02 | 1.2E-01 | 2.03 | ↑ | 1.02 |
| POS | 184.0734_5 | PC | 4.8E-02 | 1.6E-01 | 1.97 | ↑ | 0.98 | 1.4E-01 | 4.0E-01 | 2.41 | ↑ | 1.27 | 4.4E-03 | 5.2E-02 | 2.88 | ↑ | 1.53 |
| NEG | 806.5831_5.35 | PC(16:0/18:0) | 2.5E-04 | 3.1E-03 | 5.21 | ↑ | 2.38 | 3.4E-02 | 2.8E-01 | 5.62 | ↑ | 2.49 | 1.8E-04 | 1.4E-03 | 5.88 | ↑ | 2.55 |
| POS | 834.5998_5.31 | 1-Docosahexaenoyl-2-stearoyl-sn-glycero-3-p | 6.8E-02 | 1.8E-01 | 1.96 | ↑ | 0.97 | 1.6E-01 | 4.1E-01 | 2.81 | ↑ | 1.49 | 5.3E-03 | 5.8E-02 | 3.29 | ↑ | 1.72 |
| NEG | 834.5429_5.53 | PS(18:0/22:6) | 7.4E-04 | 4.7E-03 | 4.04 | ↑ | 2.01 | 1.4E-01 | 3.2E-01 | 3.22 | ↑ | 1.69 | 3.2E-03 | 1.0E-02 | 4.46 | ↑ | 2.16 |
| NEG | 742.5398_5.01 | PE(18:0/18:2) | 9.9E-05 | 3.1E-03 | 3.22 | ↑ | 1.69 | 1.1E-01 | 2.9E-01 | 2.97 | ↑ | 1.57 | 7.4E-05 | 8.6E-04 | 3.71 | ↑ | 1.89 |
| NEG | 862.5958_5.51 | C18(Plasm)-22:6 PC | 6.0E-03 | 1.6E-02 | 5.91 | ↑ | 2.56 | 1.1E-01 | 2.9E-01 | 5.15 | ↑ | 2.36 | 1.0E-04 | 1.0E-03 | 5.83 | ↑ | 2.54 |
| POS | 758.5692_5.01 | PC(16:0/18:2) | 3.1E-02 | 1.5E-01 | 1.91 | ↑ | 0.93 | 1.4E-01 | 4.0E-01 | 2.17 | ↑ | 1.12 | 4.8E-03 | 5.6E-02 | 2.49 | ↑ | 1.32 |
| POS | 760.5746_5.01 | PC(16:0/18:1) | 3.1E-02 | 1.5E-01 | 1.83 | ↑ | 0.87 | 1.5E-01 | 4.1E-01 | 2.02 | ↑ | 1.01 | 4.5E-03 | 5.3E-02 | 2.35 | ↑ | 1.23 |
| NEG | 804.576_5.36 | PC(18:1/16:0) | 6.8E-04 | 4.6E-03 | 4.06 | ↑ | 2.02 | 5.0E-02 | 2.8E-01 | 4.11 | ↑ | 2.04 | 3.3E-04 | 2.0E-03 | 4.93 | ↑ | 2.30 |
| POS | 768.5582_5.55 | PE(18:0/18:1) | 3.7E-02 | 1.6E-01 | 2.13 | ↑ | 1.09 | 2.2E-01 | 4.6E-01 | 1.73 | ↑ | 0.79 | 7.0E-02 | 2.4E-01 | 2.55 | ↑ | 1.35 |
| NEG | 738.5085_5.1 | PE(16:0/20:4) | 2.0E-03 | 7.3E-03 | 3.62 | ↑ | 1.86 | 1.3E-01 | 3.1E-01 | 3.84 | ↑ | 1.94 | 6.1E-03 | 1.7E-02 | 4.82 | ↑ | 2.27 |
| NEG | 802.5603_8.81 | PC(16:0/18:2) | 5.5E-03 | 3.5E-02 | 1.15 | ↑ | 0.21 | 1.1E-01 | 3.9E-01 | 1.13 | ↑ | 0.17 | 1.9E-02 | 1.1E-01 | 1.14 | ↑ | 0.18 |

### Supplementary Table 2: Fragmentation spectra details for metabolite identification.

| Precursor m/z | Retention Time | Name | Synonyms | Adduct | Level | CID |
|---|---|---|---|---|---|---|
| 861.5497 | 12.17 | 1,2-Dioleoyl-sn-glycero-3-phospho-(1'-myo-inositol) | PI(18:1) | [M-H]- | 1 | 861.54, 581.30, 419.24, 283.24, 241.01 |
| 790.5343 | 8.7 | 2-Docosahexaenoyl-1-stearoyl-sn-glycero-3-phosphoethanolamine | PI(18:0/22:6) | [M-H]- | 1 | 790.53, 480.30, 328.23, 327.23, 283.26 |
| 554.3457 | 5.63 | 1-Heptadecanoyl-sn-glycero-3-phosphocholine | LysoPC(17:0) | [M+HCO2]- | 1 | 554.34, 494.32, 269.24, 224.06, 153.99 |
| 826.5599 | 8.52 | 1-Palmitoyl-2-arachidonoyl-sn-glycero-3-phosphocholine | PC(16:0/20:4) | [M+HCO2]- | 1 | 826.5588, 766.5376, 528.3017, 480.3083, 462,.3005, 303.2330, 259.2424, 255.2332, 168.0423 |
| 596.3925 | 6.63 | 1-Arachidoyl-2-hydroxy-sn-glycero-3-phosphocholine | LysoPC(20:0) | [M+HCO2]- | 1 | 596.4027, 536.3759, 311.2951 |
| 861.5499 | 11.73 | 1,2-Dioleoyl-sn-glycero-3-phospho-(1'-myo-inositol) | PI(18:1) | [M-H]- | 1 | 861.5499, 581.2989, 419.2547, 283.2646, 241.0110 |
| 788.5249 | 8.71 | 2-Oleoyl-1-stearoyl-sn-glycero-3-phosphoserine | PS(18:0/18:1) | [M-H]- | 3 | 788.5262, 284.2446, 283.2432, 282.2486, 281.2482 |
| 828.5667 | 8.52 | 1-(1Z-Octadecenyl)-2-(5Z,8Z,11Z,14Z-eicosatetraenoyl)-sn-glycero-3-phosphocholine | C18(Plasm)-20:4 PC | [M+Cl]- | 2 | 828.5733, 303.2833, 259.2435, 224.0701 |
| 827.5636 | 8.52 | Lys-Arg-Thr-Leu-Arg-Arg | KRTLRR | [M-H]- | 1 | 827.5632, 508.3405, 482.3212, 304.2364, 301.2209, 283.2632, 282.2504, 279.2319, 259.2418, 224.0681 168.0414 |

| | | | | | | |
|---|---|---|---|---|---|---|
| 526.3113 | 5.06 | 1-Pentadecanoyl-sn-glycero-3-phosphocholine | LysoPC(15:0) | [M+HCO2]- | 1 | 526.3107, 466.2954, 377.2493, 224.0689, 168.0398 |
| 594.3771 | 6.08 | Pyrrolidinium, 1-[(7R)-7-(acetyloxy)-4-hydroxy-4-oxido-3,5,9-trioxa-4-phosphapentacos-1-yl]-1-methyl-, inner salt | Pyrrolidino PAF C-16 | [M+HCO2]- | 3 | 594.3757, 535.3633, 534.3548 |
| 624.4238 | 7.29 | 1-Behenoyl-2-hydroxy-sn-glycero-3-phosphocholine | LysoPC(22:0) | [M+HCO2]- | 2 | 624.42, 564.40, 339.32 |
| 512.2985 | 4.81 | 1-Myristoyl-sn-glycero-3-phosphocholine | LysoPC(14:0) | [M+HCO2]- | 1 | 512.29, 452.27, 227.20, 224.06 |
| 652.4552 | 7.9 | 1-Lignoceroyl-2-hydroxy-sn-glycero-3-phosphocholine | LysoPC(24:0) | [M+HCO2]- | 1 | 652.45, 592.43, 367.35, 349.34 |
| 226.0175 | 2.4 | 8-Methyl-2-(trifluoromethyl)quinolin-4-ol | 4-Quinolinol, 8-methyl-2-(trifluoromethyl)- | [M-H]- | 3 | 226.0162, 146.0600, 131.0356, 122.0245 |
| 816.5759 | 9.01 | 1-Palmitoyl-2-arachidonoyl-sn-glycero-3-phosphocholine | PC(16:0/20:4) | [M+Cl]- | 3 | 816.5747, 766.5404, 303.2336 |
| 229.1437 | 3.65 | 1,10-Decanedicarboxylic acid | Dodecanedioic acid | [M-H]- | 1 | 229.14, 211.13, 167.14, 83.05, 81.03 |
| 584.3554 | 4.45 | 1-O-Hexadecyl-2-deoxy-2-thio-S-acetyl-sn-glyceryl-3-phosphorylcholine | 2-Thio PAF | [M+HCO2]- | 3 | 584.34, 524.32, 168.04 |
| 263.1033 | 1.9 | Phenylacetyl-L-glutamine | L-Glutamine, N2-(2-phenylacetyl)- | [M-H]- | 1 | 263.10, 145.06, 128.04, 127.05 |
| 802.5603 | 8.81 | 1-Palmitoyl-2-linoleoyl-sn-glycero-3-phosphocholine | PC(16:0/18:2) | [M+HCO2]- | 1 | 802.5592, 742.5381, 480.3095, 391.2241, 279.2324, 255.2321, 224.0702, 168.0421 |
| 580.432 | 7.28 | 1-Behenoyl-2-hydroxy-sn-glycero-3-phosphocholine | LysoPC(22:0) | [M+H]+ | 1 | 580.43, 184.07, 104.11 |

| | | | | | | |
|---|---|---|---|---|---|---|
| 552.4031 | 6.62 | 1-Arachidoyl-2-hydroxy-sn-glycero-3-phosphocholine | LysoPC(20:0) | [M+H]+ | 1 | 552.40, 534.39, 184.07, 104.11 |
| 782.5682 | 8.51 | 2-Oleoyl-1-palmitoyl-sn-glycero-3-phosphocholine | PC(16:0/18:1) | [M+Na]+ | 1 | 782.56, 723.49, 599.50, 184.077, 125.00, 86.09 |
| 510.3561 | 5.63 | 1-Heptadecanoyl-sn-glycero-3-phosphocholine | LysoPC(17:0) | [M+H]+ | 1 | 510.35, 492.34, 433.27, 327.28, 258.11, 184.07, 125.00, 104.11, 886.09 |
| 339.3453 | 7.67 | 13,14-Dihydro-15(R)-prostaglandin E1 | 9-Oxo-11α,15R-dihydroxyprostan-1-oic acid | [M+H-H2O]+ | 2 | 339.34, 304.31, 223.13, 135.11, 121.10, 109.10 |
| 338.3423 | 7.68 | Erucamide | 13-Docosenamide, (Z)- | [M+H]+ | 1 | 338.34, 321.31, 303.30, 268.26, 149.13, 97.10 |
| 608.4631 | 7.89 | 1-Lignoceroyl-2-hydroxy-sn-glycero-3-phosphocholine | LysoPC(24:0) | [M+H]+ | 1 | 608.45, 590.45, 184.07, 104.11 |
| 770.5677 | 8.69 | 1-Hexadecyl-2-(8Z,11Z,14Z-eicosatrienoyl)-sn-glycero-3-phosphocholine | Dihomo-γ-linolenoyl PAF C-16 | [M+H]+ | 1 | 770.60, 184.07, 125.00, 104.11, 86.09 |
| 532.3383 | 5.63 | 1-Heptadecanoyl-sn-glycero-3-phosphocholine | LysoPC(17:0) | [M+Na]+ | 1 | 532.33, 473.26, 327.28, 147.00, 104.11 |
| 518.3256 | 4.86 | 1-Palmitoyl-sn-glycero-3-phosphocholine | LysoPC(16:0) | [M+Na]+ | 1 | 518.32, 459.24, 415.22, 313.27, 146.98, 104.11, 86.09 |
| 482.3248 | 5.06 | 1-Pentadecanoyl-sn-glycero-3-phosphocholine | LysoPC(15:0) | [M+H]+ | 1 | 482.31, 464.31, 299.25, 258.11, 184.07, 166.06, 125.00, 104.10, 95.09, 86.09, 71.07 |
| 524.3722 | 5.96 | 1-Stearoyl-2-hydroxy-sn-glycero-3-phosphocholine | LysoPC(18:0) | [M+H]+ | 1 | 524.377, 506.36, 447.28, 341.30, 258.11, 184.07, 125.00, 104.11, 86.09 |
| 516.3104 | 4.97 | 1-O-9Z-Octadecenylethyleneglycol-2-O-phosphorylcholine | Oleyloxyethylphosphorylcholine | [M+K]+ | 1 | 516.32, 457.24, 146.98, 104.11, 86.10, 71.08, 69.07 |

| | | | | | | |
|---|---|---|---|---|---|---|
| 162.1129 | 0.33 | (3R)-3-Hydroxy-4-(trimethylammonio)butanoate | L-Carnitine | [M+H]+ | 1 | 162.11, 103.04, 102.09, 85.02, 60.08 |
| 508.3397 | 5.21 | 1-(1Z-Octadecenyl)-sn-glycero-3-phosphocholine | C18(Plasm) LPC | [M+H]+ | 1 | 508.37, 240.09, 184.07, 181.03, 166.06, 125.00, 104.11, 86.10 |
| 468.3092 | 4.81 | 1-Myristoyl-sn-glycero-3-phosphocholine | LysoPC(14:0) | [M+H]+ | 1 | 468.31, 450.29, 285.24, 184.07, 125.00, 104.11, 86.09 |
| 530.368 | 6.36 | 1-(1Z-Octadecenyl)-sn-glycero-3-phosphocholine | C18(Plasm) LPC | [M+Na]+ | 2 | 530.35, 471.28, 147.0, 104.11, 86.10 |
| 790.6208 | 10.03 | 2-Oleoyl-1-stearoyl-sn-glycero-3-phosphoserine | PS(18:0/18:1) | [M+H]+ | 2 | 790.6, 605.54, 341.31, 184.077, 95.08 |
| 204.1141 | 0.33 | 1-Propanaminium, 2-(acetyloxy)-3-carboxy-N,N,N-trimethyl-, inner salt | Acetyl-DL-carnitine | [M+H]+ | 1 | 204.12, 146.05, 145.05, 144.10, 86.03, 85.03, 60.08 |
| 316.2485 | 3.86 | 1-Propanaminium, 3-carboxy-N,N,N-trimethyl-2-[(1-oxodecyl)oxy]-, inner salt, (2R)- | Decanoyl-L-carnitine | [M+H]+ | 2 | 316.25, 257.18, 155.14, 85.03 |
| 728.5584 | 10.1 | 1-Stearoyl-2-oleoyl-sn-glycero-3-phosphoethanolamine | PE(18:0/18:1) | [M+H-H2O]+ | 1 | 728.56, 587.55, 390.28, 339.29, 95.09 |
| 257.2267 | 3.76 | 5α-Androst-16-en-3β-ol | Androst-16-en-3-ol, (3β,5α)- | [M+H-H2O]+ | 1 | 257.22, 215.18, 201.16, 189.16, 187.14, 175.15, 173.13, 163.14, 161.13, 149.13, 147.12, 121.10, 119.08, 117.07, 109.10, 107.08, 105.07, 97.10, 83.05, 69.07, 55.05 |
| 214.1134 | 3.03 | Octanoic acid, 8-[(carboxymethyl)amino]-8-oxo- | Suberylglycine | [M+H-H2O]+ | 1 | 214.11, 157.08, 139.07, 129.09, 111.08, 95.08, 83.08, 69.07 |
| 130.0499 | 1.9 | L-Glutamic acid | Glutamic Acid | [M+H-H2O]+ | 1 | 130.05, 84.04, 83.06 |
| 421.2528 | 5.75 | Phosphoric acid, tris(2-butoxyethyl) ester | Tris(2-butoxyethyl) phosphate | [M+Na]+ | 2 | 421.23, 321.14, 146.98 |

| | | | | | | |
|---|---|---|---|---|---|---|
| 180.0972 | 0.33 | Glycine, N-(phenylmethyl)-, methyl ester | N-Benzylglycine methyl ester | [M+H]+ | 2 | 180.10, 136.07, 91.05 |
| 678.4682 | 6.87 | 1,2-Ditetradecanoyl-sn-glycero-3-phosphocholine | PC(14:0) | [M+H]+ | 2 | 678.50, 185.08, 184.07, 124.10, 86.10 |
| 554.3463 | 1.56 | 1-Heptadecanoyl-sn-glycero-3-phosphocholine | LysoPC(17:0) | [M+HCO2]- | 1 | 554.34, 494.31, 269.24, 242.08, 224.07, 168.04, 153.00, 78.96 |
| 512.299 | 0.99 | 1-Myristoyl-sn-glycero-3-phosphocholine | LysoPC(14:0) | [M+HCO2]- | 2 | 512.29, 224.07,7 168.04, 153.00 |
| 526.31 | 1.14 | 1-Pentadecanoyl-sn-glycero-3-phosphocholine | LysoPC(15:0) | [M+HCO2]- | 1 | 526.31, 466.27, 242.08, 241.21, 224.07, 1688.04, 153.00, 78.96 |
| 178.0488 | 0.62 | Hippuric acid | Hippuric acid | [M-H]- | 2 | 178.05, 134.06 |
| 452.2775 | 0.99 | 1-Palmitoyl-2-hydroxy-sn-glycero-3-phosphoethanolamine | LysoPE(16:0) | [M-H]- | 1 | 452.27, 255.23, 196.04, 153.00, 140.01, |
| 862.5542 | 4.81 | 1-(1Z-Octadecenyl)-2-(4Z,7Z,10Z,13Z,16Z,19Z-docosahexaenoyl)-sn-glycero-3-phosphocholine | C18(Plasm)-22:6 PC | [M+HCO2]- | 1 | 862.59, 502.58, 492.34, 327.23, 283.24 |
| 830.5807 | 5.1 | 1-Stearoyl-2-linoleoyl-sn-glycero-3-phosphocholine | PC(18:0/18:2) | [M+HCO2]- | 1 | 830.59, 770.57, 508.34, 504.30, 490.33, 486.29, 419.25, 283.26, 279.23, 224.07, 168.04 |
| 768.5551 | 5.11 | 1-Stearoyl-2-myristoyl-sn-glycero-3-phosphocholine | PC(18:0/18:2) | [M+Cl]- | 1 | 768.55, 718.53, 508.34, 304.23, 303.23, 283.26, 256.23, 255.23 |
| 826.5604 | 4.96 | 1-Palmitoyl-2-arachidonoyl-sn-glycero-3-phosphocholine | PC(16:0/20:4) | [M+HCO2]- | 1 | 766.53, 528.31, 480.31, 462.30, 391.22, 303.23, 259.24, 255.23, 224.07, 168.04 |
| 766.5393 | 4.96 | 2-Arachidonoyl-1-stearoyl-sn-glycero-3-phosphoethanolamine | PE(18:0/20:4) | [M-H]- | 1 | 766.54, 480.31, 305.24, 304.24, 303.23, 284.27, 283.26, 196.04, 140.01 |

| | | | | | | |
|---|---|---|---|---|---|---|
| 835.5345 | 5.54 | 1-Hexadecanoyl-2-(9Z-octadecenoyl)-sn-glycero-3-phospho-(1'-myo-inositol) | PI(16:0/18:1) | [M-H]- | 2 | 835.53, 282.25, 281.25, 255.23 |
| 850.5605 | 4.85 | 1-Palmitoyl-2-docosahexaenoyl-sn-glycero-3-phosphocholine | PC(16;0/22:6) | [M+HCO2]- | 1 | 850.56, 790.54, 552.31, 534.30, 480.31, 462.30, 327.23, 283.24, 255.23, 229.19, 168.04 |
| 770.571 | 5.48 | 1,2-Dioleoyl-sn-glycero-3-phosphoethanolamine-N,N-dimethyl | DMPE(18:1/18:1) | [M-H]- | 2 | 770.57, 281.25, 279.23 |
| 832.5981 | 5.48 | 1-Oleoyl-2-stearoyl-sn-glycero-3-phosphocholine | PC(18:1/18:0) | [M+HCO2]- | 1 | 832.60, 772.58, 508.34, 506.32, 283.26, 281.25, 224.07, 168.04 |
| 794.5685 | 5.41 | 1-Oleoyl-2-palmitoyl-sn-glycero-3-phosphocholine | PC(18:1/16:0) | [M+Cl]- | 3 | 794.55, 744.55, 281.24, 255.23 |
| 832.6078 | 5.77 | 1-Stearoyl-2-oleoyl-sn-glycero-3-phosphocholine | PC(18:1/18:0) | [M+HCO2]- | 1 | 832.60, 772.58, 508.34, 506.34, 283.26, 281.28, 224.07, 168.04 |
| 830.5917 | 5.48 | 1,2-dioleoyl-sn-glycero-3-phosphatidylcholine | PC(18:1/18:1) | [M+HCO2]- | 1 | 830.595, 770.57, 699.7, 506.3, 283.26, 281.24, 224.07, 168.04 |
| 747.5162 | 5.21 | Palmitoyl sphingomyelin | SM(d18:1/16:0) | [M+HCO2]- | 1 | 747.56, 687.54, 616.47, 449.31, 255.23, 168.04, 122.98, 78.96 |
| 762.5082 | 4.98 | 2-Docosahexaenoyl-1-palmitoyl-sn-glycero-3-phosphoethanolamine | PE(16:0/22:6) | [M-H]- | 1 | 762.50, 452.27, 434.26, 327.23, 283.24, 255.23, 229.20, 196.04, 140.01 |
| 765.5276 | 5.16 | N-(Octadecanoyl)sphing-4-enine-1-phosphocholine | SM(d18:1/18:0) | [M+Cl]- | 2 | 765.56, 715.47, 283.26, 168.04 |
| 820.577 | 5.49 | 1-Stearoyl-2-linoleoyl-sn-glycero-3-phosphocholine | PC(18:0/18:2) | [M+Cl]- | 1 | 820.56, 770.57, 508.34, 329.24, 283.26, 279.23 |
| 790.5397 | 5.44 | 2-Docosahexaenoyl-1-stearoyl-sn-glycero-3-phosphoethanolamine | PE(18:0/22:6) | [M-H]- | 1 | 790.54, 480.31, 329.24, 328.24, 327.23, 284.27, 283.26, 229.19 |

| | | | | | | |
|---|---|---|---|---|---|---|
| 806.5944 | 5.72 | 1-Palmitoyl-2-stearoyl-sn-glycero-3-phosphocholine | PC(16:0/18:0) | [M+HCO2]- | 1 | 806.59, 746.56, 480.31, 283.26, 255.23, 224.07 |
| 790.541 | 4.84 | 2-Docosahexaenoyl-1-stearoyl-sn-glycero-3-phosphoethanolamine | PE(18:0/22:6) | [M-H]- | 1 | 740.54, 480.31, 327.723, 283.26, 229.19 |
| 878.5916 | 5.31 | 1-Stearoyl-2-docosahexaenoyl-sn-glycero-3-phosphocholine | PC(18:0/22:6) | [M+HCO2]- | 1 | 878.59, 552.31, 508.34, 490.24, 327.23, 283.26, 229.20, 191.18, 168.04 |
| 766.5395 | 5.55 | 2-Arachidonoyl-1-stearoyl-sn-glycero-3-phosphoethanolamine | PE(18:0/24:0) | [M-H]- | 1 | 766.53, 481.31, 480.31, 303.23, 283.26, 259.24, 196.04 |
| 776.5592 | 5.71 | 1-Oleoyl-2-myristoyl-sn-glycero-3-phosphocholine | PC(18:1/14:0) | [M+HCO2]- | 2 | 776.54, 716.52, 281.24, 168.04 |
| 768.5463 | 5.54 | 1-Stearoyl-2-myristoyl-sn-glycero-3-phosphocholine | PC(18:0/14:0) | [M+Cl]- | 1 | 768.53, 718.53, 508.34, 304.23, 303.23, 283.26, 255.23 |
| 838.5972 | 5.6 | 1-(1Z-Octadecenyl)-2-(5Z,8Z,11Z,14Z-eicosatetraenoyl)-sn-glycero-3-phosphocholine | C18(Plasm)-20:4 PC | [M+HCO2]- | 1 | 838.59, 778.57, 492.24, 474.33, 303.23 |
| 804.5669 | 5.01 | 1-Oleoyl-2-palmitoyl-sn-glycero-3-phosphocholine | PC(18:1/16:0) | [M+HCO2]- | 1 | 804.57, 744.55, 508.32, 480.31, 462.30, 281.25, 255.23, 168.04 |
| 802.5603 | 5.01 | 1-Palmitoyl-2-linoleoyl-sn-glycero-3-phosphocholine | PC(16:0/18:2) | [M+HCO2]- | 1 | 802.56, 742.53, 504.31, 480.30, 462.29, 391.22, 279.23, 255.23, 224.07, 168.04 |
| 744.549 | 5.01 | α-Galactosyl-C16-ceramide | α-Galactosyl-N-palmitoylsphingosine | [M+HCO2]- | 2 | 744.55, 698.56, 536.50, 255.23 |
| 774.545 | 5.65 | 1-(1Z-Octadecenyl)-2-(4Z,7Z,10Z,13Z,16Z,19Z-docosahexaenoyl)-sn-glycero-3-phosphoethanolamine | C18(Plasm)-22:6 PE | [M-H]- | 1 | 774.54, 717.54, 464.31, 446.31, 327.2,3 283.24, 229.19 |

| | | | | | | |
|---|---|---|---|---|---|---|
| 714.5081 | 5.16 | 2-Linoleoyl-1-palmitoyl-sn-glycero-3-phosphoethanolamine | PE(16:0/18:2) | [M-H]- | 1 | 714.50, 452.27, 279.23, 255.23 |
| 792.5473 | 5.01 | 1-Palmitoyl-2-linoleoyl-sn-glycero-3-phosphocholine | PC(16:0/18:2) | [M+Cl]- | 1 | 792.54, 742.54, 480.31, 279.23, 255.23 |
| 806.5831 | 5.35 | 1-Palmitoyl-2-stearoyl-sn-glycero-3-phosphocholine | PC(16:0/18:0) | [M+HCO2]- | 1 | 805.68, 746.57, 480.31, 238.26, 255.23, 168.04 |
| 834.5429 | 5.53 | 2-Docosahexaenoyl-1-stearoyl-sn-glycero-3-phosphoserine | PS(18:0/22:6) | [M-H]- | 1 | 834.52, 327.23, 283.26, 255.23, 241.01 |
| 742.5398 | 5.01 | 1-Stearoyl-2-linoleoyl-sn-glycero-3-phosphoethanolamine | PE(18:0/18:2) | [M-H]- | 1 | 742.53, 480.31, 283.26, 279.23, 255.23 |
| 862.5958 | 5.51 | 1-(1Z-Octadecenyl)-2-(4Z,7Z,10Z,13Z,16Z,19Z-docosahexaenoyl)-sn-glycero-3-phosphocholine | C18(Plasm)-22:6 PC | [M+HCO2]- | 1 | 862.59, 804.57, 492.34, 327.23, 283.24 |
| 804.576 | 5.36 | 1-Oleoyl-2-palmitoyl-sn-glycero-3-phosphocholine | PC(18:1/16:0) | [M+HCO2]- | 1 | 804.57, 744.55, 506.32, 480.31, 462.29, 281.24, 255.23, 168.04 |
| 738.5085 | 5.1 | 2-Arachidonoyl-1-palmitoyl-sn-glycero-3-phosphoethanolamine | PE(16:0/20:4) | [M-H]- | 1 | 738.51, 452.28, 304.23, 303.23, 255.23, 196.04, 168.04 |
| 162.1127 | 0.57 | (3R)-3-Hydroxy-4-(trimethylammonio)butanoate | L-Carnitine | [M+H]+ | 1 | 162.11, 103.04, 102.09, 85.08, 60.08 |
| 163.1155 | 0.56 | N-Phenyldiethylenediamine | Phenylpiperazine | [M+H]+ | 2 | 163.12, 120.08, 103.04, 95.05, 79.05 |
| 548.3722 | 1.53 | 1-O-Octadecyl-sn-glyceryl-3-phosphorylcholine | Lyso-PAF C-18 | [M+K]+ | 2 | 548.34, 489.27, 385.26, 104.11, 86.10 |
| 184.0945 | 0.58 | Phosphocholine | PC | [M+H]+ | 1 | 184.07, 125.00, 97.97, 886.10 |
| 510.3564 | 1.56 | 1-Heptadecanoyl-sn-glycero-3-phosphocholine | LysoPC(17:0) | [M+H]+ | 1 | 510.35, 492.35, 327.28, 258.11, 184.07, 125.00, 104.11, 86.10 |

| | | | | | | |
|---|---|---|---|---|---|---|
| 552.4037 | 2.62 | 1-Arachidoyl-2-hydroxy-sn-glycero-3-phosphocholine | LysoPC(20:0) | [M+H]+ | 1 | 552.40, 534.39, 184.07, 104.11 |
| 204.1231 | 0.58 | 1-Propanaminium, 2-(acetyloxy)-3-carboxy-N,N,N-trimethyl-, inner salt, (2R)- | Acetyl-DL-carnitine | [M+H]+ | 1 | 204.12, 145.05, 85.02, 60.08 |
| 482.3253 | 1.13 | 1-Stearoyl-2-hydroxy-sn-glycero-3-phosphoethanolamine | LysoPE(18:0) | [M+H]+ | 1 | 482.32, 464.31, 421.24, 341.30, 310.31, 184.07, 155.01, 95.08, 885.10, 71.08, 62.06 |
| 782.5682 | 4.96 | 2-Oleoyl-1-palmitoyl-sn-glycero-3-phosphocholine | PC(16:0/18:1) | [M+Na]+ | 1 | 782.57, 723.49, 599.50, 184.07, 146.98, 125.00, 104.11, 86.10 |
| 530.3694 | 1.4 | 1-(1Z-Octadecenyl)-sn-glycero-3-phosphocholine | C18(Plasm) LPC | [M+Na]+ | 1 | 530.36, 471.29, 427.27, 104.11, 86.10 |
| 468.3099 | 0.99 | 1-Myristoyl-sn-glycero-3-phosphocholine | LysoPC(14:0) | [M+H]+ | 1 | 468.31, 450.29, 184.077, 125.00, 104.11, 86.10 |
| 806.5689 | 4.84 | Arachidonoylthiophosphorylcholine | Arachidonoylthio-PC | [M+Na]+ | 3 | 806.56, 747.48, 623.49, 203.17, 184.07, 146.98, 86.10 |
| 504.3088 | 1.14 | 1-Eicosatrienoyl-sn-glycero-3-phosphoethanolamine | Lysophosphatidylethanolamine | [M+H]+ | 1 | 504.31, 461.26, 443.25, 341.30, 195.00, 164.01, 120.97 |
| 524.3717 | 1.89 | 1-Stearoyl-2-hydroxy-sn-glycero-3-phosphocholine | LysoPC(18:0) | [M+H]+ | 1 | 524.37, 506.36, 447.28, 341.31, 258.11, 184.07, 125.00, 104.11, 86.10, 60.08 |
| 544.338 | 1.41 | 1-Oleoyl-sn-glycero-3-phosphocholine | LysoPC(18:1) | [M+Na]+ | 1 | 544.33, 485.26, 339.28, 146.98, 104.11, 86.10 |
| 490.2926 | 0.99 | 1-Myristoyl-sn-glycero-3-phosphocholine | LysoPC(14:0) | [M+Na]+ | 1 | 490.29, 431.22, 307.22, 285.22, 146.98, 104.11, 86.01 |

| | | | | | | |
|---|---|---|---|---|---|---|
| 788.6068 | 5.47 | 1-Octadecanoyl-2-octadecenoyl-sn-glycero-3-phosphocholine | PC(18:0/18:1) | [M+H]+ | 2 | 788.61, 504.33, 184.07, 125.00, 86.10 |
| 287.1015 | 0.61 | L-Glutamine, N2-(2-phenylacetyl)- | Phenylacetyl-L-glutamine | [M+Na]+ | 1 | 287.10, 270.08, 177.07, 169.06, 159.02, 123.06 |
| 786.6003 | 5.47 | 1,2-dioleoyl-sn-glycero-3-phosphatidylcholine | 1,2-Dioleoyl-sn-glycero-3-PC | [M+H]+ | 1 | 786.60, 522.35, 184.07, 125.00, 104.11, 86.10 |
| 532.3389 | 1.56 | 1-Heptadecanoyl-sn-glycero-3-phosphocholine | LysoPC(17:0) | [M+Na]+ | 1 | 532.34, 473.26, 327.29, 146.98, 104.11, 86.10 |
| 794.5989 | 5.26 | 1-(1Z-Octadecenyl)-2-(5Z,8Z,11Z,14Z-eicosatetraenoyl)-sn-glycero-3-phosphocholine | C18(Plasm)-20:4 PC | [M+H]+ | 1 | 794.60, 735.52, 526.33, 508.37, 184.07, 104.11, 96.01, 71.08 |
| 637.3064 | 1.8 | Acetylpodocarpic acid anhydride | Acetylpodocarpic acid anhydride | [M+Na]+ | 1 | 637.31, 581.24, 525.18, 393.08, 147.00, 133.07 |
| 808.5841 | 5 | 1,2-dioleoyl-sn-glycero-3-phosphatidylcholine | 1,2-Dioleoyl-sn-glycero-3-PC | [M+Na]+ | 1 | 808.58, 749.50, 625.51, 603.54, 184.07, 146.98, 137.14, 86.10, 69.07 |
| 808.5769 | 4.83 | 1,2-dioleoyl-sn-glycero-3-phosphatidylcholine | 1,2-Dioleoyl-sn-glycero-3-PC | [M+Na]+ | 1 | 808.58, 749.50, 625.51, 603.54, 184.07, 146.98, 137.14, 86.10, 69.07 |
| 502.3262 | 1.49 | 1-(1Z-Hexadecenyl)-sn-glycero-3-phosphocholine | C16(Plasm) LPC | [M+Na]+ | 1 | 502.32, 443.35, 399.22, 147.98, 104.11, 86.10 |
| 546.3536 | 1.89 | 1-Stearoyl-2-hydroxy-sn-glycero-3-phosphocholine | LysoPC(18:0) | [M+Na]+ | 1 | 546.36, 487.28, 443.26, 363.28, 341.31, 240.08, 184.08, 176.99, 146.98, 133.08, 125.00, 104.11, 86.10, 771.09 |
| 792.5895 | 5.18 | 1-Hexadecyl-2-(8Z,11Z,14Z-eicosatrienoyl)-sn-glycero-3-phosphocholine | Dihomo-γ-linolenoyl PAF C-16 | [M+Na]+ | 1 | 792.59, 734.52, 733.52, 609.52, 587.54, 184.07, 146.98, 104.11, 86.10, 71.09 |
| 750.5453 | 5.35 | 1-Palmitoyl-2-thiopalmitoyl phosphatidylcholine | Palmitoyl Thio-PC | [M+H]+ | 1 | 750.54, 567.48, 184.07, 95.08, 86.01 |

| | | | | | | |
|---|---|---|---|---|---|---|
| 810.5994 | 5.42 | 1-Octadecanoyl-2-octadecenoyl-sn-glycero-3-phosphocholine | PC(18:0/18:1) | [M+Na]+ | 1 | 810.60, 751.52, 627.59, 605.55, 467.25, 184.07, 146.98, 86.10 |
| 790.5725 | 5.24 | 1-Hexadecyl-2-(5Z,8Z,11Z,14Z-eicosatetraenoyl)-sn-glycero-3-phosphocholine | Arachidonoyl PAF C-16 | [M+Na]+ | 1 | 790.57, 731.49, 607.51, 585.52, 526.33,184.07, 146.98, 86.10, 71.08 |
| 766.5724 | 5.16 | 1-O-Hexadecyl-2-O-(5Z,8Z,11Z,14Z,17Z-eicosapentaenoyl)-sn-glyceryl-3-phosphorylcholine | Eicosapentaenoyl PAF C-16 | [M+H]+ | 2 | 766.57, 1884.07, 125.00, 104.11, 86.10 |
| 760.5848 | 5.35 | 2-Oleoyl-1-palmitoyl-sn-glycero-3-phosphocholine | PC(16:0/18:1) | [M+H]+ | 1 | 760.58, 496.34, 184.07, 125.00, 86.10 |
| 782.567 | 5.35 | 2-Oleoyl-1-palmitoyl-sn-glycero-3-phosphocholine | PC(16:0/18:1) | [M+Na]+ | 1 | 782.57, 723.49, 599.50, 184.07, 147.98, 125.00, 86.10 |
| 768.5893 | 5.23 | 1-Hexadecyl-2-(9Z-octadecenoyl)-sn-glycero-3-phosphocholine | C16-18:1 PC | [M+H]+ | 1 | 768.58, 709.51, 585.51, 563.53, 184.07, 146.98, 86.10 |
| 735.5737 | 5.29 | 1-Palmitylthio-2-palmitoylamido-1,2-dideoxy-sn-glycero-3-phosphorylcholine | Thioetheramidephosphatidylcholine | [M+H]+ | 2 | 735.57, 184.077, 125.00, 86.10 |
| 756.5531 | 5.29 | 1,2-Dihexadecanoyl-sn-glycero-3-phosphocholine | PC(16:0/16:0) | [M+Na]+ | 1 | 756.55, 697.47, 573.48, 551.50, 500.31, 478.32, 441.23, 239.23, 146.98, 86.10 |
| 184.0734 | 5 | Phosphocholine | PC | [M+H]+ | 1 | 184.07, 86.10 |
| 834.5998 | 5.31 | 1-Docosahexaenoyl-2-stearoyl-sn-glycero-3-phosphocholine | 1-Docosahexaenoyl-2-stearoyl-sn-glycero-3-phosphocholine | [M+H]+ | 1 | 834.60, 631.53, 530.32, 524.36, 506.36, 491.25, 447.26, 341.30, 184.07, 166.06, 125.00, 104.11, 91.05, 86.10 |
| 758.5692 | 5.01 | 1-Hexadecanoyl-2-octadecadienoyl-sn-glycero-3-phosphocholine | PC(16:0/18:2) | [M+H]+ | 1 | 758.57, 520.34, 496.34, 184.07, 166.07, 125.00, 104.11, 98.99, 86.10 |

| | | | | | | |
|---|---|---|---|---|---|---|
| 760.5746 | 5.01 | 2-Oleoyl-1-palmitoyl-sn-glycero-3-phosphocholine | PC(16:0/18:1) | [M+H]+ | 1 | 760.58, 496.34, 184.07, 125.00, 86.10 |
| 768.5582 | 5.55 | 1-Stearoyl-2-oleoyl-sn-glycero-3-phosphoethanolamine | PE(18:0/18:1) | [M+Na]+ | 2 | 768.55, 725.49, 627.53, 341.31, 269.22, 203.18, 164.01 |

**Supplementary Table 3: Metabolite annotations with associated test statistics across different comparative groups for targeted LC-MS/MS data set.**

## 6 Gy

| | SD 1 vs. Pre | | | | SD 2 vs. Pre | | | | SD 6 vs. Pre | | | |
|---|---|---|---|---|---|---|---|---|---|---|---|---|
| Lipid Name | p-value | FDR | FC | LOG2FC | p-value | FDR | FC | LOG2FC | p-value | FDR | FC | LOG2FC |
| PI160/201 | 2.43E-05 | 0.013266 | 0.37975 ↓ | -1.397 | 0.033006 | 0.18848 | 0.64284 ↓ | -0.637 | 0.060925 | 0.14096 | 0.73387 ↓ | -0.446 |
| LPE160 | 5.93E-05 | 0.013266 | 1.7629 ↑ | 0.818 | 0.02511 | 0.15898 | 1.3944 ↑ | 0.4797 | 0.68717 | 0.7753 | 1.0867 ↑ | 0.12 |
| PC182/203 | 6.11E-05 | 0.013266 | 0.34223 ↓ | -1.547 | 0.023264 | 0.15898 | 0.54495 ↓ | -0.876 | 0.002355 | 0.055371 | 0.41783 ↓ | -1.259 |
| PG180/182_2 | 0.000119 | 0.015817 | 0.44596 ↓ | -1.165 | 0.001475 | 0.045715 | 0.61394 ↓ | -0.704 | 0.008226 | 0.075886 | 0.57301 ↓ | -0.803 |
| PI160/204 | 0.000135 | 0.015817 | 0.50756 ↓ | -0.978 | 0.046746 | 0.21583 | 0.76645 ↓ | -0.384 | 0.078835 | 0.16356 | 1.3704 ↑ | 0.4546 |
| PG180/182 | 0.000146 | 0.015817 | 0.45886 ↓ | -1.124 | 0.001446 | 0.045715 | 0.60947 ↓ | -0.714 | 0.009182 | 0.075886 | 0.56917 ↓ | -0.813 |
| LPC202 | 0.000212 | 0.015851 | 0.60406 ↓ | -0.727 | 0.036315 | 0.19538 | 0.74957 ↓ | -0.416 | 0.010088 | 0.076615 | 0.67573 ↓ | -0.565 |
| TAG547-FA183 | 0.000285 | 0.015851 | 0.12769 ↓ | -2.969 | 0.010214 | 0.10332 | 0.25992 ↓ | -1.944 | 0.007457 | 0.075886 | 0.18029 ↓ | -2.472 |
| PI160/182 | 0.000296 | 0.015851 | 0.37015 ↓ | -1.434 | 0.11596 | 0.3282 | 0.74892 ↓ | -0.417 | 0.035683 | 0.11738 | 0.63164 ↓ | -0.663 |
| TAG547-FA182 | 0.000316 | 0.015851 | 0.15225 ↓ | -2.716 | 0.006349 | 0.095455 | 0.26476 ↓ | -1.917 | 0.005075 | 0.07096 | 0.20043 ↓ | -2.319 |
| PS161/180 | 0.000336 | 0.015851 | 0.43713 ↓ | -1.194 | 0.016569 | 0.13483 | 0.52124 ↓ | -0.94 | 0.028773 | 0.10827 | 0.59323 ↓ | -0.753 |
| TAG526-FA183 | 0.000339 | 0.015851 | 0.1193 ↓ | -3.067 | 0.00479 | 0.087253 | 0.21331 ↓ | -2.229 | 0.004444 | 0.07096 | 0.18268 ↓ | -2.453 |
| TAG483-FA183 | 0.000351 | 0.015851 | 0.14376 ↓ | -2.798 | 0.002093 | 0.059234 | 0.17754 ↓ | -2.494 | 0.026014 | 0.1039 | 0.33521 ↓ | -1.577 |
| LPE226 | 0.000363 | 0.015851 | 1.8333 ↑ | 0.8745 | 0.000505 | 0.031013 | 1.7165 ↑ | 0.7795 | 0.15119 | 0.25238 | 1.2613 ↑ | 0.3349 |
| PG160/161 | 0.000365 | 0.015851 | 0.42354 ↓ | -1.239 | 0.004825 | 0.087253 | 0.51168 ↓ | -0.967 | 0.83086 | 0.88381 | 0.97039 ↓ | -0.043 |
| TAG525-FA182 | 0.000481 | 0.016324 | 0.17512 ↓ | -2.514 | 0.019686 | 0.1449 | 0.33543 ↓ | -1.576 | 0.01217 | 0.076615 | 0.28305 ↓ | -1.821 |
| PE181/203 | 0.000486 | 0.016324 | 0.37208 ↓ | -1.426 | 0.004224 | 0.084684 | 0.46282 ↓ | -1.112 | 0.005625 | 0.07096 | 0.40347 ↓ | -1.31 |
| PI160/180 | 0.000497 | 0.016324 | 0.36742 ↓ | -1.445 | 0.44195 | 0.62141 | 0.88991 ↓ | -0.168 | 0.060477 | 0.14096 | 0.68611 ↓ | -0.543 |
| PC160/226 | 0.000499 | 0.016324 | 1.4334 ↑ | 0.5195 | 0.005289 | 0.088492 | 1.3449 ↑ | 0.4275 | 0.032055 | 0.11231 | 1.2381 ↑ | 0.3081 |
| PI160/160 | 0.000529 | 0.016324 | 0.38718 ↓ | -1.369 | 0.41762 | 0.60657 | 0.86503 ↓ | -0.209 | 0.089248 | 0.17731 | 0.68319 ↓ | -0.55 |
| PI181/203_2 | 0.00055 | 0.016324 | 0.37075 ↓ | -1.432 | 0.014821 | 0.12819 | 0.55523 ↓ | -0.849 | 0.12409 | 0.21952 | 0.71351 ↓ | -0.487 |
| TAG542-FA200 | 0.000595 | 0.016324 | 0.27055 ↓ | -1.886 | 0.025492 | 0.15898 | 0.45594 ↓ | -1.133 | 0.000389 | 0.055371 | 0.18727 ↓ | -2.417 |
| TAG546-FA183 | 0.000629 | 0.016324 | 0.15633 ↓ | -2.677 | 0.029271 | 0.17014 | 0.34324 ↓ | -1.543 | 0.00711 | 0.075886 | 0.22127 ↓ | -2.176 |
| TAG504-FA182 | 0.000645 | 0.016324 | 0.1683 ↓ | -2.571 | 0.013851 | 0.12352 | 0.29944 ↓ | -1.74 | 0.076106 | 0.16034 | 0.43988 ↓ | -1.185 |
| PC181/202 | 0.000651 | 0.016324 | 0.43703 ↓ | -1.194 | 0.002478 | 0.064528 | 0.45812 ↓ | -1.126 | 0.067362 | 0.15036 | 0.65616 ↓ | -0.608 |
| PC182/183 | 0.000667 | 0.016324 | 0.27797 ↓ | -1.847 | 0.02452 | 0.15898 | 0.44021 ↓ | -1.184 | 0.001088 | 0.055371 | 0.2618 ↓ | -1.934 |
| PI160/182_2 | 0.000703 | 0.016324 | 0.37812 ↓ | -1.403 | 0.12716 | 0.34014 | 0.74029 ↓ | -0.434 | 0.061035 | 0.14096 | 0.65303 ↓ | -0.615 |
| PI181/203 | 0.000705 | 0.016324 | 0.32505 ↓ | -1.621 | 0.050254 | 0.21956 | 0.59454 ↓ | -0.75 | 0.2068 | 0.31902 | 0.74623 ↓ | -0.422 |

| | | | | | | | | | | | | | | | |
|---|---|---|---|---|---|---|---|---|---|---|---|---|---|---|---|
| PG160/161_2 | 0.000727 | 0.016324 | 0.39369 | ↓ | -1.345 | 0.070981 | 0.25251 | 0.69132 | ↓ | -0.533 | 0.49135 | 0.6116 | 0.90551 | ↓ | -0.143 |
| LPC200 | 0.000814 | 0.01729 | 0.45973 | ↓ | -1.121 | 0.93228 | 0.97158 | 1.0199 | ↑ | 0.0285 | 0.93473 | 0.95528 | 1.0309 | ↑ | 0.0439 |
| TAG504-FA140 | 0.000832 | 0.01729 | 0.18646 | ↓ | -2.423 | 0.019091 | 0.14285 | 0.33315 | ↓ | -1.586 | 0.077217 | 0.1621 | 0.45789 | ↓ | -1.127 |
| TAG525-FA183 | 0.00085 | 0.01729 | 0.17291 | ↓ | -2.532 | 0.04032 | 0.20071 | 0.36685 | ↓ | -1.447 | 0.010492 | 0.076615 | 0.24736 | ↓ | -2.015 |
| TAG526-FA182 | 0.000927 | 0.017851 | 0.1715 | ↓ | -2.544 | 0.024116 | 0.15898 | 0.3357 | ↓ | -1.575 | 0.006268 | 0.07372 | 0.22702 | ↓ | -2.139 |
| TAG525-FA225 | 0.000932 | 0.017851 | 0.29155 | ↓ | -1.778 | 0.024538 | 0.15898 | 0.44362 | ↓ | -1.173 | 0.028103 | 0.10827 | 0.46954 | ↓ | -1.091 |
| TAG504-FA183 | 0.001003 | 0.018132 | 0.13081 | ↓ | -2.935 | 0.008932 | 0.10202 | 0.21513 | ↓ | -2.217 | 0.033282 | 0.11344 | 0.3092 | ↓ | -1.693 |
| TAG526-FA161 | 0.001038 | 0.018132 | 0.10453 | ↓ | -3.258 | 0.006807 | 0.095455 | 0.1602 | ↓ | -2.642 | 0.005396 | 0.07096 | 0.14801 | ↓ | -2.756 |
| CE204 | 0.001095 | 0.018132 | 2.4863 | ↑ | 1.314 | 0.24855 | 0.4721 | 1.3075 | ↑ | 0.3869 | 0.2541 | 0.3676 | 1.3912 | ↑ | 0.4764 |
| TAG546-FA181 | 0.001096 | 0.018132 | 0.16394 | ↓ | -2.609 | 0.042812 | 0.20555 | 0.36126 | ↓ | -1.469 | 0.01082 | 0.076615 | 0.24807 | ↓ | -2.011 |
| PI160/204_2 | 0.001114 | 0.018132 | 0.51756 | ↓ | -0.95 | 0.063214 | 0.2435 | 0.73445 | ↓ | -0.445 | 0.088896 | 0.17731 | 1.3701 | ↑ | 0.4543 |
| TAG503-FA140 | 0.001114 | 0.018132 | 0.1596 | ↓ | -2.648 | 0.047179 | 0.21629 | 0.36905 | ↓ | -1.438 | 0.31437 | 0.43164 | 0.60836 | ↓ | -0.717 |
| PI161/181 | 0.001187 | 0.018558 | 0.31151 | ↓ | -1.683 | 0.18668 | 0.40645 | 0.67239 | ↓ | -0.573 | 0.61502 | 0.72011 | 0.87873 | ↓ | -0.187 |
| LPC220 | 0.001197 | 0.018558 | 0.41438 | ↓ | -1.271 | 0.89504 | 0.94765 | 1.0169 | ↑ | 0.0242 | 0.68941 | 0.77648 | 1.0597 | ↑ | 0.0837 |
| TAG548-FA182 | 0.001279 | 0.018972 | 0.15586 | ↓ | -2.682 | 0.005202 | 0.088492 | 0.22264 | ↓ | -2.167 | 0.002239 | 0.055371 | 0.17261 | ↓ | -2.534 |
| PC182/202 | 0.001293 | 0.018972 | 0.47668 | ↓ | -1.069 | 0.025691 | 0.15898 | 0.56804 | ↓ | -0.816 | 0.020661 | 0.094255 | 0.53676 | ↓ | -0.898 |
| PI182/182 | 0.001343 | 0.018972 | 0.25208 | ↓ | -1.988 | 0.041994 | 0.20555 | 0.53921 | ↓ | -0.891 | 0.002225 | 0.055371 | 0.31953 | ↓ | -1.646 |
| TAG546-FA182 | 0.00136 | 0.018972 | 0.21773 | ↓ | -2.199 | 0.094104 | 0.29172 | 0.4902 | ↓ | -1.029 | 0.01275 | 0.076615 | 0.27721 | ↓ | -1.851 |
| PI181/182_2 | 0.00137 | 0.018972 | 0.30773 | ↓ | -1.7 | 0.98689 | 0.99607 | 1.1166 | ↑ | 0.1591 | 0.10562 | 0.1963 | 0.68203 | ↓ | -0.552 |
| PI160/180_2 | 0.001414 | 0.019164 | 0.42079 | ↓ | -1.249 | 0.55757 | 0.71173 | 0.91639 | ↓ | -0.126 | 0.074185 | 0.15927 | 0.68446 | ↓ | -0.547 |
| TAG525-FA160 | 0.001443 | 0.019164 | 0.25957 | ↓ | -1.946 | 0.025862 | 0.15898 | 0.38211 | ↓ | -1.388 | 0.011506 | 0.076615 | 0.3076 | ↓ | -1.701 |
| TAG525-FA161 | 0.001504 | 0.019229 | 0.14826 | ↓ | -2.754 | 0.047834 | 0.21776 | 0.33757 | ↓ | -1.567 | 0.020447 | 0.094255 | 0.25998 | ↓ | -1.944 |
| TAG505-FA161 | 0.001506 | 0.019229 | 0.11548 | ↓ | -3.114 | 0.004423 | 0.084684 | 0.15466 | ↓ | -2.693 | 0.009551 | 0.075886 | 0.18539 | ↓ | -2.431 |
| PG181/182 | 0.001619 | 0.019942 | 0.31717 | ↓ | -1.657 | 0.079939 | 0.27246 | 0.55081 | ↓ | -0.86 | 0.053893 | 0.13832 | 0.5095 | ↓ | -0.973 |
| TAG563-FA200 | 0.001624 | 0.019942 | 0.26398 | ↓ | -1.922 | 0.74129 | 0.85564 | 1.1544 | ↑ | 0.2071 | 0.001818 | 0.055371 | 0.23563 | ↓ | -2.085 |
| PC160/204 | 0.001688 | 0.020345 | 1.46 | ↑ | 0.5459 | 0.011013 | 0.10592 | 1.3452 | ↑ | 0.4279 | 0.039118 | 0.11987 | 1.2069 | ↑ | 0.2713 |
| PC182/182 | 0.001785 | 0.021125 | 0.37972 | ↓ | -1.397 | 0.35856 | 0.57468 | 0.82482 | ↓ | -0.278 | 0.024769 | 0.10291 | 0.5345 | ↓ | -0.904 |
| PI160/181 | 0.001838 | 0.021254 | 0.42362 | ↓ | -1.239 | 0.97312 | 0.99083 | 1.0477 | ↑ | 0.0672 | 0.94855 | 0.96335 | 1.0568 | ↑ | 0.0797 |
| TAG482-FA182 | 0.001861 | 0.021254 | 0.2698 | ↓ | -1.89 | 0.015931 | 0.13184 | 0.37109 | ↓ | -1.43 | 0.6839 | 0.77325 | 0.88546 | ↓ | -0.176 |
| PG181/181 | 0.002003 | 0.022417 | 0.46666 | ↓ | -1.1 | 0.85001 | 0.92381 | 1.0587 | ↑ | 0.0823 | 0.68008 | 0.77131 | 0.89978 | ↓ | -0.152 |
| TAG566-FA183 | 0.002032 | 0.022417 | 0.20807 | ↓ | -2.265 | 0.053181 | 0.22051 | 0.40319 | ↓ | -1.311 | 0.010494 | 0.076615 | 0.23383 | ↓ | -2.097 |
| PC182/161 | 0.002097 | 0.022642 | 0.54744 | ↓ | -0.869 | 0.1258 | 0.3396 | 0.72255 | ↓ | -0.469 | 0.18446 | 0.29302 | 0.78695 | ↓ | -0.346 |
| PI181/182 | 0.002122 | 0.022642 | 0.3291 | ↓ | -1.603 | 0.96063 | 0.98689 | 1.0979 | ↑ | 0.1348 | 0.12314 | 0.21903 | 0.69407 | ↓ | -0.527 |

| | | | | | | | | | | | | | | | |
|---|---|---|---|---|---|---|---|---|---|---|---|---|---|---|---|
| TAG524-FA221 | 0.002199 | 0.023093 | 0.26105 | ↓ | -1.938 | 0.076111 | 0.26637 | 0.49903 | ↓ | -1.003 | 0.017338 | 0.082994 | 0.34316 | ↓ | -1.543 |
| PI160/181_2 | 0.002255 | 0.023124 | 0.42054 | ↓ | -1.25 | 0.95816 | 0.98689 | 1.0371 | ↑ | 0.0525 | 0.93109 | 0.95351 | 1.0516 | ↑ | 0.0726 |
| TAG541-FA200 | 0.002307 | 0.023124 | 0.25764 | ↓ | -1.957 | 0.37639 | 0.58341 | 0.72847 | ↓ | -0.457 | 0.001874 | 0.055371 | 0.23013 | ↓ | -2.12 |
| TAG548-FA183 | 0.002309 | 0.023124 | 0.10665 | ↓ | -3.229 | 0.010317 | 0.10332 | 0.16681 | ↓ | -2.584 | 0.007803 | 0.075886 | 0.12704 | ↓ | -2.977 |
| CE226 | 0.002405 | 0.023703 | 2.5972 | ↑ | 1.3769 | 0.45046 | 0.63064 | 1.2677 | ↑ | 0.3422 | 0.046698 | 0.13104 | 1.602 | ↑ | 0.6798 |
| TAG503-FA182 | 0.002456 | 0.023703 | 0.21265 | ↓ | -2.233 | 0.042656 | 0.20555 | 0.38061 | ↓ | -1.394 | 0.094493 | 0.182 | 0.46163 | ↓ | -1.115 |
| LPC183 | 0.002476 | 0.023703 | 0.48692 | ↓ | -1.038 | 0.05261 | 0.22051 | 0.62898 | ↓ | -0.669 | 0.000971 | 0.055371 | 0.37726 | ↓ | -1.406 |
| TAG525-FA181 | 0.002563 | 0.02388 | 0.17776 | ↓ | -2.492 | 0.087912 | 0.28828 | 0.39853 | ↓ | -1.327 | 0.021191 | 0.094255 | 0.28057 | ↓ | -1.834 |
| PC140/226 | 0.002568 | 0.02388 | 0.7259 | ↓ | -0.462 | 0.1045 | 0.30369 | 0.81217 | ↓ | -0.3 | 0.050077 | 0.13527 | 2.2648 | ↑ | 1.1794 |
| LPC150 | 0.002606 | 0.023898 | 0.4855 | ↓ | -1.042 | 0.088122 | 0.28828 | 0.68633 | ↓ | -0.543 | 0.011177 | 0.076615 | 0.5778 | ↓ | -0.791 |
| PS161/181 | 0.002667 | 0.023925 | 0.47929 | ↓ | -1.061 | 0.36763 | 0.57474 | 0.82431 | ↓ | -0.279 | 0.24906 | 0.36253 | 1.6625 | ↑ | 0.7333 |
| PC180/201 | 0.002684 | 0.023925 | 0.72078 | ↓ | -0.472 | 0.000139 | 0.018452 | 0.59891 | ↓ | -0.74 | 0.019008 | 0.089946 | 0.73958 | ↓ | -0.435 |
| PI181/181 | 0.00272 | 0.023925 | 0.34498 | ↓ | -1.535 | 0.078865 | 0.27022 | 2.0243 | ↑ | 1.0174 | 0.23408 | 0.34791 | 1.5171 | ↑ | 0.6013 |
| PI160/203_2 | 0.002837 | 0.024504 | 0.4447 | ↓ | -1.169 | 0.008279 | 0.10202 | 0.51062 | ↓ | -0.97 | 0.16812 | 0.27568 | 0.71284 | ↓ | -0.488 |
| TAG562-FA200 | 0.002861 | 0.024504 | 0.32994 | ↓ | -1.6 | 0.11877 | 0.33301 | 2.0952 | ↑ | 1.0671 | 0.004656 | 0.07096 | 0.33723 | ↓ | -1.568 |
| PC180/202 | 0.00294 | 0.024558 | 0.63896 | ↓ | -0.646 | 9.3E-05 | 0.018452 | 0.47462 | ↓ | -1.075 | 0.006102 | 0.073558 | 0.62501 | ↓ | -0.678 |
| TAG547-FA181 | 0.002942 | 0.024558 | 0.18644 | ↓ | -2.423 | 0.050069 | 0.21956 | 0.34937 | ↓ | -1.517 | 0.005436 | 0.07096 | 0.18821 | ↓ | -2.41 |
| TAG503-FA181 | 0.003105 | 0.025062 | 0.18668 | ↓ | -2.421 | 0.077467 | 0.26744 | 0.40277 | ↓ | -1.312 | 0.32546 | 0.44512 | 0.60094 | ↓ | -0.735 |
| TAG484-FA120 | 0.003158 | 0.025062 | 0.20223 | ↓ | -2.306 | 0.03914 | 0.20063 | 0.36504 | ↓ | -1.454 | 0.74035 | 0.81902 | 1.2547 | ↑ | 0.3273 |
| TAG504-FA161 | 0.003188 | 0.025062 | 0.15502 | ↓ | -2.69 | 0.011064 | 0.10592 | 0.21912 | ↓ | -2.19 | 0.016938 | 0.08168 | 0.26645 | ↓ | -1.908 |
| PG160/203_2 | 0.003197 | 0.025062 | 0.47624 | ↓ | -1.07 | 0.000741 | 0.0376 | 0.35151 | ↓ | -1.508 | 0.001939 | 0.055371 | 0.3175 | ↓ | -1.655 |
| TAG524-FA161 | 0.003264 | 0.025062 | 0.16673 | ↓ | -2.584 | 0.1293 | 0.34014 | 0.4318 | ↓ | -1.212 | 0.032434 | 0.11231 | 0.2922 | ↓ | -1.775 |
| TAG563-FA182 | 0.003285 | 0.025062 | 0.32545 | ↓ | -1.62 | 0.81418 | 0.89922 | 1.1241 | ↑ | 0.1688 | 0.007292 | 0.075886 | 0.3018 | ↓ | -1.728 |
| TAG565-FA201 | 0.003305 | 0.025062 | 0.24806 | ↓ | -2.011 | 0.20104 | 0.41834 | 0.59382 | ↓ | -0.752 | 0.019882 | 0.093114 | 0.31247 | ↓ | -1.678 |
| PI160/203 | 0.003361 | 0.025062 | 0.47868 | ↓ | -1.063 | 0.018415 | 0.1416 | 0.57987 | ↓ | -0.786 | 0.35531 | 0.48033 | 0.82095 | ↓ | -0.285 |
| PI180/180 | 0.003373 | 0.025062 | 0.45499 | ↓ | -1.136 | 0.77845 | 0.87711 | 0.99154 | ↓ | -0.012 | 0.82753 | 0.88345 | 1.1552 | ↑ | 0.2082 |
| TAG541-FA160 | 0.003388 | 0.025062 | 0.32031 | ↓ | -1.643 | 0.40995 | 0.6032 | 0.76601 | ↓ | -0.385 | 0.005137 | 0.07096 | 0.29331 | ↓ | -1.77 |
| PC182/224 | 0.003646 | 0.026466 | 0.5703 | ↓ | -0.81 | 0.038232 | 0.19911 | 0.69132 | ↓ | -0.533 | 0.000566 | 0.055371 | 0.47347 | ↓ | -1.079 |
| PC182/205 | 0.003659 | 0.026466 | 0.37542 | ↓ | -1.413 | 0.28133 | 0.49607 | 0.73796 | ↓ | -0.438 | 0.013388 | 0.077008 | 0.44769 | ↓ | -1.159 |
| TAG545-FA180 | 0.003806 | 0.02681 | 0.27018 | ↓ | -1.888 | 0.025092 | 0.15898 | 0.35726 | ↓ | -1.485 | 0.008708 | 0.075886 | 0.26473 | ↓ | -1.917 |
| PI161/181_2 | 0.003812 | 0.02681 | 0.41079 | ↓ | -1.284 | 0.3525 | 0.57369 | 0.82869 | ↓ | -0.271 | 0.70444 | 0.78931 | 0.97025 | ↓ | -0.044 |
| TAG524-FA160 | 0.003845 | 0.02681 | 0.36383 | ↓ | -1.459 | 0.27482 | 0.49163 | 0.73193 | ↓ | -0.45 | 0.040368 | 0.12112 | 0.46847 | ↓ | -1.094 |
| PC160/225 | 0.003889 | 0.02681 | 1.5287 | ↑ | 0.6124 | 0.014818 | 0.12819 | 1.5398 | ↑ | 0.6227 | 0.37293 | 0.49604 | 1.1825 | ↑ | 0.2419 |

| | | | | | | | | | | | | |
|---|---|---|---|---|---|---|---|---|---|---|---|---|
| TAG524-FA183 | 0.003912 | 0.02681 | 0.27577 | ↓ -1.858 | 0.036716 | 0.19592 | 0.41037 | ↓ -1.285 | 0.016012 | 0.079367 | 0.32528 | ↓ -1.62 |
| TAG561-FA181 | 0.003976 | 0.026964 | 0.32057 | ↓ -1.641 | 0.3291 | 0.54655 | 1.6675 | ↑ 0.7377 | 0.032612 | 0.11233 | 0.41219 | ↓ -1.279 |
| TAG482-FA140 | 0.004041 | 0.027117 | 0.27851 | ↓ -1.844 | 0.025341 | 0.15898 | 0.37776 | ↓ -1.404 | 0.37055 | 0.49533 | 0.73019 | ↓ -0.454 |
| PI200/204_2 | 0.004231 | 0.028103 | 0.47184 | ↓ -1.084 | 0.045484 | 0.21302 | 0.64027 | ↓ -0.643 | 0.90684 | 0.93558 | 1.1681 | ↑ 0.2241 |
| TAG502-FA140 | 0.00443 | 0.029128 | 0.18889 | ↓ -2.404 | 0.14138 | 0.36235 | 0.46204 | ↓ -1.114 | 0.82533 | 0.88345 | 0.86566 | ↓ -0.208 |
| TAG545-FA183 | 0.004517 | 0.029407 | 0.2159 | ↓ -2.212 | 0.14624 | 0.36338 | 0.50581 | ↓ -0.983 | 0.012877 | 0.076615 | 0.25938 | ↓ -1.947 |
| PG160/203 | 0.0046 | 0.029433 | 0.52682 | ↓ -0.925 | 0.001159 | 0.044105 | 0.40378 | ↓ -1.308 | 0.001477 | 0.055371 | 0.35605 | ↓ -1.49 |
| PI160/161 | 0.004612 | 0.029433 | 0.40881 | ↓ -1.291 | 0.13456 | 0.34762 | 0.64381 | ↓ -0.635 | 0.71701 | 0.79927 | 1.1412 | ↑ 0.1906 |
| PE182/202 | 0.004747 | 0.030005 | 0.35306 | ↓ -1.502 | 0.020032 | 0.1449 | 0.41527 | ↓ -1.268 | 0.015124 | 0.079031 | 0.37791 | ↓ -1.404 |
| PI182/201 | 0.004881 | 0.030489 | 0.43923 | ↓ -1.187 | 0.63254 | 0.77799 | 0.9378 | ↓ -0.093 | 0.29017 | 0.4045 | 0.79137 | ↓ -0.338 |
| LPA180 | 0.004923 | 0.030489 | 0.68459 | ↓ -0.547 | 0.73436 | 0.85217 | 1.0584 | ↑ 0.0818 | 0.75854 | 0.83133 | 0.96375 | ↓ -0.053 |
| LPC240 | 0.004996 | 0.030489 | 0.51925 | ↓ -0.946 | 0.13011 | 0.34014 | 0.70803 | ↓ -0.498 | 0.43828 | 0.55727 | 1.1712 | ↑ 0.228 |
| TAG524-FA200 | 0.005011 | 0.030489 | 0.35258 | ↓ -1.504 | 0.28576 | 0.49982 | 0.71364 | ↓ -0.487 | 0.063945 | 0.14555 | 0.50793 | ↓ -0.977 |
| TAG566-FA202 | 0.005107 | 0.030751 | 0.28244 | ↓ -1.824 | 0.10086 | 0.30035 | 0.50092 | ↓ -0.997 | 0.011192 | 0.076615 | 0.28412 | ↓ -1.815 |
| PS181/182 | 0.005149 | 0.030751 | 0.57895 | ↓ -0.788 | 0.024352 | 0.15898 | 0.68268 | ↓ -0.551 | 0.019067 | 0.089946 | 0.66268 | ↓ -0.594 |
| TAG524-FA181 | 0.005383 | 0.031858 | 0.22741 | ↓ -2.137 | 0.093709 | 0.29172 | 0.43809 | ↓ -1.191 | 0.029091 | 0.10858 | 0.32929 | ↓ -1.603 |
| PG161/180 | 0.005755 | 0.033471 | 0.60826 | ↓ -0.717 | 0.000866 | 0.0376 | 0.48889 | ↓ -1.032 | 0.04252 | 0.12303 | 0.6902 | ↓ -0.535 |
| PI180/203_2 | 0.005795 | 0.033471 | 0.6435 | ↓ -0.636 | 0.004338 | 0.084684 | 0.65451 | ↓ -0.612 | 0.011638 | 0.076615 | 0.65024 | ↓ -0.621 |
| PI180/183_2 | 0.00583 | 0.033471 | 0.52431 | ↓ -0.932 | 0.089808 | 0.29172 | 0.72033 | ↓ -0.473 | 0.036334 | 0.11738 | 0.5969 | ↓ -0.744 |
| LPC241 | 0.005962 | 0.033471 | 0.52609 | ↓ -0.927 | 0.41836 | 0.60657 | 0.87462 | ↓ -0.193 | 0.74453 | 0.82151 | 0.96619 | ↓ -0.05 |
| CE203 | 0.005979 | 0.033471 | 1.7079 | ↑ 0.7722 | 0.40131 | 0.5989 | 0.86528 | ↓ -0.209 | 0.20821 | 0.31968 | 0.75236 | ↓ -0.411 |
| TAG524-FA182 | 0.00606 | 0.033471 | 0.38737 | ↓ -1.368 | 0.31419 | 0.52894 | 0.74307 | ↓ -0.428 | 0.040322 | 0.12112 | 0.46234 | ↓ -1.113 |
| TAG545-FA182 | 0.006066 | 0.033471 | 0.26727 | ↓ -1.904 | 0.40236 | 0.5989 | 0.72551 | ↓ -0.463 | 0.030095 | 0.11007 | 0.36606 | ↓ -1.45 |
| TAG544-FA201 | 0.0061 | 0.033471 | 0.21606 | ↓ -2.211 | 0.15953 | 0.37492 | 0.45977 | ↓ -1.121 | 0.056114 | 0.13888 | 0.36373 | ↓ -1.459 |
| TAG545-FA161 | 0.006118 | 0.033471 | 0.19999 | ↓ -2.322 | 0.10247 | 0.30256 | 0.39439 | ↓ -1.342 | 0.011141 | 0.076615 | 0.21883 | ↓ -2.192 |
| TAG502-FA181 | 0.00655 | 0.035534 | 0.22714 | ↓ -2.138 | 0.14536 | 0.36338 | 0.47938 | ↓ -1.061 | 0.37269 | 0.49604 | 0.62022 | ↓ -0.689 |
| PG181/182_2 | 0.006667 | 0.035867 | 0.35863 | -1.4794 | 0.28065 | 0.49607 | 0.65159 | -0.61797 | 0.13752 | 0.23622 | 0.56123 | -0.83334 |
| PC200/182 | 0.006868 | 0.036647 | 0.6664 | -0.58554 | 0.77521 | 0.87711 | 0.9657 | -0.05035 | 0.21294 | 0.32541 | 0.80726 | -0.30889 |
| PC140/203 | 0.007007 | 0.037088 | 0.51118 | -0.96809 | 0.004269 | 0.084684 | 0.4481 | -1.1581 | 0.71968 | 0.80088 | 1.3606 | 0.44428 |
| TAG543-FA183 | 0.00719 | 0.037632 | 0.32775 | -1.6093 | 0.086618 | 0.28624 | 0.50457 | -0.98687 | 0.010793 | 0.076615 | 0.2582 | -1.9534 |
| PC181/203 | 0.007227 | 0.037632 | 0.4995 | -1.0014 | 0.008761 | 0.10202 | 0.51163 | -0.96683 | 0.013848 | 0.077008 | 0.52401 | -0.93232 |
| TAG523-FA201 | 0.007305 | 0.037632 | 0.31498 | -1.6667 | 0.049101 | 0.21894 | 0.44466 | -1.1692 | 0.059827 | 0.14096 | 0.44835 | -1.1573 |
| LPC201 | 0.007341 | 0.037632 | 0.63596 | -0.65299 | 0.29025 | 0.50521 | 1.1987 | 0.26145 | 0.72478 | 0.80517 | 1.0874 | 0.12087 |

| | | | | | | | | | | | | |
|---|---|---|---|---|---|---|---|---|---|---|---|---|
| PE181/205 | 0.007439 | 0.037833 | 0.5074 | -0.97881 | 0.083989 | 0.28184 | 0.6566 | -0.60691 | 0.012193 | 0.076615 | 0.4338 | -1.2049 |
| LPC161 | 0.007632 | 0.038236 | 0.47067 | -1.0872 | 0.24304 | 0.46994 | 0.72089 | -0.47215 | 0.42793 | 0.54947 | 0.78839 | -0.34302 |
| TAG481-FA181 | 0.007635 | 0.038236 | 0.26004 | -1.9432 | 0.052897 | 0.22051 | 0.38247 | -1.3866 | 0.7993 | 0.86533 | 1.1536 | 0.2062 |
| PC120/181 | 0.007806 | 0.038628 | 0.5483 | -0.86695 | 0.04039 | 0.20071 | 0.59674 | -0.74484 | 0.8002 | 0.86533 | 1.1994 | 0.2623 |
| PC150/203 | 0.007832 | 0.038628 | 0.58621 | -0.7705 | 0.003281 | 0.082152 | 0.4626 | -1.1122 | 0.008964 | 0.075886 | 0.51912 | -0.94586 |
| TAG567-FA183 | 0.008085 | 0.039107 | 0.33175 | -1.5918 | 0.084905 | 0.28345 | 0.51926 | -0.94547 | 0.012403 | 0.076615 | 0.30677 | -1.7048 |
| TAG544-FA183 | 0.008117 | 0.039107 | 0.28414 | -1.8153 | 0.12974 | 0.34014 | 0.51831 | -0.94811 | 0.015426 | 0.079367 | 0.29569 | -1.7578 |
| TAG523-FA183 | 0.008132 | 0.039107 | 0.31753 | -1.655 | 0.066679 | 0.24778 | 0.46988 | -1.0896 | 0.020975 | 0.094255 | 0.36139 | -1.4684 |
| PE182/201 | 0.008197 | 0.039107 | 0.47472 | -1.0749 | 0.16727 | 0.38229 | 0.70692 | -0.50038 | 0.056533 | 0.13888 | 0.53022 | -0.91533 |
| TAG544-FA180 | 0.008264 | 0.039107 | 0.31996 | -1.644 | 0.24623 | 0.4721 | 0.64216 | -0.639 | 0.02527 | 0.10291 | 0.36587 | -1.4506 |
| TAG482-FA161 | 0.00829 | 0.039107 | 0.18625 | -2.4247 | 0.027372 | 0.16348 | 0.24731 | -2.0156 | 0.57926 | 0.68939 | 0.65109 | -0.61907 |
| TAG482-FA160 | 0.008391 | 0.0393 | 0.30991 | -1.6901 | 0.037202 | 0.1969 | 0.4061 | -1.3001 | 0.62943 | 0.73194 | 0.8583 | -0.22044 |
| PI180/181_2 | 0.008559 | 0.039799 | 0.52372 | -0.93314 | 0.46202 | 0.63455 | 1.2651 | 0.33925 | 0.47164 | 0.59046 | 1.3181 | 0.39844 |
| TAG523-FA180 | 0.008724 | 0.040277 | 0.34947 | -1.5167 | 0.06561 | 0.24689 | 0.48863 | -1.0332 | 0.01504 | 0.079031 | 0.36913 | -1.4378 |
| PG160/182_2 | 0.008945 | 0.040735 | 0.34175 | -1.549 | 0.14463 | 0.36338 | 0.54528 | -0.87492 | 0.19873 | 0.30876 | 0.57499 | -0.79838 |
| TAG503-FA161 | 0.008967 | 0.040735 | 0.25897 | -1.9492 | 0.069108 | 0.25019 | 0.41225 | -1.2784 | 0.061061 | 0.14096 | 0.40928 | -1.2889 |
| PI180/181 | 0.00901 | 0.040735 | 0.49268 | -1.0213 | 0.46172 | 0.63455 | 1.2716 | 0.34662 | 0.41736 | 0.53909 | 1.3561 | 0.43942 |
| PC182/226 | 0.009186 | 0.041179 | 0.47806 | -1.0647 | 0.69336 | 0.82368 | 0.9325 | -0.10083 | 0.051721 | 0.13622 | 0.58723 | -0.768 |
| PC181/183 | 0.009235 | 0.041179 | 0.48217 | -1.0524 | 0.099232 | 0.29948 | 0.63451 | -0.65627 | 0.001699 | 0.055371 | 0.36671 | -1.4473 |
| PC182/204 | 0.009484 | 0.041763 | 0.60018 | -0.73653 | 0.54962 | 0.70573 | 0.91312 | -0.13113 | 0.064815 | 0.14702 | 0.6986 | -0.51746 |
| TAG565-FA182 | 0.009495 | 0.041763 | 0.28104 | -1.8312 | 0.29968 | 0.5134 | 0.62996 | -0.66667 | 0.025363 | 0.10291 | 0.34002 | -1.5563 |
| TAG544-FA161 | 0.009663 | 0.042219 | 0.21285 | -2.2321 | 0.38697 | 0.58996 | 0.62345 | -0.68165 | 0.054668 | 0.13832 | 0.33056 | -1.597 |
| PC181/226 | 0.00989 | 0.042923 | 0.69427 | -0.52643 | 0.15275 | 0.36901 | 0.80011 | -0.32173 | 0.17897 | 0.28627 | 0.80386 | -0.31499 |
| TAG520-FA200 | 0.010128 | 0.043666 | 0.37064 | -1.4319 | 0.015999 | 0.13184 | 0.3865 | -1.3715 | 0.002101 | 0.055371 | 0.23925 | -2.0634 |
| SMd181/182 | 0.010422 | 0.04461 | 1.3183 | 0.39866 | 0.25179 | 0.4721 | 1.0881 | 0.12182 | 0.4324 | 0.55412 | 1.0936 | 0.12908 |
| PC140/202 | 0.010515 | 0.04461 | 0.55443 | -0.85093 | 0.018065 | 0.1416 | 0.5351 | -0.90213 | 0.40052 | 0.52357 | 1.8096 | 0.85571 |
| TAG524-FA180 | 0.010553 | 0.04461 | 0.32762 | -1.6099 | 0.051115 | 0.22032 | 0.44015 | -1.1839 | 0.009559 | 0.075886 | 0.30116 | -1.7314 |
| PG160/182 | 0.010699 | 0.044915 | 0.34566 | -1.5326 | 0.14893 | 0.3662 | 0.54011 | -0.88866 | 0.19768 | 0.30822 | 0.56649 | -0.81988 |
| PI180/182_2 | 0.010911 | 0.044915 | 0.57793 | -0.79104 | 0.36145 | 0.57468 | 0.86873 | -0.20302 | 0.074288 | 0.15927 | 0.71222 | -0.48961 |
| TAG526-FA181 | 0.010912 | 0.044915 | 0.3059 | -1.7089 | 0.24301 | 0.46994 | 0.66051 | -0.59835 | 0.009166 | 0.075886 | 0.30469 | -1.7146 |
| TAG483-FA161 | 0.010967 | 0.044915 | 0.20666 | -2.2747 | 0.028907 | 0.16954 | 0.25838 | -1.9525 | 0.56484 | 0.6747 | 0.66968 | -0.57845 |
| TAG565-FA202 | 0.010974 | 0.044915 | 0.27986 | -1.8372 | 0.13307 | 0.34515 | 0.51151 | -0.96717 | 0.031212 | 0.11103 | 0.32759 | -1.61 |
| LPC170 | 0.011039 | 0.044915 | 0.61833 | -0.69355 | 0.27776 | 0.49406 | 0.82405 | -0.2792 | 0.042168 | 0.12255 | 0.66909 | -0.57973 |

| | | | | | | | | | | | | |
|---|---|---|---|---|---|---|---|---|---|---|---|---|
| TAG545-FA181 | 0.01124 | 0.045268 | 0.27806 | -1.8465 | 0.50546 | 0.66881 | 0.77249 | -0.37241 | 0.040372 | 0.12112 | 0.37855 | -1.4014 |
| TAG523-FA202 | 0.011265 | 0.045268 | 0.29589 | -1.7569 | 0.13062 | 0.34014 | 0.49898 | -1.003 | 0.093703 | 0.1812 | 0.44265 | -1.1758 |
| PI180/182 | 0.011435 | 0.04567 | 0.59691 | -0.7444 | 0.39632 | 0.5989 | 0.88223 | -0.18078 | 0.056748 | 0.13888 | 0.69535 | -0.52419 |
| TAG540-FA160 | 0.011735 | 0.046582 | 0.3462 | -1.5303 | 0.10753 | 0.31112 | 0.50704 | -0.97982 | 0.004603 | 0.07096 | 0.25886 | -1.9498 |
| TAG526-FA160 | 0.011932 | 0.047078 | 0.28306 | -1.8208 | 0.033943 | 0.18981 | 0.33519 | -1.577 | 0.014848 | 0.079031 | 0.28481 | -1.8119 |
| TAG526-FA205 | 0.012083 | 0.047356 | 0.34294 | -1.544 | 0.064349 | 0.24498 | 0.49033 | -1.0282 | 0.067441 | 0.15036 | 0.50907 | -0.97406 |
| PS180/181 | 0.012148 | 0.047356 | 0.5521 | -0.85699 | 0.10826 | 0.31159 | 0.70285 | -0.50872 | 0.13467 | 0.23317 | 0.69827 | -0.51814 |
| TAG523-FA182 | 0.012444 | 0.047975 | 0.41507 | -1.2686 | 0.40336 | 0.5989 | 0.76646 | -0.38371 | 0.052625 | 0.13716 | 0.50174 | -0.99499 |
| TAG545-FA202 | 0.012454 | 0.047975 | 0.24579 | -2.0245 | 0.14614 | 0.36338 | 0.46847 | -1.094 | 0.015569 | 0.079367 | 0.26766 | -1.9015 |
| TAG483-FA140 | 0.012635 | 0.04816 | 0.14353 | -2.8005 | 0.035853 | 0.19538 | 0.19278 | -2.375 | 0.56336 | 0.67417 | 0.56457 | -0.82477 |
| PE180/202 | 0.01265 | 0.04816 | 0.70727 | -0.49968 | 0.000129 | 0.018452 | 0.56927 | -0.81282 | 0.002751 | 0.055371 | 0.56824 | -0.81543 |
| PI180/202_2 | 0.013325 | 0.050156 | 0.5811 | -0.78314 | 0.000205 | 0.019088 | 0.46715 | -1.098 | 0.003352 | 0.062338 | 0.5015 | -0.99567 |
| PE181/204 | 0.013329 | 0.050156 | 0.56725 | -0.81795 | 0.11589 | 0.3282 | 0.71918 | -0.47557 | 0.042967 | 0.12369 | 0.58616 | -0.77064 |
| TAG523-FA181 | 0.013653 | 0.050874 | 0.42641 | -1.2297 | 0.35818 | 0.57468 | 0.7479 | -0.41907 | 0.058228 | 0.1404 | 0.51792 | -0.94919 |
| TAG503-FA160 | 0.013676 | 0.050874 | 0.31786 | -1.6535 | 0.068158 | 0.25019 | 0.44818 | -1.1579 | 0.050842 | 0.13574 | 0.43006 | -1.2174 |
| TAG483-FA182 | 0.013924 | 0.051503 | 0.27531 | -1.8609 | 0.064949 | 0.24583 | 0.38427 | -1.3798 | 0.4439 | 0.56221 | 1.4702 | 0.55597 |
| PC140/161 | 0.014078 | 0.051777 | 0.56496 | -0.82378 | 0.01026 | 0.10332 | 0.52849 | -0.92004 | 0.37336 | 0.49604 | 1.8947 | 0.92197 |
| TAG523-FA161 | 0.014313 | 0.052348 | 0.21225 | -2.2362 | 0.34801 | 0.57071 | 0.56198 | -0.83142 | 0.073508 | 0.15919 | 0.32558 | -1.6189 |
| TAG5610-FA182 | 0.014601 | 0.052592 | 0.36046 | -1.4721 | 0.98121 | 0.99497 | 1.2261 | 0.29409 | 0.009451 | 0.075886 | 0.28621 | -1.8049 |
| PC180/183 | 0.014667 | 0.052592 | 0.615 | -0.70134 | 0.000777 | 0.0376 | 0.39217 | -1.3504 | 0.000982 | 0.055371 | 0.41643 | -1.2638 |
| TAG564-FA201 | 0.014679 | 0.052592 | 0.28032 | -1.8349 | 0.92725 | 0.96892 | 1.0589 | 0.082595 | 0.067082 | 0.15036 | 0.43081 | -1.2149 |
| PE182/161 | 0.014703 | 0.052592 | 0.52823 | -0.92076 | 0.69478 | 0.82386 | 0.96966 | -0.04445 | 0.07518 | 0.16 | 0.68232 | -0.55147 |
| PE181/226 | 0.015221 | 0.054148 | 0.57428 | -0.80017 | 0.15098 | 0.36781 | 0.73048 | -0.45309 | 0.073217 | 0.15919 | 0.62502 | -0.67803 |
| TAG481-FA140 | 0.015494 | 0.054732 | 0.31132 | -1.6835 | 0.069435 | 0.25019 | 0.42596 | -1.2312 | 0.83869 | 0.88634 | 1.1616 | 0.2161 |
| PS161/182_2 | 0.015554 | 0.054732 | 0.63724 | -0.6501 | 0.29767 | 0.5134 | 0.84488 | -0.24318 | 0.014134 | 0.077008 | 0.61774 | -0.69492 |
| PC140/181 | 0.015782 | 0.055237 | 0.57092 | -0.80863 | 0.098226 | 0.29948 | 0.6731 | -0.57111 | 0.17074 | 0.27777 | 2.3367 | 1.2244 |
| PI160/161_2 | 0.016114 | 0.056098 | 0.56146 | -0.83275 | 0.14963 | 0.3662 | 0.71768 | -0.47858 | 0.214 | 0.32627 | 1.4489 | 0.535 |
| PA180/182 | 0.016418 | 0.056391 | 0.59567 | -0.74742 | 0.25599 | 0.4721 | 1.3234 | 0.40423 | 0.38715 | 0.51019 | 0.86335 | -0.21198 |
| TAG504-FA160 | 0.016433 | 0.056391 | 0.30526 | -1.7119 | 0.060252 | 0.23917 | 0.39364 | -1.345 | 0.060497 | 0.14096 | 0.43235 | -1.2097 |
| PC182/225 | 0.016458 | 0.056391 | 0.52684 | -0.92456 | 0.9668 | 0.98804 | 1.0556 | 0.078018 | 0.10231 | 0.19305 | 0.6373 | -0.64996 |
| TAG523-FA160 | 0.016557 | 0.056431 | 0.44838 | -1.1572 | 0.46042 | 0.63455 | 0.80087 | -0.32036 | 0.053882 | 0.13832 | 0.51568 | -0.95544 |
| PI180/202 | 0.016857 | 0.057157 | 0.64306 | -0.63697 | 0.000288 | 0.023474 | 0.55462 | -0.85042 | 0.00556 | 0.07096 | 0.55644 | -0.84571 |
| PS180/181_2 | 0.017282 | 0.058295 | 0.67791 | -0.56084 | 0.051155 | 0.22032 | 0.70441 | -0.50551 | 0.13516 | 0.23339 | 0.78626 | -0.34691 |

| | | | | | | | | | | | | |
|---|---|---|---|---|---|---|---|---|---|---|---|---|
| SMd181/203 | 0.01778 | 0.05931 | 1.2819 | 0.35833 | 0.10104 | 0.30035 | 0.87559 | -0.19168 | 0.43585 | 0.55645 | 0.93945 | -0.09011 |
| PE181/202 | 0.017803 | 0.05931 | 0.519 | -0.9462 | 0.02057 | 0.14715 | 0.51625 | -0.95387 | 0.009988 | 0.076615 | 0.44953 | -1.1535 |
| TAG484-FA182 | 0.017857 | 0.05931 | 0.34904 | -1.5185 | 0.056524 | 0.23143 | 0.44381 | -1.172 | 0.67249 | 0.76672 | 1.2881 | 0.36519 |
| PI180/205 | 0.018013 | 0.059524 | 0.63849 | -0.64726 | 0.12373 | 0.3382 | 0.78423 | -0.35065 | 0.097637 | 0.1864 | 0.74176 | -0.43098 |
| PEP-180/182 | 0.018408 | 0.060524 | 0.59387 | -0.75179 | 0.7816 | 0.87879 | 1.0647 | 0.090424 | 0.038032 | 0.1179 | 0.67629 | -0.56429 |
| PE181/182 | 0.018831 | 0.061602 | 0.47342 | -1.0788 | 0.4201 | 0.60733 | 0.86338 | -0.21193 | 0.041883 | 0.12255 | 0.51383 | -0.96063 |
| PEO-160/225 | 0.019807 | 0.064077 | 1.4065 | 0.4921 | 0.046196 | 0.21481 | 1.297 | 0.37513 | 0.94663 | 0.9629 | 0.99487 | -0.00741 |
| PI180/203 | 0.01996 | 0.064077 | 0.67284 | -0.57168 | 0.026927 | 0.16231 | 0.7238 | -0.46633 | 0.031153 | 0.11103 | 0.69836 | -0.51796 |
| LPC226 | 0.020002 | 0.064077 | 1.2764 | 0.35207 | 0.021322 | 0.15087 | 1.5653 | 0.6464 | 0.10384 | 0.19463 | 1.2904 | 0.36782 |
| PEP-180/181 | 0.020017 | 0.064077 | 0.60665 | -0.72107 | 0.81369 | 0.89922 | 1.069 | 0.096329 | 0.25684 | 0.36992 | 0.8319 | -0.26552 |
| TAG569-FA204 | 0.020241 | 0.064077 | 0.51346 | -0.96169 | 0.034114 | 0.18981 | 0.50275 | -0.99208 | 0.002892 | 0.055371 | 0.28387 | -1.8167 |
| PC140/182 | 0.020309 | 0.064077 | 0.59576 | -0.74719 | 0.17365 | 0.39116 | 0.76769 | -0.38141 | 0.089734 | 0.17751 | 2.6945 | 1.43 |
| TAG544-FA182 | 0.020433 | 0.064077 | 0.33349 | -1.5843 | 0.94084 | 0.97158 | 1.0276 | 0.039248 | 0.072766 | 0.15896 | 0.46063 | -1.1183 |
| TAG522-FA181 | 0.020443 | 0.064077 | 0.41612 | -1.2649 | 0.55287 | 0.7074 | 0.82041 | -0.28557 | 0.096962 | 0.18565 | 0.54322 | -0.88038 |
| PEP-180/203 | 0.020473 | 0.064077 | 0.67567 | -0.5656 | 0.41159 | 0.60329 | 1.0851 | 0.11779 | 0.013967 | 0.077008 | 0.66808 | -0.5819 |
| PC150/182 | 0.020835 | 0.064204 | 0.57294 | -0.80354 | 0.11692 | 0.3295 | 0.68572 | -0.54431 | 0.11609 | 0.21169 | 0.69067 | -0.53394 |
| TAG566-FA182 | 0.020873 | 0.064204 | 0.36752 | -1.4441 | 0.27489 | 0.49163 | 0.68103 | -0.55422 | 0.034039 | 0.11481 | 0.39998 | -1.322 |
| PC182/201 | 0.020894 | 0.064204 | 0.55664 | -0.84518 | 0.60907 | 0.75818 | 0.90346 | -0.14647 | 0.53096 | 0.64488 | 0.89716 | -0.15656 |
| TAG567-FA182 | 0.020908 | 0.064204 | 0.41168 | -1.2804 | 0.51267 | 0.67414 | 0.87185 | -0.19785 | 0.023193 | 0.10133 | 0.37745 | -1.4056 |
| PC160/205 | 0.021046 | 0.064323 | 1.4874 | 0.57277 | 0.2063 | 0.42499 | 1.2885 | 0.36574 | 0.071505 | 0.15673 | 0.76741 | -0.38194 |
| TAG568-FA183 | 0.021365 | 0.064783 | 0.34445 | -1.5376 | 0.049866 | 0.21956 | 0.39431 | -1.3426 | 0.012938 | 0.076615 | 0.25555 | -1.9683 |
| TAG524-FA203 | 0.021395 | 0.064783 | 0.27803 | -1.8467 | 0.12416 | 0.3382 | 0.42736 | -1.2265 | 0.10997 | 0.20222 | 0.42238 | -1.2434 |
| HCER180 | 0.02163 | 0.065096 | 1.59 | 0.66905 | 0.29811 | 0.5134 | 1.2572 | 0.33019 | 0.83571 | 0.88463 | 0.97208 | -0.04085 |
| TAG567-FA203 | 0.021699 | 0.065096 | 0.41575 | -1.2662 | 0.073336 | 0.25947 | 0.48816 | -1.0346 | 0.008706 | 0.075886 | 0.29336 | -1.7693 |
| PC170/203 | 0.022119 | 0.065843 | 0.64465 | -0.63341 | 0.00842 | 0.10202 | 0.5907 | -0.7595 | 0.002819 | 0.055371 | 0.48842 | -1.0338 |
| TAG502-FA180 | 0.022303 | 0.065843 | 0.37147 | -1.4287 | 0.042008 | 0.20555 | 0.41082 | -1.2834 | 0.24019 | 0.35217 | 0.62159 | -0.68596 |
| TAG483-FA120 | 0.02233 | 0.065843 | 0.27137 | -1.8817 | 0.20114 | 0.41834 | 0.52349 | -0.93376 | 0.12219 | 0.21794 | 2.1885 | 1.13 |
| SMd181/160 | 0.022352 | 0.065843 | 1.2198 | 0.28666 | 0.81473 | 0.89922 | 1.0131 | 0.018745 | 0.39937 | 0.52312 | 1.0523 | 0.073597 |
| TAG544-FA202 | 0.02299 | 0.067254 | 0.34082 | -1.5529 | 0.15946 | 0.37492 | 0.53152 | -0.91179 | 0.050057 | 0.13527 | 0.39711 | -1.3324 |
| TAG502-FA161 | 0.023174 | 0.067254 | 0.28731 | -1.7993 | 0.12924 | 0.34014 | 0.45005 | -1.1519 | 0.13891 | 0.23681 | 0.454 | -1.1392 |
| PE140/204 | 0.023288 | 0.067254 | 0.61415 | -0.70334 | 0.022984 | 0.15898 | 0.63324 | -0.65917 | 0.91379 | 0.94126 | 1.5465 | 0.62899 |
| TAG563-FA202 | 0.023317 | 0.067254 | 0.32796 | -1.6084 | 0.2723 | 0.49105 | 0.59071 | -0.75947 | 0.027685 | 0.10792 | 0.32212 | -1.6343 |
| TAG569-FA183 | 0.023383 | 0.067254 | 0.45142 | -1.1475 | 0.064158 | 0.24498 | 0.50187 | -0.99461 | 0.002563 | 0.055371 | 0.23289 | -2.1023 |

| | | | | | | | | | | | | |
|---|---|---|---|---|---|---|---|---|---|---|---|---|
| LPE204 | 0.023451 | 0.067254 | 1.467 | 0.55282 | 0.004082 | 0.084684 | 1.5789 | 0.65891 | 0.95894 | 0.97238 | 1.0291 | 0.041413 |
| PI180/224 | 0.023774 | 0.067881 | 0.72325 | -0.46743 | 0.018488 | 0.1416 | 0.76836 | -0.38014 | 0.18911 | 0.29809 | 0.84983 | -0.23475 |
| TAG562-FA180 | 0.023967 | 0.068132 | 0.39156 | -1.3527 | 0.99554 | 0.99737 | 1.0546 | 0.076695 | 0.016916 | 0.08168 | 0.34118 | -1.5514 |
| TAG521-FA161 | 0.024354 | 0.068932 | 0.33737 | -1.5676 | 0.16053 | 0.37591 | 0.51764 | -0.94997 | 0.016093 | 0.079367 | 0.30498 | -1.7132 |
| PS181/182_2 | 0.024933 | 0.069776 | 0.66513 | -0.58829 | 0.59625 | 0.74878 | 1.0835 | 0.11566 | 0.043131 | 0.12369 | 0.71827 | -0.4774 |
| TAG503-FA183 | 0.024962 | 0.069776 | 0.37955 | -1.3977 | 0.069561 | 0.25019 | 0.47093 | -1.0864 | 0.029759 | 0.11007 | 0.38471 | -1.3782 |
| PE170/181 | 0.024979 | 0.069776 | 0.53903 | -0.89156 | 0.092129 | 0.29172 | 0.63296 | -0.65982 | 0.008019 | 0.075886 | 0.41623 | -1.2645 |
| TAG569-FA205 | 0.025174 | 0.069776 | 0.54512 | -0.87534 | 0.058225 | 0.2369 | 0.54604 | -0.87293 | 0.004397 | 0.07096 | 0.2556 | -1.968 |
| TAG563-FA181 | 0.025188 | 0.069776 | 0.32865 | -1.6054 | 0.42448 | 0.60733 | 1.4415 | 0.52753 | 0.082149 | 0.16712 | 0.4388 | -1.1884 |
| PE181/183 | 0.025319 | 0.069842 | 0.48433 | -1.0459 | 0.019055 | 0.14285 | 0.441 | -1.1811 | 0.00169 | 0.055371 | 0.25009 | -1.9995 |
| TAG481-FA160 | 0.025602 | 0.070288 | 0.35459 | -1.4958 | 0.051878 | 0.22051 | 0.40504 | -1.3039 | 0.9932 | 0.99473 | 1.0798 | 0.11071 |
| TAG542-FA160 | 0.025697 | 0.070288 | 0.39671 | -1.3339 | 0.46939 | 0.64265 | 0.78903 | -0.34185 | 0.037085 | 0.11738 | 0.40517 | -1.3034 |
| TAG546-FA203 | 0.02608 | 0.071039 | 0.30062 | -1.734 | 0.09208 | 0.29172 | 0.40924 | -1.289 | 0.012289 | 0.076615 | 0.22969 | -2.1222 |
| TAG567-FA224 | 0.026695 | 0.072206 | 0.36429 | -1.4568 | 0.14348 | 0.36338 | 0.53638 | -0.89867 | 0.011434 | 0.076615 | 0.28981 | -1.7868 |
| TAG501-FA180 | 0.026731 | 0.072206 | 0.36326 | -1.4609 | 0.086561 | 0.28624 | 0.45385 | -1.1397 | 0.334 | 0.45584 | 0.64677 | -0.62868 |
| PC181/161 | 0.026989 | 0.072602 | 0.56106 | -0.83378 | 0.20755 | 0.42623 | 0.70747 | -0.49927 | 0.83476 | 0.88463 | 0.95111 | -0.07232 |
| TAG543-FA180 | 0.027146 | 0.072725 | 0.44322 | -1.1739 | 0.68955 | 0.82101 | 0.95217 | -0.0707 | 0.037466 | 0.11738 | 0.47067 | -1.0872 |
| PC180/204 | 0.027285 | 0.072797 | 1.2301 | 0.29879 | 0.51914 | 0.67848 | 1.0775 | 0.10767 | 0.89292 | 0.92415 | 1.0083 | 0.011888 |
| TAG544-FA181 | 0.027629 | 0.073337 | 0.31825 | -1.6518 | 0.99998 | 0.99998 | 0.9974 | -0.00376 | 0.074377 | 0.15927 | 0.42294 | -1.2415 |
| TAG523-FA200 | 0.027804 | 0.073337 | 0.37286 | -1.4233 | 0.5772 | 0.73246 | 0.77438 | -0.36889 | 0.13845 | 0.23681 | 0.53051 | -0.91454 |
| PI180/224_2 | 0.027825 | 0.073337 | 0.75287 | -0.40952 | 0.007488 | 0.09705 | 0.76203 | -0.39209 | 0.18454 | 0.29302 | 0.84517 | -0.24268 |
| LPC204 | 0.027949 | 0.073366 | 1.2292 | 0.29774 | 0.006991 | 0.095455 | 1.4797 | 0.56531 | 0.096615 | 0.18553 | 1.1525 | 0.20477 |
| TAG522-FA161 | 0.028167 | 0.073642 | 0.28782 | -1.7968 | 0.28194 | 0.49607 | 0.54512 | -0.87536 | 0.078728 | 0.16356 | 0.37581 | -1.4119 |
| TAG543-FA160 | 0.030246 | 0.07876 | 0.37233 | -1.4253 | 0.3622 | 0.57468 | 0.67958 | -0.55729 | 0.057666 | 0.14008 | 0.42414 | -1.2374 |
| TAG544-FA221 | 0.03087 | 0.07983 | 0.32996 | -1.5996 | 0.79217 | 0.88609 | 1.1972 | 0.25968 | 0.043699 | 0.12369 | 0.37555 | -1.4129 |
| TAG542-FA182 | 0.030902 | 0.07983 | 0.43316 | -1.207 | 0.37255 | 0.57883 | 0.74935 | -0.41629 | 0.020304 | 0.094255 | 0.37396 | -1.4191 |
| PC170/181 | 0.031495 | 0.080612 | 0.67173 | -0.57404 | 0.16736 | 0.38229 | 0.77599 | -0.3659 | 0.11035 | 0.20236 | 0.74695 | -0.42092 |
| TAG567-FA161 | 0.031534 | 0.080612 | 0.31687 | -1.658 | 0.34304 | 0.56536 | 0.6047 | -0.72571 | 0.036542 | 0.11738 | 0.32079 | -1.6403 |
| PE180/225 | 0.031576 | 0.080612 | 1.3471 | 0.42985 | 0.76975 | 0.87606 | 0.97492 | -0.03665 | 0.085118 | 0.17237 | 0.75401 | -0.40734 |
| PG182/182 | 0.031975 | 0.081312 | 0.39357 | -1.3453 | 0.076514 | 0.26637 | 0.46026 | -1.1195 | 0.031809 | 0.11231 | 0.38176 | -1.3893 |
| PE181/225 | 0.032957 | 0.082986 | 0.66716 | -0.5839 | 0.12246 | 0.3382 | 0.74179 | -0.43091 | 0.047784 | 0.13255 | 0.60929 | -0.7148 |
| TAG543-FA182 | 0.032999 | 0.082986 | 0.41134 | -1.2816 | 0.72441 | 0.84666 | 0.90701 | -0.1408 | 0.050002 | 0.13527 | 0.45519 | -1.1354 |
| PS180/182 | 0.033016 | 0.082986 | 0.60243 | -0.73114 | 0.4234 | 0.60733 | 0.83114 | -0.26684 | 0.060203 | 0.14096 | 0.64132 | -0.64088 |

| | | | | | | | | | | | | |
|---|---|---|---|---|---|---|---|---|---|---|---|---|
| SMd181/202 | 0.033548 | 0.084 | 1.2929 | 0.3706 | 0.75711 | 0.86774 | 0.97706 | -0.03349 | 0.77363 | 0.8422 | 0.98355 | -0.02393 |
| PE160/225 | 0.034284 | 0.085513 | 1.3334 | 0.41508 | 0.31723 | 0.53108 | 0.86639 | -0.20691 | 0.21213 | 0.32494 | 0.8031 | -0.31635 |
| TAG480-FA140 | 0.034911 | 0.086744 | 0.42115 | -1.2476 | 0.033523 | 0.18977 | 0.41392 | -1.2726 | 0.88712 | 0.92255 | 1.1782 | 0.2366 |
| PC140/140 | 0.035203 | 0.087123 | 0.57634 | -0.795 | 0.007228 | 0.096028 | 0.37915 | -1.3991 | 0.059092 | 0.14096 | 9.5484 | 3.2553 |
| TAG482-FA181 | 0.035331 | 0.087123 | 0.30587 | -1.709 | 0.24098 | 0.46994 | 0.54637 | -0.87205 | 0.090254 | 0.17751 | 2.4366 | 1.2849 |
| TAG564-FA202 | 0.036264 | 0.089087 | 0.28667 | -1.8025 | 0.42179 | 0.60733 | 0.63497 | -0.65523 | 0.057942 | 0.14022 | 0.33996 | -1.5565 |
| TAG543-FA181 | 0.036461 | 0.089121 | 0.36696 | -1.4463 | 0.7727 | 0.87711 | 1.1766 | 0.23462 | 0.16888 | 0.27624 | 0.55408 | -0.85184 |
| TAG564-FA181 | 0.036689 | 0.089121 | 0.31437 | -1.6695 | 0.70637 | 0.83306 | 0.80164 | -0.31898 | 0.051179 | 0.13574 | 0.35971 | -1.4751 |
| PE140/182 | 0.036743 | 0.089121 | 0.56471 | -0.82443 | 0.059878 | 0.23917 | 0.59577 | -0.74718 | 0.94404 | 0.9629 | 1.6189 | 0.69505 |
| TAG543-FA161 | 0.036863 | 0.089121 | 0.29662 | -1.7533 | 0.23692 | 0.46738 | 0.48662 | -1.0391 | 0.056053 | 0.13888 | 0.31676 | -1.6585 |
| TAG504-FA181 | 0.036962 | 0.089121 | 0.30979 | -1.6906 | 0.1942 | 0.41237 | 0.52248 | -0.93656 | 0.23635 | 0.3489 | 0.57574 | -0.79652 |
| TAG568-FA181 | 0.037255 | 0.089494 | 0.51117 | -0.96811 | 0.52007 | 0.67848 | 0.84836 | -0.23724 | 0.004469 | 0.07096 | 0.3248 | -1.6224 |
| PEP-180/202 | 0.038281 | 0.09162 | 0.66734 | -0.58351 | 0.30361 | 0.51605 | 0.86732 | -0.20537 | 0.002594 | 0.055371 | 0.55279 | -0.85521 |
| PE160/203 | 0.038495 | 0.091795 | 0.82273 | -0.28152 | 1.18E-05 | 0.007697 | 0.5672 | -0.81807 | 0.005518 | 0.07096 | 0.57145 | -0.8073 |
| TAG565-FA181 | 0.039421 | 0.093226 | 0.33744 | -1.5673 | 0.41238 | 0.60329 | 0.68183 | -0.55252 | 0.060237 | 0.14096 | 0.39302 | -1.3473 |
| TAG564-FA182 | 0.03944 | 0.093226 | 0.32532 | -1.6201 | 0.97561 | 0.99083 | 0.9361 | -0.09526 | 0.037357 | 0.11738 | 0.33562 | -1.5751 |
| TAG566-FA203 | 0.039525 | 0.093226 | 0.37836 | -1.4022 | 0.25585 | 0.4721 | 0.59517 | -0.74862 | 0.010992 | 0.076615 | 0.27107 | -1.8833 |
| PEP-160/160 | 0.040752 | 0.095774 | 1.4213 | 0.50717 | 0.006582 | 0.095455 | 1.2487 | 0.32046 | 0.50731 | 0.62549 | 1.0647 | 0.090461 |
| TAG522-FA160 | 0.041298 | 0.096545 | 0.43074 | -1.2151 | 0.70482 | 0.83274 | 0.85832 | -0.22041 | 0.14001 | 0.23681 | 0.55796 | -0.84176 |
| LPE224 | 0.041376 | 0.096545 | 1.6263 | 0.70161 | 0.19309 | 0.41213 | 1.3068 | 0.386 | 0.49748 | 0.61805 | 1.1445 | 0.19467 |
| PI161/180_2 | 0.041848 | 0.097296 | 0.73571 | -0.44279 | 0.008797 | 0.10202 | 0.72078 | -0.47237 | 0.38453 | 0.50777 | 0.89649 | -0.15764 |
| PEO-160/205 | 0.042273 | 0.097935 | 0.69692 | -0.52095 | 0.074897 | 0.26356 | 0.73686 | -0.44054 | 0.001951 | 0.055371 | 0.49434 | -1.0164 |
| TAG568-FA204 | 0.042459 | 0.098017 | 0.63722 | -0.65013 | 0.19189 | 0.41092 | 0.75198 | -0.41124 | 0.010455 | 0.076615 | 0.40957 | -1.2878 |
| PC150/181 | 0.044227 | 0.10174 | 0.60483 | -0.72541 | 0.25517 | 0.4721 | 0.77771 | -0.36269 | 0.62963 | 0.73194 | 0.9051 | -0.14386 |
| TAG546-FA161 | 0.044451 | 0.10189 | 0.35161 | -1.5079 | 0.10369 | 0.30335 | 0.43457 | -1.2023 | 0.042017 | 0.12255 | 0.35313 | -1.5017 |
| TAG522-FA182 | 0.045501 | 0.10362 | 0.4815 | -1.0544 | 0.34102 | 0.56346 | 0.72971 | -0.45461 | 0.048283 | 0.13319 | 0.47646 | -1.0696 |
| LPC140 | 0.045524 | 0.10362 | 0.63007 | -0.66643 | 0.77374 | 0.87711 | 0.9513 | -0.07202 | 0.063411 | 0.14484 | 2.8882 | 1.5302 |
| TAG501-FA140 | 0.046052 | 0.10446 | 0.3532 | -1.5014 | 0.21075 | 0.43009 | 0.53785 | -0.89473 | 0.97679 | 0.98435 | 0.97263 | -0.04004 |
| PEP-180/183 | 0.046467 | 0.10504 | 0.63525 | -0.65461 | 0.58483 | 0.73927 | 0.91615 | -0.12634 | 0.00264 | 0.055371 | 0.36885 | -1.4389 |
| TAG543-FA202 | 0.046784 | 0.10539 | 0.35462 | -1.4957 | 0.33275 | 0.5512 | 0.63012 | -0.6663 | 0.080952 | 0.16572 | 0.40719 | -1.2962 |
| PC180/140 | 0.047313 | 0.10621 | 0.67801 | -0.56063 | 0.039836 | 0.20071 | 0.62218 | -0.6846 | 0.025231 | 0.10291 | 3.2178 | 1.6861 |
| TAG568-FA225 | 0.048112 | 0.10763 | 0.40599 | -1.3005 | 0.25152 | 0.4721 | 0.60992 | -0.71331 | 0.028429 | 0.10827 | 0.34382 | -1.5403 |
| SMd181/161 | 0.048282 | 0.10764 | 1.3079 | 0.38727 | 0.17047 | 0.38533 | 1.2235 | 0.29098 | 0.026873 | 0.10603 | 1.3966 | 0.48196 |

| | | | | | | | | | | | | |
|---|---|---|---|---|---|---|---|---|---|---|---|---|
| PEP-180/205 | 0.048933 | 0.10869 | 0.69562 | -0.52363 | 0.96444 | 0.98718 | 1.0029 | 0.00417 | 0.006662 | 0.075886 | 0.52101 | -0.94062 |
| PEP-160/182 | 0.049085 | 0.10869 | 0.65177 | -0.61757 | 0.93876 | 0.97158 | 1.0552 | 0.077577 | 0.22125 | 0.33455 | 0.80301 | -0.31652 |
| SMd181/200 | 0.049617 | 0.10949 | 1.2381 | 0.30812 | 0.066987 | 0.24778 | 0.85284 | -0.22966 | 0.68417 | 0.77325 | 1.0486 | 0.068489 |
| TAG541-FA181 | 0.050664 | 0.11143 | 0.40776 | -1.2942 | 0.81497 | 0.89922 | 0.93719 | -0.09359 | 0.035099 | 0.11738 | 0.37744 | -1.4057 |
| TAG565-FA203 | 0.051226 | 0.11202 | 0.36822 | -1.4414 | 0.3704 | 0.57687 | 0.64638 | -0.62955 | 0.028751 | 0.10827 | 0.32282 | -1.6312 |
| PG160/160 | 0.051279 | 0.11202 | 0.37031 | -1.4332 | 0.5031 | 0.66877 | 0.61186 | -0.70872 | 0.73309 | 0.81301 | 0.89284 | -0.16353 |
| PEP-160/225 | 0.052541 | 0.11367 | 1.2571 | 0.33014 | 0.043093 | 0.20555 | 1.2266 | 0.29468 | 0.4468 | 0.56451 | 1.0409 | 0.057765 |
| PC170/226 | 0.052645 | 0.11367 | 0.71674 | -0.48048 | 0.12332 | 0.3382 | 0.77055 | -0.37604 | 0.043569 | 0.12369 | 0.70259 | -0.50925 |
| TAG522-FA180 | 0.052672 | 0.11367 | 0.50686 | -0.98034 | 0.26911 | 0.488 | 0.69688 | -0.52102 | 0.033845 | 0.11476 | 0.45033 | -1.1509 |
| TAG545-FA203 | 0.05273 | 0.11367 | 0.39943 | -1.324 | 0.19662 | 0.41237 | 0.57677 | -0.79393 | 0.024547 | 0.10291 | 0.33458 | -1.5796 |
| PC181/182 | 0.053119 | 0.11413 | 0.67491 | -0.56723 | 0.9888 | 0.99645 | 1.0643 | 0.089952 | 0.43593 | 0.55645 | 0.85885 | -0.21952 |
| TAG548-FA205 | 0.054767 | 0.11722 | 0.39307 | -1.3471 | 0.060186 | 0.23917 | 0.40817 | -1.2928 | 0.007763 | 0.075886 | 0.23864 | -2.0671 |
| TAG562-FA160 | 0.054919 | 0.11722 | 0.41426 | -1.2714 | 0.80228 | 0.89432 | 1.1238 | 0.16834 | 0.047572 | 0.13255 | 0.38656 | -1.3712 |
| PG160/180_2 | 0.056935 | 0.12113 | 0.59267 | -0.7547 | 0.101 | 0.30035 | 0.6199 | -0.68988 | 0.059308 | 0.14096 | 0.57498 | -0.79842 |
| PEP-160/205 | 0.057377 | 0.12167 | 0.72809 | -0.45782 | 0.72217 | 0.8464 | 0.96289 | -0.05456 | 0.005558 | 0.07096 | 0.55659 | -0.84531 |
| TAG543-FA201 | 0.057863 | 0.12182 | 0.39868 | -1.3267 | 0.64625 | 0.78637 | 0.83245 | -0.26457 | 0.10404 | 0.19463 | 0.45917 | -1.1229 |
| PEP-181/182 | 0.057913 | 0.12182 | 0.64532 | -0.63192 | 0.84782 | 0.92381 | 1.13 | 0.17633 | 0.29444 | 0.40958 | 0.81745 | -0.2908 |
| PG160/180 | 0.058011 | 0.12182 | 0.48175 | -1.0536 | 0.14221 | 0.36306 | 0.56359 | -0.82727 | 0.090137 | 0.17751 | 0.51138 | -0.96755 |
| PEO-160/182 | 0.05894 | 0.12301 | 0.64652 | -0.62923 | 0.35362 | 0.57409 | 0.83517 | -0.25985 | 0.027205 | 0.10669 | 0.5921 | -0.75608 |
| TAG483-FA160 | 0.058953 | 0.12301 | 0.501 | -0.99711 | 0.0387 | 0.19995 | 0.42414 | -1.2374 | 0.27203 | 0.38583 | 0.77793 | -0.36229 |
| TAG547-FA161 | 0.05988 | 0.12437 | 0.44066 | -1.1823 | 0.14358 | 0.36338 | 0.52867 | -0.91957 | 0.024858 | 0.10291 | 0.35136 | -1.509 |
| TAG565-FA224 | 0.059988 | 0.12437 | 0.38198 | -1.3884 | 0.55309 | 0.7074 | 0.80551 | -0.31203 | 0.14762 | 0.24704 | 0.49174 | -1.024 |
| TAG502-FA160 | 0.060278 | 0.12458 | 0.50747 | -0.97862 | 0.2129 | 0.43313 | 0.67401 | -0.56915 | 0.10211 | 0.19305 | 0.5779 | -0.79112 |
| PE180/150 | 0.060532 | 0.1247 | 0.80424 | -0.31429 | 0.008823 | 0.10202 | 0.71847 | -0.477 | 0.00873 | 0.075886 | 0.74998 | -0.41508 |
| LPE180 | 0.061384 | 0.12602 | 1.3314 | 0.41293 | 0.15447 | 0.37014 | 1.2276 | 0.29582 | 0.31494 | 0.43164 | 0.87219 | -0.19728 |
| TAG502-FA182 | 0.061557 | 0.12602 | 0.52743 | -0.92295 | 0.25493 | 0.4721 | 0.73773 | -0.43883 | 0.10584 | 0.1963 | 0.60134 | -0.73375 |
| PEO-180/224 | 0.062582 | 0.12724 | 1.4632 | 0.54913 | 0.81906 | 0.90222 | 1.0453 | 0.063929 | 0.65449 | 0.75411 | 0.96778 | -0.04724 |
| TAG483-FA181 | 0.062779 | 0.12724 | 0.44975 | -1.1528 | 0.37763 | 0.58387 | 0.69726 | -0.52023 | 0.079306 | 0.16356 | 2.5214 | 1.3342 |
| PE182/183 | 0.062797 | 0.12724 | 0.3961 | -1.3361 | 0.015049 | 0.12819 | 0.27134 | -1.8818 | 0.002234 | 0.055371 | 0.16441 | -2.6046 |
| PG160/181 | 0.062936 | 0.12724 | 0.69567 | -0.52352 | 0.077644 | 0.26744 | 0.67915 | -0.5582 | 0.092516 | 0.18086 | 0.67295 | -0.57144 |
| PEP-180/180 | 0.06349 | 0.12739 | 0.70113 | -0.51225 | 0.75378 | 0.86545 | 1.0726 | 0.10113 | 0.28967 | 0.4045 | 0.83913 | -0.25303 |
| PS160/180 | 0.063526 | 0.12739 | 1.1892 | 0.25004 | 0.129 | 0.34014 | 0.84054 | -0.25061 | 0.2399 | 0.35217 | 0.86861 | -0.20322 |
| PG180/181_2 | 0.063598 | 0.12739 | 0.79925 | -0.32328 | 0.011735 | 0.10914 | 0.72928 | -0.45545 | 0.023558 | 0.10224 | 0.65381 | -0.61305 |

| | | | | | | | | | | | | |
|---|---|---|---|---|---|---|---|---|---|---|---|---|
| PC160/160 | 0.063837 | 0.12748 | 1.2146 | 0.28048 | 0.39255 | 0.59708 | 1.098 | 0.13486 | 0.92865 | 0.95351 | 1.0077 | 0.011106 |
| TAG562-FA201 | 0.064326 | 0.12806 | 0.39213 | -1.3506 | 0.53409 | 0.69261 | 1.3124 | 0.39225 | 0.044764 | 0.12615 | 0.3516 | -1.508 |
| PEO-160/224 | 0.064701 | 0.12842 | 1.2756 | 0.35118 | 0.89661 | 0.94765 | 0.99777 | -0.00323 | 0.1979 | 0.30822 | 0.89762 | -0.15583 |
| PE170/204 | 0.065208 | 0.12903 | 0.6554 | -0.60956 | 0.013846 | 0.12352 | 0.55704 | -0.84415 | 0.007495 | 0.075886 | 0.46668 | -1.0995 |
| TAG566-FA224 | 0.065986 | 0.12992 | 0.45476 | -1.1368 | 0.40077 | 0.5989 | 0.81264 | -0.29932 | 0.054311 | 0.13832 | 0.43267 | -1.2087 |
| PE182/182 | 0.066059 | 0.12992 | 0.40745 | -1.2953 | 0.36638 | 0.57474 | 0.67704 | -0.56268 | 0.011801 | 0.076615 | 0.2711 | -1.8831 |
| PC181/225 | 0.06685 | 0.13108 | 0.72977 | -0.45449 | 0.18184 | 0.40402 | 0.79718 | -0.32702 | 0.077441 | 0.1621 | 0.72385 | -0.46623 |
| PI161/180 | 0.067206 | 0.13128 | 0.66881 | -0.58032 | 0.081768 | 0.27725 | 0.73487 | -0.44444 | 0.85074 | 0.89636 | 0.98307 | -0.02464 |
| LPE225 | 0.067353 | 0.13128 | 1.525 | 0.60876 | 0.048672 | 0.21852 | 1.4978 | 0.58282 | 0.63178 | 0.73313 | 1.1568 | 0.21017 |
| HCER200 | 0.068662 | 0.13343 | 1.3854 | 0.47035 | 0.874 | 0.93651 | 0.96622 | -0.04958 | 0.66613 | 0.76213 | 0.93372 | -0.09894 |
| LPC203 | 0.069959 | 0.13525 | 0.76794 | -0.38093 | 0.30248 | 0.51578 | 0.85909 | -0.21913 | 0.007371 | 0.075886 | 0.68079 | -0.55471 |
| TAG544-FA160 | 0.070016 | 0.13525 | 0.43112 | -1.2138 | 0.30464 | 0.51645 | 0.64623 | -0.62988 | 0.047847 | 0.13255 | 0.39833 | -1.328 |
| PE180/203 | 0.07079 | 0.1363 | 0.7804 | -0.35771 | 0.001808 | 0.053488 | 0.59691 | -0.74441 | 0.001865 | 0.055371 | 0.50404 | -0.9884 |
| TAG546-FA204 | 0.070978 | 0.1363 | 0.64009 | -0.64365 | 0.47547 | 0.64504 | 0.9453 | -0.08116 | 0.061761 | 0.14157 | 0.58085 | -0.78376 |
| TAG568-FA161 | 0.07142 | 0.13675 | 0.4572 | -1.1291 | 0.51696 | 0.67714 | 0.71744 | -0.47907 | 0.029187 | 0.10858 | 0.37134 | -1.4292 |
| PEO-180/182 | 0.072202 | 0.13777 | 0.64534 | -0.63186 | 0.38141 | 0.58465 | 0.83219 | -0.26502 | 0.013741 | 0.077008 | 0.53305 | -0.90767 |
| PS180/180 | 0.072377 | 0.13777 | 1.2463 | 0.31761 | 0.197 | 0.41237 | 0.84169 | -0.24863 | 0.39671 | 0.52069 | 0.90381 | -0.14591 |
| TAG561-FA160 | 0.072733 | 0.13804 | 0.45515 | -1.1356 | 0.18007 | 0.40145 | 2.0242 | 1.0173 | 0.070585 | 0.15524 | 0.43583 | -1.1982 |
| PEP-160/203 | 0.073957 | 0.13996 | 0.77136 | -0.37452 | 0.90317 | 0.94765 | 1.0194 | 0.02775 | 0.12611 | 0.22235 | 0.83307 | -0.2635 |
| TAG563-FA201 | 0.074311 | 0.14022 | 0.36449 | -1.4561 | 0.38168 | 0.58465 | 1.5443 | 0.62698 | 0.1427 | 0.24005 | 0.45612 | -1.1325 |
| PE181/224 | 0.074753 | 0.14065 | 0.68752 | -0.54053 | 0.037968 | 0.19911 | 0.61321 | -0.70555 | 0.16183 | 0.26807 | 0.71516 | -0.48366 |
| TAG521-FA180 | 0.075519 | 0.14168 | 0.49859 | -1.0041 | 0.4699 | 0.64265 | 0.80323 | -0.31612 | 0.060734 | 0.14096 | 0.49594 | -1.0118 |
| TAG547-FA205 | 0.076373 | 0.14287 | 0.59467 | -0.74984 | 0.16621 | 0.38229 | 0.69376 | -0.5275 | 0.009327 | 0.075886 | 0.3477 | -1.5241 |
| PG160/181_2 | 0.076773 | 0.14321 | 0.7451 | -0.4245 | 0.061618 | 0.24108 | 0.69192 | -0.53133 | 0.051295 | 0.13574 | 0.64645 | -0.62939 |
| PE170/182 | 0.079151 | 0.14722 | 0.60871 | -0.71618 | 0.015163 | 0.12819 | 0.46239 | -1.1128 | 0.006036 | 0.073558 | 0.38616 | -1.3727 |
| PC200/203 | 0.082126 | 0.152 | 0.75955 | -0.39679 | 0.000524 | 0.031013 | 0.53447 | -0.90382 | 0.000373 | 0.055371 | 0.48784 | -1.0355 |
| TAG547-FA204 | 0.08219 | 0.152 | 0.55413 | -0.8517 | 0.18537 | 0.40645 | 0.65392 | -0.61282 | 0.008529 | 0.075886 | 0.36956 | -1.4361 |
| TAG526-FA204 | 0.082549 | 0.15224 | 0.58123 | -0.78281 | 0.048582 | 0.21852 | 0.52963 | -0.91694 | 0.16334 | 0.26852 | 0.64714 | -0.62784 |
| TAG542-FA181 | 0.0828 | 0.15227 | 0.44882 | -1.1558 | 0.62087 | 0.77135 | 1.3347 | 0.41653 | 0.14695 | 0.24656 | 0.54822 | -0.86716 |
| PI180/183 | 0.083096 | 0.15238 | 0.63949 | -0.64501 | 0.070965 | 0.25251 | 0.64665 | -0.62894 | 0.089278 | 0.17731 | 0.65427 | -0.61205 |
| TAG520-FA160 | 0.08394 | 0.15338 | 0.43942 | -1.1863 | 0.2222 | 0.44646 | 0.56282 | -0.82924 | 0.030481 | 0.11029 | 0.35415 | -1.4976 |
| TAG521-FA181 | 0.084339 | 0.15338 | 0.48252 | -1.0514 | 0.77875 | 0.87711 | 0.94354 | -0.08385 | 0.13664 | 0.23532 | 0.56412 | -0.82591 |
| TAG544-FA203 | 0.084349 | 0.15338 | 0.41721 | -1.2612 | 0.27449 | 0.49163 | 0.60989 | -0.71338 | 0.059021 | 0.14096 | 0.3973 | -1.3317 |

| | | | | | | | | | | | | |
|---|---|---|---|---|---|---|---|---|---|---|---|---|
| PEO-180/225 | 0.084661 | 0.15352 | 1.5637 | 0.64499 | 0.061845 | 0.24108 | 1.3464 | 0.4291 | 0.85092 | 0.89636 | 1.0274 | 0.039058 |
| TAG546-FA225 | 0.088376 | 0.15981 | 0.47657 | -1.0692 | 0.23619 | 0.46736 | 0.63531 | -0.65447 | 0.055963 | 0.13888 | 0.43543 | -1.1995 |
| TAG542-FA202 | 0.090967 | 0.16404 | 0.48335 | -1.0489 | 0.24996 | 0.4721 | 0.66138 | -0.59645 | 0.052673 | 0.13716 | 0.43356 | -1.2057 |
| TAG564-FA203 | 0.091215 | 0.16404 | 0.40671 | -1.2979 | 0.4998 | 0.66673 | 0.7379 | -0.4385 | 0.041678 | 0.12255 | 0.34712 | -1.5265 |
| PC160/202 | 0.092562 | 0.166 | 0.80739 | -0.30866 | 0.004062 | 0.084684 | 0.67205 | -0.57336 | 0.055859 | 0.13888 | 0.76326 | -0.38976 |
| TAG563-FA180 | 0.093737 | 0.16765 | 0.42764 | -1.2255 | 0.62534 | 0.77395 | 0.78846 | -0.3429 | 0.037558 | 0.11738 | 0.34978 | -1.5155 |
| PE180/160 | 0.094534 | 0.16861 | 1.2418 | 0.31242 | 0.14923 | 0.3662 | 0.79674 | -0.32782 | 0.17154 | 0.27777 | 0.80082 | -0.32045 |
| FFA205 | 0.095876 | 0.17053 | 1.4076 | 0.49328 | 0.061459 | 0.24108 | 1.5624 | 0.64381 | 0.82174 | 0.88276 | 0.94465 | -0.08215 |
| PE180/201 | 0.09699 | 0.17176 | 0.76907 | -0.37882 | 0.028275 | 0.16734 | 0.72584 | -0.46227 | 0.024192 | 0.10291 | 0.6637 | -0.59139 |
| PC180/203 | 0.097093 | 0.17176 | 0.78368 | -0.35166 | 0.009873 | 0.10332 | 0.63933 | -0.64536 | 0.013124 | 0.07668 | 0.6511 | -0.61905 |
| PG180/180 | 0.097365 | 0.17177 | 0.81841 | -0.2891 | 0.010067 | 0.10332 | 0.69275 | -0.52958 | 0.014398 | 0.077461 | 0.63078 | -0.6648 |
| PEP-180/204 | 0.099194 | 0.17453 | 0.81615 | -0.29309 | 0.15522 | 0.37014 | 1.1347 | 0.18231 | 0.59156 | 0.70016 | 0.95444 | -0.06728 |
| PEO-180/160 | 0.10051 | 0.17636 | 1.4498 | 0.53582 | 0.52656 | 0.68559 | 1.1219 | 0.16595 | 0.26514 | 0.3777 | 0.83965 | -0.25214 |
| TAG541-FA201 | 0.10139 | 0.17744 | 0.41414 | -1.2718 | 0.62846 | 0.77633 | 0.77239 | -0.37261 | 0.11713 | 0.21299 | 0.45091 | -1.1491 |
| PC180/161 | 0.10379 | 0.18114 | 0.67541 | -0.56617 | 0.026056 | 0.15898 | 0.5703 | -0.8102 | 0.59261 | 0.70016 | 1.1531 | 0.20551 |
| TAG568-FA205 | 0.10541 | 0.18347 | 0.66099 | -0.5973 | 0.22786 | 0.45503 | 0.73246 | -0.44918 | 0.005389 | 0.07096 | 0.30227 | -1.7261 |
| PI180/204 | 0.10599 | 0.18392 | 0.80318 | -0.3162 | 0.73232 | 0.85217 | 0.95273 | -0.06986 | 0.43724 | 0.55703 | 1.0873 | 0.12078 |
| PI180/204_2 | 0.10623 | 0.18392 | 0.81262 | -0.29935 | 0.4214 | 0.60733 | 0.92217 | -0.1169 | 0.22429 | 0.3372 | 1.1338 | 0.18114 |
| TAG546-FA205 | 0.10763 | 0.18586 | 0.60478 | -0.72551 | 0.4591 | 0.63455 | 0.87085 | -0.1995 | 0.039866 | 0.12112 | 0.50003 | -0.99992 |
| PC170/182 | 0.10948 | 0.18856 | 0.76832 | -0.38022 | 0.18647 | 0.40645 | 0.77882 | -0.36063 | 0.04153 | 0.12255 | 0.68307 | -0.5499 |
| TAG542-FA201 | 0.11086 | 0.19009 | 0.3959 | -1.3368 | 0.93489 | 0.97158 | 0.89613 | -0.15823 | 0.29936 | 0.41384 | 0.555 | -0.84943 |
| TAG521-FA160 | 0.11096 | 0.19009 | 0.48934 | -1.0311 | 0.70358 | 0.83274 | 0.87697 | -0.18939 | 0.086317 | 0.17397 | 0.4781 | -1.0646 |
| TAG482-FA120 | 0.11187 | 0.19114 | 0.46728 | -1.0977 | 0.64242 | 0.78318 | 1.0161 | 0.023043 | 0.016043 | 0.079367 | 4.4515 | 2.1543 |
| TAG564-FA180 | 0.1139 | 0.19411 | 0.48746 | -1.0367 | 0.48324 | 0.65267 | 0.77107 | -0.37507 | 0.040887 | 0.12189 | 0.39493 | -1.3403 |
| LPC180 | 0.11437 | 0.19439 | 0.84411 | -0.2445 | 0.44958 | 0.63064 | 1.0749 | 0.10417 | 0.44745 | 0.56451 | 0.92312 | -0.1154 |
| PC181/205 | 0.11495 | 0.19488 | 0.69538 | -0.52414 | 0.095667 | 0.29516 | 0.69039 | -0.53451 | 0.005668 | 0.07096 | 0.50908 | -0.97404 |
| TAG567-FA205 | 0.11647 | 0.19693 | 0.62341 | -0.68174 | 0.63725 | 0.77979 | 0.91541 | -0.12751 | 0.014006 | 0.077008 | 0.37591 | -1.4115 |
| PE181/201 | 0.11712 | 0.19752 | 0.69208 | -0.531 | 0.65719 | 0.79522 | 1.2587 | 0.33191 | 0.079477 | 0.16356 | 0.63271 | -0.66037 |
| TAG501-FA161 | 0.1183 | 0.19901 | 0.41397 | -1.2724 | 0.23842 | 0.46891 | 0.51433 | -0.95922 | 0.1712 | 0.27777 | 0.46116 | -1.1167 |
| PG180/181 | 0.12132 | 0.20311 | 0.83233 | -0.26477 | 0.00983 | 0.10332 | 0.72107 | -0.47179 | 0.02545 | 0.10291 | 0.65282 | -0.61523 |
| PI180/205_2 | 0.12136 | 0.20311 | 0.74553 | -0.42366 | 0.15305 | 0.36901 | 0.76734 | -0.38205 | 0.05065 | 0.13574 | 0.66235 | -0.59433 |
| TAG541-FA180 | 0.12239 | 0.20429 | 0.47102 | -1.0861 | 0.60508 | 0.75751 | 0.85781 | -0.22127 | 0.030903 | 0.11103 | 0.35675 | -1.487 |
| PC200/205 | 0.12344 | 0.20552 | 0.74937 | -0.41624 | 0.05318 | 0.22051 | 0.65437 | -0.61182 | 0.002404 | 0.055371 | 0.43578 | -1.1983 |

| | | | | | | | | | | | | |
|---|---|---|---|---|---|---|---|---|---|---|---|---|
| PC150/226 | 0.12599 | 0.20923 | 0.749 | -0.41697 | 0.18605 | 0.40645 | 0.77367 | -0.3702 | 0.26504 | 0.3777 | 0.80937 | -0.30513 |
| PC200/181 | 0.12671 | 0.2099 | 0.72188 | -0.47016 | 0.76041 | 0.87 | 1.0977 | 0.1345 | 0.70345 | 0.78931 | 0.95921 | -0.06009 |
| TAG527-FA160 | 0.13041 | 0.21547 | 0.50006 | -0.99981 | 0.25451 | 0.4721 | 0.61018 | -0.7127 | 0.035213 | 0.11738 | 0.38576 | -1.3742 |
| LPC181 | 0.13169 | 0.21703 | 0.7938 | -0.33315 | 0.099368 | 0.29948 | 1.4238 | 0.50975 | 0.28527 | 0.40197 | 1.1721 | 0.22904 |
| PE181/181 | 0.13234 | 0.21744 | 0.63383 | -0.65784 | 0.2769 | 0.49386 | 2.3146 | 1.2108 | 0.17959 | 0.28655 | 0.65924 | -0.60113 |
| SMd181/120 | 0.1326 | 0.21744 | 0.7205 | -0.47293 | 0.018393 | 0.1416 | 0.5463 | -0.87224 | 0.016443 | 0.080485 | 2.4055 | 1.2663 |
| PE160/161 | 0.13401 | 0.21919 | 0.61098 | -0.71081 | 0.002339 | 0.063441 | 0.34538 | -1.5337 | 0.35711 | 0.48033 | 0.82445 | -0.2785 |
| PEO-180/183 | 0.13951 | 0.22763 | 0.73979 | -0.43482 | 0.007603 | 0.09705 | 0.55847 | -0.84044 | 0.000458 | 0.055371 | 0.28722 | -1.7998 |
| TAG568-FA182 | 0.14246 | 0.23186 | 0.65861 | -0.60251 | 0.53028 | 0.68905 | 0.87829 | -0.18724 | 0.030095 | 0.11007 | 0.44597 | -1.165 |
| TAG521-FA201 | 0.1429 | 0.23193 | 0.46864 | -1.0934 | 0.34971 | 0.57071 | 0.66139 | -0.59642 | 0.26342 | 0.37689 | 0.57344 | -0.80228 |
| TAG543-FA203 | 0.14322 | 0.23193 | 0.48007 | -1.0587 | 0.2174 | 0.43952 | 0.57375 | -0.80151 | 0.037672 | 0.11738 | 0.36396 | -1.4582 |
| PE181/161 | 0.14514 | 0.23446 | 0.75029 | -0.41448 | 0.31816 | 0.53108 | 1.2335 | 0.30271 | 0.82838 | 0.88345 | 1.0076 | 0.010974 |
| TAG501-FA181 | 0.14604 | 0.23533 | 0.53629 | -0.89892 | 0.47162 | 0.64366 | 0.7944 | -0.33207 | 0.25336 | 0.36735 | 0.65663 | -0.60684 |
| TAG566-FA204 | 0.14657 | 0.23561 | 0.64461 | -0.63349 | 0.78717 | 0.88201 | 1.2276 | 0.29582 | 0.1277 | 0.22347 | 0.65147 | -0.61822 |
| LPC225 | 0.14736 | 0.23629 | 1.2536 | 0.32604 | 0.011045 | 0.10592 | 1.5359 | 0.61908 | 0.45016 | 0.56684 | 1.122 | 0.16611 |
| PE160/183 | 0.15211 | 0.2433 | 0.81482 | -0.29545 | 0.00015 | 0.018452 | 0.34037 | -1.5548 | 0.000814 | 0.055371 | 0.33089 | -1.5956 |
| HCER160 | 0.15376 | 0.24534 | 1.2182 | 0.28473 | 0.85614 | 0.92737 | 0.98735 | -0.01837 | 0.028694 | 0.10827 | 0.78111 | -0.35641 |
| TAG547-FA225 | 0.15438 | 0.24542 | 0.51397 | -0.96025 | 0.18782 | 0.40645 | 0.56628 | -0.8204 | 0.054787 | 0.13832 | 0.40069 | -1.3194 |
| TAG564-FA160 | 0.15457 | 0.24542 | 0.43082 | -1.2148 | 0.37987 | 0.58462 | 0.67018 | -0.57739 | 0.06989 | 0.15423 | 0.36417 | -1.4573 |
| PG160/202_2 | 0.15707 | 0.24879 | 0.70359 | -0.5072 | 0.053916 | 0.22215 | 0.5816 | -0.7819 | 0.039221 | 0.11987 | 0.50659 | -0.98112 |
| TAG540-FA180 | 0.15911 | 0.25141 | 0.53446 | -0.90385 | 0.1878 | 0.40645 | 0.55856 | -0.84023 | 0.055692 | 0.13888 | 0.40692 | -1.2972 |
| PE180/161 | 0.16205 | 0.25543 | 0.78677 | -0.34599 | 0.00017 | 0.018452 | 0.47386 | -1.0775 | 0.10928 | 0.20153 | 0.7303 | -0.45344 |
| PE180/226 | 0.16354 | 0.25715 | 1.1608 | 0.21509 | 0.35901 | 0.57468 | 0.8915 | -0.16569 | 0.10571 | 0.1963 | 0.78212 | -0.35455 |
| TAG563-FA160 | 0.16484 | 0.25858 | 0.52936 | -0.91768 | 0.85182 | 0.92422 | 0.92089 | -0.1189 | 0.051894 | 0.13622 | 0.41065 | -1.284 |
| SMd181/226 | 0.17075 | 0.26555 | 1.1965 | 0.25881 | 0.062589 | 0.24253 | 0.82062 | -0.28522 | 0.86229 | 0.90541 | 0.98468 | -0.02227 |
| PEO-180/205 | 0.17077 | 0.26555 | 0.78358 | -0.35185 | 0.94173 | 0.97158 | 0.98005 | -0.02907 | 0.021131 | 0.094255 | 0.58283 | -0.77884 |
| HCER220 | 0.17085 | 0.26555 | 1.1849 | 0.24474 | 0.87531 | 0.93651 | 1.003 | 0.004295 | 0.62788 | 0.73194 | 1.0734 | 0.10223 |
| TAG545-FA204 | 0.17091 | 0.26555 | 0.64238 | -0.63851 | 0.98463 | 0.99607 | 1.1252 | 0.17022 | 0.20556 | 0.31785 | 0.7215 | -0.47094 |
| SMd181/220 | 0.17186 | 0.26638 | 1.1038 | 0.1425 | 0.00084 | 0.0376 | 0.74148 | -0.43152 | 0.93154 | 0.95351 | 1.0131 | 0.018844 |
| TAG567-FA180 | 0.17235 | 0.2665 | 0.5933 | -0.75317 | 0.42442 | 0.60733 | 0.78957 | -0.34085 | 0.036277 | 0.11738 | 0.40763 | -1.2947 |
| PC160/182 | 0.17277 | 0.26653 | 1.1108 | 0.15156 | 0.19685 | 0.41237 | 1.1138 | 0.15548 | 0.2893 | 0.4045 | 0.92318 | -0.11532 |
| TAG567-FA225 | 0.17577 | 0.27051 | 0.59066 | -0.75961 | 0.7864 | 0.88201 | 1.056 | 0.078632 | 0.091244 | 0.17892 | 0.53137 | -0.91222 |
| TAG545-FA160 | 0.17769 | 0.27282 | 0.57347 | -0.80222 | 0.41795 | 0.60657 | 0.77354 | -0.37046 | 0.067387 | 0.15036 | 0.47802 | -1.0649 |

| | | | | | | | | | | | | |
|---|---|---|---|---|---|---|---|---|---|---|---|---|
| PEP-181/225 | 0.17811 | 0.27283 | 1.2113 | 0.2766 | 0.051441 | 0.22032 | 1.2523 | 0.32456 | 0.64667 | 0.74774 | 1.0493 | 0.069388 |
| PEP-182/182 | 0.17917 | 0.2738 | 0.75905 | -0.39773 | 0.67869 | 0.81368 | 1.1491 | 0.20056 | 0.38182 | 0.50521 | 0.84772 | -0.23835 |
| TAG546-FA226 | 0.17965 | 0.27389 | 1.7076 | 0.77198 | 0.6754 | 0.81123 | 1.7577 | 0.81368 | 0.54882 | 0.66163 | 0.92363 | -0.11462 |
| PE160/226 | 0.18309 | 0.27826 | 1.2275 | 0.29572 | 0.43223 | 0.61303 | 0.89174 | -0.16531 | 0.12837 | 0.22404 | 0.7722 | -0.37295 |
| FFA203 | 0.18337 | 0.27826 | 0.87573 | -0.19144 | 0.01019 | 0.10332 | 0.80453 | -0.31378 | 0.024416 | 0.10291 | 0.77812 | -0.36194 |
| LPE202 | 0.18599 | 0.28154 | 0.73341 | -0.4473 | 0.45602 | 0.63455 | 0.84062 | -0.25047 | 0.021594 | 0.094982 | 0.53194 | -0.91067 |
| SMd181/205 | 0.1864 | 0.28154 | 1.1437 | 0.19365 | 0.013147 | 0.12055 | 0.78224 | -0.35432 | 0.1729 | 0.27861 | 0.85867 | -0.21982 |
| TAG566-FA160 | 0.18999 | 0.2863 | 0.55695 | -0.84439 | 0.63339 | 0.77799 | 0.92787 | -0.108 | 0.10165 | 0.19293 | 0.50228 | -0.99344 |
| PG160/202 | 0.19055 | 0.28649 | 0.71374 | -0.48653 | 0.068681 | 0.25019 | 0.60242 | -0.73116 | 0.037684 | 0.11738 | 0.50675 | -0.98067 |
| TAG547-FA226 | 0.19109 | 0.28663 | 0.5479 | -0.86802 | 0.4766 | 0.64504 | 0.69695 | -0.52088 | 0.40923 | 0.53281 | 0.66962 | -0.57858 |
| SM241 | 0.19274 | 0.28844 | 1.1429 | 0.19267 | 0.25907 | 0.47509 | 0.92151 | -0.11793 | 0.69074 | 0.77663 | 1.0309 | 0.043859 |
| PEP-181/205 | 0.19368 | 0.28918 | 0.82937 | -0.26992 | 0.62459 | 0.77395 | 1.1151 | 0.15714 | 0.007663 | 0.075886 | 0.61684 | -0.69703 |
| HCER241 | 0.19466 | 0.28999 | 1.1259 | 0.1711 | 0.76524 | 0.87398 | 0.97345 | -0.03883 | 0.079332 | 0.16356 | 0.83814 | -0.25474 |
| FFA226 | 0.19647 | 0.29188 | 1.2052 | 0.26927 | 0.28369 | 0.49779 | 1.1224 | 0.16658 | 0.51251 | 0.63071 | 1.0625 | 0.087459 |
| FFA220 | 0.19683 | 0.29188 | 1.2025 | 0.26599 | 0.097266 | 0.29868 | 1.313 | 0.39288 | 0.22199 | 0.33455 | 1.1794 | 0.23804 |
| PEP-180/160 | 0.20072 | 0.29698 | 1.31 | 0.38954 | 0.40337 | 0.5989 | 1.088 | 0.12166 | 0.41487 | 0.53694 | 0.91817 | -0.12317 |
| PE160/201 | 0.20167 | 0.29705 | 0.78141 | -0.35584 | 0.032353 | 0.18639 | 0.65113 | -0.61899 | 0.049531 | 0.13527 | 0.63203 | -0.66193 |
| TAG565-FA180 | 0.20168 | 0.29705 | 0.60695 | -0.72035 | 0.89779 | 0.94765 | 1.0959 | 0.13214 | 0.13988 | 0.23681 | 0.63308 | -0.65954 |
| PG161/182_2 | 0.20305 | 0.29839 | 0.77301 | -0.37144 | 0.62974 | 0.77644 | 0.92052 | -0.11949 | 0.2248 | 0.3372 | 0.78219 | -0.35441 |
| TAG566-FA181 | 0.20492 | 0.30046 | 0.53853 | -0.89289 | 0.9404 | 0.97158 | 0.9972 | -0.00404 | 0.079646 | 0.16356 | 0.46951 | -1.0908 |
| PC140/225 | 0.20598 | 0.30134 | 0.81805 | -0.28975 | 0.89897 | 0.94765 | 1.0438 | 0.061889 | 0.067889 | 0.15084 | 1.9351 | 0.95241 |
| PC180/181 | 0.2117 | 0.30882 | 0.86331 | -0.21205 | 0.40569 | 0.60024 | 0.9006 | -0.15104 | 0.61849 | 0.72286 | 1.0644 | 0.089977 |
| TAG501-FA160 | 0.21227 | 0.30882 | 0.59669 | -0.74496 | 0.50823 | 0.6706 | 0.7994 | -0.32301 | 0.23546 | 0.34838 | 0.66313 | -0.59265 |
| PC160/224 | 0.21252 | 0.30882 | 1.2135 | 0.27917 | 0.41642 | 0.60657 | 1.15 | 0.20161 | 0.42423 | 0.54689 | 1.1373 | 0.18558 |
| TAG481-FA161 | 0.21308 | 0.30894 | 0.53029 | -0.91516 | 0.25697 | 0.47256 | 0.56407 | -0.82606 | 0.60169 | 0.70704 | 0.74906 | -0.41685 |
| PEP-181/183 | 0.21596 | 0.31243 | 0.76043 | -0.39512 | 0.36815 | 0.57474 | 0.82037 | -0.28566 | 0.007949 | 0.075886 | 0.45138 | -1.1476 |
| PEO-160/203 | 0.21994 | 0.31747 | 0.84514 | -0.24273 | 0.058741 | 0.23752 | 0.7673 | -0.38213 | 0.001257 | 0.055371 | 0.5715 | -0.80717 |
| PC181/201 | 0.223 | 0.32118 | 0.75515 | -0.40517 | 0.36014 | 0.57468 | 1.2595 | 0.33285 | 0.67353 | 0.76672 | 1.1846 | 0.24446 |
| TAG548-FA204 | 0.22405 | 0.32154 | 0.6809 | -0.55448 | 0.2949 | 0.51195 | 0.72912 | -0.45578 | 0.054622 | 0.13832 | 0.44179 | -1.1786 |
| PEP-181/203 | 0.22424 | 0.32154 | 0.80731 | -0.3088 | 0.97477 | 0.99083 | 1.0068 | 0.009791 | 0.075536 | 0.16003 | 0.75722 | -0.40121 |
| PE180/180 | 0.22614 | 0.32355 | 1.2306 | 0.29933 | 0.09124 | 0.29172 | 1.2765 | 0.35216 | 0.35648 | 0.48033 | 1.0959 | 0.13216 |
| TAG565-FA160 | 0.22715 | 0.32392 | 0.51607 | -0.95435 | 0.66302 | 0.79783 | 0.8758 | -0.19133 | 0.18515 | 0.29326 | 0.55151 | -0.85853 |
| SMd181/222 | 0.22739 | 0.32392 | 1.1808 | 0.23972 | 0.21816 | 0.43969 | 0.85395 | -0.22778 | 0.46136 | 0.5787 | 0.91103 | -0.13442 |

| | | | | | | | | | | | | |
|---|---|---|---|---|---|---|---|---|---|---|---|---|
| PC181/204 | 0.22989 | 0.32676 | 0.86146 | -0.21514 | 0.54131 | 0.69781 | 0.9335 | -0.09927 | 0.4266 | 0.54884 | 0.91021 | -0.13573 |
| TAG481-FA180 | 0.23273 | 0.33008 | 0.61242 | -0.70741 | 0.48603 | 0.65372 | 0.81652 | -0.29244 | 0.014195 | 0.077008 | 3.7568 | 1.9095 |
| PC181/181 | 0.23374 | 0.33079 | 0.78961 | -0.34079 | 0.18668 | 0.40645 | 1.6642 | 0.73484 | 0.37682 | 0.49961 | 1.3124 | 0.39224 |
| HCER140 | 0.23503 | 0.3319 | 0.89722 | -0.15646 | 0.010187 | 0.10332 | 0.68264 | -0.5508 | 0.48624 | 0.6064 | 1.3125 | 0.39228 |
| TAG542-FA180 | 0.23593 | 0.33245 | 0.49371 | -1.0183 | 0.74116 | 0.85564 | 1.2728 | 0.34796 | 0.16249 | 0.26843 | 0.49937 | -1.0018 |
| TAG545-FA225 | 0.24091 | 0.33872 | 1.5125 | 0.59694 | 0.5767 | 0.73246 | 1.5738 | 0.65421 | 0.54521 | 0.6585 | 0.95131 | -0.07201 |
| PE170/225 | 0.24385 | 0.34153 | 0.79552 | -0.33003 | 0.019974 | 0.1449 | 0.58756 | -0.76719 | 0.015932 | 0.079367 | 0.53479 | -0.90297 |
| PEO-180/203 | 0.24395 | 0.34153 | 0.85716 | -0.22237 | 0.38648 | 0.58996 | 0.90018 | -0.15172 | 0.004822 | 0.07096 | 0.63369 | -0.65816 |
| PA180/181 | 0.24508 | 0.34237 | 0.86426 | -0.21047 | 0.00682 | 0.095455 | 1.5784 | 0.65845 | 0.11882 | 0.21487 | 1.3443 | 0.42688 |
| PEP-160/202 | 0.24657 | 0.34373 | 0.84328 | -0.24592 | 0.34813 | 0.57071 | 0.87243 | -0.19688 | 0.056669 | 0.13888 | 0.75456 | -0.40629 |
| SMd181/224 | 0.24931 | 0.3468 | 1.1484 | 0.19966 | 0.16359 | 0.379 | 0.82813 | -0.27207 | 0.8149 | 0.87917 | 0.97244 | -0.04032 |
| FFA183 | 0.25395 | 0.3525 | 0.85663 | -0.22326 | 0.10865 | 0.31159 | 0.82931 | -0.27002 | 0.1767 | 0.28403 | 0.84794 | -0.23797 |
| TAG569-FA226 | 0.25605 | 0.35466 | 0.63876 | -0.64666 | 0.53691 | 0.69488 | 0.76598 | -0.38462 | 0.015788 | 0.079367 | 0.33872 | -1.5618 |
| SMd181/184 | 0.2572 | 0.3555 | 0.84384 | -0.24496 | 0.018214 | 0.1416 | 0.71814 | -0.47766 | 0.001781 | 0.055371 | 0.58894 | -0.76381 |
| PE180/183 | 0.26176 | 0.36102 | 0.85237 | -0.23045 | 0.001049 | 0.042695 | 0.46776 | -1.0962 | 0.000326 | 0.055371 | 0.33592 | -1.5738 |
| HCER240 | 0.26765 | 0.36777 | 1.1646 | 0.2199 | 0.80876 | 0.89922 | 1.0452 | 0.063739 | 0.52269 | 0.64082 | 1.0618 | 0.086505 |
| PEP-180/161 | 0.26778 | 0.36777 | 0.79076 | -0.33869 | 0.71573 | 0.84257 | 1.066 | 0.09219 | 0.075208 | 0.16 | 0.66762 | -0.58289 |
| PEP-180/201 | 0.27254 | 0.37352 | 0.8425 | -0.24726 | 0.71729 | 0.84288 | 0.94557 | -0.08074 | 0.28352 | 0.40037 | 0.8492 | -0.23583 |
| PEP-160/226 | 0.28037 | 0.38332 | 1.2353 | 0.30481 | 0.20524 | 0.42415 | 1.2414 | 0.31196 | 0.24929 | 0.36253 | 1.1346 | 0.18214 |
| SMd181/181 | 0.28087 | 0.38332 | 1.2094 | 0.27432 | 0.89548 | 0.94765 | 0.96647 | -0.04921 | 0.24626 | 0.36026 | 1.2537 | 0.32624 |
| LPC205 | 0.28501 | 0.38816 | 1.1483 | 0.19949 | 0.31444 | 0.52894 | 1.2326 | 0.30171 | 0.036915 | 0.11738 | 0.73299 | -0.44814 |
| PEP-160/204 | 0.29094 | 0.39542 | 0.89732 | -0.15631 | 0.26464 | 0.48258 | 1.1426 | 0.19232 | 0.82692 | 0.88345 | 1.0256 | 0.036453 |
| PE170/202 | 0.29239 | 0.39656 | 0.81685 | -0.29185 | 0.15503 | 0.37014 | 0.7747 | -0.36828 | 0.009725 | 0.07628 | 0.59647 | -0.74548 |
| HCER260 | 0.29379 | 0.39762 | 0.80252 | -0.31739 | 0.26613 | 0.48395 | 0.78485 | -0.34952 | 0.55681 | 0.66879 | 0.88801 | -0.17135 |
| PE160/160 | 0.2991 | 0.40397 | 1.1376 | 0.186 | 0.039478 | 0.20071 | 0.73415 | -0.44585 | 0.1216 | 0.21794 | 1.461 | 0.54697 |
| TAG566-FA205 | 0.30011 | 0.4045 | 0.73556 | -0.44309 | 0.93677 | 0.97158 | 1.0761 | 0.10578 | 0.02415 | 0.10291 | 0.44008 | -1.1842 |
| TAG567-FA160 | 0.31006 | 0.41704 | 0.63834 | -0.6476 | 0.99439 | 0.99737 | 1.1383 | 0.18685 | 0.17195 | 0.27777 | 0.60096 | -0.73466 |
| PEP-180/225 | 0.31945 | 0.42879 | 1.2061 | 0.27029 | 0.090543 | 0.29172 | 1.2202 | 0.28709 | 0.70857 | 0.79258 | 1.0303 | 0.043032 |
| PE180/204 | 0.32016 | 0.42885 | 1.0884 | 0.12222 | 0.11532 | 0.3282 | 0.86228 | -0.21377 | 0.008912 | 0.075886 | 0.72658 | -0.4608 |
| PC160/181 | 0.32397 | 0.43275 | 1.094 | 0.12965 | 0.35897 | 0.57468 | 1.0917 | 0.12654 | 0.21521 | 0.32734 | 1.1463 | 0.19698 |
| LPE200 | 0.325 | 0.43275 | 0.77819 | -0.36181 | 0.16343 | 0.379 | 1.4935 | 0.57868 | 0.5261 | 0.64257 | 0.85612 | -0.22411 |
| PC180/226 | 0.32506 | 0.43275 | 1.0794 | 0.11017 | 0.72289 | 0.8464 | 1.0455 | 0.064228 | 0.76836 | 0.83786 | 1.0393 | 0.05562 |
| SMd181/180 | 0.32966 | 0.43797 | 1.1376 | 0.186 | 0.24206 | 0.46994 | 0.8383 | -0.25447 | 0.89066 | 0.92338 | 0.97196 | -0.04104 |

| | | | | | | | | | | | | |
|---|---|---|---|---|---|---|---|---|---|---|---|---|
| PS160/180_2 | 0.33098 | 0.43883 | 1.1088 | 0.14903 | 0.12231 | 0.3382 | 0.79919 | -0.32339 | 0.1772 | 0.28414 | 0.83877 | -0.25366 |
| HCER181 | 0.33417 | 0.44216 | 1.0702 | 0.097851 | 0.56588 | 0.72092 | 1.0446 | 0.062999 | 0.12005 | 0.21589 | 0.87124 | -0.19886 |
| TAG566-FA225 | 0.33622 | 0.44398 | 0.62108 | -0.68715 | 0.84903 | 0.92381 | 1.2258 | 0.2937 | 0.20769 | 0.31963 | 0.62285 | -0.68305 |
| PC150/225 | 0.34323 | 0.45232 | 0.85103 | -0.23271 | 0.06689 | 0.24778 | 0.65084 | -0.61963 | 0.19445 | 0.3043 | 0.796 | -0.32916 |
| TAG546-FA160 | 0.34708 | 0.45511 | 0.75127 | -0.4126 | 0.50887 | 0.6706 | 0.92545 | -0.11178 | 0.085257 | 0.17237 | 0.57109 | -0.80822 |
| TAG567-FA204 | 0.34722 | 0.45511 | 0.66383 | -0.5911 | 0.99226 | 0.99737 | 1.0193 | 0.027547 | 0.12358 | 0.2192 | 0.55716 | -0.84383 |
| LPC182 | 0.34745 | 0.45511 | 0.87163 | -0.19822 | 0.25101 | 0.4721 | 1.2287 | 0.29709 | 0.66541 | 0.76213 | 0.931 | -0.10315 |
| PC181/224 | 0.35105 | 0.45891 | 0.81856 | -0.28883 | 0.29943 | 0.5134 | 0.79869 | -0.32429 | 0.52855 | 0.64436 | 0.86801 | -0.20422 |
| PEO-180/226 | 0.35359 | 0.4613 | 1.3659 | 0.44988 | 0.49236 | 0.66021 | 1.1991 | 0.26194 | 0.98043 | 0.98649 | 1.0031 | 0.004442 |
| SMd181/221 | 0.3614 | 0.47055 | 1.138 | 0.18652 | 0.25078 | 0.4721 | 0.88073 | -0.18322 | 0.04895 | 0.13446 | 1.3061 | 0.38524 |
| SM240 | 0.36382 | 0.47274 | 1.1078 | 0.14763 | 0.011592 | 0.10914 | 0.71552 | -0.48293 | 0.82917 | 0.88345 | 0.9833 | -0.0243 |
| DAG161/182 | 0.36494 | 0.47326 | 0.27408 | -1.8673 | 0.684 | 0.81854 | 0.54476 | -0.87631 | 0.59916 | 0.70534 | 0.60012 | -0.73667 |
| PEO-180/181 | 0.37082 | 0.47908 | 0.85828 | -0.22048 | 0.77661 | 0.87711 | 0.96827 | -0.04652 | 0.092935 | 0.18114 | 0.72459 | -0.46476 |
| PC170/224 | 0.3709 | 0.47908 | 0.83311 | -0.26342 | 0.3626 | 0.57468 | 0.83212 | -0.26514 | 0.073605 | 0.15919 | 0.68954 | -0.53629 |
| TAG520-FA180 | 0.37821 | 0.48756 | 0.62606 | -0.67564 | 0.40963 | 0.6032 | 0.66698 | -0.58428 | 0.16287 | 0.26843 | 0.51353 | -0.96148 |
| TAG565-FA204 | 0.3926 | 0.50511 | 0.74009 | -0.43423 | 0.54392 | 0.69979 | 1.4815 | 0.56707 | 0.23536 | 0.34838 | 0.71157 | -0.49093 |
| PE160/202 | 0.39464 | 0.50673 | 0.84111 | -0.24964 | 0.00557 | 0.088683 | 0.49377 | -1.0181 | 0.035831 | 0.11738 | 0.55664 | -0.84519 |
| PC140/204 | 0.39667 | 0.50817 | 0.87296 | -0.19601 | 0.87609 | 0.93651 | 0.97954 | -0.02983 | 0.11555 | 0.2113 | 2.0242 | 1.0174 |
| PEP-181/181 | 0.39783 | 0.50817 | 0.86864 | -0.20317 | 0.34979 | 0.57071 | 1.2603 | 0.33378 | 0.4132 | 0.53584 | 1.2255 | 0.29342 |
| TAG568-FA160 | 0.3981 | 0.50817 | 0.74186 | -0.43078 | 0.41048 | 0.6032 | 0.77986 | -0.35871 | 0.033021 | 0.11314 | 0.42948 | -1.2193 |
| FFA120 | 0.39951 | 0.50896 | 0.85444 | -0.22695 | 0.36512 | 0.57474 | 0.84634 | -0.24069 | 0.11784 | 0.21368 | 2.0556 | 1.0396 |
| DAG140/160 | 0.40887 | 0.51988 | 1.1806 | 0.23957 | 0.12568 | 0.3396 | 1.7947 | 0.84372 | 0.89881 | 0.92877 | 1.3072 | 0.38647 |
| TAG567-FA181 | 0.42055 | 0.53264 | 0.73143 | -0.45121 | 0.90378 | 0.94765 | 1.0966 | 0.13306 | 0.075715 | 0.16003 | 0.52254 | -0.9364 |
| CER160 | 0.42091 | 0.53264 | 0.86737 | -0.20527 | 0.73367 | 0.85217 | 0.96048 | -0.05817 | 0.6334 | 0.7337 | 0.93181 | -0.10189 |
| TAG482-FA180 | 0.42136 | 0.53264 | 0.78803 | -0.34368 | 0.58895 | 0.74304 | 0.87715 | -0.1891 | 0.026002 | 0.1039 | 3.3001 | 1.7225 |
| LPC221 | 0.42467 | 0.53578 | 0.82722 | -0.27366 | 0.1681 | 0.38263 | 1.3086 | 0.38799 | 0.27082 | 0.38494 | 1.1709 | 0.22768 |
| PEP-181/161 | 0.42903 | 0.53982 | 1.1353 | 0.18313 | 0.44078 | 0.62141 | 1.1734 | 0.23068 | 0.9738 | 0.98286 | 0.96178 | -0.05622 |
| PC160/180 | 0.42954 | 0.53982 | 1.0497 | 0.069931 | 0.48499 | 0.65368 | 1.057 | 0.079974 | 0.61478 | 0.72011 | 1.0348 | 0.049306 |
| PC170/225 | 0.44856 | 0.56264 | 0.90036 | -0.15142 | 0.15773 | 0.37475 | 0.81087 | -0.30246 | 0.021283 | 0.094255 | 0.64268 | -0.63782 |
| FFA225 | 0.45216 | 0.56607 | 1.1337 | 0.18105 | 0.64885 | 0.78806 | 1.0381 | 0.053903 | 0.66517 | 0.76213 | 1.0476 | 0.067022 |
| FFA202 | 0.45465 | 0.56809 | 0.92318 | -0.11531 | 0.021801 | 0.15261 | 0.79903 | -0.32368 | 0.26334 | 0.37689 | 0.89494 | -0.16014 |
| PEP-181/160 | 0.46911 | 0.58504 | 1.2317 | 0.30067 | 0.13005 | 0.34014 | 1.241 | 0.31153 | 0.99677 | 0.99677 | 1.0126 | 0.018057 |
| PEP-160/224 | 0.47086 | 0.5861 | 1.1647 | 0.22001 | 0.84774 | 0.92381 | 1.0547 | 0.076864 | 0.82089 | 0.88276 | 1.0335 | 0.047566 |

| | | | | | | | | | | | | |
|---|---|---|---|---|---|---|---|---|---|---|---|---|
| PEO-160/226 | 0.47287 | 0.58748 | 1.171 | 0.22774 | 0.90398 | 0.94765 | 1.0425 | 0.060109 | 0.53593 | 0.64971 | 0.92805 | -0.10772 |
| PC160/161 | 0.48307 | 0.59901 | 0.8157 | -0.29389 | 0.15899 | 0.37492 | 0.69012 | -0.53508 | 0.3564 | 0.48033 | 1.2355 | 0.3051 |
| SMd181/141 | 0.49864 | 0.61713 | 0.90258 | -0.14788 | 0.10391 | 0.30335 | 0.74446 | -0.42574 | 0.007392 | 0.075886 | 1.8465 | 0.88478 |
| SMd181/170 | 0.50558 | 0.62454 | 1.128 | 0.17379 | 0.12333 | 0.3382 | 0.80282 | -0.31686 | 0.87547 | 0.91628 | 0.97988 | -0.02932 |
| PEO-180/204 | 0.5077 | 0.62597 | 0.9278 | -0.10811 | 0.93811 | 0.97158 | 1.0077 | 0.0111 | 0.10292 | 0.19364 | 0.82291 | -0.28119 |
| FFA140 | 0.50886 | 0.62622 | 0.93364 | -0.09906 | 0.96831 | 0.98804 | 1.0075 | 0.010795 | 0.12702 | 0.22288 | 1.4959 | 0.58106 |
| SMd181/140 | 0.51504 | 0.63262 | 1.082 | 0.11369 | 0.082837 | 0.27941 | 0.80927 | -0.30531 | 0.037105 | 0.11738 | 1.2913 | 0.36878 |
| CER180 | 0.51916 | 0.63649 | 1.44 | 0.52611 | 0.45799 | 0.63455 | 0.85279 | -0.22973 | 0.18757 | 0.29638 | 0.68206 | -0.55204 |
| TAG522-FA202 | 0.52186 | 0.63859 | 0.68943 | -0.53653 | 0.37848 | 0.58387 | 0.71709 | -0.47977 | 0.23318 | 0.34737 | 0.5837 | -0.77671 |
| PE160/140 | 0.52411 | 0.64015 | 0.82121 | -0.28417 | 0.00619 | 0.095455 | 0.44862 | -1.1564 | 0.05126 | 0.13574 | 4.3721 | 2.1283 |
| PEP-160/181 | 0.52646 | 0.64181 | 0.92423 | -0.11368 | 0.18855 | 0.40645 | 1.3108 | 0.39045 | 0.30422 | 0.4187 | 1.2368 | 0.30661 |
| TAG504-FA141 | 0.53008 | 0.64431 | 0.77283 | -0.37177 | 0.66282 | 0.79783 | 1.2093 | 0.27421 | 0.58543 | 0.69547 | 0.98907 | -0.01586 |
| FFA204 | 0.53049 | 0.64431 | 0.89394 | -0.16174 | 0.49287 | 0.66021 | 1.0814 | 0.11296 | 0.76399 | 0.8355 | 0.93235 | -0.10106 |
| CE182 | 0.53166 | 0.64453 | 1.0567 | 0.079525 | 0.007038 | 0.095455 | 0.6077 | -0.71857 | 0.006342 | 0.07372 | 0.63785 | -0.64872 |
| PEP-182/226 | 0.5352 | 0.64761 | 1.1919 | 0.25324 | 0.42759 | 0.60777 | 1.2326 | 0.30169 | 0.76491 | 0.8355 | 0.96461 | -0.05198 |
| CER221 | 0.54214 | 0.65358 | 1.1926 | 0.25412 | 0.12624 | 0.3396 | 0.75993 | -0.39606 | 0.093802 | 0.1812 | 0.78955 | -0.3409 |
| TAG500-FA160 | 0.54289 | 0.65358 | 0.78916 | -0.34162 | 0.45996 | 0.63455 | 0.76582 | -0.38493 | 0.22201 | 0.33455 | 0.60762 | -0.71876 |
| TAG523-FA203 | 0.54315 | 0.65358 | 0.71219 | -0.48966 | 0.36368 | 0.57468 | 0.7152 | -0.48358 | 0.12638 | 0.22235 | 0.49983 | -1.0005 |
| PA180/203 | 0.55063 | 0.66137 | 0.92964 | -0.10526 | 0.28638 | 0.49982 | 1.1127 | 0.15409 | 0.49998 | 0.6188 | 0.91917 | -0.1216 |
| CER260 | 0.55494 | 0.66493 | 0.87981 | -0.18474 | 0.96111 | 0.98689 | 0.98761 | -0.01798 | 0.88177 | 0.9214 | 1.0096 | 0.013823 |
| TAG526-FA226 | 0.55614 | 0.66493 | 0.7249 | -0.46414 | 0.58006 | 0.73467 | 0.73975 | -0.43489 | 0.58931 | 0.6988 | 1.3083 | 0.38773 |
| TAG566-FA180 | 0.55666 | 0.66493 | 0.73969 | -0.43501 | 0.83634 | 0.91814 | 1.1663 | 0.22199 | 0.14005 | 0.23681 | 0.59399 | -0.7515 |
| LPE203 | 0.55875 | 0.6662 | 0.86537 | -0.20862 | 0.42672 | 0.60777 | 0.81195 | -0.30053 | 0.036792 | 0.11738 | 0.57155 | -0.80704 |
| PE160/182 | 0.5733 | 0.68231 | 0.90829 | -0.13877 | 0.026131 | 0.15898 | 0.66387 | -0.59102 | 0.03877 | 0.11962 | 0.62711 | -0.67321 |
| TAG500-FA180 | 0.5762 | 0.6845 | 0.82392 | -0.27942 | 0.27866 | 0.4943 | 0.61943 | -0.69098 | 0.10741 | 0.19864 | 0.48385 | -1.0474 |
| PE160/205 | 0.57843 | 0.68589 | 1.0575 | 0.080718 | 0.14568 | 0.36338 | 0.68808 | -0.53936 | 0.012946 | 0.076615 | 0.49374 | -1.0182 |
| PEP-181/224 | 0.58435 | 0.69165 | 1.121 | 0.16479 | 0.7509 | 0.86366 | 1.0999 | 0.13742 | 0.83249 | 0.88409 | 1.0235 | 0.033576 |
| SM261 | 0.58926 | 0.69546 | 0.9685 | -0.04618 | 0.001345 | 0.045715 | 0.67329 | -0.57071 | 0.35051 | 0.47539 | 0.92995 | -0.10478 |
| PC160/140 | 0.58982 | 0.69546 | 0.88279 | -0.17985 | 0.16129 | 0.37635 | 0.72963 | -0.45476 | 0.030495 | 0.11029 | 2.8011 | 1.486 |
| PC170/205 | 0.59076 | 0.69546 | 0.90421 | -0.14527 | 0.39984 | 0.5989 | 0.84795 | -0.23795 | 0.032101 | 0.11231 | 0.6051 | -0.72476 |
| HCER221 | 0.59823 | 0.70297 | 0.94082 | -0.08801 | 0.22353 | 0.44774 | 0.85641 | -0.22362 | 0.15865 | 0.26348 | 0.82607 | -0.27566 |
| SMd181/201 | 0.59956 | 0.70327 | 1.082 | 0.11374 | 0.40226 | 0.5989 | 0.88545 | -0.17552 | 0.11932 | 0.21518 | 1.3426 | 0.42503 |
| LPE240 | 0.60154 | 0.70432 | 0.91637 | -0.12599 | 0.15142 | 0.36781 | 1.1945 | 0.25643 | 0.54121 | 0.65489 | 1.1316 | 0.17834 |

| | | | | | | | | | | | | |
|---|---|---|---|---|---|---|---|---|---|---|---|---|
| HCER261 | 0.6035 | 0.70535 | 0.90049 | -0.15122 | 0.13717 | 0.35296 | 0.74868 | -0.41758 | 0.043573 | 0.12369 | 0.66781 | -0.58249 |
| PC180/225 | 0.61169 | 0.71363 | 1.0724 | 0.1009 | 0.59695 | 0.74878 | 1.0793 | 0.11006 | 0.55159 | 0.66374 | 0.93931 | -0.09032 |
| SMd181/223 | 0.61841 | 0.72019 | 1.0725 | 0.10095 | 0.093186 | 0.29172 | 0.77761 | -0.36289 | 0.28733 | 0.404 | 0.85193 | -0.2312 |
| SMd181/225 | 0.62191 | 0.72297 | 0.82886 | -0.2708 | 0.39974 | 0.5989 | 0.74486 | -0.42495 | 0.74102 | 0.81902 | 1.0573 | 0.080373 |
| PE160/204 | 0.6237 | 0.72376 | 1.0545 | 0.076574 | 0.26456 | 0.48258 | 0.86466 | -0.2098 | 0.13454 | 0.23317 | 0.78875 | -0.34236 |
| TAG545-FA205 | 0.626 | 0.72448 | 0.85797 | -0.22101 | 0.24327 | 0.46994 | 0.71412 | -0.48577 | 0.011917 | 0.076615 | 0.37243 | -1.4249 |
| PEO-160/204 | 0.62724 | 0.72448 | 0.96671 | -0.04885 | 0.59105 | 0.74425 | 1.0611 | 0.085546 | 0.081398 | 0.16611 | 0.8739 | -0.19446 |
| PEO-160/183 | 0.62766 | 0.72448 | 0.88683 | -0.17327 | 0.10271 | 0.30256 | 0.68371 | -0.54854 | 0.011712 | 0.076615 | 0.40641 | -1.299 |
| DAG160/181 | 0.63433 | 0.73088 | 0.88911 | -0.16956 | 0.90635 | 0.9486 | 1.1647 | 0.21997 | 0.19007 | 0.29889 | 0.75639 | -0.40279 |
| TAG566-FA226 | 0.63791 | 0.7337 | 1.2286 | 0.29702 | 0.81098 | 0.89922 | 1.2787 | 0.35469 | 0.1531 | 0.2549 | 0.61936 | -0.69115 |
| PC160/183 | 0.6433 | 0.7386 | 0.91889 | -0.12204 | 0.005301 | 0.088492 | 0.55668 | -0.84508 | 0.003761 | 0.068007 | 0.49074 | -1.027 |
| PC180/182 | 0.64781 | 0.74247 | 0.95539 | -0.06584 | 0.68985 | 0.82101 | 0.96031 | -0.05843 | 0.098435 | 0.18737 | 0.86834 | -0.20367 |
| FFA161 | 0.65449 | 0.74881 | 0.95337 | -0.06889 | 0.76867 | 0.87606 | 0.95335 | -0.06892 | 0.30384 | 0.4187 | 1.139 | 0.18772 |
| FFA241 | 0.65979 | 0.75355 | 1.0386 | 0.054708 | 0.65546 | 0.79461 | 0.96401 | -0.05289 | 0.0141 | 0.077008 | 0.82621 | -0.27542 |
| TAG483-FA141 | 0.66232 | 0.75511 | 0.95451 | -0.06717 | 0.45365 | 0.63375 | 0.78511 | -0.34904 | 0.52415 | 0.6414 | 1.1367 | 0.18491 |
| PEP-181/226 | 0.66853 | 0.76086 | 1.1016 | 0.13961 | 0.36708 | 0.57474 | 1.2136 | 0.27925 | 0.39168 | 0.51512 | 1.1268 | 0.17226 |
| PEP-160/180 | 0.67705 | 0.76922 | 0.96773 | -0.04732 | 0.25028 | 0.4721 | 1.1755 | 0.23332 | 0.36174 | 0.48456 | 1.1392 | 0.188 |
| FFA201 | 0.67982 | 0.77048 | 0.96176 | -0.05625 | 0.96418 | 0.98718 | 0.99674 | -0.00471 | 0.23133 | 0.3454 | 1.1234 | 0.16783 |
| FFA141 | 0.68053 | 0.77048 | 1.1525 | 0.20476 | 0.45976 | 0.63455 | 1.1474 | 0.19832 | 0.089337 | 0.17731 | 1.4956 | 0.58074 |
| PC200/226 | 0.68542 | 0.77466 | 0.99129 | -0.01262 | 0.2099 | 0.4297 | 0.85069 | -0.23329 | 0.041006 | 0.12189 | 0.78214 | -0.35449 |
| PEP-140/181 | 0.69178 | 0.77965 | 1.0901 | 0.1245 | 0.73714 | 0.85388 | 1.1994 | 0.26236 | 0.41187 | 0.53518 | 0.85913 | -0.21906 |
| LPE170 | 0.69332 | 0.77965 | 0.90774 | -0.13964 | 0.27004 | 0.48832 | 0.77103 | -0.37514 | 0.025407 | 0.10291 | 0.5912 | -0.75828 |
| FFA240 | 0.69342 | 0.77965 | 1.0764 | 0.1062 | 0.87515 | 0.93651 | 1.0522 | 0.073397 | 0.499 | 0.61876 | 1.1888 | 0.24949 |
| Cholesterol | 0.69677 | 0.78207 | 1.0239 | 0.034116 | 0.000428 | 0.030959 | 0.5462 | -0.87249 | 0.001119 | 0.055371 | 0.66525 | -0.58803 |
| FFA160 | 0.7062 | 0.79129 | 1.0249 | 0.035434 | 0.25523 | 0.4721 | 1.0534 | 0.075074 | 0.054818 | 0.13832 | 0.91607 | -0.12647 |
| PEP-181/204 | 0.71239 | 0.79567 | 0.97089 | -0.04262 | 0.19531 | 0.41237 | 1.1874 | 0.24785 | 0.53087 | 0.64488 | 1.0612 | 0.085751 |
| PE180/224 | 0.71256 | 0.79567 | 1.0667 | 0.09319 | 0.09142 | 0.29172 | 0.77441 | -0.36882 | 0.069544 | 0.15399 | 0.71823 | -0.47748 |
| CE183 | 0.71853 | 0.80097 | 1.0297 | 0.042207 | 0.004217 | 0.084684 | 0.49221 | -1.0227 | 0.015175 | 0.079031 | 0.55636 | -0.8459 |
| PEP-182/204 | 0.724 | 0.80568 | 1.0718 | 0.10006 | 0.21716 | 0.43952 | 1.2047 | 0.26873 | 0.8157 | 0.87917 | 1.0228 | 0.032561 |
| CER240 | 0.72831 | 0.80909 | 1.0873 | 0.12081 | 0.47656 | 0.64504 | 1.0912 | 0.12592 | 0.87439 | 0.91628 | 1.0198 | 0.028342 |
| PEP-160/183 | 0.73031 | 0.80994 | 0.95886 | -0.06061 | 0.23428 | 0.465 | 0.81493 | -0.29526 | 0.001033 | 0.055371 | 0.42303 | -1.2412 |
| PA181/182 | 0.73588 | 0.81354 | 0.9582 | -0.06161 | 0.005585 | 0.088683 | 1.4784 | 0.56404 | 0.093671 | 0.1812 | 1.2487 | 0.32042 |
| FFA181 | 0.73606 | 0.81354 | 1.0162 | 0.023211 | 0.63659 | 0.77979 | 1.0302 | 0.042935 | 0.057583 | 0.14008 | 0.83525 | -0.25971 |

| | | | | | | | | | | | | |
|---|---|---|---|---|---|---|---|---|---|---|---|---|
| PE180/182 | 0.74103 | 0.81764 | 0.96698 | -0.04845 | 0.30265 | 0.51578 | 0.8958 | -0.15875 | 0.012797 | 0.076615 | 0.711 | -0.49208 |
| PC160/201 | 0.74473 | 0.82002 | 0.9717 | -0.04142 | 0.043257 | 0.20555 | 0.8356 | -0.25911 | 0.94663 | 0.9629 | 1.0076 | 0.010917 |
| PEP-180/224 | 0.7457 | 0.82002 | 1.0718 | 0.099997 | 0.87114 | 0.93651 | 1.0222 | 0.031738 | 0.75809 | 0.83133 | 0.95011 | -0.07384 |
| PS160/161_2 | 0.74989 | 0.82324 | 1.0548 | 0.076976 | 0.90103 | 0.94765 | 1.0216 | 0.030879 | 0.78988 | 0.85845 | 1.0461 | 0.065002 |
| PE180/181 | 0.75533 | 0.82781 | 1.0344 | 0.048852 | 0.25578 | 0.4721 | 1.2296 | 0.29824 | 0.64797 | 0.74792 | 0.93169 | -0.10208 |
| FFA170 | 0.75811 | 0.82946 | 0.96933 | -0.04494 | 0.72919 | 0.85072 | 1.0199 | 0.028366 | 0.75485 | 0.83133 | 0.96649 | -0.04918 |
| FFA182 | 0.78119 | 0.85328 | 1.0154 | 0.022068 | 0.50432 | 0.66877 | 1.0611 | 0.085615 | 0.14081 | 0.23748 | 0.86394 | -0.211 |
| PC150/161 | 0.78896 | 0.85945 | 1.072 | 0.10029 | 0.86571 | 0.93462 | 1.0588 | 0.08243 | 0.56286 | 0.67417 | 0.9238 | -0.11434 |
| PC180/205 | 0.78948 | 0.85945 | 0.96839 | -0.04633 | 0.43665 | 0.61795 | 0.90021 | -0.15167 | 0.010853 | 0.076615 | 0.6545 | -0.61154 |
| FFA180 | 0.79152 | 0.86023 | 0.98071 | -0.02811 | 0.89688 | 0.94765 | 1.002 | 0.002817 | 0.2005 | 0.31077 | 0.92418 | -0.11375 |
| TAG524-FA204 | 0.79722 | 0.86498 | 1.0909 | 0.12547 | 0.51363 | 0.67414 | 0.96421 | -0.05258 | 0.29941 | 0.41384 | 0.80693 | -0.30949 |
| LPE182 | 0.802 | 0.86872 | 1.1271 | 0.17262 | 0.29801 | 0.5134 | 1.5533 | 0.6353 | 0.4407 | 0.55925 | 0.84691 | -0.23973 |
| TAG480-FA160 | 0.80443 | 0.86991 | 1.0178 | 0.025499 | 0.25258 | 0.4721 | 0.6509 | -0.61949 | 0.35821 | 0.48081 | 0.73669 | -0.44087 |
| CER200 | 0.8083 | 0.87164 | 1.1495 | 0.20102 | 0.66251 | 0.79783 | 0.82382 | -0.2796 | 0.67381 | 0.76672 | 0.82995 | -0.26891 |
| CER220 | 0.80871 | 0.87164 | 1.1136 | 0.15519 | 0.3637 | 0.57468 | 0.83511 | -0.25996 | 0.33885 | 0.46052 | 0.83108 | -0.26694 |
| PEO-160/160 | 0.81642 | 0.8785 | 0.92912 | -0.10606 | 0.20376 | 0.42245 | 0.74104 | -0.43237 | 0.054655 | 0.13832 | 0.62872 | -0.66951 |
| TAG564-FA204 | 0.8216 | 0.88261 | 0.9585 | -0.06115 | 0.85002 | 0.92381 | 1.2363 | 0.30608 | 0.23848 | 0.35124 | 0.77514 | -0.36748 |
| PS160/181 | 0.82505 | 0.88485 | 1.0381 | 0.053882 | 0.90208 | 0.94765 | 1.0313 | 0.044509 | 0.21704 | 0.32935 | 1.1494 | 0.20093 |
| LPE220 | 0.82696 | 0.88545 | 1.0401 | 0.056764 | 0.003854 | 0.084684 | 1.7523 | 0.80923 | 0.57539 | 0.68604 | 1.1112 | 0.15206 |
| TAG525-FA205 | 0.83009 | 0.88598 | 1.0129 | 0.018488 | 0.2423 | 0.46994 | 0.76978 | -0.37748 | 0.067143 | 0.15036 | 0.56668 | -0.81939 |
| TAG544-FA224 | 0.83018 | 0.88598 | 0.98901 | -0.01594 | 0.98656 | 0.99607 | 1.2908 | 0.36821 | 0.50216 | 0.62032 | 0.82113 | -0.28431 |
| PEO-180/161 | 0.83502 | 0.88968 | 0.96948 | -0.04471 | 0.60911 | 0.75818 | 1.0744 | 0.10349 | 0.45829 | 0.57596 | 0.84555 | -0.24203 |
| FFA221 | 0.83854 | 0.89142 | 0.95872 | -0.06083 | 0.30808 | 0.52094 | 1.3123 | 0.39211 | 0.48266 | 0.60309 | 0.89512 | -0.15984 |
| PE170/224 | 0.83939 | 0.89142 | 0.98183 | -0.02646 | 0.16614 | 0.38229 | 0.81966 | -0.2869 | 0.008506 | 0.075886 | 0.64942 | -0.62277 |
| PC180/180 | 0.84821 | 0.89933 | 0.94969 | -0.07446 | 0.8726 | 0.93651 | 0.99987 | -0.00018 | 0.98997 | 0.99302 | 0.97471 | -0.03696 |
| CE181 | 0.85124 | 0.90107 | 1.0146 | 0.020966 | 0.00122 | 0.044105 | 0.63654 | -0.65167 | 0.026238 | 0.10415 | 0.76457 | -0.38727 |
| PA220/226 | 0.85414 | 0.90267 | 0.99067 | -0.01353 | 0.044932 | 0.21196 | 1.3415 | 0.42388 | 0.6588 | 0.75773 | 0.93954 | -0.08998 |
| PEO-160/181 | 0.85637 | 0.90356 | 1.0774 | 0.10757 | 0.99584 | 0.99737 | 1.0103 | 0.014811 | 0.24948 | 0.36253 | 0.83948 | -0.25244 |
| TAG482-FA141 | 0.86822 | 0.91379 | 1.1878 | 0.24833 | 0.86297 | 0.93321 | 1.0531 | 0.074606 | 0.67486 | 0.76672 | 1.882 | 0.91226 |
| FFA150 | 0.86948 | 0.91379 | 0.99341 | -0.00954 | 0.23061 | 0.4591 | 1.1556 | 0.20866 | 0.51631 | 0.63419 | 0.90421 | -0.14527 |
| LPC160 | 0.87027 | 0.91379 | 1.0107 | 0.015371 | 0.093746 | 0.29172 | 1.1663 | 0.22193 | 0.91748 | 0.94357 | 0.98438 | -0.02271 |
| TAG568-FA226 | 0.87284 | 0.915 | 1.0144 | 0.020562 | 0.49511 | 0.66184 | 1.4615 | 0.54747 | 0.19381 | 0.30403 | 0.64185 | -0.63969 |
| LPE201 | 0.88378 | 0.92498 | 1.0205 | 0.029298 | 0.093949 | 0.29172 | 1.6892 | 0.75635 | 0.86228 | 0.90541 | 0.99192 | -0.01171 |

| | 6.5 Gy | | | | | | | | | | | |
|---|---|---|---|---|---|---|---|---|---|---|---|---|
| | SD 1 vs. Pre | | | | SD 2 vs. Pre | | | | SD 6 vs. Pre | | | |
| **Lipid Name** | **p-value** | **FDR** | **FC** | **LOG2FC** | **p-value** | **FDR** | **FC** | **LOG2FC** | **p-value** | **FDR** | **FC** | **LOG2FC** |
| PE160/224 | 0.88844 | 0.92837 | 1.0015 | 0.002126 | 0.035223 | 0.19432 | 0.58466 | -0.77434 | 0.23121 | 0.3454 | 0.75079 | -0.41352 |
| PC160/203 | 0.89696 | 0.93496 | 0.95185 | -0.07119 | 0.47342 | 0.64476 | 0.86149 | -0.2151 | 0.13453 | 0.23317 | 0.73953 | -0.43532 |
| FFA222 | 0.89762 | 0.93496 | 1.0219 | 0.031202 | 0.40386 | 0.5989 | 0.93882 | -0.09107 | 0.29812 | 0.41381 | 0.92513 | -0.11227 |
| HCER201 | 0.89968 | 0.93561 | 0.975 | -0.03653 | 0.50441 | 0.66877 | 0.9187 | -0.12234 | 0.032342 | 0.11231 | 0.75893 | -0.39797 |
| TAG544-FA204 | 0.90454 | 0.93916 | 1.0592 | 0.082962 | 0.68694 | 0.82055 | 1.0342 | 0.048538 | 0.1392 | 0.23681 | 0.68688 | -0.54187 |
| TAG567-FA226 | 0.91298 | 0.94641 | 0.9626 | -0.05499 | 0.80218 | 0.89432 | 1.2942 | 0.37208 | 0.27848 | 0.39411 | 0.71084 | -0.49241 |
| PC200/225 | 0.91825 | 0.95037 | 1.0424 | 0.059873 | 0.60901 | 0.75818 | 0.94632 | -0.0796 | 0.061333 | 0.14109 | 0.7518 | -0.41157 |
| TAG503-FA141 | 0.92164 | 0.95236 | 1.0402 | 0.056825 | 0.32736 | 0.54504 | 1.8834 | 0.91336 | 0.96403 | 0.97602 | 1.2699 | 0.34468 |
| TAG565-FA225 | 0.92413 | 0.95342 | 0.88393 | -0.178 | 0.89795 | 0.94765 | 1.1269 | 0.1723 | 0.12204 | 0.21794 | 0.56381 | -0.82671 |
| PC160/120 | 0.93081 | 0.95588 | 0.93632 | -0.09493 | 0.11919 | 0.33301 | 0.69017 | -0.53498 | 0.17109 | 0.27777 | 2.307 | 1.206 |
| FFA200 | 0.93202 | 0.95588 | 0.99051 | -0.01376 | 0.44161 | 0.62141 | 1.2573 | 0.33036 | 0.26057 | 0.37447 | 1.295 | 0.37294 |
| PC180/224 | 0.93237 | 0.95588 | 1.0088 | 0.012695 | 0.64152 | 0.78318 | 0.94355 | -0.08384 | 0.33573 | 0.45724 | 0.87433 | -0.19375 |
| PC200/141 | 0.93239 | 0.95588 | 0.99058 | -0.01365 | 0.17898 | 0.40039 | 1.1486 | 0.19985 | 0.75605 | 0.83133 | 0.95627 | -0.06451 |
| PS161/180_2 | 0.93729 | 0.95796 | 1.0251 | 0.035704 | 0.82046 | 0.90222 | 0.97622 | -0.03472 | 0.89075 | 0.92338 | 1.0395 | 0.055957 |
| LPE161 | 0.93795 | 0.95796 | 0.89152 | -0.16567 | 0.17769 | 0.39888 | 1.3564 | 0.4398 | 0.79472 | 0.86227 | 0.85063 | -0.23339 |
| PC200/204 | 0.93992 | 0.95796 | 1.0379 | 0.05366 | 0.18813 | 0.40645 | 0.84422 | -0.2443 | 0.021133 | 0.094255 | 0.76123 | -0.39359 |
| SMd181/183 | 0.94084 | 0.95796 | 0.99539 | -0.00667 | 0.036087 | 0.19538 | 0.82123 | -0.28414 | 0.013192 | 0.07668 | 0.75395 | -0.40746 |
| PC180/200 | 0.94178 | 0.95796 | 0.97589 | -0.0352 | 0.53968 | 0.69709 | 1.0739 | 0.10285 | 0.96637 | 0.97687 | 0.98235 | -0.02569 |
| PE160/181 | 0.95414 | 0.96902 | 0.98977 | -0.01484 | 0.3179 | 0.53108 | 0.83825 | -0.25455 | 0.25637 | 0.36992 | 0.80723 | -0.30895 |
| PEP-160/201 | 0.95624 | 0.96964 | 0.99447 | -0.008 | 0.74399 | 0.85724 | 0.94657 | -0.07922 | 0.88712 | 0.92255 | 1.0237 | 0.033759 |
| PC180/120 | 0.96715 | 0.97918 | 0.95695 | -0.06349 | 0.14829 | 0.3662 | 0.73109 | -0.45189 | 0.02839 | 0.10827 | 0.57703 | -0.79327 |
| PEP-180/226 | 0.9702 | 0.98075 | 1.0551 | 0.077325 | 0.39972 | 0.5989 | 1.1742 | 0.23171 | 0.88443 | 0.92255 | 1.0332 | 0.047181 |
| DAG200/200 | 0.97535 | 0.98234 | 1.0302 | 0.042946 | 0.19684 | 0.41237 | 0.76918 | -0.37861 | 0.5944 | 0.701 | 1.3282 | 0.40948 |
| PEP-160/161 | 0.97595 | 0.98234 | 1.0047 | 0.006704 | 0.8451 | 0.92381 | 0.96607 | -0.0498 | 0.71163 | 0.79463 | 1.0321 | 0.045558 |
| CER201 | 0.9763 | 0.98234 | 1.0272 | 0.038697 | 0.4262 | 0.60777 | 0.86523 | -0.20885 | 0.28905 | 0.4045 | 0.82015 | -0.28603 |
| PC170/204 | 0.98294 | 0.98749 | 0.96375 | -0.05327 | 0.18985 | 0.4079 | 0.77496 | -0.36781 | 0.086584 | 0.17397 | 0.72445 | -0.46503 |
| PE180/205 | 0.98907 | 0.99212 | 0.9641 | -0.05275 | 0.098884 | 0.29948 | 0.72771 | -0.45856 | 0.000708 | 0.055371 | 0.43466 | -1.202 |
| FFA224 | 0.99788 | 0.99935 | 1.0248 | 0.035383 | 0.3609 | 0.57468 | 0.91343 | -0.13064 | 0.98833 | 0.9929 | 1.0121 | 0.017415 |
| PC140/224 | 0.99935 | 0.99935 | 0.97785 | -0.03232 | 0.16902 | 0.3834 | 0.72936 | -0.45529 | 0.40166 | 0.524 | 1.4545 | 0.54053 |

| | | | | | | | | | | | | | | | |
|---|---|---|---|---|---|---|---|---|---|---|---|---|---|---|---|
| PI160/201 | 0.017004 | 0.090926 | 0.44375 | ↓ | -1.172 | 0.51339 | 0.74163 | 0.80192 | ↓ | -0.318 | 0.010089 | 0.038366 | 0.43559 | ↓ | -1.199 |
| LPE160 | 0.001628 | 0.070016 | 2.1085 | ↑ | 1.0762 | 0.057737 | 0.53695 | 1.4914 | ↑ | 0.5767 | 0.86188 | 0.88499 | 1.0318 | ↑ | 0.0452 |
| PC182/203 | 0.003662 | 0.070016 | 0.43345 | ↓ | -1.206 | 0.39791 | 0.70776 | 0.79781 | ↓ | -0.326 | 0.044355 | 0.088045 | 0.54607 | ↓ | -0.873 |
| PG180/182_2 | 0.002756 | 0.070016 | 0.52622 | ↓ | -0.926 | 0.081136 | 0.5914 | 0.6732 | ↓ | -0.571 | 0.005384 | 0.032569 | 0.52123 | ↓ | -0.94 |
| PI160/204 | 0.037084 | 0.11551 | 0.59533 | ↓ | -0.748 | 0.44664 | 0.72873 | 0.83102 | ↓ | -0.267 | 0.74252 | 0.79766 | 0.89897 | ↓ | -0.154 |
| PG180/182 | 0.00396 | 0.070016 | 0.53853 | ↓ | -0.893 | 0.082758 | 0.5914 | 0.67584 | ↓ | -0.565 | 0.004886 | 0.032129 | 0.51577 | ↓ | -0.955 |
| LPC202 | 0.079234 | 0.1798 | 0.71969 | ↓ | -0.475 | 0.57258 | 0.77177 | 0.89585 | ↓ | -0.159 | 0.027991 | 0.066263 | 0.60657 | ↓ | -0.721 |
| TAG547-FA183 | 0.006271 | 0.070016 | 0.15394 | ↓ | -2.7 | 0.06885 | 0.55914 | 0.41609 | ↓ | -1.265 | 0.001687 | 0.023486 | 0.1112 | ↓ | -3.169 |
| PI160/182 | 0.04477 | 0.12839 | 0.55327 | ↓ | -0.854 | 0.33189 | 0.64689 | 0.77289 | ↓ | -0.372 | 0.017847 | 0.047616 | 0.54624 | ↓ | -0.872 |
| TAG547-FA182 | 0.011688 | 0.082752 | 0.18112 | ↓ | -2.465 | 0.090663 | 0.5914 | 0.47782 | ↓ | -1.066 | 0.007994 | 0.037908 | 0.16143 | ↓ | -2.631 |
| PS161/180 | 0.002386 | 0.070016 | 0.4525 | ↓ | -1.144 | 0.014225 | 0.30735 | 0.57573 | ↓ | -0.797 | 0.020957 | 0.052273 | 0.53068 | ↓ | -0.914 |
| TAG526-FA183 | 0.00877 | 0.077743 | 0.16959 | ↓ | -2.56 | 0.10501 | 0.5914 | 0.58269 | ↓ | -0.779 | 0.001202 | 0.023486 | 0.10737 | ↓ | -3.219 |
| TAG483-FA183 | 0.001845 | 0.070016 | 0.13458 | ↓ | -2.894 | 0.063431 | 0.54498 | 0.54246 | ↓ | -0.882 | 0.001163 | 0.023486 | 0.13057 | ↓ | -2.937 |
| LPE226 | 0.014089 | 0.087814 | 1.7722 | ↑ | 0.8255 | 0.00274 | 0.28651 | 1.9562 | ↑ | 0.9681 | 0.1876 | 0.26783 | 1.2785 | ↑ | 0.3545 |
| PG160/161 | 0.069214 | 0.16335 | 0.64092 | ↓ | -0.642 | 0.21364 | 0.5914 | 0.7825 | ↓ | -0.354 | 0.098936 | 0.15595 | 0.67204 | ↓ | -0.573 |
| TAG525-FA182 | 0.002926 | 0.070016 | 0.20493 | ↓ | -2.287 | 0.10791 | 0.5914 | 0.53865 | ↓ | -0.893 | 0.00239 | 0.027585 | 0.19006 | ↓ | -2.395 |
| PE181/203 | 0.020908 | 0.09098 | 0.50323 | ↓ | -0.991 | 0.25564 | 0.59421 | 0.72104 | ↓ | -0.472 | 0.009502 | 0.038185 | 0.39042 | ↓ | -1.357 |
| PI160/180 | 0.05118 | 0.13671 | 0.56628 | ↓ | -0.82 | 0.66605 | 0.83459 | 0.8666 | ↓ | -0.207 | 0.003883 | 0.031552 | 0.481 | ↓ | -1.056 |
| PC160/226 | 0.41489 | 0.58084 | 1.1272 | ↑ | 0.1728 | 0.39678 | 0.70768 | 1.1136 | ↑ | 0.1552 | 0.51179 | 0.58969 | 1.1373 | ↑ | 0.1856 |
| PI160/160 | 0.064173 | 0.15579 | 0.54446 | ↓ | -0.877 | 0.24224 | 0.5914 | 0.69971 | ↓ | -0.515 | 0.054487 | 0.10281 | 0.56619 | ↓ | -0.821 |
| PI181/203_2 | 8.88E-05 | 0.057788 | 0.41161 | ↓ | -1.281 | 0.008058 | 0.28651 | 0.72594 | ↓ | -0.462 | 0.012263 | 0.04094 | 0.61276 | ↓ | -0.707 |
| TAG542-FA200 | 0.054718 | 0.13969 | 0.38608 | ↓ | -1.373 | 0.17407 | 0.5914 | 0.53414 | ↓ | -0.905 | 0.007672 | 0.037551 | 0.24961 | ↓ | -2.002 |
| TAG546-FA183 | 0.004317 | 0.070016 | 0.16932 | ↓ | -2.562 | 0.070657 | 0.56094 | 0.39385 | ↓ | -1.344 | 0.001775 | 0.023486 | 0.13071 | ↓ | -2.936 |
| TAG504-FA182 | 0.006216 | 0.070016 | 0.16338 | ↓ | -2.614 | 0.1367 | 0.5914 | 0.46388 | ↓ | -1.108 | 0.011292 | 0.039348 | 0.19022 | ↓ | -2.394 |
| PC181/202 | 0.005853 | 0.070016 | 0.56831 | ↓ | -0.815 | 0.045477 | 0.5194 | 0.68576 | ↓ | -0.544 | 0.023771 | 0.057959 | 0.55381 | ↓ | -0.853 |
| PC182/183 | 0.049788 | 0.13642 | 0.48347 | ↓ | -1.049 | 0.018451 | 0.33515 | 0.43985 | ↓ | -1.185 | 0.030345 | 0.07106 | 0.4197 | ↓ | -1.253 |
| PI160/182_2 | 0.047739 | 0.13281 | 0.52911 | ↓ | -0.918 | 0.4452 | 0.72846 | 0.79843 | ↓ | -0.325 | 0.031985 | 0.072804 | 0.56141 | ↓ | -0.833 |
| PI181/203 | 0.000647 | 0.070016 | 0.47286 | ↓ | -1.081 | 0.38055 | 0.68816 | 0.87977 | ↓ | -0.185 | 0.047404 | 0.092952 | 0.66521 | ↓ | -0.588 |
| PG160/161_2 | 0.2028 | 0.35206 | 0.65233 | ↓ | -0.616 | 0.82603 | 0.91298 | 0.94579 | ↓ | -0.08 | 0.49528 | 0.57473 | 0.78967 | ↓ | -0.341 |
| LPC200 | 0.011713 | 0.082752 | 0.58389 | ↓ | -0.776 | 0.66842 | 0.83459 | 1.08 | ↑ | 0.1111 | 0.0104 | 0.038366 | 0.51018 | ↓ | -0.971 |
| TAG504-FA140 | 0.011822 | 0.082752 | 0.17974 | ↓ | -2.476 | 0.20279 | 0.5914 | 0.57701 | ↓ | -0.793 | 0.012498 | 0.04151 | 0.18196 | ↓ | -2.458 |
| TAG525-FA183 | 0.020139 | 0.09098 | 0.22728 | ↓ | -2.137 | 0.1764 | 0.5914 | 0.60013 | ↓ | -0.737 | 0.008972 | 0.037926 | 0.17657 | ↓ | -2.502 |
| TAG526-FA182 | 0.007277 | 0.075267 | 0.18394 | ↓ | -2.443 | 0.10678 | 0.5914 | 0.4662 | ↓ | -1.101 | 0.00166 | 0.023486 | 0.12529 | ↓ | -2.997 |

| | | | | | | | | | | | | | | | |
|---|---|---|---|---|---|---|---|---|---|---|---|---|---|---|---|
| TAG525-FA225 | 0.002246 | 0.070016 | 0.24496 | ↓ | -2.029 | 0.10624 | 0.5914 | 0.65563 | ↓ | -0.609 | 0.011661 | 0.040165 | 0.30844 | ↓ | -1.697 |
| TAG504-FA183 | 0.004379 | 0.070016 | 0.14115 | ↓ | -2.825 | 0.17896 | 0.5914 | 0.61492 | ↓ | -0.702 | 0.004715 | 0.032129 | 0.14546 | ↓ | -2.781 |
| TAG526-FA161 | 0.00151 | 0.070016 | 0.12147 | ↓ | -3.041 | 0.052447 | 0.53349 | 0.42167 | ↓ | -1.246 | 0.001532 | 0.023486 | 0.12016 | ↓ | -3.057 |
| CE204 | 0.000302 | 0.065509 | 2.0161 | ↑ | 1.0115 | 0.87159 | 0.93786 | 0.98985 | ↓ | -0.015 | 0.008514 | 0.037908 | 1.8099 | ↑ | 0.8559 |
| TAG546-FA181 | 0.014351 | 0.088134 | 0.23004 | ↓ | -2.12 | 0.19696 | 0.5914 | 0.58686 | ↓ | -0.769 | 0.007574 | 0.037551 | 0.18834 | ↓ | -2.409 |
| PI160/204_2 | 0.042255 | 0.12471 | 0.60342 | ↓ | -0.729 | 0.36583 | 0.67841 | 0.80193 | ↓ | -0.318 | 0.81329 | 0.85001 | 0.91092 | ↓ | -0.135 |
| TAG503-FA140 | 0.011732 | 0.082752 | 0.19335 | ↓ | -2.371 | 0.45228 | 0.72879 | 0.76569 | ↓ | -0.385 | 0.01904 | 0.049067 | 0.22234 | ↓ | -2.169 |
| PI161/181 | 0.000204 | 0.065509 | 0.42157 | ↓ | -1.246 | 0.011443 | 0.28651 | 0.70911 | ↓ | -0.496 | 0.009479 | 0.038185 | 0.60039 | ↓ | -0.736 |
| LPC220 | 0.021382 | 0.09098 | 0.6033 | ↓ | -0.729 | 0.92368 | 0.96674 | 1.0693 | ↑ | 0.0967 | 0.010726 | 0.038366 | 0.48524 | ↓ | -1.043 |
| TAG548-FA182 | 0.022637 | 0.091642 | 0.17209 | ↓ | -2.539 | 0.10521 | 0.5914 | 0.44709 | ↓ | -1.161 | 0.009947 | 0.038366 | 0.12893 | ↓ | -2.955 |
| PC182/202 | 0.051127 | 0.13671 | 0.65019 | ↓ | -0.621 | 0.19526 | 0.5914 | 0.75583 | ↓ | -0.404 | 0.038622 | 0.080546 | 0.55885 | ↓ | -0.839 |
| PI182/182 | 0.051239 | 0.13671 | 0.48108 | ↓ | -1.056 | 0.13511 | 0.5914 | 0.61152 | ↓ | -0.71 | 0.030889 | 0.071649 | 0.46315 | ↓ | -1.111 |
| TAG546-FA182 | 0.013363 | 0.087814 | 0.29183 | ↓ | -1.777 | 0.099767 | 0.5914 | 0.49446 | ↓ | -1.016 | 0.003852 | 0.031552 | 0.21551 | ↓ | -2.214 |
| PI181/182_2 | 0.017916 | 0.09098 | 0.52607 | ↓ | -0.927 | 0.23869 | 0.5914 | 1.2806 | ↑ | 0.3568 | 0.018793 | 0.048741 | 0.54834 | ↓ | -0.867 |
| PI160/180_2 | 0.05349 | 0.13806 | 0.56518 | ↓ | -0.823 | 0.27083 | 0.59421 | 0.75823 | ↓ | -0.399 | 0.004151 | 0.032129 | 0.48862 | ↓ | -1.033 |
| TAG525-FA160 | 0.020164 | 0.09098 | 0.27434 | ↓ | -1.866 | 0.11191 | 0.5914 | 0.60198 | ↓ | -0.732 | 0.01006 | 0.038366 | 0.22365 | ↓ | -2.161 |
| TAG525-FA161 | 0.001839 | 0.070016 | 0.1824 | ↓ | -2.455 | 0.10419 | 0.5914 | 0.46796 | ↓ | -1.096 | 0.002161 | 0.026538 | 0.16714 | ↓ | -2.581 |
| TAG505-FA161 | 0.001382 | 0.070016 | 0.16758 | ↓ | -2.577 | 0.046638 | 0.52329 | 0.42472 | ↓ | -1.235 | 0.000415 | 0.023486 | 0.096674 | ↓ | -3.371 |
| PG181/182 | 0.017641 | 0.09098 | 0.48015 | ↓ | -1.058 | 0.2733 | 0.59421 | 0.72997 | ↓ | -0.454 | 0.05492 | 0.10311 | 0.51295 | ↓ | -0.963 |
| TAG563-FA200 | 0.021035 | 0.09098 | 0.37916 | ↓ | -1.399 | 0.64739 | 0.828 | 0.78013 | ↓ | -0.358 | 0.000919 | 0.023486 | 0.21094 | ↓ | -2.245 |
| PC160/204 | 0.036617 | 0.11516 | 1.3025 | ↑ | 0.3813 | 0.12103 | 0.5914 | 1.2051 | ↑ | 0.2691 | 0.18614 | 0.26691 | 1.1537 | ↑ | 0.2063 |
| PC182/182 | 0.030108 | 0.10164 | 0.5562 | ↓ | -0.846 | 0.93567 | 0.9746 | 0.98211 | ↓ | -0.026 | 0.077987 | 0.13256 | 0.60667 | ↓ | -0.721 |
| PI160/181 | 0.018366 | 0.09098 | 0.54967 | ↓ | -0.863 | 0.55905 | 0.76614 | 1.0754 | ↑ | 0.1049 | 0.001804 | 0.023486 | 0.52017 | ↓ | -0.943 |
| TAG482-FA182 | 0.007515 | 0.075267 | 0.22696 | ↓ | -2.14 | 0.13399 | 0.5914 | 0.61804 | ↓ | -0.694 | 0.013366 | 0.042154 | 0.25261 | ↓ | -1.985 |
| PG181/181 | 0.003811 | 0.070016 | 0.52426 | ↓ | -0.932 | 0.76125 | 0.89227 | 1.268 | ↑ | 0.3426 | 0.04187 | 0.085452 | 0.57376 | ↓ | -0.801 |
| TAG566-FA183 | 0.013581 | 0.087814 | 0.28351 | ↓ | -1.819 | 0.15539 | 0.5914 | 0.58101 | ↓ | -0.783 | 0.008735 | 0.037908 | 0.21999 | ↓ | -2.185 |
| PC182/161 | 0.011124 | 0.082341 | 0.63409 | ↓ | -0.657 | 0.014538 | 0.30735 | 0.74986 | ↓ | -0.415 | 0.047378 | 0.092952 | 0.64364 | ↓ | -0.636 |
| PI181/182 | 0.015946 | 0.090926 | 0.5248 | ↓ | -0.93 | 0.24474 | 0.5914 | 1.2728 | ↑ | 0.3481 | 0.013662 | 0.042554 | 0.5499 | ↓ | -0.863 |
| TAG524-FA221 | 0.004659 | 0.070016 | 0.31441 | ↓ | -1.669 | 0.21357 | 0.5914 | 0.69611 | ↓ | -0.523 | 0.008805 | 0.037908 | 0.34676 | ↓ | -1.528 |
| PI160/181_2 | 0.020992 | 0.09098 | 0.57266 | ↓ | -0.804 | 0.45779 | 0.73251 | 1.1048 | ↑ | 0.1438 | 0.001624 | 0.023486 | 0.50443 | ↓ | -0.987 |
| TAG541-FA200 | 0.041683 | 0.12391 | 0.35212 | ↓ | -1.506 | 0.3519 | 0.66789 | 0.58727 | ↓ | -0.768 | 0.00376 | 0.031552 | 0.20659 | ↓ | -2.275 |
| TAG548-FA183 | 0.005024 | 0.070016 | 0.13647 | ↓ | -2.873 | 0.031816 | 0.4227 | 0.37973 | ↓ | -1.397 | 0.000554 | 0.023486 | 0.075374 | ↓ | -3.73 |
| CE226 | 0.005766 | 0.070016 | 2.0884 | ↑ | 1.0624 | 0.99557 | 0.99929 | 1.0972 | ↑ | 0.1338 | 0.004339 | 0.032129 | 2.4015 | ↑ | 1.2639 |

| | | | | | | | | | | | | | | | |
|---|---|---|---|---|---|---|---|---|---|---|---|---|---|---|---|
| TAG503-FA182 | 0.002229 | 0.070016 | 0.21033 | ↓ | -2.249 | 0.19732 | 0.5914 | 0.66196 | ↓ | -0.595 | 0.0061 | 0.034 | 0.2436 | ↓ | -2.037 |
| LPC183 | 0.21895 | 0.3751 | 0.74272 | ↓ | -0.429 | 0.18278 | 0.5914 | 0.72624 | ↓ | -0.461 | 0.00796 | 0.037908 | 0.45316 | ↓ | -1.142 |
| TAG525-FA181 | 0.002463 | 0.070016 | 0.19964 | ↓ | -2.325 | 0.16257 | 0.5914 | 0.59024 | ↓ | -0.761 | 0.00169 | 0.023486 | 0.1633 | ↓ | -2.614 |
| PC140/226 | 0.034849 | 0.11176 | 0.66713 | ↓ | -0.584 | 0.49799 | 0.73497 | 0.88611 | ↓ | -0.174 | 0.038182 | 0.080296 | 0.65077 | ↓ | -0.62 |
| LPC150 | 0.16814 | 0.30678 | 0.75623 | ↓ | -0.403 | 0.62281 | 0.8109 | 0.90744 | ↓ | -0.14 | 0.004627 | 0.032129 | 0.50824 | ↓ | -0.976 |
| PS161/181 | 0.96239 | 0.97588 | 0.92711 | ↓ | -0.109 | 0.77352 | 0.89253 | 1.2533 | ↑ | 0.3258 | 0.9185 | 0.93575 | 0.96953 | ↓ | -0.045 |
| PC180/201 | 0.31461 | 0.48305 | 0.85132 | ↓ | -0.232 | 0.19691 | 0.5914 | 0.79545 | ↓ | -0.33 | 0.057888 | 0.10543 | 0.70269 | ↓ | -0.509 |
| PI181/181 | 0.006027 | 0.070016 | 0.52679 | ↓ | -0.925 | 0.016496 | 0.33253 | 2.2894 | ↑ | 1.1949 | 0.007933 | 0.037908 | 0.4766 | ↓ | -1.069 |
| PI160/203_2 | 0.031349 | 0.10307 | 0.54029 | ↓ | -0.888 | 0.061581 | 0.54498 | 0.6049 | ↓ | -0.725 | 0.10271 | 0.16039 | 0.6574 | ↓ | -0.605 |
| TAG562-FA200 | 0.031295 | 0.10307 | 0.41205 | ↓ | -1.279 | 0.42655 | 0.72444 | 1.2034 | ↑ | 0.2671 | 0.000538 | 0.023486 | 0.20242 | ↓ | -2.305 |
| PC180/202 | 0.087207 | 0.19115 | 0.75859 | ↓ | -0.399 | 0.048229 | 0.52329 | 0.67762 | ↓ | -0.561 | 0.017964 | 0.047732 | 0.61201 | ↓ | -0.708 |
| TAG547-FA181 | 0.019917 | 0.09098 | 0.23787 | ↓ | -2.072 | 0.15343 | 0.5914 | 0.58594 | ↓ | -0.771 | 0.00249 | 0.027699 | 0.13713 | ↓ | -2.866 |
| TAG503-FA181 | 0.002153 | 0.070016 | 0.20227 | ↓ | -2.306 | 0.21755 | 0.5914 | 0.6632 | ↓ | -0.592 | 0.002349 | 0.027585 | 0.19216 | ↓ | -2.38 |
| TAG484-FA120 | 0.046662 | 0.1315 | 0.11951 | ↓ | -3.065 | 0.2687 | 0.59421 | 0.3672 | ↓ | -1.445 | 0.10122 | 0.15878 | 0.18989 | ↓ | -2.397 |
| TAG504-FA161 | 0.005003 | 0.070016 | 0.18916 | ↓ | -2.402 | 0.12503 | 0.5914 | 0.53922 | ↓ | -0.891 | 0.003775 | 0.031552 | 0.1604 | ↓ | -2.64 |
| PG160/203_2 | 0.15021 | 0.28426 | 0.58938 | ↓ | -0.763 | 0.13282 | 0.5914 | 0.59167 | ↓ | -0.757 | 0.008756 | 0.037908 | 0.35435 | ↓ | -1.497 |
| TAG524-FA161 | 0.000898 | 0.070016 | 0.19574 | ↓ | -2.353 | 0.35517 | 0.67101 | 0.72425 | ↓ | -0.465 | 0.003053 | 0.029739 | 0.22848 | ↓ | -2.13 |
| TAG563-FA182 | 0.022665 | 0.091642 | 0.43566 | ↓ | -1.199 | 0.81249 | 0.90726 | 0.89739 | ↓ | -0.156 | 0.003061 | 0.029739 | 0.29235 | ↓ | -1.774 |
| TAG565-FA201 | 0.028682 | 0.10057 | 0.3953 | ↓ | -1.339 | 0.24309 | 0.5914 | 0.61905 | ↓ | -0.692 | 0.005427 | 0.032569 | 0.26535 | ↓ | -1.914 |
| PI160/203 | 0.028991 | 0.10057 | 0.53149 | ↓ | -0.912 | 0.18507 | 0.5914 | 0.70314 | ↓ | -0.508 | 0.21721 | 0.29735 | 0.72155 | ↓ | -0.471 |
| PI180/180 | 0.014164 | 0.087814 | 0.56774 | ↓ | -0.817 | 0.42024 | 0.72374 | 1.168 | ↑ | 0.224 | 0.000331 | 0.023486 | 0.42229 | ↓ | -1.244 |
| TAG541-FA160 | 0.079737 | 0.1798 | 0.45084 | ↓ | -1.149 | 0.71458 | 0.85829 | 0.84375 | ↓ | -0.245 | 0.004254 | 0.032129 | 0.2422 | ↓ | -2.046 |
| PC182/224 | 0.027979 | 0.10008 | 0.57432 | ↓ | -0.8 | 0.31402 | 0.6275 | 0.77177 | ↓ | -0.374 | 0.065646 | 0.11488 | 0.6027 | ↓ | -0.731 |
| PC182/205 | 0.19576 | 0.34443 | 0.63277 | ↓ | -0.66 | 0.40192 | 0.70798 | 1.1874 | ↑ | 0.2479 | 0.37967 | 0.4646 | 0.71387 | ↓ | -0.486 |
| TAG545-FA180 | 0.024057 | 0.093463 | 0.29214 | ↓ | -1.775 | 0.10478 | 0.5914 | 0.55294 | ↓ | -0.855 | 0.004541 | 0.032129 | 0.19227 | ↓ | -2.379 |
| PI161/181_2 | 0.029344 | 0.10107 | 0.60741 | ↓ | -0.719 | 0.47072 | 0.73251 | 1.0708 | ↑ | 0.0988 | 0.018702 | 0.0487 | 0.55262 | ↓ | -0.856 |
| TAG524-FA160 | 0.010803 | 0.082341 | 0.37795 | ↓ | -1.404 | 0.26546 | 0.59421 | 0.71832 | ↓ | -0.477 | 0.049587 | 0.094943 | 0.47255 | ↓ | -1.082 |
| PC160/225 | 0.22446 | 0.37953 | 1.2424 | ↑ | 0.3131 | 0.29335 | 0.61133 | 1.1629 | ↑ | 0.2177 | 0.43674 | 0.516 | 1.1791 | ↑ | 0.2376 |
| TAG524-FA183 | 0.008542 | 0.077743 | 0.29468 | ↓ | -1.763 | 0.17648 | 0.5914 | 0.682 | ↓ | -0.552 | 0.006219 | 0.03431 | 0.26159 | ↓ | -1.935 |
| TAG561-FA181 | 0.062573 | 0.15257 | 0.42775 | ↓ | -1.225 | 0.97763 | 0.99244 | 0.95625 | ↓ | -0.065 | 0.001412 | 0.023486 | 0.19775 | ↓ | -2.338 |
| TAG482-FA140 | 0.018005 | 0.09098 | 0.29281 | ↓ | -1.772 | 0.16984 | 0.5914 | 0.62206 | ↓ | -0.685 | 0.018179 | 0.047927 | 0.28247 | ↓ | -1.824 |
| PI200/204_2 | 0.006734 | 0.073064 | 0.54508 | ↓ | -0.875 | 0.20431 | 0.5914 | 0.80712 | ↓ | -0.309 | 0.012808 | 0.041839 | 0.56538 | ↓ | -0.823 |
| TAG502-FA140 | 0.008125 | 0.077743 | 0.26711 | ↓ | -1.905 | 0.53063 | 0.74837 | 0.88948 | ↓ | -0.169 | 0.008269 | 0.037908 | 0.24635 | ↓ | -2.021 |

| | | | | | | | | | | | | |
|---|---|---|---|---|---|---|---|---|---|---|---|---|
| TAG545-FA183 | 0.013768 | 0.087814 | 0.29264 | ↓ -1.773 | 0.17701 | 0.5914 | 0.60216 | ↓ -0.732 | 0.001447 | 0.023486 | 0.16087 | ↓ -2.636 |
| PG160/203 | 0.26199 | 0.42854 | 0.67552 | ↓ -0.566 | 0.16208 | 0.5914 | 0.63174 | ↓ -0.663 | 0.011755 | 0.040269 | 0.40491 | ↓ -1.304 |
| PI160/161 | 0.050994 | 0.13671 | 0.5008 | ↓ -0.998 | 0.3612 | 0.67183 | 0.72557 | ↓ -0.463 | 0.14941 | 0.22006 | 0.60941 | ↓ -0.715 |
| PE182/202 | 0.14309 | 0.27237 | 0.59888 | ↓ -0.74 | 0.10322 | 0.5914 | 0.57466 | ↓ -0.799 | 0.014675 | 0.043823 | 0.34376 | ↓ -1.541 |
| PI182/201 | 0.004086 | 0.070016 | 0.53106 | ↓ -0.913 | 0.77467 | 0.89253 | 1.1177 | ↑ 0.1606 | 0.034235 | 0.07554 | 0.62911 | ↓ -0.669 |
| LPA180 | 0.002691 | 0.070016 | 0.62535 | ↓ -0.677 | 0.70868 | 0.85829 | 1.0439 | ↑ 0.0619 | 0.013018 | 0.041839 | 0.67102 | ↓ -0.576 |
| LPC240 | 0.060071 | 0.14869 | 0.66282 | ↓ -0.593 | 0.69487 | 0.85482 | 0.9162 | ↓ -0.126 | 0.036374 | 0.077637 | 0.55218 | ↓ -0.857 |
| TAG524-FA200 | 0.042497 | 0.12471 | 0.38803 | ↓ -1.366 | 0.41431 | 0.71732 | 0.77489 | ↓ -0.368 | 0.082712 | 0.13842 | 0.45151 | ↓ -1.147 |
| TAG566-FA202 | 0.015347 | 0.090009 | 0.3192 | ↓ -1.648 | 0.086497 | 0.5914 | 0.48592 | ↓ -1.041 | 0.004112 | 0.032129 | 0.2333 | ↓ -2.1 |
| PS181/182 | 0.002413 | 0.070016 | 0.69699 | ↓ -0.521 | 0.005397 | 0.28651 | 0.73855 | ↓ -0.437 | 0.008221 | 0.037908 | 0.67764 | ↓ -0.561 |
| TAG524-FA181 | 0.005897 | 0.070016 | 0.25923 | ↓ -1.948 | 0.2447 | 0.5914 | 0.69476 | ↓ -0.525 | 0.003805 | 0.031552 | 0.23187 | ↓ -2.109 |
| PG161/180 | 0.91798 | 0.94708 | 1.0779 | ↑ 0.1083 | 0.098539 | 0.5914 | 0.66474 | ↓ -0.589 | 0.030592 | 0.071381 | 0.54005 | ↓ -0.889 |
| PI180/203_2 | 0.005208 | 0.070016 | 0.63663 | ↓ -0.651 | 0.022981 | 0.3649 | 0.77407 | ↓ -0.369 | 0.041533 | 0.085452 | 0.74843 | ↓ -0.418 |
| PI180/183_2 | 0.03879 | 0.11785 | 0.60365 | ↓ -0.728 | 0.23383 | 0.5914 | 0.78953 | ↓ -0.341 | 0.005079 | 0.032262 | 0.52711 | ↓ -0.924 |
| LPC241 | 0.017274 | 0.090926 | 0.56454 | ↓ -0.825 | 0.18316 | 0.5914 | 0.77348 | ↓ -0.371 | 0.027875 | 0.066228 | 0.5264 | ↓ -0.926 |
| CE203 | 0.1002 | 0.21178 | 1.3627 | ↑ 0.4465 | 0.017116 | 0.33253 | 0.6764 | ↓ -0.564 | 0.94677 | 0.95706 | 1.0328 | ↑ 0.0465 |
| TAG524-FA182 | 0.016614 | 0.090926 | 0.40387 | ↓ -1.308 | 0.27805 | 0.60136 | 0.73026 | ↓ -0.454 | 0.042562 | 0.086069 | 0.45807 | ↓ -1.126 |
| TAG545-FA182 | 0.00841 | 0.077743 | 0.30059 | ↓ -1.734 | 0.21318 | 0.5914 | 0.58613 | ↓ -0.771 | 0.004846 | 0.032129 | 0.2448 | ↓ -2.03 |
| TAG544-FA201 | 0.010046 | 0.079755 | 0.29258 | ↓ -1.773 | 0.30636 | 0.62637 | 0.64997 | ↓ -0.622 | 0.007454 | 0.037326 | 0.27123 | ↓ -1.882 |
| TAG545-FA161 | 0.006062 | 0.070016 | 0.3104 | ↓ -1.688 | 0.11842 | 0.5914 | 0.59516 | ↓ -0.749 | 0.005204 | 0.032262 | 0.19867 | ↓ -2.332 |
| TAG502-FA181 | 0.00139 | 0.070016 | 0.27402 | ↓ -1.868 | 0.43611 | 0.72846 | 0.93534 | ↓ -0.096 | 0.002415 | 0.027585 | 0.2631 | ↓ -1.926 |
| PG181/182_2 | 0.008737 | 0.077743 | 0.4691 | -1.092 | 0.32861 | 0.64559 | 0.77197 | -0.37338 | 0.057743 | 0.10543 | 0.53928 | -0.8909 |
| PC200/182 | 0.11831 | 0.2341 | 0.79671 | -0.32787 | 0.78201 | 0.8947 | 1.0641 | 0.08965 | 0.005987 | 0.034 | 0.51313 | -0.96262 |
| PC140/203 | 0.0058 | 0.070016 | 0.49465 | -1.0155 | 0.011266 | 0.28651 | 0.52343 | -0.93394 | 0.01408 | 0.043236 | 0.40054 | -1.32 |
| TAG543-FA183 | 0.031331 | 0.10307 | 0.38457 | -1.3787 | 0.21839 | 0.5914 | 0.72713 | -0.45972 | 0.003483 | 0.031053 | 0.22293 | -2.1653 |
| PC181/203 | 0.00601 | 0.070016 | 0.59604 | -0.74653 | 0.039727 | 0.48848 | 0.6818 | -0.55257 | 0.035683 | 0.077543 | 0.61709 | -0.69645 |
| TAG523-FA201 | 0.004716 | 0.070016 | 0.27432 | -1.8661 | 0.14705 | 0.5914 | 0.56154 | -0.83254 | 0.006349 | 0.034661 | 0.27708 | -1.8516 |
| LPC201 | 0.015046 | 0.089047 | 0.7106 | -0.49288 | 0.41134 | 0.71487 | 1.1382 | 0.18676 | 0.01981 | 0.050377 | 0.65345 | -0.61385 |
| PE181/205 | 0.34306 | 0.51184 | 0.72349 | -0.46695 | 0.81143 | 0.90726 | 0.86596 | -0.20763 | 0.11954 | 0.18311 | 0.56328 | -0.82808 |
| LPC161 | 0.002842 | 0.070016 | 0.5543 | -0.85126 | 0.067063 | 0.55914 | 0.80805 | -0.30749 | 0.009761 | 0.038366 | 0.58138 | -0.78246 |
| TAG481-FA181 | 0.01055 | 0.081762 | 0.2974 | -1.7495 | 0.24145 | 0.5914 | 0.75188 | -0.41143 | 0.004576 | 0.032129 | 0.24345 | -2.0383 |
| PC120/181 | 0.60183 | 0.7342 | 0.77853 | -0.36118 | 0.9425 | 0.97682 | 0.84382 | -0.24499 | 0.26469 | 0.34952 | 0.59455 | -0.75014 |
| PC150/203 | 0.032229 | 0.10543 | 0.60431 | -0.72663 | 0.007951 | 0.28651 | 0.56527 | -0.82298 | 0.008513 | 0.037908 | 0.52 | -0.94342 |

| | | | | | | | | | | | | |
|---|---|---|---|---|---|---|---|---|---|---|---|---|
| TAG567-FA183 | 0.015921 | 0.090926 | 0.26865 | -1.8962 | 0.13894 | 0.5914 | 0.56632 | -0.82031 | 0.010184 | 0.038366 | 0.23719 | -2.0759 |
| TAG544-FA183 | 0.020303 | 0.09098 | 0.34075 | -1.5532 | 0.21444 | 0.5914 | 0.68289 | -0.55028 | 0.002755 | 0.029398 | 0.22199 | -2.1715 |
| TAG523-FA183 | 0.027834 | 0.10008 | 0.36964 | -1.4358 | 0.19515 | 0.5914 | 0.7645 | -0.38741 | 0.009372 | 0.038185 | 0.27819 | -1.8459 |
| PE182/201 | 0.11015 | 0.22693 | 0.64818 | -0.62554 | 0.44931 | 0.72879 | 0.81297 | -0.29872 | 0.005417 | 0.032569 | 0.38068 | -1.3934 |
| TAG544-FA180 | 0.009136 | 0.077743 | 0.37202 | -1.4265 | 0.1771 | 0.5914 | 0.67065 | -0.57636 | 0.008629 | 0.037908 | 0.32982 | -1.6002 |
| TAG482-FA161 | 0.00072 | 0.070016 | 0.23103 | -2.1139 | 0.13138 | 0.5914 | 0.71717 | -0.47962 | 0.000869 | 0.023486 | 0.21893 | -2.1914 |
| TAG482-FA160 | 0.016473 | 0.090926 | 0.30196 | -1.7276 | 0.20042 | 0.5914 | 0.74292 | -0.42872 | 0.009606 | 0.038317 | 0.26191 | -1.9329 |
| PI180/181_2 | 0.020935 | 0.09098 | 0.63066 | -0.66507 | 0.040523 | 0.48852 | 1.4409 | 0.52695 | 0.000628 | 0.023486 | 0.49734 | -1.0077 |
| TAG523-FA180 | 0.029044 | 0.10057 | 0.40202 | -1.3147 | 0.2469 | 0.5914 | 0.8365 | -0.25756 | 0.003179 | 0.030434 | 0.24515 | -2.0283 |
| PG160/182_2 | 0.093253 | 0.20102 | 0.56552 | -0.82236 | 0.33023 | 0.64559 | 0.72911 | -0.45579 | 0.12203 | 0.18432 | 0.58093 | -0.78358 |
| TAG503-FA161 | 0.002817 | 0.070016 | 0.25343 | -1.9803 | 0.2071 | 0.5914 | 0.74209 | -0.43034 | 0.005107 | 0.032262 | 0.26497 | -1.9161 |
| PI180/181 | 0.030132 | 0.10164 | 0.65501 | -0.61042 | 0.033955 | 0.4421 | 1.574 | 0.65442 | 0.000245 | 0.023486 | 0.45014 | -1.1516 |
| PC182/226 | 0.019036 | 0.09098 | 0.55956 | -0.83764 | 0.30383 | 0.62637 | 1.1963 | 0.25857 | 0.20743 | 0.2904 | 0.75353 | -0.40826 |
| PC181/183 | 0.27696 | 0.44519 | 0.78888 | -0.34212 | 0.12258 | 0.5914 | 0.7219 | -0.47013 | 0.01003 | 0.038366 | 0.45389 | -1.1396 |
| PC182/204 | 0.047037 | 0.13199 | 0.61287 | -0.70636 | 0.90219 | 0.95366 | 1.0046 | 0.00666 | 0.19042 | 0.27093 | 0.71554 | -0.48289 |
| TAG565-FA182 | 0.023386 | 0.092269 | 0.36053 | -1.4718 | 0.25898 | 0.59421 | 0.62461 | -0.67896 | 0.017029 | 0.046633 | 0.31854 | -1.6504 |
| TAG544-FA161 | 0.018673 | 0.09098 | 0.33909 | -1.5602 | 0.41179 | 0.71487 | 0.72697 | -0.46003 | 0.006111 | 0.034 | 0.25707 | -1.9598 |
| PC181/226 | 0.057375 | 0.14421 | 0.71963 | -0.47468 | 0.16738 | 0.5914 | 0.83142 | -0.26636 | 0.042004 | 0.085452 | 0.71825 | -0.47745 |
| TAG520-FA200 | 0.027154 | 0.09931 | 0.377 | -1.4074 | 0.050925 | 0.53349 | 0.43565 | -1.1988 | 0.001569 | 0.023486 | 0.21808 | -2.1971 |
| SMd181/182 | 0.068919 | 0.16335 | 1.2567 | 0.32961 | 0.61869 | 0.80957 | 1.0676 | 0.094349 | 0.43587 | 0.516 | 1.086 | 0.11897 |
| PC140/202 | 0.059426 | 0.14812 | 0.53999 | -0.88899 | 0.11643 | 0.5914 | 0.6052 | -0.72451 | 0.037908 | 0.080123 | 0.46015 | -1.1198 |
| TAG524-FA180 | 0.087054 | 0.19115 | 0.37526 | -1.4141 | 0.24705 | 0.5914 | 0.68914 | -0.53714 | 0.016827 | 0.046417 | 0.24587 | -2.024 |
| PG160/182 | 0.092879 | 0.20088 | 0.55848 | -0.84043 | 0.31308 | 0.6275 | 0.71228 | -0.48949 | 0.1208 | 0.18375 | 0.57483 | -0.79878 |
| PI180/182_2 | 0.051704 | 0.13699 | 0.68687 | -0.54189 | 0.81912 | 0.90757 | 0.95164 | -0.07151 | 0.007076 | 0.03715 | 0.64741 | -0.62724 |
| TAG526-FA181 | 0.021809 | 0.091642 | 0.31251 | -1.678 | 0.15774 | 0.5914 | 0.59472 | -0.74972 | 0.003926 | 0.031552 | 0.18824 | -2.4093 |
| TAG483-FA161 | 0.003444 | 0.070016 | 0.20728 | -2.2703 | 0.19291 | 0.5914 | 0.63263 | -0.66058 | 0.005828 | 0.033574 | 0.23715 | -2.0761 |
| TAG565-FA202 | 0.030408 | 0.10204 | 0.36668 | -1.4474 | 0.25723 | 0.59421 | 0.59907 | -0.7392 | 0.004732 | 0.032129 | 0.24767 | -2.0135 |
| LPC170 | 0.084405 | 0.18818 | 0.66914 | -0.57962 | 0.62786 | 0.81455 | 0.87726 | -0.18892 | 0.057064 | 0.10494 | 0.65539 | -0.60956 |
| TAG545-FA181 | 0.024119 | 0.093463 | 0.3609 | -1.4703 | 0.35789 | 0.67101 | 0.69745 | -0.51983 | 0.00353 | 0.031053 | 0.2352 | -2.088 |
| TAG523-FA202 | 0.011464 | 0.082752 | 0.32697 | -1.6128 | 0.34498 | 0.66249 | 0.78724 | -0.34513 | 0.00725 | 0.037244 | 0.29035 | -1.7841 |
| PI180/182 | 0.03876 | 0.11785 | 0.67695 | -0.56287 | 0.47224 | 0.73251 | 0.90628 | -0.14198 | 0.001943 | 0.024802 | 0.62579 | -0.67626 |
| TAG540-FA160 | 0.007343 | 0.075267 | 0.31832 | -1.6515 | 0.051676 | 0.53349 | 0.46012 | -1.1199 | 0.001442 | 0.023486 | 0.23303 | -2.1014 |
| TAG526-FA160 | 0.013179 | 0.087814 | 0.23151 | -2.1108 | 0.11306 | 0.5914 | 0.66598 | -0.58644 | 0.00928 | 0.038185 | 0.20106 | -2.3143 |

| | | | | | | | | | | | | |
|---|---|---|---|---|---|---|---|---|---|---|---|---|
| TAG526-FA205 | 0.076211 | 0.1759 | 0.34193 | -1.5482 | 0.24291 | 0.5914 | 0.56377 | -0.82683 | 0.37614 | 0.46115 | 0.59028 | -0.76052 |
| PS180/181 | 0.004032 | 0.070016 | 0.52972 | -0.91669 | 0.048013 | 0.52329 | 0.70152 | -0.51145 | 0.051649 | 0.098027 | 0.67604 | -0.56482 |
| TAG523-FA182 | 0.013513 | 0.087814 | 0.48561 | -1.0421 | 0.56079 | 0.76614 | 0.91829 | -0.12298 | 0.015432 | 0.04485 | 0.45581 | -1.1335 |
| TAG545-FA202 | 0.021071 | 0.09098 | 0.30042 | -1.735 | 0.25039 | 0.5914 | 0.66689 | -0.58447 | 0.005188 | 0.032262 | 0.20899 | -2.2585 |
| TAG483-FA140 | 0.005614 | 0.070016 | 0.16477 | -2.6015 | 0.13609 | 0.5914 | 0.5795 | -0.78711 | 0.003823 | 0.031552 | 0.1386 | -2.851 |
| PE180/202 | 0.63793 | 0.75507 | 0.91628 | -0.12615 | 0.096708 | 0.5914 | 0.74793 | -0.41903 | 0.005453 | 0.032569 | 0.53871 | -0.89243 |
| PI180/202_2 | 0.028393 | 0.10046 | 0.60683 | -0.72063 | 0.008834 | 0.28651 | 0.58489 | -0.77376 | 0.01558 | 0.045078 | 0.54814 | -0.86739 |
| PE181/204 | 0.037431 | 0.11603 | 0.60975 | -0.71372 | 0.56859 | 0.77018 | 0.88129 | -0.1823 | 0.049129 | 0.094905 | 0.58983 | -0.76163 |
| TAG523-FA181 | 0.003621 | 0.070016 | 0.45097 | -1.1489 | 0.47071 | 0.73251 | 0.87949 | -0.18527 | 0.013047 | 0.041839 | 0.48475 | -1.0447 |
| TAG503-FA160 | 0.009341 | 0.077962 | 0.31564 | -1.6636 | 0.18419 | 0.5914 | 0.759 | -0.39783 | 0.006996 | 0.037025 | 0.27512 | -1.8619 |
| TAG483-FA182 | 0.008915 | 0.077743 | 0.19455 | -2.3618 | 0.20723 | 0.5914 | 0.5984 | -0.74083 | 0.02111 | 0.052453 | 0.24601 | -2.0232 |
| PC140/161 | 0.46356 | 0.62952 | 0.83825 | -0.25454 | 0.29393 | 0.61133 | 0.78384 | -0.35137 | 0.09533 | 0.15225 | 0.64295 | -0.63721 |
| TAG523-FA161 | 0.002773 | 0.070016 | 0.26105 | -1.9376 | 0.71021 | 0.85829 | 0.89617 | -0.15815 | 0.003397 | 0.03071 | 0.24739 | -2.0152 |
| TAG5610-FA182 | 0.034598 | 0.1115 | 0.367 | -1.4462 | 0.47314 | 0.73251 | 0.86815 | -0.20398 | 0.005188 | 0.032262 | 0.23141 | -2.1114 |
| PC180/183 | 0.85577 | 0.90734 | 0.95788 | -0.06209 | 0.28021 | 0.60269 | 0.75865 | -0.39849 | 0.019736 | 0.050377 | 0.46582 | -1.1022 |
| TAG564-FA201 | 0.058161 | 0.14563 | 0.44348 | -1.1731 | 0.99166 | 0.99929 | 0.90735 | -0.14027 | 0.007627 | 0.037551 | 0.28624 | -1.8047 |
| PE182/161 | 0.043243 | 0.12568 | 0.70242 | -0.5096 | 0.85119 | 0.92819 | 0.98581 | -0.02062 | 0.004884 | 0.032129 | 0.44568 | -1.1659 |
| PE181/226 | 0.019401 | 0.09098 | 0.55835 | -0.84075 | 0.40019 | 0.70794 | 0.81044 | -0.30322 | 0.043156 | 0.0867 | 0.60446 | -0.72627 |
| TAG481-FA140 | 0.013777 | 0.087814 | 0.3179 | -1.6534 | 0.24082 | 0.5914 | 0.79586 | -0.32942 | 0.006084 | 0.034 | 0.26089 | -1.9385 |
| PS161/182_2 | 0.023066 | 0.091642 | 0.75089 | -0.41333 | 0.010865 | 0.28651 | 0.76471 | -0.38701 | 0.002929 | 0.029739 | 0.60477 | -0.72554 |
| PC140/181 | 0.022501 | 0.091642 | 0.59683 | -0.74461 | 0.58961 | 0.78747 | 0.87924 | -0.18567 | 0.047805 | 0.093457 | 0.55788 | -0.84198 |
| PI160/161_2 | 0.10293 | 0.21409 | 0.62271 | -0.68337 | 0.49759 | 0.73497 | 0.82093 | -0.28467 | 0.29765 | 0.38143 | 0.74196 | -0.43058 |
| PA180/182 | 0.043579 | 0.12609 | 0.685 | -0.54582 | 0.087928 | 0.5914 | 1.4252 | 0.51117 | 0.022967 | 0.056208 | 0.58702 | -0.76852 |
| TAG504-FA160 | 0.026885 | 0.09931 | 0.31446 | -1.669 | 0.12974 | 0.5914 | 0.6271 | -0.67323 | 0.013638 | 0.042554 | 0.26113 | -1.9371 |
| PC182/225 | 0.003696 | 0.070016 | 0.57698 | -0.79339 | 0.81947 | 0.90757 | 1.0398 | 0.056356 | 0.19538 | 0.27634 | 0.79886 | -0.32398 |
| TAG523-FA160 | 0.009862 | 0.079755 | 0.47279 | -1.0807 | 0.41172 | 0.71487 | 0.84949 | -0.23533 | 0.011815 | 0.040269 | 0.45762 | -1.1278 |
| PI180/202 | 0.022624 | 0.091642 | 0.64746 | -0.62713 | 0.008141 | 0.28651 | 0.64387 | -0.63516 | 0.015176 | 0.044502 | 0.60764 | -0.71872 |
| PS180/181_2 | 0.00574 | 0.070016 | 0.56533 | -0.82282 | 0.03156 | 0.4227 | 0.67514 | -0.56675 | 0.11776 | 0.18124 | 0.74642 | -0.42195 |
| SMd181/203 | 0.35189 | 0.5202 | 1.1087 | 0.14886 | 0.005239 | 0.28651 | 0.78299 | -0.35294 | 0.96217 | 0.96812 | 1.007 | 0.010119 |
| PE181/202 | 0.23766 | 0.39384 | 0.70217 | -0.5101 | 0.26175 | 0.59421 | 0.71508 | -0.48382 | 0.019383 | 0.049679 | 0.40834 | -1.2922 |
| TAG484-FA182 | 0.033487 | 0.10846 | 0.18547 | -2.4307 | 0.16109 | 0.5914 | 0.40516 | -1.3034 | 0.041978 | 0.085452 | 0.22109 | -2.1773 |
| PI180/205 | 0.35239 | 0.5202 | 0.82224 | -0.28237 | 0.51945 | 0.74554 | 0.87191 | -0.19775 | 0.63685 | 0.70508 | 0.90561 | -0.14304 |
| PEP-180/182 | 0.013701 | 0.087814 | 0.72154 | -0.47084 | 0.76294 | 0.89227 | 1.0522 | 0.07342 | 0.000705 | 0.023486 | 0.41057 | -1.2843 |

| | | | | | | | | | | | | |
|---|---|---|---|---|---|---|---|---|---|---|---|---|
| PE181/182 | 0.24377 | 0.40176 | 0.6815 | -0.5532 | 0.70535 | 0.85829 | 0.84819 | -0.23753 | 0.021815 | 0.05396 | 0.41363 | -1.2736 |
| PEO-160/225 | 0.21224 | 0.36552 | 1.3132 | 0.39313 | 0.61935 | 0.80957 | 1.0868 | 0.12005 | 0.56788 | 0.63944 | 1.0736 | 0.10244 |
| PI180/203 | 0.011131 | 0.082341 | 0.68006 | -0.55626 | 0.068046 | 0.55914 | 0.81244 | -0.29967 | 0.093014 | 0.14988 | 0.80556 | -0.31193 |
| LPC226 | 0.67624 | 0.78333 | 1.0579 | 0.08114 | 0.47413 | 0.73251 | 1.1578 | 0.21135 | 0.82631 | 0.85521 | 1.0181 | 0.025864 |
| PEP-180/181 | 0.11127 | 0.22778 | 0.78384 | -0.35137 | 0.35916 | 0.67101 | 1.2691 | 0.3438 | 0.001402 | 0.023486 | 0.42287 | -1.2417 |
| TAG569-FA204 | 0.032446 | 0.10561 | 0.51641 | -0.9534 | 0.039769 | 0.48848 | 0.5096 | -0.97256 | 0.056525 | 0.10454 | 0.51251 | -0.96435 |
| PC140/182 | 0.01411 | 0.087814 | 0.59228 | -0.75565 | 0.79379 | 0.90076 | 0.94695 | -0.07863 | 0.073476 | 0.12688 | 0.60756 | -0.7189 |
| TAG544-FA182 | 0.026655 | 0.09931 | 0.4054 | -1.3026 | 0.61431 | 0.80628 | 0.80972 | -0.30451 | 0.012148 | 0.040763 | 0.33148 | -1.593 |
| TAG522-FA181 | 0.009918 | 0.079755 | 0.45036 | -1.1509 | 0.80978 | 0.90726 | 0.98817 | -0.01717 | 0.014304 | 0.043513 | 0.44244 | -1.1765 |
| PEP-180/203 | 0.15268 | 0.28643 | 0.8258 | -0.27614 | 0.77866 | 0.89253 | 0.96471 | -0.05184 | 0.005525 | 0.0327 | 0.54911 | -0.86483 |
| PC150/182 | 0.3297 | 0.49916 | 0.81366 | -0.2975 | 0.76491 | 0.8924 | 0.91965 | -0.12085 | 0.055117 | 0.10311 | 0.60264 | -0.73063 |
| TAG566-FA182 | 0.010291 | 0.080715 | 0.34256 | -1.5456 | 0.16963 | 0.5914 | 0.62042 | -0.68869 | 0.064524 | 0.11374 | 0.45863 | -1.1246 |
| PC182/201 | 0.15485 | 0.28796 | 0.7394 | -0.43557 | 0.60673 | 0.79956 | 1.1059 | 0.14526 | 0.055927 | 0.10402 | 0.57626 | -0.79521 |
| TAG567-FA182 | 0.024636 | 0.0949 | 0.39424 | -1.3428 | 0.18667 | 0.5914 | 0.60971 | -0.7138 | 0.059949 | 0.10781 | 0.43667 | -1.1954 |
| PC160/205 | 0.34686 | 0.51554 | 1.4028 | 0.48828 | 0.57669 | 0.77407 | 1.1984 | 0.26105 | 0.4936 | 0.57381 | 1.4259 | 0.51184 |
| TAG568-FA183 | 0.019441 | 0.09098 | 0.33622 | -1.5725 | 0.15208 | 0.5914 | 0.67396 | -0.56926 | 0.012147 | 0.040763 | 0.2854 | -1.8089 |
| TAG524-FA203 | 0.016411 | 0.090926 | 0.31669 | -1.6588 | 0.20416 | 0.5914 | 0.64034 | -0.6431 | 0.010351 | 0.038366 | 0.29186 | -1.7767 |
| HCER180 | 0.053424 | 0.13806 | 1.4036 | 0.4891 | 0.69711 | 0.85482 | 1.0665 | 0.092843 | 0.54932 | 0.62849 | 1.096 | 0.13231 |
| TAG567-FA203 | 0.013286 | 0.087814 | 0.4421 | -1.1776 | 0.057112 | 0.53695 | 0.52122 | -0.94004 | 0.001131 | 0.023486 | 0.28361 | -1.818 |
| PC170/203 | 0.2809 | 0.4504 | 0.80533 | -0.31234 | 0.11194 | 0.5914 | 0.73701 | -0.44024 | 0.036354 | 0.077637 | 0.65173 | -0.61765 |
| TAG502-FA180 | 0.050369 | 0.13671 | 0.36951 | -1.4363 | 0.23651 | 0.5914 | 0.74078 | -0.43288 | 0.004467 | 0.032129 | 0.21342 | -2.2282 |
| TAG483-FA120 | 0.075612 | 0.17517 | 0.19758 | -2.3395 | 0.39165 | 0.70045 | 0.56022 | -0.83594 | 0.077554 | 0.13217 | 0.25552 | -1.9685 |
| SMd181/160 | 0.31251 | 0.4821 | 1.0699 | 0.097535 | 0.21562 | 0.5914 | 1.0967 | 0.13316 | 0.49624 | 0.57483 | 1.0404 | 0.057097 |
| TAG544-FA202 | 0.02674 | 0.09931 | 0.39605 | -1.3362 | 0.29074 | 0.61133 | 0.74718 | -0.42046 | 0.016678 | 0.046398 | 0.35056 | -1.5123 |
| TAG502-FA161 | 0.003413 | 0.070016 | 0.32625 | -1.6159 | 0.36884 | 0.67841 | 0.92453 | -0.11321 | 0.004367 | 0.032129 | 0.30502 | -1.713 |
| PE140/204 | 0.023086 | 0.091642 | 0.58334 | -0.77758 | 0.093525 | 0.5914 | 0.64585 | -0.63074 | 0.031476 | 0.072338 | 0.56238 | -0.83039 |
| TAG563-FA202 | 0.085574 | 0.19013 | 0.47912 | -1.0615 | 0.6958 | 0.85482 | 0.90456 | -0.14471 | 0.009327 | 0.038185 | 0.29546 | -1.759 |
| TAG569-FA183 | 0.081836 | 0.18308 | 0.44142 | -1.1798 | 0.16688 | 0.5914 | 0.6901 | -0.53511 | 0.044496 | 0.088045 | 0.3739 | -1.4193 |
| LPE204 | 0.017157 | 0.090926 | 1.553 | 0.63504 | 0.001537 | 0.28651 | 1.7486 | 0.80617 | 0.76216 | 0.81205 | 1.0205 | 0.02931 |
| PI180/224 | 0.016111 | 0.090926 | 0.68072 | -0.55486 | 0.090813 | 0.5914 | 0.80692 | -0.30951 | 0.056778 | 0.10471 | 0.76392 | -0.38851 |
| TAG562-FA180 | 0.097676 | 0.20986 | 0.46091 | -1.1175 | 0.98025 | 0.99244 | 1.047 | 0.066265 | 0.002951 | 0.029739 | 0.21468 | -2.2197 |
| TAG521-FA161 | 0.018034 | 0.09098 | 0.39702 | -1.3327 | 0.2459 | 0.5914 | 0.81695 | -0.29169 | 0.00034 | 0.023486 | 0.19248 | -2.3772 |
| PS181/182_2 | 0.02887 | 0.10057 | 0.75225 | -0.41072 | 0.67853 | 0.84298 | 0.95975 | -0.05927 | 0.000152 | 0.023486 | 0.58683 | -0.76899 |

| | | | | | | | | | | | | |
|---|---|---|---|---|---|---|---|---|---|---|---|---|
| TAG503-FA183 | 0.046445 | 0.13146 | 0.36106 | -1.4697 | 0.18911 | 0.5914 | 0.79343 | -0.33382 | 0.034347 | 0.07554 | 0.33011 | -1.599 |
| PE170/181 | 0.88214 | 0.92068 | 0.85153 | -0.23188 | 0.5726 | 0.77177 | 0.74614 | -0.42249 | 0.018268 | 0.047955 | 0.36691 | -1.4465 |
| TAG569-FA205 | 0.23547 | 0.39336 | 0.61892 | -0.69217 | 0.11379 | 0.5914 | 0.53039 | -0.91488 | 0.36064 | 0.4455 | 0.74084 | -0.43277 |
| TAG563-FA181 | 0.22639 | 0.38082 | 0.56165 | -0.83226 | 0.33381 | 0.64869 | 1.2807 | 0.35697 | 0.016178 | 0.045884 | 0.30799 | -1.699 |
| PE181/183 | 0.87731 | 0.91969 | 0.93159 | -0.10223 | 0.09851 | 0.5914 | 0.5108 | -0.96916 | 0.016647 | 0.046398 | 0.32429 | -1.6246 |
| TAG481-FA160 | 0.008808 | 0.077743 | 0.38706 | -1.3694 | 0.24623 | 0.5914 | 0.90819 | -0.13894 | 0.004878 | 0.032129 | 0.31612 | -1.6614 |
| TAG542-FA160 | 0.071948 | 0.16788 | 0.52318 | -0.93461 | 0.50499 | 0.73692 | 0.84945 | -0.23539 | 0.008871 | 0.037908 | 0.33707 | -1.5689 |
| TAG546-FA203 | 0.02236 | 0.091642 | 0.26533 | -1.9142 | 0.20614 | 0.5914 | 0.54607 | -0.87284 | 0.01601 | 0.045884 | 0.23601 | -2.0831 |
| TAG567-FA224 | 0.001868 | 0.070016 | 0.28015 | -1.8357 | 0.12017 | 0.5914 | 0.64029 | -0.6432 | 0.007908 | 0.037908 | 0.33072 | -1.5963 |
| TAG501-FA180 | 0.019367 | 0.09098 | 0.38909 | -1.3618 | 0.28133 | 0.60269 | 0.91729 | -0.12455 | 0.001983 | 0.024824 | 0.23795 | -2.0713 |
| PC181/161 | 0.016719 | 0.090926 | 0.64038 | -0.643 | 0.26958 | 0.59421 | 0.86029 | -0.21711 | 0.038745 | 0.080546 | 0.6264 | -0.67484 |
| TAG543-FA180 | 0.049748 | 0.13642 | 0.52278 | -0.93572 | 0.63311 | 0.8194 | 0.92659 | -0.10999 | 0.013346 | 0.042154 | 0.39966 | -1.3232 |
| PC180/204 | 0.48508 | 0.64579 | 1.0685 | 0.095593 | 0.77874 | 0.89253 | 1.0055 | 0.007849 | 0.99078 | 0.99078 | 0.96666 | -0.04893 |
| TAG544-FA181 | 0.027723 | 0.10008 | 0.43444 | -1.2028 | 0.99135 | 0.99929 | 0.96406 | -0.0528 | 0.003343 | 0.03071 | 0.27846 | -1.8445 |
| TAG523-FA200 | 0.00471 | 0.070016 | 0.38581 | -1.3741 | 0.54452 | 0.75907 | 0.95688 | -0.06359 | 0.017577 | 0.047479 | 0.43059 | -1.2156 |
| PI180/224_2 | 0.007899 | 0.077743 | 0.67136 | -0.57484 | 0.027359 | 0.40479 | 0.77263 | -0.37215 | 0.042572 | 0.086069 | 0.74065 | -0.43314 |
| LPC204 | 0.73504 | 0.83219 | 1.0303 | 0.043063 | 0.18266 | 0.5914 | 1.247 | 0.31846 | 0.47603 | 0.55937 | 1.0818 | 0.11341 |
| TAG522-FA161 | 0.006346 | 0.070016 | 0.3367 | -1.5705 | 0.56137 | 0.76614 | 0.9369 | -0.09403 | 0.000957 | 0.023486 | 0.21904 | -2.1907 |
| TAG543-FA160 | 0.11013 | 0.22693 | 0.54017 | -0.8885 | 0.65691 | 0.83216 | 0.94694 | -0.07866 | 0.038643 | 0.080546 | 0.43647 | -1.1961 |
| TAG544-FA221 | 0.057126 | 0.14414 | 0.38258 | -1.3862 | 0.73068 | 0.86886 | 0.7988 | -0.32409 | 0.010304 | 0.038366 | 0.24816 | -2.0107 |
| TAG542-FA182 | 0.076788 | 0.17602 | 0.49429 | -1.0166 | 0.30716 | 0.62637 | 0.76296 | -0.39032 | 0.004358 | 0.032129 | 0.28299 | -1.8212 |
| PC170/181 | 0.6142 | 0.74098 | 0.90248 | -0.14803 | 0.80114 | 0.90546 | 0.99793 | -0.00299 | 0.06794 | 0.11794 | 0.70494 | -0.50444 |
| TAG567-FA161 | 0.056383 | 0.14282 | 0.36027 | -1.4729 | 0.44461 | 0.72846 | 0.69217 | -0.5308 | 0.050207 | 0.09557 | 0.35235 | -1.5049 |
| PE180/225 | 0.13552 | 0.26179 | 1.3018 | 0.38053 | 0.81618 | 0.90757 | 1.0309 | 0.043944 | 0.12317 | 0.18517 | 0.73977 | -0.43486 |
| PG182/182 | 0.11554 | 0.2333 | 0.53332 | -0.90692 | 0.13282 | 0.5914 | 0.54603 | -0.87295 | 0.09453 | 0.15195 | 0.50973 | -0.97219 |
| PE181/225 | 0.11398 | 0.23115 | 0.7285 | -0.457 | 0.14007 | 0.5914 | 0.76209 | -0.39196 | 0.008107 | 0.037908 | 0.52054 | -0.94191 |
| TAG543-FA182 | 0.025872 | 0.097922 | 0.4251 | -1.2341 | 0.56013 | 0.76614 | 0.86387 | -0.21111 | 0.012907 | 0.041839 | 0.35758 | -1.4837 |
| PS180/182 | 0.020031 | 0.09098 | 0.62597 | -0.67584 | 0.23525 | 0.5914 | 0.80363 | -0.3154 | 0.064645 | 0.11374 | 0.69109 | -0.53305 |
| SMd181/202 | 0.13189 | 0.2563 | 1.2605 | 0.33395 | 0.17216 | 0.5914 | 0.82351 | -0.28014 | 0.65081 | 0.71797 | 1.0556 | 0.078028 |
| PE160/225 | 0.089447 | 0.19475 | 1.4716 | 0.55738 | 0.52607 | 0.74605 | 0.86808 | -0.20409 | 0.10049 | 0.15802 | 0.6878 | -0.53995 |
| TAG480-FA140 | 0.007429 | 0.075267 | 0.41243 | -1.2778 | 0.2007 | 0.5914 | 0.87227 | -0.19715 | 0.000229 | 0.023486 | 0.24594 | -2.0236 |
| PC140/140 | 0.2854 | 0.45316 | 0.68492 | -0.54598 | 0.35011 | 0.66643 | 0.72129 | -0.47135 | 0.064133 | 0.11369 | 0.49249 | -1.0218 |
| TAG482-FA181 | 0.004107 | 0.070016 | 0.20907 | -2.2579 | 0.24182 | 0.5914 | 0.66377 | -0.59123 | 0.010562 | 0.038366 | 0.26441 | -1.9191 |

| | | | | | | | | | | | | |
|---|---|---|---|---|---|---|---|---|---|---|---|---|
| TAG564-FA202 | 0.023596 | 0.092536 | 0.39326 | -1.3464 | 0.34024 | 0.65726 | 0.70324 | -0.50792 | 0.006389 | 0.034661 | 0.28989 | -1.7864 |
| TAG543-FA181 | 0.052186 | 0.13754 | 0.49656 | -1.01 | 0.43281 | 0.72806 | 1.2034 | 0.26709 | 0.007266 | 0.037244 | 0.33645 | -1.5715 |
| TAG564-FA181 | 0.070885 | 0.16659 | 0.45971 | -1.1212 | 0.86108 | 0.93267 | 0.89077 | -0.16688 | 0.010661 | 0.038366 | 0.30303 | -1.7225 |
| PE140/182 | 0.15216 | 0.28643 | 0.61143 | -0.70974 | 0.21134 | 0.5914 | 0.65363 | -0.61345 | 0.011297 | 0.039348 | 0.37477 | -1.4159 |
| TAG543-FA161 | 0.007421 | 0.075267 | 0.40975 | -1.2872 | 0.50014 | 0.73497 | 0.88476 | -0.17665 | 0.000985 | 0.023486 | 0.27781 | -1.8478 |
| TAG504-FA181 | 0.003483 | 0.070016 | 0.21229 | -2.2359 | 0.15368 | 0.5914 | 0.61421 | -0.7032 | 0.001601 | 0.023486 | 0.17652 | -2.5021 |
| TAG568-FA181 | 0.086993 | 0.19115 | 0.52981 | -0.91646 | 0.36795 | 0.67841 | 0.75722 | -0.40121 | 0.065188 | 0.11439 | 0.47406 | -1.0768 |
| PEP-180/202 | 0.19703 | 0.34514 | 0.83727 | -0.25624 | 0.99767 | 0.99929 | 1.0388 | 0.054915 | 0.001532 | 0.023486 | 0.45887 | -1.1238 |
| PE160/203 | 0.97566 | 0.98169 | 0.95037 | -0.07344 | 0.38541 | 0.6931 | 0.77485 | -0.36802 | 0.037192 | 0.078866 | 0.55015 | -0.8621 |
| TAG565-FA181 | 0.016961 | 0.090926 | 0.37391 | -1.4192 | 0.49273 | 0.73497 | 0.79052 | -0.33913 | 0.034211 | 0.07554 | 0.41318 | -1.2752 |
| TAG564-FA182 | 0.079156 | 0.1798 | 0.44375 | -1.1722 | 0.66362 | 0.83459 | 0.78453 | -0.3501 | 0.008551 | 0.037908 | 0.26907 | -1.8939 |
| TAG566-FA203 | 0.022836 | 0.091642 | 0.4334 | -1.2062 | 0.091615 | 0.5914 | 0.5277 | -0.92222 | 0.001767 | 0.023486 | 0.2563 | -1.9641 |
| PEP-160/160 | 0.1327 | 0.25711 | 1.5167 | 0.60094 | 0.37881 | 0.68692 | 1.2763 | 0.35195 | 0.553 | 0.63102 | 1.1174 | 0.16017 |
| TAG522-FA160 | 0.020277 | 0.09098 | 0.47108 | -1.086 | 0.76974 | 0.89253 | 0.97728 | -0.03315 | 0.010391 | 0.038366 | 0.40146 | -1.3167 |
| LPE224 | 0.064374 | 0.15579 | 1.7283 | 0.78936 | 0.13232 | 0.5914 | 1.4728 | 0.55852 | 0.30424 | 0.38911 | 0.80883 | -0.3061 |
| PI161/180_2 | 0.11957 | 0.23517 | 0.75026 | -0.41454 | 0.036356 | 0.46408 | 0.687 | -0.54161 | 0.012933 | 0.041839 | 0.62431 | -0.67968 |
| PEO-160/205 | 0.68459 | 0.78879 | 0.94223 | -0.08584 | 0.26794 | 0.59421 | 0.80503 | -0.31288 | 0.009435 | 0.038185 | 0.56704 | -0.81848 |
| TAG568-FA204 | 0.055667 | 0.14156 | 0.68412 | -0.54768 | 0.022295 | 0.3649 | 0.55908 | -0.83887 | 0.05925 | 0.10714 | 0.61849 | -0.69317 |
| PC150/181 | 0.30647 | 0.47682 | 0.80491 | -0.31309 | 0.56068 | 0.76614 | 1.0683 | 0.095325 | 0.079165 | 0.13374 | 0.64764 | -0.62674 |
| TAG546-FA161 | 0.053469 | 0.13806 | 0.35833 | -1.4806 | 0.27235 | 0.59421 | 0.60332 | -0.72899 | 0.017785 | 0.047616 | 0.27428 | -1.8663 |
| TAG522-FA182 | 0.027125 | 0.09931 | 0.47832 | -1.0639 | 0.35912 | 0.67101 | 0.85171 | -0.23157 | 0.013936 | 0.043203 | 0.40204 | -1.3146 |
| LPC140 | 0.037956 | 0.11711 | 0.63904 | -0.64601 | 0.56118 | 0.76614 | 0.89376 | -0.16205 | 0.0145 | 0.043678 | 0.53584 | -0.90014 |
| TAG501-FA140 | 0.010963 | 0.082341 | 0.39553 | -1.3381 | 0.40409 | 0.70798 | 0.99034 | -0.01401 | 0.000606 | 0.023486 | 0.21906 | -2.1906 |
| PEP-180/183 | 0.067345 | 0.16118 | 0.6979 | -0.51891 | 0.69843 | 0.85482 | 0.92106 | -0.11863 | 0.000301 | 0.023486 | 0.32162 | -1.6366 |
| TAG543-FA202 | 0.060871 | 0.14969 | 0.43247 | -1.2093 | 0.5535 | 0.76614 | 0.8635 | -0.21173 | 0.016282 | 0.045884 | 0.32165 | -1.6364 |
| PC180/140 | 0.24056 | 0.39747 | 0.77588 | -0.3661 | 0.53835 | 0.75208 | 0.87486 | -0.19287 | 0.1087 | 0.16889 | 0.65507 | -0.61028 |
| TAG568-FA225 | 0.017319 | 0.090926 | 0.23677 | -2.0785 | 0.3083 | 0.62637 | 0.66874 | -0.58049 | 0.066949 | 0.11653 | 0.32698 | -1.6127 |
| SMd181/161 | 0.40651 | 0.57281 | 1.1168 | 0.15942 | 0.34745 | 0.66513 | 1.1163 | 0.15867 | 0.035734 | 0.077543 | 1.2702 | 0.34506 |
| PEP-180/205 | 0.77398 | 0.85545 | 0.93984 | -0.08951 | 0.71302 | 0.85829 | 0.93156 | -0.10228 | 0.026588 | 0.063634 | 0.54277 | -0.88158 |
| PEP-160/182 | 0.2817 | 0.45058 | 0.83681 | -0.25704 | 0.56636 | 0.77018 | 1.1269 | 0.17238 | 0.019069 | 0.049067 | 0.60572 | -0.72327 |
| SMd181/200 | 0.45454 | 0.62165 | 1.1056 | 0.14479 | 0.11415 | 0.5914 | 0.81138 | -0.30155 | 0.81345 | 0.85001 | 1.0234 | 0.033338 |
| TAG541-FA181 | 0.11719 | 0.2333 | 0.51799 | -0.949 | 0.97204 | 0.99185 | 1.0359 | 0.050911 | 0.001446 | 0.023486 | 0.2215 | -2.1746 |
| TAG565-FA203 | 0.025473 | 0.097548 | 0.44 | -1.1844 | 0.32878 | 0.64559 | 0.71019 | -0.49372 | 0.014944 | 0.04422 | 0.36751 | -1.4441 |

| | | | | | | | | | | | | |
|---|---|---|---|---|---|---|---|---|---|---|---|---|
| PG160/160 | 0.67015 | 0.77905 | 0.92763 | -0.10838 | 0.62055 | 0.80957 | 0.94191 | -0.08634 | 0.63475 | 0.70503 | 0.86055 | -0.21667 |
| PEP-160/225 | 0.05361 | 0.13806 | 1.3402 | 0.42245 | 0.20398 | 0.5914 | 1.2119 | 0.27732 | 0.4858 | 0.56677 | 1.0964 | 0.13283 |
| PC170/226 | 0.44441 | 0.61227 | 0.80434 | -0.31412 | 0.99608 | 0.99929 | 0.95241 | -0.07035 | 0.19758 | 0.27841 | 0.70719 | -0.49983 |
| TAG522-FA180 | 0.068649 | 0.16335 | 0.53333 | -0.9069 | 0.33921 | 0.65722 | 0.91498 | -0.12818 | 0.011954 | 0.040533 | 0.37473 | -1.4161 |
| TAG545-FA203 | 0.039284 | 0.1184 | 0.40305 | -1.311 | 0.16908 | 0.5914 | 0.62203 | -0.68495 | 0.034054 | 0.07554 | 0.3676 | -1.4438 |
| PC181/182 | 0.15362 | 0.28737 | 0.78178 | -0.35516 | 0.49746 | 0.73497 | 1.1273 | 0.17282 | 0.089178 | 0.14587 | 0.67505 | -0.56693 |
| TAG548-FA205 | 0.079819 | 0.1798 | 0.31331 | -1.6743 | 0.16241 | 0.5914 | 0.44962 | -1.1532 | 0.15561 | 0.22815 | 0.40779 | -1.2941 |
| TAG562-FA160 | 0.16555 | 0.30358 | 0.54918 | -0.86466 | 0.86321 | 0.93267 | 1.1481 | 0.19927 | 0.009957 | 0.038366 | 0.30704 | -1.7035 |
| PG160/180_2 | 0.65016 | 0.76262 | 1.0793 | 0.11012 | 0.52571 | 0.74605 | 0.81533 | -0.29454 | 0.13116 | 0.19539 | 0.62703 | -0.6734 |
| PEP-160/205 | 0.88249 | 0.92068 | 1.0206 | 0.029404 | 0.92208 | 0.96674 | 1.0363 | 0.051394 | 0.37423 | 0.45967 | 0.7811 | -0.35643 |
| TAG543-FA201 | 0.049873 | 0.13642 | 0.49968 | -1.0009 | 0.42408 | 0.72444 | 0.82318 | -0.28073 | 0.030927 | 0.071649 | 0.43307 | -1.2073 |
| PEP-181/182 | 0.12871 | 0.25161 | 0.78121 | -0.35622 | 0.52222 | 0.74554 | 1.1509 | 0.20281 | 0.017652 | 0.047485 | 0.59829 | -0.74108 |
| PG160/180 | 0.48718 | 0.6467 | 1.2463 | 0.3177 | 0.42879 | 0.72505 | 0.76261 | -0.39098 | 0.12277 | 0.185 | 0.58176 | -0.78152 |
| PEO-160/182 | 0.4402 | 0.61102 | 0.85096 | -0.23283 | 0.59646 | 0.79244 | 0.89247 | -0.16413 | 0.011503 | 0.039833 | 0.50346 | -0.99005 |
| TAG483-FA160 | 0.014723 | 0.088249 | 0.28099 | -1.8314 | 0.10951 | 0.5914 | 0.6118 | -0.70887 | 0.007144 | 0.037206 | 0.2202 | -2.1831 |
| TAG547-FA161 | 0.072747 | 0.16914 | 0.43136 | -1.213 | 0.17992 | 0.5914 | 0.59654 | -0.74531 | 0.088811 | 0.14563 | 0.44552 | -1.1664 |
| TAG565-FA224 | 0.035957 | 0.11378 | 0.49535 | -1.0135 | 0.52192 | 0.74554 | 0.93871 | -0.09125 | 0.091799 | 0.14903 | 0.56079 | -0.83448 |
| TAG502-FA160 | 0.02632 | 0.099043 | 0.48848 | -1.0336 | 0.19903 | 0.5914 | 0.75072 | -0.41366 | 0.096619 | 0.15341 | 0.56137 | -0.83298 |
| PE180/150 | 0.32298 | 0.49357 | 1.1177 | 0.1605 | 0.57592 | 0.77407 | 1.0375 | 0.053125 | 0.33754 | 0.42503 | 0.90567 | -0.14295 |
| LPE180 | 0.006105 | 0.070016 | 1.6897 | 0.7568 | 0.011225 | 0.28651 | 1.4537 | 0.53975 | 0.076713 | 0.13142 | 0.77564 | -0.36654 |
| TAG502-FA182 | 0.03066 | 0.10236 | 0.51065 | -0.96959 | 0.16157 | 0.5914 | 0.71135 | -0.49138 | 0.090615 | 0.14748 | 0.5712 | -0.80792 |
| PEO-180/224 | 0.57053 | 0.70612 | 1.2569 | 0.32991 | 0.97619 | 0.99244 | 1.0484 | 0.068137 | 0.60742 | 0.67943 | 0.8542 | -0.22735 |
| TAG483-FA181 | 0.014513 | 0.088249 | 0.20987 | -2.2524 | 0.27141 | 0.59421 | 0.57015 | -0.81059 | 0.047298 | 0.092952 | 0.33347 | -1.5844 |
| PE182/183 | 0.39015 | 0.55699 | 0.66599 | -0.58642 | 0.008092 | 0.28651 | 0.28089 | -1.8319 | 0.018184 | 0.047927 | 0.29022 | -1.7848 |
| PG160/181 | 0.37043 | 0.54191 | 0.75299 | -0.4093 | 0.65534 | 0.83216 | 0.86544 | -0.20849 | 0.098041 | 0.15491 | 0.61281 | -0.70648 |
| PEP-180/180 | 0.37778 | 0.54831 | 0.87727 | -0.1889 | 0.46096 | 0.73251 | 1.1903 | 0.2513 | 0.000969 | 0.023486 | 0.49715 | -1.0082 |
| PS160/180 | 0.53998 | 0.68562 | 1.1043 | 0.14311 | 0.22315 | 0.5914 | 0.78699 | -0.34559 | 0.39006 | 0.47375 | 0.84629 | -0.24078 |
| PG180/181_2 | 0.21698 | 0.37269 | 0.7482 | -0.41851 | 0.49421 | 0.73497 | 0.86414 | -0.21066 | 0.015812 | 0.045547 | 0.57843 | -0.78978 |
| PC160/160 | 0.018866 | 0.09098 | 1.53 | 0.61349 | 0.30747 | 0.62637 | 1.1404 | 0.18956 | 0.42011 | 0.49907 | 1.1039 | 0.14265 |
| TAG562-FA201 | 0.35172 | 0.5202 | 0.62918 | -0.66846 | 0.47483 | 0.73251 | 1.3215 | 0.40222 | 0.00808 | 0.037908 | 0.26276 | -1.9282 |
| PEO-160/224 | 0.5585 | 0.6992 | 1.2551 | 0.32778 | 0.92334 | 0.96674 | 0.97693 | -0.03368 | 0.39981 | 0.48022 | 0.79462 | -0.33167 |
| PE170/204 | 0.67957 | 0.78578 | 0.83235 | -0.26474 | 0.50392 | 0.73692 | 0.77577 | -0.3663 | 0.023934 | 0.058138 | 0.45605 | -1.1327 |
| TAG566-FA224 | 0.018865 | 0.09098 | 0.42858 | -1.2224 | 0.19616 | 0.5914 | 0.69815 | -0.51839 | 0.056467 | 0.10454 | 0.49109 | -1.0259 |

| | | | | | | | | | | | | |
|---|---|---|---|---|---|---|---|---|---|---|---|---|
| PE182/182 | 0.44486 | 0.61227 | 0.68204 | -0.55207 | 0.49097 | 0.73497 | 0.66498 | -0.58861 | 0.063555 | 0.11304 | 0.41124 | -1.2819 |
| PC181/225 | 0.11637 | 0.2333 | 0.76854 | -0.37981 | 0.24875 | 0.5914 | 0.84936 | -0.23556 | 0.079302 | 0.13374 | 0.74562 | -0.42348 |
| PI161/180 | 0.045402 | 0.12964 | 0.65136 | -0.61847 | 0.1709 | 0.5914 | 0.77947 | -0.35943 | 0.016502 | 0.046306 | 0.63344 | -0.65872 |
| LPE225 | 0.030065 | 0.10164 | 1.5888 | 0.66794 | 0.002864 | 0.28651 | 1.6784 | 0.74707 | 0.72116 | 0.77727 | 1.0631 | 0.088248 |
| HCER200 | 0.071589 | 0.16764 | 1.3179 | 0.39825 | 0.37037 | 0.67841 | 0.84046 | -0.25074 | 0.83661 | 0.86313 | 0.99658 | -0.00494 |
| LPC203 | 0.22127 | 0.37807 | 0.75766 | -0.40037 | 0.86533 | 0.93267 | 0.92991 | -0.10483 | 0.19061 | 0.27093 | 0.73948 | -0.43541 |
| TAG544-FA160 | 0.053156 | 0.13806 | 0.50264 | -0.99241 | 0.31659 | 0.62836 | 0.80927 | -0.3053 | 0.036325 | 0.077637 | 0.44756 | -1.1598 |
| PE180/203 | 0.53272 | 0.68075 | 0.88415 | -0.17764 | 0.23845 | 0.5914 | 0.79781 | -0.32589 | 0.012922 | 0.041839 | 0.60955 | -0.71417 |
| TAG546-FA204 | 0.038838 | 0.11785 | 0.53476 | -0.90303 | 0.20279 | 0.5914 | 0.72203 | -0.46987 | 0.32516 | 0.41262 | 0.75792 | -0.39989 |
| TAG568-FA161 | 0.047245 | 0.132 | 0.35292 | -1.5026 | 0.48572 | 0.73497 | 0.71148 | -0.49111 | 0.028141 | 0.066377 | 0.31197 | -1.6805 |
| PEO-180/182 | 0.26932 | 0.43722 | 0.82201 | -0.28278 | 0.5114 | 0.74148 | 0.8899 | -0.16829 | 0.001436 | 0.023486 | 0.4279 | -1.2246 |
| PS180/180 | 0.46752 | 0.63276 | 1.1263 | 0.17155 | 0.47431 | 0.73251 | 0.85436 | -0.22709 | 0.55348 | 0.63102 | 0.87255 | -0.19669 |
| TAG561-FA160 | 0.14204 | 0.27116 | 0.51289 | -0.96328 | 0.89202 | 0.94886 | 1.178 | 0.23633 | 0.003374 | 0.03071 | 0.22535 | -2.1498 |
| PEP-160/203 | 0.50909 | 0.65785 | 0.89784 | -0.15547 | 0.99505 | 0.99929 | 1.0078 | 0.011179 | 0.19524 | 0.27634 | 0.78311 | -0.35271 |
| TAG563-FA201 | 0.046281 | 0.13146 | 0.46465 | -1.1058 | 0.56049 | 0.76614 | 1.1542 | 0.20684 | 0.00937 | 0.038185 | 0.30492 | -1.7135 |
| PE181/224 | 0.1947 | 0.3435 | 0.73773 | -0.43884 | 0.16937 | 0.5914 | 0.7181 | -0.47774 | 0.024154 | 0.058413 | 0.48397 | -1.047 |
| TAG521-FA180 | 0.088753 | 0.19389 | 0.52126 | -0.93993 | 0.65064 | 0.83052 | 1.0929 | 0.12812 | 0.008899 | 0.037908 | 0.33467 | -1.5792 |
| TAG547-FA205 | 0.16823 | 0.30678 | 0.52183 | -0.93835 | 0.26675 | 0.59421 | 0.67755 | -0.56161 | 0.50119 | 0.5785 | 0.83579 | -0.25879 |
| PG160/181_2 | 0.48095 | 0.64423 | 0.79798 | -0.32558 | 0.74042 | 0.87639 | 0.90281 | -0.14751 | 0.088361 | 0.14526 | 0.61503 | -0.70128 |
| PE170/182 | 0.61354 | 0.74098 | 0.74725 | -0.42033 | 0.16211 | 0.5914 | 0.54043 | -0.88782 | 0.011061 | 0.039348 | 0.31546 | -1.6645 |
| PC200/203 | 0.1107 | 0.22733 | 0.78077 | -0.35703 | 0.003801 | 0.28651 | 0.6223 | -0.68431 | 0.000137 | 0.023486 | 0.48582 | -1.0415 |
| TAG547-FA204 | 0.020143 | 0.09098 | 0.46881 | -1.0929 | 0.11631 | 0.5914 | 0.64805 | -0.62582 | 0.029803 | 0.070043 | 0.44807 | -1.1582 |
| TAG526-FA204 | 0.03892 | 0.11785 | 0.45186 | -1.146 | 0.14165 | 0.5914 | 0.63173 | -0.66261 | 0.008591 | 0.037908 | 0.31773 | -1.6541 |
| TAG542-FA181 | 0.1029 | 0.21409 | 0.53062 | -0.91426 | 0.42638 | 0.72444 | 1.2926 | 0.37026 | 0.005817 | 0.033574 | 0.30015 | -1.7362 |
| PI180/183 | 0.076466 | 0.1759 | 0.65782 | -0.60424 | 0.20279 | 0.5914 | 0.76225 | -0.39166 | 0.004998 | 0.032262 | 0.53004 | -0.91583 |
| TAG520-FA160 | 0.027411 | 0.09969 | 0.47753 | -1.0663 | 0.15316 | 0.5914 | 0.74061 | -0.43321 | 0.000478 | 0.023486 | 0.22936 | -2.1243 |
| TAG521-FA181 | 0.051766 | 0.13699 | 0.49968 | -1.0009 | 0.90239 | 0.95366 | 1.1679 | 0.22397 | 0.010697 | 0.038366 | 0.36926 | -1.4373 |
| TAG544-FA203 | 0.01955 | 0.09098 | 0.43952 | -1.186 | 0.34214 | 0.65897 | 0.79227 | -0.33593 | 0.018543 | 0.04848 | 0.39737 | -1.3314 |
| PEO-180/225 | 0.18631 | 0.33048 | 1.3984 | 0.4838 | 0.27038 | 0.59421 | 1.2779 | 0.35381 | 0.20882 | 0.29044 | 1.2194 | 0.28615 |
| TAG546-FA225 | 0.099873 | 0.21178 | 0.34885 | -1.5193 | 0.24887 | 0.5914 | 0.62211 | -0.68477 | 0.1202 | 0.18325 | 0.38115 | -1.3916 |
| TAG542-FA202 | 0.11791 | 0.23402 | 0.56569 | -0.82193 | 0.49922 | 0.73497 | 0.90286 | -0.14743 | 0.021883 | 0.05396 | 0.41437 | -1.271 |
| TAG564-FA203 | 0.081794 | 0.18308 | 0.53884 | -0.89207 | 0.6963 | 0.85482 | 0.95842 | -0.06128 | 0.060895 | 0.10921 | 0.49721 | -1.0081 |
| PC160/202 | 0.57389 | 0.70879 | 0.91286 | -0.13153 | 0.26256 | 0.59421 | 0.83704 | -0.25664 | 0.092955 | 0.14988 | 0.77283 | -0.37177 |

| | | | | | | | | | | | | |
|---|---|---|---|---|---|---|---|---|---|---|---|---|
| TAG563-FA180 | 0.044053 | 0.1269 | 0.48098 | -1.0559 | 0.38919 | 0.69798 | 0.82461 | -0.27821 | 0.006813 | 0.036355 | 0.31976 | -1.6449 |
| PE180/160 | 0.009957 | 0.079755 | 1.7484 | 0.80604 | 0.76859 | 0.89253 | 0.92218 | -0.11687 | 0.048185 | 0.093917 | 0.65587 | -0.60852 |
| FFA205 | 0.96719 | 0.97922 | 0.99593 | -0.00589 | 0.19456 | 0.5914 | 0.73027 | -0.4535 | 0.26008 | 0.34413 | 0.75138 | -0.41239 |
| PE180/201 | 0.78109 | 0.85893 | 1.0259 | 0.036829 | 0.84769 | 0.92592 | 1.0108 | 0.015535 | 0.007334 | 0.037298 | 0.52687 | -0.92448 |
| PC180/203 | 0.30121 | 0.47136 | 0.83938 | -0.25261 | 0.21134 | 0.5914 | 0.80023 | -0.32152 | 0.21514 | 0.29548 | 0.80462 | -0.31362 |
| PG180/180 | 0.3005 | 0.47136 | 0.80089 | -0.32032 | 0.47213 | 0.73251 | 0.86954 | -0.20168 | 0.014559 | 0.043678 | 0.60346 | -0.72867 |
| PEP-180/204 | 0.50181 | 0.65634 | 0.90301 | -0.14718 | 0.79018 | 0.89932 | 1.0445 | 0.06275 | 0.17645 | 0.25499 | 0.79367 | -0.33339 |
| PEO-180/160 | 0.021209 | 0.09098 | 1.4894 | 0.5747 | 0.06177 | 0.54498 | 1.2949 | 0.37288 | 0.297 | 0.38135 | 0.84975 | -0.23489 |
| TAG541-FA201 | 0.26002 | 0.42745 | 0.56296 | -0.8289 | 0.81974 | 0.90757 | 0.98211 | -0.02605 | 0.022146 | 0.054404 | 0.33461 | -1.5795 |
| PC180/161 | 0.10237 | 0.21409 | 0.75165 | -0.41186 | 0.13593 | 0.5914 | 0.77344 | -0.37063 | 0.033706 | 0.075405 | 0.6295 | -0.66771 |
| TAG568-FA205 | 0.27106 | 0.43896 | 0.68482 | -0.5462 | 0.24295 | 0.5914 | 0.69517 | -0.52457 | 0.075068 | 0.12928 | 0.50475 | -0.98635 |
| PI180/204 | 0.00561 | 0.070016 | 0.72141 | -0.47111 | 0.08084 | 0.5914 | 0.85972 | -0.21806 | 0.63572 | 0.70503 | 0.96082 | -0.05766 |
| PI180/204_2 | 0.014776 | 0.088249 | 0.74652 | -0.42175 | 0.14671 | 0.5914 | 0.85693 | -0.22275 | 0.81836 | 0.85104 | 1.0198 | 0.028286 |
| TAG546-FA205 | 0.15425 | 0.28773 | 0.54205 | -0.88349 | 0.31423 | 0.6275 | 0.73648 | -0.44128 | 0.45159 | 0.53259 | 0.84108 | -0.24969 |
| PC170/182 | 0.22872 | 0.38376 | 0.79815 | -0.32527 | 0.3771 | 0.68573 | 0.8428 | -0.24673 | 0.049534 | 0.094943 | 0.66225 | -0.59455 |
| TAG542-FA201 | 0.051114 | 0.13671 | 0.55032 | -0.86167 | 0.89193 | 0.94886 | 1.1618 | 0.21631 | 0.017336 | 0.047025 | 0.42706 | -1.2275 |
| TAG521-FA160 | 0.1458 | 0.27673 | 0.58842 | -0.76508 | 0.97817 | 0.99244 | 1.2672 | 0.34169 | 0.014806 | 0.044013 | 0.38879 | -1.363 |
| TAG482-FA120 | 0.042529 | 0.12471 | 0.22531 | -2.15 | 0.30014 | 0.62227 | 0.49756 | -1.0071 | 0.051951 | 0.098315 | 0.27752 | -1.8493 |
| TAG564-FA180 | 0.018964 | 0.09098 | 0.41998 | -1.2516 | 0.2893 | 0.61133 | 0.79749 | -0.32645 | 0.058088 | 0.10543 | 0.48582 | -1.0415 |
| LPC180 | 0.23626 | 0.39336 | 0.84199 | -0.24813 | 0.49836 | 0.73497 | 1.0622 | 0.087075 | 0.21498 | 0.29548 | 0.83855 | -0.25403 |
| PC181/205 | 0.75598 | 0.84707 | 0.93021 | -0.10437 | 0.9482 | 0.97817 | 0.99545 | -0.00658 | 0.39448 | 0.47556 | 0.7756 | -0.36662 |
| TAG567-FA205 | 0.363 | 0.53344 | 0.69715 | -0.52047 | 0.56906 | 0.77018 | 0.85881 | -0.21959 | 0.28261 | 0.36715 | 0.68732 | -0.54096 |
| PE181/201 | 0.41645 | 0.58178 | 0.81128 | -0.30172 | 0.15581 | 0.5914 | 1.3819 | 0.46663 | 0.020076 | 0.050853 | 0.4436 | -1.1727 |
| TAG501-FA161 | 0.016778 | 0.090926 | 0.42658 | -1.2291 | 0.43858 | 0.72846 | 1.0741 | 0.10311 | 0.003017 | 0.029739 | 0.30103 | -1.732 |
| PG180/181 | 0.27294 | 0.4409 | 0.76905 | -0.37885 | 0.53548 | 0.74967 | 0.87679 | -0.18969 | 0.017048 | 0.046633 | 0.5791 | -0.78812 |
| PI180/205_2 | 0.52091 | 0.66836 | 0.86619 | -0.20724 | 0.46478 | 0.73251 | 0.83842 | -0.25425 | 0.39702 | 0.47774 | 0.80912 | -0.30558 |
| TAG541-FA180 | 0.20504 | 0.355 | 0.55591 | -0.84708 | 0.66692 | 0.83459 | 1.0198 | 0.028314 | 0.002514 | 0.027699 | 0.21214 | -2.2369 |
| PC200/205 | 0.55049 | 0.69181 | 0.79764 | -0.32619 | 0.59977 | 0.79522 | 0.82075 | -0.28499 | 0.16442 | 0.23838 | 0.57668 | -0.79415 |
| PC150/226 | 0.26105 | 0.42807 | 0.77976 | -0.3589 | 0.44179 | 0.72846 | 0.84319 | -0.24608 | 0.059544 | 0.10738 | 0.62805 | -0.67104 |
| PC200/181 | 0.35807 | 0.52739 | 0.85536 | -0.2254 | 0.12325 | 0.5914 | 1.3009 | 0.37954 | 0.009653 | 0.038317 | 0.47183 | -1.0837 |
| TAG527-FA160 | 0.099047 | 0.21178 | 0.50596 | -0.98291 | 0.23304 | 0.5914 | 0.79643 | -0.32838 | 0.003293 | 0.03071 | 0.26993 | -1.8893 |
| LPC181 | 0.31788 | 0.48691 | 0.87017 | -0.20063 | 0.01041 | 0.28651 | 1.475 | 0.56074 | 0.12015 | 0.18325 | 0.78453 | -0.3501 |
| PE181/181 | 0.76522 | 0.85145 | 1.0503 | 0.070827 | 0.15904 | 0.5914 | 1.6506 | 0.72295 | 0.014011 | 0.043227 | 0.41469 | -1.2699 |

| | | | | | | | | | | | | |
|---|---|---|---|---|---|---|---|---|---|---|---|---|
| SMd181/120 | 0.60593 | 0.73594 | 0.91904 | -0.1218 | 0.38485 | 0.6931 | 0.85379 | -0.22804 | 0.78141 | 0.82715 | 0.92646 | -0.1102 |
| PE160/161 | 0.55153 | 0.69181 | 0.80681 | -0.3097 | 0.11891 | 0.5914 | 0.63129 | -0.66362 | 0.005626 | 0.032995 | 0.3586 | -1.4795 |
| PEO-180/183 | 0.63162 | 0.75218 | 0.90584 | -0.14267 | 0.20226 | 0.5914 | 0.76362 | -0.38907 | 0.001645 | 0.023486 | 0.39944 | -1.324 |
| TAG568-FA182 | 0.16487 | 0.30319 | 0.53283 | -0.90824 | 0.27289 | 0.59421 | 0.67237 | -0.57267 | 0.28815 | 0.37155 | 0.64084 | -0.64196 |
| TAG521-FA201 | 0.13853 | 0.26682 | 0.55127 | -0.85916 | 0.60247 | 0.79674 | 1.089 | 0.12301 | 0.20964 | 0.29044 | 0.64654 | -0.62918 |
| TAG543-FA203 | 0.22577 | 0.38077 | 0.67917 | -0.55816 | 0.29298 | 0.61133 | 0.8373 | -0.25618 | 0.032652 | 0.073551 | 0.46052 | -1.1187 |
| PE181/161 | 0.73032 | 0.82973 | 0.94159 | -0.08683 | 0.46514 | 0.73251 | 1.2348 | 0.30431 | 0.038236 | 0.080296 | 0.60874 | -0.71609 |
| TAG501-FA181 | 0.035671 | 0.11378 | 0.51933 | -0.94528 | 0.4438 | 0.72846 | 0.96586 | -0.05012 | 0.05814 | 0.10543 | 0.52661 | -0.92519 |
| TAG566-FA204 | 0.012733 | 0.087814 | 0.5575 | -0.84295 | 0.37099 | 0.67841 | 0.83199 | -0.26536 | 0.086384 | 0.14273 | 0.63771 | -0.64904 |
| LPC225 | 0.64085 | 0.75579 | 1.106 | 0.14531 | 0.12453 | 0.5914 | 1.3849 | 0.46978 | 0.95265 | 0.96151 | 1.0125 | 0.017913 |
| PE160/183 | 0.34359 | 0.51184 | 1.2032 | 0.26691 | 0.13249 | 0.5914 | 0.52611 | -0.92656 | 0.034819 | 0.076065 | 0.34962 | -1.5161 |
| HCER160 | 0.74283 | 0.8352 | 1.045 | 0.063445 | 0.16543 | 0.5914 | 0.81982 | -0.28663 | 0.095475 | 0.15225 | 0.79078 | -0.33865 |
| TAG547-FA225 | 0.049057 | 0.1359 | 0.27098 | -1.8838 | 0.28144 | 0.60269 | 0.6251 | -0.67785 | 0.085601 | 0.1418 | 0.3197 | -1.6452 |
| TAG564-FA160 | 0.20595 | 0.35563 | 0.61695 | -0.69677 | 0.49226 | 0.73497 | 0.94708 | -0.07844 | 0.017165 | 0.046755 | 0.38918 | -1.3615 |
| PG160/202_2 | 0.62644 | 0.74892 | 0.81501 | -0.29511 | 0.70701 | 0.85829 | 0.85676 | -0.22303 | 0.049732 | 0.094943 | 0.51099 | -0.96865 |
| TAG540-FA180 | 0.002607 | 0.070016 | 0.29636 | -1.7546 | 0.05157 | 0.53349 | 0.51613 | -0.95418 | 8.08E-06 | 0.005258 | 0.10279 | -3.2823 |
| PE180/161 | 0.77852 | 0.85768 | 1.0104 | 0.01498 | 0.35973 | 0.67101 | 0.79704 | -0.32728 | 0.064265 | 0.11369 | 0.58172 | -0.78161 |
| PE180/226 | 0.30199 | 0.47145 | 1.2028 | 0.26642 | 0.92962 | 0.9714 | 0.97125 | -0.04208 | 0.20956 | 0.29044 | 0.77443 | -0.3688 |
| TAG563-FA160 | 0.50649 | 0.65689 | 0.72927 | -0.45547 | 0.88643 | 0.94756 | 1.0868 | 0.12009 | 0.034649 | 0.075947 | 0.43684 | -1.1948 |
| SMd181/226 | 0.45616 | 0.62209 | 1.1167 | 0.15926 | 0.078199 | 0.5914 | 0.80361 | -0.31544 | 0.92515 | 0.94105 | 1.0007 | 0.001031 |
| PEO-180/205 | 0.70225 | 0.80486 | 0.92554 | -0.11163 | 0.77616 | 0.89253 | 0.95487 | -0.06662 | 0.079882 | 0.13438 | 0.65117 | -0.61888 |
| HCER220 | 0.054019 | 0.13845 | 1.3113 | 0.39103 | 0.73812 | 0.87526 | 1.0475 | 0.066967 | 0.77096 | 0.81742 | 0.95669 | -0.06387 |
| TAG545-FA204 | 0.053653 | 0.13806 | 0.61011 | -0.71286 | 0.43103 | 0.72695 | 0.89275 | -0.16367 | 0.24502 | 0.32888 | 0.74779 | -0.4193 |
| SMd181/220 | 0.86766 | 0.91252 | 1.0192 | 0.027445 | 0.014636 | 0.30735 | 0.72792 | -0.45815 | 0.58281 | 0.65415 | 0.93924 | -0.09044 |
| TAG567-FA180 | 0.090966 | 0.1974 | 0.45873 | -1.1243 | 0.24196 | 0.5914 | 0.74622 | -0.42232 | 0.14456 | 0.21437 | 0.54099 | -0.88633 |
| PC160/182 | 0.49224 | 0.65131 | 1.0594 | 0.083285 | 0.35664 | 0.67101 | 1.0941 | 0.1297 | 0.54369 | 0.62313 | 0.90196 | -0.14887 |
| TAG567-FA225 | 0.062153 | 0.15211 | 0.38453 | -1.3788 | 0.23527 | 0.5914 | 0.6733 | -0.57067 | 0.21312 | 0.29394 | 0.55336 | -0.85371 |
| TAG545-FA160 | 0.11678 | 0.2333 | 0.53154 | -0.91174 | 0.31218 | 0.6275 | 0.7836 | -0.35181 | 0.2826 | 0.36715 | 0.69315 | -0.52876 |
| PEP-181/225 | 0.11703 | 0.2333 | 1.3193 | 0.39982 | 0.078823 | 0.5914 | 1.3845 | 0.46933 | 0.14646 | 0.2167 | 1.1924 | 0.25384 |
| PEP-182/182 | 0.28893 | 0.45433 | 0.85817 | -0.22066 | 0.23785 | 0.5914 | 1.2483 | 0.31996 | 0.03885 | 0.080546 | 0.70795 | -0.49828 |
| TAG546-FA226 | 0.75775 | 0.84758 | 0.96113 | -0.05719 | 0.88622 | 0.94756 | 1.2232 | 0.29068 | 0.71648 | 0.7748 | 1.9385 | 0.95492 |
| PE160/226 | 0.28269 | 0.45106 | 1.242 | 0.31271 | 0.8653 | 0.93267 | 0.95119 | -0.0722 | 0.25314 | 0.33839 | 0.77953 | -0.35931 |
| FFA203 | 0.19413 | 0.34343 | 0.82542 | -0.2768 | 0.19286 | 0.5914 | 0.81433 | -0.29631 | 0.027697 | 0.066047 | 0.71368 | -0.48665 |

| | | | | | | | | | | | | |
|---|---|---|---|---|---|---|---|---|---|---|---|---|
| LPE202 | 0.1027 | 0.21409 | 1.4339 | 0.51994 | 0.012807 | 0.29776 | 1.364 | 0.44779 | 0.010582 | 0.038366 | 0.61696 | -0.69675 |
| SMd181/205 | 0.99376 | 0.99376 | 0.99662 | -0.00488 | 0.053655 | 0.53471 | 0.78808 | -0.34359 | 0.71423 | 0.77386 | 0.95379 | -0.06826 |
| TAG566-FA160 | 0.060932 | 0.14969 | 0.43156 | -1.2124 | 0.3484 | 0.66513 | 0.76437 | -0.38765 | 0.10987 | 0.1703 | 0.49277 | -1.021 |
| PG160/202 | 0.57487 | 0.70879 | 0.81359 | -0.29763 | 0.69878 | 0.85482 | 0.8663 | -0.20706 | 0.032566 | 0.073551 | 0.50606 | -0.98261 |
| TAG547-FA226 | 0.17767 | 0.3195 | 0.44773 | -1.1593 | 0.60337 | 0.79674 | 1.0916 | 0.1264 | 0.2432 | 0.32779 | 0.56506 | -0.82351 |
| SM241 | 0.90206 | 0.93213 | 0.9933 | -0.0097 | 0.031578 | 0.4227 | 0.79915 | -0.32346 | 0.35697 | 0.4418 | 0.92148 | -0.11797 |
| PEP-181/205 | 0.62697 | 0.74892 | 0.90468 | -0.14452 | 0.96168 | 0.98589 | 0.9993 | -0.00101 | 0.12942 | 0.19324 | 0.71426 | -0.48547 |
| HCER241 | 0.53768 | 0.68498 | 1.07 | 0.097587 | 0.25262 | 0.59157 | 0.84344 | -0.24564 | 0.23204 | 0.31536 | 0.84317 | -0.24611 |
| FFA226 | 0.4967 | 0.65369 | 1.1034 | 0.14191 | 0.51378 | 0.74163 | 1.1316 | 0.17831 | 0.79824 | 0.84086 | 1.0127 | 0.018233 |
| FFA220 | 0.47166 | 0.63543 | 1.0708 | 0.098691 | 0.64292 | 0.828 | 0.91391 | -0.12988 | 0.035918 | 0.077637 | 0.59614 | -0.74628 |
| PEP-180/160 | 0.3949 | 0.55887 | 1.3132 | 0.3931 | 0.89524 | 0.94919 | 1.0285 | 0.04048 | 0.75759 | 0.80851 | 0.94081 | -0.08803 |
| PE160/201 | 0.56309 | 0.70359 | 1.0538 | 0.075574 | 0.74339 | 0.8783 | 0.85079 | -0.23313 | 0.025309 | 0.060798 | 0.4856 | -1.0422 |
| TAG565-FA180 | 0.040862 | 0.12202 | 0.51412 | -0.95982 | 0.55933 | 0.76614 | 0.94408 | -0.08302 | 0.23918 | 0.32371 | 0.71693 | -0.4801 |
| PG161/182_2 | 0.79963 | 0.87196 | 1.0143 | 0.020482 | 0.14746 | 0.5914 | 1.3945 | 0.47978 | 0.11929 | 0.18311 | 0.69622 | -0.52238 |
| TAG566-FA181 | 0.022927 | 0.091642 | 0.41828 | -1.2574 | 0.47773 | 0.73497 | 0.81419 | -0.29657 | 0.039489 | 0.081612 | 0.43326 | -1.2067 |
| PC140/225 | 0.23615 | 0.39336 | 0.82528 | -0.27704 | 0.83364 | 0.91828 | 1.0043 | 0.006224 | 0.23656 | 0.32083 | 0.81546 | -0.29431 |
| PC180/181 | 0.74208 | 0.8352 | 0.94079 | -0.08806 | 0.54598 | 0.75947 | 1.0538 | 0.075582 | 0.083596 | 0.13918 | 0.72558 | -0.46279 |
| TAG501-FA160 | 0.025637 | 0.0976 | 0.51611 | -0.95424 | 0.46075 | 0.73251 | 0.98928 | -0.01555 | 0.082399 | 0.13825 | 0.57927 | -0.78768 |
| PC160/224 | 0.45677 | 0.62209 | 1.0765 | 0.10641 | 0.86458 | 0.93267 | 1.0365 | 0.051663 | 0.33455 | 0.42208 | 0.84351 | -0.24552 |
| TAG481-FA161 | 0.036832 | 0.11528 | 0.50414 | -0.98811 | 0.25126 | 0.5914 | 0.95834 | -0.06139 | 0.016793 | 0.046417 | 0.41832 | -1.2573 |
| PEP-181/183 | 0.064864 | 0.15639 | 0.72811 | -0.45777 | 0.49624 | 0.73497 | 0.90723 | -0.14046 | 0.001491 | 0.023486 | 0.5191 | -0.94592 |
| PEO-160/203 | 0.2843 | 0.45252 | 0.83135 | -0.26648 | 0.21578 | 0.5914 | 0.8209 | -0.28472 | 0.014371 | 0.043513 | 0.62135 | -0.68651 |
| PC181/201 | 0.32784 | 0.49802 | 0.81789 | -0.29003 | 0.4898 | 0.73497 | 1.1332 | 0.1804 | 0.12166 | 0.18419 | 0.67856 | -0.55946 |
| TAG548-FA204 | 0.1278 | 0.25059 | 0.61464 | -0.70218 | 0.072826 | 0.5644 | 0.56481 | -0.82417 | 0.011303 | 0.039348 | 0.43007 | -1.2174 |
| PEP-181/203 | 0.34101 | 0.51173 | 0.83021 | -0.26845 | 0.67049 | 0.83459 | 0.92322 | -0.11526 | 0.096983 | 0.15362 | 0.68819 | -0.53912 |
| PE180/180 | 0.13923 | 0.26738 | 1.2587 | 0.33199 | 0.12729 | 0.5914 | 1.286 | 0.36294 | 0.21742 | 0.29735 | 1.2059 | 0.27016 |
| TAG565-FA160 | 0.13116 | 0.25565 | 0.5779 | -0.79111 | 0.53269 | 0.74837 | 0.94243 | -0.08555 | 0.077511 | 0.13217 | 0.52177 | -0.9385 |
| SMd181/222 | 0.8511 | 0.90534 | 0.9807 | -0.02812 | 0.12954 | 0.5914 | 0.7859 | -0.34758 | 0.78517 | 0.82978 | 0.96042 | -0.05826 |
| PC181/204 | 0.45015 | 0.61694 | 0.85263 | -0.23 | 0.89134 | 0.94886 | 0.95111 | -0.07232 | 0.36393 | 0.44806 | 0.82806 | -0.27219 |
| TAG481-FA180 | 0.022357 | 0.091642 | 0.25178 | -1.9897 | 0.08411 | 0.5914 | 0.50326 | -0.99063 | 0.008909 | 0.037908 | 0.27235 | -1.8765 |
| PC181/181 | 0.38381 | 0.55103 | 0.86068 | -0.21645 | 0.030074 | 0.4227 | 1.82 | 0.86391 | 0.063218 | 0.11287 | 0.63916 | -0.64576 |
| HCER140 | 0.28729 | 0.45321 | 0.88153 | -0.18192 | 0.001015 | 0.28651 | 0.61204 | -0.70831 | 0.001408 | 0.023486 | 0.63848 | -0.64729 |
| TAG542-FA180 | 0.15249 | 0.28643 | 0.58378 | -0.7765 | 0.78923 | 0.89932 | 1.2198 | 0.28669 | 0.004172 | 0.032129 | 0.29358 | -1.7682 |

| | | | | | | | | | | | | |
|---|---|---|---|---|---|---|---|---|---|---|---|---|
| TAG545-FA225 | 0.84197 | 0.90004 | 0.91459 | -0.1288 | 0.69977 | 0.85482 | 1.0409 | 0.057767 | 0.96171 | 0.96812 | 1.391 | 0.47608 |
| PE170/225 | 0.99156 | 0.99308 | 0.94007 | -0.08917 | 0.37056 | 0.67841 | 0.7324 | -0.4493 | 0.015117 | 0.044502 | 0.43453 | -1.2025 |
| PEO-180/203 | 0.65646 | 0.76587 | 0.95314 | -0.06924 | 0.52164 | 0.74554 | 0.93141 | -0.10251 | 0.002553 | 0.027699 | 0.5812 | -0.78289 |
| PA180/181 | 0.76309 | 0.85145 | 1.1001 | 0.13761 | 0.026408 | 0.3998 | 1.8594 | 0.89485 | 0.14891 | 0.21982 | 0.73992 | -0.43455 |
| PEP-160/202 | 0.57849 | 0.71191 | 0.91977 | -0.12065 | 0.5334 | 0.74837 | 0.90479 | -0.14434 | 0.092978 | 0.14988 | 0.67886 | -0.55881 |
| SMd181/224 | 0.9597 | 0.97588 | 1.033 | 0.046908 | 0.099704 | 0.5914 | 0.76294 | -0.39036 | 0.9382 | 0.95284 | 0.98801 | -0.01741 |
| FFA183 | 0.02133 | 0.09098 | 0.61269 | -0.70678 | 0.48463 | 0.73497 | 0.87655 | -0.19009 | 0.058113 | 0.10543 | 0.66133 | -0.59656 |
| TAG569-FA226 | 0.17019 | 0.30948 | 0.37012 | -1.4339 | 0.40456 | 0.70798 | 0.75114 | -0.41285 | 0.1868 | 0.26727 | 0.39177 | -1.3519 |
| SMd181/184 | 0.22323 | 0.37845 | 0.86433 | -0.21035 | 0.031469 | 0.4227 | 0.76403 | -0.3883 | 0.001249 | 0.023486 | 0.58325 | -0.7778 |
| PE180/183 | 0.19809 | 0.34554 | 1.325 | 0.40598 | 0.25889 | 0.59421 | 0.70741 | -0.49939 | 0.007403 | 0.037326 | 0.3768 | -1.4081 |
| HCER240 | 0.16268 | 0.30002 | 1.2217 | 0.28893 | 0.70896 | 0.85829 | 1.0473 | 0.066733 | 0.38132 | 0.46487 | 0.90468 | -0.14452 |
| PEP-180/161 | 0.38247 | 0.55103 | 0.82903 | -0.2705 | 0.84551 | 0.92509 | 0.96343 | -0.05375 | 0.011182 | 0.039348 | 0.48361 | -1.0481 |
| PEP-180/201 | 0.54028 | 0.68562 | 0.87293 | -0.19606 | 0.77051 | 0.89253 | 0.92392 | -0.11416 | 0.079091 | 0.13374 | 0.60445 | -0.7263 |
| PEP-160/226 | 0.11179 | 0.22781 | 1.2599 | 0.33329 | 0.17302 | 0.5914 | 1.2286 | 0.29698 | 0.095262 | 0.15225 | 1.2633 | 0.33721 |
| SMd181/181 | 0.94381 | 0.96901 | 1.0088 | 0.012709 | 0.44101 | 0.72846 | 0.87461 | -0.19329 | 0.36409 | 0.44806 | 1.1057 | 0.14492 |
| LPC205 | 0.60023 | 0.7342 | 1.2414 | 0.31194 | 0.4833 | 0.73497 | 1.2624 | 0.33615 | 0.6595 | 0.724 | 1.1986 | 0.2614 |
| PEP-160/204 | 0.77166 | 0.85433 | 1.0487 | 0.068665 | 0.59536 | 0.79244 | 1.0986 | 0.13564 | 0.6672 | 0.73123 | 0.91358 | -0.13039 |
| PE170/202 | 0.5065 | 0.65689 | 0.91412 | -0.12954 | 0.29141 | 0.61133 | 0.86172 | -0.21472 | 0.010566 | 0.038366 | 0.67157 | -0.57439 |
| HCER260 | 0.22307 | 0.37845 | 0.80855 | -0.3066 | 0.17782 | 0.5914 | 0.78405 | -0.35098 | 0.18349 | 0.26369 | 0.76572 | -0.38511 |
| PE160/160 | 0.008468 | 0.077743 | 1.6722 | 0.74176 | 0.49776 | 0.73497 | 0.8994 | -0.15296 | 0.21801 | 0.29754 | 0.83917 | -0.25296 |
| TAG566-FA205 | 0.48346 | 0.64579 | 0.74865 | -0.41764 | 0.71902 | 0.86045 | 1.0369 | 0.05233 | 0.35655 | 0.4418 | 0.77817 | -0.36184 |
| TAG567-FA160 | 0.1056 | 0.21893 | 0.3979 | -1.3295 | 0.4797 | 0.73497 | 0.76253 | -0.39113 | 0.26883 | 0.35356 | 0.5688 | -0.81401 |
| PEP-180/225 | 0.40521 | 0.57221 | 1.1566 | 0.20995 | 0.46853 | 0.73251 | 1.1332 | 0.18037 | 0.90023 | 0.92146 | 0.97156 | -0.04163 |
| PE180/204 | 0.37817 | 0.54831 | 1.1312 | 0.17791 | 0.96938 | 0.99068 | 0.98708 | -0.01876 | 0.1587 | 0.23113 | 0.7857 | -0.34794 |
| PC160/181 | 0.30689 | 0.47682 | 1.1009 | 0.13866 | 0.057165 | 0.53695 | 1.237 | 0.30682 | 0.62575 | 0.69635 | 0.92984 | -0.10495 |
| LPE200 | 0.84601 | 0.90287 | 1.01 | 0.014344 | 0.018534 | 0.33515 | 1.6417 | 0.71517 | 0.001467 | 0.023486 | 0.58955 | -0.76232 |
| PC180/226 | 0.95632 | 0.97581 | 0.96141 | -0.05677 | 0.83634 | 0.91969 | 1.0183 | 0.026127 | 0.49828 | 0.57616 | 0.87247 | -0.19683 |
| SMd181/180 | 0.89566 | 0.92699 | 1.018 | 0.025782 | 0.064662 | 0.54669 | 0.78801 | -0.34372 | 0.85411 | 0.87979 | 0.97621 | -0.03473 |
| PS160/180_2 | 0.54654 | 0.68819 | 1.1251 | 0.17007 | 0.2148 | 0.5914 | 0.81121 | -0.30186 | 0.24018 | 0.32439 | 0.83077 | -0.26748 |
| HCER181 | 0.86137 | 0.91031 | 0.99054 | -0.01371 | 0.22054 | 0.5914 | 0.8837 | -0.17837 | 0.063285 | 0.11287 | 0.80574 | -0.31161 |
| TAG566-FA225 | 0.066512 | 0.15978 | 0.44917 | -1.1547 | 0.66716 | 0.83459 | 0.93856 | -0.09148 | 0.20969 | 0.29044 | 0.6025 | -0.73096 |
| PC150/225 | 0.85819 | 0.90843 | 0.94085 | -0.08796 | 0.46738 | 0.73251 | 0.81487 | -0.29535 | 0.25853 | 0.34277 | 0.74522 | -0.42427 |
| TAG546-FA160 | 0.34205 | 0.51184 | 0.6568 | -0.60648 | 0.403 | 0.70798 | 0.87559 | -0.19168 | 0.78976 | 0.83328 | 1.078 | 0.10837 |

| | | | | | | | | | | | | |
|---|---|---|---|---|---|---|---|---|---|---|---|---|
| TAG567-FA204 | 0.040435 | 0.12131 | 0.65811 | -0.60359 | 0.26404 | 0.59421 | 0.79864 | -0.32439 | 0.009091 | 0.038182 | 0.52906 | -0.91851 |
| LPC182 | 0.79528 | 0.87013 | 0.95222 | -0.07063 | 0.18314 | 0.5914 | 1.2365 | 0.30624 | 0.25424 | 0.33916 | 0.81011 | -0.30382 |
| PC181/224 | 0.32819 | 0.49802 | 0.83401 | -0.26187 | 0.76343 | 0.89227 | 0.93737 | -0.09331 | 0.20389 | 0.28606 | 0.76662 | -0.38341 |
| PEO-180/226 | 0.37278 | 0.54412 | 1.2454 | 0.31663 | 0.62811 | 0.81455 | 1.1048 | 0.14385 | 0.35641 | 0.4418 | 1.1387 | 0.18743 |
| SMd181/221 | 0.88615 | 0.92154 | 1.0202 | 0.028896 | 0.31545 | 0.628 | 0.85677 | -0.22302 | 0.15299 | 0.22482 | 1.2177 | 0.28421 |
| SM240 | 0.70608 | 0.8066 | 1.0472 | 0.066599 | 0.007381 | 0.28651 | 0.68929 | -0.53682 | 0.3804 | 0.46461 | 0.90984 | -0.13632 |
| DAG161/182 | 0.73321 | 0.83156 | 0.46999 | -1.0893 | 0.46403 | 0.73251 | 0.82421 | -0.27891 | 0.8179 | 0.85104 | 0.60163 | -0.73306 |
| PEO-180/181 | 0.848 | 0.90351 | 1.0522 | 0.073425 | 0.58436 | 0.78276 | 1.1004 | 0.13807 | 0.001205 | 0.023486 | 0.4946 | -1.0157 |
| PC170/224 | 0.56692 | 0.70433 | 0.87372 | -0.19477 | 0.9424 | 0.97682 | 1.0087 | 0.012459 | 0.15736 | 0.2302 | 0.70017 | -0.51421 |
| TAG520-FA180 | 0.009195 | 0.077743 | 0.44077 | -1.1819 | 0.09468 | 0.5914 | 0.68319 | -0.54964 | 0.000149 | 0.023486 | 0.2211 | -2.1772 |
| TAG565-FA204 | 0.036003 | 0.11378 | 0.54704 | -0.87029 | 0.83891 | 0.92096 | 1.0835 | 0.11572 | 0.33215 | 0.42068 | 0.82281 | -0.28138 |
| PE160/202 | 0.11198 | 0.22781 | 1.3873 | 0.47226 | 0.15472 | 0.5914 | 0.65647 | -0.6072 | 0.0904 | 0.14748 | 0.58604 | -0.77093 |
| PC140/204 | 0.69333 | 0.79745 | 0.91507 | -0.12804 | 0.42702 | 0.72444 | 1.1491 | 0.20045 | 0.3334 | 0.42144 | 0.81191 | -0.30061 |
| PEP-181/181 | 0.67408 | 0.78222 | 0.93095 | -0.10322 | 0.22686 | 0.5914 | 1.4176 | 0.50348 | 0.12125 | 0.18399 | 0.69609 | -0.52264 |
| TAG568-FA160 | 0.22252 | 0.37845 | 0.46914 | -1.0919 | 0.50744 | 0.73737 | 0.80095 | -0.32022 | 0.35018 | 0.43672 | 0.64973 | -0.62208 |
| FFA120 | 0.19722 | 0.34514 | 0.72184 | -0.47024 | 0.45822 | 0.73251 | 0.85908 | -0.21914 | 0.28822 | 0.37155 | 0.85465 | -0.2266 |
| DAG140/160 | 0.38329 | 0.55103 | 0.781 | -0.35661 | 0.81029 | 0.90726 | 1.0242 | 0.034547 | 0.76529 | 0.81273 | 1.0514 | 0.072305 |
| TAG567-FA181 | 0.19852 | 0.34554 | 0.60405 | -0.72726 | 0.66838 | 0.83459 | 0.87986 | -0.18465 | 0.24646 | 0.33014 | 0.65956 | -0.60042 |
| CER160 | 0.95267 | 0.97361 | 1.0039 | 0.005601 | 0.66928 | 0.83459 | 1.0862 | 0.1193 | 0.35214 | 0.43832 | 0.83292 | -0.26376 |
| TAG482-FA180 | 0.36586 | 0.53642 | 0.32333 | -1.6289 | 0.23055 | 0.5914 | 0.31311 | -1.6752 | 0.055033 | 0.10311 | 0.17906 | -2.4815 |
| LPC221 | 0.15526 | 0.28796 | 0.77403 | -0.36954 | 0.90638 | 0.95633 | 0.98108 | -0.02756 | 0.036534 | 0.077724 | 0.69849 | -0.51769 |
| PEP-181/161 | 0.71588 | 0.81617 | 0.98667 | -0.01936 | 0.69987 | 0.85482 | 0.9722 | -0.04068 | 0.088158 | 0.14526 | 0.73099 | -0.45207 |
| PC160/180 | 0.42908 | 0.59814 | 1.0757 | 0.10527 | 0.26327 | 0.59421 | 1.1241 | 0.16882 | 0.72007 | 0.77727 | 0.94063 | -0.0883 |
| PC170/225 | 0.50655 | 0.65689 | 0.89367 | -0.16219 | 0.53299 | 0.74837 | 0.89639 | -0.15781 | 0.049473 | 0.094943 | 0.68296 | -0.55014 |
| FFA225 | 0.86451 | 0.91215 | 0.95356 | -0.06861 | 0.71625 | 0.85871 | 1.0873 | 0.12068 | 0.98384 | 0.98688 | 0.97128 | -0.04203 |
| FFA202 | 0.31208 | 0.4821 | 0.86324 | -0.21216 | 0.82999 | 0.91581 | 1.0509 | 0.071639 | 0.28255 | 0.36715 | 0.84312 | -0.24619 |
| PEP-181/160 | 0.49432 | 0.65274 | 1.2875 | 0.3646 | 0.42516 | 0.72444 | 1.2339 | 0.30324 | 0.8602 | 0.88466 | 0.96419 | -0.05261 |
| PEP-160/224 | 0.54523 | 0.68788 | 1.1883 | 0.24892 | 0.94381 | 0.97682 | 1.0216 | 0.030866 | 0.69925 | 0.75995 | 0.90528 | -0.14357 |
| PEO-160/226 | 0.51365 | 0.66215 | 1.134 | 0.18141 | 0.85586 | 0.93016 | 1.0436 | 0.061569 | 0.65481 | 0.72007 | 1.0399 | 0.056393 |
| PC160/161 | 0.77048 | 0.85433 | 0.94253 | -0.08539 | 0.76866 | 0.89253 | 0.97509 | -0.03639 | 0.41293 | 0.49324 | 0.85526 | -0.22557 |
| SMd181/141 | 0.56426 | 0.70371 | 0.87939 | -0.18542 | 0.43653 | 0.72846 | 0.84256 | -0.24715 | 0.94457 | 0.95632 | 1.0046 | 0.006565 |
| SMd181/170 | 0.89445 | 0.92699 | 0.99909 | -0.00131 | 0.022502 | 0.3649 | 0.7749 | -0.36792 | 0.98739 | 0.98891 | 0.99569 | -0.00623 |
| PEO-180/204 | 0.8818 | 0.92068 | 0.98217 | -0.02596 | 0.69034 | 0.85482 | 1.0829 | 0.11486 | 0.044114 | 0.088045 | 0.71679 | -0.48038 |

| | | | | | | | | | | | | |
|---|---|---|---|---|---|---|---|---|---|---|---|---|
| FFA140 | 0.059611 | 0.14812 | 0.74219 | -0.43013 | 0.84531 | 0.92509 | 1.0297 | 0.042184 | 0.39404 | 0.47556 | 0.89354 | -0.1624 |
| SMd181/140 | 0.43649 | 0.60716 | 0.90832 | -0.13872 | 0.062374 | 0.54498 | 0.77317 | -0.37113 | 0.16316 | 0.23709 | 0.83237 | -0.26471 |
| CER180 | 0.37512 | 0.54632 | 1.2698 | 0.34455 | 0.45029 | 0.72879 | 0.84233 | -0.24754 | 0.55632 | 0.63316 | 0.88183 | -0.18143 |
| TAG522-FA202 | 0.23776 | 0.39384 | 0.65138 | -0.61842 | 0.20668 | 0.5914 | 0.80304 | -0.31647 | 0.049105 | 0.094905 | 0.47515 | -1.0735 |
| PE160/140 | 0.60225 | 0.7342 | 1.1428 | 0.19259 | 0.7255 | 0.86502 | 1.0896 | 0.12383 | 0.24496 | 0.32888 | 0.80091 | -0.3203 |
| PEP-160/181 | 0.97065 | 0.9803 | 1.0147 | 0.021093 | 0.31213 | 0.6275 | 1.3337 | 0.41546 | 0.083481 | 0.13918 | 0.70675 | -0.50073 |
| TAG504-FA141 | 0.009157 | 0.077743 | 0.46346 | -1.1095 | 0.521 | 0.74554 | 0.85779 | -0.2213 | 0.003009 | 0.029739 | 0.33225 | -1.5897 |
| FFA204 | 0.18399 | 0.32726 | 0.78193 | -0.3549 | 0.055751 | 0.53695 | 0.68801 | -0.5395 | 0.0555 | 0.10353 | 0.67929 | -0.55789 |
| CE182 | 0.33237 | 0.50117 | 1.1707 | 0.22743 | 0.012755 | 0.29776 | 0.63866 | -0.64688 | 0.47383 | 0.5578 | 0.89225 | -0.16448 |
| PEP-182/226 | 0.18037 | 0.32348 | 1.1951 | 0.25709 | 0.24125 | 0.5914 | 1.1847 | 0.24456 | 0.58578 | 0.65635 | 1.0506 | 0.071174 |
| CER221 | 0.89097 | 0.92507 | 0.99956 | -0.00063 | 0.75788 | 0.89057 | 0.91054 | -0.1352 | 0.048945 | 0.094905 | 0.63994 | -0.644 |
| TAG500-FA160 | 0.17206 | 0.3114 | 0.62535 | -0.67727 | 0.24233 | 0.5914 | 0.77342 | -0.37068 | 0.031714 | 0.072441 | 0.45037 | -1.1508 |
| TAG523-FA203 | 0.40781 | 0.5734 | 0.72748 | -0.45902 | 0.20547 | 0.5914 | 0.73672 | -0.44081 | 0.17665 | 0.25499 | 0.6328 | -0.66019 |
| PA180/203 | 0.61317 | 0.74098 | 0.96407 | -0.0528 | 0.004247 | 0.28651 | 1.524 | 0.6079 | 0.6792 | 0.74188 | 0.92592 | -0.11103 |
| CER260 | 0.63202 | 0.75218 | 0.95442 | -0.06731 | 0.14329 | 0.5914 | 1.2819 | 0.35824 | 0.013129 | 0.041897 | 0.5961 | -0.74639 |
| TAG526-FA226 | 0.38931 | 0.55699 | 0.47191 | -1.0834 | 0.4955 | 0.73497 | 0.95391 | -0.06807 | 0.41261 | 0.49324 | 0.61455 | -0.70241 |
| TAG566-FA180 | 0.1722 | 0.3114 | 0.62078 | -0.68784 | 0.4971 | 0.73497 | 0.97747 | -0.03287 | 0.27757 | 0.36285 | 0.75076 | -0.41357 |
| LPE203 | 0.65356 | 0.76523 | 1.0875 | 0.12105 | 0.072223 | 0.5644 | 1.2777 | 0.35355 | 0.031557 | 0.072338 | 0.67809 | -0.56046 |
| PE160/182 | 0.58644 | 0.72032 | 1.0751 | 0.10452 | 0.64639 | 0.828 | 0.8258 | -0.27613 | 0.043283 | 0.0867 | 0.5163 | -0.95371 |
| TAG500-FA180 | 0.18167 | 0.32414 | 0.62419 | -0.67994 | 0.081221 | 0.5914 | 0.5599 | -0.83676 | 0.016151 | 0.045884 | 0.37818 | -1.4029 |
| PE160/205 | 0.47242 | 0.63543 | 1.275 | 0.35052 | 0.89347 | 0.94886 | 0.95703 | -0.06337 | 0.8127 | 0.85001 | 0.86813 | -0.20402 |
| PEP-181/224 | 0.88395 | 0.92072 | 1.0366 | 0.051843 | 0.99876 | 0.99929 | 1.0021 | 0.003034 | 0.73326 | 0.78901 | 0.9048 | -0.14432 |
| SM261 | 0.38428 | 0.55103 | 0.90664 | -0.1414 | 0.002122 | 0.28651 | 0.64371 | -0.63551 | 0.27497 | 0.36017 | 0.87552 | -0.19179 |
| PC160/140 | 0.41425 | 0.58084 | 1.1464 | 0.19706 | 0.9407 | 0.97682 | 0.97426 | -0.03763 | 0.25791 | 0.34277 | 0.78728 | -0.34505 |
| PC170/205 | 0.81163 | 0.88017 | 1.1248 | 0.16967 | 0.99929 | 0.99929 | 1.0645 | 0.090195 | 0.6518 | 0.71797 | 0.8348 | -0.2605 |
| HCER221 | 0.94668 | 0.96901 | 0.99768 | -0.00335 | 0.06957 | 0.55914 | 0.81864 | -0.2887 | 0.07307 | 0.12651 | 0.81565 | -0.29397 |
| SMd181/201 | 0.82566 | 0.88859 | 0.9526 | -0.07006 | 0.32953 | 0.64559 | 0.82726 | -0.27358 | 0.20953 | 0.29044 | 1.2081 | 0.2727 |
| LPE240 | 0.71724 | 0.8163 | 0.94523 | -0.08126 | 0.25147 | 0.5914 | 1.234 | 0.30332 | 0.006468 | 0.034798 | 0.59722 | -0.74366 |
| HCER261 | 0.26631 | 0.43341 | 0.82291 | -0.28119 | 0.11986 | 0.5914 | 0.76275 | -0.39071 | 0.044459 | 0.088045 | 0.70309 | -0.50821 |
| PC180/225 | 0.97277 | 0.9803 | 0.98347 | -0.02405 | 0.5013 | 0.73502 | 1.0984 | 0.13537 | 0.4809 | 0.56409 | 0.87864 | -0.18666 |
| SMd181/223 | 0.80543 | 0.87535 | 0.96605 | -0.04983 | 0.094086 | 0.5914 | 0.72973 | -0.45457 | 0.41925 | 0.49896 | 0.86782 | -0.20453 |
| SMd181/225 | 0.38379 | 0.55103 | 0.83724 | -0.25629 | 0.16832 | 0.5914 | 0.74815 | -0.4186 | 0.82165 | 0.8531 | 0.97962 | -0.0297 |
| PE160/204 | 0.59053 | 0.72399 | 1.1049 | 0.14386 | 0.94962 | 0.97817 | 0.95955 | -0.05957 | 0.2676 | 0.35264 | 0.73375 | -0.44663 |

| | | | | | | | | | | | | |
|---|---|---|---|---|---|---|---|---|---|---|---|---|
| TAG545-FA205 | 0.44116 | 0.61105 | 0.8337 | -0.2624 | 0.10261 | 0.5914 | 0.66265 | -0.59367 | 0.17819 | 0.25664 | 0.6889 | -0.53762 |
| PEO-160/204 | 0.8011 | 0.8721 | 1.0511 | 0.071968 | 0.9346 | 0.9746 | 1.0164 | 0.023417 | 0.22574 | 0.30744 | 0.79476 | -0.33142 |
| PEO-160/183 | 0.51587 | 0.6637 | 0.88019 | -0.18411 | 0.017367 | 0.33253 | 0.58029 | -0.78516 | 0.041908 | 0.085452 | 0.57649 | -0.79463 |
| DAG160/181 | 0.029557 | 0.10127 | 0.68575 | -0.54424 | 0.28741 | 0.60947 | 1.8815 | 0.91188 | 0.020388 | 0.051445 | 0.57953 | -0.78704 |
| TAG566-FA226 | 0.94656 | 0.96901 | 0.86354 | -0.21167 | 0.81096 | 0.90726 | 1.0922 | 0.12725 | 0.56826 | 0.63944 | 0.93292 | -0.10017 |
| PC160/183 | 0.27674 | 0.44519 | 1.2075 | 0.27206 | 0.42732 | 0.72444 | 0.82572 | -0.27628 | 0.08391 | 0.13935 | 0.59333 | -0.7531 |
| PC180/182 | 0.77863 | 0.85768 | 0.94605 | -0.08001 | 0.87828 | 0.9435 | 0.9959 | -0.00592 | 0.14097 | 0.20953 | 0.79332 | -0.33403 |
| FFA161 | 0.11698 | 0.2333 | 0.70576 | -0.50275 | 0.39999 | 0.70794 | 1.2812 | 0.35748 | 0.9672 | 0.97168 | 1.0165 | 0.023645 |
| FFA241 | 0.33258 | 0.50117 | 1.0851 | 0.11784 | 0.21642 | 0.5914 | 1.1187 | 0.16182 | 0.34455 | 0.43219 | 0.89662 | -0.15743 |
| TAG483-FA141 | 0.04321 | 0.12568 | 0.56874 | -0.81417 | 0.26427 | 0.59421 | 0.84323 | -0.246 | 0.012586 | 0.041592 | 0.39635 | -1.3351 |
| PEP-181/226 | 0.26366 | 0.43017 | 1.184 | 0.24368 | 0.28536 | 0.60708 | 1.2187 | 0.28537 | 0.21034 | 0.29073 | 1.1653 | 0.22075 |
| PEP-160/180 | 0.96106 | 0.97588 | 1.0304 | 0.04321 | 0.43796 | 0.72846 | 1.2096 | 0.27454 | 0.48274 | 0.56421 | 0.86978 | -0.20128 |
| FFA201 | 0.64256 | 0.75644 | 0.92585 | -0.11114 | 0.17417 | 0.5914 | 1.2748 | 0.35029 | 0.91396 | 0.93405 | 1.0549 | 0.077164 |
| FFA141 | 0.7364 | 0.83229 | 0.98042 | -0.02853 | 0.20403 | 0.5914 | 1.2866 | 0.36356 | 0.62468 | 0.69635 | 1.1277 | 0.17334 |
| PC200/226 | 0.62241 | 0.74759 | 0.90364 | -0.14618 | 0.17174 | 0.5914 | 0.78224 | -0.35431 | 0.036196 | 0.077637 | 0.6582 | -0.6034 |
| PEP-140/181 | 0.10209 | 0.21409 | 1.2634 | 0.33729 | 0.30886 | 0.62637 | 1.2633 | 0.33719 | 0.1169 | 0.18034 | 0.77164 | -0.374 |
| LPE170 | 0.39411 | 0.55887 | 1.2414 | 0.31193 | 0.91723 | 0.96465 | 0.99079 | -0.01335 | 0.008536 | 0.037908 | 0.48546 | -1.0426 |
| FFA240 | 0.08625 | 0.19098 | 0.74752 | -0.41981 | 0.73139 | 0.86886 | 1.0237 | 0.033803 | 0.11287 | 0.17454 | 0.76711 | -0.38249 |
| Cholesterol | 0.471 | 0.63543 | 1.087 | 0.12032 | 0.003448 | 0.28651 | 0.50783 | -0.97759 | 0.34136 | 0.42901 | 0.8259 | -0.27597 |
| FFA160 | 0.8258 | 0.88859 | 0.96095 | -0.05747 | 0.32003 | 0.63325 | 1.0961 | 0.1324 | 0.38848 | 0.47271 | 0.89807 | -0.1551 |
| PEP-181/204 | 0.96115 | 0.97588 | 1.0096 | 0.013738 | 0.65703 | 0.83216 | 1.0897 | 0.12397 | 0.63814 | 0.70532 | 0.90671 | -0.14129 |
| PE180/224 | 0.63579 | 0.75392 | 1.1198 | 0.16327 | 0.71395 | 0.85829 | 0.93561 | -0.09603 | 0.013404 | 0.042154 | 0.55928 | -0.83836 |
| CE183 | 0.46416 | 0.62952 | 1.2829 | 0.35939 | 0.009794 | 0.28651 | 0.58161 | -0.78189 | 0.5233 | 0.60189 | 0.88393 | -0.178 |
| PEP-182/204 | 0.65505 | 0.7656 | 1.0828 | 0.11472 | 0.50599 | 0.73692 | 1.0937 | 0.12922 | 0.8125 | 0.85001 | 0.97496 | -0.03659 |
| CER240 | 0.50331 | 0.65662 | 1.1033 | 0.14184 | 0.045442 | 0.5194 | 1.4021 | 0.48756 | 0.69531 | 0.75693 | 0.91308 | -0.13118 |
| PEP-160/183 | 0.48411 | 0.64579 | 0.86925 | -0.20215 | 0.56899 | 0.77018 | 0.88647 | -0.17386 | 0.004647 | 0.032129 | 0.50013 | -0.99963 |
| PA181/182 | 0.028306 | 0.10046 | 0.81096 | -0.30231 | 0.022659 | 0.3649 | 1.3921 | 0.47729 | 0.62158 | 0.69408 | 0.95467 | -0.06693 |
| FFA181 | 0.76644 | 0.85145 | 1.016 | 0.022935 | 0.25164 | 0.5914 | 1.1153 | 0.15749 | 0.34934 | 0.43651 | 0.88788 | -0.17157 |
| PE180/182 | 0.50931 | 0.65785 | 1.0876 | 0.12111 | 0.95548 | 0.9811 | 0.99068 | -0.01352 | 0.016281 | 0.045884 | 0.66186 | -0.59541 |
| PC160/201 | 0.49805 | 0.65369 | 1.0647 | 0.090447 | 0.90859 | 0.9571 | 0.96583 | -0.05016 | 0.48917 | 0.56967 | 0.90023 | -0.15163 |
| PEP-180/224 | 0.94079 | 0.96754 | 1.0414 | 0.058539 | 0.94913 | 0.97817 | 0.98735 | -0.01836 | 0.3471 | 0.43454 | 0.78707 | -0.34543 |
| PS160/161_2 | 0.9847 | 0.98773 | 1.0018 | 0.002648 | 0.021984 | 0.3649 | 0.78323 | -0.3525 | 0.29686 | 0.38135 | 1.0967 | 0.13322 |
| PE180/181 | 0.28752 | 0.45321 | 1.1786 | 0.23712 | 0.059743 | 0.54498 | 1.2686 | 0.3432 | 0.02075 | 0.051955 | 0.64865 | -0.62448 |

| | | | | | | | | | | | | |
|---|---|---|---|---|---|---|---|---|---|---|---|---|
| FFA170 | 0.099567 | 0.21178 | 0.80128 | -0.31963 | 0.8196 | 0.90757 | 1.0348 | 0.049309 | 0.20345 | 0.28606 | 0.83585 | -0.25868 |
| FFA182 | 0.69493 | 0.79788 | 1.0249 | 0.035473 | 0.45168 | 0.72879 | 1.0661 | 0.092357 | 0.15799 | 0.23061 | 0.83407 | -0.26177 |
| PC150/161 | 0.48 | 0.64423 | 0.91268 | -0.13183 | 0.22759 | 0.5914 | 0.86748 | -0.20509 | 0.56611 | 0.63944 | 0.93551 | -0.09618 |
| PC180/205 | 0.62185 | 0.74759 | 1.1935 | 0.25514 | 0.66761 | 0.83459 | 1.1666 | 0.22234 | 0.80141 | 0.84284 | 1.1141 | 0.15587 |
| FFA180 | 0.54488 | 0.68788 | 0.92021 | -0.11996 | 0.52716 | 0.74605 | 1.0776 | 0.10776 | 0.258 | 0.34277 | 0.86049 | -0.21677 |
| TAG524-FA204 | 0.83273 | 0.8931 | 0.92642 | -0.11026 | 0.15416 | 0.5914 | 0.74993 | -0.41518 | 0.82394 | 0.85411 | 1.199 | 0.2618 |
| LPE182 | 0.17451 | 0.31469 | 1.3578 | 0.44132 | 0.024039 | 0.37261 | 1.485 | 0.5705 | 0.034127 | 0.07554 | 0.62789 | -0.67141 |
| TAG480-FA160 | 0.44253 | 0.61165 | 0.74364 | -0.42732 | 0.089733 | 0.5914 | 0.56732 | -0.81777 | 0.10274 | 0.16039 | 0.54045 | -0.88775 |
| CER200 | 0.56954 | 0.70612 | 1.0614 | 0.085996 | 0.44352 | 0.72846 | 0.84643 | -0.24053 | 0.19569 | 0.27634 | 0.75315 | -0.40899 |
| CER220 | 0.48776 | 0.6467 | 1.0834 | 0.11555 | 0.23267 | 0.5914 | 1.1883 | 0.24893 | 0.31014 | 0.39511 | 0.82323 | -0.28063 |
| PEO-160/160 | 0.16184 | 0.2993 | 1.329 | 0.41031 | 0.80309 | 0.90608 | 0.92881 | -0.10654 | 0.31161 | 0.39621 | 0.77723 | -0.36358 |
| TAG564-FA204 | 0.31239 | 0.4821 | 0.77041 | -0.37631 | 0.63604 | 0.82155 | 1.0905 | 0.12503 | 0.56333 | 0.6389 | 0.9346 | -0.09758 |
| PS160/181 | 0.60391 | 0.73486 | 0.95951 | -0.05963 | 0.063623 | 0.54498 | 0.81424 | -0.29647 | 0.3914 | 0.47449 | 1.0803 | 0.11146 |
| LPE220 | 0.099972 | 0.21178 | 1.3013 | 0.38 | 0.002562 | 0.28651 | 1.817 | 0.86154 | 0.095655 | 0.15225 | 0.78792 | -0.34388 |
| TAG525-FA205 | 0.49722 | 0.65369 | 0.86975 | -0.20133 | 0.13838 | 0.5914 | 0.61139 | -0.70983 | 0.53915 | 0.61903 | 1.0131 | 0.018773 |
| TAG544-FA224 | 0.97181 | 0.9803 | 0.91076 | -0.13486 | 0.45185 | 0.72879 | 1.0375 | 0.053139 | 0.56838 | 0.63944 | 1.0133 | 0.019079 |
| PEO-180/161 | 0.79245 | 0.8685 | 1.0965 | 0.13291 | 0.96317 | 0.98589 | 0.99568 | -0.00625 | 0.066686 | 0.11639 | 0.67897 | -0.55858 |
| FFA221 | 0.32716 | 0.49802 | 0.80216 | -0.31804 | 0.12445 | 0.5914 | 0.72376 | -0.46642 | 0.075477 | 0.12965 | 0.66565 | -0.58717 |
| PE170/224 | 0.83967 | 0.89905 | 1.016 | 0.022931 | 0.28239 | 0.60274 | 0.83435 | -0.26127 | 0.002207 | 0.026601 | 0.52436 | -0.93136 |
| PC180/180 | 0.83206 | 0.8931 | 1.0365 | 0.051771 | 0.21144 | 0.5914 | 1.2068 | 0.27124 | 0.82928 | 0.85692 | 1.0418 | 0.059104 |
| CE181 | 0.74273 | 0.8352 | 0.90053 | -0.15115 | 0.081911 | 0.5914 | 0.67292 | -0.5715 | 0.67093 | 0.73408 | 0.90891 | -0.13778 |
| PA220/226 | 0.6408 | 0.75579 | 0.93724 | -0.09351 | 0.041678 | 0.49332 | 1.3192 | 0.39964 | 0.12483 | 0.18725 | 0.80581 | -0.31148 |
| PEO-160/181 | 0.76636 | 0.85145 | 1.1084 | 0.14844 | 0.78988 | 0.89932 | 1.0401 | 0.056753 | 0.03358 | 0.075382 | 0.61395 | -0.70382 |
| TAG482-FA141 | 0.18174 | 0.32414 | 0.79733 | -0.32675 | 0.75655 | 0.89057 | 1.2108 | 0.27595 | 0.001413 | 0.023486 | 0.44176 | -1.1787 |
| FFA150 | 0.069253 | 0.16335 | 0.8545 | -0.22685 | 0.05421 | 0.53471 | 0.84214 | -0.24787 | 0.001294 | 0.023486 | 0.73948 | -0.43542 |
| LPC160 | 0.7884 | 0.86551 | 0.96143 | -0.05674 | 0.21473 | 0.5914 | 1.1418 | 0.19128 | 0.28368 | 0.36715 | 0.87531 | -0.19214 |
| TAG568-FA226 | 0.44667 | 0.61346 | 0.56908 | -0.81329 | 0.79422 | 0.90076 | 0.97415 | -0.03778 | 0.75668 | 0.80851 | 0.9584 | -0.0613 |
| LPE201 | 0.31346 | 0.48242 | 1.2502 | 0.32213 | 0.006422 | 0.28651 | 1.7947 | 0.84378 | 0.010695 | 0.038366 | 0.62803 | -0.67109 |
| PE160/224 | 0.39194 | 0.55833 | 1.2123 | 0.27776 | 0.23379 | 0.5914 | 0.69481 | -0.52532 | 0.014225 | 0.043476 | 0.43047 | -1.216 |
| PC160/203 | 0.82065 | 0.88597 | 0.94935 | -0.07499 | 0.41924 | 0.72374 | 0.87652 | -0.19013 | 0.74944 | 0.80244 | 0.94368 | -0.08363 |
| FFA222 | 0.85504 | 0.90734 | 0.97748 | -0.03285 | 0.49452 | 0.73497 | 1.0865 | 0.11972 | 0.39375 | 0.47556 | 0.90403 | -0.14556 |
| HCER201 | 0.74824 | 0.83983 | 1.0169 | 0.024213 | 0.009722 | 0.28651 | 0.8078 | -0.30793 | 0.031245 | 0.072129 | 0.81096 | -0.30229 |
| TAG544-FA204 | 0.81256 | 0.88017 | 0.98428 | -0.02285 | 0.5903 | 0.78747 | 1.1319 | 0.17873 | 0.48208 | 0.56421 | 0.87207 | -0.19749 |

| | | | | | | | | | | | | |
|---|---|---|---|---|---|---|---|---|---|---|---|---|
| TAG567-FA226 | 0.39331 | 0.55887 | 0.57368 | -0.80168 | 0.95518 | 0.9811 | 1.046 | 0.064818 | 0.71443 | 0.77386 | 0.91371 | -0.13019 |
| PC200/225 | 0.28722 | 0.45321 | 0.87883 | -0.18634 | 0.30779 | 0.62637 | 0.88229 | -0.18067 | 0.015332 | 0.044759 | 0.66994 | -0.57789 |
| TAG503-FA141 | 0.11935 | 0.23517 | 0.64055 | -0.64262 | 0.71449 | 0.85829 | 1.3126 | 0.39241 | 0.001754 | 0.023486 | 0.34869 | -1.52 |
| TAG565-FA225 | 0.50209 | 0.65634 | 0.68106 | -0.55415 | 0.64712 | 0.828 | 0.98878 | -0.01628 | 0.2835 | 0.36715 | 0.66042 | -0.59855 |
| PC160/120 | 0.64487 | 0.75778 | 1.1359 | 0.18386 | 0.97981 | 0.99244 | 0.90565 | -0.14297 | 0.76475 | 0.81273 | 0.75615 | -0.40325 |
| FFA200 | 0.70624 | 0.8066 | 0.89098 | -0.16654 | 0.6107 | 0.80316 | 0.86904 | -0.2025 | 0.69017 | 0.7526 | 1.1056 | 0.14485 |
| PC180/224 | 0.63542 | 0.75392 | 0.90409 | -0.14546 | 0.77656 | 0.89253 | 0.94259 | -0.0853 | 0.10829 | 0.16866 | 0.72144 | -0.47105 |
| PC200/141 | 0.93383 | 0.9619 | 0.96082 | -0.05767 | 0.65586 | 0.83216 | 1.0229 | 0.032606 | 0.27024 | 0.35469 | 0.79717 | -0.32704 |
| PS161/180_2 | 0.66625 | 0.7759 | 1.0503 | 0.070777 | 0.72058 | 0.86073 | 0.95005 | -0.07392 | 0.41726 | 0.4975 | 1.1225 | 0.16672 |
| LPE161 | 0.61464 | 0.74098 | 0.89476 | -0.16042 | 0.37613 | 0.68573 | 0.83333 | -0.26305 | 0.024226 | 0.058413 | 0.57776 | -0.79147 |
| PC200/204 | 0.6267 | 0.74892 | 1.0284 | 0.04038 | 0.21212 | 0.5914 | 0.82961 | -0.26949 | 0.043137 | 0.0867 | 0.70345 | -0.50748 |
| SMd181/183 | 0.68107 | 0.78613 | 0.97595 | -0.03511 | 0.10869 | 0.5914 | 0.90026 | -0.15159 | 0.12547 | 0.18778 | 0.84544 | -0.24223 |
| PC180/200 | 0.8666 | 0.91252 | 1.0182 | 0.026086 | 0.27383 | 0.59421 | 1.245 | 0.31616 | 0.94053 | 0.95372 | 0.94936 | -0.07497 |
| PE160/181 | 0.079309 | 0.1798 | 1.4831 | 0.56861 | 0.75163 | 0.88644 | 1.0148 | 0.021201 | 0.032412 | 0.073519 | 0.52672 | -0.9249 |
| PEP-160/201 | 0.79888 | 0.87196 | 1.1024 | 0.14061 | 0.88321 | 0.94723 | 0.96453 | -0.05211 | 0.40387 | 0.4842 | 0.79673 | -0.32784 |
| PC180/120 | 0.5333 | 0.68075 | 1.1756 | 0.23344 | 0.81193 | 0.90726 | 0.86957 | -0.20163 | 0.43642 | 0.516 | 0.67711 | -0.56253 |
| PEP-180/226 | 0.56684 | 0.70433 | 1.1066 | 0.14611 | 0.44535 | 0.72846 | 1.1392 | 0.18807 | 0.91656 | 0.93524 | 0.99705 | -0.00427 |
| DAG200/200 | 0.52155 | 0.66836 | 0.92791 | -0.10795 | 0.005133 | 0.28651 | 0.56208 | -0.83116 | 0.89081 | 0.91325 | 1.1034 | 0.14196 |
| PEP-160/161 | 0.34115 | 0.51173 | 1.2307 | 0.29952 | 0.52524 | 0.74605 | 1.1718 | 0.22868 | 0.74423 | 0.79818 | 0.94146 | -0.08703 |
| CER201 | 0.81816 | 0.88475 | 1.0186 | 0.026524 | 0.85473 | 0.93016 | 1.1074 | 0.14716 | 0.56244 | 0.6389 | 0.79583 | -0.32947 |
| PC170/204 | 0.98406 | 0.98773 | 0.96749 | -0.04768 | 0.47173 | 0.73251 | 0.85216 | -0.2308 | 0.306 | 0.39061 | 0.80518 | -0.31261 |
| PE180/205 | 0.54228 | 0.68681 | 1.2509 | 0.32297 | 0.79775 | 0.90319 | 0.90177 | -0.14916 | 0.56872 | 0.63944 | 0.8039 | -0.31492 |
| FFA224 | 0.87087 | 0.91442 | 1.0024 | 0.003454 | 0.64459 | 0.828 | 1.1161 | 0.1585 | 0.81604 | 0.85104 | 0.98357 | -0.0239 |
| PC140/224 | 0.14084 | 0.26967 | 0.73638 | -0.44147 | 0.26273 | 0.59421 | 0.81022 | -0.30361 | 0.020607 | 0.051796 | 0.53696 | -0.8971 |

## 7 Gy

| | SD 1 vs. Pre | | | | SD 2 vs. Pre | | | | SD 6 vs. Pre | | | |
|---|---|---|---|---|---|---|---|---|---|---|---|---|
| Lipid Name | p-value | FDR | FC | LOG2FC | p-value | FDR | FC | LOG2FC | p-value | FDR | FC | LOG2FC |
| PI160/201 | 0.001105 | 0.007052 | 0.57792 ↓ | -0.791 | 0.002953 | 0.024037 | 0.66577 ↓ | -0.587 | 0.02242 | 0.066178 | 0.62618 ↓ | -0.675 |
| LPE160 | 0.000239 | 0.0037 | 1.8642 ↑ | 0.8986 | 0.39308 | 0.56206 | 1.1225 ↑ | 0.1667 | 0.74598 | 0.81209 | 0.94608 ↓ | -0.08 |
| PC182/203 | 0.058847 | 0.11234 | 0.53132 ↓ | -0.912 | 0.19903 | 0.36192 | 0.65912 ↓ | -0.601 | 0.18854 | 0.28023 | 0.67407 ↓ | -0.569 |
| PG180/182_2 | 0.001374 | 0.007983 | 0.56011 ↓ | -0.836 | 0.01263 | 0.060671 | 0.60178 ↓ | -0.733 | 0.010466 | 0.048666 | 0.56248 ↓ | -0.83 |
| PI160/204 | 0.006047 | 0.019316 | 0.61812 ↓ | -0.694 | 0.60779 | 0.75509 | 1.0391 ↑ | 0.0553 | 0.96539 | 0.96836 | 1.0276 ↑ | 0.0392 |

| | | | | | | | | | | | | | | | |
|---|---|---|---|---|---|---|---|---|---|---|---|---|---|---|---|
| PG180/182 | 0.001948 | 0.009607 | 0.5596 | ↓ | -0.838 | 0.013292 | 0.060936 | 0.59612 | ↓ | -0.746 | 0.010172 | 0.048666 | 0.55568 | ↓ | -0.848 |
| LPC202 | 0.048211 | 0.09657 | 0.76432 | ↓ | -0.388 | 0.002546 | 0.024019 | 0.71082 | ↓ | -0.492 | 0.08428 | 0.14962 | 0.73416 | ↓ | -0.446 |
| TAG547-FA183 | 3.3E-05 | 0.003 | 0.11024 | ↓ | -3.181 | 0.000173 | 0.011362 | 0.17035 | ↓ | -2.553 | 0.003062 | 0.046601 | 0.24915 | ↓ | -2.005 |
| PI160/182 | 0.000146 | 0.003 | 0.45286 | ↓ | -1.143 | 0.21739 | 0.38879 | 0.88381 | ↓ | -0.178 | 0.002913 | 0.046601 | 0.53889 | ↓ | -0.892 |
| TAG547-FA182 | 6.05E-05 | 0.003 | 0.1332 | ↓ | -2.908 | 0.000536 | 0.012404 | 0.21079 | ↓ | -2.246 | 0.006933 | 0.046601 | 0.32571 | ↓ | -1.618 |
| PS161/180 | 0.01024 | 0.028984 | 0.52547 | ↓ | -0.928 | 0.00328 | 0.024827 | 0.49883 | ↓ | -1.003 | 0.032064 | 0.079367 | 0.58482 | ↓ | -0.774 |
| TAG526-FA183 | 6.03E-05 | 0.003 | 0.10913 | ↓ | -3.196 | 0.000495 | 0.012404 | 0.17437 | ↓ | -2.52 | 0.002717 | 0.046601 | 0.20183 | ↓ | -2.309 |
| TAG483-FA183 | 0.000903 | 0.006413 | 0.13836 | ↓ | -2.854 | 0.004485 | 0.030414 | 0.22426 | ↓ | -2.157 | 0.008431 | 0.04704 | 0.24358 | ↓ | -2.038 |
| LPE226 | 0.001019 | 0.006767 | 2.0731 | ↑ | 1.0518 | 0.12738 | 0.25438 | 1.3441 | ↑ | 0.4266 | 0.23131 | 0.32594 | 1.1674 | ↑ | 0.2233 |
| PG160/161 | 0.038745 | 0.081365 | 0.54991 | ↓ | -0.863 | 0.073837 | 0.17167 | 0.60871 | ↓ | -0.716 | 0.079993 | 0.14506 | 0.59359 | ↓ | -0.752 |
| TAG525-FA182 | 0.000212 | 0.003464 | 0.18527 | ↓ | -2.432 | 0.001343 | 0.018197 | 0.27127 | ↓ | -1.882 | 0.005198 | 0.046601 | 0.28587 | ↓ | -1.807 |
| PE181/203 | 0.029793 | 0.065403 | 0.60754 | ↓ | -0.719 | 0.018965 | 0.074827 | 0.55501 | ↓ | -0.849 | 0.020898 | 0.062992 | 0.5118 | ↓ | -0.966 |
| PI160/180 | 0.008114 | 0.02457 | 0.56329 | ↓ | -0.828 | 0.061307 | 0.15061 | 0.79125 | ↓ | -0.338 | 0.000716 | 0.029311 | 0.51173 | ↓ | -0.967 |
| PC160/226 | 0.000647 | 0.0049 | 1.5675 | ↑ | 0.6484 | 0.047906 | 0.12572 | 1.369 | ↑ | 0.4532 | 0.002477 | 0.046601 | 1.3973 | ↑ | 0.4827 |
| PI160/160 | 0.001189 | 0.007413 | 0.58316 | ↓ | -0.778 | 0.02735 | 0.090199 | 0.8531 | ↓ | -0.229 | 0.01346 | 0.053428 | 0.67971 | ↓ | -0.557 |
| PI181/203_2 | 0.063146 | 0.11804 | 0.56151 | ↓ | -0.833 | 0.32138 | 0.50053 | 0.75305 | ↓ | -0.409 | 0.54274 | 0.63658 | 0.91654 | ↓ | -0.126 |
| TAG542-FA200 | 0.000456 | 0.004178 | 0.27108 | ↓ | -1.883 | 0.003028 | 0.024037 | 0.37345 | ↓ | -1.421 | 0.008078 | 0.04704 | 0.36114 | ↓ | -1.469 |
| TAG546-FA183 | 5.65E-05 | 0.003 | 0.13946 | ↓ | -2.842 | 0.000625 | 0.012404 | 0.25267 | ↓ | -1.985 | 0.005775 | 0.046601 | 0.30403 | ↓ | -1.718 |
| TAG504-FA182 | 7.74E-05 | 0.003 | 0.15344 | ↓ | -2.704 | 0.000723 | 0.012421 | 0.2499 | ↓ | -2.001 | 0.010895 | 0.048666 | 0.33125 | ↓ | -1.594 |
| PC181/202 | 0.040025 | 0.082981 | 0.68875 | ↓ | -0.538 | 0.010363 | 0.052705 | 0.63476 | ↓ | -0.656 | 0.15777 | 0.24223 | 0.79101 | ↓ | -0.338 |
| PC182/183 | 0.001287 | 0.007689 | 0.36904 | ↓ | -1.438 | 0.007842 | 0.043263 | 0.50576 | ↓ | -0.983 | 0.014209 | 0.054983 | 0.43805 | ↓ | -1.191 |
| PI160/182_2 | 0.000116 | 0.003 | 0.4411 | ↓ | -1.181 | 0.16836 | 0.32331 | 0.87149 | ↓ | -0.198 | 0.003085 | 0.046601 | 0.54736 | ↓ | -0.869 |
| PI181/203 | 0.039652 | 0.082702 | 0.53419 | ↓ | -0.905 | 0.23527 | 0.41035 | 0.74118 | ↓ | -0.432 | 0.36231 | 0.46248 | 0.83076 | ↓ | -0.268 |
| PG160/161_2 | 0.12925 | 0.21036 | 0.72246 | ↓ | -0.469 | 0.055594 | 0.1391 | 0.76684 | ↓ | -0.383 | 0.13052 | 0.20965 | 0.71757 | ↓ | -0.479 |
| LPC200 | 0.079812 | 0.14393 | 0.65003 | ↓ | -0.621 | 0.67389 | 0.80201 | 0.92691 | ↓ | -0.11 | 0.019109 | 0.060684 | 0.55286 | ↓ | -0.855 |
| TAG504-FA140 | 0.000138 | 0.003 | 0.15946 | ↓ | -2.649 | 0.000305 | 0.012402 | 0.21251 | ↓ | -2.234 | 0.005148 | 0.046601 | 0.26275 | ↓ | -1.928 |
| TAG525-FA183 | 0.000576 | 0.004633 | 0.23149 | ↓ | -2.111 | 0.002936 | 0.024037 | 0.32079 | ↓ | -1.64 | 0.004845 | 0.046601 | 0.30693 | ↓ | -1.704 |
| TAG526-FA182 | 9.82E-05 | 0.003 | 0.13476 | ↓ | -2.892 | 0.000728 | 0.012421 | 0.20424 | ↓ | -2.292 | 0.002713 | 0.046601 | 0.2187 | ↓ | -2.193 |
| TAG525-FA225 | 0.020617 | 0.047935 | 0.42349 | ↓ | -1.24 | 0.00491 | 0.03165 | 0.35599 | ↓ | -1.49 | 0.00759 | 0.046601 | 0.35174 | ↓ | -1.507 |
| TAG504-FA183 | 7.09E-05 | 0.003 | 0.10301 | ↓ | -3.279 | 0.001177 | 0.017416 | 0.18372 | ↓ | -2.444 | 0.003593 | 0.046601 | 0.22584 | ↓ | -2.147 |
| TAG526-FA161 | 0.000149 | 0.003 | 0.08932 | ↓ | -3.485 | 0.000336 | 0.012404 | 0.13593 | ↓ | -2.879 | 0.002715 | 0.046601 | 0.1816 | ↓ | -2.461 |
| CE204 | 0.042816 | 0.087651 | 1.7929 | ↑ | 0.8423 | 0.019416 | 0.075689 | 0.67086 | ↓ | -0.576 | 0.035282 | 0.083523 | 1.4244 | ↑ | 0.5104 |
| TAG546-FA181 | 8.15E-05 | 0.003 | 0.14826 | ↓ | -2.754 | 0.001893 | 0.019775 | 0.28932 | ↓ | -1.789 | 0.008527 | 0.04704 | 0.33052 | ↓ | -1.597 |

| | | | | | | | | | | | | | | | |
|---|---|---|---|---|---|---|---|---|---|---|---|---|---|---|---|
| PI160/204_2 | 0.003367 | 0.013364 | 0.61083 | ↓ | -0.711 | 0.48157 | 0.64908 | 1.0631 | ↑ | 0.0883 | 0.69248 | 0.76881 | 1.0969 | ↑ | 0.1335 |
| TAG503-FA140 | 3.47E-05 | 0.003 | 0.14833 | ↓ | -2.753 | 0.003504 | 0.025343 | 0.35165 | ↓ | -1.508 | 0.009109 | 0.047822 | 0.29893 | ↓ | -1.742 |
| PI161/181 | 0.013605 | 0.035569 | 0.58382 | ↓ | -0.776 | 0.79519 | 0.88188 | 0.98226 | ↓ | -0.026 | 0.22994 | 0.32542 | 0.81986 | ↓ | -0.287 |
| LPC220 | 0.062874 | 0.11796 | 0.63862 | ↓ | -0.647 | 0.69909 | 0.81561 | 0.95134 | ↓ | -0.072 | 0.005022 | 0.046601 | 0.53722 | ↓ | -0.896 |
| TAG548-FA182 | 0.000245 | 0.0037 | 0.13364 | ↓ | -2.904 | 0.000227 | 0.011362 | 0.16756 | ↓ | -2.577 | 0.002222 | 0.046601 | 0.25063 | ↓ | -1.996 |
| PC182/202 | 0.052668 | 0.10297 | 0.68106 | ↓ | -0.554 | 0.045094 | 0.12031 | 0.73773 | ↓ | -0.439 | 0.15055 | 0.23391 | 0.75473 | ↓ | -0.406 |
| PI182/182 | 0.000146 | 0.003 | 0.35546 | ↓ | -1.492 | 0.079685 | 0.17888 | 0.77671 | ↓ | -0.365 | 0.005608 | 0.046601 | 0.49289 | ↓ | -1.021 |
| TAG546-FA182 | 0.00204 | 0.009765 | 0.26861 | ↓ | -1.896 | 0.011454 | 0.056066 | 0.3592 | ↓ | -1.477 | 0.030386 | 0.077879 | 0.45112 | ↓ | -1.148 |
| PI181/182_2 | 0.003281 | 0.013186 | 0.43023 | ↓ | -1.217 | 0.77141 | 0.86883 | 1.173 | ↑ | 0.2302 | 0.020901 | 0.062992 | 0.54498 | ↓ | -0.876 |
| PI160/180_2 | 0.004566 | 0.015981 | 0.58172 | ↓ | -0.782 | 0.003213 | 0.024827 | 0.71907 | ↓ | -0.476 | 0.001552 | 0.045924 | 0.54942 | ↓ | -0.864 |
| TAG525-FA160 | 0.001573 | 0.008535 | 0.26687 | ↓ | -1.906 | 0.002082 | 0.020228 | 0.3003 | ↓ | -1.736 | 0.006654 | 0.046601 | 0.35038 | ↓ | -1.513 |
| TAG525-FA161 | 0.000152 | 0.003 | 0.15936 | ↓ | -2.65 | 0.001656 | 0.019249 | 0.26094 | ↓ | -1.938 | 0.004257 | 0.046601 | 0.24239 | ↓ | -2.045 |
| TAG505-FA161 | 0.00034 | 0.00386 | 0.11751 | ↓ | -3.089 | 0.00137 | 0.018197 | 0.16339 | ↓ | -2.614 | 0.001464 | 0.04538 | 0.12039 | ↓ | -3.054 |
| PG181/182 | 0.02635 | 0.058347 | 0.52396 | ↓ | -0.932 | 0.3681 | 0.5385 | 0.78495 | ↓ | -0.349 | 0.027921 | 0.07359 | 0.48582 | ↓ | -1.042 |
| TAG563-FA200 | 0.002001 | 0.009705 | 0.28042 | ↓ | -1.834 | 0.57793 | 0.73055 | 0.89103 | ↓ | -0.166 | 0.018304 | 0.059579 | 0.37389 | ↓ | -1.419 |
| PC160/204 | 0.015333 | 0.03854 | 1.4279 | ↑ | 0.5139 | 0.054002 | 0.13679 | 1.2098 | ↑ | 0.2748 | 0.034199 | 0.081852 | 1.3097 | ↑ | 0.3893 |
| PC182/182 | 0.001519 | 0.008309 | 0.48745 | ↓ | -1.037 | 0.24533 | 0.41918 | 0.84312 | ↓ | -0.246 | 0.04454 | 0.097302 | 0.62927 | ↓ | -0.668 |
| PI160/181 | 0.009162 | 0.026627 | 0.52421 | ↓ | -0.932 | 0.92619 | 0.95415 | 1.0452 | ↑ | 0.0637 | 0.008821 | 0.047781 | 0.53675 | ↓ | -0.898 |
| TAG482-FA182 | 0.002648 | 0.011121 | 0.24383 | ↓ | -2.036 | 0.005836 | 0.035494 | 0.29041 | ↓ | -1.784 | 0.038032 | 0.086874 | 0.39695 | ↓ | -1.333 |
| PG181/181 | 0.063888 | 0.11883 | 0.55154 | ↓ | -0.858 | 0.59403 | 0.74405 | 1.389 | ↑ | 0.4741 | 0.072251 | 0.13594 | 0.54395 | ↓ | -0.878 |
| TAG566-FA183 | 0.000278 | 0.0037 | 0.22547 | ↓ | -2.149 | 0.000183 | 0.011362 | 0.27201 | ↓ | -1.878 | 0.022555 | 0.066178 | 0.4745 | ↓ | -1.076 |
| PC182/161 | 0.17317 | 0.26714 | 0.76314 | ↓ | -0.39 | 0.26813 | 0.44303 | 0.81924 | ↓ | -0.288 | 0.13939 | 0.22025 | 0.74063 | ↓ | -0.433 |
| PI181/182 | 0.004535 | 0.015981 | 0.43062 | ↓ | -1.216 | 0.84609 | 0.90643 | 1.1069 | ↑ | 0.1465 | 0.016981 | 0.057235 | 0.51943 | ↓ | -0.945 |
| TAG524-FA221 | 0.004048 | 0.014972 | 0.31535 | ↓ | -1.665 | 0.018396 | 0.074109 | 0.40709 | ↓ | -1.297 | 0.014966 | 0.055673 | 0.36993 | ↓ | -1.435 |
| PI160/181_2 | 0.016785 | 0.041391 | 0.53146 | ↓ | -0.912 | 0.9609 | 0.97589 | 1.0372 | ↑ | 0.0527 | 0.007556 | 0.046601 | 0.51702 | ↓ | -0.952 |
| TAG541-FA200 | 0.000584 | 0.00464 | 0.2682 | ↓ | -1.899 | 0.02388 | 0.083135 | 0.50517 | ↓ | -0.985 | 0.009047 | 0.047822 | 0.35405 | ↓ | -1.498 |
| TAG548-FA183 | 5.82E-05 | 0.003 | 0.081816 | ↓ | -3.612 | 7.72E-05 | 0.011362 | 0.11673 | ↓ | -3.099 | 0.000124 | 0.009247 | 0.099394 | ↓ | -3.331 |
| CE226 | 0.000646 | 0.0049 | 1.856 | ↑ | 0.8922 | 0.31036 | 0.48797 | 0.85308 | ↓ | -0.229 | 0.003329 | 0.046601 | 2.0568 | ↑ | 1.0404 |
| TAG503-FA182 | 0.000529 | 0.004461 | 0.19836 | ↓ | -2.334 | 0.00863 | 0.046429 | 0.33793 | ↓ | -1.565 | 0.011342 | 0.049553 | 0.30772 | ↓ | -1.7 |
| LPC183 | 0.010118 | 0.028764 | 0.68889 | ↓ | -0.538 | 0.14289 | 0.28102 | 0.79816 | ↓ | -0.325 | 0.014699 | 0.055314 | 0.54911 | ↓ | -0.865 |
| TAG525-FA181 | 0.000305 | 0.0037 | 0.16439 | ↓ | -2.605 | 0.005571 | 0.034666 | 0.30401 | ↓ | -1.718 | 0.004607 | 0.046601 | 0.24287 | ↓ | -2.042 |
| PC140/226 | 0.68792 | 0.77213 | 0.94906 | ↓ | -0.075 | 0.075621 | 0.17276 | 0.78296 | ↓ | -0.353 | 0.21913 | 0.31167 | 0.83008 | ↓ | -0.269 |
| LPC150 | 0.064193 | 0.11906 | 0.75564 | ↓ | -0.404 | 0.2905 | 0.4696 | 0.8618 | ↓ | -0.215 | 0.013392 | 0.053428 | 0.60834 | ↓ | -0.717 |

| | | | | | | | | | | | | | | | |
|---|---|---|---|---|---|---|---|---|---|---|---|---|---|---|---|
| PS161/181 | 0.09439 | 0.16607 | 0.59357 | ↓ | -0.753 | 0.3358 | 0.51076 | 0.82979 | ↓ | -0.269 | 0.3992 | 0.49786 | 0.86178 | ↓ | -0.215 |
| PC180/201 | 0.84785 | 0.89081 | 1.0033 | ↑ | 0.0048 | 0.08021 | 0.17944 | 0.78327 | ↓ | -0.352 | 0.21905 | 0.31167 | 0.83794 | ↓ | -0.255 |
| PI181/181 | 0.044536 | 0.090321 | 0.49135 | ↓ | -1.025 | 0.32796 | 0.50466 | 1.7248 | ↑ | 0.7864 | 0.016441 | 0.057235 | 0.41272 | ↓ | -1.277 |
| PI160/203_2 | 0.003508 | 0.01384 | 0.55313 | ↓ | -0.854 | 0.022932 | 0.082479 | 0.76422 | ↓ | -0.388 | 0.3387 | 0.43662 | 0.88548 | ↓ | -0.175 |
| TAG562-FA200 | 0.001846 | 0.009337 | 0.31621 | ↓ | -1.661 | 0.83522 | 0.90232 | 1.0367 | ↑ | 0.0519 | 0.03376 | 0.081184 | 0.44399 | ↓ | -1.171 |
| PC180/202 | 0.78583 | 0.85121 | 1.0021 | ↑ | 0.003 | 0.046014 | 0.12215 | 0.7963 | ↓ | -0.329 | 0.26956 | 0.36408 | 0.87479 | ↓ | -0.193 |
| TAG547-FA181 | 1.14E-05 | 0.003 | 0.1585 | ↓ | -2.658 | 4.25E-05 | 0.011362 | 0.2308 | ↓ | -2.115 | 0.002765 | 0.046601 | 0.29273 | ↓ | -1.772 |
| TAG503-FA181 | 0.00021 | 0.003464 | 0.16889 | ↓ | -2.566 | 0.006616 | 0.038453 | 0.34698 | ↓ | -1.527 | 0.011703 | 0.050121 | 0.295 | ↓ | -1.761 |
| TAG484-FA120 | 0.002641 | 0.011121 | 0.13183 | ↓ | -2.923 | 0.027184 | 0.090199 | 0.33805 | ↓ | -1.565 | 0.12035 | 0.19718 | 0.62918 | ↓ | -0.668 |
| TAG504-FA161 | 0.000632 | 0.0049 | 0.1334 | ↓ | -2.906 | 0.002011 | 0.019837 | 0.16819 | ↓ | -2.572 | 0.004681 | 0.046601 | 0.18379 | ↓ | -2.444 |
| PG160/203_2 | 0.010418 | 0.029361 | 0.61839 | ↓ | -0.693 | 0.001473 | 0.018755 | 0.47703 | ↓ | -1.068 | 0.012917 | 0.052471 | 0.50766 | ↓ | -0.978 |
| TAG524-FA161 | 8.5E-05 | 0.003 | 0.15328 | ↓ | -2.706 | 0.01084 | 0.054706 | 0.36586 | ↓ | -1.451 | 0.007225 | 0.046601 | 0.27687 | ↓ | -1.853 |
| TAG563-FA182 | 0.000741 | 0.005479 | 0.2937 | ↓ | -1.768 | 0.32869 | 0.50466 | 0.77707 | ↓ | -0.364 | 0.031347 | 0.078791 | 0.47802 | ↓ | -1.065 |
| TAG565-FA201 | 0.005991 | 0.019316 | 0.35615 | ↓ | -1.489 | 0.11203 | 0.23137 | 0.57391 | ↓ | -0.801 | 0.084348 | 0.14962 | 0.61064 | ↓ | -0.712 |
| PI160/203 | 0.004792 | 0.016676 | 0.59142 | ↓ | -0.758 | 0.055767 | 0.1391 | 0.80667 | ↓ | -0.31 | 0.23908 | 0.33399 | 0.84136 | ↓ | -0.249 |
| PI180/180 | 0.057092 | 0.10964 | 0.61937 | ↓ | -0.691 | 0.97776 | 0.98727 | 1.0298 | ↑ | 0.0424 | 0.011597 | 0.050121 | 0.5081 | ↓ | -0.977 |
| TAG541-FA160 | 0.000141 | 0.003 | 0.28708 | ↓ | -1.801 | 0.024312 | 0.084188 | 0.59171 | ↓ | -0.757 | 0.007321 | 0.046601 | 0.3821 | ↓ | -1.388 |
| PC182/224 | 0.053656 | 0.10427 | 0.60462 | ↓ | -0.726 | 0.10801 | 0.22609 | 0.66705 | ↓ | -0.584 | 0.16842 | 0.2561 | 0.7008 | ↓ | -0.513 |
| PC182/205 | 0.012586 | 0.03358 | 0.49391 | ↓ | -1.018 | 0.64689 | 0.78715 | 0.94618 | ↓ | -0.08 | 0.057898 | 0.11526 | 0.61164 | ↓ | -0.709 |
| TAG545-FA180 | 0.000213 | 0.003464 | 0.27622 | ↓ | -1.856 | 0.0005 | 0.012404 | 0.30837 | ↓ | -1.697 | 0.007677 | 0.046601 | 0.42184 | ↓ | -1.245 |
| PI161/181_2 | 0.009095 | 0.026627 | 0.47113 | ↓ | -1.086 | 0.63154 | 0.77426 | 0.93003 | ↓ | -0.105 | 0.095718 | 0.16617 | 0.68345 | ↓ | -0.549 |
| TAG524-FA160 | 0.002375 | 0.01041 | 0.39153 | ↓ | -1.353 | 0.021074 | 0.078395 | 0.51343 | ↓ | -0.962 | 0.035127 | 0.083459 | 0.52805 | ↓ | -0.921 |
| PC160/225 | 0.002011 | 0.009705 | 1.7891 | ↑ | 0.8393 | 0.019392 | 0.075689 | 1.5097 | ↑ | 0.5942 | 0.069818 | 0.13213 | 1.4216 | ↑ | 0.5075 |
| TAG524-FA183 | 0.000307 | 0.0037 | 0.25541 | ↓ | -1.969 | 0.001765 | 0.019633 | 0.33922 | ↓ | -1.56 | 0.00469 | 0.046601 | 0.34653 | ↓ | -1.529 |
| TAG561-FA181 | 0.00038 | 0.003994 | 0.30746 | ↓ | -1.702 | 0.50841 | 0.67135 | 0.91298 | ↓ | -0.131 | 0.056113 | 0.11275 | 0.53426 | ↓ | -0.904 |
| TAG482-FA140 | 0.002013 | 0.009705 | 0.24047 | ↓ | -2.056 | 0.006134 | 0.036112 | 0.29571 | ↓ | -1.758 | 0.012582 | 0.051516 | 0.30077 | ↓ | -1.733 |
| PI200/204_2 | 0.35738 | 0.47096 | 0.86736 | ↓ | -0.205 | 0.75729 | 0.85888 | 0.94033 | ↓ | -0.089 | 0.096581 | 0.16677 | 0.63889 | ↓ | -0.646 |
| TAG502-FA140 | 0.000249 | 0.0037 | 0.18616 | ↓ | -2.425 | 0.038857 | 0.11203 | 0.48422 | ↓ | -1.046 | 0.03083 | 0.078096 | 0.40606 | ↓ | -1.3 |
| TAG545-FA183 | 8.52E-05 | 0.003 | 0.1913 | ↓ | -2.386 | 0.000854 | 0.013552 | 0.30016 | ↓ | -1.736 | 0.005876 | 0.046601 | 0.31446 | ↓ | -1.669 |
| PG160/203 | 0.025703 | 0.057304 | 0.70161 | ↓ | -0.511 | 0.000593 | 0.012404 | 0.50967 | ↓ | -0.972 | 0.015445 | 0.056603 | 0.54853 | ↓ | -0.866 |
| PI160/161 | 0.01108 | 0.030881 | 0.58231 | ↓ | -0.78 | 0.20531 | 0.371 | 0.85177 | ↓ | -0.231 | 0.044395 | 0.097302 | 0.67326 | ↓ | -0.571 |
| PE182/202 | 0.019455 | 0.046222 | 0.57437 | ↓ | -0.8 | 0.000726 | 0.012421 | 0.44788 | ↓ | -1.159 | 0.008376 | 0.04704 | 0.41151 | ↓ | -1.281 |
| PI182/201 | 0.035988 | 0.076813 | 0.5982 | ↓ | -0.741 | 0.82537 | 0.89936 | 0.99227 | ↓ | -0.011 | 0.022401 | 0.066178 | 0.59565 | ↓ | -0.747 |

| | | | | | | | | | | | | |
|---|---|---|---|---|---|---|---|---|---|---|---|---|
| LPA180 | 0.053847 | 0.10433 | 0.71872 | ↓ -0.477 | 0.075452 | 0.17276 | 0.75386 | ↓ -0.408 | 0.16134 | 0.24713 | 0.78592 | ↓ -0.348 |
| LPC240 | 0.10339 | 0.17854 | 0.73922 | ↓ -0.436 | 0.13996 | 0.27694 | 0.78329 | ↓ -0.352 | 0.009721 | 0.048307 | 0.63359 | ↓ -0.658 |
| TAG524-FA200 | 0.00176 | 0.009115 | 0.30579 | ↓ -1.709 | 0.020078 | 0.076891 | 0.44704 | ↓ -1.162 | 0.018907 | 0.060465 | 0.41995 | ↓ -1.252 |
| TAG566-FA202 | 0.011246 | 0.03102 | 0.38858 | ↓ -1.364 | 0.015474 | 0.067071 | 0.39976 | ↓ -1.323 | 0.082008 | 0.14789 | 0.59085 | ↓ -0.759 |
| PS181/182 | 0.010008 | 0.028575 | 0.66509 | ↓ -0.588 | 0.14184 | 0.27981 | 0.8208 | ↓ -0.285 | 0.001419 | 0.04538 | 0.59642 | ↓ -0.746 |
| TAG524-FA181 | 0.000111 | 0.003 | 0.21008 | ↓ -2.251 | 0.001574 | 0.018858 | 0.35606 | ↓ -1.49 | 0.005881 | 0.046601 | 0.31264 | ↓ -1.677 |
| PG161/180 | 0.63466 | 0.7274 | 1.1689 | ↑ 0.2252 | 0.051982 | 0.13363 | 0.60563 | ↓ -0.724 | 0.029873 | 0.0773 | 0.52945 | ↓ -0.917 |
| PI180/203_2 | 0.45797 | 0.57445 | 0.88267 | ↓ -0.18 | 0.62753 | 0.7708 | 0.90498 | ↓ -0.144 | 0.86634 | 0.91113 | 0.9977 | ↓ -0.003 |
| PI180/183_2 | 0.005444 | 0.018362 | 0.63774 | ↓ -0.649 | 0.027572 | 0.090199 | 0.80629 | ↓ -0.311 | 0.002246 | 0.046601 | 0.60155 | ↓ -0.733 |
| LPC241 | 2.47E-05 | 0.003 | 0.43379 | ↓ -1.205 | 9.45E-05 | 0.011362 | 0.58824 | ↓ -0.766 | 0.001002 | 0.038385 | 0.55388 | ↓ -0.852 |
| CE203 | 0.16031 | 0.25026 | 1.529 | ↑ 0.6126 | 0.026228 | 0.088469 | 0.57535 | ↓ -0.797 | 0.69919 | 0.77279 | 1.0557 | ↑ 0.0782 |
| TAG524-FA182 | 0.004261 | 0.015409 | 0.46577 | ↓ -1.102 | 0.016765 | 0.069514 | 0.53614 | ↓ -0.899 | 0.028842 | 0.075104 | 0.53755 | ↓ -0.896 |
| TAG545-FA182 | 0.003355 | 0.013364 | 0.30069 | ↓ -1.734 | 0.10391 | 0.21962 | 0.54298 | ↓ -0.881 | 0.02707 | 0.073032 | 0.41831 | ↓ -1.257 |
| TAG544-FA201 | 6.97E-05 | 0.003 | 0.2708 | ↓ -1.885 | 0.000463 | 0.012404 | 0.43766 | ↓ -1.192 | 0.022567 | 0.066178 | 0.49681 | ↓ -1.009 |
| TAG545-FA161 | 0.000142 | 0.003 | 0.2315 | ↓ -2.111 | 0.000429 | 0.012404 | 0.2983 | ↓ -1.745 | 0.008043 | 0.04704 | 0.35617 | ↓ -1.489 |
| TAG502-FA181 | 0.000296 | 0.0037 | 0.18234 | ↓ -2.455 | 0.022868 | 0.082479 | 0.40958 | ↓ -1.288 | 0.01674 | 0.057235 | 0.32774 | ↓ -1.609 |
| PG181/182_2 | 0.049417 | 0.097192 | 0.55488 | -0.84976 | 0.61356 | 0.75649 | 0.88247 | -0.18038 | 0.054382 | 0.11133 | 0.5321 | -0.91023 |
| PC200/182 | 0.31954 | 0.43428 | 0.8704 | -0.20026 | 0.48847 | 0.65312 | 0.91786 | -0.12366 | 0.004322 | 0.046601 | 0.59821 | -0.74128 |
| PC140/203 | 0.11853 | 0.20042 | 0.70023 | -0.51411 | 0.10131 | 0.21482 | 0.69952 | -0.51555 | 0.19196 | 0.28465 | 0.73928 | -0.4358 |
| TAG543-FA183 | 0.000573 | 0.004633 | 0.32066 | -1.6409 | 0.000187 | 0.011362 | 0.43046 | -1.2161 | 0.006455 | 0.046601 | 0.39471 | -1.3411 |
| PC181/203 | 0.30267 | 0.42012 | 0.7475 | -0.41986 | 0.074962 | 0.17276 | 0.58895 | -0.76379 | 0.37581 | 0.47598 | 0.79395 | -0.33288 |
| TAG523-FA201 | 9.75E-07 | 0.000635 | 0.22369 | -2.1604 | 9.71E-05 | 0.011362 | 0.42887 | -1.2214 | 0.011664 | 0.050121 | 0.39985 | -1.3225 |
| LPC201 | 0.10697 | 0.18325 | 0.70762 | -0.49896 | 0.29143 | 0.4696 | 0.828 | -0.27231 | 0.098683 | 0.1695 | 0.70587 | -0.50253 |
| PE181/205 | 0.30969 | 0.42533 | 0.76341 | -0.38947 | 0.41022 | 0.57679 | 0.8012 | -0.31976 | 0.11132 | 0.18487 | 0.6198 | -0.69012 |
| LPC161 | 0.030554 | 0.066747 | 0.71697 | -0.48002 | 0.42972 | 0.59775 | 0.90687 | -0.14103 | 0.071169 | 0.13429 | 0.71227 | -0.48951 |
| TAG481-FA181 | 0.002512 | 0.010829 | 0.22938 | -2.1242 | 0.029075 | 0.091778 | 0.37954 | -1.3977 | 0.052413 | 0.10901 | 0.42015 | -1.251 |
| PC120/181 | 0.23355 | 0.34014 | 0.69951 | -0.51558 | 0.44606 | 0.61523 | 0.79321 | -0.33422 | 0.24232 | 0.3378 | 0.70065 | -0.51323 |
| PC150/203 | 0.32859 | 0.44288 | 0.85359 | -0.22839 | 0.008034 | 0.043949 | 0.66568 | -0.58709 | 0.036582 | 0.084088 | 0.64034 | -0.64309 |
| TAG567-FA183 | 8.08E-05 | 0.003 | 0.21599 | -2.211 | 0.000297 | 0.012402 | 0.26639 | -1.9084 | 0.017056 | 0.057235 | 0.43232 | -1.2098 |
| TAG544-FA183 | 0.000271 | 0.0037 | 0.29643 | -1.7542 | 0.000649 | 0.012404 | 0.41948 | -1.2533 | 0.006768 | 0.046601 | 0.37935 | -1.3984 |
| TAG523-FA183 | 0.000404 | 0.004083 | 0.28485 | -1.8117 | 0.000433 | 0.012404 | 0.33917 | -1.5599 | 0.005738 | 0.046601 | 0.37447 | -1.4171 |
| PE182/201 | 0.10433 | 0.17968 | 0.61618 | -0.69858 | 0.18378 | 0.34335 | 0.67129 | -0.57499 | 0.017607 | 0.057889 | 0.43693 | -1.1945 |
| TAG544-FA180 | 0.000323 | 0.003756 | 0.36 | -1.4739 | 0.001842 | 0.019655 | 0.48377 | -1.0476 | 0.009505 | 0.047969 | 0.4126 | -1.2772 |

| | | | | | | | | | | | | |
|---|---|---|---|---|---|---|---|---|---|---|---|---|
| TAG482-FA161 | 0.000534 | 0.004461 | 0.15427 | -2.6965 | 0.10666 | 0.224 | 0.44957 | -1.1534 | 0.00534 | 0.046601 | 0.20806 | -2.265 |
| TAG482-FA160 | 0.002415 | 0.010479 | 0.24464 | -2.0313 | 0.01112 | 0.055235 | 0.32387 | -1.6265 | 0.016409 | 0.057235 | 0.32317 | -1.6296 |
| PI180/181_2 | 0.084637 | 0.15075 | 0.67589 | -0.56515 | 0.48537 | 0.65284 | 1.182 | 0.24119 | 0.005417 | 0.046601 | 0.48485 | -1.0444 |
| TAG523-FA180 | 0.000441 | 0.004178 | 0.31606 | -1.6617 | 0.000202 | 0.011362 | 0.32937 | -1.6022 | 0.00282 | 0.046601 | 0.3078 | -1.6999 |
| PG160/182_2 | 0.006023 | 0.019316 | 0.57933 | -0.78753 | 0.12079 | 0.24346 | 0.7438 | -0.42702 | 0.01496 | 0.055673 | 0.55291 | -0.85489 |
| TAG503-FA161 | 0.001778 | 0.009115 | 0.22998 | -2.1204 | 0.009963 | 0.051068 | 0.33508 | -1.5774 | 0.010027 | 0.048666 | 0.30751 | -1.7013 |
| PI180/181 | 0.10274 | 0.17789 | 0.66046 | -0.59845 | 0.64498 | 0.7863 | 1.1674 | 0.22334 | 0.010642 | 0.048666 | 0.47717 | -1.0674 |
| PC182/226 | 0.001196 | 0.007413 | 0.5556 | -0.84788 | 0.59433 | 0.74405 | 0.92379 | -0.11436 | 0.081682 | 0.14771 | 0.70124 | -0.51202 |
| PC181/183 | 0.014356 | 0.036794 | 0.61353 | -0.70479 | 0.10838 | 0.22613 | 0.72976 | -0.45451 | 0.009289 | 0.047969 | 0.45678 | -1.1304 |
| PC182/204 | 0.11228 | 0.19135 | 0.67132 | -0.57493 | 0.75461 | 0.85883 | 0.89619 | -0.15812 | 0.27085 | 0.3646 | 0.76819 | -0.38046 |
| TAG565-FA182 | 0.003281 | 0.013186 | 0.34344 | -1.5419 | 0.027941 | 0.090496 | 0.47173 | -1.084 | 0.068716 | 0.13094 | 0.5621 | -0.83111 |
| TAG544-FA161 | 0.000205 | 0.003464 | 0.26111 | -1.9373 | 0.005889 | 0.035494 | 0.43118 | -1.2136 | 0.012302 | 0.051012 | 0.39068 | -1.356 |
| PC181/226 | 0.36581 | 0.4782 | 0.86122 | -0.21555 | 0.11663 | 0.23802 | 0.78514 | -0.34898 | 0.68348 | 0.76189 | 0.94982 | -0.07428 |
| TAG520-FA200 | 0.003185 | 0.013125 | 0.38939 | -1.3607 | 0.002636 | 0.024037 | 0.34263 | -1.5453 | 0.007058 | 0.046601 | 0.36567 | -1.4514 |
| SMd181/182 | 0.029838 | 0.065403 | 1.2163 | 0.28251 | 0.33771 | 0.51128 | 1.0896 | 0.12377 | 0.40091 | 0.49903 | 1.0757 | 0.10521 |
| PC140/202 | 0.34459 | 0.45929 | 0.84745 | -0.2388 | 0.024629 | 0.084387 | 0.68019 | -0.55599 | 0.13075 | 0.20965 | 0.72701 | -0.45995 |
| TAG524-FA180 | 0.000356 | 0.00386 | 0.28785 | -1.7966 | 3.99E-05 | 0.011362 | 0.36118 | -1.4692 | 0.003391 | 0.046601 | 0.34472 | -1.5365 |
| PG160/182 | 0.00569 | 0.018804 | 0.58277 | -0.77899 | 0.12069 | 0.24346 | 0.74448 | -0.4257 | 0.017402 | 0.057505 | 0.56715 | -0.81819 |
| PI180/182_2 | 0.020244 | 0.047406 | 0.6996 | -0.5154 | 0.73815 | 0.84452 | 0.97355 | -0.03867 | 0.006143 | 0.046601 | 0.67181 | -0.57387 |
| TAG526-FA181 | 0.003775 | 0.014458 | 0.24841 | -2.0092 | 0.075631 | 0.17276 | 0.4891 | -1.0318 | 0.005547 | 0.046601 | 0.25225 | -1.9871 |
| TAG483-FA161 | 0.000484 | 0.004198 | 0.13368 | -2.9032 | 0.040215 | 0.11364 | 0.36032 | -1.4726 | 0.006121 | 0.046601 | 0.21892 | -2.1915 |
| TAG565-FA202 | 0.005588 | 0.018655 | 0.41301 | -1.2758 | 0.015792 | 0.067071 | 0.47831 | -1.064 | 0.051775 | 0.10879 | 0.53683 | -0.89745 |
| LPC170 | 0.006718 | 0.021027 | 0.75245 | -0.41033 | 0.065571 | 0.15774 | 0.85401 | -0.22767 | 0.02047 | 0.062859 | 0.75003 | -0.41498 |
| TAG545-FA181 | 0.002063 | 0.009789 | 0.2828 | -1.8221 | 0.093473 | 0.20352 | 0.53429 | -0.9043 | 0.021164 | 0.063493 | 0.39015 | -1.3579 |
| TAG523-FA202 | 0.000373 | 0.003979 | 0.27278 | -1.8742 | 0.001944 | 0.019775 | 0.39284 | -1.348 | 0.020171 | 0.062531 | 0.43127 | -1.2133 |
| PI180/182 | 0.017963 | 0.043471 | 0.68224 | -0.55165 | 0.83891 | 0.90343 | 0.9834 | -0.02415 | 0.003589 | 0.046601 | 0.63416 | -0.65709 |
| TAG540-FA160 | 0.000134 | 0.003 | 0.32431 | -1.6245 | 0.000277 | 0.012402 | 0.37136 | -1.4291 | 0.002188 | 0.046601 | 0.36297 | -1.4621 |
| TAG526-FA160 | 0.000355 | 0.00386 | 0.22212 | -2.1706 | 0.001561 | 0.018858 | 0.24821 | -2.0104 | 0.00243 | 0.046601 | 0.26065 | -1.9398 |
| TAG526-FA205 | 0.24108 | 0.34799 | 0.44037 | -1.1832 | 0.41528 | 0.58262 | 0.52439 | -0.93128 | 0.53195 | 0.62561 | 0.54668 | -0.87123 |
| PS180/181 | 0.055738 | 0.10735 | 0.6972 | -0.52036 | 0.008845 | 0.046811 | 0.59507 | -0.74886 | 0.21352 | 0.30658 | 0.78899 | -0.34192 |
| TAG523-FA182 | 0.001896 | 0.009463 | 0.46029 | -1.1194 | 0.025571 | 0.087155 | 0.61085 | -0.71112 | 0.021763 | 0.06499 | 0.53315 | -0.90738 |
| TAG545-FA202 | 0.000257 | 0.0037 | 0.24715 | -2.0165 | 0.001346 | 0.018197 | 0.34741 | -1.5253 | 0.010633 | 0.048666 | 0.38874 | -1.3631 |
| TAG483-FA140 | 0.000185 | 0.00344 | 0.10486 | -3.2535 | 0.000667 | 0.012404 | 0.1622 | -2.6242 | 0.007615 | 0.046601 | 0.22264 | -2.1672 |

| | | | | | | | | | | | | |
|---|---|---|---|---|---|---|---|---|---|---|---|---|
| PE180/202 | 0.25649 | 0.36538 | 1.1336 | 0.18094 | 0.014816 | 0.065685 | 0.73409 | -0.44597 | 0.044928 | 0.097739 | 0.78149 | -0.35569 |
| PI180/202_2 | 0.062638 | 0.11796 | 0.76087 | -0.39429 | 0.017433 | 0.071377 | 0.71801 | -0.47791 | 0.045522 | 0.098303 | 0.75655 | -0.40249 |
| PE181/204 | 0.21938 | 0.32458 | 0.7752 | -0.36737 | 0.09685 | 0.20947 | 0.69463 | -0.52568 | 0.14112 | 0.22245 | 0.72327 | -0.46739 |
| TAG523-FA181 | 0.00171 | 0.008976 | 0.45587 | -1.1333 | 0.035541 | 0.10517 | 0.64798 | -0.62599 | 0.016017 | 0.057092 | 0.49863 | -1.004 |
| TAG503-FA160 | 0.004009 | 0.014972 | 0.28726 | -1.7996 | 0.004678 | 0.031074 | 0.30111 | -1.7317 | 0.009036 | 0.047822 | 0.32245 | -1.6328 |
| TAG483-FA182 | 0.002379 | 0.01041 | 0.18041 | -2.4707 | 0.078169 | 0.1773 | 0.47515 | -1.0736 | 0.10404 | 0.17638 | 0.56421 | -0.82571 |
| PC140/161 | 0.29966 | 0.41772 | 0.76505 | -0.38637 | 0.17603 | 0.33313 | 0.70548 | -0.50333 | 0.21052 | 0.30556 | 0.72464 | -0.46466 |
| TAG523-FA161 | 0.000128 | 0.003 | 0.18459 | -2.4376 | 0.040585 | 0.11364 | 0.47346 | -1.0787 | 0.005545 | 0.046601 | 0.26225 | -1.931 |
| TAG5610-FA182 | 0.000663 | 0.004962 | 0.30131 | -1.7307 | 0.028251 | 0.090598 | 0.48821 | -1.0344 | 0.012977 | 0.052471 | 0.38577 | -1.3742 |
| PC180/183 | 0.24366 | 0.35094 | 0.80195 | -0.31842 | 0.1201 | 0.24346 | 0.73238 | -0.44934 | 0.004958 | 0.046601 | 0.50068 | -0.99804 |
| TAG564-FA201 | 0.017757 | 0.043296 | 0.3876 | -1.3674 | 0.94451 | 0.96679 | 0.94383 | -0.08341 | 0.096912 | 0.1669 | 0.58029 | -0.78516 |
| PE182/161 | 0.004816 | 0.016676 | 0.6136 | -0.70464 | 0.57758 | 0.73055 | 0.96127 | -0.05698 | 0.000547 | 0.028245 | 0.44887 | -1.1556 |
| PE181/226 | 0.084754 | 0.15075 | 0.75257 | -0.4101 | 0.021643 | 0.080054 | 0.66265 | -0.59367 | 0.076545 | 0.14077 | 0.72727 | -0.45943 |
| TAG481-FA140 | 0.001169 | 0.007391 | 0.23092 | -2.1145 | 0.014211 | 0.063963 | 0.37643 | -1.4096 | 0.015994 | 0.057092 | 0.32409 | -1.6255 |
| PS161/182_2 | 0.01516 | 0.038402 | 0.70571 | -0.50286 | 0.029465 | 0.091778 | 0.73991 | -0.43458 | 0.003414 | 0.046601 | 0.57527 | -0.7977 |
| PC140/181 | 0.30173 | 0.41971 | 0.82413 | -0.27905 | 0.68952 | 0.81319 | 0.92966 | -0.10522 | 0.17891 | 0.26775 | 0.75677 | -0.40207 |
| PI160/161_2 | 0.019929 | 0.047178 | 0.63942 | -0.64517 | 0.56762 | 0.72313 | 0.90404 | -0.14554 | 0.17157 | 0.25855 | 0.76141 | -0.39326 |
| PA180/182 | 0.043549 | 0.088874 | 0.63856 | -0.6471 | 0.24978 | 0.42456 | 1.3698 | 0.45394 | 0.055919 | 0.1127 | 0.65832 | -0.60313 |
| TAG504-FA160 | 0.003824 | 0.014559 | 0.23169 | -2.1097 | 0.005802 | 0.035494 | 0.25944 | -1.9465 | 0.006222 | 0.046601 | 0.25407 | -1.9767 |
| PC182/225 | 0.019293 | 0.046175 | 0.62001 | -0.68963 | 0.7634 | 0.86431 | 0.93651 | -0.09464 | 0.39317 | 0.49127 | 0.86617 | -0.20727 |
| TAG523-FA160 | 0.000935 | 0.006543 | 0.47097 | -1.0863 | 0.012676 | 0.060671 | 0.61446 | -0.7026 | 0.014513 | 0.055252 | 0.53111 | -0.91293 |
| PI180/202 | 0.079099 | 0.14304 | 0.78556 | -0.34821 | 0.040524 | 0.11364 | 0.77785 | -0.36244 | 0.047888 | 0.10221 | 0.79205 | -0.33633 |
| PS180/181_2 | 0.031775 | 0.069183 | 0.70179 | -0.51089 | 0.004732 | 0.031115 | 0.66487 | -0.58885 | 0.43519 | 0.5359 | 0.90126 | -0.14999 |
| SMd181/203 | 0.000108 | 0.003 | 1.2296 | 0.29815 | 0.53242 | 0.69182 | 1.0632 | 0.088347 | 0.11611 | 0.19233 | 1.0808 | 0.11204 |
| PE181/202 | 0.15181 | 0.24105 | 0.78535 | -0.34859 | 0.002654 | 0.024037 | 0.56547 | -0.82249 | 0.013704 | 0.053909 | 0.47733 | -1.0669 |
| TAG484-FA182 | 0.007301 | 0.022471 | 0.2406 | -2.0553 | 0.023366 | 0.083031 | 0.34106 | -1.5519 | 0.10344 | 0.17582 | 0.54569 | -0.87384 |
| PI180/205 | 0.20164 | 0.30176 | 0.82521 | -0.27716 | 0.76601 | 0.86576 | 0.96845 | -0.04626 | 0.63004 | 0.71084 | 0.93225 | -0.10121 |
| PEP-180/182 | 0.00734 | 0.022471 | 0.60522 | -0.72448 | 0.72898 | 0.83697 | 0.97025 | -0.04357 | 5.41E-05 | 0.00587 | 0.38816 | -1.3653 |
| PE181/182 | 0.063283 | 0.11804 | 0.70621 | -0.50184 | 0.21301 | 0.38203 | 0.79915 | -0.32346 | 0.003382 | 0.046601 | 0.46911 | -1.092 |
| PEO-160/225 | 0.002075 | 0.009789 | 1.687 | 0.75445 | 0.023668 | 0.083031 | 1.41 | 0.49573 | 0.2108 | 0.30556 | 1.3122 | 0.39199 |
| PI180/203 | 0.48909 | 0.59946 | 0.90704 | -0.14077 | 0.83579 | 0.90232 | 0.961 | -0.05739 | 0.74946 | 0.81452 | 1.0255 | 0.036283 |
| LPC226 | 0.004522 | 0.015981 | 1.4346 | 0.52061 | 0.18864 | 0.3481 | 1.2055 | 0.2696 | 0.025674 | 0.070523 | 1.2482 | 0.31984 |
| PEP-180/181 | 0.039763 | 0.082702 | 0.63331 | -0.65901 | 0.90119 | 0.94613 | 1.0434 | 0.061269 | 6.62E-05 | 0.006161 | 0.36946 | -1.4365 |

| | | | | | | | | | | | | |
|---|---|---|---|---|---|---|---|---|---|---|---|---|
| TAG569-FA204 | 0.089334 | 0.15846 | 0.5949 | -0.74928 | 0.003584 | 0.025637 | 0.39741 | -1.3313 | 0.52681 | 0.62129 | 0.86101 | -0.2159 |
| PC140/182 | 0.22277 | 0.32811 | 0.7843 | -0.35052 | 0.83959 | 0.90343 | 1.0213 | 0.03046 | 0.16943 | 0.2565 | 0.74743 | -0.42 |
| TAG544-FA182 | 0.004207 | 0.0153 | 0.39153 | -1.3528 | 0.33269 | 0.50841 | 0.73828 | -0.43776 | 0.024356 | 0.068561 | 0.44306 | -1.1744 |
| TAG522-FA181 | 0.000906 | 0.006413 | 0.40168 | -1.3159 | 0.040674 | 0.11364 | 0.65426 | -0.61207 | 0.016883 | 0.057235 | 0.4895 | -1.0306 |
| PEP-180/203 | 0.19261 | 0.29354 | 0.78322 | -0.35251 | 0.77894 | 0.87278 | 0.94771 | -0.07748 | 0.001857 | 0.046601 | 0.53382 | -0.90557 |
| PC150/182 | 0.33328 | 0.44827 | 0.85714 | -0.22239 | 0.79333 | 0.88133 | 0.96632 | -0.04943 | 0.02 | 0.062531 | 0.67575 | -0.56543 |
| TAG566-FA182 | 0.000988 | 0.006732 | 0.38327 | -1.3836 | 0.002944 | 0.024037 | 0.43381 | -1.2049 | 0.053407 | 0.11037 | 0.57546 | -0.79721 |
| PC182/201 | 0.097231 | 0.1697 | 0.67518 | -0.56665 | 0.3842 | 0.55335 | 0.83313 | -0.26338 | 0.051977 | 0.10879 | 0.59124 | -0.75818 |
| TAG567-FA182 | 0.003271 | 0.013186 | 0.38195 | -1.3885 | 0.007444 | 0.042141 | 0.39647 | -1.3347 | 0.031954 | 0.079367 | 0.50112 | -0.99676 |
| PC160/205 | 0.3044 | 0.42158 | 1.5267 | 0.61038 | 0.60492 | 0.75296 | 1.3865 | 0.47143 | 0.52232 | 0.61824 | 1.3595 | 0.44302 |
| TAG568-FA183 | 0.002708 | 0.0113 | 0.2739 | -1.8683 | 0.003127 | 0.024526 | 0.27791 | -1.8473 | 0.016988 | 0.057235 | 0.37452 | -1.4169 |
| TAG524-FA203 | 0.002606 | 0.011086 | 0.30412 | -1.7173 | 0.009236 | 0.047896 | 0.36955 | -1.4362 | 0.024303 | 0.068561 | 0.41213 | -1.2788 |
| HCER180 | 0.021042 | 0.048748 | 1.5003 | 0.58524 | 0.041942 | 0.11582 | 1.2091 | 0.27396 | 0.25803 | 0.35216 | 1.1322 | 0.17912 |
| TAG567-FA203 | 0.052669 | 0.10297 | 0.45571 | -1.1338 | 0.02187 | 0.080435 | 0.38389 | -1.3812 | 0.16877 | 0.2561 | 0.61405 | -0.70358 |
| PC170/203 | 0.69042 | 0.77361 | 0.96296 | -0.05445 | 0.052438 | 0.13387 | 0.78931 | -0.34134 | 0.24963 | 0.34301 | 0.84693 | -0.23968 |
| TAG502-FA180 | 0.00048 | 0.004198 | 0.3058 | -1.7093 | 0.000443 | 0.012404 | 0.33872 | -1.5618 | 0.00708 | 0.046601 | 0.3278 | -1.6091 |
| TAG483-FA120 | 0.020433 | 0.047677 | 0.25879 | -1.9502 | 0.29136 | 0.4696 | 0.78027 | -0.35796 | 0.24373 | 0.33903 | 1.0591 | 0.082793 |
| SMd181/160 | 0.20934 | 0.31247 | 1.0647 | 0.090394 | 0.04828 | 0.12572 | 1.1103 | 0.1509 | 0.004012 | 0.046601 | 1.1489 | 0.20022 |
| TAG544-FA202 | 0.000408 | 0.004083 | 0.35915 | -1.4773 | 0.001498 | 0.018755 | 0.42979 | -1.2183 | 0.032419 | 0.079943 | 0.52292 | -0.93534 |
| TAG502-FA161 | 0.001003 | 0.006732 | 0.23226 | -2.1062 | 0.014832 | 0.065685 | 0.38534 | -1.3758 | 0.010466 | 0.048666 | 0.31203 | -1.6802 |
| PE140/204 | 0.062204 | 0.11772 | 0.67012 | -0.5775 | 0.1715 | 0.32645 | 0.75107 | -0.41298 | 0.064844 | 0.12564 | 0.65099 | -0.6193 |
| TAG563-FA202 | 0.001442 | 0.008048 | 0.45101 | -1.1488 | 0.018303 | 0.074109 | 0.6104 | -0.71216 | 0.033555 | 0.081184 | 0.53902 | -0.8916 |
| TAG569-FA183 | 0.017933 | 0.043471 | 0.34057 | -1.554 | 0.002866 | 0.024037 | 0.34965 | -1.516 | 0.008117 | 0.04704 | 0.39634 | -1.3352 |
| LPE204 | 0.001251 | 0.00757 | 1.6017 | 0.67963 | 0.041966 | 0.11582 | 1.2696 | 0.34432 | 0.97804 | 0.97804 | 0.99607 | -0.00568 |
| PI180/224 | 0.14252 | 0.22909 | 0.83343 | -0.26286 | 0.56573 | 0.72213 | 0.93844 | -0.09166 | 0.89786 | 0.92193 | 0.97817 | -0.03185 |
| TAG562-FA180 | 0.001904 | 0.009463 | 0.37747 | -1.4056 | 0.073457 | 0.17167 | 0.64441 | -0.63396 | 0.020872 | 0.062992 | 0.45261 | -1.1437 |
| TAG521-FA161 | 0.001409 | 0.008048 | 0.30806 | -1.6987 | 0.038521 | 0.11203 | 0.50872 | -0.97506 | 0.00577 | 0.046601 | 0.31986 | -1.6445 |
| PS181/182_2 | 0.20096 | 0.30143 | 0.77566 | -0.36651 | 0.97817 | 0.98727 | 1.0002 | 0.000298 | 0.007056 | 0.046601 | 0.54946 | -0.86391 |
| TAG503-FA183 | 0.004856 | 0.016726 | 0.31936 | -1.6467 | 0.007424 | 0.042141 | 0.32248 | -1.6327 | 0.015477 | 0.056603 | 0.37733 | -1.4061 |
| PE170/181 | 0.20042 | 0.30132 | 0.69131 | -0.53259 | 0.18714 | 0.34709 | 0.68165 | -0.55291 | 0.001175 | 0.040242 | 0.33089 | -1.5956 |
| TAG569-FA205 | 0.58334 | 0.68179 | 0.98386 | -0.02347 | 0.22386 | 0.39819 | 0.74908 | -0.41681 | 0.70674 | 0.77717 | 1.1515 | 0.20351 |
| TAG563-FA181 | 0.002139 | 0.009937 | 0.32343 | -1.6285 | 0.81808 | 0.8951 | 0.96491 | -0.05153 | 0.054582 | 0.11139 | 0.47987 | -1.0593 |
| PE181/183 | 0.34408 | 0.45929 | 0.80105 | -0.32004 | 0.040281 | 0.11364 | 0.55179 | -0.85781 | 0.003941 | 0.046601 | 0.35642 | -1.4883 |

| | | | | | | | | | | | | |
|---|---|---|---|---|---|---|---|---|---|---|---|---|
| TAG481-FA160 | 0.005074 | 0.017294 | 0.27677 | -1.8532 | 0.029399 | 0.091778 | 0.38245 | -1.3867 | 0.028223 | 0.073788 | 0.36355 | -1.4598 |
| TAG542-FA160 | 6.35E-05 | 0.003 | 0.3399 | -1.5568 | 0.001278 | 0.018197 | 0.52149 | -0.93929 | 0.018824 | 0.060465 | 0.46098 | -1.1172 |
| TAG546-FA203 | 0.000641 | 0.0049 | 0.29085 | -1.7816 | 0.00284 | 0.024037 | 0.36397 | -1.4581 | 0.015669 | 0.056986 | 0.4101 | -1.2859 |
| TAG567-FA224 | 0.000143 | 0.003 | 0.28046 | -1.8341 | 0.000634 | 0.012404 | 0.29898 | -1.7419 | 0.00806 | 0.04704 | 0.39921 | -1.3248 |
| TAG501-FA180 | 0.001609 | 0.008644 | 0.30105 | -1.7319 | 0.013268 | 0.060936 | 0.40523 | -1.3032 | 0.027846 | 0.07359 | 0.41729 | -1.2609 |
| PC181/161 | 0.1981 | 0.29923 | 0.78805 | -0.34365 | 0.55359 | 0.71223 | 0.9126 | -0.13195 | 0.14628 | 0.23001 | 0.74844 | -0.41805 |
| TAG543-FA180 | 0.001413 | 0.008048 | 0.48923 | -1.0314 | 0.009156 | 0.047896 | 0.68204 | -0.55206 | 0.010166 | 0.048666 | 0.43122 | -1.2135 |
| PC180/204 | 0.060614 | 0.11504 | 1.306 | 0.38517 | 0.46256 | 0.63351 | 1.0729 | 0.10153 | 0.2087 | 0.30463 | 1.1958 | 0.25796 |
| TAG544-FA181 | 0.006988 | 0.021663 | 0.39924 | -1.3247 | 0.65067 | 0.7888 | 0.84756 | -0.23861 | 0.035002 | 0.083459 | 0.46412 | -1.1074 |
| TAG523-FA200 | 0.002831 | 0.011737 | 0.35147 | -1.5085 | 0.054296 | 0.137 | 0.56248 | -0.83013 | 0.018896 | 0.060465 | 0.4541 | -1.1389 |
| PI180/224_2 | 0.22609 | 0.3315 | 0.84902 | -0.23613 | 0.72364 | 0.83246 | 0.95293 | -0.06956 | 0.82737 | 0.87866 | 1.0128 | 0.018397 |
| LPC204 | 0.48582 | 0.59772 | 1.1326 | 0.1796 | 0.65214 | 0.78911 | 1.0374 | 0.052959 | 0.38461 | 0.4843 | 1.1385 | 0.18714 |
| TAG522-FA161 | 0.000296 | 0.0037 | 0.26658 | -1.9074 | 0.022312 | 0.081601 | 0.49818 | -1.0053 | 0.007495 | 0.046601 | 0.31391 | -1.6716 |
| TAG543-FA160 | 0.000143 | 0.003 | 0.39543 | -1.3385 | 0.000665 | 0.012404 | 0.50579 | -0.9834 | 0.03573 | 0.083671 | 0.56464 | -0.82461 |
| TAG544-FA221 | 0.011581 | 0.031812 | 0.28256 | -1.8234 | 0.51334 | 0.67252 | 0.69459 | -0.52577 | 0.026527 | 0.072255 | 0.34354 | -1.5415 |
| TAG542-FA182 | 0.000317 | 0.003756 | 0.41812 | -1.258 | 0.003025 | 0.024037 | 0.60375 | -0.72797 | 0.009326 | 0.047969 | 0.44329 | -1.1737 |
| PC170/181 | 0.87879 | 0.90665 | 0.9888 | -0.01625 | 0.78647 | 0.8767 | 1.0286 | 0.040722 | 0.1055 | 0.17839 | 0.81966 | -0.2869 |
| TAG567-FA161 | 0.000447 | 0.004178 | 0.26624 | -1.9092 | 0.005579 | 0.034666 | 0.39238 | -1.3497 | 0.024134 | 0.068561 | 0.42661 | -1.229 |
| PE180/225 | 0.044183 | 0.089884 | 1.6631 | 0.73384 | 0.95352 | 0.97448 | 1.0041 | 0.005965 | 0.48402 | 0.58459 | 0.83548 | -0.25932 |
| PG182/182 | 0.047194 | 0.094824 | 0.67946 | -0.55754 | 0.007372 | 0.042141 | 0.57089 | -0.8087 | 0.018948 | 0.060465 | 0.55635 | -0.84594 |
| PE181/225 | 0.80686 | 0.86251 | 0.95543 | -0.06578 | 0.044514 | 0.11925 | 0.70791 | -0.49836 | 0.062851 | 0.1225 | 0.6953 | -0.52429 |
| TAG543-FA182 | 0.000523 | 0.004461 | 0.41552 | -1.267 | 0.012712 | 0.060671 | 0.61423 | -0.70315 | 0.010943 | 0.048666 | 0.44551 | -1.1665 |
| PS180/182 | 0.00414 | 0.015228 | 0.75803 | -0.39967 | 0.000133 | 0.011362 | 0.73858 | -0.43718 | 0.014347 | 0.054983 | 0.68203 | -0.55208 |
| SMd181/202 | 0.003958 | 0.014895 | 1.336 | 0.41796 | 0.61305 | 0.75649 | 1.0307 | 0.043669 | 0.065144 | 0.12584 | 1.1411 | 0.19044 |
| PE160/225 | 0.1237 | 0.20609 | 1.443 | 0.52908 | 0.50034 | 0.66338 | 0.80914 | -0.30554 | 0.40965 | 0.50797 | 0.77201 | -0.37332 |
| TAG480-FA140 | 0.0105 | 0.029462 | 0.36194 | -1.4662 | 0.00602 | 0.035953 | 0.3652 | -1.4532 | 0.023126 | 0.066911 | 0.3796 | -1.3974 |
| PC140/140 | 0.30502 | 0.42158 | 0.77054 | -0.37607 | 0.35093 | 0.52518 | 0.7815 | -0.35569 | 0.16801 | 0.2561 | 0.69446 | -0.52604 |
| TAG482-FA181 | 0.00221 | 0.010059 | 0.17389 | -2.5238 | 0.40185 | 0.5687 | 0.81026 | -0.30355 | 0.25204 | 0.34542 | 1.5923 | 0.67111 |
| TAG564-FA202 | 0.00021 | 0.003464 | 0.34878 | -1.5196 | 0.038627 | 0.11203 | 0.604 | -0.72739 | 0.024968 | 0.06976 | 0.465 | -1.1047 |
| TAG543-FA181 | 0.001446 | 0.008048 | 0.37717 | -1.4067 | 0.95852 | 0.97589 | 1.0185 | 0.026449 | 0.027877 | 0.07359 | 0.46276 | -1.1117 |
| TAG564-FA181 | 0.004548 | 0.015981 | 0.35629 | -1.4889 | 0.36684 | 0.53787 | 0.72042 | -0.4731 | 0.047337 | 0.10137 | 0.48844 | -1.0337 |
| PE140/182 | 0.14544 | 0.23321 | 0.78581 | -0.34775 | 0.042454 | 0.11582 | 0.66281 | -0.59332 | 0.011858 | 0.050449 | 0.53123 | -0.9126 |
| TAG543-FA161 | 0.00029 | 0.0037 | 0.2937 | -1.7676 | 0.016117 | 0.067689 | 0.52609 | -0.92663 | 0.016266 | 0.057235 | 0.41174 | -1.2802 |

| | | | | | | | | | | | | |
|---|---|---|---|---|---|---|---|---|---|---|---|---|
| TAG504-FA181 | 0.000809 | 0.005848 | 0.19166 | -2.3833 | 0.005591 | 0.034666 | 0.30302 | -1.7225 | 0.010989 | 0.048666 | 0.30559 | -1.7103 |
| TAG568-FA181 | 0.15542 | 0.24345 | 0.71663 | -0.4807 | 0.11733 | 0.23869 | 0.70856 | -0.49703 | 0.035631 | 0.083671 | 0.61876 | -0.69254 |
| PEP-180/202 | 0.12652 | 0.20904 | 0.75644 | -0.4027 | 0.24098 | 0.41559 | 0.81602 | -0.29333 | 0.000298 | 0.017625 | 0.43556 | -1.1991 |
| PE160/203 | 0.43185 | 0.55451 | 1.1143 | 0.15614 | 0.098005 | 0.21057 | 0.74533 | -0.42404 | 0.02275 | 0.066415 | 0.66975 | -0.57831 |
| TAG565-FA181 | 0.004431 | 0.015936 | 0.40087 | -1.3188 | 0.042883 | 0.11582 | 0.54592 | -0.87323 | 0.077026 | 0.14125 | 0.57517 | -0.79794 |
| TAG564-FA182 | 0.008585 | 0.025755 | 0.40322 | -1.3104 | 0.23114 | 0.40559 | 0.68094 | -0.5544 | 0.036683 | 0.084088 | 0.48295 | -1.0501 |
| TAG566-FA203 | 0.053567 | 0.10427 | 0.51204 | -0.96567 | 0.047999 | 0.12572 | 0.50861 | -0.97537 | 0.091439 | 0.16002 | 0.55108 | -0.85966 |
| PEP-160/160 | 0.001354 | 0.007942 | 1.9395 | 0.9557 | 0.038141 | 0.11185 | 1.3541 | 0.43731 | 0.18627 | 0.27748 | 1.2515 | 0.32362 |
| TAG522-FA160 | 0.000421 | 0.004152 | 0.38275 | -1.3855 | 0.020433 | 0.076891 | 0.66204 | -0.59501 | 0.014268 | 0.054983 | 0.47857 | -1.0632 |
| LPE224 | 0.015059 | 0.038316 | 1.8983 | 0.92474 | 0.61632 | 0.75846 | 1.0935 | 0.12893 | 0.88648 | 0.91749 | 1.0847 | 0.11731 |
| PI161/180_2 | 0.44857 | 0.56682 | 0.90255 | -0.14792 | 0.33992 | 0.51224 | 0.88808 | -0.17124 | 0.041035 | 0.092757 | 0.72173 | -0.47048 |
| PEO-160/205 | 0.35953 | 0.47189 | 0.86629 | -0.20709 | 0.41942 | 0.58468 | 0.88776 | -0.17175 | 0.015963 | 0.057092 | 0.64839 | -0.62507 |
| TAG568-FA204 | 0.6206 | 0.71735 | 0.8513 | -0.23225 | 0.029264 | 0.091778 | 0.54735 | -0.86948 | 0.73057 | 0.79799 | 0.88386 | -0.1781 |
| PC150/181 | 0.85386 | 0.89081 | 1.0052 | 0.007511 | 0.46843 | 0.63531 | 1.1433 | 0.19324 | 0.035461 | 0.083642 | 0.72473 | -0.46449 |
| TAG546-FA161 | 0.000387 | 0.003997 | 0.25241 | -1.9862 | 0.002717 | 0.024037 | 0.33067 | -1.5966 | 0.009795 | 0.048307 | 0.35929 | -1.4768 |
| TAG522-FA182 | 0.00162 | 0.008644 | 0.45246 | -1.1441 | 0.000763 | 0.012421 | 0.50048 | -0.99862 | 0.007725 | 0.046601 | 0.46643 | -1.1003 |
| LPC140 | 0.34559 | 0.45929 | 0.87245 | -0.19686 | 0.41616 | 0.58262 | 0.88674 | -0.17341 | 0.042951 | 0.095431 | 0.64894 | -0.62385 |
| TAG501-FA140 | 0.000446 | 0.004178 | 0.27004 | -1.8887 | 0.01865 | 0.074484 | 0.48105 | -1.0557 | 0.05196 | 0.10879 | 0.54584 | -0.87346 |
| PEP-180/183 | 0.006053 | 0.019316 | 0.53394 | -0.90525 | 0.4552 | 0.6265 | 0.8774 | -0.18869 | 0.000128 | 0.009247 | 0.32088 | -1.6399 |
| TAG543-FA202 | 0.001072 | 0.006911 | 0.38544 | -1.3754 | 0.001402 | 0.018254 | 0.44625 | -1.1641 | 0.019597 | 0.061929 | 0.46783 | -1.0959 |
| PC180/140 | 0.99042 | 0.99348 | 0.95537 | -0.06587 | 0.80162 | 0.88477 | 1.011 | 0.015762 | 0.56554 | 0.65631 | 0.89155 | -0.16561 |
| TAG568-FA225 | 0.0037 | 0.014306 | 0.32891 | -1.6042 | 0.006157 | 0.036112 | 0.35143 | -1.5087 | 0.008716 | 0.047684 | 0.37061 | -1.432 |
| SMd181/161 | 0.17192 | 0.26584 | 1.1663 | 0.22198 | 0.43392 | 0.60192 | 1.0965 | 0.13296 | 0.077621 | 0.14194 | 1.2336 | 0.30287 |
| PEP-180/205 | 0.15467 | 0.24345 | 0.76901 | -0.37893 | 0.47701 | 0.64426 | 0.89687 | -0.15702 | 0.009729 | 0.048307 | 0.55606 | -0.84668 |
| PEP-160/182 | 0.044895 | 0.090765 | 0.67596 | -0.56499 | 0.92882 | 0.95415 | 1.0761 | 0.10583 | 0.007641 | 0.046601 | 0.60124 | -0.73398 |
| SMd181/200 | 0.000945 | 0.006546 | 1.1561 | 0.20929 | 0.63404 | 0.77441 | 1.0206 | 0.029486 | 0.033098 | 0.081003 | 1.1081 | 0.14807 |
| TAG541-FA181 | 0.000179 | 0.003424 | 0.37189 | -1.4271 | 0.035538 | 0.10517 | 0.67328 | -0.57072 | 0.014358 | 0.054983 | 0.43492 | -1.2012 |
| TAG565-FA203 | 0.011681 | 0.031952 | 0.44153 | -1.1794 | 0.042998 | 0.11582 | 0.53023 | -0.91531 | 0.042072 | 0.09412 | 0.4912 | -1.0256 |
| PG160/160 | 0.31807 | 0.43319 | 0.77362 | -0.3703 | 0.71628 | 0.83119 | 0.9447 | -0.08207 | 0.54368 | 0.63658 | 0.95307 | -0.06935 |
| PEP-160/225 | 0.006833 | 0.021283 | 1.5472 | 0.62966 | 0.06567 | 0.15774 | 1.2901 | 0.36751 | 0.2118 | 0.30556 | 1.2582 | 0.33135 |
| PC170/226 | 0.68734 | 0.77213 | 0.96823 | -0.04658 | 0.24159 | 0.41559 | 0.8821 | -0.18098 | 0.21381 | 0.30658 | 0.88725 | -0.17258 |
| TAG522-FA180 | 0.001509 | 0.008309 | 0.4455 | -1.1665 | 0.000762 | 0.012421 | 0.45248 | -1.1441 | 0.010731 | 0.048666 | 0.46106 | -1.117 |
| TAG545-FA203 | 0.001207 | 0.007414 | 0.422 | -1.2447 | 0.001109 | 0.016787 | 0.38355 | -1.3825 | 0.031267 | 0.078791 | 0.52858 | -0.9198 |

| | | | | | | | | | | | | |
|---|---|---|---|---|---|---|---|---|---|---|---|---|
| PC181/182 | 0.12811 | 0.21001 | 0.74311 | -0.42835 | 0.72377 | 0.83246 | 1.0557 | 0.078134 | 0.13273 | 0.21178 | 0.72487 | -0.4642 |
| TAG548-FA205 | 0.16814 | 0.26062 | 0.58288 | -0.77872 | 0.042813 | 0.11582 | 0.39128 | -1.3537 | 0.096013 | 0.16623 | 0.48192 | -1.0531 |
| TAG562-FA160 | 0.012548 | 0.03358 | 0.36846 | -1.4404 | 0.30377 | 0.48233 | 0.75296 | -0.40936 | 0.062506 | 0.1222 | 0.56583 | -0.82157 |
| PG160/180_2 | 0.89765 | 0.92172 | 1.0114 | 0.016346 | 0.25055 | 0.42476 | 0.82525 | -0.2771 | 0.092344 | 0.16117 | 0.71969 | -0.47454 |
| PEP-160/205 | 0.47449 | 0.58837 | 0.87208 | -0.19746 | 0.85406 | 0.91061 | 1.0649 | 0.090746 | 0.27107 | 0.3646 | 0.80529 | -0.31241 |
| TAG543-FA201 | 6.17E-05 | 0.003 | 0.38663 | -1.371 | 0.001342 | 0.018197 | 0.55524 | -0.84881 | 0.033795 | 0.081184 | 0.57477 | -0.79893 |
| PEP-181/182 | 0.12442 | 0.20609 | 0.73026 | -0.45351 | 0.99084 | 0.99094 | 1.0292 | 0.04154 | 0.008224 | 0.04704 | 0.55341 | -0.85357 |
| PG160/180 | 0.66001 | 0.74985 | 1.1028 | 0.14118 | 0.30303 | 0.48233 | 0.83643 | -0.25769 | 0.083333 | 0.14904 | 0.68478 | -0.54628 |
| PEO-160/182 | 0.005045 | 0.017285 | 0.6689 | -0.58013 | 0.23371 | 0.40899 | 0.85452 | -0.22682 | 0.00072 | 0.029311 | 0.51959 | -0.94455 |
| TAG483-FA160 | 0.002183 | 0.010006 | 0.2197 | -2.1864 | 0.001593 | 0.018858 | 0.20746 | -2.2691 | 0.010018 | 0.048666 | 0.29635 | -1.7546 |
| TAG547-FA161 | 0.01665 | 0.041213 | 0.39684 | -1.3334 | 0.0112 | 0.055235 | 0.38441 | -1.3793 | 0.038541 | 0.087422 | 0.45928 | -1.1225 |
| TAG565-FA224 | 0.013601 | 0.035569 | 0.44596 | -1.165 | 0.042343 | 0.11582 | 0.533 | -0.9078 | 0.1171 | 0.19299 | 0.60423 | -0.72684 |
| TAG502-FA160 | 0.0229 | 0.051945 | 0.50121 | -0.99651 | 0.015593 | 0.067071 | 0.4605 | -1.1187 | 0.050995 | 0.10849 | 0.54424 | -0.87768 |
| PE180/150 | 0.13698 | 0.22072 | 1.1401 | 0.18922 | 0.69511 | 0.81561 | 0.96134 | -0.05688 | 0.12179 | 0.19822 | 0.8595 | -0.21844 |
| LPE180 | 0.001777 | 0.009115 | 1.5875 | 0.66673 | 0.49445 | 0.65961 | 1.0844 | 0.11684 | 0.14827 | 0.23147 | 0.82332 | -0.28048 |
| TAG502-FA182 | 0.037913 | 0.079994 | 0.49869 | -1.0038 | 0.028973 | 0.091778 | 0.46009 | -1.12 | 0.10596 | 0.1787 | 0.58404 | -0.77587 |
| PEO-180/224 | 0.19225 | 0.29354 | 1.4877 | 0.57305 | 0.55617 | 0.71414 | 1.1527 | 0.20508 | 0.76913 | 0.83035 | 1.0751 | 0.10446 |
| TAG483-FA181 | 0.012977 | 0.034203 | 0.30227 | -1.7261 | 0.41774 | 0.58357 | 0.90449 | -0.14482 | 0.31307 | 0.41008 | 1.2594 | 0.3327 |
| PE182/183 | 0.23202 | 0.33867 | 0.75954 | -0.3968 | 0.00179 | 0.019633 | 0.30787 | -1.6996 | 0.016049 | 0.057092 | 0.40952 | -1.288 |
| PG160/181 | 0.37921 | 0.49374 | 0.86293 | -0.21269 | 0.28369 | 0.46198 | 0.83616 | -0.25816 | 0.051632 | 0.10879 | 0.68127 | -0.5537 |
| PEP-180/180 | 0.10197 | 0.17702 | 0.75426 | -0.40687 | 0.92552 | 0.95415 | 0.99202 | -0.01156 | 0.000705 | 0.029311 | 0.49702 | -1.0086 |
| PS160/180 | 0.021408 | 0.049247 | 1.4019 | 0.48738 | 0.15553 | 0.30135 | 0.80095 | -0.32022 | 0.48788 | 0.58817 | 1.0581 | 0.081493 |
| PG180/181_2 | 0.63387 | 0.7274 | 0.892 | -0.16489 | 0.31107 | 0.48797 | 0.78456 | -0.35004 | 0.075148 | 0.13938 | 0.62993 | -0.66674 |
| PC160/160 | 0.001256 | 0.00757 | 1.602 | 0.67986 | 0.049127 | 0.12742 | 1.2486 | 0.32029 | 0.058604 | 0.11599 | 1.243 | 0.31378 |
| TAG562-FA201 | 0.032737 | 0.07057 | 0.48688 | -1.0384 | 0.86205 | 0.91251 | 0.93586 | -0.09563 | 0.075657 | 0.13954 | 0.57816 | -0.79046 |
| PEO-160/224 | 0.19739 | 0.29884 | 1.4653 | 0.55115 | 0.48859 | 0.65312 | 1.1736 | 0.23098 | 0.92081 | 0.93517 | 1.0043 | 0.006123 |
| PE170/204 | 0.69276 | 0.77433 | 0.92647 | -0.11018 | 0.073572 | 0.17167 | 0.66761 | -0.58291 | 0.007172 | 0.046601 | 0.53908 | -0.89144 |
| TAG566-FA224 | 0.013734 | 0.03562 | 0.45655 | -1.1311 | 0.026164 | 0.088469 | 0.4779 | -1.0652 | 0.10842 | 0.18145 | 0.60465 | -0.72583 |
| PE182/182 | 0.34528 | 0.45929 | 0.82492 | -0.27767 | 0.40082 | 0.56848 | 0.83294 | -0.26372 | 0.028053 | 0.073639 | 0.52788 | -0.92173 |
| PC181/225 | 0.57329 | 0.67734 | 0.92815 | -0.10758 | 0.097638 | 0.21047 | 0.81044 | -0.30321 | 0.56557 | 0.65631 | 0.93843 | -0.09168 |
| PI161/180 | 0.24802 | 0.35486 | 0.83247 | -0.26453 | 0.92301 | 0.95415 | 0.98541 | -0.02121 | 0.20261 | 0.29773 | 0.82381 | -0.27963 |
| LPE225 | 0.002383 | 0.01041 | 2.1437 | 1.1001 | 0.078631 | 0.1773 | 1.484 | 0.56952 | 0.314 | 0.41047 | 1.1442 | 0.19428 |
| HCER200 | 0.012814 | 0.033911 | 1.4735 | 0.55927 | 0.58208 | 0.73437 | 1.041 | 0.057927 | 0.68027 | 0.75975 | 0.97425 | -0.03763 |

| | | | | | | | | | | | | |
|---|---|---|---|---|---|---|---|---|---|---|---|---|
| LPC203 | 0.66184 | 0.75062 | 0.92302 | -0.11557 | 0.3233 | 0.50231 | 0.82233 | -0.28221 | 0.61779 | 0.69945 | 0.91927 | -0.12144 |
| TAG544-FA160 | 0.001035 | 0.006803 | 0.45772 | -1.1275 | 0.000631 | 0.012404 | 0.43797 | -1.1911 | 0.030644 | 0.077926 | 0.55005 | -0.86237 |
| PE180/203 | 0.41083 | 0.5296 | 1.0957 | 0.13191 | 0.099909 | 0.21255 | 0.76311 | -0.39004 | 0.10683 | 0.1797 | 0.76677 | -0.38313 |
| TAG546-FA204 | 0.020109 | 0.04726 | 0.57937 | -0.78744 | 0.014247 | 0.063963 | 0.52932 | -0.91779 | 0.34798 | 0.44769 | 0.81448 | -0.29605 |
| TAG568-FA161 | 0.000356 | 0.00386 | 0.34561 | -1.5328 | 0.015528 | 0.067071 | 0.42743 | -1.2262 | 0.00205 | 0.046601 | 0.33485 | -1.5784 |
| PEO-180/182 | 0.003234 | 0.013186 | 0.69329 | -0.52846 | 0.019835 | 0.07686 | 0.77745 | -0.36317 | 3.6E-05 | 0.005865 | 0.40836 | -1.2921 |
| PS180/180 | 0.042792 | 0.087651 | 1.3952 | 0.4805 | 0.29351 | 0.47179 | 0.83737 | -0.25606 | 0.48177 | 0.58295 | 1.0744 | 0.10359 |
| TAG561-FA160 | 0.002125 | 0.009937 | 0.34955 | -1.5164 | 0.72332 | 0.83246 | 1.1422 | 0.19179 | 0.0794 | 0.14479 | 0.59034 | -0.76039 |
| PEP-160/203 | 0.46233 | 0.57659 | 0.86471 | -0.2097 | 0.78147 | 0.87412 | 1.0238 | 0.033981 | 0.30406 | 0.4007 | 0.82313 | -0.28081 |
| TAG563-FA201 | 0.009615 | 0.027573 | 0.39598 | -1.3365 | 0.78952 | 0.8786 | 1.0942 | 0.12982 | 0.10008 | 0.171 | 0.60552 | -0.72377 |
| PE181/224 | 0.97424 | 0.98331 | 0.91375 | -0.13014 | 0.35258 | 0.52644 | 0.70414 | -0.50606 | 0.23071 | 0.3258 | 0.65384 | -0.61299 |
| TAG521-FA180 | 0.001317 | 0.007796 | 0.42075 | -1.249 | 0.003334 | 0.024949 | 0.55776 | -0.84227 | 0.006439 | 0.046601 | 0.41392 | -1.2726 |
| TAG547-FA205 | 0.22065 | 0.32572 | 0.60952 | -0.71426 | 0.24214 | 0.41559 | 0.63964 | -0.64467 | 0.5324 | 0.62561 | 0.728 | -0.45799 |
| PG160/181_2 | 0.58246 | 0.68179 | 0.91622 | -0.12623 | 0.26633 | 0.44118 | 0.8352 | -0.25981 | 0.061877 | 0.12133 | 0.70258 | -0.50927 |
| PE170/182 | 0.34657 | 0.45951 | 0.7608 | -0.3944 | 0.063697 | 0.15531 | 0.57169 | -0.80669 | 0.005428 | 0.046601 | 0.3854 | -1.3756 |
| PC200/203 | 0.84685 | 0.89081 | 0.98362 | -0.02382 | 0.19458 | 0.35483 | 0.75361 | -0.40811 | 0.06899 | 0.13094 | 0.66658 | -0.58516 |
| TAG547-FA204 | 0.014216 | 0.03658 | 0.49941 | -1.0017 | 0.003806 | 0.026644 | 0.4156 | -1.2667 | 0.068823 | 0.13094 | 0.59236 | -0.75545 |
| TAG526-FA204 | 0.00403 | 0.014972 | 0.39062 | -1.3561 | 0.000613 | 0.012404 | 0.3447 | -1.5366 | 0.052137 | 0.10879 | 0.54735 | -0.86947 |
| TAG542-FA181 | 0.003714 | 0.014306 | 0.42289 | -1.2416 | 0.75611 | 0.85888 | 1.077 | 0.10702 | 0.033347 | 0.081004 | 0.47971 | -1.0598 |
| PI180/183 | 0.083751 | 0.14978 | 0.7628 | -0.39063 | 0.18407 | 0.34335 | 0.86608 | -0.20742 | 0.020329 | 0.062721 | 0.68781 | -0.53993 |
| TAG520-FA160 | 0.011101 | 0.030881 | 0.41468 | -1.2699 | 0.002688 | 0.024037 | 0.38445 | -1.3791 | 0.00469 | 0.046601 | 0.31658 | -1.6593 |
| TAG521-FA181 | 0.0003 | 0.0037 | 0.38066 | -1.3934 | 0.008169 | 0.044316 | 0.61354 | -0.70477 | 0.005945 | 0.046601 | 0.40737 | -1.2956 |
| TAG544-FA203 | 0.00057 | 0.004633 | 0.40366 | -1.3088 | 0.004831 | 0.031448 | 0.4991 | -1.0026 | 0.024243 | 0.068561 | 0.51411 | -0.95986 |
| PEO-180/225 | 0.00236 | 0.01041 | 1.6141 | 0.69074 | 0.013473 | 0.061335 | 1.3256 | 0.40667 | 0.20439 | 0.29968 | 1.3751 | 0.45959 |
| TAG546-FA225 | 0.00792 | 0.024092 | 0.2778 | -1.8479 | 0.020404 | 0.076891 | 0.31861 | -1.6501 | 0.030041 | 0.0773 | 0.34898 | -1.5188 |
| TAG542-FA202 | 0.000795 | 0.005812 | 0.43853 | -1.1893 | 0.000552 | 0.012404 | 0.49231 | -1.0224 | 0.019699 | 0.061952 | 0.51512 | -0.95702 |
| TAG564-FA203 | 0.003646 | 0.014296 | 0.41527 | -1.2679 | 0.070801 | 0.16822 | 0.62369 | -0.68111 | 0.03612 | 0.084041 | 0.52353 | -0.93366 |
| PC160/202 | 0.51468 | 0.61818 | 1.0875 | 0.12104 | 0.37702 | 0.54785 | 0.93419 | -0.09821 | 0.8359 | 0.88627 | 1.0632 | 0.088402 |
| TAG563-FA180 | 0.000477 | 0.004198 | 0.41423 | -1.2715 | 0.001017 | 0.015768 | 0.51798 | -0.94902 | 0.016769 | 0.057235 | 0.49118 | -1.0257 |
| PE180/160 | 0.003711 | 0.014306 | 1.639 | 0.71283 | 0.68116 | 0.80645 | 0.92876 | -0.10662 | 0.0315 | 0.07887 | 0.68735 | -0.54089 |
| FFA205 | 0.43411 | 0.55631 | 0.81188 | -0.30067 | 0.17825 | 0.33635 | 0.7227 | -0.46853 | 0.21123 | 0.30556 | 0.72175 | -0.47043 |
| PE180/201 | 0.26906 | 0.38127 | 1.1167 | 0.15927 | 0.2747 | 0.45274 | 0.88146 | -0.18203 | 0.017734 | 0.058013 | 0.63494 | -0.65532 |
| PC180/203 | 0.82238 | 0.87622 | 1.0538 | 0.075556 | 0.46363 | 0.63351 | 0.84624 | -0.24086 | 0.88544 | 0.91749 | 0.95942 | -0.05977 |

| | | | | | | | | | | | | |
|---|---|---|---|---|---|---|---|---|---|---|---|---|
| PG180/180 | 0.70358 | 0.78429 | 0.91724 | -0.12462 | 0.30507 | 0.48321 | 0.79381 | -0.33313 | 0.074918 | 0.13938 | 0.6504 | -0.62061 |
| PEP-180/204 | 0.39629 | 0.51495 | 0.89812 | -0.15502 | 0.63401 | 0.77441 | 1.0471 | 0.066354 | 0.054883 | 0.11165 | 0.7914 | -0.33753 |
| PEO-180/160 | 0.002152 | 0.009937 | 1.3913 | 0.47639 | 0.3272 | 0.50466 | 1.067 | 0.093525 | 0.041645 | 0.09381 | 0.84149 | -0.24898 |
| TAG541-FA201 | 0.00866 | 0.02586 | 0.4092 | -1.2891 | 0.003774 | 0.026644 | 0.5448 | -0.87621 | 0.068886 | 0.13094 | 0.63958 | -0.64481 |
| PC180/161 | 0.79386 | 0.85281 | 0.9259 | -0.11107 | 0.34132 | 0.51316 | 0.80712 | -0.30915 | 0.22121 | 0.31374 | 0.75848 | -0.39882 |
| TAG568-FA205 | 0.85252 | 0.89081 | 1.0874 | 0.12086 | 0.56388 | 0.72119 | 0.93862 | -0.09138 | 0.90642 | 0.92867 | 0.98606 | -0.02025 |
| PI180/204 | 0.3457 | 0.45929 | 0.90005 | -0.15192 | 0.91258 | 0.94903 | 1.008 | 0.011498 | 0.12627 | 0.20447 | 1.1538 | 0.20637 |
| PI180/204_2 | 0.52928 | 0.63339 | 0.93104 | -0.10309 | 0.95787 | 0.97589 | 0.99506 | -0.00714 | 0.087152 | 0.15334 | 1.2086 | 0.27332 |
| TAG546-FA205 | 0.21643 | 0.32094 | 0.618 | -0.69431 | 0.3119 | 0.4881 | 0.69416 | -0.52666 | 0.50003 | 0.59728 | 0.74096 | -0.43254 |
| PC170/182 | 0.15557 | 0.24345 | 0.86307 | -0.21245 | 0.35895 | 0.5318 | 0.91623 | -0.12621 | 0.032742 | 0.080433 | 0.74468 | -0.4253 |
| TAG542-FA201 | 0.001647 | 0.008715 | 0.42964 | -1.2188 | 0.095672 | 0.20761 | 0.73961 | -0.43516 | 0.064037 | 0.12444 | 0.62581 | -0.67621 |
| TAG521-FA160 | 0.002602 | 0.011086 | 0.43789 | -1.1914 | 0.027509 | 0.090199 | 0.62057 | -0.68834 | 0.010328 | 0.048666 | 0.43494 | -1.2011 |
| TAG482-FA120 | 0.015068 | 0.038316 | 0.29683 | -1.7523 | 0.61339 | 0.75649 | 1.281 | 0.35731 | 0.59326 | 0.68115 | 3.5657 | 1.8342 |
| TAG564-FA180 | 0.000465 | 0.004198 | 0.3955 | -1.3382 | 0.002919 | 0.024037 | 0.50268 | -0.99229 | 0.011983 | 0.050449 | 0.47204 | -1.083 |
| LPC180 | 0.51198 | 0.61608 | 0.95043 | -0.07334 | 0.45792 | 0.62891 | 1.0408 | 0.057671 | 0.78716 | 0.84701 | 0.97911 | -0.03046 |
| PC181/205 | 0.32217 | 0.43513 | 0.79624 | -0.32873 | 0.56225 | 0.72052 | 0.91573 | -0.12701 | 0.45385 | 0.55747 | 0.84496 | -0.24304 |
| TAG567-FA205 | 0.43946 | 0.55921 | 0.89789 | -0.15539 | 0.5309 | 0.69123 | 0.95995 | -0.05897 | 0.28809 | 0.38275 | 0.80956 | -0.30479 |
| PE181/201 | 0.71878 | 0.79988 | 0.93855 | -0.09149 | 0.72376 | 0.83246 | 1.1922 | 0.25358 | 0.042738 | 0.095283 | 0.56869 | -0.81428 |
| TAG501-FA161 | 0.004489 | 0.015981 | 0.31194 | -1.6807 | 0.029229 | 0.091778 | 0.43823 | -1.1902 | 0.004599 | 0.046601 | 0.27931 | -1.84 |
| PG180/181 | 0.72129 | 0.80129 | 0.91749 | -0.12424 | 0.33716 | 0.51128 | 0.7891 | -0.34173 | 0.075124 | 0.13938 | 0.62375 | -0.68095 |
| PI180/205_2 | 0.15077 | 0.23998 | 0.76973 | -0.37758 | 0.35982 | 0.5318 | 0.83987 | -0.25177 | 0.17412 | 0.26179 | 0.75785 | -0.40001 |
| TAG541-FA180 | 0.001063 | 0.006911 | 0.41407 | -1.2721 | 0.023215 | 0.083031 | 0.56176 | -0.83198 | 0.008526 | 0.04704 | 0.38382 | -1.3815 |
| PC200/205 | 0.62754 | 0.72305 | 0.92849 | -0.10704 | 0.40016 | 0.56848 | 0.85954 | -0.21837 | 0.21927 | 0.31167 | 0.73077 | -0.45251 |
| PC150/226 | 0.34938 | 0.46229 | 0.88032 | -0.1839 | 0.0273 | 0.090199 | 0.74241 | -0.42971 | 0.054284 | 0.11133 | 0.75118 | -0.41278 |
| PC200/181 | 0.57968 | 0.68117 | 0.95222 | -0.07063 | 0.70828 | 0.82485 | 1.0727 | 0.10131 | 0.004141 | 0.046601 | 0.57773 | -0.79153 |
| TAG527-FA160 | 0.006124 | 0.019448 | 0.3847 | -1.3782 | 0.003411 | 0.025005 | 0.35671 | -1.4872 | 0.007614 | 0.046601 | 0.34236 | -1.5464 |
| LPC181 | 0.45493 | 0.57173 | 0.90011 | -0.15183 | 0.11532 | 0.23637 | 1.2622 | 0.33598 | 0.3242 | 0.42043 | 0.86256 | -0.2133 |
| PE181/181 | 0.43095 | 0.55445 | 0.81535 | -0.2945 | 0.36025 | 0.5318 | 1.98 | 0.98549 | 0.030611 | 0.077926 | 0.4825 | -1.0514 |
| SMd181/120 | 0.57495 | 0.67807 | 0.84438 | -0.24404 | 0.33967 | 0.51224 | 0.71918 | -0.47557 | 0.69323 | 0.76881 | 0.91427 | -0.12931 |
| PE160/161 | 0.97307 | 0.98331 | 0.89705 | -0.15673 | 0.20573 | 0.371 | 0.49476 | -1.0152 | 0.08363 | 0.14916 | 0.38672 | -1.3706 |
| PEO-180/183 | 0.015885 | 0.039471 | 0.71677 | -0.48042 | 0.002011 | 0.019837 | 0.59764 | -0.74264 | 3.25E-05 | 0.005865 | 0.38591 | -1.3737 |
| TAG568-FA182 | 0.02514 | 0.056241 | 0.60002 | -0.73692 | 0.04616 | 0.12215 | 0.56599 | -0.82116 | 0.19485 | 0.28829 | 0.76664 | -0.38339 |
| TAG521-FA201 | 0.001436 | 0.008048 | 0.34157 | -1.5498 | 0.001761 | 0.019633 | 0.44838 | -1.1572 | 0.053349 | 0.11037 | 0.58202 | -0.78086 |

| | | | | | | | | | | | | |
|---|---|---|---|---|---|---|---|---|---|---|---|---|
| TAG543-FA203 | 0.005275 | 0.017885 | 0.603 | -0.72976 | 0.001933 | 0.019775 | 0.4865 | -1.0395 | 0.045816 | 0.098436 | 0.63058 | -0.66524 |
| PE181/161 | 0.30773 | 0.42354 | 0.89389 | -0.16184 | 0.80586 | 0.88755 | 1.084 | 0.11636 | 4.98E-05 | 0.00587 | 0.57366 | -0.80173 |
| TAG501-FA181 | 0.015794 | 0.039393 | 0.47111 | -1.0859 | 0.027889 | 0.090496 | 0.48494 | -1.0441 | 0.031707 | 0.079084 | 0.47265 | -1.0812 |
| TAG566-FA204 | 0.021495 | 0.049273 | 0.59365 | -0.7523 | 0.1812 | 0.33995 | 0.75896 | -0.3979 | 0.23898 | 0.33399 | 0.77974 | -0.35894 |
| LPC225 | 0.07806 | 0.14195 | 1.3376 | 0.41964 | 0.30668 | 0.48458 | 1.1496 | 0.20111 | 0.3274 | 0.42289 | 1.1213 | 0.1652 |
| PE160/183 | 0.74139 | 0.81665 | 1.1471 | 0.19798 | 0.056929 | 0.14145 | 0.49296 | -1.0205 | 0.005483 | 0.046601 | 0.37373 | -1.4199 |
| HCER160 | 0.64619 | 0.73673 | 1.0642 | 0.089702 | 0.10521 | 0.22165 | 0.79219 | -0.33608 | 0.025357 | 0.070244 | 0.71783 | -0.47829 |
| TAG547-FA225 | 0.004167 | 0.015241 | 0.3009 | -1.7326 | 0.004383 | 0.030036 | 0.27898 | -1.8418 | 0.009455 | 0.047969 | 0.32417 | -1.6252 |
| TAG564-FA160 | 0.027745 | 0.061227 | 0.55083 | -0.86033 | 0.02087 | 0.078084 | 0.51269 | -0.96385 | 0.055817 | 0.1127 | 0.55898 | -0.83913 |
| PG160/202_2 | 0.85811 | 0.8938 | 0.94657 | -0.07922 | 0.13132 | 0.26144 | 0.71952 | -0.47489 | 0.086599 | 0.15278 | 0.65709 | -0.60583 |
| TAG540-FA180 | 0.012481 | 0.033575 | 0.38337 | -1.3832 | 0.003419 | 0.025005 | 0.34956 | -1.5164 | 0.005266 | 0.046601 | 0.27095 | -1.8839 |
| PE180/161 | 0.98536 | 0.99071 | 1.0668 | 0.093255 | 0.73026 | 0.83697 | 0.87221 | -0.19725 | 0.045041 | 0.097739 | 0.53899 | -0.89166 |
| PE180/226 | 0.017464 | 0.042742 | 1.5393 | 0.62227 | 0.77036 | 0.86883 | 0.9523 | -0.07051 | 0.92048 | 0.93517 | 0.96061 | -0.05799 |
| TAG563-FA160 | 0.033748 | 0.072509 | 0.58637 | -0.77011 | 0.043055 | 0.11582 | 0.64808 | -0.62576 | 0.10311 | 0.17572 | 0.69878 | -0.51709 |
| SMd181/226 | 0.03764 | 0.079816 | 1.2379 | 0.3079 | 0.15538 | 0.30135 | 0.86944 | -0.20184 | 0.74222 | 0.80935 | 0.97242 | -0.04035 |
| PEO-180/205 | 0.5764 | 0.67855 | 0.9095 | -0.13686 | 0.8181 | 0.8951 | 0.98158 | -0.02682 | 0.099083 | 0.16975 | 0.7001 | -0.51436 |
| HCER220 | 0.15555 | 0.24345 | 1.3072 | 0.38645 | 0.26248 | 0.43899 | 1.1493 | 0.20071 | 0.38592 | 0.48501 | 0.90835 | -0.13867 |
| TAG545-FA204 | 0.036602 | 0.077868 | 0.67263 | -0.57212 | 0.065076 | 0.15749 | 0.69264 | -0.52982 | 0.63767 | 0.71697 | 0.93811 | -0.09217 |
| SMd181/220 | 0.43604 | 0.55769 | 1.0757 | 0.10534 | 0.088728 | 0.19514 | 0.87369 | -0.19481 | 0.62255 | 0.70361 | 0.97445 | -0.03733 |
| TAG567-FA180 | 0.018102 | 0.043486 | 0.53324 | -0.90715 | 0.033112 | 0.099335 | 0.5394 | -0.89058 | 0.0333 | 0.081004 | 0.57011 | -0.81068 |
| PC160/182 | 0.032121 | 0.069703 | 1.1919 | 0.25331 | 0.12314 | 0.24667 | 1.127 | 0.17246 | 0.96038 | 0.96528 | 0.991 | -0.01304 |
| TAG567-FA225 | 0.003921 | 0.01484 | 0.40354 | -1.3092 | 0.0203 | 0.076891 | 0.45864 | -1.1246 | 0.027796 | 0.07359 | 0.51531 | -0.95649 |
| TAG545-FA160 | 0.006657 | 0.020937 | 0.52667 | -0.92502 | 0.007575 | 0.042511 | 0.49967 | -1.001 | 0.053781 | 0.11079 | 0.62834 | -0.67038 |
| PEP-181/225 | 0.007352 | 0.022471 | 1.621 | 0.69686 | 0.098344 | 0.2106 | 1.2741 | 0.34951 | 0.13755 | 0.21788 | 1.2093 | 0.27417 |
| PEP-182/182 | 0.10733 | 0.1834 | 0.78509 | -0.34908 | 0.65788 | 0.79035 | 1.1293 | 0.17546 | 0.010921 | 0.048666 | 0.65871 | -0.60229 |
| TAG546-FA226 | 0.77682 | 0.84701 | 1.1484 | 0.19962 | 0.49639 | 0.65986 | 0.89586 | -0.15866 | 0.91143 | 0.92867 | 1.0713 | 0.099426 |
| PE160/226 | 0.024199 | 0.054443 | 1.4764 | 0.56212 | 0.54261 | 0.70087 | 0.9096 | -0.13669 | 0.85824 | 0.90589 | 0.95908 | -0.06028 |
| FFA203 | 0.86681 | 0.89999 | 1.0224 | 0.031948 | 0.004567 | 0.030649 | 0.67283 | -0.5717 | 0.9286 | 0.94015 | 1.0083 | 0.011917 |
| LPE202 | 0.24716 | 0.35441 | 1.1674 | 0.22331 | 0.58613 | 0.73804 | 0.93039 | -0.10409 | 0.15702 | 0.24166 | 0.78488 | -0.34946 |
| SMd181/205 | 0.59318 | 0.69205 | 1.0311 | 0.044196 | 0.24259 | 0.41559 | 0.93523 | -0.09661 | 0.96964 | 0.97113 | 1.0106 | 0.015194 |
| TAG566-FA160 | 0.005828 | 0.019123 | 0.38318 | -1.3839 | 0.018442 | 0.074109 | 0.42857 | -1.2224 | 0.027115 | 0.073032 | 0.45261 | -1.1437 |
| PG160/202 | 0.97386 | 0.98331 | 0.97805 | -0.03202 | 0.093258 | 0.20352 | 0.71588 | -0.48221 | 0.091374 | 0.16002 | 0.68297 | -0.5501 |
| TAG547-FA226 | 0.012814 | 0.033911 | 0.3449 | -1.5357 | 0.023648 | 0.083031 | 0.44913 | -1.1548 | 0.01523 | 0.056335 | 0.40797 | -1.2935 |

| | | | | | | | | | | | | |
|---|---|---|---|---|---|---|---|---|---|---|---|---|
| SM241 | 0.14769 | 0.23565 | 1.0604 | 0.084614 | 0.36565 | 0.53733 | 0.94966 | -0.07452 | 0.49513 | 0.5947 | 1.0261 | 0.037157 |
| PEP-181/205 | 0.65697 | 0.7477 | 0.93618 | -0.09514 | 0.54871 | 0.70735 | 0.91407 | -0.12963 | 0.13374 | 0.21263 | 0.76541 | -0.3857 |
| HCER241 | 0.45186 | 0.56897 | 1.1084 | 0.14842 | 0.28386 | 0.46198 | 0.92191 | -0.11731 | 0.044318 | 0.097302 | 0.85489 | -0.22619 |
| FFA226 | 0.00942 | 0.027134 | 1.4695 | 0.55535 | 0.68134 | 0.80645 | 0.93694 | -0.09397 | 0.10949 | 0.18276 | 1.3458 | 0.42844 |
| FFA220 | 0.23751 | 0.3436 | 1.2528 | 0.32519 | 0.92113 | 0.95415 | 0.98863 | -0.0165 | 0.148 | 0.23147 | 0.71203 | -0.48999 |
| PEP-180/160 | 0.024253 | 0.054443 | 1.4296 | 0.51557 | 0.51016 | 0.6723 | 1.0707 | 0.098528 | 0.82406 | 0.87657 | 1.0287 | 0.040879 |
| PE160/201 | 0.7853 | 0.85121 | 1.0355 | 0.050322 | 0.086965 | 0.19259 | 0.72652 | -0.46092 | 0.036147 | 0.084041 | 0.58923 | -0.7631 |
| TAG565-FA180 | 0.005651 | 0.01877 | 0.53867 | -0.89252 | 0.022725 | 0.082479 | 0.63577 | -0.65343 | 0.082846 | 0.14857 | 0.67734 | -0.56204 |
| PG161/182_2 | 0.4653 | 0.57918 | 1.1284 | 0.17434 | 0.32835 | 0.50466 | 1.1568 | 0.21019 | 0.21215 | 0.30556 | 0.70816 | -0.49785 |
| TAG566-FA181 | 0.002304 | 0.01041 | 0.41006 | -1.2861 | 0.01293 | 0.060671 | 0.49883 | -1.0034 | 0.02755 | 0.07359 | 0.50192 | -0.99446 |
| PC140/225 | 0.66365 | 0.75136 | 1.0668 | 0.093297 | 0.53717 | 0.6966 | 1.0688 | 0.096048 | 0.31783 | 0.41464 | 0.82728 | -0.27355 |
| PC180/181 | 0.78953 | 0.85272 | 1.0362 | 0.051349 | 0.26299 | 0.43899 | 1.1068 | 0.14641 | 0.21707 | 0.31058 | 0.84236 | -0.24749 |
| TAG501-FA160 | 0.015713 | 0.039342 | 0.50076 | -0.99781 | 0.033041 | 0.099335 | 0.55681 | -0.84473 | 0.03827 | 0.08711 | 0.53628 | -0.89894 |
| PC160/224 | 0.092313 | 0.1633 | 1.4732 | 0.559 | 0.30269 | 0.48233 | 1.1823 | 0.24163 | 0.69646 | 0.77108 | 1.0937 | 0.12926 |
| TAG481-FA161 | 0.017231 | 0.04233 | 0.30876 | -1.6955 | 0.071071 | 0.16824 | 0.4147 | -1.2699 | 0.022922 | 0.066617 | 0.3219 | -1.6353 |
| PEP-181/183 | 0.60169 | 0.70071 | 0.87521 | -0.1923 | 0.96967 | 0.98326 | 0.96999 | -0.04396 | 0.067207 | 0.12906 | 0.63132 | -0.66355 |
| PEO-160/203 | 0.99395 | 0.99548 | 0.97548 | -0.03581 | 0.57079 | 0.72575 | 0.89353 | -0.16241 | 0.061267 | 0.12064 | 0.70866 | -0.49683 |
| PC181/201 | 0.44661 | 0.56564 | 0.86889 | -0.20275 | 0.90706 | 0.94613 | 1.0137 | 0.0197 | 0.13046 | 0.20965 | 0.70007 | -0.51442 |
| TAG548-FA204 | 0.10476 | 0.17994 | 0.6739 | -0.5694 | 0.003251 | 0.024827 | 0.44325 | -1.1738 | 0.1314 | 0.21018 | 0.69284 | -0.52941 |
| PEP-181/203 | 0.94727 | 0.96657 | 0.9879 | -0.01756 | 0.84671 | 0.90643 | 0.93769 | -0.09283 | 0.24489 | 0.33992 | 0.77473 | -0.36824 |
| PE180/180 | 0.006557 | 0.020722 | 1.6141 | 0.69071 | 0.055224 | 0.13881 | 1.3944 | 0.47969 | 0.045603 | 0.098303 | 1.3302 | 0.41161 |
| TAG565-FA160 | 0.013715 | 0.03562 | 0.47275 | -1.0809 | 0.023723 | 0.083031 | 0.50375 | -0.98922 | 0.055169 | 0.11188 | 0.54911 | -0.86485 |
| SMd181/222 | 0.44928 | 0.56682 | 1.0355 | 0.050306 | 0.032946 | 0.099335 | 0.88847 | -0.1706 | 0.35623 | 0.45741 | 1.0475 | 0.066944 |
| PC181/204 | 0.96824 | 0.98331 | 0.97926 | -0.03023 | 0.51282 | 0.67252 | 0.84184 | -0.24838 | 0.81571 | 0.87197 | 0.95314 | -0.06923 |
| TAG481-FA180 | 0.009135 | 0.026627 | 0.33342 | -1.5846 | 0.9299 | 0.95415 | 1.2084 | 0.27308 | 0.51404 | 0.61065 | 3.5617 | 1.8326 |
| PC181/181 | 0.43981 | 0.55921 | 0.84131 | -0.24928 | 0.18983 | 0.34909 | 1.6012 | 0.67917 | 0.12658 | 0.20447 | 0.68557 | -0.54463 |
| HCER140 | 0.2694 | 0.38127 | 0.80668 | -0.30994 | 0.031626 | 0.096661 | 0.62077 | -0.68786 | 0.003673 | 0.046601 | 0.53844 | -0.89315 |
| TAG542-FA180 | 0.000303 | 0.0037 | 0.44877 | -1.1559 | 0.40433 | 0.57097 | 0.88709 | -0.17285 | 0.017307 | 0.057485 | 0.45784 | -1.1271 |
| TAG545-FA225 | 0.85185 | 0.89081 | 0.9847 | -0.02224 | 0.30157 | 0.48233 | 0.73112 | -0.45182 | 0.50227 | 0.59777 | 0.74969 | -0.41563 |
| PE170/225 | 0.93043 | 0.95237 | 0.96199 | -0.0559 | 0.22905 | 0.40519 | 0.63704 | -0.65055 | 0.041865 | 0.093981 | 0.48213 | -1.0525 |
| PEO-180/203 | 0.62148 | 0.71735 | 0.93186 | -0.10182 | 0.21302 | 0.38203 | 0.84696 | -0.23964 | 0.003878 | 0.046601 | 0.60318 | -0.72935 |
| PA180/181 | 0.58328 | 0.68179 | 0.92564 | -0.11147 | 0.061913 | 0.15152 | 1.5432 | 0.62592 | 0.067067 | 0.12906 | 0.71917 | -0.47559 |
| PEP-160/202 | 0.96405 | 0.98216 | 1.0085 | 0.012149 | 0.65021 | 0.7888 | 1.0845 | 0.11709 | 0.24658 | 0.34153 | 0.83112 | -0.26687 |

| | | | | | | | | | | | | |
|---|---|---|---|---|---|---|---|---|---|---|---|---|
| SMd181/224 | 0.22476 | 0.33029 | 1.0864 | 0.11952 | 0.015806 | 0.067071 | 0.83579 | -0.25878 | 0.58517 | 0.67305 | 1.0483 | 0.068068 |
| FFA183 | 0.98614 | 0.99071 | 0.96247 | -0.05518 | 0.23775 | 0.41164 | 0.71645 | -0.48106 | 0.2481 | 0.34292 | 1.2558 | 0.32863 |
| TAG569-FA226 | 0.006041 | 0.019316 | 0.40445 | -1.306 | 0.00927 | 0.047896 | 0.42533 | -1.2333 | 0.012089 | 0.050449 | 0.44996 | -1.1521 |
| SMd181/184 | 0.00045 | 0.004178 | 0.80894 | -0.30589 | 0.065905 | 0.15774 | 0.87924 | -0.18568 | 1.31E-05 | 0.005865 | 0.63841 | -0.64744 |
| PE180/183 | 0.48988 | 0.59946 | 1.2596 | 0.33298 | 0.059479 | 0.14667 | 0.62266 | -0.68349 | 0.001085 | 0.039244 | 0.40366 | -1.3088 |
| HCER240 | 0.21431 | 0.31852 | 1.2329 | 0.30211 | 0.7519 | 0.85725 | 1.0415 | 0.05866 | 0.35855 | 0.45948 | 0.89281 | -0.16357 |
| PEP-180/161 | 0.048809 | 0.096579 | 0.72822 | -0.45755 | 0.79715 | 0.88256 | 0.99845 | -0.00224 | 0.000256 | 0.016638 | 0.42132 | -1.247 |
| PEP-180/201 | 0.74467 | 0.81889 | 1.1122 | 0.15342 | 0.65802 | 0.79035 | 1.1374 | 0.18573 | 0.15287 | 0.23638 | 0.70823 | -0.49772 |
| PEP-160/226 | 0.00185 | 0.009337 | 1.5758 | 0.65611 | 0.035435 | 0.10517 | 1.3757 | 0.46021 | 0.01127 | 0.049553 | 1.4263 | 0.51233 |
| SMd181/181 | 0.46658 | 0.57966 | 1.1127 | 0.15408 | 0.90507 | 0.94613 | 1.0264 | 0.037586 | 0.11653 | 0.19253 | 1.261 | 0.33457 |
| LPC205 | 0.72905 | 0.80854 | 1.1239 | 0.16849 | 0.9307 | 0.95415 | 1.1768 | 0.23482 | 0.8961 | 0.92158 | 1.0679 | 0.094829 |
| PEP-160/204 | 0.49858 | 0.60533 | 1.1202 | 0.16382 | 0.33241 | 0.50841 | 1.1423 | 0.192 | 0.72495 | 0.79452 | 1.0311 | 0.044159 |
| PE170/202 | 0.072571 | 0.13233 | 0.82322 | -0.28065 | 0.082616 | 0.18419 | 0.79707 | -0.32723 | 0.004428 | 0.046601 | 0.67759 | -0.56151 |
| HCER260 | 0.048486 | 0.096579 | 0.77704 | -0.36394 | 0.25679 | 0.43308 | 0.86101 | -0.21589 | 0.029974 | 0.0773 | 0.77904 | -0.36023 |
| PE160/160 | 0.008262 | 0.024901 | 1.6259 | 0.70123 | 0.43932 | 0.60722 | 0.86083 | -0.2162 | 0.4781 | 0.5808 | 0.87003 | -0.20087 |
| TAG566-FA205 | 0.50465 | 0.60951 | 0.92355 | -0.11474 | 0.84795 | 0.90643 | 1.1909 | 0.25206 | 0.52363 | 0.61867 | 0.89082 | -0.16679 |
| TAG567-FA160 | 0.011852 | 0.032262 | 0.37457 | -1.4167 | 0.024447 | 0.084207 | 0.46007 | -1.1201 | 0.024342 | 0.068561 | 0.4801 | -1.0586 |
| PEP-180/225 | 0.008739 | 0.025912 | 1.3484 | 0.43126 | 0.075024 | 0.17276 | 1.2247 | 0.29243 | 0.75138 | 0.81525 | 1.0573 | 0.080323 |
| PE180/204 | 0.068708 | 0.12635 | 1.2995 | 0.37791 | 0.69749 | 0.81561 | 0.94228 | -0.08577 | 0.49965 | 0.59728 | 0.90832 | -0.13873 |
| PC160/181 | 0.060439 | 0.11504 | 1.2556 | 0.3284 | 0.016662 | 0.069514 | 1.3057 | 0.3848 | 0.4025 | 0.50005 | 1.0887 | 0.12263 |
| LPE200 | 0.04861 | 0.096579 | 0.7537 | -0.40793 | 0.98973 | 0.99094 | 1.0378 | 0.053491 | 0.001729 | 0.046601 | 0.48627 | -1.0402 |
| PC180/226 | 0.071935 | 0.13154 | 1.1994 | 0.26233 | 0.1731 | 0.32854 | 1.1658 | 0.22129 | 0.28351 | 0.37743 | 1.0996 | 0.13696 |
| SMd181/180 | 0.46041 | 0.5753 | 1.0521 | 0.073262 | 0.23077 | 0.40559 | 0.89595 | -0.15852 | 0.70649 | 0.77717 | 1.0284 | 0.040342 |
| PS160/180_2 | 0.026217 | 0.05825 | 1.5057 | 0.59045 | 0.19043 | 0.34921 | 0.79706 | -0.32724 | 0.45647 | 0.55963 | 1.0652 | 0.091122 |
| HCER181 | 0.49932 | 0.60533 | 0.90767 | -0.13977 | 0.069942 | 0.16679 | 0.7752 | -0.36737 | 0.084889 | 0.15017 | 0.78559 | -0.34815 |
| TAG566-FA225 | 0.011894 | 0.032262 | 0.45948 | -1.1219 | 0.086976 | 0.19259 | 0.58985 | -0.76157 | 0.073436 | 0.13777 | 0.57445 | -0.79976 |
| PC150/225 | 0.87612 | 0.90665 | 0.99667 | -0.00481 | 0.61123 | 0.75649 | 0.80293 | -0.31665 | 0.87822 | 0.91329 | 0.88372 | -0.17834 |
| TAG546-FA160 | 0.1357 | 0.21921 | 0.65425 | -0.61209 | 0.072882 | 0.17129 | 0.58145 | -0.78227 | 0.17139 | 0.25855 | 0.69255 | -0.53001 |
| TAG567-FA204 | 0.53655 | 0.64091 | 0.8212 | -0.28419 | 0.59553 | 0.74413 | 0.83141 | -0.26636 | 0.78352 | 0.84449 | 0.91853 | -0.12261 |
| LPC182 | 0.7494 | 0.82132 | 0.97603 | -0.035 | 0.042492 | 0.11582 | 1.1936 | 0.25538 | 0.278 | 0.37238 | 0.89299 | -0.16329 |
| PC181/224 | 0.84733 | 0.89081 | 0.95268 | -0.06994 | 0.65477 | 0.78937 | 0.86213 | -0.21403 | 0.5988 | 0.6842 | 0.88441 | -0.17721 |
| PEO-180/226 | 0.002337 | 0.01041 | 1.4722 | 0.55802 | 0.20028 | 0.36318 | 1.2149 | 0.28084 | 0.082589 | 0.14852 | 1.2873 | 0.36436 |
| SMd181/221 | 0.77936 | 0.84701 | 1.0432 | 0.060958 | 0.83422 | 0.90232 | 0.98233 | -0.02572 | 0.10761 | 0.18056 | 1.1945 | 0.25643 |

| | | | | | | | | | | | | |
|---|---|---|---|---|---|---|---|---|---|---|---|---|
| SM240 | 0.6449 | 0.73673 | 1.0432 | 0.061024 | 0.008837 | 0.046811 | 0.82277 | -0.28144 | 0.18603 | 0.27748 | 0.90841 | -0.13858 |
| DAG161/182 | 0.062726 | 0.11796 | 0.49687 | -1.009 | 0.51343 | 0.67252 | 0.79744 | -0.32655 | 0.4782 | 0.5808 | 0.82679 | -0.2744 |
| PEO-180/181 | 0.13455 | 0.21789 | 0.82573 | -0.27626 | 0.28201 | 0.46127 | 0.87801 | -0.18769 | 3.45E-05 | 0.005865 | 0.43427 | -1.2033 |
| PC170/224 | 0.60761 | 0.70383 | 1.0761 | 0.10579 | 0.47133 | 0.63792 | 0.82607 | -0.27567 | 0.89458 | 0.92148 | 0.94696 | -0.07863 |
| TAG520-FA180 | 0.019439 | 0.046222 | 0.4099 | -1.2866 | 0.003995 | 0.027667 | 0.35168 | -1.5077 | 0.008495 | 0.04704 | 0.34363 | -1.5411 |
| TAG565-FA204 | 0.008794 | 0.025912 | 0.55863 | -0.84004 | 0.21854 | 0.38977 | 0.78769 | -0.3443 | 0.19914 | 0.29331 | 0.77811 | -0.36195 |
| PE160/202 | 0.054243 | 0.10478 | 1.3322 | 0.41383 | 0.00181 | 0.019633 | 0.5891 | -0.76342 | 0.036386 | 0.084078 | 0.65071 | -0.6199 |
| PC140/204 | 0.77262 | 0.84392 | 0.96018 | -0.05862 | 0.98166 | 0.98799 | 0.98904 | -0.01589 | 0.59718 | 0.6842 | 0.94458 | -0.08225 |
| PEP-181/181 | 0.79223 | 0.85272 | 0.97882 | -0.03088 | 0.50232 | 0.66465 | 1.2208 | 0.28779 | 0.058618 | 0.11599 | 0.66565 | -0.58717 |
| TAG568-FA160 | 0.011148 | 0.030881 | 0.47199 | -1.0832 | 0.030769 | 0.095385 | 0.4992 | -1.0023 | 0.044116 | 0.097302 | 0.57663 | -0.79429 |
| FFA120 | 0.86175 | 0.89616 | 1.2156 | 0.28163 | 0.46565 | 0.63418 | 1.1691 | 0.22535 | 0.90823 | 0.92867 | 1.5074 | 0.59209 |
| DAG140/160 | 0.90394 | 0.92672 | 0.94425 | -0.08276 | 0.3641 | 0.53627 | 0.78086 | -0.35687 | 0.37229 | 0.47336 | 0.901 | -0.1504 |
| TAG567-FA181 | 0.024034 | 0.054326 | 0.60509 | -0.72479 | 0.17998 | 0.33864 | 0.74757 | -0.41971 | 0.11026 | 0.18358 | 0.6423 | -0.63868 |
| CER160 | 0.099149 | 0.17258 | 1.3347 | 0.41651 | 0.48738 | 0.65312 | 1.1005 | 0.13815 | 0.89318 | 0.92148 | 1.0382 | 0.054067 |
| TAG482-FA180 | 0.083215 | 0.14924 | 0.58468 | -0.77429 | 0.89167 | 0.9408 | 1.2314 | 0.30029 | 0.32394 | 0.42043 | 1.3582 | 0.4417 |
| LPC221 | 0.021135 | 0.04879 | 0.6583 | -0.60319 | 0.037163 | 0.10947 | 0.76467 | -0.3871 | 0.020099 | 0.062531 | 0.68692 | -0.54178 |
| PEP-181/161 | 0.247 | 0.35441 | 1.2792 | 0.35529 | 0.30758 | 0.48482 | 1.2449 | 0.31598 | 0.28324 | 0.37743 | 0.83829 | -0.25447 |
| PC160/180 | 0.021901 | 0.050027 | 1.264 | 0.338 | 0.040082 | 0.11364 | 1.1812 | 0.2402 | 0.26666 | 0.3609 | 1.0935 | 0.12894 |
| PC170/225 | 0.87753 | 0.90665 | 0.99304 | -0.01007 | 0.16668 | 0.32103 | 0.81762 | -0.2905 | 0.025277 | 0.070244 | 0.7444 | -0.42585 |
| FFA225 | 0.045835 | 0.09238 | 1.3592 | 0.44274 | 0.37656 | 0.54785 | 0.84773 | -0.23832 | 0.17815 | 0.26723 | 1.4554 | 0.54147 |
| FFA202 | 0.28611 | 0.40055 | 1.2172 | 0.28361 | 0.37799 | 0.54805 | 0.84994 | -0.23457 | 0.36137 | 0.46219 | 1.2563 | 0.32923 |
| PEP-181/160 | 0.066278 | 0.12223 | 1.5679 | 0.64883 | 0.34565 | 0.51848 | 1.2905 | 0.36793 | 0.57134 | 0.66064 | 1.1101 | 0.15071 |
| PEP-160/224 | 0.3352 | 0.44992 | 1.304 | 0.38292 | 0.66733 | 0.79713 | 1.1058 | 0.14515 | 0.79248 | 0.85133 | 1.0289 | 0.041113 |
| PEO-160/226 | 0.005846 | 0.019123 | 1.4236 | 0.50958 | 0.099459 | 0.21229 | 1.226 | 0.29393 | 0.14767 | 0.23147 | 1.2122 | 0.27765 |
| PC160/161 | 0.54848 | 0.65158 | 1.1292 | 0.17528 | 0.6693 | 0.79801 | 1.0978 | 0.13461 | 0.87032 | 0.9124 | 0.95568 | -0.0654 |
| SMd181/141 | 0.11978 | 0.20149 | 0.75778 | -0.40015 | 0.28162 | 0.46127 | 0.82295 | -0.28113 | 0.37798 | 0.47687 | 0.85481 | -0.22632 |
| SMd181/170 | 0.97015 | 0.98331 | 0.98941 | -0.01536 | 0.14772 | 0.28879 | 0.86187 | -0.21446 | 0.71484 | 0.78475 | 1.034 | 0.048178 |
| PEO-180/204 | 0.84954 | 0.89081 | 1.0217 | 0.031015 | 0.74369 | 0.84938 | 1.0241 | 0.034321 | 0.027148 | 0.073032 | 0.73943 | -0.43551 |
| FFA140 | 0.81896 | 0.874 | 1.0566 | 0.07938 | 0.4073 | 0.57392 | 1.1208 | 0.16453 | 0.55128 | 0.64432 | 1.2933 | 0.37102 |
| SMd181/140 | 0.73949 | 0.81665 | 1.0341 | 0.048365 | 0.80733 | 0.88755 | 0.94648 | -0.07935 | 0.95686 | 0.96426 | 0.99139 | -0.01248 |
| CER180 | 0.23147 | 0.33862 | 1.3019 | 0.38063 | 0.80848 | 0.88755 | 1.0118 | 0.016958 | 0.81173 | 0.86914 | 0.99449 | -0.00797 |
| TAG522-FA202 | 0.015299 | 0.03854 | 0.44646 | -1.1634 | 0.012954 | 0.060671 | 0.41216 | -1.2787 | 0.079701 | 0.14493 | 0.56091 | -0.83417 |
| PE160/140 | 0.85161 | 0.89081 | 1.0595 | 0.083406 | 0.78574 | 0.8767 | 1.0289 | 0.041134 | 0.4119 | 0.50978 | 0.92831 | -0.10732 |

| | | | | | | | | | | | | |
|---|---|---|---|---|---|---|---|---|---|---|---|---|
| PEP-160/181 | 0.60381 | 0.70192 | 0.94142 | -0.08709 | 0.26428 | 0.43942 | 1.2793 | 0.35538 | 0.006856 | 0.046601 | 0.68275 | -0.55056 |
| TAG504-FA141 | 0.0656 | 0.12132 | 0.45924 | -1.1227 | 0.14551 | 0.28531 | 0.54669 | -0.8712 | 0.02553 | 0.070423 | 0.38437 | -1.3794 |
| FFA204 | 0.032407 | 0.07009 | 0.75469 | -0.40605 | 0.000363 | 0.012404 | 0.61099 | -0.71077 | 0.052074 | 0.10879 | 0.77408 | -0.36945 |
| CE182 | 0.28086 | 0.3949 | 1.1872 | 0.24754 | 0.000117 | 0.011362 | 0.51074 | -0.96935 | 0.14907 | 0.23217 | 0.80837 | -0.30691 |
| PEP-182/226 | 0.005538 | 0.018582 | 1.5443 | 0.62694 | 0.31361 | 0.48959 | 1.211 | 0.27616 | 0.30127 | 0.39783 | 1.1509 | 0.20278 |
| CER221 | 0.11696 | 0.19828 | 1.3596 | 0.44322 | 0.98192 | 0.98799 | 0.96613 | -0.04971 | 0.87378 | 0.9124 | 1.0055 | 0.007951 |
| TAG500-FA160 | 0.057738 | 0.11055 | 0.47641 | -1.0697 | 0.007646 | 0.042544 | 0.34348 | -1.5417 | 0.020826 | 0.062992 | 0.37859 | -1.4013 |
| TAG523-FA203 | 0.1194 | 0.20137 | 0.62256 | -0.68371 | 0.026404 | 0.088602 | 0.44379 | -1.172 | 0.24961 | 0.34301 | 0.7121 | -0.48985 |
| PA180/203 | 0.75294 | 0.82381 | 0.90625 | -0.14202 | 0.71394 | 0.82996 | 1.0431 | 0.060828 | 0.49205 | 0.5921 | 0.85584 | -0.22458 |
| CER260 | 0.28335 | 0.39755 | 1.3032 | 0.38206 | 0.23036 | 0.40559 | 1.2577 | 0.33077 | 0.056954 | 0.11408 | 0.74569 | -0.42336 |
| TAG526-FA226 | 0.014137 | 0.03652 | 0.41061 | -1.2841 | 0.03151 | 0.096661 | 0.40865 | -1.2911 | 0.036421 | 0.084078 | 0.46118 | -1.1166 |
| TAG566-FA180 | 0.022505 | 0.051226 | 0.58669 | -0.76933 | 0.017103 | 0.070468 | 0.57384 | -0.80128 | 0.075665 | 0.13954 | 0.66522 | -0.58809 |
| LPE203 | 0.18315 | 0.28145 | 1.162 | 0.21656 | 0.77488 | 0.87124 | 0.96147 | -0.05669 | 0.06134 | 0.12064 | 0.73845 | -0.43743 |
| PE160/182 | 0.5415 | 0.64483 | 1.1582 | 0.21191 | 0.51678 | 0.67555 | 0.88016 | -0.18417 | 0.016137 | 0.057092 | 0.62138 | -0.68646 |
| TAG500-FA180 | 0.040406 | 0.083507 | 0.42245 | -1.2431 | 0.003003 | 0.024037 | 0.25947 | -1.9464 | 0.010768 | 0.048666 | 0.3142 | -1.6703 |
| PE160/205 | 0.55308 | 0.65584 | 1.4176 | 0.50343 | 0.89997 | 0.94613 | 0.96796 | -0.04698 | 0.72905 | 0.79766 | 0.86079 | -0.21626 |
| PEP-181/224 | 0.322 | 0.43513 | 1.3751 | 0.45958 | 0.86854 | 0.91789 | 1.0737 | 0.10254 | 0.92287 | 0.93581 | 1.0312 | 0.044272 |
| SM261 | 0.48283 | 0.5953 | 0.94887 | -0.07571 | 0.012867 | 0.060671 | 0.81062 | -0.3029 | 0.64928 | 0.72876 | 0.96833 | -0.04644 |
| PC160/140 | 0.55639 | 0.65856 | 1.1998 | 0.26283 | 0.80186 | 0.88477 | 1.0308 | 0.043753 | 0.26273 | 0.35633 | 0.76297 | -0.3903 |
| PC170/205 | 0.73243 | 0.8109 | 1.1477 | 0.1987 | 0.86113 | 0.91251 | 1.1244 | 0.16912 | 0.63137 | 0.71111 | 0.81886 | -0.28831 |
| HCER221 | 0.48001 | 0.59296 | 0.93943 | -0.09014 | 0.015386 | 0.067071 | 0.7984 | -0.32481 | 0.007731 | 0.046601 | 0.82278 | -0.28142 |
| SMd181/201 | 0.67167 | 0.75913 | 1.0468 | 0.066018 | 0.95953 | 0.97589 | 1.0076 | 0.010887 | 0.044494 | 0.097302 | 1.3107 | 0.39031 |
| LPE240 | 0.88257 | 0.9091 | 0.99172 | -0.01199 | 0.22516 | 0.3994 | 1.1314 | 0.17809 | 0.024433 | 0.068561 | 0.71803 | -0.47788 |
| HCER261 | 0.12406 | 0.20609 | 0.81709 | -0.29144 | 0.018957 | 0.074827 | 0.72091 | -0.47212 | 0.000564 | 0.028245 | 0.60691 | -0.72045 |
| PC180/225 | 0.43836 | 0.55921 | 1.1141 | 0.15589 | 0.37291 | 0.54431 | 1.1149 | 0.1569 | 0.87639 | 0.91284 | 1.0147 | 0.021016 |
| SMd181/223 | 0.88422 | 0.90937 | 1.0072 | 0.010347 | 0.078209 | 0.1773 | 0.82904 | -0.27049 | 0.8814 | 0.91513 | 0.98475 | -0.02217 |
| SMd181/225 | 0.12839 | 0.21001 | 0.85732 | -0.2221 | 0.43456 | 0.60192 | 0.92536 | -0.11191 | 0.20686 | 0.30261 | 0.88282 | -0.1798 |
| PE160/204 | 0.1233 | 0.20609 | 1.3574 | 0.44083 | 0.85808 | 0.91127 | 0.96824 | -0.04656 | 0.45906 | 0.56069 | 0.87692 | -0.18948 |
| TAG545-FA205 | 0.82588 | 0.87851 | 1.0718 | 0.10001 | 0.18875 | 0.3481 | 0.67858 | -0.55942 | 0.57606 | 0.66492 | 0.85094 | -0.23287 |
| PEO-160/204 | 0.54181 | 0.64483 | 1.0791 | 0.10979 | 0.33548 | 0.51076 | 1.1511 | 0.20301 | 0.68039 | 0.75975 | 0.90245 | -0.14809 |
| PEO-160/183 | 0.50259 | 0.60815 | 0.89486 | -0.16026 | 0.23638 | 0.41035 | 0.82076 | -0.28496 | 0.017172 | 0.057326 | 0.62843 | -0.67018 |
| DAG160/181 | 0.15226 | 0.24118 | 0.69662 | -0.52156 | 0.27856 | 0.45793 | 0.75089 | -0.41332 | 0.012049 | 0.050449 | 0.43429 | -1.2033 |
| TAG566-FA226 | 0.12781 | 0.21001 | 0.71445 | -0.48509 | 0.091512 | 0.20059 | 0.66811 | -0.58184 | 0.074049 | 0.13852 | 0.65414 | -0.61233 |

| | | | | | | | | | | | | |
|---|---|---|---|---|---|---|---|---|---|---|---|---|
| PC160/183 | 0.30689 | 0.42328 | 1.1898 | 0.25068 | 0.49667 | 0.65986 | 0.90957 | -0.13674 | 0.013746 | 0.053909 | 0.6629 | -0.59313 |
| PC180/182 | 0.44488 | 0.56455 | 1.0621 | 0.08695 | 0.67579 | 0.8028 | 1.0211 | 0.030156 | 0.60536 | 0.69018 | 0.95439 | -0.06735 |
| FFA161 | 0.80129 | 0.85798 | 1.2231 | 0.2906 | 0.40022 | 0.56848 | 1.2142 | 0.28 | 0.23269 | 0.32647 | 1.7032 | 0.76821 |
| FFA241 | 0.12437 | 0.20609 | 1.1881 | 0.24864 | 0.36025 | 0.5318 | 1.0754 | 0.1049 | 0.38874 | 0.48762 | 0.9199 | -0.12044 |
| TAG483-FA141 | 0.049051 | 0.096765 | 0.43169 | -1.2119 | 0.02014 | 0.076891 | 0.3813 | -1.391 | 0.009427 | 0.047969 | 0.31087 | -1.6856 |
| PEP-181/226 | 0.039552 | 0.082702 | 1.4712 | 0.55703 | 0.2541 | 0.42966 | 1.2909 | 0.36833 | 0.19742 | 0.29143 | 1.2298 | 0.29837 |
| PEP-160/180 | 0.9979 | 0.9979 | 1.019 | 0.027113 | 0.13772 | 0.27334 | 1.3025 | 0.38126 | 0.13391 | 0.21263 | 0.84677 | -0.23995 |
| FFA201 | 0.28889 | 0.40357 | 1.3426 | 0.42508 | 0.12191 | 0.24496 | 1.1967 | 0.25901 | 0.1569 | 0.24166 | 1.5229 | 0.60682 |
| FFA141 | 0.40019 | 0.51897 | 1.4602 | 0.54615 | 0.2646 | 0.43942 | 1.3655 | 0.44939 | 0.26056 | 0.35412 | 2.0057 | 1.0041 |
| PC200/226 | 0.14656 | 0.23442 | 1.2987 | 0.37709 | 0.71858 | 0.83238 | 0.9829 | -0.02488 | 0.29472 | 0.39075 | 0.89294 | -0.16336 |
| PEP-140/181 | 0.31474 | 0.42966 | 1.2549 | 0.32753 | 0.11143 | 0.23137 | 1.3094 | 0.38888 | 0.16682 | 0.25493 | 0.82403 | -0.27923 |
| LPE170 | 0.60684 | 0.70383 | 1.0483 | 0.068033 | 0.078709 | 0.1773 | 0.72063 | -0.47267 | 0.002175 | 0.046601 | 0.48599 | -1.041 |
| FFA240 | 0.93856 | 0.95919 | 0.98702 | -0.01884 | 0.89763 | 0.94556 | 1.0232 | 0.033126 | 0.75444 | 0.8172 | 0.94481 | -0.0819 |
| Cholesterol | 0.35247 | 0.46543 | 1.1643 | 0.21942 | 0.000214 | 0.011362 | 0.52431 | -0.9315 | 0.61748 | 0.69945 | 0.93408 | -0.09839 |
| FFA160 | 0.095809 | 0.16812 | 1.191 | 0.25217 | 0.32503 | 0.5038 | 1.0713 | 0.099376 | 0.2324 | 0.32647 | 1.0994 | 0.13678 |
| PEP-181/204 | 0.50633 | 0.61041 | 1.1547 | 0.20751 | 0.68714 | 0.81184 | 1.0642 | 0.089765 | 0.91155 | 0.92867 | 0.95459 | -0.06705 |
| PE180/224 | 0.20042 | 0.30132 | 1.3081 | 0.38743 | 0.65427 | 0.78937 | 0.84949 | -0.23534 | 0.25883 | 0.35251 | 0.71131 | -0.49144 |
| CE183 | 0.3656 | 0.4782 | 1.1953 | 0.25738 | 0.0112 | 0.055235 | 0.63777 | -0.64889 | 0.32571 | 0.42155 | 0.84209 | -0.24796 |
| PEP-182/204 | 0.18331 | 0.28145 | 1.235 | 0.30453 | 0.38275 | 0.5532 | 1.122 | 0.16603 | 0.61465 | 0.69832 | 1.0465 | 0.065608 |
| CER240 | 0.000999 | 0.006732 | 1.4898 | 0.57508 | 0.032268 | 0.098161 | 1.3592 | 0.44279 | 0.43547 | 0.5359 | 1.1203 | 0.16394 |
| PEP-160/183 | 0.12023 | 0.20173 | 0.82365 | -0.2799 | 0.69825 | 0.81561 | 0.98224 | -0.02585 | 0.002004 | 0.046601 | 0.61695 | -0.69678 |
| PA181/182 | 0.18603 | 0.28496 | 0.84832 | -0.23732 | 0.000528 | 0.012404 | 1.6138 | 0.69044 | 0.87456 | 0.9124 | 1.0186 | 0.026657 |
| FFA181 | 0.037969 | 0.079994 | 1.2249 | 0.29264 | 0.088126 | 0.19447 | 1.1603 | 0.21452 | 0.81911 | 0.87273 | 1.0122 | 0.017427 |
| PE180/182 | 0.40151 | 0.51965 | 1.1342 | 0.18164 | 0.66331 | 0.79377 | 0.94751 | -0.07779 | 0.008881 | 0.047781 | 0.73519 | -0.44381 |
| PC160/201 | 0.6861 | 0.77213 | 1.1009 | 0.13868 | 0.97742 | 0.98727 | 0.99416 | -0.00845 | 0.79895 | 0.85686 | 0.97661 | -0.03414 |
| PEP-180/224 | 0.51608 | 0.61873 | 1.1774 | 0.23565 | 0.84619 | 0.90643 | 1.0294 | 0.041863 | 0.8647 | 0.91088 | 0.92306 | -0.11551 |
| PS160/161_2 | 0.2497 | 0.35647 | 1.1486 | 0.19989 | 0.38324 | 0.5532 | 0.89485 | -0.16029 | 0.3056 | 0.40191 | 1.1854 | 0.24542 |
| PE180/181 | 0.078422 | 0.14221 | 1.2462 | 0.31757 | 0.053454 | 0.13593 | 1.3073 | 0.38658 | 0.016641 | 0.057235 | 0.72578 | -0.4624 |
| FFA170 | 0.979 | 0.98657 | 0.99776 | -0.00323 | 0.57258 | 0.7266 | 1.0572 | 0.08027 | 0.65181 | 0.73034 | 1.0598 | 0.083852 |
| FFA182 | 0.009381 | 0.027134 | 1.2142 | 0.28006 | 0.039064 | 0.11203 | 1.1421 | 0.19172 | 0.55894 | 0.65093 | 0.95856 | -0.06107 |
| PC150/161 | 0.40365 | 0.52138 | 1.1501 | 0.20174 | 0.99094 | 0.99094 | 0.99516 | -0.00701 | 0.56674 | 0.65649 | 1.0833 | 0.11547 |
| PC180/205 | 0.74044 | 0.81665 | 1.1308 | 0.17735 | 0.82474 | 0.89936 | 1.2092 | 0.27404 | 0.76619 | 0.82855 | 1.1229 | 0.16727 |
| FFA180 | 0.20975 | 0.31247 | 1.1257 | 0.17081 | 0.83424 | 0.90232 | 0.97504 | -0.03647 | 0.48121 | 0.58295 | 1.053 | 0.074526 |

| | | | | | | | | | | | | |
|---|---|---|---|---|---|---|---|---|---|---|---|---|
| TAG524-FA204 | 0.85039 | 0.89081 | 0.9169 | -0.12517 | 0.11546 | 0.23637 | 0.61241 | -0.70744 | 0.55784 | 0.65081 | 1.1657 | 0.22123 |
| LPE182 | 0.081839 | 0.14717 | 1.3491 | 0.43195 | 0.064099 | 0.1557 | 1.47 | 0.55582 | 0.013447 | 0.053428 | 0.69455 | -0.52585 |
| TAG480-FA160 | 0.13131 | 0.21317 | 0.59794 | -0.74192 | 0.013215 | 0.060936 | 0.37463 | -1.4164 | 0.025854 | 0.070718 | 0.43977 | -1.1852 |
| CER200 | 0.12923 | 0.21036 | 1.3708 | 0.45501 | 0.66128 | 0.7928 | 1.0491 | 0.069112 | 0.45746 | 0.55979 | 0.8506 | -0.23344 |
| CER220 | 0.093889 | 0.16564 | 1.2474 | 0.31896 | 0.60333 | 0.75242 | 1.0374 | 0.053011 | 0.3919 | 0.49063 | 0.82556 | -0.27656 |
| PEO-160/160 | 0.000298 | 0.0037 | 1.7005 | 0.76594 | 0.38518 | 0.55353 | 1.1022 | 0.14045 | 0.84597 | 0.89549 | 1.0006 | 0.000868 |
| TAG564-FA204 | 0.008797 | 0.025912 | 0.65666 | -0.60678 | 0.005083 | 0.032441 | 0.63811 | -0.64812 | 0.11892 | 0.1955 | 0.7762 | -0.36549 |
| PS160/181 | 0.79007 | 0.85272 | 0.96584 | -0.05015 | 0.11231 | 0.23137 | 0.79097 | -0.33831 | 0.37711 | 0.4767 | 1.1537 | 0.20632 |
| LPE220 | 0.11364 | 0.19316 | 1.186 | 0.24608 | 0.015866 | 0.067071 | 1.6092 | 0.68638 | 0.094766 | 0.16495 | 0.83248 | -0.26451 |
| TAG525-FA205 | 0.97311 | 0.98331 | 0.98557 | -0.02096 | 0.28529 | 0.46315 | 0.60531 | -0.72424 | 0.91098 | 0.92867 | 0.90957 | -0.13675 |
| TAG544-FA224 | 0.79246 | 0.85272 | 0.88924 | -0.16936 | 0.18487 | 0.34386 | 0.66107 | -0.59712 | 0.68575 | 0.76312 | 0.92241 | -0.11652 |
| PEO-180/161 | 0.1973 | 0.29884 | 0.87545 | -0.19191 | 0.90746 | 0.94613 | 1.0038 | 0.005466 | 0.002741 | 0.046601 | 0.63722 | -0.65013 |
| FFA221 | 0.84048 | 0.89081 | 0.91206 | -0.13279 | 0.3937 | 0.56206 | 1.3057 | 0.38486 | 0.29831 | 0.39471 | 0.75475 | -0.40592 |
| PE170/224 | 0.83666 | 0.88852 | 1.0223 | 0.031752 | 0.028161 | 0.090598 | 0.66885 | -0.58025 | 0.15272 | 0.23638 | 0.77115 | -0.37491 |
| PC180/180 | 0.36918 | 0.48163 | 1.1165 | 0.15904 | 0.98872 | 0.99094 | 1.0033 | 0.0048 | 0.50095 | 0.59729 | 1.06 | 0.084089 |
| CE181 | 0.74785 | 0.821 | 1.1209 | 0.16464 | 0.031512 | 0.096661 | 0.72541 | -0.46313 | 0.58065 | 0.66903 | 0.91363 | -0.13031 |
| PA220/226 | 0.15286 | 0.24154 | 1.2087 | 0.27349 | 0.048156 | 0.12572 | 1.3982 | 0.48355 | 0.49933 | 0.59728 | 1.0938 | 0.1294 |
| PEO-160/181 | 0.7781 | 0.84701 | 1.0388 | 0.054878 | 0.82614 | 0.89936 | 1.0704 | 0.098114 | 0.002454 | 0.046601 | 0.65224 | -0.61653 |
| TAG482-FA141 | 0.048699 | 0.096579 | 0.49041 | -1.0279 | 0.052137 | 0.13363 | 0.54454 | -0.87689 | 0.00407 | 0.046601 | 0.30394 | -1.7181 |
| FFA150 | 0.035965 | 0.076813 | 0.78766 | -0.34435 | 0.54244 | 0.70087 | 1.1262 | 0.17143 | 0.003119 | 0.046601 | 0.68888 | -0.53768 |
| LPC160 | 0.2349 | 0.34134 | 1.101 | 0.1388 | 0.072132 | 0.17014 | 1.1608 | 0.21518 | 0.81761 | 0.87257 | 1.0188 | 0.02684 |
| TAG568-FA226 | 0.069497 | 0.12744 | 0.58606 | -0.77089 | 0.17144 | 0.32645 | 0.67854 | -0.5595 | 0.31289 | 0.41008 | 0.78767 | -0.34434 |
| LPE201 | 0.32215 | 0.43513 | 1.1876 | 0.24801 | 0.25922 | 0.43498 | 1.2167 | 0.28298 | 0.014653 | 0.055314 | 0.62657 | -0.67446 |
| PE160/224 | 0.19299 | 0.29354 | 1.3895 | 0.47456 | 0.39252 | 0.56206 | 0.66813 | -0.58179 | 0.27959 | 0.37374 | 0.60615 | -0.72225 |
| PC160/203 | 0.23726 | 0.3436 | 1.2213 | 0.28844 | 0.77732 | 0.87248 | 1.0232 | 0.033076 | 0.24975 | 0.34301 | 1.1734 | 0.23071 |
| FFA222 | 0.31482 | 0.42966 | 1.1575 | 0.21107 | 0.59421 | 0.74405 | 1.0332 | 0.047132 | 0.59907 | 0.6842 | 1.115 | 0.15708 |
| HCER201 | 0.35869 | 0.47173 | 0.91547 | -0.12742 | 0.002275 | 0.021783 | 0.71716 | -0.47963 | 0.023679 | 0.068209 | 0.80703 | -0.30931 |
| TAG544-FA204 | 0.67508 | 0.76166 | 0.94796 | -0.07711 | 0.038898 | 0.11203 | 0.71361 | -0.4868 | 0.87039 | 0.9124 | 1.0177 | 0.025292 |
| TAG567-FA226 | 0.041552 | 0.085602 | 0.55826 | -0.84099 | 0.24711 | 0.42113 | 0.77544 | -0.36691 | 0.21182 | 0.30556 | 0.7751 | -0.36755 |
| PC200/225 | 0.49159 | 0.6003 | 1.1022 | 0.14043 | 0.83064 | 0.90232 | 0.9548 | -0.06673 | 0.12537 | 0.20352 | 0.76736 | -0.38202 |
| TAG503-FA141 | 0.0201 | 0.04726 | 0.47721 | -1.0673 | 0.52255 | 0.68173 | 0.86125 | -0.21549 | 0.012418 | 0.051166 | 0.38356 | -1.3825 |
| TAG565-FA225 | 0.27844 | 0.39234 | 0.68386 | -0.54823 | 0.05748 | 0.14228 | 0.52916 | -0.91823 | 0.057451 | 0.11473 | 0.52462 | -0.93067 |
| PC160/120 | 0.49791 | 0.60533 | 1.4093 | 0.495 | 0.91574 | 0.95079 | 1.128 | 0.17372 | 0.88896 | 0.9186 | 1.0275 | 0.039175 |

| | | | | | | | | | | | | |
|---|---|---|---|---|---|---|---|---|---|---|---|---|
| FFA200 | 0.16292 | 0.25314 | 1.2449 | 0.31603 | 0.16909 | 0.32376 | 2.0349 | 1.025 | 0.96083 | 0.96528 | 1.0007 | 0.000958 |
| PC180/224 | 0.63351 | 0.7274 | 1.0696 | 0.097029 | 0.92886 | 0.95415 | 0.98402 | -0.02324 | 0.70503 | 0.77717 | 0.92083 | -0.119 |
| PC200/141 | 0.86898 | 0.90081 | 1.0121 | 0.017295 | 0.19393 | 0.35463 | 1.1307 | 0.17726 | 0.36602 | 0.4663 | 0.90341 | -0.14654 |
| PS161/180_2 | 0.012122 | 0.032744 | 1.413 | 0.49875 | 0.050227 | 0.12975 | 1.2541 | 0.3267 | 0.006301 | 0.046601 | 1.4756 | 0.56126 |
| LPE161 | 0.84266 | 0.89081 | 0.94211 | -0.08604 | 0.3031 | 0.48233 | 0.82001 | -0.28628 | 0.12112 | 0.19762 | 0.70563 | -0.50302 |
| PC200/204 | 0.12807 | 0.21001 | 1.32 | 0.40055 | 0.35354 | 0.52666 | 0.86049 | -0.21677 | 0.37388 | 0.47446 | 0.86021 | -0.21724 |
| SMd181/183 | 0.69345 | 0.77433 | 0.97291 | -0.03962 | 0.46419 | 0.63351 | 0.94242 | -0.08556 | 0.31851 | 0.4147 | 0.90581 | -0.14272 |
| PC180/200 | 0.49241 | 0.6003 | 1.0961 | 0.13232 | 0.23616 | 0.41035 | 1.1774 | 0.23564 | 0.95451 | 0.96339 | 1.027 | 0.038437 |
| PE160/181 | 0.27483 | 0.3881 | 1.2913 | 0.36879 | 0.69815 | 0.81561 | 0.93561 | -0.09602 | 0.004764 | 0.046601 | 0.52696 | -0.92424 |
| PEP-160/201 | 0.45951 | 0.57527 | 1.268 | 0.34256 | 0.46709 | 0.63482 | 1.2283 | 0.29664 | 0.93399 | 0.94414 | 0.97456 | -0.03718 |
| PC180/120 | 0.48662 | 0.59772 | 1.3437 | 0.42624 | 0.90517 | 0.94613 | 1.0478 | 0.067313 | 0.85858 | 0.90589 | 0.91201 | -0.13288 |
| PEP-180/226 | 0.018038 | 0.043486 | 1.3117 | 0.39139 | 0.16185 | 0.31266 | 1.2462 | 0.31751 | 0.2779 | 0.37238 | 1.1274 | 0.17297 |
| DAG200/200 | 0.096067 | 0.16812 | 1.6318 | 0.70651 | 0.85606 | 0.91061 | 0.9565 | -0.06416 | 0.60866 | 0.69272 | 0.98626 | -0.01996 |
| PEP-160/161 | 0.47559 | 0.58861 | 1.1395 | 0.1884 | 0.25925 | 0.43498 | 1.1899 | 0.2508 | 0.5195 | 0.61602 | 0.92444 | -0.11335 |
| CER201 | 0.265 | 0.37667 | 1.3935 | 0.47871 | 0.85474 | 0.91061 | 0.89192 | -0.16502 | 0.87385 | 0.9124 | 0.91601 | -0.12657 |
| PC170/204 | 0.31056 | 0.42562 | 1.1195 | 0.1628 | 0.11207 | 0.23137 | 0.84875 | -0.23658 | 0.42894 | 0.52986 | 0.91924 | -0.12148 |
| PE180/205 | 0.80131 | 0.85798 | 1.1651 | 0.22051 | 0.69411 | 0.81561 | 0.91074 | -0.13488 | 0.4704 | 0.57346 | 0.76894 | -0.37905 |
| FFA224 | 0.16271 | 0.25314 | 1.3761 | 0.46059 | 0.90834 | 0.94613 | 0.96393 | -0.053 | 0.2544 | 0.34793 | 1.4314 | 0.51743 |
| PC140/224 | 0.64614 | 0.73673 | 0.90162 | -0.14941 | 0.15013 | 0.29263 | 0.69341 | -0.52823 | 0.12055 | 0.19718 | 0.66298 | -0.59297 |

## 7.5 Gy

| | SD 1 vs. Pre | | | | SD 2 vs. Pre | | | | SD 6 vs. Pre | | | |
|---|---|---|---|---|---|---|---|---|---|---|---|---|
| Lipid Name | p-value | FDR | FC | LOG2FC | p-value | FDR | FC | LOG2FC | p-value | FDR | FC | LOG2FC |
| PI160/201 | 0.010648 | 0.050968 | 0.44724 ↓ | -1.161 | 0.45902 | 0.63118 | 0.79342 ↓ | -0.334 | 0.007514 | 0.024609 | 0.38969 ↓ | -1.36 |
| LPE160 | 0.021999 | 0.069893 | 1.4562 ↑ | 0.5423 | 0.41861 | 0.58985 | 0.85597 ↓ | -0.224 | 0.21408 | 0.28854 | 0.79183 ↓ | -0.337 |
| PC182/203 | 0.000135 | 0.009734 | 0.346 ↓ | -1.531 | 0.020414 | 0.11428 | 0.61399 ↓ | -0.704 | 0.001212 | 0.012765 | 0.36658 ↓ | -1.448 |
| PG180/182_2 | 0.030522 | 0.082447 | 0.6445 ↓ | -0.634 | 0.011853 | 0.11428 | 0.52071 ↓ | -0.941 | 0.011282 | 0.031369 | 0.50332 ↓ | -0.99 |
| PI160/204 | 0.025315 | 0.074571 | 0.62172 ↓ | -0.686 | 0.84631 | 0.90526 | 0.97783 ↓ | -0.032 | 0.10536 | 0.16101 | 0.74748 ↓ | -0.42 |
| PG180/182 | 0.033156 | 0.086518 | 0.64305 ↓ | -0.637 | 0.011553 | 0.11428 | 0.50948 ↓ | -0.973 | 0.01351 | 0.035002 | 0.51087 ↓ | -0.969 |
| LPC202 | 0.016268 | 0.06088 | 0.61009 ↓ | -0.713 | 0.033052 | 0.12787 | 0.63492 ↓ | -0.655 | 0.00307 | 0.01797 | 0.47263 ↓ | -1.081 |
| TAG547-FA183 | 0.000751 | 0.018074 | 0.086378 ↓ | -3.533 | 0.004496 | 0.11015 | 0.11402 ↓ | -3.133 | 0.000272 | 0.008616 | 0.065829 ↓ | -3.925 |
| PI160/182 | 0.010069 | 0.049771 | 0.49607 ↓ | -1.011 | 0.61465 | 0.73826 | 0.86989 ↓ | -0.201 | 0.017354 | 0.041541 | 0.51106 ↓ | -0.968 |
| TAG547-FA182 | 0.008388 | 0.044966 | 0.12757 ↓ | -2.971 | 0.009644 | 0.11015 | 0.12079 ↓ | -3.049 | 0.003529 | 0.018829 | 0.10236 ↓ | -3.288 |

| | | | | | | | | | | | | | | | |
|---|---|---|---|---|---|---|---|---|---|---|---|---|---|---|---|
| PS161/180 | 0.010237 | 0.049771 | 0.50313 | ↓ | -0.991 | 0.10668 | 0.21302 | 0.69807 | ↓ | -0.519 | 0.000899 | 0.012194 | 0.39899 | ↓ | -1.326 |
| TAG526-FA183 | 0.00051 | 0.018074 | 0.12401 | ↓ | -3.012 | 0.003541 | 0.11015 | 0.15457 | ↓ | -2.694 | 0.000149 | 0.007659 | 0.090672 | ↓ | -3.463 |
| TAG483-FA183 | 0.003648 | 0.030709 | 0.15572 | ↓ | -2.683 | 0.014398 | 0.11428 | 0.20478 | ↓ | -2.288 | 0.004857 | 0.019885 | 0.16204 | ↓ | -2.626 |
| LPE226 | 9.26E-05 | 0.008861 | 1.597 | ↑ | 0.6754 | 0.73071 | 0.82729 | 1.0724 | ↑ | 0.1008 | 0.25316 | 0.32625 | 1.2781 | ↑ | 0.3541 |
| PG160/161 | 0.17875 | 0.28313 | 0.6192 | ↓ | -0.692 | 0.19715 | 0.34043 | 0.65511 | ↓ | -0.61 | 0.043573 | 0.081279 | 0.45517 | ↓ | -1.136 |
| TAG525-FA182 | 0.000554 | 0.018074 | 0.20466 | ↓ | -2.289 | 0.006234 | 0.11015 | 0.26088 | ↓ | -1.939 | 0.000131 | 0.007659 | 0.16596 | ↓ | -2.591 |
| PE181/203 | 0.009146 | 0.046514 | 0.48995 | ↓ | -1.029 | 0.035879 | 0.12787 | 0.59445 | ↓ | -0.75 | 0.003878 | 0.018938 | 0.35017 | ↓ | -1.514 |
| PI160/180 | 0.012351 | 0.053913 | 0.50432 | ↓ | -0.988 | 0.82038 | 0.88727 | 0.94026 | ↓ | -0.089 | 0.015839 | 0.039244 | 0.50093 | ↓ | -0.997 |
| PC160/226 | 0.11655 | 0.21135 | 1.2244 | ↑ | 0.2921 | 0.81652 | 0.88727 | 1.0344 | ↑ | 0.0488 | 0.026587 | 0.055298 | 1.5881 | ↑ | 0.6673 |
| PI160/160 | 0.031723 | 0.084435 | 0.51779 | ↓ | -0.95 | 0.69071 | 0.79444 | 0.86995 | ↓ | -0.201 | 0.12401 | 0.18431 | 0.63633 | ↓ | -0.652 |
| PI181/203_2 | 0.000931 | 0.018192 | 0.47553 | ↓ | -1.072 | 0.33894 | 0.50262 | 0.86419 | ↓ | -0.211 | 0.01725 | 0.041541 | 0.53001 | ↓ | -0.916 |
| TAG542-FA200 | 0.0018 | 0.023374 | 0.18545 | ↓ | -2.431 | 0.005245 | 0.11015 | 0.22691 | ↓ | -2.14 | 0.000218 | 0.008255 | 0.10522 | ↓ | -3.249 |
| TAG546-FA183 | 0.001831 | 0.023374 | 0.11189 | ↓ | -3.16 | 0.00961 | 0.11015 | 0.15959 | ↓ | -2.648 | 0.000828 | 0.011465 | 0.08642 | ↓ | -3.533 |
| TAG504-FA182 | 0.000437 | 0.018074 | 0.17652 | ↓ | -2.502 | 0.005874 | 0.11015 | 0.26819 | ↓ | -1.899 | 0.000334 | 0.008616 | 0.15067 | ↓ | -2.731 |
| PC181/202 | 0.00157 | 0.023374 | 0.50109 | ↓ | -0.997 | 0.002942 | 0.11015 | 0.53683 | ↓ | -0.897 | 0.001212 | 0.012765 | 0.43712 | ↓ | -1.194 |
| PC182/183 | 4.08E-05 | 0.007869 | 0.32626 | ↓ | -1.616 | 4.32E-05 | 0.028101 | 0.41247 | ↓ | -1.278 | 0.000382 | 0.008651 | 0.24554 | ↓ | -2.026 |
| PI160/182_2 | 0.012303 | 0.053913 | 0.49134 | ↓ | -1.025 | 0.79562 | 0.87197 | 0.9075 | ↓ | -0.14 | 0.019605 | 0.044159 | 0.49706 | ↓ | -1.009 |
| PI181/203 | 0.001746 | 0.023374 | 0.45795 | ↓ | -1.127 | 0.28959 | 0.4551 | 0.86424 | ↓ | -0.211 | 0.017502 | 0.041736 | 0.54822 | ↓ | -0.867 |
| PG160/161_2 | 0.10776 | 0.19874 | 0.63546 | ↓ | -0.654 | 0.10203 | 0.20688 | 0.62906 | ↓ | -0.669 | 0.030038 | 0.060547 | 0.62351 | ↓ | -0.682 |
| LPC200 | 0.001109 | 0.019512 | 0.49213 | ↓ | -1.023 | 0.16019 | 0.29211 | 0.75981 | ↓ | -0.396 | 0.011958 | 0.031843 | 0.45931 | ↓ | -1.122 |
| TAG504-FA140 | 0.000711 | 0.018074 | 0.18366 | ↓ | -2.445 | 0.005296 | 0.11015 | 0.25649 | ↓ | -1.963 | 0.000143 | 0.007659 | 0.1243 | ↓ | -3.008 |
| TAG525-FA183 | 0.003003 | 0.029174 | 0.21027 | ↓ | -2.25 | 0.010935 | 0.11428 | 0.22857 | ↓ | -2.129 | 0.000501 | 0.009864 | 0.14361 | ↓ | -2.8 |
| TAG526-FA182 | 0.000281 | 0.018074 | 0.16951 | ↓ | -2.561 | 0.004499 | 0.11015 | 0.21369 | ↓ | -2.226 | 0.000102 | 0.007659 | 0.12581 | ↓ | -2.991 |
| TAG525-FA225 | 0.008147 | 0.044693 | 0.28288 | ↓ | -1.822 | 0.019488 | 0.11428 | 0.29585 | ↓ | -1.757 | 0.006381 | 0.022215 | 0.24713 | ↓ | -2.017 |
| TAG504-FA183 | 0.000478 | 0.018074 | 0.1261 | ↓ | -2.987 | 0.014347 | 0.11428 | 0.25459 | ↓ | -1.974 | 0.001012 | 0.012765 | 0.1272 | ↓ | -2.975 |
| TAG526-FA161 | 0.000106 | 0.008861 | 0.11438 | ↓ | -3.128 | 0.002992 | 0.11015 | 0.17379 | ↓ | -2.525 | 0.000329 | 0.008616 | 0.11783 | ↓ | -3.085 |
| CE204 | 0.19883 | 0.30819 | 1.5394 | ↑ | 0.6224 | 0.5976 | 0.73128 | 0.84416 | ↓ | -0.244 | 0.073584 | 0.11946 | 1.5894 | ↑ | 0.6685 |
| TAG546-FA181 | 0.001599 | 0.023374 | 0.12189 | ↓ | -3.036 | 0.008801 | 0.11015 | 0.16581 | ↓ | -2.592 | 0.001553 | 0.013948 | 0.11761 | ↓ | -3.088 |
| PI160/204_2 | 0.021616 | 0.069854 | 0.64021 | ↓ | -0.643 | 0.72658 | 0.82549 | 0.94968 | ↓ | -0.074 | 0.10442 | 0.15994 | 0.74952 | ↓ | -0.416 |
| TAG503-FA140 | 0.000888 | 0.018074 | 0.20881 | ↓ | -2.26 | 0.025991 | 0.12086 | 0.37409 | ↓ | -1.419 | 0.001408 | 0.013481 | 0.19057 | ↓ | -2.392 |
| PI161/181 | 0.018443 | 0.063783 | 0.44984 | ↓ | -1.153 | 0.97024 | 0.98818 | 0.95673 | ↓ | -0.064 | 0.007205 | 0.024429 | 0.43541 | ↓ | -1.2 |
| LPC220 | 0.000734 | 0.018074 | 0.487 | ↓ | -1.038 | 0.29242 | 0.45542 | 0.82126 | ↓ | -0.284 | 0.017892 | 0.042501 | 0.52664 | ↓ | -0.925 |
| TAG548-FA182 | 0.001326 | 0.021581 | 0.11024 | ↓ | -3.181 | 0.00652 | 0.11015 | 0.13157 | ↓ | -2.926 | 0.000352 | 0.008616 | 0.075455 | ↓ | -3.728 |

| | | | | | | | | | | | | | | | |
|---|---|---|---|---|---|---|---|---|---|---|---|---|---|---|---|
| PC182/202 | 0.000373 | 0.018074 | 0.50402 | ↓ | -0.988 | 0.000893 | 0.10917 | 0.61774 | ↓ | -0.695 | 0.000466 | 0.009864 | 0.444 | ↓ | -1.171 |
| PI182/182 | 0.003127 | 0.029502 | 0.38076 | ↓ | -1.393 | 0.33734 | 0.50254 | 0.76948 | ↓ | -0.378 | 0.000544 | 0.009864 | 0.26326 | ↓ | -1.926 |
| TAG546-FA182 | 0.012491 | 0.053913 | 0.15762 | ↓ | -2.665 | 0.028807 | 0.12307 | 0.19058 | ↓ | -2.392 | 0.005786 | 0.021403 | 0.1262 | ↓ | -2.986 |
| PI181/182_2 | 0.000663 | 0.018074 | 0.37876 | ↓ | -1.401 | 0.46281 | 0.63163 | 1.3218 | ↑ | 0.4025 | 0.019618 | 0.044159 | 0.46627 | ↓ | -1.101 |
| PI160/180_2 | 0.02849 | 0.079931 | 0.55196 | ↓ | -0.857 | 0.63817 | 0.75536 | 0.87963 | ↓ | -0.185 | 0.022358 | 0.049352 | 0.53777 | ↓ | -0.895 |
| TAG525-FA160 | 0.002532 | 0.02658 | 0.21455 | ↓ | -2.221 | 0.004616 | 0.11015 | 0.18305 | ↓ | -2.45 | 0.001482 | 0.013784 | 0.18041 | ↓ | -2.471 |
| TAG525-FA161 | 4.84E-05 | 0.007869 | 0.16261 | ↓ | -2.621 | 0.005106 | 0.11015 | 0.29376 | ↓ | -1.767 | 0.000164 | 0.007659 | 0.17501 | ↓ | -2.515 |
| TAG505-FA161 | 0.003406 | 0.03053 | 0.12984 | ↓ | -2.945 | 0.016616 | 0.11428 | 0.19993 | ↓ | -2.322 | 0.001897 | 0.015371 | 0.099684 | ↓ | -3.327 |
| PG181/182 | 0.012734 | 0.053913 | 0.45381 | ↓ | -1.14 | 0.032946 | 0.12787 | 0.51254 | ↓ | -0.964 | 0.006155 | 0.021941 | 0.35111 | ↓ | -1.51 |
| TAG563-FA200 | 0.004585 | 0.033919 | 0.17413 | ↓ | -2.522 | 0.40463 | 0.57766 | 0.62989 | ↓ | -0.667 | 0.000982 | 0.012765 | 0.10584 | ↓ | -3.24 |
| PC160/204 | 0.24465 | 0.36446 | 1.2693 | ↑ | 0.3441 | 0.97654 | 0.98857 | 0.98452 | ↓ | -0.023 | 0.051785 | 0.092871 | 1.485 | ↑ | 0.5705 |
| PC182/182 | 3.21E-05 | 0.007869 | 0.40078 | ↓ | -1.319 | 0.036496 | 0.1282 | 0.71776 | ↓ | -0.478 | 0.002081 | 0.015416 | 0.40861 | ↓ | -1.291 |
| PI160/181 | 0.028333 | 0.079849 | 0.54792 | ↓ | -0.868 | 0.59561 | 0.73128 | 1.1197 | ↑ | 0.1632 | 0.032559 | 0.064621 | 0.53563 | ↓ | -0.901 |
| TAG482-FA182 | 0.00659 | 0.039785 | 0.26497 | ↓ | -1.916 | 0.021693 | 0.11636 | 0.31372 | ↓ | -1.673 | 0.004032 | 0.019163 | 0.23725 | ↓ | -2.076 |
| PG181/181 | 0.018041 | 0.063783 | 0.54278 | ↓ | -0.882 | 0.21087 | 0.35878 | 0.72144 | ↓ | -0.471 | 0.057831 | 0.10095 | 0.55479 | ↓ | -0.85 |
| TAG566-FA183 | 0.003456 | 0.03053 | 0.17583 | ↓ | -2.508 | 0.004973 | 0.11015 | 0.1801 | ↓ | -2.473 | 0.00075 | 0.010853 | 0.12052 | ↓ | -3.053 |
| PC182/161 | 0.00045 | 0.018074 | 0.50464 | ↓ | -0.987 | 0.12574 | 0.23894 | 0.77164 | ↓ | -0.374 | 0.031005 | 0.062104 | 0.59612 | ↓ | -0.746 |
| PI181/182 | 0.002068 | 0.024926 | 0.3987 | ↓ | -1.327 | 0.59399 | 0.73097 | 1.1932 | ↑ | 0.2548 | 0.019343 | 0.044159 | 0.46324 | ↓ | -1.11 |
| TAG524-FA221 | 0.003666 | 0.030709 | 0.29675 | ↓ | -1.753 | 0.005591 | 0.11015 | 0.29324 | ↓ | -1.77 | 0.000569 | 0.009864 | 0.23412 | ↓ | -2.095 |
| PI160/181_2 | 0.033364 | 0.086518 | 0.52956 | ↓ | -0.917 | 0.57896 | 0.72203 | 1.144 | ↑ | 0.194 | 0.029483 | 0.05998 | 0.51103 | ↓ | -0.969 |
| TAG541-FA200 | 0.002679 | 0.027255 | 0.16809 | ↓ | -2.573 | 0.036985 | 0.1282 | 0.30274 | ↓ | -1.724 | 0.000385 | 0.008651 | 0.09905 | ↓ | -3.336 |
| TAG548-FA183 | 0.001217 | 0.020849 | 0.082465 | ↓ | -3.6 | 0.006764 | 0.11015 | 0.10416 | ↓ | -3.263 | 0.000209 | 0.008255 | 0.046918 | ↓ | -4.414 |
| CE226 | 0.13292 | 0.23062 | 1.5942 | ↑ | 0.6728 | 0.62167 | 0.74289 | 1.0687 | ↑ | 0.0959 | 0.011914 | 0.031843 | 2.2842 | ↑ | 1.1917 |
| TAG503-FA182 | 0.000866 | 0.018074 | 0.21648 | ↓ | -2.208 | 0.013426 | 0.11428 | 0.32209 | ↓ | -1.635 | 0.003279 | 0.018829 | 0.24587 | ↓ | -2.024 |
| LPC183 | 0.021041 | 0.06918 | 0.56015 | ↓ | -0.836 | 0.024792 | 0.12086 | 0.59157 | ↓ | -0.757 | 0.003481 | 0.018829 | 0.34061 | ↓ | -1.554 |
| TAG525-FA181 | 0.000878 | 0.018074 | 0.1742 | ↓ | -2.521 | 0.007489 | 0.11015 | 0.23942 | ↓ | -2.062 | 0.00109 | 0.012765 | 0.1494 | ↓ | -2.743 |
| PC140/226 | 0.053889 | 0.12181 | 0.72179 | ↓ | -0.47 | 0.68225 | 0.78749 | 1.0535 | ↑ | 0.0752 | 0.68758 | 0.74478 | 0.91949 | ↓ | -0.121 |
| LPC150 | 0.014537 | 0.056667 | 0.56201 | ↓ | -0.831 | 0.038632 | 0.12835 | 0.61283 | ↓ | -0.706 | 0.001987 | 0.015416 | 0.39366 | ↓ | -1.345 |
| PS161/181 | 0.10919 | 0.19911 | 0.59349 | ↓ | -0.753 | 0.043405 | 0.13022 | 1.4048 | ↑ | 0.4904 | 0.059443 | 0.10265 | 0.60173 | ↓ | -0.733 |
| PC180/201 | 0.1081 | 0.1988 | 0.72113 | ↓ | -0.472 | 0.030518 | 0.12449 | 0.6398 | ↓ | -0.644 | 0.010015 | 0.029105 | 0.58876 | ↓ | -0.764 |
| PI181/181 | 0.009074 | 0.046514 | 0.4053 | ↓ | -1.303 | 0.29149 | 0.45514 | 2.119 | ↑ | 1.0834 | 0.1137 | 0.17254 | 0.60258 | ↓ | -0.731 |
| PI160/203_2 | 0.00749 | 0.042113 | 0.55967 | ↓ | -0.837 | 0.18402 | 0.32395 | 0.77924 | ↓ | -0.36 | 0.005418 | 0.020746 | 0.55899 | ↓ | -0.839 |
| TAG562-FA200 | 0.010841 | 0.051143 | 0.22205 | ↓ | -2.171 | 0.46054 | 0.63118 | 0.72014 | ↓ | -0.474 | 0.002606 | 0.016964 | 0.12656 | ↓ | -2.982 |

| | | | | | | | | | | | | | | | |
|---|---|---|---|---|---|---|---|---|---|---|---|---|---|---|---|
| PC180/202 | 0.028985 | 0.080294 | 0.65364 | ↓ | -0.613 | 0.027404 | 0.12263 | 0.60257 | ↓ | -0.731 | 0.001238 | 0.012788 | 0.53234 | ↓ | -0.91 |
| TAG547-FA181 | 0.001811 | 0.023374 | 0.12339 | ↓ | -3.019 | 0.006181 | 0.11015 | 0.14449 | ↓ | -2.791 | 0.001298 | 0.013002 | 0.10659 | ↓ | -3.23 |
| TAG503-FA181 | 0.001759 | 0.023374 | 0.20184 | ↓ | -2.309 | 0.035275 | 0.12787 | 0.36177 | ↓ | -1.467 | 0.001586 | 0.013948 | 0.1736 | ↓ | -2.526 |
| TAG484-FA120 | 0.022847 | 0.071165 | 0.3526 | ↓ | -1.504 | 0.22771 | 0.37913 | 0.71879 | ↓ | -0.476 | 0.003716 | 0.018938 | 0.22115 | ↓ | -2.177 |
| TAG504-FA161 | 0.00086 | 0.018074 | 0.1633 | ↓ | -2.614 | 0.006182 | 0.11015 | 0.23325 | ↓ | -2.1 | 0.002661 | 0.016986 | 0.17037 | ↓ | -2.553 |
| PG160/203_2 | 0.24991 | 0.36976 | 0.82551 | ↓ | -0.277 | 0.022463 | 0.11722 | 0.54481 | ↓ | -0.876 | 0.000132 | 0.007659 | 0.35659 | ↓ | -1.488 |
| TAG524-FA161 | 8.81E-05 | 0.008861 | 0.19164 | ↓ | -2.384 | 0.007424 | 0.11015 | 0.36846 | ↓ | -1.44 | 0.000744 | 0.010853 | 0.22488 | ↓ | -2.153 |
| TAG563-FA182 | 0.006365 | 0.03946 | 0.20477 | ↓ | -2.288 | 0.35551 | 0.52125 | 0.58539 | ↓ | -0.773 | 0.001722 | 0.014554 | 0.13965 | ↓ | -2.84 |
| TAG565-FA201 | 0.025085 | 0.074569 | 0.18748 | ↓ | -2.415 | 0.084519 | 0.18219 | 0.27476 | ↓ | -1.864 | 0.010398 | 0.02956 | 0.14673 | ↓ | -2.769 |
| PI160/203 | 0.018566 | 0.063783 | 0.61821 | ↓ | -0.694 | 0.3064 | 0.47155 | 0.83417 | ↓ | -0.262 | 0.00588 | 0.021627 | 0.5624 | ↓ | -0.83 |
| PI180/180 | 0.028708 | 0.079931 | 0.57145 | ↓ | -0.807 | 0.56819 | 0.71133 | 1.2369 | ↑ | 0.3067 | 0.039678 | 0.075088 | 0.55341 | ↓ | -0.854 |
| TAG541-FA160 | 0.007323 | 0.041817 | 0.17039 | ↓ | -2.553 | 0.074346 | 0.17078 | 0.30214 | ↓ | -1.727 | 0.002031 | 0.015416 | 0.12063 | ↓ | -3.051 |
| PC182/224 | 0.017014 | 0.062716 | 0.54803 | ↓ | -0.868 | 0.26722 | 0.4268 | 0.76851 | ↓ | -0.38 | 0.017002 | 0.041249 | 0.53414 | ↓ | -0.905 |
| PC182/205 | 0.006076 | 0.0384 | 0.42534 | ↓ | -1.233 | 0.081232 | 0.17949 | 0.67098 | ↓ | -0.576 | 0.006003 | 0.021904 | 0.41444 | ↓ | -1.271 |
| TAG545-FA180 | 0.014253 | 0.056129 | 0.1742 | ↓ | -2.521 | 0.006883 | 0.11015 | 0.12607 | ↓ | -2.988 | 0.00224 | 0.016124 | 0.10758 | ↓ | -3.217 |
| PI161/181_2 | 0.010416 | 0.050227 | 0.48557 | ↓ | -1.042 | 0.87181 | 0.9154 | 1.0424 | ↑ | 0.0598 | 0.014539 | 0.036545 | 0.52409 | ↓ | -0.932 |
| TAG524-FA160 | 0.01017 | 0.049771 | 0.38375 | ↓ | -1.382 | 0.019028 | 0.11428 | 0.36888 | ↓ | -1.439 | 0.008886 | 0.027287 | 0.37065 | ↓ | -1.432 |
| PC160/225 | 0.001673 | 0.023374 | 1.4385 | ↑ | 0.5246 | 0.46223 | 0.63163 | 1.1068 | ↑ | 0.1464 | 0.0297 | 0.060232 | 1.5087 | ↑ | 0.5934 |
| TAG524-FA183 | 0.002215 | 0.02531 | 0.22 | ↓ | -2.184 | 0.00813 | 0.11015 | 0.2443 | ↓ | -2.033 | 0.001071 | 0.012765 | 0.18652 | ↓ | -2.423 |
| TAG561-FA181 | 0.023488 | 0.072469 | 0.18181 | ↓ | -2.46 | 0.48428 | 0.65066 | 0.60664 | ↓ | -0.721 | 0.003709 | 0.018938 | 0.1079 | ↓ | -3.212 |
| TAG482-FA140 | 0.013634 | 0.055199 | 0.26612 | ↓ | -1.91 | 0.025833 | 0.12086 | 0.29081 | ↓ | -1.782 | 0.00531 | 0.020577 | 0.20018 | ↓ | -2.321 |
| PI200/204_2 | 0.048502 | 0.11197 | 0.61579 | ↓ | -0.699 | 0.84906 | 0.90613 | 0.98448 | ↓ | -0.023 | 0.060897 | 0.10433 | 0.64036 | ↓ | -0.643 |
| TAG502-FA140 | 0.002996 | 0.029174 | 0.22438 | ↓ | -2.156 | 0.12589 | 0.23894 | 0.48138 | ↓ | -1.055 | 0.002845 | 0.017482 | 0.17768 | ↓ | -2.493 |
| TAG545-FA183 | 0.003107 | 0.029502 | 0.14799 | ↓ | -2.756 | 0.017505 | 0.11428 | 0.21288 | ↓ | -2.232 | 0.000698 | 0.010812 | 0.098158 | ↓ | -3.349 |
| PG160/203 | 0.32376 | 0.45036 | 0.87043 | ↓ | -0.2 | 0.015717 | 0.11428 | 0.56497 | ↓ | -0.824 | 7.31E-05 | 0.007659 | 0.38151 | ↓ | -1.39 |
| PI160/161 | 0.046831 | 0.10967 | 0.51834 | ↓ | -0.948 | 0.8531 | 0.90746 | 1.0234 | ↑ | 0.0334 | 0.063307 | 0.10677 | 0.56068 | ↓ | -0.835 |
| PE182/202 | 0.003513 | 0.03053 | 0.49643 | ↓ | -1.01 | 0.001006 | 0.10917 | 0.51306 | ↓ | -0.963 | 0.00024 | 0.008255 | 0.24907 | ↓ | -2.005 |
| PI182/201 | 0.022706 | 0.071165 | 0.50748 | ↓ | -0.979 | 0.18379 | 0.32395 | 1.3413 | ↑ | 0.4237 | 0.079523 | 0.1272 | 0.57933 | ↓ | -0.788 |
| LPA180 | 0.42142 | 0.55089 | 0.86346 | ↓ | -0.212 | 0.51122 | 0.66694 | 0.89067 | ↓ | -0.167 | 0.97498 | 0.97798 | 1.0141 | ↑ | 0.0202 |
| LPC240 | 0.000435 | 0.018074 | 0.49076 | ↓ | -1.027 | 0.002959 | 0.11015 | 0.61934 | ↓ | -0.691 | 0.004906 | 0.01996 | 0.51762 | ↓ | -0.95 |
| TAG524-FA200 | 0.014312 | 0.056129 | 0.35913 | ↓ | -1.477 | 0.025893 | 0.12086 | 0.348 | ↓ | -1.523 | 0.010628 | 0.030082 | 0.32716 | ↓ | -1.612 |
| TAG566-FA202 | 0.006213 | 0.038888 | 0.24289 | ↓ | -2.042 | 0.014822 | 0.11428 | 0.26027 | ↓ | -1.942 | 0.002254 | 0.016124 | 0.18278 | ↓ | -2.452 |
| PS181/182 | 0.092542 | 0.17581 | 0.77287 | ↓ | -0.372 | 0.31081 | 0.47609 | 1.1834 | ↑ | 0.2429 | 0.005345 | 0.020588 | 0.57795 | ↓ | -0.791 |

| | | | | | | | | | | | | |
|---|---|---|---|---|---|---|---|---|---|---|---|---|
| TAG524-FA181 | 0.000577 | 0.018074 | 0.21488 | ↓ -2.218 | 0.009025 | 0.11015 | 0.30253 | ↓ -1.725 | 0.001123 | 0.012765 | 0.22159 | ↓ -2.174 |
| PG161/180 | 0.81432 | 0.89096 | 1.2883 | ↑ 0.3655 | 0.039668 | 0.12837 | 0.4837 | ↓ -1.048 | 0.009824 | 0.02868 | 0.35839 | ↓ -1.48 |
| PI180/203_2 | 0.32006 | 0.44616 | 0.89742 | ↓ -0.156 | 0.98501 | 0.98981 | 1.0123 | ↑ 0.0176 | 0.46791 | 0.54589 | 0.92978 | ↓ -0.105 |
| PI180/183_2 | 0.047675 | 0.11094 | 0.67923 | ↓ -0.558 | 0.46712 | 0.63485 | 0.89485 | ↓ -0.16 | 0.008942 | 0.027307 | 0.55132 | ↓ -0.859 |
| LPC241 | 0.000109 | 0.008861 | 0.36149 | ↓ -1.468 | 0.001289 | 0.11015 | 0.53767 | ↓ -0.895 | 0.000483 | 0.009864 | 0.39288 | ↓ -1.348 |
| CE203 | 0.65042 | 0.75994 | 1.0868 | ↑ 0.12 | 0.25206 | 0.40819 | 0.68439 | ↓ -0.547 | 0.56662 | 0.63598 | 1.1136 | ↑ 0.1552 |
| TAG524-FA182 | 0.008914 | 0.046514 | 0.37522 | ↓ -1.414 | 0.022576 | 0.11722 | 0.39031 | ↓ -1.357 | 0.008735 | 0.026949 | 0.37585 | ↓ -1.412 |
| TAG545-FA182 | 0.011595 | 0.05269 | 0.20074 | ↓ -2.317 | 0.091437 | 0.19389 | 0.34612 | ↓ -1.531 | 0.002839 | 0.017482 | 0.14128 | ↓ -2.823 |
| TAG544-FA201 | 0.002416 | 0.025783 | 0.2222 | ↓ -2.17 | 0.027924 | 0.12263 | 0.33736 | ↓ -1.568 | 0.004134 | 0.019361 | 0.22876 | ↓ -2.128 |
| TAG545-FA161 | 0.000798 | 0.018074 | 0.22713 | ↓ -2.138 | 0.004348 | 0.11015 | 0.25593 | ↓ -1.966 | 0.001444 | 0.013627 | 0.20961 | ↓ -2.254 |
| TAG502-FA181 | 0.004674 | 0.034185 | 0.21272 | ↓ -2.233 | 0.073199 | 0.17078 | 0.38414 | ↓ -1.38 | 0.00637 | 0.022215 | 0.19827 | ↓ -2.335 |
| PG181/182_2 | 0.011453 | 0.05269 | 0.43997 | -1.1845 | 0.056297 | 0.14838 | 0.54 | -0.88897 | 0.009127 | 0.027507 | 0.37652 | -1.4092 |
| PC200/182 | 0.00264 | 0.027255 | 0.6454 | -0.63173 | 0.33879 | 0.50262 | 0.87994 | -0.18453 | 0.00118 | 0.012765 | 0.50554 | -0.98412 |
| PC140/203 | 0.034879 | 0.088847 | 0.59363 | -0.75236 | 0.31424 | 0.47908 | 0.81287 | -0.29889 | 0.004702 | 0.019874 | 0.4669 | -1.0988 |
| TAG543-FA183 | 0.013774 | 0.055199 | 0.18042 | -2.4706 | 0.013142 | 0.11428 | 0.1741 | -2.522 | 0.001913 | 0.015371 | 0.10567 | -3.2424 |
| PC181/203 | 0.029501 | 0.081034 | 0.56787 | -0.81638 | 0.027979 | 0.12263 | 0.57071 | -0.80916 | 0.042487 | 0.079708 | 0.57788 | -0.79116 |
| TAG523-FA201 | 0.003198 | 0.029583 | 0.22953 | -2.1232 | 0.018709 | 0.11428 | 0.30899 | -1.6944 | 0.001997 | 0.015416 | 0.20571 | -2.2813 |
| LPC201 | 0.065775 | 0.13768 | 0.70337 | -0.50764 | 0.25419 | 0.4096 | 0.81669 | -0.29214 | 0.051998 | 0.092997 | 0.63652 | -0.65173 |
| PE181/205 | 0.20503 | 0.3148 | 0.73407 | -0.44602 | 0.21985 | 0.36868 | 0.74499 | -0.4247 | 0.014053 | 0.035598 | 0.46118 | -1.1166 |
| LPC161 | 0.005359 | 0.035933 | 0.56358 | -0.82731 | 0.1076 | 0.21421 | 0.72899 | -0.45603 | 0.0274 | 0.056626 | 0.55981 | -0.83698 |
| TAG481-FA181 | 0.013149 | 0.054523 | 0.25543 | -1.969 | 0.066154 | 0.16251 | 0.35777 | -1.4829 | 0.008122 | 0.025793 | 0.21474 | -2.2193 |
| PC120/181 | 0.19088 | 0.29943 | 0.6919 | -0.53137 | 0.33547 | 0.5009 | 0.77042 | -0.37628 | 0.045365 | 0.08332 | 0.51805 | -0.94885 |
| PC150/203 | 0.024566 | 0.074384 | 0.61314 | -0.70572 | 0.014157 | 0.11428 | 0.54101 | -0.88628 | 0.0011 | 0.012765 | 0.46153 | -1.1155 |
| TAG567-FA183 | 0.001551 | 0.023374 | 0.24522 | -2.0278 | 0.006735 | 0.11015 | 0.26136 | -1.9359 | 0.001842 | 0.015268 | 0.2509 | -1.9948 |
| TAG544-FA183 | 0.013475 | 0.055199 | 0.19188 | -2.3817 | 0.019852 | 0.11428 | 0.20798 | -2.2655 | 0.003775 | 0.018938 | 0.13945 | -2.8421 |
| TAG523-FA183 | 0.022009 | 0.069893 | 0.237 | -2.0771 | 0.016471 | 0.11428 | 0.2021 | -2.3068 | 0.005482 | 0.020761 | 0.17054 | -2.5518 |
| PE182/201 | 0.068743 | 0.14073 | 0.58752 | -0.76728 | 0.45189 | 0.62459 | 0.80709 | -0.30919 | 0.004761 | 0.019874 | 0.3544 | -1.4966 |
| TAG544-FA180 | 0.025437 | 0.074593 | 0.26694 | -1.9054 | 0.017605 | 0.11428 | 0.23057 | -2.1167 | 0.00523 | 0.020577 | 0.18787 | -2.4122 |
| TAG482-FA161 | 0.018912 | 0.064124 | 0.22795 | -2.1332 | 0.063474 | 0.16016 | 0.34082 | -1.5529 | 0.008714 | 0.026949 | 0.1911 | -2.3876 |
| TAG482-FA160 | 0.012815 | 0.053913 | 0.28762 | -1.7978 | 0.021807 | 0.11636 | 0.30553 | -1.7106 | 0.009303 | 0.027527 | 0.2535 | -1.98 |
| PI180/181_2 | 0.031612 | 0.084435 | 0.61778 | -0.69483 | 0.33031 | 0.49661 | 1.4053 | 0.49087 | 0.035363 | 0.06872 | 0.56216 | -0.83095 |
| TAG523-FA180 | 0.021357 | 0.069517 | 0.2505 | -1.9971 | 0.018301 | 0.11428 | 0.22641 | -2.143 | 0.005272 | 0.020577 | 0.17525 | -2.5125 |
| PG160/182_2 | 0.098583 | 0.18491 | 0.59344 | -0.75283 | 0.049556 | 0.14066 | 0.51165 | -0.96676 | 0.008205 | 0.025825 | 0.41182 | -1.2799 |

| | | | | | | | | | | | | |
|---|---|---|---|---|---|---|---|---|---|---|---|---|
| TAG503-FA161 | 0.001413 | 0.022442 | 0.22575 | -2.1472 | 0.015146 | 0.11428 | 0.3152 | -1.6657 | 0.0041 | 0.019339 | 0.2568 | -1.9613 |
| PI180/181 | 0.02332 | 0.072291 | 0.56076 | -0.83454 | 0.48881 | 0.65238 | 1.3821 | 0.4669 | 0.022776 | 0.049925 | 0.49713 | -1.0083 |
| PC182/226 | 0.000499 | 0.018074 | 0.42751 | -1.226 | 0.0201 | 0.11428 | 0.70523 | -0.50384 | 0.00069 | 0.010812 | 0.49468 | -1.0154 |
| PC181/183 | 0.008427 | 0.044966 | 0.56151 | -0.83262 | 0.020804 | 0.11478 | 0.62984 | -0.66693 | 0.003447 | 0.018829 | 0.38324 | -1.3837 |
| PC182/204 | 0.00368 | 0.030709 | 0.51709 | -0.9515 | 0.12864 | 0.24264 | 0.75923 | -0.39739 | 0.042759 | 0.07999 | 0.61722 | -0.69614 |
| TAG565-FA182 | 0.001939 | 0.023817 | 0.19075 | -2.3902 | 0.022868 | 0.11722 | 0.29289 | -1.7716 | 0.001216 | 0.012765 | 0.16289 | -2.618 |
| TAG544-FA161 | 0.001023 | 0.019029 | 0.19469 | -2.3607 | 0.048866 | 0.14014 | 0.38295 | -1.3848 | 0.003453 | 0.018829 | 0.19968 | -2.3243 |
| PC181/226 | 0.029253 | 0.080692 | 0.67016 | -0.57742 | 0.016336 | 0.11428 | 0.64957 | -0.62245 | 0.15916 | 0.22722 | 0.78633 | -0.34679 |
| TAG520-FA200 | 0.027291 | 0.077923 | 0.20196 | -2.3079 | 0.008422 | 0.11015 | 0.13995 | -2.837 | 0.00204 | 0.015416 | 0.09268 | -3.4316 |
| SMd181/182 | 0.48398 | 0.60942 | 1.0656 | 0.091706 | 0.44385 | 0.61741 | 0.93892 | -0.09092 | 0.082806 | 0.13084 | 1.2723 | 0.34742 |
| PC140/202 | 0.031776 | 0.084435 | 0.57823 | -0.79028 | 0.179 | 0.32029 | 0.70921 | -0.49571 | 0.00348 | 0.018829 | 0.38986 | -1.359 |
| TAG524-FA180 | 0.008285 | 0.044945 | 0.2216 | -2.174 | 0.012457 | 0.11428 | 0.222 | -2.1714 | 0.004202 | 0.019527 | 0.18405 | -2.4418 |
| PG160/182 | 0.1075 | 0.19874 | 0.58647 | -0.76987 | 0.063801 | 0.16036 | 0.5241 | -0.93207 | 0.010124 | 0.029199 | 0.41224 | -1.2784 |
| PI180/182_2 | 0.090361 | 0.17404 | 0.73909 | -0.43618 | 0.49975 | 0.66531 | 1.1136 | 0.15527 | 0.049247 | 0.089303 | 0.70064 | -0.51326 |
| TAG526-FA181 | 0.005215 | 0.035521 | 0.20673 | -2.2742 | 0.019729 | 0.11428 | 0.24797 | -2.0118 | 0.007679 | 0.02474 | 0.19737 | -2.341 |
| TAG483-FA161 | 0.005238 | 0.035521 | 0.21359 | -2.2271 | 0.030978 | 0.12449 | 0.33387 | -1.5827 | 0.004033 | 0.019163 | 0.19747 | -2.3403 |
| TAG565-FA202 | 0.004261 | 0.033421 | 0.25852 | -1.9516 | 0.016594 | 0.11428 | 0.30668 | -1.7052 | 0.002547 | 0.016786 | 0.22046 | -2.1814 |
| LPC170 | 0.007971 | 0.04435 | 0.59601 | -0.74659 | 0.013631 | 0.11428 | 0.60175 | -0.73277 | 0.011523 | 0.031387 | 0.54762 | -0.86876 |
| TAG545-FA181 | 0.011655 | 0.05269 | 0.20536 | -2.2838 | 0.098066 | 0.20259 | 0.35457 | -1.4959 | 0.003927 | 0.018938 | 0.15633 | -2.6773 |
| TAG523-FA202 | 0.004756 | 0.034394 | 0.26491 | -1.9164 | 0.025073 | 0.12086 | 0.3189 | -1.6488 | 0.005173 | 0.020577 | 0.2511 | -1.9936 |
| PI180/182 | 0.059731 | 0.13005 | 0.71288 | -0.48827 | 0.70478 | 0.80776 | 1.0651 | 0.090941 | 0.025259 | 0.053562 | 0.64322 | -0.63661 |
| TAG540-FA160 | 0.012574 | 0.053913 | 0.20325 | -2.2987 | 0.00921 | 0.11015 | 0.18573 | -2.4287 | 0.002494 | 0.016786 | 0.13463 | -2.8929 |
| TAG526-FA160 | 0.004028 | 0.032779 | 0.17252 | -2.5351 | 0.008646 | 0.11015 | 0.1931 | -2.3726 | 0.003364 | 0.018829 | 0.16055 | -2.6389 |
| TAG526-FA205 | 0.039298 | 0.095816 | 0.38313 | -1.3841 | 0.051913 | 0.14281 | 0.42724 | -1.2269 | 0.19954 | 0.27463 | 0.81824 | -0.28941 |
| PS180/181 | 0.11773 | 0.21231 | 0.70868 | -0.4968 | 0.44736 | 0.62096 | 0.86648 | -0.20677 | 0.056174 | 0.098836 | 0.64035 | -0.64307 |
| TAG523-FA182 | 0.015036 | 0.057788 | 0.39091 | -1.3551 | 0.053284 | 0.14281 | 0.46461 | -1.1059 | 0.010226 | 0.029199 | 0.37153 | -1.4285 |
| TAG545-FA202 | 0.001289 | 0.021522 | 0.20617 | -2.2781 | 0.017156 | 0.11428 | 0.29904 | -1.7416 | 0.000787 | 0.011134 | 0.16635 | -2.5877 |
| TAG483-FA140 | 0.004886 | 0.034573 | 0.17718 | -2.4967 | 0.009644 | 0.11015 | 0.20093 | -2.3152 | 0.002507 | 0.016786 | 0.1175 | -3.0893 |
| PE180/202 | 0.6505 | 0.75994 | 0.92676 | -0.10973 | 0.034853 | 0.12787 | 0.61332 | -0.70529 | 0.004349 | 0.019527 | 0.55475 | -0.8501 |
| PI180/202_2 | 0.091282 | 0.17478 | 0.73778 | -0.43873 | 0.16405 | 0.29748 | 0.77368 | -0.37018 | 0.004279 | 0.019527 | 0.60918 | -0.71507 |
| PE181/204 | 0.056774 | 0.12614 | 0.65029 | -0.62085 | 0.088815 | 0.19082 | 0.70488 | -0.50456 | 0.061606 | 0.10471 | 0.6102 | -0.71265 |
| TAG523-FA181 | 0.01362 | 0.055199 | 0.38654 | -1.3713 | 0.059759 | 0.15256 | 0.47396 | -1.0772 | 0.011502 | 0.031387 | 0.38297 | -1.3847 |
| TAG503-FA160 | 0.005834 | 0.037604 | 0.2633 | -1.9252 | 0.012467 | 0.11428 | 0.26676 | -1.9064 | 0.004707 | 0.019874 | 0.25016 | -1.9991 |

| | | | | | | | | | | | | |
|---|---|---|---|---|---|---|---|---|---|---|---|---|
| TAG483-FA182 | 0.00912 | 0.046514 | 0.26644 | -1.9081 | 0.56036 | 0.70696 | 0.80923 | -0.30538 | 0.009624 | 0.02835 | 0.23594 | -2.0835 |
| PC140/161 | 0.033491 | 0.086518 | 0.62494 | -0.67821 | 0.099799 | 0.20367 | 0.6781 | -0.56044 | 0.011034 | 0.030873 | 0.55548 | -0.84819 |
| TAG523-FA161 | 0.00034 | 0.018074 | 0.17221 | -2.5378 | 0.030823 | 0.12449 | 0.37912 | -1.3993 | 0.002872 | 0.017482 | 0.21503 | -2.2174 |
| TAG5610-FA182 | 0.012685 | 0.053913 | 0.15061 | -2.7311 | 0.21409 | 0.36201 | 0.44196 | -1.178 | 0.00577 | 0.021403 | 0.11982 | -3.0611 |
| PC180/183 | 0.12706 | 0.22236 | 0.65529 | -0.6098 | 0.093977 | 0.19665 | 0.62808 | -0.67098 | 0.011324 | 0.031369 | 0.41362 | -1.2736 |
| TAG564-FA201 | 0.018082 | 0.063783 | 0.20662 | -2.2749 | 0.32545 | 0.49157 | 0.53796 | -0.89442 | 0.011476 | 0.031387 | 0.18365 | -2.445 |
| PE182/161 | 0.005064 | 0.035072 | 0.53934 | -0.89074 | 0.5888 | 0.72734 | 0.90031 | -0.15151 | 0.00029 | 0.008616 | 0.35691 | -1.4864 |
| PE181/226 | 0.030283 | 0.082447 | 0.62175 | -0.6856 | 0.037434 | 0.1282 | 0.65815 | -0.60351 | 0.079207 | 0.1272 | 0.67664 | -0.56354 |
| TAG481-FA140 | 0.0128 | 0.053913 | 0.26697 | -1.9052 | 0.025804 | 0.12086 | 0.30751 | -1.7013 | 0.004713 | 0.019874 | 0.20297 | -2.3007 |
| PS161/182_2 | 0.012316 | 0.053913 | 0.72891 | -0.45618 | 0.85286 | 0.90746 | 1.0342 | 0.048478 | 0.000109 | 0.007659 | 0.48786 | -1.0355 |
| PC140/181 | 0.006793 | 0.039785 | 0.58623 | -0.77045 | 0.62497 | 0.74379 | 0.93934 | -0.09027 | 0.018265 | 0.042618 | 0.55147 | -0.85865 |
| PI160/161_2 | 0.055569 | 0.12483 | 0.57735 | -0.79248 | 0.87745 | 0.91757 | 0.99156 | -0.01223 | 0.079406 | 0.1272 | 0.6096 | -0.71406 |
| PA180/182 | 0.003226 | 0.029583 | 0.61746 | -0.69558 | 0.86915 | 0.9154 | 1.107 | 0.14669 | 0.083466 | 0.13156 | 0.75065 | -0.4138 |
| TAG504-FA160 | 0.00225 | 0.02531 | 0.21156 | -2.2409 | 0.01514 | 0.11428 | 0.26439 | -1.9192 | 0.002425 | 0.016786 | 0.19079 | -2.39 |
| PC182/225 | 3.19E-05 | 0.007869 | 0.48961 | -1.0303 | 0.014487 | 0.11428 | 0.71637 | -0.48123 | 0.00351 | 0.018829 | 0.65452 | -0.61149 |
| TAG523-FA160 | 0.015959 | 0.060754 | 0.39644 | -1.3348 | 0.049695 | 0.14066 | 0.46015 | -1.1198 | 0.022364 | 0.049352 | 0.42756 | -1.2258 |
| PI180/202 | 0.15097 | 0.25007 | 0.81319 | -0.29833 | 0.23994 | 0.39275 | 0.83656 | -0.25746 | 0.013603 | 0.035002 | 0.68739 | -0.54079 |
| PS180/181_2 | 0.20224 | 0.31124 | 0.79429 | -0.33226 | 0.62986 | 0.74816 | 0.92614 | -0.1107 | 0.12753 | 0.18784 | 0.7637 | -0.38893 |
| SMd181/203 | 0.13256 | 0.23062 | 1.1361 | 0.18405 | 0.17928 | 0.32029 | 0.91443 | -0.12906 | 0.4512 | 0.5302 | 1.0939 | 0.12945 |
| PE181/202 | 0.061069 | 0.13121 | 0.65567 | -0.60896 | 0.010017 | 0.11053 | 0.60266 | -0.73057 | 0.002553 | 0.016786 | 0.37866 | -1.401 |
| TAG484-FA182 | 0.018216 | 0.063783 | 0.36743 | -1.4445 | 0.50736 | 0.66694 | 0.92654 | -0.11008 | 0.013265 | 0.034542 | 0.29595 | -1.7566 |
| PI180/205 | 0.83563 | 0.90365 | 0.97122 | -0.04213 | 0.82904 | 0.89503 | 1.0915 | 0.12633 | 0.92626 | 0.94365 | 1.1501 | 0.20175 |
| PEP-180/182 | 0.00211 | 0.024976 | 0.5984 | -0.74081 | 0.7744 | 0.85592 | 0.97605 | -0.03498 | 7.2E-05 | 0.007659 | 0.36508 | -1.4537 |
| PE181/182 | 0.020391 | 0.067383 | 0.61095 | -0.71086 | 0.37625 | 0.54674 | 0.83372 | -0.26237 | 0.002873 | 0.017482 | 0.35536 | -1.4927 |
| PEO-160/225 | 0.21618 | 0.32729 | 1.2291 | 0.29764 | 0.94393 | 0.96594 | 1.007 | 0.010125 | 0.11524 | 0.17402 | 1.3846 | 0.46946 |
| PI180/203 | 0.40565 | 0.53457 | 0.92161 | -0.11777 | 0.97794 | 0.98857 | 1.0127 | 0.018152 | 0.37617 | 0.45349 | 0.91113 | -0.13428 |
| LPC226 | 0.37822 | 0.50662 | 1.1972 | 0.25963 | 0.46007 | 0.63118 | 1.1302 | 0.17653 | 0.25236 | 0.32625 | 1.3617 | 0.44538 |
| PEP-180/181 | 0.024725 | 0.074517 | 0.64612 | -0.63013 | 0.71993 | 0.8208 | 1.1838 | 0.24344 | 0.000616 | 0.010024 | 0.40354 | -1.3092 |
| TAG569-FA204 | 0.049295 | 0.11333 | 0.42256 | -1.2428 | 0.027481 | 0.12263 | 0.31736 | -1.6558 | 0.22915 | 0.3032 | 0.62474 | -0.67866 |
| PC140/182 | 0.001702 | 0.023374 | 0.58887 | -0.76399 | 0.57687 | 0.72081 | 1.2329 | 0.30206 | 0.090045 | 0.14125 | 0.6714 | -0.57476 |
| TAG544-FA182 | 0.014107 | 0.055998 | 0.27784 | -1.8477 | 0.22987 | 0.38176 | 0.54766 | -0.86864 | 0.006432 | 0.022274 | 0.23549 | -2.0862 |
| TAG522-FA181 | 0.01504 | 0.057788 | 0.35155 | -1.5082 | 0.11717 | 0.22578 | 0.50934 | -0.9733 | 0.017357 | 0.041541 | 0.36307 | -1.4617 |
| PEP-180/203 | 0.13983 | 0.23644 | 0.7909 | -0.33844 | 0.66801 | 0.77934 | 0.94119 | -0.08745 | 0.009107 | 0.027507 | 0.57064 | -0.80934 |

| | | | | | | | | | | | | |
|---|---|---|---|---|---|---|---|---|---|---|---|---|
| PC150/182 | 0.001903 | 0.023817 | 0.53633 | -0.8988 | 0.026759 | 0.12263 | 0.67289 | -0.57156 | 0.000342 | 0.008616 | 0.44125 | -1.1803 |
| TAG566-FA182 | 0.006679 | 0.039785 | 0.31771 | -1.6542 | 0.019883 | 0.11428 | 0.34382 | -1.5403 | 0.018876 | 0.043421 | 0.36913 | -1.4378 |
| PC182/201 | 0.012919 | 0.053913 | 0.60291 | -0.73 | 0.50684 | 0.66694 | 0.88565 | -0.17519 | 0.018161 | 0.042564 | 0.52325 | -0.93443 |
| TAG567-FA182 | 0.005956 | 0.038012 | 0.41769 | -1.2595 | 0.027722 | 0.12263 | 0.40958 | -1.2878 | 0.057843 | 0.10095 | 0.50731 | -0.97906 |
| PC160/205 | 0.25726 | 0.37551 | 1.3563 | 0.4397 | 0.65745 | 0.76979 | 0.99189 | -0.01175 | 0.45231 | 0.53054 | 2.2733 | 1.1848 |
| TAG568-FA183 | 0.012888 | 0.053913 | 0.31079 | -1.686 | 0.051254 | 0.14259 | 0.37199 | -1.4267 | 0.018458 | 0.042916 | 0.31919 | -1.6475 |
| TAG524-FA203 | 0.011595 | 0.05269 | 0.28833 | -1.7942 | 0.037612 | 0.1282 | 0.33644 | -1.5716 | 0.004814 | 0.019885 | 0.2375 | -2.074 |
| HCER180 | 0.060988 | 0.13121 | 1.2946 | 0.37253 | 0.54832 | 0.69719 | 0.94152 | -0.08693 | 0.58999 | 0.65543 | 1.1206 | 0.16422 |
| TAG567-FA203 | 0.004509 | 0.033873 | 0.31338 | -1.674 | 0.009986 | 0.11053 | 0.30421 | -1.7169 | 0.005297 | 0.020577 | 0.26879 | -1.8955 |
| PC170/203 | 0.20077 | 0.31004 | 0.73295 | -0.44821 | 0.040577 | 0.12837 | 0.59199 | -0.75634 | 0.006365 | 0.022215 | 0.48548 | -1.0425 |
| TAG502-FA180 | 0.046702 | 0.10967 | 0.27889 | -1.8423 | 0.020342 | 0.11428 | 0.22071 | -2.1798 | 0.0057 | 0.021327 | 0.15825 | -2.6597 |
| TAG483-FA120 | 0.16143 | 0.26404 | 0.54154 | -0.88486 | 0.74887 | 0.83909 | 2.0753 | 1.0533 | 0.015703 | 0.039168 | 0.28917 | -1.79 |
| SMd181/160 | 0.1251 | 0.22011 | 1.0552 | 0.077506 | 0.002952 | 0.11015 | 0.84754 | -0.23864 | 0.000737 | 0.010853 | 1.2523 | 0.32458 |
| TAG544-FA202 | 0.004513 | 0.033873 | 0.31727 | -1.6562 | 0.018704 | 0.11428 | 0.34942 | -1.517 | 0.002917 | 0.017584 | 0.27235 | -1.8764 |
| TAG502-FA161 | 0.005716 | 0.03721 | 0.22722 | -2.1378 | 0.020538 | 0.11428 | 0.28719 | -1.7999 | 0.011452 | 0.031387 | 0.24719 | -2.0163 |
| PE140/204 | 0.14625 | 0.24476 | 0.79498 | -0.331 | 0.078285 | 0.17696 | 0.7266 | -0.46078 | 0.010184 | 0.029199 | 0.473 | -1.0801 |
| TAG563-FA202 | 0.006837 | 0.039785 | 0.2725 | -1.8757 | 0.025317 | 0.12086 | 0.32507 | -1.6212 | 0.001569 | 0.013948 | 0.18946 | -2.4001 |
| TAG569-FA183 | 0.036061 | 0.089945 | 0.35557 | -1.4918 | 0.041017 | 0.12837 | 0.32284 | -1.6311 | 0.025122 | 0.053447 | 0.31915 | -1.6477 |
| LPE204 | 0.20631 | 0.31527 | 1.2576 | 0.33073 | 0.88246 | 0.91757 | 1.0148 | 0.021233 | 0.53305 | 0.60667 | 0.88979 | -0.16846 |
| PI180/224 | 0.19662 | 0.30549 | 0.82079 | -0.28492 | 0.81738 | 0.88727 | 0.98161 | -0.02677 | 0.1648 | 0.23425 | 0.8134 | -0.29797 |
| TAG562-FA180 | 0.018226 | 0.063783 | 0.21327 | -2.2293 | 0.11595 | 0.22478 | 0.3447 | -1.5366 | 0.004379 | 0.019527 | 0.14469 | -2.7889 |
| TAG521-FA161 | 0.017637 | 0.063783 | 0.24191 | -2.0474 | 0.023343 | 0.1178 | 0.25886 | -1.9497 | 0.002124 | 0.015539 | 0.13822 | -2.855 |
| PS181/182_2 | 0.42611 | 0.55369 | 0.84708 | -0.23942 | 0.1979 | 0.34083 | 1.5384 | 0.62146 | 0.011737 | 0.031704 | 0.53674 | -0.89769 |
| TAG503-FA183 | 0.026374 | 0.076523 | 0.28666 | -1.8026 | 0.013885 | 0.11428 | 0.21474 | -2.2194 | 0.007389 | 0.024452 | 0.21119 | -2.2434 |
| PE170/181 | 0.15285 | 0.25256 | 0.60272 | -0.73045 | 0.053042 | 0.14281 | 0.50595 | -0.98294 | 0.000567 | 0.009864 | 0.23029 | -2.1185 |
| TAG569-FA205 | 0.10481 | 0.19495 | 0.47124 | -1.0855 | 0.050253 | 0.14161 | 0.33278 | -1.5873 | 0.22119 | 0.29568 | 0.66701 | -0.58422 |
| TAG563-FA181 | 0.010245 | 0.049771 | 0.2441 | -2.0344 | 0.73405 | 0.82963 | 0.88774 | -0.1718 | 0.004665 | 0.019874 | 0.19214 | -2.3798 |
| PE181/183 | 0.30295 | 0.42782 | 0.73528 | -0.44363 | 0.021282 | 0.11545 | 0.47307 | -1.0799 | 0.003725 | 0.018938 | 0.29288 | -1.7716 |
| TAG481-FA160 | 0.046212 | 0.109 | 0.28854 | -1.7932 | 0.049527 | 0.14066 | 0.29199 | -1.776 | 0.031662 | 0.063227 | 0.26302 | -1.9267 |
| TAG542-FA160 | 0.017333 | 0.063037 | 0.2729 | -1.8736 | 0.039554 | 0.12837 | 0.31392 | -1.6715 | 0.004606 | 0.019874 | 0.20206 | -2.3071 |
| TAG546-FA203 | 0.002255 | 0.02531 | 0.23877 | -2.0663 | 0.012849 | 0.11428 | 0.28205 | -1.826 | 0.002084 | 0.015416 | 0.20045 | -2.3187 |
| TAG567-FA224 | 0.002358 | 0.02563 | 0.26199 | -1.9324 | 0.010677 | 0.11395 | 0.2785 | -1.8443 | 0.020457 | 0.045765 | 0.3367 | -1.5704 |
| TAG501-FA180 | 0.035796 | 0.089627 | 0.24699 | -2.0175 | 0.0373 | 0.1282 | 0.24642 | -2.0208 | 0.0074 | 0.024452 | 0.15796 | -2.6624 |

| | | | | | | | | | | | | |
|---|---|---|---|---|---|---|---|---|---|---|---|---|
| PC181/161 | 0.00378 | 0.031148 | 0.53835 | -0.89339 | 0.21163 | 0.35878 | 0.81429 | -0.29638 | 0.021384 | 0.047512 | 0.55646 | -0.84566 |
| TAG543-FA180 | 0.051696 | 0.11726 | 0.33423 | -1.5811 | 0.092522 | 0.1943 | 0.3789 | -1.4001 | 0.009261 | 0.027527 | 0.2326 | -2.1041 |
| PC180/204 | 0.61073 | 0.7242 | 1.0763 | 0.1061 | 0.67418 | 0.78281 | 0.92058 | -0.11939 | 0.18464 | 0.25961 | 1.2442 | 0.31527 |
| TAG544-FA181 | 0.013821 | 0.055199 | 0.2723 | -1.8768 | 0.28271 | 0.44562 | 0.57625 | -0.79523 | 0.007377 | 0.024452 | 0.22846 | -2.13 |
| TAG523-FA200 | 0.006752 | 0.039785 | 0.34841 | -1.5211 | 0.035355 | 0.12787 | 0.41529 | -1.2678 | 0.004833 | 0.019885 | 0.32913 | -1.6033 |
| PI180/224_2 | 0.22351 | 0.33682 | 0.80876 | -0.30622 | 0.45335 | 0.62528 | 0.88197 | -0.1812 | 0.052833 | 0.093973 | 0.70932 | -0.49549 |
| LPC204 | 0.73806 | 0.82841 | 1.1481 | 0.19931 | 0.75871 | 0.84287 | 1.0808 | 0.11211 | 0.41726 | 0.49478 | 1.3129 | 0.39272 |
| TAG522-FA161 | 0.007504 | 0.042113 | 0.23907 | -2.0645 | 0.038408 | 0.12835 | 0.33572 | -1.5747 | 0.007559 | 0.024609 | 0.21781 | -2.1989 |
| TAG543-FA160 | 0.029804 | 0.081522 | 0.32913 | -1.6033 | 0.065313 | 0.16251 | 0.37231 | -1.4254 | 0.025713 | 0.053932 | 0.31485 | -1.6672 |
| TAG544-FA221 | 0.008548 | 0.045241 | 0.21104 | -2.2444 | 0.2203 | 0.36868 | 0.50111 | -0.99679 | 0.005485 | 0.020761 | 0.1489 | -2.7476 |
| TAG542-FA182 | 0.040717 | 0.098174 | 0.26626 | -1.9091 | 0.053445 | 0.14281 | 0.27888 | -1.8423 | 0.003613 | 0.018938 | 0.14577 | -2.7782 |
| PC170/181 | 0.012026 | 0.053913 | 0.62002 | -0.68962 | 0.052838 | 0.14281 | 0.70907 | -0.496 | 0.002995 | 0.01784 | 0.5296 | -0.91703 |
| TAG567-FA161 | 0.002362 | 0.02563 | 0.34649 | -1.5291 | 0.058535 | 0.15121 | 0.47575 | -1.0717 | 0.007561 | 0.024609 | 0.31549 | -1.6643 |
| PE180/225 | 0.000663 | 0.018074 | 1.4193 | 0.50518 | 0.20644 | 0.35367 | 0.83469 | -0.26069 | 0.45118 | 0.5302 | 0.92907 | -0.10614 |
| PG182/182 | 0.080192 | 0.15898 | 0.49148 | -1.0248 | 0.028115 | 0.12263 | 0.41301 | -1.2757 | 0.001292 | 0.013002 | 0.23714 | -2.0762 |
| PE181/225 | 0.026802 | 0.077203 | 0.7487 | -0.41754 | 0.003498 | 0.11015 | 0.64928 | -0.62309 | 0.00553 | 0.02081 | 0.58879 | -0.76418 |
| TAG543-FA182 | 0.019653 | 0.065823 | 0.2925 | -1.7735 | 0.10931 | 0.21696 | 0.42031 | -1.2505 | 0.008706 | 0.026949 | 0.24643 | -2.0208 |
| PS180/182 | 0.044146 | 0.10527 | 0.73988 | -0.43463 | 0.88024 | 0.91757 | 1.0131 | 0.018768 | 0.000603 | 0.010024 | 0.52074 | -0.94137 |
| SMd181/202 | 0.06104 | 0.13121 | 1.2179 | 0.28437 | 0.18585 | 0.32612 | 0.90971 | -0.13651 | 0.21675 | 0.29094 | 1.2591 | 0.33241 |
| PE160/225 | 0.003517 | 0.03053 | 1.4434 | 0.52943 | 0.079787 | 0.17911 | 0.74282 | -0.42891 | 0.010173 | 0.029199 | 0.762 | -0.39214 |
| TAG480-FA140 | 0.11895 | 0.21333 | 0.28281 | -1.8221 | 0.070893 | 0.16843 | 0.22915 | -2.1256 | 0.045436 | 0.08332 | 0.20132 | -2.3124 |
| PC140/140 | 0.022743 | 0.071165 | 0.57561 | -0.79683 | 0.62216 | 0.74289 | 0.97743 | -0.03293 | 0.004327 | 0.019527 | 0.38717 | -1.3689 |
| TAG482-FA181 | 0.038126 | 0.094015 | 0.33489 | -1.5782 | 0.97675 | 0.98857 | 1.2623 | 0.33602 | 0.014182 | 0.035784 | 0.24442 | -2.0325 |
| TAG564-FA202 | 0.000874 | 0.018074 | 0.26403 | -1.9212 | 0.013021 | 0.11428 | 0.37859 | -1.4013 | 0.001357 | 0.013389 | 0.23136 | -2.1118 |
| TAG543-FA181 | 0.021675 | 0.069854 | 0.31279 | -1.6767 | 0.60705 | 0.73675 | 0.74191 | -0.43069 | 0.023786 | 0.051617 | 0.30704 | -1.7035 |
| TAG564-FA181 | 0.015091 | 0.057788 | 0.24695 | -2.0177 | 0.21822 | 0.36709 | 0.48064 | -1.057 | 0.01105 | 0.030873 | 0.21732 | -2.2021 |
| PE140/182 | 0.005711 | 0.03721 | 0.55579 | -0.84739 | 0.17634 | 0.318 | 0.79347 | -0.33376 | 0.002016 | 0.015416 | 0.3732 | -1.422 |
| TAG543-FA161 | 0.010929 | 0.051187 | 0.25188 | -1.9892 | 0.038803 | 0.12835 | 0.32964 | -1.601 | 0.003586 | 0.018938 | 0.18406 | -2.4417 |
| TAG504-FA181 | 0.004087 | 0.032816 | 0.23008 | -2.1198 | 0.017216 | 0.11428 | 0.288 | -1.7959 | 0.006201 | 0.021941 | 0.2203 | -2.1825 |
| TAG568-FA181 | 0.045795 | 0.10841 | 0.42319 | -1.2406 | 0.09202 | 0.1943 | 0.46998 | -1.0893 | 0.054729 | 0.096816 | 0.39887 | -1.326 |
| PEP-180/202 | 0.02765 | 0.078602 | 0.66755 | -0.58305 | 0.23423 | 0.38604 | 0.80347 | -0.31569 | 0.000141 | 0.007659 | 0.41398 | -1.2724 |
| PE160/203 | 0.98244 | 0.98883 | 1.0072 | 0.01033 | 0.057133 | 0.14877 | 0.66829 | -0.58146 | 0.002648 | 0.016986 | 0.48303 | -1.0498 |
| TAG565-FA181 | 0.005409 | 0.035933 | 0.28323 | -1.8199 | 0.028782 | 0.12307 | 0.3574 | -1.4844 | 0.007702 | 0.02474 | 0.28745 | -1.7986 |

| | | | | | | | | | | | | |
|---|---|---|---|---|---|---|---|---|---|---|---|---|
| TAG564-FA182 | 0.016272 | 0.06088 | 0.24556 | -2.0258 | 0.18787 | 0.32877 | 0.46262 | -1.1121 | 0.00334 | 0.018829 | 0.16437 | -2.605 |
| TAG566-FA203 | 0.004424 | 0.033873 | 0.31489 | -1.6671 | 0.008603 | 0.11015 | 0.32761 | -1.61 | 0.003014 | 0.01784 | 0.26392 | -1.9218 |
| PEP-160/160 | 0.1413 | 0.23831 | 1.2943 | 0.37213 | 0.61166 | 0.73687 | 1.1049 | 0.14398 | 0.41657 | 0.49478 | 1.1637 | 0.21871 |
| TAG522-FA160 | 0.021958 | 0.069893 | 0.34253 | -1.5457 | 0.097531 | 0.20259 | 0.45401 | -1.1392 | 0.024445 | 0.052348 | 0.34841 | -1.5211 |
| LPE224 | 0.78816 | 0.86965 | 1.0602 | 0.084289 | 0.34739 | 0.51165 | 0.74106 | -0.43233 | 0.19323 | 0.26652 | 0.65833 | -0.60313 |
| PI161/180_2 | 0.66323 | 0.76614 | 0.93661 | -0.09448 | 0.7429 | 0.83673 | 0.95524 | -0.06607 | 0.058907 | 0.10199 | 0.68218 | -0.55178 |
| PEO-160/205 | 0.19386 | 0.30262 | 0.85216 | -0.23081 | 0.059617 | 0.15256 | 0.77361 | -0.37031 | 0.006811 | 0.023335 | 0.60703 | -0.72015 |
| TAG568-FA204 | 0.12566 | 0.2205 | 0.52156 | -0.93909 | 0.033823 | 0.12787 | 0.37731 | -1.4062 | 0.18901 | 0.26292 | 0.55809 | -0.84144 |
| PC150/181 | 0.004973 | 0.034808 | 0.53559 | -0.90079 | 0.073204 | 0.17078 | 0.66893 | -0.58007 | 0.001204 | 0.012765 | 0.41902 | -1.2549 |
| TAG546-FA161 | 0.001074 | 0.019426 | 0.26719 | -1.904 | 0.008304 | 0.11015 | 0.29746 | -1.7493 | 0.009746 | 0.02858 | 0.34421 | -1.5386 |
| TAG522-FA182 | 0.026995 | 0.077418 | 0.30886 | -1.695 | 0.024026 | 0.11849 | 0.2876 | -1.7979 | 0.022854 | 0.049925 | 0.29567 | -1.7579 |
| LPC140 | 0.035029 | 0.088847 | 0.71268 | -0.48868 | 0.88019 | 0.91757 | 1.0181 | 0.025836 | 0.015868 | 0.039244 | 0.54379 | -0.87889 |
| TAG501-FA140 | 0.028731 | 0.079931 | 0.25836 | -1.9526 | 0.066122 | 0.16251 | 0.31528 | -1.6653 | 0.003911 | 0.018938 | 0.15113 | -2.7261 |
| PEP-180/183 | 0.019443 | 0.065582 | 0.62421 | -0.67989 | 0.59728 | 0.73128 | 0.90064 | -0.15097 | 7.73E-05 | 0.007659 | 0.28693 | -1.8012 |
| TAG543-FA202 | 0.021232 | 0.069457 | 0.33445 | -1.5802 | 0.053528 | 0.14281 | 0.3609 | -1.4703 | 0.012887 | 0.033829 | 0.29024 | -1.7847 |
| PC180/140 | 0.016243 | 0.06088 | 0.62366 | -0.68118 | 0.43048 | 0.60138 | 0.89139 | -0.16587 | 0.012061 | 0.031918 | 0.5801 | -0.78563 |
| TAG568-FA225 | 0.034703 | 0.088847 | 0.39372 | -1.3448 | 0.08155 | 0.17949 | 0.42932 | -1.2199 | 0.08168 | 0.12938 | 0.44621 | -1.1642 |
| SMd181/161 | 0.43445 | 0.56006 | 1.0826 | 0.1145 | 0.31305 | 0.47839 | 0.92354 | -0.11475 | 0.00796 | 0.025402 | 1.3645 | 0.44834 |
| PEP-180/205 | 0.37549 | 0.504 | 0.84104 | -0.24975 | 0.23349 | 0.38579 | 0.78487 | -0.34947 | 0.004316 | 0.019527 | 0.48923 | -1.0314 |
| PEP-160/182 | 0.009617 | 0.048275 | 0.63255 | -0.66074 | 0.86353 | 0.91538 | 0.98585 | -0.02056 | 0.00451 | 0.019874 | 0.54039 | -0.88792 |
| SMd181/200 | 0.48859 | 0.61256 | 1.0567 | 0.079558 | 0.094245 | 0.19665 | 0.84836 | -0.23725 | 0.12671 | 0.18705 | 1.1465 | 0.19727 |
| TAG541-FA181 | 0.018328 | 0.063783 | 0.21691 | -2.2049 | 0.11129 | 0.21955 | 0.34708 | -1.5266 | 0.006038 | 0.021904 | 0.16449 | -2.6039 |
| TAG565-FA203 | 0.006725 | 0.039785 | 0.33294 | -1.5866 | 0.042827 | 0.12968 | 0.436 | -1.1976 | 0.008977 | 0.027307 | 0.3228 | -1.6313 |
| PG160/160 | 0.46443 | 0.59167 | 0.74841 | -0.4181 | 0.19212 | 0.33441 | 0.5943 | -0.75073 | 0.67095 | 0.73134 | 0.79011 | -0.33988 |
| PEP-160/225 | 0.26372 | 0.38237 | 1.1521 | 0.20423 | 0.84163 | 0.90484 | 0.97605 | -0.03497 | 0.15567 | 0.22272 | 1.3406 | 0.42289 |
| PC170/226 | 0.088162 | 0.17132 | 0.66986 | -0.57807 | 0.08278 | 0.18034 | 0.66409 | -0.59055 | 0.1761 | 0.24868 | 0.73336 | -0.44741 |
| TAG522-FA180 | 0.062561 | 0.13227 | 0.30185 | -1.7281 | 0.030811 | 0.12449 | 0.24091 | -2.0535 | 0.016001 | 0.039244 | 0.21442 | -2.2215 |
| TAG545-FA203 | 0.017145 | 0.062716 | 0.36805 | -1.442 | 0.030149 | 0.12449 | 0.35959 | -1.4756 | 0.016843 | 0.041066 | 0.34797 | -1.523 |
| PC181/182 | 0.006906 | 0.039785 | 0.65049 | -0.6204 | 0.7355 | 0.82983 | 0.97326 | -0.0391 | 0.062597 | 0.10585 | 0.67057 | -0.57653 |
| TAG548-FA205 | 0.018714 | 0.063783 | 0.35326 | -1.5012 | 0.023932 | 0.11849 | 0.33712 | -1.5687 | 0.047702 | 0.086985 | 0.45556 | -1.1343 |
| TAG562-FA160 | 0.098844 | 0.18491 | 0.19071 | -2.3905 | 0.25409 | 0.4096 | 0.32894 | -1.6041 | 0.02349 | 0.051143 | 0.12017 | -3.0568 |
| PG160/180_2 | 0.86041 | 0.91975 | 0.89788 | -0.15541 | 0.031394 | 0.12538 | 0.46658 | -1.0998 | 0.028667 | 0.05851 | 0.47746 | -1.0665 |
| PEP-160/205 | 0.3825 | 0.51131 | 0.85403 | -0.22764 | 0.30473 | 0.47009 | 0.82483 | -0.27784 | 0.071934 | 0.11825 | 0.67936 | -0.55775 |

| | | | | | | | | | | | | |
|---|---|---|---|---|---|---|---|---|---|---|---|---|
| TAG543-FA201 | 0.022175 | 0.070077 | 0.29037 | -1.7841 | 0.07397 | 0.17078 | 0.36823 | -1.4413 | 0.018769 | 0.043328 | 0.27883 | -1.8425 |
| PEP-181/182 | 0.013737 | 0.055199 | 0.68054 | -0.55524 | 0.80561 | 0.88143 | 0.98946 | -0.01528 | 0.003092 | 0.01797 | 0.56414 | -0.82589 |
| PG160/180 | 0.88874 | 0.9362 | 0.87331 | -0.19544 | 0.023092 | 0.11744 | 0.35953 | -1.4758 | 0.018032 | 0.042533 | 0.35492 | -1.4944 |
| PEO-160/182 | 0.006905 | 0.039785 | 0.5967 | -0.74493 | 0.17988 | 0.32029 | 0.78567 | -0.348 | 0.000357 | 0.008616 | 0.41492 | -1.2691 |
| TAG483-FA160 | 0.019717 | 0.065823 | 0.27009 | -1.8885 | 0.012383 | 0.11428 | 0.22855 | -2.1294 | 0.019671 | 0.044159 | 0.27177 | -1.8795 |
| TAG547-FA161 | 0.024039 | 0.073128 | 0.36715 | -1.4456 | 0.035841 | 0.12787 | 0.39131 | -1.3536 | 0.06364 | 0.10677 | 0.43794 | -1.1912 |
| TAG565-FA224 | 0.071359 | 0.14427 | 0.36072 | -1.471 | 0.042635 | 0.12968 | 0.30329 | -1.7212 | 0.10268 | 0.15915 | 0.39071 | -1.3558 |
| TAG502-FA160 | 0.037201 | 0.092433 | 0.37111 | -1.4301 | 0.022784 | 0.11722 | 0.3038 | -1.7188 | 0.081487 | 0.12938 | 0.42423 | -1.2371 |
| PE180/150 | 0.66564 | 0.76614 | 0.91506 | -0.12806 | 0.47327 | 0.64187 | 0.86629 | -0.20707 | 0.2114 | 0.28553 | 0.78936 | -0.34124 |
| LPE180 | 0.011297 | 0.052531 | 1.4208 | 0.5067 | 0.33306 | 0.49843 | 0.85545 | -0.22524 | 0.057478 | 0.10086 | 0.70453 | -0.50526 |
| TAG502-FA182 | 0.03556 | 0.089381 | 0.3659 | -1.4505 | 0.023805 | 0.11849 | 0.29771 | -1.748 | 0.1641 | 0.23376 | 0.4899 | -1.0295 |
| PEO-180/224 | 0.89117 | 0.93724 | 1.0325 | 0.046203 | 0.53564 | 0.68642 | 0.83962 | -0.2522 | 0.96741 | 0.97189 | 0.94 | -0.08927 |
| TAG483-FA181 | 0.075467 | 0.15163 | 0.46394 | -1.108 | 0.78545 | 0.86516 | 1.8545 | 0.89106 | 0.058404 | 0.10166 | 0.43093 | -1.2145 |
| PE182/183 | 0.13933 | 0.2362 | 0.50318 | -0.99086 | 0.0015 | 0.11015 | 0.19315 | -2.3722 | 0.000217 | 0.008255 | 0.1113 | -3.1675 |
| PG160/181 | 0.8544 | 0.91784 | 0.93249 | -0.10083 | 0.04429 | 0.13226 | 0.58926 | -0.76303 | 0.030041 | 0.060547 | 0.59236 | -0.75545 |
| PEP-180/180 | 0.12004 | 0.21469 | 0.71967 | -0.4746 | 0.79031 | 0.86761 | 1.078 | 0.10839 | 0.0025 | 0.016786 | 0.48786 | -1.0355 |
| PS160/180 | 0.023723 | 0.072847 | 1.3827 | 0.46751 | 0.84685 | 0.90526 | 1.0098 | 0.014058 | 0.9469 | 0.95571 | 1.026 | 0.037 |
| PG180/181_2 | 0.7677 | 0.85431 | 0.92315 | -0.11536 | 0.041961 | 0.12885 | 0.57419 | -0.8004 | 0.048006 | 0.087295 | 0.5989 | -0.7396 |
| PC160/160 | 0.068967 | 0.14074 | 1.1896 | 0.25053 | 0.869 | 0.9154 | 0.98561 | -0.02091 | 0.10325 | 0.15935 | 1.2599 | 0.33328 |
| TAG562-FA201 | 0.03049 | 0.082447 | 0.2516 | -1.9908 | 0.34342 | 0.50811 | 0.51718 | -0.95128 | 0.004752 | 0.019874 | 0.13482 | -2.8909 |
| PEO-160/224 | 0.86816 | 0.92198 | 1.0591 | 0.082884 | 0.73064 | 0.82729 | 0.86863 | -0.20319 | 0.79508 | 0.84162 | 0.8768 | -0.18969 |
| PE170/204 | 0.092007 | 0.17565 | 0.67149 | -0.57456 | 0.006625 | 0.11015 | 0.46997 | -1.0893 | 0.000561 | 0.009864 | 0.33645 | -1.5715 |
| TAG566-FA224 | 0.07398 | 0.1491 | 0.41451 | -1.2705 | 0.020182 | 0.11428 | 0.28389 | -1.8166 | 0.070245 | 0.11606 | 0.40206 | -1.3145 |
| PE182/182 | 0.06258 | 0.13227 | 0.52436 | -0.93138 | 0.041399 | 0.12885 | 0.50498 | -0.9857 | 0.000165 | 0.007659 | 0.17098 | -2.5481 |
| PC181/225 | 0.084771 | 0.16523 | 0.78606 | -0.34728 | 0.067079 | 0.16355 | 0.76001 | -0.39591 | 0.20061 | 0.27494 | 0.83486 | -0.26039 |
| PI161/180 | 0.43657 | 0.56168 | 0.86748 | -0.2051 | 0.84369 | 0.90484 | 1.0708 | 0.09868 | 0.11409 | 0.17273 | 0.73009 | -0.45386 |
| LPE225 | 0.000671 | 0.018074 | 1.492 | 0.57727 | 0.5043 | 0.66694 | 1.1586 | 0.21242 | 0.81495 | 0.8557 | 1.0487 | 0.068586 |
| HCER200 | 0.13488 | 0.23062 | 1.2584 | 0.33154 | 0.93798 | 0.96161 | 0.98442 | -0.02266 | 0.84417 | 0.87789 | 1.0271 | 0.038507 |
| LPC203 | 0.34514 | 0.47325 | 0.78069 | -0.35719 | 0.54165 | 0.69141 | 0.84074 | -0.25026 | 0.2415 | 0.31507 | 0.71205 | -0.48995 |
| TAG544-FA160 | 0.032434 | 0.085485 | 0.39115 | -1.3542 | 0.035818 | 0.12787 | 0.35775 | -1.483 | 0.019481 | 0.044159 | 0.34221 | -1.5471 |
| PE180/203 | 0.85974 | 0.91975 | 0.96826 | -0.04653 | 0.04703 | 0.13653 | 0.65982 | -0.59986 | 0.037254 | 0.07133 | 0.68032 | -0.55572 |
| TAG546-FA204 | 0.11426 | 0.20777 | 0.50051 | -0.99854 | 0.046234 | 0.13557 | 0.39263 | -1.3488 | 0.27903 | 0.35341 | 0.60337 | -0.72888 |
| TAG568-FA161 | 0.061042 | 0.13121 | 0.39383 | -1.3444 | 0.14375 | 0.26662 | 0.47243 | -1.0818 | 0.047228 | 0.086363 | 0.35477 | -1.4951 |

| | | | | | | | | | | | | |
|---|---|---|---|---|---|---|---|---|---|---|---|---|
| PEO-180/182 | 0.010782 | 0.051143 | 0.5837 | -0.7767 | 0.13588 | 0.25419 | 0.73713 | -0.44001 | 5.74E-05 | 0.007659 | 0.31717 | -1.6567 |
| PS180/180 | 0.004808 | 0.034394 | 1.5694 | 0.65023 | 0.82049 | 0.88727 | 1.0153 | 0.021878 | 0.52841 | 0.60244 | 1.1509 | 0.20272 |
| TAG561-FA160 | 0.18278 | 0.28812 | 0.20708 | -2.2717 | 0.68006 | 0.78635 | 0.5999 | -0.73722 | 0.025651 | 0.053932 | 0.096164 | -3.3784 |
| PEP-160/203 | 0.34575 | 0.47325 | 0.85033 | -0.23391 | 0.7442 | 0.83674 | 0.95061 | -0.07307 | 0.40092 | 0.47977 | 0.866 | -0.20756 |
| TAG563-FA201 | 0.030849 | 0.082987 | 0.26044 | -1.941 | 0.64856 | 0.76349 | 0.80267 | -0.31713 | 0.015926 | 0.039244 | 0.21988 | -2.1852 |
| PE181/224 | 0.083887 | 0.16499 | 0.68764 | -0.54028 | 0.011924 | 0.11428 | 0.56175 | -0.832 | 0.044151 | 0.08212 | 0.55314 | -0.85429 |
| TAG521-FA180 | 0.063722 | 0.13425 | 0.24581 | -2.0244 | 0.051744 | 0.14281 | 0.2239 | -2.1591 | 0.024889 | 0.053124 | 0.18357 | -2.4456 |
| TAG547-FA205 | 0.08459 | 0.16523 | 0.50266 | -0.99235 | 0.079246 | 0.17851 | 0.4219 | -1.245 | 0.27629 | 0.35062 | 0.79953 | -0.32278 |
| PG160/181_2 | 0.92246 | 0.95532 | 0.98298 | -0.02476 | 0.053138 | 0.14281 | 0.61506 | -0.70119 | 0.02536 | 0.053601 | 0.59541 | -0.74805 |
| PE170/182 | 0.038483 | 0.094538 | 0.55775 | -0.84231 | 0.004467 | 0.11015 | 0.40848 | -1.2917 | 7.59E-05 | 0.007659 | 0.20708 | -2.2717 |
| PC200/203 | 0.2516 | 0.37068 | 0.77706 | -0.36391 | 0.004862 | 0.11015 | 0.51996 | -0.94353 | 0.00519 | 0.020577 | 0.53596 | -0.89981 |
| TAG547-FA204 | 0.024987 | 0.074569 | 0.40373 | -1.3085 | 0.021273 | 0.11545 | 0.36069 | -1.4712 | 0.030949 | 0.062104 | 0.39955 | -1.3236 |
| TAG526-FA204 | 0.062473 | 0.13227 | 0.37698 | -1.4074 | 0.065952 | 0.16251 | 0.37579 | -1.412 | 0.065529 | 0.1091 | 0.38356 | -1.3825 |
| TAG542-FA181 | 0.047717 | 0.11094 | 0.31544 | -1.6646 | 0.40216 | 0.5754 | 0.57164 | -0.80683 | 0.025765 | 0.053932 | 0.2662 | -1.9094 |
| PI180/183 | 0.075855 | 0.15194 | 0.74063 | -0.43318 | 0.92759 | 0.95246 | 1.0495 | 0.069682 | 0.013734 | 0.035061 | 0.59298 | -0.75394 |
| TAG520-FA160 | 0.077458 | 0.15421 | 0.17481 | -2.5161 | 0.04166 | 0.12885 | 0.14136 | -2.8226 | 0.021018 | 0.046859 | 0.11461 | -3.1252 |
| TAG521-FA181 | 0.045673 | 0.10841 | 0.26806 | -1.8994 | 0.081519 | 0.17949 | 0.3072 | -1.7028 | 0.045136 | 0.08332 | 0.25979 | -1.9446 |
| TAG544-FA203 | 0.016832 | 0.062615 | 0.35663 | -1.4875 | 0.040961 | 0.12837 | 0.38083 | -1.3928 | 0.013572 | 0.035002 | 0.32347 | -1.6283 |
| PEO-180/225 | 0.088649 | 0.17176 | 1.2812 | 0.35751 | 0.85907 | 0.91233 | 1.0244 | 0.034757 | 0.074825 | 0.12087 | 1.5424 | 0.62514 |
| TAG546-FA225 | 0.026448 | 0.076523 | 0.31187 | -1.681 | 0.050468 | 0.14161 | 0.33179 | -1.5917 | 0.073327 | 0.11934 | 0.36562 | -1.4516 |
| TAG542-FA202 | 0.066375 | 0.13805 | 0.31502 | -1.6665 | 0.068776 | 0.1667 | 0.30611 | -1.7079 | 0.028162 | 0.058016 | 0.2508 | -1.9954 |
| TAG564-FA203 | 0.008995 | 0.046514 | 0.39307 | -1.3471 | 0.040405 | 0.12837 | 0.46101 | -1.1171 | 0.011944 | 0.031843 | 0.36994 | -1.4346 |
| PC160/202 | 0.37077 | 0.49871 | 0.84938 | -0.23552 | 0.18412 | 0.32395 | 0.77164 | -0.374 | 0.13672 | 0.20001 | 0.7713 | -0.37463 |
| TAG563-FA180 | 0.017148 | 0.062716 | 0.26788 | -1.9004 | 0.036778 | 0.1282 | 0.30778 | -1.7 | 0.004379 | 0.019527 | 0.19397 | -2.3661 |
| PE180/160 | 0.25608 | 0.37547 | 1.1945 | 0.25639 | 0.17831 | 0.32029 | 0.76064 | -0.39472 | 0.007332 | 0.024452 | 0.57354 | -0.80203 |
| FFA205 | 0.4031 | 0.53228 | 1.2426 | 0.31332 | 0.78069 | 0.86141 | 0.97969 | -0.0296 | 0.50319 | 0.57978 | 0.92953 | -0.10543 |
| PE180/201 | 0.57014 | 0.68479 | 0.86598 | -0.2076 | 0.11601 | 0.22478 | 0.69599 | -0.52286 | 0.006056 | 0.021904 | 0.44553 | -1.1664 |
| PC180/203 | 0.34603 | 0.47325 | 0.81841 | -0.28911 | 0.091109 | 0.19389 | 0.69397 | -0.52705 | 0.28787 | 0.36108 | 0.80193 | -0.31845 |
| PG180/180 | 0.695 | 0.79237 | 0.89957 | -0.15269 | 0.035917 | 0.12787 | 0.56293 | -0.82897 | 0.050801 | 0.091358 | 0.61135 | -0.70993 |
| PEP-180/204 | 0.73435 | 0.82758 | 0.96422 | -0.05257 | 0.98335 | 0.98981 | 1.0203 | 0.028945 | 0.45794 | 0.53523 | 0.90723 | -0.14046 |
| PEO-180/160 | 0.71005 | 0.8067 | 1.0296 | 0.042034 | 0.12397 | 0.23667 | 0.84984 | -0.23474 | 0.013126 | 0.034318 | 0.745 | -0.42469 |
| TAG541-FA201 | 0.034491 | 0.088751 | 0.17639 | -2.5031 | 0.1024 | 0.20688 | 0.24623 | -2.0219 | 0.028671 | 0.05851 | 0.16564 | -2.5939 |
| PC180/161 | 0.067978 | 0.14004 | 0.63614 | -0.65259 | 0.13862 | 0.25858 | 0.71012 | -0.49386 | 0.019406 | 0.044159 | 0.53123 | -0.91259 |

| | | | | | | | | | | | | |
|---|---|---|---|---|---|---|---|---|---|---|---|---|
| TAG568-FA205 | 0.12373 | 0.21888 | 0.54536 | -0.87471 | 0.036142 | 0.12787 | 0.38914 | -1.3616 | 0.13389 | 0.19675 | 0.56455 | -0.82483 |
| PI180/204 | 0.51573 | 0.63467 | 0.93003 | -0.10466 | 0.67678 | 0.78395 | 1.0544 | 0.076405 | 0.85825 | 0.8911 | 1.0291 | 0.041429 |
| PI180/204_2 | 0.50905 | 0.63003 | 0.92451 | -0.11323 | 0.50933 | 0.66694 | 1.1007 | 0.13841 | 0.59373 | 0.65847 | 1.083 | 0.11503 |
| TAG546-FA205 | 0.25676 | 0.37551 | 0.59825 | -0.74118 | 0.112 | 0.22027 | 0.47886 | -1.0623 | 0.53537 | 0.60825 | 1.0241 | 0.034383 |
| PC170/182 | 0.00095 | 0.018192 | 0.61748 | -0.69554 | 0.005505 | 0.11015 | 0.68102 | -0.55424 | 0.000299 | 0.008616 | 0.55458 | -0.85054 |
| TAG542-FA201 | 0.040241 | 0.09775 | 0.30458 | -1.7151 | 0.20747 | 0.3545 | 0.45803 | -1.1265 | 0.035555 | 0.068887 | 0.29514 | -1.7606 |
| TAG521-FA160 | 0.093405 | 0.17676 | 0.28133 | -1.8297 | 0.098652 | 0.20259 | 0.2812 | -1.8303 | 0.036274 | 0.069754 | 0.21273 | -2.2329 |
| TAG482-FA120 | 0.33581 | 0.46284 | 0.61855 | -0.69303 | 0.56026 | 0.70696 | 2.7737 | 1.4718 | 0.024361 | 0.052341 | 0.30801 | -1.699 |
| TAG564-FA180 | 0.01791 | 0.063783 | 0.34361 | -1.5412 | 0.035262 | 0.12787 | 0.36097 | -1.47 | 0.017045 | 0.041249 | 0.31522 | -1.6656 |
| LPC180 | 0.42532 | 0.55369 | 0.90553 | -0.14316 | 0.22526 | 0.37601 | 0.85678 | -0.22301 | 0.6238 | 0.68713 | 0.93805 | -0.09226 |
| PC181/205 | 0.1755 | 0.28003 | 0.69839 | -0.5179 | 0.19604 | 0.34033 | 0.72504 | -0.46386 | 0.33466 | 0.40952 | 0.82062 | -0.28521 |
| TAG567-FA205 | 0.10204 | 0.19033 | 0.53511 | -0.90209 | 0.12143 | 0.2325 | 0.52288 | -0.93545 | 0.22558 | 0.2997 | 0.68971 | -0.53594 |
| PE181/201 | 0.54757 | 0.66381 | 0.84314 | -0.24616 | 0.58807 | 0.72734 | 1.1796 | 0.23833 | 0.018176 | 0.042564 | 0.4628 | -1.1115 |
| TAG501-FA161 | 0.020331 | 0.067383 | 0.23727 | -2.0754 | 0.017842 | 0.11428 | 0.22774 | -2.1346 | 0.012221 | 0.032209 | 0.20865 | -2.2609 |
| PG180/181 | 0.8313 | 0.90214 | 0.9381 | -0.09219 | 0.040073 | 0.12837 | 0.56434 | -0.82537 | 0.054069 | 0.095909 | 0.60472 | -0.72567 |
| PI180/205_2 | 0.79967 | 0.87817 | 1.0546 | 0.076727 | 0.98701 | 0.99005 | 1.0225 | 0.032103 | 0.64927 | 0.70919 | 0.99874 | -0.00182 |
| TAG541-FA180 | 0.066849 | 0.13859 | 0.20868 | -2.2606 | 0.090883 | 0.19389 | 0.22631 | -2.1436 | 0.01398 | 0.035551 | 0.1261 | -2.9874 |
| PC200/205 | 0.28744 | 0.41125 | 0.74105 | -0.43236 | 0.056744 | 0.14841 | 0.54532 | -0.87484 | 0.11644 | 0.17465 | 0.60787 | -0.71816 |
| PC150/226 | 0.027886 | 0.078928 | 0.58255 | -0.77955 | 0.057961 | 0.15033 | 0.5984 | -0.74081 | 0.01372 | 0.035061 | 0.54581 | -0.87352 |
| PC200/181 | 0.047946 | 0.11108 | 0.7132 | -0.48763 | 0.69266 | 0.79527 | 0.96766 | -0.04743 | 0.001199 | 0.012765 | 0.45717 | -1.1292 |
| TAG527-FA160 | 0.051522 | 0.11726 | 0.1648 | -2.6012 | 0.039574 | 0.12837 | 0.15064 | -2.7308 | 0.022829 | 0.049925 | 0.12823 | -2.9632 |
| LPC181 | 0.2192 | 0.33108 | 0.79977 | -0.32235 | 0.71492 | 0.81652 | 1.0943 | 0.13007 | 0.19147 | 0.26521 | 0.75119 | -0.41274 |
| PE181/181 | 0.53633 | 0.65384 | 0.76846 | -0.37996 | 0.38278 | 0.55498 | 1.836 | 0.8766 | 0.087698 | 0.1379 | 0.52605 | -0.92674 |
| SMd181/120 | 0.22585 | 0.33956 | 0.69052 | -0.53425 | 0.39811 | 0.57086 | 0.79479 | -0.33135 | 0.71027 | 0.76487 | 0.9992 | -0.00116 |
| PE160/161 | 0.25908 | 0.37732 | 0.66638 | -0.58558 | 0.076489 | 0.17411 | 0.50713 | -0.97958 | 0.003782 | 0.018938 | 0.24264 | -2.0431 |
| PEO-180/183 | 0.068387 | 0.14044 | 0.62187 | -0.68532 | 0.10265 | 0.20688 | 0.66949 | -0.57887 | 0.000241 | 0.008255 | 0.27581 | -1.8583 |
| TAG568-FA182 | 0.065587 | 0.13768 | 0.50771 | -0.97792 | 0.07398 | 0.17078 | 0.46723 | -1.0978 | 0.19317 | 0.26652 | 0.63738 | -0.64978 |
| TAG521-FA201 | 0.059713 | 0.13005 | 0.21459 | -2.2204 | 0.034424 | 0.12787 | 0.17855 | -2.4856 | 0.10379 | 0.15935 | 0.24841 | -2.0092 |
| TAG543-FA203 | 0.049439 | 0.11333 | 0.40333 | -1.31 | 0.044613 | 0.13262 | 0.36048 | -1.472 | 0.03446 | 0.067368 | 0.36398 | -1.4581 |
| PE181/161 | 0.033384 | 0.086518 | 0.72756 | -0.45886 | 0.97193 | 0.98818 | 1.0346 | 0.049123 | 0.006679 | 0.023006 | 0.58858 | -0.7647 |
| TAG501-FA181 | 0.081459 | 0.1607 | 0.33523 | -1.5768 | 0.070587 | 0.16832 | 0.31414 | -1.6705 | 0.17061 | 0.24197 | 0.3986 | -1.327 |
| TAG566-FA204 | 0.0847 | 0.16523 | 0.54759 | -0.86884 | 0.084091 | 0.18187 | 0.54491 | -0.8759 | 0.23361 | 0.30662 | 0.65451 | -0.61152 |
| LPC225 | 0.22779 | 0.34095 | 1.2072 | 0.27163 | 0.36989 | 0.53991 | 1.1862 | 0.24637 | 0.6368 | 0.70026 | 1.1181 | 0.16102 |

| | | | | | | | | | | | | |
|---|---|---|---|---|---|---|---|---|---|---|---|---|
| PE160/183 | 0.94835 | 0.97225 | 0.98308 | -0.02462 | 0.028941 | 0.12307 | 0.45629 | -1.132 | 0.000991 | 0.012765 | 0.27598 | -1.8574 |
| HCER160 | 0.21262 | 0.32416 | 1.1589 | 0.21276 | 0.067011 | 0.16355 | 0.84065 | -0.25042 | 0.094478 | 0.14785 | 0.82416 | -0.27901 |
| TAG547-FA225 | 0.00964 | 0.048275 | 0.26225 | -1.931 | 0.050695 | 0.14164 | 0.36279 | -1.4628 | 0.036242 | 0.069754 | 0.32485 | -1.6222 |
| TAG564-FA160 | 0.16431 | 0.26675 | 0.35518 | -1.4934 | 0.10475 | 0.21048 | 0.3134 | -1.6739 | 0.080794 | 0.12891 | 0.29327 | -1.7697 |
| PG160/202_2 | 0.91381 | 0.95122 | 1.0346 | 0.049068 | 0.069233 | 0.1667 | 0.63939 | -0.64523 | 0.036324 | 0.069754 | 0.59662 | -0.74512 |
| TAG540-FA180 | 0.032908 | 0.086385 | 0.16112 | -2.6338 | 0.016639 | 0.11428 | 0.12883 | -2.9564 | 0.003328 | 0.018829 | 0.078062 | -3.6792 |
| PE180/161 | 0.92696 | 0.95786 | 1.0077 | 0.011069 | 0.43161 | 0.60166 | 0.77873 | -0.3608 | 0.017954 | 0.042501 | 0.40683 | -1.2975 |
| PE180/226 | 0.040437 | 0.097862 | 1.3124 | 0.39221 | 0.14797 | 0.27288 | 0.76811 | -0.38062 | 0.92152 | 0.9403 | 1.0402 | 0.056828 |
| TAG563-FA160 | 0.12941 | 0.22585 | 0.35729 | -1.4848 | 0.10019 | 0.20383 | 0.33108 | -1.5948 | 0.034078 | 0.067002 | 0.25578 | -1.967 |
| SMd181/226 | 0.58144 | 0.69709 | 0.96109 | -0.05726 | 0.032895 | 0.12787 | 0.8157 | -0.2939 | 0.89669 | 0.92364 | 1.0394 | 0.055724 |
| PEO-180/205 | 0.43366 | 0.56006 | 0.84128 | -0.24935 | 0.35136 | 0.51633 | 0.81195 | -0.30054 | 0.052711 | 0.093973 | 0.60946 | -0.71441 |
| HCER220 | 0.52969 | 0.64818 | 1.0897 | 0.1239 | 0.81979 | 0.88727 | 0.97839 | -0.03152 | 0.64102 | 0.70371 | 0.93908 | -0.09068 |
| TAG545-FA204 | 0.14447 | 0.24267 | 0.51371 | -0.96098 | 0.082828 | 0.18034 | 0.44457 | -1.1695 | 0.38852 | 0.46751 | 0.6557 | -0.6089 |
| SMd181/220 | 0.8402 | 0.90708 | 1.0255 | 0.036266 | 0.025083 | 0.12086 | 0.78154 | -0.35561 | 0.78248 | 0.83234 | 1.0346 | 0.049131 |
| TAG567-FA180 | 0.024958 | 0.074569 | 0.38633 | -1.3721 | 0.030497 | 0.12449 | 0.35307 | -1.502 | 0.061249 | 0.10456 | 0.45108 | -1.1485 |
| PC160/182 | 0.69631 | 0.79248 | 1.0266 | 0.037939 | 0.60959 | 0.73675 | 0.95482 | -0.0667 | 0.91782 | 0.93947 | 1.0186 | 0.026624 |
| TAG567-FA225 | 0.10614 | 0.19686 | 0.48491 | -1.0442 | 0.08161 | 0.17949 | 0.42061 | -1.2495 | 0.18845 | 0.2627 | 0.56222 | -0.83079 |
| TAG545-FA160 | 0.035075 | 0.088847 | 0.43747 | -1.1928 | 0.038839 | 0.12835 | 0.3809 | -1.3925 | 0.13961 | 0.20378 | 0.53797 | -0.89439 |
| PEP-181/225 | 0.11888 | 0.21333 | 1.1461 | 0.19667 | 0.99532 | 0.99532 | 0.99642 | -0.00517 | 0.12566 | 0.18592 | 1.2965 | 0.37459 |
| PEP-182/182 | 0.004134 | 0.032816 | 0.66275 | -0.59346 | 0.53361 | 0.68539 | 0.92204 | -0.11709 | 0.001683 | 0.014415 | 0.59302 | -0.75385 |
| TAG546-FA226 | 0.98605 | 0.98909 | 0.93506 | -0.09687 | 0.33136 | 0.49703 | 0.66706 | -0.5841 | 0.83597 | 0.87214 | 1.7912 | 0.8409 |
| PE160/226 | 0.12284 | 0.2179 | 1.1852 | 0.24515 | 0.077696 | 0.17624 | 0.72752 | -0.45894 | 0.26612 | 0.34036 | 0.85091 | -0.23292 |
| FFA203 | 0.64677 | 0.75994 | 0.88843 | -0.17067 | 0.11126 | 0.21955 | 0.69316 | -0.52874 | 0.92132 | 0.9403 | 1.0249 | 0.035435 |
| LPE202 | 0.87884 | 0.9318 | 0.97899 | -0.03064 | 0.16262 | 0.29571 | 0.81947 | -0.28724 | 0.004711 | 0.019874 | 0.48271 | -1.0508 |
| SMd181/205 | 0.61225 | 0.72468 | 0.95907 | -0.06029 | 0.000747 | 0.10917 | 0.67627 | -0.56433 | 0.58934 | 0.65543 | 1.0978 | 0.13462 |
| TAG566-FA160 | 0.018651 | 0.063783 | 0.47299 | -1.0801 | 0.029114 | 0.12307 | 0.39107 | -1.3545 | 0.11941 | 0.17829 | 0.55257 | -0.85577 |
| PG160/202 | 0.97922 | 0.98883 | 1.0062 | 0.008868 | 0.074306 | 0.17078 | 0.64745 | -0.62717 | 0.03417 | 0.067002 | 0.59456 | -0.75011 |
| TAG547-FA226 | 0.174 | 0.27832 | 0.36309 | -1.4616 | 0.27482 | 0.43635 | 0.47522 | -1.0733 | 0.32403 | 0.40027 | 0.53895 | -0.89178 |
| SM241 | 0.4757 | 0.60366 | 1.0471 | 0.066437 | 0.014359 | 0.11428 | 0.84368 | -0.24524 | 0.099317 | 0.15468 | 1.1367 | 0.1849 |
| PEP-181/205 | 0.29996 | 0.42451 | 0.80886 | -0.30604 | 0.27006 | 0.42985 | 0.8001 | -0.32175 | 0.0638 | 0.10677 | 0.65012 | -0.62123 |
| HCER241 | 0.89468 | 0.93941 | 0.98769 | -0.01787 | 0.21123 | 0.35878 | 0.89021 | -0.16778 | 0.14376 | 0.20844 | 0.8569 | -0.22281 |
| FFA226 | 0.55897 | 0.6759 | 1.071 | 0.098999 | 0.88935 | 0.92192 | 0.94024 | -0.08889 | 0.32803 | 0.40368 | 1.3417 | 0.42411 |
| FFA220 | 0.055607 | 0.12483 | 1.3139 | 0.39382 | 0.64731 | 0.7634 | 1.0161 | 0.023094 | 0.02863 | 0.05851 | 1.3661 | 0.45003 |

| | | | | | | | | | | | | |
|---|---|---|---|---|---|---|---|---|---|---|---|---|
| PEP-180/160 | 0.7694 | 0.85475 | 0.96163 | -0.05645 | 0.60077 | 0.73225 | 0.92879 | -0.10658 | 0.72687 | 0.77827 | 0.95241 | -0.07034 |
| PE160/201 | 0.81799 | 0.89347 | 0.97829 | -0.03167 | 0.19695 | 0.34043 | 0.71915 | -0.47564 | 0.033732 | 0.066656 | 0.49995 | -1.0002 |
| TAG565-FA180 | 0.023869 | 0.072951 | 0.42679 | -1.2284 | 0.028255 | 0.12263 | 0.40649 | -1.2987 | 0.074474 | 0.1206 | 0.49015 | -1.0287 |
| PG161/182_2 | 0.80375 | 0.88088 | 0.91342 | -0.13066 | 0.59034 | 0.72786 | 1.1903 | 0.25133 | 0.041674 | 0.07841 | 0.56515 | -0.82329 |
| TAG566-FA181 | 0.004527 | 0.033873 | 0.39167 | -1.3523 | 0.018263 | 0.11428 | 0.41915 | -1.2545 | 0.035061 | 0.068338 | 0.4669 | -1.0988 |
| PC140/225 | 0.91469 | 0.95122 | 1.0215 | 0.030645 | 0.23206 | 0.38441 | 1.1714 | 0.22825 | 0.007307 | 0.024452 | 0.75353 | -0.40827 |
| PC180/181 | 0.056363 | 0.1258 | 0.77066 | -0.37584 | 0.36101 | 0.52812 | 0.88191 | -0.18129 | 0.044601 | 0.082721 | 0.73072 | -0.45262 |
| TAG501-FA160 | 0.061989 | 0.13227 | 0.36079 | -1.4708 | 0.041933 | 0.12885 | 0.32098 | -1.6395 | 0.11338 | 0.17245 | 0.40659 | -1.2984 |
| PC160/224 | 0.48925 | 0.61256 | 1.2184 | 0.28493 | 0.74603 | 0.83735 | 0.97239 | -0.0404 | 0.83057 | 0.86877 | 0.97353 | -0.03871 |
| TAG481-FA161 | 0.090249 | 0.17404 | 0.33493 | -1.578 | 0.043306 | 0.13022 | 0.28356 | -1.8183 | 0.072243 | 0.11833 | 0.35588 | -1.4905 |
| PEP-181/183 | 0.10871 | 0.19906 | 0.76353 | -0.38924 | 0.29012 | 0.4551 | 0.84136 | -0.24921 | 0.03919 | 0.074381 | 0.62775 | -0.67173 |
| PEO-160/203 | 0.60822 | 0.72254 | 0.88417 | -0.17761 | 0.39637 | 0.56962 | 0.82293 | -0.28116 | 0.076057 | 0.12256 | 0.63899 | -0.64613 |
| PC181/201 | 0.076702 | 0.15317 | 0.68965 | -0.53607 | 0.51998 | 0.67432 | 0.88932 | -0.16922 | 0.058738 | 0.10197 | 0.60892 | -0.71567 |
| TAG548-FA204 | 0.051663 | 0.11726 | 0.49795 | -1.0059 | 0.060632 | 0.15419 | 0.48405 | -1.0468 | 0.09879 | 0.15423 | 0.54489 | -0.87597 |
| PEP-181/203 | 0.53473 | 0.65311 | 0.8946 | -0.16069 | 0.48061 | 0.64778 | 0.88217 | -0.18087 | 0.23481 | 0.30757 | 0.79427 | -0.33229 |
| PE180/180 | 0.092633 | 0.17581 | 1.2295 | 0.29808 | 0.77129 | 0.85472 | 0.9795 | -0.02988 | 0.072381 | 0.11833 | 1.4856 | 0.57108 |
| TAG565-FA160 | 0.057536 | 0.1274 | 0.42488 | -1.2349 | 0.046301 | 0.13557 | 0.35461 | -1.4957 | 0.1155 | 0.17402 | 0.47516 | -1.0735 |
| SMd181/222 | 0.88491 | 0.93519 | 1.019 | 0.027209 | 0.026307 | 0.12146 | 0.7847 | -0.34979 | 0.56566 | 0.63598 | 1.0924 | 0.1275 |
| PC181/204 | 0.36243 | 0.49052 | 0.7889 | -0.34208 | 0.41188 | 0.58389 | 0.80341 | -0.3158 | 0.68203 | 0.74012 | 0.90871 | -0.13811 |
| TAG481-FA180 | 0.47829 | 0.60577 | 0.61802 | -0.69428 | 0.77201 | 0.85472 | 0.97292 | -0.03961 | 0.03774 | 0.071839 | 0.30564 | -1.7101 |
| PC181/181 | 0.13841 | 0.23525 | 0.78673 | -0.34606 | 0.31543 | 0.47977 | 1.5644 | 0.64563 | 0.26159 | 0.33522 | 0.81667 | -0.29218 |
| HCER140 | 0.041617 | 0.099973 | 0.73185 | -0.45038 | 0.052709 | 0.14281 | 0.71402 | -0.48597 | 0.000576 | 0.009864 | 0.5188 | -0.94675 |
| TAG542-FA180 | 0.066058 | 0.13783 | 0.30762 | -1.7008 | 0.29902 | 0.46238 | 0.47049 | -1.0878 | 0.016035 | 0.039244 | 0.21545 | -2.2146 |
| TAG545-FA225 | 0.45211 | 0.57823 | 0.6242 | -0.67993 | 0.092512 | 0.1943 | 0.42397 | -1.238 | 0.605 | 0.66982 | 0.88921 | -0.16941 |
| PE170/225 | 0.25434 | 0.37376 | 0.75237 | -0.41049 | 0.022159 | 0.11722 | 0.52029 | -0.94261 | 0.001379 | 0.013395 | 0.37156 | -1.4283 |
| PEO-180/203 | 0.18751 | 0.29485 | 0.77576 | -0.36632 | 0.1507 | 0.27636 | 0.75469 | -0.40604 | 0.00706 | 0.024064 | 0.52135 | -0.93968 |
| PA180/181 | 0.33618 | 0.46284 | 0.84961 | -0.23513 | 0.51532 | 0.66961 | 1.2659 | 0.34012 | 0.71083 | 0.76487 | 0.94163 | -0.08676 |
| PEP-160/202 | 0.31941 | 0.44616 | 0.84088 | -0.25003 | 0.39301 | 0.56603 | 0.86616 | -0.2073 | 0.072526 | 0.11833 | 0.72274 | -0.46844 |
| SMd181/224 | 0.83431 | 0.90365 | 0.98976 | -0.01486 | 0.055091 | 0.14579 | 0.81486 | -0.29538 | 0.83129 | 0.86877 | 1.0456 | 0.064307 |
| FFA183 | 0.39386 | 0.52434 | 0.76027 | -0.39543 | 0.15873 | 0.29026 | 0.63037 | -0.66573 | 0.68886 | 0.74493 | 1.1778 | 0.23605 |
| TAG569-FA226 | 0.056424 | 0.1258 | 0.38638 | -1.3719 | 0.11984 | 0.23013 | 0.44881 | -1.1558 | 0.15352 | 0.22013 | 0.48345 | -1.0486 |
| SMd181/184 | 0.025636 | 0.074839 | 0.72904 | -0.45593 | 0.000717 | 0.10917 | 0.59357 | -0.75252 | 0.005215 | 0.020577 | 0.62126 | -0.68672 |
| PE180/183 | 0.65255 | 0.75994 | 1.0911 | 0.12583 | 0.072992 | 0.17078 | 0.60618 | -0.72218 | 0.00531 | 0.020577 | 0.38505 | -1.3769 |

| | | | | | | | | | | | | |
|---|---|---|---|---|---|---|---|---|---|---|---|---|
| HCER240 | 0.66836 | 0.76628 | 1.0448 | 0.063208 | 0.60178 | 0.73225 | 0.96567 | -0.0504 | 0.003737 | 0.018938 | 0.8035 | -0.31563 |
| PEP-180/161 | 0.42418 | 0.55339 | 0.81348 | -0.29783 | 0.56593 | 0.70986 | 1.1719 | 0.22884 | 0.023981 | 0.051693 | 0.50441 | -0.98732 |
| PEP-180/201 | 0.080346 | 0.15898 | 0.80699 | -0.30938 | 0.52372 | 0.67749 | 0.9347 | -0.09743 | 0.011673 | 0.031663 | 0.63859 | -0.64704 |
| PEP-160/226 | 0.43914 | 0.56387 | 1.0982 | 0.13513 | 0.61236 | 0.73687 | 0.94812 | -0.07686 | 0.063722 | 0.10677 | 1.4744 | 0.56015 |
| SMd181/181 | 0.97125 | 0.98525 | 1.0093 | 0.013333 | 0.008764 | 0.11015 | 0.85087 | -0.23299 | 0.00844 | 0.026417 | 1.3817 | 0.46644 |
| LPC205 | 0.98184 | 0.98883 | 0.9951 | -0.00708 | 0.62307 | 0.74289 | 0.95961 | -0.05948 | 0.95556 | 0.96296 | 1.5408 | 0.6237 |
| PEP-160/204 | 0.9072 | 0.94712 | 1.0263 | 0.037429 | 0.81378 | 0.88727 | 0.96686 | -0.04862 | 0.80104 | 0.84518 | 1.103 | 0.14138 |
| PE170/202 | 0.00817 | 0.044693 | 0.57828 | -0.79017 | 0.009188 | 0.11015 | 0.60108 | -0.73437 | 0.001853 | 0.015268 | 0.534 | -0.90508 |
| HCER260 | 0.067574 | 0.13965 | 0.77354 | -0.37044 | 0.065755 | 0.16251 | 0.76737 | -0.38201 | 0.008212 | 0.025825 | 0.66856 | -0.58088 |
| PE160/160 | 0.14756 | 0.24632 | 1.2492 | 0.32096 | 0.047186 | 0.13653 | 0.7638 | -0.38874 | 0.33721 | 0.4113 | 0.91156 | -0.13358 |
| TAG566-FA205 | 0.17638 | 0.28075 | 0.58792 | -0.76632 | 0.080858 | 0.17949 | 0.48076 | -1.0566 | 0.20202 | 0.27571 | 0.66022 | -0.59898 |
| TAG567-FA160 | 0.061927 | 0.13227 | 0.48133 | -1.0549 | 0.069238 | 0.1667 | 0.44423 | -1.1706 | 0.28585 | 0.35924 | 0.67075 | -0.57615 |
| PEP-180/225 | 0.659 | 0.76472 | 1.0395 | 0.055875 | 0.60827 | 0.73675 | 0.94817 | -0.07679 | 0.41453 | 0.49334 | 1.1448 | 0.19506 |
| PE180/204 | 0.33629 | 0.46284 | 1.1154 | 0.15762 | 0.070366 | 0.16832 | 0.79195 | -0.33652 | 0.3427 | 0.41701 | 0.88834 | -0.17082 |
| PC160/181 | 0.8836 | 0.93519 | 0.97934 | -0.03011 | 0.94516 | 0.96594 | 0.99757 | -0.00352 | 0.90988 | 0.93369 | 0.98966 | -0.01499 |
| LPE200 | 0.25168 | 0.37068 | 0.82285 | -0.2813 | 0.64219 | 0.75874 | 1.1139 | 0.15558 | 0.003839 | 0.018938 | 0.41365 | -1.2735 |
| PC180/226 | 0.90784 | 0.94712 | 1.0075 | 0.010757 | 0.42267 | 0.59302 | 0.89109 | -0.16636 | 0.29924 | 0.37391 | 1.2046 | 0.26853 |
| SMd181/180 | 0.77557 | 0.85902 | 1.0275 | 0.039124 | 0.017525 | 0.11428 | 0.79291 | -0.33476 | 0.23359 | 0.30662 | 1.1568 | 0.21015 |
| PS160/180_2 | 0.035548 | 0.089381 | 1.3631 | 0.4469 | 0.81194 | 0.88686 | 0.98069 | -0.02813 | 0.94125 | 0.95149 | 0.99786 | -0.0031 |
| HCER181 | 0.069864 | 0.14213 | 0.87625 | -0.19059 | 0.037228 | 0.1282 | 0.79579 | -0.32955 | 0.29125 | 0.36462 | 0.89581 | -0.15874 |
| TAG566-FA225 | 0.037766 | 0.093481 | 0.46459 | -1.106 | 0.13402 | 0.25143 | 0.53079 | -0.91377 | 0.22213 | 0.29633 | 0.6181 | -0.69408 |
| PC150/225 | 0.26892 | 0.38904 | 0.72156 | -0.4708 | 0.11271 | 0.221 | 0.6329 | -0.65995 | 0.22884 | 0.3032 | 0.70522 | -0.50386 |
| TAG546-FA160 | 0.1786 | 0.28313 | 0.57595 | -0.79599 | 0.064627 | 0.16182 | 0.43835 | -1.1899 | 0.5573 | 0.62987 | 0.86026 | -0.21716 |
| TAG567-FA204 | 0.20568 | 0.31505 | 0.59917 | -0.73896 | 0.21499 | 0.36258 | 0.59684 | -0.74457 | 0.20039 | 0.27494 | 0.59373 | -0.75212 |
| LPC182 | 0.4893 | 0.61256 | 0.88383 | -0.17815 | 0.65172 | 0.76583 | 1.0953 | 0.13139 | 0.17893 | 0.25213 | 0.73918 | -0.43601 |
| PC181/224 | 0.36395 | 0.49155 | 0.84137 | -0.24918 | 0.27581 | 0.43687 | 0.81186 | -0.30069 | 0.33132 | 0.40696 | 0.82921 | -0.27019 |
| PEO-180/226 | 0.50759 | 0.62942 | 1.1395 | 0.18836 | 0.45006 | 0.62339 | 0.91711 | -0.12483 | 0.13561 | 0.19883 | 1.4373 | 0.52336 |
| SMd181/221 | 0.98805 | 0.98957 | 1.0126 | 0.01811 | 0.11409 | 0.22304 | 0.83309 | -0.26346 | 0.054949 | 0.096943 | 1.2172 | 0.28357 |
| SM240 | 0.97734 | 0.98883 | 1.0078 | 0.011224 | 0.005909 | 0.11015 | 0.77228 | -0.37281 | 0.91074 | 0.93369 | 1.007 | 0.010056 |
| DAG161/182 | 0.26052 | 0.37857 | 0.71568 | -0.48262 | 0.29712 | 0.46053 | 0.74774 | -0.41938 | 0.96413 | 0.97009 | 1.0887 | 0.12257 |
| PEO-180/181 | 0.12456 | 0.21975 | 0.70943 | -0.49526 | 0.50997 | 0.66694 | 0.84873 | -0.23662 | 0.001187 | 0.012765 | 0.41366 | -1.2735 |
| PC170/224 | 0.13478 | 0.23062 | 0.7558 | -0.40393 | 0.03082 | 0.12449 | 0.64519 | -0.63221 | 0.002299 | 0.01627 | 0.53319 | -0.90728 |
| TAG520-FA180 | 0.090749 | 0.17427 | 0.17479 | -2.5163 | 0.03342 | 0.12787 | 0.12016 | -3.0569 | 0.032194 | 0.064092 | 0.1202 | -3.0565 |

| | | | | | | | | | | | | |
|---|---|---|---|---|---|---|---|---|---|---|---|---|
| TAG565-FA204 | 0.11761 | 0.21231 | 0.56556 | -0.82225 | 0.12814 | 0.2425 | 0.56872 | -0.8142 | 0.20769 | 0.28286 | 0.62468 | -0.67882 |
| PE160/202 | 0.38362 | 0.51175 | 1.1897 | 0.25062 | 0.017747 | 0.11428 | 0.63011 | -0.66633 | 0.00921 | 0.027527 | 0.53429 | -0.90431 |
| PC140/204 | 0.19431 | 0.30262 | 0.78925 | -0.34145 | 0.83751 | 0.90268 | 0.97484 | -0.03676 | 0.25292 | 0.32625 | 0.79298 | -0.33464 |
| PEP-181/181 | 0.24773 | 0.3682 | 0.80105 | -0.32004 | 0.973 | 0.98818 | 1.0503 | 0.070755 | 0.067581 | 0.11223 | 0.6734 | -0.57046 |
| TAG568-FA160 | 0.19267 | 0.30152 | 0.5564 | -0.84582 | 0.11722 | 0.22578 | 0.47887 | -1.0623 | 0.28006 | 0.35402 | 0.68598 | -0.54376 |
| FFA120 | 0.54604 | 0.66319 | 1.358 | 0.44149 | 0.38704 | 0.55867 | 0.83239 | -0.26467 | 0.87342 | 0.90397 | 1.0617 | 0.086342 |
| DAG140/160 | 0.39932 | 0.52944 | 1.6713 | 0.74098 | 0.26361 | 0.42269 | 2.1429 | 1.0996 | 0.49398 | 0.5728 | 2.3371 | 1.2247 |
| TAG567-FA181 | 0.070448 | 0.14287 | 0.53196 | -0.9106 | 0.18007 | 0.32029 | 0.59595 | -0.74673 | 0.20913 | 0.28363 | 0.64472 | -0.63325 |
| CER160 | 0.90653 | 0.94712 | 0.96367 | -0.05339 | 0.67459 | 0.78281 | 1.0691 | 0.096419 | 0.23143 | 0.30499 | 1.5931 | 0.6718 |
| TAG482-FA180 | 0.84781 | 0.91378 | 0.74739 | -0.42006 | 0.84262 | 0.90484 | 0.99903 | -0.0014 | 0.045275 | 0.08332 | 0.32861 | -1.6055 |
| LPC221 | 0.032164 | 0.085118 | 0.709 | -0.49615 | 0.47972 | 0.64778 | 0.90147 | -0.14964 | 0.026749 | 0.055458 | 0.60281 | -0.73022 |
| PEP-181/161 | 0.65141 | 0.75994 | 1.0693 | 0.096614 | 0.71074 | 0.81317 | 1.1198 | 0.16329 | 0.83141 | 0.86877 | 0.93817 | -0.09207 |
| PC160/180 | 0.98518 | 0.98909 | 0.98897 | -0.016 | 0.48904 | 0.65238 | 0.9232 | -0.11529 | 0.47605 | 0.55341 | 1.0692 | 0.09651 |
| PC170/225 | 0.12271 | 0.2179 | 0.70831 | -0.49756 | 0.035071 | 0.12787 | 0.62893 | -0.66904 | 0.018624 | 0.043147 | 0.57095 | -0.80855 |
| FFA225 | 0.6661 | 0.76614 | 1.007 | 0.01007 | 0.92491 | 0.95122 | 0.86869 | -0.20308 | 0.21081 | 0.28532 | 1.7166 | 0.77957 |
| FFA202 | 0.92303 | 0.95532 | 0.9761 | -0.03489 | 0.25143 | 0.40819 | 0.69556 | -0.52376 | 0.45421 | 0.53181 | 1.2219 | 0.28907 |
| PEP-181/160 | 0.77589 | 0.85902 | 1.0857 | 0.11858 | 0.97971 | 0.98883 | 0.99528 | -0.00683 | 0.80246 | 0.84531 | 1.058 | 0.081388 |
| PEP-160/224 | 0.66165 | 0.76614 | 1.1357 | 0.18363 | 0.53379 | 0.68539 | 0.88409 | -0.17773 | 0.87521 | 0.90438 | 1.0576 | 0.080791 |
| PEO-160/226 | 0.82069 | 0.89492 | 0.98676 | -0.01922 | 0.4243 | 0.59402 | 0.90717 | -0.14056 | 0.2365 | 0.30916 | 1.287 | 0.36399 |
| PC160/161 | 0.24953 | 0.36976 | 0.81832 | -0.28927 | 0.41389 | 0.58448 | 0.92836 | -0.10725 | 0.26957 | 0.34342 | 0.81187 | -0.30067 |
| SMd181/141 | 0.00292 | 0.029174 | 0.7264 | -0.46117 | 0.004149 | 0.11015 | 0.71754 | -0.47886 | 0.451 | 0.5302 | 0.92128 | -0.11828 |
| SMd181/170 | 0.95915 | 0.97743 | 1.0014 | 0.00206 | 0.003456 | 0.11015 | 0.72191 | -0.47011 | 0.37546 | 0.45348 | 1.1343 | 0.18181 |
| PEO-180/204 | 0.44956 | 0.5761 | 0.85622 | -0.22394 | 0.51298 | 0.6679 | 0.87115 | -0.199 | 0.11272 | 0.17186 | 0.70244 | -0.50955 |
| FFA140 | 0.79048 | 0.87073 | 1.2601 | 0.33353 | 0.40715 | 0.57999 | 0.79252 | -0.33549 | 0.32592 | 0.40184 | 1.2808 | 0.357 |
| SMd181/140 | 0.018712 | 0.063783 | 0.78892 | -0.34204 | 0.005614 | 0.11015 | 0.73414 | -0.44586 | 0.71849 | 0.77058 | 0.97061 | -0.04304 |
| CER180 | 0.22783 | 0.34095 | 1.2244 | 0.2921 | 0.53677 | 0.68652 | 0.93572 | -0.09586 | 0.35083 | 0.4253 | 2.1009 | 1.071 |
| TAG522-FA202 | 0.058538 | 0.12874 | 0.30665 | -1.7054 | 0.032249 | 0.12724 | 0.24564 | -2.0254 | 0.071313 | 0.11753 | 0.31467 | -1.6681 |
| PE160/140 | 0.35614 | 0.48606 | 0.83096 | -0.26715 | 0.86617 | 0.91538 | 0.9783 | -0.03166 | 0.011984 | 0.031843 | 0.51396 | -0.96028 |
| PEP-160/181 | 0.15522 | 0.25517 | 0.76208 | -0.39199 | 0.67285 | 0.78281 | 1.1282 | 0.17402 | 0.10039 | 0.15598 | 0.71586 | -0.48225 |
| TAG504-FA141 | 0.18005 | 0.2845 | 0.42637 | -1.2298 | 0.75609 | 0.84139 | 0.78303 | -0.35287 | 0.14722 | 0.2125 | 0.52749 | -0.92278 |
| FFA204 | 0.32903 | 0.45575 | 0.81047 | -0.30318 | 0.028039 | 0.12263 | 0.585 | -0.7735 | 0.6125 | 0.67698 | 0.90158 | -0.14947 |
| CE182 | 0.98276 | 0.98883 | 0.94443 | -0.08248 | 0.061284 | 0.15524 | 0.57064 | -0.80935 | 0.93875 | 0.95043 | 0.91493 | -0.12827 |
| PEP-182/226 | 0.30499 | 0.42976 | 1.1449 | 0.19519 | 0.99059 | 0.99211 | 1.0079 | 0.011344 | 0.38992 | 0.46833 | 1.2645 | 0.33852 |

| | | | | | | | | | | | | |
|---|---|---|---|---|---|---|---|---|---|---|---|---|
| CER221 | 0.54021 | 0.65734 | 0.91328 | -0.13087 | 0.071777 | 0.16992 | 0.72435 | -0.46524 | 0.49449 | 0.5728 | 0.93291 | -0.10019 |
| TAG500-FA160 | 0.13433 | 0.23062 | 0.24444 | -2.0325 | 0.035843 | 0.12787 | 0.14782 | -2.7581 | 0.10354 | 0.15935 | 0.21051 | -2.248 |
| TAG523-FA203 | 0.059634 | 0.13005 | 0.38888 | -1.3626 | 0.038346 | 0.12835 | 0.32665 | -1.6142 | 0.14622 | 0.21153 | 0.47141 | -1.0849 |
| PA180/203 | 0.88666 | 0.93552 | 0.99611 | -0.00562 | 0.63093 | 0.74816 | 0.92856 | -0.10693 | 0.49709 | 0.57479 | 0.90301 | -0.14719 |
| CER260 | 0.55962 | 0.6759 | 0.84295 | -0.24648 | 0.053768 | 0.14287 | 0.53886 | -0.89203 | 0.05974 | 0.10289 | 0.58745 | -0.76747 |
| TAG526-FA226 | 0.16035 | 0.26294 | 0.35558 | -1.4918 | 0.098408 | 0.20259 | 0.32742 | -1.6108 | 0.14032 | 0.20435 | 0.37321 | -1.4219 |
| TAG566-FA180 | 0.13366 | 0.23062 | 0.58715 | -0.7682 | 0.083386 | 0.18095 | 0.49383 | -1.0179 | 0.21568 | 0.29009 | 0.63472 | -0.65582 |
| LPE203 | 0.49426 | 0.6164 | 0.85275 | -0.22981 | 0.325 | 0.49157 | 0.80114 | -0.31988 | 0.009274 | 0.027527 | 0.47859 | -1.0631 |
| PE160/182 | 0.72177 | 0.8186 | 1.0272 | 0.038719 | 0.072912 | 0.17078 | 0.71731 | -0.47932 | 0.000528 | 0.009864 | 0.45122 | -1.1481 |
| TAG500-FA180 | 0.13497 | 0.23062 | 0.20588 | -2.2802 | 0.01487 | 0.11428 | 0.088634 | -3.496 | 0.06075 | 0.10433 | 0.14542 | -2.7817 |
| PE160/205 | 0.51673 | 0.6347 | 1.3286 | 0.40995 | 0.2466 | 0.40134 | 0.77103 | -0.37514 | 0.30815 | 0.38263 | 0.82028 | -0.28582 |
| PEP-181/224 | 0.94793 | 0.97225 | 1.0427 | 0.060388 | 0.26749 | 0.4268 | 0.78616 | -0.3471 | 0.75367 | 0.80565 | 0.94033 | -0.08877 |
| SM261 | 0.36211 | 0.49052 | 0.92525 | -0.11209 | 0.010603 | 0.11395 | 0.72064 | -0.47265 | 0.52184 | 0.59809 | 0.94589 | -0.08025 |
| PC160/140 | 0.14989 | 0.24892 | 0.80593 | -0.31127 | 0.65707 | 0.76979 | 0.97595 | -0.03512 | 0.049721 | 0.089912 | 0.6783 | -0.56 |
| PC170/205 | 0.56418 | 0.67889 | 0.88851 | -0.17054 | 0.18878 | 0.32948 | 0.66422 | -0.59026 | 0.37199 | 0.45012 | 0.83981 | -0.25187 |
| HCER221 | 0.039073 | 0.095625 | 0.83088 | -0.26729 | 0.007284 | 0.11015 | 0.71761 | -0.47872 | 0.037391 | 0.071382 | 0.80053 | -0.32098 |
| SMd181/201 | 0.5625 | 0.67813 | 0.95087 | -0.07268 | 0.097797 | 0.20259 | 0.84342 | -0.24568 | 0.061354 | 0.10456 | 1.2634 | 0.33737 |
| LPE240 | 0.13483 | 0.23062 | 0.77571 | -0.36641 | 0.9157 | 0.94426 | 0.99946 | -0.00078 | 0.00388 | 0.018938 | 0.53506 | -0.90222 |
| HCER261 | 0.095665 | 0.17999 | 0.78765 | -0.34437 | 0.007355 | 0.11015 | 0.64094 | -0.64174 | 0.007715 | 0.02474 | 0.60228 | -0.73149 |
| PC180/225 | 0.51325 | 0.63281 | 1.0543 | 0.076283 | 0.98525 | 0.98981 | 1.0147 | 0.021071 | 0.31678 | 0.39206 | 1.1036 | 0.14226 |
| SMd181/223 | 0.28877 | 0.41136 | 0.90787 | -0.13945 | 0.012286 | 0.11428 | 0.71587 | -0.48224 | 0.54129 | 0.6139 | 0.94725 | -0.07819 |
| SMd181/225 | 0.042208 | 0.10102 | 0.81586 | -0.2936 | 0.000348 | 0.10917 | 0.73758 | -0.43914 | 0.68214 | 0.74012 | 0.97925 | -0.03025 |
| PE160/204 | 0.35932 | 0.48936 | 1.1495 | 0.20098 | 0.18333 | 0.32395 | 0.80224 | -0.3179 | 0.081019 | 0.12896 | 0.7436 | -0.4274 |
| TAG545-FA205 | 0.31618 | 0.4436 | 0.68409 | -0.54773 | 0.039582 | 0.12837 | 0.36377 | -1.4589 | 0.30177 | 0.37635 | 0.86151 | -0.21506 |
| PEO-160/204 | 0.9566 | 0.97743 | 1.0113 | 0.016274 | 0.8829 | 0.91757 | 0.99991 | -0.00013 | 0.57809 | 0.64662 | 0.89001 | -0.1681 |
| PEO-160/183 | 0.058049 | 0.1281 | 0.67014 | -0.57746 | 0.036023 | 0.12787 | 0.60542 | -0.724 | 0.001674 | 0.014415 | 0.39252 | -1.3492 |
| DAG160/181 | 0.83146 | 0.90214 | 0.86287 | -0.21278 | 0.55472 | 0.70394 | 0.76673 | -0.38322 | 0.040586 | 0.076584 | 0.48194 | -1.0531 |
| TAG566-FA226 | 0.62251 | 0.73415 | 0.70326 | -0.50787 | 0.19935 | 0.34242 | 0.52286 | -0.9355 | 0.49995 | 0.57707 | 0.82403 | -0.27924 |
| PC160/183 | 0.49908 | 0.62123 | 0.8672 | -0.20557 | 0.12896 | 0.24264 | 0.71467 | -0.48464 | 0.065405 | 0.1091 | 0.62241 | -0.68407 |
| PC180/182 | 0.1689 | 0.27216 | 0.88932 | -0.16923 | 0.29154 | 0.45514 | 0.90077 | -0.15077 | 0.27428 | 0.34874 | 0.89303 | -0.16321 |
| FFA161 | 0.94665 | 0.97225 | 1.1712 | 0.22794 | 0.58525 | 0.72709 | 0.74201 | -0.4305 | 0.2234 | 0.29741 | 1.6153 | 0.69183 |
| FFA241 | 0.28294 | 0.40661 | 1.1629 | 0.2177 | 0.48575 | 0.65066 | 0.92896 | -0.1063 | 0.069901 | 0.11579 | 0.83205 | -0.26525 |
| TAG483-FA141 | 0.13622 | 0.23215 | 0.43182 | -1.2115 | 0.059238 | 0.15243 | 0.35531 | -1.4928 | 0.15343 | 0.22013 | 0.61042 | -0.71212 |

| | | | | | | | | | | | | |
|---|---|---|---|---|---|---|---|---|---|---|---|---|
| PEP-181/226 | 0.73544 | 0.82758 | 1.0465 | 0.065633 | 0.58806 | 0.72734 | 0.95745 | -0.06273 | 0.24866 | 0.32246 | 1.2054 | 0.26951 |
| PEP-160/180 | 0.14463 | 0.24267 | 0.77686 | -0.36428 | 0.41969 | 0.59011 | 1.1796 | 0.23827 | 0.2084 | 0.28324 | 0.81912 | -0.28785 |
| FFA201 | 0.64956 | 0.75994 | 1.1399 | 0.18889 | 0.865 | 0.91538 | 0.93732 | -0.09339 | 0.20125 | 0.27524 | 1.3283 | 0.40959 |
| FFA141 | 0.48054 | 0.60744 | 1.815 | 0.85999 | 0.51074 | 0.66694 | 0.7828 | -0.35328 | 0.23032 | 0.30413 | 1.8818 | 0.91209 |
| PC200/226 | 0.60327 | 0.71929 | 0.90827 | -0.13881 | 0.029794 | 0.12449 | 0.60978 | -0.71364 | 0.33738 | 0.4113 | 0.83175 | -0.26577 |
| PEP-140/181 | 0.30672 | 0.43126 | 0.86966 | -0.20148 | 0.16903 | 0.30567 | 0.79082 | -0.33858 | 0.00151 | 0.013843 | 0.51196 | -0.9659 |
| LPE170 | 0.32532 | 0.45156 | 0.75372 | -0.40789 | 0.015688 | 0.11428 | 0.50174 | -0.99499 | 0.002422 | 0.016786 | 0.35884 | -1.4786 |
| FFA240 | 0.40805 | 0.53665 | 0.80259 | -0.31727 | 0.55797 | 0.70668 | 0.84843 | -0.23713 | 0.76315 | 0.81311 | 1.1729 | 0.23013 |
| Cholesterol | 0.95941 | 0.97743 | 0.94292 | -0.08479 | 0.082638 | 0.18034 | 0.5787 | -0.78911 | 0.93362 | 0.94818 | 0.9076 | -0.13987 |
| FFA160 | 0.97163 | 0.98525 | 0.99836 | -0.00237 | 0.32154 | 0.48793 | 0.89346 | -0.16253 | 0.64437 | 0.70501 | 1.0487 | 0.068632 |
| PEP-181/204 | 0.91644 | 0.95151 | 1.0322 | 0.045727 | 0.75205 | 0.83935 | 0.95978 | -0.05923 | 0.93717 | 0.95031 | 1.0254 | 0.036232 |
| PE180/224 | 0.62142 | 0.73415 | 1.0656 | 0.091656 | 0.14972 | 0.27533 | 0.66748 | -0.5832 | 0.10363 | 0.15935 | 0.63212 | -0.66173 |
| CE183 | 0.75024 | 0.83918 | 0.92546 | -0.11175 | 0.056765 | 0.14841 | 0.53706 | -0.89684 | 0.30857 | 0.38263 | 0.73663 | -0.44098 |
| PEP-182/204 | 0.66589 | 0.76614 | 1.0877 | 0.12122 | 0.68614 | 0.79058 | 0.94454 | -0.08231 | 0.79968 | 0.84512 | 1.0685 | 0.095527 |
| CER240 | 0.46857 | 0.59578 | 1.1194 | 0.16273 | 0.5988 | 0.73137 | 1.0592 | 0.082977 | 0.47533 | 0.55341 | 1.1104 | 0.15109 |
| PEP-160/183 | 0.16784 | 0.27113 | 0.70632 | -0.50161 | 0.61 | 0.73675 | 0.86081 | -0.21624 | 0.049866 | 0.089925 | 0.5813 | -0.78264 |
| PA181/182 | 0.20098 | 0.31004 | 0.83082 | -0.26739 | 0.46487 | 0.63311 | 1.1484 | 0.19964 | 0.30449 | 0.37901 | 1.1882 | 0.24884 |
| FFA181 | 0.28357 | 0.40661 | 1.1074 | 0.14719 | 0.66218 | 0.77393 | 1.0336 | 0.047742 | 0.79353 | 0.84135 | 1.0101 | 0.014431 |
| PE180/182 | 0.86343 | 0.91995 | 0.96309 | -0.05425 | 0.14693 | 0.27174 | 0.82708 | -0.27391 | 0.006153 | 0.021941 | 0.65121 | -0.61881 |
| PC160/201 | 0.52069 | 0.63836 | 0.9122 | -0.13258 | 0.38452 | 0.55628 | 0.88211 | -0.18096 | 0.6718 | 0.73134 | 0.93766 | -0.09286 |
| PEP-180/224 | 0.90387 | 0.94712 | 0.9957 | -0.00621 | 0.24012 | 0.39275 | 0.8037 | -0.31527 | 0.8414 | 0.87641 | 0.96831 | -0.04645 |
| PS160/161_2 | 0.10886 | 0.19906 | 1.1379 | 0.18635 | 0.019185 | 0.11428 | 1.4213 | 0.50718 | 0.50687 | 0.58299 | 1.1447 | 0.19496 |
| PE180/181 | 0.50669 | 0.62942 | 1.0771 | 0.10715 | 0.52763 | 0.68018 | 1.1186 | 0.16165 | 0.06231 | 0.10564 | 0.65583 | -0.6086 |
| FFA170 | 0.72702 | 0.82168 | 1.4575 | 0.54352 | 0.074501 | 0.17078 | 0.77227 | -0.37282 | 0.90419 | 0.9299 | 1.0703 | 0.098 |
| FFA182 | 0.41325 | 0.5413 | 1.0733 | 0.10208 | 0.90881 | 0.9406 | 0.99731 | -0.00388 | 0.80494 | 0.84655 | 0.95841 | -0.06128 |
| PC150/161 | 0.93553 | 0.96366 | 0.98613 | -0.02016 | 0.88332 | 0.91757 | 1.0092 | 0.013251 | 0.52426 | 0.59876 | 1.2198 | 0.28661 |
| PC180/205 | 0.79993 | 0.87817 | 0.96661 | -0.049 | 0.56246 | 0.70824 | 0.92184 | -0.11741 | 0.98895 | 0.98895 | 1.3884 | 0.47343 |
| FFA180 | 0.36702 | 0.49468 | 0.91253 | -0.13206 | 0.038038 | 0.12835 | 0.79511 | -0.33078 | 0.78711 | 0.8359 | 1.0399 | 0.056383 |
| TAG524-FA204 | 0.36158 | 0.49052 | 0.56921 | -0.81296 | 0.046439 | 0.13557 | 0.32971 | -1.6007 | 0.89275 | 0.92105 | 0.86228 | -0.21376 |
| LPE182 | 0.99175 | 0.99175 | 0.91455 | -0.12887 | 0.87045 | 0.9154 | #N/A | #N/A | 0.019238 | 0.044098 | 0.44463 | -1.1693 |
| TAG480-FA160 | 0.27041 | 0.39032 | 0.27978 | -1.8376 | 0.042718 | 0.12968 | 0.13744 | -2.8631 | 0.24745 | 0.32154 | 0.27606 | -1.8569 |
| CER200 | 0.29562 | 0.41928 | 0.79481 | -0.33133 | 0.72633 | 0.82549 | 0.98036 | -0.02862 | 0.54917 | 0.62176 | 1.1938 | 0.25562 |
| CER220 | 0.96704 | 0.98366 | 0.9294 | -0.10563 | 0.58231 | 0.72483 | 0.8456 | -0.24196 | 0.55927 | 0.63099 | 1.1844 | 0.24414 |

| | | | | | | | | | | | | |
|---|---|---|---|---|---|---|---|---|---|---|---|---|
| PEO-160/160 | 0.82717 | 0.90048 | 0.98639 | -0.01977 | 0.036069 | 0.12787 | 0.71975 | -0.47444 | 0.015173 | 0.037991 | 0.67442 | -0.56828 |
| TAG564-FA204 | 0.17129 | 0.27465 | 0.54941 | -0.86404 | 0.080553 | 0.17949 | 0.45823 | -1.1259 | 0.21816 | 0.29222 | 0.58006 | -0.78572 |
| PS160/181 | 0.21446 | 0.32544 | 1.1701 | 0.22669 | 0.14093 | 0.26212 | 1.1621 | 0.21677 | 0.28093 | 0.35443 | 1.2871 | 0.3641 |
| LPE220 | 0.43427 | 0.56006 | 1.1058 | 0.14507 | 0.041009 | 0.12837 | 1.4016 | 0.48712 | 0.19092 | 0.265 | 0.76168 | -0.39275 |
| TAG525-FA205 | 0.40125 | 0.53093 | 0.72942 | -0.45518 | 0.052029 | 0.14281 | 0.3904 | -1.357 | 0.70067 | 0.75645 | 1.4348 | 0.52085 |
| TAG544-FA224 | 0.58638 | 0.70171 | 0.57105 | -0.80832 | 0.10509 | 0.21051 | 0.33066 | -1.5966 | 0.52298 | 0.59835 | 0.56822 | -0.81547 |
| PEO-180/161 | 0.094668 | 0.17863 | 0.68376 | -0.54843 | 0.50738 | 0.66694 | 0.8752 | -0.19232 | 0.002816 | 0.017482 | 0.46802 | -1.0954 |
| FFA221 | 0.4111 | 0.53957 | 1.7844 | 0.83545 | 0.099689 | 0.20367 | 0.62362 | -0.68127 | 0.18811 | 0.2627 | 0.68207 | -0.552 |
| PE170/224 | 0.60788 | 0.72254 | 0.85257 | -0.23011 | 0.019555 | 0.11428 | 0.50475 | -0.98635 | 0.023947 | 0.051693 | 0.53069 | -0.91405 |
| PC180/180 | 0.164 | 0.26675 | 0.8747 | -0.19314 | 0.091232 | 0.19389 | 0.85373 | -0.22814 | 0.98603 | 0.98754 | 1.0379 | 0.053622 |
| CE181 | 0.73605 | 0.82758 | 0.92593 | -0.11102 | 0.074831 | 0.17093 | 0.63408 | -0.65727 | 0.92864 | 0.94461 | 0.9977 | -0.00332 |
| PA220/226 | 0.77799 | 0.85989 | 0.95743 | -0.06276 | 0.56572 | 0.70986 | 0.91319 | -0.13101 | 0.28577 | 0.35924 | 1.2107 | 0.27587 |
| PEO-160/181 | 0.23208 | 0.34653 | 0.80016 | -0.32164 | 0.3734 | 0.54381 | 0.84955 | -0.23522 | 0.004763 | 0.019874 | 0.56785 | -0.81643 |
| TAG482-FA141 | 0.46363 | 0.59167 | 0.50139 | -0.99601 | 0.27737 | 0.43828 | 0.43419 | -1.2036 | 0.25692 | 0.32989 | 0.62875 | -0.66945 |
| FFA150 | 0.49348 | 0.6164 | 2.1843 | 1.1272 | 0.11572 | 0.22478 | 0.81208 | -0.30031 | 0.026179 | 0.054623 | 0.74762 | -0.41963 |
| LPC160 | 0.76314 | 0.85069 | 0.96406 | -0.0528 | 0.34586 | 0.51056 | 0.89483 | -0.16032 | 0.58384 | 0.65194 | 0.93862 | -0.09138 |
| TAG568-FA226 | 0.31696 | 0.44374 | 0.62659 | -0.67441 | 0.25615 | 0.41174 | 0.61939 | -0.69109 | 0.52116 | 0.59809 | 0.87418 | -0.194 |
| LPE201 | 0.86257 | 0.91995 | 0.95201 | -0.07095 | 0.54806 | 0.69719 | 1.1142 | 0.15607 | 0.006185 | 0.021941 | 0.48946 | -1.0307 |
| PE160/224 | 0.76078 | 0.84952 | 1.0019 | 0.002771 | 0.069396 | 0.1667 | 0.4797 | -1.0598 | 0.033789 | 0.066656 | 0.42094 | -1.2483 |
| PC160/203 | 0.64802 | 0.75994 | 1.0994 | 0.13674 | 0.50341 | 0.66694 | 0.85652 | -0.22344 | 0.56531 | 0.63598 | 1.0755 | 0.10497 |
| FFA222 | 0.16646 | 0.26957 | 1.5577 | 0.63939 | 0.61762 | 0.74046 | 0.99174 | -0.01196 | 0.25358 | 0.32625 | 1.2961 | 0.37415 |
| HCER201 | 0.025275 | 0.074571 | 0.87435 | -0.19371 | 0.002956 | 0.11015 | 0.72095 | -0.47202 | 0.12496 | 0.1853 | 0.87719 | -0.18903 |
| TAG544-FA204 | 0.39499 | 0.52478 | 0.61117 | -0.71035 | 0.045777 | 0.13546 | 0.34857 | -1.5205 | 0.40619 | 0.48519 | 0.64315 | -0.63677 |
| TAG567-FA226 | 0.29534 | 0.41928 | 0.58422 | -0.77541 | 0.32721 | 0.49309 | 0.63833 | -0.64763 | 0.61778 | 0.68166 | 0.88579 | -0.17497 |
| PC200/225 | 0.43366 | 0.56006 | 0.89311 | -0.16309 | 0.014752 | 0.11428 | 0.60788 | -0.71815 | 0.17184 | 0.24319 | 0.80245 | -0.31752 |
| TAG503-FA141 | 0.21315 | 0.32421 | 0.46074 | -1.118 | 0.9167 | 0.94426 | 0.88943 | -0.16904 | 0.12096 | 0.18019 | 0.5367 | -0.89781 |
| TAG565-FA225 | 0.28867 | 0.41136 | 0.581 | -0.78339 | 0.032105 | 0.12724 | 0.34636 | -1.5296 | 0.18739 | 0.26235 | 0.53033 | -0.91504 |
| PC160/120 | 0.67154 | 0.76832 | 1.1926 | 0.25409 | 0.75267 | 0.83935 | 0.95654 | -0.0641 | 0.39489 | 0.47343 | 0.87537 | -0.19204 |
| FFA200 | 0.15408 | 0.25394 | 1.4524 | 0.53841 | 0.40898 | 0.58132 | 1.1555 | 0.20852 | 0.18515 | 0.25977 | 1.3879 | 0.47289 |
| PC180/224 | 0.74995 | 0.83918 | 0.94051 | -0.08849 | 0.48548 | 0.65066 | 0.8778 | -0.18803 | 0.41037 | 0.48928 | 0.8524 | -0.23039 |
| PC200/141 | 0.28335 | 0.40661 | 0.91233 | -0.13238 | 0.31024 | 0.47609 | 0.89534 | -0.15949 | 0.64348 | 0.70501 | 0.97356 | -0.03866 |
| PS161/180_2 | 0.12106 | 0.21592 | 1.2078 | 0.27235 | 0.04084 | 0.12837 | 1.3202 | 0.40076 | 0.24512 | 0.31915 | 1.399 | 0.48444 |
| LPE161 | 0.67903 | 0.77553 | 1.0916 | 0.1264 | 0.91588 | 0.94426 | 1.0302 | 0.042925 | 0.26697 | 0.34079 | 0.79614 | -0.32891 |

| | | | | | | | | | | | | |
|---|---|---|---|---|---|---|---|---|---|---|---|---|
| PC200/204 | 0.85711 | 0.91924 | 0.96948 | -0.04471 | 0.048533 | 0.1398 | 0.65639 | -0.60737 | 0.34558 | 0.41973 | 0.81697 | -0.29164 |
| SMd181/183 | 0.14863 | 0.24746 | 0.82811 | -0.2721 | 0.014531 | 0.11428 | 0.73735 | -0.43958 | 0.1523 | 0.21935 | 0.80835 | -0.30696 |
| PC180/200 | 0.92973 | 0.95919 | 0.9861 | -0.0202 | 0.50426 | 0.66694 | 0.93525 | -0.09657 | 0.76093 | 0.81207 | 1.0535 | 0.075248 |
| PE160/181 | 0.48302 | 0.6094 | 1.109 | 0.14921 | 0.52451 | 0.67749 | 0.79834 | -0.32492 | 0.01099 | 0.030873 | 0.44706 | -1.1615 |
| PEP-160/201 | 0.51151 | 0.63187 | 0.88808 | -0.17124 | 0.4761 | 0.64437 | 0.87623 | -0.19062 | 0.7144 | 0.76745 | 0.94874 | -0.07592 |
| PC180/120 | 0.65598 | 0.76258 | 1.1462 | 0.19683 | 0.24391 | 0.39796 | 0.76855 | -0.37979 | 0.33369 | 0.4091 | 0.83271 | -0.26411 |
| PEP-180/226 | 0.95541 | 0.97743 | 1.0059 | 0.008512 | 0.41258 | 0.58389 | 0.92412 | -0.11385 | 0.58832 | 0.65543 | 1.1025 | 0.14082 |
| DAG200/200 | 0.72572 | 0.82164 | 1.0006 | 0.000909 | 0.78675 | 0.86516 | 1.1222 | 0.16632 | 0.86765 | 0.89943 | 0.99711 | -0.00417 |
| PEP-160/161 | 0.86579 | 0.92097 | 1.0464 | 0.065393 | 0.88374 | 0.91757 | 1.1246 | 0.16939 | 0.5773 | 0.64662 | 0.95048 | -0.07327 |
| CER201 | 0.59025 | 0.70505 | 1.1002 | 0.13775 | 0.23881 | 0.39259 | 0.80934 | -0.30518 | 0.42247 | 0.50005 | 0.99822 | -0.00257 |
| PC170/204 | 0.16416 | 0.26675 | 0.78411 | -0.35087 | 0.015647 | 0.11428 | 0.60815 | -0.71751 | 0.14063 | 0.20435 | 0.75063 | -0.41383 |
| PE180/205 | 0.85331 | 0.91784 | 1.1504 | 0.20208 | 0.29644 | 0.46053 | 0.8177 | -0.29035 | 0.44152 | 0.52165 | 0.94068 | -0.08823 |
| FFA224 | 0.66858 | 0.76628 | 1.0673 | 0.093948 | 0.75297 | 0.83935 | 0.91006 | -0.13596 | 0.11575 | 0.17402 | 1.6474 | 0.72023 |
| PC140/224 | 0.17081 | 0.27456 | 0.78791 | -0.34389 | 0.95325 | 0.97268 | 1.0572 | 0.080188 | 0.11742 | 0.17572 | 0.70259 | -0.50925 |

## 8 Gy

| | SD 1 vs. Pre | | | | SD 2 vs. Pre | | | | SD 6 vs. Pre | | | |
|---|---|---|---|---|---|---|---|---|---|---|---|---|
| **Lipid Name** | **p-value** | **FDR** | **FC** | **LOG2FC** | **p-value** | **FDR** | **FC** | **LOG2FC** | **p-value** | **FDR** | **FC** | **LOG2FC** |
| PI160/201 | 0.000556 | 0.002782 | 0.36913 ↓ | -1.438 | 0.02435 | 0.10361 | 0.62176 ↓ | -0.686 | 0.006962 | 0.017297 | 0.44981 ↓ | -1.153 |
| LPE160 | 0.001568 | 0.005971 | 1.825 ↑ | 0.8679 | 0.1703 | 0.31347 | 1.1576 ↑ | 0.2112 | 0.94473 | 0.95352 | 1.0069 ↑ | 0.01 |
| PC182/203 | 7.76E-05 | 0.000924 | 0.37185 ↓ | -1.427 | 0.012919 | 0.070673 | 0.58322 ↓ | -0.778 | 0.003153 | 0.009822 | 0.43677 ↓ | -1.195 |
| PG180/182_2 | 0.001654 | 0.006188 | 0.54071 ↓ | -0.887 | 0.012359 | 0.069961 | 0.6014 ↓ | -0.734 | 0.005289 | 0.013784 | 0.48428 ↓ | -1.046 |
| PI160/204 | 0.002438 | 0.008001 | 0.5781 ↓ | -0.791 | 0.94057 | 0.97192 | 1.0021 ↑ | 0.003 | 0.29272 | 0.37512 | 0.85506 ↓ | -0.226 |
| PG180/182 | 0.001262 | 0.005081 | 0.53555 ↓ | -0.901 | 0.008197 | 0.058932 | 0.58311 ↓ | -0.778 | 0.004232 | 0.012084 | 0.47649 ↓ | -1.07 |
| LPC202 | 0.039668 | 0.077519 | 0.70887 ↓ | -0.496 | 0.052332 | 0.15346 | 0.73862 ↓ | -0.437 | 0.026122 | 0.051311 | 0.64232 ↓ | -0.639 |
| TAG547-FA183 | 3.5E-05 | 0.0006 | 0.17623 ↓ | -2.505 | 0.000405 | 0.032039 | 0.21546 ↓ | -2.215 | 1.01E-05 | 0.000398 | 0.088648 ↓ | -3.496 |
| PI160/182 | 0.002444 | 0.008001 | 0.50863 ↓ | -0.975 | 0.16315 | 0.30871 | 0.81567 ↓ | -0.294 | 0.003572 | 0.010867 | 0.48945 ↓ | -1.031 |
| TAG547-FA182 | 3.24E-05 | 0.000586 | 0.20138 ↓ | -2.312 | 0.001181 | 0.032039 | 0.28343 ↓ | -1.819 | 2.08E-05 | 0.000411 | 0.12896 ↓ | -2.955 |
| PS161/180 | 0.030691 | 0.062831 | 0.50115 ↓ | -0.997 | 0.21093 | 0.36216 | 0.66443 ↓ | -0.59 | 0.011859 | 0.026853 | 0.43911 ↓ | -1.187 |
| TAG526-FA183 | 4.18E-05 | 0.000681 | 0.17075 ↓ | -2.55 | 0.001663 | 0.033246 | 0.27775 ↓ | -1.848 | 1.29E-05 | 0.000408 | 0.089639 ↓ | -3.48 |
| TAG483-FA183 | 0.000324 | 0.001922 | 0.23067 ↓ | -2.116 | 0.49117 | 0.63569 | 1.1606 ↑ | 0.2149 | 0.0002 | 0.001345 | 0.18882 ↓ | -2.405 |
| LPE226 | 0.038867 | 0.076494 | 1.6176 ↑ | 0.6939 | 0.95159 | 0.97403 | 0.94914 ↓ | -0.075 | 0.17848 | 0.24827 | 1.3477 ↑ | 0.4305 |
| PG160/161 | 0.00542 | 0.01446 | 0.44749 ↓ | -1.16 | 0.84513 | 0.91518 | 1.0885 ↑ | 0.1224 | 0.005762 | 0.014769 | 0.42479 ↓ | -1.235 |

| | | | | | | | | | | | | | | | |
|---|---|---|---|---|---|---|---|---|---|---|---|---|---|---|---|
| TAG525-FA182 | 1.62E-05 | 0.000481 | 0.20622 | ↓ | -2.278 | 0.000956 | 0.032039 | 0.30626 | ↓ | -1.707 | 3.11E-05 | 0.00043 | 0.14752 | ↓ | -2.761 |
| PE181/203 | 0.005729 | 0.015223 | 0.55167 | ↓ | -0.858 | 0.005616 | 0.050778 | 0.46075 | ↓ | -1.118 | 0.014091 | 0.030577 | 0.47096 | ↓ | -1.086 |
| PI160/180 | 0.000459 | 0.002467 | 0.37246 | ↓ | -1.425 | 0.050212 | 0.14926 | 0.68937 | ↓ | -0.537 | 0.00058 | 0.002841 | 0.40091 | ↓ | -1.319 |
| PC160/226 | 0.59388 | 0.67827 | 1.0676 | ↑ | 0.0944 | 0.82552 | 0.90019 | 1.0532 | ↑ | 0.0748 | 0.019078 | 0.039303 | 1.4407 | ↑ | 0.5268 |
| PI160/160 | 0.002068 | 0.0072 | 0.40437 | ↓ | -1.306 | 0.60943 | 0.73335 | 0.9576 | ↓ | -0.063 | 0.012588 | 0.02816 | 0.57272 | ↓ | -0.804 |
| PI181/203_2 | 0.008597 | 0.021524 | 0.52363 | ↓ | -0.933 | 0.082666 | 0.20462 | 0.66365 | ↓ | -0.592 | 0.022953 | 0.046098 | 0.54683 | ↓ | -0.871 |
| TAG542-FA200 | 0.000146 | 0.001232 | 0.24988 | ↓ | -2.001 | 0.013514 | 0.071528 | 0.46617 | ↓ | -1.101 | 1.54E-05 | 0.000408 | 0.1435 | ↓ | -2.801 |
| TAG546-FA183 | 0.000228 | 0.001533 | 0.19104 | ↓ | -2.388 | 0.000815 | 0.032039 | 0.26857 | ↓ | -1.897 | 3.2E-06 | 0.000277 | 0.10615 | ↓ | -3.236 |
| TAG504-FA182 | 0.00014 | 0.001201 | 0.25747 | ↓ | -1.958 | 0.036519 | 0.12919 | 0.5265 | ↓ | -0.926 | 0.00013 | 0.001007 | 0.18018 | ↓ | -2.473 |
| PC181/202 | 0.002667 | 0.00846 | 0.48639 | ↓ | -1.04 | 0.009143 | 0.062074 | 0.56465 | ↓ | -0.825 | 0.047522 | 0.084067 | 0.61041 | ↓ | -0.712 |
| PC182/183 | 0.000885 | 0.003947 | 0.40872 | ↓ | -1.291 | 0.01487 | 0.07683 | 0.5437 | ↓ | -0.879 | 0.000989 | 0.004074 | 0.23737 | ↓ | -2.075 |
| PI160/182_2 | 0.0014 | 0.00549 | 0.47358 | ↓ | -1.078 | 0.13682 | 0.27522 | 0.79477 | ↓ | -0.331 | 0.003695 | 0.011006 | 0.47656 | ↓ | -1.069 |
| PI181/203 | 0.000271 | 0.00173 | 0.44512 | ↓ | -1.168 | 0.009505 | 0.06288 | 0.57317 | ↓ | -0.803 | 0.011058 | 0.025437 | 0.48871 | ↓ | -1.033 |
| PG160/161_2 | 0.039771 | 0.077519 | 0.60759 | ↓ | -0.719 | 0.9154 | 0.95348 | 0.9818 | ↓ | -0.027 | 0.056165 | 0.096728 | 0.62285 | ↓ | -0.683 |
| LPC200 | 0.001621 | 0.006098 | 0.46105 | ↓ | -1.117 | 0.012891 | 0.070673 | 0.6015 | ↓ | -0.733 | 0.000567 | 0.002797 | 0.42778 | ↓ | -1.225 |
| TAG504-FA140 | 0.000498 | 0.002614 | 0.27991 | ↓ | -1.837 | 0.03237 | 0.12391 | 0.50528 | ↓ | -0.985 | 0.000124 | 0.000982 | 0.16821 | ↓ | -2.572 |
| TAG525-FA183 | 2.96E-05 | 0.000551 | 0.26869 | ↓ | -1.896 | 0.003475 | 0.037082 | 0.41435 | ↓ | -1.271 | 4.03E-05 | 0.000495 | 0.16291 | ↓ | -2.618 |
| TAG526-FA182 | 0.000194 | 0.001435 | 0.19204 | ↓ | -2.381 | 0.001528 | 0.033246 | 0.26059 | ↓ | -1.94 | 2.4E-05 | 0.00043 | 0.12091 | ↓ | -3.048 |
| TAG525-FA225 | 0.000148 | 0.001234 | 0.38457 | ↓ | -1.379 | 0.13978 | 0.27743 | 0.72721 | ↓ | -0.46 | 0.000306 | 0.00172 | 0.41209 | ↓ | -1.279 |
| TAG504-FA183 | 2.52E-05 | 0.000496 | 0.17328 | ↓ | -2.529 | 0.12326 | 0.25718 | 0.63516 | ↓ | -0.655 | 1.91E-05 | 0.000411 | 0.1265 | ↓ | -2.983 |
| TAG526-FA161 | 4.98E-05 | 0.000718 | 0.15068 | ↓ | -2.731 | 0.00117 | 0.032039 | 0.2359 | ↓ | -2.084 | 2.71E-05 | 0.00043 | 0.11251 | ↓ | -3.152 |
| CE204 | 0.035317 | 0.070526 | 1.5449 | ↑ | 0.6275 | 0.75085 | 0.84862 | 0.91478 | ↓ | -0.129 | 0.16252 | 0.2305 | 1.4813 | ↑ | 0.5669 |
| TAG546-FA181 | 8.02E-05 | 0.000924 | 0.22142 | ↓ | -2.175 | 0.002413 | 0.033393 | 0.32104 | ↓ | -1.639 | 1.33E-05 | 0.000408 | 0.13833 | ↓ | -2.854 |
| PI160/204_2 | 0.001412 | 0.005505 | 0.57779 | ↓ | -0.791 | 0.9992 | 0.9992 | 1.0186 | ↑ | 0.0266 | 0.3733 | 0.45087 | 0.89766 | ↓ | -0.156 |
| TAG503-FA140 | 0.000517 | 0.00267 | 0.28337 | ↓ | -1.819 | 0.19592 | 0.34659 | 0.7839 | ↓ | -0.351 | 0.000566 | 0.002797 | 0.23758 | ↓ | -2.074 |
| PI161/181 | 2.08E-06 | 0.00019 | 0.38751 | ↓ | -1.368 | 0.097233 | 0.22536 | 0.75933 | ↓ | -0.397 | 0.002036 | 0.006961 | 0.48422 | ↓ | -1.046 |
| LPC220 | 0.000395 | 0.002178 | 0.42014 | ↓ | -1.251 | 0.006541 | 0.0539 | 0.5777 | ↓ | -0.792 | 0.00066 | 0.003046 | 0.39101 | ↓ | -1.355 |
| TAG548-FA182 | 1.87E-05 | 0.000481 | 0.18636 | ↓ | -2.424 | 0.000129 | 0.032039 | 0.2337 | ↓ | -2.097 | 2.17E-06 | 0.000277 | 0.11292 | ↓ | -3.147 |
| PC182/202 | 0.003288 | 0.009728 | 0.54185 | ↓ | -0.884 | 0.006697 | 0.054494 | 0.59006 | ↓ | -0.761 | 0.03683 | 0.068114 | 0.61258 | ↓ | -0.707 |
| PI182/182 | 0.014907 | 0.033933 | 0.52067 | ↓ | -0.942 | 0.11869 | 0.25088 | 0.75462 | ↓ | -0.406 | 0.0003 | 0.001698 | 0.24837 | ↓ | -2.009 |
| TAG546-FA182 | 0.000173 | 0.001327 | 0.25582 | ↓ | -1.967 | 0.01095 | 0.066005 | 0.39416 | ↓ | -1.343 | 4.35E-05 | 0.000506 | 0.14514 | ↓ | -2.784 |
| PI181/182_2 | 0.002916 | 0.008955 | 0.4881 | ↓ | -1.035 | 0.32402 | 0.48159 | 0.86948 | ↓ | -0.202 | 0.005904 | 0.015072 | 0.46145 | ↓ | -1.116 |
| PI160/180_2 | 0.000187 | 0.001415 | 0.38075 | ↓ | -1.393 | 0.035326 | 0.12706 | 0.68031 | ↓ | -0.556 | 0.000929 | 0.003928 | 0.44692 | ↓ | -1.162 |

| | | | | | | | | | | | | | | | |
|---|---|---|---|---|---|---|---|---|---|---|---|---|---|---|---|
| TAG525-FA160 | 2.07E-05 | 0.000496 | 0.27707 | ↓ | -1.852 | 0.001904 | 0.033246 | 0.39494 | ↓ | -1.34 | 0.000141 | 0.001054 | 0.21352 | ↓ | -2.228 |
| TAG525-FA161 | 8.12E-05 | 0.000924 | 0.1812 | ↓ | -2.464 | 0.001139 | 0.032039 | 0.26501 | ↓ | -1.916 | 6.12E-05 | 0.000633 | 0.14915 | ↓ | -2.745 |
| TAG505-FA161 | 0.000224 | 0.001533 | 0.13118 | ↓ | -2.93 | 0.029853 | 0.1185 | 0.39517 | ↓ | -1.34 | 6.95E-06 | 0.000323 | 0.089327 | ↓ | -3.485 |
| PG181/182 | 0.005364 | 0.014392 | 0.47129 | ↓ | -1.085 | 0.001084 | 0.032039 | 0.38481 | ↓ | -1.378 | 0.001094 | 0.004374 | 0.30714 | ↓ | -1.703 |
| TAG563-FA200 | 0.000369 | 0.002124 | 0.26868 | ↓ | -1.896 | 0.045739 | 0.14229 | 0.56038 | ↓ | -0.836 | 5.11E-06 | 0.000277 | 0.12777 | ↓ | -2.968 |
| PC160/204 | 0.17422 | 0.24549 | 1.2012 | ↑ | 0.2645 | 0.42822 | 0.57245 | 1.121 | ↑ | 0.1647 | 0.015591 | 0.032953 | 1.4211 | ↑ | 0.507 |
| PC182/182 | 0.000117 | 0.001053 | 0.4476 | ↓ | -1.16 | 0.006416 | 0.053546 | 0.63256 | ↓ | -0.661 | 0.000113 | 0.000934 | 0.3859 | ↓ | -1.374 |
| PI160/181 | 0.001583 | 0.00599 | 0.48993 | ↓ | -1.029 | 0.21251 | 0.36216 | 0.82547 | ↓ | -0.277 | 0.001533 | 0.005703 | 0.46663 | ↓ | -1.1 |
| TAG482-FA182 | 0.001216 | 0.005044 | 0.33824 | ↓ | -1.564 | 0.50893 | 0.64964 | 1.2587 | ↑ | 0.3319 | 0.000719 | 0.003229 | 0.31208 | ↓ | -1.68 |
| PG181/181 | 0.003849 | 0.010708 | 0.512 | ↓ | -0.966 | 0.002175 | 0.033246 | 0.43917 | ↓ | -1.187 | 0.004694 | 0.012786 | 0.44138 | ↓ | -1.18 |
| TAG566-FA183 | 8.21E-05 | 0.000924 | 0.22665 | ↓ | -2.142 | 0.000458 | 0.032039 | 0.26083 | ↓ | -1.939 | 0.003377 | 0.010467 | 0.16542 | ↓ | -2.596 |
| PC182/161 | 0.00057 | 0.002832 | 0.52778 | ↓ | -0.922 | 0.022243 | 0.099535 | 0.69591 | ↓ | -0.523 | 0.003008 | 0.009461 | 0.56648 | ↓ | -0.82 |
| PI181/182 | 0.002185 | 0.007462 | 0.49681 | ↓ | -1.009 | 0.30625 | 0.4669 | 0.86736 | ↓ | -0.205 | 0.003709 | 0.011006 | 0.45126 | ↓ | -1.148 |
| TAG524-FA221 | 0.000162 | 0.001288 | 0.31136 | ↓ | -1.683 | 0.01156 | 0.066967 | 0.49175 | ↓ | -1.024 | 0.000766 | 0.00337 | 0.32059 | ↓ | -1.641 |
| PI160/181_2 | 0.002912 | 0.008955 | 0.49006 | ↓ | -1.029 | 0.14565 | 0.2856 | 0.78649 | ↓ | -0.347 | 0.001108 | 0.004374 | 0.43952 | ↓ | -1.186 |
| TAG541-FA200 | 0.000107 | 0.001001 | 0.22149 | ↓ | -2.175 | 0.011054 | 0.066017 | 0.44522 | ↓ | -1.167 | 4.35E-07 | 9.43E-05 | 0.11082 | ↓ | -3.174 |
| TAG548-FA183 | 4.19E-05 | 0.000681 | 0.15113 | ↓ | -2.726 | 0.000664 | 0.032039 | 0.20611 | ↓ | -2.279 | 3.1E-07 | 9.43E-05 | 0.060102 | ↓ | -4.056 |
| CE226 | 0.057252 | 0.10268 | 1.563 | ↑ | 0.6443 | 0.73201 | 0.83311 | 1.1371 | ↑ | 0.1853 | 0.026325 | 0.051311 | 1.9901 | ↑ | 0.9928 |
| TAG503-FA182 | 4.81E-05 | 0.000718 | 0.26835 | ↓ | -1.898 | 0.16813 | 0.31297 | 0.71981 | ↓ | -0.474 | 0.000194 | 0.001319 | 0.25293 | ↓ | -1.983 |
| LPC183 | 0.074045 | 0.12719 | 0.74234 | ↓ | -0.43 | 0.649 | 0.76401 | 0.95298 | ↓ | -0.069 | 0.003427 | 0.010522 | 0.41605 | ↓ | -1.265 |
| TAG525-FA181 | 0.000152 | 0.00125 | 0.21681 | ↓ | -2.206 | 0.007961 | 0.058424 | 0.37186 | ↓ | -1.427 | 9.46E-05 | 0.000833 | 0.17077 | ↓ | -2.55 |
| PC140/226 | 0.003221 | 0.009575 | 0.67746 | ↓ | -0.562 | 0.52283 | 0.66476 | 1.1586 | ↑ | 0.2124 | 0.34051 | 0.41983 | 0.88603 | ↓ | -0.175 |
| LPC150 | 0.011944 | 0.028379 | 0.67897 | ↓ | -0.559 | 0.67748 | 0.78758 | 0.98019 | ↓ | -0.029 | 0.00642 | 0.016076 | 0.58452 | ↓ | -0.775 |
| PS161/181 | 0.060171 | 0.10685 | 0.54866 | ↓ | -0.866 | 0.48123 | 0.62407 | 0.80413 | ↓ | -0.315 | 0.4624 | 0.53658 | 0.82215 | ↓ | -0.283 |
| PC180/201 | 0.006266 | 0.016381 | 0.6985 | ↓ | -0.518 | 0.18272 | 0.3295 | 0.85291 | ↓ | -0.23 | 0.039624 | 0.071653 | 0.74115 | ↓ | -0.432 |
| PI181/181 | 0.000752 | 0.003511 | 0.44674 | ↓ | -1.163 | 0.47026 | 0.6135 | 0.99079 | ↓ | -0.013 | 0.037369 | 0.068721 | 0.57088 | ↓ | -0.809 |
| PI160/203_2 | 0.003365 | 0.009868 | 0.51452 | ↓ | -0.959 | 0.17046 | 0.31347 | 0.74582 | ↓ | -0.423 | 0.14578 | 0.2095 | 0.70817 | ↓ | -0.498 |
| TAG562-FA200 | 0.000526 | 0.002686 | 0.2924 | ↓ | -1.774 | 0.048654 | 0.14664 | 0.57639 | ↓ | -0.795 | 2.66E-05 | 0.00043 | 0.14924 | ↓ | -2.744 |
| PC180/202 | 0.052488 | 0.095632 | 0.69242 | ↓ | -0.53 | 0.31281 | 0.47249 | 0.84054 | ↓ | -0.251 | 0.22253 | 0.29931 | 0.79139 | ↓ | -0.338 |
| TAG547-FA181 | 5.16E-05 | 0.000718 | 0.22105 | ↓ | -2.178 | 0.001872 | 0.033246 | 0.3246 | ↓ | -1.623 | 4.29E-06 | 0.000277 | 0.13841 | ↓ | -2.853 |
| TAG503-FA181 | 0.000258 | 0.001684 | 0.26648 | ↓ | -1.908 | 0.22022 | 0.37286 | 0.80518 | ↓ | -0.313 | 9.29E-05 | 0.000828 | 0.19116 | ↓ | -2.387 |
| TAG484-FA120 | 0.23339 | 0.31654 | 1.1594 | ↑ | 0.2134 | 0.68575 | 0.79293 | 3.5382 | ↑ | 1.823 | 0.0546 | 0.094785 | 0.57322 | ↓ | -0.803 |
| TAG504-FA161 | 8.3E-05 | 0.000924 | 0.13765 | ↓ | -2.861 | 0.039785 | 0.13704 | 0.43957 | ↓ | -1.186 | 4.91E-05 | 0.00056 | 0.11298 | ↓ | -3.146 |

| | | | | | | | | | | | | | | | |
|---|---|---|---|---|---|---|---|---|---|---|---|---|---|---|---|
| PG160/203_2 | 0.12621 | 0.19333 | 0.73571 | ↓ | -0.443 | 0.02068 | 0.097256 | 0.59019 | ↓ | -0.761 | 0.004957 | 0.013279 | 0.40609 | ↓ | -1.3 |
| TAG524-FA161 | 9.05E-05 | 0.000969 | 0.20913 | ↓ | -2.258 | 0.031517 | 0.12213 | 0.47222 | ↓ | -1.083 | 6.41E-05 | 0.000652 | 0.17758 | ↓ | -2.494 |
| TAG563-FA182 | 0.000876 | 0.003935 | 0.3289 | ↓ | -1.604 | 0.099882 | 0.22536 | 0.66465 | ↓ | -0.589 | 7.06E-05 | 0.000696 | 0.19384 | ↓ | -2.367 |
| TAG565-FA201 | 0.000384 | 0.002144 | 0.32761 | ↓ | -1.61 | 0.022476 | 0.099535 | 0.47004 | ↓ | -1.089 | 2.98E-05 | 0.00043 | 0.23342 | ↓ | -2.099 |
| PI160/203 | 0.000538 | 0.002716 | 0.51736 | ↓ | -0.951 | 0.10154 | 0.22536 | 0.78072 | ↓ | -0.357 | 0.098451 | 0.1494 | 0.71851 | ↓ | -0.477 |
| PI180/180 | 0.000294 | 0.001821 | 0.50512 | ↓ | -0.985 | 0.076877 | 0.19691 | 0.75091 | ↓ | -0.413 | 0.000911 | 0.003926 | 0.44616 | ↓ | -1.164 |
| TAG541-FA160 | 1.78E-05 | 0.000481 | 0.23132 | ↓ | -2.112 | 0.042807 | 0.14075 | 0.59832 | ↓ | -0.741 | 2.33E-06 | 0.000277 | 0.17571 | ↓ | -2.509 |
| PC182/224 | 0.013747 | 0.031962 | 0.56451 | ↓ | -0.825 | 0.31039 | 0.46992 | 0.81759 | ↓ | -0.291 | 0.069712 | 0.11374 | 0.66096 | ↓ | -0.597 |
| PC182/205 | 0.029211 | 0.060177 | 0.47411 | ↓ | -1.077 | 0.13276 | 0.26841 | 0.63582 | ↓ | -0.653 | 0.010046 | 0.023441 | 0.40248 | ↓ | -1.313 |
| TAG545-FA180 | 0.001045 | 0.004533 | 0.31399 | ↓ | -1.671 | 0.01027 | 0.063932 | 0.41522 | ↓ | -1.268 | 1.31E-05 | 0.000408 | 0.14222 | ↓ | -2.814 |
| PI161/181_2 | 0.003005 | 0.009142 | 0.34346 | ↓ | -1.542 | 0.085061 | 0.20831 | 0.76313 | ↓ | -0.39 | 0.006012 | 0.015288 | 0.53776 | ↓ | -0.895 |
| TAG524-FA160 | 1.72E-05 | 0.000481 | 0.39527 | ↓ | -1.339 | 0.003194 | 0.037082 | 0.52298 | ↓ | -0.935 | 0.000187 | 0.00131 | 0.38669 | ↓ | -1.371 |
| PC160/225 | 0.043526 | 0.083585 | 1.2365 | ↑ | 0.3062 | 0.191 | 0.33973 | 1.1982 | ↑ | 0.2609 | 0.017298 | 0.035978 | 1.4445 | ↑ | 0.5306 |
| TAG524-FA183 | 7.82E-06 | 0.000411 | 0.26922 | ↓ | -1.893 | 0.01068 | 0.064975 | 0.49241 | ↓ | -1.022 | 3E-05 | 0.00043 | 0.21389 | ↓ | -2.225 |
| TAG561-FA181 | 8.28E-06 | 0.000411 | 0.21598 | ↓ | -2.211 | 0.053276 | 0.15553 | 0.58406 | ↓ | -0.776 | 3.98E-07 | 9.43E-05 | 0.16438 | ↓ | -2.605 |
| TAG482-FA140 | 0.000331 | 0.001941 | 0.32316 | ↓ | -1.63 | 0.36381 | 0.51264 | 1.0632 | ↑ | 0.0884 | 0.000516 | 0.002602 | 0.26373 | ↓ | -1.923 |
| PI200/204_2 | 0.005919 | 0.0156 | 0.56251 | ↓ | -0.83 | 0.10332 | 0.22578 | 0.7385 | ↓ | -0.437 | 0.029934 | 0.056814 | 0.57205 | ↓ | -0.806 |
| TAG502-FA140 | 0.002603 | 0.00837 | 0.34051 | ↓ | -1.554 | 0.60065 | 0.72952 | 1.2122 | ↑ | 0.2776 | 0.00104 | 0.004233 | 0.25647 | ↓ | -1.963 |
| TAG545-FA183 | 0.000325 | 0.001922 | 0.24283 | ↓ | -2.042 | 0.009006 | 0.062074 | 0.37286 | ↓ | -1.423 | 2.02E-05 | 0.000411 | 0.13144 | ↓ | -2.928 |
| PG160/203 | 0.17156 | 0.2428 | 0.75801 | ↓ | -0.4 | 0.019672 | 0.094862 | 0.60162 | ↓ | -0.733 | 0.005293 | 0.013784 | 0.41968 | ↓ | -1.253 |
| PI160/161 | 2.14E-05 | 0.000496 | 0.40723 | ↓ | -1.296 | 0.5943 | 0.72588 | 1.1737 | ↑ | 0.2311 | 0.090273 | 0.13926 | 0.6833 | ↓ | -0.549 |
| PE182/202 | 0.028911 | 0.059749 | 0.57531 | ↓ | -0.798 | 0.000406 | 0.032039 | 0.39201 | ↓ | -1.351 | 0.003902 | 0.011442 | 0.33282 | ↓ | -1.587 |
| PI182/201 | 0.008277 | 0.020803 | 0.5406 | ↓ | -0.887 | 0.049231 | 0.14769 | 0.66429 | ↓ | -0.59 | 0.012784 | 0.028357 | 0.49992 | ↓ | -1 |
| LPA180 | 0.62653 | 0.70811 | 0.9418 | ↓ | -0.087 | 0.28334 | 0.44661 | 0.88607 | ↓ | -0.175 | 0.94852 | 0.95586 | 0.99963 | ↓ | -5E-04 |
| LPC240 | 0.002142 | 0.007378 | 0.45783 | ↓ | -1.127 | 0.020766 | 0.097256 | 0.59437 | ↓ | -0.751 | 0.020828 | 0.042372 | 0.54604 | ↓ | -0.873 |
| TAG524-FA200 | 0.000206 | 0.001473 | 0.38608 | ↓ | -1.373 | 0.00718 | 0.056927 | 0.51917 | ↓ | -0.946 | 0.000657 | 0.003046 | 0.37987 | ↓ | -1.396 |
| TAG566-FA202 | 0.000732 | 0.003455 | 0.34709 | ↓ | -1.527 | 0.002906 | 0.035692 | 0.34615 | ↓ | -1.531 | 0.000464 | 0.002361 | 0.28471 | ↓ | -1.812 |
| PS181/182 | 0.048302 | 0.089332 | 0.73161 | ↓ | -0.451 | 0.33105 | 0.48321 | 0.87034 | ↓ | -0.2 | 0.000462 | 0.002361 | 0.4855 | ↓ | -1.042 |
| TAG524-FA181 | 1.13E-05 | 0.000459 | 0.23101 | ↓ | -2.114 | 0.013348 | 0.071228 | 0.46332 | ↓ | -1.11 | 9.24E-05 | 0.000828 | 0.21219 | ↓ | -2.237 |
| PG161/180 | 0.41547 | 0.50555 | 0.88087 | ↓ | -0.183 | 0.012534 | 0.07034 | 0.62415 | ↓ | -0.68 | 0.00151 | 0.005651 | 0.42672 | ↓ | -1.229 |
| PI180/203_2 | 0.10203 | 0.16481 | 0.77604 | ↓ | -0.366 | 0.94897 | 0.97403 | 0.97687 | ↓ | -0.034 | 0.18207 | 0.25272 | 0.81581 | ↓ | -0.294 |
| PI180/183_2 | 0.003433 | 0.009932 | 0.6213 | ↓ | -0.687 | 0.27075 | 0.43094 | 0.86337 | ↓ | -0.212 | 0.001733 | 0.006194 | 0.52326 | ↓ | -0.934 |
| LPC241 | 0.000868 | 0.003924 | 0.43784 | ↓ | -1.192 | 0.098614 | 0.22536 | 0.73677 | ↓ | -0.441 | 0.004496 | 0.012425 | 0.46063 | ↓ | -1.118 |

| | | | | | | | | | | | | |
|---|---|---|---|---|---|---|---|---|---|---|---|---|
| CE203 | 0.43034 | 0.5217 | 1.1223 | ↑ 0.1664 | 0.19756 | 0.3476 | 0.72964 | ↓ -0.455 | 0.23677 | 0.3165 | 1.2442 | ↑ 0.3152 |
| TAG524-FA182 | 9.93E-05 | 0.000987 | 0.40841 | ↓ -1.292 | 0.0034 | 0.037082 | 0.50584 | ↓ -0.983 | 0.000286 | 0.001694 | 0.41264 | ↓ -1.277 |
| TAG545-FA182 | 0.000755 | 0.003511 | 0.3236 | ↓ -1.628 | 0.022322 | 0.099535 | 0.46249 | ↓ -1.113 | 0.000135 | 0.001025 | 0.22641 | ↓ -2.143 |
| TAG544-FA201 | 0.000108 | 0.001001 | 0.28025 | ↓ -1.835 | 0.023481 | 0.10123 | 0.53047 | ↓ -0.915 | 0.000292 | 0.001696 | 0.31357 | ↓ -1.673 |
| TAG545-FA161 | 0.00028 | 0.001754 | 0.24734 | ↓ -2.015 | 0.006224 | 0.052991 | 0.34854 | ↓ -1.521 | 0.000426 | 0.002219 | 0.23354 | ↓ -2.098 |
| TAG502-FA181 | 0.000189 | 0.001415 | 0.27908 | ↓ -1.841 | 0.62612 | 0.74789 | 1.0948 | ↑ 0.1307 | 0.000124 | 0.000982 | 0.24762 | ↓ -2.014 |
| PG181/182_2 | 0.004855 | 0.013115 | 0.43171 | -1.2119 | 0.003362 | 0.037082 | 0.38777 | -1.3667 | 0.001657 | 0.005993 | 0.32268 | -1.6318 |
| PC200/182 | 0.015092 | 0.034233 | 0.61271 | -0.70673 | 0.043687 | 0.14149 | 0.67292 | -0.57149 | 0.003711 | 0.011006 | 0.52026 | -0.9427 |
| PC140/203 | 0.001263 | 0.005081 | 0.4787 | -1.0628 | 0.25379 | 0.41202 | 0.76479 | -0.38686 | 0.13137 | 0.19223 | 0.6901 | -0.53513 |
| TAG543-FA183 | 0.001837 | 0.006635 | 0.36492 | -1.4543 | 0.04359 | 0.14149 | 0.56021 | -0.83595 | 5.69E-06 | 0.000285 | 0.17907 | -2.4814 |
| PC181/203 | 0.006891 | 0.017661 | 0.55458 | -0.85054 | 0.008954 | 0.062074 | 0.5475 | -0.86907 | 0.066014 | 0.10935 | 0.67027 | -0.57719 |
| TAG523-FA201 | 0.000157 | 0.001261 | 0.3037 | -1.7193 | 0.034741 | 0.12706 | 0.48291 | -1.0502 | 3.17E-05 | 0.00043 | 0.29135 | -1.7792 |
| LPC201 | 0.014719 | 0.033754 | 0.6461 | -0.63017 | 0.007345 | 0.056927 | 0.67464 | -0.56781 | 0.001809 | 0.00633 | 0.59353 | -0.75262 |
| PE181/205 | 0.34904 | 0.44121 | 0.74428 | -0.42609 | 0.076743 | 0.19691 | 0.55711 | -0.84396 | 0.074764 | 0.12009 | 0.53164 | -0.91147 |
| LPC161 | 0.000629 | 0.003102 | 0.5509 | -0.86014 | 0.35626 | 0.50528 | 0.89303 | -0.16323 | 0.01124 | 0.025765 | 0.60709 | -0.72002 |
| TAG481-FA181 | 0.001106 | 0.004736 | 0.32516 | -1.6208 | 0.85049 | 0.9182 | 1.598 | 0.67626 | 0.000193 | 0.001319 | 0.28159 | -1.8284 |
| PC120/181 | 0.37783 | 0.46963 | 0.75424 | -0.40691 | 0.35225 | 0.50206 | 0.74099 | -0.43248 | 0.1653 | 0.23343 | 0.64157 | -0.64033 |
| PC150/203 | 0.19263 | 0.26853 | 0.71876 | -0.47642 | 0.13782 | 0.27522 | 0.70899 | -0.49617 | 0.083824 | 0.13149 | 0.64377 | -0.63539 |
| TAG567-FA183 | 5.29E-05 | 0.000718 | 0.2755 | -1.8599 | 0.000742 | 0.032039 | 0.31336 | -1.6741 | 0.00037 | 0.001974 | 0.31944 | -1.6464 |
| TAG544-FA183 | 0.001524 | 0.005859 | 0.35101 | -1.5104 | 0.041229 | 0.13979 | 0.57557 | -0.79693 | 1.76E-05 | 0.00041 | 0.21203 | -2.2376 |
| TAG523-FA183 | 1.8E-05 | 0.000481 | 0.26053 | -1.9405 | 0.020939 | 0.097368 | 0.54481 | -0.87619 | 2.86E-05 | 0.00043 | 0.22487 | -2.1528 |
| PE182/201 | 0.000859 | 0.003912 | 0.53766 | -0.89524 | 0.002314 | 0.033393 | 0.43506 | -1.2007 | 0.000981 | 0.004069 | 0.34608 | -1.5308 |
| TAG544-FA180 | 0.002297 | 0.007673 | 0.46186 | -1.1145 | 0.020217 | 0.096066 | 0.53811 | -0.89402 | 5.13E-05 | 0.000566 | 0.28037 | -1.8346 |
| TAG482-FA161 | 5.42E-05 | 0.000718 | 0.22725 | -2.1376 | 0.84629 | 0.91518 | 1.3306 | 0.41209 | 1.63E-05 | 0.000408 | 0.20175 | -2.3094 |
| TAG482-FA160 | 0.000259 | 0.001684 | 0.32389 | -1.6264 | 0.54911 | 0.68925 | 1.1594 | 0.21339 | 0.000278 | 0.001659 | 0.2947 | -1.7627 |
| PI180/181_2 | 0.002289 | 0.007673 | 0.57308 | -0.8032 | 0.19259 | 0.34163 | 0.81615 | -0.2931 | 0.001466 | 0.005549 | 0.47809 | -1.0646 |
| TAG523-FA180 | 0.000476 | 0.002529 | 0.3524 | -1.5047 | 0.08844 | 0.21245 | 0.65874 | -0.60222 | 1.04E-05 | 0.000398 | 0.19513 | -2.3575 |
| PG160/182_2 | 0.047058 | 0.087529 | 0.53349 | -0.90646 | 0.09285 | 0.21901 | 0.57456 | -0.79948 | 0.004293 | 0.012151 | 0.34043 | -1.5546 |
| TAG503-FA161 | 8.25E-05 | 0.000924 | 0.24466 | -2.0312 | 0.21122 | 0.36216 | 0.72927 | -0.45548 | 0.000274 | 0.001653 | 0.25094 | -1.9946 |
| PI180/181 | 0.002114 | 0.007322 | 0.51674 | -0.95248 | 0.1468 | 0.28699 | 0.77555 | -0.3667 | 0.001722 | 0.006194 | 0.42027 | -1.2506 |
| PC182/226 | 0.000154 | 0.001255 | 0.45112 | -1.1484 | 0.023444 | 0.10123 | 0.69653 | -0.52174 | 0.003676 | 0.011006 | 0.51153 | -0.96711 |
| PC181/183 | 0.024884 | 0.052256 | 0.60959 | -0.71409 | 0.3772 | 0.52505 | 0.86013 | -0.21738 | 0.002487 | 0.008135 | 0.3855 | -1.3752 |
| PC182/204 | 0.000437 | 0.002393 | 0.5907 | -0.75951 | 0.016493 | 0.082636 | 0.75543 | -0.40463 | 0.022801 | 0.045955 | 0.65671 | -0.60667 |

| | | | | | | | | | | | | |
|---|---|---|---|---|---|---|---|---|---|---|---|---|
| TAG565-FA182 | 0.000478 | 0.002529 | 0.32316 | -1.6297 | 0.004721 | 0.045867 | 0.40708 | -1.2966 | 0.000216 | 0.001421 | 0.26352 | -1.924 |
| TAG544-FA161 | 0.000929 | 0.004088 | 0.32268 | -1.6318 | 0.038899 | 0.13542 | 0.54945 | -0.86393 | 0.00017 | 0.00123 | 0.28543 | -1.8088 |
| PC181/226 | 0.025832 | 0.054073 | 0.65601 | -0.60821 | 0.009252 | 0.062095 | 0.58265 | -0.7793 | 0.10082 | 0.15264 | 0.7366 | -0.44105 |
| TAG520-FA200 | 8.67E-06 | 0.000411 | 0.21151 | -2.2412 | 0.010537 | 0.064715 | 0.4656 | -1.1028 | 1.44E-05 | 0.000408 | 0.14526 | -2.7833 |
| SMd181/182 | 0.17708 | 0.24791 | 1.1112 | 0.15215 | 0.056159 | 0.16035 | 0.8289 | -0.27072 | 0.036762 | 0.068114 | 1.1839 | 0.24357 |
| PC140/202 | 0.043684 | 0.083642 | 0.64788 | -0.6262 | 0.053954 | 0.15611 | 0.6917 | -0.53178 | 0.17406 | 0.24369 | 0.75957 | -0.39674 |
| TAG524-FA180 | 0.000167 | 0.001312 | 0.311 | -1.685 | 0.073282 | 0.19509 | 0.64626 | -0.62981 | 3.56E-06 | 0.000277 | 0.24174 | -2.0485 |
| PG160/182 | 0.046415 | 0.087171 | 0.51714 | -0.95137 | 0.10045 | 0.22536 | 0.56796 | -0.81614 | 0.004349 | 0.012256 | 0.33414 | -1.5815 |
| PI180/182_2 | 0.01339 | 0.031355 | 0.68655 | -0.54257 | 0.25247 | 0.41121 | 0.87649 | -0.19018 | 0.002571 | 0.008367 | 0.58428 | -0.77527 |
| TAG526-FA181 | 0.0002 | 0.001465 | 0.33523 | -1.5768 | 0.024514 | 0.10363 | 0.4917 | -1.0241 | 0.000146 | 0.001077 | 0.18955 | -2.3993 |
| TAG483-FA161 | 0.00017 | 0.001319 | 0.25678 | -1.9614 | 0.44678 | 0.58759 | 0.94052 | -0.08846 | 3.31E-05 | 0.00044 | 0.21772 | -2.1994 |
| TAG565-FA202 | 0.001357 | 0.005354 | 0.38733 | -1.3684 | 0.002001 | 0.033246 | 0.39089 | -1.3552 | 0.000955 | 0.004011 | 0.38246 | -1.3866 |
| LPC170 | 0.003482 | 0.010029 | 0.64819 | -0.62552 | 0.046039 | 0.14229 | 0.76094 | -0.39415 | 0.003392 | 0.010467 | 0.61528 | -0.7007 |
| TAG545-FA181 | 0.000512 | 0.002667 | 0.33747 | -1.5672 | 0.01931 | 0.093814 | 0.46374 | -1.1086 | 5.99E-05 | 0.000629 | 0.19775 | -2.3383 |
| TAG523-FA202 | 5.7E-05 | 0.000727 | 0.30563 | -1.7101 | 0.063645 | 0.17782 | 0.64094 | -0.64174 | 0.000234 | 0.00148 | 0.3272 | -1.6118 |
| PI180/182 | 0.020151 | 0.043296 | 0.70776 | -0.49867 | 0.56394 | 0.70466 | 0.92829 | -0.10735 | 0.002955 | 0.009339 | 0.56496 | -0.82377 |
| TAG540-FA160 | 1.23E-06 | 0.000134 | 0.21466 | -2.2199 | 0.000451 | 0.032039 | 0.35832 | -1.4807 | 4.83E-06 | 0.000277 | 0.20379 | -2.2949 |
| TAG526-FA160 | 2.42E-05 | 0.000496 | 0.2031 | -2.2998 | 0.007562 | 0.057912 | 0.3924 | -1.3496 | 8.45E-06 | 0.000367 | 0.17884 | -2.4832 |
| TAG526-FA205 | 0.004712 | 0.012781 | 0.44686 | -1.1621 | 0.23553 | 0.38917 | 0.77716 | -0.36371 | 0.19028 | 0.26133 | 0.71785 | -0.47825 |
| PS180/181 | 0.15619 | 0.22752 | 0.73318 | -0.44775 | 0.02682 | 0.10965 | 0.58331 | -0.77767 | 0.19865 | 0.27054 | 0.74578 | -0.42317 |
| TAG523-FA182 | 0.000112 | 0.00103 | 0.43968 | -1.1855 | 0.05736 | 0.16235 | 0.67255 | -0.5723 | 0.000298 | 0.001698 | 0.44373 | -1.1723 |
| TAG545-FA202 | 9.59E-05 | 0.000987 | 0.25711 | -1.9595 | 0.001093 | 0.032039 | 0.31248 | -1.6782 | 2.51E-05 | 0.00043 | 0.20554 | -2.2825 |
| TAG483-FA140 | 0.000322 | 0.001922 | 0.20643 | -2.2763 | 0.17522 | 0.31862 | 0.74658 | -0.42163 | 0.000246 | 0.00151 | 0.15088 | -2.7286 |
| PE180/202 | 0.71524 | 0.77638 | 0.92537 | -0.11189 | 0.048469 | 0.14664 | 0.69243 | -0.53026 | 0.10228 | 0.15413 | 0.68473 | -0.54638 |
| PI180/202_2 | 0.038157 | 0.075503 | 0.57855 | -0.78948 | 0.16134 | 0.30711 | 0.69396 | -0.52708 | 0.074761 | 0.12009 | 0.61323 | -0.7055 |
| PE181/204 | 0.02225 | 0.047181 | 0.77637 | -0.36518 | 0.000571 | 0.032039 | 0.52329 | -0.93433 | 0.14227 | 0.20536 | 0.79599 | -0.32917 |
| TAG523-FA181 | 0.000101 | 0.000987 | 0.4294 | -1.2196 | 0.089666 | 0.21382 | 0.71731 | -0.47933 | 0.000357 | 0.001936 | 0.46444 | -1.1064 |
| TAG503-FA160 | 1.75E-05 | 0.000481 | 0.25335 | -1.9808 | 0.10922 | 0.23311 | 0.6943 | -0.52637 | 0.000245 | 0.00151 | 0.29785 | -1.7474 |
| TAG483-FA182 | 0.11784 | 0.1857 | 0.7085 | -0.49715 | 0.79423 | 0.88383 | 2.5031 | 1.3237 | 0.01512 | 0.032166 | 0.4117 | -1.2803 |
| PC140/161 | 0.00112 | 0.004764 | 0.5792 | -0.78787 | 0.99612 | 0.99919 | 1.0152 | 0.021818 | 0.11228 | 0.16881 | 0.72282 | -0.4683 |
| TAG523-FA161 | 0.000528 | 0.002686 | 0.23181 | -2.109 | 0.16392 | 0.30871 | 0.64299 | -0.63712 | 0.000294 | 0.001696 | 0.19486 | -2.3595 |
| TAG5610-FA182 | 0.000383 | 0.002144 | 0.30636 | -1.7067 | 0.049794 | 0.1487 | 0.58196 | -0.781 | 1.69E-05 | 0.000408 | 0.23947 | -2.0621 |
| PC180/183 | 0.036652 | 0.072746 | 0.70372 | -0.50693 | 0.78429 | 0.87426 | 1.1854 | 0.24542 | 0.013154 | 0.029029 | 0.50135 | -0.99611 |

| | | | | | | | | | | | | |
|---|---|---|---|---|---|---|---|---|---|---|---|---|
| TAG564-FA201 | 0.00378 | 0.010636 | 0.38522 | -1.3762 | 0.087777 | 0.21243 | 0.63667 | -0.65139 | 0.000563 | 0.002797 | 0.29324 | -1.7699 |
| PE182/161 | 0.020013 | 0.043141 | 0.55461 | -0.85044 | 0.075205 | 0.19583 | 0.67936 | -0.55775 | 0.002208 | 0.007296 | 0.38077 | -1.393 |
| PE181/226 | 0.046465 | 0.087171 | 0.68024 | -0.55588 | 0.001321 | 0.033246 | 0.47335 | -1.079 | 0.11799 | 0.17616 | 0.73075 | -0.45254 |
| TAG481-FA140 | 0.000222 | 0.001533 | 0.30242 | -1.7254 | 0.80904 | 0.88967 | 1.4889 | 0.57425 | 0.000649 | 0.00304 | 0.32424 | -1.6249 |
| PS161/182_2 | 0.10041 | 0.16341 | 0.80988 | -0.30422 | 0.39951 | 0.54754 | 0.89432 | -0.16114 | 0.002071 | 0.007024 | 0.5254 | -0.92851 |
| PC140/181 | 0.004518 | 0.012306 | 0.68535 | -0.54508 | 0.76107 | 0.85571 | 0.99065 | -0.01355 | 0.084272 | 0.13158 | 0.73518 | -0.44383 |
| PI160/161_2 | 0.002824 | 0.008839 | 0.50222 | -0.99361 | 0.58305 | 0.71887 | 1.1183 | 0.16137 | 0.2516 | 0.33023 | 0.80452 | -0.31381 |
| PA180/182 | 0.19822 | 0.27455 | 0.69368 | -0.52765 | 0.43536 | 0.57606 | 0.79203 | -0.33637 | 0.30282 | 0.38678 | 0.75025 | -0.41456 |
| TAG504-FA160 | 0.000102 | 0.000987 | 0.29726 | -1.7502 | 0.070179 | 0.18801 | 0.6113 | -0.71005 | 9.2E-05 | 0.000828 | 0.2496 | -2.0023 |
| PC182/225 | 0.006692 | 0.017288 | 0.60575 | -0.72321 | 0.06464 | 0.17831 | 0.73673 | -0.44079 | 0.032164 | 0.060691 | 0.65439 | -0.61177 |
| TAG523-FA160 | 8.38E-05 | 0.000924 | 0.46113 | -1.1168 | 0.035145 | 0.12706 | 0.66117 | -0.5969 | 0.000179 | 0.001271 | 0.47842 | -1.0636 |
| PI180/202 | 0.053062 | 0.095687 | 0.65304 | -0.61476 | 0.15614 | 0.30155 | 0.74677 | -0.42126 | 0.060893 | 0.10243 | 0.66093 | -0.59742 |
| PS180/181_2 | 0.14259 | 0.21248 | 0.76635 | -0.38393 | 0.03659 | 0.12919 | 0.68232 | -0.55148 | 0.3301 | 0.41168 | 0.82738 | -0.27338 |
| SMd181/203 | 0.67198 | 0.74398 | 1.0545 | 0.07658 | 0.55056 | 0.68925 | 0.94101 | -0.08773 | 0.2395 | 0.3195 | 1.0869 | 0.1202 |
| PE181/202 | 0.1724 | 0.24346 | 0.757 | -0.40163 | 0.004577 | 0.045147 | 0.48561 | -1.0421 | 0.014248 | 0.030815 | 0.46189 | -1.1144 |
| TAG484-FA182 | 0.65944 | 0.73493 | 1.5106 | 0.59513 | 0.90729 | 0.94655 | 3.4939 | 1.8048 | 0.12674 | 0.18709 | 0.67806 | -0.56053 |
| PI180/205 | 0.2007 | 0.27682 | 0.79538 | -0.33028 | 0.95081 | 0.97403 | 0.99002 | -0.01447 | 0.49717 | 0.56782 | 0.89053 | -0.16727 |
| PEP-180/182 | 0.1348 | 0.20267 | 0.65044 | -0.6205 | 0.20545 | 0.3559 | 0.70858 | -0.49701 | 0.005115 | 0.013536 | 0.39239 | -1.3496 |
| PE181/182 | 0.009037 | 0.022425 | 0.70246 | -0.5095 | 0.005493 | 0.050778 | 0.55762 | -0.84264 | 0.002864 | 0.009138 | 0.40678 | -1.2977 |
| PEO-160/225 | 0.37373 | 0.46609 | 1.1529 | 0.20524 | 0.82069 | 0.89793 | 0.94386 | -0.08336 | 0.076933 | 0.12186 | 1.4738 | 0.55955 |
| PI180/203 | 0.10017 | 0.16341 | 0.80229 | -0.31781 | 0.87269 | 0.9342 | 0.97207 | -0.04087 | 0.12344 | 0.18264 | 0.82167 | -0.28337 |
| LPC226 | 0.067589 | 0.1186 | 1.2797 | 0.35579 | 0.58133 | 0.71811 | 1.0995 | 0.1369 | 0.20959 | 0.28426 | 1.2373 | 0.30718 |
| PEP-180/181 | 0.050876 | 0.093297 | 0.59446 | -0.75036 | 0.15845 | 0.30429 | 0.70129 | -0.51192 | 0.00119 | 0.004667 | 0.37922 | -1.3989 |
| TAG569-FA204 | 0.018242 | 0.040121 | 0.49891 | -1.0032 | 0.00187 | 0.033246 | 0.49416 | -1.017 | 0.11349 | 0.17023 | 0.73316 | -0.44781 |
| PC140/182 | 0.057648 | 0.10282 | 0.70994 | -0.49424 | 0.57021 | 0.70818 | 1.2258 | 0.29373 | 0.27814 | 0.35997 | 0.8531 | -0.22922 |
| TAG544-FA182 | 0.002405 | 0.007987 | 0.42599 | -1.2311 | 0.066719 | 0.1825 | 0.61539 | -0.70042 | 0.000209 | 0.001387 | 0.29693 | -1.7518 |
| TAG522-FA181 | 0.000697 | 0.003377 | 0.39953 | -1.3236 | 0.32948 | 0.48293 | 0.84591 | -0.24142 | 0.000928 | 0.003928 | 0.463 | -1.1109 |
| PEP-180/203 | 0.2689 | 0.35293 | 0.81681 | -0.29193 | 0.099003 | 0.22536 | 0.73192 | -0.45023 | 0.010169 | 0.023642 | 0.55878 | -0.83965 |
| PC150/182 | 0.000795 | 0.003645 | 0.6111 | -0.71051 | 0.10276 | 0.22578 | 0.79322 | -0.33421 | 0.027707 | 0.053443 | 0.63371 | -0.6581 |
| TAG566-FA182 | 9.23E-05 | 0.000969 | 0.40133 | -1.3171 | 0.002145 | 0.033246 | 0.43991 | -1.1847 | 0.003538 | 0.010812 | 0.49779 | -1.0064 |
| PC182/201 | 0.000904 | 0.004005 | 0.53712 | -0.89668 | 0.002701 | 0.03448 | 0.5761 | -0.7956 | 0.008474 | 0.020504 | 0.53079 | -0.91379 |
| TAG567-FA182 | 0.002696 | 0.008478 | 0.38864 | -1.3635 | 0.000557 | 0.032039 | 0.33677 | -1.5702 | 0.009559 | 0.022546 | 0.47189 | -1.0835 |
| PC160/205 | 0.51527 | 0.6044 | 1.2554 | 0.32813 | 0.889 | 0.93952 | 1.0768 | 0.1068 | 0.30404 | 0.38734 | 1.6464 | 0.71929 |

| | | | | | | | | | | | | |
|---|---|---|---|---|---|---|---|---|---|---|---|---|
| TAG568-FA183 | 0.001434 | 0.005558 | 0.31607 | -1.6617 | 0.00267 | 0.03448 | 0.34129 | -1.5509 | 0.001982 | 0.006863 | 0.30008 | -1.7366 |
| TAG524-FA203 | 0.000125 | 0.001098 | 0.31925 | -1.6472 | 0.076669 | 0.19691 | 0.60009 | -0.73674 | 0.003614 | 0.010942 | 0.39604 | -1.3363 |
| HCER180 | 0.034673 | 0.069453 | 1.4101 | 0.4958 | 0.6851 | 0.79293 | 1.084 | 0.11636 | 0.92664 | 0.94257 | 1.0172 | 0.024595 |
| TAG567-FA203 | 0.002557 | 0.008283 | 0.41093 | -1.283 | 0.002411 | 0.033393 | 0.34234 | -1.5465 | 0.00848 | 0.020504 | 0.42588 | -1.2315 |
| PC170/203 | 0.07771 | 0.13243 | 0.72456 | -0.46482 | 0.067837 | 0.18398 | 0.7118 | -0.49046 | 0.009785 | 0.022996 | 0.64241 | -0.63842 |
| TAG502-FA180 | 0.003102 | 0.009307 | 0.41307 | -1.2755 | 0.44529 | 0.5868 | 1.0589 | 0.082597 | 1.85E-05 | 0.000411 | 0.2228 | -2.1662 |
| TAG483-FA120 | 0.63892 | 0.71467 | 1.9688 | 0.97733 | 0.90404 | 0.94655 | 6.6652 | 2.7366 | 0.17141 | 0.24049 | 0.80661 | -0.31006 |
| SMd181/160 | 0.79861 | 0.84262 | 1.0203 | 0.028992 | 0.4352 | 0.57606 | 0.94775 | -0.07741 | 0.005247 | 0.013772 | 1.2352 | 0.30469 |
| TAG544-FA202 | 0.000775 | 0.003578 | 0.44958 | -1.1533 | 0.00301 | 0.036284 | 0.47278 | -1.0808 | 0.0042 | 0.012046 | 0.41891 | -1.2553 |
| TAG502-FA161 | 0.000217 | 0.001533 | 0.30934 | -1.6927 | 0.6601 | 0.7715 | 1.0634 | 0.088651 | 8.48E-05 | 0.0008 | 0.28939 | -1.7889 |
| PE140/204 | 0.94315 | 0.96152 | 1.1581 | 0.21179 | 0.68389 | 0.79293 | 1.2243 | 0.29196 | 0.43787 | 0.5136 | 0.87371 | -0.19477 |
| TAG563-FA202 | 0.005852 | 0.015485 | 0.44623 | -1.1641 | 0.062571 | 0.17558 | 0.61733 | -0.69588 | 0.002901 | 0.009214 | 0.4102 | -1.2856 |
| TAG569-FA183 | 0.00153 | 0.005859 | 0.3701 | -1.434 | 0.00215 | 0.033246 | 0.36063 | -1.4714 | 0.001656 | 0.005993 | 0.34691 | -1.5274 |
| LPE204 | 0.002026 | 0.007129 | 1.4808 | 0.56634 | 0.63485 | 0.75417 | 0.94789 | -0.07721 | 0.49447 | 0.56673 | 1.1029 | 0.14128 |
| PI180/224 | 0.10531 | 0.16886 | 0.74601 | -0.42274 | 0.95495 | 0.97441 | 0.98364 | -0.0238 | 0.41716 | 0.49524 | 0.86397 | -0.21095 |
| TAG562-FA180 | 9.72E-05 | 0.000987 | 0.30454 | -1.7153 | 0.047867 | 0.14561 | 0.58446 | -0.77481 | 3.59E-05 | 0.000467 | 0.20271 | -2.3025 |
| TAG521-FA161 | 0.000711 | 0.003377 | 0.36302 | -1.4619 | 0.41069 | 0.55996 | 0.93566 | -0.09595 | 3.71E-05 | 0.000474 | 0.22492 | -2.1525 |
| PS181/182_2 | 0.23545 | 0.318 | 0.76838 | -0.3801 | 0.84425 | 0.91518 | 1.0908 | 0.12535 | 0.001079 | 0.004361 | 0.44029 | -1.1835 |
| TAG503-FA183 | 1.22E-06 | 0.000134 | 0.22879 | -2.1279 | 0.044909 | 0.14157 | 0.63357 | -0.65842 | 3.17E-05 | 0.00043 | 0.29258 | -1.7731 |
| PE170/181 | 0.019481 | 0.042273 | 0.71899 | -0.47595 | 0.12042 | 0.25324 | 0.7417 | -0.43108 | 0.000239 | 0.001499 | 0.25724 | -1.9588 |
| TAG569-FA205 | 0.048825 | 0.090043 | 0.59745 | -0.74311 | 0.002079 | 0.033246 | 0.3624 | -1.4643 | 0.10178 | 0.15373 | 0.67692 | -0.56295 |
| TAG563-FA181 | 0.002907 | 0.008955 | 0.41672 | -1.2628 | 0.23406 | 0.38846 | 0.8314 | -0.26639 | 0.000342 | 0.001869 | 0.29223 | -1.7748 |
| PE181/183 | 0.93154 | 0.95803 | 1.0344 | 0.048826 | 0.026686 | 0.10965 | 0.54839 | -0.86672 | 0.001621 | 0.00596 | 0.25665 | -1.9621 |
| TAG481-FA160 | 4.47E-05 | 0.00071 | 0.31617 | -1.6612 | 0.81143 | 0.8908 | 1.6354 | 0.70965 | 9.75E-05 | 0.000846 | 0.35222 | -1.5055 |
| TAG542-FA160 | 6.24E-05 | 0.000782 | 0.34121 | -1.5513 | 0.082478 | 0.20462 | 0.67647 | -0.5639 | 2.38E-05 | 0.00043 | 0.29818 | -1.7458 |
| TAG546-FA203 | 0.003045 | 0.009221 | 0.3879 | -1.3663 | 0.001048 | 0.032039 | 0.31961 | -1.6456 | 0.000293 | 0.001696 | 0.27644 | -1.855 |
| TAG567-FA224 | 0.001253 | 0.005081 | 0.36458 | -1.4557 | 0.001018 | 0.032039 | 0.33397 | -1.5822 | 0.000607 | 0.002864 | 0.31908 | -1.648 |
| TAG501-FA180 | 0.000269 | 0.00173 | 0.30746 | -1.7015 | 0.74656 | 0.84671 | 1.2102 | 0.27525 | 2.76E-05 | 0.00043 | 0.24657 | -2.0199 |
| PC181/161 | 0.00189 | 0.006725 | 0.48331 | -1.049 | 0.17624 | 0.3196 | 0.75651 | -0.40257 | 0.007969 | 0.019577 | 0.50639 | -0.98168 |
| TAG543-FA180 | 0.012802 | 0.030195 | 0.53412 | -0.90478 | 0.16293 | 0.30871 | 0.74712 | -0.42059 | 3.11E-05 | 0.00043 | 0.34026 | -1.5553 |
| PC180/204 | 0.62784 | 0.70836 | 1.0482 | 0.067895 | 0.99867 | 0.9992 | 0.99091 | -0.01317 | 0.07607 | 0.12108 | 1.2175 | 0.28391 |
| TAG544-FA181 | 0.003612 | 0.010224 | 0.4105 | -1.2845 | 0.064246 | 0.17831 | 0.59552 | -0.74777 | 0.000389 | 0.002061 | 0.29185 | -1.7767 |
| TAG523-FA200 | 9.11E-05 | 0.000969 | 0.42049 | -1.2498 | 0.077728 | 0.19691 | 0.69457 | -0.52581 | 0.001482 | 0.005575 | 0.43565 | -1.1988 |

| | | | | | | | | | | | | |
|---|---|---|---|---|---|---|---|---|---|---|---|---|
| PI180/224_2 | 0.08899 | 0.14854 | 0.73321 | -0.4477 | 0.85512 | 0.92009 | 0.951 | -0.07248 | 0.20881 | 0.28379 | 0.79865 | -0.32437 |
| LPC204 | 0.61515 | 0.69888 | 1.0849 | 0.11757 | 0.91943 | 0.95614 | 1.0013 | 0.001918 | 0.2817 | 0.36314 | 1.1586 | 0.21244 |
| TAG522-FA161 | 0.003563 | 0.010129 | 0.38602 | -1.3733 | 0.47423 | 0.61745 | 0.9389 | -0.09096 | 7.62E-05 | 0.00074 | 0.24833 | -2.0097 |
| TAG543-FA160 | 0.000248 | 0.001646 | 0.4104 | -1.2849 | 0.035267 | 0.12706 | 0.62253 | -0.68379 | 0.001109 | 0.004374 | 0.4089 | -1.2902 |
| TAG544-FA221 | 0.00124 | 0.005081 | 0.35826 | -1.4809 | 0.068109 | 0.18398 | 0.58751 | -0.7673 | 9.25E-05 | 0.000828 | 0.2026 | -2.3033 |
| TAG542-FA182 | 0.002189 | 0.007462 | 0.36615 | -1.4495 | 0.077734 | 0.19691 | 0.64169 | -0.64005 | 1.48E-05 | 0.000408 | 0.23125 | -2.1124 |
| PC170/181 | 0.006428 | 0.016672 | 0.65897 | -0.60172 | 0.49395 | 0.63675 | 0.90779 | -0.13957 | 0.001219 | 0.004753 | 0.55116 | -0.85946 |
| TAG567-FA161 | 0.001806 | 0.006606 | 0.29551 | -1.7587 | 0.001699 | 0.033246 | 0.31038 | -1.6879 | 0.004129 | 0.011894 | 0.35981 | -1.4747 |
| PE180/225 | 0.033352 | 0.06764 | 1.3875 | 0.47247 | 0.384 | 0.53301 | 0.88617 | -0.17434 | 0.86375 | 0.88972 | 1.009 | 0.012927 |
| PG182/182 | 0.022987 | 0.048587 | 0.4648 | -1.1053 | 0.011293 | 0.066836 | 0.39774 | -1.3301 | 0.00067 | 0.003071 | 0.23201 | -2.1078 |
| PE181/225 | 0.17517 | 0.24629 | 0.83391 | -0.26204 | 0.002176 | 0.033246 | 0.56566 | -0.82199 | 0.074897 | 0.12009 | 0.72913 | -0.45575 |
| TAG543-FA182 | 0.00308 | 0.009282 | 0.41371 | -1.2733 | 0.080069 | 0.20048 | 0.64913 | -0.62341 | 5.58E-05 | 0.000596 | 0.30141 | -1.7302 |
| PS180/182 | 0.10251 | 0.16518 | 0.83177 | -0.26573 | 0.043314 | 0.14149 | 0.78764 | -0.34439 | 0.008436 | 0.020504 | 0.55454 | -0.85063 |
| SMd181/202 | 0.084108 | 0.14112 | 1.3001 | 0.3786 | 0.69091 | 0.79608 | 0.95538 | -0.06585 | 0.074135 | 0.11976 | 1.2534 | 0.32581 |
| PE160/225 | 0.018806 | 0.041104 | 1.5417 | 0.6245 | 0.88393 | 0.93719 | 0.94492 | -0.08173 | 0.60275 | 0.66733 | 0.90364 | -0.14618 |
| TAG480-FA140 | 2.59E-05 | 0.000496 | 0.27473 | -1.8639 | 0.67271 | 0.78342 | 1.4503 | 0.53638 | 0.001741 | 0.006194 | 0.41904 | -1.2549 |
| PC140/140 | 0.15948 | 0.23072 | 0.66334 | -0.59218 | 0.52479 | 0.66597 | 2.1157 | 1.0811 | 0.35723 | 0.43632 | 0.9071 | -0.14067 |
| TAG482-FA181 | 0.19673 | 0.27307 | 0.87624 | -0.19061 | 0.83003 | 0.90359 | 3.6998 | 1.8874 | 0.012806 | 0.028357 | 0.44883 | -1.1558 |
| TAG564-FA202 | 0.002202 | 0.007467 | 0.44151 | -1.1795 | 0.039532 | 0.13689 | 0.58858 | -0.76468 | 0.000595 | 0.002864 | 0.3438 | -1.5403 |
| TAG543-FA181 | 0.006781 | 0.017449 | 0.46461 | -1.1059 | 0.27219 | 0.43218 | 0.83953 | -0.25235 | 0.002456 | 0.008076 | 0.40952 | -1.288 |
| TAG564-FA181 | 0.001677 | 0.006237 | 0.37071 | -1.4317 | 0.04181 | 0.14021 | 0.54745 | -0.86919 | 0.000782 | 0.003419 | 0.30775 | -1.7002 |
| PE140/182 | 0.76345 | 0.81343 | 1.0884 | 0.12227 | 0.96381 | 0.97885 | 1.1232 | 0.16755 | 0.071617 | 0.11627 | 0.62334 | -0.68191 |
| TAG543-FA161 | 0.00028 | 0.001754 | 0.26037 | -1.9414 | 0.047044 | 0.14422 | 0.54231 | -0.88282 | 3.86E-05 | 0.000483 | 0.18722 | -2.4172 |
| TAG504-FA181 | 0.001952 | 0.006905 | 0.35876 | -1.4789 | 0.086307 | 0.20965 | 0.60725 | -0.71963 | 9.99E-05 | 0.000855 | 0.23613 | -2.0823 |
| TAG568-FA181 | 0.002663 | 0.00846 | 0.48554 | -1.0423 | 0.001734 | 0.033246 | 0.45298 | -1.1425 | 0.004739 | 0.012854 | 0.47487 | -1.0744 |
| PEP-180/202 | 0.10156 | 0.16446 | 0.67456 | -0.56798 | 0.10104 | 0.22536 | 0.67684 | -0.56312 | 0.002042 | 0.006961 | 0.41075 | -1.2837 |
| PE160/203 | 0.90108 | 0.92964 | 1.0041 | 0.005917 | 0.085396 | 0.20831 | 0.72131 | -0.47131 | 0.063466 | 0.10621 | 0.67646 | -0.56393 |
| TAG565-FA181 | 0.001834 | 0.006635 | 0.36306 | -1.4617 | 0.013127 | 0.071113 | 0.44872 | -1.1561 | 0.005211 | 0.013735 | 0.42575 | -1.2319 |
| TAG564-FA182 | 0.001212 | 0.005044 | 0.36817 | -1.4416 | 0.050766 | 0.14954 | 0.55528 | -0.84872 | 8.44E-05 | 0.0008 | 0.24014 | -2.058 |
| TAG566-FA203 | 0.003563 | 0.010129 | 0.40239 | -1.3133 | 0.008238 | 0.058932 | 0.40164 | -1.316 | 0.00537 | 0.013873 | 0.39547 | -1.3384 |
| PEP-160/160 | 0.93543 | 0.95876 | 1.014 | 0.020088 | 0.97182 | 0.98464 | 0.98996 | -0.01455 | 0.43934 | 0.51441 | 1.2034 | 0.26709 |
| TAG522-FA160 | 0.000314 | 0.00191 | 0.40119 | -1.3177 | 0.30147 | 0.46287 | 0.84221 | -0.24775 | 0.001008 | 0.004127 | 0.4554 | -1.1348 |
| LPE224 | 0.15546 | 0.22752 | 1.3591 | 0.44264 | 0.96148 | 0.97801 | 0.96367 | -0.05338 | 0.70165 | 0.75751 | 0.89998 | -0.15203 |

| | | | | | | | | | | | | |
|---|---|---|---|---|---|---|---|---|---|---|---|---|
| PI161/180_2 | 0.04583 | 0.086731 | 0.75883 | -0.39815 | 0.16123 | 0.30711 | 0.82497 | -0.27759 | 0.059417 | 0.10076 | 0.71495 | -0.48409 |
| PEO-160/205 | 0.2964 | 0.38362 | 0.77204 | -0.37325 | 0.08475 | 0.20831 | 0.63636 | -0.65208 | 0.038671 | 0.070518 | 0.58825 | -0.7655 |
| TAG568-FA204 | 0.066742 | 0.11743 | 0.77033 | -0.37644 | 0.00622 | 0.052991 | 0.586 | -0.77103 | 0.41791 | 0.49524 | 0.91033 | -0.13554 |
| PC150/181 | 0.001052 | 0.004537 | 0.56194 | -0.83151 | 0.32158 | 0.48016 | 0.85745 | -0.22188 | 0.00616 | 0.015544 | 0.53052 | -0.91453 |
| TAG546-FA161 | 0.00135 | 0.005354 | 0.28229 | -1.8248 | 0.01022 | 0.063932 | 0.38574 | -1.3743 | 0.001796 | 0.00632 | 0.32286 | -1.631 |
| TAG522-FA182 | 1.25E-05 | 0.000463 | 0.34685 | -1.5276 | 0.027485 | 0.11045 | 0.6365 | -0.65177 | 4.25E-05 | 0.000506 | 0.34511 | -1.5349 |
| LPC140 | 0.069001 | 0.12011 | 0.7691 | -0.37876 | 0.36814 | 0.5154 | 1.2673 | 0.34179 | 0.15338 | 0.2185 | 0.78118 | -0.35628 |
| TAG501-FA140 | 0.003428 | 0.009932 | 0.37088 | -1.431 | 0.78016 | 0.87116 | 1.3845 | 0.46933 | 0.000762 | 0.00337 | 0.28532 | -1.8094 |
| PEP-180/183 | 0.15137 | 0.22346 | 0.61554 | -0.70008 | 0.39655 | 0.54577 | 0.74855 | -0.41784 | 0.004525 | 0.01243 | 0.32125 | -1.6382 |
| TAG543-FA202 | 0.000128 | 0.001107 | 0.39194 | -1.3513 | 0.010008 | 0.063932 | 0.56569 | -0.82192 | 0.000922 | 0.003928 | 0.38862 | -1.3636 |
| PC180/140 | 0.16755 | 0.23973 | 0.76179 | -0.39254 | 0.23432 | 0.38846 | 1.6459 | 0.71887 | 0.73426 | 0.7809 | 1.033 | 0.0468 |
| TAG568-FA225 | 0.002985 | 0.009122 | 0.36063 | -1.4714 | 0.000692 | 0.032039 | 0.30128 | -1.7308 | 0.002184 | 0.007278 | 0.3371 | -1.5688 |
| SMd181/161 | 0.25007 | 0.33428 | 1.1113 | 0.15222 | 0.97254 | 0.98464 | 0.98435 | -0.02275 | 0.004994 | 0.013288 | 1.393 | 0.47823 |
| PEP-180/205 | 0.39076 | 0.48088 | 0.75068 | -0.41374 | 0.15554 | 0.30136 | 0.6175 | -0.69549 | 0.034571 | 0.064486 | 0.4658 | -1.1022 |
| PEP-160/182 | 0.055881 | 0.10049 | 0.59856 | -0.74043 | 0.073421 | 0.19509 | 0.65118 | -0.61888 | 0.042281 | 0.075617 | 0.57989 | -0.78616 |
| SMd181/200 | 0.79597 | 0.84119 | 0.97437 | -0.03746 | 0.57289 | 0.70903 | 0.94405 | -0.08306 | 0.33607 | 0.41673 | 1.0756 | 0.1051 |
| TAG541-FA181 | 0.001279 | 0.00511 | 0.36652 | -1.448 | 0.29496 | 0.45483 | 0.84864 | -0.23678 | 2.64E-05 | 0.00043 | 0.2695 | -1.8916 |
| TAG565-FA203 | 0.003388 | 0.009891 | 0.41045 | -1.2847 | 0.033578 | 0.12553 | 0.51981 | -0.94393 | 0.011921 | 0.026854 | 0.49708 | -1.0084 |
| PG160/160 | 0.22979 | 0.3123 | 0.56346 | -0.82763 | 0.58477 | 0.71963 | 0.73951 | -0.43536 | 0.13853 | 0.20085 | 0.50793 | -0.9773 |
| PEP-160/225 | 0.26365 | 0.34674 | 1.191 | 0.25211 | 0.59724 | 0.72673 | 0.91343 | -0.13064 | 0.15655 | 0.22252 | 1.316 | 0.39621 |
| PC170/226 | 0.017757 | 0.03932 | 0.69625 | -0.52232 | 0.13197 | 0.26764 | 0.78617 | -0.34709 | 0.058848 | 0.10055 | 0.76286 | -0.39052 |
| TAG522-FA180 | 0.000119 | 0.001059 | 0.35065 | -1.5119 | 0.12803 | 0.26128 | 0.71911 | -0.47572 | 5.29E-05 | 0.000574 | 0.32028 | -1.6426 |
| TAG545-FA203 | 0.003139 | 0.009375 | 0.43076 | -1.215 | 0.010201 | 0.063932 | 0.44582 | -1.1655 | 0.015921 | 0.033542 | 0.48569 | -1.0419 |
| PC181/182 | 0.009895 | 0.024127 | 0.68571 | -0.54434 | 0.026193 | 0.10861 | 0.70636 | -0.50153 | 0.011745 | 0.026827 | 0.64614 | -0.63008 |
| TAG548-FA205 | 0.009077 | 0.022425 | 0.40027 | -1.3209 | 0.003859 | 0.039249 | 0.35819 | -1.4812 | 0.025221 | 0.049755 | 0.46957 | -1.0906 |
| TAG562-FA160 | 2.39E-05 | 0.000496 | 0.18176 | -2.4599 | 0.11159 | 0.2374 | 0.64106 | -0.64148 | 0.000101 | 0.000856 | 0.25189 | -1.9892 |
| PG160/180_2 | 0.47931 | 0.57148 | 0.84527 | -0.24251 | 0.064448 | 0.17831 | 0.64012 | -0.64358 | 0.002728 | 0.008791 | 0.44455 | -1.1696 |
| PEP-160/205 | 0.61952 | 0.70141 | 0.88524 | -0.17585 | 0.22809 | 0.38074 | 0.67162 | -0.57429 | 0.1847 | 0.25513 | 0.64671 | -0.62881 |
| TAG543-FA201 | 8.84E-06 | 0.000411 | 0.35466 | -1.4955 | 0.033419 | 0.12553 | 0.61032 | -0.71237 | 0.000133 | 0.001018 | 0.32869 | -1.6052 |
| PEP-181/182 | 0.12385 | 0.19105 | 0.69057 | -0.53414 | 0.097257 | 0.22536 | 0.67954 | -0.55737 | 0.028533 | 0.054632 | 0.57048 | -0.80976 |
| PG160/180 | 0.53192 | 0.62212 | 0.82217 | -0.28248 | 0.042446 | 0.14026 | 0.53024 | -0.91528 | 0.001433 | 0.00546 | 0.33314 | -1.5858 |
| PEO-160/182 | 0.046757 | 0.087217 | 0.63667 | -0.65137 | 0.025937 | 0.10844 | 0.63399 | -0.65748 | 0.004585 | 0.01254 | 0.49686 | -1.0091 |
| TAG483-FA160 | 0.000227 | 0.001533 | 0.34683 | -1.5277 | 0.25056 | 0.40983 | 0.84733 | -0.239 | 0.000813 | 0.003528 | 0.3389 | -1.5611 |

| | | | | | | | | | | | | |
|---|---|---|---|---|---|---|---|---|---|---|---|---|
| TAG547-FA161 | 0.008879 | 0.022148 | 0.39972 | -1.3229 | 0.011624 | 0.066967 | 0.42269 | -1.2423 | 0.05414 | 0.094238 | 0.50753 | -0.97843 |
| TAG565-FA224 | 0.007968 | 0.020182 | 0.50293 | -0.99157 | 0.12495 | 0.25987 | 0.68979 | -0.53577 | 0.048935 | 0.086333 | 0.6242 | -0.67993 |
| TAG502-FA160 | 2.33E-06 | 0.00019 | 0.35854 | -1.4798 | 0.10212 | 0.22536 | 0.73767 | -0.43894 | 0.000363 | 0.001955 | 0.53294 | -0.90795 |
| PE180/150 | 0.67407 | 0.74469 | 1.0433 | 0.061175 | 0.16816 | 0.31297 | 1.1569 | 0.21027 | 0.18537 | 0.25513 | 0.8798 | -0.18476 |
| LPE180 | 0.002915 | 0.008955 | 1.5917 | 0.67061 | 0.95149 | 0.97403 | 0.98489 | -0.02196 | 0.58561 | 0.65391 | 0.93679 | -0.0942 |
| TAG502-FA182 | 1.24E-06 | 0.000134 | 0.2918 | -1.7769 | 0.033799 | 0.12553 | 0.64791 | -0.62613 | 0.001586 | 0.005866 | 0.56393 | -0.82641 |
| PEO-180/224 | 0.94781 | 0.96345 | 1.0794 | 0.1102 | 0.44031 | 0.58142 | 0.85647 | -0.22353 | 0.92605 | 0.94257 | 1.0373 | 0.052901 |
| TAG483-FA181 | 0.64298 | 0.71797 | 1.6507 | 0.72305 | 0.846 | 0.91518 | 5.5638 | 2.4761 | 0.24213 | 0.32234 | 0.79455 | -0.3318 |
| PE182/183 | 0.37296 | 0.46602 | 0.73634 | -0.44155 | 0.001964 | 0.033246 | 0.27717 | -1.8511 | 0.00013 | 0.001007 | 0.11537 | -3.1156 |
| PG160/181 | 0.15396 | 0.22677 | 0.78424 | -0.35064 | 0.042038 | 0.14021 | 0.6598 | -0.59989 | 0.004383 | 0.012299 | 0.51009 | -0.97119 |
| PEP-180/180 | 0.040388 | 0.078485 | 0.61919 | -0.69154 | 0.20305 | 0.35438 | 0.75557 | -0.40437 | 0.002617 | 0.008475 | 0.48775 | -1.0358 |
| PS160/180 | 0.079562 | 0.13523 | 1.3925 | 0.47773 | 0.80896 | 0.88967 | 0.98393 | -0.02337 | 0.59297 | 0.65875 | 1.0531 | 0.0746 |
| PG180/181_2 | 0.12862 | 0.19518 | 0.72474 | -0.46446 | 0.044682 | 0.14157 | 0.62136 | -0.6865 | 0.009132 | 0.021617 | 0.5045 | -0.98708 |
| PC160/160 | 0.16846 | 0.24039 | 1.1784 | 0.23684 | 0.16922 | 0.31297 | 1.1833 | 0.24285 | 0.33514 | 0.41636 | 1.1312 | 0.17785 |
| TAG562-FA201 | 0.005372 | 0.014392 | 0.40587 | -1.3009 | 0.25266 | 0.41121 | 0.80687 | -0.3096 | 0.000232 | 0.00148 | 0.27845 | -1.8445 |
| PEO-160/224 | 0.96592 | 0.97794 | 1.07 | 0.097618 | 0.22108 | 0.37286 | 0.80826 | -0.30712 | 0.68812 | 0.74661 | 1.1181 | 0.16101 |
| PE170/204 | 0.021574 | 0.046048 | 0.71933 | -0.47528 | 0.000371 | 0.032039 | 0.57421 | -0.80036 | 0.003052 | 0.009552 | 0.48836 | -1.034 |
| TAG566-FA224 | 0.007816 | 0.019876 | 0.52765 | -0.92235 | 0.007274 | 0.056927 | 0.48971 | -1.03 | 0.004258 | 0.012105 | 0.48549 | -1.0425 |
| PE182/182 | 0.052872 | 0.095687 | 0.61649 | -0.69785 | 0.007331 | 0.056927 | 0.46534 | -1.1036 | 0.000234 | 0.00148 | 0.21626 | -2.2091 |
| PC181/225 | 0.14263 | 0.21248 | 0.77183 | -0.37365 | 0.021268 | 0.097504 | 0.64079 | -0.64208 | 0.24574 | 0.32374 | 0.80434 | -0.31412 |
| PI161/180 | 0.009576 | 0.023466 | 0.69506 | -0.5248 | 0.42387 | 0.5712 | 0.89415 | -0.16141 | 0.069503 | 0.11368 | 0.73599 | -0.44224 |
| LPE225 | 0.001168 | 0.004905 | 2.0109 | 1.0079 | 0.023114 | 0.10099 | 1.3732 | 0.45757 | 0.017531 | 0.036231 | 1.5238 | 0.60765 |
| HCER200 | 0.19948 | 0.27571 | 1.2783 | 0.35425 | 0.97912 | 0.98823 | 1.0006 | 0.000888 | 0.39052 | 0.46906 | 0.85464 | -0.22662 |
| LPC203 | 0.11806 | 0.1857 | 0.80005 | -0.32184 | 0.29553 | 0.45483 | 0.85088 | -0.23297 | 0.31272 | 0.39607 | 0.86632 | -0.20703 |
| TAG544-FA160 | 0.000385 | 0.002144 | 0.43782 | -1.1916 | 0.016756 | 0.082636 | 0.56873 | -0.81418 | 0.004425 | 0.012363 | 0.47665 | -1.069 |
| PE180/203 | 0.71774 | 0.77745 | 0.92927 | -0.10583 | 0.028093 | 0.1122 | 0.69114 | -0.53295 | 0.26437 | 0.34421 | 0.83039 | -0.26813 |
| TAG546-FA204 | 0.018101 | 0.039945 | 0.61411 | -0.70342 | 0.14499 | 0.28517 | 0.72783 | -0.45832 | 0.35841 | 0.43694 | 0.84038 | -0.25089 |
| TAG568-FA161 | 0.022105 | 0.047027 | 0.38441 | -1.3793 | 0.005505 | 0.050778 | 0.27973 | -1.8379 | 0.028672 | 0.054737 | 0.40614 | -1.2999 |
| PEO-180/182 | 0.023305 | 0.049099 | 0.60261 | -0.7307 | 0.016592 | 0.082636 | 0.58713 | -0.76825 | 0.000965 | 0.004026 | 0.37104 | -1.4304 |
| PS180/180 | 0.031466 | 0.064116 | 1.4958 | 0.58094 | 0.9489 | 0.97403 | 0.96295 | -0.05446 | 0.39515 | 0.47287 | 1.1932 | 0.2548 |
| TAG561-FA160 | 1.92E-05 | 0.000481 | 0.15671 | -2.6738 | 0.13761 | 0.27522 | 0.66818 | -0.58169 | 7.06E-05 | 0.000696 | 0.18279 | -2.4517 |
| PEP-160/203 | 0.30708 | 0.39508 | 0.84161 | -0.24878 | 0.13832 | 0.27536 | 0.7899 | -0.34026 | 0.76302 | 0.80376 | 0.94649 | -0.07934 |
| TAG563-FA201 | 0.004212 | 0.011617 | 0.36943 | -1.4366 | 0.28987 | 0.45037 | 0.8385 | -0.25411 | 0.002015 | 0.006941 | 0.30546 | -1.711 |

| | | | | | | | | | | | | |
|---|---|---|---|---|---|---|---|---|---|---|---|---|
| PE181/224 | 0.038993 | 0.076494 | 0.75603 | -0.40348 | 0.001329 | 0.033246 | 0.53048 | -0.91462 | 0.026201 | 0.051311 | 0.62903 | -0.6688 |
| TAG521-FA180 | 5.29E-05 | 0.000718 | 0.27848 | -1.8443 | 0.30583 | 0.4669 | 0.84601 | -0.24125 | 4.99E-05 | 0.00056 | 0.30315 | -1.7219 |
| TAG547-FA205 | 0.029443 | 0.060464 | 0.52314 | -0.93474 | 0.011526 | 0.066967 | 0.48637 | -1.0399 | 0.14846 | 0.21241 | 0.69144 | -0.53233 |
| PG160/181_2 | 0.31983 | 0.40986 | 0.85899 | -0.21929 | 0.074925 | 0.19583 | 0.71854 | -0.47686 | 0.005001 | 0.013288 | 0.54168 | -0.88448 |
| PE170/182 | 0.002298 | 0.007673 | 0.57873 | -0.78903 | 0.002405 | 0.033393 | 0.52245 | -0.93663 | 0.000194 | 0.001319 | 0.27624 | -1.856 |
| PC200/203 | 0.034397 | 0.069453 | 0.71839 | -0.47716 | 0.001079 | 0.032039 | 0.58089 | -0.78366 | 0.02239 | 0.045267 | 0.71858 | -0.47678 |
| TAG547-FA204 | 0.00816 | 0.02059 | 0.5818 | -0.78141 | 0.010178 | 0.063932 | 0.54841 | -0.86666 | 0.01471 | 0.0315 | 0.55529 | -0.84869 |
| TAG526-FA204 | 0.01434 | 0.033222 | 0.57826 | -0.7902 | 0.32934 | 0.48293 | 0.88579 | -0.17496 | 0.28908 | 0.37119 | 0.84895 | -0.23625 |
| TAG542-FA181 | 0.015476 | 0.034981 | 0.49644 | -1.0103 | 0.60885 | 0.73335 | 1.0283 | 0.040329 | 0.00018 | 0.001271 | 0.33338 | -1.5848 |
| PI180/183 | 0.002446 | 0.008001 | 0.63662 | -0.65149 | 0.98891 | 0.99349 | 1.0244 | 0.034772 | 0.024066 | 0.047765 | 0.57768 | -0.79165 |
| TAG520-FA160 | 1.28E-05 | 0.000463 | 0.2416 | -2.0493 | 0.082154 | 0.20462 | 0.64212 | -0.63908 | 1.32E-05 | 0.000408 | 0.23076 | -2.1155 |
| TAG521-FA181 | 0.000204 | 0.001472 | 0.32194 | -1.6352 | 0.39596 | 0.54577 | 0.88691 | -0.17314 | 0.000112 | 0.000934 | 0.38854 | -1.3639 |
| TAG544-FA203 | 0.003797 | 0.010636 | 0.40114 | -1.3178 | 0.075166 | 0.19583 | 0.58563 | -0.77195 | 0.019854 | 0.040772 | 0.48496 | -1.0441 |
| PEO-180/225 | 0.2621 | 0.3454 | 1.2951 | 0.37304 | 0.95708 | 0.97505 | 0.95412 | -0.06775 | 0.24617 | 0.32374 | 1.3486 | 0.43148 |
| TAG546-FA225 | 0.000704 | 0.003377 | 0.36872 | -1.4394 | 0.017432 | 0.085325 | 0.46814 | -1.095 | 0.006816 | 0.017 | 0.42864 | -1.2222 |
| TAG542-FA202 | 0.000224 | 0.001533 | 0.44075 | -1.182 | 0.17476 | 0.31862 | 0.77643 | -0.36507 | 0.005442 | 0.014004 | 0.46713 | -1.0981 |
| TAG564-FA203 | 0.00701 | 0.017896 | 0.53045 | -0.9147 | 0.058777 | 0.16564 | 0.64398 | -0.6349 | 0.00857 | 0.020586 | 0.58149 | -0.78218 |
| PC160/202 | 0.10776 | 0.17152 | 0.79556 | -0.32995 | 0.86194 | 0.92442 | 1.0034 | 0.004856 | 0.64903 | 0.71493 | 0.95646 | -0.06423 |
| TAG563-FA180 | 0.000708 | 0.003377 | 0.39621 | -1.3357 | 0.025985 | 0.10844 | 0.58973 | -0.76187 | 0.000337 | 0.001858 | 0.32905 | -1.6036 |
| PE180/160 | 0.56131 | 0.64905 | 1.0505 | 0.071047 | 0.59411 | 0.72588 | 0.9061 | -0.14225 | 0.032159 | 0.060691 | 0.6222 | -0.68454 |
| FFA205 | 0.12338 | 0.19078 | 0.65386 | -0.61295 | 0.89286 | 0.94206 | 0.95236 | -0.07042 | 0.28787 | 0.37036 | 0.75631 | -0.40294 |
| PE180/201 | 0.3631 | 0.45721 | 0.84009 | -0.25138 | 0.016548 | 0.082636 | 0.63313 | -0.65942 | 0.001918 | 0.006677 | 0.48023 | -1.0582 |
| PC180/203 | 0.044183 | 0.084349 | 0.81976 | -0.28673 | 0.1274 | 0.26081 | 0.8539 | -0.22787 | 0.83004 | 0.86319 | 0.98208 | -0.02609 |
| PG180/180 | 0.12147 | 0.18919 | 0.72643 | -0.4611 | 0.036701 | 0.12919 | 0.61959 | -0.69062 | 0.008077 | 0.019767 | 0.5233 | -0.93429 |
| PEP-180/204 | 0.63819 | 0.71467 | 0.92107 | -0.11862 | 0.32145 | 0.48016 | 0.8458 | -0.24161 | 0.30882 | 0.3919 | 0.83992 | -0.25167 |
| PEO-180/160 | 0.84195 | 0.8812 | 0.97474 | -0.03691 | 0.36507 | 0.51282 | 0.88365 | -0.17845 | 0.020026 | 0.040869 | 0.71784 | -0.47827 |
| TAG541-FA201 | 5.51E-05 | 0.000718 | 0.25601 | -1.9657 | 0.10423 | 0.22694 | 0.7074 | -0.49941 | 0.000158 | 0.001153 | 0.30016 | -1.7362 |
| PC180/161 | 0.001133 | 0.004789 | 0.56897 | -0.81357 | 0.61348 | 0.73686 | 1.1568 | 0.2101 | 0.014305 | 0.030836 | 0.64137 | -0.64076 |
| TAG568-FA205 | 0.070292 | 0.1217 | 0.64003 | -0.64378 | 0.013218 | 0.071113 | 0.51028 | -0.97063 | 0.13354 | 0.19492 | 0.71672 | -0.48051 |
| PI180/204 | 0.14867 | 0.22047 | 0.85294 | -0.22948 | 0.3616 | 0.51063 | 1.0767 | 0.10667 | 0.70998 | 0.76396 | 1.0408 | 0.057658 |
| PI180/204_2 | 0.11921 | 0.187 | 0.84321 | -0.24603 | 0.24594 | 0.4033 | 1.0919 | 0.12685 | 0.96136 | 0.96432 | 1.0055 | 0.007913 |
| TAG546-FA205 | 0.031516 | 0.064116 | 0.57237 | -0.80497 | 0.042213 | 0.14021 | 0.61903 | -0.69193 | 0.45698 | 0.53219 | 0.91171 | -0.13335 |
| PC170/182 | 0.004457 | 0.01219 | 0.69781 | -0.51909 | 0.10025 | 0.22536 | 0.83139 | -0.2664 | 0.002191 | 0.007278 | 0.63542 | -0.65422 |

| | | | | | | | | | | | | |
|---|---|---|---|---|---|---|---|---|---|---|---|---|
| TAG542-FA201 | 0.000352 | 0.002045 | 0.37898 | -1.3998 | 0.40612 | 0.55543 | 0.88349 | -0.17872 | 0.001098 | 0.004374 | 0.42078 | -1.2489 |
| TAG521-FA160 | 0.000372 | 0.002124 | 0.33077 | -1.5961 | 0.64617 | 0.76206 | 1.0135 | 0.019339 | 0.00073 | 0.003253 | 0.36108 | -1.4696 |
| TAG482-FA120 | 0.76201 | 0.81343 | 2.0419 | 1.0299 | 0.61557 | 0.738 | 8.2809 | 3.0498 | 0.30805 | 0.39168 | 0.85462 | -0.22665 |
| TAG564-FA180 | 0.001855 | 0.006635 | 0.44978 | -1.1527 | 0.003251 | 0.037082 | 0.48874 | -1.0329 | 0.002818 | 0.009037 | 0.47154 | -1.0846 |
| LPC180 | 0.091654 | 0.15161 | 0.83161 | -0.26602 | 0.41372 | 0.56041 | 0.91249 | -0.13213 | 0.49999 | 0.56905 | 0.92584 | -0.11117 |
| PC181/205 | 0.21431 | 0.29372 | 0.65791 | -0.60404 | 0.15657 | 0.30155 | 0.61127 | -0.71011 | 0.4424 | 0.51706 | 0.77265 | -0.37212 |
| TAG567-FA205 | 0.080312 | 0.13615 | 0.68543 | -0.54493 | 0.030589 | 0.12054 | 0.60616 | -0.72224 | 0.18516 | 0.25513 | 0.74422 | -0.42619 |
| PE181/201 | 0.092401 | 0.15229 | 0.78617 | -0.34708 | 0.10504 | 0.22719 | 0.709 | -0.49615 | 0.004504 | 0.012425 | 0.44839 | -1.1572 |
| TAG501-FA161 | 4.87E-05 | 0.000718 | 0.28002 | -1.8364 | 0.73148 | 0.83311 | 1.1064 | 0.14592 | 2.08E-05 | 0.000411 | 0.26388 | -1.9221 |
| PG180/181 | 0.12683 | 0.19337 | 0.7181 | -0.47773 | 0.04194 | 0.14021 | 0.60979 | -0.71363 | 0.008831 | 0.021023 | 0.49473 | -1.0153 |
| PI180/205_2 | 0.23543 | 0.318 | 0.77154 | -0.37418 | 0.7291 | 0.83271 | 0.93803 | -0.09229 | 0.14927 | 0.21311 | 0.69873 | -0.51719 |
| TAG541-FA180 | 0.000446 | 0.002418 | 0.27092 | -1.8841 | 0.21246 | 0.36216 | 0.74914 | -0.41668 | 2.37E-05 | 0.00043 | 0.24029 | -2.0571 |
| PC200/205 | 0.25886 | 0.34182 | 0.72141 | -0.47111 | 0.18516 | 0.33206 | 0.66758 | -0.58299 | 0.050832 | 0.089168 | 0.50688 | -0.98028 |
| PC150/226 | 0.026786 | 0.05589 | 0.66338 | -0.5921 | 0.027255 | 0.11021 | 0.65803 | -0.60376 | 0.075471 | 0.12072 | 0.71667 | -0.48062 |
| PC200/181 | 0.016931 | 0.037619 | 0.67035 | -0.57701 | 0.031019 | 0.12092 | 0.73892 | -0.43652 | 0.001315 | 0.005065 | 0.511 | -0.9686 |
| TAG527-FA160 | 5.82E-06 | 0.000379 | 0.21273 | -2.2329 | 0.20029 | 0.35051 | 0.77979 | -0.35884 | 0.000123 | 0.000982 | 0.2903 | -1.7844 |
| LPC181 | 0.045206 | 0.086051 | 0.81 | -0.304 | 0.32822 | 0.48293 | 0.91657 | -0.12569 | 0.026661 | 0.051809 | 0.75574 | -0.40403 |
| PE181/181 | 0.97006 | 0.98061 | 1.0119 | 0.017083 | 0.33977 | 0.48963 | 0.89534 | -0.15948 | 0.047161 | 0.083862 | 0.60993 | -0.71329 |
| SMd181/120 | 0.38797 | 0.47886 | 0.80452 | -0.3138 | 0.79792 | 0.88462 | 1.2047 | 0.26863 | 0.3259 | 0.40836 | 2.2998 | 1.2015 |
| PE160/161 | 0.051078 | 0.093404 | 0.58618 | -0.77058 | 0.89742 | 0.94534 | 1.0215 | 0.030705 | 0.002131 | 0.007149 | 0.34628 | -1.53 |
| PEO-180/183 | 0.21164 | 0.29068 | 0.74909 | -0.41679 | 0.21018 | 0.36216 | 0.74535 | -0.424 | 0.001434 | 0.00546 | 0.34832 | -1.5215 |
| TAG568-FA182 | 0.012084 | 0.028606 | 0.54726 | -0.8697 | 0.002786 | 0.034877 | 0.43414 | -1.2038 | 0.066571 | 0.10976 | 0.63719 | -0.65021 |
| TAG521-FA201 | 3.2E-06 | 0.000231 | 0.18979 | -2.3975 | 0.1688 | 0.31297 | 0.77201 | -0.37331 | 0.000257 | 0.001561 | 0.44081 | -1.1818 |
| TAG543-FA203 | 0.00395 | 0.010942 | 0.51374 | -0.96088 | 0.1266 | 0.26081 | 0.71766 | -0.47864 | 0.008849 | 0.021023 | 0.56153 | -0.83256 |
| PE181/161 | 0.096311 | 0.15793 | 0.6908 | -0.53366 | 0.13723 | 0.27522 | 0.73094 | -0.45218 | 0.012724 | 0.028357 | 0.56339 | -0.82781 |
| TAG501-FA181 | 1.1E-05 | 0.000459 | 0.29732 | -1.7499 | 0.37745 | 0.52505 | 0.89117 | -0.16622 | 0.001653 | 0.005993 | 0.51553 | -0.95588 |
| TAG566-FA204 | 0.020811 | 0.044565 | 0.65538 | -0.60959 | 0.32948 | 0.48293 | 0.84585 | -0.24153 | 0.86036 | 0.88884 | 0.98625 | -0.01997 |
| LPC225 | 0.16302 | 0.23531 | 1.1876 | 0.24807 | 0.36552 | 0.51282 | 1.1441 | 0.19428 | 0.61899 | 0.68414 | 1.0775 | 0.10766 |
| PE160/183 | 0.23665 | 0.31896 | 1.2087 | 0.27342 | 0.43465 | 0.57606 | 0.82363 | -0.27994 | 0.006113 | 0.015485 | 0.3597 | -1.4751 |
| HCER160 | 0.99028 | 0.9918 | 0.98251 | -0.02545 | 0.1594 | 0.30521 | 0.78291 | -0.35308 | 0.039221 | 0.071322 | 0.70847 | -0.49722 |
| TAG547-FA225 | 0.001798 | 0.006606 | 0.43028 | -1.2167 | 0.00898 | 0.062074 | 0.40416 | -1.307 | 0.000607 | 0.002864 | 0.36906 | -1.4381 |
| TAG564-FA160 | 0.00261 | 0.00837 | 0.54201 | -0.8836 | 0.28922 | 0.45037 | 0.81717 | -0.29129 | 0.014904 | 0.031811 | 0.55708 | -0.84405 |
| PG160/202_2 | 0.59351 | 0.67827 | 1.1079 | 0.14789 | 0.22074 | 0.37286 | 0.77117 | -0.37488 | 0.06136 | 0.10295 | 0.62537 | -0.67721 |

| | | | | | | | | | | | | |
|---|---|---|---|---|---|---|---|---|---|---|---|---|
| TAG540-FA180 | 0.000103 | 0.000987 | 0.19666 | -2.3462 | 0.032548 | 0.12391 | 0.49772 | -1.0066 | 3.86E-06 | 0.000277 | 0.15181 | -2.7197 |
| PE180/161 | 0.56361 | 0.6494 | 0.92554 | -0.11164 | 0.75527 | 0.85066 | 1.1305 | 0.17691 | 0.017223 | 0.035937 | 0.46947 | -1.0909 |
| PE180/226 | 0.1314 | 0.19848 | 1.2866 | 0.36359 | 0.24056 | 0.39547 | 0.80978 | -0.3044 | 0.65464 | 0.71867 | 1.0804 | 0.11156 |
| TAG563-FA160 | 0.006178 | 0.016218 | 0.58171 | -0.78162 | 0.9497 | 0.97403 | 1.0828 | 0.11475 | 0.000601 | 0.002864 | 0.42176 | -1.2455 |
| SMd181/226 | 0.51282 | 0.6026 | 0.93014 | -0.10449 | 0.079093 | 0.1988 | 0.78759 | -0.34447 | 0.96024 | 0.96432 | 1.0038 | 0.005408 |
| PEO-180/205 | 0.40279 | 0.49382 | 0.76713 | -0.38247 | 0.22736 | 0.3805 | 0.6857 | -0.54436 | 0.095618 | 0.14646 | 0.58796 | -0.76621 |
| HCER220 | 0.34675 | 0.44003 | 1.1109 | 0.15169 | 0.32889 | 0.48293 | 0.88301 | -0.17949 | 0.0291 | 0.055392 | 0.73388 | -0.44638 |
| TAG545-FA204 | 0.026887 | 0.055921 | 0.61056 | -0.7118 | 0.50084 | 0.64182 | 0.92791 | -0.10795 | 0.73531 | 0.7809 | 1.0678 | 0.094622 |
| SMd181/220 | 0.16875 | 0.24039 | 0.89751 | -0.156 | 0.16868 | 0.31297 | 0.85755 | -0.22171 | 0.47268 | 0.54463 | 0.9445 | -0.08238 |
| TAG567-FA180 | 0.000964 | 0.004211 | 0.47432 | -1.0761 | 0.002577 | 0.034236 | 0.49194 | -1.0235 | 0.012216 | 0.027423 | 0.528 | -0.92139 |
| PC160/182 | 0.74163 | 0.79539 | 0.97488 | -0.03671 | 0.85579 | 0.92009 | 1.0088 | 0.012634 | 0.85285 | 0.88398 | 0.98505 | -0.02173 |
| TAG567-FA225 | 0.011447 | 0.027397 | 0.45629 | -1.132 | 0.001723 | 0.033246 | 0.34823 | -1.5219 | 0.060838 | 0.10243 | 0.55831 | -0.84085 |
| TAG545-FA160 | 0.001705 | 0.006308 | 0.4811 | -1.0556 | 0.036713 | 0.12919 | 0.60728 | -0.71956 | 0.073017 | 0.11824 | 0.66449 | -0.58969 |
| PEP-181/225 | 0.034593 | 0.069453 | 1.3254 | 0.4064 | 0.35594 | 0.50528 | 0.90291 | -0.14735 | 0.1183 | 0.17616 | 1.3233 | 0.40413 |
| PEP-182/182 | 0.12575 | 0.19307 | 0.69379 | -0.52744 | 0.09133 | 0.2162 | 0.67458 | -0.56794 | 0.047277 | 0.083862 | 0.60864 | -0.71633 |
| TAG546-FA226 | 0.98088 | 0.9839 | 0.92108 | -0.11861 | 0.76698 | 0.86087 | 0.90189 | -0.14898 | 0.54748 | 0.61769 | 1.2441 | 0.31511 |
| PE160/226 | 0.15598 | 0.22752 | 1.2822 | 0.35867 | 0.35035 | 0.50126 | 0.79702 | -0.32732 | 0.87419 | 0.89783 | 0.92623 | -0.11055 |
| FFA203 | 0.25548 | 0.33943 | 0.85463 | -0.22662 | 0.012861 | 0.070673 | 0.69743 | -0.51987 | 0.46793 | 0.54178 | 1.1357 | 0.18364 |
| LPE202 | 0.21054 | 0.28977 | 1.3223 | 0.4031 | 0.10004 | 0.22536 | 0.71799 | -0.47796 | 0.064496 | 0.10738 | 0.64271 | -0.63777 |
| SMd181/205 | 0.15622 | 0.22752 | 0.87239 | -0.19696 | 0.14406 | 0.28419 | 0.83096 | -0.26715 | 0.96085 | 0.96432 | 0.99474 | -0.00761 |
| TAG566-FA160 | 0.002047 | 0.007166 | 0.42973 | -1.2185 | 0.002196 | 0.033246 | 0.40523 | -1.3032 | 0.033741 | 0.063119 | 0.56346 | -0.82762 |
| PG160/202 | 0.66773 | 0.74053 | 1.0793 | 0.11003 | 0.26616 | 0.42678 | 0.8045 | -0.31383 | 0.051298 | 0.089532 | 0.62122 | -0.68683 |
| TAG547-FA226 | 0.011731 | 0.027974 | 0.42207 | -1.2444 | 0.10014 | 0.22536 | 0.57687 | -0.79369 | 0.037296 | 0.068721 | 0.46503 | -1.1046 |
| SM241 | 0.68911 | 0.75524 | 1.0307 | 0.043694 | 0.85649 | 0.92009 | 0.98932 | -0.0155 | 0.088355 | 0.13695 | 1.1401 | 0.1892 |
| PEP-181/205 | 0.78961 | 0.83583 | 0.90598 | -0.14245 | 0.22385 | 0.37655 | 0.68508 | -0.54565 | 0.19464 | 0.2662 | 0.6646 | -0.58944 |
| HCER241 | 0.53589 | 0.62521 | 1.0817 | 0.1133 | 0.88196 | 0.9371 | 0.96297 | -0.05444 | 0.35944 | 0.43738 | 0.85192 | -0.23121 |
| FFA226 | 0.31397 | 0.40315 | 1.1842 | 0.24393 | 0.28044 | 0.44421 | 0.81084 | -0.30251 | 0.11769 | 0.17612 | 1.4971 | 0.58218 |
| FFA220 | 0.42261 | 0.51328 | 0.7804 | -0.35771 | 0.7713 | 0.86423 | 0.90735 | -0.14026 | 0.72447 | 0.77571 | 1.1482 | 0.19933 |
| PEP-180/160 | 0.37981 | 0.47096 | 0.87373 | -0.19474 | 0.11819 | 0.25062 | 0.79273 | -0.3351 | 0.56692 | 0.63852 | 0.91493 | -0.12826 |
| PE160/201 | 0.92641 | 0.95427 | 0.97163 | -0.04153 | 0.35757 | 0.50604 | 0.81615 | -0.29309 | 0.01188 | 0.026853 | 0.51672 | -0.95255 |
| TAG565-FA180 | 0.001265 | 0.005081 | 0.48976 | -1.0298 | 0.005809 | 0.051807 | 0.56916 | -0.81308 | 0.070443 | 0.11465 | 0.70813 | -0.49792 |
| PG161/182_2 | 0.70881 | 0.77293 | 1.025 | 0.035611 | 0.95313 | 0.97408 | 0.9816 | -0.0268 | 0.50326 | 0.57077 | 0.86649 | -0.20675 |
| TAG566-FA181 | 0.003316 | 0.009769 | 0.38665 | -1.3709 | 0.00766 | 0.057982 | 0.41671 | -1.2629 | 0.011788 | 0.026833 | 0.46905 | -1.0922 |

| | | | | | | | | | | | | |
|---|---|---|---|---|---|---|---|---|---|---|---|---|
| PC140/225 | 0.68669 | 0.75385 | 0.92847 | -0.10708 | 0.41407 | 0.56041 | 1.1297 | 0.1759 | 0.74786 | 0.79164 | 0.95259 | -0.07008 |
| PC180/181 | 0.01357 | 0.031663 | 0.73322 | -0.44769 | 0.70052 | 0.80557 | 0.95419 | -0.06765 | 0.050953 | 0.089168 | 0.77124 | -0.37475 |
| TAG501-FA160 | 1.5E-05 | 0.000481 | 0.30874 | -1.6956 | 0.63233 | 0.75393 | 0.99784 | -0.00312 | 0.000414 | 0.002173 | 0.52199 | -0.9379 |
| PC160/224 | 0.87306 | 0.90647 | 1.036 | 0.050958 | 0.87599 | 0.93487 | 1.0427 | 0.060344 | 0.84958 | 0.8821 | 1.0235 | 0.033536 |
| TAG481-FA161 | 2.48E-05 | 0.000496 | 0.31731 | -1.656 | 0.66919 | 0.78073 | 1.5675 | 0.64846 | 0.000449 | 0.00232 | 0.41252 | -1.2775 |
| PEP-181/183 | 0.60794 | 0.69191 | 0.85793 | -0.22106 | 0.65844 | 0.77094 | 0.88183 | -0.18143 | 0.33868 | 0.41837 | 0.76966 | -0.3777 |
| PEO-160/203 | 0.49286 | 0.58337 | 0.94249 | -0.08546 | 0.022975 | 0.10099 | 0.77664 | -0.36468 | 0.059434 | 0.10076 | 0.77813 | -0.36191 |
| PC181/201 | 0.009588 | 0.023466 | 0.66655 | -0.58522 | 0.10202 | 0.22536 | 0.76737 | -0.382 | 0.017185 | 0.035937 | 0.64125 | -0.64104 |
| TAG548-FA204 | 0.010302 | 0.024932 | 0.58974 | -0.76184 | 0.010312 | 0.063932 | 0.60347 | -0.72864 | 0.22799 | 0.30539 | 0.81815 | -0.28957 |
| PEP-181/203 | 0.73027 | 0.7884 | 0.96881 | -0.04571 | 0.055163 | 0.1582 | 0.79573 | -0.32965 | 0.37771 | 0.45535 | 0.90416 | -0.14536 |
| PE180/180 | 0.25491 | 0.33935 | 1.2099 | 0.27491 | 0.42912 | 0.57245 | 0.86858 | -0.20327 | 0.068586 | 0.11247 | 1.4476 | 0.53369 |
| TAG565-FA160 | 0.000301 | 0.001848 | 0.47382 | -1.0776 | 0.021233 | 0.097504 | 0.6155 | -0.70016 | 0.008504 | 0.020504 | 0.54616 | -0.8726 |
| SMd181/222 | 0.47479 | 0.56714 | 1.104 | 0.14276 | 0.65654 | 0.77094 | 0.93782 | -0.09261 | 0.42787 | 0.50369 | 1.0951 | 0.13109 |
| PC181/204 | 0.014728 | 0.033754 | 0.80014 | -0.32168 | 0.000675 | 0.032039 | 0.73065 | -0.45275 | 0.25465 | 0.33222 | 0.89106 | -0.1664 |
| TAG481-FA180 | 0.66043 | 0.73493 | 1.3776 | 0.46211 | 0.42876 | 0.57245 | 5.5882 | 2.4824 | 0.06029 | 0.10195 | 0.59581 | -0.74706 |
| PC181/181 | 0.034655 | 0.069453 | 0.71964 | -0.47465 | 0.28616 | 0.44888 | 0.83763 | -0.25562 | 0.076738 | 0.12185 | 0.71311 | -0.48781 |
| HCER140 | 0.43934 | 0.53161 | 0.85787 | -0.22117 | 0.12732 | 0.26081 | 0.72535 | -0.46325 | 0.1419 | 0.20528 | 0.72103 | -0.47188 |
| TAG542-FA180 | 0.015693 | 0.035227 | 0.47331 | -1.0792 | 0.42482 | 0.5712 | 0.89479 | -0.16038 | 0.001231 | 0.004768 | 0.37194 | -1.4269 |
| TAG545-FA225 | 0.041208 | 0.079367 | 0.67741 | -0.56189 | 0.12059 | 0.25324 | 0.73123 | -0.4516 | 0.79307 | 0.83004 | 0.96446 | -0.05221 |
| PE170/225 | 0.71556 | 0.77638 | 0.9411 | -0.08759 | 0.00596 | 0.052432 | 0.59639 | -0.74567 | 0.000682 | 0.003093 | 0.451 | -1.1488 |
| PEO-180/203 | 0.10653 | 0.16997 | 0.82057 | -0.28531 | 0.007987 | 0.058424 | 0.67952 | -0.55742 | 0.009974 | 0.023357 | 0.63519 | -0.65474 |
| PA180/181 | 0.56305 | 0.6494 | 0.87676 | -0.18975 | 0.9008 | 0.94655 | 0.94934 | -0.075 | 0.85411 | 0.88398 | 0.95708 | -0.06329 |
| PEP-160/202 | 0.091758 | 0.15161 | 0.75405 | -0.40727 | 0.33965 | 0.48963 | 0.8581 | -0.22078 | 0.11852 | 0.17616 | 0.76651 | -0.38363 |
| SMd181/224 | 0.55948 | 0.64808 | 1.0625 | 0.087522 | 0.10053 | 0.22536 | 0.81398 | -0.29694 | 0.46935 | 0.54178 | 1.1132 | 0.15474 |
| FFA183 | 0.067882 | 0.11879 | 0.66594 | -0.58653 | 0.011892 | 0.067908 | 0.51063 | -0.96964 | 0.77212 | 0.81161 | 1.0826 | 0.11456 |
| TAG569-FA226 | 0.012963 | 0.030466 | 0.40096 | -1.3185 | 0.001072 | 0.032039 | 0.27308 | -1.8726 | 0.007558 | 0.018636 | 0.34983 | -1.5153 |
| SMd181/184 | 0.047482 | 0.088064 | 0.72225 | -0.46944 | 0.014371 | 0.074845 | 0.66417 | -0.59038 | 0.001773 | 0.006273 | 0.59396 | -0.75157 |
| PE180/183 | 0.10615 | 0.16978 | 1.2474 | 0.31896 | 0.39828 | 0.547 | 0.86752 | -0.20503 | 0.004474 | 0.012425 | 0.40932 | -1.2887 |
| HCER240 | 0.50707 | 0.59802 | 1.0664 | 0.092777 | 0.41248 | 0.56041 | 0.88365 | -0.17845 | 0.023029 | 0.046098 | 0.68135 | -0.55353 |
| PEP-180/161 | 0.12841 | 0.19518 | 0.68172 | -0.55276 | 0.1826 | 0.3295 | 0.72208 | -0.46978 | 0.020912 | 0.04241 | 0.5493 | -0.86433 |
| PEP-180/201 | 0.18916 | 0.26426 | 0.74385 | -0.42692 | 0.073754 | 0.19518 | 0.66411 | -0.5905 | 0.015471 | 0.032808 | 0.56691 | -0.81882 |
| PEP-160/226 | 0.34503 | 0.43956 | 1.1739 | 0.23136 | 0.52885 | 0.66941 | 0.88077 | -0.18316 | 0.080459 | 0.12652 | 1.3865 | 0.47145 |
| SMd181/181 | 0.70325 | 0.76815 | 1.0218 | 0.03106 | 0.21239 | 0.36216 | 0.88704 | -0.17292 | 0.02749 | 0.053261 | 1.2475 | 0.31909 |

| | | | | | | | | | | | | |
|---|---|---|---|---|---|---|---|---|---|---|---|---|
| LPC205 | 0.84051 | 0.8812 | 0.99862 | -0.00199 | 0.57112 | 0.70818 | 0.89437 | -0.16105 | 0.9978 | 0.9978 | 1.1047 | 0.14365 |
| PEP-160/204 | 0.62964 | 0.70916 | 1.0801 | 0.11112 | 0.33996 | 0.48963 | 0.84058 | -0.25055 | 0.46158 | 0.53658 | 1.1359 | 0.18378 |
| PE170/202 | 0.004239 | 0.011643 | 0.60973 | -0.71376 | 0.005424 | 0.050778 | 0.58046 | -0.78474 | 0.085253 | 0.13278 | 0.7106 | -0.49288 |
| HCER260 | 0.16514 | 0.23785 | 0.78555 | -0.34823 | 0.37196 | 0.51962 | 0.85574 | -0.22476 | 0.017139 | 0.035937 | 0.66243 | -0.59417 |
| PE160/160 | 0.34667 | 0.44003 | 1.1848 | 0.24465 | 0.62276 | 0.74525 | 1.11 | 0.15054 | 0.81965 | 0.85375 | 0.99828 | -0.00249 |
| TAG566-FA205 | 0.040688 | 0.078598 | 0.64823 | -0.62543 | 0.041223 | 0.13979 | 0.67853 | -0.55953 | 0.091285 | 0.14049 | 0.70051 | -0.51352 |
| TAG567-FA160 | 0.006338 | 0.016504 | 0.40077 | -1.3192 | 0.003754 | 0.038795 | 0.34774 | -1.5239 | 0.027747 | 0.053443 | 0.46802 | -1.0954 |
| PEP-180/225 | 0.76847 | 0.81744 | 1.0306 | 0.043498 | 0.33634 | 0.48766 | 0.86338 | -0.21193 | 0.67748 | 0.73876 | 1.0733 | 0.10205 |
| PE180/204 | 0.075686 | 0.12966 | 1.2327 | 0.30184 | 0.044004 | 0.14157 | 0.77678 | -0.36443 | 0.67232 | 0.7356 | 1.0497 | 0.069975 |
| PC160/181 | 0.089843 | 0.14958 | 0.88294 | -0.17961 | 0.047186 | 0.14422 | 1.1599 | 0.21405 | 0.74918 | 0.79175 | 0.97533 | -0.03604 |
| LPE200 | 0.12494 | 0.19228 | 0.80218 | -0.318 | 0.026949 | 0.10965 | 0.76389 | -0.38856 | 0.000323 | 0.001795 | 0.44869 | -1.1562 |
| PC180/226 | 0.55322 | 0.64197 | 0.92321 | -0.11528 | 0.59636 | 0.72673 | 1.054 | 0.075843 | 0.21427 | 0.29 | 1.2052 | 0.26933 |
| SMd181/180 | 0.2893 | 0.37591 | 0.91433 | -0.12921 | 0.18759 | 0.33458 | 0.87901 | -0.18605 | 0.77647 | 0.81398 | 0.98171 | -0.02663 |
| PS160/180_2 | 0.052917 | 0.095687 | 1.4847 | 0.57015 | 0.63899 | 0.75633 | 1.0324 | 0.045935 | 0.48407 | 0.55579 | 1.0922 | 0.12718 |
| HCER181 | 0.41486 | 0.50555 | 0.86939 | -0.20192 | 0.030736 | 0.12054 | 0.69737 | -0.51999 | 0.24534 | 0.32374 | 0.82774 | -0.27276 |
| TAG566-FA225 | 0.0035 | 0.010037 | 0.47133 | -1.0852 | 0.002093 | 0.033246 | 0.43165 | -1.2121 | 0.063817 | 0.10653 | 0.61842 | -0.69335 |
| PC150/225 | 0.003807 | 0.010636 | 0.64545 | -0.63163 | 0.007814 | 0.058424 | 0.61982 | -0.69007 | 0.004849 | 0.013098 | 0.64055 | -0.64261 |
| TAG546-FA160 | 0.018816 | 0.041104 | 0.61081 | -0.7112 | 0.069352 | 0.18656 | 0.63532 | -0.65444 | 0.508 | 0.57515 | 0.87795 | -0.18779 |
| TAG567-FA204 | 0.040511 | 0.078489 | 0.71106 | -0.49195 | 0.08813 | 0.21245 | 0.7167 | -0.48057 | 0.35555 | 0.43509 | 0.85832 | -0.22042 |
| LPC182 | 0.30058 | 0.38748 | 0.87076 | -0.19966 | 0.63485 | 0.75417 | 0.93673 | -0.09429 | 0.013606 | 0.029824 | 0.70709 | -0.50003 |
| PC181/224 | 0.15658 | 0.22753 | 0.79527 | -0.33048 | 0.085438 | 0.20831 | 0.76768 | -0.38142 | 0.32525 | 0.40836 | 0.85552 | -0.22513 |
| PEO-180/226 | 0.44322 | 0.53464 | 1.1617 | 0.2162 | 0.26238 | 0.42369 | 0.8329 | -0.26379 | 0.24558 | 0.32374 | 1.278 | 0.35387 |
| SMd181/221 | 0.54536 | 0.63501 | 1.0429 | 0.060633 | 0.6368 | 0.75511 | 0.95407 | -0.06783 | 0.13071 | 0.19223 | 1.1668 | 0.22251 |
| SM240 | 0.57095 | 0.6567 | 0.93763 | -0.09292 | 0.39131 | 0.54086 | 0.89291 | -0.16342 | 0.27915 | 0.36057 | 0.89179 | -0.16522 |
| DAG161/182 | 0.12229 | 0.19 | 0.43643 | -1.1962 | 0.29516 | 0.45483 | 0.51594 | -0.95473 | 0.40394 | 0.48251 | 1.0303 | 0.043026 |
| PEO-180/181 | 0.045774 | 0.086731 | 0.67022 | -0.57729 | 0.074926 | 0.19583 | 0.69501 | -0.5249 | 0.000233 | 0.00148 | 0.40671 | -1.2979 |
| PC170/224 | 0.05759 | 0.10282 | 0.73772 | -0.43885 | 0.10203 | 0.22536 | 0.81935 | -0.28744 | 0.033172 | 0.062413 | 0.75022 | -0.41461 |
| TAG520-FA180 | 2.26E-05 | 0.000496 | 0.22378 | -2.1599 | 0.067682 | 0.18398 | 0.62846 | -0.6701 | 1.58E-05 | 0.000408 | 0.2516 | -1.9908 |
| TAG565-FA204 | 0.010429 | 0.025146 | 0.60095 | -0.73469 | 0.035298 | 0.12706 | 0.69978 | -0.51503 | 0.22757 | 0.30539 | 0.82524 | -0.27712 |
| PE160/202 | 0.073076 | 0.12619 | 1.5685 | 0.64939 | 0.17099 | 0.31356 | 0.70504 | -0.50422 | 0.088718 | 0.13719 | 0.61615 | -0.69864 |
| PC140/204 | 0.50901 | 0.59921 | 0.88344 | -0.1788 | 0.72553 | 0.83009 | 1.1331 | 0.18032 | 0.39474 | 0.47287 | 0.87199 | -0.19762 |
| PEP-181/181 | 0.15448 | 0.22701 | 0.78083 | -0.35691 | 0.099148 | 0.22536 | 0.74802 | -0.41885 | 0.039616 | 0.071653 | 0.70146 | -0.51157 |
| TAG568-FA160 | 0.009094 | 0.022425 | 0.44931 | -1.1542 | 0.003472 | 0.037082 | 0.379 | -1.3997 | 0.023644 | 0.047072 | 0.47507 | -1.0738 |

| | | | | | | | | | | | | |
|---|---|---|---|---|---|---|---|---|---|---|---|---|
| FFA120 | 0.73993 | 0.79487 | 1.4595 | 0.54544 | 0.82352 | 0.89952 | 1.4533 | 0.53933 | 0.056116 | 0.096728 | 2.9922 | 1.5812 |
| DAG140/160 | 0.25642 | 0.33997 | 0.5971 | -0.74395 | 0.033938 | 0.12553 | 2.4108 | 1.2695 | 0.098399 | 0.1494 | 1.8576 | 0.8934 |
| TAG567-FA181 | 0.027709 | 0.057447 | 0.62228 | -0.68436 | 0.003954 | 0.039596 | 0.50238 | -0.99314 | 0.084285 | 0.13158 | 0.67728 | -0.56217 |
| CER160 | 0.7149 | 0.77638 | 1.0159 | 0.02273 | 0.90701 | 0.94655 | 0.95914 | -0.06019 | 0.75143 | 0.79284 | 1.2868 | 0.36373 |
| TAG482-FA180 | 0.6344 | 0.71206 | 2.8149 | 1.4931 | 0.7527 | 0.84923 | 6.0668 | 2.6009 | 0.11888 | 0.17629 | 0.66107 | -0.59712 |
| LPC221 | 0.10069 | 0.16346 | 0.74158 | -0.43133 | 0.31634 | 0.47671 | 0.88469 | -0.17676 | 0.013439 | 0.029556 | 0.65643 | -0.60729 |
| PEP-181/161 | 0.97632 | 0.9818 | 1.022 | 0.031442 | 0.33449 | 0.48706 | 0.87598 | -0.19103 | 0.79534 | 0.83108 | 0.98004 | -0.02908 |
| PC160/180 | 0.09659 | 0.15799 | 0.86646 | -0.2068 | 0.9886 | 0.99349 | 0.99159 | -0.01219 | 0.93262 | 0.9454 | 1.0149 | 0.021309 |
| PC170/225 | 0.039011 | 0.076494 | 0.771 | -0.37519 | 0.038738 | 0.13542 | 0.74662 | -0.42156 | 0.007031 | 0.017404 | 0.64154 | -0.64039 |
| FFA225 | 0.59946 | 0.68345 | 1.0999 | 0.13736 | 0.33518 | 0.48706 | 0.77712 | -0.36379 | 0.06543 | 0.10866 | 1.7561 | 0.81234 |
| FFA202 | 0.53229 | 0.62212 | 0.89504 | -0.15998 | 0.10335 | 0.22578 | 0.73754 | -0.43921 | 0.096764 | 0.14787 | 1.4603 | 0.54623 |
| PEP-181/160 | 0.75616 | 0.80964 | 1.0581 | 0.081421 | 0.60183 | 0.72959 | 0.9138 | -0.13005 | 0.69116 | 0.74866 | 1.0809 | 0.11218 |
| PEP-160/224 | 0.80432 | 0.84727 | 1.0556 | 0.078073 | 0.34445 | 0.49501 | 0.79974 | -0.3224 | 0.67662 | 0.73876 | 1.0596 | 0.083525 |
| PEO-160/226 | 0.66748 | 0.74053 | 1.0615 | 0.086158 | 0.28117 | 0.44427 | 0.86404 | -0.21084 | 0.055565 | 0.096205 | 1.3992 | 0.48456 |
| PC160/161 | 0.019138 | 0.041668 | 0.69696 | -0.52085 | 0.053764 | 0.15611 | 1.3066 | 0.38579 | 0.16999 | 0.23901 | 0.85194 | -0.23118 |
| SMd181/141 | 0.014771 | 0.033754 | 0.69576 | -0.52333 | 0.12637 | 0.26081 | 0.78239 | -0.35404 | 0.46937 | 0.54178 | 1.1423 | 0.19191 |
| SMd181/170 | 0.58571 | 0.67248 | 0.94027 | -0.08885 | 0.26294 | 0.42369 | 0.85948 | -0.21847 | 0.90804 | 0.92655 | 0.9984 | -0.00231 |
| PEO-180/204 | 0.36627 | 0.45943 | 0.8832 | -0.17919 | 0.090277 | 0.21449 | 0.77526 | -0.36725 | 0.066601 | 0.10976 | 0.74036 | -0.4337 |
| FFA140 | 0.22525 | 0.30742 | 0.80232 | -0.31775 | 0.7347 | 0.83471 | 0.95541 | -0.06581 | 0.014416 | 0.030973 | 1.9006 | 0.92642 |
| SMd181/140 | 0.14653 | 0.21778 | 0.77956 | -0.35927 | 0.31732 | 0.47708 | 0.85126 | -0.23233 | 0.58334 | 0.6525 | 1.1343 | 0.18177 |
| CER180 | 0.28691 | 0.37356 | 1.3588 | 0.44233 | 0.87393 | 0.9342 | 0.96782 | -0.04719 | 0.2747 | 0.35624 | 1.6761 | 0.74509 |
| TAG522-FA202 | 1.56E-07 | 5.1E-05 | 0.31514 | -1.6659 | 0.045014 | 0.14157 | 0.70322 | -0.50796 | 0.002125 | 0.007149 | 0.54249 | -0.88232 |
| PE160/140 | 0.88833 | 0.92087 | 1.4137 | 0.49946 | 0.64304 | 0.75975 | 1.9686 | 0.97716 | 0.96508 | 0.96656 | 1.0413 | 0.058436 |
| PEP-160/181 | 0.12026 | 0.1882 | 0.72769 | -0.45861 | 0.29345 | 0.45483 | 0.81079 | -0.3026 | 0.18491 | 0.25513 | 0.76914 | -0.37869 |
| TAG504-FA141 | 0.009993 | 0.024275 | 0.53094 | -0.91338 | 0.022189 | 0.099535 | 0.50492 | -0.98588 | 0.058252 | 0.099795 | 0.66546 | -0.58757 |
| FFA204 | 0.014777 | 0.033754 | 0.68284 | -0.55039 | 0.0201 | 0.096066 | 0.67136 | -0.57485 | 0.58318 | 0.6525 | 0.9318 | -0.10191 |
| CE182 | 0.1408 | 0.21121 | 0.70571 | -0.50285 | 0.014304 | 0.074845 | 0.55691 | -0.84448 | 0.59295 | 0.65875 | 0.88685 | -0.17323 |
| PEP-182/226 | 0.082795 | 0.14 | 1.2991 | 0.37751 | 0.50873 | 0.64964 | 0.90309 | -0.14706 | 0.13138 | 0.19223 | 1.282 | 0.35843 |
| CER221 | 0.073924 | 0.12719 | 0.70526 | -0.50377 | 0.39075 | 0.54086 | 0.85639 | -0.22367 | 0.63207 | 0.69742 | 0.96236 | -0.05536 |
| TAG500-FA160 | 6.71E-07 | 0.000134 | 0.22737 | -2.1369 | 0.089174 | 0.21343 | 0.68172 | -0.55276 | 4.29E-05 | 0.000506 | 0.38248 | -1.3865 |
| TAG523-FA203 | 5.46E-05 | 0.000718 | 0.44442 | -1.17 | 0.18744 | 0.33458 | 0.78891 | -0.34207 | 0.31339 | 0.39615 | 0.85033 | -0.2339 |
| PA180/203 | 0.24048 | 0.32345 | 0.90438 | -0.145 | 0.49244 | 0.63607 | 0.94538 | -0.08103 | 0.372 | 0.45013 | 0.92515 | -0.11224 |
| CER260 | 0.37801 | 0.46963 | 0.85284 | -0.22965 | 0.19824 | 0.34786 | 0.78209 | -0.3546 | 0.033383 | 0.06263 | 0.60592 | -0.72279 |

| | | | | | | | | | | | | |
|---|---|---|---|---|---|---|---|---|---|---|---|---|
| TAG526-FA226 | 0.046692 | 0.087217 | 0.63066 | -0.66506 | 0.70163 | 0.80557 | 1.3162 | 0.39636 | 0.0041 | 0.011876 | 0.44482 | -1.1687 |
| TAG566-FA180 | 0.002555 | 0.008283 | 0.58898 | -0.76372 | 0.002462 | 0.033393 | 0.56888 | -0.81381 | 0.077735 | 0.12253 | 0.79747 | -0.3265 |
| LPE203 | 0.95146 | 0.9648 | 0.96517 | -0.05115 | 0.14942 | 0.29123 | 0.75883 | -0.39816 | 0.097424 | 0.14853 | 0.73362 | -0.4469 |
| PE160/182 | 0.84781 | 0.88591 | 1.0212 | 0.030203 | 0.18406 | 0.331 | 0.80785 | -0.30783 | 0.014029 | 0.030545 | 0.57802 | -0.79081 |
| TAG500-FA180 | 1.57E-07 | 5.1E-05 | 0.20156 | -2.3107 | 0.032827 | 0.12425 | 0.56021 | -0.83597 | 3.06E-06 | 0.000277 | 0.3097 | -1.691 |
| PE160/205 | 0.73148 | 0.7884 | 1.1205 | 0.16414 | 0.47953 | 0.6231 | 0.77826 | -0.36167 | 0.35111 | 0.43127 | 0.73794 | -0.43842 |
| PEP-181/224 | 0.3923 | 0.48187 | 1.1674 | 0.22325 | 0.076297 | 0.19691 | 0.73519 | -0.44381 | 0.41388 | 0.49257 | 1.1331 | 0.18031 |
| SM261 | 0.40973 | 0.50044 | 0.88663 | -0.17359 | 0.42555 | 0.5712 | 0.88375 | -0.17829 | 0.58979 | 0.65745 | 0.92439 | -0.11342 |
| PC160/140 | 0.48729 | 0.57994 | 0.8744 | -0.19364 | 0.26844 | 0.42937 | 1.6668 | 0.7371 | 0.47413 | 0.54533 | 0.90349 | -0.14642 |
| PC170/205 | 0.81831 | 0.86062 | 0.97057 | -0.0431 | 0.41115 | 0.55996 | 0.75216 | -0.41089 | 0.36187 | 0.43869 | 0.72772 | -0.45854 |
| HCER221 | 0.076755 | 0.13115 | 0.77149 | -0.37428 | 0.015605 | 0.079992 | 0.69059 | -0.53409 | 0.024265 | 0.048014 | 0.69622 | -0.52238 |
| SMd181/201 | 0.84065 | 0.8812 | 0.97831 | -0.03164 | 0.45599 | 0.59849 | 0.92405 | -0.11396 | 0.056959 | 0.097837 | 1.1881 | 0.24871 |
| LPE240 | 0.45584 | 0.54651 | 0.87236 | -0.19701 | 0.68935 | 0.79569 | 0.91577 | -0.12694 | 0.045195 | 0.080608 | 0.65767 | -0.60457 |
| HCER261 | 0.36723 | 0.45974 | 0.80336 | -0.31589 | 0.17355 | 0.31737 | 0.7183 | -0.47734 | 0.14488 | 0.20867 | 0.70784 | -0.49851 |
| PC180/225 | 0.59172 | 0.67819 | 0.92857 | -0.10691 | 0.32356 | 0.48159 | 1.1195 | 0.16283 | 0.50211 | 0.57046 | 1.0921 | 0.12715 |
| SMd181/223 | 0.93423 | 0.95876 | 1.0141 | 0.020202 | 0.2637 | 0.42387 | 0.83853 | -0.25406 | 0.81454 | 0.84979 | 0.96354 | -0.05358 |
| SMd181/225 | 0.2488 | 0.33326 | 0.8088 | -0.30615 | 0.30804 | 0.46744 | 0.81508 | -0.29498 | 0.42408 | 0.50014 | 0.86262 | -0.2132 |
| PE160/204 | 0.29282 | 0.37974 | 1.2025 | 0.26609 | 0.35245 | 0.50206 | 0.8602 | -0.21725 | 0.43017 | 0.50549 | 0.87972 | -0.18489 |
| TAG545-FA205 | 0.016724 | 0.037285 | 0.53384 | -0.90553 | 0.003345 | 0.037082 | 0.52371 | -0.93317 | 0.27343 | 0.35529 | 0.81338 | -0.29799 |
| PEO-160/204 | 0.9724 | 0.98145 | 0.98653 | -0.01957 | 0.55012 | 0.68925 | 0.91932 | -0.12136 | 0.73506 | 0.7809 | 0.97142 | -0.04183 |
| PEO-160/183 | 0.85057 | 0.88685 | 1.0022 | 0.003123 | 0.30219 | 0.46289 | 0.79673 | -0.32783 | 0.041796 | 0.075164 | 0.5843 | -0.77523 |
| DAG160/181 | 0.083527 | 0.14051 | 0.60875 | -0.71607 | 0.20499 | 0.3559 | 0.64379 | -0.63533 | 0.093203 | 0.1431 | 0.59788 | -0.74207 |
| TAG566-FA226 | 0.019938 | 0.043121 | 0.62043 | -0.68866 | 0.009562 | 0.06288 | 0.53851 | -0.89295 | 0.16993 | 0.23901 | 0.73662 | -0.441 |
| PC160/183 | 0.45389 | 0.54517 | 0.88396 | -0.17795 | 0.71974 | 0.82491 | 1.116 | 0.15835 | 0.1777 | 0.24772 | 0.7349 | -0.44439 |
| PC180/182 | 0.17017 | 0.24136 | 0.88301 | -0.1795 | 0.65801 | 0.77094 | 0.95555 | -0.0656 | 0.42213 | 0.49874 | 0.92202 | -0.11712 |
| FFA161 | 0.32299 | 0.4131 | 0.70772 | -0.49875 | 0.56901 | 0.70818 | 0.81465 | -0.29574 | 0.026282 | 0.051311 | 1.9088 | 0.93269 |
| FFA241 | 0.97526 | 0.9818 | 0.98986 | -0.0147 | 0.90593 | 0.94655 | 1.0061 | 0.008754 | 0.68059 | 0.73967 | 0.96324 | -0.05403 |
| TAG483-FA141 | 0.00064 | 0.003133 | 0.46963 | -1.0904 | 0.28978 | 0.45037 | 0.87388 | -0.19449 | 0.019989 | 0.040869 | 0.54674 | -0.87106 |
| PEP-181/226 | 0.066733 | 0.11743 | 1.2232 | 0.29069 | 0.23451 | 0.38846 | 0.87558 | -0.19169 | 0.067679 | 0.11126 | 1.2905 | 0.3679 |
| PEP-160/180 | 0.1303 | 0.19727 | 0.7155 | -0.48298 | 0.33011 | 0.48293 | 0.80191 | -0.31849 | 0.35227 | 0.43188 | 0.81064 | -0.30287 |
| FFA201 | 0.35576 | 0.44884 | 0.90027 | -0.15157 | 0.34649 | 0.49684 | 0.89066 | -0.16706 | 0.028401 | 0.054541 | 1.5137 | 0.59809 |
| FFA141 | 0.76221 | 0.81343 | 0.92572 | -0.11135 | 0.88017 | 0.9371 | 0.93462 | -0.09754 | 0.003793 | 0.011173 | 2.4323 | 1.2823 |
| PC200/226 | 0.14911 | 0.22061 | 0.77209 | -0.37316 | 0.065191 | 0.17907 | 0.68109 | -0.55408 | 0.49712 | 0.56782 | 0.89452 | -0.16082 |

| | | | | | | | | | | | | |
|---|---|---|---|---|---|---|---|---|---|---|---|---|
| PEP-140/181 | 0.61725 | 0.70005 | 0.86321 | -0.21221 | 0.077352 | 0.19691 | 0.64778 | -0.62643 | 0.32544 | 0.40836 | 0.77727 | -0.36352 |
| LPE170 | 0.99245 | 0.99245 | 1.0175 | 0.024959 | 0.10775 | 0.2315 | 0.78015 | -0.35817 | 0.000684 | 0.003093 | 0.50557 | -0.98401 |
| FFA240 | 0.24675 | 0.3312 | 0.79801 | -0.32552 | 0.28508 | 0.44828 | 0.80626 | -0.31068 | 0.1943 | 0.2662 | 1.2691 | 0.34384 |
| Cholesterol | 0.091131 | 0.15134 | 0.69967 | -0.51524 | 0.046118 | 0.14229 | 0.63267 | -0.66048 | 0.65312 | 0.71822 | 0.91228 | -0.13244 |
| FFA160 | 0.12101 | 0.18891 | 0.89718 | -0.15654 | 0.054716 | 0.15761 | 0.87913 | -0.18585 | 0.77297 | 0.81161 | 1.0261 | 0.037189 |
| PEP-181/204 | 0.50186 | 0.59294 | 1.0925 | 0.1276 | 0.19653 | 0.34673 | 0.83867 | -0.25382 | 0.72158 | 0.77388 | 1.0593 | 0.083084 |
| PE180/224 | 0.46956 | 0.56192 | 1.1101 | 0.15072 | 0.15493 | 0.30108 | 0.83193 | -0.26546 | 0.16512 | 0.23343 | 0.83597 | -0.25848 |
| CE183 | 0.28094 | 0.36651 | 0.73753 | -0.43922 | 0.32489 | 0.48178 | 0.75356 | -0.40821 | 0.41841 | 0.49524 | 0.79489 | -0.33117 |
| PEP-182/204 | 0.29757 | 0.38437 | 1.1748 | 0.23244 | 0.32158 | 0.48016 | 0.86054 | -0.21669 | 0.30301 | 0.38678 | 1.1937 | 0.25547 |
| CER240 | 0.11391 | 0.18087 | 0.74828 | -0.41834 | 0.54249 | 0.68309 | 0.90638 | -0.14181 | 0.32972 | 0.41168 | 0.82956 | -0.26958 |
| PEP-160/183 | 0.15875 | 0.23016 | 0.72207 | -0.46978 | 0.30743 | 0.46744 | 0.7933 | -0.33406 | 0.04999 | 0.087955 | 0.60583 | -0.72301 |
| PA181/182 | 0.89644 | 0.92632 | 0.94153 | -0.08692 | 0.43449 | 0.57606 | 1.1157 | 0.15797 | 0.73129 | 0.7809 | 1.0586 | 0.082177 |
| FFA181 | 0.11585 | 0.1835 | 0.93149 | -0.10238 | 0.045427 | 0.14218 | 0.83797 | -0.25503 | 0.004098 | 0.011876 | 0.82909 | -0.27041 |
| PE180/182 | 0.93666 | 0.95876 | 0.99293 | -0.01023 | 0.02411 | 0.10326 | 0.78621 | -0.34701 | 0.023084 | 0.046098 | 0.72183 | -0.47027 |
| PC160/201 | 0.036621 | 0.072746 | 0.81227 | -0.29997 | 0.8824 | 0.9371 | 1.0145 | 0.020777 | 0.21561 | 0.29121 | 0.88015 | -0.18417 |
| PEP-180/224 | 0.67491 | 0.74469 | 0.94354 | -0.08385 | 0.12646 | 0.26081 | 0.77058 | -0.37598 | 0.94453 | 0.95352 | 0.99834 | -0.00239 |
| PS160/161_2 | 0.49094 | 0.58321 | 1.0596 | 0.083542 | 0.13039 | 0.26527 | 1.1866 | 0.24684 | 0.93378 | 0.9454 | 0.97777 | -0.03243 |
| PE180/181 | 0.38354 | 0.47469 | 1.087 | 0.12031 | 0.4987 | 0.64035 | 0.92923 | -0.10589 | 0.004105 | 0.011876 | 0.68086 | -0.55457 |
| FFA170 | 0.1768 | 0.24791 | 0.83291 | -0.26377 | 0.14374 | 0.28419 | 0.83275 | -0.26404 | 0.077477 | 0.12242 | 1.2648 | 0.3389 |
| FFA182 | 0.25702 | 0.34008 | 0.9306 | -0.10377 | 0.044671 | 0.14157 | 0.8587 | -0.21978 | 0.01738 | 0.036033 | 0.84257 | -0.24714 |
| PC150/161 | 0.68665 | 0.75385 | 0.94295 | -0.08474 | 0.59002 | 0.72336 | 0.93877 | -0.09116 | 0.58013 | 0.65115 | 1.0725 | 0.10092 |
| PC180/205 | 0.69293 | 0.75815 | 0.91315 | -0.13107 | 0.79913 | 0.88462 | 0.9534 | -0.06885 | 0.74182 | 0.78652 | 1.1868 | 0.24706 |
| FFA180 | 0.093937 | 0.15443 | 0.85848 | -0.22014 | 0.050497 | 0.14942 | 0.84264 | -0.24701 | 0.21778 | 0.29353 | 1.1295 | 0.17574 |
| TAG524-FA204 | 0.12681 | 0.19337 | 0.67162 | -0.57427 | 0.74847 | 0.8474 | 1.312 | 0.39179 | 0.33683 | 0.41688 | 1.2138 | 0.2795 |
| LPE182 | 0.97727 | 0.9818 | 0.98134 | -0.02718 | 0.27068 | 0.43094 | 0.81396 | -0.29697 | 0.004925 | 0.013247 | 0.52104 | -0.94052 |
| TAG480-FA160 | 3.38E-05 | 0.000594 | 0.28972 | -1.7873 | 0.1664 | 0.31217 | 0.76463 | -0.38717 | 0.000604 | 0.002864 | 0.45153 | -1.1471 |
| CER200 | 0.76994 | 0.81767 | 1.0441 | 0.062323 | 0.60457 | 0.73019 | 0.86833 | -0.20368 | 0.89738 | 0.9171 | 1.0626 | 0.087618 |
| CER220 | 0.083224 | 0.14036 | 0.62704 | -0.67336 | 0.20556 | 0.3559 | 0.70449 | -0.50534 | 0.49837 | 0.56819 | 0.83175 | -0.26577 |
| PEO-160/160 | 0.16725 | 0.23973 | 0.77945 | -0.35948 | 0.078921 | 0.1988 | 0.73532 | -0.44356 | 0.34141 | 0.42014 | 0.84501 | -0.24296 |
| TAG564-FA204 | 0.015597 | 0.035134 | 0.60539 | -0.72406 | 0.25606 | 0.41466 | 0.79998 | -0.32196 | 0.40952 | 0.48827 | 0.85871 | -0.21975 |
| PS160/181 | 0.088705 | 0.14845 | 1.2865 | 0.36341 | 0.10548 | 0.22737 | 1.1832 | 0.24265 | 0.03646 | 0.067815 | 1.2855 | 0.36232 |
| LPE220 | 0.63355 | 0.71206 | 1.0935 | 0.129 | 0.80497 | 0.8882 | 0.96712 | -0.04823 | 0.059058 | 0.10065 | 0.71113 | -0.49182 |
| TAG525-FA205 | 0.016047 | 0.035899 | 0.54342 | -0.87987 | 0.003688 | 0.038729 | 0.51906 | -0.94602 | 0.86153 | 0.88884 | 1.2497 | 0.32153 |

| | | | | | | | | | | | | |
|---|---|---|---|---|---|---|---|---|---|---|---|---|
| TAG544-FA224 | 0.25252 | 0.33686 | 0.72955 | -0.45491 | 0.93632 | 0.97035 | 1.2106 | 0.27574 | 0.87438 | 0.89783 | 1.064 | 0.089521 |
| PEO-180/161 | 0.060236 | 0.10685 | 0.76299 | -0.39026 | 0.46997 | 0.6135 | 0.91407 | -0.12962 | 0.008792 | 0.021023 | 0.60769 | -0.7186 |
| FFA221 | 0.12301 | 0.19066 | 1.1631 | 0.21799 | 0.21652 | 0.36802 | 0.85955 | -0.21835 | 0.44715 | 0.52167 | 1.2901 | 0.36746 |
| PE170/224 | 0.10413 | 0.16738 | 0.78196 | -0.35484 | 0.00017 | 0.032039 | 0.50035 | -0.999 | 0.00372 | 0.011006 | 0.59272 | -0.75457 |
| PC180/180 | 0.89041 | 0.92155 | 1.0202 | 0.02885 | 0.56954 | 0.70818 | 0.92548 | -0.11173 | 0.50999 | 0.5764 | 0.91656 | -0.12569 |
| CE181 | 0.069876 | 0.12131 | 0.66762 | -0.5829 | 0.022462 | 0.099535 | 0.59026 | -0.76057 | 0.2455 | 0.32374 | 0.75052 | -0.41404 |
| PA220/226 | 0.49211 | 0.58337 | 1.1 | 0.13756 | 0.49669 | 0.63902 | 0.89215 | -0.16464 | 0.33402 | 0.41576 | 1.2072 | 0.27165 |
| PEO-160/181 | 0.14248 | 0.21248 | 0.79133 | -0.33765 | 0.1048 | 0.22719 | 0.78595 | -0.3475 | 0.013846 | 0.030248 | 0.64523 | -0.63211 |
| TAG482-FA141 | 0.002677 | 0.00846 | 0.52382 | -0.93285 | 0.77399 | 0.86575 | 1.2345 | 0.30389 | 0.041932 | 0.075201 | 0.63644 | -0.65189 |
| FFA150 | 0.21694 | 0.29669 | 0.91096 | -0.13454 | 0.45823 | 0.60021 | 0.93425 | -0.09813 | 0.17566 | 0.24539 | 0.89566 | -0.15898 |
| LPC160 | 0.13416 | 0.20218 | 0.90913 | -0.13744 | 0.80037 | 0.88462 | 1.0213 | 0.030435 | 0.71883 | 0.77221 | 0.97886 | -0.03083 |
| TAG568-FA226 | 0.1181 | 0.1857 | 0.58791 | -0.76634 | 0.01666 | 0.082636 | 0.4306 | -1.2156 | 0.075952 | 0.12108 | 0.53459 | -0.90348 |
| LPE201 | 0.32915 | 0.42015 | 1.1574 | 0.21086 | 0.09563 | 0.22475 | 0.75763 | -0.40043 | 0.00535 | 0.013873 | 0.58417 | -0.77554 |
| PE160/224 | 0.73723 | 0.79329 | 1.0968 | 0.13324 | 0.12316 | 0.25718 | 0.76311 | -0.39003 | 0.010331 | 0.023934 | 0.57593 | -0.79603 |
| PC160/203 | 0.72897 | 0.7883 | 0.95799 | -0.06192 | 0.88722 | 0.93916 | 0.98567 | -0.02083 | 0.037926 | 0.069549 | 1.2269 | 0.29506 |
| FFA222 | 0.85226 | 0.88685 | 0.93466 | -0.09748 | 0.52957 | 0.66941 | 1.0542 | 0.076201 | 0.13596 | 0.19757 | 1.4536 | 0.5396 |
| HCER201 | 0.38838 | 0.47886 | 0.84332 | -0.24585 | 0.10919 | 0.23311 | 0.71433 | -0.48535 | 0.24588 | 0.32374 | 0.80003 | -0.32187 |
| TAG544-FA204 | 0.062102 | 0.10986 | 0.71805 | -0.47785 | 0.97744 | 0.98806 | 1.0938 | 0.1293 | 0.89643 | 0.9171 | 1.0254 | 0.036198 |
| TAG567-FA226 | 0.05259 | 0.095632 | 0.56391 | -0.82646 | 0.044872 | 0.14157 | 0.55346 | -0.85346 | 0.19785 | 0.27002 | 0.67559 | -0.56577 |
| PC200/225 | 0.1697 | 0.24121 | 0.80034 | -0.32131 | 0.056448 | 0.16047 | 0.70293 | -0.50856 | 0.70604 | 0.76098 | 0.94207 | -0.0861 |
| TAG503-FA141 | 0.046267 | 0.087171 | 0.60899 | -0.71551 | 0.51012 | 0.64988 | 1.0074 | 0.010616 | 0.14649 | 0.21005 | 0.77873 | -0.36081 |
| TAG565-FA225 | 0.001846 | 0.006635 | 0.52253 | -0.93641 | 0.005563 | 0.050778 | 0.55861 | -0.84008 | 0.087743 | 0.13633 | 0.70543 | -0.50343 |
| PC160/120 | 0.45241 | 0.5444 | 1.2039 | 0.26771 | 0.16408 | 0.30871 | 1.3048 | 0.38384 | 0.69316 | 0.74958 | 0.89982 | -0.15229 |
| FFA200 | 0.54625 | 0.63501 | 1.0753 | 0.10471 | 0.098765 | 0.22536 | 1.2013 | 0.26462 | 0.006286 | 0.015799 | 1.5509 | 0.63311 |
| PC180/224 | 0.36463 | 0.45825 | 0.89967 | -0.15254 | 0.98099 | 0.98858 | 1.0066 | 0.009489 | 0.88567 | 0.90799 | 0.99113 | -0.01285 |
| PC200/141 | 0.16681 | 0.23972 | 0.86084 | -0.21618 | 0.53495 | 0.67491 | 0.93743 | -0.09321 | 0.73516 | 0.7809 | 1.0648 | 0.090593 |
| PS161/180_2 | 0.40952 | 0.50044 | 1.1069 | 0.1465 | 0.81795 | 0.89644 | 1.0169 | 0.024204 | 0.2544 | 0.33222 | 1.0976 | 0.13429 |
| LPE161 | 0.9438 | 0.96152 | 1.0508 | 0.071497 | 0.040451 | 0.1386 | 0.75889 | -0.39804 | 0.038477 | 0.070361 | 0.78357 | -0.35187 |
| PC200/204 | 0.27591 | 0.3614 | 0.88754 | -0.17211 | 0.032498 | 0.12391 | 0.73554 | -0.44313 | 0.6801 | 0.73967 | 0.95835 | -0.06137 |
| SMd181/183 | 0.44348 | 0.53464 | 0.92854 | -0.10696 | 0.09922 | 0.22536 | 0.82348 | -0.2802 | 0.13564 | 0.19754 | 0.87156 | -0.19833 |
| PC180/200 | 0.85279 | 0.88685 | 0.97535 | -0.03601 | 0.93756 | 0.97035 | 0.98745 | -0.01822 | 0.38367 | 0.46169 | 0.9076 | -0.13987 |
| PE160/181 | 0.28078 | 0.36651 | 1.1614 | 0.2159 | 0.90622 | 0.94655 | 1.0171 | 0.024442 | 0.010968 | 0.025319 | 0.51081 | -0.96914 |
| PEP-160/201 | 0.010945 | 0.026293 | 0.68882 | -0.5378 | 0.006268 | 0.052991 | 0.64303 | -0.63704 | 0.32618 | 0.40836 | 0.88048 | -0.18363 |

| | | | | | | | | | | | | |
|---|---|---|---|---|---|---|---|---|---|---|---|---|
| PC180/120 | 0.77202 | 0.81855 | 1.0741 | 0.10308 | 0.80021 | 0.88462 | 0.90141 | -0.14974 | 0.6668 | 0.73078 | 0.89587 | -0.15864 |
| PEP-180/226 | 0.68245 | 0.75174 | 1.0445 | 0.062748 | 0.23846 | 0.39301 | 0.82114 | -0.2843 | 0.93032 | 0.94483 | 0.96664 | -0.04895 |
| DAG200/200 | 0.049994 | 0.091938 | 0.55836 | -0.84074 | 0.60415 | 0.73019 | 0.89429 | -0.16118 | 0.31696 | 0.39988 | 1.7116 | 0.77536 |
| PEP-160/161 | 0.3486 | 0.44121 | 0.85969 | -0.21811 | 0.58587 | 0.71963 | 0.91353 | -0.13047 | 0.59824 | 0.66346 | 1.0916 | 0.12645 |
| CER201 | 0.22651 | 0.3085 | 0.79209 | -0.33626 | 0.41663 | 0.56271 | 0.86217 | -0.21395 | 0.36034 | 0.43765 | 0.84714 | -0.23932 |
| PC170/204 | 0.068283 | 0.11917 | 0.85167 | -0.23164 | 0.009154 | 0.062074 | 0.74305 | -0.42847 | 0.25326 | 0.33173 | 0.89774 | -0.15564 |
| PE180/205 | 0.94865 | 0.96345 | 1.0472 | 0.066531 | 0.22581 | 0.37888 | 0.69984 | -0.51491 | 0.56869 | 0.63941 | 0.87143 | -0.19854 |
| FFA224 | 0.93828 | 0.9589 | 0.98723 | -0.01854 | 0.28789 | 0.45037 | 0.82051 | -0.28541 | 0.040034 | 0.072194 | 1.7007 | 0.76611 |
| PC140/224 | 0.19632 | 0.27307 | 0.8244 | -0.27857 | 0.92308 | 0.95841 | 1.1183 | 0.16134 | 0.1314 | 0.19223 | 0.75173 | -0.41171 |

## 8.5 Gy

| | SD 1 vs. Pre | | | | SD 2 vs. Pre | | | | SD 6 vs. Pre | | | |
|---|---|---|---|---|---|---|---|---|---|---|---|---|
| Lipid Name | p-value | FDR | FC | LOG2FC | p-value | FDR | FC | LOG2FC | p-value | FDR | FC | LOG2FC |
| PI160/201 | 0.000116 | 0.018752 | 0.42195 ↓ | -1.245 | 0.001978 | 0.042361 | 0.50991 ↓ | -0.972 | 0.001172 | 0.010394 | 0.48935 ↓ | -1.031 |
| LPE160 | 0.003914 | 0.030702 | 1.6618 ↑ | 0.7327 | 0.04967 | 0.16331 | 1.4067 ↑ | 0.4923 | 0.26651 | 0.35192 | 1.3018 ↑ | 0.3805 |
| PC182/203 | 0.000973 | 0.021852 | 0.36226 ↓ | -1.465 | 0.011229 | 0.081295 | 0.4901 ↓ | -1.029 | 0.004643 | 0.021435 | 0.46377 ↓ | -1.109 |
| PG180/182_2 | 0.000654 | 0.018752 | 0.42706 ↓ | -1.228 | 0.015154 | 0.086538 | 0.61792 ↓ | -0.695 | 0.000442 | 0.005708 | 0.43236 ↓ | -1.21 |
| PI160/204 | 0.003872 | 0.030702 | 0.50807 ↓ | -0.977 | 0.26843 | 0.48143 | 0.8351 ↓ | -0.26 | 0.046841 | 0.093315 | 0.75567 ↓ | -0.404 |
| PG180/182 | 0.000582 | 0.018752 | 0.41762 ↓ | -1.26 | 0.014815 | 0.085768 | 0.60917 ↓ | -0.715 | 0.000402 | 0.005336 | 0.42846 ↓ | -1.223 |
| LPC202 | 0.004444 | 0.031329 | 0.64966 ↓ | -0.622 | 0.008144 | 0.072759 | 0.68932 ↓ | -0.537 | 0.052626 | 0.10076 | 0.79468 ↓ | -0.332 |
| TAG547-FA183 | 0.000972 | 0.021852 | 0.31318 ↓ | -1.675 | 0.002088 | 0.042361 | 0.43399 ↓ | -1.204 | 0.000238 | 0.00465 | 0.2099 ↓ | -2.252 |
| PI160/182 | 0.002753 | 0.027101 | 0.39034 ↓ | -1.357 | 0.025054 | 0.1066 | 0.57492 ↓ | -0.799 | 0.000124 | 0.004481 | 0.45962 ↓ | -1.122 |
| TAG547-FA182 | 0.004481 | 0.031329 | 0.38567 ↓ | -1.375 | 0.011238 | 0.081295 | 0.51583 ↓ | -0.955 | 0.000497 | 0.005989 | 0.25348 ↓ | -1.98 |
| PS161/180 | 0.005129 | 0.033057 | 0.38096 ↓ | -1.392 | 0.036322 | 0.13107 | 0.48644 ↓ | -1.04 | 0.006192 | 0.024537 | 0.37306 ↓ | -1.423 |
| TAG526-FA183 | 0.004487 | 0.031329 | 0.26566 ↓ | -1.912 | 0.030801 | 0.11943 | 0.44247 ↓ | -1.176 | 0.000447 | 0.005708 | 0.14596 ↓ | -2.776 |
| TAG483-FA183 | 0.014842 | 0.053381 | 0.28521 ↓ | -1.81 | 0.58064 | 0.7575 | 0.87649 ↓ | -0.19 | 0.003375 | 0.017863 | 0.19956 ↓ | -2.325 |
| LPE226 | 0.21587 | 0.33602 | 1.3992 ↑ | 0.4846 | 0.16941 | 0.3677 | 1.3226 ↑ | 0.4034 | 0.092899 | 0.15527 | 1.6702 ↑ | 0.74 |
| PG160/161 | 0.12363 | 0.22671 | 0.59545 ↓ | -0.748 | 0.48526 | 0.69736 | 0.79795 ↓ | -0.326 | 0.0929 | 0.15527 | 0.55642 ↓ | -0.846 |
| TAG525-FA182 | 0.002418 | 0.026861 | 0.33904 ↓ | -1.561 | 0.016383 | 0.090212 | 0.51936 ↓ | -0.945 | 0.00024 | 0.00465 | 0.23651 ↓ | -2.08 |
| PE181/203 | 0.001941 | 0.023844 | 0.33423 ↓ | -1.581 | 0.005876 | 0.068309 | 0.36155 ↓ | -1.468 | 0.005128 | 0.022185 | 0.36625 ↓ | -1.449 |
| PI160/180 | 0.000379 | 0.018752 | 0.36939 ↓ | -1.437 | 0.011784 | 0.081295 | 0.54005 ↓ | -0.889 | 3.75E-05 | 0.004338 | 0.45102 ↓ | -1.149 |
| PC160/226 | 0.52851 | 0.65562 | 1.0811 ↑ | 0.1126 | 0.76563 | 0.87906 | 1.049 ↑ | 0.069 | 0.019479 | 0.050677 | 1.508 ↑ | 0.5927 |
| PI160/160 | 0.001568 | 0.022041 | 0.38802 ↓ | -1.366 | 0.02855 | 0.11763 | 0.58876 ↓ | -0.764 | 0.001086 | 0.00982 | 0.49582 ↓ | -1.012 |

| | | | | | | | | | | | | |
|---|---|---|---|---|---|---|---|---|---|---|---|---|
| PI181/203_2 | 0.001601 | 0.022041 | 0.38083 ↓ | -1.393 | 0.009235 | 0.076103 | 0.49356 ↓ | -1.019 | 0.010457 | 0.033533 | 0.49716 ↓ | -1.008 |
| TAG542-FA200 | 0.023146 | 0.071111 | 0.56879 ↓ | -0.814 | 0.053987 | 0.17228 | 0.65099 ↓ | -0.619 | 6.84E-05 | 0.00445 | 0.32776 ↓ | -1.609 |
| TAG546-FA183 | 0.003295 | 0.028391 | 0.3293 ↓ | -1.603 | 0.007253 | 0.072759 | 0.42924 ↓ | -1.22 | 0.000486 | 0.005989 | 0.22135 ↓ | -2.176 |
| TAG504-FA182 | 0.010593 | 0.045027 | 0.38393 ↓ | -1.381 | 0.33583 | 0.54824 | 0.7853 ↓ | -0.349 | 0.000272 | 0.004927 | 0.2355 ↓ | -2.086 |
| PC181/202 | 0.000609 | 0.018752 | 0.53975 ↓ | -0.89 | 0.001455 | 0.037304 | 0.54398 ↓ | -0.878 | 0.01867 | 0.049393 | 0.73399 ↓ | -0.446 |
| PC182/183 | 0.000691 | 0.018752 | 0.34025 ↓ | -1.555 | 0.002572 | 0.045251 | 0.44437 ↓ | -1.17 | 0.002379 | 0.014339 | 0.41071 ↓ | -1.284 |
| PI160/182_2 | 0.002749 | 0.027101 | 0.39856 ↓ | -1.327 | 0.034705 | 0.12837 | 0.6078 ↓ | -0.718 | 8.43E-05 | 0.004481 | 0.4666 ↓ | -1.1 |
| PI181/203 | 0.000242 | 0.018752 | 0.33523 ↓ | -1.577 | 0.005426 | 0.066642 | 0.47246 ↓ | -1.082 | 0.010167 | 0.033052 | 0.53572 ↓ | -0.9 |
| PG160/161_2 | 0.038486 | 0.099421 | 0.53248 ↓ | -0.909 | 0.077918 | 0.22346 | 0.59651 ↓ | -0.745 | 0.077565 | 0.1343 | 0.62167 ↓ | -0.686 |
| LPC200 | 0.001319 | 0.022041 | 0.40828 ↓ | -1.292 | 0.007987 | 0.072759 | 0.4919 ↓ | -1.024 | 0.10549 | 0.17083 | 0.69068 ↓ | -0.534 |
| TAG504-FA140 | 0.011571 | 0.0465 | 0.34047 ↓ | -1.554 | 0.13098 | 0.30342 | 0.59861 ↓ | -0.74 | 0.000331 | 0.00519 | 0.19831 ↓ | -2.334 |
| TAG525-FA183 | 0.003664 | 0.029813 | 0.42998 ↓ | -1.218 | 0.059221 | 0.18185 | 0.64317 ↓ | -0.637 | 0.000726 | 0.007499 | 0.27216 ↓ | -1.878 |
| TAG526-FA182 | 0.001601 | 0.022041 | 0.32992 ↓ | -1.6 | 0.008159 | 0.072759 | 0.45109 ↓ | -1.149 | 0.000123 | 0.004481 | 0.18572 ↓ | -2.429 |
| TAG525-FA225 | 0.01028 | 0.044614 | 0.50599 ↓ | -0.983 | 0.86839 | 0.92865 | 0.99776 ↓ | -0.003 | 0.024326 | 0.059092 | 0.50288 ↓ | -0.992 |
| TAG504-FA183 | 0.013138 | 0.049727 | 0.26838 ↓ | -1.898 | 0.55427 | 0.73639 | 0.80871 ↓ | -0.306 | 0.000911 | 0.008725 | 0.14852 ↓ | -2.751 |
| TAG526-FA161 | 0.002089 | 0.025179 | 0.25622 ↓ | -1.965 | 0.01018 | 0.0783 | 0.38357 ↓ | -1.382 | 0.000272 | 0.004927 | 0.1506 ↓ | -2.731 |
| CE204 | 0.59746 | 0.70589 | 1.0897 ↑ | 0.1239 | 0.014808 | 0.085768 | 0.65878 ↓ | -0.602 | 0.091888 | 0.15497 | 1.4381 ↑ | 0.5242 |
| TAG546-FA181 | 0.009014 | 0.042492 | 0.35311 ↓ | -1.502 | 0.03898 | 0.13717 | 0.5226 ↓ | -0.936 | 0.001653 | 0.012198 | 0.27518 ↓ | -1.862 |
| PI160/204_2 | 0.006491 | 0.035833 | 0.51574 ↓ | -0.955 | 0.20525 | 0.40863 | 0.81422 ↓ | -0.297 | 0.092364 | 0.15527 | 0.798 ↓ | -0.326 |
| TAG503-FA140 | 0.015759 | 0.055757 | 0.34816 ↓ | -1.522 | 0.53236 | 0.72202 | 0.78908 ↓ | -0.342 | 0.002681 | 0.015751 | 0.26132 ↓ | -1.936 |
| PI161/181 | 0.00114 | 0.022041 | 0.45643 ↓ | -1.132 | 0.093621 | 0.25003 | 0.69597 ↓ | -0.523 | 0.009832 | 0.032328 | 0.55772 ↓ | -0.842 |
| LPC220 | 0.001322 | 0.022041 | 0.37472 ↓ | -1.416 | 0.014206 | 0.085768 | 0.46438 ↓ | -1.107 | 0.30506 | 0.39404 | 0.73911 ↓ | -0.436 |
| TAG548-FA182 | 0.001178 | 0.022041 | 0.27606 ↓ | -1.857 | 0.001264 | 0.035765 | 0.40709 ↓ | -1.297 | 3.13E-05 | 0.004338 | 0.16345 ↓ | -2.613 |
| PC182/202 | 0.000502 | 0.018752 | 0.52631 ↓ | -0.926 | 0.001072 | 0.033228 | 0.53905 ↓ | -0.892 | 0.016069 | 0.045089 | 0.67173 ↓ | -0.574 |
| PI182/182 | 0.009243 | 0.042981 | 0.36815 ↓ | -1.442 | 0.029386 | 0.11815 | 0.48721 ↓ | -1.037 | 0.00084 | 0.008163 | 0.31837 ↓ | -1.651 |
| TAG546-FA182 | 0.05612 | 0.13326 | 0.65231 ↓ | -0.616 | 0.053632 | 0.17199 | 0.66257 ↓ | -0.594 | 0.002848 | 0.016406 | 0.42583 ↓ | -1.232 |
| PI181/182_2 | 0.002527 | 0.026861 | 0.34364 ↓ | -1.541 | 0.029945 | 0.11815 | 0.5313 ↓ | -0.912 | 0.008973 | 0.030364 | 0.52335 ↓ | -0.934 |
| PI160/180_2 | 0.000415 | 0.018752 | 0.4062 ↓ | -1.3 | 0.010223 | 0.0783 | 0.55593 ↓ | -0.847 | 8.85E-05 | 0.004481 | 0.4846 ↓ | -1.045 |
| TAG525-FA160 | 0.004085 | 0.031329 | 0.43033 ↓ | -1.217 | 0.093927 | 0.25003 | 0.65765 ↓ | -0.605 | 0.001372 | 0.011166 | 0.31927 ↓ | -1.647 |
| TAG525-FA161 | 0.001892 | 0.023843 | 0.29144 ↓ | -1.779 | 0.023019 | 0.10446 | 0.48692 ↓ | -1.038 | 0.000537 | 0.006244 | 0.23697 ↓ | -2.077 |
| TAG505-FA161 | 0.006605 | 0.035833 | 0.23556 ↓ | -2.086 | 0.10753 | 0.26899 | 0.47356 ↓ | -1.078 | 0.000229 | 0.00465 | 0.090969 ↓ | -3.459 |
| PG181/182 | 0.001334 | 0.022041 | 0.3816 ↓ | -1.39 | 0.004909 | 0.06511 | 0.39886 ↓ | -1.326 | 0.000391 | 0.005336 | 0.33211 ↓ | -1.59 |
| TAG563-FA200 | 0.036291 | 0.094976 | 0.61053 ↓ | -0.712 | 0.021815 | 0.10366 | 0.59352 ↓ | -0.753 | 0.000178 | 0.00465 | 0.26801 ↓ | -1.9 |

| | | | | | | | | | | | | | | | |
|---|---|---|---|---|---|---|---|---|---|---|---|---|---|---|---|
| PC160/204 | 0.050715 | 0.12412 | 1.173 | ↑ | 0.2302 | 0.75536 | 0.87498 | 1.0431 | ↑ | 0.0609 | 0.010882 | 0.034557 | 1.4322 | ↑ | 0.5183 |
| PC182/182 | 0.000491 | 0.018752 | 0.40605 | ↓ | -1.3 | 0.000507 | 0.027482 | 0.48309 | ↓ | -1.05 | 0.020044 | 0.05097 | 0.65265 | ↓ | -0.616 |
| PI160/181 | 0.001758 | 0.02334 | 0.38229 | ↓ | -1.387 | 0.022401 | 0.10446 | 0.5573 | ↓ | -0.843 | 0.002929 | 0.016578 | 0.54255 | ↓ | -0.882 |
| TAG482-FA182 | 0.013379 | 0.050289 | 0.33498 | ↓ | -1.578 | 0.71397 | 0.8506 | 0.93228 | ↓ | -0.101 | 0.009155 | 0.030563 | 0.33062 | ↓ | -1.597 |
| PG181/181 | 0.004195 | 0.031329 | 0.44904 | ↓ | -1.155 | 0.005757 | 0.068143 | 0.44163 | ↓ | -1.179 | 0.002317 | 0.014229 | 0.43589 | ↓ | -1.198 |
| TAG566-FA183 | 0.002574 | 0.026861 | 0.49099 | ↓ | -1.026 | 0.017015 | 0.090528 | 0.54872 | ↓ | -0.866 | 0.000394 | 0.005336 | 0.28559 | ↓ | -1.808 |
| PC182/161 | 0.00128 | 0.022041 | 0.48866 | ↓ | -1.033 | 0.010157 | 0.0783 | 0.63726 | ↓ | -0.65 | 0.037784 | 0.079347 | 0.68377 | ↓ | -0.548 |
| PI181/182 | 0.002864 | 0.027101 | 0.35166 | ↓ | -1.508 | 0.023895 | 0.10511 | 0.52224 | ↓ | -0.937 | 0.006391 | 0.025065 | 0.5032 | ↓ | -0.991 |
| TAG524-FA221 | 0.012367 | 0.047921 | 0.44566 | ↓ | -1.166 | 0.2633 | 0.47745 | 0.71643 | ↓ | -0.481 | 0.007397 | 0.026754 | 0.38794 | ↓ | -1.366 |
| PI160/181_2 | 0.001334 | 0.022041 | 0.38063 | ↓ | -1.394 | 0.014047 | 0.085768 | 0.5472 | ↓ | -0.87 | 0.001703 | 0.01232 | 0.52722 | ↓ | -0.924 |
| TAG541-FA200 | 0.012927 | 0.049505 | 0.48682 | ↓ | -1.039 | 0.05724 | 0.17829 | 0.64142 | ↓ | -0.641 | 3.62E-05 | 0.004338 | 0.25398 | ↓ | -1.977 |
| TAG548-FA183 | 0.000443 | 0.018752 | 0.21506 | ↓ | -2.217 | 0.002979 | 0.049724 | 0.36874 | ↓ | -1.439 | 2.48E-05 | 0.004338 | 0.11544 | ↓ | -3.115 |
| CE226 | 0.26636 | 0.3932 | 1.1765 | ↑ | 0.2345 | 0.15999 | 0.35548 | 0.69365 | ↓ | -0.528 | 0.049781 | 0.097029 | 1.5183 | ↑ | 0.6024 |
| TAG503-FA182 | 0.022326 | 0.070853 | 0.38882 | ↓ | -1.363 | 0.56212 | 0.74076 | 0.83626 | ↓ | -0.258 | 0.00809 | 0.028406 | 0.32833 | ↓ | -1.607 |
| LPC183 | 0.034364 | 0.091686 | 0.66951 | ↓ | -0.579 | 0.16636 | 0.36464 | 0.78306 | ↓ | -0.353 | 0.002036 | 0.013386 | 0.5208 | ↓ | -0.941 |
| TAG525-FA181 | 0.006212 | 0.035167 | 0.3135 | ↓ | -1.674 | 0.049372 | 0.16315 | 0.47346 | ↓ | -1.079 | 0.000582 | 0.006418 | 0.1865 | ↓ | -2.423 |
| PC140/226 | 0.010997 | 0.045027 | 0.54062 | ↓ | -0.887 | 0.17677 | 0.37606 | 0.73307 | ↓ | -0.448 | 0.19639 | 0.27734 | 0.74514 | ↓ | -0.424 |
| LPC150 | 0.012162 | 0.047921 | 0.64558 | ↓ | -0.631 | 0.049159 | 0.16315 | 0.70897 | ↓ | -0.496 | 0.004086 | 0.020451 | 0.62379 | ↓ | -0.681 |
| PS161/181 | 0.056291 | 0.13326 | 0.47591 | ↓ | -1.071 | 0.1319 | 0.30342 | 0.55874 | ↓ | -0.84 | 0.77446 | 0.82759 | 1.0513 | ↑ | 0.0722 |
| PC180/201 | 0.000669 | 0.018752 | 0.7085 | ↓ | -0.497 | 0.000579 | 0.029009 | 0.68382 | ↓ | -0.548 | 0.001213 | 0.010394 | 0.812 | ↓ | -0.3 |
| PI181/181 | 0.002312 | 0.026861 | 0.32803 | ↓ | -1.608 | 0.013386 | 0.085427 | 0.44913 | ↓ | -1.155 | 0.22204 | 0.30785 | 0.73945 | ↓ | -0.435 |
| PI160/203_2 | 0.003049 | 0.027956 | 0.45047 | ↓ | -1.151 | 0.013516 | 0.085427 | 0.55826 | ↓ | -0.841 | 0.003224 | 0.017847 | 0.53037 | ↓ | -0.915 |
| TAG562-FA200 | 0.071717 | 0.1559 | 0.62387 | ↓ | -0.681 | 0.011672 | 0.081295 | 0.52201 | ↓ | -0.938 | 0.000357 | 0.005277 | 0.30423 | ↓ | -1.717 |
| PC180/202 | 0.010363 | 0.044676 | 0.72004 | ↓ | -0.474 | 0.000855 | 0.030924 | 0.68981 | ↓ | -0.536 | 0.005649 | 0.023333 | 0.75942 | ↓ | -0.397 |
| TAG547-FA181 | 0.003272 | 0.028391 | 0.34014 | ↓ | -1.556 | 0.023434 | 0.10446 | 0.47452 | ↓ | -1.076 | 0.000343 | 0.00519 | 0.22798 | ↓ | -2.133 |
| TAG503-FA181 | 0.051662 | 0.12596 | 0.41176 | ↓ | -1.28 | 0.56838 | 0.7475 | 0.77101 | ↓ | -0.375 | 0.001668 | 0.012198 | 0.22895 | ↓ | -2.127 |
| TAG484-FA120 | 0.00245 | 0.026861 | 0.28326 | ↓ | -1.82 | 0.12307 | 0.29347 | 0.57406 | ↓ | -0.801 | 0.007214 | 0.026533 | 0.31239 | ↓ | -1.679 |
| TAG504-FA161 | 0.009771 | 0.043868 | 0.25046 | ↓ | -1.997 | 0.25806 | 0.47238 | 0.60522 | ↓ | -0.724 | 0.002113 | 0.013753 | 0.15002 | ↓ | -2.737 |
| PG160/203_2 | 0.001426 | 0.022041 | 0.45915 | ↓ | -1.123 | 0.013501 | 0.085427 | 0.47818 | ↓ | -1.064 | 0.000138 | 0.004481 | 0.30714 | ↓ | -1.703 |
| TAG524-FA161 | 0.004743 | 0.031829 | 0.32016 | ↓ | -1.643 | 0.19661 | 0.40376 | 0.63634 | ↓ | -0.652 | 0.001529 | 0.01199 | 0.27203 | ↓ | -1.878 |
| TAG563-FA182 | 0.12666 | 0.23032 | 0.72893 | ↓ | -0.456 | 0.17377 | 0.37173 | 0.76402 | ↓ | -0.388 | 0.004996 | 0.022058 | 0.45202 | ↓ | -1.146 |
| TAG565-FA201 | 0.20357 | 0.32408 | 0.70808 | ↓ | -0.498 | 0.29841 | 0.51122 | 0.75353 | ↓ | -0.408 | 0.009002 | 0.030364 | 0.47158 | ↓ | -1.084 |
| PI160/203 | 0.001793 | 0.02334 | 0.43712 | ↓ | -1.194 | 0.00763 | 0.072759 | 0.58549 | ↓ | -0.772 | 0.000213 | 0.00465 | 0.48825 | ↓ | -1.034 |

| | | | | | | | | | | | | | | | |
|---|---|---|---|---|---|---|---|---|---|---|---|---|---|---|---|
| PI180/180 | 0.00063 | 0.018752 | 0.41087 | ↓ | -1.283 | 0.008658 | 0.074165 | 0.54393 | ↓ | -0.879 | 0.005735 | 0.023333 | 0.53687 | ↓ | -0.897 |
| TAG541-FA160 | 0.05833 | 0.13554 | 0.58866 | ↓ | -0.765 | 0.51188 | 0.71313 | 0.88384 | ↓ | -0.178 | 0.005223 | 0.022225 | 0.42667 | ↓ | -1.229 |
| PC182/224 | 0.008691 | 0.041601 | 0.54408 | ↓ | -0.878 | 0.018065 | 0.091167 | 0.59942 | ↓ | -0.738 | 0.025691 | 0.06104 | 0.61241 | ↓ | -0.707 |
| PC182/205 | 0.009935 | 0.044164 | 0.45193 | ↓ | -1.146 | 0.001991 | 0.042361 | 0.43073 | ↓ | -1.215 | 0.014662 | 0.041865 | 0.47608 | ↓ | -1.071 |
| TAG545-FA180 | 0.058504 | 0.13554 | 0.57997 | ↓ | -0.786 | 0.22766 | 0.43719 | 0.7447 | ↓ | -0.425 | 0.004252 | 0.02052 | 0.31917 | ↓ | -1.648 |
| PI161/181_2 | 0.017401 | 0.059002 | 0.43713 | ↓ | -1.194 | 0.037174 | 0.13248 | 0.56545 | ↓ | -0.823 | 0.005723 | 0.023333 | 0.54684 | ↓ | -0.871 |
| TAG524-FA160 | 0.041686 | 0.10588 | 0.67437 | ↓ | -0.568 | 0.40196 | 0.62155 | 0.8984 | ↓ | -0.155 | 0.019892 | 0.050783 | 0.61629 | ↓ | -0.698 |
| PC160/225 | 0.060902 | 0.13939 | 1.2778 | ↑ | 0.3536 | 0.23238 | 0.443 | 1.2107 | ↑ | 0.2758 | 0.1084 | 0.17468 | 1.3262 | ↑ | 0.4073 |
| TAG524-FA183 | 0.016049 | 0.056477 | 0.47862 | ↓ | -1.063 | 0.23934 | 0.45162 | 0.75285 | ↓ | -0.41 | 0.002919 | 0.016578 | 0.37931 | ↓ | -1.399 |
| TAG561-FA181 | 0.02084 | 0.067782 | 0.5208 | ↓ | -0.941 | 0.091213 | 0.24741 | 0.6535 | ↓ | -0.614 | 0.000799 | 0.008006 | 0.34709 | ↓ | -1.527 |
| TAG482-FA140 | 0.060674 | 0.13939 | 0.45009 | ↓ | -1.152 | 0.6795 | 0.82838 | 0.93775 | ↓ | -0.093 | 0.006679 | 0.025472 | 0.29555 | ↓ | -1.759 |
| PI200/204_2 | 0.000881 | 0.021852 | 0.3613 | ↓ | -1.469 | 0.010189 | 0.0783 | 0.48216 | ↓ | -1.052 | 0.064191 | 0.1164 | 0.624 | ↓ | -0.68 |
| TAG502-FA140 | 0.089448 | 0.18197 | 0.46511 | ↓ | -1.104 | 0.86215 | 0.92865 | 0.89458 | ↓ | -0.161 | 0.005775 | 0.023352 | 0.27558 | ↓ | -1.859 |
| TAG545-FA183 | 0.004653 | 0.031829 | 0.37833 | ↓ | -1.402 | 0.051857 | 0.16795 | 0.54471 | ↓ | -0.876 | 0.001202 | 0.010394 | 0.2554 | ↓ | -1.969 |
| PG160/203 | 0.002579 | 0.026861 | 0.51745 | ↓ | -0.951 | 0.01235 | 0.082041 | 0.50822 | ↓ | -0.976 | 0.000292 | 0.005048 | 0.34849 | ↓ | -1.521 |
| PI160/161 | 0.000397 | 0.018752 | 0.38355 | ↓ | -1.383 | 0.15356 | 0.34353 | 0.74733 | ↓ | -0.42 | 0.005015 | 0.022058 | 0.52831 | ↓ | -0.921 |
| PE182/202 | 0.004261 | 0.031329 | 0.43543 | ↓ | -1.2 | 0.003669 | 0.058253 | 0.39864 | ↓ | -1.327 | 0.000137 | 0.004481 | 0.29933 | ↓ | -1.74 |
| PI182/201 | 0.084542 | 0.1767 | 0.63217 | ↓ | -0.662 | 0.14532 | 0.32848 | 0.6607 | ↓ | -0.598 | 0.21941 | 0.3052 | 0.71597 | ↓ | -0.482 |
| LPA180 | 0.86309 | 0.92066 | 1.0714 | ↑ | 0.0994 | 0.005006 | 0.06511 | 0.83502 | ↓ | -0.26 | 0.64182 | 0.72039 | 0.96862 | ↓ | -0.046 |
| LPC240 | 0.003432 | 0.028641 | 0.46604 | ↓ | -1.102 | 0.012088 | 0.081295 | 0.49955 | ↓ | -1.001 | 0.33571 | 0.42601 | 0.78152 | ↓ | -0.356 |
| TAG524-FA200 | 0.010917 | 0.045027 | 0.56162 | ↓ | -0.832 | 0.29155 | 0.50613 | 0.84965 | ↓ | -0.235 | 0.024663 | 0.059686 | 0.56667 | ↓ | -0.819 |
| TAG566-FA202 | 0.01736 | 0.059002 | 0.5765 | ↓ | -0.795 | 0.014684 | 0.085768 | 0.61307 | ↓ | -0.706 | 0.001331 | 0.011037 | 0.42657 | ↓ | -1.229 |
| PS181/182 | 0.2787 | 0.4068 | 0.79555 | ↓ | -0.33 | 0.30423 | 0.51709 | 0.81081 | ↓ | -0.303 | 0.021654 | 0.054448 | 0.58271 | ↓ | -0.779 |
| TAG524-FA181 | 0.015464 | 0.05501 | 0.41775 | ↓ | -1.259 | 0.2031 | 0.40863 | 0.67356 | ↓ | -0.57 | 0.002726 | 0.015843 | 0.32882 | ↓ | -1.605 |
| PG161/180 | 0.5039 | 0.63821 | 1.1562 | ↑ | 0.2094 | 0.096772 | 0.25426 | 0.68097 | ↓ | -0.554 | 0.000183 | 0.00465 | 0.34864 | ↓ | -1.52 |
| PI180/203_2 | 0.017248 | 0.059002 | 0.62875 | ↓ | -0.669 | 0.16571 | 0.36445 | 0.78867 | ↓ | -0.343 | 0.022907 | 0.056832 | 0.6269 | ↓ | -0.674 |
| PI180/183_2 | 0.007514 | 0.038822 | 0.57458 | ↓ | -0.799 | 0.041276 | 0.14369 | 0.70573 | ↓ | -0.503 | 0.000041 | 0.004338 | 0.43695 | ↓ | -1.195 |
| LPC241 | 0.001534 | 0.022041 | 0.43127 | ↓ | -1.213 | 0.082239 | 0.22977 | 0.63451 | ↓ | -0.656 | 0.070775 | 0.12506 | 0.66288 | ↓ | -0.593 |
| CE203 | 0.9755 | 0.9861 | 0.89779 | ↓ | -0.156 | 0.017469 | 0.090528 | 0.45192 | ↓ | -1.146 | 0.54334 | 0.63504 | 1.0929 | ↑ | 0.1282 |
| TAG524-FA182 | 0.060928 | 0.13939 | 0.70077 | ↓ | -0.513 | 0.26607 | 0.4798 | 0.86106 | ↓ | -0.216 | 0.018741 | 0.049393 | 0.61791 | ↓ | -0.695 |
| TAG545-FA182 | 0.061568 | 0.14014 | 0.63545 | ↓ | -0.654 | 0.076562 | 0.22152 | 0.67987 | ↓ | -0.557 | 0.005832 | 0.023436 | 0.42797 | ↓ | -1.224 |
| TAG544-FA201 | 0.078369 | 0.16564 | 0.5671 | ↓ | -0.818 | 0.53187 | 0.72202 | 0.83036 | ↓ | -0.268 | 0.006971 | 0.025783 | 0.42138 | ↓ | -1.247 |
| TAG545-FA161 | 0.005843 | 0.033965 | 0.36513 | ↓ | -1.454 | 0.15466 | 0.3448 | 0.60826 | ↓ | -0.717 | 0.000529 | 0.006244 | 0.24239 | ↓ | -2.045 |

| | | | | | | | | | | | |
|---|---|---|---|---|---|---|---|---|---|---|---|
| TAG502-FA181 | 0.069647 | 0.15215 | 0.42803 | -1.224 | 0.67764 | 0.82838 | 0.86006 | -0.217 | 0.01382 | 0.04115 | 0.29566 | -1.758 |
| PG181/182_2 | 0.001169 | 0.022041 | 0.39813 | -1.3287 | 0.004323 | 0.063963 | 0.40584 | -1.301 | 0.000243 | 0.00465 | 0.34415 | -1.5389 |
| PC200/182 | 0.038345 | 0.099421 | 0.69348 | -0.52808 | 0.00149 | 0.037304 | 0.552 | -0.85727 | 0.031214 | 0.070556 | 0.69505 | -0.5248 |
| PC140/203 | 0.00608 | 0.034722 | 0.46507 | -1.1045 | 0.00681 | 0.071506 | 0.47237 | -1.082 | 0.015571 | 0.043881 | 0.54347 | -0.87973 |
| TAG543-FA183 | 0.095555 | 0.19037 | 0.69233 | -0.53047 | 0.50695 | 0.71153 | 0.9059 | -0.14257 | 0.002178 | 0.013864 | 0.32305 | -1.6302 |
| PC181/203 | 0.010824 | 0.045027 | 0.53049 | -0.9146 | 0.008079 | 0.072759 | 0.49372 | -1.0182 | 0.2737 | 0.35995 | 0.75768 | -0.40034 |
| TAG523-FA201 | 0.018663 | 0.061989 | 0.45314 | -1.142 | 0.29523 | 0.50846 | 0.71519 | -0.48361 | 0.063667 | 0.1162 | 0.52947 | -0.91739 |
| LPC201 | 0.002914 | 0.027101 | 0.62415 | -0.68004 | 0.03177 | 0.12238 | 0.72376 | -0.46641 | 0.86653 | 0.90258 | 0.99817 | -0.00264 |
| PE181/205 | 0.054514 | 0.13098 | 0.47602 | -1.0709 | 0.015509 | 0.087039 | 0.39865 | -1.3268 | 0.02681 | 0.062781 | 0.41258 | -1.2773 |
| LPC161 | 0.004965 | 0.032648 | 0.57153 | -0.8071 | 0.40678 | 0.62642 | 0.88129 | -0.18232 | 0.069942 | 0.12407 | 0.75574 | -0.40404 |
| TAG481-FA181 | 0.066244 | 0.14668 | 0.4652 | -1.1041 | 0.7643 | 0.87906 | 0.99247 | -0.0109 | 0.014307 | 0.041214 | 0.34858 | -1.5204 |
| PC120/181 | 0.054448 | 0.13098 | 0.51515 | -0.95695 | 0.017782 | 0.090528 | 0.46009 | -1.12 | 0.038755 | 0.081069 | 0.49212 | -1.0229 |
| PC150/203 | 0.024525 | 0.073807 | 0.62469 | -0.67879 | 0.008623 | 0.074165 | 0.53699 | -0.89703 | 0.001782 | 0.012683 | 0.50797 | -0.97719 |
| TAG567-FA183 | 0.000638 | 0.018752 | 0.41133 | -1.2816 | 0.028912 | 0.11792 | 0.62475 | -0.67865 | 0.001958 | 0.013144 | 0.44869 | -1.1562 |
| TAG544-FA183 | 0.11401 | 0.21085 | 0.67125 | -0.57507 | 0.53605 | 0.724 | 0.87183 | -0.19788 | 0.006013 | 0.024014 | 0.39857 | -1.3271 |
| TAG523-FA183 | 0.024402 | 0.073807 | 0.50219 | -0.99368 | 0.70044 | 0.84131 | 0.94147 | -0.08701 | 0.006219 | 0.024537 | 0.41249 | -1.2776 |
| PE182/201 | 0.000939 | 0.021852 | 0.45184 | -1.1461 | 0.001647 | 0.039715 | 0.37661 | -1.4089 | 0.000155 | 0.004573 | 0.39487 | -1.3405 |
| TAG544-FA180 | 0.21627 | 0.33602 | 0.78941 | -0.34115 | 0.52957 | 0.72202 | 0.90345 | -0.14649 | 0.014558 | 0.041749 | 0.48774 | -1.0358 |
| TAG482-FA161 | 0.15909 | 0.27254 | 0.4119 | -1.2796 | 0.98213 | 0.9928 | 1.2208 | 0.28778 | 0.01808 | 0.049041 | 0.22004 | -2.1841 |
| TAG482-FA160 | 0.063888 | 0.14391 | 0.48533 | -1.043 | 0.82235 | 0.90904 | 1.068 | 0.094883 | 0.00498 | 0.022058 | 0.30247 | -1.7251 |
| PI180/181_2 | 0.000588 | 0.018752 | 0.42702 | -1.2276 | 0.007513 | 0.072759 | 0.54521 | -0.87512 | 0.012747 | 0.039142 | 0.55729 | -0.8435 |
| TAG523-FA180 | 0.34644 | 0.47986 | 0.78074 | -0.35709 | 0.78907 | 0.89286 | 0.97657 | -0.0342 | 0.003255 | 0.017847 | 0.3441 | -1.5391 |
| PG160/182_2 | 0.011239 | 0.045727 | 0.45732 | -1.1287 | 0.050572 | 0.16461 | 0.54311 | -0.88069 | 0.00518 | 0.022185 | 0.42702 | -1.2276 |
| TAG503-FA161 | 0.039028 | 0.10003 | 0.42331 | -1.2402 | 0.63268 | 0.79666 | 0.8712 | -0.19892 | 0.006691 | 0.025472 | 0.29599 | -1.7564 |
| PI180/181 | 0.001176 | 0.022041 | 0.43493 | -1.2012 | 0.011862 | 0.081295 | 0.53659 | -0.8981 | 0.024033 | 0.058598 | 0.56627 | -0.82045 |
| PC182/226 | 0.000239 | 0.018752 | 0.43117 | -1.2137 | 0.016398 | 0.090212 | 0.60798 | -0.71791 | 0.01325 | 0.040495 | 0.66044 | -0.5985 |
| PC181/183 | 0.008378 | 0.041003 | 0.55457 | -0.85055 | 0.00187 | 0.042361 | 0.563 | -0.82879 | 0.006794 | 0.025565 | 0.47422 | -1.0764 |
| PC182/204 | 0.000214 | 0.018752 | 0.52658 | -0.92528 | 0.003237 | 0.052682 | 0.63804 | -0.64828 | 0.003898 | 0.019919 | 0.72959 | -0.45484 |
| TAG565-FA182 | 0.085029 | 0.17685 | 0.68507 | -0.54567 | 0.11941 | 0.28791 | 0.72002 | -0.47388 | 0.013853 | 0.04115 | 0.52621 | -0.9263 |
| TAG544-FA161 | 0.035179 | 0.093096 | 0.49429 | -1.0166 | 0.25664 | 0.47238 | 0.6875 | -0.54057 | 0.002029 | 0.013386 | 0.27487 | -1.8632 |
| PC181/226 | 0.005791 | 0.033965 | 0.59024 | -0.76062 | 0.000638 | 0.029687 | 0.5117 | -0.96663 | 0.22226 | 0.30785 | 0.82651 | -0.27489 |
| TAG520-FA200 | 0.00844 | 0.041003 | 0.47567 | -1.072 | 0.022806 | 0.10446 | 0.56318 | -0.82833 | 0.000224 | 0.00465 | 0.28538 | -1.8091 |
| SMd181/182 | 0.74276 | 0.83225 | 1.0531 | 0.074581 | 0.20313 | 0.40863 | 0.8964 | -0.15778 | 0.023429 | 0.057453 | 1.2228 | 0.29015 |

| | | | | | | | | | | | | |
|---|---|---|---|---|---|---|---|---|---|---|---|---|
| PC140/202 | 0.005712 | 0.033965 | 0.41872 | -1.256 | 0.001375 | 0.037304 | 0.48403 | -1.0468 | 0.00012 | 0.004481 | 0.38024 | -1.395 |
| TAG524-FA180 | 0.021347 | 0.068797 | 0.47136 | -1.0851 | 0.88734 | 0.93672 | 0.98014 | -0.02894 | 0.001958 | 0.013144 | 0.30421 | -1.7169 |
| PG160/182 | 0.012269 | 0.047921 | 0.45242 | -1.1443 | 0.055346 | 0.17576 | 0.54575 | -0.87369 | 0.007296 | 0.026682 | 0.43342 | -1.2062 |
| PI180/182_2 | 0.006661 | 0.035838 | 0.55367 | -0.8529 | 0.094376 | 0.25003 | 0.73393 | -0.44628 | 0.000689 | 0.007239 | 0.5196 | -0.94453 |
| TAG526-FA181 | 0.032293 | 0.088331 | 0.42698 | -1.2278 | 0.079929 | 0.22623 | 0.51659 | -0.9529 | 0.004705 | 0.021438 | 0.20246 | -2.3043 |
| TAG483-FA161 | 0.096921 | 0.19037 | 0.40397 | -1.3077 | 0.98675 | 0.99439 | 1.0406 | 0.057477 | 0.012534 | 0.038672 | 0.24768 | -2.0135 |
| TAG565-FA202 | 0.15443 | 0.26596 | 0.7176 | -0.47875 | 0.042978 | 0.14803 | 0.6464 | -0.6295 | 0.004031 | 0.020341 | 0.45484 | -1.1366 |
| LPC170 | 0.006541 | 0.035833 | 0.66303 | -0.59286 | 0.056636 | 0.17733 | 0.74855 | -0.41783 | 0.028869 | 0.065943 | 0.77385 | -0.36986 |
| TAG545-FA181 | 0.036327 | 0.094976 | 0.58256 | -0.77951 | 0.07582 | 0.22035 | 0.66473 | -0.58915 | 0.002351 | 0.014305 | 0.37085 | -1.4311 |
| TAG523-FA202 | 0.14558 | 0.25615 | 0.63854 | -0.64715 | 0.94131 | 0.9789 | 1.023 | 0.03274 | 0.017423 | 0.047656 | 0.46684 | -1.099 |
| PI180/182 | 0.011966 | 0.047497 | 0.55822 | -0.8411 | 0.12024 | 0.28883 | 0.72625 | -0.46146 | 0.000829 | 0.008163 | 0.48565 | -1.042 |
| TAG540-FA160 | 0.005107 | 0.033057 | 0.48706 | -1.0378 | 0.018425 | 0.092267 | 0.57124 | -0.80784 | 0.001012 | 0.009411 | 0.39819 | -1.3285 |
| TAG526-FA160 | 0.009581 | 0.043341 | 0.43367 | -1.2053 | 0.19084 | 0.39565 | 0.73437 | -0.44543 | 0.000385 | 0.005336 | 0.2355 | -2.0862 |
| TAG526-FA205 | 0.60872 | 0.71518 | 0.86837 | -0.20361 | 0.83494 | 0.9166 | 1.0531 | 0.074587 | 0.082968 | 0.14251 | 0.53502 | -0.90234 |
| PS180/181 | 0.005518 | 0.033946 | 0.47969 | -1.0598 | 0.011948 | 0.081295 | 0.51171 | -0.9666 | 0.055168 | 0.10407 | 0.59402 | -0.75143 |
| TAG523-FA182 | 0.11931 | 0.22003 | 0.73054 | -0.45297 | 0.54035 | 0.7283 | 0.90005 | -0.15193 | 0.025432 | 0.060645 | 0.61874 | -0.69261 |
| TAG545-FA202 | 0.014696 | 0.053151 | 0.46057 | -1.1185 | 0.052596 | 0.16951 | 0.56436 | -0.82531 | 0.001055 | 0.009669 | 0.27108 | -1.8832 |
| TAG483-FA140 | 0.032871 | 0.088792 | 0.27389 | -1.8683 | 0.4522 | 0.66512 | 0.69284 | -0.52941 | 0.003345 | 0.017847 | 0.15306 | -2.7078 |
| PE180/202 | 0.22114 | 0.34114 | 0.86976 | -0.2013 | 0.023364 | 0.10446 | 0.74253 | -0.42948 | 0.047191 | 0.093379 | 0.7324 | -0.44929 |
| PI180/202_2 | 0.006911 | 0.036578 | 0.49264 | -1.0214 | 0.017362 | 0.090528 | 0.57928 | -0.78767 | 0.001662 | 0.012198 | 0.45046 | -1.1505 |
| PE181/204 | 0.002831 | 0.027101 | 0.4691 | -1.092 | 0.004785 | 0.06511 | 0.45384 | -1.1397 | 0.048329 | 0.094519 | 0.62316 | -0.68233 |
| TAG523-FA181 | 0.10189 | 0.19394 | 0.72148 | -0.47097 | 0.60789 | 0.78054 | 0.91078 | -0.13483 | 0.026074 | 0.061725 | 0.6226 | -0.68362 |
| TAG503-FA160 | 0.027932 | 0.08046 | 0.43141 | -1.2129 | 0.55295 | 0.73614 | 0.85594 | -0.22441 | 0.011914 | 0.037109 | 0.36627 | -1.449 |
| TAG483-FA182 | 0.048294 | 0.12 | 0.36239 | -1.4644 | 0.49785 | 0.7061 | 0.73733 | -0.43962 | 0.013746 | 0.04115 | 0.25843 | -1.9521 |
| PC140/161 | 0.009484 | 0.043341 | 0.52155 | -0.93912 | 0.056658 | 0.17733 | 0.62675 | -0.67404 | 0.028324 | 0.065155 | 0.60928 | -0.71482 |
| TAG523-FA161 | 0.049259 | 0.12183 | 0.43709 | -1.194 | 0.45669 | 0.6666 | 0.72952 | -0.45498 | 0.003917 | 0.019919 | 0.29322 | -1.77 |
| TAG5610-FA182 | 0.20991 | 0.32949 | 0.79487 | -0.33121 | 0.33602 | 0.54824 | 0.84976 | -0.23488 | 0.00214 | 0.013791 | 0.46535 | -1.1036 |
| PC180/183 | 0.26315 | 0.39201 | 0.82646 | -0.27499 | 0.27439 | 0.48408 | 0.82473 | -0.278 | 0.017289 | 0.04749 | 0.52424 | -0.9317 |
| TAG564-FA201 | 0.55247 | 0.67226 | 0.81608 | -0.29322 | 0.57557 | 0.7524 | 0.8108 | -0.30259 | 0.024896 | 0.0597 | 0.50713 | -0.97958 |
| PE182/161 | 0.010952 | 0.045027 | 0.51279 | -0.96356 | 0.029882 | 0.11815 | 0.56814 | -0.81567 | 0.024944 | 0.0597 | 0.5235 | -0.93373 |
| PE181/226 | 0.006985 | 0.03667 | 0.44208 | -1.1776 | 0.005277 | 0.066068 | 0.42891 | -1.2212 | 0.088992 | 0.15126 | 0.61086 | -0.71108 |
| TAG481-FA140 | 0.10085 | 0.19341 | 0.47292 | -1.0803 | 0.87159 | 0.92865 | 1.1532 | 0.20568 | 0.030887 | 0.07006 | 0.37313 | -1.4223 |
| PS161/182_2 | 0.13879 | 0.24619 | 0.73804 | -0.43823 | 0.17829 | 0.37685 | 0.75532 | -0.40485 | 0.004261 | 0.02052 | 0.53238 | -0.90948 |

| | | | | | | | | | | | | |
|---|---|---|---|---|---|---|---|---|---|---|---|---|
| PC140/181 | 0.005579 | 0.033946 | 0.55434 | -0.85115 | 0.022644 | 0.10446 | 0.66249 | -0.59402 | 0.014308 | 0.041214 | 0.63673 | -0.65124 |
| PI160/161_2 | 0.000921 | 0.021852 | 0.48458 | -1.0452 | 0.19233 | 0.39748 | 0.81445 | -0.2961 | 0.015316 | 0.04354 | 0.63908 | -0.64594 |
| PA180/182 | 0.56601 | 0.6811 | 0.8804 | -0.18377 | 0.3443 | 0.55756 | 0.78654 | -0.3464 | 0.8718 | 0.90594 | 1.005 | 0.007171 |
| TAG504-FA160 | 0.04649 | 0.11596 | 0.45969 | -1.1213 | 0.53172 | 0.72202 | 0.81705 | -0.29151 | 0.001919 | 0.013144 | 0.25183 | -1.9895 |
| PC182/225 | 0.001559 | 0.022041 | 0.5069 | -0.98022 | 0.048799 | 0.16291 | 0.70052 | -0.5135 | 0.1173 | 0.184 | 0.7724 | -0.37257 |
| TAG523-FA160 | 0.087298 | 0.18042 | 0.70468 | -0.50495 | 0.36733 | 0.58325 | 0.86403 | -0.21084 | 0.053154 | 0.10118 | 0.69332 | -0.52841 |
| PI180/202 | 0.006596 | 0.035833 | 0.50798 | -0.97716 | 0.015304 | 0.086632 | 0.59086 | -0.75911 | 0.002215 | 0.013864 | 0.46476 | -1.1054 |
| PS180/181_2 | 0.010084 | 0.044356 | 0.5664 | -0.82011 | 0.018986 | 0.092363 | 0.6122 | -0.70793 | 0.052495 | 0.10076 | 0.63342 | -0.65877 |
| SMd181/203 | 0.006439 | 0.035833 | 1.1317 | 0.17843 | 0.42261 | 0.63832 | 1.0531 | 0.074631 | 0.97703 | 0.9846 | 0.99764 | -0.00341 |
| PE181/202 | 0.012315 | 0.047921 | 0.4856 | -1.0422 | 0.004662 | 0.06511 | 0.41332 | -1.2747 | 0.001184 | 0.010394 | 0.36132 | -1.4687 |
| TAG484-FA182 | 0.17668 | 0.2972 | 0.6904 | -0.53449 | 0.25832 | 0.47238 | 0.71915 | -0.47563 | 0.004115 | 0.020451 | 0.34134 | -1.5507 |
| PI180/205 | 0.088675 | 0.18096 | 0.70977 | -0.49457 | 0.19483 | 0.40138 | 0.77021 | -0.37667 | 0.008391 | 0.028936 | 0.58291 | -0.77865 |
| PEP-180/182 | 0.025079 | 0.074497 | 0.54938 | -0.86412 | 0.033166 | 0.12553 | 0.57455 | -0.7995 | 0.017144 | 0.047291 | 0.51198 | -0.96583 |
| PE181/182 | 0.005715 | 0.033965 | 0.4535 | -1.1408 | 0.006641 | 0.071506 | 0.44074 | -1.182 | 0.000563 | 0.006392 | 0.38385 | -1.3814 |
| PEO-160/225 | 0.26622 | 0.3932 | 1.2178 | 0.28428 | 0.64518 | 0.80955 | 1.0706 | 0.098392 | 0.044116 | 0.088915 | 1.2847 | 0.36139 |
| PI180/203 | 0.004348 | 0.031329 | 0.61648 | -0.69787 | 0.047426 | 0.15915 | 0.76548 | -0.38557 | 0.00487 | 0.022017 | 0.63237 | -0.66115 |
| LPC226 | 0.68172 | 0.77903 | 0.91005 | -0.13599 | 0.6454 | 0.80955 | 1.1122 | 0.15336 | 0.18074 | 0.26031 | 1.3199 | 0.40039 |
| PEP-180/181 | 0.097378 | 0.19037 | 0.60837 | -0.71698 | 0.094481 | 0.25003 | 0.61012 | -0.71285 | 0.15216 | 0.22513 | 0.65602 | -0.60819 |
| TAG569-FA204 | 0.058293 | 0.13554 | 0.76125 | -0.39356 | 0.025032 | 0.1066 | 0.68336 | -0.54929 | 0.031328 | 0.07057 | 0.62525 | -0.67751 |
| PC140/182 | 0.004524 | 0.031329 | 0.55188 | -0.85757 | 0.039223 | 0.13728 | 0.74345 | -0.42769 | 0.026697 | 0.062743 | 0.69851 | -0.51765 |
| TAG544-FA182 | 0.1931 | 0.31194 | 0.76249 | -0.39121 | 0.36155 | 0.57689 | 0.83923 | -0.25286 | 0.01051 | 0.033539 | 0.50162 | -0.99532 |
| TAG522-FA181 | 0.1601 | 0.27285 | 0.71505 | -0.48388 | 0.8025 | 0.90229 | 1.0353 | 0.050087 | 0.090239 | 0.15298 | 0.65564 | -0.60902 |
| PEP-180/203 | 0.002875 | 0.027101 | 0.70311 | -0.50817 | 0.000964 | 0.03137 | 0.67292 | -0.5715 | 0.005734 | 0.023333 | 0.68233 | -0.55145 |
| PC150/182 | 0.001322 | 0.022041 | 0.53371 | -0.90587 | 0.000281 | 0.019518 | 0.56813 | -0.81571 | 0.000977 | 0.009218 | 0.58542 | -0.77245 |
| TAG566-FA182 | 0.00774 | 0.039366 | 0.63992 | -0.64404 | 0.25243 | 0.46819 | 0.83335 | -0.263 | 0.074986 | 0.13123 | 0.69971 | -0.51517 |
| PC182/201 | 0.00041 | 0.018752 | 0.56568 | -0.82193 | 0.000693 | 0.029996 | 0.53248 | -0.90921 | 0.1218 | 0.18925 | 0.78574 | -0.34788 |
| TAG567-FA182 | 0.065324 | 0.14564 | 0.67088 | -0.57587 | 0.10667 | 0.26811 | 0.72342 | -0.4671 | 0.034194 | 0.074175 | 0.57954 | -0.78702 |
| PC160/205 | 0.75558 | 0.84082 | 1.2253 | 0.29315 | 0.78369 | 0.89118 | 0.98632 | -0.01987 | 0.84546 | 0.88773 | 1.1008 | 0.1386 |
| TAG568-FA183 | 0.003878 | 0.030702 | 0.39127 | -1.3538 | 0.007228 | 0.072759 | 0.58063 | -0.7843 | 0.000222 | 0.00465 | 0.34831 | -1.5216 |
| TAG524-FA203 | 0.068256 | 0.14961 | 0.53523 | -0.90178 | 0.48401 | 0.6971 | 0.79652 | -0.32823 | 0.007849 | 0.027849 | 0.38716 | -1.369 |
| HCER180 | 0.024579 | 0.073807 | 1.6368 | 0.7109 | 0.1268 | 0.29795 | 1.2389 | 0.309 | 0.87854 | 0.91071 | 0.99435 | -0.00818 |
| TAG567-FA203 | 0.013929 | 0.051358 | 0.66446 | -0.58975 | 0.014456 | 0.085768 | 0.62911 | -0.66862 | 0.000605 | 0.006467 | 0.49944 | -1.0016 |
| PC170/203 | 0.043592 | 0.10957 | 0.66251 | -0.59398 | 0.006529 | 0.071506 | 0.54714 | -0.87001 | 0.038854 | 0.081069 | 0.65768 | -0.60454 |

| | | | | | | | | | | | | |
|---|---|---|---|---|---|---|---|---|---|---|---|---|
| TAG502-FA180 | 0.29884 | 0.42851 | 0.74466 | -0.42534 | 0.98984 | 0.99442 | 1.1642 | 0.21938 | 0.003448 | 0.0181 | 0.34068 | -1.5535 |
| TAG483-FA120 | 0.073555 | 0.15856 | 0.41598 | -1.2654 | 0.33048 | 0.54192 | 0.61345 | -0.70499 | 0.035215 | 0.075163 | 0.36115 | -1.4693 |
| SMd181/160 | 0.70877 | 0.80106 | 0.98365 | -0.02378 | 0.036441 | 0.13107 | 0.85838 | -0.22031 | 0.0351 | 0.075163 | 1.1494 | 0.20089 |
| TAG544-FA202 | 0.03054 | 0.085421 | 0.65817 | -0.60347 | 0.49714 | 0.7061 | 0.88323 | -0.17914 | 0.017095 | 0.047291 | 0.55236 | -0.85632 |
| TAG502-FA161 | 0.088418 | 0.18096 | 0.45504 | -1.1359 | 0.83162 | 0.9145 | 0.95156 | -0.07164 | 0.034524 | 0.074175 | 0.37003 | -1.4343 |
| PE140/204 | 0.017078 | 0.058825 | 0.6687 | -0.58058 | 0.05656 | 0.17733 | 0.7258 | -0.46235 | 0.027106 | 0.063022 | 0.62704 | -0.67337 |
| TAG563-FA202 | 0.37007 | 0.50507 | 0.80075 | -0.32058 | 0.99736 | 0.99736 | 0.9783 | -0.03165 | 0.11458 | 0.18344 | 0.64168 | -0.64008 |
| TAG569-FA183 | 0.001412 | 0.022041 | 0.52274 | -0.93584 | 0.00244 | 0.044126 | 0.53474 | -0.9031 | 0.000162 | 0.004573 | 0.34706 | -1.5267 |
| LPE204 | 0.26032 | 0.38869 | 1.1822 | 0.24151 | 0.22684 | 0.43719 | 1.1899 | 0.2509 | 0.24273 | 0.32852 | 1.2472 | 0.31866 |
| PI180/224 | 0.014565 | 0.053151 | 0.65894 | -0.60178 | 0.20388 | 0.40863 | 0.84613 | -0.24106 | 0.00145 | 0.011513 | 0.61203 | -0.70833 |
| TAG562-FA180 | 0.09676 | 0.19037 | 0.73889 | -0.43657 | 0.27141 | 0.48143 | 0.82393 | -0.2794 | 0.011434 | 0.035944 | 0.47804 | -1.0648 |
| TAG521-FA161 | 0.28311 | 0.41047 | 0.71568 | -0.48261 | 0.77548 | 0.88426 | 0.95615 | -0.06469 | 0.009071 | 0.03044 | 0.34035 | -1.5549 |
| PS181/182_2 | 0.18094 | 0.30052 | 0.72119 | -0.47154 | 0.24584 | 0.46122 | 0.75155 | -0.41205 | 0.10356 | 0.16859 | 0.61163 | -0.70928 |
| TAG503-FA183 | 0.02268 | 0.070853 | 0.44332 | -1.1736 | 0.434 | 0.651 | 0.86543 | -0.2085 | 0.014069 | 0.04115 | 0.40954 | -1.2879 |
| PE170/181 | 0.041798 | 0.10588 | 0.62047 | -0.68858 | 0.10784 | 0.26899 | 0.64711 | -0.62791 | 0.00023 | 0.00465 | 0.31453 | -1.6687 |
| TAG569-FA205 | 0.6573 | 0.76139 | 0.97554 | -0.03573 | 0.044229 | 0.15075 | 0.56239 | -0.83035 | 0.052508 | 0.10076 | 0.52343 | -0.93394 |
| TAG563-FA181 | 0.19787 | 0.31885 | 0.72165 | -0.47062 | 0.4637 | 0.67532 | 0.82218 | -0.28248 | 0.006456 | 0.025166 | 0.44819 | -1.1578 |
| PE181/183 | 0.18677 | 0.30781 | 0.65823 | -0.60334 | 0.017026 | 0.090528 | 0.4495 | -1.1536 | 0.00144 | 0.011513 | 0.30001 | -1.7369 |
| TAG481-FA160 | 0.13465 | 0.24082 | 0.52405 | -0.93223 | 0.73077 | 0.85872 | 1.3709 | 0.4551 | 0.048218 | 0.094519 | 0.42034 | -1.2504 |
| TAG542-FA160 | 0.028875 | 0.081728 | 0.62957 | -0.66756 | 0.8593 | 0.92865 | 0.96928 | -0.04502 | 0.026438 | 0.06236 | 0.57924 | -0.78776 |
| TAG546-FA203 | 0.024803 | 0.074069 | 0.5504 | -0.86145 | 0.079027 | 0.22535 | 0.63545 | -0.65414 | 0.000127 | 0.004481 | 0.30355 | -1.72 |
| TAG567-FA224 | 0.00957 | 0.043341 | 0.49561 | -1.0127 | 0.52855 | 0.72202 | 0.87041 | -0.20023 | 0.018627 | 0.049393 | 0.51814 | -0.94858 |
| TAG501-FA180 | 0.33528 | 0.46738 | 0.64649 | -0.6293 | 0.88391 | 0.93652 | 1.1906 | 0.2517 | 0.029084 | 0.066201 | 0.35995 | -1.4741 |
| PC181/161 | 0.003543 | 0.029195 | 0.50172 | -0.99504 | 0.024466 | 0.10618 | 0.6421 | -0.63912 | 0.016356 | 0.0457 | 0.60679 | -0.72073 |
| TAG543-FA180 | 0.57729 | 0.69196 | 0.91793 | -0.12354 | 0.98614 | 0.99439 | 1.0023 | 0.003378 | 0.04471 | 0.089834 | 0.5949 | -0.74928 |
| PC180/204 | 0.19999 | 0.32102 | 1.0872 | 0.12058 | 0.65997 | 0.81992 | 0.97199 | -0.04099 | 0.014066 | 0.04115 | 1.2768 | 0.35253 |
| TAG544-FA181 | 0.15054 | 0.26133 | 0.69092 | -0.53342 | 0.22334 | 0.43401 | 0.74655 | -0.42168 | 0.005014 | 0.022058 | 0.43324 | -1.2067 |
| TAG523-FA200 | 0.040009 | 0.10214 | 0.59726 | -0.74357 | 0.99318 | 0.99624 | 0.99512 | -0.00705 | 0.11649 | 0.184 | 0.66761 | -0.58293 |
| PI180/224_2 | 0.028473 | 0.081298 | 0.6382 | -0.64792 | 0.29343 | 0.50804 | 0.83206 | -0.26523 | 0.00779 | 0.027849 | 0.61149 | -0.70959 |
| LPC204 | 0.66267 | 0.76625 | 1.0251 | 0.035756 | 0.50823 | 0.71153 | 1.0906 | 0.12518 | 0.033793 | 0.074038 | 1.4831 | 0.56859 |
| TAG522-FA161 | 0.28204 | 0.40984 | 0.6533 | -0.61418 | 0.97769 | 0.9914 | 0.96433 | -0.05241 | 0.010162 | 0.033052 | 0.33359 | -1.5839 |
| TAG543-FA160 | 0.10174 | 0.19394 | 0.71922 | -0.47549 | 0.95002 | 0.98327 | 1.0288 | 0.04098 | 0.048348 | 0.094519 | 0.62977 | -0.66711 |
| TAG544-FA221 | 0.39244 | 0.53004 | 0.72892 | -0.45618 | 0.56984 | 0.74792 | 0.76877 | -0.37938 | 0.01252 | 0.038672 | 0.38743 | -1.368 |

| | | | | | | | | | | | | |
|---|---|---|---|---|---|---|---|---|---|---|---|---|
| TAG542-FA182 | 0.38942 | 0.52705 | 0.85097 | -0.23281 | 0.99639 | 0.99736 | 1.014 | 0.02001 | 0.018274 | 0.049363 | 0.51219 | -0.96525 |
| PC170/181 | 0.008582 | 0.041386 | 0.67769 | -0.5613 | 0.008915 | 0.07537 | 0.74998 | -0.41507 | 0.004709 | 0.021438 | 0.75194 | -0.41131 |
| TAG567-FA161 | 0.03286 | 0.088792 | 0.47658 | -1.0692 | 0.50784 | 0.71153 | 0.77316 | -0.37116 | 0.062154 | 0.1143 | 0.52835 | -0.92044 |
| PE180/225 | 0.80427 | 0.87324 | 1.0409 | 0.057842 | 0.28008 | 0.49015 | 0.89149 | -0.1657 | 0.17371 | 0.25175 | 0.84379 | -0.24504 |
| PG182/182 | 0.002241 | 0.026528 | 0.46283 | -1.1114 | 0.00738 | 0.072759 | 0.43541 | -1.1996 | 6E-05 | 0.004338 | 0.31231 | -1.679 |
| PE181/225 | 0.002371 | 0.026861 | 0.57074 | -0.8091 | 0.000935 | 0.03137 | 0.50559 | -0.98397 | 0.003072 | 0.017238 | 0.56492 | -0.82387 |
| TAG543-FA182 | 0.18938 | 0.309 | 0.75034 | -0.41437 | 0.8383 | 0.91875 | 1.0236 | 0.03364 | 0.032806 | 0.072641 | 0.57327 | -0.80272 |
| PS180/182 | 0.008982 | 0.042492 | 0.63025 | -0.666 | 0.011273 | 0.081295 | 0.71447 | -0.48505 | 0.000606 | 0.006467 | 0.5519 | -0.85753 |
| SMd181/202 | 0.057318 | 0.13471 | 1.2425 | 0.31324 | 0.30432 | 0.51709 | 1.1287 | 0.17472 | 0.052937 | 0.10106 | 1.1625 | 0.21726 |
| PE160/225 | 0.75391 | 0.84041 | 1.0584 | 0.081872 | 0.32359 | 0.5387 | 0.86701 | -0.20588 | 0.24987 | 0.33748 | 0.84137 | -0.2492 |
| TAG480-FA140 | 0.21004 | 0.32949 | 0.5678 | -0.81656 | 0.9225 | 0.96263 | 1.27 | 0.34487 | 0.25683 | 0.34202 | 0.58981 | -0.76167 |
| PC140/140 | 0.030742 | 0.085525 | 0.53123 | -0.9126 | 0.21201 | 0.41824 | 0.74958 | -0.41585 | 0.034242 | 0.074175 | 0.59662 | -0.74511 |
| TAG482-FA181 | 0.17929 | 0.30052 | 0.47298 | -1.0802 | 0.62967 | 0.79524 | 0.78337 | -0.35224 | 0.019652 | 0.050677 | 0.25994 | -1.9437 |
| TAG564-FA202 | 0.10101 | 0.19341 | 0.74647 | -0.42185 | 0.23011 | 0.44059 | 0.80754 | -0.3084 | 0.008187 | 0.028503 | 0.50955 | -0.97271 |
| TAG543-FA181 | 0.31772 | 0.4477 | 0.74618 | -0.4224 | 0.8677 | 0.92865 | 0.98606 | -0.02026 | 0.050998 | 0.098808 | 0.54032 | -0.88812 |
| TAG564-FA181 | 0.0973 | 0.19037 | 0.66118 | -0.59688 | 0.23671 | 0.44796 | 0.76745 | -0.38185 | 0.007524 | 0.027062 | 0.49352 | -1.0188 |
| PE140/182 | 0.011564 | 0.0465 | 0.58901 | -0.76364 | 0.03082 | 0.11943 | 0.61573 | -0.69964 | 0.002645 | 0.015751 | 0.45601 | -1.1328 |
| TAG543-FA161 | 0.018068 | 0.060631 | 0.46538 | -1.1035 | 0.22587 | 0.43719 | 0.71338 | -0.48725 | 0.000569 | 0.006392 | 0.30998 | -1.6898 |
| TAG504-FA181 | 0.095061 | 0.19037 | 0.52496 | -0.92971 | 0.39348 | 0.60989 | 0.73427 | -0.44563 | 0.001863 | 0.013044 | 0.26147 | -1.9353 |
| TAG568-FA181 | 0.43413 | 0.567 | 0.88855 | -0.17048 | 0.071908 | 0.21182 | 0.70271 | -0.509 | 0.005576 | 0.023333 | 0.44103 | -1.1811 |
| PEP-180/202 | 0.10779 | 0.20222 | 0.67024 | -0.57725 | 0.023427 | 0.10446 | 0.56233 | -0.8305 | 0.01394 | 0.04115 | 0.51591 | -0.95481 |
| PE160/203 | 0.16605 | 0.28224 | 0.651 | -0.61926 | 0.18767 | 0.3941 | 0.65584 | -0.60859 | 0.05531 | 0.10407 | 0.52892 | -0.91887 |
| TAG565-FA181 | 0.084684 | 0.1767 | 0.65436 | -0.61185 | 0.43511 | 0.65116 | 0.82183 | -0.28309 | 0.065542 | 0.11819 | 0.60255 | -0.73086 |
| TAG564-FA182 | 0.087614 | 0.1805 | 0.70214 | -0.51017 | 0.12236 | 0.29286 | 0.75623 | -0.40311 | 0.002686 | 0.015751 | 0.47037 | -1.0881 |
| TAG566-FA203 | 0.022841 | 0.070853 | 0.60467 | -0.72577 | 0.014003 | 0.085768 | 0.59581 | -0.74708 | 0.000337 | 0.00519 | 0.42721 | -1.227 |
| PEP-160/160 | 0.59726 | 0.70589 | 1.211 | 0.27624 | 0.8452 | 0.92475 | 0.9768 | -0.03387 | 0.69452 | 0.76503 | 1.2152 | 0.28125 |
| TAG522-FA160 | 0.18058 | 0.30052 | 0.71581 | -0.48236 | 0.90489 | 0.95013 | 0.97877 | -0.03096 | 0.06954 | 0.12369 | 0.62929 | -0.6682 |
| LPE224 | 0.40787 | 0.54411 | 1.1453 | 0.19575 | 0.71602 | 0.8506 | 1.055 | 0.077217 | 0.27343 | 0.35995 | 1.2159 | 0.28204 |
| PI161/180_2 | 0.010786 | 0.045027 | 0.68104 | -0.55418 | 0.067857 | 0.20451 | 0.742 | -0.43051 | 0.000399 | 0.005336 | 0.53201 | -0.91047 |
| PEO-160/205 | 0.27134 | 0.39965 | 0.74705 | -0.42073 | 0.13184 | 0.30342 | 0.65815 | -0.6035 | 0.050719 | 0.098561 | 0.53808 | -0.8941 |
| TAG568-FA204 | 0.96293 | 0.97895 | 1.038 | 0.053786 | 0.49407 | 0.7038 | 0.94752 | -0.07777 | 0.46748 | 0.5615 | 0.93504 | -0.0969 |
| PC150/181 | 0.001905 | 0.023843 | 0.55732 | -0.84342 | 0.005101 | 0.06511 | 0.63795 | -0.64849 | 0.004274 | 0.02052 | 0.59567 | -0.74742 |
| TAG546-FA161 | 0.010154 | 0.044365 | 0.44163 | -1.1791 | 0.19065 | 0.39565 | 0.69083 | -0.53359 | 0.015407 | 0.043609 | 0.41453 | -1.2704 |

| | | | | | | | | | | | | |
|---|---|---|---|---|---|---|---|---|---|---|---|---|
| TAG522-FA182 | 0.14663 | 0.25729 | 0.73025 | -0.45353 | 0.88976 | 0.93727 | 1.059 | 0.082725 | 0.061097 | 0.11299 | 0.6287 | -0.66956 |
| LPC140 | 0.026909 | 0.078204 | 0.70018 | -0.5142 | 0.45365 | 0.66512 | 0.91721 | -0.12468 | 0.003275 | 0.017847 | 0.6843 | -0.54729 |
| TAG501-FA140 | 0.2007 | 0.32102 | 0.58452 | -0.77468 | 0.94448 | 0.98063 | 1.0417 | 0.058996 | 0.014159 | 0.04115 | 0.32888 | -1.6044 |
| PEP-180/183 | 0.095796 | 0.19037 | 0.59779 | -0.74229 | 0.11135 | 0.27458 | 0.60927 | -0.71486 | 0.010205 | 0.033052 | 0.38079 | -1.3929 |
| TAG543-FA202 | 0.03229 | 0.088331 | 0.61086 | -0.71108 | 0.57264 | 0.75007 | 0.89694 | -0.15691 | 0.023047 | 0.056832 | 0.56074 | -0.83459 |
| PC180/140 | 0.025721 | 0.075765 | 0.70642 | -0.50139 | 0.096859 | 0.25426 | 0.81686 | -0.29184 | 0.006643 | 0.025472 | 0.79805 | -0.32546 |
| TAG568-FA225 | 0.011882 | 0.047456 | 0.56754 | -0.8172 | 0.11199 | 0.27512 | 0.74646 | -0.42186 | 0.004354 | 0.020539 | 0.45167 | -1.1467 |
| SMd181/161 | 0.5507 | 0.67226 | 1.0695 | 0.096908 | 0.79735 | 0.89805 | 1.0273 | 0.038867 | 0.021746 | 0.054448 | 1.3944 | 0.47969 |
| PEP-180/205 | 0.067073 | 0.14751 | 0.57591 | -0.79607 | 0.032099 | 0.12292 | 0.52803 | -0.9213 | 0.043295 | 0.087532 | 0.49337 | -1.0193 |
| PEP-160/182 | 0.12828 | 0.23198 | 0.65529 | -0.60978 | 0.14783 | 0.33301 | 0.67355 | -0.57015 | 0.40295 | 0.49941 | 0.81266 | -0.29927 |
| SMd181/200 | 0.22409 | 0.34406 | 0.8976 | -0.15585 | 0.53164 | 0.72202 | 0.94058 | -0.08838 | 0.11553 | 0.18344 | 1.1241 | 0.16875 |
| TAG541-FA181 | 0.40329 | 0.54021 | 0.82144 | -0.28377 | 0.81216 | 0.90706 | 1.0394 | 0.055696 | 0.033417 | 0.073743 | 0.54472 | -0.87641 |
| TAG565-FA203 | 0.021839 | 0.069692 | 0.7055 | -0.50328 | 0.26101 | 0.47464 | 0.82666 | -0.27464 | 0.045144 | 0.090427 | 0.63137 | -0.66345 |
| PG160/160 | 0.15355 | 0.26573 | 0.57864 | -0.78927 | 0.3006 | 0.51363 | 0.65237 | -0.61625 | 0.35167 | 0.44282 | 0.80179 | -0.3187 |
| PEP-160/225 | 0.11091 | 0.2064 | 1.1682 | 0.22425 | 0.61114 | 0.78233 | 1.0567 | 0.079558 | 0.019701 | 0.050677 | 1.3285 | 0.40975 |
| PC170/226 | 0.031353 | 0.086854 | 0.71393 | -0.48615 | 0.011176 | 0.081295 | 0.67768 | -0.56133 | 0.11493 | 0.18344 | 0.79288 | -0.33483 |
| TAG522-FA180 | 0.53836 | 0.66503 | 0.87151 | -0.19842 | 0.58335 | 0.75952 | 1.2121 | 0.2775 | 0.13736 | 0.20892 | 0.67169 | -0.57414 |
| TAG545-FA203 | 0.005578 | 0.033946 | 0.6024 | -0.73119 | 0.10127 | 0.2603 | 0.71935 | -0.47524 | 0.00695 | 0.025783 | 0.52454 | -0.93088 |
| PC181/182 | 0.004519 | 0.031329 | 0.64634 | -0.62964 | 7.9E-05 | 0.017442 | 0.56118 | -0.83346 | 0.17128 | 0.24889 | 0.82603 | -0.27574 |
| TAG548-FA205 | 0.20361 | 0.32408 | 0.71899 | -0.47595 | 0.050466 | 0.16461 | 0.57918 | -0.78791 | 0.004552 | 0.021321 | 0.36459 | -1.4556 |
| TAG562-FA160 | 0.20021 | 0.32102 | 0.7371 | -0.44006 | 0.96967 | 0.98629 | 1.106 | 0.14539 | 0.04317 | 0.087532 | 0.61481 | -0.70179 |
| PG160/180_2 | 0.41773 | 0.55386 | 0.90621 | -0.14209 | 0.069264 | 0.20652 | 0.67524 | -0.56653 | 0.014147 | 0.04115 | 0.58633 | -0.77021 |
| PEP-160/205 | 0.50333 | 0.63821 | 0.80155 | -0.31913 | 0.32552 | 0.53921 | 0.72785 | -0.45829 | 0.25278 | 0.34 | 0.71028 | -0.49354 |
| TAG543-FA201 | 0.071844 | 0.1559 | 0.67775 | -0.56118 | 0.82386 | 0.90904 | 0.9715 | -0.04171 | 0.091495 | 0.15471 | 0.67679 | -0.56322 |
| PEP-181/182 | 0.054727 | 0.13098 | 0.69744 | -0.51986 | 0.044603 | 0.15123 | 0.69131 | -0.5326 | 0.21333 | 0.29802 | 0.79324 | -0.33416 |
| PG160/180 | 0.51784 | 0.64842 | 1.006 | 0.00866 | 0.042314 | 0.14652 | 0.59202 | -0.75628 | 0.008401 | 0.028936 | 0.51065 | -0.96959 |
| PEO-160/182 | 0.003267 | 0.028391 | 0.59084 | -0.75915 | 0.004455 | 0.064444 | 0.58276 | -0.77903 | 0.004332 | 0.020539 | 0.54928 | -0.86437 |
| TAG483-FA160 | 0.32387 | 0.45538 | 0.66284 | -0.59326 | 0.85487 | 0.92847 | 0.99787 | -0.00308 | 0.023475 | 0.057453 | 0.31622 | -1.661 |
| TAG547-FA161 | 0.026511 | 0.077393 | 0.57045 | -0.80983 | 0.41634 | 0.63532 | 0.85775 | -0.22137 | 0.003719 | 0.019213 | 0.41389 | -1.2727 |
| TAG565-FA224 | 0.30282 | 0.43231 | 0.71259 | -0.48886 | 0.65097 | 0.8134 | 1.1253 | 0.17025 | 0.35636 | 0.44786 | 0.7407 | -0.43304 |
| TAG502-FA160 | 0.022615 | 0.070853 | 0.5441 | -0.87806 | 0.67477 | 0.82838 | 0.95411 | -0.06777 | 0.22916 | 0.3154 | 0.75481 | -0.40581 |
| PE180/150 | 0.045054 | 0.11281 | 0.79002 | -0.34003 | 0.03724 | 0.13248 | 0.79943 | -0.32296 | 0.036302 | 0.076979 | 0.7365 | -0.44125 |
| LPE180 | 0.017037 | 0.058825 | 1.3852 | 0.47011 | 0.11113 | 0.27458 | 1.1774 | 0.23561 | 0.99881 | 0.99881 | 1.028 | 0.039816 |

| | | | | | | | | | | | | |
|---|---|---|---|---|---|---|---|---|---|---|---|---|
| TAG502-FA182 | 0.008416 | 0.041003 | 0.54625 | -0.87238 | 0.84997 | 0.92819 | 0.98493 | -0.02191 | 0.27897 | 0.36472 | 0.80386 | -0.31499 |
| PEO-180/224 | 0.74939 | 0.8368 | 1.0804 | 0.11155 | 0.87008 | 0.92865 | 0.96102 | -0.05737 | 0.63828 | 0.71889 | 1.0728 | 0.10139 |
| TAG483-FA181 | 0.29656 | 0.42619 | 0.77209 | -0.37316 | 0.95121 | 0.98327 | 1.0248 | 0.035391 | 0.031791 | 0.070877 | 0.44308 | -1.1744 |
| PE182/183 | 0.030573 | 0.085421 | 0.42253 | -1.2429 | 0.006701 | 0.071506 | 0.28438 | -1.8141 | 0.000737 | 0.007501 | 0.2001 | -2.3212 |
| PG160/181 | 0.055763 | 0.13297 | 0.65408 | -0.61247 | 0.069792 | 0.20652 | 0.61868 | -0.69273 | 0.024878 | 0.0597 | 0.56791 | -0.81627 |
| PEP-180/180 | 0.43461 | 0.567 | 0.78346 | -0.35207 | 0.30864 | 0.52052 | 0.71545 | -0.48308 | 0.51179 | 0.61021 | 0.82566 | -0.27639 |
| PS160/180 | 0.74768 | 0.83632 | 1.0251 | 0.035781 | 0.44125 | 0.65696 | 0.86496 | -0.2093 | 0.36556 | 0.45853 | 0.85495 | -0.22609 |
| PG180/181_2 | 0.004446 | 0.031329 | 0.62763 | -0.67201 | 0.028982 | 0.11792 | 0.68067 | -0.55498 | 0.002197 | 0.013864 | 0.51265 | -0.96395 |
| PC160/160 | 0.13105 | 0.23567 | 1.265 | 0.33919 | 0.3376 | 0.54945 | 1.1496 | 0.20111 | 0.075368 | 0.13154 | 1.3412 | 0.4235 |
| TAG562-FA201 | 0.25821 | 0.38643 | 0.71784 | -0.47827 | 0.67655 | 0.82838 | 0.85658 | -0.22333 | 0.027088 | 0.063022 | 0.47392 | -1.0773 |
| PEO-160/224 | 0.52082 | 0.65077 | 1.1415 | 0.19092 | 0.96135 | 0.98327 | 0.95307 | -0.06935 | 0.95436 | 0.97228 | 0.97696 | -0.03363 |
| PE170/204 | 0.000435 | 0.018752 | 0.51901 | -0.94616 | 0.003949 | 0.059992 | 0.48399 | -1.0469 | 0.000295 | 0.005048 | 0.40978 | -1.2871 |
| TAG566-FA224 | 0.20905 | 0.32949 | 0.74023 | -0.43395 | 0.86427 | 0.92865 | 1.0162 | 0.023221 | 0.11723 | 0.184 | 0.68179 | -0.55259 |
| PE182/182 | 0.018561 | 0.061964 | 0.46912 | -1.092 | 0.017286 | 0.090528 | 0.45008 | -1.1518 | 0.001339 | 0.011037 | 0.31965 | -1.6455 |
| PC181/225 | 0.0026 | 0.026861 | 0.71635 | -0.48127 | 0.000236 | 0.019518 | 0.62857 | -0.66984 | 0.53166 | 0.62588 | 0.94767 | -0.07754 |
| PI161/180 | 0.010801 | 0.045027 | 0.66539 | -0.58772 | 0.36625 | 0.58296 | 0.88704 | -0.17293 | 0.001603 | 0.012198 | 0.58618 | -0.7706 |
| LPE225 | 0.034096 | 0.091343 | 1.5883 | 0.6675 | 0.034547 | 0.12837 | 1.6122 | 0.68904 | 0.077262 | 0.13421 | 1.5347 | 0.61799 |
| HCER200 | 0.016329 | 0.057153 | 1.5117 | 0.59619 | 0.26857 | 0.48143 | 1.1457 | 0.19624 | 0.55476 | 0.64288 | 0.90374 | -0.14601 |
| LPC203 | 0.013964 | 0.051358 | 0.6923 | -0.53053 | 0.13425 | 0.30773 | 0.77921 | -0.35992 | 0.99867 | 0.99881 | 1.025 | 0.035667 |
| TAG544-FA160 | 0.031646 | 0.087293 | 0.66318 | -0.59253 | 0.85384 | 0.92847 | 0.99565 | -0.00629 | 0.010359 | 0.033386 | 0.56512 | -0.82338 |
| PE180/203 | 0.10623 | 0.20044 | 0.72228 | -0.46937 | 0.034608 | 0.12837 | 0.61419 | -0.70324 | 0.19169 | 0.27307 | 0.73943 | -0.43551 |
| TAG546-FA204 | 0.80332 | 0.87324 | 0.95759 | -0.06252 | 0.68308 | 0.83119 | 1.1965 | 0.25885 | 0.67235 | 0.75077 | 0.97193 | -0.04107 |
| TAG568-FA161 | 0.020686 | 0.067672 | 0.52511 | -0.92932 | 0.064918 | 0.19712 | 0.67945 | -0.55756 | 0.001604 | 0.012198 | 0.38729 | -1.3685 |
| PEO-180/182 | 0.001594 | 0.022041 | 0.52257 | -0.9363 | 0.005587 | 0.067354 | 0.54696 | -0.87048 | 0.000152 | 0.004573 | 0.40848 | -1.2917 |
| PS180/180 | 0.55126 | 0.67226 | 1.0928 | 0.128 | 0.66989 | 0.82751 | 0.95293 | -0.06956 | 0.33113 | 0.42267 | 0.87689 | -0.18953 |
| TAG561-FA160 | 0.025176 | 0.074497 | 0.52142 | -0.93949 | 0.38066 | 0.59778 | 0.85205 | -0.231 | 0.005169 | 0.022185 | 0.47483 | -1.0745 |
| PEP-160/203 | 0.60344 | 0.71163 | 0.93754 | -0.09305 | 0.37671 | 0.59523 | 0.87376 | -0.19469 | 0.98459 | 0.99068 | 1.0632 | 0.088478 |
| TAG563-FA201 | 0.60972 | 0.71518 | 0.80255 | -0.31735 | 0.95246 | 0.98327 | 0.89869 | -0.15411 | 0.14819 | 0.22126 | 0.60545 | -0.72393 |
| PE181/224 | 0.076169 | 0.16205 | 0.63629 | -0.65223 | 0.026402 | 0.11161 | 0.53012 | -0.91561 | 0.063916 | 0.11623 | 0.61629 | -0.69833 |
| TAG521-FA180 | 0.27358 | 0.40163 | 0.73185 | -0.45038 | 0.81279 | 0.90706 | 1.1373 | 0.18566 | 0.13815 | 0.20964 | 0.63771 | -0.64903 |
| TAG547-FA205 | 0.56075 | 0.67727 | 0.95332 | -0.06897 | 0.27021 | 0.48143 | 0.76747 | -0.38182 | 0.079673 | 0.13721 | 0.57637 | -0.79493 |
| PG160/181_2 | 0.061025 | 0.13939 | 0.69095 | -0.53335 | 0.081584 | 0.22893 | 0.65046 | -0.62047 | 0.019102 | 0.050143 | 0.56924 | -0.8129 |
| PE170/182 | 0.000106 | 0.018752 | 0.44112 | -1.1808 | 0.002138 | 0.042361 | 0.43541 | -1.1996 | 1.3E-06 | 0.000847 | 0.27462 | -1.8645 |

| | | | | | | | | | | | | |
|---|---|---|---|---|---|---|---|---|---|---|---|---|
| PC200/203 | 0.0389 | 0.10003 | 0.74371 | -0.42719 | 0.000288 | 0.019518 | 0.56152 | -0.83258 | 0.027837 | 0.06449 | 0.7217 | -0.47053 |
| TAG547-FA204 | 0.022856 | 0.070853 | 0.60888 | -0.71577 | 0.098966 | 0.25629 | 0.6844 | -0.54708 | 0.004287 | 0.02052 | 0.4991 | -1.0026 |
| TAG526-FA204 | 0.54653 | 0.6713 | 0.87284 | -0.19622 | 0.66156 | 0.82034 | 0.96891 | -0.04556 | 0.028263 | 0.065155 | 0.52149 | -0.93928 |
| TAG542-FA181 | 0.99605 | 0.99758 | 0.98586 | -0.02055 | 0.25983 | 0.4738 | 1.2496 | 0.32141 | 0.084935 | 0.14512 | 0.58599 | -0.77105 |
| PI180/183 | 0.004709 | 0.031829 | 0.57074 | -0.80909 | 0.022542 | 0.10446 | 0.67409 | -0.56899 | 5.36E-05 | 0.004338 | 0.40723 | -1.2961 |
| TAG520-FA160 | 0.14877 | 0.26004 | 0.63672 | -0.65127 | 0.75702 | 0.87534 | 0.93481 | -0.09726 | 0.033891 | 0.074038 | 0.46025 | -1.1195 |
| TAG521-FA181 | 0.27392 | 0.40163 | 0.71848 | -0.47699 | 0.76996 | 0.88247 | 1.0917 | 0.12663 | 0.26143 | 0.34732 | 0.71454 | -0.48491 |
| TAG544-FA203 | 0.063634 | 0.14384 | 0.62326 | -0.6821 | 0.4836 | 0.6971 | 0.83486 | -0.2604 | 0.019757 | 0.050677 | 0.502 | -0.99425 |
| PEO-180/225 | 0.24362 | 0.36797 | 1.1935 | 0.25521 | 0.81407 | 0.90706 | 1.0314 | 0.044671 | 0.059551 | 0.11076 | 1.2907 | 0.36819 |
| TAG546-FA225 | 0.23633 | 0.35946 | 0.71705 | -0.47985 | 0.95635 | 0.98327 | 1.0267 | 0.037985 | 0.09489 | 0.15778 | 0.55457 | -0.85055 |
| TAG542-FA202 | 0.7395 | 0.83003 | 0.91257 | -0.13199 | 0.35275 | 0.56883 | 1.2524 | 0.32475 | 0.1873 | 0.26798 | 0.69403 | -0.52692 |
| TAG564-FA203 | 0.36297 | 0.49957 | 0.83614 | -0.25819 | 0.67887 | 0.82838 | 0.93756 | -0.09301 | 0.17743 | 0.25612 | 0.7492 | -0.41658 |
| PC160/202 | 0.12772 | 0.23159 | 0.86312 | -0.21237 | 0.20463 | 0.40863 | 0.88516 | -0.176 | 0.68508 | 0.75965 | 0.96191 | -0.05603 |
| TAG563-FA180 | 0.1745 | 0.29429 | 0.76791 | -0.38099 | 0.86769 | 0.92865 | 1.0086 | 0.012417 | 0.031559 | 0.070601 | 0.57542 | -0.7973 |
| PE180/160 | 0.99898 | 0.99898 | 1.0071 | 0.010195 | 0.12751 | 0.29795 | 0.80715 | -0.3091 | 0.02202 | 0.054923 | 0.6892 | -0.537 |
| FFA205 | 0.15051 | 0.26133 | 0.69549 | -0.52389 | 0.12559 | 0.29729 | 0.67957 | -0.5573 | 0.2412 | 0.32712 | 0.75121 | -0.4127 |
| PE180/201 | 0.07657 | 0.16237 | 0.7427 | -0.42915 | 0.035391 | 0.12943 | 0.65459 | -0.61133 | 0.008676 | 0.029726 | 0.57715 | -0.79298 |
| PC180/203 | 0.013441 | 0.050289 | 0.72085 | -0.47223 | 0.019887 | 0.095897 | 0.70431 | -0.50571 | 0.54144 | 0.63395 | 0.91484 | -0.12842 |
| PG180/180 | 0.002759 | 0.027101 | 0.67338 | -0.5705 | 0.022953 | 0.10446 | 0.69606 | -0.52271 | 0.001287 | 0.010884 | 0.54503 | -0.87559 |
| PEP-180/204 | 0.29094 | 0.41997 | 0.85087 | -0.23298 | 0.37034 | 0.5866 | 0.87076 | -0.19966 | 0.87254 | 0.90594 | 1.0183 | 0.026226 |
| PEO-180/160 | 0.42592 | 0.56061 | 0.92991 | -0.10484 | 0.19978 | 0.40771 | 0.88461 | -0.17688 | 0.04689 | 0.093315 | 0.80924 | -0.30536 |
| TAG541-FA201 | 0.18162 | 0.30086 | 0.70422 | -0.5059 | 0.62232 | 0.78973 | 1.1398 | 0.18878 | 0.12082 | 0.18861 | 0.64702 | -0.62812 |
| PC180/161 | 0.09074 | 0.18357 | 0.69874 | -0.51718 | 0.38199 | 0.59778 | 0.85961 | -0.21825 | 0.096553 | 0.15994 | 0.71768 | -0.47858 |
| TAG568-FA205 | 0.8984 | 0.94484 | 1.1707 | 0.2274 | 0.068977 | 0.20652 | 0.61396 | -0.70377 | 0.077776 | 0.1343 | 0.55164 | -0.85819 |
| PI180/204 | 0.006761 | 0.036078 | 0.69936 | -0.51589 | 0.22721 | 0.43719 | 0.87632 | -0.19047 | 0.048034 | 0.094519 | 0.86711 | -0.20571 |
| PI180/204_2 | 0.003092 | 0.027956 | 0.66597 | -0.58648 | 0.27627 | 0.48608 | 0.87843 | -0.187 | 0.034415 | 0.074175 | 0.82634 | -0.2752 |
| TAG546-FA205 | 0.67455 | 0.77448 | 1.028 | 0.039826 | 0.47698 | 0.69157 | 0.84673 | -0.24002 | 0.23112 | 0.317 | 0.76009 | -0.39576 |
| PC170/182 | 0.010763 | 0.045027 | 0.67146 | -0.57463 | 0.002157 | 0.042361 | 0.62818 | -0.67075 | 0.008902 | 0.030342 | 0.67993 | -0.55653 |
| TAG542-FA201 | 0.50177 | 0.63799 | 0.80582 | -0.31147 | 0.35747 | 0.57451 | 1.2665 | 0.34086 | 0.51442 | 0.61193 | 0.81464 | -0.29577 |
| TAG521-FA160 | 0.25743 | 0.38614 | 0.73537 | -0.44346 | 0.7426 | 0.86758 | 1.1231 | 0.16749 | 0.13586 | 0.20712 | 0.65209 | -0.61685 |
| TAG482-FA120 | 0.22555 | 0.34548 | 0.70172 | -0.51103 | 0.59399 | 0.76723 | 0.8535 | -0.22853 | 0.011485 | 0.035944 | 0.38497 | -1.3772 |
| TAG564-FA180 | 0.049404 | 0.12183 | 0.68513 | -0.54554 | 0.44365 | 0.65696 | 0.87523 | -0.19226 | 0.14478 | 0.21834 | 0.72782 | -0.45836 |
| LPC180 | 0.17 | 0.28746 | 0.88377 | -0.17826 | 0.54621 | 0.73316 | 0.94756 | -0.07771 | 0.41417 | 0.50984 | 1.1019 | 0.13995 |

| | | | | | | | | | | | | |
|---|---|---|---|---|---|---|---|---|---|---|---|---|
| PC181/205 | 0.17976 | 0.30052 | 0.67332 | -0.57063 | 0.032631 | 0.12423 | 0.50697 | -0.98004 | 0.16753 | 0.24508 | 0.66519 | -0.58816 |
| TAG567-FA205 | 0.96665 | 0.9802 | 1.1778 | 0.23612 | 0.25629 | 0.47238 | 0.76152 | -0.39306 | 0.071548 | 0.12589 | 0.58818 | -0.76567 |
| PE181/201 | 0.026355 | 0.077283 | 0.51566 | -0.9555 | 0.018692 | 0.092363 | 0.48408 | -1.0467 | 0.001792 | 0.012683 | 0.37225 | -1.4256 |
| TAG501-FA161 | 0.20713 | 0.32808 | 0.56768 | -0.81684 | 0.8512 | 0.92819 | 1.2131 | 0.27871 | 0.018658 | 0.049393 | 0.33026 | -1.5983 |
| PG180/181 | 0.004193 | 0.031329 | 0.62475 | -0.67864 | 0.027438 | 0.11524 | 0.66684 | -0.58459 | 0.002287 | 0.014177 | 0.50795 | -0.97725 |
| PI180/205_2 | 0.19308 | 0.31194 | 0.73614 | -0.44194 | 0.26991 | 0.48143 | 0.77132 | -0.3746 | 0.011171 | 0.035304 | 0.5053 | -0.98478 |
| TAG541-FA180 | 0.40296 | 0.54021 | 0.79403 | -0.33274 | 0.88473 | 0.93652 | 1.0271 | 0.038547 | 0.051591 | 0.099661 | 0.53612 | -0.89938 |
| PC200/205 | 0.22352 | 0.344 | 0.6964 | -0.52202 | 0.023588 | 0.10446 | 0.48275 | -1.0506 | 0.036692 | 0.077302 | 0.48445 | -1.0456 |
| PC150/226 | 0.000329 | 0.018752 | 0.45945 | -1.122 | 0.000769 | 0.029996 | 0.4844 | -1.0457 | 0.023034 | 0.056832 | 0.67919 | -0.55811 |
| PC200/181 | 0.074961 | 0.16052 | 0.73301 | -0.4481 | 0.007987 | 0.072759 | 0.62 | -0.68967 | 0.019773 | 0.050677 | 0.65812 | -0.60358 |
| TAG527-FA160 | 0.03703 | 0.096427 | 0.58053 | -0.78455 | 0.54512 | 0.73316 | 0.93623 | -0.09507 | 0.009713 | 0.032098 | 0.43378 | -1.205 |
| LPC181 | 0.12058 | 0.22174 | 0.81011 | -0.3038 | 0.42851 | 0.64424 | 0.90184 | -0.14905 | 0.51511 | 0.61193 | 1.0582 | 0.081588 |
| PE181/181 | 0.33038 | 0.46253 | 0.72563 | -0.4627 | 0.077822 | 0.22346 | 0.55219 | -0.85677 | 0.009534 | 0.031667 | 0.42656 | -1.2292 |
| SMd181/120 | 0.009073 | 0.042492 | 0.51633 | -0.95363 | 0.035886 | 0.13051 | 0.66282 | -0.5933 | 0.019451 | 0.050677 | 1.5848 | 0.66428 |
| PE160/161 | 0.099183 | 0.1916 | 0.54339 | -0.87994 | 0.2658 | 0.4798 | 0.73601 | -0.44221 | 0.007871 | 0.027849 | 0.3632 | -1.4612 |
| PEO-180/183 | 0.023157 | 0.071111 | 0.66488 | -0.58884 | 0.010955 | 0.081295 | 0.5944 | -0.7505 | 0.000331 | 0.00519 | 0.32372 | -1.6272 |
| TAG568-FA182 | 0.5539 | 0.67274 | 0.95217 | -0.07071 | 0.71964 | 0.85248 | 0.95377 | -0.06828 | 0.093017 | 0.15527 | 0.71802 | -0.47791 |
| TAG521-FA201 | 0.14301 | 0.25299 | 0.60889 | -0.71576 | 0.7022 | 0.84186 | 1.1953 | 0.25743 | 0.70237 | 0.77078 | 0.88027 | -0.18399 |
| TAG543-FA203 | 0.26385 | 0.39215 | 0.7937 | -0.33334 | 0.87031 | 0.92865 | 1.0113 | 0.016225 | 0.061664 | 0.11372 | 0.64227 | -0.63876 |
| PE181/161 | 0.34748 | 0.48027 | 0.79861 | -0.32444 | 0.31714 | 0.53074 | 0.78358 | -0.35185 | 0.76061 | 0.81845 | 0.96699 | -0.04842 |
| TAG501-FA181 | 0.11097 | 0.2064 | 0.59103 | -0.7587 | 0.96021 | 0.98327 | 0.99426 | -0.0083 | 0.45884 | 0.55521 | 0.8248 | -0.27789 |
| TAG566-FA204 | 0.57929 | 0.69196 | 0.92355 | -0.11474 | 0.63014 | 0.79524 | 1.1544 | 0.2071 | 0.86286 | 0.90019 | 1.0374 | 0.052988 |
| LPC225 | 0.94439 | 0.96971 | 0.96266 | -0.0549 | 0.68952 | 0.83435 | 1.1018 | 0.13992 | 0.25516 | 0.34109 | 1.2259 | 0.2938 |
| PE160/183 | 0.42318 | 0.5588 | 0.89426 | -0.16124 | 0.082912 | 0.23067 | 0.66511 | -0.58835 | 0.00012 | 0.004481 | 0.34301 | -1.5437 |
| HCER160 | 0.49915 | 0.6359 | 1.0774 | 0.10756 | 0.3817 | 0.59778 | 0.86531 | -0.2087 | 0.039439 | 0.081879 | 0.71193 | -0.49019 |
| TAG547-FA225 | 0.18893 | 0.309 | 0.69251 | -0.53009 | 0.44425 | 0.65696 | 0.81997 | -0.28636 | 0.000488 | 0.005989 | 0.2881 | -1.7954 |
| TAG564-FA160 | 0.52326 | 0.65133 | 0.82839 | -0.27162 | 0.50202 | 0.7097 | 1.273 | 0.34823 | 0.34129 | 0.43226 | 0.78438 | -0.35037 |
| PG160/202_2 | 0.008418 | 0.041003 | 0.62429 | -0.67972 | 0.011982 | 0.081295 | 0.57289 | -0.80367 | 0.006914 | 0.025783 | 0.52367 | -0.93326 |
| TAG540-FA180 | 0.028664 | 0.081487 | 0.48738 | -1.0369 | 0.093379 | 0.25003 | 0.57932 | -0.78756 | 0.001888 | 0.013073 | 0.28252 | -1.8236 |
| PE180/161 | 0.36937 | 0.50507 | 0.83906 | -0.25316 | 0.58703 | 0.76127 | 0.95425 | -0.06756 | 0.01359 | 0.04115 | 0.5074 | -0.97879 |
| PE180/226 | 0.96391 | 0.97895 | 1.0119 | 0.017017 | 0.24765 | 0.46195 | 0.82619 | -0.27546 | 0.58552 | 0.66756 | 0.9428 | -0.08498 |
| TAG563-FA160 | 0.93204 | 0.95995 | 0.98403 | -0.02322 | 0.29791 | 0.51122 | 1.4529 | 0.53897 | 0.1214 | 0.18908 | 0.7227 | -0.46853 |
| SMd181/226 | 0.005395 | 0.033771 | 0.71725 | -0.47944 | 0.074143 | 0.21742 | 0.78997 | -0.34013 | 0.23725 | 0.32244 | 0.8637 | -0.2114 |

| | | | | | | | | | | | | |
|---|---|---|---|---|---|---|---|---|---|---|---|---|
| PEO-180/205 | 0.40777 | 0.54411 | 0.75499 | -0.40547 | 0.23661 | 0.44796 | 0.68534 | -0.54511 | 0.054412 | 0.10305 | 0.52067 | -0.94155 |
| HCER220 | 0.028197 | 0.080863 | 1.2922 | 0.36987 | 0.93792 | 0.97694 | 0.99547 | -0.00654 | 0.56741 | 0.65262 | 0.93784 | -0.09259 |
| TAG545-FA204 | 0.72181 | 0.81297 | 0.95365 | -0.06847 | 0.66382 | 0.82157 | 1.1872 | 0.2476 | 0.95226 | 0.97228 | 1.1598 | 0.21387 |
| SMd181/220 | 0.10402 | 0.19686 | 0.84696 | -0.23963 | 0.090516 | 0.24741 | 0.83295 | -0.26371 | 0.96487 | 0.97749 | 0.99281 | -0.01041 |
| TAG567-FA180 | 0.28199 | 0.40984 | 0.84173 | -0.24857 | 0.21737 | 0.42666 | 0.8315 | -0.26621 | 0.086168 | 0.14685 | 0.63365 | -0.65824 |
| PC160/182 | 0.69019 | 0.78689 | 0.97406 | -0.03791 | 0.18982 | 0.39565 | 0.89656 | -0.15753 | 0.69329 | 0.76497 | 1.0468 | 0.06601 |
| TAG567-FA225 | 0.064296 | 0.14433 | 0.69999 | -0.51459 | 0.32979 | 0.54192 | 0.85858 | -0.21998 | 0.11705 | 0.184 | 0.71914 | -0.47566 |
| TAG545-FA160 | 0.096311 | 0.19037 | 0.73621 | -0.44181 | 0.80533 | 0.90391 | 0.97551 | -0.03578 | 0.13081 | 0.1999 | 0.71804 | -0.47785 |
| PEP-181/225 | 0.096767 | 0.19037 | 1.1391 | 0.18788 | 0.9698 | 0.98629 | 0.99348 | -0.00944 | 0.004212 | 0.02052 | 1.3185 | 0.39893 |
| PEP-182/182 | 0.14899 | 0.26004 | 0.75398 | -0.4074 | 0.14955 | 0.33571 | 0.75772 | -0.40027 | 0.40352 | 0.49941 | 0.85769 | -0.22148 |
| TAG546-FA226 | 0.050002 | 0.12283 | 1.5934 | 0.67215 | 0.065102 | 0.19712 | 2.072 | 1.0511 | 0.82217 | 0.87087 | 1.4959 | 0.58106 |
| PE160/226 | 0.47467 | 0.60948 | 0.87034 | -0.20035 | 0.1242 | 0.29509 | 0.71829 | -0.47737 | 0.33236 | 0.42341 | 0.83171 | -0.26584 |
| FFA203 | 0.016502 | 0.057447 | 0.76672 | -0.38323 | 0.003963 | 0.059992 | 0.61328 | -0.70537 | 0.44367 | 0.53886 | 0.9199 | -0.12045 |
| LPE202 | 0.86432 | 0.92066 | 1.029 | 0.041237 | 0.42722 | 0.6438 | 0.84398 | -0.24471 | 0.039779 | 0.081949 | 0.63316 | -0.65936 |
| SMd181/205 | 0.087916 | 0.18055 | 0.85415 | -0.22744 | 0.098718 | 0.25629 | 0.8485 | -0.23701 | 0.9431 | 0.96839 | 1.0031 | 0.004483 |
| TAG566-FA160 | 0.058377 | 0.13554 | 0.62078 | -0.68785 | 0.50593 | 0.71153 | 0.85143 | -0.23204 | 0.14634 | 0.22001 | 0.67187 | -0.57375 |
| PG160/202 | 0.005835 | 0.033965 | 0.63927 | -0.6455 | 0.013979 | 0.085768 | 0.57825 | -0.79023 | 0.004585 | 0.021321 | 0.53543 | -0.90123 |
| TAG547-FA226 | 0.55014 | 0.67226 | 0.81568 | -0.29393 | 0.49264 | 0.7038 | 1.2997 | 0.37817 | 0.042331 | 0.086186 | 0.43611 | -1.1972 |
| SM241 | 0.93341 | 0.95995 | 0.99103 | -0.013 | 0.61292 | 0.78237 | 0.96415 | -0.05268 | 0.10421 | 0.16918 | 1.1132 | 0.15466 |
| PEP-181/205 | 0.18898 | 0.309 | 0.74705 | -0.42072 | 0.089518 | 0.24589 | 0.67509 | -0.56685 | 0.12395 | 0.19168 | 0.66954 | -0.57876 |
| HCER241 | 0.086789 | 0.17993 | 1.2332 | 0.30239 | 0.80943 | 0.90695 | 0.97639 | -0.03447 | 0.069247 | 0.12351 | 0.83529 | -0.25966 |
| FFA226 | 0.43805 | 0.56929 | 0.92622 | -0.11057 | 0.11657 | 0.28421 | 0.76482 | -0.38682 | 0.43861 | 0.53371 | 1.147 | 0.1979 |
| FFA220 | 0.812 | 0.87664 | 0.93317 | -0.09978 | 0.35829 | 0.57451 | 0.79038 | -0.33938 | 0.37535 | 0.46811 | 0.8022 | -0.31797 |
| PEP-180/160 | 0.85282 | 0.91314 | 0.97241 | -0.04037 | 0.16864 | 0.3677 | 0.77466 | -0.36837 | 0.70927 | 0.77343 | 1.0841 | 0.11643 |
| PE160/201 | 0.3161 | 0.44638 | 0.75191 | -0.41137 | 0.13843 | 0.3162 | 0.6461 | -0.63018 | 0.067505 | 0.12073 | 0.59326 | -0.75327 |
| TAG565-FA180 | 0.095058 | 0.19037 | 0.77404 | -0.36951 | 0.8878 | 0.93672 | 1.0413 | 0.058389 | 0.46921 | 0.56253 | 0.9434 | -0.08406 |
| PG161/182_2 | 0.42748 | 0.56077 | 1.2052 | 0.26932 | 0.98938 | 0.99442 | 1.025 | 0.035665 | 0.45791 | 0.55511 | 0.90016 | -0.15175 |
| TAG566-FA181 | 0.098354 | 0.19142 | 0.68985 | -0.53564 | 0.18864 | 0.39487 | 0.7587 | -0.39839 | 0.097657 | 0.16108 | 0.6637 | -0.59139 |
| PC140/225 | 0.21495 | 0.33558 | 0.86747 | -0.20512 | 0.14135 | 0.32175 | 0.85311 | -0.2292 | 0.23526 | 0.32041 | 0.85565 | -0.2249 |
| PC180/181 | 0.065785 | 0.14616 | 0.79768 | -0.32611 | 0.16073 | 0.35591 | 0.86568 | -0.20809 | 0.72546 | 0.78581 | 0.97063 | -0.04301 |
| TAG501-FA160 | 0.23933 | 0.36233 | 0.67481 | -0.56745 | 0.7515 | 0.87206 | 1.096 | 0.13228 | 0.55467 | 0.64288 | 0.84805 | -0.23779 |
| PC160/224 | 0.42812 | 0.56077 | 1.1147 | 0.15661 | 0.6481 | 0.81137 | 1.0717 | 0.099966 | 0.42895 | 0.52475 | 1.1184 | 0.16143 |
| TAG481-FA161 | 0.36531 | 0.50113 | 0.62708 | -0.67327 | 0.51595 | 0.71313 | 1.6623 | 0.73314 | 0.15004 | 0.22352 | 0.5017 | -0.9951 |

| | | | | | | | | | | | | |
|---|---|---|---|---|---|---|---|---|---|---|---|---|
| PEP-181/183 | 0.32769 | 0.45976 | 0.80691 | -0.30952 | 0.17354 | 0.37173 | 0.73541 | -0.44339 | 0.20208 | 0.28413 | 0.74218 | -0.43016 |
| PEO-160/203 | 0.01272 | 0.049 | 0.77403 | -0.36954 | 0.014888 | 0.085768 | 0.73618 | -0.44186 | 0.0217 | 0.054448 | 0.70823 | -0.4977 |
| PC181/201 | 0.013079 | 0.049727 | 0.66305 | -0.59281 | 0.012113 | 0.081295 | 0.65602 | -0.6082 | 0.85627 | 0.89619 | 0.98052 | -0.02838 |
| TAG548-FA204 | 0.12652 | 0.23032 | 0.72383 | -0.46627 | 0.099211 | 0.25629 | 0.67161 | -0.5743 | 0.005713 | 0.023333 | 0.48041 | -1.0577 |
| PEP-181/203 | 0.024265 | 0.073807 | 0.83049 | -0.26796 | 0.030185 | 0.11838 | 0.80104 | -0.32005 | 0.56646 | 0.65262 | 0.96134 | -0.05688 |
| PE180/180 | 0.47043 | 0.60643 | 1.2007 | 0.2639 | 0.90201 | 0.94864 | 1.0369 | 0.052255 | 0.14784 | 0.22125 | 1.5453 | 0.62793 |
| TAG565-FA160 | 0.38094 | 0.51881 | 0.78681 | -0.34591 | 0.70452 | 0.8431 | 1.0553 | 0.077601 | 0.31252 | 0.40207 | 0.76098 | -0.39408 |
| SMd181/222 | 0.2509 | 0.37721 | 0.94562 | -0.08066 | 0.68682 | 0.83418 | 0.98041 | -0.02855 | 0.96842 | 0.97894 | 1.0076 | 0.010908 |
| PC181/204 | 0.017878 | 0.060302 | 0.76379 | -0.38874 | 0.001238 | 0.035765 | 0.62633 | -0.675 | 0.72462 | 0.78581 | 1.0604 | 0.084646 |
| TAG481-FA180 | 0.93262 | 0.95995 | 1.0074 | 0.010708 | 0.353 | 0.56883 | 1.943 | 0.95827 | 0.10359 | 0.16859 | 0.47506 | -1.0738 |
| PC181/181 | 0.13061 | 0.23553 | 0.75484 | -0.40576 | 0.024921 | 0.1066 | 0.67291 | -0.57151 | 0.70847 | 0.77343 | 0.94379 | -0.08346 |
| HCER140 | 0.42627 | 0.56061 | 0.88702 | -0.17295 | 0.034945 | 0.12853 | 0.73417 | -0.44582 | 0.018433 | 0.049393 | 0.64466 | -0.63338 |
| TAG542-FA180 | 0.81852 | 0.88221 | 0.9694 | -0.04484 | 0.44503 | 0.65696 | 1.1654 | 0.22077 | 0.097738 | 0.16108 | 0.61478 | -0.70185 |
| TAG545-FA225 | 0.99516 | 0.99758 | 1.0522 | 0.073477 | 0.21825 | 0.42666 | 1.4307 | 0.51671 | 0.57532 | 0.65961 | 0.98578 | -0.02066 |
| PE170/225 | 0.003315 | 0.028391 | 0.5816 | -0.78189 | 0.000783 | 0.029996 | 0.48054 | -1.0573 | 5.79E-05 | 0.004338 | 0.42739 | -1.2264 |
| PEO-180/203 | 0.001625 | 0.022041 | 0.74629 | -0.42219 | 0.000153 | 0.019518 | 0.63402 | -0.6574 | 0.000334 | 0.00519 | 0.60378 | -0.72791 |
| PA180/181 | 0.79281 | 0.86452 | 1.2241 | 0.29172 | 0.82317 | 0.90904 | 0.95656 | -0.06408 | 0.61433 | 0.69674 | 1.2993 | 0.37772 |
| PEP-160/202 | 0.73697 | 0.82862 | 1.0051 | 0.007268 | 0.72564 | 0.85435 | 0.98195 | -0.02628 | 0.69164 | 0.76445 | 0.98415 | -0.02306 |
| SMd181/224 | 0.23309 | 0.35611 | 0.88296 | -0.17958 | 0.21932 | 0.42748 | 0.87464 | -0.19324 | 0.68613 | 0.75965 | 0.95674 | -0.06381 |
| FFA183 | 0.000267 | 0.018752 | 0.52299 | -0.93513 | 0.01649 | 0.090212 | 0.49407 | -1.0172 | 0.067418 | 0.12073 | 0.81406 | -0.29679 |
| TAG569-FA226 | 0.15853 | 0.2723 | 0.74943 | -0.41613 | 0.24031 | 0.45214 | 0.81599 | -0.29338 | 0.008116 | 0.028406 | 0.51518 | -0.95686 |
| SMd181/184 | 0.009973 | 0.044164 | 0.70703 | -0.50015 | 0.002751 | 0.047127 | 0.66086 | -0.59758 | 0.000211 | 0.00465 | 0.64602 | -0.63035 |
| PE180/183 | 0.43812 | 0.56929 | 1.1857 | 0.24568 | 0.10201 | 0.26042 | 0.7534 | -0.40851 | 0.00013 | 0.004481 | 0.39006 | -1.3582 |
| HCER240 | 0.30372 | 0.43265 | 1.1327 | 0.1798 | 0.98023 | 0.99242 | 0.99173 | -0.01198 | 0.060558 | 0.11232 | 0.7889 | -0.34209 |
| PEP-180/161 | 0.51009 | 0.6448 | 0.9076 | -0.13987 | 0.39133 | 0.60911 | 0.82788 | -0.2725 | 0.26423 | 0.35034 | 0.75092 | -0.41326 |
| PEP-180/201 | 0.63179 | 0.73577 | 0.89036 | -0.16754 | 0.39204 | 0.60911 | 0.7238 | -0.46634 | 0.9764 | 0.9846 | 0.95267 | -0.06996 |
| PEP-160/226 | 0.6821 | 0.77903 | 1.0786 | 0.10921 | 0.85573 | 0.92847 | 1.0299 | 0.04252 | 0.12425 | 0.19168 | 1.5242 | 0.60802 |
| SMd181/181 | 0.98139 | 0.99052 | 0.99717 | -0.00409 | 0.68938 | 0.83435 | 1.0578 | 0.081016 | 0.034418 | 0.074175 | 1.3485 | 0.43134 |
| LPC205 | 0.76883 | 0.85165 | 0.9228 | -0.11591 | 0.7756 | 0.88426 | 0.91547 | -0.12741 | 0.82572 | 0.87122 | 1.0104 | 0.014923 |
| PEP-160/204 | 0.92377 | 0.95644 | 1.0003 | 0.000409 | 0.9175 | 0.96028 | 0.97448 | -0.0373 | 0.3256 | 0.41643 | 1.2636 | 0.33758 |
| PE170/202 | 0.009587 | 0.043341 | 0.62009 | -0.68946 | 0.009104 | 0.075987 | 0.62876 | -0.66941 | 0.1013 | 0.16611 | 0.76529 | -0.38593 |
| HCER260 | 0.21495 | 0.33558 | 0.80471 | -0.31345 | 0.40799 | 0.62642 | 0.86781 | -0.20455 | 0.041861 | 0.085697 | 0.66528 | -0.58797 |
| PE160/160 | 0.78495 | 0.86173 | 0.94742 | -0.07793 | 0.32263 | 0.53854 | 0.85186 | -0.23131 | 0.53151 | 0.62588 | 0.91005 | -0.13598 |

| | | | | | | | | | | | | |
|---|---|---|---|---|---|---|---|---|---|---|---|---|
| TAG566-FA205 | 0.58301 | 0.69385 | 1.2935 | 0.37128 | 0.78577 | 0.89118 | 0.99596 | -0.00584 | 0.12647 | 0.19464 | 0.62284 | -0.68306 |
| TAG567-FA160 | 0.075495 | 0.16114 | 0.6828 | -0.55046 | 0.52231 | 0.71929 | 0.88327 | -0.17907 | 0.11542 | 0.18344 | 0.67454 | -0.56801 |
| PEP-180/225 | 0.78966 | 0.86253 | 1.0159 | 0.022694 | 0.4813 | 0.69628 | 0.89238 | -0.16427 | 0.14719 | 0.22079 | 1.181 | 0.24006 |
| PE180/204 | 0.44556 | 0.57781 | 0.9351 | -0.0968 | 0.10993 | 0.27314 | 0.81103 | -0.30218 | 0.4143 | 0.50984 | 0.91675 | -0.1254 |
| PC160/181 | 0.77602 | 0.85336 | 0.98139 | -0.0271 | 0.41769 | 0.63532 | 1.0998 | 0.1373 | 0.19446 | 0.27521 | 1.1528 | 0.2051 |
| LPE200 | 0.33474 | 0.46738 | 0.79486 | -0.33123 | 0.099003 | 0.25629 | 0.66728 | -0.58363 | 0.005288 | 0.022354 | 0.52294 | -0.93527 |
| PC180/226 | 0.18777 | 0.30868 | 0.87332 | -0.19542 | 0.54927 | 0.73575 | 0.93664 | -0.09443 | 0.17058 | 0.24843 | 1.2178 | 0.28428 |
| SMd181/180 | 0.53462 | 0.66167 | 0.94566 | -0.0806 | 0.29482 | 0.50846 | 0.90625 | -0.14203 | 0.57787 | 0.66115 | 1.0432 | 0.061073 |
| PS160/180_2 | 0.90818 | 0.94584 | 0.96984 | -0.04418 | 0.51426 | 0.71313 | 0.90424 | -0.14523 | 0.23301 | 0.31801 | 0.81239 | -0.29975 |
| HCER181 | 0.94972 | 0.97212 | 0.97495 | -0.03661 | 0.017717 | 0.090528 | 0.72065 | -0.47263 | 0.43457 | 0.52978 | 0.89428 | -0.16121 |
| TAG566-FA225 | 0.23358 | 0.35611 | 0.78752 | -0.34462 | 0.74988 | 0.87173 | 1.0675 | 0.094212 | 0.32116 | 0.41156 | 0.82572 | -0.27627 |
| PC150/225 | 0.032709 | 0.088792 | 0.70434 | -0.50565 | 0.012653 | 0.083205 | 0.61447 | -0.70258 | 0.20017 | 0.28206 | 0.80524 | -0.3125 |
| TAG546-FA160 | 0.95702 | 0.97652 | 1.0421 | 0.059507 | 0.52273 | 0.71929 | 1.2367 | 0.30655 | 0.37495 | 0.46811 | 0.90115 | -0.15016 |
| TAG567-FA204 | 0.70611 | 0.79944 | 1.0689 | 0.096117 | 0.95714 | 0.98327 | 1.0725 | 0.10093 | 0.22721 | 0.31338 | 0.81183 | -0.30075 |
| LPC182 | 0.10661 | 0.20059 | 0.77512 | -0.36751 | 0.28558 | 0.49842 | 0.84954 | -0.23525 | 0.76736 | 0.82298 | 1.0239 | 0.034085 |
| PC181/224 | 0.39813 | 0.5358 | 0.88397 | -0.17793 | 0.057558 | 0.17843 | 0.7231 | -0.46774 | 0.76263 | 0.81926 | 0.96878 | -0.04576 |
| PEO-180/226 | 0.9863 | 0.99087 | 0.98437 | -0.02273 | 0.41316 | 0.63286 | 0.8704 | -0.20024 | 0.11603 | 0.18379 | 1.28 | 0.3562 |
| SMd181/221 | 0.57625 | 0.69196 | 0.93134 | -0.10261 | 0.78999 | 0.89286 | 0.96654 | -0.04909 | 0.028536 | 0.065411 | 1.2508 | 0.32288 |
| SM240 | 0.10028 | 0.19314 | 0.85714 | -0.22239 | 0.12769 | 0.29795 | 0.87653 | -0.19013 | 0.10102 | 0.16607 | 0.88311 | -0.17933 |
| DAG161/182 | 0.090883 | 0.18357 | 0.44938 | -1.154 | 0.21775 | 0.42666 | 0.42023 | -1.2507 | 0.17402 | 0.25175 | 0.37794 | -1.4038 |
| PEO-180/181 | 0.034981 | 0.092948 | 0.67193 | -0.57361 | 0.018782 | 0.092363 | 0.62922 | -0.66836 | 0.001656 | 0.012198 | 0.52807 | -0.9212 |
| PC170/224 | 0.66739 | 0.76855 | 0.96294 | -0.05448 | 0.10156 | 0.2603 | 0.84289 | -0.24658 | 0.10983 | 0.17654 | 0.84662 | -0.24021 |
| TAG520-FA180 | 0.14366 | 0.25345 | 0.57694 | -0.79352 | 0.51563 | 0.71313 | 0.7641 | -0.38816 | 0.0366 | 0.077302 | 0.43733 | -1.1932 |
| TAG565-FA204 | 0.55796 | 0.67641 | 0.90818 | -0.13895 | 0.31132 | 0.52234 | 1.2746 | 0.35003 | 0.92797 | 0.95587 | 1.1139 | 0.15556 |
| PE160/202 | 0.53989 | 0.66566 | 0.88881 | -0.17006 | 0.079271 | 0.22535 | 0.66701 | -0.58421 | 0.042365 | 0.086186 | 0.61472 | -0.70199 |
| PC140/204 | 0.007703 | 0.039366 | 0.77416 | -0.3693 | 0.086019 | 0.23728 | 0.8524 | -0.2304 | 0.016696 | 0.046448 | 0.80322 | -0.31613 |
| PEP-181/181 | 0.39835 | 0.5358 | 0.87101 | -0.19924 | 0.20965 | 0.41485 | 0.79286 | -0.33485 | 0.94583 | 0.96848 | 1.0248 | 0.035392 |
| TAG568-FA160 | 0.36565 | 0.50113 | 0.85666 | -0.22321 | 0.81649 | 0.90706 | 0.95332 | -0.06897 | 0.18558 | 0.26669 | 0.74462 | -0.42542 |
| FFA120 | 0.10974 | 0.2053 | 0.65581 | -0.60864 | 0.037646 | 0.13319 | 0.57551 | -0.79709 | 0.58138 | 0.664 | 1.0687 | 0.095805 |
| DAG140/160 | 0.94909 | 0.97212 | 0.92293 | -0.11571 | 0.5554 | 0.73639 | 1.5466 | 0.62914 | 0.27825 | 0.36472 | 1.7991 | 0.84725 |
| TAG567-FA181 | 0.4876 | 0.62363 | 0.87796 | -0.18778 | 0.72022 | 0.85248 | 1.0545 | 0.076591 | 0.19409 | 0.27521 | 0.74945 | -0.4161 |
| CER160 | 0.86723 | 0.92099 | 1.0483 | 0.068028 | 0.30501 | 0.51709 | 0.86191 | -0.21439 | 0.83438 | 0.87893 | 0.98916 | -0.01572 |
| TAG482-FA180 | 0.78822 | 0.86253 | 1.4177 | 0.50358 | 0.87099 | 0.92865 | 1.3242 | 0.40509 | 0.066911 | 0.12033 | 0.56933 | -0.81267 |

| | | | | | | | | | | | | |
|---|---|---|---|---|---|---|---|---|---|---|---|---|
| LPC221 | 0.008016 | 0.040453 | 0.62131 | -0.68663 | 0.20365 | 0.40863 | 0.79965 | -0.32256 | 0.6779 | 0.7531 | 0.92745 | -0.10865 |
| PEP-181/161 | 0.90596 | 0.94516 | 1.2758 | 0.35136 | 0.73544 | 0.8611 | 1.0775 | 0.10765 | 0.93161 | 0.9581 | 1.1489 | 0.20031 |
| PC160/180 | 0.31348 | 0.44461 | 0.91463 | -0.12874 | 0.69147 | 0.83515 | 0.97007 | -0.04383 | 0.29513 | 0.38273 | 1.0964 | 0.1328 |
| PC170/225 | 0.019004 | 0.062801 | 0.67121 | -0.57517 | 0.010075 | 0.0783 | 0.67612 | -0.56464 | 0.003331 | 0.017847 | 0.60656 | -0.72127 |
| FFA225 | 0.15389 | 0.26573 | 0.81888 | -0.28827 | 0.10505 | 0.26714 | 0.71886 | -0.47622 | 0.279 | 0.36472 | 1.2248 | 0.29252 |
| FFA202 | 0.020545 | 0.067548 | 0.72432 | -0.46531 | 0.010126 | 0.0783 | 0.57929 | -0.78765 | 0.61157 | 0.69482 | 1.0561 | 0.078707 |
| PEP-181/160 | 0.42301 | 0.5588 | 1.2643 | 0.33838 | 0.72346 | 0.85435 | 0.95044 | -0.07333 | 0.31079 | 0.40064 | 1.3331 | 0.41474 |
| PEP-160/224 | 0.82639 | 0.88922 | 1.0551 | 0.077388 | 0.83036 | 0.9145 | 0.92733 | -0.10885 | 0.70923 | 0.77343 | 1.0816 | 0.1132 |
| PEO-160/226 | 0.90952 | 0.94584 | 0.99328 | -0.00972 | 0.55923 | 0.73996 | 0.9252 | -0.11216 | 0.10757 | 0.17377 | 1.2391 | 0.30929 |
| PC160/161 | 0.33964 | 0.47144 | 0.82796 | -0.27237 | 0.95776 | 0.98327 | 1.0331 | 0.047044 | 0.70329 | 0.77078 | 0.94025 | -0.08888 |
| SMd181/141 | 0.024602 | 0.073807 | 0.65925 | -0.60111 | 0.074524 | 0.21756 | 0.79972 | -0.32243 | 0.64175 | 0.72039 | 0.95181 | -0.07126 |
| SMd181/170 | 0.062003 | 0.14064 | 0.8473 | -0.23905 | 0.1313 | 0.30342 | 0.86797 | -0.20429 | 0.82271 | 0.87087 | 1.0099 | 0.014253 |
| PEO-180/204 | 0.033042 | 0.088885 | 0.85337 | -0.22876 | 0.043492 | 0.14902 | 0.83044 | -0.26805 | 0.063346 | 0.11616 | 0.8158 | -0.2937 |
| FFA140 | 7.43E-05 | 0.018752 | 0.65819 | -0.60342 | 0.028118 | 0.11734 | 0.70822 | -0.49774 | 0.39355 | 0.48894 | 1.08 | 0.111 |
| SMd181/140 | 0.005394 | 0.033771 | 0.71705 | -0.47985 | 0.006604 | 0.071506 | 0.75955 | -0.39678 | 0.014009 | 0.04115 | 0.81313 | -0.29845 |
| CER180 | 0.24965 | 0.37621 | 1.3968 | 0.48213 | 0.9227 | 0.96263 | 1.1526 | 0.20489 | 0.12408 | 0.19168 | 1.212 | 0.2774 |
| TAG522-FA202 | 0.081878 | 0.17194 | 0.61443 | -0.70267 | 0.97114 | 0.98629 | 1.0307 | 0.043662 | 0.1398 | 0.21165 | 0.65658 | -0.60696 |
| PE160/140 | 0.10275 | 0.19502 | 0.5886 | -0.76465 | 0.17777 | 0.37685 | 0.66981 | -0.57817 | 0.040051 | 0.08225 | 0.57951 | -0.78709 |
| PEP-160/181 | 0.62478 | 0.73022 | 0.96239 | -0.05531 | 0.61169 | 0.78233 | 0.93949 | -0.09006 | 0.81436 | 0.86491 | 1.181 | 0.24003 |
| TAG504-FA141 | 0.76033 | 0.84467 | 0.91932 | -0.12136 | 0.87655 | 0.9324 | 0.91879 | -0.1222 | 0.013739 | 0.04115 | 0.37322 | -1.4219 |
| FFA204 | 0.1338 | 0.23996 | 0.83507 | -0.26003 | 0.033379 | 0.1256 | 0.68027 | -0.55583 | 0.46418 | 0.55856 | 0.9222 | -0.11684 |
| CE182 | 0.014627 | 0.053151 | 0.63397 | -0.65752 | 0.0003 | 0.019518 | 0.43724 | -1.1935 | 0.23273 | 0.31801 | 0.80376 | -0.31516 |
| PEP-182/226 | 0.47202 | 0.60728 | 1.1106 | 0.1514 | 0.78494 | 0.89118 | 0.9614 | -0.05679 | 0.14489 | 0.21834 | 1.3907 | 0.47584 |
| CER221 | 0.13812 | 0.24567 | 0.74662 | -0.42156 | 0.27097 | 0.48143 | 0.82091 | -0.28471 | 0.65475 | 0.73364 | 0.95462 | -0.067 |
| TAG500-FA160 | 0.098503 | 0.19142 | 0.60038 | -0.73606 | 0.32811 | 0.54077 | 0.79022 | -0.33967 | 0.22462 | 0.31046 | 0.70352 | -0.50734 |
| TAG523-FA203 | 0.20777 | 0.3283 | 0.719 | -0.47594 | 0.90856 | 0.95246 | 0.98668 | -0.01935 | 0.20372 | 0.28583 | 0.71189 | -0.49027 |
| PA180/203 | 0.18096 | 0.30052 | 1.3097 | 0.38924 | 0.20153 | 0.40863 | 1.2809 | 0.35712 | 0.55863 | 0.64595 | 1.1462 | 0.19687 |
| CER260 | 0.054657 | 0.13098 | 0.64573 | -0.631 | 0.064562 | 0.19712 | 0.66452 | -0.58962 | 0.16906 | 0.24677 | 0.7301 | -0.45384 |
| TAG526-FA226 | 0.56344 | 0.67925 | 0.8776 | -0.18837 | 0.3072 | 0.51945 | 1.8634 | 0.89791 | 0.039493 | 0.081879 | 0.54179 | -0.88421 |
| TAG566-FA180 | 0.38896 | 0.52705 | 0.89582 | -0.15872 | 0.61561 | 0.78427 | 1.1084 | 0.14852 | 0.52538 | 0.62073 | 0.96097 | -0.05743 |
| LPE203 | 0.66617 | 0.76855 | 0.91209 | -0.13275 | 0.45465 | 0.66512 | 0.8575 | -0.22179 | 0.42033 | 0.51541 | 0.85106 | -0.23267 |
| PE160/182 | 0.20708 | 0.32808 | 0.82768 | -0.27285 | 0.090923 | 0.24741 | 0.70718 | -0.49986 | 0.006632 | 0.025472 | 0.5763 | -0.7951 |
| TAG500-FA180 | 0.079497 | 0.16748 | 0.49822 | -1.0051 | 0.31074 | 0.52234 | 0.69702 | -0.52073 | 0.16051 | 0.23588 | 0.58505 | -0.77336 |

| | | | | | | | | | | | | |
|---|---|---|---|---|---|---|---|---|---|---|---|---|
| PE160/205 | 0.6682 | 0.76855 | 0.88168 | -0.18167 | 0.18048 | 0.38023 | 0.66219 | -0.59468 | 0.12805 | 0.1966 | 0.61793 | -0.69448 |
| PEP-181/224 | 0.98503 | 0.99087 | 0.99339 | -0.00957 | 0.49399 | 0.7038 | 0.83279 | -0.26398 | 0.76911 | 0.8235 | 1.0642 | 0.089809 |
| SM261 | 0.035701 | 0.094094 | 0.79941 | -0.323 | 0.32438 | 0.5387 | 0.89317 | -0.16299 | 0.55499 | 0.64288 | 0.94551 | -0.08084 |
| PC160/140 | 0.12617 | 0.23032 | 0.77974 | -0.35893 | 0.41713 | 0.63532 | 0.89189 | -0.16507 | 0.2313 | 0.317 | 0.83311 | -0.26341 |
| PC170/205 | 0.54312 | 0.66837 | 0.89318 | -0.16297 | 0.10612 | 0.26811 | 0.62452 | -0.67918 | 0.072731 | 0.12762 | 0.57105 | -0.80832 |
| HCER221 | 0.55156 | 0.67226 | 0.9448 | -0.08191 | 0.013422 | 0.085427 | 0.76994 | -0.37718 | 0.16598 | 0.24336 | 0.86911 | -0.20238 |
| SMd181/201 | 0.80483 | 0.87324 | 0.94157 | -0.08686 | 0.55044 | 0.73581 | 1.0925 | 0.12769 | 0.033648 | 0.074002 | 1.3759 | 0.4604 |
| LPE240 | 0.006032 | 0.034722 | 0.68225 | -0.55162 | 0.42077 | 0.63702 | 0.89184 | -0.16514 | 0.65832 | 0.73637 | 0.94046 | -0.08856 |
| HCER261 | 0.29014 | 0.41974 | 0.81894 | -0.28817 | 0.20616 | 0.40917 | 0.78156 | -0.35557 | 0.006761 | 0.025565 | 0.58532 | -0.77271 |
| PC180/225 | 0.46846 | 0.60509 | 0.9366 | -0.0945 | 0.67632 | 0.82838 | 1.0518 | 0.072918 | 0.6754 | 0.7516 | 1.0555 | 0.077968 |
| SMd181/223 | 0.16752 | 0.284 | 0.90631 | -0.14192 | 0.029924 | 0.11815 | 0.82517 | -0.27724 | 0.054452 | 0.10305 | 0.83945 | -0.25249 |
| SMd181/225 | 0.098796 | 0.19142 | 0.7593 | -0.39726 | 0.58643 | 0.76127 | 0.92015 | -0.12005 | 0.29132 | 0.37855 | 0.84715 | -0.23931 |
| PE160/204 | 0.30761 | 0.43723 | 0.85748 | -0.22183 | 0.33977 | 0.55159 | 0.85252 | -0.2302 | 0.31956 | 0.41032 | 0.86321 | -0.21221 |
| TAG545-FA205 | 0.86246 | 0.92066 | 0.9853 | -0.02137 | 0.38426 | 0.59988 | 0.77489 | -0.36794 | 0.15588 | 0.22959 | 0.64936 | -0.62292 |
| PEO-160/204 | 0.52873 | 0.65562 | 0.92024 | -0.11992 | 0.79396 | 0.89579 | 0.97127 | -0.04206 | 0.52035 | 0.61591 | 0.92081 | -0.11903 |
| PEO-160/183 | 0.3148 | 0.4455 | 0.86324 | -0.21216 | 0.17131 | 0.3705 | 0.79701 | -0.32733 | 0.003597 | 0.01873 | 0.5037 | -0.98936 |
| DAG160/181 | 0.4104 | 0.54636 | 0.81999 | -0.28633 | 0.96212 | 0.98327 | 0.98614 | -0.02014 | 0.063722 | 0.1162 | 0.58492 | -0.7737 |
| TAG566-FA226 | 0.45923 | 0.59435 | 1.2821 | 0.35849 | 0.25806 | 0.47238 | 1.2476 | 0.31911 | 0.4615 | 0.5574 | 0.94061 | -0.08833 |
| PC160/183 | 0.90285 | 0.94516 | 1.0192 | 0.027423 | 0.60186 | 0.77434 | 0.93484 | -0.09722 | 0.070884 | 0.12506 | 0.67767 | -0.56135 |
| PC180/182 | 0.22062 | 0.34114 | 0.8988 | -0.15392 | 0.080929 | 0.22807 | 0.85698 | -0.22267 | 0.94617 | 0.96848 | 1.0081 | 0.011623 |
| FFA161 | 0.015125 | 0.054102 | 0.62695 | -0.67357 | 0.028353 | 0.11757 | 0.61308 | -0.70586 | 0.63708 | 0.71879 | 1.0534 | 0.075114 |
| FFA241 | 0.13798 | 0.24567 | 0.90836 | -0.13867 | 0.020986 | 0.10045 | 0.82074 | -0.285 | 0.72426 | 0.78581 | 0.98447 | -0.02258 |
| TAG483-FA141 | 0.88771 | 0.93662 | 1.2186 | 0.28518 | 0.69361 | 0.83619 | 1.2652 | 0.33938 | 0.03601 | 0.07661 | 0.41576 | -1.2662 |
| PEP-181/226 | 0.77449 | 0.85315 | 1.0499 | 0.070226 | 0.74364 | 0.86758 | 0.94563 | -0.08066 | 0.1901 | 0.2714 | 1.3694 | 0.4535 |
| PEP-160/180 | 0.90405 | 0.94516 | 0.98683 | -0.01913 | 0.71568 | 0.8506 | 0.93027 | -0.10428 | 0.74284 | 0.8033 | 1.2401 | 0.31042 |
| FFA201 | 0.00529 | 0.033762 | 0.69302 | -0.52904 | 0.00602 | 0.068757 | 0.61842 | -0.69333 | 0.065147 | 0.11781 | 1.1667 | 0.22248 |
| FFA141 | 0.066529 | 0.14681 | 0.78785 | -0.34401 | 0.058086 | 0.17921 | 0.72165 | -0.47063 | 0.15396 | 0.22728 | 1.2336 | 0.30288 |
| PC200/226 | 0.35625 | 0.49136 | 0.84832 | -0.23732 | 0.084913 | 0.23523 | 0.71829 | -0.47736 | 0.9226 | 0.95184 | 0.97869 | -0.03108 |
| PEP-140/181 | 0.87039 | 0.92284 | 1.1126 | 0.15389 | 0.09275 | 0.25003 | 0.71566 | -0.48265 | 0.34658 | 0.43792 | 0.84467 | -0.24354 |
| LPE170 | 0.58105 | 0.6928 | 0.8916 | -0.16553 | 0.12641 | 0.29795 | 0.75334 | -0.40863 | 0.03225 | 0.071655 | 0.64765 | -0.62671 |
| FFA240 | 0.027274 | 0.078913 | 0.70321 | -0.50798 | 0.014754 | 0.085768 | 0.62747 | -0.67237 | 0.6224 | 0.70467 | 1.0892 | 0.12327 |
| Cholesterol | 0.013569 | 0.050477 | 0.69904 | -0.51656 | 4.99E-05 | 0.017442 | 0.42388 | -1.2383 | 0.21935 | 0.3052 | 0.85101 | -0.23276 |
| FFA160 | 0.021479 | 0.06888 | 0.87459 | -0.19332 | 0.008357 | 0.073521 | 0.76252 | -0.39115 | 0.95336 | 0.97228 | 1.0041 | 0.005879 |

| | | | | | | | | | | | | |
|---|---|---|---|---|---|---|---|---|---|---|---|---|
| PEP-181/204 | 0.77281 | 0.85315 | 0.96902 | -0.0454 | 0.53534 | 0.724 | 0.9337 | -0.09897 | 0.51639 | 0.61233 | 1.1225 | 0.16677 |
| PE180/224 | 0.8945 | 0.94227 | 1.0244 | 0.034718 | 0.41899 | 0.63582 | 0.86821 | -0.20388 | 0.208 | 0.2912 | 0.79508 | -0.33083 |
| CE183 | 0.022599 | 0.070853 | 0.59486 | -0.74937 | 8.04E-05 | 0.017442 | 0.34517 | -1.5346 | 0.017694 | 0.048195 | 0.61271 | -0.70672 |
| PEP-182/204 | 0.62951 | 0.73442 | 1.0438 | 0.061827 | 0.63033 | 0.79524 | 0.95369 | -0.06841 | 0.25512 | 0.34109 | 1.1713 | 0.22812 |
| CER240 | 0.33895 | 0.47144 | 0.87199 | -0.19762 | 0.20452 | 0.40863 | 0.81792 | -0.28996 | 0.96132 | 0.97632 | 1.004 | 0.005811 |
| PEP-160/183 | 0.51145 | 0.64493 | 0.84677 | -0.23996 | 0.51096 | 0.71313 | 0.87115 | -0.19901 | 0.28092 | 0.36649 | 0.7739 | -0.36978 |
| PA181/182 | 0.020928 | 0.067782 | 1.7147 | 0.77799 | 0.1161 | 0.28415 | 1.3294 | 0.41078 | 0.077309 | 0.13421 | 1.4556 | 0.54161 |
| FFA181 | 0.77452 | 0.85315 | 1.0201 | 0.028656 | 0.11802 | 0.28562 | 0.85034 | -0.23389 | 0.57551 | 0.65961 | 0.96097 | -0.05744 |
| PE180/182 | 0.09108 | 0.18357 | 0.85022 | -0.2341 | 0.029603 | 0.11815 | 0.75555 | -0.40441 | 0.005107 | 0.022185 | 0.70779 | -0.4986 |
| PC160/201 | 0.064985 | 0.14538 | 0.83825 | -0.25455 | 0.17416 | 0.37173 | 0.88289 | -0.17969 | 0.5951 | 0.67729 | 1.0397 | 0.056147 |
| PEP-180/224 | 0.69832 | 0.79337 | 0.92863 | -0.10682 | 0.37903 | 0.59746 | 0.84287 | -0.24661 | 0.69999 | 0.76975 | 1.061 | 0.085413 |
| PS160/161_2 | 0.29959 | 0.42864 | 1.1651 | 0.22044 | 0.24707 | 0.46195 | 1.2037 | 0.26744 | 0.99081 | 0.9954 | 1.0023 | 0.003363 |
| PE180/181 | 0.4847 | 0.62114 | 1.0734 | 0.10214 | 0.86018 | 0.92865 | 1.031 | 0.044027 | 0.047016 | 0.093315 | 0.76502 | -0.38643 |
| FFA170 | 0.004922 | 0.032648 | 0.77673 | -0.36452 | 0.004958 | 0.06511 | 0.6923 | -0.53053 | 0.91268 | 0.9446 | 1.0154 | 0.022009 |
| FFA182 | 0.5787 | 0.69196 | 1.0473 | 0.066648 | 0.16251 | 0.35862 | 0.84859 | -0.23686 | 0.85371 | 0.89495 | 0.98965 | -0.01501 |
| PC150/161 | 0.072611 | 0.15704 | 0.7287 | -0.45659 | 0.0178 | 0.090528 | 0.61952 | -0.69078 | 0.19236 | 0.27342 | 0.79626 | -0.3287 |
| PC180/205 | 0.92412 | 0.95644 | 0.99311 | -0.00997 | 0.69684 | 0.83852 | 0.89791 | -0.15536 | 0.55359 | 0.64288 | 0.85042 | -0.23375 |
| FFA180 | 0.029033 | 0.081821 | 0.8449 | -0.24315 | 0.017505 | 0.090528 | 0.73281 | -0.44849 | 0.99345 | 0.99651 | 1.0152 | 0.021818 |
| TAG524-FA204 | 0.67654 | 0.7754 | 0.87072 | -0.19972 | 0.59365 | 0.76723 | 1.3504 | 0.43342 | 0.74807 | 0.80761 | 1.2587 | 0.33189 |
| LPE182 | 0.71657 | 0.80847 | 0.87068 | -0.19979 | 0.95683 | 0.98327 | 0.93367 | -0.09902 | 0.25691 | 0.34202 | 0.74796 | -0.41897 |
| TAG480-FA160 | 0.19058 | 0.31017 | 0.64972 | -0.62211 | 0.73394 | 0.86089 | 1.0035 | 0.005096 | 0.41112 | 0.50786 | 0.78372 | -0.3516 |
| CER200 | 0.90564 | 0.94516 | 1.056 | 0.078562 | 0.40775 | 0.62642 | 0.89236 | -0.16431 | 0.77547 | 0.82759 | 1.0769 | 0.10687 |
| CER220 | 0.074172 | 0.15936 | 0.72613 | -0.4617 | 0.069674 | 0.20652 | 0.60606 | -0.72248 | 0.72184 | 0.78581 | 0.95928 | -0.05997 |
| PEO-160/160 | 0.043157 | 0.1089 | 0.73897 | -0.43642 | 0.002389 | 0.044126 | 0.58105 | -0.78326 | 0.10181 | 0.16653 | 0.76663 | -0.38339 |
| TAG564-FA204 | 0.9833 | 0.99087 | 0.95725 | -0.06303 | 0.3613 | 0.57689 | 1.2475 | 0.31908 | 0.81442 | 0.86491 | 1.1704 | 0.22702 |
| PS160/181 | 0.70303 | 0.79734 | 1.0715 | 0.099604 | 0.74521 | 0.86786 | 1.0188 | 0.026904 | 0.3026 | 0.39163 | 1.2034 | 0.26715 |
| LPE220 | 0.76923 | 0.85165 | 1.0488 | 0.068712 | 0.50256 | 0.7097 | 1.0915 | 0.12627 | 0.91607 | 0.94661 | 0.99534 | -0.00674 |
| TAG525-FA205 | 0.55919 | 0.67663 | 0.91083 | -0.13475 | 0.2903 | 0.50531 | 0.74136 | -0.43176 | 0.3471 | 0.43792 | 0.84111 | -0.24963 |
| TAG544-FA224 | 0.93053 | 0.95995 | 1.0122 | 0.01747 | 0.24872 | 0.46261 | 1.6357 | 0.70992 | 0.75243 | 0.81098 | 1.1706 | 0.22725 |
| PEO-180/161 | 0.19256 | 0.31194 | 0.81978 | -0.28668 | 0.32693 | 0.54019 | 0.87896 | -0.18613 | 0.003326 | 0.017847 | 0.61323 | -0.70551 |
| FFA221 | 0.29238 | 0.4211 | 1.1169 | 0.15956 | 0.23273 | 0.443 | 0.84866 | -0.23674 | 0.567 | 0.65262 | 1.0465 | 0.065511 |
| PE170/224 | 0.00841 | 0.041003 | 0.71377 | -0.48647 | 0.002212 | 0.042361 | 0.58302 | -0.77839 | 0.00737 | 0.026754 | 0.64724 | -0.62764 |
| PC180/180 | 0.91579 | 0.95085 | 0.9889 | -0.0161 | 0.49284 | 0.7038 | 0.86427 | -0.21045 | 0.54465 | 0.63543 | 1.1876 | 0.24801 |

| | | | | | | | | | | | | |
|---|---|---|---|---|---|---|---|---|---|---|---|---|
| CE181 | 0.003394 | 0.028641 | 0.64396 | -0.63496 | 0.000206 | 0.019518 | 0.4663 | -1.1007 | 0.95975 | 0.97625 | 0.99392 | -0.0088 |
| PA220/226 | 0.59175 | 0.70169 | 1.1917 | 0.25302 | 0.51453 | 0.71313 | 0.86345 | -0.21182 | 0.53988 | 0.63395 | 1.2338 | 0.30312 |
| PEO-160/181 | 0.26521 | 0.3932 | 0.86522 | -0.20886 | 0.10658 | 0.26811 | 0.77403 | -0.36954 | 0.039663 | 0.081949 | 0.74723 | -0.42038 |
| TAG482-FA141 | 0.87205 | 0.92309 | 1.599 | 0.67715 | 0.75843 | 0.87542 | 1.2943 | 0.37212 | 0.12018 | 0.18808 | 0.51086 | -0.96899 |
| FFA150 | 0.21905 | 0.33952 | 1.0461 | 0.065028 | 0.67329 | 0.82838 | 0.9795 | -0.02988 | 0.42041 | 0.51541 | 1.0822 | 0.11399 |
| LPC160 | 0.95997 | 0.978 | 0.99167 | -0.01207 | 0.96454 | 0.98419 | 0.99666 | -0.00483 | 0.33331 | 0.42379 | 1.1369 | 0.18512 |
| TAG568-FA226 | 0.84361 | 0.90476 | 1.0454 | 0.064053 | 0.52372 | 0.71929 | 1.1152 | 0.15728 | 0.15182 | 0.22513 | 0.74326 | -0.42805 |
| LPE201 | 0.78903 | 0.86253 | 1.0093 | 0.013341 | 0.77253 | 0.88386 | 0.91175 | -0.13329 | 0.26587 | 0.35179 | 0.78214 | -0.35449 |
| PE160/224 | 0.51314 | 0.64493 | 0.83825 | -0.25455 | 0.16945 | 0.3677 | 0.67284 | -0.57167 | 0.056476 | 0.10565 | 0.5858 | -0.77151 |
| PC160/203 | 0.8412 | 0.90367 | 0.94172 | -0.08663 | 0.60106 | 0.77434 | 0.89926 | -0.15318 | 0.83573 | 0.87894 | 1.0573 | 0.080365 |
| FFA222 | 0.90052 | 0.94516 | 0.937 | -0.09387 | 0.71486 | 0.8506 | 0.84499 | -0.24299 | 0.25487 | 0.34109 | 1.2269 | 0.29498 |
| HCER201 | 0.49227 | 0.62837 | 0.94451 | -0.08236 | 0.019012 | 0.092363 | 0.80838 | -0.3069 | 0.12858 | 0.19696 | 0.85606 | -0.22421 |
| TAG544-FA204 | 0.52265 | 0.65133 | 1.0886 | 0.12251 | 0.65783 | 0.81883 | 1.2099 | 0.27492 | 0.96548 | 0.97749 | 1.0965 | 0.1329 |
| TAG567-FA226 | 0.9726 | 0.9847 | 1.0532 | 0.074749 | 0.27221 | 0.48154 | 1.2564 | 0.32928 | 0.38391 | 0.47787 | 0.86387 | -0.21111 |
| PC200/225 | 0.23798 | 0.36113 | 0.81597 | -0.29342 | 0.047385 | 0.15915 | 0.68595 | -0.54383 | 0.78697 | 0.83849 | 0.95514 | -0.06622 |
| TAG503-FA141 | 0.69712 | 0.79337 | 1.4197 | 0.50558 | 0.8196 | 0.90896 | 1.0759 | 0.10555 | 0.084213 | 0.14427 | 0.47133 | -1.0852 |
| TAG565-FA225 | 0.64818 | 0.75217 | 0.91714 | -0.12478 | 0.7614 | 0.8773 | 1.0614 | 0.085997 | 0.25171 | 0.33927 | 0.7851 | -0.34905 |
| PC160/120 | 0.58481 | 0.69473 | 0.89619 | -0.15813 | 0.44504 | 0.65696 | 0.86506 | -0.20913 | 0.18619 | 0.26698 | 0.71081 | -0.49247 |
| FFA200 | 0.18552 | 0.30653 | 1.3297 | 0.41115 | 0.4376 | 0.65339 | 1.1226 | 0.1668 | 0.031528 | 0.070601 | 1.53 | 0.61351 |
| PC180/224 | 0.38808 | 0.52705 | 0.87737 | -0.18874 | 0.47654 | 0.69157 | 0.89612 | -0.15824 | 0.46269 | 0.5578 | 0.89554 | -0.15917 |
| PC200/141 | 0.27826 | 0.4068 | 0.85576 | -0.22472 | 0.14476 | 0.32836 | 0.8204 | -0.2856 | 0.28596 | 0.37232 | 1.145 | 0.19538 |
| PS161/180_2 | 0.41226 | 0.54772 | 1.1979 | 0.26055 | 0.88031 | 0.93488 | 1.1014 | 0.13937 | 0.42963 | 0.52475 | 1.3036 | 0.38247 |
| LPE161 | 0.81037 | 0.87664 | 1.058 | 0.081283 | 0.72573 | 0.85435 | 0.89749 | -0.15603 | 0.36855 | 0.4614 | 0.77394 | -0.36971 |
| PC200/204 | 0.16 | 0.27285 | 0.85326 | -0.22893 | 0.000423 | 0.025021 | 0.65813 | -0.60355 | 0.54102 | 0.63395 | 0.94315 | -0.08444 |
| SMd181/183 | 0.056835 | 0.13406 | 0.88746 | -0.17224 | 0.007692 | 0.072759 | 0.79431 | -0.33223 | 0.020473 | 0.05186 | 0.85014 | -0.23424 |
| PC180/200 | 0.95317 | 0.97412 | 0.9826 | -0.02533 | 0.65753 | 0.81883 | 0.92022 | -0.11994 | 0.47769 | 0.57165 | 1.2083 | 0.27294 |
| PE160/181 | 0.60451 | 0.71163 | 0.90157 | -0.14948 | 0.2796 | 0.49015 | 0.79029 | -0.33955 | 0.018451 | 0.049393 | 0.54807 | -0.86757 |
| PEP-160/201 | 0.64135 | 0.74556 | 0.94581 | -0.08038 | 0.45457 | 0.66512 | 0.8219 | -0.28296 | 0.67419 | 0.75153 | 1.2244 | 0.29207 |
| PC180/120 | 0.51317 | 0.64493 | 0.86236 | -0.21363 | 0.19786 | 0.40506 | 0.7058 | -0.50267 | 0.095007 | 0.15778 | 0.60557 | -0.72365 |
| PEP-180/226 | 0.8655 | 0.92066 | 0.94427 | -0.08273 | 0.56183 | 0.74076 | 0.85715 | -0.22238 | 0.63449 | 0.7171 | 1.1316 | 0.17838 |
| DAG200/200 | 0.052847 | 0.12837 | 0.26447 | -1.9188 | 0.17225 | 0.37132 | 0.35473 | -1.4952 | 0.85845 | 0.89703 | 0.63697 | -0.6507 |
| PEP-160/161 | 0.81178 | 0.87664 | 1.2997 | 0.3782 | 0.81546 | 0.90706 | 1.3549 | 0.43822 | 0.82495 | 0.87122 | 1.2903 | 0.36775 |
| CER201 | 0.51794 | 0.64842 | 0.87031 | -0.2004 | 0.62195 | 0.78973 | 0.86169 | -0.21476 | 0.05829 | 0.10873 | 0.57564 | -0.79676 |

| | | | | | | | | | | | | |
|---|---|---|---|---|---|---|---|---|---|---|---|---|
| PC170/204 | 0.007292 | 0.037978 | 0.78883 | -0.34221 | 0.000148 | 0.019518 | 0.63958 | -0.6448 | 0.4847 | 0.57898 | 0.95267 | -0.06995 |
| PE180/205 | 0.6127 | 0.71739 | 0.87687 | -0.18957 | 0.11732 | 0.28499 | 0.61631 | -0.69827 | 0.11524 | 0.18344 | 0.60101 | -0.73454 |
| FFA224 | 0.11286 | 0.20933 | 0.78716 | -0.34528 | 0.024437 | 0.10618 | 0.66145 | -0.59629 | 0.15163 | 0.22513 | 1.1693 | 0.2257 |
| PC140/224 | 0.88739 | 0.93662 | 0.9477 | -0.0775 | 0.55197 | 0.73614 | 0.88221 | -0.18081 | 0.05623 | 0.10549 | 0.62573 | -0.67639 |

**Supplementary Table 4: Pathway perturbation upon irradiation in NHPs.**

| Lipidomics Mode | 6 Gy | | | | | | 6.5 Gy | | | | | | 7 Gy | | | | | | 7.5 Gy | | | | | | 8 Gy | | | | | | 8.5 Gy | | | | | |
|---|---|---|---|---|---|---|---|---|---|---|---|---|---|---|---|---|---|---|---|---|---|---|---|---|---|---|---|---|---|---|---|---|---|---|---|---|
| | SD 1 vs. Pre | | SD 2 vs. Pre | | SD 6 vs. Pre | | SD 1 vs. Pre | | SD 2 vs. Pre | | SD 6 vs. Pre | | SD 1 vs. Pre | | SD 2 vs. Pre | | SD 6 vs. Pre | | SD 1 vs. Pre | | SD 2 vs. Pre | | SD 6 vs. Pre | | SD 1 vs. Pre | | SD 2 vs. Pre | | SD 6 vs. Pre | | SD 1 vs. Pre | | SD 2 vs. Pre | | SD 6 vs. Pre | |
| Pathway | *overlap size* | *p-value* | *overlap size* | *p-value* | *overlap size* | *p-value* | *overlap size* | *p-value* | *overlap size* | *p-value* | *overlap size* | *p-value* | *overlap size* | *p-value* | *overlap size* | *p-value* | *overlap size* | *p-value* | *overlap size* | *p-value* | *overlap size* | *p-value* | *overlap size* | *p-value* | *overlap size* | *p-value* | *overlap size* | *p-value* | *overlap size* | *p-value* | *overlap size* | *p-value* | *overlap size* | *p-value* | *overlap size* | *p-value* |
| **C21-steroid hormone biosynthesis and metabolism** | 5(6) | 8.40E-05 | 5(6) | 8.40E-05 | 4(6) | 1.01E-03 | 3(6) | 4.60E-02 | 4(6) | 3.58E-02 | 3(6) | 2.98E-02 | 3(6) | 1.18E-02 | - | - | - | - | 4(6) | 4.20E-04 | 3(6) | 4.62E-02 | - | - | 4(6) | 8.15E-03 | 3(6) | 3.60E-02 | - | - | 4(6) | 1.48E-02 | 3(6) | 1.49E-02 | - | - |
| **Carnitine shuttle** | - | - | - | - | - | - | 6(16) | 4.27E-02 | - | - | - | - | 12(16) | 1.68E-04 | - | - | - | - | 12(16) | 1.68E-04 | - | - | 11(16) | 7.39E-03 | - | - | - | - | 12(16) | 1.87E-02 | - | - | - | - | - | - |
| **Tyrosine metabolism** | - | - | - | - | - | - | - | - | 3(5) | 1.99E-02 | 4(5) | 5.04E-03 | 4(5) | 1.62E-02 | 3(5) | 2.29E-02 | - | - | 4(5) | 7.48E-03 | 3(5) | 6.55E-03 | 4(5) | 2.05E-02 | 4(5) | 2.86E-03 | 3(5) | 1.68E-03 | 5(5) | 4.29E-03 | 4(5) | 6.13E-03 | 4(5) | 4.20E-04 | 4(5) | 1.17E-02 |
| **Pyruvate Metabolism** | 3(3) | 6.72E-04 | - | - | - | - | - | - | 3(3) | 1.85E-03 | 3(3) | 7.56E-04 | - | - | - | - | 3(3) | 3.70E-03 | - | - | - | - | 3(3) | 5.88E-04 | - | - | 2(3) | 1.63E-02 | - | - | - | - | - | - | 3(3) | 4.20E-04 |
| **Glycolysis and Gluconeogenesis** | 2(3) | 1.95E-02 | 1(1) | 4.80E-02 | 1(1) | 4.85E-02 | - | - | 3(3) | 1.85E-03 | 2(3) | 2.61E-02 | - | - | - | - | 3(3) | 8.40E-04 | - | - | 2(3) | 2.64E-02 | 2(3) | 2.63E-02 | - | - | 3(3) | 4.20E-04 | - | - | - | - | - | - | 2(3) | 1.70E-02 |
| **Glycine, serine, alanine and threonine metabolism** | 4(6) | 1.51E-03 | - | - | 3(3) | 5.88E-04 | 2(3) | 2.94E-02 | - | - | 4(6) | 4.12E-03 | - | - | - | - | 4(6) | 1.91E-02 | - | - | - | - | 4(6) | 3.45E-03 | - | - | 3(6) | 1.34E-02 | 3(3) | 3.06E-02 | - | - | - | - | 4(6) | 2.10E-03 |
| **Butanoate metabolism** | 2(3) | 2.85E-02 | - | - | 3(3) | 5.88E-04 | 2(3) | 2.94E-02 | - | - | 3(3) | 4.20E-03 | - | - | - | - | 3(3) | 1.68E-03 | - | - | - | - | 3(3) | 9.33E-03 | 3(3) | 5.21E-03 | - | - | 3(3) | 3.06E-02 | - | - | - | - | 3(3) | 3.02E-03 |
| **Phosphatidylinositol phosphate metabolism** | - | - | - | - | 2(4) | 2.54E-02 | - | - | 3(4) | 1.07E-02 | - | - | - | - | - | - | 4(4) | 5.88E-04 | - | - | - | - | - | - | - | - | 2(4) | 3.16E-02 | 3(4) | 1.08E-02 | - | - | - | - | 3(4) | 3.11E-03 |
| **Bile acid biosynthesis** | - | - | - | - | - | - | - | - | 5(6) | 7.56E-04 | - | - | - | - | - | - | - | - | - | - | - | - | 3(6) | 3.71E-02 | - | - | - | - | 4(6) | 1.08E-02 | - | - | - | - | - | - |
| **Glycerophospholipid metabolism** | - | - | 7(15) | 4.79E-03 | - | - | 9(15) | 2.32E-02 | 7(10) | 7.56E-04 | 7(15) | 4.35E-02 | - | - | 3(10) | 4.54E-02 | 5(10) | 7.31E-03 | - | - | 5(10) | 6.89E-03 | 9(15) | 3.67E-02 | - | - | 5(10) | 1.30E-02 | - | - | 9(15) | 4.80E-02 | - | - | 4(10) | 3.47E-02 |
| **Prostaglandin formation from arachidonate** | - | - | - | - | - | - | - | - | 4(4) | 1.43E-03 | 3(4) | 4.20E-03 | 3(4) | 2.35E-03 | 3(4) | 9.24E-04 | 3(4) | 1.88E-02 | - | - | - | - | - | - | - | - | - | - | - | - | 3(5) | 4.85E-02 | - | - | 3(4) | 5.97E-03 |
| **De novo fatty acid biosynthesis** | - | - | - | - | - | - | - | - | - | - | - | - | - | - | - | - | - | - | - | - | - | - | - | - | - | - | - | - | 5(7) | 2.44E-03 | 5(7) | 1.01E-03 | - | - | - | - |
| **Fatty acid activation** | - | - | - | - | - | - | - | - | - | - | - | - | - | - | - | - | 3(3) | 3.70E-03 | - | - | - | - | 2(3) | 2.63E-02 | - | - | - | - | 3(3) | 1.34E-03 | 3(3) | 1.01E-03 | - | - | - | - |
| **Vitamin E metabolism** | - | - | - | - | - | - | - | - | - | - | 1(1) | 4.86E-02 | 1(1) | 3.59E-02 | 5(7) | 1.18E-03 | - | - | 1(1) | 2.60E-02 | 4(7) | 1.44E-02 | 5(7) | 4.39E-02 | 1(1) | 4.99E-02 | - | - | - | - | 1(1) | 4.78E-02 | 1(1) | 3.07E-02 | 5(7) | 1.59E-02 |
| **Linoleate metabolism** | - | - | - | - | - | - | - | - | - | - | - | - | - | - | - | - | - | - | - | - | - | - | - | - | 3(5) | 3.74E-02 | - | - | - | - | 5(5) | 1.26E-03 | - | - | - | - |
| **Arachidonic acid metabolism** | - | - | - | - | - | - | 3(5) | 2.73E-02 | 3(5) | 3.55E-02 | 3(5) | 1.40E-02 | 3(5) | 5.71E-03 | 3(5) | 1.51E-03 | - | - | - | - | 3(5) | 1.21E-02 | - | - | 3(5) | 1.48E-02 | - | - | - | - | 2(2) | 2.34E-02 | - | - | - | - |
| **Urea cycle/amino group metabolism** | 3(6) | 3.10E-02 | - | - | - | - | - | - | - | - | 3(6) | 3.87E-02 | - | - | - | - | - | - | 2(5) | 4.76E-02 | - | - | 2(5) | 3.90E-02 | 3(5) | 1.48E-02 | 2(6) | 2.66E-02 | 5(6) | 3.06E-02 | - | - | 4(6) | 1.76E-03 | - | - |
| **Androgen and estrogen biosynthesis and metabolism** | 2(2) | 6.22E-03 | 2(2) | 4.12E-03 | 2(2) | 3.70E-03 | - | - | 2(2) | 8.74E-03 | - | - | - | - | - | - | 2(2) | 1.01E-02 | 2(2) | 1.76E-03 | - | - | - | - | - | - | 2(2) | 1.07E-02 | - | - | 2(2) | 1.29E-02 | - | - | 2(2) | 2.78E-02 |
| **Fatty Acid Metabolism** | - | - | 2(2) | 4.54E-03 | - | - | 2(2) | 1.05E-02 | - | - | - | - | - | - | - | - | 2(2) | 1.48E-02 | - | - | 2(2) | 9.33E-03 | 2(2) | 3.68E-02 | - | - | - | - | 2(2) | 8.82E-03 | 2(2) | 4.79E-03 | 2(2) | 6.97E-03 | 2(2) | 1.66E-02 |
| **Fructose and mannose metabolism** | - | - | - | - | - | - | - | - | 2(2) | 9.24E-03 | - | - | - | - | - | - | 2(2) | 6.30E-03 | - | - | 2(2) | 5.38E-03 | - | - | - | - | 2(2) | 4.71E-03 | - | - | - | - | - | - | - | - |
| **Galactose metabolism** | - | - | - | - | - | - | - | - | 2(2) | 9.24E-03 | - | - | - | - | - | - | 2(2) | 6.30E-03 | - | - | 2(2) | 5.38E-03 | - | - | - | - | 2(2) | 4.71E-03 | - | - | - | - | - | - | - | - |
| **Xenobiotics metabolism** | - | - | - | - | - | - | - | - | - | - | - | - | - | - | - | - | - | - | - | - | - | - | - | - | - | - | 1(1) | 3.82E-02 | - | - | 2(2) | 4.79E-03 | - | - | - | - |
| **Tryptophan metabolism** | - | - | - | - | - | - | - | - | - | - | - | - | - | - | - | - | - | - | - | - | - | - | - | - | - | - | 2(4) | 2.55E-02 | - | - | 5(7) | 4.96E-03 | 2(4) | 4.28E-02 | - | - |
| **Valine, leucine and isoleucine degradation** | - | - | - | - | 2(2) | 6.55E-03 | - | - | - | - | 2(2) | 2.07E-02 | - | - | - | - | 2(2) | 1.01E-02 | - | - | - | - | 2(2) | 3.68E-02 | 2(2) | 1.93E-02 | - | - | - | - | - | - | - | - | 2(2) | 1.66E-02 |
| **Ascorbate (Vitamin C) and Aldarate Metabolism** | 1(1) | 4.55E-02 | 1(1) | 4.10E-02 | 1(1) | 3.61E-02 | - | - | - | - | 1(1) | 4.86E-02 | 1(1) | 4.22E-02 | 2(2) | 7.81E-03 | - | - | 1(1) | 2.60E-02 | - | - | 1(1) | 3.34E-02 | - | - | 1(1) | 4.25E-02 | - | - | 1(1) | 4.78E-02 | 1(1) | 3.15E-02 | 1(1) | 3.60E-02 |
| **Aspartate and asparagine metabolism** | - | - | - | - | - | - | - | - | - | - | 2(2) | 2.07E-02 | - | - | - | - | 2(2) | 3.02E-02 | - | - | - | - | 3(6) | 8.91E-03 | 2(2) | 1.33E-02 | - | - | - | - | 4(6) | 1.48E-02 | - | - | 2(2) | 2.78E-02 |
| **Sialic acid metabolism** | 2(3) | 2.85E-02 | 2(3) | 2.05E-02 | - | - | - | - | 2(2) | 9.24E-03 | - | - | - | - | - | - | - | - | - | - | - | - | - | - | - | - | - | - | - | - | - | - | - | - | 2(3) | 4.62E-02 |
| **Aminosugars metabolism** | - | - | - | - | - | - | - | - | - | - | - | - | 2(3) | 1.34E-02 | - | - | - | - | 2(3) | 9.50E-03 | - | - | - | - | 2(3) | 4.28E-02 | - | - | - | - | - | - | - | - | - | - |
| **Squalene and cholesterol biosynthesis** | - | - | - | - | - | - | 2(3) | 2.94E-02 | - | - | - | - | - | - | - | - | - | - | - | - | - | - | 3(3) | 1.13E-02 | - | - | - | - | - | - | - | - | - | - | 2(3) | 4.62E-02 |
| **Vitamin B3 (nicotinate and nicotinamide) metabolism** | - | - | - | - | - | - | - | - | - | - | - | - | - | - | - | - | - | - | - | - | - | - | - | - | - | - | - | - | - | - | 3(4) | 1.48E-02 | - | - | - | - |
| **Alanine and Aspartate Metabolism** | - | - | - | - | - | - | - | - | - | - | - | - | 2(4) | 3.80E-02 | - | - | - | - | 2(4) | 2.74E-02 | - | - | 2(4) | 2.44E-02 | - | - | - | - | - | - | 3(4) | 1.48E-02 | - | - | - | - |
| **Glycosphingolipid metabolism** | - | - | - | - | - | - | - | - | - | - | - | - | - | - | - | - | 2(2) | 1.48E-02 | 3(4) | 2.70E-02 | - | - | - | - | - | - | - | - | - | - | - | - | - | - | - | - |
| **Polyunsaturated fatty acid biosynthesis** | - | - | - | - | - | - | - | - | - | - | - | - | - | - | 1(1) | 1.72E-02 | - | - | 1(1) | 2.60E-02 | 1(1) | 3.39E-02 | - | - | - | - | 1(1) | 4.25E-02 | - | - | 1(1) | 4.78E-02 | 1(1) | 3.07E-02 | 1(1) | 3.60E-02 |
| **Omega-6 fatty acid metabolism** | - | - | - | - | - | - | - | - | - | - | 1(1) | 4.86E-02 | 1(1) | 3.59E-02 | 1(1) | 1.72E-02 | - | - | - | - | 1(1) | 3.39E-02 | - | - | 1(1) | 4.99E-02 | - | - | - | - | 1(1) | 4.78E-02 | 1(1) | 3.07E-02 | - | - |
| **Sphingolipid metabolism** | - | - | 1(1) | 4.80E-02 | 1(1) | 4.85E-02 | - | - | - | - | - | - | - | - | 1(1) | 4.09E-02 | - | - | - | - | 1(1) | 3.63E-02 | - | - | - | - | 1(1) | 2.24E-02 | - | - | - | - | - | - | - | - |
| **Arginine and Proline Metabolism** | - | - | - | - | - | - | - | - | - | - | - | - | - | - | - | - | - | - | - | - | - | - | - | - | 2(3) | 2.60E-02 | 1(1) | 2.24E-02 | - | - | - | - | - | - | - | - |
| **Lysine metabolism** | - | - | 1(1) | 4.80E-02 | 1(1) | 4.85E-02 | - | - | - | - | - | - | - | - | 1(1) | 4.09E-02 | - | - | - | - | 1(1) | 3.63E-02 | - | - | - | - | 1(1) | 2.24E-02 | - | - | - | - | - | - | - | - |
| **Vitamin A (retinol) metabolism** | - | - | - | - | - | - | - | - | - | - | - | - | - | - | 2(4) | 4.03E-02 | - | - | 3(4) | 2.70E-02 | 2(4) | 3.39E-02 | - | - | - | - | 2(4) | 2.55E-02 | - | - | 3(4) | 2.60E-02 | - | - | - | - |
| **Glyoxylate and Dicarboxylate Metabolism** | 1(1) | 4.55E-02 | 1(1) | 4.10E-02 | 1(1) | 3.61E-02 | - | - | - | - | 1(1) | 4.86E-02 | 1(1) | 4.22E-02 | - | - | - | - | 1(1) | 2.60E-02 | - | - | 1(1) | 3.34E-02 | - | - | 1(1) | 4.25E-02 | - | - | 1(1) | 4.78E-02 | 1(1) | 3.15E-02 | 1(1) | 3.60E-02 |
| **TCA cycle** | 1(1) | 4.55E-02 | 1(1) | 4.10E-02 | 1(1) | 3.61E-02 | - | - | - | - | 1(1) | 4.86E-02 | 1(1) | 4.22E-02 | - | - | - | - | 1(1) | 2.60E-02 | - | - | 1(1) | 3.34E-02 | - | - | 1(1) | 4.25E-02 | - | - | 1(1) | 4.78E-02 | 1(1) | 3.15E-02 | 1(1) | 3.60E-02 |
| **Proteoglycan biosynthesis** | 1(1) | 4.55E-02 | 1(1) | 4.10E-02 | 1(1) | 3.61E-02 | - | - | - | - | 1(1) | 4.86E-02 | 1(1) | 4.22E-02 | - | - | - | - | 1(1) | 2.60E-02 | - | - | 1(1) | 3.34E-02 | - | - | 1(1) | 4.25E-02 | - | - | 1(1) | 4.78E-02 | 1(1) | 3.15E-02 | 1(1) | 3.60E-02 |
| **Beta-Alanine metabolism** | - | - | - | - | - | - | - | - | - | - | - | - | - | - | - | - | - | - | - | - | 2(3) | 2.64E-02 | - | - | - | - | - | - | - | - | - | - | - | - | - | - |
| **Saturated fatty acids beta-oxidation** | - | - | - | - | 1(1) | 3.61E-02 | - | - | - | - | - | - | 2(2) | 2.71E-02 | - | - | - | - | 2(2) | 3.12E-02 | - | - | - | - | - | - | - | - | - | - | 1(1) | 4.78E-02 | - | - | 1(1) | 3.60E-02 |
| **Propanoate metabolism** | 1(1) | 4.55E-02 | - | - | 1(1) | 4.85E-02 | - | - | - | - | 1(1) | 4.86E-02 | - | - | - | - | - | - | - | - | - | - | - | - | - | - | 1(1) | 4.25E-02 | - | - | - | - | - | - | 1(1) | 3.60E-02 |
| **Histidine metabolism** | - | - | - | - | - | - | - | - | - | - | - | - | - | - | - | - | - | - | - | - | 1(1) | 3.63E-02 | - | - | - | - | - | - | - | - | - | - | - | - | - | - |
| **Porphyrin metabolism** | 1(1) | 4.55E-02 | 1(1) | 4.10E-02 | - | - | - | - | - | - | - | - | - | - | - | - | - | - | - | - | - | - | - | - | 1(1) | 4.99E-02 | - | - | - | - | - | - | - | - | - | - |
| **Heparan sulfate degradation** | - | - | - | - | - | - | - | - | - | - | - | - | - | - | - | - | - | - | - | - | - | - | - | - | - | - | 1(1) | 4.25E-02 | - | - | - | - | - | - | - | - |
| **N-Glycan Degradation** | - | - | - | - | - | - | - | - | - | - | - | - | - | - | - | - | - | - | - | - | - | - | - | - | - | - | 1(1) | 4.25E-02 | - | - | - | - | - | - | - | - |
| **Starch and Sucrose Metabolism** | - | - | 1(1) | 4.80E-02 | 1(1) | 4.85E-02 | - | - | - | - | - | - | - | - | - | - | - | - | - | - | - | - | - | - | - | - | 1(1) | 4.25E-02 | - | - | - | - | - | - | - | - |
| **Pentose phosphate pathway** | - | - | 1(1) | 4.80E-02 | 1(1) | 4.85E-02 | - | - | - | - | - | - | - | - | - | - | - | - | - | - | - | - | - | - | - | - | 1(1) | 4.25E-02 | - | - | - | - | - | - | - | - |
| **N-Glycan biosynthesis** | - | - | 1(1) | 4.80E-02 | 1(1) | 4.85E-02 | - | - | - | - | - | - | - | - | - | - | - | - | - | - | - | - | - | - | - | - | 1(1) | 4.25E-02 | - | - | - | - | - | - | - | - |
| **Chondroitin sulfate degradation** | - | - | - | - | - | - | - | - | - | - | - | - | - | - | - | - | - | - | - | - | - | - | - | - | - | - | 1(1) | 4.25E-02 | - | - | - | - | - | - | - | - |
| **Hexose phosphorylation** | - | - | - | - | - | - | - | - | - | - | - | - | - | - | - | - | - | - | - | - | - | - | - | - | - | - | 1(1) | 4.25E-02 | - | - | - | - | - | - | - | - |
| **Glycosphingolipid biosynthesis - ganglioseries** | - | - | - | - | - | - | - | - | - | - | - | - | - | - | - | - | - | - | - | - | - | - | - | - | - | - | 1(1) | 4.25E-02 | - | - | - | - | - | - | - | - |
| **Glycosphingolipid biosynthesis - globoseries** | - | - | 1(1) | 4.80E-02 | 1(1) | 4.85E-02 | - | - | - | - | - | - | - | - | - | - | - | - | - | - | - | - | - | - | - | - | 1(1) | 4.25E-02 | - | - | - | - | - | - | - | - |
| **Keratan sulfate degradation** | - | - | - | - | - | - | - | - | - | - | - | - | - | - | - | - | - | - | - | - | - | - | - | - | - | - | 1(1) | 4.25E-02 | - | - | - | - | - | - | - | - |
| **Parathio degradation** | - | - | - | - | - | - | - | - | - | - | - | - | - | - | - | - | - | - | - | - | - | - | - | - | - | - | - | - | - | - | 1(1) | 4.78E-02 | - | - | - | - |
| **Di-unsaturated fatty acid beta-oxidation** | - | - | - | - | - | - | - | - | - | - | - | - | - | - | - | - | - | - | - | - | - | - | - | - | - | - | - | - | - | - | 1(1) | 4.78E-02 | - | - | - | - |
| **Glycosylphosphatidylinositol(GPI)-anchor biosynthesis** | - | - | 1(1) | 4.80E-02 | - | - | - | - | - | - | - | - | - | - | - | - | - | - | - | - | - | - | - | - | - | - | - | - | - | - | - | - | - | - | - | - |
| **Vitamin K metabolism** | - | - | 1(1) | 4.80E-02 | 1(1) | 4.85E-02 | - | - | - | - | - | - | - | - | - | - | - | - | - | - | - | - | - | - | - | - | - | - | - | - | - | - | - | - | - | - |
| **Glutamate metabolism** | - | - | - | - | 1(1) | 4.85E-02 | - | - | - | - | - | - | - | - | - | - | - | - | - | - | - | - | - | - | - | - | - | - | - | - | - | - | - | - | - | - |
| **Pyrimidine metabolism** | - | - | - | - | 1(1) | 4.85E-02 | - | - | - | - | - | - | - | - | - | - | - | - | - | - | - | - | - | - | - | - | - | - | - | - | - | - | - | - | - | - |
| **Methionine and cysteine metabolism** | - | - | - | - | 1(1) | 4.85E-02 | - | - | 2(3) | 4.97E-02 | - | - | - | - | - | - | - | - | - | - | - | - | - | - | - | - | - | - | - | - | - | - | - | - | - | - |
| **Selenoamino acid metabolism** | - | - | - | - | 1(1) | 4.85E-02 | - | - | - | - | - | - | - | - | - | - | - | - | - | - | - | - | - | - | - | - | - | - | - | - | - | - | - | - | - | - |
| **Drug metabolism - other enzymes** | - | - | - | - | - | - | - | - | - | - | - | - | - | - | - | - | - | - | - | - | - | - | - | - | 1(1) | 4.99E-02 | - | - | - | - | - | - | - | - | - | - |

| Metabolomics Mode | 6 Gy | | | | | | 6.5 Gy | | | | | | 7 Gy | | | | | | 7.5 Gy | | | | | | 8 Gy | | | | | | 8.5 Gy | | | | | |
|---|---|---|---|---|---|---|---|---|---|---|---|---|---|---|---|---|---|---|---|---|---|---|---|---|---|---|---|---|---|---|---|---|---|---|---|---|
| | SD 1 vs. Pre | | SD 2 vs. Pre | | SD 6 vs. Pre | | SD 1 vs. Pre | | SD 2 vs. Pre | | SD 6 vs. Pre | | SD 1 vs. Pre | | SD 2 vs. Pre | | SD 6 vs. Pre | | SD 1 vs. Pre | | SD 2 vs. Pre | | SD 6 vs. Pre | | SD 1 vs. Pre | | SD 2 vs. Pre | | SD 6 vs. Pre | | SD 1 vs. Pre | | SD 2 vs. Pre | | SD 6 vs. Pre | |
| Pathway | *overlap size* | *p-value* | *overlap size* | *p-value* | *overlap size* | *p-value* | *overlap size* | *p-value* | *overlap size* | *p-value* | *overlap size* | *p-value* | *overlap size* | *p-value* | *overlap size* | *p-value* | *overlap size* | *p-value* | *overlap size* | *p-value* | *overlap size* | *p-value* | *overlap size* | *p-value* | *overlap size* | *p-value* | *overlap size* | *p-value* | *overlap size* | *p-value* | *overlap size* | *p-value* | *overlap size* | *p-value* | *overlap size* | *p-value* |
| **Aspartate and asparagine metabolism** | - | - | 4(11) | 0.0484 | - | - | 3(4) | 0.02286 | 8(11) | 2.52E-04 | 8(11) | 0.0026 | 8(11) | 0.00101 | - | - | 8(11) | 0.00126 | 6(11) | 0.00723 | 3(11) | 0.01924 | 8(11) | 5.88E-04 | 8(11) | 0.00252 | 7(11) | 5.04E-04 | 9(11) | 0.00269 | 8(11) | 0.00882 | 9(11) | 8.40E-05 | 4(4) | 8.40E-04 |
| **Bile acid biosynthesis** | - | - | - | - | 11(20) | 1.68E-04 | - | - | - | - | 8(20) | 0.04605 | - | - | - | - | 9(20) | 0.02201 | - | - | - | - | 11(20) | 0.00193 | - | - | - | - | 12(14) | 8.40E-05 | - | - | - | - | - | - |
| **Tyrosine metabolism** | - | - | 6(9) | 2.52E-04 | 4(9) | 0.04748 | 7(9) | 9.24E-04 | 8(9) | 4.20E-04 | 6(9) | 0.00941 | 6(9) | 0.0016 | 7(9) | 2.52E-04 | - | - | 5(9) | 0.00412 | 6(9) | 1.68E-04 | 6(9) | 0.00807 | 7(9) | 8.40E-05 | 7(9) | 1.68E-04 | 6(9) | 0.03252 | 5(9) | 0.01479 | 7(9) | 2.52E-04 | - | - |
| **Arachidonic acid metabolism** | - | - | - | - | - | - | 7(8) | 8.40E-05 | 5(8) | 3.87E-02 | 4(8) | 0.02143 | 6(8) | 5.88E-04 | 5(8) | 0.00429 | 5(8) | 0.00496 | - | - | - | - | - | - | - | - | 4(8) | 0.01286 | - | - | 6(8) | 0.0163 | 4(8) | 2.45E-02 | - | - |
| **Arginine and Proline Metabolism** | - | - | 4(8) | 0.01706 | - | - | 2(2) | 0.02344 | 6(8) | 7.56E-04 | 6(8) | 0.0058 | 6(8) | 0.00227 | - | - | 6(8) | 0.0026 | 5(8) | 0.00655 | - | - | 6(8) | 0.00202 | 6(8) | 0.00479 | 5(8) | 0.00269 | 6(8) | 2.29E-02 | - | - | 7(8) | 1.68E-04 | 4(8) | 0.03974 |
| **De novo fatty acid biosynthesis** | - | - | - | - | - | - | - | - | - | - | - | - | - | - | - | - | - | - | - | - | - | - | - | - | - | - | - | - | 20(24) | 1.68E-04 | 12(24) | 0.01891 | 8(24) | 0.03991 | - | - |
| **Androgen and estrogen biosynthesis and metabolism** | 8(10) | 2.52E-04 | 6(10) | 0.00168 | 4(10) | 0.02731 | 9(10) | 1.68E-04 | 7(10) | 6.72E-04 | 8(10) | 0.0016 | 6(10) | 0.01067 | - | - | - | - | 6(10) | 0.00395 | 3(10) | 0.04495 | - | - | 8(10) | 4.20E-04 | - | - | 8(10) | 9.50E-03 | 8(10) | 0.00429 | 4(10) | 0.04663 | 7(10) | 0.01168 |
| **Carbon fixation** | 4(4) | 1.68E-04 | - | - | - | - | - | - | - | - | - | - | - | - | - | - | 4(4) | 0.00765 | - | - | - | - | 4(4) | 2.27E-03 | - | - | - | - | 4(4) | 0.00723 | - | - | - | - | - | - |
| **C21-steroid hormone biosynthesis and metabolism** | 17(34) | 4.20E-04 | 10(34) | 0.03781 | 6(13) | 0.01849 | 18(34) | 2.52E-04 | 16(34) | 3.36E-04 | 16(34) | 0.04 | 18(34) | 5.88E-04 | - | - | 8(13) | 0.03655 | 16(34) | 0.00101 | - | - | 7(13) | 0.04571 | 20(34) | 7.56E-04 | - | - | 25(34) | 2.52E-04 | 8(13) | 0.0042 | - | - | 21(34) | 0.00109 |
| **Carnitine shuttle** | - | - | - | - | - | - | - | - | - | - | - | - | 13(20) | 5.04E-04 | - | - | 15(20) | 0.00445 | 12(20) | 0.00176 | 5(20) | 0.00891 | 15(20) | 3.36E-04 | - | - | - | - | - | - | - | - | - | - | 13(20) | 0.00311 |
| **Glycine, serine, alanine and threonine metabolism** | - | - | 4(7) | 0.01151 | 3(6) | 0.04823 | - | - | 5(7) | 0.00328 | 4(7) | 0.03033 | 4(7) | 0.03327 | 6(7) | 5.88E-04 | - | - | 5(7) | 0.00924 | 3(7) | 0.0063 | - | - | - | - | 4(7) | 0.00487 | - | - | - | - | - | - | - | - |
| **Butanoate metabolism** | 4(7) | 0.02412 | 3(7) | 0.04512 | - | - | 6(7) | 0.00218 | - | - | 5(7) | 0.00555 | 6(7) | 0.00101 | 5(7) | 0.00328 | 4(7) | 0.03025 | 5(7) | 0.00924 | 2(7) | 0.04327 | 4(7) | 0.02857 | 6(7) | 9.24E-04 | 4(7) | 0.00933 | - | - | 6(7) | 0.00588 | 4(7) | 0.01991 | 5(7) | 0.00403 |
| **Galactose metabolism** | 4(7) | 0.00429 | - | - | - | - | - | - | 5(7) | 0.01445 | - | - | - | - | 4(7) | 0.01798 | 6(7) | 0.00571 | - | - | - | - | 6(7) | 0.00134 | - | - | 3(7) | 0.04865 | 5(7) | 0.03899 | - | - | 4(7) | 0.02521 | - | - |
| **Fructose and mannose metabolism** | 4(7) | 0.00429 | - | - | - | - | - | - | 5(7) | 0.01445 | - | - | - | - | 4(7) | 0.01798 | 6(7) | 0.00571 | - | - | - | - | 6(7) | 0.00134 | - | - | 3(7) | 0.04865 | 5(7) | 0.03899 | - | - | 4(7) | 0.02521 | - | - |
| **Valine, leucine and isoleucine degradation** | - | - | 2(3) | 0.03243 | - | - | - | - | 3(3) | 0.00454 | 3(3) | 0.00597 | 3(3) | 0.00739 | 3(3) | 0.00454 | 3(3) | 0.00773 | 3(3) | 0.00924 | 2(3) | 0.00773 | 3(3) | 0.00487 | 3(3) | 0.00546 | 3(3) | 0.00151 | 3(3) | 0.02722 | 3(3) | 0.00891 | 3(3) | 0.00353 | 3(3) | 0.00445 |
| **Purine metabolism** | - | - | 2(3) | 0.03243 | 2(3) | 0.02756 | 2(2) | 0.01328 | 2(2) | 0.04327 | - | - | 2(2) | 0.01218 | 2(2) | 0.01672 | - | - | 3(3) | 0.00924 | 2(2) | 0.00555 | 3(3) | 0.00487 | 3(3) | 0.00546 | 3(3) | 0.00151 | 3(3) | 0.02722 | 2(2) | 0.01613 | 3(3) | 0.00353 | - | - |
| **Omega-6 fatty acid metabolism** | 3(4) | 0.01697 | - | - | - | - | 3(4) | 0.0358 | - | - | - | - | - | - | 3(4) | 0.01353 | - | - | 3(4) | 0.01395 | 3(4) | 0.00151 | - | - | 3(4) | 0.02084 | 3(4) | 0.00487 | 4(4) | 0.01319 | 4(4) | 0.00244 | 3(4) | 0.0126 | - | - |
| **Methionine and cysteine metabolism** | - | - | - | - | - | - | 4(4) | 0.00328 | 3(4) | 0.01546 | 4(4) | 0.00328 | 3(4) | 0.03428 | 3(4) | 0.01353 | - | - | 4(4) | 0.0016 | 2(4) | 0.01269 | - | - | 4(4) | 0.00328 | 3(4) | 0.00487 | - | - | 4(4) | 0.00739 | 3(4) | 0.0126 | - | - |
| **Alanine and Aspartate Metabolism** | - | - | 3(3) | 0.0021 | - | - | - | - | 2(3) | 0.04369 | - | - | - | - | 2(3) | 0.04075 | - | - | 3(3) | 0.0042 | 1(1) | 0.04084 | - | - | - | - | 2(3) | 0.02201 | - | - | - | - | 2(3) | 0.04025 | - | - |
| **Omega-3 fatty acid metabolism** | - | - | - | - | - | - | 3(3) | 0.01092 | 2(3) | 0.04369 | - | - | - | - | 2(3) | 0.04958 | - | - | - | - | - | - | - | - | - | - | 3(3) | 0.0026 | - | - | - | - | 2(3) | 0.04025 | - | - |
| **Fatty acid activation** | - | - | - | - | - | - | - | - | - | - | - | - | - | - | - | - | 7(11) | 0.03605 | - | - | - | - | - | - | - | - | - | - | 13(17) | 0.00269 | 6(11) | 0.02025 | 6(17) | 0.04277 | - | - |
| **Vitamin A (retinol) metabolism** | - | - | - | - | - | - | - | - | - | - | - | - | - | - | 3(5) | 0.02739 | - | - | - | - | 3(5) | 0.00277 | - | - | 3(5) | 0.04008 | - | - | - | - | - | - | - | - | - | - |
| **Linoleate metabolism** | - | - | - | - | - | - | - | - | - | - | - | - | - | - | - | - | 7(11) | 0.03605 | - | - | - | - | - | - | 5(11) | 0.00748 | - | - | - | - | 7(11) | 0.00563 | 6(11) | 0.00294 | - | - |
| **Tryptophan metabolism** | - | - | - | - | - | - | - | - | - | - | - | - | - | - | - | - | - | - | - | - | - | - | - | - | - | - | 6(9) | 0.00319 | - | - | - | - | - | - | - | - |
| **Porphyrin metabolism** | - | - | - | - | - | - | - | - | - | - | - | - | - | - | - | - | 4(5) | 0.02874 | - | - | - | - | 4(5) | 0.01445 | - | - | - | - | 5(5) | 0.00328 | - | - | - | - | 3(3) | 0.01664 |
| **Glutamate metabolism** | - | - | - | - | - | - | 2(2) | 0.02143 | 2(2) | 0.0179 | 2(2) | 0.02638 | 2(2) | 0.02672 | 2(2) | 0.01773 | 2(2) | 0.02613 | 2(2) | 0.02067 | 2(2) | 0.00353 | 2(2) | 0.0226 | 2(2) | 0.02386 | 2(2) | 0.00807 | - | - | 2(2) | 0.03117 | 2(2) | 0.01639 | 2(2) | 0.01462 |
| **Glycolysis and Gluconeogenesis** | 4(8) | 0.00857 | - | - | - | - | - | - | 6(8) | 0.00605 | - | - | - | - | 5(8) | 0.00487 | 6(8) | 0.01714 | - | - | - | - | 6(8) | 0.00387 | - | - | 4(8) | 0.02176 | - | - | - | - | 5(8) | 0.00739 | - | - |
| **Glycosphingolipid biosynthesis - globoseries** | - | - | 3(4) | 0.00513 | - | - | - | - | 4(4) | 0.00387 | - | - | - | - | - | - | - | - | - | - | - | - | - | - | - | - | - | - | - | - | - | - | - | - | 3(4) | 0.01554 |
| **Lysine metabolism** | - | - | - | - | - | - | - | - | 4(5) | 0.0042 | - | - | - | - | 3(5) | 0.02739 | - | - | - | - | 2(5) | 0.01891 | - | - | 4(5) | 0.01193 | 3(5) | 0.00992 | - | - | - | - | - | - | 3(5) | 0.03353 |
| **Urea cycle/amino group metabolism** | - | - | 4(8) | 0.01706 | - | - | - | - | 4(8) | 0.03033 | 5(8) | 0.01185 | 5(8) | 0.02487 | 4(8) | 0.03529 | - | - | 4(8) | 0.02983 | - | - | 5(8) | 0.00975 | 2(2) | 0.01008 | 4(8) | 0.01286 | - | - | 2(2) | 0.01613 | 5(8) | 0.00429 | 2(2) | 0.01773 |
| **Ascorbate (Vitamin C) and Aldarate Metabolism** | - | - | - | - | 2(2) | 0.01512 | - | - | 2(2) | 0.04327 | 2(2) | 0.03134 | - | - | 2(2) | 0.01672 | 2(2) | 0.01538 | 2(2) | 0.00429 | - | - | 2(2) | 0.03361 | - | - | 2(2) | 0.0137 | - | - | - | - | 2(2) | 0.01891 | 3(4) | 0.0163 |
| **TCA cycle** | - | - | - | - | - | - | 2(2) | 0.02344 | 2(2) | 0.04605 | - | - | 2(2) | 0.01218 | 2(2) | 0.01672 | - | - | 2(2) | 0.0095 | 2(2) | 0.00555 | - | - | - | - | 2(2) | 0.0137 | - | - | 2(2) | 0.01613 | 2(2) | 0.01151 | - | - |
| **Aminosugars metabolism** | - | - | - | - | - | - | - | - | - | - | 5(7) | 0.01706 | - | - | - | - | 4(7) | 0.03025 | 1(1) | 0.04311 | 1(1) | 0.04084 | - | - | - | - | - | - | - | - | 6(7) | 0.00588 | - | - | - | - |
| **Sialic acid metabolism** | - | - | 4(7) | 0.00655 | - | - | - | - | - | - | - | - | - | - | - | - | - | - | - | - | - | - | - | - | - | - | - | - | - | - | 4(7) | 0.01983 | - | - | 4(7) | 0.02378 |
| **Leukotriene metabolism** | - | - | - | - | - | - | 2(2) | 0.01328 | 2(2) | 0.04327 | - | - | 2(2) | 0.01218 | 2(2) | 0.01672 | - | - | - | - | - | - | - | - | 3(4) | 0.03235 | - | - | - | - | 4(4) | 0.00739 | - | - | - | - |
| **Glyoxylate and Dicarboxylate Metabolism** | 2(2) | 0.00739 | - | - | - | - | - | - | - | - | - | - | - | - | - | - | - | - | - | - | - | - | 2(2) | 0.03319 | - | - | - | - | - | - | - | - | - | - | - | - |
| **Fatty Acid Metabolism** | - | - | - | - | - | - | - | - | - | - | - | - | - | - | - | - | 4(4) | 0.00765 | - | - | - | - | - | - | - | - | - | - | 9(14) | 0.04311 | 3(4) | 0.02286 | - | - | - | - |
| **Pyrimidine metabolism** | - | - | - | - | - | - | - | - | 2(3) | 0.04369 | - | - | - | - | 2(3) | 0.04075 | - | - | - | - | 2(3) | 0.00773 | - | - | - | - | 2(3) | 0.02201 | - | - | - | - | 2(3) | 0.04025 | 3(4) | 0.01126 |
| **Pentose phosphate pathway** | - | - | - | - | - | - | - | - | 4(5) | 0.01428 | - | - | - | - | - | - | 4(5) | 0.02874 | - | - | - | - | 4(5) | 0.00924 | - | - | - | - | 4(5) | 0.03319 | - | - | - | - | 3(5) | 0.03134 |
| **Beta-Alanine metabolism** | - | - | - | - | - | - | - | - | 2(2) | 0.04327 | 3(3) | 0.01218 | - | - | 2(3) | 0.04958 | - | - | - | - | - | - | 2(2) | 0.03361 | 3(3) | 0.0095 | 2(3) | 0.02764 | - | - | - | - | 2(2) | 0.01891 | - | - |
| **Glycerophospholipid metabolism** | - | - | - | - | - | - | 12(29) | 0.01218 | - | - | - | - | - | - | - | - | 7(18) | 0.03832 | - | - | - | - | - | - | - | - | 8(29) | 0.03949 | - | - | - | - | - | - | - | - |
| **Glutathione Metabolism** | - | - | - | - | - | - | 3(4) | 0.01874 | - | - | 3(4) | 0.04504 | 3(4) | 0.02302 | - | - | 3(4) | 0.02302 | - | - | 2(4) | 0.01269 | - | - | 3(4) | 0.02084 | 2(4) | 0.03832 | - | - | 3(4) | 0.03235 | 3(4) | 0.01588 | - | - |
| **Vitamin B3 (nicotinate and nicotinamide) metabolism** | - | - | 2(2) | 0.01344 | - | - | - | - | - | - | - | - | - | - | - | - | - | - | 2(2) | 0.02067 | - | - | - | - | - | - | - | - | - | - | - | - | 2(3) | 0.03697 | - | - |
| **N-Glycan biosynthesis** | - | - | - | - | - | - | 2(2) | 0.02344 | 2(2) | 0.04327 | - | - | - | - | - | - | - | - | - | - | - | - | - | - | - | - | 2(2) | 0.0137 | - | - | - | - | - | - | - | - |
| **Chondroitin sulfate degradation** | - | - | - | - | - | - | 2(2) | 0.02344 | 2(2) | 0.04327 | - | - | - | - | - | - | - | - | - | - | - | - | - | - | - | - | 2(2) | 0.0137 | - | - | - | - | - | - | - | - |
| **Hexose phosphorylation** | - | - | - | - | - | - | 2(2) | 0.02344 | 2(2) | 0.04327 | - | - | - | - | - | - | - | - | - | - | - | - | - | - | - | - | 2(2) | 0.0137 | - | - | - | - | - | - | - | - |
| **Keratan sulfate degradation** | - | - | - | - | - | - | 2(2) | 0.02344 | 2(2) | 0.04327 | - | - | - | - | - | - | - | - | - | - | - | - | - | - | - | - | 2(2) | 0.0137 | - | - | - | - | - | - | - | - |
| **Heparan sulfate degradation** | - | - | - | - | - | - | 2(2) | 0.02344 | 2(2) | 0.04327 | - | - | - | - | - | - | - | - | - | - | - | - | - | - | - | - | 2(2) | 0.0137 | - | - | - | - | - | - | - | - |
| **N-Glycan Degradation** | - | - | - | - | - | - | 2(2) | 0.02344 | 2(2) | 0.04327 | - | - | - | - | - | - | - | - | - | - | - | - | - | - | - | - | 2(2) | 0.0137 | - | - | - | - | - | - | - | - |
| **Starch and Sucrose Metabolism** | - | - | - | - | - | - | 2(2) | 0.02344 | 2(2) | 0.04327 | - | - | - | - | - | - | - | - | - | - | - | - | - | - | - | - | 2(2) | 0.0137 | - | - | - | - | - | - | 2(2) | 0.02025 |
| **Drug metabolism - cytochrome P450** | - | - | - | - | - | - | - | - | - | - | 2(2) | 0.03134 | - | - | - | - | - | - | - | - | - | - | - | - | - | - | - | - | - | - | 2(2) | 0.01613 | - | - | - | - |
| **Vitamin D3 (cholecalciferol) metabolism** | - | - | - | - | - | - | - | - | - | - | - | - | - | - | - | - | - | - | - | - | - | - | - | - | - | - | - | - | 3(3) | 0.01639 | - | - | - | - | - | - |
| **Phosphatidylinositol phosphate metabolism** | - | - | - | - | - | - | - | - | 4(6) | 0.04663 | - | - | - | - | - | - | - | - | - | - | - | - | 4(6) | 0.0347 | - | - | - | - | 5(6) | 0.01664 | - | - | - | - | - | - |
| **Squalene and cholesterol biosynthesis** | - | - | - | - | - | - | - | - | - | - | - | - | - | - | - | - | - | - | 5(8) | 0.01723 | - | - | - | - | - | - | - | - | - | - | - | - | - | - | - | - |
| **Glycosphingolipid biosynthesis - ganglioseries** | - | - | - | - | - | - | - | - | - | - | - | - | - | - | - | - | - | - | - | - | - | - | - | - | - | - | 3(4) | 0.01773 | - | - | - | - | - | - | - | - |
| **Fatty acid oxidation** | - | - | - | - | - | - | - | - | 2(2) | 0.0179 | - | - | - | - | 2(2) | 0.01991 | - | - | 2(2) | 0.02067 | - | - | - | - | - | - | - | - | - | - | - | - | - | - | - | - |
| **Saturated fatty acids beta-oxidation** | - | - | - | - | 2(3) | 0.0463 | - | - | - | - | 2(3) | 0.04378 | - | - | 2(3) | 0.03949 | 3(3) | 0.01933 | - | - | - | - | - | - | - | - | 3(7) | 0.03571 | - | - | - | - | - | - | - | - |
| **Histidine metabolism** | - | - | - | - | - | - | - | - | - | - | 2(2) | 0.04277 | - | - | 2(2) | 0.01991 | - | - | - | - | - | - | - | - | 2(2) | 0.03227 | - | - | - | - | - | - | - | - | - | - |
| **Biopterin metabolism** | - | - | - | - | - | - | - | - | - | - | 2(2) | 0.02638 | - | - | 2(2) | 0.01991 | - | - | - | - | - | - | - | - | 2(2) | 0.02386 | - | - | - | - | - | - | - | - | - | - |
| **Di-unsaturated fatty acid beta-oxidation** | - | - | - | - | - | - | - | - | - | - | - | - | - | - | - | - | - | - | - | - | - | - | - | - | - | - | 2(3) | 0.02201 | - | - | - | - | - | - | - | - |
| **Vitamin B1 (thiamin) metabolism** | - | - | - | - | - | - | - | - | - | - | - | - | - | - | - | - | - | - | 1(1) | 0.04311 | 1(1) | 0.02252 | - | - | - | - | - | - | - | - | - | - | - | - | - | - |
| **Selenoamino acid metabolism** | - | - | - | - | - | - | 2(2) | 0.02344 | - | - | - | - | - | - | - | - | - | - | - | - | 1(1) | 0.02622 | - | - | - | - | - | - | - | - | - | - | - | - | - | - |
| **Drug metabolism - other enzymes** | - | - | - | - | - | - | - | - | - | - | - | - | - | - | - | - | - | - | - | - | - | - | - | - | - | - | 2(2) | 0.02487 | - | - | - | - | - | - | - | - |
| **Xenobiotics metabolism** | - | - | - | - | - | - | - | - | - | - | - | - | - | - | - | - | - | - | - | - | - | - | - | - | - | - | - | - | - | - | 2(2) | 0.02496 | - | - | 4(6) | 0.04924 |
| **Vitamin E metabolism** | - | - | - | - | - | - | - | - | - | - | - | - | - | - | - | - | - | - | - | - | - | - | - | - | - | - | - | - | - | - | - | - | - | - | 7(16) | 0.02512 |
| **Putative anti-Inflammatory metabolites formation from EPA** | - | - | - | - | - | - | - | - | - | - | - | - | - | - | - | - | - | - | - | - | 1(1) | 0.02622 | - | - | - | - | - | - | - | - | - | - | - | - | - | - |
| **Nitrogen metabolism** | - | - | - | - | - | - | - | - | - | - | - | - | - | - | - | - | - | - | 1(1) | 0.04311 | 1(1) | 0.02622 | - | - | - | - | - | - | - | - | - | - | - | - | - | - |
| **Propanoate metabolism** | - | - | - | - | - | - | - | - | - | - | - | - | - | - | - | - | - | - | - | - | 1(1) | 0.02622 | - | - | - | - | - | - | - | - | - | - | - | - | - | - |
| **Fatty acid oxidation, peroxisome** | - | - | - | - | - | - | - | - | - | - | - | - | - | - | - | - | - | - | - | - | 1(1) | 0.02622 | - | - | - | - | - | - | - | - | - | - | - | - | - | - |
| **Limonene and pinene degradation** | - | - | - | - | - | - | - | - | - | - | - | - | - | - | - | - | 5(6) | 0.02756 | - | - | - | - | - | - | - | - | - | - | - | - | - | - | 3(6) | 0.03991 | - | - |
| **Pyruvate Metabolism** | - | - | - | - | - | - | - | - | - | - | - | - | - | - | 3(5) | 3.18E-02 | - | - | 2(5) | 4.26E-02 | - | - | - | - | - | - | - | - | - | - | - | - | 3(5) | 3.66E-02 | - | - |
| **Polyunsaturated fatty acid biosynthesis** | - | - | - | - | - | - | - | - | - | - | - | - | - | - | - | - | - | - | - | - | - | - | - | - | - | - | - | - | - | - | 3(4) | 3.24E-02 | - | - | - | - |
| **Prostaglandin formation from arachidonate** | - | - | - | - | - | - | - | - | 4(9) | 0.04126 | - | - | - | - | - | - | - | - | - | - | - | - | - | - | - | - | - | - | - | - | - | - | - | - | - | - |
| **Glycosylphosphatidylinositol(GPI)-anchor biosynthesis** | - | - | - | - | - | - | - | - | - | - | - | - | - | - | - | - | - | - | 1(1) | 0.04311 | - | - | - | - | - | - | - | - | - | - | - | - | - | - | - | - |

**Supplementary Table 5**: **Comparative multivariate analysis for NHP serum proteomics profiling.**

| | | 6 Gy | | | | | | | | | | | |
|---|---|---|---|---|---|---|---|---|---|---|---|---|---|
| | | SD 1 vs. Pre | | | | SD 2 vs. Pre | | | | SD 6 vs. Pre | | | |
| **UniProtKB** | **Protein Name** | *p-value* | FDR | FC | LOG2FC | *p-value* | FDR | FC | LOG2FC | *p-value* | FDR | FC | LOG2FC |
| **H2R974** | Serpin family A member 9 | 3.91E-05 | 0.006769 | 3.1566 ↑ | 1.6584 | 4.43E-06 | 0.000766 | 3.5186 ↑ | 1.815 | 0.000604 | 0.048326 | 1.6025 ↑ | 0.6803 |
| **P02741** | C-reactive protein | 0.000198 | 0.017156 | 18.123 ↑ | 4.1798 | 0.005935 | 0.17114 | 4.882 ↑ | 2.2875 | 0.48412 | 0.81313 | 0.73299 ↓ | -0.448 |
| **B3KS79** | cDNA FLJ35730 fis, clone TESTI2003131, highly similar to ALPHA-1-/ | 0.000386 | 0.022245 | 3.8691 ↑ | 1.952 | 0.001893 | 0.10917 | 3.4693 ↑ | 1.7946 | 0.42073 | 0.79297 | 1.2718 ↑ | 0.3469 |
| **P07589** | Fibronectin | 0.001268 | 0.050824 | 0.47213 ↓ | -1.083 | 0.004685 | 0.16211 | 0.61826 ↓ | -0.694 | 0.035338 | 0.40081 | 0.65826 ↓ | -0.603 |
| **Q5VY30** | Retinol-binding protein | 0.001469 | 0.050824 | 0.49275 ↓ | -1.021 | 0.099628 | 0.52336 | 0.70986 ↓ | -0.494 | 0.93549 | 0.98232 | 0.98878 ↓ | -0.016 |
| **G1SQ27** | Clusterin | 0.001824 | 0.052591 | 0.52681 ↓ | -0.925 | 0.008026 | 0.18647 | 0.567 ↓ | -0.819 | 0.002477 | 0.10714 | 0.55567 ↓ | -0.848 |
| **P18659** | Apolipoprotein C-III | 0.002372 | 0.056028 | 0.35338 ↓ | -1.501 | 0.044346 | 0.48939 | 0.59721 ↓ | -0.744 | 0.000196 | 0.033825 | 0.3123 ↓ | -1.679 |
| **Q4R955** | Testis cDNA clone: QtsA-10685, similar to human I factor (complem | 0.002591 | 0.056028 | 0.73133 ↓ | -0.451 | 0.001676 | 0.10917 | 0.75527 ↓ | -0.405 | 0.48198 | 0.81313 | 0.94002 ↓ | -0.089 |
| **H2QIN8** | Protein C, inactivator of coagulation factors Va and VIIIa | 0.003363 | 0.06464 | 0.6422 ↓ | -0.639 | 0.008623 | 0.18647 | 0.70469 ↓ | -0.505 | 0.19446 | 0.62478 | 0.86277 ↓ | -0.213 |
| **K6ZY85** | Endonuclease domain containing 1 | 0.004505 | 0.077933 | 0.18513 ↓ | -2.433 | 0.77526 | 0.89413 | 0.89698 ↓ | -0.157 | 0.10916 | 0.48787 | 0.5194 ↓ | -0.945 |
| **I7G968** | Macaca fascicularis brain cDNA clone: QtrA-15720, similar to huma | 0.008314 | 0.13076 | 0.68542 ↓ | -0.545 | 0.18896 | 0.67042 | 0.84943 ↓ | -0.235 | 0.45266 | 0.79909 | 0.91747 ↓ | -0.124 |
| **G1U9R4** | Apolipoprotein B | 0.009855 | 0.14207 | 0.18658 ↓ | -2.422 | 0.1853 | 0.67042 | 0.59078 ↓ | -0.759 | 0.013289 | 0.34605 | 0.18642 ↓ | -2.423 |
| **P02750** | Leucine-rich alpha-2-glycoprotein | 0.011211 | 0.14919 | 2.0758 ↑ | 1.0537 | 0.684 | 0.86577 | 1.1186 ↑ | 0.1616 | 0.029431 | 0.40081 | 0.46162 ↓ | -1.115 |
| **A2NUT2** | Lambda-chain (AA -20 to 215) | 0.01645 | 0.18746 | 0.74527 ↓ | -0.424 | 0.078101 | 0.52336 | 0.80504 ↓ | -0.313 | 0.073114 | 0.42162 | 0.76108 ↓ | -0.394 |
| **G1PCZ3** | Inter-alpha-trypsin inhibitor heavy chain 1 | 0.01671 | 0.18746 | 0.33296 ↓ | -1.587 | 0.30057 | 0.70269 | 0.6285 ↓ | -0.67 | 0.28566 | 0.70599 | 0.64257 ↓ | -0.638 |
| **G1SJM1** | Apolipoprotein H | 0.017337 | 0.18746 | 0.7956 ↓ | -0.33 | 0.062878 | 0.518 | 0.86174 ↓ | -0.215 | 0.3654 | 0.7709 | 1.1295 ↑ | 0.1757 |
| **Q4R6T2** | Testis cDNA, clone: QtsA-17169, similar to human complement con | 0.019503 | 0.1912 | 0.79973 ↓ | -0.322 | 0.020272 | 0.35071 | 0.8168 ↓ | -0.292 | 0.30679 | 0.72704 | 0.88376 ↓ | -0.178 |
| **Q4R4P0** | Cathepsin D | 0.02089 | 0.1912 | 0.34728 ↓ | -1.526 | 0.074653 | 0.52336 | 0.54537 ↓ | -0.875 | 0.034556 | 0.40081 | 0.35535 ↓ | -1.493 |
| **H2QNX1** | Histidine rich glycoprotein | 0.020999 | 0.1912 | 1.8333 ↑ | 0.8745 | 0.77122 | 0.89413 | 1.2062 ↑ | 0.2705 | 0.10303 | 0.48542 | 1.7441 ↑ | 0.8025 |
| **A2RTY6** | Inter-alpha (Globulin) inhibitor H2 | 0.024264 | 0.20689 | 0.76482 ↓ | -0.387 | 0.082622 | 0.52336 | 0.75295 ↓ | -0.409 | 0.55611 | 0.81459 | 0.92729 ↓ | -0.109 |
| **P18428** | Lipopolysaccharide-binding protein | 0.025114 | 0.20689 | 4.4768 ↑ | 2.1625 | 0.30747 | 0.70574 | 1.9717 ↑ | 0.9794 | 0.30367 | 0.72704 | 1.6076 ↑ | 0.6849 |
| **G1NXF7** | Inter-alpha-trypsin inhibitor heavy chain 2 | 0.027672 | 0.2176 | 0.76387 ↓ | -0.389 | 0.14816 | 0.58252 | 0.8281 ↓ | -0.272 | 0.64758 | 0.86178 | 1.3174 ↑ | 0.3977 |
| **H2QPN7** | C-X-C motif chemokine | 0.034984 | 0.25105 | 0.59264 ↓ | -0.755 | 0.24466 | 0.67042 | 0.7393 ↓ | -0.436 | 0.066523 | 0.42162 | 0.60367 ↓ | -0.728 |
| **A0A2R9AGL7** | Hexosyltransferase | 0.036405 | 0.25105 | 0.38101 ↓ | -1.392 | 0.092762 | 0.52336 | 0.53524 ↓ | -0.902 | 0.15185 | 0.55893 | 0.68785 ↓ | -0.54 |
| **P02776** | Platelet factor 4 | 0.03651 | 0.25105 | 0.37012 ↓ | -1.434 | 0.14071 | 0.58252 | 0.50031 ↓ | -0.999 | 0.054177 | 0.4075 | 0.407 ↓ | -1.297 |
| **O14502** | Alpha-2-HS-glycoprotein | 0.03773 | 0.25105 | 0.5975 ↓ | -0.743 | 0.016003 | 0.30761 | 0.58569 ↓ | -0.772 | 0.99076 | 0.99076 | 0.96962 ↓ | -0.045 |
| **Q96PD5** | N-acetylmuramoyl-L-alanine amidase | 0.04422 | 0.28334 | 0.72031 ↓ | -0.473 | 0.074348 | 0.52336 | 0.74445 ↓ | -0.426 | 0.39397 | 0.78796 | 1.3536 ↑ | 0.4368 |
| **G1PA91** | Vitellogenin domain-containing protein | 0.049652 | 0.30678 | 0.69372 ↓ | -0.528 | 0.12714 | 0.57518 | 0.81499 ↓ | -0.295 | 0.028647 | 0.40081 | 0.71003 ↓ | -0.494 |
| **G1NVY1** | Apolipoprotein H | 0.05227 | 0.31182 | 0.56786 ↓ | -0.816 | 0.2619 | 0.67042 | 0.79848 ↓ | -0.325 | 0.043243 | 0.40081 | 0.49628 ↓ | -1.011 |
| **G1PYJ2** | C3-beta-c | 0.062666 | 0.3332 | 0.58153 ↓ | -0.782 | 0.023668 | 0.37224 | 0.53093 ↓ | -0.913 | 0.083861 | 0.45194 | 0.59777 ↓ | -0.742 |
| **H2QMY8** | Vitamin K-dependent protein S | 0.063384 | 0.3332 | 0.55909 ↓ | -0.839 | 0.47485 | 0.75366 | 0.83046 ↓ | -0.268 | 0.47865 | 0.81313 | 0.792 ↓ | -0.336 |
| **P11276** | Fibronectin | 0.063457 | 0.3332 | 0.66482 ↓ | -0.589 | 0.14185 | 0.58252 | 0.74272 ↓ | -0.429 | 0.021704 | 0.40081 | 0.51926 ↓ | -0.945 |
| **H2QU06** | Plasminogen | 0.063875 | 0.3332 | 0.70136 ↓ | -0.512 | 0.10704 | 0.52336 | 0.75428 ↓ | -0.407 | 0.10998 | 0.48787 | 0.76828 ↓ | -0.38 |
| **P03952** | Plasma kallikrein | 0.065485 | 0.3332 | 0.81212 ↓ | -0.3 | 0.004354 | 0.16211 | 0.77167 ↓ | -0.374 | 0.98753 | 0.99076 | 1.0334 ↑ | 0.0473 |
| **Q4R6W8** | Testis cDNA, clone: QtsA-16984, similar to human sex hormone-bin | 0.074084 | 0.35571 | 0.66909 ↓ | -0.58 | 0.035901 | 0.47776 | 0.4693 ↓ | -1.091 | 0.18949 | 0.62478 | 0.72537 ↓ | -0.463 |
| **P06868** | Plasminogen | 0.07427 | 0.35571 | 0.78077 ↓ | -0.357 | 0.030868 | 0.44502 | 0.72449 ↓ | -0.465 | 0.18878 | 0.62478 | 0.82885 ↓ | -0.271 |
| **G1SU82** | Gc-globulin | 0.076076 | 0.35571 | 0.73844 ↓ | -0.437 | 0.10685 | 0.52336 | 0.77378 ↓ | -0.37 | 0.56414 | 0.81459 | 0.92912 ↓ | -0.106 |
| **H2RC06** | Alpha-2-macroglobulin | 0.08469 | 0.37707 | 0.3929 ↓ | -1.348 | 0.24365 | 0.67042 | 0.50528 ↓ | -0.985 | 0.19502 | 0.62478 | 0.48325 ↓ | -1.049 |
| **Q8IZZ5** | Coagulation factor XII-Mie | 0.085147 | 0.37707 | 0.69304 ↓ | -0.529 | 0.050194 | 0.48939 | 0.69913 ↓ | -0.516 | 0.43706 | 0.79693 | 0.87536 ↓ | -0.192 |
| **H2QV12** | Alpha-2-glycoprotein 1, zinc-binding | 0.089659 | 0.37707 | 0.7259 ↓ | -0.462 | 0.19497 | 0.67042 | 0.77237 ↓ | -0.373 | 0.072398 | 0.42162 | 0.71707 ↓ | -0.48 |
| **A0A024R462** | Fibronectin | 0.091012 | 0.37707 | 0.54684 ↓ | -0.871 | 0.63459 | 0.84449 | 0.95283 ↓ | -0.07 | 0.28029 | 0.70599 | 0.7351 ↓ | -0.444 |
| **A0A2R9BXC9** | SERPIN domain-containing protein | 0.091544 | 0.37707 | 0.80353 ↓ | -0.316 | 0.1054 | 0.52336 | 0.81524 ↓ | -0.295 | 0.66982 | 0.87787 | 0.96399 ↓ | -0.053 |
| **Q59HE0** | Receptor protein-tyrosine kinase | 0.11078 | 0.44568 | 3.4854 ↑ | 1.8013 | 0.14155 | 0.58252 | 3.6148 ↑ | 1.8539 | 0.1696 | 0.59878 | 4.2242 ↑ | 2.0787 |
| **B4E1D8** | cDNA FLJ51597, highly similar to C4b-binding protein alpha chain | 0.11375 | 0.44723 | 0.64358 ↓ | -0.636 | 0.5552 | 0.79544 | 1.4203 ↑ | 0.5062 | 0.28549 | 0.70599 | 0.81323 ↓ | -0.298 |
| **A0A286XIX4** | Fibronectin | 0.13133 | 0.5049 | 0.40388 ↓ | -1.308 | 0.56618 | 0.79544 | 0.73873 ↓ | -0.437 | 0.91745 | 0.98232 | 1.0097 ↑ | 0.014 |
| **P13645** | Keratin, type I cytoskeletal 10 | 0.13851 | 0.52093 | 0.66934 ↓ | -0.579 | 0.25535 | 0.67042 | 0.73382 ↓ | -0.446 | 0.014002 | 0.34605 | 0.50159 ↓ | -0.995 |
| **P00735** | Prothrombin | 0.14515 | 0.53085 | 0.71582 ↓ | -0.482 | 0.099586 | 0.52336 | 0.70268 ↓ | -0.509 | 0.048653 | 0.40081 | 0.66772 ↓ | -0.583 |

| | | | | | | | | | | | | | |
|---|---|---|---|---|---|---|---|---|---|---|---|---|---|
| **H2QMF6** | C-type lectin domain family 3 member B | 0.14729 | 0.53085 | 0.66164 ↓ | -0.596 | 0.10555 | 0.52336 | 0.62889 ↓ | -0.669 | 0.33228 | 0.73697 | 0.75225 ↓ | -0.411 |
| **B4E1Z4** | C3/C5 convertase | 0.15448 | 0.53757 | 0.87334 ↓ | -0.195 | 0.37061 | 0.70688 | 0.90922 ↓ | -0.137 | 0.033205 | 0.40081 | 0.72587 ↓ | -0.462 |
| **H2Q321** | Hemopexin | 0.15682 | 0.53757 | 1.2766 ↑ | 0.3524 | 0.092245 | 0.52336 | 1.3798 ↑ | 0.4644 | 0.35681 | 0.7709 | 1.1901 ↑ | 0.2511 |
| **A0A2R9BCB8** | SERPIN domain-containing protein | 0.15847 | 0.53757 | 0.6391 ↓ | -0.646 | 0.049667 | 0.48939 | 0.78258 ↓ | -0.354 | 0.051796 | 0.4073 | 0.53259 ↓ | -0.909 |
| **Q14520** | Hyaluronan-binding protein 2 | 0.16897 | 0.56216 | 0.4749 ↓ | -1.074 | 0.6566 | 0.85271 | 0.89155 ↓ | -0.166 | 0.82929 | 0.95405 | 0.86277 ↓ | -0.213 |
| **A0A2R8ZZK5** | SERPIN domain-containing protein | 0.17363 | 0.56675 | 0.6983 ↓ | -0.518 | 0.28262 | 0.67042 | 0.75044 ↓ | -0.414 | 0.20473 | 0.64153 | 0.70481 ↓ | -0.505 |
| **P48770** | Complement component C9 | 0.1814 | 0.57142 | 0.82897 ↓ | -0.271 | 0.27689 | 0.67042 | 0.87473 ↓ | -0.193 | 0.046446 | 0.40081 | 0.75559 ↓ | -0.404 |
| **A0A2R9BYA2** | Uncharacterized protein | 0.18166 | 0.57142 | 1.3934 ↑ | 0.4786 | 0.93851 | 0.97809 | 0.98898 ↓ | -0.016 | 0.98773 | 0.99076 | 1.1085 ↑ | 0.1487 |
| **G1PHQ7** | Uncharacterized protein | 0.18683 | 0.57527 | 0.44 ↓ | -1.184 | 0.25523 | 0.67042 | 0.48438 ↓ | -1.046 | 0.7575 | 0.95325 | 0.78755 ↓ | -0.345 |
| **Q28508** | Haptoglobin | 0.18954 | 0.57527 | 1.7674 ↑ | 0.8217 | 0.65288 | 0.85271 | 1.4537 ↑ | 0.5397 | 0.03166 | 0.40081 | 1.7966 ↑ | 0.8453 |
| **H2QPM1** | Joining chain of multimeric IgA and IgM | 0.19733 | 0.586 | 0.38518 ↓ | -1.376 | 0.24167 | 0.67042 | 0.43202 ↓ | -1.211 | 0.23119 | 0.65566 | 0.44277 ↓ | -1.175 |
| **Q8IVC0** | Serpin peptidase inhibitor, clade D (Heparin cofactor), member 1 | 0.20393 | 0.586 | 0.65089 ↓ | -0.62 | 0.31894 | 0.70574 | 0.7187 ↓ | -0.477 | 0.32579 | 0.73697 | 0.72813 ↓ | -0.458 |
| **H0W536** | Complement C3 alpha chain | 0.2052 | 0.586 | 0.75409 ↓ | -0.407 | 0.2586 | 0.67042 | 0.78912 ↓ | -0.342 | 0.78233 | 0.95325 | 1.3091 ↑ | 0.3886 |
| **A0A2R9B5S5** | Uncharacterized protein | 0.20662 | 0.586 | 0.68841 ↓ | -0.539 | 0.57107 | 0.79544 | 0.90979 ↓ | -0.136 | 0.57102 | 0.81459 | 1.1623 ↑ | 0.217 |
| **P05160** | Coagulation factor XIII B chain | 0.21099 | 0.58872 | 0.64072 ↓ | -0.642 | 0.47324 | 0.75366 | 0.73938 ↓ | -0.436 | 0.579 | 0.81459 | 0.78792 ↓ | -0.344 |
| **A0A2R9BFI7** | C3-beta-c | 0.21943 | 0.60255 | 0.36602 ↓ | -1.45 | 0.19769 | 0.67042 | 0.35842 ↓ | -1.48 | 0.1156 | 0.49112 | 0.30635 ↓ | -1.707 |
| **H0UT45** | Uncharacterized protein | 0.22736 | 0.61028 | 0.53329 ↓ | -0.907 | 0.46915 | 0.75366 | 0.64028 ↓ | -0.643 | 0.42285 | 0.79297 | 0.72282 ↓ | -0.468 |
| **H2QCZ6** | Keratin 9 | 0.2293 | 0.61028 | 0.55948 ↓ | -0.838 | 0.15641 | 0.60131 | 0.51801 ↓ | -0.949 | 0.004046 | 0.13998 | 0.22221 ↓ | -2.17 |
| **Q8HXR9** | Soluble type II IL-1 receptor | 0.23349 | 0.61202 | 0.67697 ↓ | -0.563 | 0.34982 | 0.70688 | 0.7468 ↓ | -0.421 | 0.058618 | 0.42162 | 0.42005 ↓ | -1.251 |
| **V9HWD8** | Epididymis secretory sperm binding protein Li 163pA | 0.24604 | 0.62453 | 0.77414 ↓ | -0.369 | 0.32742 | 0.70574 | 0.79898 ↓ | -0.324 | 0.38905 | 0.78796 | 0.82432 ↓ | -0.279 |
| **P01031** | Complement C5 | 0.24921 | 0.62453 | 0.7115 ↓ | -0.491 | 0.34604 | 0.70688 | 0.75365 ↓ | -0.408 | 0.47415 | 0.81313 | 0.79835 ↓ | -0.325 |
| **V9HWA9** | C3-beta-c | 0.25229 | 0.62453 | 0.90578 ↓ | -0.143 | 0.050919 | 0.48939 | 0.84473 ↓ | -0.243 | 0.000838 | 0.048326 | 0.71484 ↓ | -0.484 |
| **A0A2R9A2B8** | C1q domain-containing protein | 0.2527 | 0.62453 | 1.3789 ↑ | 0.4635 | 0.66048 | 0.85271 | 0.57993 ↓ | -0.786 | 0.63198 | 0.84754 | 1.3395 ↑ | 0.4216 |
| **G1P5R5** | Gc-globulin | 0.27331 | 0.63544 | 0.89865 ↓ | -0.154 | 0.26804 | 0.67042 | 0.88548 ↓ | -0.175 | 0.24876 | 0.67651 | 0.85995 ↓ | -0.218 |
| **H0V4A3** | Apolipoprotein B | 0.27467 | 0.63544 | 0.69115 ↓ | -0.533 | 0.66587 | 0.8533 | 0.83811 ↓ | -0.255 | 0.078182 | 0.43631 | 0.50037 ↓ | -0.999 |
| **P48740** | Mannan-binding lectin serine protease 1 | 0.27684 | 0.63544 | 0.85218 ↓ | -0.231 | 0.12966 | 0.57518 | 0.79124 ↓ | -0.338 | 0.064316 | 0.42162 | 0.75739 ↓ | -0.401 |
| **A0A2J8QA94** | Complement component C6 | 0.28178 | 0.63544 | 0.83186 ↓ | -0.266 | 0.51375 | 0.77963 | 0.89885 ↓ | -0.154 | 0.48911 | 0.81361 | 1.1963 ↑ | 0.2585 |
| **A0A2R9AC29** | Clusterin | 0.2838 | 0.63544 | 0.83287 ↓ | -0.264 | 0.058043 | 0.50207 | 0.68146 ↓ | -0.553 | 0.84393 | 0.95405 | 1.0581 ↑ | 0.0815 |
| **P02768** | Albumin | 0.28861 | 0.63544 | 0.79635 ↓ | -0.329 | 0.44616 | 0.74218 | 0.85205 ↓ | -0.231 | 0.86152 | 0.95719 | 0.96614 ↓ | -0.05 |
| **H2QXT3** | Actin-depolymerizing factor | 0.29105 | 0.63544 | 0.6957 ↓ | -0.523 | 0.49118 | 0.7587 | 0.85923 ↓ | -0.219 | 0.091815 | 0.46718 | 0.54072 ↓ | -0.887 |
| **H2QCH3** | Vitronectin | 0.29289 | 0.63544 | 0.78823 ↓ | -0.343 | 0.8322 | 0.93746 | 0.96134 ↓ | -0.057 | 0.91831 | 0.98232 | 1.0472 ↑ | 0.0665 |
| **G1SS91** | C4a anaphylatoxin | 0.29294 | 0.63544 | 0.88785 ↓ | -0.172 | 0.054804 | 0.499 | 0.83531 ↓ | -0.26 | 0.56622 | 0.81459 | 1.1258 ↑ | 0.1709 |
| **Q9NP10** | Insulin-like growth factor I | 0.29727 | 0.63544 | 0.25402 ↓ | -1.977 | 0.10891 | 0.52336 | 0.17881 ↓ | -2.484 | 0.14676 | 0.55196 | 0.18002 ↓ | -2.474 |
| **Q04756** | Hepatocyte growth factor activator | 0.31089 | 0.63544 | 0.72782 ↓ | -0.458 | 0.49028 | 0.7587 | 0.79886 ↓ | -0.324 | 0.52809 | 0.81459 | 0.80672 ↓ | -0.31 |
| **V9HWE3** | Carbonic anhydrase | 0.31596 | 0.63544 | 1.9501 ↑ | 0.9635 | 0.59321 | 0.80807 | 1.0114 ↑ | 0.0163 | 0.36256 | 0.7709 | 1.612 ↑ | 0.6888 |
| **Q53G99** | Beta actin variant | 0.31639 | 0.63544 | 0.58904 ↓ | -0.764 | 0.69874 | 0.86577 | 0.82196 ↓ | -0.283 | 0.7617 | 0.95325 | 0.80042 ↓ | -0.321 |
| **A0A2I3RER0** | Alpha-1-microglobulin | 0.31743 | 0.63544 | 0.86234 ↓ | -0.214 | 0.26127 | 0.67042 | 0.84632 ↓ | -0.241 | 0.73396 | 0.94758 | 0.96687 ↓ | -0.049 |
| **H2QPM5** | Gc-globulin | 0.32668 | 0.63544 | 0.85119 ↓ | -0.232 | 0.4351 | 0.73713 | 0.87916 ↓ | -0.186 | 0.79755 | 0.95325 | 0.97152 ↓ | -0.042 |
| **H2QDM3** | Mannose receptor C type 2 | 0.32795 | 0.63544 | 2.0531 ↑ | 1.0378 | 0.70711 | 0.86759 | 1.0024 ↑ | 0.0035 | 0.040551 | 0.40081 | 2.753 ↑ | 1.461 |
| **I7GPF3** | Macaca fascicularis brain cDNA clone: QtrA-15786, similar to huma | 0.33093 | 0.63544 | 0.72891 ↓ | -0.456 | 0.26483 | 0.67042 | 0.68291 ↓ | -0.55 | 0.99065 | 0.99076 | 0.97481 ↓ | -0.037 |
| **D1MGQ2** | Alpha-1-globin | 0.33163 | 0.63544 | 1.7569 ↑ | 0.8131 | 0.62322 | 0.8358 | 1.3046 ↑ | 0.3836 | 0.79106 | 0.95325 | 1.6523 ↑ | 0.7245 |
| **Q5JVE7** | Coagulation factor X | 0.33454 | 0.63544 | 0.73404 ↓ | -0.446 | 0.55283 | 0.79544 | 0.81995 ↓ | -0.286 | 0.8802 | 0.9699 | 0.8915 ↓ | -0.166 |
| **H0VMS1** | Ceruloplasmin | 0.33491 | 0.63544 | 0.88076 ↓ | -0.183 | 0.043873 | 0.48939 | 0.70788 ↓ | -0.498 | 0.069485 | 0.42162 | 0.64062 ↓ | -0.642 |
| **Q9Y6R7** | IgGFc-binding protein | 0.33916 | 0.63544 | 0.75367 ↓ | -0.408 | 0.33043 | 0.70574 | 0.7518 ↓ | -0.412 | 0.28334 | 0.70599 | 0.72977 ↓ | -0.454 |
| **H2QBZ3** | Profilin | 0.34362 | 0.63544 | 0.45374 ↓ | -1.14 | 0.99727 | 0.99888 | 0.68939 ↓ | -0.537 | 0.3066 | 0.72704 | 0.43845 ↓ | -1.19 |
| **K7BH37** | Serpin family G member 1 | 0.3472 | 0.63544 | 0.84062 ↓ | -0.25 | 0.56658 | 0.79544 | 0.93011 ↓ | -0.105 | 0.069845 | 0.42162 | 0.63937 ↓ | -0.645 |
| **A0A024R962** | HCG40889, isoform CRA_b | 0.35022 | 0.63544 | 0.88598 ↓ | -0.175 | 0.99167 | 0.99888 | 0.9791 ↓ | -0.03 | 0.86313 | 0.95719 | 1.0187 ↑ | 0.0267 |
| **A0A2I3TF97** | C4a anaphylatoxin | 0.35138 | 0.63544 | 0.91377 ↓ | -0.13 | 0.83992 | 0.93746 | 1.0408 ↑ | 0.0576 | 0.25582 | 0.68087 | 1.2825 ↑ | 0.359 |
| **H0VJK2** | Plasminogen | 0.3536 | 0.63544 | 0.87867 ↓ | -0.187 | 0.22834 | 0.67042 | 0.82939 ↓ | -0.27 | 0.14261 | 0.54825 | 1.3893 ↑ | 0.4743 |
| **A0A2R9C7E2** | Protein S100 | 0.35629 | 0.63544 | 0.54054 ↓ | -0.888 | 0.99888 | 0.99888 | 2.422 ↑ | 1.2762 | 0.33227 | 0.73697 | 0.53707 ↓ | -0.897 |
| **H2QX94** | Reversion inducing cysteine rich protein with kazal motifs | 0.36124 | 0.63769 | 0.7652 ↓ | -0.386 | 0.14638 | 0.58252 | 0.45008 ↓ | -1.152 | 0.23846 | 0.66539 | 0.47977 ↓ | -1.06 |
| **A0A024R035** | Complement component C9 | 0.37641 | 0.64807 | 0.89478 ↓ | -0.16 | 0.7582 | 0.89413 | 0.96517 ↓ | -0.051 | 0.10382 | 0.48542 | 0.84361 ↓ | -0.245 |
| **V9HVZ4** | Glyceraldehyde-3-phosphate dehydrogenase | 0.38148 | 0.64807 | 1.0186 ↑ | 0.0266 | 0.39529 | 0.71234 | 3.4016 ↑ | 1.7662 | 0.61046 | 0.82564 | 0.99814 ↓ | -0.003 |

| | | | | | | | | | | | | | | | | |
|---|---|---|---|---|---|---|---|---|---|---|---|---|---|---|---|---|
| **B4DPQ0** | Complement subcomponent C1r | 0.38463 | 0.64807 | 0.91642 | ↓ | -0.126 | 0.10035 | 0.52336 | 0.77115 | ↓ | -0.375 | 0.025001 | 0.40081 | 0.69518 | ↓ | -0.525 |
| **V9HVY1** | Fibrinogen beta chain | 0.38546 | 0.64807 | 0.83065 | ↓ | -0.268 | 0.98431 | 0.99888 | 1.0157 | ↑ | 0.0225 | 0.13565 | 0.53336 | 7.3275 | ↑ | 2.8733 |
| **P11717** | Cation-independent mannose-6-phosphate receptor | 0.38809 | 0.64807 | 1.5973 | ↑ | 0.6757 | 0.61662 | 0.8334 | 0.92428 | ↓ | -0.114 | 0.8458 | 0.95405 | 1.007 | ↑ | 0.0101 |
| **Q9NZP8** | Complement C1r subcomponent-like protein | 0.39327 | 0.64807 | 0.87216 | ↓ | -0.197 | 0.52961 | 0.78514 | 0.93735 | ↓ | -0.093 | 0.12916 | 0.51964 | 0.77157 | ↓ | -0.374 |
| **P00738** | Haptoglobin | 0.39334 | 0.64807 | 1.3324 | ↑ | 0.414 | 0.36564 | 0.70688 | 1.8774 | ↑ | 0.9087 | 0.086209 | 0.45194 | 1.8532 | ↑ | 0.89 |
| **P18656** | Apolipoprotein A-II | 0.41185 | 0.67175 | 0.59201 | ↓ | -0.756 | 0.375 | 0.70688 | 0.49282 | ↓ | -1.021 | 0.40992 | 0.78796 | 0.6669 | ↓ | -0.584 |
| **B7Z8Q2** | Alpha-2-HS-glycoprotein | 0.41591 | 0.67175 | 0.62416 | ↓ | -0.68 | 0.21178 | 0.67042 | 0.54592 | ↓ | -0.873 | 0.51323 | 0.81459 | 0.69331 | ↓ | -0.528 |
| **P68292** | Apolipoprotein A-I | 0.42034 | 0.67175 | 0.46806 | ↓ | -1.095 | 0.34626 | 0.70688 | 0.44376 | ↓ | -1.172 | 0.77264 | 0.95325 | 1.0212 | ↑ | 0.0302 |
| **P68223** | Hemoglobin subunit beta | 0.43425 | 0.67175 | 2.8958 | ↑ | 1.534 | 0.94525 | 0.97921 | 1.2312 | ↑ | 0.3001 | 0.22835 | 0.65566 | 2.7058 | ↑ | 1.4361 |
| **H2QZH0** | Angiotensin-converting enzyme | 0.43452 | 0.67175 | 1.092 | ↑ | 0.1269 | 0.83639 | 0.93746 | 1.0367 | ↑ | 0.0519 | 0.56336 | 0.81459 | 1.1654 | ↑ | 0.2209 |
| **H2QY75** | Complement C8 gamma chain | 0.43481 | 0.67175 | 0.92656 | ↓ | -0.11 | 0.077239 | 0.52336 | 0.86436 | ↓ | -0.21 | 0.58858 | 0.81459 | 0.95832 | ↓ | -0.061 |
| **A0A2S0BDD1** | Antithrombin-III | 0.44015 | 0.67175 | 0.86596 | ↓ | -0.208 | 0.21863 | 0.67042 | 0.76446 | ↓ | -0.387 | 0.79473 | 0.95325 | 1.1584 | ↑ | 0.2122 |
| **A0A2R9CM19** | C1q domain-containing protein | 0.44114 | 0.67175 | 0.86887 | ↓ | -0.203 | 0.42938 | 0.73713 | 0.86732 | ↓ | -0.205 | 0.93184 | 0.98232 | 1.1041 | ↑ | 0.1429 |
| **V9HWJ7** | Epididymis secretory protein Li 37 | 0.44266 | 0.67175 | 1.6002 | ↑ | 0.6783 | 0.9961 | 0.99888 | 1.0062 | ↑ | 0.009 | 0.21508 | 0.64153 | 0.60889 | ↓ | -0.716 |
| **C9JEU5** | Fibrinogen gamma chain | 0.46242 | 0.69564 | 0.74044 | ↓ | -0.434 | 0.76564 | 0.89413 | 1.084 | ↑ | 0.1164 | 0.40791 | 0.78796 | 4.568 | ↑ | 2.1916 |
| **P34929** | Apolipoprotein C-I, basic form | 0.48358 | 0.7212 | 0.42102 | ↓ | -1.248 | 0.69314 | 0.86577 | 0.46766 | ↓ | -1.097 | 0.52062 | 0.81459 | 0.4642 | ↓ | -1.107 |
| **A0A2R9A8Q1** | Uncharacterized protein | 0.49207 | 0.72759 | 0.86264 | ↓ | -0.213 | 0.4337 | 0.73713 | 0.84833 | ↓ | -0.237 | 0.19223 | 0.62478 | 0.75442 | ↓ | -0.407 |
| **P0C0L5** | Complement C4-B | 0.49934 | 0.72792 | 1.1382 | ↑ | 0.1867 | 0.57934 | 0.79544 | 0.97226 | ↓ | -0.041 | 0.042776 | 0.40081 | 1.7047 | ↑ | 0.7695 |
| **A0A2R9CH07** | SERPIN domain-containing protein | 0.50805 | 0.72792 | 0.80916 | ↓ | -0.306 | 0.1223 | 0.57183 | 0.59161 | ↓ | -0.757 | 0.90304 | 0.98232 | 0.99093 | ↓ | -0.013 |
| **Q4R4R3** | Lumican | 0.51042 | 0.72792 | 0.7532 | ↓ | -0.409 | 0.41825 | 0.73713 | 0.70181 | ↓ | -0.511 | 0.53292 | 0.81459 | 0.75565 | ↓ | -0.404 |
| **K7DFV8** | Cadherin-13 | 0.51084 | 0.72792 | 0.46621 | ↓ | -1.101 | 0.56494 | 0.79544 | 0.53426 | ↓ | -0.904 | 0.54044 | 0.81459 | 2.0384 | ↑ | 1.0274 |
| **I7GH44** | Angiotensin 1-10 | 0.51333 | 0.72792 | 1.2256 | ↑ | 0.2935 | 0.92847 | 0.97809 | 1.0179 | ↑ | 0.0256 | 0.47223 | 0.81313 | 1.1769 | ↑ | 0.235 |
| **P35908** | Keratin, type II cytoskeletal 2 epidermal | 0.52253 | 0.73495 | 0.87483 | ↓ | -0.193 | 0.57883 | 0.79544 | 0.88884 | ↓ | -0.17 | 0.045192 | 0.40081 | 0.63105 | ↓ | -0.664 |
| **K7BA97** | Fibulin-1 | 0.53266 | 0.74314 | 0.82533 | ↓ | -0.277 | 0.32469 | 0.70574 | 0.69437 | ↓ | -0.526 | 0.90501 | 0.98232 | 1.0797 | ↑ | 0.1107 |
| **P00761** | Trypsin | 0.54734 | 0.75332 | 0.91669 | ↓ | -0.125 | 0.28289 | 0.67042 | 0.79471 | ↓ | -0.332 | 0.52164 | 0.81459 | 1.47 | ↑ | 0.5558 |
| **B2R4C5** | 1,4-beta-N-acetylmuramidase C | 0.55051 | 0.75332 | 1.3575 | ↑ | 0.4409 | 0.38817 | 0.70688 | 1.038 | ↑ | 0.0538 | 0.32816 | 0.73697 | 0.5485 | ↓ | -0.866 |
| **I7GA03** | Macaca fascicularis brain cDNA clone: QorA-13734, similar to huma | 0.55714 | 0.75332 | 0.74402 | ↓ | -0.427 | 0.6482 | 0.85271 | 0.75723 | ↓ | -0.401 | 0.2245 | 0.65566 | 0.58507 | ↓ | -0.773 |
| **A0A140VJI7** | Testicular tissue protein Li 61 | 0.55737 | 0.75332 | 0.92302 | ↓ | -0.116 | 0.32233 | 0.70574 | 0.84305 | ↓ | -0.246 | 0.34568 | 0.75699 | 0.85274 | ↓ | -0.23 |
| **A0A2R8ZJ60** | Uncharacterized protein | 0.56364 | 0.75589 | 0.86994 | ↓ | -0.201 | 0.38458 | 0.70688 | 0.82916 | ↓ | -0.27 | 0.11639 | 0.49112 | 0.71978 | ↓ | -0.474 |
| **A0A2R8ZJE1** | Vitellogenin domain-containing protein | 0.57243 | 0.76177 | 0.90811 | ↓ | -0.139 | 0.38807 | 0.70688 | 0.85954 | ↓ | -0.218 | 0.16376 | 0.59022 | 0.77471 | ↓ | -0.368 |
| **A0A2R8ZLB9** | Uncharacterized protein | 0.60709 | 0.79238 | 0.89248 | ↓ | -0.164 | 0.43739 | 0.73713 | 0.83558 | ↓ | -0.259 | 0.39089 | 0.78796 | 0.81581 | ↓ | -0.294 |
| **H2Q8V0** | Serpin family A member 10 | 0.60809 | 0.79238 | 0.57312 | ↓ | -0.803 | 0.38415 | 0.70688 | 0.55004 | ↓ | -0.862 | 0.61088 | 0.82564 | 0.60491 | ↓ | -0.725 |
| **A8K9M5** | Complement component 8 subunit beta | 0.60917 | 0.79238 | 1.3404 | ↑ | 0.4227 | 0.4549 | 0.7454 | 1.4747 | ↑ | 0.5605 | 0.32043 | 0.73697 | 1.9504 | ↑ | 0.9638 |
| **H2QJP2** | Collagen type VI alpha 3 chain | 0.61454 | 0.79339 | 1.4147 | ↑ | 0.5005 | 0.7363 | 0.89077 | 0.95546 | ↓ | -0.066 | 0.84509 | 0.95405 | 1.0881 | ↑ | 0.1218 |
| **H2R580** | Glutathione peroxidase | 0.63266 | 0.80141 | 0.84861 | ↓ | -0.237 | 0.69796 | 0.86577 | 0.83664 | ↓ | -0.257 | 0.92594 | 0.98232 | 1.0781 | ↑ | 0.1085 |
| **G1P212** | Mannan binding lectin serine peptidase 2 | 0.63462 | 0.80141 | 1.1617 | ↑ | 0.2162 | 0.26287 | 0.67042 | 0.73033 | ↓ | -0.453 | 0.80526 | 0.95405 | 1.0798 | ↑ | 0.1107 |
| **B2R4M6** | Protein S100 | 0.63464 | 0.80141 | 0.87903 | ↓ | -0.186 | 0.53095 | 0.78514 | 2.7395 | ↑ | 1.4539 | 0.76484 | 0.95325 | 1.1246 | ↑ | 0.1694 |
| **B4E1C2** | Bradykinin | 0.6548 | 0.82088 | 0.8994 | ↓ | -0.153 | 0.90052 | 0.96412 | 1.0618 | ↑ | 0.0865 | 0.25027 | 0.67651 | 0.71506 | ↓ | -0.484 |
| **H2R909** | Protein C receptor | 0.68652 | 0.84391 | 1.0885 | ↑ | 0.1223 | 0.96508 | 0.9938 | 0.94519 | ↓ | -0.081 | 0.44223 | 0.79693 | 1.1007 | ↑ | 0.1384 |
| **K7B5N7** | Protein-tyrosine-phosphatase | 0.68744 | 0.84391 | 0.9669 | ↓ | -0.049 | 0.21644 | 0.67042 | 0.86088 | ↓ | -0.216 | 0.12298 | 0.50656 | 0.76684 | ↓ | -0.383 |
| **A0A2R9BZ89** | Uncharacterized protein | 0.68781 | 0.84391 | 0.92988 | ↓ | -0.105 | 0.76125 | 0.89413 | 0.89466 | ↓ | -0.161 | 0.58671 | 0.81459 | 0.70826 | ↓ | -0.498 |
| **Q60FE6** | Filamin A | 0.69588 | 0.84725 | 0.83747 | ↓ | -0.256 | 0.37933 | 0.70688 | 0.58116 | ↓ | -0.783 | 0.39987 | 0.78796 | 0.55509 | ↓ | -0.849 |
| **H0VH18** | Inter-alpha-trypsin inhibitor heavy chain 2 | 0.70682 | 0.84725 | 0.87004 | ↓ | -0.201 | 0.43887 | 0.73713 | 0.81403 | ↓ | -0.297 | 0.78662 | 0.95325 | 0.92751 | ↓ | -0.109 |
| **P02671** | Fibrinogen alpha chain | 0.70864 | 0.84725 | 0.87166 | ↓ | -0.198 | 0.90282 | 0.96412 | 0.93979 | ↓ | -0.09 | 0.55276 | 0.81459 | 1.4769 | ↑ | 0.5626 |
| **A0A2I3T5B0** | Complement C1s | 0.71012 | 0.84725 | 0.97882 | ↓ | -0.031 | 0.48105 | 0.75655 | 0.95697 | ↓ | -0.063 | 0.94258 | 0.98232 | 1.0447 | ↑ | 0.0631 |
| **P33908** | Mannosyl-oligosaccharide 1,2-alpha-mannosidase IA | 0.72221 | 0.85475 | 0.97729 | ↓ | -0.033 | 0.50852 | 0.77852 | 0.91055 | ↓ | -0.135 | 0.65611 | 0.86646 | 1.0872 | ↑ | 0.1206 |
| **C0JYY2** | Apolipoprotein B (Including Ag(X) antigen) | 0.72629 | 0.85475 | 0.96628 | ↓ | -0.049 | 0.38096 | 0.70688 | 0.7814 | ↓ | -0.356 | 0.21148 | 0.64153 | 0.73897 | ↓ | -0.436 |
| **P07357** | Complement component C8 alpha chain | 0.74283 | 0.86831 | 1.0661 | ↑ | 0.0923 | 0.41286 | 0.73634 | 0.88096 | ↓ | -0.183 | 0.9731 | 0.99076 | 1.0263 | ↑ | 0.0375 |
| **B3KX75** | cDNA FLJ44930 fis, clone BRAMY3015549, highly similar to Neural c | 0.75883 | 0.88105 | 1.0158 | ↑ | 0.0226 | 0.9235 | 0.97809 | 0.93606 | ↓ | -0.095 | 0.42628 | 0.79297 | 0.70764 | ↓ | -0.499 |
| **V9HWK2** | Metavinculin | 0.77014 | 0.88758 | 0.9565 | ↓ | -0.064 | 0.31857 | 0.70574 | 1.22 | ↑ | 0.2869 | 0.40245 | 0.78796 | 1.3361 | ↑ | 0.418 |
| **B7ZKJ8** | ITIH4 protein | 0.77552 | 0.88758 | 1.4762 | ↑ | 0.5619 | 0.3688 | 0.70688 | 0.56885 | ↓ | -0.814 | 0.26959 | 0.70599 | 2.4919 | ↑ | 1.3172 |
| **P18658** | Apolipoprotein C-II | 0.77986 | 0.88758 | 0.41906 | ↓ | -1.255 | 0.89297 | 0.96412 | 0.42633 | ↓ | -1.23 | 0.60527 | 0.82564 | 0.70097 | ↓ | -0.513 |
| **P00734** | Prothrombin | 0.78497 | 0.88758 | 0.89811 | ↓ | -0.155 | 0.85818 | 0.94564 | 1.1045 | ↑ | 0.1434 | 0.84927 | 0.95405 | 0.89319 | ↓ | -0.163 |

| | | | | | | | | | | | | | |
|---|---|---|---|---|---|---|---|---|---|---|---|---|---|
| **A0A024R6I7** | Alpha-1-antitrypsin | 0.82257 | 0.91663 | 0.99479 ↓ | -0.008 | 0.87392 | 0.95087 | 1.3794 ↑ | 0.4641 | 0.44177 | 0.79693 | 1.8837 ↑ | 0.9135 |
| **D6R934** | Complement C1q subcomponent subunit B | 0.82702 | 0.91663 | 0.914 ↓ | -0.13 | 0.19116 | 0.67042 | 0.70632 ↓ | -0.502 | 0.93806 | 0.98232 | 0.97019 ↓ | -0.044 |
| **Q4LE41** | SEMA4D variant protein | 0.83037 | 0.91663 | 0.79602 ↓ | -0.329 | 0.7984 | 0.91472 | 0.81304 ↓ | -0.299 | 0.58664 | 0.81459 | 1.1386 ↑ | 0.1873 |
| **A5PL27** | CP protein | 0.83311 | 0.91663 | 1.0207 ↑ | 0.0296 | 0.71579 | 0.87206 | 1.0457 ↑ | 0.0644 | 0.38754 | 0.78796 | 2.7407 ↑ | 1.4545 |
| **K7C3E2** | Attractin | 0.83715 | 0.91663 | 0.93634 ↓ | -0.095 | 0.34417 | 0.70688 | 0.70239 ↓ | -0.51 | 0.84369 | 0.95405 | 1.1065 ↑ | 0.146 |
| **V9HW80** | 15S Mg(2+)-ATPase p97 subunit | 0.87881 | 0.95619 | 1.0201 ↑ | 0.0287 | 0.22775 | 0.67042 | 0.75579 ↓ | -0.404 | 0.57918 | 0.81459 | 1.359 ↑ | 0.4426 |
| **Q6IBD2** | CST6 protein | 0.89021 | 0.95918 | 1.4721 ↑ | 0.5579 | 0.17738 | 0.66711 | 1.4109 ↑ | 0.4966 | 0.82047 | 0.95405 | 1.5826 ↑ | 0.6623 |
| **B2R888** | Monocyte differentiation antigen CD14 | 0.89336 | 0.95918 | 0.9793 ↓ | -0.03 | 0.83618 | 0.93746 | 0.90898 ↓ | -0.138 | 0.78491 | 0.95325 | 1.0248 ↑ | 0.0353 |
| **A0A2R9BCI1** | Uncharacterized protein | 0.89819 | 0.95918 | 0.64802 ↓ | -0.626 | 0.76444 | 0.89413 | 0.68926 ↓ | -0.537 | 0.10082 | 0.48542 | 1.6351 ↑ | 0.7094 |
| **G1THZ6** | Immunoglobulin heavy constant mu | 0.91858 | 0.97463 | 1.1582 ↑ | 0.2119 | 0.24234 | 0.67042 | 1.5882 ↑ | 0.6674 | 0.83315 | 0.95405 | 1.4079 ↑ | 0.4935 |
| **H2Q4U2** | Apolipoprotein A4 | 0.92901 | 0.97463 | 1.3505 ↑ | 0.4335 | 0.70062 | 0.86577 | 1.1863 ↑ | 0.2465 | 0.58513 | 0.81459 | 1.066 ↑ | 0.0922 |
| **Q6UXM4** | Ficolin (Collagen/fibrinogen domain containing) 3 (Hakata antigen), | 0.93399 | 0.97463 | 0.62077 ↓ | -0.688 | 0.87028 | 0.95087 | 0.86467 ↓ | -0.21 | 0.68199 | 0.88709 | 0.91126 ↓ | -0.134 |
| **H2PYU7** | Receptor protein-tyrosine kinase | 0.93519 | 0.97463 | 1.1066 ↑ | 0.1461 | 0.26379 | 0.67042 | 0.76158 ↓ | -0.393 | 0.44974 | 0.79909 | 1.383 ↑ | 0.4679 |
| **H2QUJ6** | Insulin-like growth factor-binding protein 3 | 0.95946 | 0.98961 | 0.97242 ↓ | -0.04 | 0.85149 | 0.94428 | 0.92412 ↓ | -0.114 | 0.54573 | 0.81459 | 1.1023 ↑ | 0.1406 |
| **Q6N096** | Uncharacterized protein DKFZp686I15196 | 0.96101 | 0.98961 | 1.0685 ↑ | 0.0956 | 0.57212 | 0.79544 | 2.1237 ↑ | 1.0865 | 0.79897 | 0.95325 | 1.0058 ↑ | 0.0084 |
| **B2R8I2** | cDNA, FLJ93914, highly similar to Homo sapiens histidine-rich glyco | 0.96742 | 0.99031 | 0.94231 ↓ | -0.086 | 0.93712 | 0.97809 | 0.94388 ↓ | -0.083 | 0.50233 | 0.81459 | 0.77385 ↓ | -0.37 |
| **G1SIB7** | Coagulation factor XII | 0.98701 | 0.99865 | 0.80618 ↓ | -0.311 | 0.27235 | 0.67042 | 1.1372 ↑ | 0.1855 | 0.51277 | 0.81459 | 1.2646 ↑ | 0.3387 |
| **P31151** | Protein S100-A7 | 0.99282 | 0.99865 | 0.98209 ↓ | -0.026 | 0.75581 | 0.89413 | 5.6356 ↑ | 2.4946 | 0.21077 | 0.64153 | 0.4691 ↓ | -1.092 |
| **Q8TAY0** | Insulin-like growth factor binding protein, acid labile subunit | 0.99466 | 0.99865 | 0.96973 ↓ | -0.044 | 0.53099 | 0.78514 | 0.82626 ↓ | -0.275 | 0.97997 | 0.99076 | 0.98425 ↓ | -0.023 |
| **D3DNN4** | Carboxylic ester hydrolase | 0.99865 | 0.99865 | 0.9781 ↓ | -0.032 | 0.45672 | 0.7454 | 0.74631 ↓ | -0.422 | 0.97726 | 0.99076 | 0.98189 ↓ | -0.026 |

## 6.5 Gy

| | SD 1 vs. Pre | | | | SD 2 vs. Pre | | | | SD 6 vs. Pre | | | |
|---|---|---|---|---|---|---|---|---|---|---|---|---|
| **Protein Name** | ***p-value*** | **FDR** | **FC** | **LOG2FC** | ***p-value*** | **FDR** | **FC** | **LOG2FC** | ***p-value*** | **FDR** | **FC** | **LOG2FC** |
| Serpin family A member 9 | 5.69E-06 | 0.000984 | 4.955 ↑ | 2.3089 | 7.86E-06 | 0.001359 | 5.2917 ↑ | 2.4037 | 0.001011 | 0.065456 | 3.2216 ↑ | 1.6878 |
| C-reactive protein | 1.63E-05 | 0.001414 | 52.333 ↑ | 5.7096 | 0.003185 | 0.063892 | 10.763 ↑ | 3.428 | 0.89701 | 0.9661 | 1.282 ↑ | 0.3584 |
| cDNA FLJ35730 fis, clone TESTI2003131, highly similar to ALPHA-1-/ | 0.000296 | 0.017094 | 3.5422 ↑ | 1.8247 | 0.001978 | 0.048874 | 3.5711 ↑ | 1.8364 | 0.16062 | 0.66162 | 1.654 ↑ | 0.726 |
| Fibronectin | 0.008603 | 0.10104 | 0.46024 ↓ | -1.12 | 0.001111 | 0.048874 | 0.57148 ↓ | -0.807 | 0.026478 | 0.41982 | 0.53043 ↓ | -0.915 |
| Retinol-binding protein | 0.003316 | 0.070548 | 0.47909 ↓ | -1.062 | 0.003324 | 0.063892 | 0.57795 ↓ | -0.791 | 0.14973 | 0.65772 | 0.75983 ↓ | -0.396 |
| Clusterin | 0.23846 | 0.5219 | 0.76553 ↓ | -0.385 | 0.09669 | 0.35391 | 0.51062 ↓ | -0.97 | 0.080602 | 0.52081 | 0.53961 ↓ | -0.89 |
| Apolipoprotein C-III | 0.16673 | 0.43705 | 0.66193 ↓ | -0.595 | 0.13999 | 0.40364 | 0.65562 ↓ | -0.609 | 0.01409 | 0.40319 | 0.44934 ↓ | -1.154 |
| Testis cDNA clone: QtsA-10685, similar to human I factor (complem | 0.025142 | 0.16109 | 0.71456 ↓ | -0.485 | 0.009548 | 0.13764 | 0.72496 ↓ | -0.464 | 0.12871 | 0.63618 | 0.80586 ↓ | -0.311 |
| Protein C, inactivator of coagulation factors Va and VIIIa | 0.002977 | 0.070548 | 0.6237 ↓ | -0.681 | 0.00052 | 0.045015 | 0.59675 ↓ | -0.745 | 0.12691 | 0.63618 | 0.83057 ↓ | -0.268 |
| Endonuclease domain containing 1 | 0.13517 | 0.38973 | 4.314 ↑ | 2.109 | 0.20441 | 0.49806 | 3.2524 ↑ | 1.7015 | 0.17505 | 0.66464 | 3.5049 ↑ | 1.8094 |
| Macaca fascicularis brain cDNA clone: QtrA-15720, similar to huma | 0.28957 | 0.57581 | 0.77859 ↓ | -0.361 | 0.97385 | 0.97951 | 1.0051 ↑ | 0.0073 | 0.8722 | 0.9661 | 1.076 ↑ | 0.1056 |
| Apolipoprotein B | 0.37824 | 0.66096 | 0.68196 ↓ | -0.552 | 0.90922 | 0.94188 | 0.66878 ↓ | -0.58 | 0.48045 | 0.81205 | 0.98835 ↓ | -0.017 |
| Leucine-rich alpha-2-glycoprotein | 0.069059 | 0.3229 | 3.1734 ↑ | 1.666 | 0.18347 | 0.46677 | 3.047 ↑ | 1.6074 | 0.78455 | 0.94915 | 1.7201 ↑ | 0.7825 |
| Lambda-chain (AA -20 to 215) | 0.093645 | 0.33751 | 0.54187 ↓ | -0.884 | 0.05758 | 0.24904 | 0.68002 ↓ | -0.556 | 0.25646 | 0.70426 | 0.77704 ↓ | -0.364 |
| Inter-alpha-trypsin inhibitor heavy chain 1 | 0.73377 | 0.87909 | 1.2106 ↑ | 0.2757 | 0.51791 | 0.73314 | 0.98999 ↓ | -0.015 | 0.28636 | 0.75822 | 0.63664 ↓ | -0.651 |
| Apolipoprotein H | 0.009009 | 0.10104 | 0.62827 ↓ | -0.671 | 0.034212 | 0.21141 | 0.72638 ↓ | -0.461 | 0.35591 | 0.78693 | 0.85835 ↓ | -0.22 |
| Testis cDNA, clone: QtsA-17169, similar to human complement con | 0.00367 | 0.070548 | 0.68035 ↓ | -0.556 | 0.031295 | 0.21141 | 0.74368 ↓ | -0.427 | 0.43978 | 0.79819 | 0.92047 ↓ | -0.12 |
| Cathepsin D | 0.29761 | 0.58508 | 0.86101 ↓ | -0.216 | 0.36 | 0.64153 | 0.88976 ↓ | -0.169 | 0.77885 | 0.94915 | 1.029 ↑ | 0.0412 |
| Histidine rich glycoprotein | 0.051729 | 0.25569 | 0.52125 ↓ | -0.94 | 0.045212 | 0.24443 | 0.52367 ↓ | -0.933 | 0.15375 | 0.65772 | 0.69789 ↓ | -0.519 |
| Inter-alpha (Globulin) inhibitor H2 | 0.061776 | 0.29687 | 0.79224 ↓ | -0.336 | 0.016368 | 0.18878 | 0.67111 ↓ | -0.575 | 0.25212 | 0.70426 | 0.85416 ↓ | -0.227 |
| Lipopolysaccharide-binding protein | 0.55779 | 0.77826 | 2.15 ↑ | 1.1043 | 0.54668 | 0.73314 | 1.2371 ↑ | 0.307 | 0.77276 | 0.94915 | 1.14 ↑ | 0.189 |
| Inter-alpha-trypsin inhibitor heavy chain 2 | 0.702 | 0.87371 | 2.2182 ↑ | 1.1494 | 0.22581 | 0.52087 | 0.79193 ↓ | -0.337 | 0.72972 | 0.94915 | 1.2271 ↑ | 0.2953 |
| C-X-C motif chemokine | 0.011852 | 0.11391 | 0.60828 ↓ | -0.717 | 0.035439 | 0.21141 | 0.51918 ↓ | -0.946 | 0.016314 | 0.40319 | 0.4756 ↓ | -1.072 |
| Hexosyltransferase | 0.80803 | 0.90803 | 0.87758 ↓ | -0.188 | 0.39858 | 0.65598 | 0.44284 ↓ | -1.175 | 0.69706 | 0.94915 | 1.0204 ↑ | 0.0291 |
| Platelet factor 4 | 0.18603 | 0.47052 | 0.61066 ↓ | -0.712 | 0.10643 | 0.35936 | 0.54511 ↓ | -0.875 | 0.038437 | 0.41982 | 0.39518 ↓ | -1.339 |
| Alpha-2-HS-glycoprotein | 0.004831 | 0.083568 | 0.57709 ↓ | -0.793 | 0.006152 | 0.096756 | 0.54774 ↓ | -0.868 | 0.23491 | 0.70169 | 0.80442 ↓ | -0.314 |
| N-acetylmuramoyl-L-alanine amidase | 0.78739 | 0.89618 | 1.0158 ↑ | 0.0226 | 0.97026 | 0.97951 | 0.91983 ↓ | -0.121 | 0.72394 | 0.94915 | 1.1736 ↑ | 0.2309 |
| Vitellogenin domain-containing protein | 0.3317 | 0.61779 | 0.79474 ↓ | -0.331 | 0.26889 | 0.58194 | 0.77568 ↓ | -0.366 | 0.86159 | 0.9661 | 1.1558 ↑ | 0.2089 |
| Apolipoprotein H | 0.011064 | 0.1126 | 0.38545 ↓ | -1.375 | 0.089039 | 0.35009 | 0.54173 ↓ | -0.884 | 0.063886 | 0.44209 | 0.57802 ↓ | -0.791 |

| | | | | | | | | | | | | |
|---|---|---|---|---|---|---|---|---|---|---|---|---|
| C3-beta-c | 0.83774 | 0.91727 | 1.0772 ↑ | 0.1073 | 0.16672 | 0.45068 | 0.64687 ↓ | -0.628 | 0.83335 | 0.9661 | 0.86128 ↓ | -0.215 |
| Vitamin K-dependent protein S | 0.73729 | 0.87909 | 1.0812 ↑ | 0.1126 | 0.5404 | 0.73314 | 0.79577 ↓ | -0.33 | 0.9285 | 0.9661 | 1.0294 ↑ | 0.0419 |
| Fibronectin | 0.074707 | 0.32828 | 0.67596 ↓ | -0.565 | 0.11735 | 0.3604 | 0.63243 ↓ | -0.661 | 0.09729 | 0.58039 | 0.55793 ↓ | -0.842 |
| Plasminogen | 0.84856 | 0.92328 | 1.0018 ↑ | 0.0026 | 0.54205 | 0.73314 | 0.88157 ↓ | -0.182 | 0.96127 | 0.98403 | 0.97766 ↓ | -0.033 |
| Plasma kallikrein | 0.005476 | 0.086117 | 0.6737 ↓ | -0.57 | 0.001299 | 0.048874 | 0.71538 ↓ | -0.483 | 0.021969 | 0.41982 | 0.74623 ↓ | -0.422 |
| Testis cDNA, clone: QtsA-16984, similar to human sex hormone-bin | 0.78638 | 0.89618 | 1.0689 ↑ | 0.0961 | 0.23612 | 0.53051 | 0.75788 ↓ | -0.4 | 0.48928 | 0.81205 | 0.89227 ↓ | -0.164 |
| Plasminogen | 0.099286 | 0.34127 | 0.81107 ↓ | -0.302 | 0.032865 | 0.21141 | 0.76875 ↓ | -0.379 | 0.34042 | 0.78693 | 0.85185 ↓ | -0.231 |
| Gc-globulin | 0.16347 | 0.43705 | 0.75182 ↓ | -0.412 | 0.17324 | 0.45102 | 0.77271 ↓ | -0.372 | 0.32251 | 0.78693 | 0.82467 ↓ | -0.278 |
| Alpha-2-macroglobulin | 0.43678 | 0.67468 | 1.7739 ↑ | 0.827 | 0.47628 | 0.70425 | 0.62329 ↓ | -0.682 | 0.60249 | 0.91423 | 0.70541 ↓ | -0.503 |
| Coagulation factor XII-Mie | 0.56232 | 0.77826 | 0.9367 ↓ | -0.094 | 0.039307 | 0.22245 | 0.73364 ↓ | -0.447 | 0.43552 | 0.79819 | 0.91915 ↓ | -0.122 |
| Alpha-2-glycoprotein 1, zinc-binding | 0.017603 | 0.15198 | 0.59883 ↓ | -0.74 | 0.02774 | 0.21141 | 0.58469 ↓ | -0.774 | 0.24505 | 0.70426 | 0.77772 ↓ | -0.363 |
| Fibronectin | 0.08339 | 0.32828 | 0.60171 ↓ | -0.733 | 0.19808 | 0.48955 | 0.69554 ↓ | -0.524 | 0.47738 | 0.81205 | 0.94278 ↓ | -0.085 |
| SERPIN domain-containing protein | 0.036961 | 0.21007 | 0.83433 ↓ | -0.261 | 0.4737 | 0.70425 | 0.93726 ↓ | -0.093 | 0.18051 | 0.66464 | 1.1239 ↑ | 0.1685 |
| Receptor protein-tyrosine kinase | 0.91962 | 0.97167 | 0.8962 ↓ | -0.158 | 0.49105 | 0.71992 | 0.18766 ↓ | -2.414 | 0.3513 | 0.78693 | 0.99115 ↓ | -0.013 |
| cDNA FLJ51597, highly similar to C4b-binding protein alpha chain | 0.38831 | 0.67036 | 0.74874 ↓ | -0.417 | 0.736 | 0.87212 | 0.96901 ↓ | -0.045 | 0.4607 | 0.81205 | 1.095 ↑ | 0.1309 |
| Fibronectin | 0.049413 | 0.25143 | 0.20876 ↓ | -2.26 | 0.27708 | 0.58923 | 0.53252 ↓ | -0.909 | 0.06187 | 0.44209 | 0.23131 ↓ | -2.112 |
| Keratin, type I cytoskeletal 10 | 0.20762 | 0.48781 | 0.66336 ↓ | -0.592 | 0.12151 | 0.3604 | 0.58328 ↓ | -0.778 | 0.92243 | 0.9661 | 0.9545 ↓ | -0.067 |
| Prothrombin | 0.50549 | 0.73765 | 0.84062 ↓ | -0.25 | 0.61785 | 0.78322 | 0.8784 ↓ | -0.187 | 0.49286 | 0.81205 | 0.83576 ↓ | -0.259 |
| C-type lectin domain family 3 member B | 0.18766 | 0.47052 | 0.76904 ↓ | -0.379 | 0.027783 | 0.21141 | 0.67543 ↓ | -0.566 | 0.008614 | 0.37257 | 0.55287 ↓ | -0.855 |
| C3/C5 convertase | 0.13044 | 0.38249 | 1.2299 ↑ | 0.2986 | 0.58529 | 0.76708 | 0.93921 ↓ | -0.09 | 0.17454 | 0.66464 | 1.2722 ↑ | 0.3474 |
| Hemopexin | 0.24436 | 0.5219 | 1.215 ↑ | 0.281 | 0.019346 | 0.19687 | 1.4204 ↑ | 0.5063 | 0.96065 | 0.98403 | 1.1158 ↑ | 0.1581 |
| SERPIN domain-containing protein | 0.47774 | 0.71018 | 1.084 ↑ | 0.1164 | 0.46365 | 0.70425 | 0.72403 ↓ | -0.466 | 0.35809 | 0.78693 | 0.69911 ↓ | -0.516 |
| Hyaluronan-binding protein 2 | 0.98039 | 0.98609 | 1.0216 ↑ | 0.0308 | 0.4952 | 0.71992 | 1.0312 ↑ | 0.0444 | 0.82125 | 0.9661 | 0.90687 ↓ | -0.141 |
| SERPIN domain-containing protein | 0.85394 | 0.92332 | 0.92226 ↓ | -0.117 | 0.86415 | 0.92558 | 0.91847 ↓ | -0.123 | 0.43493 | 0.79819 | 1.1985 ↑ | 0.2612 |
| Complement component C9 | 0.75622 | 0.88997 | 1.0488 ↑ | 0.0687 | 0.87078 | 0.92558 | 1.039 ↑ | 0.0552 | 0.73635 | 0.94915 | 1.2032 ↑ | 0.2669 |
| Uncharacterized protein | 0.94279 | 0.97167 | 0.96749 ↓ | -0.048 | 0.11548 | 0.3604 | 1.5225 ↑ | 0.6064 | 0.14903 | 0.65772 | 1.4869 ↑ | 0.5723 |
| Uncharacterized protein | 0.24241 | 0.5219 | 1.7502 ↑ | 0.8075 | 0.831 | 0.9197 | 0.71847 ↓ | -0.477 | 0.72858 | 0.94915 | 0.96258 ↓ | -0.055 |
| Haptoglobin | 0.37116 | 0.65521 | 4.4804 ↑ | 2.1636 | 0.29732 | 0.61234 | 4.0076 ↑ | 2.0027 | 0.22031 | 0.69362 | 3.1034 ↑ | 1.6339 |
| Joining chain of multimeric IgA and IgM | 0.16574 | 0.43705 | 3.2732 ↑ | 1.7107 | 0.63524 | 0.79062 | 1.2832 ↑ | 0.3598 | 0.84662 | 0.9661 | 0.96189 ↓ | -0.056 |
| Serpin peptidase inhibitor, clade D (Heparin cofactor), member 1 | 0.36108 | 0.6507 | 0.68281 ↓ | -0.55 | 0.60196 | 0.77933 | 0.78842 ↓ | -0.343 | 0.54437 | 0.85615 | 0.79364 ↓ | -0.333 |
| Complement C3 alpha chain | 0.92372 | 0.97167 | 0.97452 ↓ | -0.037 | 0.87743 | 0.92558 | 0.94677 ↓ | -0.079 | 0.57705 | 0.88922 | 1.1772 ↑ | 0.2353 |
| Uncharacterized protein | 0.66567 | 0.85534 | 0.96716 ↓ | -0.048 | 0.8836 | 0.92645 | 1.0465 ↑ | 0.0656 | 0.74974 | 0.94915 | 1.1461 ↑ | 0.1968 |
| Coagulation factor XIII B chain | 0.97919 | 0.98609 | 0.96748 ↓ | -0.048 | 0.19696 | 0.48955 | 0.64489 ↓ | -0.633 | 0.73666 | 0.94915 | 0.8352 ↓ | -0.26 |
| C3-beta-c | 0.26902 | 0.55406 | 0.4447 ↓ | -1.169 | 0.11809 | 0.3604 | 0.38032 ↓ | -1.395 | 0.16457 | 0.66211 | 0.50966 ↓ | -0.972 |
| Uncharacterized protein | 0.15077 | 0.4157 | 1.8081 ↑ | 0.8545 | 0.53649 | 0.73314 | 0.89547 ↓ | -0.159 | 0.35913 | 0.78693 | 1.1252 ↑ | 0.1702 |
| Keratin 9 | 0.20974 | 0.48781 | 0.48166 ↓ | -1.054 | 0.056891 | 0.24904 | 0.34038 ↓ | -1.555 | 0.37747 | 0.78693 | 0.57355 ↓ | -0.802 |
| Soluble type II IL-1 receptor | 0.7224 | 0.87909 | 0.96369 ↓ | -0.053 | 0.17467 | 0.45102 | 0.71062 ↓ | -0.493 | 0.1124 | 0.62727 | 0.73932 ↓ | -0.436 |
| Epididymis secretory sperm binding protein Li 163pA | 0.081715 | 0.32828 | 0.66436 ↓ | -0.59 | 0.061639 | 0.26009 | 0.61641 ↓ | -0.698 | 0.5098 | 0.82949 | 0.84802 ↓ | -0.238 |
| Complement C5 | 0.12813 | 0.38219 | 1.3007 ↑ | 0.3793 | 0.40073 | 0.65598 | 1.0808 ↑ | 0.1121 | 0.31147 | 0.78094 | 1.2418 ↑ | 0.3124 |
| C3-beta-c | 0.24403 | 0.5219 | 0.85185 ↓ | -0.231 | 0.031242 | 0.21141 | 0.72901 ↓ | -0.456 | 0.23525 | 0.70169 | 0.85714 ↓ | -0.222 |
| C1q domain-containing protein | 0.95327 | 0.97167 | 1.5182 ↑ | 0.6024 | 0.025767 | 0.21141 | 0.074977 ↓ | -3.737 | 0.3777 | 0.78693 | 0.55081 ↓ | -0.86 |
| Gc-globulin | 0.000923 | 0.039927 | 0.6425 ↓ | -0.638 | 0.05696 | 0.24904 | 0.65513 ↓ | -0.61 | 0.055815 | 0.41982 | 0.71271 ↓ | -0.489 |
| Apolipoprotein B | 0.021866 | 0.15198 | 0.59937 ↓ | -0.738 | 0.098549 | 0.35391 | 0.65321 ↓ | -0.614 | 0.37727 | 0.78693 | 0.86849 ↓ | -0.203 |
| Mannan-binding lectin serine protease 1 | 0.41781 | 0.67329 | 0.86853 ↓ | -0.203 | 0.22078 | 0.52087 | 0.81809 ↓ | -0.29 | 0.30824 | 0.78094 | 0.83181 ↓ | -0.266 |
| Complement component C6 | 0.24812 | 0.52348 | 0.81173 ↓ | -0.301 | 0.38419 | 0.6453 | 0.8591 ↓ | -0.219 | 0.77423 | 0.94915 | 1.0954 ↑ | 0.1315 |
| Clusterin | 0.022114 | 0.15198 | 0.63777 ↓ | -0.649 | 0.04939 | 0.24778 | 0.66436 ↓ | -0.59 | 0.38209 | 0.78693 | 0.84671 ↓ | -0.24 |
| Albumin | 0.93771 | 0.97167 | 0.79263 ↓ | -0.335 | 0.4644 | 0.70425 | 10.065 ↑ | 3.3313 | 0.77129 | 0.94915 | 0.90427 ↓ | -0.145 |
| Actin-depolymerizing factor | 0.77754 | 0.89618 | 0.86982 ↓ | -0.201 | 0.75862 | 0.87568 | 1.0465 ↑ | 0.0656 | 0.97818 | 0.9917 | 0.77314 ↓ | -0.371 |
| Vitronectin | 0.41691 | 0.67329 | 0.81443 ↓ | -0.296 | 0.69226 | 0.84339 | 0.91429 ↓ | -0.129 | 0.98614 | 0.9917 | 0.96568 ↓ | -0.05 |
| C4a anaphylatoxin | 0.90927 | 0.97102 | 0.92429 ↓ | -0.114 | 0.36712 | 0.64153 | 0.75111 ↓ | -0.413 | 0.49142 | 0.81205 | 1.0749 ↑ | 0.1042 |
| Insulin-like growth factor I | 0.2739 | 0.55537 | 3.4386 ↑ | 1.7818 | 0.33634 | 0.63239 | 1.5062 ↑ | 0.5909 | 0.6752 | 0.94915 | 1.8434 ↑ | 0.8824 |
| Hepatocyte growth factor activator | 0.082328 | 0.32828 | 0.71511 ↓ | -0.484 | 0.01105 | 0.14705 | 0.6383 ↓ | -0.648 | 0.041517 | 0.41982 | 0.64491 ↓ | -0.633 |
| Carbonic anhydrase | 0.12321 | 0.38064 | 1.7275 ↑ | 0.7887 | 0.89886 | 0.93676 | 1.1851 ↑ | 0.2451 | 0.93114 | 0.9661 | 1.4764 ↑ | 0.5621 |

| | | | | | | | | | | | | |
|---|---|---|---|---|---|---|---|---|---|---|---|---|
| Beta actin variant | 0.81356 | 0.90803 | 1.0085 ↑ | 0.0122 | 0.47581 | 0.70425 | 1.1396 ↑ | 0.1885 | 0.74342 | 0.94915 | 0.99885 ↓ | -0.002 |
| Alpha-1-microglobulin | 0.47803 | 0.71018 | 0.93012 ↓ | -0.105 | 0.014219 | 0.17571 | 0.77731 ↓ | -0.363 | 0.75514 | 0.94915 | 1.1202 ↑ | 0.1638 |
| Gc-globulin | 0.022576 | 0.15198 | 0.62175 ↓ | -0.686 | 0.10802 | 0.35936 | 0.71034 ↓ | -0.493 | 0.2503 | 0.70426 | 0.78503 ↓ | -0.349 |
| Mannose receptor C type 2 | 0.54813 | 0.77726 | 1.77 ↑ | 0.8237 | 0.87582 | 0.92558 | 1.1864 ↑ | 0.2466 | 0.34913 | 0.78693 | 2.0798 ↑ | 1.0565 |
| Macaca fascicularis brain cDNA clone: QtrA-15786, similar to huma | 0.95482 | 0.97167 | 1.0069 ↑ | 0.0099 | 0.47357 | 0.70425 | 0.88048 ↓ | -0.184 | 0.69357 | 0.94915 | 0.99152 ↓ | -0.012 |
| Alpha-1-globin | 0.7387 | 0.87909 | 1.1149 ↑ | 0.157 | 0.83465 | 0.9197 | 0.40792 ↓ | -1.294 | 0.62934 | 0.92268 | 0.9791 ↓ | -0.03 |
| Coagulation factor X | 0.20963 | 0.48781 | 0.7613 ↓ | -0.393 | 0.60497 | 0.77933 | 1.1244 ↑ | 0.1691 | 0.88296 | 0.9661 | 1.004 ↑ | 0.0057 |
| Ceruloplasmin | 0.33779 | 0.62169 | 0.81506 ↓ | -0.295 | 0.048498 | 0.24778 | 0.57925 ↓ | -0.788 | 0.15588 | 0.65772 | 0.68249 ↓ | -0.551 |
| IgGFc-binding protein | 0.6359 | 0.84017 | 1.0085 ↑ | 0.0122 | 0.47525 | 0.70425 | 1.2518 ↑ | 0.324 | 0.39247 | 0.79819 | 1.1477 ↑ | 0.1988 |
| Profilin | 0.30771 | 0.59723 | 0.74478 ↓ | -0.425 | 0.3124 | 0.62843 | 0.91037 ↓ | -0.135 | 0.11179 | 0.62727 | 0.56818 ↓ | -0.816 |
| Serpin family G member 1 | 0.41882 | 0.67329 | 0.72182 ↓ | -0.47 | 0.10523 | 0.35936 | 0.56606 ↓ | -0.821 | 0.9326 | 0.9661 | 1.0109 ↑ | 0.0156 |
| HCG40889, isoform CRA_b | 0.77249 | 0.89618 | 0.89446 ↓ | -0.161 | 0.85924 | 0.92558 | 0.97709 ↓ | -0.033 | 0.33726 | 0.78693 | 0.77955 ↓ | -0.359 |
| C4a anaphylatoxin | 0.52263 | 0.74723 | 1.1529 ↑ | 0.2053 | 0.51517 | 0.73314 | 1.1675 ↑ | 0.2234 | 0.016227 | 0.40319 | 1.7669 ↑ | 0.8213 |
| Plasminogen | 0.40399 | 0.67329 | 0.92743 ↓ | -0.109 | 0.60815 | 0.77933 | 1.2663 ↑ | 0.3406 | 0.13847 | 0.64746 | 1.2982 ↑ | 0.3765 |
| Protein S100 | 0.47037 | 0.71018 | 0.87101 ↓ | -0.199 | 0.92172 | 0.94353 | 2.6902 ↑ | 1.4277 | 0.47824 | 0.81205 | 0.6272 ↓ | -0.673 |
| Reversion inducing cysteine rich protein with kazal motifs | 0.64887 | 0.84402 | 0.92066 ↓ | -0.119 | 0.38238 | 0.6453 | 0.63106 ↓ | -0.664 | 0.40164 | 0.79819 | 0.5221 ↓ | -0.938 |
| Complement component C9 | 0.1755 | 0.45315 | 1.3141 ↑ | 0.3941 | 0.2326 | 0.52946 | 1.1786 ↑ | 0.2371 | 0.18825 | 0.66464 | 1.3053 ↑ | 0.3844 |
| Glyceraldehyde-3-phosphate dehydrogenase | 0.63242 | 0.84017 | 0.39762 ↓ | -1.331 | 0.71181 | 0.85516 | 1.2954 ↑ | 0.3734 | 0.37874 | 0.78693 | 0.32348 ↓ | -1.628 |
| Complement subcomponent C1r | 0.022148 | 0.15198 | 0.75071 ↓ | -0.414 | 0.15965 | 0.44548 | 0.83681 ↓ | -0.257 | 0.04078 | 0.41982 | 0.668 ↓ | -0.582 |
| Fibrinogen beta chain | 0.039431 | 0.21007 | 0.69043 ↓ | -0.534 | 0.7695 | 0.87581 | 1.0467 ↑ | 0.0658 | 0.12384 | 0.63618 | 0.7621 ↓ | -0.392 |
| Cation-independent mannose-6-phosphate receptor | 0.56878 | 0.78094 | 0.80245 ↓ | -0.318 | 0.29072 | 0.60596 | 0.68365 ↓ | -0.549 | 0.71573 | 0.94915 | 1.0141 ↑ | 0.0202 |
| Complement C1r subcomponent-like protein | 0.022841 | 0.15198 | 0.4889 ↓ | -1.032 | 0.2691 | 0.58194 | 0.74298 ↓ | -0.429 | 0.081283 | 0.52081 | 0.64167 ↓ | -0.64 |
| Haptoglobin | 0.085964 | 0.32828 | 4.1373 ↑ | 2.0487 | 0.12069 | 0.3604 | 4.6575 ↑ | 2.2196 | 0.26234 | 0.70913 | 3.7586 ↑ | 1.9102 |
| Apolipoprotein A-II | 0.21438 | 0.48781 | 2.9549 ↑ | 1.5631 | 0.54501 | 0.73314 | 1.0764 ↑ | 0.1063 | 0.89156 | 0.9661 | 0.77474 ↓ | -0.368 |
| Alpha-2-HS-glycoprotein | 0.002817 | 0.070548 | 0.42228 ↓ | -1.244 | 0.001978 | 0.048874 | 0.43272 ↓ | -1.209 | 7.46E-05 | 0.012909 | 0.26981 ↓ | -1.89 |
| Apolipoprotein A-I | 0.10061 | 0.34127 | 3.7933 ↑ | 1.9235 | 0.91781 | 0.94353 | 1.0012 ↑ | 0.0018 | 0.89014 | 0.9661 | 0.76037 ↓ | -0.395 |
| Hemoglobin subunit beta | 0.015336 | 0.13964 | 5.196 ↑ | 2.3774 | 0.11043 | 0.3604 | 2.4455 ↑ | 1.2901 | 0.13779 | 0.64746 | 2.4777 ↑ | 1.309 |
| Angiotensin-converting enzyme | 0.3107 | 0.59723 | 0.67106 ↓ | -0.575 | 0.5265 | 0.73314 | 0.78179 ↓ | -0.355 | 0.52196 | 0.82949 | 0.79075 ↓ | -0.339 |
| Complement C8 gamma chain | 0.57766 | 0.78689 | 0.87911 ↓ | -0.186 | 0.33048 | 0.63239 | 0.82091 ↓ | -0.285 | 0.98673 | 0.9917 | 0.97992 ↓ | -0.029 |
| Antithrombin-III | 0.78719 | 0.89618 | 0.86081 ↓ | -0.216 | 0.40193 | 0.65598 | 1.1659 ↑ | 0.2214 | 0.87578 | 0.9661 | 1.1476 ↑ | 0.1987 |
| C1q domain-containing protein | 0.6895 | 0.86438 | 1.2 ↑ | 0.263 | 0.17446 | 0.45102 | 0.7982 ↓ | -0.325 | 0.64969 | 0.9387 | 1.0875 ↑ | 0.121 |
| Epididymis secretory protein Li 37 | 0.006839 | 0.09101 | 6.5535 ↑ | 2.7123 | 0.017687 | 0.19124 | 4.8207 ↑ | 2.2692 | 0.42556 | 0.79819 | 2.0703 ↑ | 1.0498 |
| Fibrinogen gamma chain | 0.27608 | 0.55537 | 1.4319 ↑ | 0.5179 | 0.42571 | 0.68193 | 1.3506 ↑ | 0.4336 | 0.04875 | 0.41982 | 1.9352 ↑ | 0.9525 |
| Apolipoprotein C-I, basic form | 0.64857 | 0.84402 | 0.83183 ↓ | -0.266 | 0.16403 | 0.45043 | 0.55287 ↓ | -0.855 | 0.36382 | 0.78693 | 0.70665 ↓ | -0.501 |
| Uncharacterized protein | 0.21505 | 0.48781 | 1.1278 ↑ | 0.1735 | 0.76433 | 0.87568 | 0.97684 ↓ | -0.034 | 0.18335 | 0.66464 | 1.2315 ↑ | 0.3004 |
| Complement C4-B | 0.88049 | 0.94611 | 0.97618 ↓ | -0.035 | 0.86205 | 0.92558 | 0.97448 ↓ | -0.037 | 0.21084 | 0.69362 | 1.3315 ↑ | 0.4131 |
| SERPIN domain-containing protein | 0.81015 | 0.90803 | 0.8456 ↓ | -0.242 | 0.44632 | 0.70425 | 0.72605 ↓ | -0.462 | 0.43057 | 0.79819 | 0.74804 ↓ | -0.419 |
| Lumican | 0.038431 | 0.21007 | 0.51112 ↓ | -0.968 | 0.030583 | 0.21141 | 0.49474 ↓ | -1.015 | 0.049238 | 0.41982 | 0.52129 ↓ | -0.94 |
| Cadherin-13 | 0.41577 | 0.67329 | 0.61237 ↓ | -0.708 | 0.58438 | 0.76708 | 0.72694 ↓ | -0.46 | 0.34713 | 0.78693 | 1.3666 ↑ | 0.4506 |
| Angiotensin 1-10 | 0.006171 | 0.088961 | 1.571 ↑ | 0.6517 | 0.12291 | 0.3604 | 1.2178 ↑ | 0.2843 | 0.051445 | 0.41982 | 2.0049 ↑ | 1.0035 |
| Keratin, type II cytoskeletal 2 epidermal | 0.021199 | 0.15198 | 0.52694 ↓ | -0.924 | 0.0344 | 0.21141 | 0.55523 ↓ | -0.849 | 0.8244 | 0.9661 | 0.92158 ↓ | -0.118 |
| Fibulin-1 | 0.82298 | 0.91098 | 0.90553 ↓ | -0.143 | 0.36441 | 0.64153 | 0.74093 ↓ | -0.433 | 0.42847 | 0.79819 | 0.76583 ↓ | -0.385 |
| Trypsin | 0.42032 | 0.67329 | 1.5427 ↑ | 0.6255 | 0.78599 | 0.88011 | 0.97717 ↓ | -0.033 | 0.92346 | 0.9661 | 1.1385 ↑ | 0.1872 |
| 1,4-beta-N-acetylmuramidase C | 0.66902 | 0.85534 | 0.86344 ↓ | -0.212 | 0.70658 | 0.85481 | 5.9498 ↑ | 2.5728 | 0.12641 | 0.63618 | 2.728 ↑ | 1.4479 |
| Macaca fascicularis brain cDNA clone: QorA-13734, similar to huma | 0.15138 | 0.4157 | 1.3753 ↑ | 0.4598 | 0.27929 | 0.58923 | 1.2177 ↑ | 0.2842 | 0.43267 | 0.79819 | 1.1733 ↑ | 0.2306 |
| Testicular tissue protein Li 61 | 0.5074 | 0.73765 | 0.80243 ↓ | -0.318 | 0.67621 | 0.82967 | 0.85446 ↓ | -0.227 | 0.30576 | 0.78094 | 0.72495 ↓ | -0.464 |
| Uncharacterized protein | 0.4803 | 0.71018 | 1.16 ↑ | 0.2142 | 0.84712 | 0.92558 | 0.91142 ↓ | -0.134 | 0.92384 | 0.9661 | 0.98527 ↓ | -0.021 |
| Vitellogenin domain-containing protein | 0.47607 | 0.71018 | 0.90373 ↓ | -0.146 | 0.22519 | 0.52087 | 0.83613 ↓ | -0.258 | 0.83825 | 0.9661 | 1.257 ↑ | 0.33 |
| Uncharacterized protein | 0.040071 | 0.21007 | 0.58841 ↓ | -0.765 | 0.039861 | 0.22245 | 0.57418 ↓ | -0.8 | 0.055117 | 0.41982 | 0.60455 ↓ | -0.726 |
| Serpin family A member 10 | 0.12581 | 0.38186 | 0.266 ↓ | -1.911 | 0.58214 | 0.76708 | 0.73591 ↓ | -0.442 | 0.81602 | 0.9661 | 1.0759 ↑ | 0.1055 |
| Complement component 8 subunit beta | 0.11655 | 0.37984 | 0.21641 ↓ | -2.208 | 0.056822 | 0.24904 | 0.17668 ↓ | -2.501 | 0.22453 | 0.69362 | 0.29071 ↓ | -1.782 |
| Collagen type VI alpha 3 chain | 0.087287 | 0.32828 | 0.62227 ↓ | -0.684 | 0.069764 | 0.28068 | 0.5278 ↓ | -0.922 | 0.22104 | 0.69362 | 0.75006 ↓ | -0.415 |
| Glutathione peroxidase | 0.096224 | 0.33973 | 1.5286 ↑ | 0.6122 | 0.45847 | 0.70425 | 1.1275 ↑ | 0.1731 | 0.86902 | 0.9661 | 1.0215 ↑ | 0.0307 |

| | | | | | | | | | | | | |
|---|---|---|---|---|---|---|---|---|---|---|---|---|
| Mannan binding lectin serine peptidase 2 | 0.11856 | 0.37984 | 0.63476 ↓ | -0.656 | 0.052447 | 0.24904 | 0.55671 ↓ | -0.845 | 0.097261 | 0.58039 | 0.60706 ↓ | -0.72 |
| Protein S100 | 0.36492 | 0.65083 | 6.1658 ↑ | 2.6243 | 0.20883 | 0.50178 | 5.0187 ↑ | 2.3273 | 0.035888 | 0.41982 | 3.1131 ↑ | 1.6384 |
| Bradykinin | 0.028273 | 0.17469 | 0.7168 ↓ | -0.48 | 0.35348 | 0.64153 | 0.88942 ↓ | -0.169 | 0.20227 | 0.69362 | 0.76458 ↓ | -0.387 |
| Protein C receptor | 0.33211 | 0.61779 | 0.78615 ↓ | -0.347 | 0.36063 | 0.64153 | 0.79776 ↓ | -0.326 | 0.39973 | 0.79819 | 0.81282 ↓ | -0.299 |
| Protein-tyrosine-phosphatase | 0.21503 | 0.48781 | 0.67401 ↓ | -0.569 | 0.3014 | 0.61344 | 0.75042 ↓ | -0.414 | 0.68305 | 0.94915 | 1.061 ↑ | 0.0854 |
| Uncharacterized protein | 0.090646 | 0.33365 | 0.7812 ↓ | -0.356 | 0.31655 | 0.62945 | 0.82076 ↓ | -0.285 | 0.77916 | 0.94915 | 1.0502 ↑ | 0.0707 |
| Filamin A | 0.43446 | 0.67468 | 1.575 ↑ | 0.6553 | 0.35759 | 0.64153 | 0.67001 ↓ | -0.578 | 0.85967 | 0.9661 | 1.1752 ↑ | 0.2329 |
| Inter-alpha-trypsin inhibitor heavy chain 2 | 0.74189 | 0.87909 | 0.9085 ↓ | -0.138 | 0.78854 | 0.88011 | 0.7393 ↓ | -0.436 | 0.65407 | 0.9387 | 0.87795 ↓ | -0.188 |
| Fibrinogen alpha chain | 0.67241 | 0.85534 | 0.90005 ↓ | -0.152 | 0.94482 | 0.96149 | 1.1867 ↑ | 0.247 | 0.61669 | 0.91423 | 0.84983 ↓ | -0.235 |
| Complement C1s | 0.009345 | 0.10104 | 0.73243 ↓ | -0.449 | 0.10024 | 0.35391 | 0.84078 ↓ | -0.25 | 0.28926 | 0.75822 | 0.86461 ↓ | -0.21 |
| Mannosyl-oligosaccharide 1,2-alpha-mannosidase IA | 0.11504 | 0.37984 | 0.52116 ↓ | -0.94 | 0.73551 | 0.87212 | 1.0442 ↑ | 0.0624 | 0.44293 | 0.79819 | 0.92962 ↓ | -0.105 |
| Apolipoprotein B (Including Ag(X) antigen) | 0.73677 | 0.87909 | 1.2547 ↑ | 0.3273 | 0.020927 | 0.20114 | 0.50528 ↓ | -0.985 | 0.22009 | 0.69362 | 0.73693 ↓ | -0.44 |
| Complement component C8 alpha chain | 0.55839 | 0.77826 | 1.0324 ↑ | 0.046 | 0.78138 | 0.88011 | 0.95848 ↓ | -0.061 | 0.61829 | 0.91423 | 1.04 ↑ | 0.0566 |
| cDNA FLJ44930 fis, clone BRAMY3015549, highly similar to Neural c | 0.12104 | 0.38064 | 0.51404 ↓ | -0.96 | 0.050128 | 0.24778 | 0.38212 ↓ | -1.388 | 0.02553 | 0.41982 | 0.48066 ↓ | -1.057 |
| Metavinculin | 0.67879 | 0.85715 | 0.88409 ↓ | -0.178 | 0.74799 | 0.87568 | 0.94311 ↓ | -0.085 | 0.72636 | 0.94915 | 1.5503 ↑ | 0.6326 |
| ITIH4 protein | 0.71145 | 0.87909 | 0.96484 ↓ | -0.052 | 0.7637 | 0.87568 | 1.3983 ↑ | 0.4837 | 0.18584 | 0.66464 | 1.9672 ↑ | 0.9761 |
| Apolipoprotein C-II | 0.42803 | 0.67468 | 0.51797 ↓ | -0.949 | 0.62268 | 0.78322 | 1.16 ↑ | 0.2141 | 0.61605 | 0.91423 | 0.66487 ↓ | -0.589 |
| Prothrombin | 0.58792 | 0.79462 | 1.0543 ↑ | 0.0762 | 0.66519 | 0.82198 | 0.6731 ↓ | -0.571 | 0.41151 | 0.79819 | 1.1039 ↑ | 0.1426 |
| Alpha-1-antitrypsin | 0.029505 | 0.17601 | 2.5014 ↑ | 1.3228 | 0.14586 | 0.41367 | 2.5151 ↑ | 1.3306 | 0.22217 | 0.69362 | 2.7069 ↑ | 1.4366 |
| Complement C1q subcomponent subunit B | 0.078677 | 0.32828 | 0.56193 ↓ | -0.832 | 0.51057 | 0.73314 | 0.81727 ↓ | -0.291 | 0.055365 | 0.41982 | 0.57389 ↓ | -0.801 |
| SEMA4D variant protein | 0.13971 | 0.39624 | 2.3135 ↑ | 1.2101 | 0.62476 | 0.78322 | 1.4696 ↑ | 0.5554 | 0.56781 | 0.88496 | 1.3249 ↑ | 0.4059 |
| CP protein | 0.39137 | 0.67036 | 1.103 ↑ | 0.1415 | 0.53366 | 0.73314 | 0.98406 ↓ | -0.023 | 0.70078 | 0.94915 | 1.0917 ↑ | 0.1266 |
| Attractin | 0.26004 | 0.542 | 0.70436 ↓ | -0.506 | 0.09453 | 0.35391 | 0.5908 ↓ | -0.759 | 0.25409 | 0.70426 | 0.72012 ↓ | -0.474 |
| 15S Mg(2+)-ATPase p97 subunit | 0.086031 | 0.32828 | 0.58785 ↓ | -0.766 | 0.003916 | 0.067745 | 0.34844 ↓ | -1.521 | 0.52263 | 0.82949 | 1.017 ↑ | 0.0243 |
| CST6 protein | 0.34367 | 0.62583 | 1.4989 ↑ | 0.5839 | 0.38032 | 0.6453 | 1.4554 ↑ | 0.5414 | 0.049908 | 0.41982 | 2.0621 ↑ | 1.0441 |
| Monocyte differentiation antigen CD14 | 0.82673 | 0.91098 | 0.96524 ↓ | -0.051 | 0.32924 | 0.63239 | 0.71352 ↓ | -0.487 | 0.65654 | 0.9387 | 1.0639 ↑ | 0.0894 |
| Uncharacterized protein | 0.99604 | 0.99604 | 0.86409 ↓ | -0.211 | 0.33996 | 0.63239 | 0.65799 ↓ | -0.604 | 0.85901 | 0.9661 | 0.80188 ↓ | -0.319 |
| Immunoglobulin heavy constant mu | 0.93759 | 0.97167 | 1.0134 ↑ | 0.0192 | 0.33872 | 0.63239 | 0.70614 ↓ | -0.502 | 0.51998 | 0.82949 | 0.59814 ↓ | -0.741 |
| Apolipoprotein A4 | 0.32036 | 0.60904 | 1.6065 ↑ | 0.6839 | 0.028331 | 0.21141 | 1.8439 ↑ | 0.8828 | 0.045644 | 0.41982 | 0.4798 ↓ | -1.06 |
| Ficolin (Collagen/fibrinogen domain containing) 3 (Hakata antigen), | 0.6362 | 0.84017 | 1.1693 ↑ | 0.2256 | 0.097474 | 0.35391 | 0.34124 ↓ | -1.551 | 0.46004 | 0.81205 | 0.72898 ↓ | -0.456 |
| Receptor protein-tyrosine kinase | 0.51907 | 0.74723 | 1.0594 ↑ | 0.0832 | 0.75228 | 0.87568 | 0.87366 ↓ | -0.195 | 0.8056 | 0.9661 | 1.0028 ↑ | 0.0041 |
| Insulin-like growth factor-binding protein 3 | 0.95245 | 0.97167 | 1.0296 ↑ | 0.0421 | 0.41574 | 0.67218 | 0.88226 ↓ | -0.181 | 0.9917 | 0.9917 | 1.0384 ↑ | 0.0543 |
| Uncharacterized protein DKFZp686I15196 | 0.42936 | 0.67468 | 0.53195 ↓ | -0.911 | 0.99218 | 0.99218 | 4.9698 ↑ | 2.3132 | 0.91336 | 0.9661 | 0.73242 ↓ | -0.449 |
| cDNA, FLJ93914, highly similar to Homo sapiens histidine-rich glyco | 0.001358 | 0.046997 | 0.4973 ↓ | -1.008 | 0.001749 | 0.048874 | 0.51569 ↓ | -0.955 | 0.001135 | 0.065456 | 0.5962 ↓ | -0.746 |
| Coagulation factor XII | 0.084494 | 0.32828 | 3.3346 ↑ | 1.7375 | 0.37428 | 0.6453 | 1.5992 ↑ | 0.6774 | 0.58082 | 0.88922 | 1.0861 ↑ | 0.1191 |
| Protein S100-A7 | 0.40077 | 0.67329 | 0.78153 ↓ | -0.356 | 0.26773 | 0.58194 | 0.66309 ↓ | -0.593 | 0.039528 | 0.41982 | 0.33418 ↓ | -1.581 |
| Insulin-like growth factor binding protein, acid labile subunit | 0.21712 | 0.48781 | 0.81299 ↓ | -0.299 | 0.33622 | 0.63239 | 0.84063 ↓ | -0.25 | 0.75302 | 0.94915 | 0.95961 ↓ | -0.059 |
| Carboxylic ester hydrolase | 0.20337 | 0.48781 | 0.73497 ↓ | -0.444 | 0.066387 | 0.27345 | 0.62288 ↓ | -0.683 | 0.46905 | 0.81205 | 0.85758 ↓ | -0.222 |

## 7 Gy

| | SD 1 vs. Pre | | | | SD 2 vs. Pre | | | | SD 6 vs. Pre | | | |
|---|---|---|---|---|---|---|---|---|---|---|---|---|
| **Protein Name** | ***p-value*** | **FDR** | **FC** | **LOG2FC** | ***p-value*** | **FDR** | **FC** | **LOG2FC** | ***p-value*** | **FDR** | **FC** | **LOG2FC** |
| Serpin family A member 9 | 1.58E-05 | 0.00137 | 4.8905 ↑ | 2.29 | 3.51E-07 | 6.07E-05 | 4.95 ↑ | 2.3074 | 0.000955 | 0.16371 | 2.7726 ↑ | 1.4712 |
| C-reactive protein | 1.83E-07 | 3.16E-05 | 97.944 ↑ | 6.6139 | 8.3E-06 | 0.000718 | 14.558 ↑ | 3.8638 | 0.14522 | 0.66686 | 2.008 ↑ | 1.0057 |
| cDNA FLJ35730 fis, clone TESTI2003131, highly similar to ALPHA-1-/ | 0.004638 | 0.1003 | 3.5267 ↑ | 1.8183 | 0.000135 | 0.005818 | 5.0222 ↑ | 2.3283 | 0.043613 | 0.59163 | 2.3627 ↑ | 1.2404 |
| Fibronectin | 0.034512 | 0.24877 | 0.35132 ↓ | -1.509 | 0.070526 | 0.2542 | 0.70666 ↓ | -0.501 | 0.0976 | 0.59163 | 0.60618 ↓ | -0.722 |
| Retinol-binding protein | 0.00165 | 0.047582 | 0.41509 ↓ | -1.269 | 0.038541 | 0.23368 | 0.59224 ↓ | -0.756 | 0.28921 | 0.69405 | 0.76848 ↓ | -0.38 |
| Clusterin | 0.071341 | 0.3757 | 0.47093 ↓ | -1.086 | 0.08018 | 0.26761 | 0.46623 ↓ | -1.101 | 0.07414 | 0.59163 | 0.65719 ↓ | -0.606 |
| Apolipoprotein C-III | 0.008303 | 0.12691 | 0.35711 ↓ | -1.486 | 0.004634 | 0.061313 | 0.37504 ↓ | -1.415 | 0.005541 | 0.23963 | 0.35956 ↓ | -1.476 |
| Testis cDNA clone: QtsA-10685, similar to human I factor (complem | 0.059921 | 0.34554 | 0.79104 ↓ | -0.338 | 0.044406 | 0.23368 | 0.78893 ↓ | -0.342 | 0.76078 | 0.95254 | 0.99509 ↓ | -0.007 |
| Protein C, inactivator of coagulation factors Va and VIIIa | 0.000997 | 0.04689 | 0.63342 ↓ | -0.659 | 0.014452 | 0.11906 | 0.53018 ↓ | -0.915 | 0.090273 | 0.59163 | 0.60278 ↓ | -0.73 |
| Endonuclease domain containing 1 | 0.68268 | 0.84967 | 0.72668 ↓ | -0.461 | 0.75952 | 0.92424 | 0.87474 ↓ | -0.193 | 0.8939 | 0.99526 | 0.60554 ↓ | -0.724 |
| Macaca fascicularis brain cDNA clone: QtrA-15720, similar to huma | 0.023885 | 0.2066 | 0.58988 ↓ | -0.762 | 0.18821 | 0.41908 | 0.77061 ↓ | -0.376 | 0.071739 | 0.59163 | 0.61237 ↓ | -0.708 |

| | | | | | | | | | | | | |
|---|---|---|---|---|---|---|---|---|---|---|---|---|
| Apolipoprotein B | 0.0545 | 0.32512 | 0.53856 ↓ | -0.893 | 0.13837 | 0.3627 | 0.74378 ↓ | -0.427 | 0.061845 | 0.59163 | 0.56641 ↓ | -0.82 |
| Leucine-rich alpha-2-glycoprotein | 0.013574 | 0.16644 | 2.8809 ↑ | 1.5265 | 0.11652 | 0.32291 | 1.702 ↑ | 0.7672 | 0.34691 | 0.71818 | 1.3867 ↑ | 0.4717 |
| Lambda-chain (AA -20 to 215) | 0.41113 | 0.74517 | 0.76432 ↓ | -0.388 | 0.35016 | 0.56615 | 0.74017 ↓ | -0.434 | 0.66439 | 0.92693 | 0.83614 ↓ | -0.258 |
| Inter-alpha-trypsin inhibitor heavy chain 1 | 0.022059 | 0.20085 | 0.48542 ↓ | -1.043 | 0.84265 | 0.94661 | 1.3259 ↑ | 0.407 | 0.38309 | 0.74547 | 1.2934 ↑ | 0.3712 |
| Apolipoprotein H | 0.17859 | 0.50649 | 0.82894 ↓ | -0.271 | 0.21807 | 0.43867 | 0.85433 ↓ | -0.227 | 0.65201 | 0.92693 | 0.95259 ↓ | -0.07 |
| Testis cDNA, clone: QtsA-17169, similar to human complement con | 0.086469 | 0.41553 | 0.83271 ↓ | -0.264 | 0.042814 | 0.23368 | 0.8267 ↓ | -0.275 | 0.75981 | 0.95254 | 0.97119 ↓ | -0.042 |
| Cathepsin D | 0.3806 | 0.74476 | 1.1298 ↑ | 0.1761 | 0.41674 | 0.63243 | 0.61622 ↓ | -0.698 | 0.89072 | 0.99526 | 0.70624 ↓ | -0.502 |
| Histidine rich glycoprotein | 0.81722 | 0.916 | 0.85966 ↓ | -0.218 | 0.87403 | 0.95144 | 0.93229 ↓ | -0.101 | 0.52991 | 0.84638 | 1.2663 ↑ | 0.3406 |
| Inter-alpha (Globulin) inhibitor H2 | 0.002097 | 0.051832 | 0.69235 ↓ | -0.53 | 3.29E-05 | 0.0019 | 0.59474 ↓ | -0.75 | 0.056517 | 0.59163 | 0.76389 ↓ | -0.389 |
| Lipopolysaccharide-binding protein | 0.14418 | 0.50569 | 6.3907 ↑ | 2.676 | 0.11275 | 0.32291 | 2.8586 ↑ | 1.5153 | 0.18488 | 0.66686 | 2.3862 ↑ | 1.2547 |
| Inter-alpha-trypsin inhibitor heavy chain 2 | 0.45411 | 0.74953 | 0.92711 ↓ | -0.109 | 0.76459 | 0.92424 | 1.799 ↑ | 0.8472 | 0.79486 | 0.96414 | 1.9733 ↑ | 0.9806 |
| C-X-C motif chemokine | 0.16068 | 0.50649 | 0.74734 ↓ | -0.42 | 0.009701 | 0.094391 | 0.56168 ↓ | -0.832 | 0.17367 | 0.66686 | 0.60769 ↓ | -0.719 |
| Hexosyltransferase | 0.70891 | 0.85397 | 0.96943 ↓ | -0.045 | 0.80208 | 0.93383 | 0.93696 ↓ | -0.094 | 0.65357 | 0.92693 | 0.92891 ↓ | -0.106 |
| Platelet factor 4 | 0.020944 | 0.20085 | 0.44178 ↓ | -1.179 | 0.005302 | 0.061313 | 0.42201 ↓ | -1.245 | 0.031812 | 0.59163 | 0.43676 ↓ | -1.195 |
| Alpha-2-HS-glycoprotein | 0.082223 | 0.41553 | 0.73505 ↓ | -0.444 | 0.010881 | 0.099076 | 0.57779 ↓ | -0.791 | 0.12804 | 0.66686 | 0.75646 ↓ | -0.403 |
| N-acetylmuramoyl-L-alanine amidase | 0.46997 | 0.74953 | 0.89184 ↓ | -0.165 | 0.014134 | 0.11906 | 0.58701 ↓ | -0.769 | 0.41262 | 0.77491 | 0.90554 ↓ | -0.143 |
| Vitellogenin domain-containing protein | 0.60259 | 0.83736 | 0.91327 ↓ | -0.131 | 0.7584 | 0.92424 | 0.93586 ↓ | -0.096 | 0.70427 | 0.94578 | 0.97099 ↓ | -0.042 |
| Apolipoprotein H | 0.026613 | 0.2131 | 0.56404 ↓ | -0.826 | 0.26819 | 0.50149 | 0.82572 ↓ | -0.276 | 0.077664 | 0.59163 | 0.6548 ↓ | -0.611 |
| C3-beta-c | 0.44771 | 0.74953 | 0.88368 ↓ | -0.178 | 0.037839 | 0.23368 | 0.69411 ↓ | -0.527 | 0.30323 | 0.70889 | 0.83853 ↓ | -0.254 |
| Vitamin K-dependent protein S | 0.15087 | 0.50569 | 0.5583 ↓ | -0.841 | 0.37036 | 0.58247 | 0.79684 ↓ | -0.328 | 0.25738 | 0.69405 | 0.69779 ↓ | -0.519 |
| Fibronectin | 0.007687 | 0.12691 | 0.28946 ↓ | -1.789 | 0.11343 | 0.32291 | 0.70574 ↓ | -0.503 | 0.08737 | 0.59163 | 0.67751 ↓ | -0.562 |
| Plasminogen | 0.21193 | 0.53917 | 0.79888 ↓ | -0.324 | 0.038505 | 0.23368 | 0.66673 ↓ | -0.585 | 0.13837 | 0.66686 | 0.7547 ↓ | -0.406 |
| Plasma kallikrein | 0.019411 | 0.19754 | 0.78287 ↓ | -0.353 | 0.051623 | 0.23368 | 0.74938 ↓ | -0.416 | 0.13178 | 0.66686 | 0.77751 ↓ | -0.363 |
| Testis cDNA, clone: QtsA-16984, similar to human sex hormone-bin | 0.53732 | 0.76509 | 0.9103 ↓ | -0.136 | 0.004959 | 0.061313 | 0.57347 ↓ | -0.802 | 0.4694 | 0.80403 | 0.87936 ↓ | -0.185 |
| Plasminogen | 0.10249 | 0.45464 | 0.57643 ↓ | -0.795 | 0.11169 | 0.32291 | 0.55991 ↓ | -0.837 | 0.90374 | 0.99526 | 0.9755 ↓ | -0.036 |
| Gc-globulin | 0.001179 | 0.04689 | 0.72027 ↓ | -0.473 | 0.002622 | 0.056554 | 0.64587 ↓ | -0.631 | 0.00711 | 0.24601 | 0.83254 ↓ | -0.264 |
| Alpha-2-macroglobulin | 0.48302 | 0.74953 | 0.78562 ↓ | -0.348 | 0.003307 | 0.057211 | 0.31962 ↓ | -1.646 | 0.031468 | 0.59163 | 0.43317 ↓ | -1.207 |
| Coagulation factor XII-Mie | 0.20676 | 0.53917 | 0.8224 ↓ | -0.282 | 0.005316 | 0.061313 | 0.59959 ↓ | -0.738 | 0.17122 | 0.66686 | 0.78236 ↓ | -0.354 |
| Alpha-2-glycoprotein 1, zinc-binding | 0.014431 | 0.16644 | 0.70674 ↓ | -0.501 | 0.001099 | 0.02716 | 0.68415 ↓ | -0.548 | 0.028346 | 0.59163 | 0.71781 ↓ | -0.478 |
| Fibronectin | 0.017475 | 0.18895 | 0.29288 ↓ | -1.772 | 0.33791 | 0.56244 | 0.65848 ↓ | -0.603 | 0.26028 | 0.69405 | 0.62346 ↓ | -0.682 |
| SERPIN domain-containing protein | 0.73343 | 0.86315 | 0.97697 ↓ | -0.034 | 0.031208 | 0.22496 | 0.8196 ↓ | -0.287 | 0.1748 | 0.66686 | 0.85682 ↓ | -0.223 |
| Receptor protein-tyrosine kinase | 0.61898 | 0.83736 | 1.6756 ↑ | 0.7446 | 0.46302 | 0.65658 | 1.4967 ↑ | 0.5818 | 0.38351 | 0.74547 | 0.56881 ↓ | -0.814 |
| cDNA FLJ51597, highly similar to C4b-binding protein alpha chain | 0.53672 | 0.76509 | 0.78109 ↓ | -0.356 | 0.91672 | 0.95537 | 1.0134 ↑ | 0.0192 | 0.38905 | 0.74784 | 1.3315 ↑ | 0.413 |
| Fibronectin | 0.001355 | 0.04689 | 0.19064 ↓ | -2.391 | 0.157 | 0.39943 | 0.56632 ↓ | -0.82 | 0.26922 | 0.69405 | 0.65804 ↓ | -0.604 |
| Keratin, type I cytoskeletal 10 | 0.042299 | 0.2924 | 0.3846 ↓ | -1.379 | 0.050936 | 0.23368 | 0.40378 ↓ | -1.308 | 0.44736 | 0.79787 | 0.80046 ↓ | -0.321 |
| Prothrombin | 0.85746 | 0.9426 | 0.92765 ↓ | -0.108 | 0.77408 | 0.92424 | 0.80328 ↓ | -0.316 | 0.53327 | 0.84638 | 0.9635 ↓ | -0.054 |
| C-type lectin domain family 3 member B | 0.053953 | 0.32512 | 0.76573 ↓ | -0.385 | 0.045481 | 0.23368 | 0.73626 ↓ | -0.442 | 0.010123 | 0.29188 | 0.61249 ↓ | -0.707 |
| C3/C5 convertase | 0.96595 | 0.98299 | 0.954 ↓ | -0.068 | 0.86962 | 0.95144 | 0.9343 ↓ | -0.098 | 0.81366 | 0.96414 | 0.99821 ↓ | -0.003 |
| Hemopexin | 0.1271 | 0.47237 | 1.2223 ↑ | 0.2896 | 0.000293 | 0.010131 | 1.5974 ↑ | 0.6758 | 0.067131 | 0.59163 | 1.2923 ↑ | 0.3699 |
| SERPIN domain-containing protein | 0.11382 | 0.46732 | 0.6702 ↓ | -0.577 | 0.048864 | 0.23368 | 0.72163 ↓ | -0.471 | 0.24894 | 0.69405 | 0.72878 ↓ | -0.456 |
| Hyaluronan-binding protein 2 | 0.35984 | 0.73488 | 0.81243 ↓ | -0.3 | 0.17979 | 0.41908 | 0.75959 ↓ | -0.397 | 0.181 | 0.66686 | 0.70724 ↓ | -0.5 |
| SERPIN domain-containing protein | 0.36476 | 0.73488 | 0.80421 ↓ | -0.314 | 0.17753 | 0.41908 | 0.73243 ↓ | -0.449 | 0.82116 | 0.9664 | 0.96133 ↓ | -0.057 |
| Complement component C9 | 0.95971 | 0.98243 | 0.94385 ↓ | -0.083 | 0.71412 | 0.90177 | 0.89269 ↓ | -0.164 | 0.51328 | 0.8377 | 0.84636 ↓ | -0.241 |
| Uncharacterized protein | 0.99995 | 0.99995 | 0.93807 ↓ | -0.092 | 0.20155 | 0.4287 | 1.344 ↑ | 0.4265 | 0.45942 | 0.80403 | 1.3375 ↑ | 0.4195 |
| Uncharacterized protein | 0.67154 | 0.84186 | 1.186 ↑ | 0.2461 | 0.018792 | 0.14135 | 0.40743 ↓ | -1.295 | 0.21872 | 0.69405 | 0.64491 ↓ | -0.633 |
| Haptoglobin | 0.086201 | 0.41553 | 2.1767 ↑ | 1.1222 | 0.31743 | 0.54916 | 1.2244 ↑ | 0.2921 | 0.70422 | 0.94578 | 0.83187 ↓ | -0.266 |
| Joining chain of multimeric IgA and IgM | 0.62641 | 0.83736 | 1.034 ↑ | 0.0482 | 0.20334 | 0.4287 | 0.372 ↓ | -1.427 | 0.37799 | 0.74547 | 0.48681 ↓ | -1.039 |
| Serpin peptidase inhibitor, clade D (Heparin cofactor), member 1 | 0.044462 | 0.2924 | 0.52654 ↓ | -0.925 | 0.009821 | 0.094391 | 0.65425 ↓ | -0.612 | 0.13309 | 0.66686 | 0.75864 ↓ | -0.399 |
| Complement C3 alpha chain | 0.43073 | 0.74517 | 1.1273 ↑ | 0.1729 | 0.28349 | 0.5215 | 0.80389 ↓ | -0.315 | 0.62117 | 0.92641 | 1.1192 ↑ | 0.1624 |
| Uncharacterized protein | 0.093127 | 0.42397 | 2.1763 ↑ | 1.1218 | 0.5426 | 0.72767 | 0.76472 ↓ | -0.387 | 0.42105 | 0.77491 | 1.1915 ↑ | 0.2527 |
| Coagulation factor XIII B chain | 0.75402 | 0.88138 | 0.90654 ↓ | -0.142 | 0.04808 | 0.23368 | 0.60551 ↓ | -0.724 | 0.525 | 0.84638 | 0.88209 ↓ | -0.181 |
| C3-beta-c | 0.78511 | 0.89985 | 0.92115 ↓ | -0.119 | 0.11025 | 0.32291 | 0.36595 ↓ | -1.45 | 0.83249 | 0.97312 | 1.3739 ↑ | 0.4583 |
| Uncharacterized protein | 0.45369 | 0.74953 | 1.2417 ↑ | 0.3123 | 0.66928 | 0.85767 | 0.60847 ↓ | -0.717 | 0.96074 | 0.99526 | 0.80878 ↓ | -0.306 |

| | | | | | | | | | | | | |
|---|---|---|---|---|---|---|---|---|---|---|---|---|
| Keratin 9 | 0.1508 | 0.50569 | 0.39093 ↓ | -1.355 | 0.28637 | 0.5215 | 0.43363 ↓ | -1.206 | 0.25553 | 0.69405 | 0.43096 ↓ | -1.214 |
| Soluble type II IL-1 receptor | 0.19473 | 0.53184 | 0.76296 ↓ | -0.39 | 0.11294 | 0.32291 | 0.65073 ↓ | -0.62 | 0.26376 | 0.69405 | 0.78489 ↓ | -0.349 |
| Epididymis secretory sperm binding protein Li 163pA | 0.089569 | 0.4188 | 0.67845 ↓ | -0.56 | 0.62661 | 0.81507 | 0.89523 ↓ | -0.16 | 0.77457 | 0.95715 | 0.96381 ↓ | -0.053 |
| Complement C5 | 0.9922 | 0.99995 | 0.99535 ↓ | -0.007 | 0.47469 | 0.66765 | 0.8623 ↓ | -0.214 | 0.80337 | 0.96414 | 1.0204 ↑ | 0.0291 |
| C3-beta-c | 0.13106 | 0.47237 | 0.80914 ↓ | -0.306 | 0.2027 | 0.4287 | 0.86746 ↓ | -0.205 | 0.94406 | 0.99526 | 1.0463 ↑ | 0.0652 |
| C1q domain-containing protein | 0.11886 | 0.46732 | 0.087154 ↓ | -3.52 | 0.30106 | 0.53146 | 0.46407 ↓ | -1.108 | 0.60401 | 0.92472 | 1.2045 ↑ | 0.2685 |
| Gc-globulin | 0.027099 | 0.2131 | 0.82337 ↓ | -0.28 | 0.018609 | 0.14135 | 0.82636 ↓ | -0.275 | 0.51096 | 0.8377 | 0.94778 ↓ | -0.077 |
| Apolipoprotein B | 0.21046 | 0.53917 | 0.66388 ↓ | -0.591 | 0.838 | 0.94661 | 0.94194 ↓ | -0.086 | 0.91619 | 0.99526 | 0.97245 ↓ | -0.04 |
| Mannan-binding lectin serine protease 1 | 0.53347 | 0.76509 | 0.94264 ↓ | -0.085 | 0.080438 | 0.26761 | 0.80536 ↓ | -0.312 | 0.058192 | 0.59163 | 0.78361 ↓ | -0.352 |
| Complement component C6 | 0.93783 | 0.98243 | 1.0547 ↑ | 0.0769 | 0.16183 | 0.40575 | 0.87407 ↓ | -0.194 | 0.48864 | 0.82877 | 0.96659 ↓ | -0.049 |
| Clusterin | 0.32509 | 0.70441 | 0.83377 ↓ | -0.262 | 0.19137 | 0.41908 | 0.78547 ↓ | -0.348 | 0.3229 | 0.71107 | 0.89295 ↓ | -0.163 |
| Albumin | 0.91085 | 0.96673 | 1.1537 ↑ | 0.2062 | 0.054617 | 0.23616 | 0.50014 ↓ | -1 | 0.17672 | 0.66686 | 0.6342 ↓ | -0.657 |
| Actin-depolymerizing factor | 0.014302 | 0.16644 | 0.43378 ↓ | -1.205 | 0.30458 | 0.53225 | 0.79273 ↓ | -0.335 | 0.001893 | 0.16371 | 0.27658 ↓ | -1.854 |
| Vitronectin | 0.42402 | 0.74517 | 0.86441 ↓ | -0.21 | 0.18762 | 0.41908 | 0.79558 ↓ | -0.33 | 0.23321 | 0.69405 | 0.78618 ↓ | -0.347 |
| C4a anaphylatoxin | 0.47596 | 0.74953 | 0.89132 ↓ | -0.166 | 0.25421 | 0.49414 | 0.79548 ↓ | -0.33 | 0.63617 | 0.92693 | 1.1185 ↑ | 0.1616 |
| Insulin-like growth factor I | 0.35476 | 0.73488 | 0.39629 ↓ | -1.335 | 0.048377 | 0.23368 | 0.15775 ↓ | -2.664 | 0.13537 | 0.66686 | 0.23316 ↓ | -2.101 |
| Hepatocyte growth factor activator | 0.12476 | 0.47237 | 0.79364 ↓ | -0.333 | 0.009127 | 0.094391 | 0.67648 ↓ | -0.564 | 0.059354 | 0.59163 | 0.71863 ↓ | -0.477 |
| Carbonic anhydrase | 0.48932 | 0.74953 | 1.3409 ↑ | 0.4232 | 0.50273 | 0.69026 | 0.68024 ↓ | -0.556 | 0.19554 | 0.66686 | 0.5242 ↓ | -0.932 |
| Beta actin variant | 0.72069 | 0.85397 | 0.95558 ↓ | -0.066 | 0.06285 | 0.24712 | 0.64942 ↓ | -0.623 | 0.19288 | 0.66686 | 0.76005 ↓ | -0.396 |
| Alpha-1-microglobulin | 0.90703 | 0.96673 | 0.99639 ↓ | -0.005 | 0.070529 | 0.2542 | 0.82288 ↓ | -0.281 | 0.92613 | 0.99526 | 1.0064 ↑ | 0.0092 |
| Gc-globulin | 0.008803 | 0.12691 | 0.80352 ↓ | -0.316 | 0.003946 | 0.061313 | 0.80077 ↓ | -0.321 | 0.49869 | 0.83761 | 0.96072 ↓ | -0.058 |
| Mannose receptor C type 2 | 0.17567 | 0.50649 | 1.2635 ↑ | 0.3374 | 0.82221 | 0.93581 | 1.1527 ↑ | 0.2051 | 0.68356 | 0.93877 | 1.1101 ↑ | 0.1507 |
| Macaca fascicularis brain cDNA clone: QtrA-15786, similar to huma | 0.8878 | 0.95398 | 0.9692 ↓ | -0.045 | 0.067596 | 0.2542 | 0.6616 ↓ | -0.596 | 0.18701 | 0.66686 | 0.73378 ↓ | -0.447 |
| Alpha-1-globin | 0.31755 | 0.70431 | 1.7355 ↑ | 0.7953 | 0.098039 | 0.31409 | 0.083586 ↓ | -3.581 | 0.41593 | 0.77491 | 1.8578 ↑ | 0.8936 |
| Coagulation factor X | 0.42698 | 0.74517 | 0.80358 ↓ | -0.315 | 0.3374 | 0.56244 | 1.1494 ↑ | 0.2009 | 0.86388 | 0.98975 | 1.1397 ↑ | 0.1886 |
| Ceruloplasmin | 0.029259 | 0.22008 | 0.59168 ↓ | -0.757 | 0.1297 | 0.3452 | 0.73578 ↓ | -0.443 | 0.20044 | 0.66686 | 0.78597 ↓ | -0.347 |
| IgGFc-binding protein | 0.40455 | 0.74517 | 1.9978 ↑ | 0.9984 | 0.18961 | 0.41908 | 0.74082 ↓ | -0.433 | 0.94777 | 0.99526 | 1.1259 ↑ | 0.1711 |
| Profilin | 0.47176 | 0.74953 | 0.7049 ↓ | -0.505 | 0.062113 | 0.24712 | 0.42799 ↓ | -1.224 | 0.4208 | 0.77491 | 0.73585 ↓ | -0.443 |
| Serpin family G member 1 | 0.53017 | 0.76509 | 0.91533 ↓ | -0.128 | 0.48615 | 0.67339 | 0.84644 ↓ | -0.241 | 0.50748 | 0.8377 | 0.88487 ↓ | -0.176 |
| HCG40889, isoform CRA_b | 0.66292 | 0.83736 | 0.88836 ↓ | -0.171 | 0.89055 | 0.95144 | 0.98521 ↓ | -0.022 | 0.19839 | 0.66686 | 0.6816 ↓ | -0.553 |
| C4a anaphylatoxin | 0.66311 | 0.83736 | 1.1365 ↑ | 0.1846 | 0.77999 | 0.92424 | 0.93196 ↓ | -0.102 | 0.28501 | 0.69405 | 1.3757 ↑ | 0.4601 |
| Plasminogen | 0.19675 | 0.53184 | 1.323 ↑ | 0.4038 | 0.43913 | 0.64381 | 0.87506 ↓ | -0.193 | 0.80292 | 0.96414 | 1.0208 ↑ | 0.0297 |
| Protein S100 | 0.87177 | 0.9426 | 0.97771 ↓ | -0.033 | 0.16446 | 0.40646 | 0.15689 ↓ | -2.672 | 0.95613 | 0.99526 | 2.3599 ↑ | 1.2387 |
| Reversion inducing cysteine rich protein with kazal motifs | 0.16727 | 0.50649 | 1.6701 ↑ | 0.7399 | 0.22754 | 0.45246 | 0.49069 ↓ | -1.027 | 0.70523 | 0.94578 | 1.3969 ↑ | 0.4822 |
| Complement component C9 | 0.31304 | 0.70431 | 1.155 ↑ | 0.2079 | 0.7385 | 0.91257 | 1.0189 ↑ | 0.027 | 0.65987 | 0.92693 | 1.0542 ↑ | 0.0761 |
| Glyceraldehyde-3-phosphate dehydrogenase | 0.38314 | 0.74476 | 0.38751 ↓ | -1.368 | 0.55324 | 0.73623 | 1.47 ↑ | 0.5559 | 0.75188 | 0.95254 | 0.74986 ↓ | -0.415 |
| Complement subcomponent C1r | 0.17583 | 0.50649 | 0.80378 ↓ | -0.315 | 0.34237 | 0.56244 | 0.86761 ↓ | -0.205 | 0.046835 | 0.59163 | 0.74938 ↓ | -0.416 |
| Fibrinogen beta chain | 0.42793 | 0.74517 | 1.2393 ↑ | 0.3096 | 0.35559 | 0.5696 | 0.78461 ↓ | -0.35 | 0.15642 | 0.66686 | 1.2847 ↑ | 0.3615 |
| Cation-independent mannose-6-phosphate receptor | 0.25549 | 0.61388 | 1.2843 ↑ | 0.361 | 0.34462 | 0.56244 | 1.2119 ↑ | 0.2773 | 0.97391 | 0.99696 | 1.1983 ↑ | 0.261 |
| Complement C1r subcomponent-like protein | 0.63084 | 0.83736 | 0.96218 ↓ | -0.056 | 0.80346 | 0.93383 | 1.0904 ↑ | 0.1249 | 0.35355 | 0.71818 | 0.83721 ↓ | -0.256 |
| Haptoglobin | 0.045635 | 0.2924 | 2.9447 ↑ | 1.5581 | 0.055969 | 0.23616 | 2.2145 ↑ | 1.147 | 0.092367 | 0.59163 | 1.499 ↑ | 0.584 |
| Apolipoprotein A-II | 0.87011 | 0.9426 | 1.0698 ↑ | 0.0973 | 0.91007 | 0.95537 | 0.46304 ↓ | -1.111 | 0.27818 | 0.69405 | 0.3117 ↓ | -1.682 |
| Alpha-2-HS-glycoprotein | 0.00531 | 0.10207 | 0.51555 ↓ | -0.956 | 0.064341 | 0.24736 | 0.55399 ↓ | -0.852 | 0.24097 | 0.69405 | 0.76781 ↓ | -0.381 |
| Apolipoprotein A-I | 0.49945 | 0.75135 | 2.058 ↑ | 1.0413 | 0.079319 | 0.26761 | 0.33574 ↓ | -1.575 | 0.35477 | 0.71818 | 0.52893 ↓ | -0.919 |
| Hemoglobin subunit beta | 0.10933 | 0.46732 | 1.9397 ↑ | 0.9558 | 0.52751 | 0.71296 | 0.43339 ↓ | -1.206 | 0.68373 | 0.93877 | 0.40397 ↓ | -1.308 |
| Angiotensin-converting enzyme | 0.23778 | 0.58766 | 1.1954 ↑ | 0.2575 | 0.87153 | 0.95144 | 0.9267 ↓ | -0.11 | 0.98139 | 0.99871 | 1.0117 ↑ | 0.0168 |
| Complement C8 gamma chain | 0.41144 | 0.74517 | 0.90515 ↓ | -0.144 | 0.32804 | 0.56188 | 0.89325 ↓ | -0.163 | 0.90211 | 0.99526 | 0.99144 ↓ | -0.012 |
| Antithrombin-III | 0.71291 | 0.85397 | 0.95436 ↓ | -0.067 | 0.69893 | 0.88908 | 0.882 ↓ | -0.181 | 0.19142 | 0.66686 | 0.69482 ↓ | -0.525 |
| C1q domain-containing protein | 0.55644 | 0.78264 | 0.85485 ↓ | -0.226 | 0.96463 | 0.98131 | 1.119 ↑ | 0.1623 | 0.66328 | 0.92693 | 1.1066 ↑ | 0.1462 |
| Epididymis secretory protein Li 37 | 0.62799 | 0.83736 | 3.3046 ↑ | 1.7245 | 0.42818 | 0.63312 | 1.5857 ↑ | 0.6652 | 0.081229 | 0.59163 | 0.61602 ↓ | -0.699 |
| Fibrinogen gamma chain | 0.43604 | 0.74689 | 1.4275 ↑ | 0.5135 | 0.21369 | 0.43493 | 2.0707 ↑ | 1.0501 | 0.59109 | 0.91302 | 1.1253 ↑ | 0.1703 |
| Apolipoprotein C-I, basic form | 0.22373 | 0.56094 | 0.62252 ↓ | -0.684 | 0.057621 | 0.23734 | 0.48131 ↓ | -1.055 | 0.06283 | 0.59163 | 0.46363 ↓ | -1.109 |
| Uncharacterized protein | 0.71496 | 0.85397 | 0.92195 ↓ | -0.117 | 0.45391 | 0.65062 | 0.86867 ↓ | -0.203 | 0.6172 | 0.92641 | 0.90455 ↓ | -0.145 |

| | | | | | | | | | | | | |
|---|---|---|---|---|---|---|---|---|---|---|---|---|
| Complement C4-B | 0.4717 | 0.74953 | 1.2687 ↑ | 0.3433 | 0.9118 | 0.95537 | 1.1354 ↑ | 0.1832 | 0.077538 | 0.59163 | 1.5727 ↑ | 0.6533 |
| SERPIN domain-containing protein | 0.65574 | 0.83736 | 0.99009 ↓ | -0.014 | 0.002942 | 0.056554 | 0.57585 ↓ | -0.796 | 0.094326 | 0.59163 | 0.70216 ↓ | -0.51 |
| Lumican | 0.12896 | 0.47237 | 0.65421 ↓ | -0.612 | 0.14902 | 0.38479 | 0.67561 ↓ | -0.566 | 0.14678 | 0.66686 | 0.68388 ↓ | -0.548 |
| Cadherin-13 | 0.95369 | 0.98243 | 1.47 ↑ | 0.5558 | 0.077815 | 0.26761 | 0.5192 ↓ | -0.946 | 0.74548 | 0.95254 | 0.85559 ↓ | -0.225 |
| Angiotensin 1-10 | 0.424 | 0.74517 | 1.155 ↑ | 0.2079 | 0.48655 | 0.67339 | 0.90143 ↓ | -0.15 | 0.29193 | 0.69405 | 1.1493 ↑ | 0.2007 |
| Keratin, type II cytoskeletal 2 epidermal | 0.152 | 0.50569 | 0.48678 ↓ | -1.039 | 0.37398 | 0.58287 | 0.62174 ↓ | -0.686 | 0.81036 | 0.96414 | 0.98913 ↓ | -0.016 |
| Fibulin-1 | 0.87005 | 0.9426 | 0.89973 ↓ | -0.152 | 0.11347 | 0.32291 | 0.60911 ↓ | -0.715 | 0.58628 | 0.91302 | 0.87532 ↓ | -0.192 |
| Trypsin | 0.32981 | 0.70441 | 1.4243 ↑ | 0.5102 | 0.81508 | 0.93383 | 1.193 ↑ | 0.2546 | 0.469 | 0.80403 | 1.8342 ↑ | 0.8751 |
| 1,4-beta-N-acetylmuramidase C | 0.26307 | 0.62344 | 1.8798 ↑ | 0.9106 | 0.92277 | 0.95593 | 1.1948 ↑ | 0.2568 | 0.94787 | 0.99526 | 0.8421 ↓ | -0.248 |
| Macaca fascicularis brain cDNA clone: QorA-13734, similar to huma | 0.17749 | 0.50649 | 1.2906 ↑ | 0.368 | 0.19027 | 0.41908 | 1.2248 ↑ | 0.2926 | 0.95319 | 0.99526 | 0.53653 ↓ | -0.898 |
| Testicular tissue protein Li 61 | 0.41764 | 0.74517 | 0.89127 ↓ | -0.166 | 0.047704 | 0.23368 | 0.7353 ↓ | -0.444 | 0.31437 | 0.71107 | 0.85938 ↓ | -0.219 |
| Uncharacterized protein | 0.99915 | 0.99995 | 1.0432 ↑ | 0.0611 | 0.12675 | 0.34263 | 0.82498 ↓ | -0.278 | 0.26623 | 0.69405 | 0.8653 ↓ | -0.209 |
| Vitellogenin domain-containing protein | 0.63833 | 0.83736 | 0.9399 ↓ | -0.089 | 0.88306 | 0.95144 | 0.94973 ↓ | -0.074 | 0.23486 | 0.69405 | 1.1927 ↑ | 0.2542 |
| Uncharacterized protein | 0.17162 | 0.50649 | 0.68029 ↓ | -0.556 | 0.3908 | 0.5983 | 0.78205 ↓ | -0.355 | 0.33938 | 0.71818 | 0.758 ↑ | -0.4 |
| Serpin family A member 10 | 0.11697 | 0.46732 | 3.1096 ↑ | 1.6367 | 0.001047 | 0.02716 | 4.2829 ↑ | 2.0986 | 0.4656 | 0.80403 | 1.5925 ↑ | 0.6713 |
| Complement component 8 subunit beta | 0.63547 | 0.83736 | 0.56633 ↓ | -0.82 | 0.81219 | 0.93383 | 0.69108 ↓ | -0.533 | 0.75089 | 0.95254 | 2.3703 ↑ | 1.2451 |
| Collagen type VI alpha 3 chain | 0.52999 | 0.76509 | 0.8709 ↓ | -0.199 | 0.18823 | 0.41908 | 0.66857 ↓ | -0.581 | 0.17969 | 0.66686 | 0.66002 ↓ | -0.599 |
| Glutathione peroxidase | 0.24149 | 0.58842 | 1.3306 ↑ | 0.4121 | 0.87742 | 0.95144 | 0.96025 ↓ | -0.059 | 0.80571 | 0.96414 | 0.91721 ↓ | -0.125 |
| Mannan binding lectin serine peptidase 2 | 0.20672 | 0.53917 | 0.70604 ↓ | -0.502 | 0.36703 | 0.58247 | 0.77939 ↓ | -0.36 | 0.96885 | 0.99696 | 0.98113 ↓ | -0.027 |
| Protein S100 | 0.63537 | 0.83736 | 1.1605 ↑ | 0.2147 | 0.29937 | 0.53146 | 0.29044 ↓ | -1.784 | 0.92443 | 0.99526 | 1.5417 ↑ | 0.6245 |
| Bradykinin | 0.39777 | 0.74517 | 0.88701 ↓ | -0.173 | 0.23667 | 0.46527 | 0.73676 ↓ | -0.441 | 0.34179 | 0.71818 | 0.84566 ↓ | -0.242 |
| Protein C receptor | 0.49797 | 0.75135 | 0.95631 ↓ | -0.064 | 0.96997 | 0.98131 | 1.0312 ↑ | 0.0443 | 0.663 | 0.92693 | 0.8982 ↓ | -0.155 |
| Protein-tyrosine-phosphatase | 0.65349 | 0.83736 | 0.85592 ↓ | -0.224 | 0.73418 | 0.91257 | 0.98937 ↓ | -0.015 | 0.85847 | 0.98975 | 1.0363 ↑ | 0.0514 |
| Uncharacterized protein | 0.47112 | 0.74953 | 1.1064 ↑ | 0.1458 | 0.6348 | 0.81955 | 1.1414 ↑ | 0.1908 | 0.32882 | 0.71107 | 0.71145 ↓ | -0.491 |
| Filamin A | 0.69068 | 0.85348 | 0.75956 ↓ | -0.397 | 0.45506 | 0.65062 | 1.089 ↑ | 0.1231 | 0.55556 | 0.87375 | 0.61658 ↓ | -0.698 |
| Inter-alpha-trypsin inhibitor heavy chain 2 | 0.48957 | 0.74953 | 1.1582 ↑ | 0.2119 | 0.9517 | 0.97423 | 0.93871 ↓ | -0.091 | 0.76534 | 0.95254 | 1.0732 ↑ | 0.102 |
| Fibrinogen alpha chain | 0.71973 | 0.85397 | 0.84745 ↓ | -0.239 | 0.052679 | 0.23368 | 0.4583 ↓ | -1.126 | 0.72807 | 0.95254 | 0.81076 ↓ | -0.303 |
| Complement C1s | 0.53954 | 0.76509 | 0.95145 ↓ | -0.072 | 0.86943 | 0.95144 | 0.98399 ↓ | -0.023 | 0.60942 | 0.92482 | 0.95553 ↓ | -0.066 |
| Mannosyl-oligosaccharide 1,2-alpha-mannosidase IA | 0.78542 | 0.89985 | 0.6005 ↓ | -0.736 | 0.090239 | 0.29455 | 1.6157 ↑ | 0.6921 | 0.27421 | 0.69405 | 1.451 ↑ | 0.537 |
| Apolipoprotein B (Including Ag(X) antigen) | 0.11511 | 0.46732 | 0.66624 ↓ | -0.586 | 0.26959 | 0.50149 | 0.8314 ↓ | -0.266 | 0.099175 | 0.59163 | 0.63211 ↓ | -0.662 |
| Complement component C8 alpha chain | 0.76725 | 0.89084 | 0.93691 ↓ | -0.094 | 0.29911 | 0.53146 | 0.73415 ↓ | -0.446 | 0.35701 | 0.71818 | 0.8107 ↓ | -0.303 |
| cDNA FLJ44930 fis, clone BRAMY3015549, highly similar to Neural c | 0.16113 | 0.50649 | 0.59479 ↓ | -0.75 | 0.81255 | 0.93383 | 0.88811 ↓ | -0.171 | 0.99528 | 0.99902 | 0.98636 ↓ | -0.02 |
| Metavinculin | 0.40219 | 0.74517 | 0.65903 ↓ | -0.602 | 0.11658 | 0.32291 | 0.46718 ↓ | -1.098 | 0.23406 | 0.69405 | 0.58027 ↓ | -0.785 |
| ITIH4 protein | 0.34698 | 0.73205 | 1.6687 ↑ | 0.7388 | 0.99389 | 0.99389 | 1.4563 ↑ | 0.5423 | 0.44059 | 0.79398 | 1.9052 ↑ | 0.93 |
| Apolipoprotein C-II | 0.81173 | 0.916 | 0.57826 ↓ | -0.79 | 0.37753 | 0.58315 | 0.34268 ↓ | -1.545 | 0.73626 | 0.95254 | 0.42603 ↓ | -1.231 |
| Prothrombin | 0.71126 | 0.85397 | 0.73986 ↓ | -0.435 | 0.44539 | 0.6475 | 1.1891 ↑ | 0.2499 | 0.99902 | 0.99902 | 0.877 ↓ | -0.189 |
| Alpha-1-antitrypsin | 0.48503 | 0.74953 | 2.2627 ↑ | 1.178 | 0.97858 | 0.98427 | 1.1051 ↑ | 0.1442 | 0.25555 | 0.69405 | 1.0575 ↑ | 0.0806 |
| Complement C1q subcomponent subunit B | 0.8207 | 0.916 | 1.2367 ↑ | 0.3066 | 0.89094 | 0.95144 | 0.93984 ↓ | -0.09 | 0.63959 | 0.92693 | 0.89881 ↓ | -0.154 |
| SEMA4D variant protein | 0.36532 | 0.73488 | 1.7867 ↑ | 0.8373 | 0.95118 | 0.97423 | 1.4663 ↑ | 0.5522 | 0.25292 | 0.69405 | 1.4368 ↑ | 0.5229 |
| CP protein | 0.65175 | 0.83736 | 2.2994 ↑ | 1.2012 | 0.21269 | 0.43493 | 1.6625 ↑ | 0.7334 | 0.99652 | 0.99902 | 0.98681 ↓ | -0.019 |
| Attractin | 0.37198 | 0.73969 | 0.86902 ↓ | -0.203 | 0.035301 | 0.23368 | 0.65407 ↓ | -0.612 | 0.13159 | 0.66686 | 0.7379 ↓ | -0.439 |
| 15S Mg(2+)-ATPase p97 subunit | 0.071666 | 0.3757 | 1.8067 ↑ | 0.8533 | 0.91341 | 0.95537 | 1.0927 ↑ | 0.1279 | 0.080939 | 0.59163 | 1.6702 ↑ | 0.7401 |
| CST6 protein | 0.19359 | 0.53184 | 1.3847 ↑ | 0.4696 | 0.42173 | 0.63312 | 0.90657 ↓ | -0.142 | 0.32767 | 0.71107 | 0.8375 ↓ | -0.256 |
| Monocyte differentiation antigen CD14 | 0.53592 | 0.76509 | 0.90132 ↓ | -0.15 | 0.20568 | 0.4287 | 0.79925 ↓ | -0.323 | 0.32769 | 0.71107 | 0.83811 ↓ | -0.255 |
| Uncharacterized protein | 0.29992 | 0.69181 | 2.1105 ↑ | 1.0776 | 0.42623 | 0.63312 | 0.78254 ↓ | -0.354 | 0.91846 | 0.99526 | 1.2293 ↑ | 0.2978 |
| Immunoglobulin heavy constant mu | 0.93108 | 0.98217 | 0.86875 ↓ | -0.203 | 0.73186 | 0.91257 | 1.359 ↑ | 0.4426 | 0.87493 | 0.99526 | 1.3246 ↑ | 0.4055 |
| Apolipoprotein A4 | 0.32679 | 0.70441 | 0.89973 ↓ | -0.152 | 0.26409 | 0.50149 | 0.75923 ↓ | -0.397 | 0.005119 | 0.23963 | 0.35291 ↓ | -1.503 |
| Ficolin (Collagen/fibrinogen domain containing) 3 (Hakata antigen), | 0.94958 | 0.98243 | 1.3498 ↑ | 0.4328 | 0.18445 | 0.41908 | 0.72922 ↓ | -0.456 | 0.29287 | 0.69405 | 0.61249 ↓ | -0.707 |
| Receptor protein-tyrosine kinase | 0.16555 | 0.50649 | 1.4184 ↑ | 0.5043 | 0.60692 | 0.79543 | 0.95614 ↓ | -0.065 | 0.86388 | 0.98975 | 1.0982 ↑ | 0.1351 |
| Insulin-like growth factor-binding protein 3 | 0.81231 | 0.916 | 1.02 ↑ | 0.0286 | 0.34394 | 0.56244 | 0.89229 ↓ | -0.164 | 0.93743 | 0.99526 | 1.0889 ↑ | 0.1228 |
| Uncharacterized protein DKFZp686I15196 | 0.65995 | 0.83736 | 3.7581 ↑ | 1.91 | 0.77747 | 0.92424 | 1.1703 ↑ | 0.2269 | 0.75281 | 0.95254 | 0.9578 ↓ | -0.062 |
| cDNA, FLJ93914, highly similar to Homo sapiens histidine-rich glyco | 0.066517 | 0.37121 | 0.54908 ↓ | -0.865 | 0.26032 | 0.5004 | 0.75214 ↓ | -0.411 | 0.7368 | 0.95254 | 0.9004 ↓ | -0.151 |
| Coagulation factor XII | 0.28648 | 0.66975 | 1.3114 ↑ | 0.3911 | 0.59391 | 0.78433 | 0.76902 ↓ | -0.379 | 0.14359 | 0.66686 | 2.1126 ↑ | 1.079 |

| | | | | | | | | | | | | |
|---|---|---|---|---|---|---|---|---|---|---|---|---|
| Protein S100-A7 | 0.31472 | 0.70431 | 0.40306 ↓ | -1.311 | 0.51614 | 0.70309 | 0.21455 ↓ | -2.221 | 0.4393 | 0.79398 | 0.52678 ↓ | -0.925 |
| Insulin-like growth factor binding protein, acid labile subunit | 0.95851 | 0.98243 | 1.0022 ↑ | 0.0031 | 0.11759 | 0.32291 | 0.76688 ↓ | -0.383 | 0.27915 | 0.69405 | 0.85978 ↓ | -0.218 |
| Carboxylic ester hydrolase | 0.84362 | 0.93555 | 0.96716 ↓ | -0.048 | 0.045743 | 0.23368 | 0.64494 ↓ | -0.633 | 0.31791 | 0.71107 | 0.81498 ↓ | -0.295 |

## 7.5 Gy

| | SD 1 vs. Pre | | | | SD 2 vs. Pre | | | | SD 6 vs. Pre | | | |
|---|---|---|---|---|---|---|---|---|---|---|---|---|
| **Protein Name** | ***p-value*** | **FDR** | **FC** | **LOG2FC** | ***p-value*** | **FDR** | **FC** | **LOG2FC** | ***p-value*** | **FDR** | **FC** | **LOG2FC** |
| Serpin family A member 9 | 1.28E-06 | 0.000111 | 5.9761 ↑ | 2.5792 | 1.15E-06 | 0.000199 | 5.1405 ↑ | 2.3619 | 0.000246 | 0.014182 | 3.1119 ↑ | 1.6378 |
| C-reactive protein | 6.64E-08 | 1.15E-05 | 62.786 ↑ | 5.9724 | 3.98E-05 | 0.003442 | 14.88 ↑ | 3.8953 | 0.18663 | 0.54723 | 9.0193 ↑ | 3.173 |
| cDNA FLJ35730 fis, clone TESTI2003131, highly similar to ALPHA-1-/ | 0.000474 | 0.011717 | 5.0689 ↑ | 2.3417 | 0.00228 | 0.065725 | 3.9238 ↑ | 1.9723 | 0.006452 | 0.11126 | 2.3957 ↑ | 1.2605 |
| Fibronectin | 0.089727 | 0.32527 | 0.58059 ↓ | -0.784 | 0.078459 | 0.33113 | 0.49853 ↓ | -1.004 | 0.26192 | 0.63441 | 0.6932 ↓ | -0.529 |
| Retinol-binding protein | 0.005383 | 0.066521 | 0.51777 ↓ | -0.95 | 0.032582 | 0.25621 | 0.66424 ↓ | -0.59 | 0.26403 | 0.63441 | 0.82073 ↓ | -0.285 |
| Clusterin | 0.31158 | 0.5555 | 0.48861 ↓ | -1.033 | 0.18817 | 0.5248 | 0.43629 ↓ | -1.197 | 0.60681 | 0.83547 | 0.61955 ↓ | -0.691 |
| Apolipoprotein C-III | 3.69E-05 | 0.002127 | 0.34177 ↓ | -1.549 | 0.000489 | 0.024761 | 0.2625 ↓ | -1.93 | 8.91E-05 | 0.007703 | 0.31954 ↓ | -1.646 |
| Testis cDNA clone: QtsA-10685, similar to human I factor (complem | 0.3665 | 0.60385 | 0.89131 ↓ | -0.166 | 0.51998 | 0.75594 | 0.9242 ↓ | -0.114 | 0.7024 | 0.86363 | 1.0391 ↑ | 0.0554 |
| Protein C, inactivator of coagulation factors Va and VIIIa | 0.003675 | 0.052987 | 0.59177 ↓ | -0.757 | 0.011127 | 0.14807 | 0.58074 ↓ | -0.784 | 0.05517 | 0.31812 | 0.76616 ↓ | -0.384 |
| Endonuclease domain containing 1 | 0.001829 | 0.03164 | 0.20496 ↓ | -2.287 | 0.10446 | 0.37297 | 0.59117 ↓ | -0.758 | 0.65999 | 0.84217 | 0.85419 ↓ | -0.227 |
| Macaca fascicularis brain cDNA clone: QtrA-15720, similar to huma | 0.18802 | 0.4582 | 0.70854 ↓ | -0.497 | 0.24699 | 0.55492 | 0.71728 ↓ | -0.479 | 0.42706 | 0.73707 | 0.80128 ↓ | -0.32 |
| Apolipoprotein B | 0.58801 | 0.74659 | 0.94094 ↓ | -0.088 | 0.4874 | 0.73463 | 0.63333 ↓ | -0.659 | 0.58811 | 0.83547 | 0.79273 ↓ | -0.335 |
| Leucine-rich alpha-2-glycoprotein | 0.007288 | 0.084054 | 2.6893 ↑ | 1.4272 | 0.080389 | 0.33113 | 2.4746 ↑ | 1.3072 | 0.22606 | 0.59971 | 1.5899 ↑ | 0.669 |
| Lambda-chain (AA -20 to 215) | 0.921 | 0.95983 | 0.85895 ↓ | -0.219 | 0.94475 | 0.95932 | 0.88287 ↓ | -0.18 | 0.64451 | 0.84217 | 0.82441 ↓ | -0.279 |
| Inter-alpha-trypsin inhibitor heavy chain 1 | 0.55349 | 0.72442 | 0.76213 ↓ | -0.392 | 0.42147 | 0.67514 | 0.62318 ↓ | -0.682 | 0.32547 | 0.6784 | 0.44297 ↓ | -1.175 |
| Apolipoprotein H | 0.087858 | 0.32527 | 0.85203 ↓ | -0.231 | 0.072351 | 0.33113 | 0.85509 ↓ | -0.226 | 0.81949 | 0.92866 | 0.9912 ↓ | -0.013 |
| Testis cDNA, clone: QtsA-17169, similar to human complement con | 0.73104 | 0.83428 | 1.0265 ↑ | 0.0377 | 0.39126 | 0.65116 | 1.0813 ↑ | 0.1128 | 0.89499 | 0.93811 | 1.0102 ↑ | 0.0147 |
| Cathepsin D | 0.82692 | 0.89411 | 0.68014 ↓ | -0.556 | 0.51773 | 0.75594 | 0.66213 ↓ | -0.595 | 0.14899 | 0.46865 | 0.42382 ↓ | -1.239 |
| Histidine rich glycoprotein | 0.43819 | 0.65864 | 0.70255 ↓ | -0.509 | 0.20235 | 0.53106 | 0.54812 ↓ | -0.867 | 0.82235 | 0.92866 | 0.8228 ↓ | -0.281 |
| Inter-alpha (Globulin) inhibitor H2 | 0.027453 | 0.18997 | 0.77571 ↓ | -0.366 | 0.026657 | 0.23383 | 0.74051 ↓ | -0.433 | 0.099744 | 0.41085 | 0.85111 ↓ | -0.233 |
| Lipopolysaccharide-binding protein | 0.070264 | 0.31989 | 5.6626 ↑ | 2.5015 | 0.29959 | 0.5913 | 3.1104 ↑ | 1.6371 | 0.23133 | 0.59971 | 2.7297 ↑ | 1.4488 |
| Inter-alpha-trypsin inhibitor heavy chain 2 | 0.20331 | 0.4582 | 0.33136 ↓ | -1.594 | 0.2308 | 0.55144 | 0.34844 ↓ | -1.521 | 0.3603 | 0.70832 | 0.39396 ↓ | -1.344 |
| C-X-C motif chemokine | 0.034783 | 0.21883 | 0.63078 ↓ | -0.665 | 0.010913 | 0.14807 | 0.46586 ↓ | -1.102 | 0.007718 | 0.11126 | 0.45634 ↓ | -1.132 |
| Hexosyltransferase | 0.056974 | 0.2899 | 2.5105 ↑ | 1.328 | 0.84304 | 0.92895 | 1.4716 ↑ | 0.5574 | 0.64135 | 0.84217 | 1.0622 ↑ | 0.0871 |
| Platelet factor 4 | 0.090249 | 0.32527 | 0.58301 ↓ | -0.778 | 0.001478 | 0.051128 | 0.20238 ↓ | -2.305 | 0.001095 | 0.027068 | 0.40361 ↓ | -1.309 |
| Alpha-2-HS-glycoprotein | 0.000409 | 0.011717 | 0.45788 ↓ | -1.127 | 0.005942 | 0.10937 | 0.52904 ↓ | -0.919 | 0.025542 | 0.24549 | 0.65269 ↓ | -0.616 |
| N-acetylmuramoyl-L-alanine amidase | 0.066849 | 0.31951 | 0.6731 ↓ | -0.571 | 0.26362 | 0.58469 | 0.79442 ↓ | -0.332 | 0.04777 | 0.31812 | 0.67366 ↓ | -0.57 |
| Vitellogenin domain-containing protein | 0.047136 | 0.25483 | 0.70946 ↓ | -0.495 | 0.008918 | 0.14025 | 0.56502 ↓ | -0.824 | 0.38665 | 0.71925 | 0.88001 ↓ | -0.184 |
| Apolipoprotein H | 0.87125 | 0.92976 | 0.86507 ↓ | -0.209 | 0.40965 | 0.67495 | 0.76805 ↓ | -0.381 | 0.78201 | 0.9141 | 1.1332 ↑ | 0.1804 |
| C3-beta-c | 0.064119 | 0.31693 | 0.6694 ↓ | -0.579 | 0.21987 | 0.54755 | 0.75158 ↓ | -0.412 | 0.46079 | 0.76433 | 0.84052 ↓ | -0.251 |
| Vitamin K-dependent protein S | 0.025531 | 0.18404 | 0.56344 ↓ | -0.828 | 0.12818 | 0.41587 | 0.55935 ↓ | -0.838 | 0.53514 | 0.7981 | 0.86292 ↓ | -0.213 |
| Fibronectin | 0.46541 | 0.6766 | 0.77288 ↓ | -0.372 | 0.028384 | 0.23383 | 0.38071 ↓ | -1.393 | 0.62298 | 0.83547 | 0.82305 ↓ | -0.281 |
| Plasminogen | 0.072812 | 0.32298 | 0.72916 ↓ | -0.456 | 0.096836 | 0.36419 | 0.7473 ↓ | -0.42 | 0.056318 | 0.31812 | 0.74728 ↓ | -0.42 |
| Plasma kallikrein | 0.16294 | 0.42577 | 0.81603 ↓ | -0.293 | 0.013452 | 0.16622 | 0.77437 ↓ | -0.369 | 0.21278 | 0.58568 | 0.87556 ↓ | -0.192 |
| Testis cDNA, clone: QtsA-16984, similar to human sex hormone-bin | 0.13362 | 0.38564 | 0.75144 ↓ | -0.412 | 0.002695 | 0.066613 | 0.55777 ↓ | -0.842 | 0.84423 | 0.92866 | 0.96664 ↓ | -0.049 |
| Plasminogen | 0.11331 | 0.35954 | 0.81575 ↓ | -0.294 | 0.057445 | 0.33113 | 0.79986 ↓ | -0.322 | 0.34241 | 0.6916 | 0.90671 ↓ | -0.141 |
| Gc-globulin | 0.025225 | 0.18404 | 0.70973 ↓ | -0.495 | 0.08026 | 0.33113 | 0.74888 ↓ | -0.417 | 0.32139 | 0.6784 | 0.85883 ↓ | -0.22 |
| Alpha-2-macroglobulin | 0.20327 | 0.4582 | 0.51272 ↓ | -0.964 | 0.10068 | 0.37059 | 0.39451 ↓ | -1.342 | 0.056723 | 0.31812 | 0.33164 ↓ | -1.592 |
| Coagulation factor XII-Mie | 0.079259 | 0.32527 | 0.6937 ↓ | -0.528 | 0.027609 | 0.23383 | 0.62454 ↓ | -0.679 | 0.090847 | 0.40299 | 0.7066 ↓ | -0.501 |
| Alpha-2-glycoprotein 1, zinc-binding | 0.041283 | 0.23562 | 0.78657 ↓ | -0.346 | 0.039653 | 0.28303 | 0.68199 ↓ | -0.552 | 0.093824 | 0.40579 | 0.80412 ↓ | -0.315 |
| Fibronectin | 0.53219 | 0.72442 | 0.78047 ↓ | -0.358 | 0.075078 | 0.33113 | 0.5003 ↓ | -0.999 | 0.99169 | 0.99169 | 0.98125 ↓ | -0.027 |
| SERPIN domain-containing protein | 0.35117 | 0.59561 | 0.88481 ↓ | -0.177 | 0.88345 | 0.9493 | 1.0238 ↑ | 0.0339 | 0.2458 | 0.60843 | 0.85438 ↓ | -0.227 |
| Receptor protein-tyrosine kinase | 0.53782 | 0.72442 | 1.2228 ↑ | 0.2902 | 0.92829 | 0.95592 | 0.89912 ↓ | -0.153 | 0.5093 | 0.78668 | 0.54696 ↓ | -0.87 |
| cDNA FLJ51597, highly similar to C4b-binding protein alpha chain | 0.37995 | 0.60572 | 0.65053 ↓ | -0.62 | 0.66229 | 0.86703 | 0.85473 ↓ | -0.226 | 0.49969 | 0.78587 | 0.67375 ↓ | -0.57 |
| Fibronectin | 0.13897 | 0.39414 | 0.39159 ↓ | -1.353 | 0.44792 | 0.69187 | 0.68782 ↓ | -0.54 | 0.3478 | 0.6916 | 0.49199 ↓ | -1.023 |
| Keratin, type I cytoskeletal 10 | 0.10702 | 0.34934 | 0.6495 ↓ | -0.623 | 0.17142 | 0.50265 | 0.68214 ↓ | -0.552 | 0.13029 | 0.42486 | 0.6885 ↓ | -0.538 |

| | | | | | | | | | | | | | | | |
|---|---|---|---|---|---|---|---|---|---|---|---|---|---|---|---|
| Prothrombin | 0.4567 | 0.66956 | 0.69084 | ↓ | -0.534 | 0.063439 | 0.33113 | 0.4367 | ↓ | -1.195 | 0.65735 | 0.84217 | 0.87067 | ↓ | -0.2 |
| C-type lectin domain family 3 member B | 8.66E-05 | 0.002998 | 0.63945 | ↓ | -0.645 | 0.006322 | 0.10937 | 0.67313 | ↓ | -0.571 | 0.00038 | 0.014755 | 0.51494 | ↓ | -0.958 |
| C3/C5 convertase | 0.10569 | 0.34934 | 0.86985 | ↓ | -0.201 | 0.75358 | 0.89828 | 0.9762 | ↓ | -0.035 | 0.43388 | 0.73707 | 1.1355 | ↑ | 0.1833 |
| Hemopexin | 0.00077 | 0.016657 | 1.6654 | ↑ | 0.7359 | 0.004301 | 0.093003 | 1.6559 | ↑ | 0.7276 | 0.000426 | 0.014755 | 1.6335 | ↑ | 0.708 |
| SERPIN domain-containing protein | 0.084433 | 0.32527 | 0.78259 | ↓ | -0.354 | 0.33727 | 0.60779 | 0.83148 | ↓ | -0.266 | 0.86961 | 0.92866 | 0.97347 | ↓ | -0.039 |
| Hyaluronan-binding protein 2 | 0.49932 | 0.70875 | 0.62677 | ↓ | -0.674 | 0.68356 | 0.86703 | 0.97224 | ↓ | -0.041 | 0.85205 | 0.92866 | 0.69741 | ↓ | -0.52 |
| SERPIN domain-containing protein | 0.20128 | 0.4582 | 0.71985 | ↓ | -0.474 | 0.37702 | 0.63945 | 0.8145 | ↓ | -0.296 | 0.51605 | 0.78865 | 0.8739 | ↓ | -0.194 |
| Complement component C9 | 0.81131 | 0.88833 | 1.016 | ↑ | 0.0229 | 0.2815 | 0.59084 | 1.3607 | ↑ | 0.4443 | 0.46628 | 0.76433 | 1.1125 | ↑ | 0.1539 |
| Uncharacterized protein | 0.25128 | 0.52375 | 1.5264 | ↑ | 0.6102 | 0.08015 | 0.33113 | 1.6035 | ↑ | 0.6812 | 0.070325 | 0.34284 | 1.6725 | ↑ | 0.742 |
| Uncharacterized protein | 0.30029 | 0.55267 | 0.55198 | ↓ | -0.857 | 0.19667 | 0.53106 | 0.42591 | ↓ | -1.231 | 0.12231 | 0.42486 | 0.35622 | ↓ | -1.489 |
| Haptoglobin | 0.14778 | 0.41236 | 1.8619 | ↑ | 0.8968 | 0.084624 | 0.33273 | 2.0911 | ↑ | 1.0642 | 0.34513 | 0.6916 | 1.7184 | ↑ | 0.7811 |
| Joining chain of multimeric IgA and IgM | 0.32932 | 0.56758 | 0.56713 | ↓ | -0.818 | 0.14473 | 0.4317 | 0.26723 | ↓ | -1.904 | 0.058738 | 0.31812 | 0.19654 | ↓ | -2.347 |
| Serpin peptidase inhibitor, clade D (Heparin cofactor), member 1 | 0.79577 | 0.87687 | 0.96751 | ↓ | -0.048 | 0.68985 | 0.86703 | 0.8331 | ↓ | -0.263 | 0.39543 | 0.72777 | 1.1508 | ↑ | 0.2026 |
| Complement C3 alpha chain | 0.075316 | 0.32527 | 0.72339 | ↓ | -0.467 | 0.62287 | 0.84848 | 0.98796 | ↓ | -0.017 | 0.33517 | 0.6903 | 0.85335 | ↓ | -0.229 |
| Uncharacterized protein | 0.55692 | 0.72442 | 1.571 | ↑ | 0.6517 | 0.90478 | 0.95592 | 1.2377 | ↑ | 0.3077 | 0.58023 | 0.83547 | 0.84612 | ↓ | -0.241 |
| Coagulation factor XIII B chain | 0.015418 | 0.14819 | 0.56383 | ↓ | -0.827 | 0.2219 | 0.54755 | 0.77333 | ↓ | -0.371 | 0.04433 | 0.31812 | 0.60519 | ↓ | -0.725 |
| C3-beta-c | 0.19861 | 0.4582 | 0.43316 | ↓ | -1.207 | 0.75136 | 0.89828 | 1.0205 | ↑ | 0.0293 | 0.86851 | 0.92866 | 1.0155 | ↑ | 0.0222 |
| Uncharacterized protein | 0.91295 | 0.95721 | 0.89801 | ↓ | -0.155 | 0.56235 | 0.79508 | 0.83725 | ↓ | -0.256 | 0.97337 | 0.98475 | 0.74457 | ↓ | -0.426 |
| Keratin 9 | 0.13375 | 0.38564 | 0.62404 | ↓ | -0.68 | 0.065727 | 0.33113 | 0.49393 | ↓ | -1.018 | 0.16399 | 0.50479 | 0.64648 | ↓ | -0.629 |
| Soluble type II IL-1 receptor | 0.99372 | 0.99372 | 0.94123 | ↓ | -0.087 | 0.69162 | 0.86703 | 0.8982 | ↓ | -0.155 | 0.911 | 0.93811 | 1.0044 | ↑ | 0.0064 |
| Epididymis secretory sperm binding protein Li 163pA | 0.54994 | 0.72442 | 1.085 | ↑ | 0.1177 | 0.92645 | 0.95592 | 0.95738 | ↓ | -0.063 | 0.53236 | 0.7981 | 1.1051 | ↑ | 0.1442 |
| Complement C5 | 0.58923 | 0.74659 | 0.9218 | ↓ | -0.117 | 0.91241 | 0.95592 | 1.0074 | ↑ | 0.0107 | 0.61139 | 0.83547 | 0.92589 | ↓ | -0.111 |
| C3-beta-c | 0.62148 | 0.76797 | 0.91538 | ↓ | -0.128 | 0.29694 | 0.5913 | 0.83882 | ↓ | -0.254 | 0.8945 | 0.93811 | 1.0157 | ↑ | 0.0224 |
| C1q domain-containing protein | 0.64084 | 0.78628 | 1.464 | ↑ | 0.55 | 0.23318 | 0.55144 | 0.26726 | ↓ | -1.904 | 0.28346 | 0.65385 | 0.48302 | ↓ | -1.05 |
| Gc-globulin | 0.25824 | 0.52636 | 0.80678 | ↓ | -0.31 | 0.19111 | 0.5248 | 0.77318 | ↓ | -0.371 | 0.5769 | 0.83547 | 0.88958 | ↓ | -0.169 |
| Apolipoprotein B | 0.298 | 0.55267 | 0.8344 | ↓ | -0.261 | 0.12981 | 0.41587 | 0.6546 | ↓ | -0.611 | 0.59116 | 0.83547 | 1.3269 | ↑ | 0.408 |
| Mannan-binding lectin serine protease 1 | 0.086242 | 0.32527 | 0.81018 | ↓ | -0.304 | 0.78544 | 0.90587 | 0.968 | ↓ | -0.047 | 0.037178 | 0.29236 | 0.77485 | ↓ | -0.368 |
| Complement component C6 | 0.5373 | 0.72442 | 0.91376 | ↓ | -0.13 | 0.59835 | 0.82155 | 1.1161 | ↑ | 0.1584 | 0.66527 | 0.84217 | 1.0779 | ↑ | 0.1082 |
| Clusterin | 0.18083 | 0.45339 | 0.78743 | ↓ | -0.345 | 0.069969 | 0.33113 | 0.60877 | ↓ | -0.716 | 0.13132 | 0.42486 | 0.67022 | ↓ | -0.577 |
| Albumin | 0.068335 | 0.31951 | 0.58298 | ↓ | -0.778 | 0.015669 | 0.18072 | 0.44763 | ↓ | -1.16 | 6.29E-05 | 0.007703 | 0.26626 | ↓ | -1.909 |
| Actin-depolymerizing factor | 0.087211 | 0.32527 | 1.4836 | ↑ | 0.5691 | 0.79399 | 0.90967 | 1.0764 | ↑ | 0.1063 | 0.75077 | 0.89403 | 1.1908 | ↑ | 0.252 |
| Vitronectin | 0.032793 | 0.2182 | 0.56827 | ↓ | -0.815 | 0.14078 | 0.4317 | 0.67069 | ↓ | -0.576 | 0.077577 | 0.36272 | 0.64766 | ↓ | -0.627 |
| C4a anaphylatoxin | 0.9899 | 0.99372 | 0.99139 | ↓ | -0.012 | 0.9204 | 0.95592 | 1.0238 | ↑ | 0.034 | 0.11952 | 0.42486 | 1.2924 | ↑ | 0.3701 |
| Insulin-like growth factor I | 0.27648 | 0.53744 | 0.45726 | ↓ | -1.129 | 0.077612 | 0.33113 | 0.24526 | ↓ | -2.028 | 0.014197 | 0.18893 | 0.14524 | ↓ | -2.784 |
| Hepatocyte growth factor activator | 0.28996 | 0.54526 | 0.82247 | ↓ | -0.282 | 0.14471 | 0.4317 | 0.75301 | ↓ | -0.409 | 0.44549 | 0.74824 | 0.8749 | ↓ | -0.193 |
| Carbonic anhydrase | 0.36565 | 0.60385 | 1.4075 | ↑ | 0.4931 | 0.95377 | 0.95932 | 1.6126 | ↑ | 0.6894 | 0.65882 | 0.84217 | 0.9387 | ↓ | -0.091 |
| Beta actin variant | 0.1143 | 0.35954 | 0.64644 | ↓ | -0.629 | 0.41805 | 0.67514 | 0.8061 | ↓ | -0.311 | 0.84386 | 0.92866 | 0.91826 | ↓ | -0.123 |
| Alpha-1-microglobulin | 0.002351 | 0.036978 | 0.62529 | ↓ | -0.677 | 0.14129 | 0.4317 | 0.77282 | ↓ | -0.372 | 0.060683 | 0.31812 | 0.76425 | ↓ | -0.388 |
| Gc-globulin | 0.10054 | 0.34104 | 0.75628 | ↓ | -0.403 | 0.2858 | 0.59084 | 0.82978 | ↓ | -0.269 | 0.50744 | 0.78668 | 0.89609 | ↓ | -0.158 |
| Mannose receptor C type 2 | 0.42988 | 0.65237 | 0.72136 | ↓ | -0.471 | 0.95367 | 0.95932 | 0.97073 | ↓ | -0.043 | 0.42087 | 0.73707 | 0.82533 | ↓ | -0.277 |
| Macaca fascicularis brain cDNA clone: QtrA-15786, similar to huma | 0.017279 | 0.15733 | 0.63881 | ↓ | -0.647 | 0.05648 | 0.33113 | 0.69849 | ↓ | -0.518 | 0.004702 | 0.10168 | 0.56092 | ↓ | -0.834 |
| Alpha-1-globin | 0.38917 | 0.60655 | 0.67748 | ↓ | -0.562 | 0.11229 | 0.38089 | 0.26689 | ↓ | -1.906 | 0.29531 | 0.67222 | 0.57722 | ↓ | -0.793 |
| Coagulation factor X | 0.97223 | 0.99372 | 0.95572 | ↓ | -0.065 | 0.32276 | 0.59403 | 0.7821 | ↓ | -0.355 | 0.43447 | 0.73707 | 0.82236 | ↓ | -0.282 |
| Ceruloplasmin | 0.48945 | 0.70562 | 1.0896 | ↑ | 0.1238 | 0.77712 | 0.90587 | 1.0879 | ↑ | 0.1215 | 0.43274 | 0.73707 | 1.2223 | ↑ | 0.2896 |
| IgGFc-binding protein | 0.16123 | 0.42577 | 0.62215 | ↓ | -0.685 | 0.54426 | 0.77816 | 1.3898 | ↑ | 0.4749 | 0.41289 | 0.73707 | 0.86265 | ↓ | -0.213 |
| Profilin | 0.51597 | 0.72404 | 1.9954 | ↑ | 0.9967 | 0.6681 | 0.86703 | 1.068 | ↑ | 0.095 | 0.30353 | 0.67322 | 1.8899 | ↑ | 0.9183 |
| Serpin family G member 1 | 0.74748 | 0.83428 | 0.95343 | ↓ | -0.069 | 0.67559 | 0.86703 | 0.79799 | ↓ | -0.326 | 0.41308 | 0.73707 | 1.1303 | ↑ | 0.1767 |
| HCG40889, isoform CRA_b | 0.82573 | 0.89411 | 0.97673 | ↓ | -0.034 | 0.53059 | 0.76493 | 0.83598 | ↓ | -0.258 | 0.72702 | 0.87954 | 0.88855 | ↓ | -0.17 |
| C4a anaphylatoxin | 0.21401 | 0.47466 | 0.75524 | ↓ | -0.405 | 0.39145 | 0.65116 | 0.84858 | ↓ | -0.237 | 0.21667 | 0.58568 | 1.2277 | ↑ | 0.296 |
| Plasminogen | 0.87602 | 0.92976 | 1.0286 | ↑ | 0.0407 | 0.41439 | 0.67514 | 0.97807 | ↓ | -0.032 | 0.81506 | 0.92866 | 0.99099 | ↓ | -0.013 |
| Protein S100 | 0.020998 | 0.18163 | 0.22762 | ↓ | -2.135 | 0.057422 | 0.33113 | 0.23657 | ↓ | -2.08 | 0.057284 | 0.31812 | 0.29927 | ↓ | -1.741 |
| Reversion inducing cysteine rich protein with kazal motifs | 0.27607 | 0.53744 | 0.57574 | ↓ | -0.797 | 0.56529 | 0.79508 | 1.0116 | ↑ | 0.0167 | 0.47071 | 0.76433 | 1.8417 | ↑ | 0.881 |
| Complement component C9 | 0.56694 | 0.73194 | 1.1056 | ↑ | 0.1448 | 0.20818 | 0.53753 | 1.2321 | ↑ | 0.3012 | 0.59509 | 0.83547 | 1.1018 | ↑ | 0.1399 |

| | | | | | | | | | | | | |
|---|---|---|---|---|---|---|---|---|---|---|---|---|
| Glyceraldehyde-3-phosphate dehydrogenase | 0.83471 | 0.89692 | 2.2318 ↑ | 1.1582 | 0.50825 | 0.75151 | 1.0476 ↑ | 0.067 | 0.86703 | 0.92866 | 2.5605 ↑ | 1.3564 |
| Complement subcomponent C1r | 0.23171 | 0.50106 | 0.86903 ↓ | -0.203 | 0.021715 | 0.22098 | 0.67717 ↓ | -0.562 | 0.12782 | 0.42486 | 0.82579 ↓ | -0.276 |
| Fibrinogen beta chain | 0.5477 | 0.72442 | 0.93029 ↓ | -0.104 | 0.7508 | 0.89828 | 1.0397 ↑ | 0.0561 | 0.69746 | 0.86363 | 1.2776 ↑ | 0.3535 |
| Cation-independent mannose-6-phosphate receptor | 0.31151 | 0.5555 | 1.3191 ↑ | 0.3996 | 0.89808 | 0.95592 | 0.78762 ↓ | -0.344 | 0.8773 | 0.93113 | 0.77666 ↓ | -0.365 |
| Complement C1r subcomponent-like protein | 0.39737 | 0.60836 | 0.77237 ↓ | -0.373 | 0.50559 | 0.75151 | 0.81985 ↓ | -0.287 | 0.90326 | 0.93811 | 0.95662 ↓ | -0.064 |
| Haptoglobin | 0.28853 | 0.54526 | 2.1282 ↑ | 1.0896 | 0.11015 | 0.38089 | 2.3548 ↑ | 1.2356 | 0.74813 | 0.89403 | 1.774 ↑ | 0.827 |
| Apolipoprotein A-II | 0.17525 | 0.44586 | 0.4275 ↓ | -1.226 | 0.026698 | 0.23383 | 0.20422 ↓ | -2.292 | 0.007582 | 0.11126 | 0.15225 ↓ | -2.716 |
| Alpha-2-HS-glycoprotein | 0.51896 | 0.72404 | 0.71553 ↓ | -0.483 | 0.35407 | 0.62278 | 0.65921 ↓ | -0.601 | 0.30155 | 0.67322 | 0.70601 ↓ | -0.502 |
| Apolipoprotein A-I | 0.20394 | 0.4582 | 0.53773 ↓ | -0.895 | 0.018481 | 0.19983 | 0.13151 ↓ | -2.927 | 0.024418 | 0.24549 | 0.14822 ↓ | -2.754 |
| Hemoglobin subunit beta | 0.99252 | 0.99372 | 1.0986 ↑ | 0.1357 | 0.70721 | 0.8739 | 1.3917 ↑ | 0.4769 | 0.32256 | 0.6784 | 0.50955 ↓ | -0.973 |
| Angiotensin-converting enzyme | 0.0997 | 0.34104 | 0.61169 ↓ | -0.709 | 0.68539 | 0.86703 | 0.88708 ↓ | -0.173 | 0.62003 | 0.83547 | 0.8762 ↓ | -0.191 |
| Complement C8 gamma chain | 0.37308 | 0.60572 | 0.92745 ↓ | -0.109 | 0.44423 | 0.69187 | 0.937 ↓ | -0.094 | 0.77577 | 0.91298 | 0.97878 ↓ | -0.031 |
| Antithrombin-III | 0.60902 | 0.75897 | 0.72619 ↓ | -0.462 | 0.48834 | 0.73463 | 0.75291 ↓ | -0.409 | 0.24618 | 0.60843 | 0.54139 ↓ | -0.885 |
| C1q domain-containing protein | 0.20045 | 0.4582 | 0.6901 ↓ | -0.535 | 0.18991 | 0.5248 | 0.67818 ↓ | -0.56 | 0.12246 | 0.42486 | 0.64147 ↓ | -0.641 |
| Epididymis secretory protein Li 37 | 0.004 | 0.053233 | 4.5552 ↑ | 2.1875 | 0.12073 | 0.40165 | 2.1308 ↑ | 1.0914 | 0.27138 | 0.64163 | 0.41648 ↓ | -1.264 |
| Fibrinogen gamma chain | 0.27063 | 0.53744 | 1.3361 ↑ | 0.418 | 0.21891 | 0.54755 | 1.3145 ↑ | 0.3946 | 0.94034 | 0.9626 | 1.1577 ↑ | 0.2112 |
| Apolipoprotein C-I, basic form | 0.31468 | 0.5555 | 0.4724 ↓ | -1.082 | 0.23864 | 0.55144 | 0.4455 ↓ | -1.167 | 0.19707 | 0.56822 | 0.40955 ↓ | -1.288 |
| Uncharacterized protein | 0.69919 | 0.80639 | 0.97085 ↓ | -0.043 | 0.8318 | 0.9284 | 0.99672 ↓ | -0.005 | 0.86417 | 0.92866 | 0.99755 ↓ | -0.004 |
| Complement C4-B | 0.76137 | 0.84434 | 1.0487 ↑ | 0.0686 | 0.35639 | 0.62278 | 1.209 ↑ | 0.2738 | 0.36894 | 0.71558 | 1.2062 ↑ | 0.2705 |
| SERPIN domain-containing protein | 0.094866 | 0.33493 | 0.57762 ↓ | -0.792 | 0.080315 | 0.33113 | 0.55101 ↓ | -0.86 | 0.007385 | 0.11126 | 0.40067 ↓ | -1.32 |
| Lumican | 7.47E-05 | 0.002998 | 0.68605 ↓ | -0.544 | 0.000573 | 0.024761 | 0.58848 ↓ | -0.765 | 0.001028 | 0.027068 | 0.62077 ↓ | -0.688 |
| Cadherin-13 | 0.74739 | 0.83428 | 0.97229 ↓ | -0.041 | 0.65865 | 0.86703 | 2.5305 ↑ | 1.3394 | 0.70887 | 0.86363 | 1.1395 ↑ | 0.1884 |
| Angiotensin 1-10 | 0.38164 | 0.60572 | 1.1346 ↑ | 0.1822 | 0.65885 | 0.86703 | 1.088 ↑ | 0.1217 | 0.37227 | 0.71558 | 1.224 ↑ | 0.2917 |
| Keratin, type II cytoskeletal 2 epidermal | 0.37941 | 0.60572 | 0.74494 ↓ | -0.425 | 0.27438 | 0.59084 | 0.70161 ↓ | -0.511 | 0.099478 | 0.41085 | 0.5995 ↓ | -0.738 |
| Fibulin-1 | 0.1258 | 0.3818 | 0.68587 ↓ | -0.544 | 0.31189 | 0.59403 | 0.81729 ↓ | -0.291 | 0.016089 | 0.19881 | 0.54416 ↓ | -0.878 |
| Trypsin | 0.013012 | 0.13242 | 0.38161 ↓ | -1.39 | 0.28298 | 0.59084 | 0.67023 ↓ | -0.577 | 0.05983 | 0.31812 | 0.47974 ↓ | -1.06 |
| 1,4-beta-N-acetylmuramidase C | 0.33136 | 0.56758 | 0.46232 ↓ | -1.113 | 0.64402 | 0.86703 | 0.53171 ↓ | -0.911 | 0.022542 | 0.24374 | 0.20497 ↓ | -2.287 |
| Macaca fascicularis brain cDNA clone: QorA-13734, similar to huma | 0.22209 | 0.48636 | 1.3024 ↑ | 0.3812 | 0.69705 | 0.86755 | 0.98713 ↓ | -0.019 | 0.51969 | 0.78865 | 0.89589 ↓ | -0.159 |
| Testicular tissue protein Li 61 | 0.25861 | 0.52636 | 0.855 ↓ | -0.226 | 0.31379 | 0.59403 | 0.85966 ↓ | -0.218 | 0.086385 | 0.39328 | 0.75763 ↓ | -0.4 |
| Uncharacterized protein | 0.65344 | 0.78885 | 0.93017 ↓ | -0.104 | 0.28983 | 0.59084 | 0.8448 ↓ | -0.243 | 0.27445 | 0.64163 | 0.84113 ↓ | -0.25 |
| Vitellogenin domain-containing protein | 0.59123 | 0.74659 | 0.88989 ↓ | -0.168 | 0.082657 | 0.33255 | 0.67953 ↓ | -0.557 | 0.96711 | 0.98418 | 1.026 ↑ | 0.037 |
| Uncharacterized protein | 0.3889 | 0.60655 | 1.19 ↑ | 0.251 | 0.37198 | 0.63715 | 0.85217 ↓ | -0.231 | 0.70857 | 0.86363 | 0.96509 ↓ | -0.051 |
| Serpin family A member 10 | 0.20242 | 0.4582 | 0.69464 ↓ | -0.526 | 0.86128 | 0.93712 | 0.90364 ↓ | -0.146 | 0.86916 | 0.92866 | 1.1302 ↑ | 0.1766 |
| Complement component 8 subunit beta | 0.96503 | 0.99372 | 0.88831 ↓ | -0.171 | 0.067059 | 0.33113 | 3.0904 ↑ | 1.6278 | 0.8498 | 0.92866 | 1.0063 ↑ | 0.009 |
| Collagen type VI alpha 3 chain | 0.68992 | 0.80318 | 1.0926 ↑ | 0.1277 | 0.88143 | 0.9493 | 0.83843 ↓ | -0.254 | 0.38128 | 0.71925 | 1.1128 ↑ | 0.1542 |
| Glutathione peroxidase | 0.39528 | 0.60836 | 0.82112 ↓ | -0.284 | 0.48553 | 0.73463 | 0.89278 ↓ | -0.164 | 0.017318 | 0.19973 | 0.60302 ↓ | -0.73 |
| Mannan binding lectin serine peptidase 2 | 0.89832 | 0.94762 | 0.954 ↓ | -0.068 | 0.97681 | 0.97681 | 0.97685 ↓ | -0.034 | 0.47447 | 0.76433 | 0.74167 ↓ | -0.431 |
| Protein S100 | 0.35991 | 0.60385 | 0.63675 ↓ | -0.651 | 0.3409 | 0.608 | 0.60477 ↓ | -0.726 | 0.43457 | 0.73707 | 0.64561 ↓ | -0.631 |
| Bradykinin | 0.24505 | 0.51937 | 0.7688 ↓ | -0.379 | 0.59692 | 0.82155 | 0.87462 ↓ | -0.193 | 0.11401 | 0.42486 | 0.63975 ↓ | -0.644 |
| Protein C receptor | 0.69176 | 0.80318 | 0.98016 ↓ | -0.029 | 0.039075 | 0.28303 | 0.78931 ↓ | -0.341 | 0.23226 | 0.59971 | 0.89675 ↓ | -0.157 |
| Protein-tyrosine-phosphatase | 0.012117 | 0.13102 | 1.4074 ↑ | 0.4931 | 0.052101 | 0.33113 | 1.5264 ↑ | 0.6102 | 0.048236 | 0.31812 | 1.5977 ↑ | 0.676 |
| Uncharacterized protein | 0.65268 | 0.78885 | 0.7229 ↓ | -0.468 | 0.80262 | 0.91351 | 0.76721 ↓ | -0.382 | 0.59945 | 0.83547 | 1.1334 ↑ | 0.1806 |
| Filamin A | 0.037814 | 0.22558 | 1.6251 ↑ | 0.7006 | 0.095961 | 0.36419 | 2.5208 ↑ | 1.3339 | 0.49948 | 0.78587 | 1.4815 ↑ | 0.5671 |
| Inter-alpha-trypsin inhibitor heavy chain 2 | 0.30597 | 0.5555 | 0.72109 ↓ | -0.472 | 0.26948 | 0.59014 | 0.74376 ↓ | -0.427 | 0.56777 | 0.8324 | 0.82109 ↓ | -0.284 |
| Fibrinogen alpha chain | 0.001296 | 0.024902 | 0.48812 ↓ | -1.035 | 0.32277 | 0.59403 | 0.8426 ↓ | -0.247 | 0.052728 | 0.31812 | 0.65785 ↓ | -0.604 |
| Complement C1s | 0.11971 | 0.36981 | 0.85098 ↓ | -0.233 | 0.82071 | 0.92196 | 1.0342 ↑ | 0.0486 | 0.064248 | 0.32691 | 0.81541 ↓ | -0.294 |
| Mannosyl-oligosaccharide 1,2-alpha-mannosidase IA | 0.7392 | 0.83428 | 1.3162 ↑ | 0.3964 | 0.18143 | 0.52311 | 2.2036 ↑ | 1.1399 | 0.81703 | 0.92866 | 1.7749 ↑ | 0.8278 |
| Apolipoprotein B (Including Ag(X) antigen) | 0.54712 | 0.72442 | 0.81247 ↓ | -0.3 | 0.74805 | 0.89828 | 1.0207 ↑ | 0.0295 | 0.21485 | 0.58568 | 0.62527 ↓ | -0.677 |
| Complement component C8 alpha chain | 0.96601 | 0.99372 | 0.93948 ↓ | -0.09 | 0.36294 | 0.62789 | 0.75748 ↓ | -0.401 | 0.67561 | 0.84696 | 0.86011 ↓ | -0.217 |
| cDNA FLJ44930 fis, clone BRAMY3015549, highly similar to Neural c | 0.67556 | 0.80049 | 0.64582 ↓ | -0.631 | 0.78137 | 0.90587 | 0.68426 ↓ | -0.547 | 0.7545 | 0.89403 | 0.71597 ↓ | -0.482 |
| Metavinculin | 0.24618 | 0.51937 | 0.51032 ↓ | -0.971 | 0.8602 | 0.93712 | 0.82141 ↓ | -0.284 | 0.66692 | 0.84217 | 0.67365 ↓ | -0.57 |
| ITIH4 protein | 0.66872 | 0.79785 | 1.0037 ↑ | 0.0053 | 0.74338 | 0.89828 | 1.0887 ↑ | 0.1226 | 0.97913 | 0.98483 | 1.2908 ↑ | 0.3682 |
| Apolipoprotein C-II | 0.2853 | 0.54526 | 0.65298 ↓ | -0.615 | 0.10564 | 0.37297 | 0.44475 ↓ | -1.169 | 0.033101 | 0.28632 | 0.33641 ↓ | -1.572 |

| | | | | | | | | | | | | |
|---|---|---|---|---|---|---|---|---|---|---|---|---|
| Prothrombin | 0.16045 | 0.42577 | 0.56251 ↓ | -0.83 | 0.81156 | 0.91765 | 0.92994 ↓ | -0.105 | 0.17616 | 0.52544 | 0.57099 ↓ | -0.808 |
| Alpha-1-antitrypsin | 0.98694 | 0.99372 | 0.79009 ↓ | -0.34 | 0.59513 | 0.82155 | 0.7407 ↓ | -0.433 | 0.38326 | 0.71925 | 0.50841 ↓ | -0.976 |
| Complement C1q subcomponent subunit B | 0.65661 | 0.78885 | 0.86183 ↓ | -0.215 | 0.042536 | 0.28303 | 0.54327 ↓ | -0.88 | 0.31243 | 0.6778 | 0.73853 ↓ | -0.437 |
| SEMA4D variant protein | 0.025137 | 0.18404 | 2.0802 ↑ | 1.0567 | 0.2903 | 0.59084 | 1.417 ↑ | 0.5028 | 0.62182 | 0.83547 | 1.2519 ↑ | 0.3241 |
| CP protein | 0.085727 | 0.32527 | 0.43873 ↓ | -1.189 | 0.9101 | 0.95592 | 1.7987 ↑ | 0.8469 | 0.55938 | 0.82712 | 0.76144 ↓ | -0.393 |
| Attractin | 0.050601 | 0.26527 | 0.76256 ↓ | -0.391 | 0.23906 | 0.55144 | 0.83528 ↓ | -0.26 | 0.035778 | 0.29236 | 0.67684 ↓ | -0.563 |
| 15S Mg(2+)-ATPase p97 subunit | 0.68948 | 0.80318 | 0.83708 ↓ | -0.257 | 0.76967 | 0.9058 | 1.0845 ↑ | 0.117 | 0.80579 | 0.92866 | 1.0376 ↑ | 0.0533 |
| CST6 protein | 0.022083 | 0.18192 | 0.67348 ↓ | -0.57 | 0.3325 | 0.6055 | 0.86253 ↓ | -0.213 | 0.20684 | 0.58568 | 0.65867 ↓ | -0.602 |
| Monocyte differentiation antigen CD14 | 0.26929 | 0.53744 | 0.83738 ↓ | -0.256 | 0.30078 | 0.5913 | 0.85766 ↓ | -0.222 | 0.1259 | 0.42486 | 0.80622 ↓ | -0.311 |
| Uncharacterized protein | 0.60981 | 0.75897 | 0.86733 ↓ | -0.205 | 0.84154 | 0.92895 | 1.6916 ↑ | 0.7584 | 0.47716 | 0.76433 | 2.2993 ↑ | 1.2012 |
| Immunoglobulin heavy constant mu | 0.53844 | 0.72442 | 0.99178 ↓ | -0.012 | 0.94955 | 0.95932 | 1.1137 ↑ | 0.1554 | 0.16632 | 0.50479 | 0.5485 ↓ | -0.866 |
| Apolipoprotein A4 | 0.49981 | 0.70875 | 0.64296 ↓ | -0.637 | 0.24676 | 0.55492 | 0.54071 ↓ | -0.887 | 0.027603 | 0.25133 | 0.24627 ↓ | -2.022 |
| Ficolin (Collagen/fibrinogen domain containing) 3 (Hakata antigen), | 0.32039 | 0.55987 | 0.56011 ↓ | -0.836 | 0.31878 | 0.59403 | 0.55721 ↓ | -0.844 | 0.12809 | 0.42486 | 0.40407 ↓ | -1.307 |
| Receptor protein-tyrosine kinase | 0.035417 | 0.21883 | 0.65608 ↓ | -0.608 | 0.4413 | 0.69187 | 0.87849 ↓ | -0.187 | 0.13262 | 0.42486 | 0.74445 ↓ | -0.426 |
| Insulin-like growth factor-binding protein 3 | 0.12873 | 0.38396 | 0.63977 ↓ | -0.644 | 0.2026 | 0.53106 | 0.66083 ↓ | -0.598 | 0.11614 | 0.42486 | 0.64324 ↓ | -0.637 |
| Uncharacterized protein DKFZp686I15196 | 0.44344 | 0.65864 | 0.663 ↓ | -0.593 | 0.43606 | 0.69187 | 0.48441 ↓ | -1.046 | 0.31343 | 0.6778 | 0.4574 ↓ | -1.129 |
| cDNA, FLJ93914, highly similar to Homo sapiens histidine-rich glyco | 0.74566 | 0.83428 | 0.87163 ↓ | -0.198 | 0.22472 | 0.54755 | 0.67386 ↓ | -0.569 | 0.90571 | 0.93811 | 0.90161 ↓ | -0.149 |
| Coagulation factor XII | 0.44544 | 0.65864 | 0.64439 ↓ | -0.634 | 0.3161 | 0.59403 | 0.92983 ↓ | -0.105 | 0.24477 | 0.60843 | 0.55184 ↓ | -0.858 |
| Protein S100-A7 | 0.16489 | 0.42577 | 1.1312 ↑ | 0.1778 | 0.68581 | 0.86703 | 0.77659 ↓ | -0.365 | 0.65536 | 0.84217 | 0.95708 ↓ | -0.063 |
| Insulin-like growth factor binding protein, acid labile subunit | 0.15291 | 0.4199 | 0.74283 ↓ | -0.429 | 0.75809 | 0.89828 | 0.94964 ↓ | -0.075 | 0.12255 | 0.42486 | 0.72987 ↓ | -0.454 |
| Carboxylic ester hydrolase | 0.042221 | 0.23562 | 0.68294 ↓ | -0.55 | 0.042086 | 0.28303 | 0.70427 ↓ | -0.506 | 0.071341 | 0.34284 | 0.70147 ↓ | -0.512 |

## 8 Gy

| | SD 1 vs. Pre | | | | SD 2 vs. Pre | | | | SD 6 vs. Pre | | | |
|---|---|---|---|---|---|---|---|---|---|---|---|---|
| Protein Name | *p-value* | FDR | FC | LOG2FC | *p-value* | FDR | FC | LOG2FC | *p-value* | FDR | FC | LOG2FC |
| Serpin family A member 9 | 5.9E-06 | 0.001021 | 3.8626 ↑ | 1.9496 | 1.04E-05 | 0.000899 | 4.1026 ↑ | 2.0365 | 0.000566 | 0.009785 | 2.4403 ↑ | 1.2871 |
| C-reactive protein | 1.28E-05 | 0.001108 | 44.624 ↑ | 5.4797 | 0.041907 | 0.1652 | 7.5118 ↑ | 2.9092 | 0.44614 | 0.74214 | 2.1151 ↑ | 1.0807 |
| cDNA FLJ35730 fis, clone TESTI2003131, highly similar to ALPHA-1-/ | 0.00268 | 0.057948 | 3.7541 ↑ | 1.9085 | 0.002659 | 0.041813 | 3.3142 ↑ | 1.7287 | 0.000367 | 0.007931 | 2.7281 ↑ | 1.4479 |
| Fibronectin | 0.026657 | 0.24868 | 0.52019 ↓ | -0.943 | 0.088422 | 0.24151 | 0.61533 ↓ | -0.701 | 0.047183 | 0.20751 | 0.59504 ↓ | -0.749 |
| Retinol-binding protein | 0.002411 | 0.057948 | 0.48931 ↓ | -1.031 | 0.010287 | 0.088558 | 0.63591 ↓ | -0.653 | 0.23534 | 0.53644 | 0.83915 ↓ | -0.253 |
| Clusterin | 0.033273 | 0.24868 | 0.43738 ↓ | -1.193 | 0.073386 | 0.21366 | 0.44864 ↓ | -1.156 | 0.43176 | 0.74214 | 0.81322 ↓ | -0.298 |
| Apolipoprotein C-III | 0.000635 | 0.020674 | 0.27821 ↓ | -1.846 | 0.000143 | 0.007454 | 0.22092 ↓ | -2.178 | 0.000138 | 0.004763 | 0.25953 ↓ | -1.946 |
| Testis cDNA clone: QtsA-10685, similar to human I factor (complem | 0.30903 | 0.6694 | 0.87683 ↓ | -0.19 | 0.029919 | 0.14378 | 0.7145 ↓ | -0.485 | 0.87146 | 0.94374 | 0.98244 ↓ | -0.026 |
| Protein C, inactivator of coagulation factors Va and VIIIa | 0.000338 | 0.014627 | 0.55681 ↓ | -0.845 | 0.000504 | 0.017047 | 0.55021 ↓ | -0.862 | 0.014916 | 0.12902 | 0.75147 ↓ | -0.412 |
| Endonuclease domain containing 1 | 0.0737 | 0.41129 | 3.2502 ↑ | 1.7005 | 0.12544 | 0.28554 | 2.2179 ↑ | 1.1492 | 0.013732 | 0.12503 | 4.4772 ↑ | 2.1626 |
| Macaca fascicularis brain cDNA clone: QtrA-15720, similar to huma | 0.032062 | 0.24868 | 0.7037 ↓ | -0.507 | 0.23565 | 0.43267 | 0.83474 ↓ | -0.261 | 0.53436 | 0.81683 | 0.9091 ↓ | -0.137 |
| Apolipoprotein B | 0.58411 | 0.88642 | 0.54007 ↓ | -0.889 | 0.50878 | 0.67707 | 0.39222 ↓ | -1.35 | 0.65847 | 0.88307 | 0.77186 ↓ | -0.374 |
| Leucine-rich alpha-2-glycoprotein | 0.006098 | 0.097991 | 2.2104 ↑ | 1.1443 | 0.061731 | 0.19688 | 1.3838 ↑ | 0.4687 | 0.26179 | 0.5608 | 1.2283 ↑ | 0.2966 |
| Lambda-chain (AA -20 to 215) | 0.31081 | 0.6694 | 0.89034 ↓ | -0.168 | 0.013203 | 0.090595 | 0.68164 ↓ | -0.553 | 0.022216 | 0.16014 | 0.82671 ↓ | -0.275 |
| Inter-alpha-trypsin inhibitor heavy chain 1 | 0.019985 | 0.22701 | 0.30203 ↓ | -1.727 | 0.014139 | 0.090595 | 0.29379 ↓ | -1.767 | 0.18584 | 0.50153 | 0.48186 ↓ | -1.053 |
| Apolipoprotein H | 0.003077 | 0.059142 | 0.75061 ↓ | -0.414 | 0.065665 | 0.20142 | 0.78818 ↓ | -0.343 | 0.011687 | 0.11893 | 0.7567 ↓ | -0.402 |
| Testis cDNA, clone: QtsA-17169, similar to human complement con | 0.2477 | 0.6437 | 0.77647 ↓ | -0.365 | 0.29937 | 0.49991 | 0.88059 ↓ | -0.183 | 0.70499 | 0.89024 | 1.0371 ↑ | 0.0525 |
| Cathepsin D | 0.28701 | 0.6694 | 0.8098 ↓ | -0.304 | 0.006209 | 0.071634 | 0.30027 ↓ | -1.736 | 0.000921 | 0.014492 | 0.38609 ↓ | -1.373 |
| Histidine rich glycoprotein | 0.91112 | 0.94448 | 0.87502 ↓ | -0.193 | 0.58135 | 0.71838 | 0.80309 ↓ | -0.316 | 0.11678 | 0.35443 | 0.53077 ↓ | -0.914 |
| Inter-alpha (Globulin) inhibitor H2 | 0.12558 | 0.49374 | 0.77784 ↓ | -0.362 | 0.029814 | 0.14378 | 0.70667 ↓ | -0.501 | 0.56055 | 0.81683 | 0.90441 ↓ | -0.145 |
| Lipopolysaccharide-binding protein | 0.19427 | 0.56942 | 2.3127 ↑ | 1.2096 | 0.62645 | 0.73725 | 0.91909 ↓ | -0.122 | 0.13372 | 0.37354 | 2.4944 ↑ | 1.3187 |
| Inter-alpha-trypsin inhibitor heavy chain 2 | 0.058421 | 0.36429 | 0.73586 ↓ | -0.443 | 0.16571 | 0.3454 | 0.80637 ↓ | -0.31 | 0.40671 | 0.71926 | 0.8782 ↓ | -0.187 |
| C-X-C motif chemokine | 0.020995 | 0.22701 | 0.51344 ↓ | -0.962 | 0.021995 | 0.12275 | 0.51327 ↓ | -0.962 | 0.013247 | 0.12503 | 0.51527 ↓ | -0.957 |
| Hexosyltransferase | 0.15831 | 0.52453 | 0.53888 ↓ | -0.892 | 0.97694 | 0.98262 | 0.81833 ↓ | -0.289 | 0.44192 | 0.74214 | 1.2569 ↑ | 0.3299 |
| Platelet factor 4 | 0.35008 | 0.68398 | 0.43082 ↓ | -1.215 | 0.25225 | 0.4453 | 0.35265 ↓ | -1.504 | 0.64844 | 0.87641 | 0.52167 ↓ | -0.939 |
| Alpha-2-HS-glycoprotein | 0.031317 | 0.24868 | 0.65445 ↓ | -0.612 | 0.038493 | 0.1652 | 0.66362 ↓ | -0.592 | 0.019625 | 0.15432 | 0.64716 ↓ | -0.628 |
| N-acetylmuramoyl-L-alanine amidase | 0.8967 | 0.94448 | 0.86828 ↓ | -0.204 | 0.88395 | 0.92681 | 0.909 ↓ | -0.138 | 0.80226 | 0.93148 | 0.71458 ↓ | -0.485 |
| Vitellogenin domain-containing protein | 0.10373 | 0.44349 | 0.72112 ↓ | -0.472 | 0.066364 | 0.20142 | 0.69236 ↓ | -0.53 | 0.89589 | 0.94553 | 0.98275 ↓ | -0.025 |

| | | | | | | | | | | | | | | | |
|---|---|---|---|---|---|---|---|---|---|---|---|---|---|---|---|
| Apolipoprotein H | 0.33266 | 0.68348 | 0.62491 | ↓ | -0.678 | 0.17759 | 0.36371 | 0.49307 | ↓ | -1.02 | 0.33218 | 0.63151 | 0.68737 | ↓ | -0.541 |
| C3-beta-c | 0.85821 | 0.94448 | 0.95677 | ↓ | -0.064 | 0.82514 | 0.89218 | 1.0084 | ↑ | 0.0121 | 0.79632 | 0.93083 | 1.0193 | ↑ | 0.0275 |
| Vitamin K-dependent protein S | 0.69622 | 0.91424 | 0.87244 | ↓ | -0.197 | 0.09726 | 0.25055 | 0.63762 | ↓ | -0.649 | 0.39799 | 0.71926 | 0.79441 | ↓ | -0.332 |
| Fibronectin | 0.031944 | 0.24868 | 0.41107 | ↓ | -1.283 | 0.020038 | 0.11555 | 0.44952 | ↓ | -1.154 | 0.033792 | 0.18831 | 0.52515 | ↓ | -0.929 |
| Plasminogen | 0.39255 | 0.72636 | 0.88028 | ↓ | -0.184 | 0.13763 | 0.30923 | 0.8021 | ↓ | -0.318 | 0.2346 | 0.53644 | 0.85165 | ↓ | -0.232 |
| Plasma kallikrein | 0.034499 | 0.24868 | 0.85328 | ↓ | -0.229 | 0.12101 | 0.27912 | 0.86424 | ↓ | -0.21 | 0.057555 | 0.22127 | 0.84914 | ↓ | -0.236 |
| Testis cDNA, clone: QtsA-16984, similar to human sex hormone-bin | 0.25301 | 0.6437 | 0.73308 | ↓ | -0.448 | 0.16289 | 0.3451 | 0.78754 | ↓ | -0.345 | 0.75411 | 0.90598 | 0.94797 | ↓ | -0.077 |
| Plasminogen | 0.2649 | 0.66417 | 0.9119 | ↓ | -0.133 | 0.1787 | 0.36371 | 0.69484 | ↓ | -0.525 | 0.034831 | 0.18831 | 0.88775 | ↓ | -0.172 |
| Gc-globulin | 0.000717 | 0.020674 | 0.66542 | ↓ | -0.588 | 0.000591 | 0.017047 | 0.66993 | ↓ | -0.578 | 0.001384 | 0.019949 | 0.69805 | ↓ | -0.519 |
| Alpha-2-macroglobulin | 0.72763 | 0.91424 | 0.85212 | ↓ | -0.231 | 0.44357 | 0.64394 | 0.64438 | ↓ | -0.634 | 0.22068 | 0.53035 | 0.51473 | ↓ | -0.958 |
| Coagulation factor XII-Mie | 0.073146 | 0.41129 | 0.8379 | ↓ | -0.255 | 0.015937 | 0.095074 | 0.75851 | ↓ | -0.399 | 0.10259 | 0.33183 | 0.86524 | ↓ | -0.209 |
| Alpha-2-glycoprotein 1, zinc-binding | 0.000214 | 0.012352 | 0.59392 | ↓ | -0.752 | 4.11E-07 | 7.11E-05 | 0.45175 | ↓ | -1.146 | 3.66E-05 | 0.002111 | 0.55215 | ↓ | -0.857 |
| Fibronectin | 0.006231 | 0.097991 | 0.48142 | ↓ | -1.055 | 0.006515 | 0.071634 | 0.45701 | ↓ | -1.13 | 0.007421 | 0.080237 | 0.41678 | ↓ | -1.263 |
| SERPIN domain-containing protein | 0.63257 | 0.89701 | 1.0386 | ↑ | 0.0546 | 0.30394 | 0.49991 | 1.1085 | ↑ | 0.1486 | 0.13387 | 0.37354 | 0.80714 | ↓ | -0.309 |
| Receptor protein-tyrosine kinase | 0.77651 | 0.91651 | 0.91679 | ↓ | -0.125 | 0.10118 | 0.25055 | 0.39437 | ↓ | -1.342 | 0.27075 | 0.56434 | 0.47616 | ↓ | -1.071 |
| cDNA FLJ51597, highly similar to C4b-binding protein alpha chain | 0.22464 | 0.63229 | 1.9281 | ↑ | 0.9472 | 0.27154 | 0.47451 | 1.7969 | ↑ | 0.8455 | 0.74654 | 0.90598 | 1.4658 | ↑ | 0.5517 |
| Fibronectin | 0.23652 | 0.63229 | 0.3956 | ↓ | -1.338 | 0.22869 | 0.43004 | 0.47058 | ↓ | -1.088 | 0.54459 | 0.81683 | 0.75047 | ↓ | -0.414 |
| Keratin, type I cytoskeletal 10 | 0.45022 | 0.77888 | 0.91061 | ↓ | -0.135 | 0.56649 | 0.7108 | 0.89346 | ↓ | -0.163 | 0.78932 | 0.92893 | 1.1252 | ↑ | 0.1702 |
| Prothrombin | 0.48468 | 0.79103 | 1.1715 | ↑ | 0.2283 | 0.009656 | 0.088558 | 0.39974 | ↓ | -1.323 | 0.29 | 0.58603 | 0.69039 | ↓ | -0.535 |
| C-type lectin domain family 3 member B | 0.1051 | 0.44349 | 0.8186 | ↓ | -0.289 | 0.11268 | 0.26705 | 0.80693 | ↓ | -0.309 | 8.8E-06 | 0.001523 | 0.39958 | ↓ | -1.324 |
| C3/C5 convertase | 0.48042 | 0.79103 | 1.0507 | ↑ | 0.0714 | 0.47209 | 0.64503 | 0.95615 | ↓ | -0.065 | 0.041481 | 0.19781 | 1.2098 | ↑ | 0.2747 |
| Hemopexin | 0.16069 | 0.52453 | 1.2358 | ↑ | 0.3054 | 0.04111 | 0.1652 | 1.2644 | ↑ | 0.3385 | 0.060145 | 0.2262 | 1.3234 | ↑ | 0.4043 |
| SERPIN domain-containing protein | 0.14093 | 0.51875 | 0.64309 | ↓ | -0.637 | 0.044902 | 0.16887 | 0.71747 | ↓ | -0.479 | 0.816 | 0.93489 | 0.9488 | ↓ | -0.076 |
| Hyaluronan-binding protein 2 | 0.88467 | 0.94448 | 1.0506 | ↑ | 0.0712 | 0.47305 | 0.64503 | 1.2114 | ↑ | 0.2767 | 0.22473 | 0.53257 | 0.48959 | ↓ | -1.03 |
| SERPIN domain-containing protein | 0.68847 | 0.91424 | 0.89649 | ↓ | -0.158 | 0.44858 | 0.64394 | 0.84894 | ↓ | -0.236 | 0.49088 | 0.7894 | 0.8632 | ↓ | -0.212 |
| Complement component C9 | 0.29973 | 0.6694 | 1.1125 | ↑ | 0.1538 | 0.10789 | 0.25925 | 1.1878 | ↑ | 0.2483 | 0.042306 | 0.19781 | 1.2612 | ↑ | 0.3348 |
| Uncharacterized protein | 0.535 | 0.82639 | 1.1681 | ↑ | 0.2242 | 0.099807 | 0.25055 | 1.4177 | ↑ | 0.5036 | 0.026606 | 0.17893 | 1.8203 | ↑ | 0.8642 |
| Uncharacterized protein | 0.76627 | 0.91424 | 0.91412 | ↓ | -0.13 | 0.58846 | 0.72201 | 0.71027 | ↓ | -0.494 | 0.10215 | 0.33183 | 0.42046 | ↓ | -1.25 |
| Haptoglobin | 0.23757 | 0.63229 | 1.7423 | ↑ | 0.801 | 0.000172 | 0.007454 | 3.0917 | ↑ | 1.6284 | 0.002067 | 0.027502 | 2.3367 | ↑ | 1.2244 |
| Joining chain of multimeric IgA and IgM | 0.59595 | 0.89651 | 1.1561 | ↑ | 0.2093 | 0.72062 | 0.80431 | 0.77013 | ↓ | -0.377 | 0.1055 | 0.33183 | 0.29586 | ↓ | -1.757 |
| Serpin peptidase inhibitor, clade D (Heparin cofactor), member 1 | 0.039836 | 0.27567 | 0.50847 | ↓ | -0.976 | 0.012903 | 0.090595 | 0.60469 | ↓ | -0.726 | 0.87828 | 0.94374 | 1.0026 | ↑ | 0.0038 |
| Complement C3 alpha chain | 0.50653 | 0.79663 | 1.2747 | ↑ | 0.3501 | 0.22152 | 0.42113 | 1.3364 | ↑ | 0.4183 | 0.033427 | 0.18831 | 0.79779 | ↓ | -0.326 |
| Uncharacterized protein | 0.19749 | 0.56942 | 1.8021 | ↑ | 0.8497 | 0.21436 | 0.41205 | 1.3302 | ↑ | 0.4116 | 0.22033 | 0.53035 | 0.67565 | ↓ | -0.566 |
| Coagulation factor XIII B chain | 0.41146 | 0.72636 | 0.87879 | ↓ | -0.186 | 0.059904 | 0.19554 | 0.65051 | ↓ | -0.62 | 0.018504 | 0.15244 | 0.60622 | ↓ | -0.722 |
| C3-beta-c | 0.23589 | 0.63229 | 0.69289 | ↓ | -0.529 | 0.39075 | 0.59299 | 0.77254 | ↓ | -0.372 | 0.70089 | 0.89024 | 1.0252 | ↑ | 0.0359 |
| Uncharacterized protein | 0.64549 | 0.90199 | 1.045 | ↑ | 0.0636 | 0.86132 | 0.91416 | 0.84055 | ↓ | -0.251 | 0.92021 | 0.95638 | 0.77778 | ↓ | -0.363 |
| Keratin 9 | 0.39697 | 0.72636 | 0.94486 | ↓ | -0.082 | 0.046775 | 0.17166 | 0.44812 | ↓ | -1.158 | 0.44547 | 0.74214 | 0.848 | ↓ | -0.238 |
| Soluble type II IL-1 receptor | 0.098665 | 0.44349 | 0.67611 | ↓ | -0.565 | 0.006625 | 0.071634 | 0.43083 | ↓ | -1.215 | 0.20442 | 0.53035 | 0.82046 | ↓ | -0.286 |
| Epididymis secretory sperm binding protein Li 163pA | 0.022499 | 0.22896 | 0.69873 | ↓ | -0.517 | 0.002079 | 0.035963 | 0.54026 | ↓ | -0.888 | 0.049922 | 0.20751 | 0.74603 | ↓ | -0.423 |
| Complement C5 | 0.99122 | 0.99217 | 0.99128 | ↓ | -0.013 | 0.41892 | 0.61418 | 1.0714 | ↑ | 0.0995 | 0.43724 | 0.74214 | 1.1252 | ↑ | 0.1702 |
| C3-beta-c | 0.33234 | 0.68348 | 0.83111 | ↓ | -0.267 | 0.087138 | 0.24151 | 0.73642 | ↓ | -0.441 | 0.81046 | 0.93473 | 1.0149 | ↑ | 0.0213 |
| C1q domain-containing protein | 0.09644 | 0.44349 | 0.13789 | ↓ | -2.859 | 0.7015 | 0.78996 | 1.3228 | ↑ | 0.4036 | 0.40744 | 0.71926 | 2.115 | ↑ | 1.0806 |
| Gc-globulin | 0.060766 | 0.36429 | 0.63596 | ↓ | -0.653 | 0.001561 | 0.03199 | 0.60892 | ↓ | -0.716 | 0.020807 | 0.1565 | 0.74432 | ↓ | -0.426 |
| Apolipoprotein B | 0.16042 | 0.52453 | 0.57211 | ↓ | -0.806 | 0.059834 | 0.19554 | 0.46146 | ↓ | -1.116 | 0.58006 | 0.81683 | 1.0417 | ↑ | 0.0589 |
| Mannan-binding lectin serine protease 1 | 0.76243 | 0.91424 | 0.97841 | ↓ | -0.031 | 0.60423 | 0.73233 | 0.94998 | ↓ | -0.074 | 0.099599 | 0.33183 | 0.80673 | ↓ | -0.31 |
| Complement component C6 | 0.50621 | 0.79663 | 0.90375 | ↓ | -0.146 | 0.70093 | 0.78996 | 0.96742 | ↓ | -0.048 | 0.40513 | 0.71926 | 1.1285 | ↑ | 0.1743 |
| Clusterin | 0.099674 | 0.44349 | 0.72902 | ↓ | -0.456 | 0.3063 | 0.49991 | 0.82986 | ↓ | -0.269 | 0.6736 | 0.89024 | 0.94901 | ↓ | -0.075 |
| Albumin | 0.71256 | 0.91424 | 2.4472 | ↑ | 1.2911 | 0.47352 | 0.64503 | 1.8415 | ↑ | 0.8809 | 0.049102 | 0.20751 | 0.43774 | ↓ | -1.192 |
| Actin-depolymerizing factor | 0.17435 | 0.55491 | 0.60632 | ↓ | -0.722 | 0.074101 | 0.21366 | 0.50022 | ↓ | -0.999 | 0.054314 | 0.21355 | 0.44352 | ↓ | -1.173 |
| Vitronectin | 0.50067 | 0.79663 | 0.91619 | ↓ | -0.126 | 0.65573 | 0.75723 | 0.9604 | ↓ | -0.058 | 0.10389 | 0.33183 | 0.76365 | ↓ | -0.389 |
| C4a anaphylatoxin | 0.7637 | 0.91424 | 1.0618 | ↑ | 0.0865 | 0.65812 | 0.75723 | 1.0725 | ↑ | 0.101 | 0.027601 | 0.17893 | 1.765 | ↑ | 0.8197 |
| Insulin-like growth factor I | 0.6116 | 0.89675 | 0.38618 | ↓ | -1.373 | 0.66094 | 0.75723 | 0.41306 | ↓ | -1.276 | 0.22051 | 0.53035 | 0.25749 | ↓ | -1.957 |
| Hepatocyte growth factor activator | 0.47533 | 0.79103 | 0.89863 | ↓ | -0.154 | 0.041353 | 0.1652 | 0.7446 | ↓ | -0.425 | 0.088187 | 0.31135 | 0.77438 | ↓ | -0.369 |

| | | | | | | | | | | | | |
|---|---|---|---|---|---|---|---|---|---|---|---|---|
| Carbonic anhydrase | 0.34491 | 0.68398 | 0.79469 ↓ | -0.332 | 0.062593 | 0.19688 | 1.9618 ↑ | 0.9722 | 0.88578 | 0.94553 | 1.5705 ↑ | 0.6513 |
| Beta actin variant | 0.77877 | 0.91651 | 1.2142 ↑ | 0.28 | 0.85843 | 0.91416 | 0.96979 ↓ | -0.044 | 0.11413 | 0.35259 | 0.7054 ↓ | -0.503 |
| Alpha-1-microglobulin | 0.80258 | 0.91787 | 0.97298 ↓ | -0.04 | 0.23759 | 0.43267 | 0.8405 ↓ | -0.251 | 0.63036 | 0.8621 | 0.93823 ↓ | -0.092 |
| Gc-globulin | 0.33977 | 0.68348 | 0.80434 ↓ | -0.314 | 0.10138 | 0.25055 | 0.66731 ↓ | -0.584 | 0.31478 | 0.61187 | 0.79435 ↓ | -0.332 |
| Mannose receptor C type 2 | 0.12116 | 0.48744 | 0.57668 ↓ | -0.794 | 0.7032 | 0.78996 | 0.88385 ↓ | -0.178 | 0.69235 | 0.89024 | 0.86175 ↓ | -0.215 |
| Macaca fascicularis brain cDNA clone: QtrA-15786, similar to huma | 0.88563 | 0.94448 | 0.92959 ↓ | -0.105 | 0.95447 | 0.97775 | 0.98009 ↓ | -0.029 | 0.053294 | 0.21355 | 0.63283 ↓ | -0.66 |
| Alpha-1-globin | 0.67258 | 0.90199 | 0.62688 ↓ | -0.674 | 0.39474 | 0.59382 | 1.4026 ↑ | 0.4881 | 0.74086 | 0.90598 | 1.125 ↑ | 0.1699 |
| Coagulation factor X | 0.10069 | 0.44349 | 0.75561 ↓ | -0.404 | 0.039862 | 0.1652 | 0.70397 ↓ | -0.506 | 0.040763 | 0.19781 | 0.73284 ↓ | -0.448 |
| Ceruloplasmin | 0.28327 | 0.6694 | 0.83895 ↓ | -0.253 | 0.12059 | 0.27912 | 0.57955 ↓ | -0.787 | 0.007087 | 0.080237 | 1.3908 ↑ | 0.4759 |
| IgGFc-binding protein | 0.30052 | 0.6694 | 1.1903 ↑ | 0.2513 | 0.20283 | 0.39874 | 1.9414 ↑ | 0.9571 | 0.50193 | 0.7894 | 0.78192 ↓ | -0.355 |
| Profilin | 0.60942 | 0.89675 | 2.3529 ↑ | 1.2344 | 0.52663 | 0.69547 | 1.2338 ↑ | 0.3031 | 0.49832 | 0.7894 | 0.67795 ↓ | -0.561 |
| Serpin family G member 1 | 0.76388 | 0.91424 | 0.98129 ↓ | -0.027 | 0.044545 | 0.16887 | 0.64604 ↓ | -0.63 | 0.57491 | 0.81683 | 1.2 ↑ | 0.263 |
| HCG40889, isoform CRA_b | 0.016262 | 0.22701 | 0.78639 ↓ | -0.347 | 0.024036 | 0.12681 | 0.78718 ↓ | -0.345 | 0.23876 | 0.53644 | 0.89776 ↓ | -0.156 |
| C4a anaphylatoxin | 0.44785 | 0.77888 | 1.3876 ↑ | 0.4726 | 0.31739 | 0.50842 | 1.4793 ↑ | 0.5649 | 0.079471 | 0.29252 | 1.6575 ↑ | 0.729 |
| Plasminogen | 0.66742 | 0.90199 | 1.1164 ↑ | 0.1588 | 0.16118 | 0.3451 | 1.272 ↑ | 0.3471 | 0.87043 | 0.94374 | 1.0103 ↑ | 0.0148 |
| Protein S100 | 0.18295 | 0.55934 | 1.842 ↑ | 0.8813 | 0.047629 | 0.17166 | 5.5382 ↑ | 2.4694 | 0.51711 | 0.80594 | 2.2122 ↑ | 1.1455 |
| Reversion inducing cysteine rich protein with kazal motifs | 0.90131 | 0.94448 | 0.54453 ↓ | -0.877 | 0.41782 | 0.61418 | 1.0501 ↑ | 0.0705 | 0.56765 | 0.81683 | 1.1739 ↑ | 0.2313 |
| Complement component C9 | 0.93574 | 0.95789 | 1.001 ↑ | 0.0015 | 0.567 | 0.7108 | 1.0516 ↑ | 0.0726 | 0.038565 | 0.19651 | 1.178 ↑ | 0.2364 |
| Glyceraldehyde-3-phosphate dehydrogenase | 0.090302 | 0.44349 | 3.996 ↑ | 1.9986 | 0.152 | 0.33287 | 2.8412 ↑ | 1.5065 | 0.22006 | 0.53035 | 0.56545 ↓ | -0.823 |
| Complement subcomponent C1r | 0.020626 | 0.22701 | 0.69078 ↓ | -0.534 | 0.011262 | 0.088558 | 0.69786 ↓ | -0.519 | 0.74812 | 0.90598 | 0.9711 ↓ | -0.042 |
| Fibrinogen beta chain | 0.28575 | 0.6694 | 0.73733 ↓ | -0.44 | 0.04131 | 0.1652 | 0.57396 ↓ | -0.801 | 0.57274 | 0.81683 | 1.5548 ↑ | 0.6367 |
| Cation-independent mannose-6-phosphate receptor | 0.91154 | 0.94448 | 1.1279 ↑ | 0.1737 | 0.55883 | 0.7108 | 0.77701 ↓ | -0.364 | 0.74936 | 0.90598 | 0.75051 ↓ | -0.414 |
| Complement C1r subcomponent-like protein | 0.49503 | 0.79663 | 0.90212 ↓ | -0.149 | 0.10381 | 0.25295 | 0.72354 ↓ | -0.467 | 0.32069 | 0.61644 | 0.8545 ↓ | -0.227 |
| Haptoglobin | 0.28527 | 0.6694 | 1.6732 ↑ | 0.7427 | 0.003144 | 0.043602 | 3.1328 ↑ | 1.6475 | 0.028959 | 0.17893 | 2.1619 ↑ | 1.1123 |
| Apolipoprotein A-II | 0.56257 | 0.86127 | 0.61127 ↓ | -0.71 | 0.61812 | 0.73725 | 0.3859 ↓ | -1.374 | 0.60155 | 0.83254 | 0.36797 ↓ | -1.442 |
| Alpha-2-HS-glycoprotein | 0.3723 | 0.70009 | 0.71993 ↓ | -0.474 | 0.89437 | 0.92957 | 0.97165 ↓ | -0.041 | 0.95169 | 0.97248 | 0.99085 ↓ | -0.013 |
| Apolipoprotein A-I | 0.83485 | 0.94164 | 0.71787 ↓ | -0.478 | 0.95515 | 0.97775 | 0.52918 ↓ | -0.918 | 0.083658 | 0.30152 | 0.11772 ↓ | -3.087 |
| Hemoglobin subunit beta | 0.80053 | 0.91787 | 1.3241 ↑ | 0.405 | 0.003277 | 0.043602 | 5.6588 ↑ | 2.5005 | 0.94864 | 0.97248 | 1.2381 ↑ | 0.3081 |
| Angiotensin-converting enzyme | 0.18744 | 0.55934 | 0.58617 ↓ | -0.771 | 0.28885 | 0.49003 | 0.62177 ↓ | -0.686 | 0.4161 | 0.72712 | 0.71096 ↓ | -0.492 |
| Complement C8 gamma chain | 0.76114 | 0.91424 | 0.94218 ↓ | -0.086 | 0.45707 | 0.64394 | 0.88117 ↓ | -0.183 | 0.58547 | 0.81683 | 1.0696 ↑ | 0.097 |
| Antithrombin-III | 0.89036 | 0.94448 | 0.97349 ↓ | -0.039 | 0.014721 | 0.090954 | 0.71051 ↓ | -0.493 | 0.000549 | 0.009785 | 0.43668 ↓ | -1.195 |
| C1q domain-containing protein | 0.61195 | 0.89675 | 0.85839 ↓ | -0.22 | 0.62555 | 0.73725 | 0.83339 ↓ | -0.263 | 0.35443 | 0.66511 | 0.73406 ↓ | -0.446 |
| Epididymis secretory protein Li 37 | 0.13744 | 0.51688 | 1.8809 ↑ | 0.9115 | 0.089347 | 0.24151 | 1.7764 ↑ | 0.829 | 0.12167 | 0.35677 | 1.6533 ↑ | 0.7254 |
| Fibrinogen gamma chain | 0.66334 | 0.90199 | 0.86431 ↓ | -0.21 | 0.16357 | 0.3451 | 0.5529 ↓ | -0.855 | 0.70433 | 0.89024 | 0.89128 ↓ | -0.166 |
| Apolipoprotein C-I, basic form | 0.66727 | 0.90199 | 0.3292 ↓ | -1.603 | 0.63434 | 0.74149 | 0.31661 ↓ | -1.659 | 0.98868 | 0.98868 | 0.48072 ↓ | -1.057 |
| Uncharacterized protein | 0.30406 | 0.6694 | 0.8994 ↓ | -0.153 | 0.02419 | 0.12681 | 0.78372 ↓ | -0.352 | 0.23729 | 0.53644 | 1.0977 ↑ | 0.1345 |
| Complement C4-B | 0.48073 | 0.79103 | 1.291 ↑ | 0.3685 | 0.3649 | 0.55865 | 1.3652 ↑ | 0.4491 | 0.03862 | 0.19651 | 1.5576 ↑ | 0.6394 |
| SERPIN domain-containing protein | 0.73077 | 0.91424 | 0.93563 ↓ | -0.096 | 0.28151 | 0.48701 | 0.7941 ↓ | -0.333 | 0.002695 | 0.033306 | 0.44923 ↓ | -1.155 |
| Lumican | 0.018111 | 0.22701 | 0.75845 ↓ | -0.399 | 0.00113 | 0.027918 | 0.66129 ↓ | -0.597 | 0.000306 | 0.007931 | 0.51327 ↓ | -0.962 |
| Cadherin-13 | 0.7847 | 0.91725 | 0.94484 ↓ | -0.082 | 0.057576 | 0.19531 | 1.5901 ↑ | 0.6691 | 0.54929 | 0.81683 | 1.3528 ↑ | 0.4359 |
| Angiotensin 1-10 | 0.23345 | 0.63229 | 1.204 ↑ | 0.2679 | 0.46081 | 0.64394 | 1.0722 ↑ | 0.1006 | 0.25452 | 0.55736 | 1.165 ↑ | 0.2203 |
| Keratin, type II cytoskeletal 2 epidermal | 0.40367 | 0.72636 | 0.76699 ↓ | -0.383 | 0.54767 | 0.70183 | 0.79522 ↓ | -0.331 | 0.9655 | 0.9768 | 1.0109 ↑ | 0.0156 |
| Fibulin-1 | 0.3655 | 0.69485 | 1.1917 ↑ | 0.253 | 0.32738 | 0.5196 | 1.2639 ↑ | 0.3379 | 0.049019 | 0.20751 | 0.71197 ↓ | -0.49 |
| Trypsin | 0.40942 | 0.72636 | 1.3569 ↑ | 0.4403 | 0.054904 | 0.18997 | 1.5284 ↑ | 0.612 | 0.050378 | 0.20751 | 0.79691 ↓ | -0.328 |
| 1,4-beta-N-acetylmuramidase C | 0.47118 | 0.79103 | 13.507 ↑ | 3.7557 | 0.013798 | 0.090595 | 7.2307 ↑ | 2.8541 | 0.73008 | 0.90598 | 0.92942 ↓ | -0.106 |
| Macaca fascicularis brain cDNA clone: QorA-13734, similar to huma | 0.91172 | 0.94448 | 0.63253 ↓ | -0.661 | 0.48457 | 0.65493 | 0.45113 ↓ | -1.148 | 0.92322 | 0.95638 | 0.71383 ↓ | -0.486 |
| Testicular tissue protein Li 61 | 0.17642 | 0.55491 | 0.82667 ↓ | -0.275 | 0.31426 | 0.50811 | 0.87546 ↓ | -0.192 | 0.000354 | 0.007931 | 0.5913 ↓ | -0.758 |
| Uncharacterized protein | 0.90023 | 0.94448 | 1.0039 ↑ | 0.0056 | 0.28892 | 0.49003 | 0.8854 ↓ | -0.176 | 0.24543 | 0.54435 | 0.86877 ↓ | -0.203 |
| Vitellogenin domain-containing protein | 0.7326 | 0.91424 | 0.94229 ↓ | -0.086 | 0.24691 | 0.44037 | 0.76171 ↓ | -0.393 | 0.89713 | 0.94553 | 1.0212 ↑ | 0.0303 |
| Uncharacterized protein | 0.1021 | 0.44349 | 0.66861 ↓ | -0.581 | 0.010225 | 0.088558 | 0.501 ↓ | -0.997 | 0.22072 | 0.53035 | 0.75488 ↓ | -0.406 |
| Serpin family A member 10 | 0.96662 | 0.97792 | 0.72888 ↓ | -0.456 | 0.60607 | 0.73233 | 0.63197 ↓ | -0.662 | 0.21979 | 0.53035 | 1.3131 ↑ | 0.393 |
| Complement component 8 subunit beta | 0.40051 | 0.72636 | 0.78038 ↓ | -0.358 | 0.33799 | 0.53157 | 0.65754 ↓ | -0.605 | 0.38473 | 0.70807 | 0.7259 ↓ | -0.462 |
| Collagen type VI alpha 3 chain | 0.76126 | 0.91424 | 0.73355 ↓ | -0.447 | 0.46156 | 0.64394 | 1.1999 ↑ | 0.2629 | 0.70174 | 0.89024 | 1.086 ↑ | 0.119 |

| | | | | | | | | | | | | |
|---|---|---|---|---|---|---|---|---|---|---|---|---|
| Glutathione peroxidase | 0.14484 | 0.52204 | 1.6087 ↑ | 0.6859 | 0.09283 | 0.24707 | 1.6725 ↑ | 0.742 | 0.16066 | 0.44119 | 0.77496 ↓ | -0.368 |
| Mannan binding lectin serine peptidase 2 | 0.83822 | 0.94164 | 1.0797 ↑ | 0.1106 | 0.86905 | 0.91674 | 1.0427 ↑ | 0.0603 | 0.95562 | 0.97248 | 1.0763 ↑ | 0.1062 |
| Protein S100 | 0.15863 | 0.52453 | 13.031 ↑ | 3.7039 | 0.001664 | 0.03199 | 5.4617 ↑ | 2.4493 | 0.54537 | 0.81683 | 1.9663 ↑ | 0.9755 |
| Bradykinin | 0.042972 | 0.28593 | 0.47487 ↓ | -1.074 | 0.010784 | 0.088558 | 0.39869 ↓ | -1.327 | 0.29176 | 0.58603 | 0.81966 ↓ | -0.287 |
| Protein C receptor | 0.46646 | 0.79103 | 1.1678 ↑ | 0.2239 | 0.54435 | 0.70183 | 0.89559 ↓ | -0.159 | 0.094662 | 0.32753 | 0.74523 ↓ | -0.424 |
| Protein-tyrosine-phosphatase | 0.52533 | 0.81876 | 1.0562 ↑ | 0.0789 | 0.30235 | 0.49991 | 0.75147 ↓ | -0.412 | 0.58037 | 0.81683 | 0.90313 ↓ | -0.147 |
| Uncharacterized protein | 0.66816 | 0.90199 | 0.92741 ↓ | -0.109 | 0.54309 | 0.70183 | 0.79554 ↓ | -0.33 | 0.83937 | 0.94374 | 0.91053 ↓ | -0.135 |
| Filamin A | 0.79034 | 0.91765 | 1.1036 ↑ | 0.1422 | 0.4184 | 0.61418 | 1.1977 ↑ | 0.2602 | 0.55957 | 0.81683 | 1.2646 ↑ | 0.3386 |
| Inter-alpha-trypsin inhibitor heavy chain 2 | 0.70129 | 0.91424 | 0.89719 ↓ | -0.157 | 0.82426 | 0.89218 | 0.64382 ↓ | -0.635 | 0.85617 | 0.94374 | 0.89127 ↓ | -0.166 |
| Fibrinogen alpha chain | 0.18752 | 0.55934 | 0.69626 ↓ | -0.522 | 0.83156 | 0.89354 | 1.0259 ↑ | 0.0369 | 0.98264 | 0.98835 | 1.0183 ↑ | 0.0262 |
| Complement C1s | 0.12914 | 0.49646 | 0.80751 ↓ | -0.308 | 0.50818 | 0.67707 | 0.90443 ↓ | -0.145 | 0.13071 | 0.37354 | 0.81417 ↓ | -0.297 |
| Mannosyl-oligosaccharide 1,2-alpha-mannosidase IA | 0.35569 | 0.68398 | 1.2813 ↑ | 0.3576 | 0.098929 | 0.25055 | 1.4967 ↑ | 0.5817 | 0.77115 | 0.91377 | 1.099 ↑ | 0.1362 |
| Apolipoprotein B (Including Ag(X) antigen) | 0.15627 | 0.52453 | 0.699 ↓ | -0.517 | 0.073848 | 0.21366 | 1.5885 ↑ | 0.6677 | 0.63287 | 0.8621 | 0.91494 ↓ | -0.128 |
| Complement component C8 alpha chain | 0.75829 | 0.91424 | 0.94376 ↓ | -0.084 | 0.18675 | 0.37567 | 0.76448 ↓ | -0.387 | 0.9018 | 0.94553 | 0.96166 ↓ | -0.056 |
| cDNA FLJ44930 fis, clone BRAMY3015549, highly similar to Neural c | 0.632 | 0.89701 | 0.87618 ↓ | -0.191 | 0.34604 | 0.53932 | 0.69424 ↓ | -0.526 | 0.46321 | 0.756 | 0.77919 ↓ | -0.36 |
| Metavinculin | 0.27702 | 0.6694 | 0.65548 ↓ | -0.609 | 0.20621 | 0.40084 | 0.62198 ↓ | -0.685 | 0.30724 | 0.60402 | 0.68972 ↓ | -0.536 |
| ITIH4 protein | 0.084597 | 0.44349 | 4.3979 ↑ | 2.1368 | 0.039808 | 0.1652 | 5.9841 ↑ | 2.5811 | 0.45917 | 0.756 | 2.7636 ↑ | 1.4666 |
| Apolipoprotein C-II | 0.89262 | 0.94448 | 0.94206 ↓ | -0.086 | 0.46137 | 0.64394 | 1.4955 ↑ | 0.5806 | 0.27535 | 0.56708 | 2.6278 ↑ | 1.3939 |
| Prothrombin | 0.094008 | 0.44349 | 0.41389 ↓ | -1.273 | 0.24185 | 0.43583 | 0.51605 ↓ | -0.954 | 0.29471 | 0.58603 | 0.5441 ↓ | -0.878 |
| Alpha-1-antitrypsin | 0.31285 | 0.6694 | 1.5578 ↑ | 0.6395 | 0.23208 | 0.43171 | 1.5304 ↑ | 0.6139 | 0.84049 | 0.94374 | 0.81059 ↓ | -0.303 |
| Complement C1q subcomponent subunit B | 0.31342 | 0.6694 | 0.80404 ↓ | -0.315 | 0.73616 | 0.81633 | 1.1221 ↑ | 0.1661 | 0.58448 | 0.81683 | 1.0445 ↑ | 0.0628 |
| SEMA4D variant protein | 0.90825 | 0.94448 | 1.9804 ↑ | 0.9858 | 0.96911 | 0.98183 | 1.4259 ↑ | 0.5119 | 0.8768 | 0.94374 | 1.2371 ↑ | 0.307 |
| CP protein | 0.2521 | 0.6437 | 1.8205 ↑ | 0.8644 | 0.14782 | 0.32785 | 1.7013 ↑ | 0.7666 | 0.74446 | 0.90598 | 1.1387 ↑ | 0.1874 |
| Attractin | 0.65078 | 0.90199 | 0.91936 ↓ | -0.121 | 0.083331 | 0.23633 | 0.73594 ↓ | -0.442 | 0.032287 | 0.18831 | 0.65613 ↓ | -0.608 |
| 15S Mg(2+)-ATPase p97 subunit | 0.061066 | 0.36429 | 2.0334 ↑ | 1.0239 | 0.025345 | 0.12896 | 1.7812 ↑ | 0.8328 | 0.76846 | 0.91377 | 1.1935 ↑ | 0.2552 |
| CST6 protein | 0.80645 | 0.91787 | 1.3604 ↑ | 0.4441 | 0.53535 | 0.70164 | 1.1971 ↑ | 0.2596 | 0.86395 | 0.94374 | 0.82896 ↓ | -0.271 |
| Monocyte differentiation antigen CD14 | 0.35583 | 0.68398 | 0.74085 ↓ | -0.433 | 0.97048 | 0.98183 | 0.91666 ↓ | -0.126 | 0.87094 | 0.94374 | 0.90567 ↓ | -0.143 |
| Uncharacterized protein | 0.70774 | 0.91424 | 0.81635 ↓ | -0.293 | 0.60957 | 0.73233 | 0.74114 ↓ | -0.432 | 0.87404 | 0.94374 | 1.0375 ↑ | 0.0531 |
| Immunoglobulin heavy constant mu | 0.3274 | 0.68348 | 0.5429 ↓ | -0.881 | 0.36413 | 0.55865 | 0.44907 ↓ | -1.155 | 0.35754 | 0.66511 | 0.509 ↓ | -0.974 |
| Apolipoprotein A4 | 0.99217 | 0.99217 | 0.97701 ↓ | -0.034 | 0.57198 | 0.71189 | 0.85973 ↓ | -0.218 | 3.26E-05 | 0.002111 | 0.11747 ↓ | -3.09 |
| Ficolin (Collagen/fibrinogen domain containing) 3 (Hakata antigen), | 0.11415 | 0.47017 | 0.55205 ↓ | -0.857 | 0.012853 | 0.090595 | 0.28943 ↓ | -1.789 | 5.22E-05 | 0.002259 | 0.079424 ↓ | -3.654 |
| Receptor protein-tyrosine kinase | 0.96132 | 0.97792 | 1.0749 ↑ | 0.1043 | 0.19566 | 0.38907 | 1.5026 ↑ | 0.5874 | 0.6927 | 0.89024 | 0.89069 ↓ | -0.167 |
| Insulin-like growth factor-binding protein 3 | 0.85214 | 0.94448 | 0.96275 ↓ | -0.055 | 0.89733 | 0.92957 | 1.0042 ↑ | 0.0061 | 0.2702 | 0.56434 | 0.72488 ↓ | -0.464 |
| Uncharacterized protein DKFZp686I15196 | 0.33696 | 0.68348 | 5.0231 ↑ | 2.3286 | 0.054786 | 0.18997 | 4.0579 ↑ | 2.0207 | 0.49706 | 0.7894 | 0.75639 ↓ | -0.403 |
| cDNA, FLJ93914, highly similar to Homo sapiens histidine-rich glyco | 0.033976 | 0.24868 | 0.57584 ↓ | -0.796 | 0.009635 | 0.088558 | 0.50281 ↓ | -0.992 | 0.12084 | 0.35677 | 0.67269 ↓ | -0.572 |
| Coagulation factor XII | 0.72366 | 0.91424 | 1.4152 ↑ | 0.501 | 0.79142 | 0.86656 | 1.1531 ↑ | 0.2055 | 0.68104 | 0.89024 | 0.68778 ↓ | -0.54 |
| Protein S100-A7 | 0.61684 | 0.89675 | 0.64613 ↓ | -0.63 | 0.74083 | 0.81633 | 0.77103 ↓ | -0.375 | 0.18844 | 0.50153 | 1.5362 ↑ | 0.6194 |
| Insulin-like growth factor binding protein, acid labile subunit | 0.92423 | 0.95174 | 0.95447 ↓ | -0.067 | 0.99395 | 0.99395 | 0.93617 ↓ | -0.095 | 0.26257 | 0.5608 | 0.72798 ↓ | -0.458 |
| Carboxylic ester hydrolase | 0.62973 | 0.89701 | 0.92492 ↓ | -0.113 | 0.042017 | 0.1652 | 0.74318 ↓ | -0.428 | 0.028756 | 0.17893 | 0.67744 ↓ | -0.562 |

## 8.5 Gy

| Protein Name | SD 1 vs. Pre *p-value* | FDR | FC | LOG2FC | SD 2 vs. Pre *p-value* | FDR | FC | LOG2FC | SD 6 vs. Pre *p-value* | FDR | FC | LOG2FC |
|---|---|---|---|---|---|---|---|---|---|---|---|---|
| Serpin family A member 9 | 1.87E-05 | 0.001574 | 4.6172 ↑ | 2.207 | 0.000685 | 0.024528 | 3.853 ↑ | 1.946 | 0.098235 | 0.30826 | 1.6148 ↑ | 0.6913 |
| C-reactive protein | 9.12E-06 | 0.001574 | 20.553 ↑ | 4.3613 | 0.024391 | 0.16229 | 8.6617 ↑ | 3.1146 | 0.90572 | 0.98222 | 1.1006 ↑ | 0.1383 |
| cDNA FLJ35730 fis, clone TESTI2003131, highly similar to ALPHA-1-/ | 0.000779 | 0.01685 | 4.3236 ↑ | 2.1122 | 0.000356 | 0.024036 | 4.3943 ↑ | 2.1356 | 0.26774 | 0.53241 | 1.5007 ↑ | 0.5857 |
| Fibronectin | 0.028015 | 0.21982 | 0.48569 ↓ | -1.042 | 0.17986 | 0.47109 | 0.63142 ↓ | -0.663 | 0.11264 | 0.31429 | 0.57553 ↓ | -0.797 |
| Retinol-binding protein | 0.000658 | 0.016266 | 0.49363 ↓ | -1.019 | 0.034808 | 0.1917 | 0.62138 ↓ | -0.686 | 0.45581 | 0.70234 | 0.87715 ↓ | -0.189 |
| Clusterin | 0.029224 | 0.21982 | 0.34991 ↓ | -1.515 | 0.000417 | 0.024036 | 0.39255 ↓ | -1.349 | 0.010827 | 0.14357 | 0.60834 ↓ | -0.717 |
| Apolipoprotein C-III | 0.00018 | 0.006215 | 0.24421 ↓ | -2.034 | 0.006108 | 0.08128 | 0.36788 ↓ | -1.443 | 0.12028 | 0.33029 | 0.56196 ↓ | -0.831 |
| Testis cDNA clone: QtsA-10685, similar to human I factor (complem | 0.67327 | 0.87576 | 0.97385 ↓ | -0.038 | 0.001023 | 0.025811 | 0.77777 ↓ | -0.363 | 0.91855 | 0.98222 | 1.0057 ↑ | 0.0082 |
| Protein C, inactivator of coagulation factors Va and VIIIa | 0.000435 | 0.012532 | 0.56727 ↓ | -0.818 | 0.004673 | 0.068905 | 0.54453 ↓ | -0.877 | 0.16039 | 0.37124 | 0.79855 ↓ | -0.325 |
| Endonuclease domain containing 1 | 0.34228 | 0.64789 | 0.39883 ↓ | -1.326 | 0.63418 | 0.81875 | 0.65672 ↓ | -0.607 | 0.36694 | 0.63692 | 0.53439 ↓ | -0.904 |

| | | | | | | | | | | | | |
|---|---|---|---|---|---|---|---|---|---|---|---|---|
| Macaca fascicularis brain cDNA clone: QtrA-15720, similar to huma | 0.48641 | 0.78645 | 0.83028 ↓ | -0.268 | 0.78131 | 0.91329 | 0.94863 ↓ | -0.076 | 0.32919 | 0.61901 | 1.1462 ↑ | 0.1969 |
| Apolipoprotein B | 0.050146 | 0.25005 | 0.43876 ↓ | -1.189 | 0.10144 | 0.33747 | 0.59145 ↓ | -0.758 | 0.016659 | 0.14528 | 0.29251 ↓ | -1.774 |
| Leucine-rich alpha-2-glycoprotein | 0.1853 | 0.51182 | 2.3593 ↑ | 1.2383 | 0.28889 | 0.57074 | 1.5518 ↑ | 0.6339 | 0.16094 | 0.37124 | 0.64655 ↓ | -0.629 |
| Lambda-chain (AA -20 to 215) | 3.64E-05 | 0.001574 | 0.57134 ↓ | -0.808 | 0.000143 | 0.024036 | 0.58205 ↓ | -0.781 | 0.002951 | 0.091633 | 0.5926 ↓ | -0.755 |
| Inter-alpha-trypsin inhibitor heavy chain 1 | 0.20379 | 0.51182 | 0.75433 ↓ | -0.407 | 0.76201 | 0.91261 | 1.47 ↑ | 0.5558 | 0.76051 | 0.89703 | 0.97717 ↓ | -0.033 |
| Apolipoprotein H | 0.49769 | 0.78993 | 0.91574 ↓ | -0.127 | 0.5577 | 0.77649 | 0.94465 ↓ | -0.082 | 0.36816 | 0.63692 | 0.8616 ↓ | -0.215 |
| Testis cDNA, clone: QtsA-17169, similar to human complement com | 0.11706 | 0.39774 | 0.86245 ↓ | -0.213 | 0.099224 | 0.33658 | 0.85318 ↓ | -0.229 | 0.024933 | 0.1659 | 0.80159 ↓ | -0.319 |
| Cathepsin D | 0.005891 | 0.072263 | 0.59684 ↓ | -0.745 | 0.13422 | 0.39184 | 0.6768 ↓ | -0.563 | 0.028674 | 0.17106 | 0.50899 ↓ | -0.974 |
| Histidine rich glycoprotein | 0.73224 | 0.8797 | 0.82033 ↓ | -0.286 | 0.37128 | 0.62972 | 0.67884 ↓ | -0.559 | 0.14759 | 0.36968 | 0.55958 ↓ | -0.838 |
| Inter-alpha (Globulin) inhibitor H2 | 0.01709 | 0.15561 | 0.79504 ↓ | -0.331 | 0.008004 | 0.082654 | 0.71207 ↓ | -0.49 | 0.17075 | 0.38364 | 0.86204 ↓ | -0.214 |
| Lipopolysaccharide-binding protein | 0.005421 | 0.072263 | 7.6493 ↑ | 2.9353 | 0.098217 | 0.33658 | 2.6746 ↑ | 1.4193 | 0.15172 | 0.36968 | 2.9197 ↑ | 1.5458 |
| Inter-alpha-trypsin inhibitor heavy chain 2 | 0.9416 | 0.9855 | 0.98709 ↓ | -0.019 | 0.74763 | 0.91261 | 2.1988 ↑ | 1.1367 | 0.77259 | 0.89703 | 1.1336 ↑ | 0.1809 |
| C-X-C motif chemokine | 0.042512 | 0.25005 | 0.71361 ↓ | -0.487 | 0.0086 | 0.082654 | 0.60318 ↓ | -0.729 | 0.093618 | 0.29992 | 0.45936 ↓ | -1.122 |
| Hexosyltransferase | 0.4678 | 0.76349 | 0.73454 ↓ | -0.445 | 0.38784 | 0.64457 | 0.67079 ↓ | -0.576 | 0.43133 | 0.70234 | 0.78788 ↓ | -0.344 |
| Platelet factor 4 | 0.00861 | 0.087617 | 0.34759 ↓ | -1.525 | 0.039891 | 0.1917 | 0.42176 ↓ | -1.246 | 0.011419 | 0.14357 | 0.31387 ↓ | -1.672 |
| Alpha-2-HS-glycoprotein | 0.010397 | 0.099931 | 0.56023 ↓ | -0.836 | 0.004468 | 0.068905 | 0.49935 ↓ | -1.002 | 0.027189 | 0.16799 | 0.61539 ↓ | -0.7 |
| N-acetylmuramoyl-L-alanine amidase | 0.70994 | 0.8797 | 0.93447 ↓ | -0.098 | 0.7518 | 0.91261 | 0.97298 ↓ | -0.04 | 0.81793 | 0.93094 | 1.1144 ↑ | 0.1563 |
| Vitellogenin domain-containing protein | 0.10547 | 0.39117 | 0.62671 ↓ | -0.674 | 0.12382 | 0.3825 | 0.64124 ↓ | -0.641 | 0.39223 | 0.66657 | 0.79519 ↓ | -0.331 |
| Apolipoprotein H | 0.005719 | 0.072263 | 0.36532 ↓ | -1.453 | 0.028692 | 0.18036 | 0.43046 ↓ | -1.216 | 0.021557 | 0.15539 | 0.42134 ↓ | -1.247 |
| C3-beta-c | 0.14022 | 0.42557 | 0.69805 ↓ | -0.519 | 0.19458 | 0.47109 | 0.71553 ↓ | -0.483 | 0.22058 | 0.46244 | 0.74073 ↓ | -0.433 |
| Vitamin K-dependent protein S | 0.97507 | 0.9855 | 0.93562 ↓ | -0.096 | 0.29511 | 0.57365 | 0.69889 ↓ | -0.517 | 0.73152 | 0.87885 | 1.0408 ↑ | 0.0577 |
| Fibronectin | 0.032603 | 0.22561 | 0.42891 ↓ | -1.221 | 0.081774 | 0.30754 | 0.52298 ↓ | -0.935 | 0.016796 | 0.14528 | 0.29107 ↓ | -1.781 |
| Plasminogen | 0.09365 | 0.36821 | 0.81914 ↓ | -0.288 | 0.051429 | 0.22243 | 0.75362 ↓ | -0.408 | 0.053306 | 0.22459 | 0.76592 ↓ | -0.385 |
| Plasma kallikrein | 0.3343 | 0.64789 | 0.90975 ↓ | -0.136 | 0.19606 | 0.47109 | 0.84949 ↓ | -0.235 | 0.27686 | 0.53884 | 0.84407 ↓ | -0.245 |
| Testis cDNA, clone: QtsA-16984, similar to human sex hormone-bin | 0.033929 | 0.22576 | 0.6162 ↓ | -0.699 | 0.008212 | 0.082654 | 0.52848 ↓ | -0.92 | 0.24984 | 0.50849 | 0.77436 ↓ | -0.369 |
| Plasminogen | 0.005877 | 0.072263 | 0.78099 ↓ | -0.357 | 0.18525 | 0.47109 | 0.63593 ↓ | -0.653 | 0.16665 | 0.37936 | 0.67706 ↓ | -0.563 |
| Gc-globulin | 2.85E-05 | 0.001574 | 0.59589 ↓ | -0.747 | 0.001044 | 0.025811 | 0.57918 ↓ | -0.788 | 0.003659 | 0.091633 | 0.60143 ↓ | -0.734 |
| Alpha-2-macroglobulin | 0.52111 | 0.7978 | 0.56567 ↓ | -0.822 | 0.42432 | 0.66133 | 0.53847 ↓ | -0.893 | 0.62879 | 0.79986 | 0.77671 ↓ | -0.365 |
| Coagulation factor XII-Mie | 0.038449 | 0.24636 | 0.77217 ↓ | -0.373 | 0.007257 | 0.082654 | 0.64508 ↓ | -0.632 | 0.050953 | 0.22459 | 0.77728 ↓ | -0.363 |
| Alpha-2-glycoprotein 1, zinc-binding | 0.005084 | 0.072263 | 0.65952 ↓ | -0.601 | 0.002331 | 0.0504 | 0.61772 ↓ | -0.695 | 0.00028 | 0.024181 | 0.54889 ↓ | -0.865 |
| Fibronectin | 0.9855 | 0.9855 | 0.95086 ↓ | -0.073 | 0.97899 | 0.98723 | 1.0413 ↑ | 0.0583 | 0.76297 | 0.89703 | 1.1309 ↑ | 0.1775 |
| SERPIN domain-containing protein | 0.5028 | 0.79077 | 1.0897 ↑ | 0.124 | 0.95296 | 0.9841 | 1.0174 ↑ | 0.0248 | 0.52519 | 0.73869 | 0.92917 ↓ | -0.106 |
| Receptor protein-tyrosine kinase | 0.51975 | 0.7978 | 1.025 ↑ | 0.0356 | 0.832 | 0.91678 | 0.68241 ↓ | -0.551 | 0.47406 | 0.70234 | 0.33588 ↓ | -1.574 |
| cDNA FLJ51597, highly similar to C4b-binding protein alpha chain | 0.55438 | 0.81174 | 1.0627 ↑ | 0.0877 | 0.983 | 0.98723 | 0.92989 ↓ | -0.105 | 0.67187 | 0.82792 | 0.82831 ↓ | -0.272 |
| Fibronectin | 0.59763 | 0.82317 | 0.61314 ↓ | -0.706 | 0.73592 | 0.91261 | 0.58692 ↓ | -0.769 | 0.52183 | 0.73869 | 0.5303 ↓ | -0.915 |
| Keratin, type I cytoskeletal 10 | 0.29983 | 0.62899 | 0.66278 ↓ | -0.593 | 0.57055 | 0.7772 | 0.7701 ↓ | -0.377 | 0.46728 | 0.70234 | 2.5416 ↑ | 1.3457 |
| Prothrombin | 0.7291 | 0.8797 | 0.9385 ↓ | -0.092 | 0.49848 | 0.72973 | 0.63684 ↓ | -0.651 | 0.58077 | 0.75915 | 1.0588 ↑ | 0.0825 |
| C-type lectin domain family 3 member B | 0.045848 | 0.25005 | 0.78727 ↓ | -0.345 | 0.47784 | 0.71263 | 0.9274 ↓ | -0.109 | 0.009228 | 0.14357 | 0.51625 ↓ | -0.954 |
| C3/C5 convertase | 0.59954 | 0.82317 | 0.92509 ↓ | -0.112 | 0.17363 | 0.46212 | 0.81918 ↓ | -0.288 | 0.092999 | 0.29992 | 0.76291 ↓ | -0.39 |
| Hemopexin | 0.006266 | 0.072263 | 1.3813 ↑ | 0.466 | 0.036213 | 0.1917 | 1.5191 ↑ | 0.6032 | 0.018546 | 0.15278 | 1.2878 ↑ | 0.3649 |
| SERPIN domain-containing protein | 0.061264 | 0.27712 | 0.48588 ↓ | -1.041 | 0.021881 | 0.16229 | 0.46324 ↓ | -1.11 | 0.10521 | 0.31381 | 0.57425 ↓ | -0.8 |
| Hyaluronan-binding protein 2 | 0.81184 | 0.92647 | 0.87085 ↓ | -0.2 | 0.39443 | 0.64457 | 0.65458 ↓ | -0.611 | 0.93225 | 0.98222 | 0.88184 ↓ | -0.181 |
| SERPIN domain-containing protein | 0.079408 | 0.32789 | 0.80355 ↓ | -0.316 | 0.00478 | 0.068905 | 0.68318 ↓ | -0.55 | 0.003708 | 0.091633 | 0.64029 ↓ | -0.643 |
| Complement component C9 | 0.35921 | 0.65415 | 1.1002 ↑ | 0.1378 | 0.60953 | 0.81078 | 0.94393 ↓ | -0.083 | 0.047918 | 0.22459 | 1.2621 ↑ | 0.3358 |
| Uncharacterized protein | 0.58141 | 0.82256 | 1.2286 ↑ | 0.297 | 0.11067 | 0.35456 | 1.3806 ↑ | 0.4653 | 0.15963 | 0.37124 | 1.695 ↑ | 0.7613 |
| Uncharacterized protein | 0.38495 | 0.67955 | 0.61311 ↓ | -0.706 | 0.2847 | 0.57074 | 0.59867 ↓ | -0.74 | 0.48311 | 0.70234 | 0.73627 ↓ | -0.442 |
| Haptoglobin | 0.024749 | 0.20388 | 3.2769 ↑ | 1.7123 | 0.003569 | 0.068601 | 3.3753 ↑ | 1.755 | 0.000225 | 0.024181 | 6.1019 ↑ | 2.6092 |
| Joining chain of multimeric IgA and IgM | 0.81936 | 0.92647 | 0.6582 ↓ | -0.603 | 0.78936 | 0.91522 | 0.63531 ↓ | -0.654 | 0.93391 | 0.98222 | 0.76175 ↓ | -0.393 |
| Serpin peptidase inhibitor, clade D (Heparin cofactor), member 1 | 0.002963 | 0.056953 | 0.43399 ↓ | -1.204 | 0.000709 | 0.024528 | 0.60472 ↓ | -0.726 | 0.2772 | 0.53884 | 0.82058 ↓ | -0.285 |
| Complement C3 alpha chain | 0.33934 | 0.64789 | 1.4198 ↑ | 0.5057 | 0.9002 | 0.95917 | 1.2013 ↑ | 0.2647 | 0.96583 | 0.98288 | 1.1457 ↑ | 0.1963 |
| Uncharacterized protein | 0.11951 | 0.39774 | 2.026 ↑ | 1.0186 | 0.07773 | 0.30058 | 1.796 ↑ | 0.8448 | 0.012888 | 0.14357 | 2.5671 ↑ | 1.3601 |
| Coagulation factor XIII B chain | 0.11631 | 0.39774 | 0.67803 ↓ | -0.561 | 0.25992 | 0.56081 | 0.78818 ↓ | -0.343 | 0.20573 | 0.44488 | 0.75162 ↓ | -0.412 |
| C3-beta-c | 0.11955 | 0.39774 | 0.33133 ↓ | -1.594 | 0.095841 | 0.33658 | 0.31833 ↓ | -1.651 | 0.052294 | 0.22459 | 0.27234 ↓ | -1.877 |

| | | | | | | | | | | | | |
|---|---|---|---|---|---|---|---|---|---|---|---|---|
| Uncharacterized protein | 0.8911 | 0.96956 | 0.8346 ↓ | -0.261 | 0.76962 | 0.91261 | 0.96453 ↓ | -0.052 | 0.87458 | 0.97462 | 0.8924 ↓ | -0.164 |
| Keratin 9 | 0.047079 | 0.25005 | 0.29135 ↓ | -1.779 | 0.13153 | 0.39184 | 0.39666 ↓ | -1.334 | 0.445 | 0.70234 | 0.60636 ↓ | -0.722 |
| Soluble type II IL-1 receptor | 0.35792 | 0.65415 | 0.58824 ↓ | -0.766 | 0.21767 | 0.50209 | 0.4322 ↓ | -1.21 | 0.10905 | 0.31429 | 0.35027 ↓ | -1.514 |
| Epididymis secretory sperm binding protein Li 163pA | 0.58397 | 0.82256 | 0.90302 ↓ | -0.147 | 0.36104 | 0.62897 | 0.84556 ↓ | -0.242 | 0.13666 | 0.35304 | 0.75827 ↓ | -0.399 |
| Complement C5 | 0.45484 | 0.74941 | 0.88836 ↓ | -0.171 | 0.84661 | 0.92698 | 0.99857 ↓ | -0.002 | 0.96037 | 0.98288 | 0.9996 ↓ | -6E-04 |
| C3-beta-c | 0.24854 | 0.58105 | 0.87942 ↓ | -0.185 | 0.1293 | 0.39184 | 0.87124 ↓ | -0.199 | 0.030239 | 0.17438 | 0.79391 ↓ | -0.333 |
| C1q domain-containing protein | 0.30475 | 0.62899 | 2.2872 ↑ | 1.1936 | 0.26258 | 0.56081 | 0.11638 ↓ | -3.103 | 0.46685 | 0.70234 | 0.51755 ↓ | -0.95 |
| Gc-globulin | 0.00735 | 0.079472 | 0.69747 ↓ | -0.52 | 0.033455 | 0.1917 | 0.751 ↓ | -0.413 | 0.00676 | 0.14357 | 0.57474 ↓ | -0.799 |
| Apolipoprotein B | 0.079603 | 0.32789 | 0.60364 ↓ | -0.728 | 0.078186 | 0.30058 | 0.62047 ↓ | -0.689 | 0.054524 | 0.22459 | 0.62161 ↓ | -0.686 |
| Mannan-binding lectin serine protease 1 | 0.052511 | 0.25235 | 0.80616 ↓ | -0.311 | 0.037577 | 0.1917 | 0.74893 ↓ | -0.417 | 0.013429 | 0.14357 | 0.65109 ↓ | -0.619 |
| Complement component C6 | 0.86974 | 0.95231 | 1.0075 ↑ | 0.0107 | 0.40374 | 0.64673 | 0.88627 ↓ | -0.174 | 0.57563 | 0.75915 | 1.1176 ↑ | 0.1604 |
| Clusterin | 0.58241 | 0.82256 | 0.85491 ↓ | -0.226 | 0.094341 | 0.33658 | 0.62503 ↓ | -0.678 | 0.55328 | 0.75915 | 1.0733 ↑ | 0.1021 |
| Albumin | 0.36319 | 0.65451 | 0.58444 ↓ | -0.775 | 0.96225 | 0.98503 | 1.7066 ↑ | 0.7711 | 0.83618 | 0.94548 | 0.85794 ↓ | -0.221 |
| Actin-depolymerizing factor | 0.25205 | 0.58139 | 0.58949 ↓ | -0.762 | 0.76132 | 0.91261 | 0.88174 ↓ | -0.182 | 0.035831 | 0.18784 | 0.30448 ↓ | -1.716 |
| Vitronectin | 0.86478 | 0.95231 | 1.0175 ↑ | 0.025 | 0.82529 | 0.91522 | 0.9354 ↓ | -0.096 | 0.95725 | 0.98288 | 1.0303 ↑ | 0.043 |
| C4a anaphylatoxin | 0.85311 | 0.94607 | 1.0451 ↑ | 0.0636 | 0.80418 | 0.91522 | 1.0415 ↑ | 0.0587 | 0.11191 | 0.31429 | 1.3789 ↑ | 0.4635 |
| Insulin-like growth factor I | 0.3054 | 0.62899 | 0.45348 ↓ | -1.141 | 0.1359 | 0.39184 | 0.28477 ↓ | -1.812 | 0.95359 | 0.98288 | 0.70155 ↓ | -0.511 |
| Hepatocyte growth factor activator | 0.2578 | 0.58682 | 0.83994 ↓ | -0.252 | 0.54989 | 0.77649 | 0.91548 ↓ | -0.127 | 0.082071 | 0.2784 | 0.73678 ↓ | -0.441 |
| Carbonic anhydrase | 0.35071 | 0.6524 | 0.74448 ↓ | -0.426 | 0.90373 | 0.95917 | 0.88838 ↓ | -0.171 | 0.98445 | 0.99179 | 1.3388 ↑ | 0.4209 |
| Beta actin variant | 0.91485 | 0.98303 | 1.1469 ↑ | 0.1977 | 0.017901 | 0.15484 | 0.5175 ↓ | -0.95 | 0.94248 | 0.98222 | 1.4482 ↑ | 0.5343 |
| Alpha-1-microglobulin | 0.11724 | 0.39774 | 0.86448 ↓ | -0.21 | 0.064787 | 0.26065 | 0.79345 ↓ | -0.334 | 0.15142 | 0.36968 | 0.84845 ↓ | -0.237 |
| Gc-globulin | 0.18725 | 0.51182 | 0.77479 ↓ | -0.368 | 0.22329 | 0.50828 | 0.79613 ↓ | -0.329 | 0.13504 | 0.35304 | 0.73823 ↓ | -0.438 |
| Mannose receptor C type 2 | 0.22769 | 0.5396 | 2.3428 ↑ | 1.2283 | 0.81588 | 0.91522 | 2.1366 ↑ | 1.0953 | 0.099785 | 0.30826 | 3.6761 ↑ | 1.8782 |
| Macaca fascicularis brain cDNA clone: QtrA-15786, similar to huma | 0.76765 | 0.89732 | 0.9356 ↓ | -0.096 | 0.45175 | 0.68053 | 0.80487 ↓ | -0.313 | 0.65067 | 0.82013 | 0.8889 ↓ | -0.17 |
| Alpha-1-globin | 0.34454 | 0.64789 | 0.7767 ↓ | -0.365 | 0.9731 | 0.98723 | 1.4723 ↑ | 0.5581 | 0.58999 | 0.7617 | 1.5102 ↑ | 0.5947 |
| Coagulation factor X | 0.20305 | 0.51182 | 0.79375 ↓ | -0.333 | 0.34561 | 0.62049 | 0.84301 ↓ | -0.246 | 0.22187 | 0.46244 | 0.80432 ↓ | -0.314 |
| Ceruloplasmin | 0.047851 | 0.25005 | 0.64359 ↓ | -0.636 | 0.022204 | 0.16229 | 0.61878 ↓ | -0.693 | 0.14915 | 0.36968 | 0.72807 ↓ | -0.458 |
| IgGFc-binding protein | 0.92684 | 0.9855 | 1.4509 ↑ | 0.5369 | 0.14954 | 0.41725 | 0.68611 ↓ | -0.543 | 0.065038 | 0.2408 | 0.61634 ↓ | -0.698 |
| Profilin | 0.69318 | 0.8797 | 1.5908 ↑ | 0.6698 | 0.16045 | 0.43372 | 0.6661 ↓ | -0.586 | 0.36038 | 0.63692 | 1.4508 ↑ | 0.5369 |
| Serpin family G member 1 | 0.12533 | 0.40152 | 0.60006 ↓ | -0.737 | 0.4202 | 0.66085 | 0.73939 ↓ | -0.436 | 0.068779 | 0.24789 | 0.5465 ↓ | -0.872 |
| HCG40889, isoform CRA_b | 0.41554 | 0.70923 | 1.1197 ↑ | 0.1632 | 0.81528 | 0.91522 | 0.95481 ↓ | -0.067 | 0.51841 | 0.73869 | 0.91714 ↓ | -0.125 |
| C4a anaphylatoxin | 0.69203 | 0.8797 | 1.1818 ↑ | 0.241 | 0.94853 | 0.9841 | 1.0947 ↑ | 0.1306 | 0.071805 | 0.25351 | 1.5396 ↑ | 0.6225 |
| Plasminogen | 0.72423 | 0.8797 | 1.148 ↑ | 0.1991 | 0.61994 | 0.8125 | 0.97688 ↓ | -0.034 | 0.65421 | 0.82013 | 1.311 ↑ | 0.3907 |
| Protein S100 | 0.049178 | 0.25005 | 0.24245 ↓ | -2.044 | 0.022673 | 0.16229 | 0.19412 ↓ | -2.365 | 0.55102 | 0.75915 | 20.666 ↑ | 4.3692 |
| Reversion inducing cysteine rich protein with kazal motifs | 0.51064 | 0.79586 | 0.78805 ↓ | -0.344 | 0.77187 | 0.91261 | 0.85964 ↓ | -0.218 | 0.7886 | 0.90952 | 0.81838 ↓ | -0.289 |
| Complement component C9 | 0.078099 | 0.32789 | 1.155 ↑ | 0.2079 | 0.34668 | 0.62049 | 1.1622 ↑ | 0.2169 | 0.0462 | 0.22459 | 1.1892 ↑ | 0.2499 |
| Glyceraldehyde-3-phosphate dehydrogenase | 0.19633 | 0.51182 | 0.44346 ↓ | -1.173 | 0.60623 | 0.81078 | 0.73949 ↓ | -0.435 | 0.47783 | 0.70234 | 3.3351 ↑ | 1.7377 |
| Complement subcomponent C1r | 0.10183 | 0.39117 | 0.82478 ↓ | -0.278 | 0.061252 | 0.2523 | 0.80642 ↓ | -0.31 | 0.001126 | 0.063979 | 0.67197 ↓ | -0.574 |
| Fibrinogen beta chain | 0.28884 | 0.62899 | 0.79399 ↓ | -0.333 | 0.024206 | 0.16229 | 0.61203 ↓ | -0.708 | 0.056317 | 0.22658 | 0.62171 ↓ | -0.686 |
| Cation-independent mannose-6-phosphate receptor | 0.74218 | 0.88393 | 1.0007 ↑ | 0.001 | 0.56868 | 0.7772 | 0.77791 ↓ | -0.362 | 0.18224 | 0.39908 | 0.5708 ↓ | -0.809 |
| Complement C1r subcomponent-like protein | 0.325 | 0.64626 | 0.60635 ↓ | -0.722 | 0.86636 | 0.94264 | 0.90205 ↓ | -0.149 | 0.39301 | 0.66657 | 0.60191 ↓ | -0.732 |
| Haptoglobin | 0.031479 | 0.22561 | 4.5519 ↑ | 2.1865 | 0.029192 | 0.18036 | 3.6676 ↑ | 1.8749 | 0.001479 | 0.063979 | 8.7884 ↑ | 3.1356 |
| Apolipoprotein A-II | 0.66671 | 0.8738 | 0.39897 ↓ | -1.326 | 0.33941 | 0.62049 | 0.37187 ↓ | -1.427 | 0.62097 | 0.79577 | 0.62395 ↓ | -0.681 |
| Alpha-2-HS-glycoprotein | 0.41272 | 0.70923 | 0.68739 ↓ | -0.541 | 0.7435 | 0.91261 | 0.80841 ↓ | -0.307 | 0.34636 | 0.63692 | 0.66327 ↓ | -0.592 |
| Apolipoprotein A-I | 0.7909 | 0.91829 | 0.39487 ↓ | -1.341 | 0.3544 | 0.62562 | 0.19466 ↓ | -2.361 | 0.5656 | 0.75915 | 0.84018 ↓ | -0.251 |
| Hemoglobin subunit beta | 0.82715 | 0.9292 | 0.40872 ↓ | -1.291 | 0.93767 | 0.9841 | 0.28592 ↓ | -1.806 | 0.4829 | 0.70234 | 0.88262 ↓ | -0.18 |
| Angiotensin-converting enzyme | 0.43437 | 0.72256 | 1.0441 ↑ | 0.0623 | 0.63103 | 0.81875 | 0.98396 ↓ | -0.023 | 0.76839 | 0.89703 | 0.7465 ↓ | -0.422 |
| Complement C8 gamma chain | 0.64411 | 0.85716 | 0.91642 ↓ | -0.126 | 0.59522 | 0.80448 | 0.90882 ↓ | -0.138 | 0.35002 | 0.63692 | 0.85632 ↓ | -0.224 |
| Antithrombin-III | 0.53529 | 0.81174 | 1.111 ↑ | 0.1518 | 0.8913 | 0.95773 | 1.0027 ↑ | 0.0038 | 0.92055 | 0.98222 | 1.0201 ↑ | 0.0287 |
| C1q domain-containing protein | 0.98201 | 0.9855 | 1.0545 ↑ | 0.0765 | 0.61394 | 0.81078 | 0.98994 ↓ | -0.015 | 0.13673 | 0.35304 | 0.6797 ↓ | -0.557 |
| Epididymis secretory protein Li 37 | 0.091532 | 0.36821 | 1.5159 ↑ | 0.6002 | 0.25443 | 0.55936 | 0.54902 ↓ | -0.865 | 0.99179 | 0.99179 | 1.4656 ↑ | 0.5515 |
| Fibrinogen gamma chain | 0.28205 | 0.62899 | 0.71503 ↓ | -0.484 | 0.20824 | 0.48682 | 1.3922 ↑ | 0.4774 | 0.46025 | 0.70234 | 0.80127 ↓ | -0.32 |
| Apolipoprotein C-I, basic form | 0.022832 | 0.1975 | 0.25893 ↓ | -1.949 | 0.316 | 0.60074 | 0.69551 ↓ | -0.524 | 0.36245 | 0.63692 | 0.9212 ↓ | -0.118 |

| | | | | | | | | | | | | |
|---|---|---|---|---|---|---|---|---|---|---|---|---|
| Uncharacterized protein | 0.73039 | 0.8797 | 1.0328 ↑ | 0.0466 | 0.41351 | 0.6563 | 1.0781 ↑ | 0.1085 | 0.10863 | 0.31429 | 1.1839 ↑ | 0.2435 |
| Complement C4-B | 0.29721 | 0.62899 | 1.381 ↑ | 0.4657 | 0.49867 | 0.72973 | 1.2775 ↑ | 0.3533 | 0.022493 | 0.15565 | 1.853 ↑ | 0.8899 |
| SERPIN domain-containing protein | 0.74598 | 0.88393 | 0.87235 ↓ | -0.197 | 0.27081 | 0.56446 | 0.69224 ↓ | -0.531 | 0.22564 | 0.4647 | 0.65622 ↓ | -0.608 |
| Lumican | 0.059057 | 0.27613 | 0.57986 ↓ | -0.786 | 0.057324 | 0.24188 | 0.56629 ↓ | -0.82 | 0.014108 | 0.14357 | 0.41528 ↓ | -1.268 |
| Cadherin-13 | 0.67924 | 0.87693 | 0.8942 ↓ | -0.161 | 0.36356 | 0.62897 | 0.70442 ↓ | -0.505 | 0.31352 | 0.59602 | 1.3794 ↑ | 0.4641 |
| Angiotensin 1-10 | 0.29337 | 0.62899 | 1.1678 ↑ | 0.2238 | 0.51452 | 0.74177 | 1.1358 ↑ | 0.1838 | 0.46497 | 0.70234 | 1.2057 ↑ | 0.2698 |
| Keratin, type II cytoskeletal 2 epidermal | 0.61213 | 0.83384 | 0.733 ↓ | -0.448 | 0.98723 | 0.98723 | 0.87589 ↓ | -0.191 | 0.98796 | 0.99179 | 0.87578 ↓ | -0.191 |
| Fibulin-1 | 0.98279 | 0.9855 | 0.98677 ↓ | -0.019 | 0.87714 | 0.94841 | 1.0155 ↑ | 0.0223 | 0.29927 | 0.57525 | 0.753 ↓ | -0.409 |
| Trypsin | 0.58958 | 0.82256 | 0.92488 ↓ | -0.113 | 0.76883 | 0.91261 | 1.4581 ↑ | 0.5441 | 0.42774 | 0.70234 | 0.84749 ↓ | -0.239 |
| 1,4-beta-N-acetylmuramidase C | 0.37628 | 0.6711 | 1.3961 ↑ | 0.4814 | 0.95072 | 0.9841 | 1.2233 ↑ | 0.2908 | 0.13538 | 0.35304 | 1.7085 ↑ | 0.7728 |
| Macaca fascicularis brain cDNA clone: QorA-13734, similar to huma | 0.72734 | 0.8797 | 0.78044 ↓ | -0.358 | 0.81811 | 0.91522 | 0.86443 ↓ | -0.21 | 0.88448 | 0.97462 | 0.72603 ↓ | -0.462 |
| Testicular tissue protein Li 61 | 0.10627 | 0.39117 | 0.78046 ↓ | -0.358 | 0.039047 | 0.1917 | 0.66684 ↓ | -0.585 | 0.033827 | 0.18343 | 0.57687 ↓ | -0.794 |
| Uncharacterized protein | 0.54999 | 0.81174 | 0.93352 ↓ | -0.099 | 0.03851 | 0.1917 | 0.7335 ↓ | -0.447 | 0.026226 | 0.16799 | 0.69211 ↓ | -0.531 |
| Vitellogenin domain-containing protein | 0.13789 | 0.42557 | 0.60829 ↓ | -0.717 | 0.15255 | 0.41889 | 0.56571 ↓ | -0.822 | 0.06451 | 0.2408 | 0.75037 ↓ | -0.414 |
| Uncharacterized protein | 0.062472 | 0.27712 | 0.59792 ↓ | -0.742 | 0.007752 | 0.082654 | 0.4616 ↓ | -1.115 | 0.010688 | 0.14357 | 0.47505 ↓ | -1.074 |
| Serpin family A member 10 | 0.58694 | 0.82256 | 0.49149 ↓ | -1.025 | 0.77545 | 0.91261 | 0.65573 ↓ | -0.609 | 0.044473 | 0.22459 | 0.19613 ↓ | -2.35 |
| Complement component 8 subunit beta | 0.15549 | 0.46378 | 0.41186 ↓ | -1.28 | 0.11779 | 0.37051 | 0.38107 ↓ | -1.392 | 0.01043 | 0.14357 | 0.20884 ↓ | -2.26 |
| Collagen type VI alpha 3 chain | 0.62044 | 0.83857 | 0.96017 ↓ | -0.059 | 0.69897 | 0.88913 | 0.89447 ↓ | -0.161 | 0.155 | 0.37124 | 0.66685 ↓ | -0.585 |
| Glutathione peroxidase | 0.19466 | 0.51182 | 1.3625 ↑ | 0.4463 | 0.32525 | 0.61161 | 1.4856 ↑ | 0.5711 | 0.68143 | 0.83019 | 0.9407 ↓ | -0.088 |
| Mannan binding lectin serine peptidase 2 | 0.69749 | 0.8797 | 0.91791 ↓ | -0.124 | 0.14056 | 0.39864 | 0.73573 ↓ | -0.443 | 0.21155 | 0.45183 | 0.78834 ↓ | -0.343 |
| Protein S100 | 0.54458 | 0.81174 | 0.85546 ↓ | -0.225 | 0.20599 | 0.48682 | 0.72846 ↓ | -0.457 | 0.42743 | 0.70234 | 8.8616 ↑ | 3.1476 |
| Bradykinin | 0.42226 | 0.70923 | 0.82045 ↓ | -0.286 | 0.021971 | 0.16229 | 0.37623 ↓ | -1.41 | 0.40998 | 0.6886 | 0.79338 ↓ | -0.334 |
| Protein C receptor | 0.18926 | 0.51182 | 0.82216 ↓ | -0.283 | 0.26959 | 0.56446 | 0.81492 ↓ | -0.295 | 0.015683 | 0.14528 | 0.59736 ↓ | -0.743 |
| Protein-tyrosine-phosphatase | 0.81911 | 0.92647 | 1.0857 ↑ | 0.1186 | 0.25543 | 0.55936 | 1.3495 ↑ | 0.4324 | 0.49376 | 0.71183 | 1.169 ↑ | 0.2253 |
| Uncharacterized protein | 0.81191 | 0.92647 | 0.97124 ↓ | -0.042 | 0.82444 | 0.91522 | 0.9702 ↓ | -0.044 | 0.55907 | 0.75915 | 1.1825 ↑ | 0.2418 |
| Filamin A | 0.22275 | 0.5396 | 0.73135 ↓ | -0.451 | 0.34791 | 0.62049 | 0.77859 ↓ | -0.361 | 0.93966 | 0.98222 | 1.0469 ↑ | 0.0661 |
| Inter-alpha-trypsin inhibitor heavy chain 2 | 0.63218 | 0.84781 | 0.86257 ↓ | -0.213 | 0.10535 | 0.34388 | 0.61614 ↓ | -0.699 | 0.67478 | 0.82792 | 1.016 ↑ | 0.023 |
| Fibrinogen alpha chain | 0.98388 | 0.9855 | 0.96591 ↓ | -0.05 | 0.68999 | 0.88421 | 1.161 ↑ | 0.2154 | 0.88241 | 0.97462 | 0.98234 ↓ | -0.026 |
| Complement C1s | 0.96877 | 0.9855 | 1.0021 ↑ | 0.003 | 0.74284 | 0.91261 | 1.0301 ↑ | 0.0428 | 0.080765 | 0.2784 | 0.80474 ↓ | -0.313 |
| Mannosyl-oligosaccharide 1,2-alpha-mannosidase IA | 0.050306 | 0.25005 | 2.4703 ↑ | 1.3047 | 0.43311 | 0.66899 | 1.8654 ↑ | 0.8995 | 0.35579 | 0.63692 | 1.9625 ↑ | 0.9727 |
| Apolipoprotein B (Including Ag(X) antigen) | 0.55837 | 0.81174 | 0.83001 ↓ | -0.269 | 0.039646 | 0.1917 | 0.4888 ↓ | -1.033 | 0.085263 | 0.28366 | 0.53643 ↓ | -0.899 |
| Complement component C8 alpha chain | 0.34118 | 0.64789 | 1.1598 ↑ | 0.2139 | 0.50195 | 0.72973 | 0.8834 ↓ | -0.179 | 0.58362 | 0.75915 | 1.057 ↑ | 0.08 |
| cDNA FLJ44930 fis, clone BRAMY3015549, highly similar to Neural c | 0.29715 | 0.62899 | 0.61486 ↓ | -0.702 | 0.45237 | 0.68053 | 0.86425 ↓ | -0.21 | 0.76026 | 0.89703 | 0.65543 ↓ | -0.609 |
| Metavinculin | 0.71936 | 0.8797 | 0.69389 ↓ | -0.527 | 0.19231 | 0.47109 | 0.47766 ↓ | -1.066 | 0.56998 | 0.75915 | 0.67026 ↓ | -0.577 |
| ITIH4 protein | 0.1239 | 0.40152 | 3.7287 ↑ | 1.8987 | 0.048559 | 0.22107 | 3.6647 ↑ | 1.8737 | 0.050135 | 0.22459 | 3.7348 ↑ | 1.901 |
| Apolipoprotein C-II | 0.90878 | 0.98262 | 1.0818 ↑ | 0.1134 | 0.19465 | 0.47109 | 0.5142 ↓ | -0.96 | 0.17672 | 0.39195 | 0.46076 ↓ | -1.118 |
| Prothrombin | 0.96622 | 0.9855 | 0.60607 ↓ | -0.722 | 0.95566 | 0.9841 | 0.78059 ↓ | -0.357 | 0.58288 | 0.75915 | 0.51014 ↓ | -0.971 |
| Alpha-1-antitrypsin | 0.1362 | 0.42557 | 2.3374 ↑ | 1.2249 | 0.33549 | 0.62049 | 0.91583 ↓ | -0.127 | 0.033929 | 0.18343 | 2.1704 ↑ | 1.1179 |
| Complement C1q subcomponent subunit B | 0.22536 | 0.5396 | 0.76539 ↓ | -0.386 | 0.37578 | 0.63116 | 0.84314 ↓ | -0.246 | 0.92525 | 0.98222 | 0.97109 ↓ | -0.042 |
| SEMA4D variant protein | 0.19749 | 0.51182 | 0.77204 ↓ | -0.373 | 0.40151 | 0.64673 | 0.85337 ↓ | -0.229 | 0.46147 | 0.70234 | 0.92453 ↓ | -0.113 |
| CP protein | 0.42151 | 0.70923 | 1.1106 ↑ | 0.1513 | 0.30636 | 0.58889 | 0.75198 ↓ | -0.411 | 0.8552 | 0.96072 | 1.165 ↑ | 0.2204 |
| Attractin | 0.1866 | 0.51182 | 0.79368 ↓ | -0.333 | 0.050201 | 0.22243 | 0.67212 ↓ | -0.573 | 0.020631 | 0.15518 | 0.58448 ↓ | -0.775 |
| 15S Mg(2+)-ATPase p97 subunit | 0.84995 | 0.94607 | 0.87002 ↓ | -0.201 | 0.25101 | 0.55936 | 0.58549 ↓ | -0.772 | 0.94158 | 0.98222 | 1.2767 ↑ | 0.3525 |
| CST6 protein | 0.4977 | 0.78993 | 1.2262 ↑ | 0.2942 | 0.56105 | 0.77649 | 1.2335 ↑ | 0.3028 | 0.36192 | 0.63692 | 0.85264 ↓ | -0.23 |
| Monocyte differentiation antigen CD14 | 0.65727 | 0.868 | 1.0161 ↑ | 0.023 | 0.28993 | 0.57074 | 0.86172 ↓ | -0.215 | 0.66169 | 0.82354 | 0.95888 ↓ | -0.061 |
| Uncharacterized protein | 0.16146 | 0.47344 | 0.61739 ↓ | -0.696 | 0.090534 | 0.33324 | 0.54635 ↓ | -0.872 | 0.065419 | 0.2408 | 0.52782 ↓ | -0.922 |
| Immunoglobulin heavy constant mu | 0.20414 | 0.51182 | 1.2331 ↑ | 0.3023 | 0.37052 | 0.62972 | 0.28966 ↓ | -1.788 | 0.80884 | 0.92669 | 0.4664 ↓ | -1.1 |
| Apolipoprotein A4 | 0.54323 | 0.81174 | 1.1011 ↑ | 0.139 | 0.81067 | 0.91522 | 0.96767 ↓ | -0.047 | 0.048092 | 0.22459 | 0.43164 ↓ | -1.212 |
| Ficolin (Collagen/fibrinogen domain containing) 3 (Hakata antigen), | 0.9504 | 0.9855 | 0.90136 ↓ | -0.15 | 0.44171 | 0.67624 | 0.69764 ↓ | -0.519 | 0.2601 | 0.52322 | 0.5192 ↓ | -0.946 |
| Receptor protein-tyrosine kinase | 0.95919 | 0.9855 | 1.0617 ↑ | 0.0864 | 0.28711 | 0.57074 | 0.86752 ↓ | -0.205 | 0.10423 | 0.31381 | 0.62087 ↓ | -0.688 |
| Insulin-like growth factor-binding protein 3 | 0.32326 | 0.64626 | 0.78905 ↓ | -0.342 | 0.18871 | 0.47109 | 0.66749 ↓ | -0.583 | 0.43832 | 0.70234 | 0.81376 ↓ | -0.297 |
| Uncharacterized protein DKFZp686I15196 | 0.76157 | 0.89627 | 0.96169 ↓ | -0.056 | 0.55506 | 0.77649 | 0.72906 ↓ | -0.456 | 0.45038 | 0.70234 | 2.386 ↑ | 1.2546 |
| cDNA, FLJ93914, highly similar to Homo sapiens histidine-rich glyco | 0.050587 | 0.25005 | 0.56998 ↓ | -0.811 | 0.015201 | 0.13841 | 0.52484 ↓ | -0.93 | 0.012503 | 0.14357 | 0.37397 ↓ | -1.419 |

| | | | | | | | | | | | | |
|---|---|---|---|---|---|---|---|---|---|---|---|---|
| Coagulation factor XII | 0.9538 | 0.9855 | 0.81204 ↓ | -0.3 | 0.39494 | 0.64457 | 1.026 ↑ | 0.0371 | 0.58173 | 0.75915 | 0.66297 ↓ | -0.593 |
| Protein S100-A7 | 0.21312 | 0.5267 | 0.3094 ↓ | -1.693 | 0.56079 | 0.77649 | 0.4452 ↓ | -1.168 | 0.69474 | 0.84049 | 19.969 ↑ | 4.3197 |
| Insulin-like growth factor binding protein, acid labile subunit | 0.41958 | 0.70923 | 0.87135 ↓ | -0.199 | 0.29032 | 0.57074 | 0.82232 ↓ | -0.282 | 0.02 | 0.15518 | 0.64472 ↓ | -0.633 |
| Carboxylic ester hydrolase | 0.31864 | 0.64626 | 0.78071 ↓ | -0.357 | 0.043641 | 0.20405 | 0.58952 ↓ | -0.762 | 0.062301 | 0.2408 | 0.60791 ↓ | -0.718 |

**Supplementary Table 6**: **List of putative serum biomarkers of radiation exposure in NHPs.**

| Name | 6 Gy | | | | 6.5 Gy | | | | 7 Gy | | | | 7.5 Gy | | | | 8 Gy | | | | 8.5 Gy | | | |
|---|---|---|---|---|---|---|---|---|---|---|---|---|---|---|---|---|---|---|---|---|---|---|---|---|
| | p.value | FDR | Fold Change | log2(FC) | p.value | FDR | Fold Change | log2(FC) | p.value | FDR | Fold Change | log2(FC) | p.value | FDR | Fold Change | log2(FC) | p.value | FDR | Fold Change | log2(FC) | p.value | FDR | Fold Change | log2(FC) |
| H2R974 | 3.89E-16 | 9.88E-14 | 3.61 | ↑1.85 | 1.17E-17 | 2.98E-15 | 3.89 | ↑1.96 | 6.24E-17 | 1.59E-14 | 3.82 | ↑1.93 | 6.10E-18 | 1.55E-15 | 4.00 | ↑2.00 | 3.47E-17 | 8.81E-15 | 3.53 | ↑1.82 | 1.54E-11 | 2.79E-10 | 3.54 | ↑1.82 |
| L-Carnitine | 7.72E-14 | 9.80E-12 | 2.41 | ↑1.27 | 2.99E-13 | 2.54E-11 | 2.42 | ↑1.28 | 3.52E-15 | 4.47E-13 | 2.51 | ↑1.33 | 2.10E-10 | 5.92E-09 | 2.11 | ↑1.08 | 3.52E-11 | 1.62E-09 | 2.12 | ↑1.08 | 3.57E-16 | 9.08E-14 | 2.98 | ↑1.57 |
| LysoPC160 | 5.49E-13 | 4.65E-11 | 0.40 | ↓-1.33 | 1.20E-07 | 2.53E-06 | 0.56 | ↓-0.84 | 2.29E-07 | 2.77E-06 | 0.56 | ↓-0.83 | 1.46E-08 | 2.07E-07 | 0.50 | ↓-1.01 | 4.29E-02 | 7.52E-02 | 0.88 | ↓-0.18 | 9.22E-10 | 9.36E-09 | 0.48 | ↓-1.05 |
| Phenylpiperazine | 1.58E-12 | 1.01E-10 | 2.38 | ↑1.25 | 4.07E-12 | 2.58E-10 | 2.36 | ↑1.24 | 1.56E-14 | 1.32E-12 | 2.44 | ↑1.29 | 3.12E-09 | 6.09E-08 | 2.03 | ↑1.02 | 1.49E-10 | 4.74E-09 | 2.07 | ↑1.05 | 3.52E-15 | 4.47E-13 | 2.89 | ↑1.53 |
| Lyso-PAF C-18 | 2.06E-12 | 1.04E-10 | 0.55 | ↓-0.87 | 1.37E-11 | 6.98E-10 | 0.55 | ↓-0.86 | 1.39E-08 | 2.53E-07 | 0.60 | ↓-0.73 | 5.19E-16 | 6.59E-14 | 0.45 | ↓-1.15 | 1.95E-07 | 2.36E-06 | 0.63 | ↓-0.67 | 5.16E-08 | 2.73E-07 | 0.61 | ↓-0.71 |
| 4-Quinolinol, 8-methyl-2-trifluoromethyl- | 1.56E-11 | 6.60E-10 | 0.26 | ↓-1.96 | 7.50E-06 | 6.15E-05 | 0.44 | ↓-1.20 | 1.51E-12 | 9.58E-11 | 0.27 | ↓-1.89 | 1.96E-12 | 1.66E-10 | 0.20 | ↓-2.33 | 1.79E-07 | 2.27E-06 | 0.37 | ↓-1.42 | 2.39E-13 | 1.52E-11 | 0.25 | ↓-1.97 |
| PC160/180 | 1.11E-10 | 3.97E-09 | 3.99 | ↑2.00 | 2.23E-04 | 8.19E-04 | 2.82 | ↑1.49 | 2.01E-04 | 7.35E-04 | 2.72 | ↑1.44 | 3.64E-05 | 1.44E-04 | 2.62 | ↑1.39 | 2.31E-05 | 9.94E-05 | 2.92 | ↑1.55 | 1.91E-08 | 1.21E-07 | 3.82 | ↑1.94 |
| PC160/204 | 1.25E-10 | 3.97E-09 | 3.34 | ↑1.74 | 5.90E-05 | 3.06E-04 | 2.27 | ↑1.18 | 4.04E-05 | 1.83E-04 | 2.33 | ↑1.22 | 2.29E-07 | 2.15E-06 | 2.62 | ↑1.39 | 7.30E-05 | 2.51E-04 | 2.53 | ↑1.34 | 9.75E-09 | 6.69E-08 | 3.06 | ↑1.61 |
| PE180/204 | 3.89E-10 | 1.10E-08 | 3.38 | ↑1.76 | 1.29E-04 | 5.45E-04 | 2.24 | ↑1.16 | 1.04E-04 | 4.27E-04 | 2.30 | ↑1.20 | 1.23E-06 | 8.65E-06 | 2.70 | ↑1.43 | 1.42E-04 | 4.57E-04 | 2.57 | ↑1.36 | 4.40E-09 | 3.72E-08 | 3.18 | ↑1.67 |
| PS180/181 | 8.68E-10 | 2.20E-08 | 0.48 | ↓-1.05 | 2.85E-05 | 1.80E-04 | 0.67 | ↓-0.57 | 1.22E-08 | 2.38E-07 | 0.58 | ↓-0.79 | 1.08E-07 | 1.05E-06 | 0.58 | ↓-0.79 | 4.91E-06 | 2.97E-05 | 0.65 | ↓-0.63 | 5.74E-09 | 4.56E-08 | 0.56 | ↓-0.84 |
| PC181/160 | 1.20E-09 | 2.57E-08 | 3.89 | ↑1.96 | 3.42E-04 | 1.14E-03 | 2.47 | ↑1.30 | 2.10E-04 | 7.48E-04 | 2.42 | ↑1.28 | 8.89E-06 | 4.99E-05 | 2.82 | ↑1.49 | 8.33E-04 | 2.25E-03 | 2.89 | ↑1.53 | 2.89E-08 | 1.71E-07 | 3.86 | ↑1.95 |
| LysoPE180 | 1.26E-09 | 2.57E-08 | 0.52 | ↓-0.93 | 2.45E-07 | 4.44E-06 | 0.60 | ↓-0.73 | 2.84E-07 | 3.28E-06 | 0.59 | ↓-0.77 | 3.46E-11 | 1.91E-09 | 0.44 | ↓-1.20 | 4.70E-06 | 2.91E-05 | 0.61 | ↓-0.72 | 4.44E-08 | 2.45E-07 | 0.55 | ↓-0.86 |
| SMd181/160 | 1.31E-09 | 2.57E-08 | 5.68 | ↑2.51 | 8.71E-06 | 6.90E-05 | 4.96 | ↑2.31 | 3.63E-06 | 2.56E-05 | 3.90 | ↑1.96 | 2.28E-06 | 1.57E-05 | 5.63 | ↑2.49 | 5.38E-06 | 3.18E-05 | 5.45 | ↑2.45 | 4.29E-10 | 5.19E-09 | 8.87 | ↑3.15 |
| -Galactosyl-N-palmitoylsphingosine | 2.29E-09 | 4.15E-08 | 2.65 | ↑1.40 | 1.20E-03 | 3.28E-03 | 1.83 | ↑0.87 | 1.41E-04 | 5.43E-04 | 2.08 | ↑1.06 | 2.37E-04 | 7.24E-04 | 1.96 | ↑0.97 | 5.13E-04 | 1.43E-03 | 2.03 | ↑1.02 | 2.13E-07 | 9.65E-07 | 2.78 | ↑1.48 |
| PC180/181 | 2.66E-09 | 4.44E-08 | 3.52 | ↑1.81 | 1.66E-05 | 1.21E-04 | 2.23 | ↑1.16 | 1.16E-04 | 4.63E-04 | 3.41 | ↑1.77 | 1.03E-07 | 1.05E-06 | 3.15 | ↑1.66 | 2.54E-05 | 1.08E-04 | 2.57 | ↑1.36 | 4.25E-12 | 1.08E-10 | 4.38 | ↑2.13 |
| PI180/226 | 2.84E-09 | 4.44E-08 | 0.59 | ↓-0.76 | 6.34E-07 | 9.11E-06 | 0.67 | ↓-0.57 | 1.02E-08 | 2.38E-07 | 0.64 | ↓-0.65 | 2.83E-05 | 1.20E-04 | 0.72 | ↓-0.48 | 1.98E-06 | 1.74E-05 | 0.69 | ↓-0.53 | 6.11E-09 | 4.70E-08 | 0.64 | ↓-0.65 |
| PE180/182 | 3.07E-09 | 4.44E-08 | 2.53 | ↑1.34 | 5.42E-04 | 1.68E-03 | 1.85 | ↑0.89 | 4.04E-04 | 1.32E-03 | 1.92 | ↑0.94 | 2.67E-04 | 8.07E-04 | 1.90 | ↑0.92 | 5.18E-05 | 1.88E-04 | 2.11 | ↑1.07 | 2.69E-07 | 1.18E-06 | 2.53 | ↑1.34 |
| C18Plasm-204 PC | 3.17E-09 | 4.44E-08 | 5.38 | ↑2.43 | 7.72E-04 | 2.25E-03 | 3.62 | ↑1.86 | 5.42E-04 | 1.70E-03 | 2.87 | ↑1.52 | 6.28E-05 | 2.28E-04 | 2.65 | ↑1.41 | 3.95E-04 | 1.16E-03 | 3.46 | ↑1.79 | 1.90E-08 | 1.21E-07 | 4.82 | ↑2.27 |
| Acetyl-DL-carnitine | 3.47E-09 | 4.44E-08 | 4.02 | ↑2.01 | 1.13E-04 | 4.94E-04 | 2.04 | ↑1.03 | 1.11E-08 | 2.38E-07 | 2.34 | ↑1.23 | 6.01E-04 | 1.64E-03 | 2.12 | ↑1.09 | 3.32E-04 | 9.91E-04 | 2.47 | ↑1.31 | 6.63E-07 | 2.63E-06 | 3.00 | ↑1.58 |
| P18659 | 3.50E-09 | 4.44E-08 | 0.41 | ↓-1.30 | 2.06E-08 | 5.80E-07 | 0.43 | ↓-1.20 | 1.50E-09 | 5.46E-08 | 0.38 | ↓-1.38 | 5.85E-10 | 1.35E-08 | 0.37 | ↓-1.44 | 4.43E-14 | 5.63E-12 | 0.28 | ↓-1.83 | 1.36E-09 | 1.28E-08 | 0.37 | ↓-1.45 |
| B3KS79 | 4.21E-09 | 5.09E-08 | 3.30 | ↑1.72 | 8.04E-10 | 2.92E-08 | 3.27 | ↑1.71 | 4.79E-10 | 2.03E-08 | 3.47 | ↑1.79 | 4.22E-11 | 1.91E-09 | 3.37 | ↑1.75 | 2.84E-12 | 2.40E-10 | 3.34 | ↑1.74 | 4.24E-07 | 1.79E-06 | 2.94 | ↑1.56 |
| PI160/181 | 5.76E-09 | 6.65E-08 | 3.51 | ↑1.81 | 2.36E-04 | 8.58E-04 | 2.44 | ↑1.29 | 2.12E-04 | 7.48E-04 | 2.79 | ↑1.48 | 3.31E-05 | 1.33E-04 | 3.30 | ↑1.72 | 4.13E-05 | 1.59E-04 | 3.70 | ↑1.89 | 1.55E-07 | 7.42E-07 | 3.47 | ↑1.80 |
| Lysophosphatidylethanolamine | 1.36E-08 | 1.50E-07 | 0.56 | ↓-0.85 | 7.29E-06 | 6.15E-05 | 0.64 | ↓-0.65 | 7.22E-06 | 4.27E-05 | 0.63 | ↓-0.67 | 6.40E-10 | 1.36E-08 | 0.49 | ↓-1.04 | 8.58E-06 | 4.52E-05 | 0.63 | ↓-0.67 | 4.93E-07 | 1.99E-06 | 0.59 | ↓-0.77 |
| C16-181 PC | 2.07E-08 | 2.19E-07 | 4.35 | ↑2.12 | 2.50E-04 | 8.95E-04 | 2.47 | ↑1.30 | 5.34E-04 | 1.70E-03 | 4.23 | ↑2.08 | 6.26E-08 | 6.92E-07 | 3.30 | ↑1.72 | 2.32E-06 | 1.90E-05 | 2.97 | ↑1.57 | 1.72E-12 | 6.22E-11 | 5.21 | ↑2.38 |
| LysoPC150 | 2.82E-08 | 2.87E-07 | 0.62 | ↓-0.68 | 5.37E-05 | 2.84E-04 | 0.71 | ↓-0.49 | 3.14E-06 | 2.28E-05 | 0.66 | ↓-0.59 | 6.98E-09 | 1.04E-07 | 0.57 | ↓-0.82 | 2.32E-04 | 7.20E-04 | 0.73 | ↓-0.44 | 3.99E-05 | 1.17E-04 | 0.72 | ↓-0.47 |
| PE180/240 | 3.51E-08 | 3.43E-07 | 4.46 | ↑2.16 | 2.62E-04 | 9.10E-04 | 2.95 | ↑1.56 | 3.47E-04 | 1.15E-03 | 3.25 | ↑1.70 | 8.80E-05 | 3.15E-04 | 4.05 | ↑2.02 | 2.51E-04 | 7.70E-04 | 4.70 | ↑2.23 | 2.00E-07 | 9.40E-07 | 4.45 | ↑2.15 |
| PC181/140 | 3.78E-08 | 3.55E-07 | 5.19 | ↑2.38 | 2.91E-05 | 1.80E-04 | 4.63 | ↑2.21 | 9.52E-06 | 5.38E-05 | 3.43 | ↑1.78 | 9.04E-06 | 4.99E-05 | 4.87 | ↑2.28 | 1.42E-05 | 6.57E-05 | 5.58 | ↑2.48 | 6.48E-09 | 4.84E-08 | 6.40 | ↑2.68 |
| H2QIN8 | 6.64E-08 | 5.92E-07 | 0.73 | ↓-0.45 | 8.00E-10 | 2.92E-08 | 0.67 | ↓-0.57 | 1.39E-07 | 2.07E-06 | 0.54 | ↓-0.90 | 3.35E-07 | 3.03E-06 | 0.69 | ↓-0.53 | 3.83E-11 | 1.62E-09 | 0.63 | ↓-0.66 | 4.70E-08 | 2.54E-07 | 0.65 | ↓-0.63 |
| Decanoyl-L-carnitine | 6.76E-08 | 5.92E-07 | 4.68 | ↑2.23 | 3.12E-05 | 1.87E-04 | 4.05 | ↑2.02 | 9.25E-05 | 3.85E-04 | 3.72 | ↑1.89 | 2.27E-05 | 1.04E-04 | 3.50 | ↑1.81 | 5.18E-03 | 1.11E-02 | 2.80 | ↑1.49 | 1.30E-03 | 2.93E-03 | 2.65 | ↑1.41 |
| Arachidonoyl PAF C-16 | 7.44E-08 | 6.30E-07 | 4.62 | ↑2.21 | 1.10E-04 | 4.94E-04 | 2.73 | ↑1.45 | 8.89E-04 | 2.57E-03 | 3.51 | ↑1.81 | 7.68E-08 | 8.13E-07 | 3.46 | ↑1.79 | 3.49E-06 | 2.53E-05 | 3.02 | ↑1.60 | 3.96E-12 | 1.08E-10 | 4.85 | ↑2.28 |
| Suberylglycine | 8.36E-08 | 6.85E-07 | 4.78 | ↑2.26 | 2.80E-13 | 2.54E-11 | 5.88 | ↑2.56 | 3.25E-12 | 1.65E-10 | 6.28 | ↑2.65 | 1.05E-10 | 3.34E-09 | 6.01 | ↑2.59 | 2.74E-10 | 7.73E-09 | 5.42 | ↑2.44 | 7.07E-12 | 1.63E-10 | 5.67 | ↑2.50 |
| Glutamic Acid | 8.98E-08 | 7.12E-07 | 0.42 | ↓-1.25 | 8.97E-05 | 4.30E-04 | 0.54 | ↓-0.89 | 1.16E-08 | 2.38E-07 | 0.39 | ↓-1.36 | 4.50E-11 | 1.91E-09 | 0.29 | ↓-1.80 | 7.19E-07 | 7.31E-06 | 0.45 | ↓-1.15 | 9.71E-12 | 2.05E-10 | 0.31 | ↓-1.71 |
| Thioetheramidephosphatidylcholine | 1.15E-07 | 8.83E-07 | 3.82 | ↑1.93 | 2.35E-06 | 2.49E-05 | 2.69 | ↑1.43 | 3.46E-05 | 1.63E-04 | 5.09 | ↑2.35 | 2.05E-05 | 9.66E-05 | 3.52 | ↑1.82 | 1.07E-06 | 1.03E-05 | 3.14 | ↑1.65 | 2.83E-12 | 8.99E-11 | 5.90 | ↑2.56 |
| PC160/160 | 1.32E-07 | 9.20E-07 | 2.94 | ↑1.56 | 7.60E-07 | 1.01E-05 | 2.23 | ↑1.16 | 1.08E-05 | 5.94E-05 | 3.35 | ↑1.74 | 2.40E-05 | 1.05E-04 | 2.58 | ↑1.37 | 6.17E-07 | 6.53E-06 | 2.45 | ↑1.29 | 1.45E-12 | 6.15E-11 | 3.73 | ↑1.90 |
| Eicosapentaenoyl PAF C-16 | 1.33E-07 | 9.20E-07 | 3.84 | ↑1.94 | 1.73E-03 | 4.45E-03 | 2.24 | ↑1.17 | 2.01E-03 | 5.05E-03 | 4.19 | ↑2.07 | 2.41E-06 | 1.61E-05 | 2.84 | ↑1.51 | 1.10E-05 | 5.49E-05 | 2.90 | ↑1.54 | 3.15E-11 | 5.33E-10 | 5.18 | ↑2.37 |
| PC | 1.33E-07 | 9.20E-07 | 2.91 | ↑1.54 | 4.11E-04 | 1.34E-03 | 1.96 | ↑0.97 | 8.51E-05 | 3.60E-04 | 2.73 | ↑1.45 | 4.46E-06 | 2.70E-05 | 2.47 | ↑1.31 | 7.07E-06 | 3.92E-05 | 2.30 | ↑1.20 | 1.77E-10 | 2.36E-09 | 3.46 | ↑1.79 |
| C18Plasm-226 PE | 1.34E-07 | 9.20E-07 | 5.31 | ↑2.41 | 5.02E-05 | 2.77E-04 | 4.88 | ↑2.29 | 1.88E-05 | 8.99E-05 | 3.32 | ↑1.73 | 2.29E-05 | 1.04E-04 | 5.15 | ↑2.37 | 9.89E-06 | 5.02E-05 | 5.01 | ↑2.33 | 1.11E-08 | 7.44E-08 | 8.41 | ↑3.07 |
| Tris2-butoxyethyl phosphate | 1.38E-07 | 9.20E-07 | 2.20 | ↑1.14 | 2.13E-09 | 6.76E-08 | 2.44 | ↑1.28 | 1.80E-09 | 5.72E-08 | 2.33 | ↑1.22 | 3.25E-06 | 2.07E-05 | 2.81 | ↑1.49 | 1.49E-09 | 3.43E-08 | 2.13 | ↑1.09 | 2.45E-08 | 1.52E-07 | 2.22 | ↑1.15 |
| PC181/181 | 2.07E-07 | 1.33E-06 | 2.51 | ↑1.33 | 2.07E-03 | 5.16E-03 | 2.01 | ↑1.01 | 2.43E-03 | 5.89E-03 | 1.88 | ↑0.91 | 2.48E-03 | 5.83E-03 | 1.77 | ↑0.82 | 2.42E-03 | 5.81E-03 | 2.04 | ↑1.03 | 1.15E-06 | 4.34E-06 | 2.73 | ↑1.45 |
| L-Glutamine, N2-2-phenylacetyl- | 2.14E-07 | 1.33E-06 | 0.44 | ↓-1.20 | 3.08E-04 | 1.06E-03 | 0.57 | ↓-0.81 | 9.33E-09 | 2.38E-07 | 0.39 | ↓-1.38 | 5.64E-11 | 2.05E-09 | 0.29 | ↓-1.79 | 2.53E-06 | 2.01E-05 | 0.47 | ↓-1.08 | 9.54E-11 | 1.35E-09 | 0.33 | ↓-1.62 |
| PC160/226 | 2.14E-07 | 1.33E-06 | 3.07 | ↑1.62 | 2.11E-05 | 1.37E-04 | 2.73 | ↑1.45 | 2.72E-06 | 2.09E-05 | 2.52 | ↑1.33 | 2.60E-06 | 1.69E-05 | 3.28 | ↑1.71 | 3.86E-05 | 1.52E-04 | 3.12 | ↑1.64 | 6.43E-10 | 6.96E-09 | 4.08 | ↑2.03 |
| PC160/181 | 2.83E-07 | 1.71E-06 | 2.39 | ↑1.26 | 1.48E-04 | 5.98E-04 | 1.81 | ↑0.86 | 1.14E-03 | 3.18E-03 | 2.01 | ↑1.01 | 4.04E-05 | 1.58E-04 | 1.90 | ↑0.93 | 8.02E-06 | 4.34E-05 | 1.83 | ↑0.87 | 2.71E-10 | 3.44E-09 | 2.50 | ↑1.32 |
| Arachidonoylthio-PC | 2.96E-07 | 1.75E-06 | 2.38 | ↑1.25 | 4.13E-06 | 3.74E-05 | 2.13 | ↑1.09 | 3.87E-07 | 4.28E-06 | 2.55 | ↑1.35 | 4.12E-08 | 5.51E-07 | 2.93 | ↑1.55 | 4.99E-07 | 5.52E-06 | 2.49 | ↑1.32 | 1.44E-14 | 1.22E-12 | 3.67 | ↑1.87 |
| PS180/226 | 3.05E-07 | 1.76E-06 | 3.01 | ↑1.59 | 1.65E-03 | 4.31E-03 | 2.13 | ↑1.09 | 2.26E-03 | 5.55E-03 | 2.42 | ↑1.28 | 4.05E-04 | 1.16E-03 | 2.77 | ↑1.47 | 1.09E-03 | 2.77E-03 | 3.09 | ↑1.63 | 1.67E-06 | 5.73E-06 | 3.20 | ↑1.68 |
| PC181/180 | 3.21E-07 | 1.81E-06 | 2.79 | ↑1.48 | 1.73E-03 | 4.45E-03 | 2.19 | ↑1.13 | 2.75E-03 | 6.47E-03 | 2.04 | ↑1.03 | 3.06E-03 | 7.01E-03 | 1.91 | ↑0.93 | 2.83E-03 | 6.41E-03 | 2.26 | ↑1.18 | 1.08E-06 | 4.15E-06 | 3.10 | ↑1.63 |
| C18Plasm LPC | 3.46E-07 | 1.91E-06 | 3.80 | ↑1.93 | 2.10E-06 | 2.32E-05 | 3.78 | ↑1.92 | 3.06E-08 | 4.85E-07 | 4.08 | ↑2.03 | 8.92E-04 | 2.36E-03 | 2.27 | ↑1.19 | 3.85E-06 | 2.65E-05 | 2.38 | ↑1.25 | 3.56E-06 | 1.16E-05 | 2.99 | ↑1.58 |
| PE160/204 | 4.51E-07 | 2.44E-06 | 3.40 | ↑1.76 | 1.77E-03 | 4.50E-03 | 2.54 | ↑1.34 | 2.27E-03 | 5.55E-03 | 2.68 | ↑1.42 | 2.03E-04 | 6.53E-04 | 3.24 | ↑1.69 | 2.93E-04 | 8.87E-04 | 3.87 | ↑1.95 | 2.42E-06 | 7.98E-06 | 3.52 | ↑1.82 |
| PC160/182 | 6.05E-07 | 3.20E-06 | 2.09 | ↑1.07 | 1.50E-03 | 3.96E-03 | 1.86 | ↑0.90 | 1.17E-04 | 4.63E-04 | 1.88 | ↑0.91 | 2.01E-04 | 6.53E-04 | 2.09 | ↑1.06 | 8.66E-04 | 2.29E-03 | 2.16 | ↑1.11 | 2.07E-07 | 9.58E-07 | 2.69 | ↑1.43 |
| Acetylpodocarpic acid anhydride | 6.30E-07 | 3.26E-06 | 1.65 | ↑0.72 | 1.54E-04 | 6.10E-04 | 1.47 | ↑0.56 | 6.23E-07 | 5.86E-06 | 1.67 | ↑0.74 | 8.95E-05 | 3.16E-04 | 1.65 | ↑0.72 | 3.70E-05 | 1.49E-04 | 1.59 | ↑0.66 | 4.28E-05 | 1.24E-04 | 1.69 | ↑0.76 |
| DMPE181/181 | 7.09E-07 | 3.60E-06 | 2.73 | ↑1.45 | 2.74E-03 | 6.50E-03 | 2.15 | ↑1.10 | 4.26E-03 | 9.75E-03 | 1.97 | ↑0.98 | 4.45E-03 | 9.73E-03 | 1.91 | ↑0.93 | 2.46E-03 | 5.85E-03 | 2.23 | ↑1.16 | 1.17E-06 | 4.34E-06 | 3.23 | ↑1.69 |
| PC180/226 | 1.02E-06 | 5.07E-06 | 3.26 | ↑1.70 | 1.45E-04 | 5.98E-04 | 2.96 | ↑1.57 | 5.82E-05 | 2.55E-04 | 2.64 | ↑1.40 | 5.71E-05 | 2.13E-04 | 3.43 | ↑1.78 | 2.01E-04 | 6.37E-04 | 3.21 | ↑1.68 | 4.80E-09 | 3.93E-08 | 4.69 | ↑2.23 |
| LysoPC170 | 1.11E-06 | 5.42E-06 | 0.67 | ↓-0.58 | 1.88E-06 | 2.17E-05 | 0.67 | ↓-0.58 | 5.21E-07 | 5.44E-06 | 0.65 | ↓-0.63 | 4.95E-09 | 7.86E-08 | 0.58 | ↓-0.79 | 1.18E-05 | 5.74E-05 | 0.69 | ↓-0.54 | 1.84E-02 | 3.30E-02 | 0.84 | ↓-0.24 |
| PE180/181 | 1.16E-06 | 5.56E-06 | 3.74 | ↑1.90 | 8.52E-04 | 2.43E-03 | 2.03 | ↑1.02 | 1.38E-03 | 3.65E-03 | 4.98 | ↑2.32 | 3.19E-05 | 1.31E-04 | 3.69 | ↑1.88 | 5.20E-05 | 1.88E-04 | 3.57 | ↑1.84 | 6.91E-11 | 1.03E-09 | 4.45 | ↑2.15 |
| H2QPN7 | 1.20E-06 | 5.65E-06 | 0.60 | ↓-0.74 | 2.55E-06 | 2.59E-05 | 0.54 | ↓-0.88 | 4.32E-03 | 9.80E-03 | 0.70 | ↓-0.52 | 6.83E-07 | 5.98E-06 | 0.57 | ↓-0.81 | 1.25E-10 | 4.55E-09 | 0.46 | ↓-1.13 | 3.38E-04 | 8.18E-04 | 0.57 | ↓-0.80 |

| | | | | | | | | | | | | | | | | | | | | | | | | |
|---|---|---|---|---|---|---|---|---|---|---|---|---|---|---|---|---|---|---|---|---|---|---|---|---|
| PE180/226 | 1.33E-06 | 6.13E-06 | 2.78 | ↑1.47 | 1.85E-05 | 1.31E-04 | 2.63 | ↑1.39 | 4.83E-06 | 3.15E-05 | 2.28 | ↑1.19 | 1.49E-05 | 7.73E-05 | 3.02 | ↑1.59 | 4.51E-05 | 1.71E-04 | 2.89 | ↑1.53 | 7.43E-09 | 5.24E-08 | 3.58 | ↑1.84 |
| PC180/140 | 1.36E-06 | 6.17E-06 | 3.24 | ↑1.70 | 2.74E-03 | 6.50E-03 | 2.23 | ↑1.16 | 6.40E-04 | 1.96E-03 | 2.54 | ↑1.34 | 5.88E-04 | 1.62E-03 | 2.92 | ↑1.55 | 1.05E-03 | 2.69E-03 | 3.20 | ↑1.68 | 1.36E-06 | 4.81E-06 | 3.26 | ↑1.70 |
| Palmitoyl Thio-PC | 1.45E-06 | 6.38E-06 | 3.06 | ↑1.61 | 5.21E-04 | 1.63E-03 | 2.36 | ↑1.24 | 6.64E-04 | 2.01E-03 | 3.94 | ↑1.98 | 7.74E-07 | 6.55E-06 | 3.99 | ↑2.00 | 1.90E-06 | 1.73E-05 | 3.33 | ↑1.74 | 3.43E-11 | 5.45E-10 | 5.49 | ↑2.46 |
| PE160/226 | 1.46E-06 | 6.38E-06 | 4.35 | ↑2.12 | 1.18E-04 | 5.08E-04 | 3.62 | ↑1.86 | 2.03E-04 | 7.35E-04 | 3.54 | ↑1.82 | 1.51E-04 | 4.99E-04 | 5.89 | ↑2.56 | 4.85E-05 | 1.81E-04 | 6.45 | ↑2.69 | 2.68E-08 | 1.62E-07 | 6.05 | ↑2.60 |
| H2QMF6 | 1.98E-06 | 8.51E-06 | 0.65 | ↓-0.63 | 1.05E-05 | 7.82E-05 | 0.69 | ↓-0.53 | 1.08E-06 | 9.79E-06 | 0.65 | ↓-0.62 | 1.18E-06 | 8.64E-06 | 0.67 | ↓-0.59 | 3.51E-05 | 1.44E-04 | 0.70 | ↓-0.52 | 2.00E-04 | 4.99E-04 | 0.72 | ↓-0.48 |
| Hippuric acid | 2.04E-06 | 8.66E-06 | 0.46 | ↓-1.11 | 1.05E-04 | 4.93E-04 | 0.53 | ↓-0.92 | 3.14E-06 | 2.28E-05 | 0.48 | ↓-1.06 | 9.42E-07 | 7.48E-06 | 0.43 | ↓-1.21 | 8.44E-05 | 2.86E-04 | 0.52 | ↓-0.94 | 4.83E-05 | 1.36E-04 | 0.58 | ↓-0.78 |
| PI181 | 2.65E-06 | 1.10E-05 | 0.67 | ↓-0.58 | 2.08E-05 | 1.37E-04 | 0.72 | ↓-0.47 | 1.81E-05 | 8.87E-05 | 0.72 | ↓-0.48 | 2.82E-04 | 8.34E-04 | 0.78 | ↓-0.36 | 1.24E-05 | 5.94E-05 | 0.74 | ↓-0.44 | 3.60E-09 | 3.16E-08 | 0.58 | ↓-0.78 |
| PC180/182 | 2.79E-06 | 1.13E-05 | 2.25 | ↑1.17 | 4.82E-02 | 8.06E-02 | 1.72 | ↑0.78 | 4.46E-02 | 7.45E-02 | 1.56 | ↑0.65 | 5.46E-03 | 1.15E-02 | 1.56 | ↑0.64 | 3.06E-02 | 5.55E-02 | 1.80 | ↑0.85 | 7.99E-05 | 2.18E-04 | 2.18 | ↑1.12 |
| PE160/182 | 2.79E-06 | 1.13E-05 | 2.74 | ↑1.45 | 1.24E-03 | 3.33E-03 | 2.39 | ↑1.26 | 1.88E-03 | 4.82E-03 | 2.44 | ↑1.29 | 2.45E-03 | 5.82E-03 | 2.69 | ↑1.43 | 1.45E-05 | 6.59E-05 | 3.35 | ↑1.75 | 1.24E-06 | 4.50E-06 | 3.10 | ↑1.63 |
| Phenylacetyl-L-glutamine | 2.92E-06 | 1.16E-05 | 0.45 | ↓-1.15 | 1.96E-04 | 7.56E-04 | 0.53 | ↓-0.91 | 1.47E-07 | 2.07E-06 | 0.40 | ↓-1.31 | 4.31E-10 | 1.10E-08 | 0.32 | ↓-1.65 | 2.18E-06 | 1.84E-05 | 0.44 | ↓-1.17 | 1.14E-09 | 1.11E-08 | 0.34 | ↓-1.57 |
| Oleyloxyethylphosphorylcholine | 3.82E-06 | 1.49E-05 | 0.66 | ↓-0.60 | 5.40E-08 | 1.27E-06 | 0.60 | ↓-0.75 | 2.15E-07 | 2.73E-06 | 0.62 | ↓-0.70 | 5.56E-08 | 6.42E-07 | 0.59 | ↓-0.75 | 4.13E-02 | 7.28E-02 | 1.11 | ↑0.14 | 1.26E-07 | 6.14E-07 | 0.60 | ↓-0.74 |
| KRTLRR | 4.27E-06 | 1.64E-05 | 0.71 | ↓-0.50 | 1.46E-04 | 5.98E-04 | 0.79 | ↓-0.33 | 1.59E-06 | 1.30E-05 | 0.73 | ↓-0.46 | 7.51E-06 | 4.34E-05 | 0.72 | ↓-0.47 | 3.57E-09 | 6.98E-08 | 0.68 | ↓-0.56 | 1.48E-09 | 1.34E-08 | 0.63 | ↓-0.67 |
| Dihomo--linolenoyl PAF C-16 | 4.35E-06 | 1.65E-05 | 3.34 | ↑1.74 | 4.30E-04 | 1.38E-03 | 2.45 | ↑1.29 | 2.01E-04 | 7.35E-04 | 3.79 | ↑1.92 | 1.19E-06 | 8.64E-06 | 3.51 | ↑1.81 | 6.59E-06 | 3.80E-05 | 3.10 | ↑1.63 | 1.09E-11 | 2.13E-10 | 5.53 | ↑2.47 |
| O14502 | 4.89E-06 | 1.83E-05 | 0.65 | ↓-0.62 | 5.22E-07 | 8.29E-06 | 0.62 | ↓-0.68 | 4.53E-06 | 3.03E-05 | 0.65 | ↓-0.63 | 1.94E-05 | 9.31E-05 | 0.68 | ↓-0.56 | 5.58E-09 | 1.01E-07 | 0.58 | ↓-0.80 | 2.37E-07 | 1.06E-06 | 0.59 | ↓-0.76 |
| 1,2-Dioleoyl-sn-glycero-3-PC | 5.52E-06 | 2.03E-05 | 2.20 | ↑1.14 | 2.60E-04 | 9.10E-04 | 1.85 | ↑0.89 | 2.52E-04 | 8.64E-04 | 2.25 | ↑1.17 | 8.57E-07 | 7.02E-06 | 2.43 | ↑1.28 | 6.09E-05 | 2.15E-04 | 2.13 | ↑1.09 | 6.57E-10 | 6.96E-09 | 3.25 | ↑1.70 |
| Androst-16-en-3-ol, 3,5- | 6.97E-06 | 2.53E-05 | 2.09 | ↑1.06 | 5.50E-08 | 1.27E-06 | 2.48 | ↑1.31 | 5.63E-05 | 2.51E-04 | 2.22 | ↑1.15 | 5.85E-04 | 1.62E-03 | 1.62 | ↑0.70 | 3.72E-06 | 2.62E-05 | 2.03 | ↑1.02 | 3.36E-07 | 1.45E-06 | 2.59 | ↑1.37 |
| Q4R6W8 | 8.65E-06 | 3.09E-05 | 0.56 | ↓-0.83 | 5.04E-03 | 1.06E-02 | 0.71 | ↓-0.48 | 6.66E-03 | 1.41E-02 | 0.75 | ↓-0.41 | 4.02E-02 | 6.94E-02 | 0.81 | ↓-0.31 | 1.65E-01 | 2.30E-01 | 0.88 | ↓-0.18 | 1.42E-02 | 2.62E-02 | 0.77 | ↓-0.37 |
| P06868 | 1.53E-05 | 5.38E-05 | 0.78 | ↓-0.36 | 2.08E-04 | 7.88E-04 | 0.78 | ↓-0.36 | 7.46E-04 | 2.20E-03 | 0.69 | ↓-0.54 | 1.87E-03 | 4.62E-03 | 0.85 | ↓-0.24 | 1.81E-02 | 3.56E-02 | 0.83 | ↓-0.27 | 4.32E-03 | 8.92E-03 | 0.73 | ↓-0.45 |
| C18Plasm-226 PC | 1.61E-05 | 5.62E-05 | 3.87 | ↑1.95 | 5.88E-03 | 1.19E-02 | 3.53 | ↑1.82 | 1.52E-03 | 3.98E-03 | 2.79 | ↑1.48 | 4.23E-03 | 9.50E-03 | 2.31 | ↑1.21 | 1.97E-03 | 4.75E-03 | 2.48 | ↑1.31 | 1.20E-05 | 3.76E-05 | 3.53 | ↑1.82 |
| H2Q321 | 1.73E-05 | 5.92E-05 | 1.41 | ↑0.50 | 1.83E-02 | 3.34E-02 | 1.33 | ↑0.41 | 1.42E-05 | 7.21E-05 | 1.40 | ↑0.48 | 1.04E-04 | 3.53E-04 | 1.33 | ↑0.42 | 5.25E-05 | 1.88E-04 | 1.34 | ↑0.42 | 1.18E-04 | 3.12E-04 | 1.34 | ↑0.43 |
| LysoPC240 | 1.99E-05 | 6.73E-05 | 0.69 | ↓-0.55 | 1.12E-04 | 4.94E-04 | 0.73 | ↓-0.45 | 4.43E-03 | 9.97E-03 | 0.81 | ↓-0.30 | 2.67E-05 | 1.15E-04 | 0.70 | ↓-0.51 | 1.68E-08 | 2.85E-07 | 0.59 | ↓-0.75 | 4.29E-07 | 1.79E-06 | 0.64 | ↓-0.65 |
| P07589 | 2.80E-05 | 9.37E-05 | 0.60 | ↓-0.73 | 7.95E-07 | 1.01E-05 | 0.51 | ↓-0.96 | 1.12E-05 | 6.06E-05 | 0.48 | ↓-1.05 | 5.89E-05 | 2.17E-04 | 0.58 | ↓-0.78 | 1.02E-03 | 2.64E-03 | 0.69 | ↓-0.54 | 7.69E-07 | 3.01E-06 | 0.52 | ↓-0.93 |
| 2-Thio PAF | 3.65E-05 | 1.20E-04 | 0.63 | ↓-0.67 | 3.91E-03 | 8.43E-03 | 0.73 | ↓-0.45 | 6.19E-02 | 1.01E-01 | 0.83 | ↓-0.28 | 2.21E-04 | 6.92E-04 | 0.55 | ↓-0.87 | 3.89E-05 | 1.52E-04 | 0.63 | ↓-0.68 | 9.28E-02 | 1.40E-01 | 0.85 | ↓-0.23 |
| PC140 | 4.22E-05 | 1.37E-04 | 0.54 | ↓-0.88 | 9.04E-07 | 1.09E-05 | 0.47 | ↓-1.08 | 1.82E-08 | 3.08E-07 | 0.40 | ↓-1.33 | 4.31E-09 | 7.83E-08 | 0.37 | ↓-1.43 | 3.68E-08 | 5.50E-07 | 0.42 | ↓-1.25 | 6.99E-09 | 5.07E-08 | 0.39 | ↓-1.34 |
| 1-Docosahexaenoyl-2-stearoyl-sn-glycero-3-phosphocholine | 4.60E-05 | 1.48E-04 | 2.82 | ↑1.50 | 3.17E-05 | 1.87E-04 | 2.48 | ↑1.31 | 1.22E-05 | 6.48E-05 | 3.22 | ↑1.69 | 3.73E-06 | 2.31E-05 | 3.65 | ↑1.87 | 7.09E-06 | 3.92E-05 | 2.88 | ↑1.53 | 9.98E-13 | 5.07E-11 | 5.61 | ↑2.49 |
| P48740 | 5.28E-05 | 1.68E-04 | 0.76 | ↓-0.39 | 2.53E-03 | 6.18E-03 | 0.82 | ↓-0.29 | 5.16E-03 | 1.14E-02 | 0.84 | ↓-0.26 | 2.20E-01 | 2.86E-01 | 0.93 | ↓-0.11 | 3.09E-03 | 6.95E-03 | 0.82 | ↓-0.28 | 8.39E-04 | 1.94E-03 | 0.80 | ↓-0.32 |
| Q4R955 | 6.02E-05 | 1.89E-04 | 0.80 | ↓-0.33 | 4.95E-04 | 1.57E-03 | 0.83 | ↓-0.27 | 1.06E-03 | 2.99E-03 | 0.83 | ↓-0.28 | 1.31E-01 | 1.94E-01 | 0.93 | ↓-0.11 | 4.61E-03 | 1.01E-02 | 0.85 | ↓-0.23 | 3.67E-02 | 6.09E-02 | 0.90 | ↓-0.15 |
| G1U9R4 | 6.42E-05 | 1.99E-04 | 0.40 | ↓-1.32 | 6.50E-02 | 1.04E-01 | 0.62 | ↓-0.68 | 1.19E-02 | 2.36E-02 | 0.64 | ↓-0.65 | 1.77E-01 | 2.42E-01 | 0.83 | ↓-0.27 | 8.69E-02 | 1.39E-01 | 0.53 | ↓-0.92 | 1.47E-04 | 3.77E-04 | 0.41 | ↓-1.28 |
| B4DPQ0 | 9.08E-05 | 2.78E-04 | 0.77 | ↓-0.38 | 6.74E-04 | 2.01E-03 | 0.80 | ↓-0.31 | 6.65E-06 | 4.02E-05 | 0.75 | ↓-0.42 | 2.07E-04 | 6.56E-04 | 0.79 | ↓-0.33 | 5.72E-04 | 1.58E-03 | 0.81 | ↓-0.30 | 1.11E-05 | 3.57E-05 | 0.77 | ↓-0.38 |
| A2RTY6 | 1.31E-04 | 3.97E-04 | 0.78 | ↓-0.36 | 6.46E-07 | 9.11E-06 | 0.71 | ↓-0.50 | 9.23E-06 | 5.33E-05 | 0.75 | ↓-0.41 | 1.04E-04 | 3.53E-04 | 0.80 | ↓-0.33 | 6.36E-04 | 1.74E-03 | 0.82 | ↓-0.28 | 2.23E-05 | 6.82E-05 | 0.78 | ↓-0.36 |
| Q5VY30 | 1.45E-04 | 4.34E-04 | 0.71 | ↓-0.50 | 4.12E-06 | 3.74E-05 | 0.65 | ↓-0.63 | 5.36E-07 | 5.44E-06 | 0.57 | ↓-0.80 | 5.37E-06 | 3.17E-05 | 0.64 | ↓-0.64 | 4.41E-06 | 2.87E-05 | 0.64 | ↓-0.65 | 1.38E-04 | 3.62E-04 | 0.71 | ↓-0.50 |
| P02741 | 1.62E-04 | 4.79E-04 | 9.69 | ↑3.28 | 4.31E-07 | 7.30E-06 | 28.10 | ↑4.81 | 1.96E-07 | 2.62E-06 | 19.26 | ↑4.27 | 4.49E-08 | 5.70E-07 | 16.54 | ↑4.05 | 4.66E-06 | 2.91E-05 | 15.59 | ↑3.96 | 3.36E-08 | 1.94E-07 | 15.43 | ↑3.95 |
| V9HWA9 | 2.17E-04 | 6.32E-04 | 0.79 | ↓-0.33 | 3.18E-03 | 7.34E-03 | 0.83 | ↓-0.26 | 5.93E-03 | 1.30E-02 | 0.84 | ↓-0.25 | 6.01E-02 | 9.72E-02 | 0.89 | ↓-0.16 | 9.05E-02 | 1.39E-01 | 0.91 | ↓-0.14 | 6.81E-02 | 1.07E-01 | 0.90 | ↓-0.16 |
| H2QCZ6 | 2.19E-04 | 6.32E-04 | 0.40 | ↓-1.33 | 1.29E-02 | 2.47E-02 | 0.52 | ↓-0.94 | 6.94E-03 | 1.46E-02 | 0.50 | ↓-0.99 | 1.17E-03 | 2.94E-03 | 0.49 | ↓-1.04 | 2.78E-03 | 6.36E-03 | 0.54 | ↓-0.88 | 4.70E-03 | 9.56E-03 | 0.52 | ↓-0.95 |
| H2QV12 | 3.75E-04 | 1.07E-03 | 0.77 | ↓-0.38 | 5.19E-05 | 2.80E-04 | 0.73 | ↓-0.45 | 5.61E-07 | 5.48E-06 | 0.68 | ↓-0.56 | 5.02E-08 | 6.08E-07 | 0.63 | ↓-0.66 | 3.79E-12 | 2.41E-10 | 0.56 | ↓-0.83 | 4.83E-10 | 5.58E-09 | 0.60 | ↓-0.73 |
| SMd181/180 | 4.34E-04 | 1.23E-03 | 2.26 | ↑1.18 | 4.40E-03 | 9.30E-03 | 2.11 | ↑1.08 | 4.94E-03 | 1.10E-02 | 2.08 | ↑1.06 | 9.00E-04 | 2.36E-03 | 2.84 | ↑1.51 | 2.76E-03 | 6.36E-03 | 3.07 | ↑1.62 | 1.23E-05 | 3.80E-05 | 2.91 | ↑1.54 |
| G1PYJ2 | 4.76E-04 | 1.33E-03 | 0.65 | ↓-0.61 | 8.26E-02 | 1.27E-01 | 0.82 | ↓-0.29 | 7.39E-03 | 1.54E-02 | 0.72 | ↓-0.47 | 3.30E-02 | 5.83E-02 | 0.77 | ↓-0.38 | 1.59E-01 | 2.26E-01 | 0.82 | ↓-0.29 | 1.70E-01 | 2.27E-01 | 0.84 | ↓-0.26 |
| LysoPC220 | 7.02E-04 | 1.94E-03 | 0.74 | ↓-0.43 | 8.53E-05 | 4.17E-04 | 0.71 | ↓-0.50 | 1.37E-05 | 7.11E-05 | 0.67 | ↓-0.59 | 3.21E-04 | 9.37E-04 | 0.73 | ↓-0.45 | 9.10E-10 | 2.31E-08 | 0.54 | ↓-0.89 | 1.48E-06 | 5.16E-06 | 0.61 | ↓-0.72 |
| P02768 | 7.46E-04 | 2.04E-03 | 0.48 | ↓-1.05 | 9.41E-01 | 9.56E-01 | 3.42 | ↑1.77 | 1.02E-01 | 1.54E-01 | 0.75 | ↓-0.42 | 1.70E-02 | 3.29E-02 | 0.64 | ↓-0.64 | 9.80E-01 | 9.88E-01 | 1.64 | ↑0.71 | 6.35E-01 | 7.00E-01 | 1.12 | ↑0.17 |
| N-Benzylglycine methyl ester | 8.01E-04 | 2.16E-03 | 0.72 | ↓-0.48 | 2.03E-05 | 1.37E-04 | 0.55 | ↓-0.86 | 6.03E-06 | 3.73E-05 | 0.53 | ↓-0.92 | 3.55E-04 | 1.03E-03 | 0.61 | ↓-0.72 | 3.98E-04 | 1.16E-03 | 0.62 | ↓-0.69 | 2.80E-05 | 8.36E-05 | 0.56 | ↓-0.83 |
| Q8IZZ5 | 8.40E-04 | 2.25E-03 | 0.76 | ↓-0.39 | 7.79E-05 | 3.88E-04 | 0.72 | ↓-0.48 | 6.12E-03 | 1.33E-02 | 0.81 | ↓-0.30 | 1.16E-03 | 2.94E-03 | 0.78 | ↓-0.36 | 1.28E-04 | 4.21E-04 | 0.75 | ↓-0.42 | 4.77E-05 | 1.36E-04 | 0.72 | ↓-0.47 |
| G1SQ27 | 1.08E-03 | 2.86E-03 | 0.60 | ↓-0.75 | 3.58E-03 | 8.13E-03 | 0.63 | ↓-0.66 | 7.28E-05 | 3.13E-04 | 0.48 | ↓-1.05 | 7.98E-04 | 2.13E-03 | 0.58 | ↓-0.79 | 1.55E-05 | 6.88E-05 | 0.47 | ↓-1.09 | 2.39E-05 | 7.23E-05 | 0.45 | ↓-1.14 |
| P02776 | 1.10E-03 | 2.89E-03 | 0.47 | ↓-1.09 | 6.03E-04 | 1.85E-03 | 0.47 | ↓-1.08 | 6.96E-04 | 2.08E-03 | 0.47 | ↓-1.09 | 9.96E-06 | 5.38E-05 | 0.35 | ↓-1.49 | 8.73E-06 | 4.52E-05 | 0.36 | ↓-1.47 | 1.56E-04 | 3.97E-04 | 0.42 | ↓-1.24 |
| P11276 | 1.13E-03 | 2.93E-03 | 0.65 | ↓-0.61 | 4.30E-05 | 2.48E-04 | 0.51 | ↓-0.96 | 5.75E-06 | 3.65E-05 | 0.46 | ↓-1.12 | 2.33E-04 | 7.22E-04 | 0.58 | ↓-0.78 | 1.15E-03 | 2.90E-03 | 0.64 | ↓-0.65 | 1.36E-06 | 4.81E-06 | 0.44 | ↓-1.18 |
| H2RC06 | 1.17E-03 | 3.00E-03 | 0.39 | ↓-1.35 | 1.50E-01 | 2.12E-01 | 0.73 | ↓-0.46 | 2.07E-01 | 2.84E-01 | 0.67 | ↓-0.57 | 4.45E-02 | 7.49E-02 | 0.60 | ↓-0.73 | 7.55E-02 | 1.21E-01 | 0.59 | ↓-0.76 | 8.28E-03 | 1.61E-02 | 0.49 | ↓-1.02 |
| Q4R4P0 | 1.46E-03 | 3.70E-03 | 0.52 | ↓-0.94 | 5.36E-01 | 6.00E-01 | 0.80 | ↓-0.32 | 1.52E-02 | 2.89E-02 | 0.62 | ↓-0.70 | 5.30E-03 | 1.12E-02 | 0.59 | ↓-0.75 | 5.12E-03 | 1.11E-02 | 0.56 | ↓-0.85 | 1.47E-02 | 2.70E-02 | 0.60 | ↓-0.74 |
| H2QU06 | 1.69E-03 | 4.26E-03 | 0.77 | ↓-0.39 | 9.22E-04 | 2.57E-03 | 0.75 | ↓-0.41 | 7.98E-03 | 1.63E-02 | 0.80 | ↓-0.33 | 7.35E-03 | 1.51E-02 | 0.80 | ↓-0.33 | 3.49E-02 | 6.24E-02 | 0.84 | ↓-0.26 | 1.03E-02 | 1.96E-02 | 0.81 | ↓-0.31 |
| A0A2R9CH07 | 2.02E-03 | 5.02E-03 | 0.64 | ↓-0.65 | 3.51E-03 | 8.02E-03 | 0.65 | ↓-0.63 | 2.36E-02 | 4.17E-02 | 0.79 | ↓-0.35 | 1.57E-03 | 3.90E-03 | 0.64 | ↓-0.65 | 1.29E-02 | 2.60E-02 | 0.69 | ↓-0.54 | 1.60E-01 | 2.17E-01 | 0.84 | ↓-0.26 |
| G1SU82 | 2.77E-03 | 6.82E-03 | 0.82 | ↓-0.29 | 8.96E-06 | 6.90E-05 | 0.73 | ↓-0.45 | 4.00E-06 | 2.74E-05 | 0.73 | ↓-0.46 | 1.88E-05 | 9.27E-05 | 0.75 | ↓-0.42 | 1.77E-09 | 3.74E-08 | 0.66 | ↓-0.61 | 6.31E-08 | 3.27E-07 | 0.67 | ↓-0.58 |
| P18428 | 2.92E-03 | 7.13E-03 | 2.81 | ↑1.49 | 2.06E-01 | 2.70E-01 | 2.04 | ↑1.03 | 3.68E-02 | 6.27E-02 | 2.34 | ↑1.23 | 1.90E-05 | 9.27E-05 | 5.51 | ↑2.46 | 7.74E-03 | 1.61E-02 | 2.12 | ↑1.09 | 4.33E-02 | 7.10E-02 | 1.99 | ↑0.99 |
| Q9NP10 | 3.14E-03 | 7.60E-03 | 0.23 | ↓-2.11 | 1.83E-01 | 2.48E-01 | 0.52 | ↓-0.93 | 2.43E-02 | 4.25E-02 | 0.34 | ↓-1.56 | 5.14E-02 | 8.37E-02 | 0.39 | ↓-1.38 | 6.22E-02 | 1.04E-01 | 0.33 | ↓-1.59 | 2.32E-01 | 2.93E-01 | 0.50 | ↓-1.01 |
| A0A2R9BCB8 | 3.35E-03 | 8.02E-03 | 0.69 | ↓-0.53 | 8.18E-03 | 1.62E-02 | 0.74 | ↓-0.43 | 1.97E-02 | 3.56E-02 | 0.76 | ↓-0.39 | 2.60E-02 | 4.90E-02 | 0.77 | ↓-0.38 | 7.65E-03 | 1.61E-02 | 0.71 | ↓-0.48 | 1.42E-04 | 3.67E-04 | 0.59 | ↓-0.76 |
| G1P5R5 | 3.72E-03 | 8.84E-03 | 0.84 | ↓-0.26 | 1.07E-04 | 4.93E-04 | 0.75 | ↓-0.41 | 1.61E-03 | 4.17E-03 | 0.83 | ↓-0.27 | 2.80E-04 | 8.34E-04 | 0.80 | ↓-0.32 | 3.54E-07 | 4.09E-06 | 0.64 | ↓-0.65 | 4.59E-07 | 1.88E-06 | 0.70 | ↓-0.51 |
| Q6UXM4 | 3.95E-03 | 9.29E-03 | 0.38 | ↓-1.40 | 1.03E-01 | 1.53E-01 | 0.58 | ↓-0.78 | 1.30E-02 | 2.51E-02 | 0.51 | ↓-0.97 | 3.18E-02 | 5.73E-02 | 0.55 | ↓-0.86 | 2.55E-02 | 4.76E-02 | 0.57 | ↓-0.82 | 9.31E-01 | 9.38E-01 | 0.88 | ↓-0.18 |
| Q4R4R3 | 4.03E-03 | 9.38E-03 | 0.70 | ↓-0.51 | 6.08E-05 | 3.09E-04 | 0.58 | ↓-0.77 | 1.82E-05 | 8.87E-05 | 0.57 | ↓-0.82 | 6.49E-04 | 1.75E-03 | 0.64 | ↓-0.64 | 1.18E-04 | 3.95E-04 | 0.60 | ↓-0.73 | 4.96E-05 | 1.39E-04 | 0.58 | ↓-0.80 |
| 13-Docosenamide, Z- | 4.12E-03 | 9.51E-03 | 0.27 | ↓-1.87 | 3.81E-03 | 8.42E-03 | 0.27 | ↓-1.88 | 1.88E-01 | 2.61E-01 | 0.47 | ↓-1.09 | 1.35E-01 | 1.97E-01 | 0.49 | ↓-1.02 | 5.41E-02 | 9.21E-02 | 0.49 | ↓-1.03 | 1.19E-01 | 1.72E-01 | 0.65 | ↓-0.62 |
| G1PA91 | 4.26E-03 | 9.75E-03 | 0.75 | ↓-0.41 | 6.30E-03 | 1.26E-02 | 0.78 | ↓-0.36 | 6.83E-02 | 1.08E-01 | 0.85 | ↓-0.24 | 1.75E-01 | 2.40E-01 | 0.90 | ↓-0.15 | 1.50E-02 | 2.99E-02 | 0.79 | ↓-0.34 | 1.75E-04 | 4.39E-04 | 0.67 | ↓-0.57 |
| G1PHQ7 | 4.86E-03 | 1.10E-02 | 0.46 | ↓-1.12 | 3.91E-01 | 4.71E-01 | 0.77 | ↓-0.38 | 8.51E-01 | 8.85E-01 | 0.88 | ↓-0.18 | 3.14E-02 | 5.73E-02 | 0.60 | ↓-0.73 | 8.93E-02 | 1.39E-01 | 0.62 | ↓-0.68 | 2.28E-01 | 2.90E-01 | 0.69 | ↓-0.53 |
| H0VMS1 | 5.82E-03 | 1.30E-02 | 0.77 | ↓-0.37 | 4.77E-02 | 8.02E-02 | 0.85 | ↓-0.23 | 1.20E-03 | 3.24E-03 | 0.72 | ↓-0.46 | 1.17E-01 | 1.76E-01 | 0.89 | ↓-0.16 | 2.11E-02 | 4.09E-02 | 0.81 | ↓-0.30 | 3.66E-04 | 8.78E-04 | 0.70 | ↓-0.52 |
| 9-Oxo-11,15R-dihydroxyprostan-1-oic acid | 5.85E-03 | 1.30E-02 | 0.26 | ↓-1.97 | 3.88E-03 | 8.43E-03 | 0.25 | ↓-2.01 | 2.21E-01 | 2.96E-01 | 0.46 | ↓-1.13 | 1.43E-01 | 2.05E-01 | 0.48 | ↓-1.07 | 6.60E-02 | 1.08E-01 | 0.49 | ↓-1.04 | 1.56E-01 | 2.14E-01 | 0.67 | ↓-0.58 |
| P00738 | 5.88E-03 | 1.30E-02 | 2.03 | ↑1.02 | 1.32E-02 | 2.51E-02 | 2.57 | ↑1.36 | 1.15E-03 | 3.18E-03 | 1.79 | ↑0.84 | 4.29E-04 | 1.21E-03 | 3.25 | ↑1.70 | 1.39E-07 | 1.86E-06 | 3.36 | ↑1.75 | 8.13E-03 | 1.59E-02 | 2.10 | ↑1.07 |

| | | | | | | | | | | | | | | | | | | | | | | | | |
|---|---|---|---|---|---|---|---|---|---|---|---|---|---|---|---|---|---|---|---|---|---|---|---|---|
| A0A2I3RER0 | 6.27E-03 | 1.37E-02 | 0.85 | ↓-0.24 | 1.36E-01 | 1.94E-01 | 0.93 | ↓-0.11 | 1.32E-02 | 2.52E-02 | 0.86 | ↓-0.21 | 7.20E-03 | 1.49E-02 | 0.85 | ↓-0.24 | 6.99E-02 | 1.14E-01 | 0.90 | ↓-0.15 | 3.24E-03 | 6.87E-03 | 0.83 | ↓-0.26 |
| G1NVY1 | 6.71E-03 | 1.46E-02 | 0.72 | ↓-0.48 | 1.02E-03 | 2.81E-03 | 0.66 | ↓-0.61 | 1.26E-03 | 3.37E-03 | 0.64 | ↓-0.64 | 2.16E-02 | 4.10E-02 | 0.73 | ↓-0.45 | 3.47E-05 | 1.44E-04 | 0.52 | ↓-0.96 | 1.00E-07 | 5.10E-07 | 0.38 | ↓-1.38 |
| Q28508 | 7.56E-03 | 1.63E-02 | 1.69 | ↑0.76 | 1.91E-01 | 2.55E-01 | 2.29 | ↑1.20 | 1.92E-03 | 4.88E-03 | 1.87 | ↑0.91 | 3.05E-05 | 1.27E-04 | 2.62 | ↑1.39 | 1.09E-06 | 1.03E-05 | 2.65 | ↑1.41 | 1.18E-05 | 3.76E-05 | 2.43 | ↑1.28 |
| A0A2R8ZJ60 | 9.26E-03 | 1.98E-02 | 0.81 | ↓-0.31 | 2.94E-01 | 3.71E-01 | 0.92 | ↓-0.12 | 1.82E-02 | 3.33E-02 | 0.82 | ↓-0.28 | 5.62E-01 | 6.43E-01 | 1.04 | ↑0.06 | 5.16E-01 | 6.13E-01 | 0.93 | ↓-0.11 | 3.16E-03 | 6.74E-03 | 0.78 | ↓-0.36 |
| Pyrrolidino PAF C-16 | 1.11E-02 | 2.35E-02 | 0.84 | ↓-0.25 | 2.89E-03 | 6.79E-03 | 0.81 | ↓-0.30 | 1.67E-04 | 6.33E-04 | 0.74 | ↓-0.44 | 1.27E-05 | 6.72E-05 | 0.69 | ↓-0.54 | 6.24E-08 | 8.81E-07 | 0.65 | ↓-0.61 | 1.18E-06 | 4.34E-06 | 0.68 | ↓-0.56 |
| B7Z8Q2 | 1.19E-02 | 2.50E-02 | 0.60 | ↓-0.73 | 7.04E-04 | 2.08E-03 | 0.51 | ↓-0.97 | 7.36E-02 | 1.15E-01 | 0.69 | ↓-0.53 | 9.12E-02 | 1.42E-01 | 0.73 | ↓-0.45 | 5.36E-03 | 1.14E-02 | 0.59 | ↓-0.77 | 1.13E-02 | 2.13E-02 | 0.60 | ↓-0.74 |
| B4E1C2 | 1.22E-02 | 2.54E-02 | 0.77 | ↓-0.37 | 1.48E-02 | 2.78E-02 | 0.78 | ↓-0.36 | 3.16E-02 | 5.51E-02 | 0.80 | ↓-0.32 | 2.57E-01 | 3.33E-01 | 0.88 | ↓-0.19 | 2.31E-04 | 7.20E-04 | 0.62 | ↓-0.69 | 7.53E-05 | 2.08E-04 | 0.62 | ↓-0.70 |
| P03952 | 1.30E-02 | 2.68E-02 | 0.89 | ↓-0.17 | 1.89E-07 | 3.70E-06 | 0.78 | ↓-0.37 | 1.25E-06 | 1.06E-05 | 0.75 | ↓-0.41 | 9.90E-07 | 7.62E-06 | 0.76 | ↓-0.40 | 2.78E-03 | 6.36E-03 | 0.88 | ↓-0.19 | 2.17E-03 | 4.70E-03 | 0.84 | ↓-0.25 |
| Q8IVC0 | 1.35E-02 | 2.77E-02 | 0.71 | ↓-0.50 | 3.55E-02 | 6.05E-02 | 0.74 | ↓-0.43 | 6.33E-03 | 1.36E-02 | 0.68 | ↓-0.55 | 1.85E-01 | 2.50E-01 | 0.83 | ↓-0.26 | 7.10E-02 | 1.15E-01 | 0.79 | ↓-0.34 | 5.87E-04 | 1.38E-03 | 0.61 | ↓-0.71 |
| A0A2R9AC29 | 1.49E-02 | 3.04E-02 | 0.79 | ↓-0.33 | 5.42E-03 | 1.12E-02 | 0.77 | ↓-0.38 | 2.52E-03 | 6.03E-03 | 0.74 | ↓-0.44 | 1.29E-02 | 2.54E-02 | 0.79 | ↓-0.34 | 1.59E-02 | 3.15E-02 | 0.80 | ↓-0.32 | 8.81E-02 | 1.34E-01 | 0.86 | ↓-0.21 |
| V9HWD8 | 1.53E-02 | 3.09E-02 | 0.75 | ↓-0.42 | 3.11E-02 | 5.44E-02 | 0.78 | ↓-0.36 | 2.87E-01 | 3.69E-01 | 0.89 | ↓-0.17 | 3.08E-01 | 3.90E-01 | 0.90 | ↓-0.16 | 2.70E-02 | 4.96E-02 | 0.76 | ↓-0.39 | 2.03E-02 | 3.60E-02 | 0.76 | ↓-0.40 |
| A0A2R9A8Q1 | 1.57E-02 | 3.14E-02 | 0.85 | ↓-0.24 | 1.71E-01 | 2.36E-01 | 1.09 | ↑0.13 | 1.95E-01 | 2.70E-01 | 0.91 | ↓-0.13 | 7.11E-01 | 7.81E-01 | 0.98 | ↓-0.03 | 5.38E-01 | 6.35E-01 | 1.03 | ↑0.05 | 5.36E-01 | 6.13E-01 | 0.95 | ↓-0.07 |
| A0A2R9BFI7 | 1.59E-02 | 3.15E-02 | 0.42 | ↓-1.24 | 1.06E-02 | 2.04E-02 | 0.50 | ↓-1.00 | 3.64E-01 | 4.49E-01 | 0.89 | ↓-0.17 | 1.92E-01 | 2.58E-01 | 0.76 | ↓-0.39 | 1.97E-02 | 3.84E-02 | 0.45 | ↓-1.16 | 1.15E-03 | 2.60E-03 | 0.36 | ↓-1.47 |
| P13645 | 1.60E-02 | 3.15E-02 | 0.65 | ↓-0.63 | 2.50E-02 | 4.44E-02 | 0.68 | ↓-0.55 | 1.44E-01 | 2.08E-01 | 0.86 | ↓-0.22 | 3.46E-03 | 7.85E-03 | 0.61 | ↓-0.72 | 4.57E-02 | 7.90E-02 | 0.73 | ↓-0.45 | 1.26E-01 | 1.78E-01 | 1.03 | ↑0.04 |
| Q04756 | 1.75E-02 | 3.42E-02 | 0.80 | ↓-0.31 | 3.72E-03 | 8.29E-03 | 0.76 | ↓-0.39 | 9.92E-04 | 2.83E-03 | 0.73 | ↓-0.45 | 3.98E-02 | 6.94E-02 | 0.83 | ↓-0.27 | 1.20E-03 | 2.99E-03 | 0.74 | ↓-0.43 | 1.03E-03 | 2.35E-03 | 0.73 | ↓-0.45 |
| Q4R6T2 | 2.29E-02 | 4.45E-02 | 0.88 | ↓-0.19 | 1.96E-03 | 4.92E-03 | 0.83 | ↓-0.26 | 3.39E-04 | 1.15E-03 | 0.81 | ↓-0.30 | 1.52E-01 | 2.17E-01 | 0.92 | ↓-0.12 | 1.09E-01 | 1.65E-01 | 0.91 | ↓-0.13 | 2.22E-02 | 3.88E-02 | 0.88 | ↓-0.18 |
| H0V4A3 | 2.96E-02 | 5.70E-02 | 0.77 | ↓-0.37 | 6.16E-03 | 1.24E-02 | 0.69 | ↓-0.53 | 3.54E-02 | 6.11E-02 | 0.78 | ↓-0.36 | 2.65E-02 | 4.95E-02 | 0.82 | ↓-0.29 | 4.64E-02 | 7.97E-02 | 0.81 | ↓-0.31 | 1.12E-04 | 3.00E-04 | 0.58 | ↓-0.79 |
| G1PCZ3 | 3.09E-02 | 5.89E-02 | 0.63 | ↓-0.67 | 2.23E-02 | 4.02E-02 | 0.69 | ↓-0.54 | 7.23E-02 | 1.14E-01 | 0.91 | ↓-0.13 | 1.69E-01 | 2.33E-01 | 0.82 | ↓-0.29 | 8.42E-04 | 2.25E-03 | 0.46 | ↓-1.12 | 4.62E-02 | 7.48E-02 | 0.64 | ↓-0.65 |
| A0A2R8ZLB9 | 3.32E-02 | 6.29E-02 | 0.78 | ↓-0.37 | 3.70E-04 | 1.22E-03 | 0.65 | ↓-0.63 | 6.68E-03 | 1.41E-02 | 0.73 | ↓-0.46 | 4.95E-02 | 8.12E-02 | 0.80 | ↓-0.33 | 1.36E-04 | 4.42E-04 | 0.63 | ↓-0.67 | 1.02E-04 | 2.75E-04 | 0.63 | ↓-0.67 |
| B2R8I2 | 3.43E-02 | 6.45E-02 | 0.76 | ↓-0.39 | 1.85E-04 | 7.24E-04 | 0.61 | ↓-0.72 | 8.92E-03 | 1.81E-02 | 0.71 | ↓-0.49 | 4.30E-03 | 9.50E-03 | 0.68 | ↓-0.55 | 4.01E-04 | 1.16E-03 | 0.63 | ↓-0.68 | 3.74E-05 | 1.10E-04 | 0.55 | ↓-0.86 |
| A2NUT2 | 3.73E-02 | 6.97E-02 | 0.79 | ↓-0.34 | 5.27E-03 | 1.10E-02 | 0.72 | ↓-0.48 | 2.07E-02 | 3.71E-02 | 0.77 | ↓-0.38 | 3.22E-02 | 5.76E-02 | 0.79 | ↓-0.35 | 9.72E-04 | 2.55E-03 | 0.69 | ↓-0.54 | 2.13E-04 | 5.25E-04 | 0.66 | ↓-0.60 |
| Q8HXR9 | 3.96E-02 | 7.35E-02 | 0.76 | ↓-0.39 | 7.95E-02 | 1.23E-01 | 0.88 | ↓-0.19 | 1.56E-02 | 2.94E-02 | 0.70 | ↓-0.51 | 4.03E-02 | 6.94E-02 | 0.73 | ↓-0.45 | 4.93E-04 | 1.39E-03 | 0.52 | ↓-0.93 | 2.59E-04 | 6.32E-04 | 0.51 | ↓-0.96 |
| P01031 | 4.31E-02 | 7.94E-02 | 0.81 | ↓-0.30 | 6.74E-01 | 7.38E-01 | 1.02 | ↑0.02 | 3.03E-01 | 3.85E-01 | 1.07 | ↑0.10 | 4.68E-01 | 5.55E-01 | 1.02 | ↑0.03 | 9.10E-01 | 9.43E-01 | 0.98 | ↓-0.03 | 5.36E-01 | 6.13E-01 | 0.93 | ↓-0.10 |
| P68292 | 4.54E-02 | 8.30E-02 | 0.36 | ↓-1.46 | 6.01E-01 | 6.64E-01 | 0.79 | ↓-0.34 | 5.25E-01 | 6.15E-01 | 0.74 | ↓-0.43 | 4.04E-02 | 6.94E-02 | 0.42 | ↓-1.26 | 1.59E-01 | 2.26E-01 | 0.50 | ↓-0.99 | 6.49E-01 | 7.10E-01 | 0.77 | ↓-0.39 |
| A0A2R8ZZK5 | 4.87E-02 | 8.83E-02 | 0.79 | ↓-0.33 | 3.83E-01 | 4.65E-01 | 0.89 | ↓-0.17 | 3.21E-03 | 7.49E-03 | 0.71 | ↓-0.50 | 4.52E-03 | 9.80E-03 | 0.72 | ↓-0.47 | 1.58E-01 | 2.26E-01 | 0.83 | ↓-0.26 | 6.96E-01 | 7.49E-01 | 0.92 | ↓-0.11 |
| H2QPM1 | 5.15E-02 | 9.28E-02 | 0.42 | ↓-1.26 | 4.51E-01 | 5.28E-01 | 1.24 | ↑0.31 | 9.68E-01 | 9.75E-01 | 0.76 | ↓-0.40 | 4.16E-02 | 7.05E-02 | 0.56 | ↓-0.83 | 8.87E-02 | 1.39E-01 | 0.56 | ↓-0.84 | 1.24E-01 | 1.77E-01 | 0.50 | ↓-0.99 |
| A0A024R462 | 5.64E-02 | 1.01E-01 | 0.74 | ↓-0.43 | 1.24E-03 | 3.33E-03 | 0.60 | ↓-0.73 | 6.18E-04 | 1.91E-03 | 0.57 | ↓-0.81 | 4.86E-03 | 1.05E-02 | 0.62 | ↓-0.69 | 6.16E-02 | 1.04E-01 | 0.72 | ↓-0.47 | 3.12E-02 | 5.32E-02 | 0.72 | ↓-0.47 |
| A0A2R9BXC9 | 6.00E-02 | 1.07E-01 | 0.90 | ↓-0.15 | 3.91E-03 | 8.43E-03 | 0.84 | ↓-0.24 | 1.68E-01 | 2.35E-01 | 0.93 | ↓-0.10 | 6.10E-01 | 6.88E-01 | 0.98 | ↓-0.03 | 5.45E-01 | 6.41E-01 | 0.97 | ↓-0.04 | 5.38E-01 | 6.13E-01 | 0.98 | ↓-0.02 |
| A0A2I3T5B0 | 6.07E-02 | 1.07E-01 | 0.91 | ↓-0.14 | 5.71E-03 | 1.17E-02 | 0.86 | ↓-0.21 | 9.95E-03 | 2.01E-02 | 0.88 | ↓-0.19 | 1.31E-01 | 1.94E-01 | 0.93 | ↓-0.11 | 1.80E-01 | 2.48E-01 | 0.93 | ↓-0.10 | 1.30E-01 | 1.83E-01 | 0.93 | ↓-0.11 |
| P0C0L5 | 6.71E-02 | 1.17E-01 | 1.28 | ↑0.35 | 1.88E-02 | 3.41E-02 | 1.32 | ↑0.40 | 5.05E-01 | 5.97E-01 | 1.14 | ↑0.18 | 1.54E-01 | 2.19E-01 | 1.15 | ↑0.20 | 1.57E-01 | 2.26E-01 | 1.22 | ↑0.29 | 8.10E-04 | 1.89E-03 | 1.60 | ↑0.68 |
| P34929 | 6.71E-02 | 1.17E-01 | 0.44 | ↓-1.18 | 3.50E-02 | 6.01E-02 | 0.49 | ↓-1.03 | 7.75E-02 | 1.19E-01 | 0.48 | ↓-1.05 | 3.05E-01 | 3.89E-01 | 0.53 | ↓-0.90 | 4.45E-02 | 7.74E-02 | 0.45 | ↓-1.16 | 3.31E-02 | 5.60E-02 | 0.62 | ↓-0.69 |
| H2QCH3 | 6.81E-02 | 1.18E-01 | 0.82 | ↓-0.29 | 1.45E-01 | 2.06E-01 | 0.84 | ↓-0.26 | 4.58E-01 | 5.46E-01 | 0.93 | ↓-0.10 | 9.58E-02 | 1.48E-01 | 0.83 | ↓-0.27 | 1.73E-01 | 2.41E-01 | 0.85 | ↓-0.23 | 1.67E-02 | 3.01E-02 | 0.76 | ↓-0.40 |
| P18656 | 8.06E-02 | 1.38E-01 | 0.36 | ↓-1.47 | 5.32E-01 | 5.98E-01 | 0.70 | ↓-0.52 | 1.58E-01 | 2.24E-01 | 0.48 | ↓-1.05 | 3.52E-01 | 4.34E-01 | 0.47 | ↓-1.09 | 1.62E-01 | 2.29E-01 | 0.43 | ↓-1.22 | 4.79E-01 | 5.61E-01 | 0.66 | ↓-0.60 |
| A0A2R9C7E2 | 8.89E-02 | 1.52E-01 | 0.90 | ↓-0.16 | 3.90E-01 | 4.71E-01 | 1.10 | ↑0.13 | 6.81E-01 | 7.48E-01 | 2.21 | ↑1.15 | 9.81E-04 | 2.54E-03 | 0.30 | ↓-1.72 | 2.35E-01 | 3.14E-01 | 1.47 | ↑0.55 | 1.17E-01 | 1.72E-01 | 6.27 | ↑2.65 |
| P00761 | 9.03E-02 | 1.53E-01 | 0.85 | ↓-0.24 | 8.40E-01 | 8.78E-01 | 1.08 | ↑0.11 | 5.16E-01 | 6.06E-01 | 1.25 | ↑0.32 | 2.04E-01 | 2.68E-01 | 0.87 | ↓-0.20 | 6.21E-01 | 7.07E-01 | 0.94 | ↓-0.10 | 8.51E-01 | 8.75E-01 | 1.08 | ↑0.11 |
| D3DNN4 | 9.55E-02 | 1.61E-01 | 0.83 | ↓-0.27 | 9.71E-03 | 1.90E-02 | 0.72 | ↓-0.47 | 3.45E-04 | 1.15E-03 | 0.63 | ↓-0.67 | 2.33E-05 | 1.04E-04 | 0.57 | ↓-0.80 | 9.04E-01 | 9.43E-01 | 0.96 | ↓-0.06 | 1.79E-01 | 2.38E-01 | 0.84 | ↓-0.25 |
| A0A2R9AGL7 | 9.66E-02 | 1.61E-01 | 0.63 | ↓-0.66 | 5.19E-02 | 8.50E-02 | 0.61 | ↓-0.71 | 2.88E-01 | 3.69E-01 | 1.16 | ↑0.21 | 6.93E-01 | 7.65E-01 | 0.93 | ↓-0.10 | 4.25E-01 | 5.30E-01 | 0.83 | ↓-0.27 | 6.64E-01 | 7.23E-01 | 0.95 | ↓-0.07 |
| I7GH44 | 1.12E-01 | 1.86E-01 | 1.20 | ↑0.26 | 2.35E-03 | 5.79E-03 | 1.39 | ↑0.48 | 8.06E-02 | 1.23E-01 | 1.14 | ↑0.19 | 3.27E-02 | 5.81E-02 | 1.21 | ↑0.27 | 8.96E-02 | 1.39E-01 | 1.11 | ↑0.15 | 9.48E-02 | 1.42E-01 | 1.16 | ↑0.21 |
| D6R934 | 1.15E-01 | 1.90E-01 | 0.82 | ↓-0.28 | 1.18E-01 | 1.72E-01 | 0.84 | ↓-0.26 | 7.19E-01 | 7.77E-01 | 0.97 | ↓-0.04 | 6.28E-03 | 1.31E-02 | 0.72 | ↓-0.48 | 9.81E-02 | 1.50E-01 | 0.81 | ↓-0.30 | 1.36E-01 | 1.90E-01 | 0.85 | ↓-0.23 |
| P35908 | 1.19E-01 | 1.95E-01 | 0.74 | ↓-0.44 | 5.71E-02 | 9.24E-02 | 0.71 | ↓-0.49 | 2.37E-01 | 3.13E-01 | 0.90 | ↓-0.15 | 1.45E-02 | 2.83E-02 | 0.65 | ↓-0.62 | 5.60E-02 | 9.48E-02 | 0.71 | ↓-0.50 | 2.09E-01 | 2.67E-01 | 0.78 | ↓-0.37 |
| A0A286XIX4 | 1.20E-01 | 1.96E-01 | 0.62 | ↓-0.69 | 6.71E-04 | 2.01E-03 | 0.35 | ↓-1.53 | 1.64E-02 | 3.06E-02 | 0.44 | ↓-1.19 | 6.79E-02 | 1.09E-01 | 0.56 | ↓-0.82 | 1.04E-02 | 2.12E-02 | 0.63 | ↓-0.68 | 6.26E-02 | 9.95E-02 | 0.52 | ↓-0.94 |
| H2QPM5 | 1.24E-01 | 2.01E-01 | 0.88 | ↓-0.19 | 3.71E-03 | 8.29E-03 | 0.78 | ↓-0.36 | 2.12E-01 | 2.89E-01 | 0.89 | ↓-0.17 | 4.08E-02 | 6.95E-02 | 0.84 | ↓-0.25 | 1.88E-05 | 8.22E-05 | 0.66 | ↓-0.61 | 2.07E-03 | 4.53E-03 | 0.77 | ↓-0.38 |
| A0A2R8ZJE1 | 1.27E-01 | 2.05E-01 | 0.87 | ↓-0.19 | 7.64E-02 | 1.19E-01 | 0.88 | ↓-0.18 | 1.08E-01 | 1.63E-01 | 0.87 | ↓-0.20 | 4.47E-01 | 5.33E-01 | 0.96 | ↓-0.06 | 2.70E-01 | 3.56E-01 | 0.93 | ↓-0.11 | 4.93E-03 | 9.87E-03 | 0.71 | ↓-0.49 |
| C0JYY2 | 1.29E-01 | 2.06E-01 | 0.88 | ↓-0.19 | 8.75E-04 | 2.47E-03 | 0.64 | ↓-0.63 | 4.30E-02 | 7.29E-02 | 0.85 | ↓-0.23 | 4.30E-03 | 9.50E-03 | 0.70 | ↓-0.52 | 4.64E-01 | 5.64E-01 | 1.00 | ↓-0.01 | 1.56E-02 | 2.86E-02 | 0.74 | ↓-0.44 |
| A0A2I3TF97 | 1.30E-01 | 2.06E-01 | 1.20 | ↑0.26 | 1.58E-02 | 2.91E-02 | 1.34 | ↑0.43 | 7.99E-01 | 8.43E-01 | 1.00 | ↓0.00 | 5.66E-01 | 6.45E-01 | 1.08 | ↑0.11 | 6.24E-01 | 7.08E-01 | 1.14 | ↑0.18 | 1.36E-02 | 2.53E-02 | 1.44 | ↑0.53 |
| H2QXT3 | 1.34E-01 | 2.11E-01 | 0.83 | ↓-0.27 | 8.35E-02 | 1.28E-01 | 0.66 | ↓-0.60 | 1.82E-02 | 3.33E-02 | 0.61 | ↓-0.70 | 1.36E-01 | 1.97E-01 | 0.72 | ↓-0.47 | 2.50E-02 | 4.70E-02 | 0.64 | ↓-0.64 | 1.66E-02 | 3.01E-02 | 0.62 | ↓-0.70 |
| A0A140VJI7 | 1.49E-01 | 2.33E-01 | 0.89 | ↓-0.17 | 3.04E-03 | 7.09E-03 | 0.77 | ↓-0.38 | 2.18E-04 | 7.58E-04 | 0.72 | ↓-0.48 | 7.73E-03 | 1.57E-02 | 0.79 | ↓-0.34 | 2.58E-02 | 4.78E-02 | 0.82 | ↓-0.28 | 2.26E-03 | 4.86E-03 | 0.75 | ↓-0.42 |
| P05160 | 1.50E-01 | 2.33E-01 | 0.83 | ↓-0.26 | 5.17E-02 | 8.50E-02 | 0.79 | ↓-0.34 | 2.69E-03 | 6.39E-03 | 0.68 | ↓-0.55 | 2.61E-03 | 6.09E-03 | 0.70 | ↓-0.52 | 1.01E-02 | 2.09E-02 | 0.74 | ↓-0.44 | 1.48E-03 | 3.27E-03 | 0.67 | ↓-0.57 |
| Q9Y6R7 | 1.54E-01 | 2.39E-01 | 0.76 | ↓-0.40 | 8.23E-01 | 8.68E-01 | 1.01 | ↑0.01 | 4.65E-01 | 5.51E-01 | 1.05 | ↑0.08 | 3.81E-01 | 4.65E-01 | 1.00 | ↑0.01 | 5.53E-01 | 6.44E-01 | 1.15 | ↑0.21 | 7.53E-01 | 7.93E-01 | 1.23 | ↑0.30 |
| H2QDM3 | 1.64E-01 | 2.53E-01 | 1.42 | ↑0.51 | 5.26E-01 | 5.97E-01 | 1.35 | ↑0.44 | 2.57E-01 | 3.35E-01 | 1.23 | ↑0.30 | 1.13E-01 | 1.71E-01 | 1.27 | ↑0.34 | 4.63E-01 | 5.64E-01 | 0.97 | ↓-0.04 | 5.64E-01 | 6.37E-01 | 1.74 | ↑0.80 |
| Q53G99 | 1.82E-01 | 2.78E-01 | 0.86 | ↓-0.22 | 3.30E-01 | 4.11E-01 | 0.85 | ↓-0.23 | 1.23E-02 | 2.43E-02 | 0.73 | ↓-0.46 | 1.63E-01 | 2.27E-01 | 0.82 | ↓-0.28 | 2.60E-01 | 3.44E-01 | 0.88 | ↓-0.18 | 7.58E-01 | 7.93E-01 | 1.18 | ↑0.24 |
| I7GPF3 | 1.86E-01 | 2.83E-01 | 0.88 | ↓-0.19 | 1.98E-01 | 2.62E-01 | 0.86 | ↓-0.22 | 6.61E-02 | 1.06E-01 | 0.81 | ↓-0.30 | 1.94E-02 | 3.71E-02 | 0.75 | ↓-0.42 | 4.49E-03 | 1.00E-02 | 0.71 | ↓-0.50 | 8.75E-02 | 1.34E-01 | 0.84 | ↓-0.26 |
| H2QY75 | 1.91E-01 | 2.89E-01 | 0.92 | ↓-0.13 | 3.54E-01 | 4.36E-01 | 0.93 | ↓-0.10 | 4.84E-02 | 7.98E-02 | 0.87 | ↓-0.20 | 4.12E-01 | 4.96E-01 | 1.04 | ↑0.05 | 3.75E-01 | 4.74E-01 | 0.94 | ↓-0.09 | 2.19E-02 | 3.87E-02 | 0.85 | ↓-0.23 |
| H0VH18 | 2.03E-01 | 3.03E-01 | 0.78 | ↓-0.36 | 3.15E-01 | 3.94E-01 | 0.78 | ↓-0.36 | 5.30E-01 | 6.17E-01 | 0.82 | ↓-0.29 | 9.01E-01 | 9.37E-01 | 0.88 | ↓-0.19 | 7.96E-01 | 8.57E-01 | 0.94 | ↓-0.09 | 6.99E-01 | 7.49E-01 | 0.92 | ↓-0.12 |
| G1P212 | 2.03E-01 | 3.03E-01 | 0.84 | ↓-0.25 | 1.52E-02 | 2.83E-02 | 0.70 | ↓-0.51 | 3.36E-01 | 4.24E-01 | 0.86 | ↓-0.22 | 3.16E-02 | 5.73E-02 | 0.75 | ↓-0.42 | 4.41E-01 | 5.47E-01 | 0.89 | ↓-0.18 | 4.50E-01 | 5.29E-01 | 1.03 | ↑0.05 |
| A8K9M5 | 2.20E-01 | 3.27E-01 | 0.51 | ↓-0.96 | 3.44E-01 | 4.26E-01 | 0.51 | ↓-0.97 | 9.35E-01 | 9.50E-01 | 1.09 | ↑0.13 | 5.47E-01 | 6.31E-01 | 0.70 | ↓-0.51 | 3.73E-02 | 6.62E-02 | 0.39 | ↓-1.36 | 1.82E-01 | 2.38E-01 | 0.52 | ↓-0.94 |
| V9HW80 | 2.21E-01 | 3.27E-01 | 0.79 | ↓-0.34 | 4.66E-01 | 5.43E-01 | 0.97 | ↓-0.05 | 5.65E-01 | 6.38E-01 | 1.08 | ↑0.11 | 6.16E-01 | 6.92E-01 | 1.14 | ↑0.18 | 6.98E-01 | 7.71E-01 | 0.88 | ↓-0.18 | 4.14E-01 | 4.91E-01 | 1.25 | ↑0.32 |
| G1NXF7 | 2.24E-01 | 3.29E-01 | 0.79 | ↓-0.34 | 2.95E-01 | 3.71E-01 | 1.03 | ↑0.04 | 8.15E-01 | 8.55E-01 | 1.20 | ↑0.27 | 7.20E-02 | 1.14E-01 | 0.71 | ↓-0.50 | 2.96E-01 | 3.83E-01 | 0.76 | ↓-0.40 | 3.77E-01 | 4.60E-01 | 1.10 | ↑0.14 |
| I7GA03 | 2.25E-01 | 3.29E-01 | 0.92 | ↓-0.12 | 2.59E-01 | 3.32E-01 | 1.02 | ↑0.02 | 5.53E-01 | 6.30E-01 | 0.93 | ↓-0.11 | 9.04E-01 | 9.37E-01 | 0.85 | ↓-0.24 | 4.00E-01 | 5.00E-01 | 0.72 | ↓-0.47 | 9.24E-01 | 9.35E-01 | 0.75 | ↓-0.41 |
| H2R909 | 2.27E-01 | 3.30E-01 | 1.14 | ↑0.19 | 7.09E-01 | 7.66E-01 | 0.95 | ↓-0.07 | 6.56E-02 | 1.05E-01 | 0.83 | ↓-0.27 | 4.95E-02 | 8.12E-02 | 0.79 | ↓-0.34 | 1.51E-01 | 2.23E-01 | 0.84 | ↓-0.25 | 3.76E-02 | 6.20E-02 | 0.79 | ↓-0.35 |
| V9HWJ7 | 2.29E-01 | 3.30E-01 | 1.16 | ↑0.21 | 5.08E-02 | 8.44E-02 | 2.04 | ↑1.03 | 5.52E-01 | 6.30E-01 | 1.57 | ↑0.65 | 1.41E-01 | 2.03E-01 | 1.92 | ↑0.94 | 4.44E-04 | 1.27E-03 | 2.05 | ↑1.03 | 7.37E-02 | 1.15E-01 | 1.94 | ↑0.95 |

| | | | | | | | | | | | | | | | | | | | | | | | | |
|---|---|---|---|---|---|---|---|---|---|---|---|---|---|---|---|---|---|---|---|---|---|---|---|---|
| H2QMY8 | 2.30E-01 | 3.30E-01 | 0.86 | ↓-0.23 | 6.11E-02 | 9.82E-02 | 0.77 | ↓-0.38 | 1.32E-02 | 2.52E-02 | 0.69 | ↓-0.54 | 2.71E-02 | 5.03E-02 | 0.73 | ↓-0.46 | 3.50E-01 | 4.44E-01 | 0.86 | ↓-0.23 | 3.40E-02 | 5.71E-02 | 0.72 | ↓-0.47 |
| I7G968 | 2.33E-01 | 3.33E-01 | 0.87 | ↓-0.20 | 2.30E-02 | 4.12E-02 | 0.79 | ↓-0.34 | 4.53E-04 | 1.46E-03 | 0.68 | ↓-0.56 | 4.92E-02 | 8.12E-02 | 0.81 | ↓-0.30 | 4.68E-01 | 5.67E-01 | 0.91 | ↓-0.14 | 1.25E-01 | 1.77E-01 | 0.85 | ↓-0.24 |
| Q6N096 | 2.41E-01 | 3.42E-01 | 0.79 | ↓-0.34 | 9.71E-01 | 9.77E-01 | 2.45 | ↑1.29 | 2.38E-01 | 3.13E-01 | 2.28 | ↑1.19 | 3.67E-01 | 4.51E-01 | 0.74 | ↓-0.43 | 1.46E-01 | 2.17E-01 | 2.84 | ↑1.51 | 8.63E-01 | 8.80E-01 | 1.15 | ↑0.20 |
| P02671 | 2.42E-01 | 3.42E-01 | 0.88 | ↓-0.18 | 4.78E-01 | 5.55E-01 | 0.97 | ↓-0.05 | 2.11E-02 | 3.74E-02 | 0.72 | ↓-0.47 | 1.11E-01 | 1.68E-01 | 0.82 | ↓-0.28 | 8.25E-01 | 8.77E-01 | 0.99 | ↓-0.02 | 2.04E-01 | 2.61E-01 | 0.90 | ↓-0.15 |
| K7BA97 | 2.47E-01 | 3.46E-01 | 0.92 | ↓-0.13 | 1.76E-01 | 2.40E-01 | 0.86 | ↓-0.22 | 4.07E-03 | 9.39E-03 | 0.69 | ↓-0.54 | 9.89E-04 | 2.54E-03 | 0.65 | ↓-0.62 | 6.70E-01 | 7.49E-01 | 1.05 | ↑0.07 | 6.36E-01 | 7.00E-01 | 0.97 | ↓-0.04 |
| Q96PD5 | 2.48E-01 | 3.47E-01 | 0.86 | ↓-0.22 | 8.11E-01 | 8.59E-01 | 0.95 | ↓-0.07 | 1.17E-02 | 2.34E-02 | 0.71 | ↓-0.49 | 1.21E-01 | 1.81E-01 | 0.78 | ↓-0.35 | 2.95E-01 | 3.83E-01 | 0.89 | ↓-0.17 | 3.99E-01 | 4.80E-01 | 1.11 | ↑0.15 |
| G1SIB7 | 2.54E-01 | 3.53E-01 | 1.08 | ↑0.11 | 4.27E-01 | 5.02E-01 | 1.24 | ↑0.31 | 3.59E-01 | 4.48E-01 | 1.04 | ↑0.06 | 2.64E-01 | 3.41E-01 | 1.30 | ↑0.38 | 6.19E-01 | 7.07E-01 | 0.93 | ↓-0.10 | 8.26E-01 | 8.57E-01 | 0.78 | ↓-0.35 |
| H2Q8V0 | 2.69E-01 | 3.70E-01 | 0.65 | ↓-0.63 | 7.37E-02 | 1.16E-01 | 0.69 | ↓-0.53 | 7.61E-01 | 8.16E-01 | 0.95 | ↓-0.08 | 9.58E-01 | 9.69E-01 | 0.96 | ↓-0.06 | 2.11E-01 | 2.88E-01 | 1.05 | ↑0.07 | 3.21E-01 | 3.95E-01 | 0.60 | ↓-0.75 |
| P31151 | 2.69E-01 | 3.70E-01 | 1.10 | ↑0.14 | 1.14E-01 | 1.68E-01 | 0.43 | ↓-1.23 | 7.80E-01 | 8.29E-01 | 1.04 | ↑0.06 | 8.28E-03 | 1.67E-02 | 0.40 | ↓-1.31 | 1.52E-01 | 2.23E-01 | 1.24 | ↑0.31 | 2.25E-02 | 3.91E-02 | 2.74 | ↑1.45 |
| H0UT45 | 2.81E-01 | 3.84E-01 | 0.64 | ↓-0.64 | 2.69E-01 | 3.43E-01 | 1.03 | ↑0.04 | 2.13E-01 | 2.90E-01 | 1.09 | ↑0.12 | 4.77E-01 | 5.64E-01 | 0.75 | ↓-0.41 | 4.63E-01 | 5.64E-01 | 0.89 | ↓-0.16 | 5.95E-01 | 6.66E-01 | 0.93 | ↓-0.11 |
| B2R4C5 | 2.90E-01 | 3.94E-01 | 0.75 | ↓-0.42 | 1.64E-01 | 2.28E-01 | 2.81 | ↑1.49 | 8.63E-01 | 8.91E-01 | 1.01 | ↑0.02 | 7.99E-01 | 8.60E-01 | 0.86 | ↓-0.21 | 2.26E-02 | 4.35E-02 | 5.91 | ↑2.56 | 7.57E-01 | 7.93E-01 | 0.87 | ↓-0.21 |
| H2R580 | 2.91E-01 | 3.94E-01 | 0.92 | ↓-0.12 | 9.22E-01 | 9.44E-01 | 1.00 | ↓0.00 | 2.28E-01 | 3.04E-01 | 1.13 | ↑0.18 | 6.57E-01 | 7.35E-01 | 0.94 | ↓-0.10 | 5.05E-01 | 6.02E-01 | 1.15 | ↑0.20 | 4.66E-02 | 7.49E-02 | 1.34 | ↑0.43 |
| H0W536 | 3.12E-01 | 4.18E-01 | 0.99 | ↓-0.02 | 5.80E-01 | 6.46E-01 | 0.99 | ↓-0.01 | 1.58E-01 | 2.24E-01 | 0.91 | ↓-0.13 | 5.18E-01 | 6.10E-01 | 0.95 | ↓-0.07 | 9.05E-01 | 9.43E-01 | 1.04 | ↑0.05 | 1.83E-01 | 2.38E-01 | 1.35 | ↑0.44 |
| V9HWE3 | 3.13E-01 | 4.18E-01 | 1.38 | ↑0.47 | 6.92E-01 | 7.54E-01 | 1.22 | ↑0.28 | 4.16E-01 | 5.03E-01 | #N/A | #N/A | 9.38E-01 | 9.57E-01 | 1.06 | ↑0.08 | 3.99E-01 | 5.00E-01 | 1.45 | ↑0.54 | 2.57E-01 | 3.21E-01 | 1.27 | ↑0.34 |
| B4E1D8 | 3.39E-01 | 4.50E-01 | 0.88 | ↓-0.18 | 7.95E-01 | 8.52E-01 | 1.05 | ↑0.07 | 8.75E-01 | 8.99E-01 | 1.08 | ↑0.11 | 8.73E-01 | 9.18E-01 | 1.00 | ↓0.00 | 2.25E-01 | 3.02E-01 | 1.02 | ↑0.03 | 4.28E-01 | 5.06E-01 | 0.91 | ↓-0.14 |
| A0A2R9BYA2 | 3.50E-01 | 4.63E-01 | 1.29 | ↑0.37 | 2.94E-02 | 5.19E-02 | 1.40 | ↑0.49 | 1.55E-01 | 2.21E-01 | 1.25 | ↑0.32 | 1.81E-02 | 3.47E-02 | 1.34 | ↑0.43 | 5.20E-03 | 1.11E-02 | 1.49 | ↑0.57 | 4.94E-02 | 7.89E-02 | 1.35 | ↑0.44 |
| Dodecanedioic acid | 3.52E-01 | 4.63E-01 | 0.92 | ↓-0.12 | 3.83E-06 | 3.74E-05 | 0.57 | ↓-0.81 | 1.16E-06 | 1.02E-05 | 0.56 | ↓-0.84 | 2.38E-03 | 5.69E-03 | 0.85 | ↓-0.24 | 7.24E-05 | 2.51E-04 | 0.63 | ↓-0.66 | 3.86E-08 | 2.18E-07 | 0.51 | ↓-0.99 |
| A0A2J8QA94 | 3.67E-01 | 4.79E-01 | 0.96 | ↓-0.06 | 8.36E-01 | 8.77E-01 | 1.01 | ↑0.02 | 2.28E-01 | 3.04E-01 | 0.95 | ↓-0.07 | 6.92E-01 | 7.65E-01 | 0.99 | ↓-0.02 | 6.69E-01 | 7.49E-01 | 1.06 | ↑0.08 | 1.81E-01 | 2.38E-01 | 0.92 | ↓-0.11 |
| H2QJP2 | 3.68E-01 | 4.79E-01 | 0.89 | ↓-0.17 | 1.80E-01 | 2.44E-01 | 0.82 | ↓-0.28 | 2.59E-01 | 3.36E-01 | 0.86 | ↓-0.22 | 5.56E-01 | 6.39E-01 | 0.88 | ↓-0.18 | 9.47E-01 | 9.59E-01 | 0.99 | ↓-0.01 | 1.61E-01 | 2.17E-01 | 0.76 | ↓-0.40 |
| H2QX94 | 3.76E-01 | 4.87E-01 | 0.82 | ↓-0.28 | 1.31E-01 | 1.90E-01 | 0.68 | ↓-0.56 | 6.27E-01 | 6.99E-01 | 0.90 | ↓-0.15 | 4.11E-01 | 4.96E-01 | 0.93 | ↓-0.11 | 9.28E-01 | 9.50E-01 | 1.00 | ↓0.00 | 5.30E-01 | 6.11E-01 | 0.74 | ↓-0.43 |
| K7BH37 | 4.00E-01 | 5.14E-01 | 0.89 | ↓-0.17 | 9.25E-02 | 1.40E-01 | 0.80 | ↓-0.31 | 9.11E-02 | 1.39E-01 | 0.84 | ↓-0.26 | 9.18E-01 | 9.44E-01 | 0.94 | ↓-0.08 | 2.42E-02 | 4.60E-02 | 0.74 | ↓-0.44 | 2.51E-02 | 4.31E-02 | 0.71 | ↓-0.49 |
| K7C3E2 | 4.01E-01 | 5.14E-01 | 0.98 | ↓-0.03 | 2.21E-04 | 8.19E-04 | 0.66 | ↓-0.60 | 2.16E-03 | 5.37E-03 | 0.72 | ↓-0.48 | 2.19E-03 | 5.30E-03 | 0.72 | ↓-0.47 | 4.53E-03 | 1.00E-02 | 0.74 | ↓-0.44 | 5.61E-03 | 1.11E-02 | 0.75 | ↓-0.42 |
| K7B5N7 | 4.36E-01 | 5.54E-01 | 1.06 | ↑0.08 | 1.11E-01 | 1.65E-01 | 0.89 | ↓-0.16 | 9.66E-01 | 9.75E-01 | 0.98 | ↓-0.03 | 5.28E-01 | 6.18E-01 | 1.03 | ↑0.04 | 9.45E-01 | 9.59E-01 | 0.98 | ↓-0.02 | 5.80E-01 | 6.52E-01 | 1.10 | ↑0.13 |
| A5PL27 | 4.36E-01 | 5.54E-01 | 1.41 | ↑0.50 | 2.84E-01 | 3.61E-01 | 1.17 | ↑0.23 | 4.54E-01 | 5.45E-01 | 1.59 | ↑0.67 | 7.54E-01 | 8.18E-01 | 1.30 | ↑0.38 | 9.26E-01 | 9.50E-01 | 0.98 | ↓-0.03 | 3.82E-01 | 4.64E-01 | 1.12 | ↑0.17 |
| C16Plasm LPC | 4.41E-01 | 5.57E-01 | 0.90 | ↓-0.15 | 6.84E-02 | 1.08E-01 | 0.78 | ↓-0.36 | 1.83E-02 | 3.33E-02 | 0.72 | ↓-0.47 | 1.49E-04 | 4.96E-04 | 0.59 | ↓-0.76 | 2.58E-03 | 6.06E-03 | 0.67 | ↓-0.58 | 5.63E-01 | 6.37E-01 | 0.94 | ↓-0.10 |
| P18658 | 4.44E-01 | 5.58E-01 | 0.55 | ↓-0.86 | 8.09E-01 | 8.59E-01 | 0.75 | ↓-0.41 | 5.32E-01 | 6.17E-01 | 0.66 | ↓-0.60 | 4.74E-02 | 7.92E-02 | 0.41 | ↓-1.29 | 6.14E-01 | 7.07E-01 | 0.79 | ↓-0.34 | 9.63E-01 | 9.63E-01 | 0.80 | ↓-0.32 |
| G1SS91 | 4.47E-01 | 5.59E-01 | 1.06 | ↑0.08 | 1.56E-01 | 2.19E-01 | 1.14 | ↑0.19 | 1.70E-01 | 2.38E-01 | 0.87 | ↓-0.21 | 7.13E-01 | 7.81E-01 | 1.02 | ↑0.03 | 9.39E-01 | 9.58E-01 | 1.03 | ↑0.04 | 1.41E-01 | 1.96E-01 | 1.15 | ↑0.21 |
| K6ZY85 | 4.65E-01 | 5.80E-01 | 0.78 | ↓-0.35 | 4.98E-01 | 5.72E-01 | 1.32 | ↑0.40 | 7.80E-01 | 8.29E-01 | 0.73 | ↓-0.45 | 7.98E-01 | 8.60E-01 | 0.94 | ↓-0.09 | 2.84E-01 | 3.72E-01 | 1.14 | ↑0.19 | 2.35E-01 | 2.95E-01 | 0.60 | ↓-0.75 |
| A0A024R035 | 4.75E-01 | 5.88E-01 | 1.04 | ↑0.06 | 3.35E-02 | 5.82E-02 | 1.18 | ↑0.24 | 1.11E-01 | 1.66E-01 | 1.09 | ↑0.13 | 5.93E-01 | 6.73E-01 | 1.05 | ↑0.07 | 8.97E-02 | 1.39E-01 | 1.09 | ↑0.12 | 3.47E-02 | 5.81E-02 | 1.13 | ↑0.18 |
| A0A2R9CM19 | 4.93E-01 | 6.08E-01 | 0.90 | ↓-0.15 | 5.59E-02 | 9.11E-02 | 0.76 | ↓-0.39 | 4.55E-01 | 5.45E-01 | 0.93 | ↓-0.11 | 3.93E-01 | 4.78E-01 | 0.90 | ↓-0.16 | 8.25E-01 | 8.77E-01 | 1.06 | ↑0.08 | 3.76E-03 | 7.84E-03 | 0.69 | ↓-0.54 |
| A0A2R9B5S5 | 5.08E-01 | 6.23E-01 | 1.00 | ↑0.01 | 8.49E-01 | 8.84E-01 | 1.19 | ↑0.25 | 9.92E-01 | 9.92E-01 | 1.16 | ↑0.21 | 1.94E-01 | 2.60E-01 | 1.43 | ↑0.51 | 2.10E-01 | 2.88E-01 | 1.28 | ↑0.36 | 8.38E-03 | 1.61E-02 | 1.61 | ↑0.69 |
| V9HWK2 | 5.11E-01 | 6.24E-01 | 0.76 | ↓-0.40 | 8.50E-02 | 1.29E-01 | 0.68 | ↓-0.56 | 3.48E-01 | 4.37E-01 | 0.76 | ↓-0.40 | 3.35E-01 | 4.19E-01 | 0.72 | ↓-0.48 | 3.16E-01 | 4.04E-01 | 0.68 | ↓-0.55 | 7.15E-01 | 7.63E-01 | 0.80 | ↓-0.32 |
| LysoPC200 | 5.37E-01 | 6.52E-01 | 1.13 | ↑0.18 | 9.19E-03 | 1.81E-02 | 0.80 | ↓-0.33 | 1.78E-02 | 3.30E-02 | 0.85 | ↓-0.23 | 5.27E-05 | 2.00E-04 | 0.65 | ↓-0.63 | 2.83E-06 | 2.18E-05 | 0.58 | ↓-0.78 | 8.29E-02 | 1.28E-01 | 0.93 | ↓-0.10 |
| A0A024R962 | 5.56E-01 | 6.73E-01 | 1.00 | ↑0.00 | 4.01E-01 | 4.76E-01 | 0.89 | ↓-0.17 | 7.32E-02 | 1.15E-01 | 0.84 | ↓-0.25 | 1.07E-01 | 1.63E-01 | 0.83 | ↓-0.27 | 9.13E-01 | 9.43E-01 | 0.96 | ↓-0.07 | 1.85E-01 | 2.39E-01 | 0.87 | ↓-0.20 |
| P00734 | 5.70E-01 | 6.86E-01 | 0.79 | ↓-0.34 | 3.99E-01 | 4.76E-01 | 0.71 | ↓-0.50 | 3.61E-01 | 4.48E-01 | 0.99 | ↓-0.02 | 8.10E-02 | 1.27E-01 | 0.64 | ↓-0.64 | 3.03E-01 | 3.89E-01 | 0.65 | ↓-0.63 | 2.81E-01 | 3.49E-01 | 0.71 | ↓-0.49 |
| H2PYU7 | 5.87E-01 | 6.98E-01 | 0.99 | ↓-0.01 | 2.53E-01 | 3.27E-01 | 0.89 | ↓-0.17 | 3.93E-01 | 4.80E-01 | 1.12 | ↑0.16 | 9.18E-01 | 9.44E-01 | 0.96 | ↓-0.06 | 6.89E-01 | 7.64E-01 | 0.98 | ↓-0.03 | 1.18E-01 | 1.72E-01 | 0.93 | ↓-0.10 |
| A0A2S0BDD1 | 5.89E-01 | 6.98E-01 | 0.92 | ↓-0.12 | 2.12E-01 | 2.77E-01 | 0.81 | ↓-0.30 | 1.35E-01 | 1.95E-01 | 0.85 | ↓-0.24 | 7.66E-02 | 1.21E-01 | 0.80 | ↓-0.33 | 5.48E-01 | 6.42E-01 | 0.88 | ↓-0.18 | 2.80E-01 | 3.49E-01 | 0.85 | ↓-0.24 |
| P68223 | 5.92E-01 | 6.98E-01 | 1.25 | ↑0.33 | 9.48E-02 | 1.42E-01 | 1.28 | ↑0.36 | 5.40E-01 | 6.24E-01 | 0.87 | ↓-0.20 | 3.43E-01 | 4.25E-01 | 0.99 | ↓-0.01 | 6.49E-02 | 1.07E-01 | 1.76 | ↑0.81 | 4.37E-02 | 7.12E-02 | 1.31 | ↑0.39 |
| H2QUJ6 | 5.93E-01 | 6.98E-01 | 0.94 | ↓-0.09 | 9.12E-01 | 9.37E-01 | 1.00 | ↑0.01 | 7.98E-01 | 8.43E-01 | 0.97 | ↓-0.04 | 8.82E-03 | 1.76E-02 | 0.73 | ↓-0.46 | 1.84E-03 | 4.50E-03 | 0.69 | ↓-0.53 | 9.94E-02 | 1.48E-01 | 0.89 | ↓-0.18 |
| V9HVZ4 | 5.95E-01 | 6.98E-01 | 2.52 | ↑1.34 | 9.89E-01 | 9.89E-01 | 0.96 | ↓-0.05 | 6.48E-01 | 7.18E-01 | 1.10 | ↑0.13 | 5.35E-01 | 6.22E-01 | 1.13 | ↑0.18 | 2.40E-01 | 3.19E-01 | 1.64 | ↑0.71 | 1.85E-01 | 2.39E-01 | 0.99 | ↓-0.01 |
| H0VJK2 | 5.96E-01 | 6.98E-01 | 1.01 | ↑0.01 | 2.03E-01 | 2.67E-01 | 1.00 | ↓0.00 | 9.04E-01 | 9.26E-01 | 1.08 | ↑0.11 | 4.18E-01 | 5.01E-01 | 1.10 | ↑0.13 | 2.42E-02 | 4.60E-02 | 1.20 | ↑0.26 | 6.13E-01 | 6.83E-01 | 1.15 | ↑0.20 |
| LysoPC180 | 6.00E-01 | 6.99E-01 | 0.99 | ↓-0.02 | 4.94E-05 | 2.77E-04 | 0.82 | ↓-0.29 | 7.63E-04 | 2.23E-03 | 0.86 | ↓-0.22 | 4.88E-09 | 7.86E-08 | 0.74 | ↓-0.44 | 4.15E-06 | 2.77E-05 | 0.76 | ↓-0.39 | 9.12E-02 | 1.38E-01 | 0.94 | ↓-0.09 |
| A0A2R9BZ89 | 6.10E-01 | 7.06E-01 | 0.88 | ↓-0.18 | 4.86E-01 | 5.61E-01 | 1.10 | ↑0.13 | 7.24E-01 | 7.79E-01 | 0.88 | ↓-0.19 | 2.95E-01 | 3.78E-01 | 0.79 | ↓-0.33 | 8.50E-01 | 8.96E-01 | 0.98 | ↓-0.03 | 4.10E-01 | 4.91E-01 | 0.84 | ↓-0.26 |
| B7ZKJ8 | 6.12E-01 | 7.06E-01 | 1.55 | ↑0.64 | 1.02E-02 | 1.98E-02 | 2.32 | ↑1.22 | 1.30E-01 | 1.90E-01 | 1.77 | ↑0.82 | 6.63E-01 | 7.39E-01 | 1.21 | ↑0.27 | 6.44E-02 | 1.07E-01 | 2.12 | ↑1.09 | 4.00E-04 | 9.50E-04 | 2.78 | ↑1.47 |
| H2QNX1 | 6.36E-01 | 7.31E-01 | 1.04 | ↑0.06 | 1.75E-01 | 2.40E-01 | 0.78 | ↓-0.37 | 4.41E-02 | 7.42E-02 | 0.68 | ↓-0.56 | 6.26E-02 | 1.01E-01 | 0.65 | ↓-0.63 | 6.00E-01 | 6.96E-01 | 0.83 | ↓-0.26 | 9.50E-01 | 9.54E-01 | 0.88 | ↓-0.19 |
| Q4LE41 | 6.40E-01 | 7.32E-01 | 1.05 | ↑0.06 | 9.01E-01 | 9.31E-01 | 1.05 | ↑0.07 | 5.63E-01 | 6.38E-01 | 1.22 | ↑0.29 | 1.05E-01 | 1.61E-01 | 1.25 | ↑0.32 | 6.17E-01 | 7.07E-01 | 1.40 | ↑0.49 | 5.07E-01 | 5.88E-01 | 1.50 | ↑0.58 |
| Q6IBD2 | 6.46E-01 | 7.36E-01 | 0.99 | ↓-0.02 | 5.12E-01 | 5.83E-01 | 1.09 | ↑0.13 | 2.51E-01 | 3.28E-01 | 1.19 | ↑0.25 | 9.33E-01 | 9.55E-01 | 1.07 | ↑0.10 | 9.94E-01 | 9.94E-01 | 1.07 | ↑0.10 | 5.02E-01 | 5.85E-01 | 1.25 | ↑0.32 |
| G1THZ6 | 6.72E-01 | 7.62E-01 | 0.75 | ↓-0.41 | 1.91E-01 | 2.55E-01 | 0.64 | ↓-0.64 | 3.74E-01 | 4.60E-01 | 0.76 | ↓-0.39 | 8.75E-01 | 9.18E-01 | 0.89 | ↓-0.17 | 8.24E-01 | 8.77E-01 | 0.87 | ↓-0.21 | 6.29E-01 | 6.98E-01 | 0.83 | ↓-0.28 |
| Q9NZP8 | 6.83E-01 | 7.71E-01 | 1.10 | ↑0.13 | 2.66E-03 | 6.45E-03 | 0.64 | ↓-0.64 | 7.71E-02 | 1.19E-01 | 0.79 | ↓-0.34 | 1.89E-03 | 4.62E-03 | 0.64 | ↓-0.65 | 8.31E-01 | 8.80E-01 | 0.98 | ↓-0.03 | 4.42E-03 | 9.05E-03 | 0.66 | ↓-0.59 |
| D1MGQ2 | 6.89E-01 | 7.75E-01 | 1.16 | ↑0.21 | 8.08E-01 | 8.59E-01 | 0.93 | ↓-0.11 | 7.10E-01 | 7.70E-01 | 0.85 | ↓-0.23 | 5.37E-01 | 6.22E-01 | 0.80 | ↓-0.33 | 7.14E-01 | 7.85E-01 | 0.80 | ↓-0.32 | 4.12E-01 | 4.91E-01 | 1.41 | ↑0.49 |
| Q59HE0 | 6.95E-01 | 7.75E-01 | 0.95 | ↓-0.08 | 3.99E-01 | 4.76E-01 | 0.70 | ↓-0.51 | 6.27E-01 | 6.99E-01 | 0.68 | ↓-0.55 | 3.38E-01 | 4.21E-01 | 1.07 | ↑0.09 | 9.08E-01 | 9.43E-01 | 0.81 | ↓-0.30 | 1.60E-01 | 2.17E-01 | 1.11 | ↑0.15 |
| LysoPC140 | 6.98E-01 | 7.75E-01 | 1.38 | ↑0.46 | 5.32E-06 | 4.66E-05 | 0.66 | ↓-0.60 | 2.48E-06 | 1.97E-05 | 0.64 | ↓-0.64 | 4.94E-05 | 1.90E-04 | 0.68 | ↓-0.56 | 1.54E-01 | 2.25E-01 | 1.00 | ↓0.00 | 1.20E-07 | 5.98E-07 | 0.62 | ↓-0.70 |
| P02750 | 6.99E-01 | 7.75E-01 | 1.17 | ↑0.23 | 2.53E-01 | 3.27E-01 | 2.12 | ↑1.09 | 1.22E-04 | 4.76E-04 | 2.00 | ↑1.00 | 1.25E-02 | 2.49E-02 | 1.66 | ↑0.73 | 1.93E-08 | 3.07E-07 | 2.20 | ↑1.14 | 6.93E-02 | 1.09E-01 | 1.71 | ↑0.78 |
| Q14520 | 7.08E-01 | 7.80E-01 | 0.86 | ↓-0.22 | 6.99E-01 | 7.59E-01 | 0.97 | ↓-0.05 | 6.17E-01 | 6.94E-01 | 0.89 | ↓-0.17 | 1.59E-01 | 2.23E-01 | 0.70 | ↓-0.51 | 1.57E-01 | 2.26E-01 | 0.71 | ↓-0.48 | 3.87E-01 | 4.68E-01 | 0.81 | ↓-0.31 |
| K7DFV8 | 7.09E-01 | 7.80E-01 | 1.28 | ↑0.36 | 6.14E-01 | 6.75E-01 | 0.83 | ↓-0.27 | 2.16E-01 | 2.92E-01 | 1.37 | ↑0.45 | 8.42E-01 | 8.92E-01 | 1.25 | ↑0.32 | 1.65E-01 | 2.30E-01 | 0.67 | ↓-0.58 | 8.76E-01 | 8.90E-01 | 1.04 | ↑0.06 |
| A0A2R9A2B8 | 7.34E-01 | 8.03E-01 | 1.28 | ↑0.36 | 1.17E-01 | 1.72E-01 | 0.57 | ↓-0.82 | 3.61E-01 | 4.48E-01 | 0.74 | ↓-0.43 | 3.09E-01 | 3.90E-01 | 0.98 | ↓-0.04 | 7.67E-01 | 8.36E-01 | 0.78 | ↓-0.35 | 3.53E-01 | 4.33E-01 | 0.77 | ↓-0.38 |
| H2Q4U2 | 7.39E-01 | 8.06E-01 | 1.12 | ↑0.16 | 1.23E-01 | 1.79E-01 | 0.88 | ↓-0.18 | 3.58E-02 | 6.14E-02 | 0.79 | ↓-0.35 | 5.16E-03 | 1.10E-02 | 0.64 | ↓-0.65 | 2.94E-02 | 5.37E-02 | 0.79 | ↓-0.35 | 1.42E-03 | 3.16E-03 | 0.57 | ↓-0.82 |
| LysoPC181 | 7.60E-01 | 8.23E-01 | 1.02 | ↑0.03 | 3.32E-04 | 1.12E-03 | 0.79 | ↓-0.34 | 7.67E-03 | 1.58E-02 | 0.85 | ↓-0.23 | 1.55E-05 | 7.86E-05 | 0.72 | ↓-0.46 | 3.29E-06 | 2.46E-05 | 0.69 | ↓-0.54 | 2.34E-06 | 7.82E-06 | 0.72 | ↓-0.47 |
| P00735 | 7.61E-01 | 8.23E-01 | 0.85 | ↓-0.24 | 3.40E-02 | 5.87E-02 | 0.64 | ↓-0.65 | 4.80E-02 | 7.97E-02 | 0.72 | ↓-0.48 | 1.56E-01 | 2.20E-01 | 0.80 | ↓-0.32 | 7.17E-01 | 7.85E-01 | 0.93 | ↓-0.11 | 1.02E-01 | 1.50E-01 | 0.70 | ↓-0.51 |
| B4E1Z4 | 7.75E-01 | 8.30E-01 | 0.98 | ↓-0.03 | 8.86E-01 | 9.19E-01 | 1.02 | ↑0.03 | 1.52E-01 | 2.19E-01 | 0.90 | ↓-0.15 | 8.23E-01 | 8.79E-01 | 0.99 | ↓-0.02 | 9.91E-01 | 9.94E-01 | 0.99 | ↓-0.02 | 1.57E-01 | 2.14E-01 | 0.91 | ↓-0.14 |
| H2QZH0 | 7.75E-01 | 8.30E-01 | 0.89 | ↓-0.17 | 5.07E-01 | 5.80E-01 | 0.84 | ↓-0.25 | 7.08E-01 | 7.70E-01 | 0.86 | ↓-0.22 | 8.20E-01 | 8.79E-01 | 0.90 | ↓-0.15 | 2.23E-01 | 3.01E-01 | 0.78 | ↓-0.36 | 6.94E-01 | 7.49E-01 | 0.82 | ↓-0.28 |

| | | | | | | | | | | | | | | | | | | | | | | | | |
|---|---|---|---|---|---|---|---|---|---|---|---|---|---|---|---|---|---|---|---|---|---|---|---|---|
| B3KX75 | 7.78E-01 | 8.30E-01 | 0.92 | ↓-0.12 | 6.64E-02 | 1.05E-01 | 0.66 | ↓-0.59 | 1.28E-02 | 2.51E-02 | 0.61 | ↓-0.72 | 2.00E-01 | 2.65E-01 | 0.73 | ↓-0.46 | 2.15E-01 | 2.92E-01 | 0.75 | ↓-0.41 | 1.03E-02 | 1.96E-02 | 0.53 | ↓-0.91 |
| B2R888 | 8.09E-01 | 8.60E-01 | 0.99 | ↓-0.01 | 5.93E-01 | 6.58E-01 | 0.93 | ↓-0.11 | 1.18E-01 | 1.74E-01 | 0.84 | ↓-0.25 | 3.16E-01 | 3.97E-01 | 0.87 | ↓-0.20 | 6.60E-01 | 7.45E-01 | 0.94 | ↓-0.09 | 2.29E-02 | 3.95E-02 | 0.78 | ↓-0.35 |
| C9JEU5 | 8.15E-01 | 8.63E-01 | 1.82 | ↑0.87 | 3.60E-01 | 4.40E-01 | 1.10 | ↑0.13 | 9.91E-01 | 9.92E-01 | 1.08 | ↑0.12 | 1.62E-01 | 2.25E-01 | 1.31 | ↑0.39 | 4.78E-01 | 5.75E-01 | 1.14 | ↑0.19 | 1.18E-01 | 1.72E-01 | 1.19 | ↑0.25 |
| P11717 | 8.19E-01 | 8.64E-01 | 1.03 | ↑0.05 | 5.31E-01 | 5.98E-01 | 1.04 | ↑0.06 | 8.22E-01 | 8.59E-01 | 0.95 | ↓-0.08 | 8.39E-01 | 8.91E-01 | 0.93 | ↓-0.11 | 6.82E-01 | 7.60E-01 | 0.88 | ↓-0.18 | 8.34E-01 | 8.61E-01 | 0.90 | ↓-0.16 |
| B2R4M6 | 8.48E-01 | 8.90E-01 | 1.34 | ↑0.43 | 4.44E-02 | 7.52E-02 | 2.91 | ↑1.54 | 3.02E-01 | 3.85E-01 | 2.14 | ↑1.10 | 1.79E-01 | 2.43E-01 | 0.62 | ↓-0.70 | 1.09E-02 | 2.21E-02 | 3.99 | ↑2.00 | 8.57E-01 | 8.78E-01 | 2.90 | ↑1.54 |
| A0A024R6I7 | 8.52E-01 | 8.91E-01 | 0.91 | ↓-0.14 | 1.35E-01 | 1.94E-01 | 1.38 | ↑0.46 | 5.46E-01 | 6.27E-01 | 1.79 | ↑0.84 | 8.98E-01 | 9.37E-01 | 1.05 | ↑0.07 | 9.05E-02 | 1.39E-01 | 1.38 | ↑0.46 | 7.32E-03 | 1.44E-02 | 1.83 | ↑0.87 |
| G1SJM1 | 8.62E-01 | 8.98E-01 | 1.00 | ↓0.00 | 4.18E-03 | 8.92E-03 | 0.84 | ↓-0.24 | 1.19E-03 | 3.24E-03 | 0.82 | ↓-0.28 | 2.90E-03 | 6.70E-03 | 0.84 | ↓-0.25 | 1.31E-05 | 6.15E-05 | 0.77 | ↓-0.38 | 4.90E-03 | 9.87E-03 | 0.82 | ↓-0.29 |
| P48770 | 8.82E-01 | 9.14E-01 | 0.99 | ↓-0.02 | 7.68E-01 | 8.26E-01 | 1.08 | ↑0.11 | 8.53E-01 | 8.85E-01 | 0.98 | ↓-0.04 | 7.22E-01 | 7.88E-01 | 1.05 | ↑0.06 | 3.02E-01 | 3.89E-01 | 1.05 | ↑0.07 | 6.80E-01 | 7.38E-01 | 1.03 | ↑0.04 |
| H2QBZ3 | 8.88E-01 | 9.14E-01 | 0.96 | ↓-0.06 | 1.54E-02 | 2.85E-02 | 0.57 | ↓-0.80 | 4.12E-01 | 5.00E-01 | 0.74 | ↓-0.43 | 2.02E-01 | 2.68E-01 | 0.83 | ↓-0.28 | 1.45E-01 | 2.17E-01 | 0.84 | ↓-0.26 | 7.50E-01 | 7.93E-01 | 1.40 | ↑0.48 |
| Q5JVE7 | 8.89E-01 | 9.14E-01 | 0.98 | ↓-0.03 | 1.60E-01 | 2.24E-01 | 0.86 | ↓-0.22 | 6.51E-02 | 1.05E-01 | 0.82 | ↓-0.28 | 2.14E-01 | 2.80E-01 | 0.87 | ↓-0.20 | 4.80E-01 | 5.76E-01 | 0.89 | ↓-0.16 | 3.52E-03 | 7.39E-03 | 0.71 | ↓-0.50 |
| V9HVY1 | 8.96E-01 | 9.14E-01 | 2.44 | ↑1.29 | 4.17E-01 | 4.93E-01 | 0.92 | ↓-0.13 | 1.16E-01 | 1.72E-01 | 0.83 | ↓-0.27 | 1.34E-01 | 1.97E-01 | 0.86 | ↓-0.22 | 4.58E-01 | 5.64E-01 | 1.27 | ↑0.35 | 1.22E-01 | 1.75E-01 | 0.83 | ↓-0.27 |
| LysoPE160 | 8.96E-01 | 9.14E-01 | 1.46 | ↑0.54 | 8.09E-04 | 2.34E-03 | 0.77 | ↓-0.38 | 3.53E-05 | 1.63E-04 | 0.71 | ↓-0.50 | 9.07E-05 | 3.16E-04 | 0.70 | ↓-0.51 | 1.34E-01 | 2.02E-01 | 0.99 | ↓-0.01 | 2.23E-06 | 7.54E-06 | 0.68 | ↓-0.56 |
| P07357 | 9.01E-01 | 9.15E-01 | 0.97 | ↓-0.04 | 3.59E-01 | 4.40E-01 | 0.86 | ↓-0.22 | 6.10E-02 | 9.99E-02 | 0.83 | ↓-0.27 | 9.90E-01 | 9.90E-01 | 0.97 | ↓-0.04 | 7.90E-01 | 8.54E-01 | 0.99 | ↓-0.01 | 8.23E-01 | 8.56E-01 | 0.94 | ↓-0.08 |
| Q8TAY0 | 9.68E-01 | 9.80E-01 | 1.00 | ↑0.01 | 1.92E-01 | 2.55E-01 | 0.87 | ↓-0.20 | 1.28E-01 | 1.88E-01 | 0.85 | ↓-0.23 | 2.85E-02 | 5.24E-02 | 0.80 | ↓-0.32 | 1.43E-03 | 3.54E-03 | 0.72 | ↓-0.48 | 1.53E-01 | 2.11E-01 | 0.88 | ↓-0.18 |
| Q60FE6 | 9.85E-01 | 9.89E-01 | 1.00 | ↓-0.01 | 9.41E-01 | 9.56E-01 | 1.17 | ↑0.23 | 6.54E-01 | 7.22E-01 | 0.90 | ↓-0.15 | 9.79E-01 | 9.83E-01 | 0.92 | ↓-0.12 | 7.79E-01 | 8.46E-01 | 0.87 | ↓-0.20 | 7.55E-01 | 7.93E-01 | 0.97 | ↓-0.04 |
| A0A2R9BCI1 | 9.88E-01 | 9.89E-01 | 0.96 | ↓-0.06 | 9.53E-01 | 9.64E-01 | 0.98 | ↓-0.03 | 9.30E-01 | 9.49E-01 | 1.13 | ↑0.17 | 9.56E-01 | 9.69E-01 | 1.42 | ↑0.51 | 9.99E-02 | 1.52E-01 | 0.68 | ↓-0.56 | 1.81E-01 | 2.38E-01 | 0.72 | ↓-0.47 |
| P33908 | 9.89E-01 | 9.89E-01 | 1.15 | ↑0.20 | 9.73E-01 | 9.77E-01 | 1.05 | ↑0.07 | 6.89E-01 | 7.54E-01 | 1.04 | ↑0.06 | 9.77E-01 | 9.83E-01 | 1.11 | ↑0.15 | 3.29E-02 | 5.92E-02 | 1.62 | ↑0.70 | 8.57E-02 | 1.32E-01 | 1.78 | ↑0.83 |
